# Genome-wide association study in two cohorts from a multi-generational mouse advanced intercross line highlights the difficulty of replication

**DOI:** 10.1101/387613

**Authors:** Xinzhu Zhou, Celine L. St. Pierre, Natalia M. Gonzales, Riyan Cheng, Apurva Chitre, Greta Sokoloff, Abraham A. Palmer

**Affiliations:** Biomedical Sciences Graduate Program, University of California San Diego, La Jolla, CA, USA; Department of Genetics, Washington University School of Medicine, St. Louis, MO, USA; Department of Human Genetics, University of Chicago, Chicago, IL, USA; Department of Psychiatry, University of California San Diego, La Jolla, CA, USA; Department of Psychological & Brain Sciences, University of Iowa, Iowa City, IO, USA; Institute for Genomic Medicine, University of California San Diego, La Jolla, CA, USA

## Abstract

Replication is considered to be critical for genome-wide association studies (**GWAS**) in humans, but is not routinely performed in model organisms. We explored replication using an advanced intercross line (**AIL**) which is the simplest possible multigenerational intercross. We re-genotyped a previously published cohort of LG/J x SM/J AIL mice (F_34_; n=428) using a denser marker set and also genotyped a novel cohort of AIL mice (F_39-43_; n=600) for the first time. We identified 110 significant loci in the F_34_ cohort, 36 of which were new discoveries attributable to the denser marker set; we also identified 27 novel significant loci in the F_39-43_ cohort. For traits measured in both cohorts (locomotor activity, body weight, and coat color), the genetic correlations were high, although, the F_39-43_ cohort showed systematically lower SNP-heritability estimates. We then attempted to replicate loci identified in either F_34_ or F_39-43_ in the other cohort. Albino coat color was robustly replicated; we observed only partial replication of associations for locomotor activity and body weight. Finally, we performed a mega-analysis of locomotor activity and body weight by combining F_34_ and F_39-43_ cohorts (n=1,028), which identified four novel loci. The incomplete replication was inconsistent with simulations we performed to estimate our power to replicate. This may reflect: 1) false positives errors in the discovery cohort, 2) environmental or genetic heterogeneity between the two samples, or 3) the systematic over estimation of the effect sizes at significant loci (“Winner’s Curse”). Our results demonstrate that it is difficult to replicate GWAS results even when using similarly sized discovery and replication cohorts drawn from the same population.

## Introduction

Despite its importance in human GWAS, replication has been infrequently used for GWAS in model organisms. Challenges in replicating human GWAS findings include genetic, demographic or environmental differences between cohorts. In contrast, model organism GWAS can use genetically identical cohorts phenotyped under extremely similar conditions, which would be expected to enhance the success of replication. We set out to investigate replication in model organism GWAS using an intercross mouse population. The use of multi-parental crosses and commercially available outbred populations for GWAS in model organisms such as mice (1–17), rats (18), chickens (19,20), zebrafish (21,22), fruit flies (23–27), *C. elegans* (28) and various plant species (29–31) has become increasingly common over the last decade. These mapping populations can further be categorized as multi-parental crosses, which are created by interbreeding two or more inbred strains, and commercially available outbred populations, in which the founders are of unknown provenance. An F_2_ cross between two inbred strains is the prototypical mapping population; however, F_2_s provide poor mapping resolution (32). To improve mapping resolution, Darvasi and Soller (33) proposed the creation of advanced intercross lines (**AILs**), which are produced by intercrossing F_2_ mice for additional generations. AILs accumulate additional crossovers with every successive generation, leading to a population with shorter LD blocks, which improves mapping precision, albeit at the expense of power (32,34).

The longest running mouse AIL was generated by crossing LG/J and SM/J inbred strains, which were selectively bred for large and small body size. We obtained this AIL in 2006 at generation 33 from Dr. James Cheverud (Jmc: LG,SM-G_33_). Since then, we have collected genotype and phenotype information from multiple generations, including F_34_ (16,35–38), F_39_-F_43_ and F_50-56_ (39). Our previous publications using the F_34_ generation employed a custom Illumina Infinium genotyping microarray to obtain genotypes for 4,593 SNPs (35,36), we refer to this set of SNPs as the ‘sparse markers’. Those genotypes were used to identify significant associations for numerous traits, including methamphetamine sensitivity (35), pre-pulse inhibition (16), musculoskeletal measurements (17), muscle weight (37), body weight (40), open field (36), conditioned fear (36), red blood cell parameters (41), and murine soleus muscle (38). Although not previously published, we also collected phenotype information from the F_39_-F_43_ generations, including body weight, fear conditioning, locomotor activity in response to methamphetamine, and the light dark test for anxiety.

While the prior GWAS using the F_34_ generation detected many significant loci, the sparsity of the markers likely precluded the discovery of some true loci, and also made it difficult to clearly define the boundaries of the loci that we did identify. For example, Parker et al conducted an integrated analysis of F_2_ and F_34_ AILs (40). One of their body weight loci spanned from 87.93–102.70 Mb on chromosome 14. Denser markers might have more clearly defined implicated region. In the present study, we used genotyping-by-sequencing (**GBS**), which is a reduced-representation sequencing method (42–44), to obtain a much denser set of SNPs in the F_34_ and to genotype mice from the F_39_-F_43_ generations for the first time. With this denser set of SNPs, we attempted to identify novel loci in the F_34_s that were not detected using the sparse SNPs. We also performed a GWAS using the mice from the F_39_-F_43_ AILs. We explored whether imputation from the array SNPs could have provided the additional coverage we obtained using the denser GBS genotypes. Because F_39_-F_43_ AILs are direct descendants of the F_34_, they are uniquely suited to serve as a replication population for GWAS in the F_34_ generation. Using multiple cohorts of the same mouse intercross strain, we attempted to examine which loci are replicable between the F_34_ and F_39-43_ LG/J x SM/J AILs and also performed simulations to estimate the power for these replica on studies. In addition to their use as a replication sample, the F_39_-F_43_ can also provide improved resolution and allow for the discovery of novel loci not detected in the F_34_ generation. Therefore, we performed a mega-analysis of F_34_ mice and F_39_-F_43_ mice to identify loci that were not identified in either individual dataset. Apart from identifying novel loci with a range of physiological and behavioral traits in mice, the present study explores replication in a mouse intercross line by comparing association findings of 1) the same cohort with two genotype panels, 2) two cohorts in the same intercross line, 3) mega-analysis of combined cohorts, and 4) two cohorts with imputed genotypes.

## Results

We used 214 males and 214 females from generation F_34_ (Aap:LG,SM-G34) and 305 males and 295 females from generations F_39-43_. For the F_34_ AIL 79 traits were available from previous published and unpublished, for the F_39-43_ AIL 49 unpublished traits were available (S1 Table). F_34_ mice had been previously genotyped on a custom SNP array (35,36). The average minor allele frequency (**MAF**) of those 4,593 array SNPs was 0.388 (Fig 1). To obtain a denser set of SNP markers, we used genotyping-by-sequencing in F_34_ and F_39-43_ AIL mice. Since data about the F_39-43_ AIL mice had been collected over the span of approximately two years, we carefully considered the possibility of sample contamination and sample mislabeling (45). We removed samples based on four major features: heterozygosity distribution, number of reads aligned to sex chromosomes, discrepancies between pedigree and genetic kinship relatedness, and coat color genotype to phenotype mismatch (see Methods; S1 and S2 Figs). The final SNP sets included 60,392 GBS-derived SNPs in 428 F_34_ AIL mice, 59,790 GBS-derived SNPs in 600 F_39-43_ AIL mice, and 58,461 GBS-derived SNPs that existed in both F_34_ and F_39-43_ AIL mice (S2 Table). The MAF for the GBS SNPs was 0.382 in F_34_, 0.358 in F_39-43_, and 0.370 in F_34_ and F_39-43_ (Fig 1). There were 66 SNPs called from our GBS data that were also present on the genotyping array. The genotype concordance rate for those 66 SNPs, which reflects the sum of errors from both sets of genotypes, was 95.4% (S3 Fig). We found that LD decay rates using F_34_ array, F_34_ GBS, F_39-43_ GBS, and F_34_ and F_39-43_ GBS genotypes were generally similar to one another, though levels of LD using the GBS genotypes appear to be slightly reduced in the later generations of AILs (S4 Fig).

**Fig 1.**
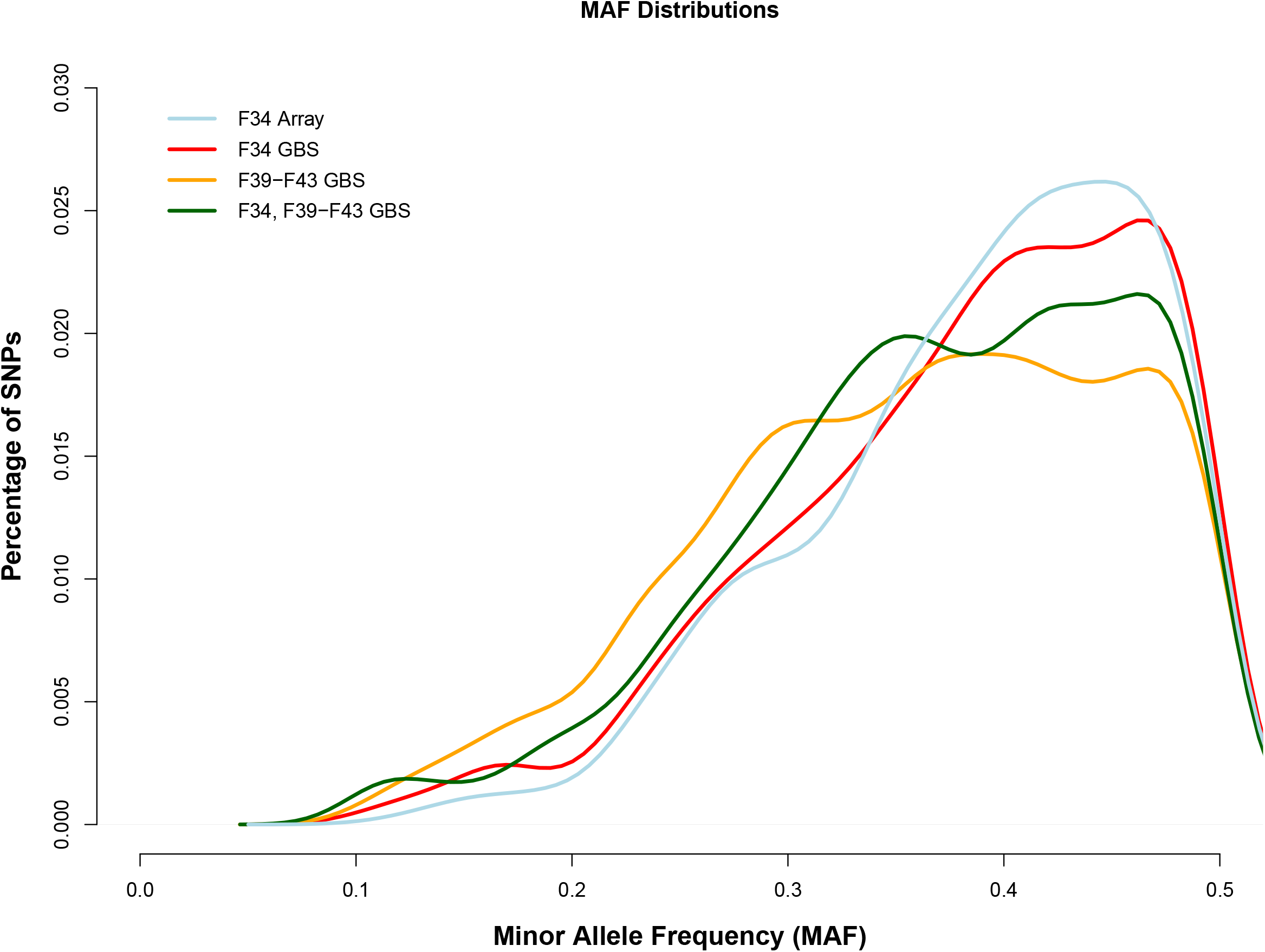
Minor allele frequency distributions for F_34_ array, F_34_ GBS, F_39_-F_43_ GBS, and F_34_ and F_39_-F_43_ GBS SNP sets. MAF distributions are highly comparable between AIL generations.

### GBS genotypes produced more significant associations than array genotypes in F_34_

We used a linear mixed model (**LMM**) as implemented in GEMMA (46) to perform GWAS. We used the leave-one-chromosome-out (**LOCO**) approach to address the problem of proximal contamination, as previously described (39,47–49). We performed GWAS using both the sparse array SNPs and the dense GBS SNPs to determine whether additional SNPs would produce additional genome-wide significant associations. Autosomal and X chromosome SNPs were included in all GWAS. We obtained a significance threshold for each SNP set using MultiTrans and SLIDES (50,51).To select independently associated loci (“lead loci”), we used a LD-based clumping method implemented in PLINK to group SNPs that passed the adjusted genome-wide significance thresholds over a large genomic region flanking the index SNP (52). Applying the most stringent clumping parameters (*r*^2^ = 0.1 and sliding window size = 12,150kb, S3 Table), we identified 110 significant lead loci in 49 out of 79 F_34_ phenotypes using the GBS SNPs. Table 1 contains the significant associations that contained 5 or less genes; additional significant associations that contained more than 5 genes are listed in S4 Table. In contrast, we identified 83 significant lead loci in 45 out of 79 F_34_ phenotypes using the sparse array SNPs (Table 1, S4 Table). Among the loci identified in the F_34_, 36 were uniquely identified using the GBS genotypes, whereas 11 were uniquely identified using the array genotypes. GBS SNPs consistently yielded more significant lead loci compared to array SNPs regardless of the clumping parameter values (S3 Table), indicating that a dense marker panel was able to detect more association signals compared to a sparse marker panel.

**Table 1.**
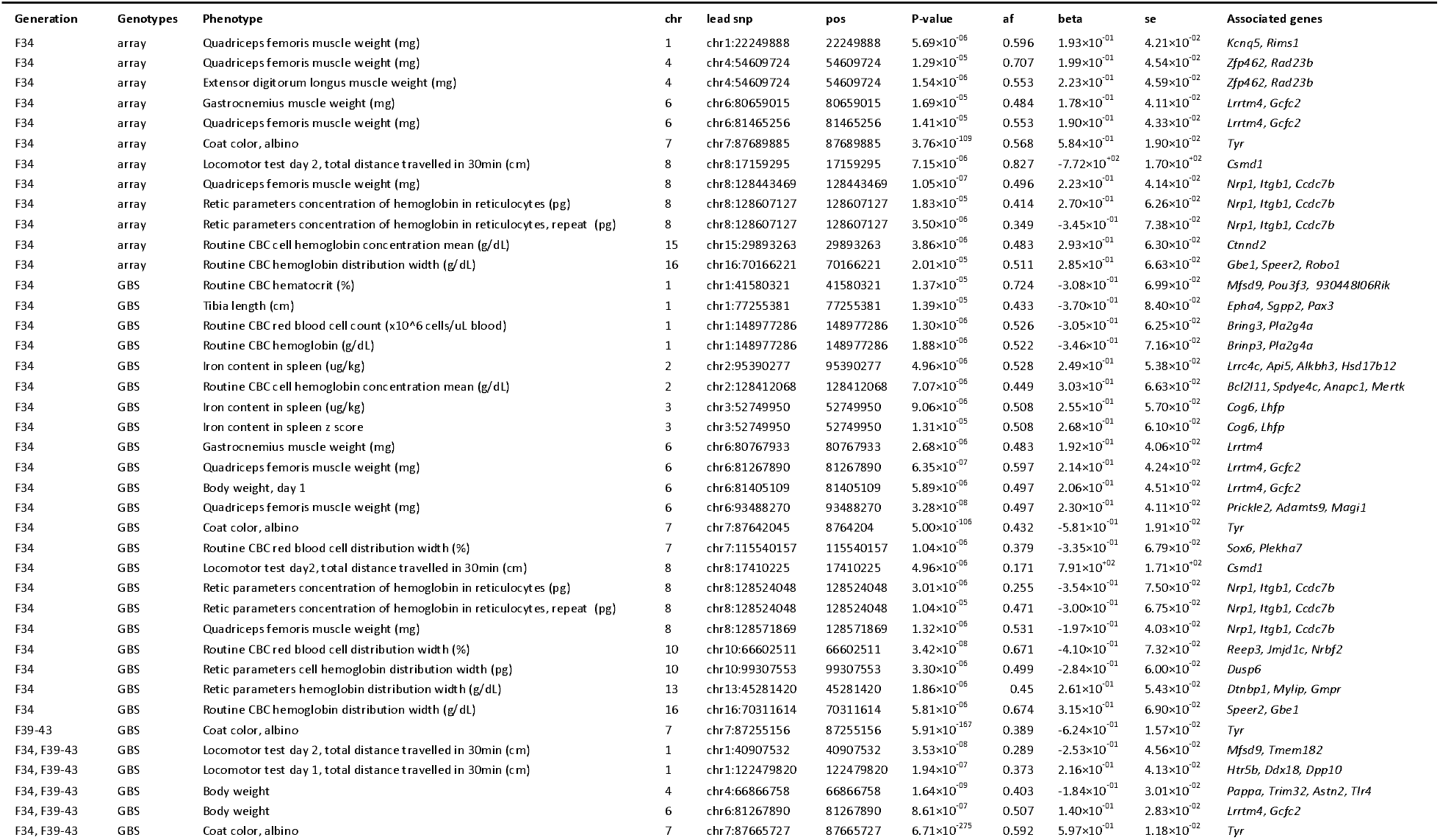
Select lead SNPs with association regions containing less than 5 coding genes. Credible set analysis was performed to define the boundaries of the locus (*r*^2^ threshold = 0.8, posterior probability threshold = 0.99). Genes contained in and/or immediately downstream of the credible set interval were included as associated genes.

To determine the boundaries of each locus, we performed a Bayesian-framework credible set analysis, which estimated a posterior probability for association at each SNP (*r*^2^ threshold = 0.8, posterior probability threshold = 0.99; (53)). The physical positions of the SNPs in the credible set were used to determine the boundaries of each locus. As expected, the greater density of the GBS genotypes allowed us to better define each interval. For instance, the lead locus at chr17:27130383 was associated with distance travelled in periphery in the open filed test in F_34_ AILs (Fig 2). However, no SNPs were genotyped between 26.7 and 28.7 Mb in the array SNPs, which makes the size of this LD block ambiguous. In contrast, the LocusZoom plot portraying GBS SNPs in the same region shows that SNPs in high LD with chr17:27130383 are between 27 Mb and 28.3 Mb. The more accurate definition of the implicated intervals allowed us to better refine the list of the coding genes and non-coding variants associated with the phenotype (Table 1).

**Fig 2.**
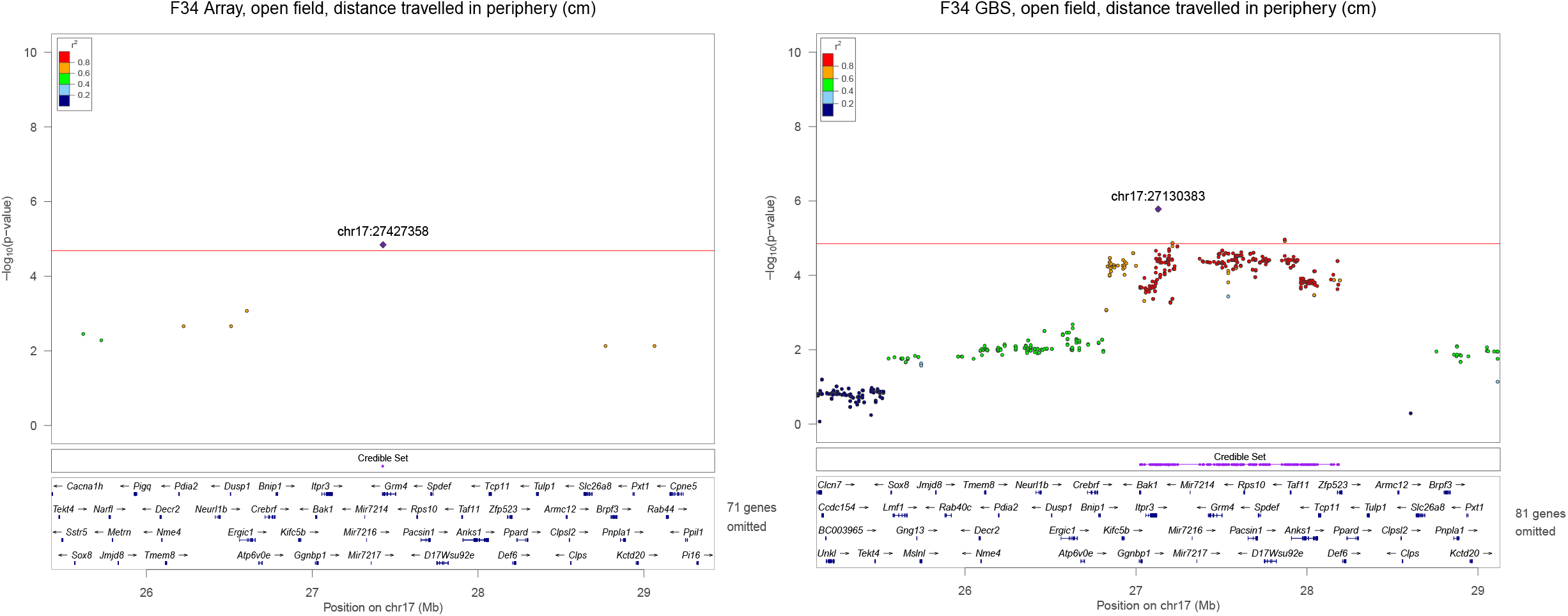
Significant loci on chromosome 17 for open field, distance traveled in periphery in F_34_ AIL. As exemplified in this pair of LocusZoom plots, GBS SNPs defined the boundaries of the loci much more precisely than array SNPs. GBS SNPs that are in high LD (*r*_2_ > 0.8, red dots) with lead SNP chr17:27130383 resides between 27 ~ 28.3 Mb. In contrast, too few SNPs are present in the array plot to draw any definitive conclusion about the boundaries or LD pattern in this region. Purple track shows the credible set interval. LocusZoom plots for all loci identified in this paper are in Fig S7.

In our prior studies using the sparse marker set, we did not attempt to increase the number of available markers by using imputation. Therefore, we examined whether the disparity between the numbers of loci identified by the two SNP sets could be resolved by imputation, which should increase the number of markers available for GWAS. We used LG/J and SM/J whole genome sequencing data as reference panels (54) and performed imputation on array and GBS SNPs using Beagle v4.1 (55). After QC filtering, we obtained 4.3M SNPs imputed from the array SNPs and 4.1M SNPs imputed from the GBS SNPs. More imputed GBS SNPs were filtered out because GBS SNPs were called from genotype probabilities, thus introducing uncertainty in imputed SNPs. We found that imputed array genotypes and imputed GBS genotypes did not meaningfully increase the number of loci discovered (Fig S5).

Under a polygenic model where a large number of additive common variants contribute to a complex trait, heritability estimates could be higher when more SNPs are considered (56). Given that there were more GBS SNPs than array SNPs, we used autosomal SNPs to examine whether GBS SNPs would generate higher SNP heritability estimates compared to the sparse array SNPs. Heritability estimates were similar for the two SNP sets, with the exception of agouti coat color, which showed marginally greater heritability for the GBS SNPs (S6 Fig; S5 Table). Our results show that while the denser GBS SNP set was able to identify more genome-wide significant loci, greater SNP density did not improve the polygenic signal.

### Partial replication of loci indented in F_34_ or F_39-43_ and mega-analysis

We identified 27 genome-wide significant loci for 21 phenotypes in the F_39-43_ cohort. A subset of those traits, including coat color, body weight, and locomotor activity, were also phenotyped in the F_34_ AILs (Table 2; S8 Table). To assess replication, we determined whether loci that were significant in one cohort (either F_34_ or F_39-43_) would also be significant in the other. We termed the cohort in which a locus was initially discovered as its “discovery set” and the cohort we attempted replication in as the “replication set” (Table 2). Coat color phenotypes (both albino and agouti) are Mendelian traits and thus served as positive control. As expected, all coat color and body weight loci were replicated. The three body weight loci identified in the F_34_ were replicated at nominal levels of significance (p<0.05) in F_39-43_; similarly, the one body weight locus identified in F_39-43_ was replicated in F_34_ (p<0.05). However, none of the five locomotor activity loci were replicated in the reciprocal (replication) cohorts. We then considered the more liberal “sign test” to determine whether the directions of the effect (beta) of the coat color, body weight and activity loci were in the same direction between the discovery and replication cohorts. We found that 11 of 13 loci passed this much less stringent test of replication. The two loci that did not pass the sign test were the two locomotor loci “discovered” in F_39-43_ (Table 2).

**Table 2.**
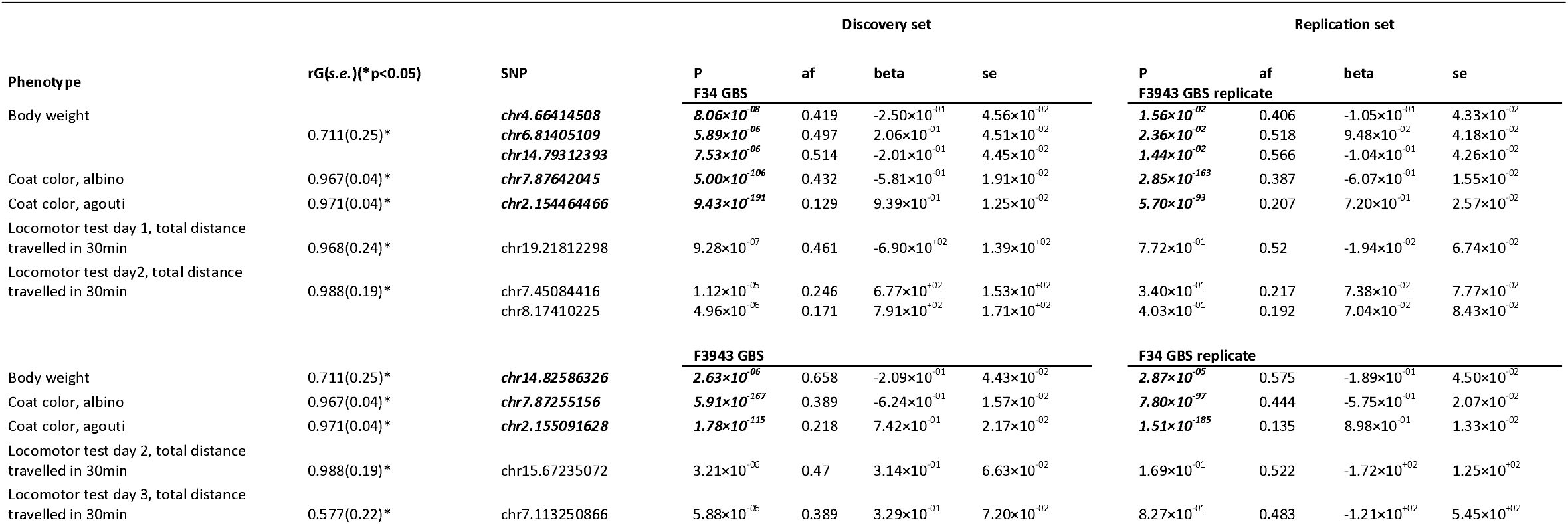
Replication of significant SNPs between F_34_ and F_39-43_ AIL association analyses. “Discovery set” indicates the AIL generation that significant SNPs were identified. “Replication set” shows the association p-value, β estimates, etc. of the “discovery set” significant SNPs in the replication AIL generation. SNPs that replicated (p<0.05, same sign for the beta) between F_34_ and F_39-43_ are highlighted in bold italics. Genetic correlations for phenotypes measured in both F_34_ and F_39-43_ are listed (see also Supplementary Table 6).

In light of the failure to replicate the locomotor activity findings, we conducted a series of 2,500 simulations per trait to estimate the expected power of our replication cohorts. For each phenotype we used the kinship relatedness matrix and variance components estimated from the replication set. For the coat color traits, we found that we had 100% power to replicated the association at either genome-wide significant levels or the more liberal p<0.05 threshold (S8 Fig). For body weight and locomotor activity, power to replicate at a genome-wide significance threshold ranged from 50% to 80%, whereas power to replicate at the p<0.05 threshold was nearly 100% (S8 Fig). These power estimates were clearly inconsistent with our empirical observations for the locomotor activity traits, none of which replicated at even the p<0.05 threshold, where we should have had almost 100% power (Table 2). However, our power simulations did not account for “Winner’s Curse”, which would be expected to systematically overestimate of effect size estimates used in our simulations (57).

We also evaluated whether or not the traits showed genetic correlations across the two cohorts; low genetic correlations between the two cohorts could indicate that environmental sources of heterogeneity had limited the potential for replication (F_34_ and F_39-43_). We used autosomal SNPs to calculate genetic correlations between the F_34_ and F_39-43_ generations for body weight, coat color, and locomotor activity phenotypes (S6 Table), using GCTA-GREML (58). Albino and agouti coat color, body weight and locomotor activity on days 1 and 2 were highly genetically correlated (r_G_s >0.7; S6 Table). In contrast, locomotor activity on day 3 showed a significant but weaker genetic correlation (r_G_=0.577), perhaps reflecting variability in the quality of the methamphetamine injection, which were only given on day 3. Overall, these results suggest that genetic influences on these traits were broadly similar in the two cohorts; however, the genetic correlations were less than 1, suggesting an additional barrier to replication that was not accounted for in our power simulations.

We also calculated the SNP heritabilities for all traits using GCTA. SNP heritability was consistently lower in the F_39-43_ compared to the F_34_, possibly a result of increased experimental variance introduced by our extended phenotype collection period (Fig 3; S7 Table).

**Fig 3.**
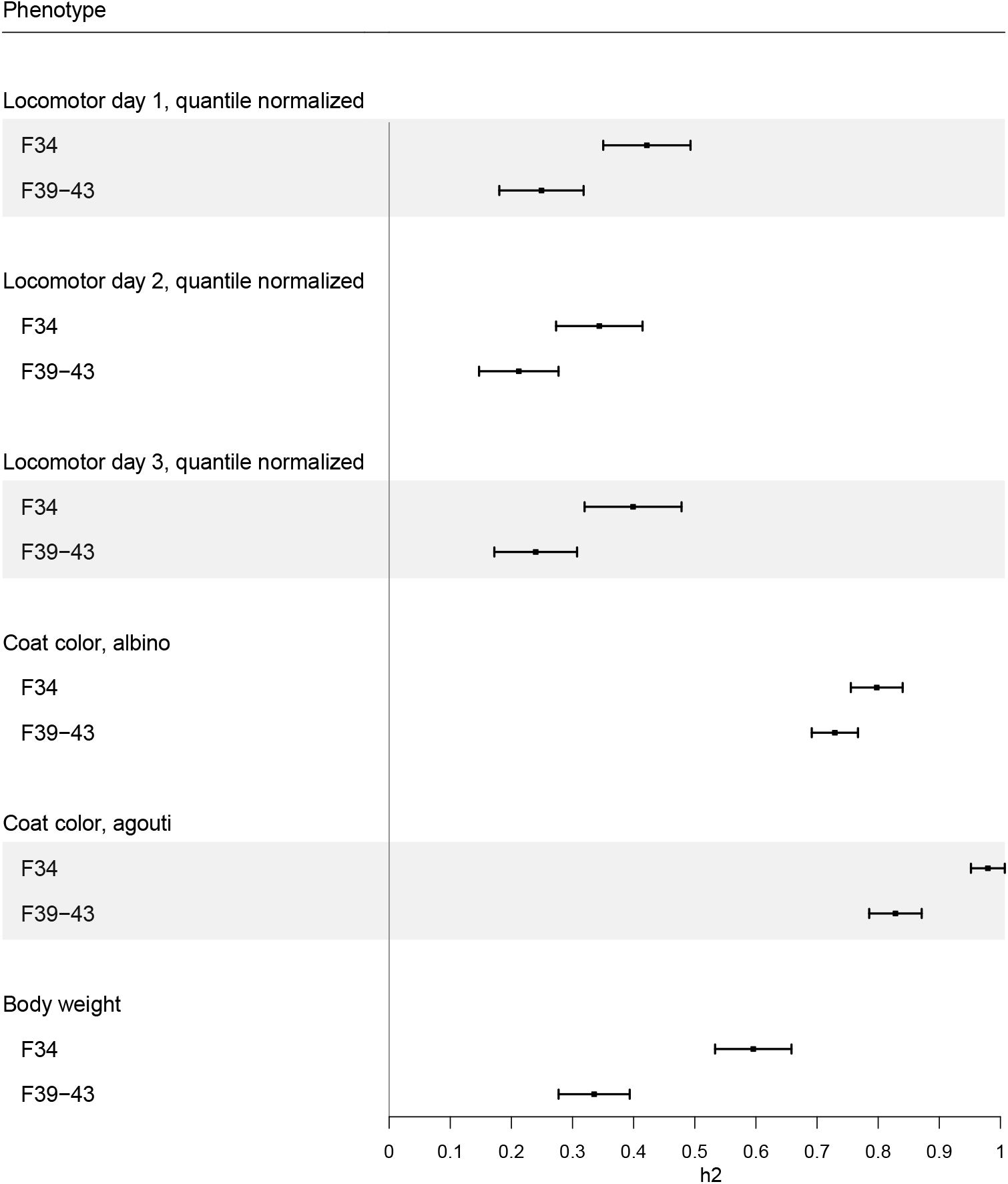
Chip-heritability estimates in F_34_ and F_39-43_ AILs. All heritability estimates are highly significant (p < 1.0×10-05; see S7 Table).

Due to the relatively high genetic correlations (S6 Table), we suspected that a mega-analysis using the combined sample set would allow for the identification of additional loci; indeed, mega-analysis identified four novel genome-wide significant associations (Fig 4; S9 Table). The significance of the associations identified by the mega-analysis was often greater than in either individual cohort. For instance, the p-values obtained by mega-analysis for chr4:66866758 (p = 1.64×10^−9^) and chr14:82672838 (p = 2.06×10^−10^) for body weight were lower than the corresponding p-values for the same loci for F_34_ (chr4:65246120, p = 9.06×10^−8^; chr14:78926547, p = 6.24×10^−6^) and F_39-43_ (chr4:66414508, p = 8.06×10^−8^; chr14:79312393, p = 7.53×10^−6^; S7 Fig).

**Fig 4.**
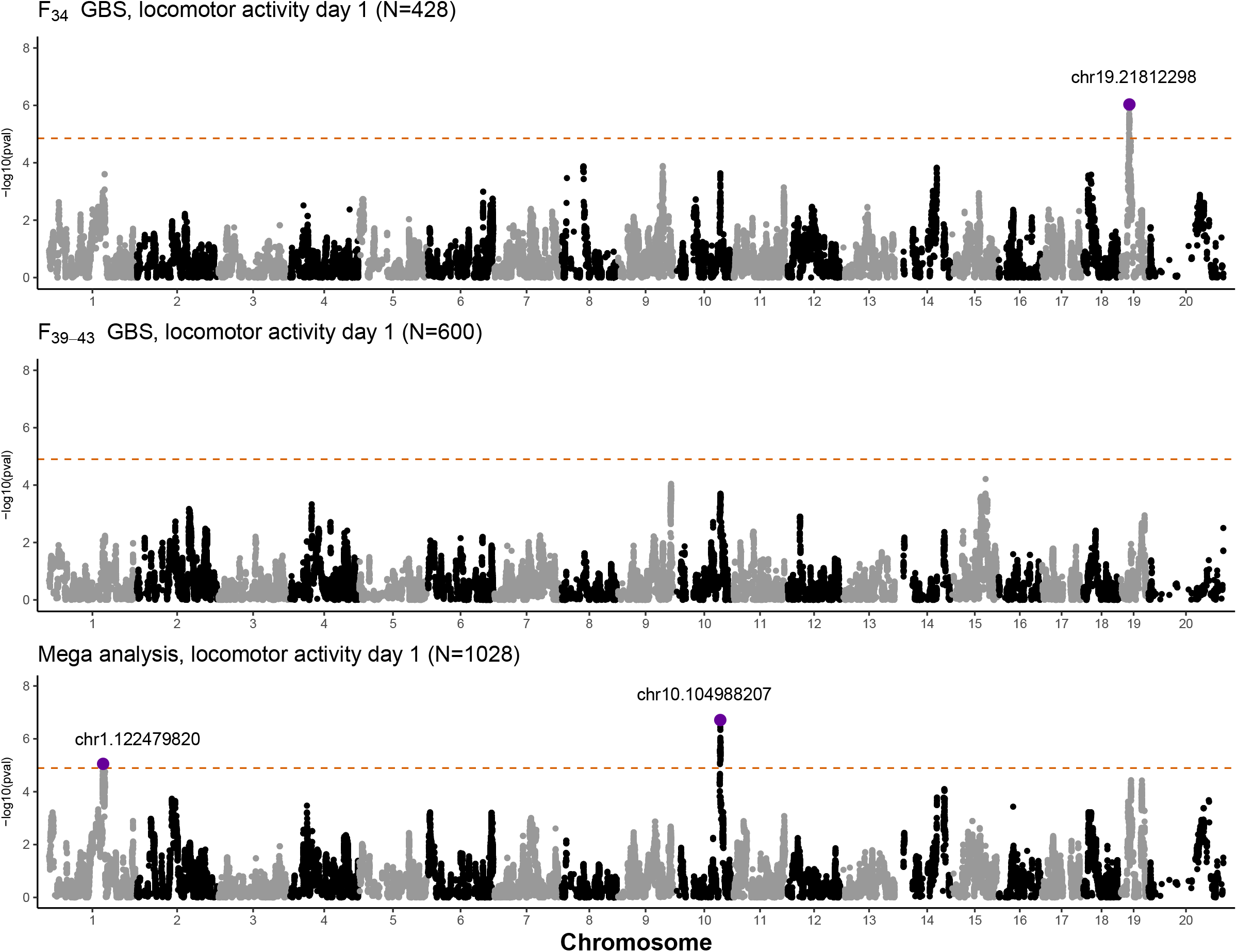
Manhattan plots comparing F_34_ GBS, F_39-43_ GBS, and mega-analysis on locomotor day 1 test using 57,170 shared SNPs in all AIL generations. Mega-analysis identified a locus on chromosome 10 (chr10.104988207) that was not detected in the F_34_ or F_39_-F_43_ alone, suggesting that mega-analysis enhanced power to detect some loci.

## Discussion

We used F_34_ and F_39-43_ generations of a LG/J x SM/J AIL to perform GWAS, SNP heritability estimates, genetic correlations, replication and mega-analysis. We had previously performed several GWAS using a sparse marker set in the F_34_ cohort. In this study we used a denser set of SNPs, obtained using GBS, to reanalyze the F_34_ cohort. We found 110 significant loci, 36 of which had not been identified in our prior studies using the sparse marker set. We used a new, previously unpublished F_39-43_ cohort and showed that genetic correlations were high for the subset of traits that were measured in both cohorts. Despite this, we found that some but not all loci replicated between cohorts, even when we used a relatively liberal definition of replication (p<0.05). The failure to replicate some of our findings was not predicted by our power simulations. Imputation to reference panels increased the number of SNPs available for analysis but did not meaningfully enhance the number of loci we discovered presumably because it did not improve our ability to capture recombination events. Finally, mega-analysis of the two cohorts allowed us to discover 4 additional loci. Taken together, we have identified refined regions of associations for numerous physiological and behavioral traits in multiple generations of AILs.

Previous publications from our lab used a sparse set of array genotypes for GWAS of various behavioral and physiological traits in 688 F_34_ AILs (16,17,35–38,40,41). In this study we obtained a much denser marker set for 428 of the initial 688 AIL mice using GBS. The denser genotypes allowed us to identify most of the loci obtained using the sparse set, as well as many additional loci. For instance, using the sparse markers we identified significant locus on chromosome 8 for locomotor day 2 activity that contained only one gene: *Csmd1* (CUB and sushi multiple domains 1). Gonzales et al. (39) replicated this finding in F_50-56_ AIL and identified a *cis*-eQTL mapped to the same region. *Csmd1* mutant mice showed increased locomotor activity compared to wild-type and heterozygous mice, indicating that *Csmd1* is likely a causal gene for locomotor and related traits (39). We replicated this locus in the analysis of the F_34_ cohort that used the denser marker set (S7 Fig). We also replicated a locus on chromosome 17 for distance traveled in the periphery in the open field test (Fig 4; (36,39)), three loci on chromosomes 4, 6, and 14 for body weight (Supplementary Fig 7; (40)), one locus on chromosome 7 for mean corpuscular hemoglobin concentrations (MCHC, complete blood count; S7 Fig; (41)), and numerous loci on chromosome 4, 6, 7, 8, and 11 for muscle weights (Supplementary Fig 7; (37)). We noticed that even using original spares markers, some previously published loci were not replicated in the current GWAS. The most likely explanation is that we had only 428 of the 688 mice used in the previous publications. Methodological differences between prior studies and the current study, such as the use of QTLRel rather than GEMMA and the choice of pedigree rather than genotypes for estimating relatedness, may also lead to lack of complete replication (56).

F_39-43_ AILs replicated some, but not all, significant loci identified in F_34_, despite generally high genetic correlations between the two cohorts. Significant loci for coat color, which are monogenic and served as positive controls, were consistent between the two cohorts. Loci for body weight were fully replicated (p<0.05) between F_34_ and F_39-43_, while loci for locomotor activity were not. Nevertheless, the beta estimates for all but two loci shared the same sign, which provides modest evidence for replication. Several possibilities may cause the lack of replication for locomotor activity. Firstly, the loci for locomotor activity identified in F_34_ could be false positives. We controlled the genome-wide false positive error rate at 5% using permutation, however, our replication study considered six phenotypes, which was not accounted for by our permutations, thus somewhat increasing the chances that at least one of our significant associations could have been a false positive. Second, unlike the F_34_ dataset, the F_39-43_ used multiple technicians to conduct the behavioral tests and occurred over a prolonged period of time in which numerous environmental factors may have changed. The circumstances under which the F_39-43_ data were collected may have introduced greater environmental heterogeneity, possibly contributing to their lower heritability and the limited replication. The genetic correlations (S6 Table) were high but still less than 1, suggesting a role for heterogeneity.

Finally, our simulation did not account for the systematic overestimation of the effect sizes (“the Winner’s Curse” (57)). The failure to replicate the locomotor activity loci is likely due to a combination of these factors: false positives, heterogeneity, and Winner’s Curse. Thus, while it may seem intuitive that a genome-wide significant result should be reliable at a nominal threshold of p<0.05, when a similarly sized cohort drawn from the same population is used, our results show that this is not the case.

Finally, we performed a mega-analysis using F_34_ and F_39-43_ AIL mice. The combined dataset increased our power and allowed us to identify four novel genome-wide significant associations that were not detected in either the F_34_ or the F_39-43_ cohorts. For example, the mega-analysis identified a locus for body weight on chromosome 2 (Fig S7). Parker et al. (40) identified the same locus using an integrated analysis of LG/J x SM/J F_2_ and F_34_ AILs.

QTL mapping studies have traditionally used a 1.0~2.0 LOD support interval to approximate the size of the association region (see (72,73)). The LOD support interval, proposed by Conneally et al. (74) and Lander & Botstein (75), is a simple confidence interval method involving converting the p-value of the peak locus into a LOD score, subtracting “drop size” from the peak locus LOD score, and finding the two physical positions to the left and to the right of the peak locus location that correspond to the subtracted LOD score. Although Mangin et al. (76) showed via simulation that the boundaries of LOD support intervals depend on effect size, others observed that a 1.0 ~ 2.0 LOD support interval accurately captures ~95% coverage of the true location of the loci when using a dense set of markers (75,77,78). In the present study, we considered using LOD support intervals but found that the sparse array SNPs produced misleadingly large support intervals. Various methods have been proposed for calculating confidence intervals in analogous situations (e.g. (12,79)). We performed credible set analysis and compared LocusZoom plots of the same locus region between array SNPs and the GBS SNPs (S7 Fig; (80)). For example, the benefit of the denser SNP coverage is easily observed in the locus on chromosome 7 (array lead SNP chr7:44560350; GBS lead SNP chr7:44630890) for the complete blood count trait “retic parameters cell hemoglobin concentration mean, repeat” (Fig S7). Thus, there are advantages of dense SNP sets that go beyond the ability to discover additional loci.

LD in the LG/J x SM/J AIL mice is more extensive than in the Diversity Outbred mice and Carworth Farms White mice (1). Some of the loci that we identified are relatively large, making it difficult to infer which genes are responsible for the association. We focused on loci that contained 5 or fewer genes (Table 1). We highlight a few genes that are supported by the existing literature for their role in the corresponding traits. The lead SNP at chr1:77255381 is associated with tibia length in F_34_ AILs (Table 1; S7 Fig). One gene at this locus, *EphA4*, codes for a receptor for membrane-bound ephrins. EphA4 plays an important role in the activation of the tyrosine kinase Jak2 and the signal transducer and transcriptional activator Stat5B in muscle, promoting the synthesis of insulin-like growth factor 1 (IGF-1) (64–66). Mice with mutated EphA4 shows significant defect in body growth (66). Curiously, another gene at this locus, *Pax3*, has been shown as a transcription factor expressed in resident muscle progenitor cells and is essential for the formation of skeletal muscle in mice (67). It is possible that both *EphA4* and *Pax3* are associated with the trait tibia length because they are both involved in organismal growth. Another region of interest is the locus at chr4:66866758, which is associated with body weight in all AIL cohorts (Table 1; S4 Table; S8 Table; S9 Table). The lead SNP is immediately upstream of *Tlr4*, Toll-like receptor 4, which recognizes Gram-negative bacteria by its cell wall component, lipopolysaccharide (68,69). TLR4 responds to the high circulating level of fatty acids and induces inflammatory signaling, which leads to insulin resistance (70). Kim et al showed TLR4-deficient mice were protected from the increase in proinflammatory cytokine level and gained less weight than wild-type mice when fed on high fat diet (71). The association between *Tlr4* and body weight in the AILs corroborates these findings.

Our study has notable limitations. First, not all F_34_ and F_39-43_ animals that were phenotyped were later genotyped by GBS due to missing DNA samples, which in turn lowered our sample size and reduced the power of association analyses. Second, F_39-43_ traits have been collected by different technicians over the span of several years, which introduced environmental heterogeneity and diminished trait heritability (Fig 2). Finally, our power simulations did not account for common factors that can limit replication, including false positives errors, heterogeneity and the Winner’s Curse.

The present study explored replication of GWAS results, the role of marker density, and imputation in GWAS. The combination of denser marker coverage and the addition of 600 F_39-43_ AIL mice allowed us to identify novel loci for locomotor activity, open field test, fear conditioning, light dark test for anxiety, complete blood count, iron content in liver and spleen, and muscle weight. An important conclusion is that power for replication is modest even when a similarly sized replication cohort is used in a genetically identical population tested under conditions that are designed to be as similar as possible.

## Materials and Methods

### Animals

All mice used in this study were members of the advanced intercross line (**AIL**) between LG/J and SM/J that was originally created by Dr. James Cheverud (Loyola University Chicago, Chicago, IL). The AIL line has been maintained in the Palmer laboratory since generation F_33_. Age and exact number of animals tested in each phenotype are described in S1 Table. Several previous publications (16,35–38,40,41) have reported on association analyses of the F_34_ mice (n=428). No prior publications have described the F_39-43_ generations (n=600). The sample size of F_34_ mice reported in this study (n=428) is smaller than that in previous publications of F_34_ (n=688) because we only sequenced a subset of F_34_ animals using GBS. With the exception of coat color, we quantile normalized all phenotypes. Coat color traits were coded in binary numbers (albino: 1 = white, 0 = non-white; agouti: 1 = tan, 0 = black, NA=white). Locomotor activity traits in F_34_ were not quantile transformed in order to follow the guideline described in Cheng et al. (35) for direct comparison.

### F_34_, F_39-43_ Phenotypes

We have previously described the phenotyping of F_34_ animals for locomotor activity (35), fear conditioning (36), open field (36), coat color, body weight (40), complete blood counts (41), heart and tibia measurements (37), muscle weight (37). Iron content in liver and spleen, which have not been previously reported in these mice, was measured by atomic absorption spectrophotometry, as described in Gardenghi et al. (59) and Graziano, Grady and Cerami (60). Although the phenotyping of F_39-43_ animals has not been previously reported, we used method that were identical to those previously reported for locomotor activity (35), open field (36), coat color, body weight (40), and light/dark test for anxiety (15).

### F_34_ AIL Array Genotypes

F_34_ animals had been genotyped on a custom SNP array on the Illumina Infinium platform (35,36), which yielded a set of 4,593 SNPs on autosomes and X chromosome that we refer to as ‘array SNPs’.

### F_34_ and F_39-43_ GBS Genotypes

F_34_ and F_39-43_ animals were genotyped using genotyping-by-sequencing (**GBS**), which is a reduced-representation genome sequencing method (1,39). We used the same protocol for GBS library preparation that was described in Gonzales et al (39). We called GBS genotype probabilities using ANGSD (61). GBS identified 1,667,920 autosomal and 43,015 X-chromosome SNPs. To fill in missing genotypes at SNPs where some but not all mice had calls, we ran within-sample imputation using Beagle v4.1, which generated hard call genotypes as well as genotype probabilities (55). After imputation, only SNPs that had dosage *r*^2^ > 0.9 were retained. We removed SNPs with minor allele frequency < 0.1 and SNPs with p < 1.0×10^−6^ in the Chi-square test of Hardy–Weinberg Equilibrium (**HWE**) (S2 Table). All phenotype and GBS genotype data are deposited in GeneNetwork (http://www.genenetwork.org).

### QC of individuals

We have found that large genetic studies are often hampered by cross-contamination between samples and sample mix-ups. We used four features of the data to identify problematic samples: heterozygosity distribution, proportion of reads aligned to sex chromosomes, pedigree/kinship, and coat color. We first examined heterozygosity across autosomes and removed animals where the proportion of heterozygosity that was more than 3 standard deviations from the mean (S1 Fig). Next, we sought to identify animals in which the recorded sex did not agree with the sequencing data. We compared the ratio of reads mapped to the X and Y chromosomes. The 95% CI for this ratio was 196.84 to 214.3 in females and 2.13 to 2.18 in males. Twenty-two F_34_ and F_39-43_ animals were removed because their sex (as determined by reads ratio) did not agree with their recorded sex; we assumed this discrepancy was due to sample mix-ups. To further identify mislabeled samples, we calculated kinship coefficients based on the full AIL pedigree using QTLRel. We then calculated a genetic relatedness matrix (**GRM**) using IBDLD, which estimates identity by descent using genotype data. The comparison between pedigree kinship relatedness and genetic kinship relatedness identified 7 pairs of animals that showed obvious disagreement between kinship coefficients and the GRM. Lastly, we excluded 14 F_39-43_ animals that showed discordance between their recorded coat color and their genotypes at markers flanking *Tyr*, which causes albinism in mice. The numbers of animals filtered at each step are listed in S2 Table. Some animals were detected by more than one QC step, substantiating our belief that these samples were erroneous.

At the end of SNP and sample filtering, we had 59,561 autosomal and 831 X chromosome SNPs in F_34_, 58,966 autosomal and 824 X chromosome SNPs in F_39-43_, and 57,635 autosomal and 826 X chromosome SNPs in the combined F_34_ and F_39-43_ set (S2 Table). GBS genotype quality was estimated by examining concordance between the 66 SNPs that were present in both the array and GBS genotyping results.

### LD decay

Average LD (*r*^2^) was calculated using allele frequency matched SNPs (MAF difference < 0.05) within 100,000bp distance, as described in Parker et al. (1).

### Imputation to LG/J and SM/J reference panels

F_34_ array genotypes (n=428) and F_34_ GBS genotypes (n=428) were imputed to LG/J and SM/J whole genome sequence data (54) using BEAGLE. For F_34_ array imputation, we used a large window size (100,000 SNPs and 45,000 SNPs overlap). Imputation to reference panels yielded 4.3 million SNPs for F_34_ array and F_34_ GBS imputed sets. Imputed SNPs with DR^2^ > 0.9, MAF > 0.1, HWE p value > 1.0×10^−6^ were retained, resulting in 4.1M imputed F34 GBS SNPs and 4.3M imputed F_34_ array SNPs.

### Genome-wide association analysis (GWAS)

We used the linear mixed model, as implemented in GEMMA (46), to perform a GWAS that accounted for the complex familial relationships among the AIL mice (35,39). We used the leave-one-chromosome-out (**LOCO**) approach to calculate genetic relatedness matrix, which effectively circumvented the problem of proximal contamination (48). Separate GWAS were performed using F_34_ array genotypes, F_34_ GBS genotypes, and F_39-43_ GBS genotypes. Apart from coat color (binary trait) and locomotor activity, raw phenotypes were quantile normalized prior to analysis. Locomotor activity was not quantile normalized because the trait was reasonably normally distributed already and because we wanted our analysis to match those performed in Cheng et al (35). Because F_34_ AIL had already been studied using array genotypes (35) and mapped using QTLRel (62), we used the same covariates as described in Cheng et al. (35) in order to examine whether our array and GBS GWAS would replicate their findings. We included sex and body weight as covariates for locomotor activity traits (see covariates used in (35))and sex, age, and coat color as covariates for fear conditioning and open field test in F_34_ AILs (see covariates used in (36)). We used sex and age as covariates for all other phenotypes. Covariates for each analysis are shown in S1 Table. Finally, we performed mega-analysis of F_34_ and F_39-43_ animals (n=1,028) for body weight, coat color, and locomotor activity, since these traits were measured in the same way in both cohorts. For the mega-analyses, locomotor activity was quantile normalized after the combination of the two datasets to ensure that data were normally distributed across generations.

### Identifying suspicious SNPs

Some significant SNPs in F_34_ GWAS and in the mega-analysis of F_34_ and F_39-43_ were suspicious because nearby SNPs, which would have been expected to be in high LD (a very strong assumption in an AIL), did not have high −log10 values. We only examined SNPs that obtained significant p-values; these examinations reveled that these SNPs had suspicious ratios of heterozygotes to homozygotes calls and had corresponding HWE p-values that were close to our 1.0×10^−6^ threshold (S10 and S11 Tables). To avoid counting these as novel loci, we removed those SNPs prior to summarizing our results as they likely reflected genotyping errors.

### Selecting independent significant SNPs

To identify independent “lead loci” among significant GWAS SNPs that surpass the significance threshold, we used the LD-based clumping method in PLINK v1.9. We empirically chose clumping parameters (*r*^2^ = 0.1 and sliding window size = 12,150kb) that gave us a conservative set of independent SNPs (S3 Table). For the coat color phenotypes, we found that multiple SNPs remained significant even after LD-based clumping, presumably due to the extremely significant associations at these Mendelian loci. In these cases, we used a stepwise model selection procedure in GCTA (58) and performed association analyses conditioning on the most significant SNPs.

### Significance thresholds

We used MultiTrans and SLIDE to set significance thresholds for the GWAS (50,51). MultiTrans and SLIDE are methods that assume multivariate normal distribution of the phenotypes, which in LMM models, contain a covariance structure due to various degrees of relatedness among individuals. We were curious to see whether MultiTrans/SLIDE produces significance thresholds drastically different from the threshold we obtained from a standard permutation test (‘naïve permutation’ as per Cheng et al. (48)). We performed 1,000 permutations using the F_34_ GBS genotypes and the phenotypic data from locomotor activity (days 1, 2, and 3). We found that the 95^th^ percentile values for these permutations were 4.65, 4.79, and 4.85, respectively, which were very similar to 4.85, the threshold obtained from MultiTrans using the same data. Thus, the thresholds presented here were obtained from MultiTrans but are similar (if anything slightly more conservative) than thresholds we would have obtained had we used permutation. Because the effective number of tests depends on the number of SNPs and the specific animals used in GWAS, we obtained a unique adjusted significance threshold for each SNP set in each animal cohort (S12 Table).

### Power analysis

To estimate the power of replication of a SNP from the discovery set in the replication set, we simulated GWAS with 50 varying effect sizes for the discovery SNP using the LMM model. We first fit the trait in a null model (i.e., no genotype effect), and obtained estimates of model parameters including the intercept and the genetic variance component. Using these model parameters, we added the genotype effect to the random numbers generated from the null model to recreate a trait. For each simulated effect size, we scanned every simulated trait 2,500 times and examined the ratio of association tests whose test statistics surpassed the significance thresholds (both the genome-wide significance threshold for the cohort and the nominal P value of 0.05). In order to compare effect sizes measured in phenotypes with different scales and units, we converted beta estimates (of discovery SNPs and simulated effect sizes) to z scores based on the standard errors of the beta estimates of discovery SNPs in the simulation cohort.

### Credible set analysis

We followed the method described in (53). The R script could be found on GitHub: https://github.com/hailianghuang/FM-summary/blob/master/getCredible.r

### Genetic correlation and heritability estimates between F_34_ and F_39-43_ phenotypes

Locomotor activity, body weight, and coat color had been measured in both F_34_ and F_39-43_ populations. We calculated both SNP heritability and genetic correlations between F_34_ and F_39-43_ animals using GCTA bivariate GREML analysis (58). Because F_39-43_ day 1 locomotor activity data were not normally distributed, we quantile normalized locomotor activity data when estimating SNP heritabilities and genetic correlations.

### LocusZoom Plots

LocusZoom plots were generated using the standalone implementation of LocusZoom (63), using LD scores calculated from PLINK v.1.9 --ld option and mm10 gene annotation file downloaded from USCS genome browser.

## Ethics Statement

All procedures were approved by the Institutional Animal Care and Use Committee (IACUC protocol: S15226) Euthanasia was accomplished using CO_2_ asphyxiation followed by cervical dislocation.

## Acknowledgements

We would like to recognize Jackie Lim and Kaitlin Samocha for collecting F_34_ AIL phenotype data and Ryan Walters for collecting F_39-43_ AIL phenotype data. We wish to acknowledge Alex Gileta for input on a draft of this manuscript.

## Supporting information

**S1 Fig. Autosomal heterozygosity distribution in F_34_, F_39-43_ AILs.** Animals with excessive or insufficient heterozygosity (3 *s.d.* away from mean) were removed from further analysis. As controls, we have sequenced two F_2_s of LG and SM, four LG mice and four SM mice (see annotated data points with 1 and 0 heterozygosity).

**S2 Fig. Kinship coefficients in F_34_ and F_39-43_ AILs calculated from pedigree against genetic relatedness matrix calculated using IBDLD [49].** Each circle represents a pair of animals, which their genetic kinship relatedness on the x-axis and pedigree kinship relatedness on the y-axis. Color signifies relatedness based on AIIL pedigree. Blue circles represent identical twins, red full siblings, yellow parent-offspring pairs, grey other relationships. Seven animal pairs that deviate from the pedigree relationship clusters were excluded (see black arrows).

**S3 Fig. Heatmap showing F_34_ array and F_34_ GBS genotype concordance in percentages, using 66 shared SNPs.** “A” codes for the LG/J allele, and “B” codes for the SM/J allele. “AA” genotype concordance between array and GBS is 24.54%, “AB” 43.23%, “BB” 27.60%.

**S4 Fig. LD decay in F_34_ array, F_34_ GBS, F_39-43_ GBS, and F_34_ and F_39-43_ GBS SNP sets.**

**S5 Fig. Manhattan plots comparing 4,593 F_34_ array, 60.3K F_34_ GBS, 4.3M imputed F_34_ array, and 4.1M imputed F_34_ GBS (N=428) SNPs on day 2 locomotor activity.** Adjusted significance thresholds for imputed array and GBS SNPs were estimated using LD pruned SNPs (r2=0.1, window size=20kb; PLINK v1.9). Notice that even though the imputed sets have more SNPs (the two right panels), they are frequently blocks of many SNPs with almost identical position and LD=1, therefore making it hard to visualize the additional SNPs.

**S6 Fig. SNP heritability using F_34_ GBS and F_34_ array SNPs (slope=1).**

**S7 Fig. LocusZoom for F_34_ array, F_34_ GBS, F_39-43_ GBS, and mega-analysis QTLs.** Purple track shows the credible set interval (*r*^2^ threshold = 0.8, posterior probability threshold = 0.99).

**S8 Fig. Power simulations for discovery SNPs in the replication set.** Power was simulated at both the genome-wide significance level for the cohort and the nominal P value of 0.05. Each data point represents the estimated power at the simulated beta (plotted as Z score). The vertical dashed line in orange indicates the effect size of the discovery SNP.

**S1 Table. List of phenotypes used in GWAS.**

**S2 Table. SNP and individual QC filter table.** Numbers of animals and SNPs remained after each step of filtering are shown per GBS SNP set.

**S3 Table. Effect of PLINK v1.9 clump-based pruning parameters on number of independent SNPs remained.** At all *r*_2_ values examined, a sliding window size of 12150kb was the first smallest window that yield the most stringent number of clumped SNPs in both array and GBS GWAS.

**S4 Table. Lead QTL in F_34_ GBS and F_34_ array GWAS studies across phenotypes.** Significant SNPs are clumped using parameters *r*_2_=0.1, 12150kb.

**S5 Table. F_34_ GBS and array SNP heritability estimates.**

**S6 Table. F_34_ and F_39-43_ genetic correlations in locomotor activity, coat color, and body weight.**

**S7 Table. SNP-heritability comparison between F_34_ and F_39-43_ GBS.**

**S8 Table. Lead QTL in F_39-43_ N=600 GBS GWAS studies across phenotypes.** Significant SNPs are clumped using parameters *r*_2_=0.1, 12150kb.

**S9 Table. Lead QTL in F_34_ and F_39-43_ (N=1028) mega-analysis across phenotypes.** Significant SNPs are clumped using parameters *r*_2_=0.1, 12150kb.

**S10 Table. SNPs in F_34_ GBS set with HWE p-values close to 1.0×10-6 cutoff threshold.** These SNPs are removed from QTL summary tables.

**S11 Table. SNPs in F_34_ and F_39-43_ mega-analysis GBS set with HWE p values close to 1.0×10-6 cutoff threshold.** These SNPs are removed from QTL summary tables.

**S12 Table. Adjusted significance threshold for each SNP set and GWAS cohort.**

