## Supplementary figures and images for "Genome-wide association study in two cohorts from a multi-generational mouse advanced intercross line highlights the difficulty of replication"

### S1 Fig

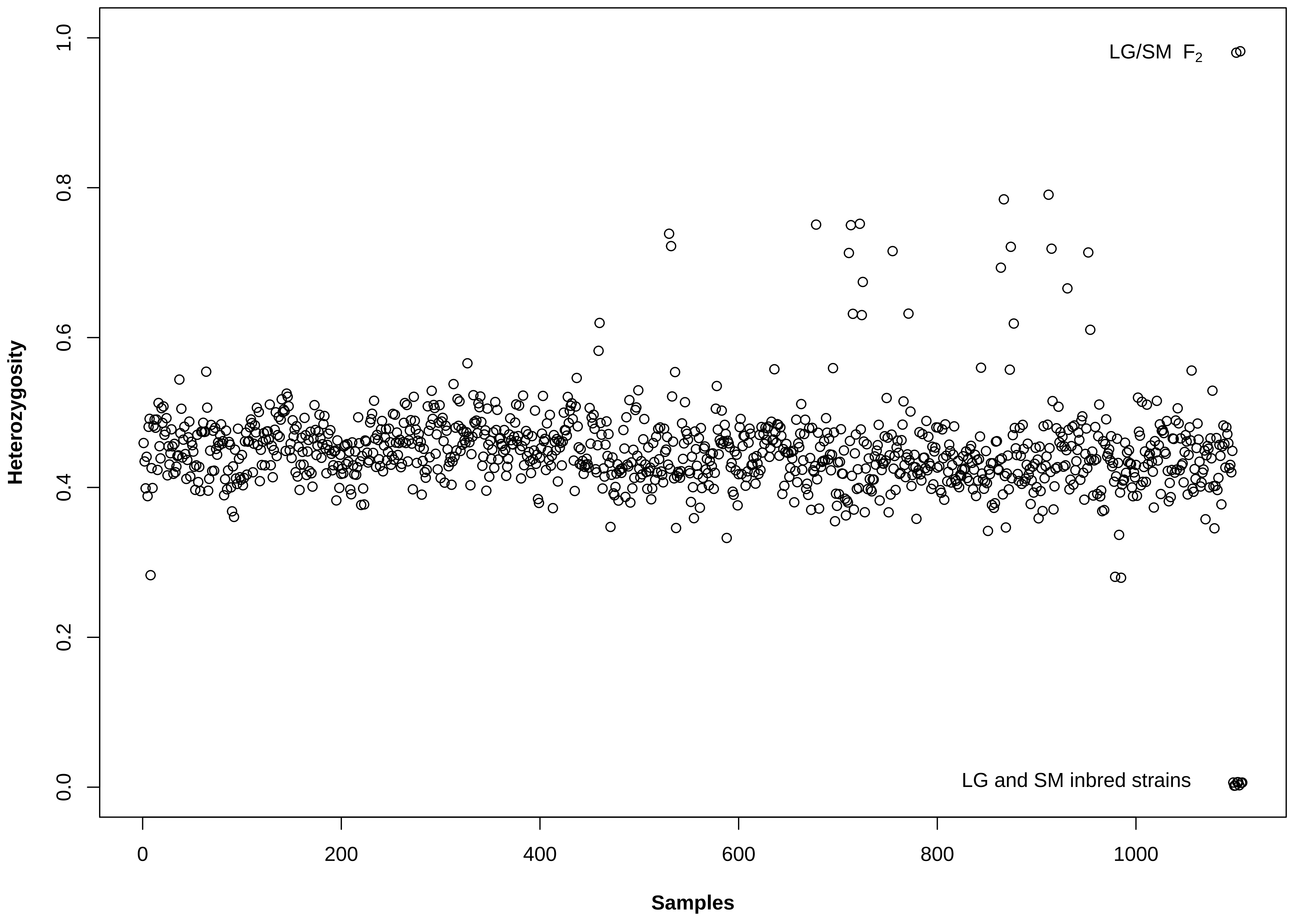

### S2 Fig

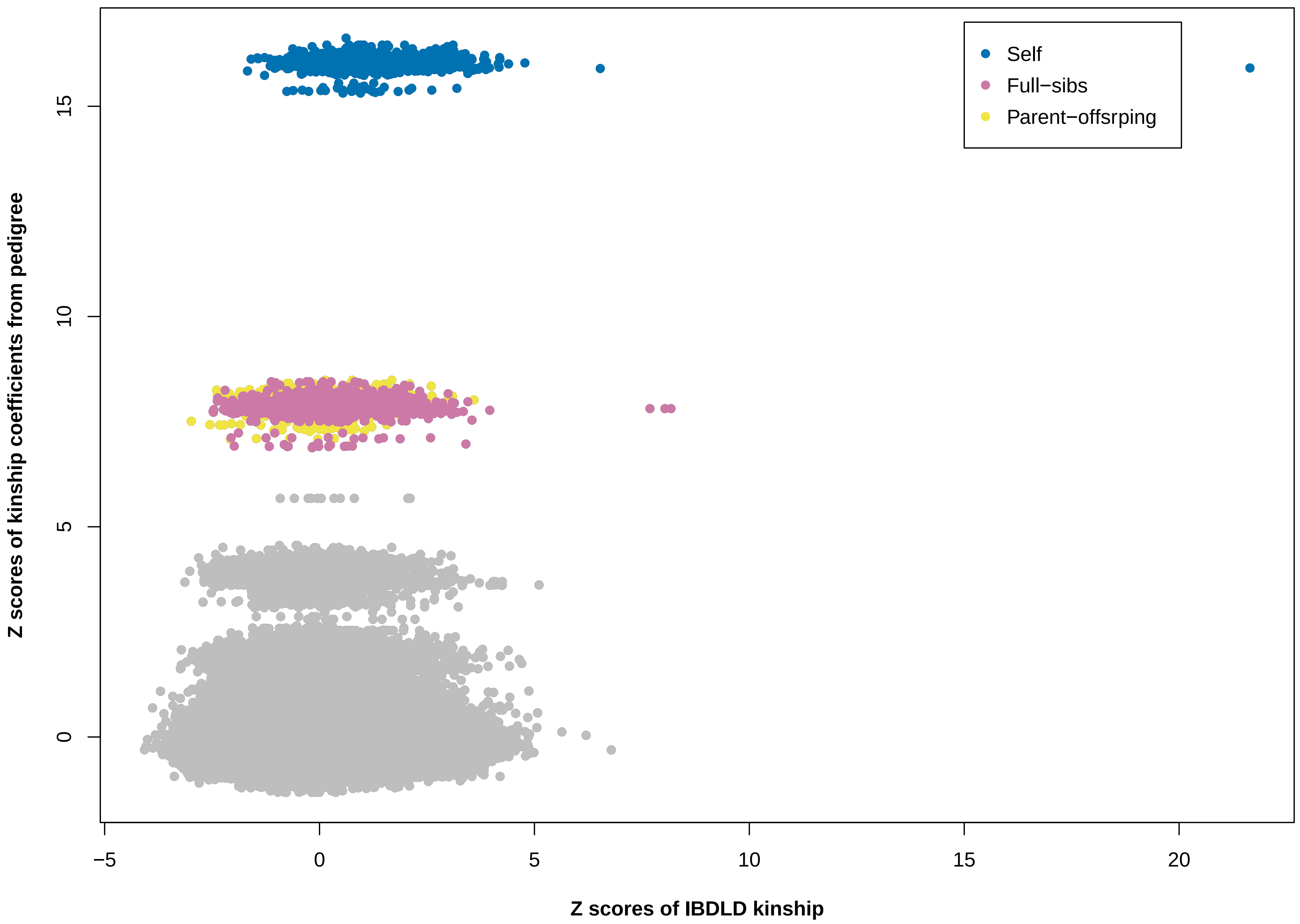

### S3 Fig

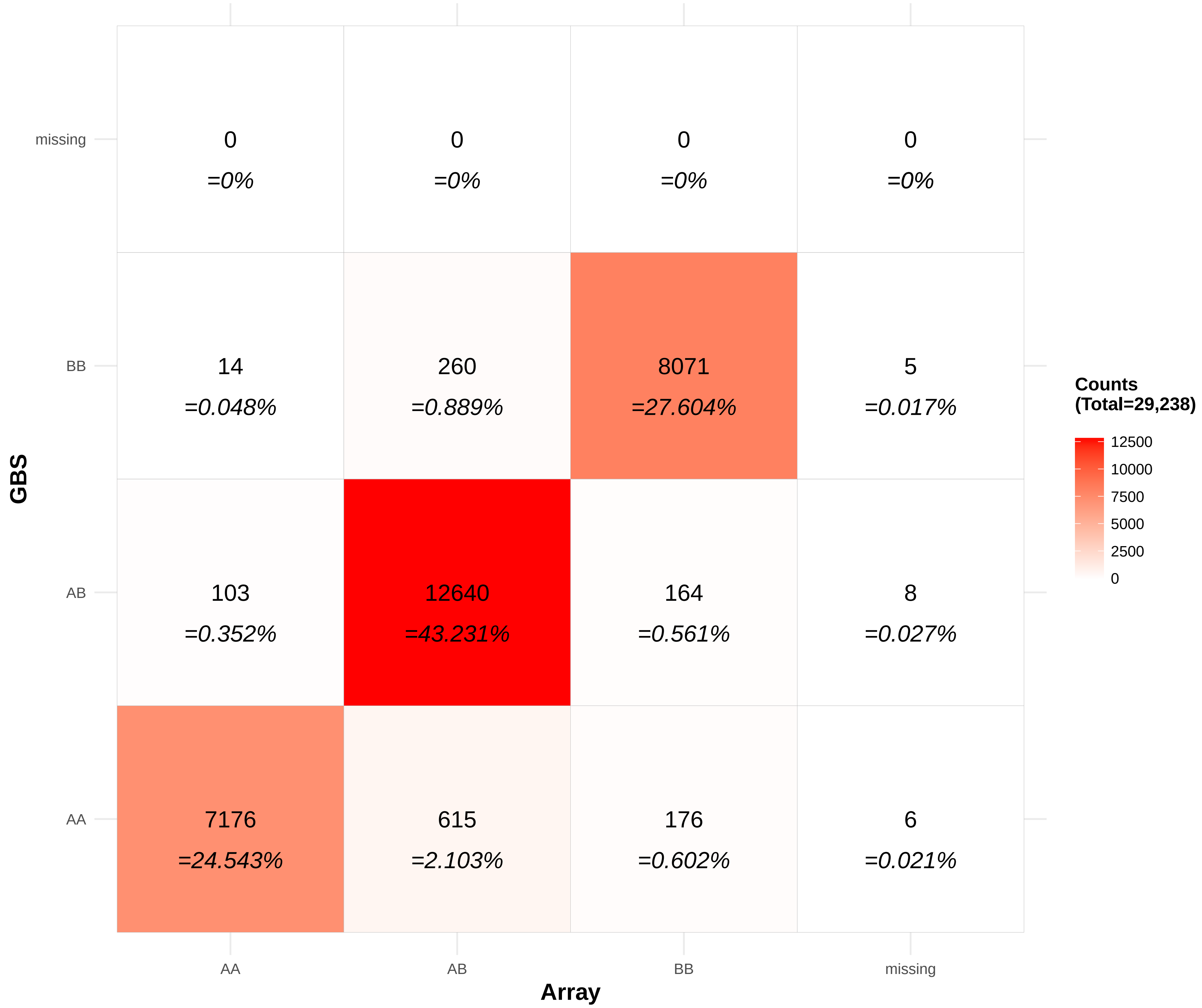

### S4 Fig

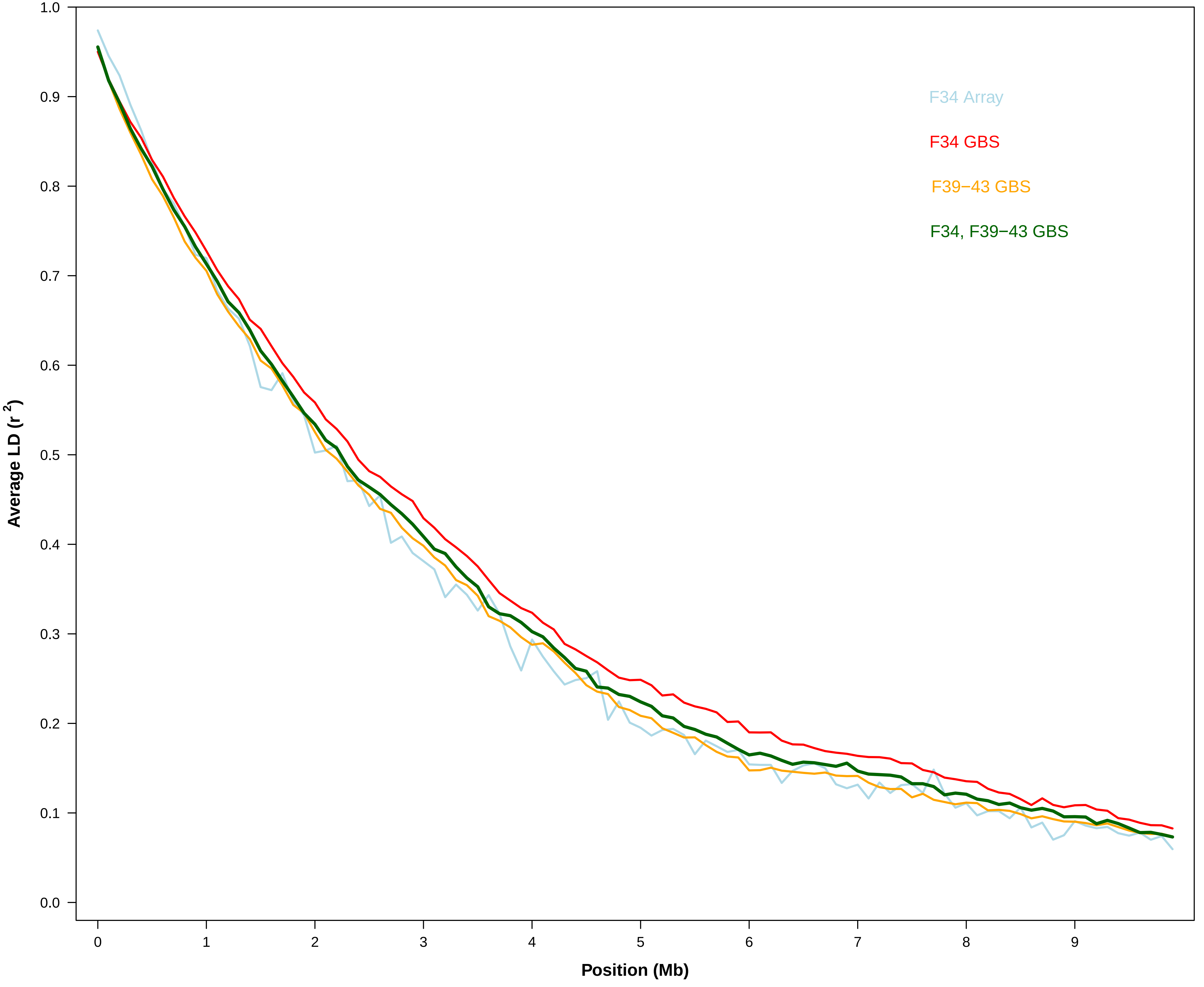

### S5 Fig

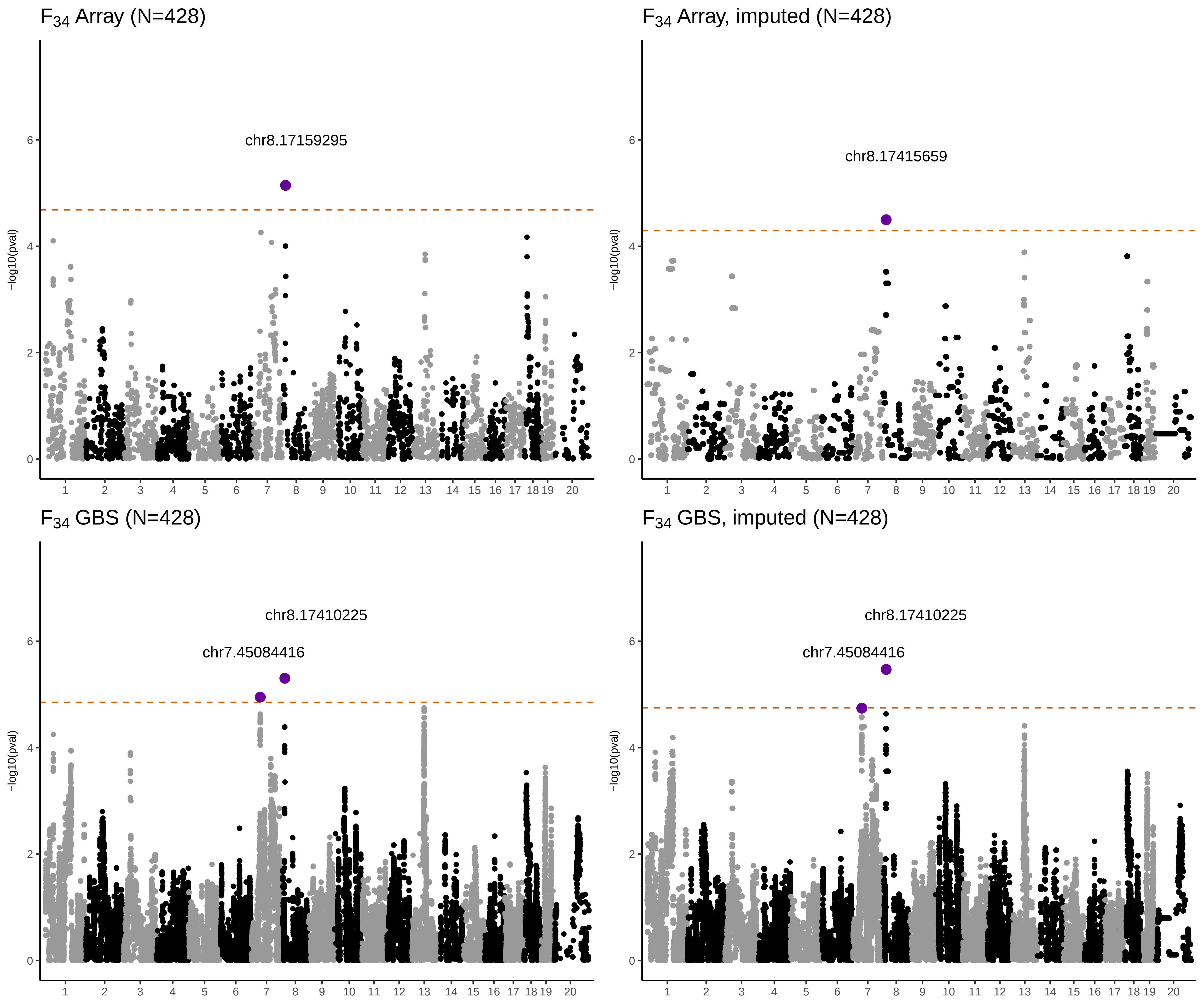

### S6 Fig

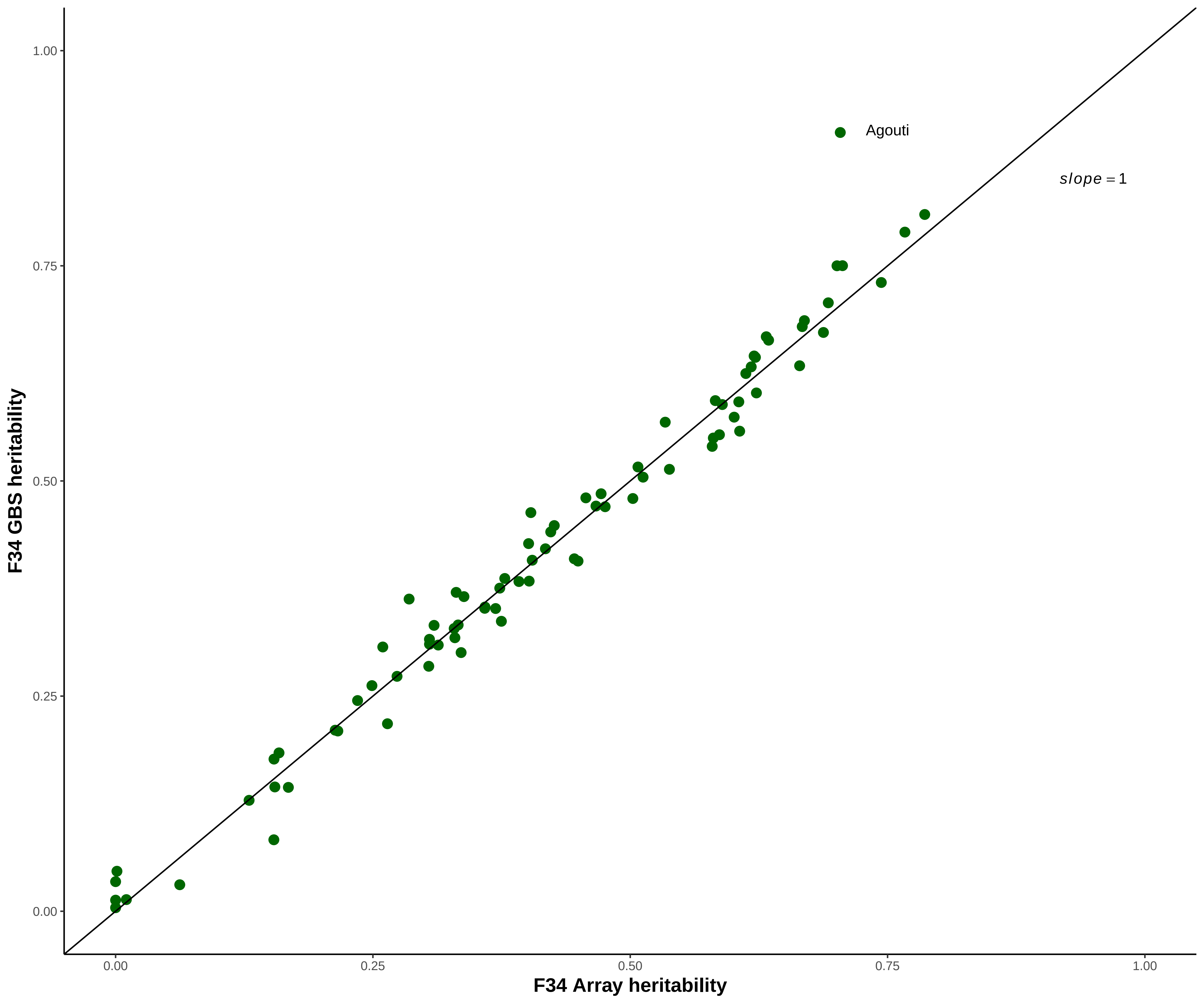
