## Supplementary material for "Genome-wide association study in two cohorts from a multi-generational mouse advanced intercross line highlights the difficulty of replication": S7 Fig

### F34 Array, body weight day 1

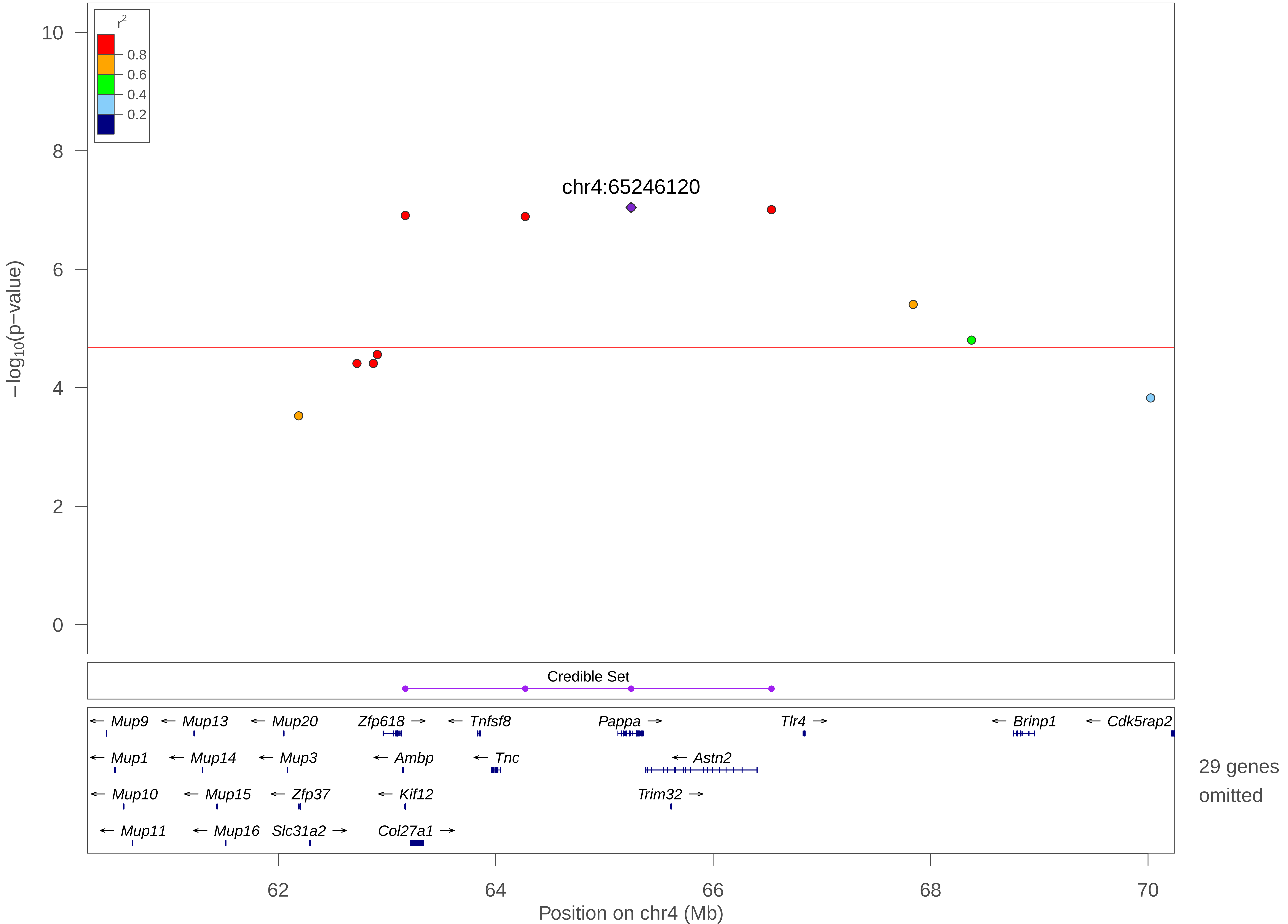

| | |
|---|---|
| date: | Thu Oct 25 16:44:34 2018 |
| build: | F34Array |
| display range: | chr4:60246120−70246120 [60246120−70246120] |
| hilite range: | 0 − 0 [ 0 − 0 ] |
| reference SNP: | chr4:65246120 |
| number of SNPs plotted: | 12 |
| min P−value: | 9.06E−8 [chr4:65246120] |
| max P−value: | 2.99E−4 [chr4:62188670] |
| omitted Genes: | Mup12, Mup17, Mup18 |
| omitted Genes: | Mup19, Mup5, Mup21 |
| omitted Genes: | Fkbp15, Slc31a1, Cdc26 |
| omitted Genes: | Prpf4, Rnf183, Wdr31 |
| omitted Genes: | Bspry, Hdhd3, Alad |
| omitted Genes: | Pole3, 4933430I17Rik, Rgs3 |
| omitted Genes: | Mir455, Orm1, Orm3 |
| omitted Genes: | Orm2, Akna, Whrn |
| omitted Genes: | Atp6v1g1, 6330416G13Rik, Gm11213 |
| omitted Genes: | 1700018C11Rik, Tnfsf15 |

Make more plots at http://csg.sph.umich.edu/locuszoom/

Fine−mapping Regions

| snp | chr | pos (Mb) | pp | group | color |
|---|---|---|---|---|---|
| chr4:63169190 | 4 | 63.16919 | 0.2189622 | Credible Set | purple |
| chr4:64272026 | 4 | 64.27203 | 0.2102491 | Credible Set | purple |
| chr4:65246120 | 4 | 65.24612 | 0.2955478 | Credible Set | purple |
| chr4:66536974 | 4 | 66.53697 | 0.2722694 | Credible Set | purple |

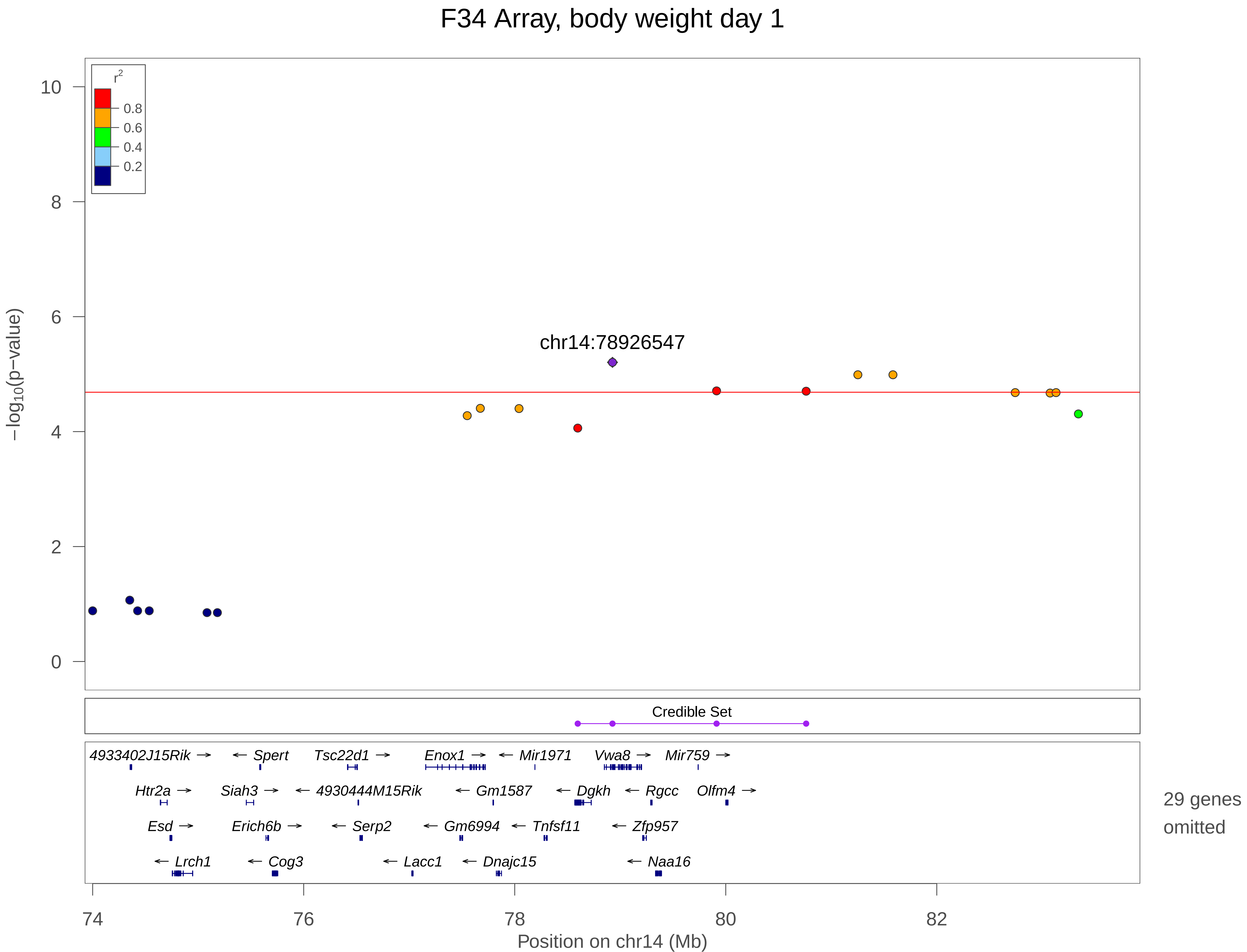

F34 Array, body weight day 1
r2
0.8
0.6
0.4
0.2
chr14:78926547
−log10(p−value)
10
8
6
4
2
0
Credible Set
4933402J15Rik →
← Spert
Tsc22d1 →
Enox1 →
← Mir1971
Vwa8 →
Mir759 →
Htr2a →
Siah3 →
← 4930444M15Rik
← Gm1587
← Dgkh
← Rgcc
Olfm4 →
Esd →
Erich6b →
← Serp2
← Gm6994
← Tnfsf11
← Zfp957
← Lrch1
← Cog3
← Lacc1
← Dnajc15
← Naa16
29 genes omitted
74
76
78
80
82
Position on chr14 (Mb)

| | |
|---|---|
| date: | Thu Oct 25 16:44:27 2018 |
| build: | F34Array |
| display range: | chr14:73926547−83926547 [73926547−83926547] |
| hilite range: | 0 − 0 [ 0 − 0 ] |
| reference SNP: | chr14:78926547 |
| number of SNPs plotted: | 20 |
| min P−value: | 6.24E−6 [chr14:78926547] |
| max P−value: | 1.41E−1 [chr14:75083773] |
| omitted Genes: | Gm4278, 5031414D18Rik, Lrrc63 |
| omitted Genes: | Lcp1, Cpb2, Zc3h13 |
| omitted Genes: | Slc25a30, Gm4285, Tpt1 |
| omitted Genes: | Snora31, Gtf2f2, Kctd4 |
| omitted Genes: | Gpalpp1, Nufip1, 1700108F19Rik |
| omitted Genes: | Ccdc122, Epsti1, Fam216b |
| omitted Genes: | Akap11, Mtrf1, Kbtbd7 |
| omitted Genes: | Kbtbd6, Wbp4, Elf1 |
| omitted Genes: | Sugt1, Lect1, 4930452G13Rik |
| omitted Genes: | Pcdh8, Gm10845 |

Make more plots at http://csg.sph.umich.edu/locuszoom/

Fine−mapping Regions

| snp | chr | pos (Mb) | pp | group | color |
|---|---|---|---|---|---|
| chr14:78597230 | 14 | 78.59723 | 0.04671101 | Credible Set | purple |
| chr14:78926547 | 14 | 78.92655 | 0.57169961 | Credible Set | purple |
| chr14:79913027 | 14 | 79.91303 | 0.19194230 | Credible Set | purple |
| chr14:80762011 | 14 | 80.76201 | 0.18964708 | Credible Set | purple |

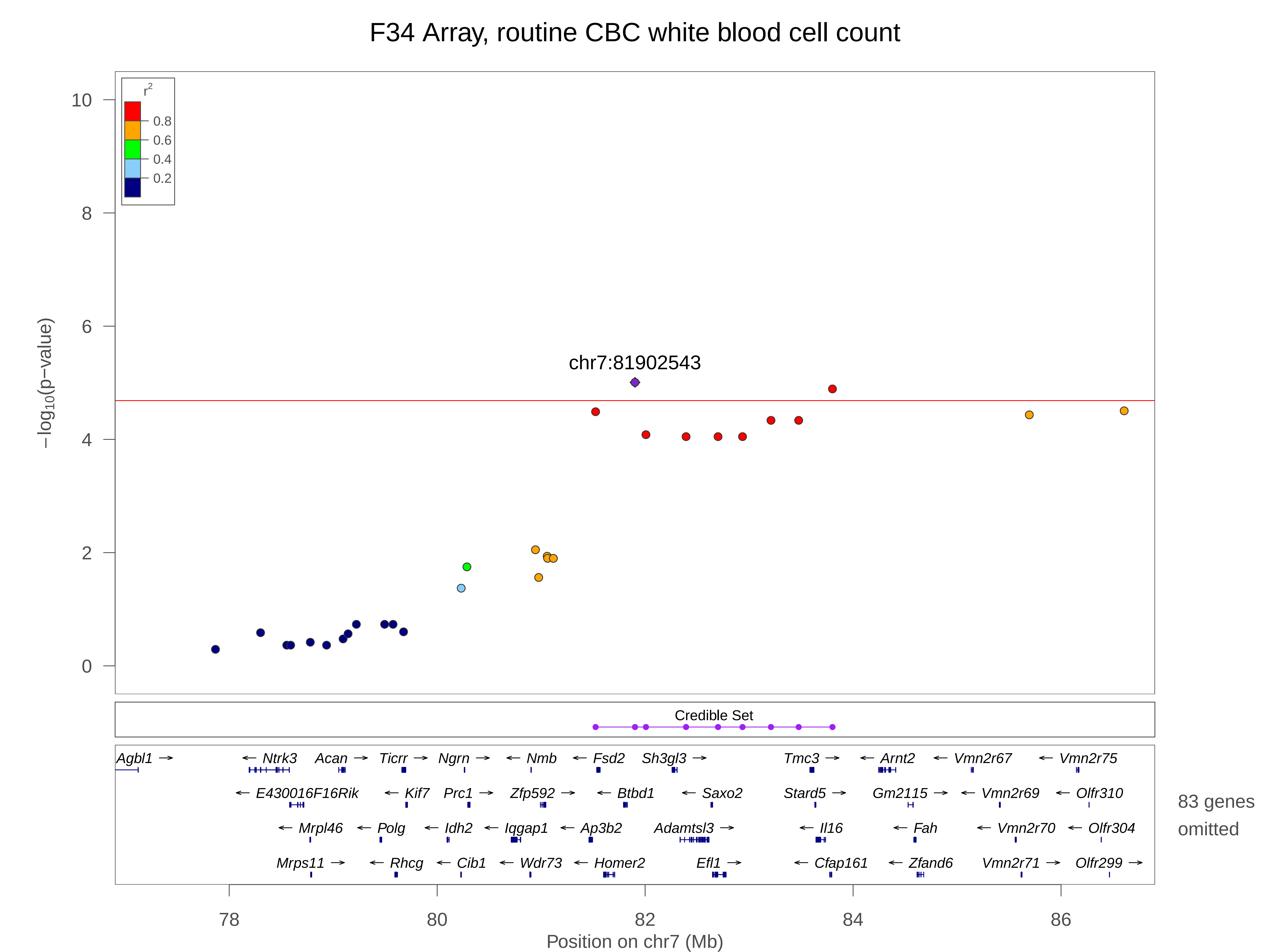

F34 Array, routine CBC white blood cell count
r2
0.8
0.6
0.4
0.2
10
8
6
4
2
0
−log10(p−value)
chr7:81902543
Credible Set
Agbl1 →
← Ntrk3
Acan →
Ticrr →
Ngrn →
← Nmb
← Fsd2
Sh3gl3 →
Tmc3 →
← Arnt2
← Vmn2r67
← Vmn2r75
← E430016F16Rik
← Kif7
Prc1 →
Zfp592 →
← Btbd1
← Saxo2
Stard5 →
Gm2115 →
← Vmn2r69
← Olfr310
← Mrpl46
← Polg
← Idh2
← Iqgap1
← Ap3b2
Adamtsl3 →
← Il16
← Fah
← Vmn2r70
← Olfr304
Mrps11 →
← Rhcg
← Cib1
← Wdr73
← Homer2
Efl1 →
← Cfap161
← Zfand6
Vmn2r71 →
Olfr299 →
83 genes omitted
78
80
82
84
86
Position on chr7 (Mb)

| | |
|---|---|
| date: | Thu Oct 25 16:44:49 2018 |
| build: | F34Array |
| display range: | chr7:76902543−86902543 [76902543−86902543] |
| hilite range: | 0 − 0 [ 0 − 0 ] |
| reference SNP: | chr7:81902543 |
| number of SNPs plotted: | 31 |
| min P−value: | 9.83E−6 [chr7:81902543] |
| max P−value: | 5.11E−1 [chr7:77868097] |
| omitted Genes: | Det1, Mir7−2, Aen |
| omitted Genes: | Isg20, Gm26633, Hapln3 |
| omitted Genes: | Mfge8, Abhd2, Rlbp1 |
| omitted Genes: | Fanci, Mir9−3hg, Mir9−3 |
| omitted Genes: | Plin1, Pex11a, Wdr93 |
| omitted Genes: | Mesp1, Mesp2, Anpep |
| omitted Genes: | Ap3s2, Arpin, Zfp710 |
| omitted Genes: | Gm21057, Sema4b, Gdpgp1 |
| omitted Genes: | Ttll13, Vps33b, Rccd1 |
| omitted Genes: | Unc45a, Hddc3, Man2a2 |
| omitted Genes: | Fes, Furin, Blm |
| omitted Genes: | Crtc3, Gm15880, Zscan2 |
| omitted Genes: | Sec11a, Alpk3, Slc28a1 |
| omitted Genes: | Pde8a, Rps17, Cpeb1 |
| omitted Genes: | BC048679, 2900076A07Rik, Mir1839 |
| omitted Genes: | Whamm, Fam103a1, 3110040N11Rik |
| omitted Genes: | Tm6sf1, Hdgfrp3, Bnc1 |
| omitted Genes: | Gm20744, 4933406J10Rik, Mex3b |
| omitted Genes: | 4933430H16Rik, Mesdc1, Mesdc2 |
| omitted Genes: | Cemip, Abhd17c, 2610206C17Rik |
| omitted Genes: | Olfr291, Olfr290, Vmn2r65 |
| omitted Genes: | Vmn2r66, Vmn2r68, Vmn2r72 |
| omitted Genes: | Vmn2r73, Vmn2r74, Vmn2r76 |
| omitted Genes: | Olfr309, Olfr308, Olfr307 |
| omitted Genes: | Olfr305, Olfr303, Olfr301 |
| omitted Genes: | Olfr298, Olfr297, Olfr295 |
| omitted Genes: | Olfr294, Olfr293, Olfr292 |
| omitted Genes: | Folh1, Vmn2r77 |

Make more plots at http://csg.sph.umich.edu/locuszoom/

Fine−mapping Regions

| snp | chr | pos (Mb) | pp | group | color |
|---|---|---|---|---|---|
| chr7:81523675 | 7 | 81.52367 | 0.10491650 | Credible Set | purple |
| chr7:81902543 | 7 | 81.90254 | 0.32716949 | Credible Set | purple |
| chr7:82007699 | 7 | 82.00770 | 0.04331532 | Credible Set | purple |
| chr7:82393693 | 7 | 82.39369 | 0.04018994 | Credible Set | purple |
| chr7:82701107 | 7 | 82.70111 | 0.04018994 | Credible Set | purple |
| chr7:82936666 | 7 | 82.93667 | 0.04018994 | Credible Set | purple |
| chr7:83210268 | 7 | 83.21027 | 0.07521401 | Credible Set | purple |
| chr7:83476529 | 7 | 83.47653 | 0.07521401 | Credible Set | purple |
| chr7:83801589 | 7 | 83.80159 | 0.25360084 | Credible Set | purple |

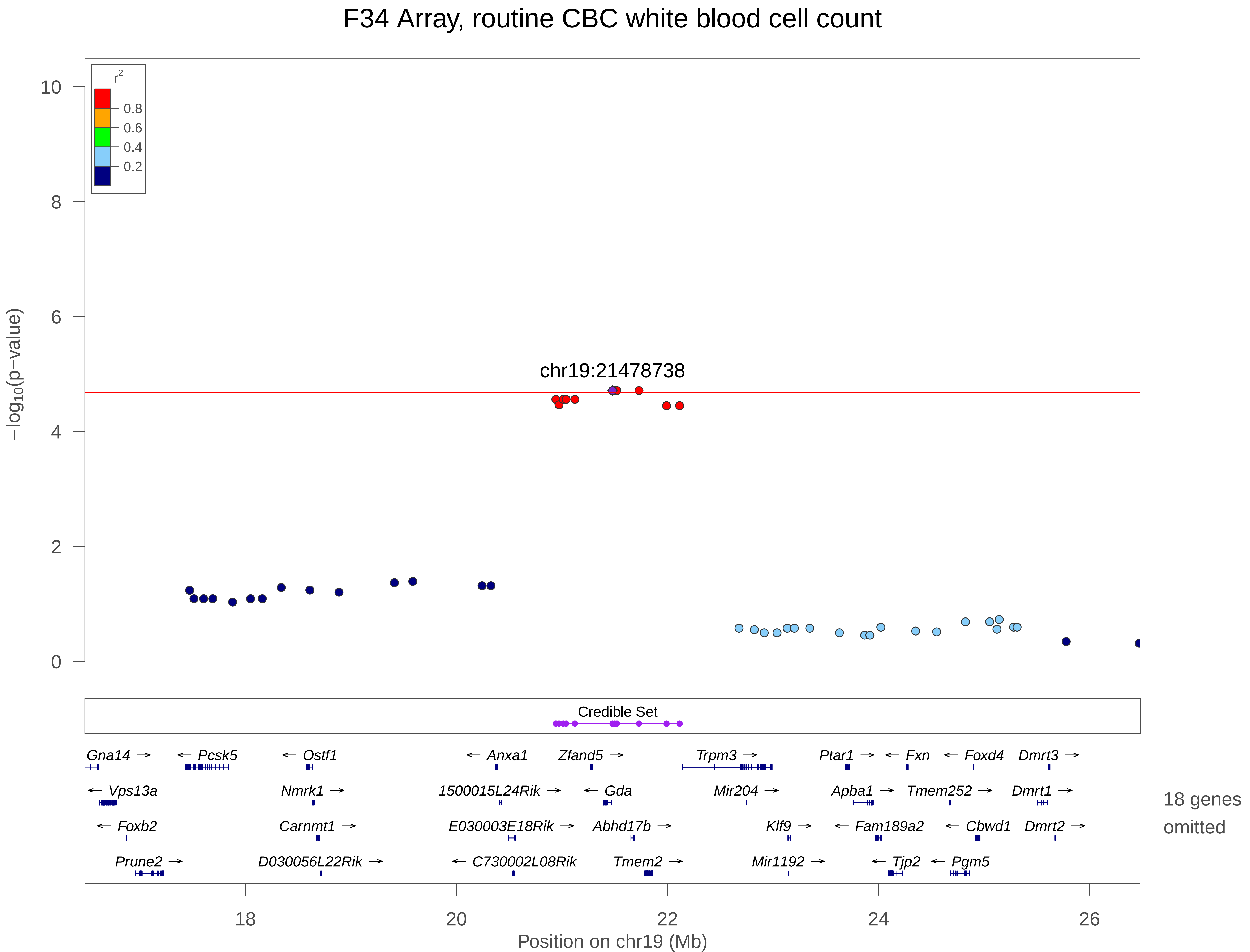
F34 Array, routine CBC white blood cell count
r2
0.8
0.6
0.4
0.2
−log10(p−value)
chr19:21478738
Credible Set
Gna14
Pcsk5
Ostf1
Anxa1
Zfand5
Trpm3
Ptar1
Fxn
Foxd4
Dmrt3
Vps13a
Nmrk1
1500015L24Rik
Gda
Mir204
Apba1
Tmem252
Dmrt1
Foxb2
Carnmt1
E030003E18Rik
Abhd17b
Klf9
Fam189a2
Cbwd1
Dmrt2
Prune2
D030056L22Rik
C730002L08Rik
Tmem2
Mir1192
Tjp2
Pgm5
18 genes omitted
Position on chr19 (Mb)

| | |
|---|---|
| date: | Thu Oct 25 16:44:41 2018 |
| build: | F34Array |
| display range: | chr19:16478738−26478738 [16478738−26478738] |
| hilite range: | 0 − 0 [ 0 − 0 ] |
| reference SNP: | chr19:21478738 |
| number of SNPs plotted: | 47 |
| min P−value: | 1.93E−5 [chr19:21478738] |
| max P−value: | 4.8E−1 [chr19:26470094] |
| omitted Genes: | Gcnt1, Rfk, Trpm6 |
| omitted Genes: | Rorb, Aldh1a1, Aldh1a7 |
| omitted Genes: | Tmc1, 4930554I06Rik, 1110059E24Rik |
| omitted Genes: | Smc5, Mamdc2, 1700028P14Rik |
| omitted Genes: | Pip5k1b, Fam122a, Mir3084−1 |
| omitted Genes: | Dock8, Kank1, 2610016A17Rik |

Make more plots at http://csg.sph.umich.edu/locuszoom/

Fine−mapping Regions

| snp | chr | pos (Mb) | pp | group | color |
|---|---|---|---|---|---|
| chr19:20941927 | 19 | 20.94193 | 0.08362408 | Credible Set | purple |
| chr19:20971640 | 19 | 20.97164 | 0.06780032 | Credible Set | purple |
| chr19:21011731 | 19 | 21.01173 | 0.08362408 | Credible Set | purple |
| chr19:21038918 | 19 | 21.03892 | 0.08362408 | Credible Set | purple |
| chr19:21122136 | 19 | 21.12214 | 0.08362408 | Credible Set | purple |
| chr19:21478738 | 19 | 21.47874 | 0.11662427 | Credible Set | purple |
| chr19:21500356 | 19 | 21.50036 | 0.11662427 | Credible Set | purple |
| chr19:21520189 | 19 | 21.52019 | 0.11662427 | Credible Set | purple |
| chr19:21729752 | 19 | 21.72975 | 0.11662427 | Credible Set | purple |
| chr19:21991002 | 19 | 21.99100 | 0.06560313 | Credible Set | purple |
| chr19:22115053 | 19 | 22.11505 | 0.06560313 | Credible Set | purple |

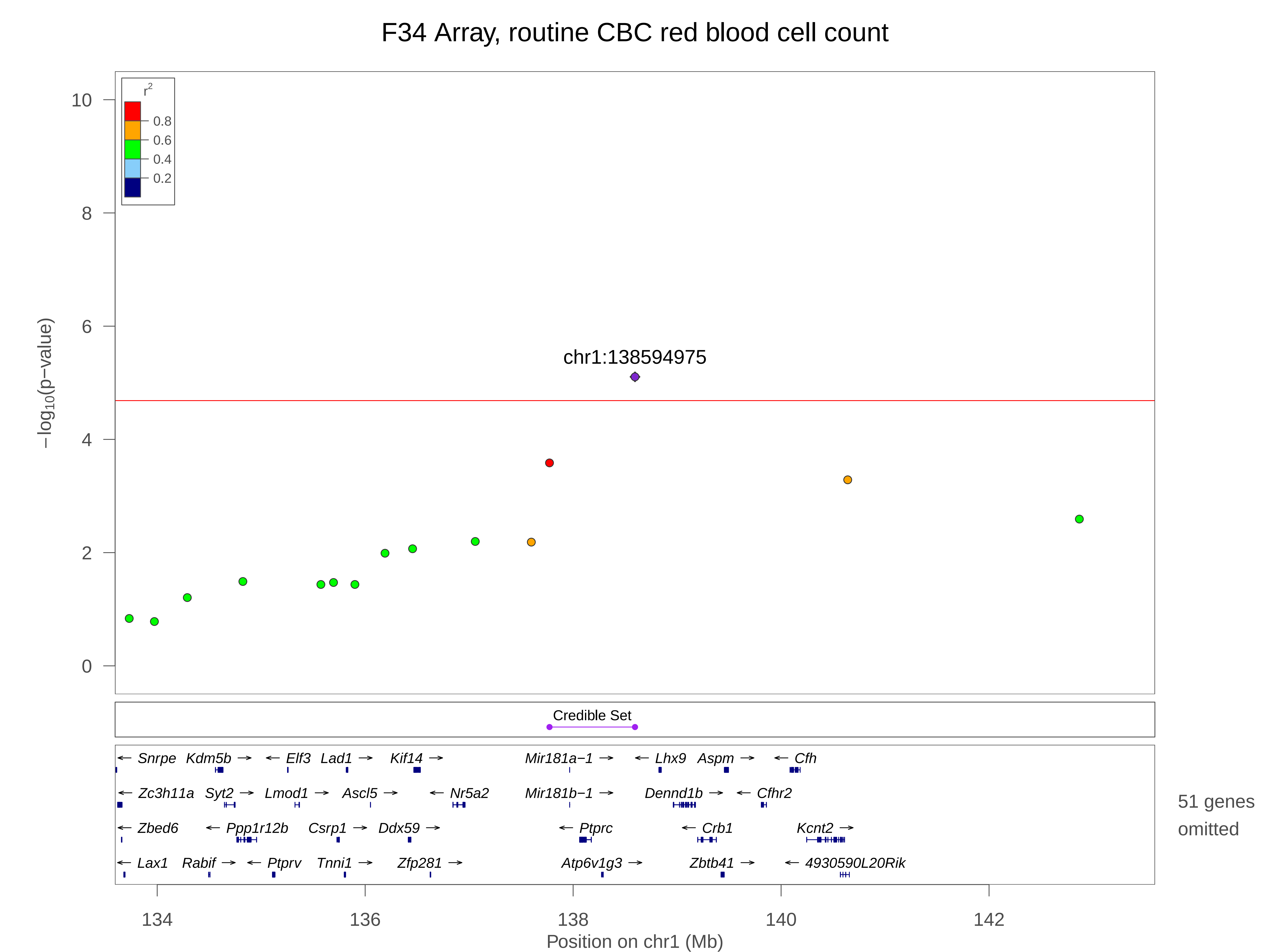

F34 Array, routine CBC red blood cell count
r2
0.8
0.6
0.4
0.2
10
8
6
4
2
0
−log10(p−value)
chr1:138594975
Credible Set
← Snrpe Kdm5b → ← Elf3 Lad1 → Kif14 → Mir181a−1 → ← Lhx9 Aspm → ← Cfh
← Zc3h11a Syt2 → Lmod1 → Ascl5 → ← Nr5a2 Mir181b−1 → Dennd1b → ← Cfhr2
← Zbed6 ← Ppp1r12b Csrp1 → Ddx59 → ← Ptprc ← Crb1 Kcnt2 →
← Lax1 Rabif → ← Ptprv Tnni1 → Zfp281 → Atp6v1g3 → Zbtb41 → ← 4930590L20Rik
51 genes omitted
134
136
138
140
142
Position on chr1 (Mb)

| | |
|---|---|
| date: | Thu Oct 25 16:46:11 2018 |
| build: | F34Array |
| display range: | chr1:133594975−143594975 [133594975−143594975] |
| hilite range: | 0 − 0 [ 0 − 0 ] |
| reference SNP: | chr1:138594975 |
| number of SNPs plotted: | 16 |
| min P−value: | 7.85E−6 [chr1:138594975] |
| max P−value: | 1.65E−1 [chr1:133972959] |
| omitted Genes: | Atp2b4, Mir6903, Optc |
| omitted Genes: | Prelp, Fmod, Btg2 |
| omitted Genes: | Chit1, Chil1, Mybph |
| omitted Genes: | Adora1, Myog, Ppfia4 |
| omitted Genes: | Tmem183a, 4933406M09Rik, Cyb5r1 |
| omitted Genes: | Adipor1, Klhl12, Mgat4e |
| omitted Genes: | Ube2t, Lgr6, Ptpn7 |
| omitted Genes: | Arl8a, Gpr37l1, Rnpep |
| omitted Genes: | Timm17a, Shisa4, Ipo9 |
| omitted Genes: | Nav1, Mir1231, Phlda3 |
| omitted Genes: | Tnnt2, Pkp1, Igfn1 |
| omitted Genes: | Tmem9, Cacna1s, Gm15850 |
| omitted Genes: | Kif21b, 5730559C18Rik, Gpr25 |
| omitted Genes: | Camsap2, 9230116N13Rik, Gm19705 |
| omitted Genes: | Platr22, Nek7, Gm5833 |
| omitted Genes: | 2310009B15Rik, 4930596I21Rik, 4933436E23Rik |
| omitted Genes: | F13b, Cfhr1, Gm4788 |

Make more plots at http://csg.sph.umich.edu/locuszoom/

Fine−mapping Regions

| snp | chr | pos (Mb) | pp | group | color |
|---|---|---|---|---|---|
| chr1:137772769 | 1 | 137.7728 | 0.03498787 | Credible Set | purple |
| chr1:138594975 | 1 | 138.5950 | 0.96501213 | Credible Set | purple |

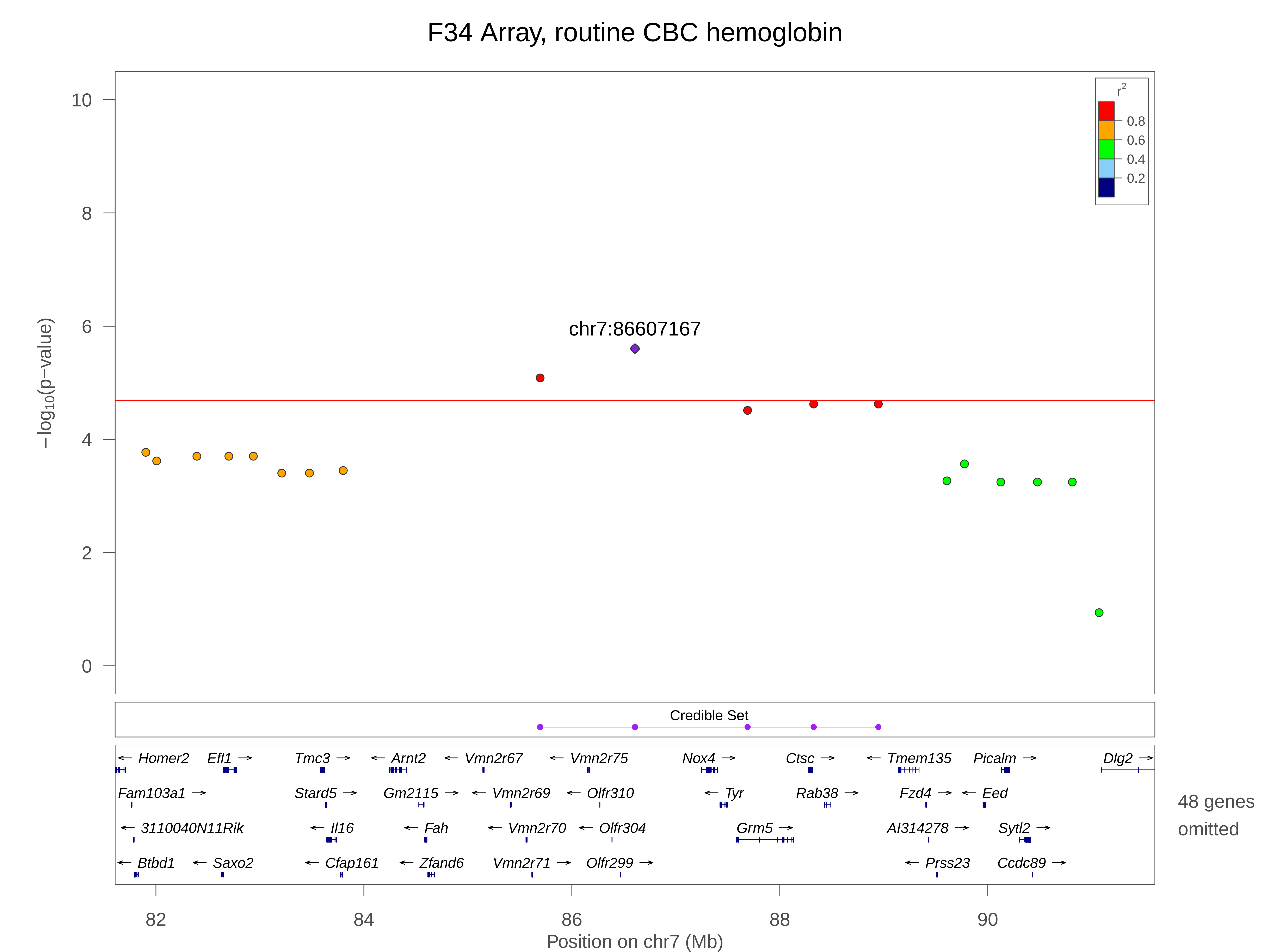
F34 Array, routine CBC hemoglobin
−log10(p−value)
10
8
6
4
2
0
chr7:86607167
r2
0.8
0.6
0.4
0.2
Credible Set
← Homer2
Efl1 →
Tmc3 →
← Arnt2
← Vmn2r67
← Vmn2r75
Nox4 →
Ctsc →
← Tmem135
Picalm →
Dlg2 →
Fam103a1 →
Stard5 →
Gm2115 →
← Vmn2r69
← Olfr310
← Tyr
Rab38 →
Fzd4 →
← Eed
← 3110040N11Rik
← Il16
← Fah
← Vmn2r70
← Olfr304
Grm5 →
AI314278 →
Sytl2 →
← Btbd1
← Saxo2
← Cfap161
← Zfand6
Vmn2r71 →
Olfr299 →
← Prss23
Ccdc89 →
48 genes omitted
82
84
86
88
90
Position on chr7 (Mb)

| | |
|---|---|
| date: | Thu Oct 25 16:47:30 2018 |
| build: | F34Array |
| display range: | chr7:81607167−91607167 [81607167−91607167] |
| hilite range: | 0 − 0 [ 0 − 0 ] |
| reference SNP: | chr7:86607167 |
| number of SNPs plotted: | 20 |
| min P−value: | 2.49E−6 [chr7:86607167] |
| max P−value: | 1.15E−1 [chr7:91070939] |
| omitted Genes: | Tm6sf1, Hdgfrp3, Bnc1 |
| omitted Genes: | Gm20744, Sh3gl3, Adamtsl3 |
| omitted Genes: | 4933406J10Rik, Mex3b, 4933430H16Rik |
| omitted Genes: | Mesdc1, Mesdc2, Cemip |
| omitted Genes: | Abhd17c, 2610206C17Rik, Olfr291 |
| omitted Genes: | Olfr290, Vmn2r65, Vmn2r66 |
| omitted Genes: | Vmn2r68, Vmn2r72, Vmn2r73 |
| omitted Genes: | Vmn2r74, Vmn2r76, Olfr309 |
| omitted Genes: | Olfr308, Olfr307, Olfr305 |
| omitted Genes: | Olfr303, Olfr301, Olfr298 |
| omitted Genes: | Olfr297, Olfr295, Olfr294 |
| omitted Genes: | Olfr293, Olfr292, Folh1 |
| omitted Genes: | Vmn2r77, Vmn2r78, Vmn2r79 |
| omitted Genes: | Me3, Ccdc81, I7Rn6 |
| omitted Genes: | E230029C05Rik, 2310010J17Rik, Ccdc83 |
| omitted Genes: | Crebzf, Tmem126a, Tmem126b |

Make more plots at http://csg.sph.umich.edu/locuszoom/

Fine−mapping Regions

| snp | chr | pos (Mb) | pp | group | color |
|---|---|---|---|---|---|
| chr7:85694603 | 7 | 85.69460 | 0.19438670 | Credible Set | purple |
| chr7:86607167 | 7 | 86.60717 | 0.60948915 | Credible Set | purple |
| chr7:87689885 | 7 | 87.68989 | 0.05528989 | Credible Set | purple |
| chr7:88325650 | 7 | 88.32565 | 0.07041713 | Credible Set | purple |
| chr7:88947234 | 7 | 88.94723 | 0.07041713 | Credible Set | purple |

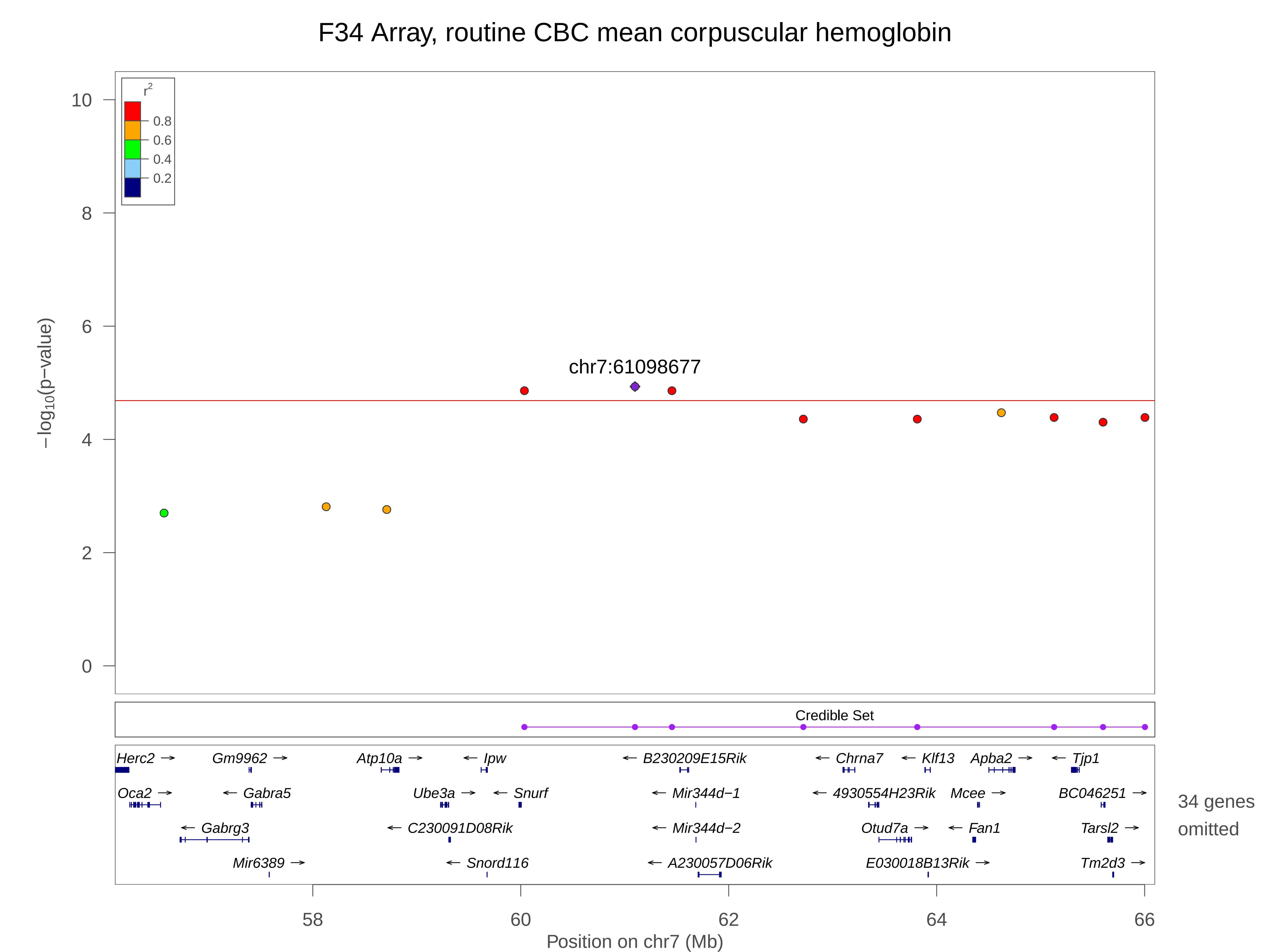

F34 Array, routine CBC mean corpuscular hemoglobin
r2
0.8
0.6
0.4
0.2
−log10(p−value)
10
8
6
4
2
0
chr7:61098677
Credible Set
Herc2 →
Gm9962 →
Atp10a →
← Ipw
← B230209E15Rik
← Chrna7
← Klf13
Apba2 →
← Tjp1
Oca2 →
← Gabra5
Ube3a →
← Snurf
← Mir344d−1
← 4930554H23Rik
Mcee →
BC046251 →
← Gabrg3
← C230091D08Rik
← Mir344d−2
Otud7a →
← Fan1
Tarsl2 →
Mir6389 →
← Snord116
← A230057D06Rik
E030018B13Rik →
Tm2d3 →
34 genes omitted
58
60
62
64
66
Position on chr7 (Mb)

| | |
|---|---|
| date: | Thu Oct 25 16:52:02 2018 |
| build: | F34Array |
| display range: | chr7:56098677−66098677 [56098677−66098677] |
| hilite range: | 0 − 0 [ 0 − 0 ] |
| reference SNP: | chr7:61098677 |
| number of SNPs plotted: | 13 |
| min P−value: | 1.17E−5 [chr7:61098677] |
| max P−value: | 2E−3 [chr7:56569907] |
| omitted Genes: | Gabrb3, Snord116l2, Snord116l1 |
| omitted Genes: | Snhg14, Snord64, Snrpn |
| omitted Genes: | Gm7367, Mir344d−3, Mir344e |
| omitted Genes: | Mir344h−1, Mir344h−2, Mir344b |
| omitted Genes: | Mir344c, Mir344, Mir344−2 |
| omitted Genes: | A330076H08Rik, Mir344g, Mir344f |
| omitted Genes: | Mir344i, Ndn, Magel2 |
| omitted Genes: | Mkrn3, Peg12, Trpm1 |
| omitted Genes: | Mir211, Mir6416, Mtmr10 |
| omitted Genes: | Mphosph10, Fam189a1, Ndnl2 |
| omitted Genes: | 1810008I18Rik, Pcsk6, Snrpa1 |
| omitted Genes: | Vimp |

Make more plots at http://csg.sph.umich.edu/locuszoom/

Fine−mapping Regions

| snp | chr | pos (Mb) | pp | group | color |
|---|---|---|---|---|---|
| chr7:60035047 | 7 | 60.03505 | 0.20630661 | Credible Set | purple |
| chr7:61098677 | 7 | 61.09868 | 0.24234896 | Credible Set | purple |
| chr7:61453986 | 7 | 61.45399 | 0.20630661 | Credible Set | purple |
| chr7:62717634 | 7 | 62.71763 | 0.06883805 | Credible Set | purple |
| chr7:63813305 | 7 | 63.81330 | 0.06883805 | Credible Set | purple |
| chr7:65129296 | 7 | 65.12930 | 0.07316961 | Credible Set | purple |
| chr7:65600162 | 7 | 65.60016 | 0.06102250 | Credible Set | purple |
| chr7:66002947 | 7 | 66.00295 | 0.07316961 | Credible Set | purple |

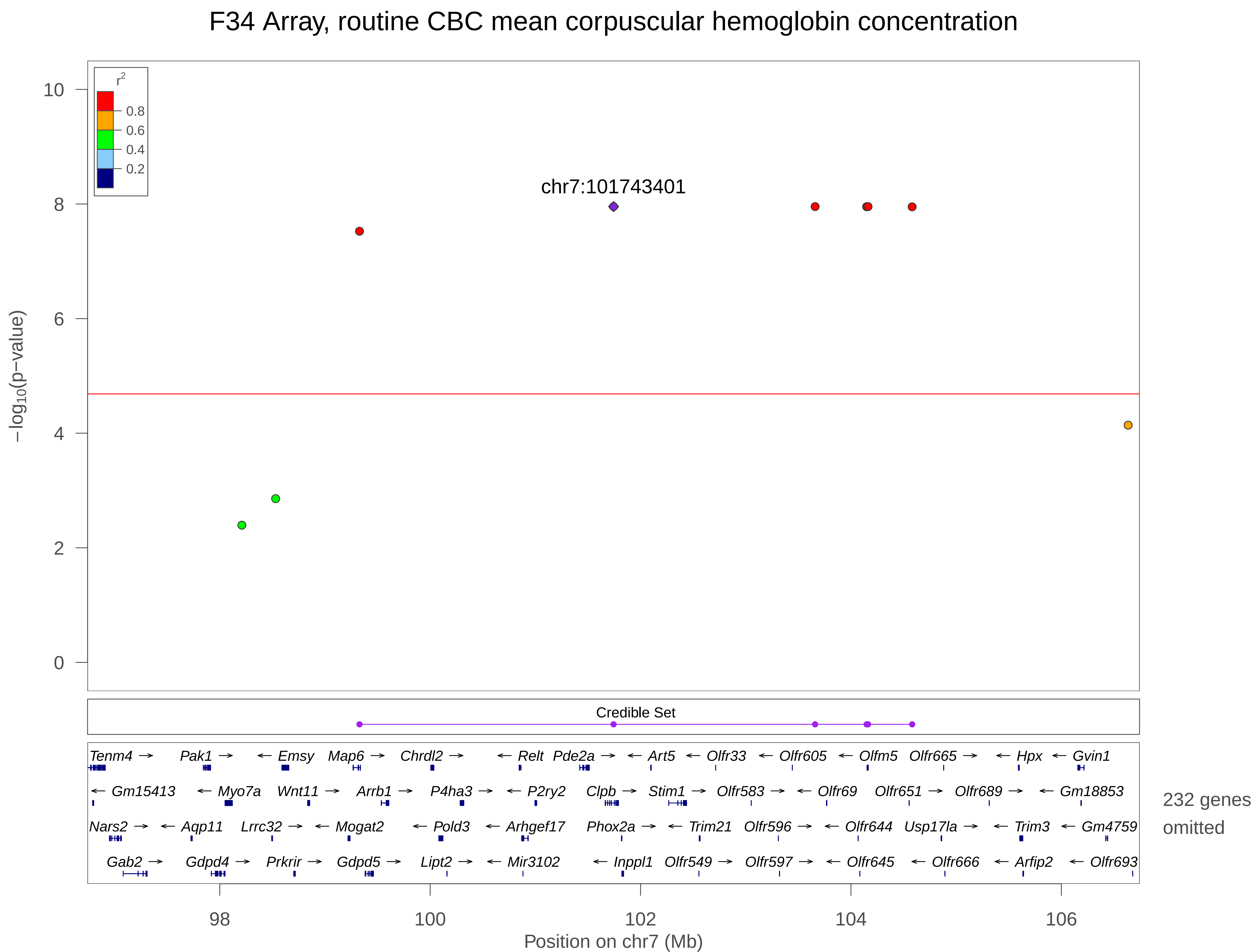

F34 Array, routine CBC mean corpuscular hemoglobin concentration
r2
0.8
0.6
0.4
0.2
chr7:101743401
−log10(p−value)
10
8
6
4
2
0
Credible Set
Tenm4 → Pak1 → ← Emsy Map6 → Chrdl2 → ← Relt Pde2a → ← Art5 ← Olfr33 ← Olfr605 ← Olfm5 Olfr665 → ← Hpx ← Gvin1
← Gm15413 ← Myo7a Wnt11 → Arrb1 → P4ha3 → ← P2ry2 Clpb → Stim1 → Olfr583 → ← Olfr69 Olfr651 → Olfr689 → ← Gm18853
Nars2 → ← Aqp11 Lrrc32 → ← Mogat2 ← Pold3 ← Arhgef17 Phox2a → ← Trim21 Olfr596 → ← Olfr644 Usp17la → ← Trim3 ← Gm4759
Gab2 → Gdpd4 → Prkrir → Gdpd5 → Lipt2 → ← Mir3102 ← Inppl1 Olfr549 → Olfr597 → ← Olfr645 ← Olfr666 ← Arfip2 ← Olfr693
232 genes omitted
98
100
102
104
106
Position on chr7 (Mb)

| | |
|---|---|
| date: | Thu Oct 25 16:52:13 2018 |
| build: | F34Array |
| display range: | chr7:96743401−106743401 [96743401−106743401] |
| hilite range: | 0 − 0 [ 0 − 0 ] |
| reference SNP: | chr7:101743401 |
| number of SNPs plotted: | 10 |
| min P−value: | 1.11E−8 [chr7:101743401] |
| max P−value: | 4.03E−3 [chr7:98210049] |
| omitted Genes: | Usp35, Kctd21, Alg8 |
| omitted Genes: | Ndufc2, Thrsp, Kctd14 |
| omitted Genes: | Ints4, Aamdc, Rsf1 |
| omitted Genes: | Rsf1os1, Clns1a, Capn5 |
| omitted Genes: | Omp, Gm16938, B3gnt6 |
| omitted Genes: | Acer3, Tsku, Gucy2d |
| omitted Genes: | Uvrag, Dgat2, Gm26705 |
| omitted Genes: | Serpinh1, Klhl35, Rps3 |
| omitted Genes: | Snord15b, Snord15a, Mir326 |
| omitted Genes: | Tpbgl, Slco2b1, Olfr520 |
| omitted Genes: | Olfr521, Neu3, Spcs2 |
| omitted Genes: | Xrra1, Rnf169, Kcne3 |
| omitted Genes: | Pgm2l1, Gpx2−ps1, Ppme1 |
| omitted Genes: | C2cd3, Ucp3, Ucp2 |
| omitted Genes: | Dnajb13, Coa4, Mrpl48 |
| omitted Genes: | Rab6a, Plekhb1, Fam168a |
| omitted Genes: | P2ry6, Fchsd2, Atg16l2 |
| omitted Genes: | Stard10, Arap1, Mir139 |
| omitted Genes: | Art2a−ps, Art2b, Folr2 |
| omitted Genes: | Folr1, Anapc15, Tomt |
| omitted Genes: | Lamtor1, Lrrc51, Numa1 |
| omitted Genes: | Il18bp, Rnf121, Xndc1 |
| omitted Genes: | Xntrpc, Trpc2, Art1 |
| omitted Genes: | Chrna10, Nup98, Pgap2 |
| omitted Genes: | Rhog, Rrm1, Olfr543 |
| omitted Genes: | Olfr544, Olfr545, Olfr547 |
| omitted Genes: | Olfr550, Olfr551, Olfr552 |
| omitted Genes: | Olfr553, Olfr554, Olfr555 |
| omitted Genes: | Olfr556, Trim68, Olfr557 |
| omitted Genes: | Olfr558, Olfr559, Olfr78 |
| omitted Genes: | Olfr560, Olfr561, Olfr564 |
| omitted Genes: | Olfr566, Olfr568, Olfr569 |

Make more plots at http://csg.sph.umich.edu/locuszoom/

Fine−mapping Regions

| snp | chr | pos (Mb) | pp | group | color |
|---|---|---|---|---|---|
| chr7:99328376 | 7 | 99.32838 | 0.07118739 | Credible Set | purple |
| chr7:101743401 | 7 | 101.74340 | 0.18627943 | Credible Set | purple |
| chr7:103659355 | 7 | 103.65936 | 0.18627943 | Credible Set | purple |
| chr7:104149022 | 7 | 104.14902 | 0.18547207 | Credible Set | purple |
| chr7:104162444 | 7 | 104.16244 | 0.18627943 | Credible Set | purple |
| chr7:104581631 | 7 | 104.58163 | 0.18450225 | Credible Set | purple |

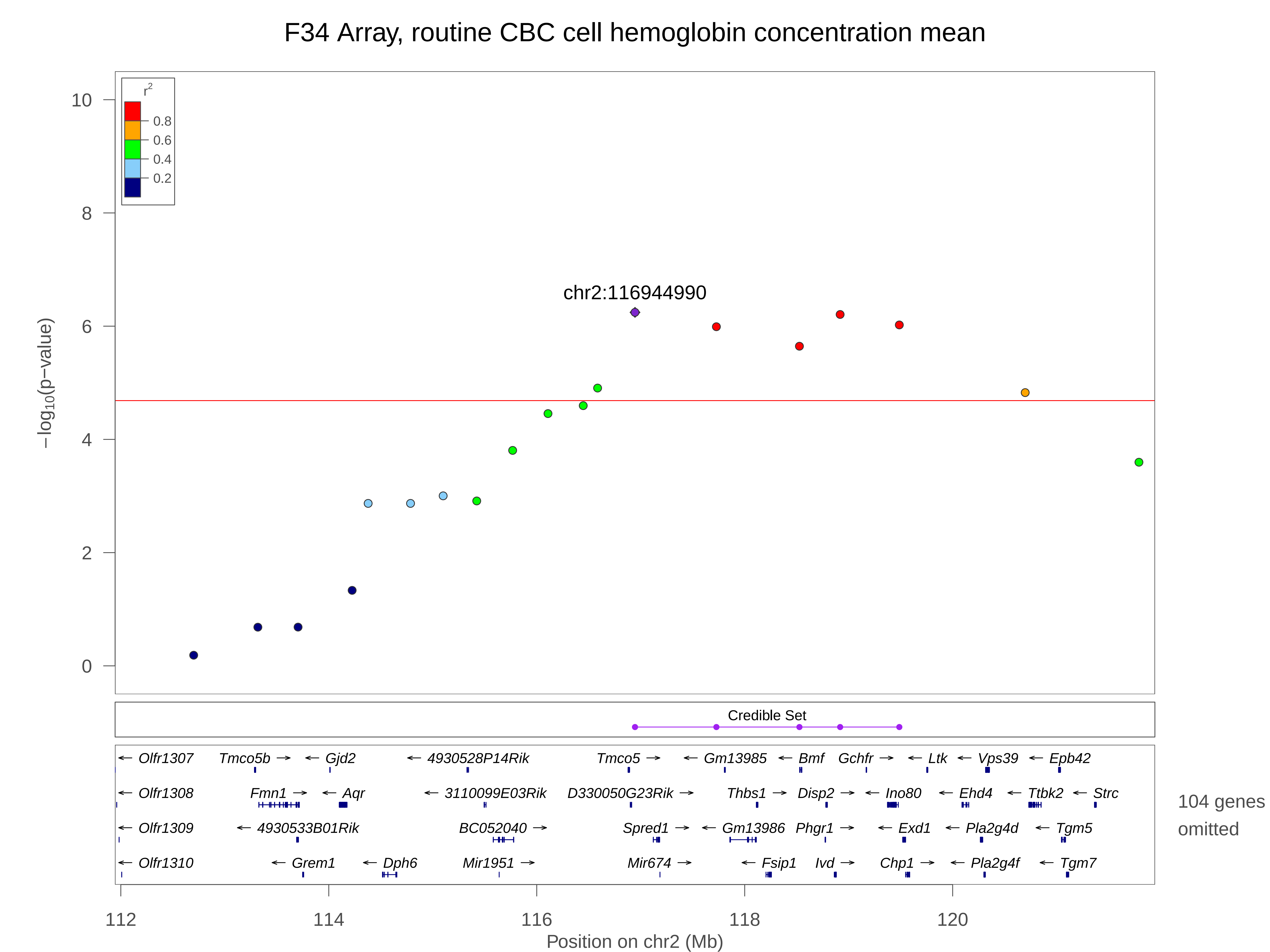
F34 Array, routine CBC cell hemoglobin concentration mean
r2
0.8
0.6
0.4
0.2
10
8
6
4
2
0
−log10(p−value)
chr2:116944990
Credible Set
← Olfr1307
Tmco5b →
← Gjd2
← 4930528P14Rik
Tmco5 →
← Gm13985
← Bmf
Gchfr →
← Ltk
← Vps39
← Epb42
← Olfr1308
Fmn1 →
← Aqr
← 3110099E03Rik
D330050G23Rik →
Thbs1 →
Disp2 →
← Ino80
← Ehd4
← Ttbk2
← Strc
← Olfr1309
← 4930533B01Rik
BC052040 →
Spred1 →
← Gm13986
Phgr1 →
← Exd1
← Pla2g4d
← Tgm5
← Olfr1310
← Grem1
← Dph6
Mir1951 →
Mir674 →
← Fsip1
Ivd →
Chp1 →
← Pla2g4f
← Tgm7
104 genes omitted
112
114
116
118
120
Position on chr2 (Mb)

date: Thu Oct 25 16:52:29 2018
build: F34Array
display range: chr2:111944990−121944990 [111944990−121944990]
hilite range: 0 − 0 [ 0 − 0 ]
reference SNP: chr2:116944990
number of SNPs plotted: 20
min P−value: 5.7E−7 [chr2:116944990]
max P−value: 6.48E−1 [chr2:112700792]
omitted Genes: Olfr1311, Olfr1312, Olfr1313
omitted Genes: Olfr1314, Olfr1316, Olfr1317
omitted Genes: Olfr1318, Lpcat4, Nutm1
omitted Genes: Nop10, Slc12a6, Emc4
omitted Genes: Katnbl1, Emc7, Chrm5
omitted Genes: Aven, Ryr3, Scg5
omitted Genes: Arhgap11a, A530058N18Rik, Actc1
omitted Genes: C130080G10Rik, Mir7002, Zfp770
omitted Genes: Meis2, 2810405F15Rik, Fam98b
omitted Genes: Rasgrp1, 4930412B13Rik, Gpr176
omitted Genes: 1700054M17Rik, Eif2ak4, Srp14
omitted Genes: Bub1b, Pak6, Ankrd63
omitted Genes: Plcb2, Inafm2, A430105I19Rik
omitted Genes: Knstrn, Bahd1, Chst14
omitted Genes: Ccdc32, Rpusd2, Casc5
omitted Genes: Rad51, Rmdn3, Dnajc17
omitted Genes: Gm14137, Zfyve19, Ppp1r14d
omitted Genes: Spint1, Rhov, Vps18
omitted Genes: Gm14207, Dll4, Chac1
omitted Genes: 1700020I14Rik, Oip5, Nusap1
omitted Genes: Ndufaf1, Rtf1, Itpka
omitted Genes: Rpap1, Tyro3, Mga
omitted Genes: Mapkbp1, Mir7665, Jmjd7
omitted Genes: Gm28042, Pla2g4b, Pla2g4e
omitted Genes: Tmem87a, Ganc, 4931402G19Rik
omitted Genes: Capn3, Zfp106, Snap23
omitted Genes: Lrrc57, Haus2, Cdan1
omitted Genes: AV039307, Ubr1, Tmem62
omitted Genes: Ccndbp1, Lcmt2, Adal
omitted Genes: Zscan29, Tubgcp4, Trp53bp1
omitted Genes: Map1a, Ppip5k1, Ckmt1
Make more plots at http://csg.sph.umich.edu/locuszoom/
omitted Genes: Catsper2, Pdia3, Ell3

Fine−mapping Regions

| snp | chr | pos (Mb) | pp | group | color |
|---|---|---|---|---|---|
| chr2:116944990 | 2 | 116.9450 | 0.29685060 | Credible Set | purple |
| chr2:117727776 | 2 | 117.7278 | 0.16923215 | Credible Set | purple |
| chr2:118526291 | 2 | 118.5263 | 0.07893283 | Credible Set | purple |
| chr2:118917838 | 2 | 118.9178 | 0.27342629 | Credible Set | purple |
| chr2:119486992 | 2 | 119.4870 | 0.18155814 | Credible Set | purple |

F34 Array, routine CBC cell hemoglobin concentration mean

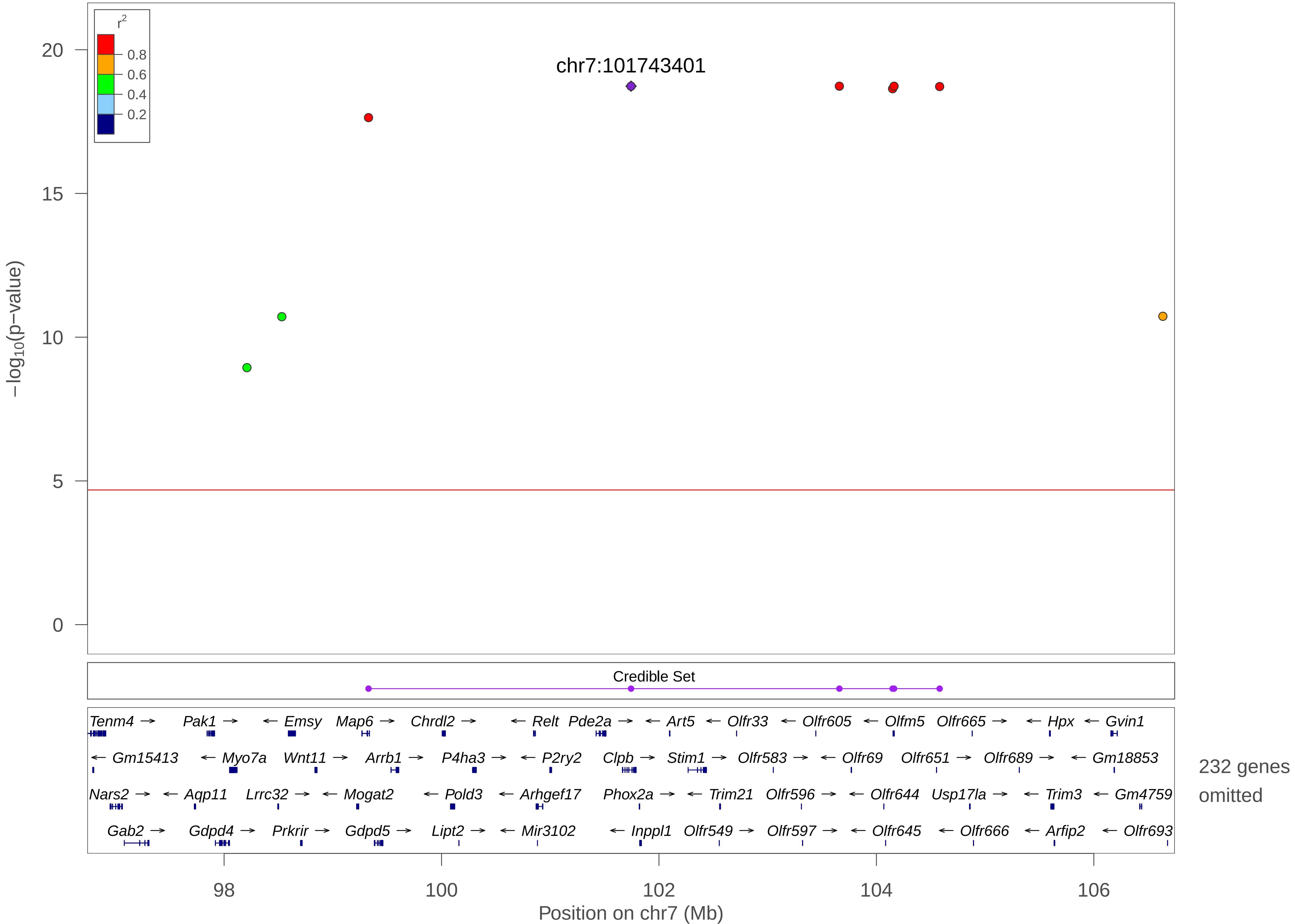

date: Thu Oct 25 16:52:40 2018
build: F34Array
display range: chr7:96743401−106743401 [96743401−106743401]
hilite range: 0 − 0 [ 0 − 0 ]
reference SNP: chr7:101743401
number of SNPs plotted: 10
min P−value: 1.85E−19 [chr7:101743401]
max P−value: 1.15E−9 [chr7:98210049]
omitted Genes: Usp35, Kctd21, Alg8
omitted Genes: Ndufc2, Thrsp, Kctd14
omitted Genes: Ints4, Aamdc, Rsf1
omitted Genes: Rsf1os1, Clns1a, Capn5
omitted Genes: Omp, Gm16938, B3gnt6
omitted Genes: Acer3, Tsku, Gucy2d
omitted Genes: Uvrag, Dgat2, Gm26705
omitted Genes: Serpinh1, Klhl35, Rps3
omitted Genes: Snord15b, Snord15a, Mir326
omitted Genes: Tpbgl, Slco2b1, Olfr520
omitted Genes: Olfr521, Neu3, Spcs2
omitted Genes: Xrra1, Rnf169, Kcne3
omitted Genes: Pgm2l1, Gpx2−ps1, Ppme1
omitted Genes: C2cd3, Ucp3, Ucp2
omitted Genes: Dnajb13, Coa4, Mrpl48
omitted Genes: Rab6a, Plekhb1, Fam168a
omitted Genes: P2ry6, Fchsd2, Atg16l2
omitted Genes: Stard10, Arap1, Mir139
omitted Genes: Art2a−ps, Art2b, Folr2
omitted Genes: Folr1, Anapc15, Tomt
omitted Genes: Lamtor1, Lrrc51, Numa1
omitted Genes: Il18bp, Rnf121, Xndc1
omitted Genes: Xntrpc, Trpc2, Art1
omitted Genes: Chrna10, Nup98, Pgap2
omitted Genes: Rhog, Rrm1, Olfr543
omitted Genes: Olfr544, Olfr545, Olfr547
omitted Genes: Olfr550, Olfr551, Olfr552
omitted Genes: Olfr553, Olfr554, Olfr555
omitted Genes: Olfr556, Trim68, Olfr557
omitted Genes: Olfr558, Olfr559, Olfr78
omitted Genes: Olfr560, Olfr561, Olfr564
Make more plots at http://csg.sph.umich.edu/locuszoom/
omitted Genes: Olfr566, Olfr568, Olfr569

Fine−mapping Regions

| snp | chr | pos (Mb) | pp | group | color |
|---|---|---|---|---|---|
| chr7:99328376 | 7 | 99.32838 | 0.01707747 | Credible Set | purple |
| chr7:101743401 | 7 | 101.74340 | 0.20517644 | Credible Set | purple |
| chr7:103659355 | 7 | 103.65936 | 0.20517644 | Credible Set | purple |
| chr7:104149022 | 7 | 104.14902 | 0.16778480 | Credible Set | purple |
| chr7:104162444 | 7 | 104.16244 | 0.20517644 | Credible Set | purple |
| chr7:104581631 | 7 | 104.58163 | 0.19960843 | Credible Set | purple |

### F34 Array, routine CBC cell hemoglobin concentration mean

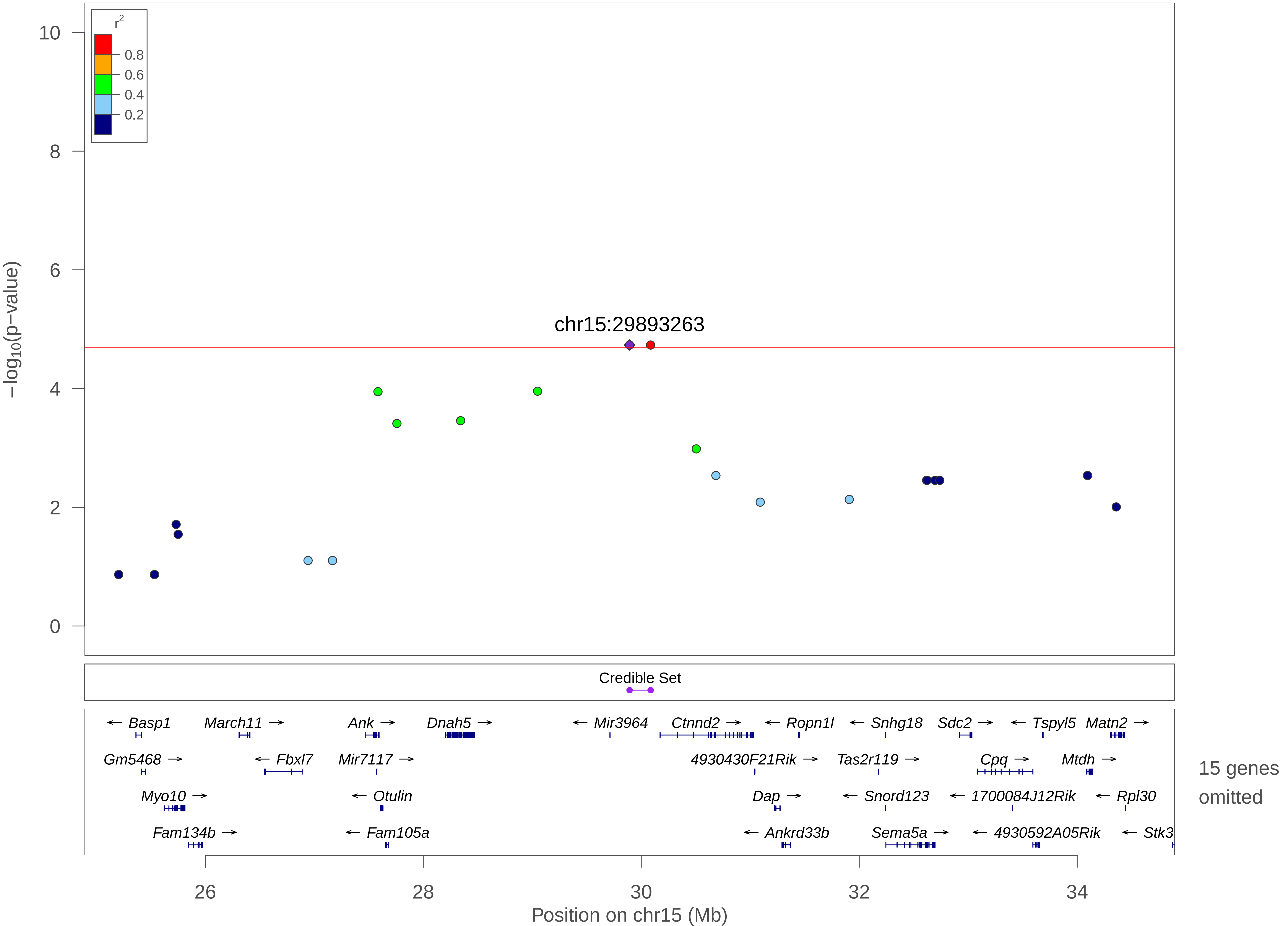

| | |
|---|---|
| date: | Thu Oct 25 16:52:20 2018 |
| build: | F34Array |
| display range: | chr15:24893263−34893263 [24893263−34893263] |
| hilite range: | 0 − 0 [ 0 − 0 ] |
| reference SNP: | chr15:29893263 |
| number of SNPs plotted: | 23 |
| min P−value: | 1.85E−5 [chr15:29893263] |
| max P−value: | 1.36E−1 [chr15:25205426] |
| omitted Genes: | Mir7212, Zfp622, Trio |
| omitted Genes: | March6, Cmbl, Cct5 |
| omitted Genes: | Fam173b, Laptm4b, 9430069I07Rik |
| omitted Genes: | Erich5, Rida, Pop1 |
| omitted Genes: | Nipal2, 4930413F20Rik, Kcns2 |

Make more plots at http://csg.sph.umich.edu/locuszoom/

Fine−mapping Regions

| snp | chr | pos (Mb) | pp | group | color |
|---|---|---|---|---|---|
| chr15:29893263 | 15 | 29.89326 | 0.5 | Credible Set | purple |
| chr15:30086139 | 15 | 30.08614 | 0.5 | Credible Set | purple |

F34 Array, routine CBC red blood cell distribution width

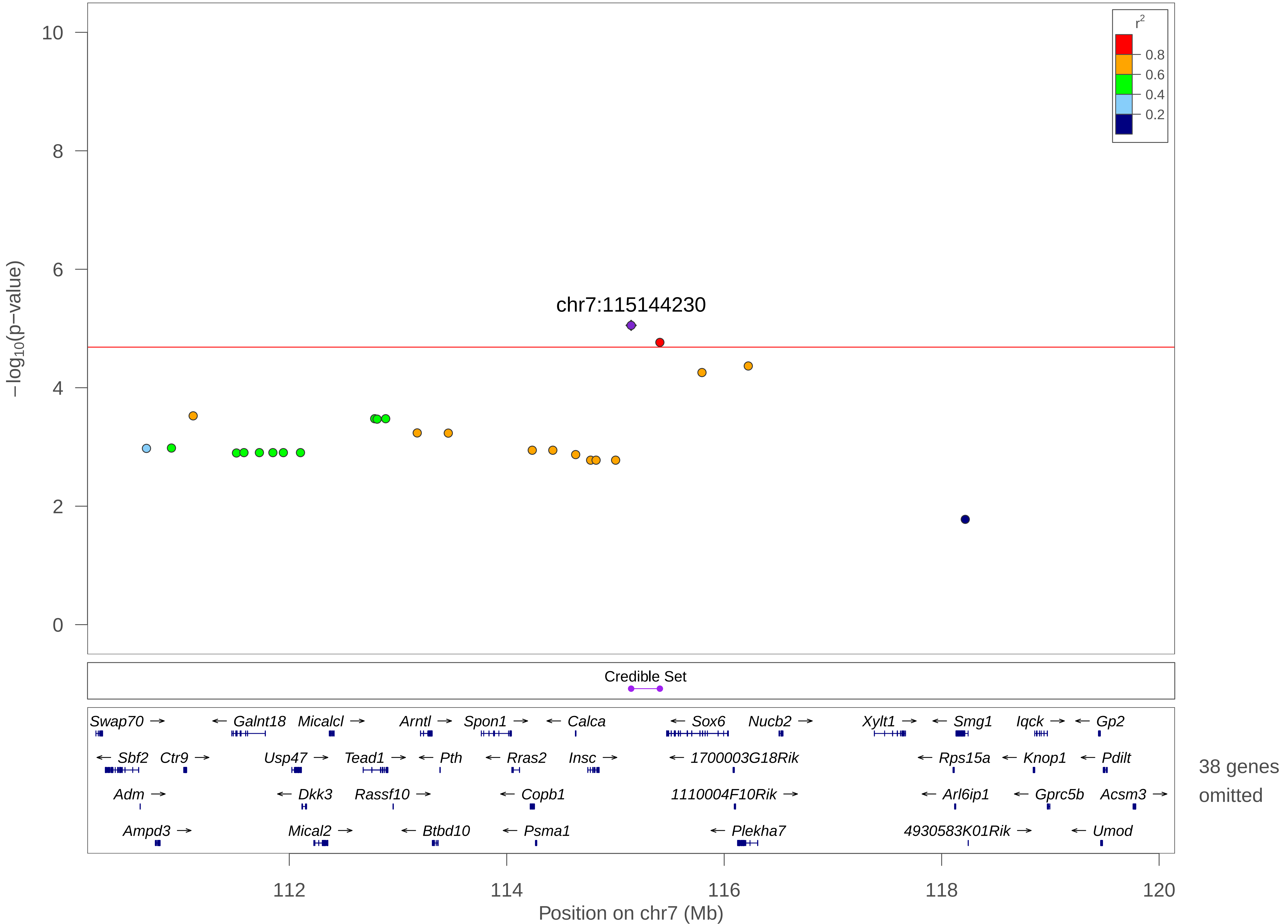

| | |
|---|---|
| date: | Thu Oct 25 16:53:02 2018 |
| build: | F34Array |
| display range: | chr7:110144230−120144230 [110144230−120144230] |
| hilite range: | 0 − 0 [ 0 − 0 ] |
| reference SNP: | chr7:115144230 |
| number of SNPs plotted: | 26 |
| min P−value: | 8.86E−6 [chr7:115144230] |
| max P−value: | 1.67E−2 [chr7:118217303] |
| omitted Genes: | Rnf141, Lyve1, Mrvi1 |
| omitted Genes: | Gm16336, Eif4g2, 1700012D14Rik |
| omitted Genes: | Parva, 4930543E12Rik, Far1 |
| omitted Genes: | Far1os, 4933406I18Rik, Pde3b |
| omitted Genes: | Cyp2r1, Calcb, A730082K24Rik |
| omitted Genes: | Rps13, Snord14a, Pik3c2a |
| omitted Genes: | Syt17, Itpripl2, Coq7 |
| omitted Genes: | Tmc7, Tmc5, Gde1 |
| omitted Genes: | Ccp110, 9030624J02Rik, Gpr139 |
| omitted Genes: | Acsm5, Acsm2, Acsm1 |
| omitted Genes: | Acsm4, Thumpd1, Eri2 |
| omitted Genes: | 2610020H08Rik, Dcun1d3, Lyrm1 |
| omitted Genes: | Tmem159, Zp2 |

Make more plots at http://csg.sph.umich.edu/locuszoom/

Fine−mapping Regions

| snp | chr | pos (Mb) | pp | group | color |
|---|---|---|---|---|---|
| chr7:115144230 | 7 | 115.1442 | 0.6525872 | Credible Set | purple |
| chr7:115408492 | 7 | 115.4085 | 0.3474128 | Credible Set | purple |

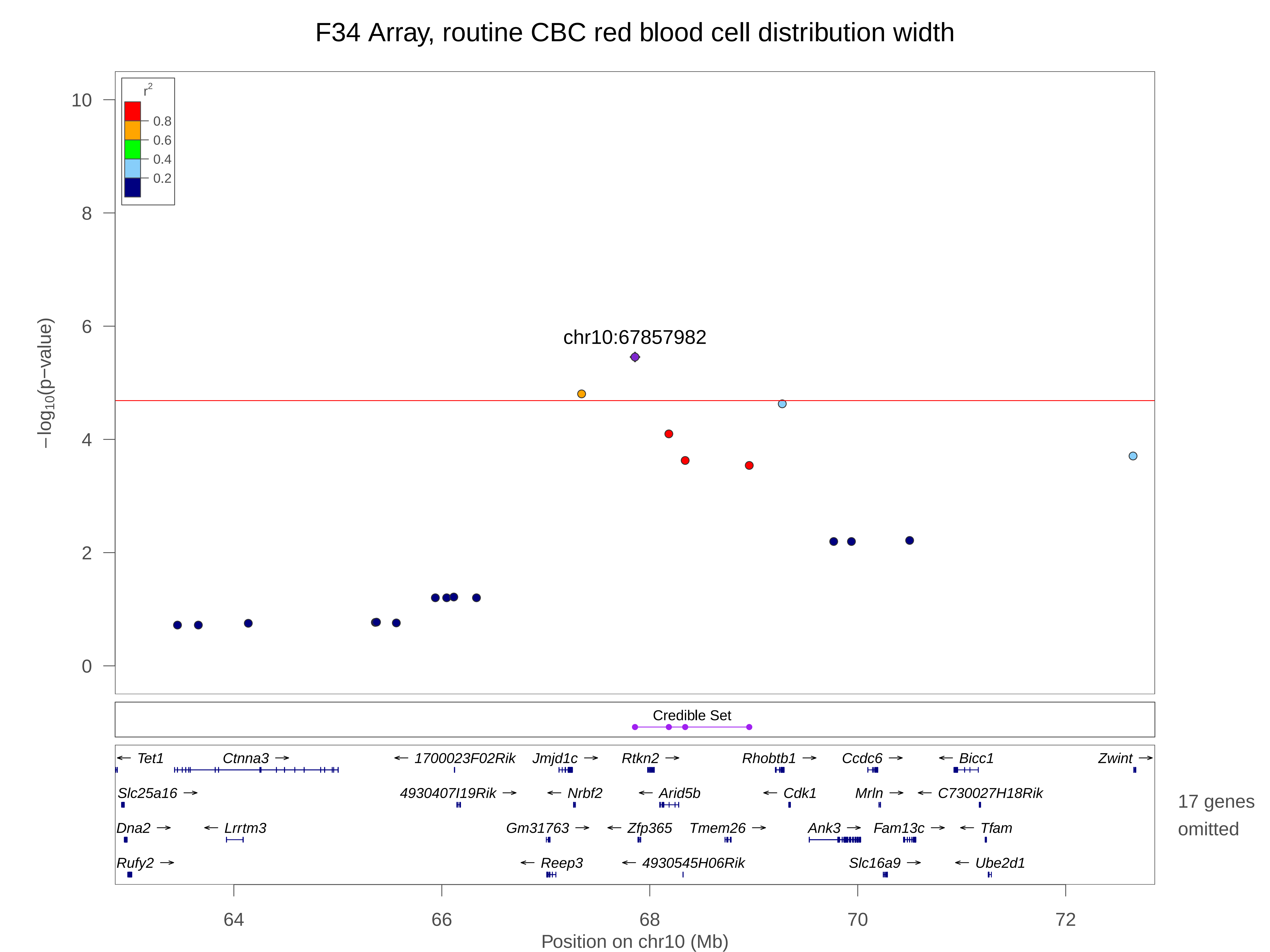
F34 Array, routine CBC red blood cell distribution width
r2
0.8
0.6
0.4
0.2
−log10(p−value)
10
8
6
4
2
0
chr10:67857982
Credible Set
Tet1
Ctnna3
1700023F02Rik
Jmjd1c
Rtkn2
Rhobtb1
Ccdc6
Bicc1
Zwint
Slc25a16
4930407I19Rik
Nrbf2
Arid5b
Cdk1
Mrln
C730027H18Rik
Dna2
Lrrtm3
Gm31763
Zfp365
Tmem26
Ank3
Fam13c
Tfam
Rufy2
Reep3
4930545H06Rik
Slc16a9
Ube2d1
17 genes omitted
64
66
68
70
72
Position on chr10 (Mb)

| | |
|---|---|
| date: | Thu Oct 25 16:52:55 2018 |
| build: | F34Array |
| display range: | chr10:62857982−72857982 [62857982−72857982] |
| hilite range: | 0 − 0 [ 0 − 0 ] |
| reference SNP: | chr10:67857982 |
| number of SNPs plotted: | 21 |
| min P−value: | 3.5E−6 [chr10:67857982] |
| max P−value: | 1.9E−1 [chr10:63458094] |
| omitted Genes: | Hnrnph3, Pbld2, Pbld1 |
| omitted Genes: | Atoh7, Mypn, Herc4 |
| omitted Genes: | Sirt1, Dnajc12, Egr2 |
| omitted Genes: | Ado, 1700040L02Rik, Phyhipl |
| omitted Genes: | D630013N20Rik, 1700113B09Rik, Cisd1 |
| omitted Genes: | Ipmk, 1700049L16Rik |

Make more plots at http://csg.sph.umich.edu/locuszoom/

Fine−mapping Regions

| snp | chr | pos (Mb) | pp | group | color |
|---|---|---|---|---|---|
| chr10:67857982 | 10 | 67.85798 | 0.92212713 | Credible Set | purple |
| chr10:68183490 | 10 | 68.18349 | 0.04691722 | Credible Set | purple |
| chr10:68340621 | 10 | 68.34062 | 0.01693441 | Credible Set | purple |
| chr10:68956755 | 10 | 68.95676 | 0.01402124 | Credible Set | purple |

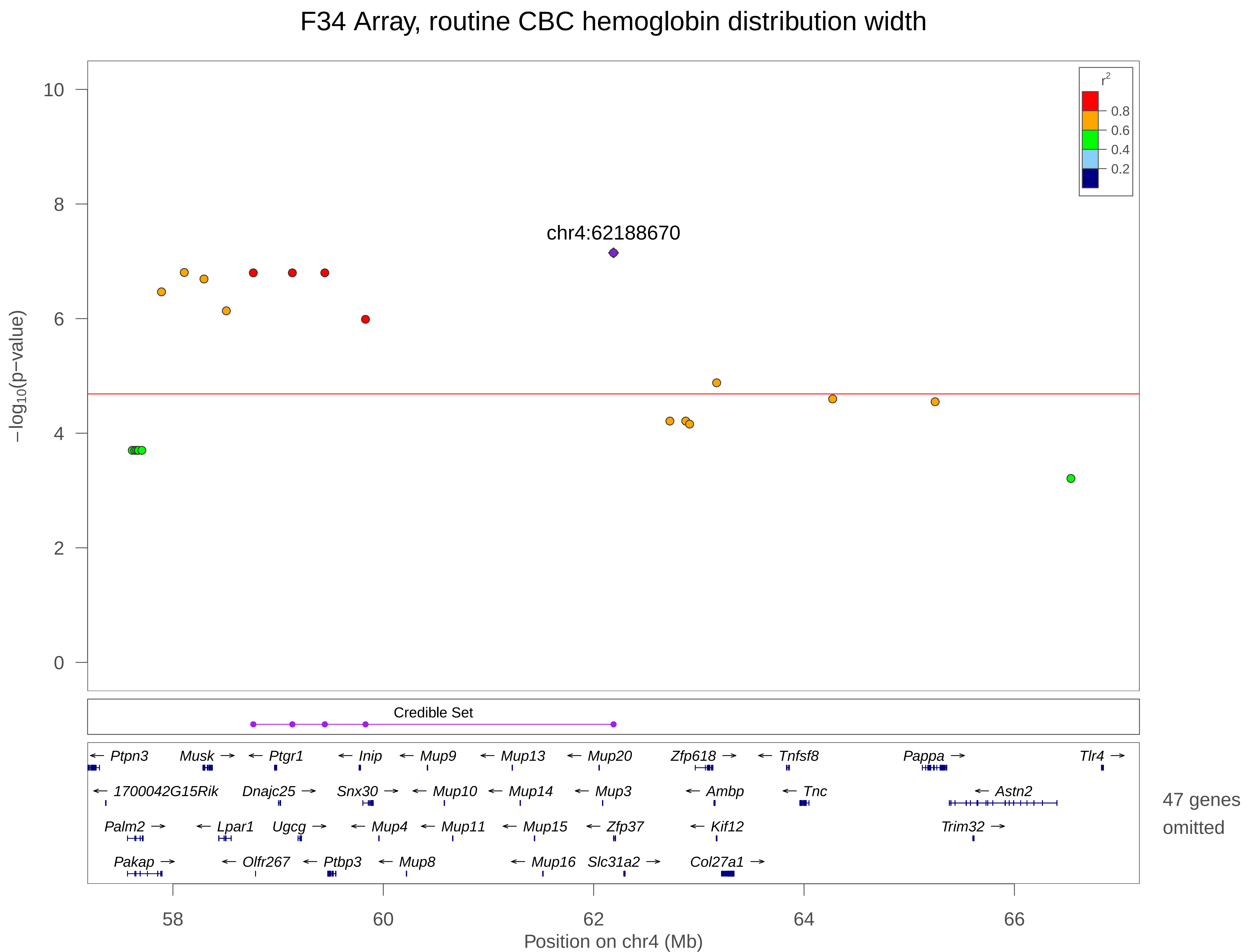

F34 Array, routine CBC hemoglobin distribution width
−log10(p−value)
10
8
6
4
2
0
r2
0.8
0.6
0.4
0.2
chr4:62188670
Credible Set
← Ptpn3
Musk →
← Ptgr1
← Inip
← Mup9
← Mup13
← Mup20
Zfp618 →
← Tnfsf8
Pappa →
Tlr4 →
← 1700042G15Rik
Dnajc25 →
Snx30 →
← Mup10
← Mup14
← Mup3
← Ambp
← Tnc
← Astn2
Palm2 →
← Lpar1
Ugcg →
← Mup4
← Mup11
← Mup15
← Zfp37
← Kif12
Trim32 →
Pakap →
← Olfr267
← Ptbp3
← Mup8
← Mup16
Slc31a2 →
Col27a1 →
47 genes omitted
58
60
62
64
66
Position on chr4 (Mb)

| | |
|---|---|
| date: | Thu Oct 25 16:45:10 2018 |
| build: | F34Array |
| display range: | chr4:57188670−67188670 [57188670−67188670] |
| hilite range: | 0 − 0 [ 0 − 0 ] |
| reference SNP: | chr4:62188670 |
| number of SNPs plotted: | 23 |
| min P−value: | 7.11E−8 [chr4:62188670] |
| max P−value: | 6.17E−4 [chr4:66536974] |
| omitted Genes: | Akap2, D630039A03Rik, Txn1 |
| omitted Genes: | Txndc8, Svep1, Mir3095 |
| omitted Genes: | AI314180, Zkscan16, Gng10 |
| omitted Genes: | Gm12596, Susd1, Hsdl2 |
| omitted Genes: | E130308A19Rik, Slc46a2, Mup6 |
| omitted Genes: | Mup7, Mup2, Mup1 |
| omitted Genes: | Mup12, Mup17, Mup18 |
| omitted Genes: | Mup19, Mup5, Mup21 |
| omitted Genes: | Fkbp15, Slc31a1, Cdc26 |
| omitted Genes: | Prpf4, Rnf183, Wdr31 |
| omitted Genes: | Bspry, Hdhd3, Alad |
| omitted Genes: | Pole3, 4933430I17Rik, Rgs3 |
| omitted Genes: | Mir455, Orm1, Orm3 |
| omitted Genes: | Orm2, Akna, Whrn |
| omitted Genes: | Atp6v1g1, 6330416G13Rik, Gm11213 |
| omitted Genes: | 1700018C11Rik, Tnfsf15 |

Make more plots at http://csg.sph.umich.edu/locuszoom/

Fine−mapping Regions

| snp | chr | pos (Mb) | pp | group | color |
|---|---|---|---|---|---|
| chr4:58764292 | 4 | 58.76429 | 0.18717006 | Credible Set | purple |
| chr4:59135384 | 4 | 59.13538 | 0.18717006 | Credible Set | purple |
| chr4:59443996 | 4 | 59.44400 | 0.18717006 | Credible Set | purple |
| chr4:59830858 | 4 | 59.83086 | 0.03088785 | Credible Set | purple |
| chr4:62188670 | 4 | 62.18867 | 0.40760197 | Credible Set | purple |

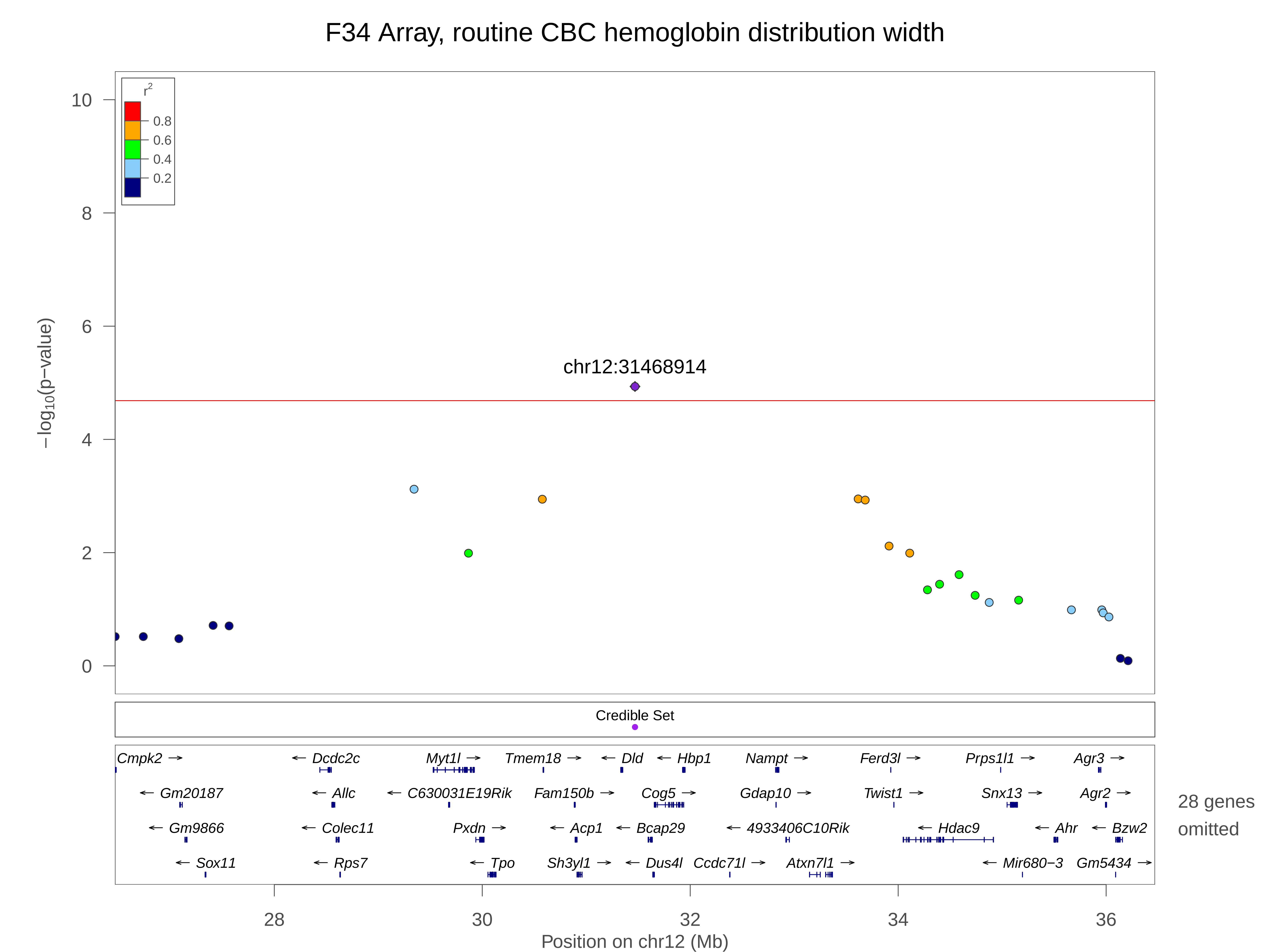
F34 Array, routine CBC hemoglobin distribution width
r2
0.8
0.6
0.4
0.2
10
8
6
4
2
0
−log10(p−value)
chr12:31468914
Credible Set
Cmpk2
Dcdc2c
Myt1l
Tmem18
Dld
Hbp1
Nampt
Ferd3l
Prps1l1
Agr3
Gm20187
Allc
C630031E19Rik
Fam150b
Cog5
Gdap10
Twist1
Snx13
Agr2
Gm9866
Colec11
Pxdn
Acp1
Bcap29
4933406C10Rik
Hdac9
Ahr
Bzw2
Sox11
Rps7
Tpo
Sh3yl1
Dus4l
Ccdc71l
Atxn7l1
Mir680−3
Gm5434
28 genes omitted
28
30
32
34
36
Position on chr12 (Mb)

| | |
|---|---|
| date: | Thu Oct 25 16:44:56 2018 |
| build: | F34Array |
| display range: | chr12:26468914−36468914 [26468914−36468914] |
| hilite range: | 0 − 0 [ 0 − 0 ] |
| reference SNP: | chr12:31468914 |
| number of SNPs plotted: | 26 |
| min P−value: | 1.16E−5 [chr12:31468914] |
| max P−value: | 8.11E−1 [chr12:36210830] |
| omitted Genes: | Rnaseh1, Adi1, Mir6937 |
| omitted Genes: | Trappc12, Tssc1, Sntg2 |
| omitted Genes: | 2310016D03Rik, Fam110c, 6030469F06Rik |
| omitted Genes: | Lamb1, Slc26a3, Cbll1 |
| omitted Genes: | Slc26a4, Gpr22, Prkar2b |
| omitted Genes: | Pik3cg, Sypl, Cdhr3 |
| omitted Genes: | F730043M19Rik, Atxn7l1os2, Efcab10 |
| omitted Genes: | Twistnb, 9130015A21Rik, Tspan13 |
| omitted Genes: | Ankmy2, Lrrc72, Sostdc1 |
| omitted Genes: | Ispd |

Make more plots at http://csg.sph.umich.edu/locuszoom/

Fine−mapping Regions

| snp | chr | pos (Mb) | pp | group | color |
|---|---|---|---|---|---|
| chr12:31468914 | 12 | 31.46891 | 1 | Credible Set | purple |

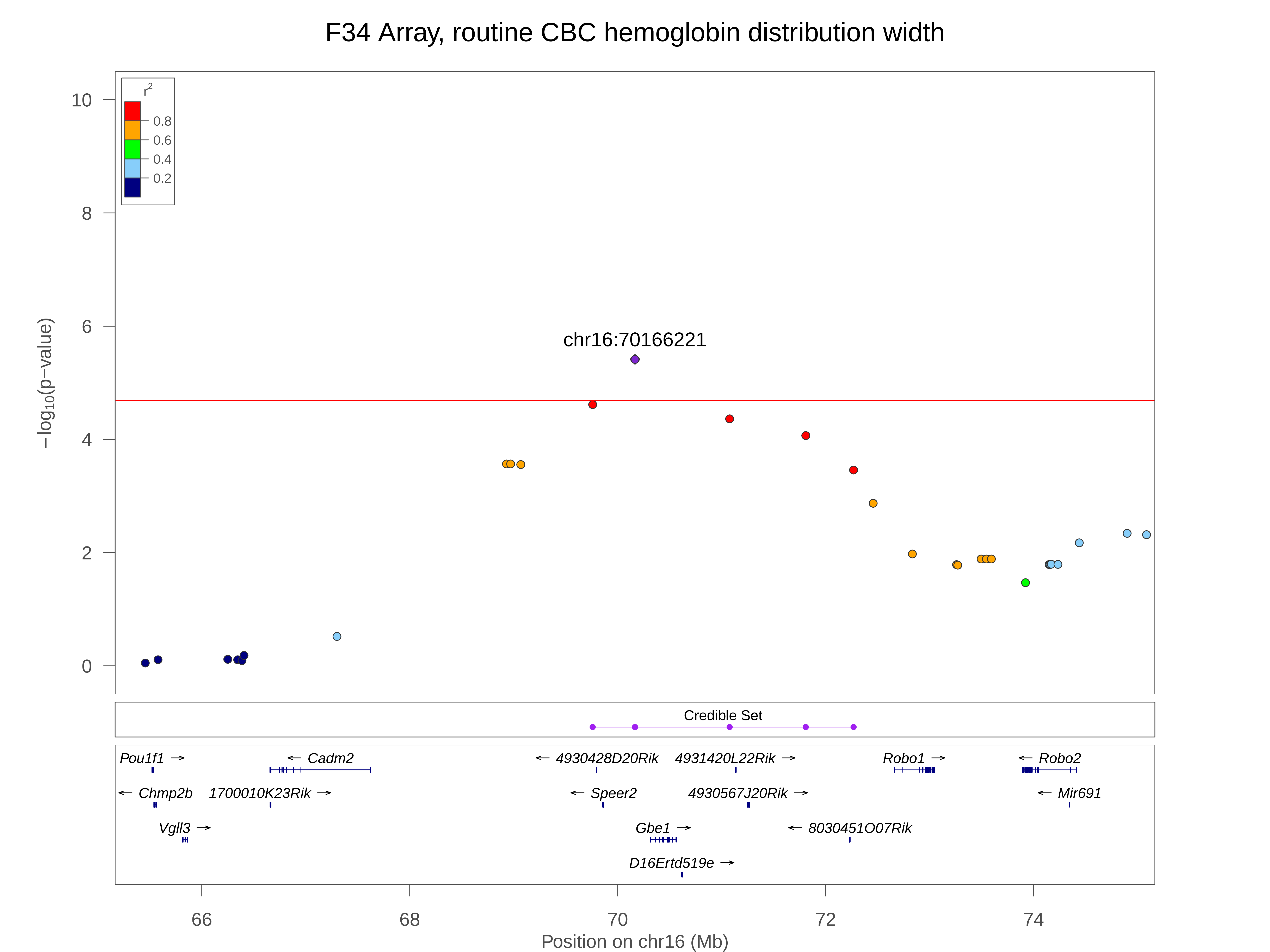

F34 Array, routine CBC hemoglobin distribution width
r2
0.8
0.6
0.4
0.2
10
8
6
4
2
0
−log10(p−value)
chr16:70166221
Credible Set
Pou1f1
Cadm2
4930428D20Rik
4931420L22Rik
Robo1
Robo2
Chmp2b
1700010K23Rik
Speer2
4930567J20Rik
Mir691
Vgll3
Gbe1
8030451O07Rik
D16Ertd519e
66
68
70
72
74
Position on chr16 (Mb)

| | |
|---|---|
| date: | Thu Oct 25 16:45:02 2018 |
| build: | F34Array |
| display range: | chr16:65166221−75166221 [65166221−75166221] |
| hilite range: | 0 − 0 [ 0 − 0 ] |
| reference SNP: | chr16:70166221 |
| number of SNPs plotted: | 31 |
| min P−value: | 3.86E−6 [chr16:70166221] |
| max P−value: | 8.91E−1 [chr16:65456201] |

Make more plots at http://csg.sph.umich.edu/locuszoom/

Fine−mapping Regions

| snp | chr | pos (Mb) | pp | group | color |
|---|---|---|---|---|---|
| chr16:69759143 | 16 | 69.75914 | 0.12904633 | Credible Set | purple |
| chr16:70166221 | 16 | 70.16622 | 0.74694638 | Credible Set | purple |
| chr16:71076501 | 16 | 71.07650 | 0.07445709 | Credible Set | purple |
| chr16:71808739 | 16 | 71.80874 | 0.03909493 | Credible Set | purple |
| chr16:72268105 | 16 | 72.26811 | 0.01045527 | Credible Set | purple |

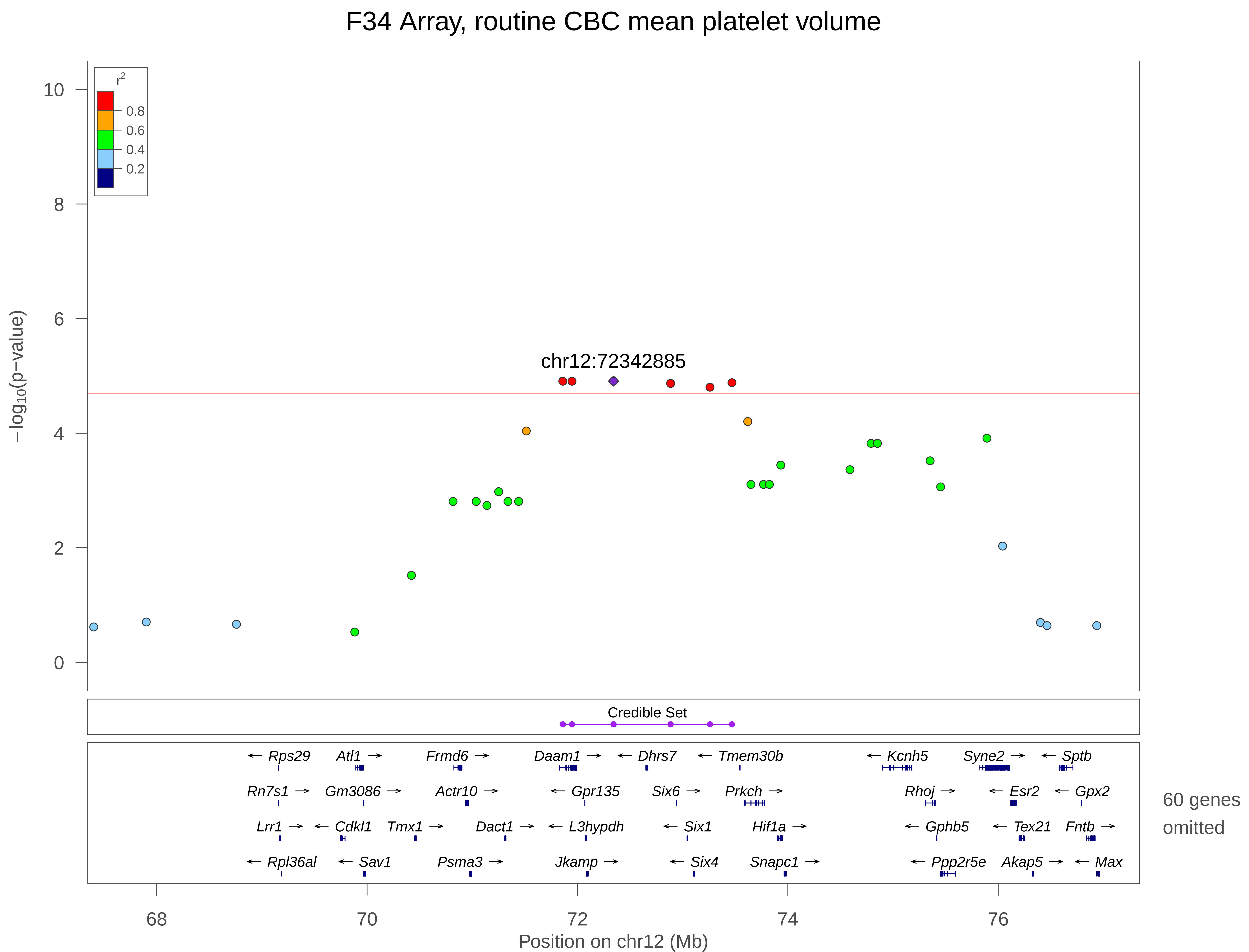
F34 Array, routine CBC mean platelet volume
r2
0.8
0.6
0.4
0.2
10
8
6
4
2
0
−log10(p−value)
chr12:72342885
Credible Set
← Rps29
Atl1 →
Frmd6 →
Daam1 →
← Dhrs7
← Tmem30b
← Kcnh5
Syne2 →
← Sptb
Rn7s1 →
Gm3086 →
Actr10 →
← Gpr135
Six6 →
Prkch →
Rhoj →
← Esr2
← Gpx2
Lrr1 →
← Cdkl1
Tmx1 →
Dact1 →
← L3hypdh
← Six1
Hif1a →
← Gphb5
← Tex21
Fntb →
← Rpl36al
← Sav1
Psma3 →
Jkamp →
← Six4
Snapc1 →
← Ppp2r5e
Akap5 →
← Max
60 genes omitted
68
70
72
74
76
Position on chr12 (Mb)

| | |
|---|---|
| date: | Thu Oct 25 16:45:17 2018 |
| build: | F34Array |
| display range: | chr12:67342885−77342885 [67342885−77342885] |
| hilite range: | 0 − 0 [ 0 − 0 ] |
| reference SNP: | chr12:72342885 |
| number of SNPs plotted: | 34 |
| min P−value: | 1.23E−5 [chr12:72342885] |
| max P−value: | 2.95E−1 [chr12:69883335] |
| omitted Genes: | Mgat2, Dnaaf2, 9330151L19Rik |
| omitted Genes: | Pole2, Klhdc1, Klhdc2 |
| omitted Genes: | Nemf, Rn7s2, Arf6 |
| omitted Genes: | 5830428M24Rik, Vcpkmt, Sos2 |
| omitted Genes: | L2hgdh, Atp5s, Mir681 |
| omitted Genes: | 4930512B01Rik, Map4k5, 4931403G20Rik |
| omitted Genes: | Nin, Abhd12b, Pygl |
| omitted Genes: | F730035M05Rik, Trim9, 3110056K07Rik |
| omitted Genes: | Arid4a, Tomm20l, Timm9 |
| omitted Genes: | 2700049A03Rik, 1700083H02Rik, 4930404H11Rik |
| omitted Genes: | Ccdc175, Rtn1, Lrrc9 |
| omitted Genes: | Pcnx4, Ppm1a, 4930447C04Rik |
| omitted Genes: | Mnat1, Trmt5, Slc38a6 |
| omitted Genes: | D830013O20Rik, 2210039B01Rik, Syt16 |
| omitted Genes: | 1700086L19Rik, Dbpht2, Wdr89 |
| omitted Genes: | Rplp2−ps1, Sgpp1, Mir5101 |
| omitted Genes: | Mthfd1, Zbtb25, Zbtb1 |
| omitted Genes: | Hspa2, Ppp1r36, Gm10451 |
| omitted Genes: | Mir5135, Plekhg3, AI463170 |
| omitted Genes: | Churc1, Rab15, Fut8 |

Make more plots at http://csg.sph.umich.edu/locuszoom/

Fine−mapping Regions

| snp | chr | pos (Mb) | pp | group | color |
|---|---|---|---|---|---|
| chr12:71860639 | 12 | 71.86064 | 0.1766683 | Credible Set | purple |
| chr12:71948169 | 12 | 71.94817 | 0.1766683 | Credible Set | purple |
| chr12:72342885 | 12 | 72.34288 | 0.1773017 | Credible Set | purple |
| chr12:72885260 | 12 | 72.88526 | 0.1621440 | Credible Set | purple |
| chr12:73260123 | 12 | 73.26012 | 0.1405488 | Credible Set | purple |
| chr12:73468910 | 12 | 73.46891 | 0.1666690 | Credible Set | purple |

#### F34 Array, routine WBC differential white blood cell count

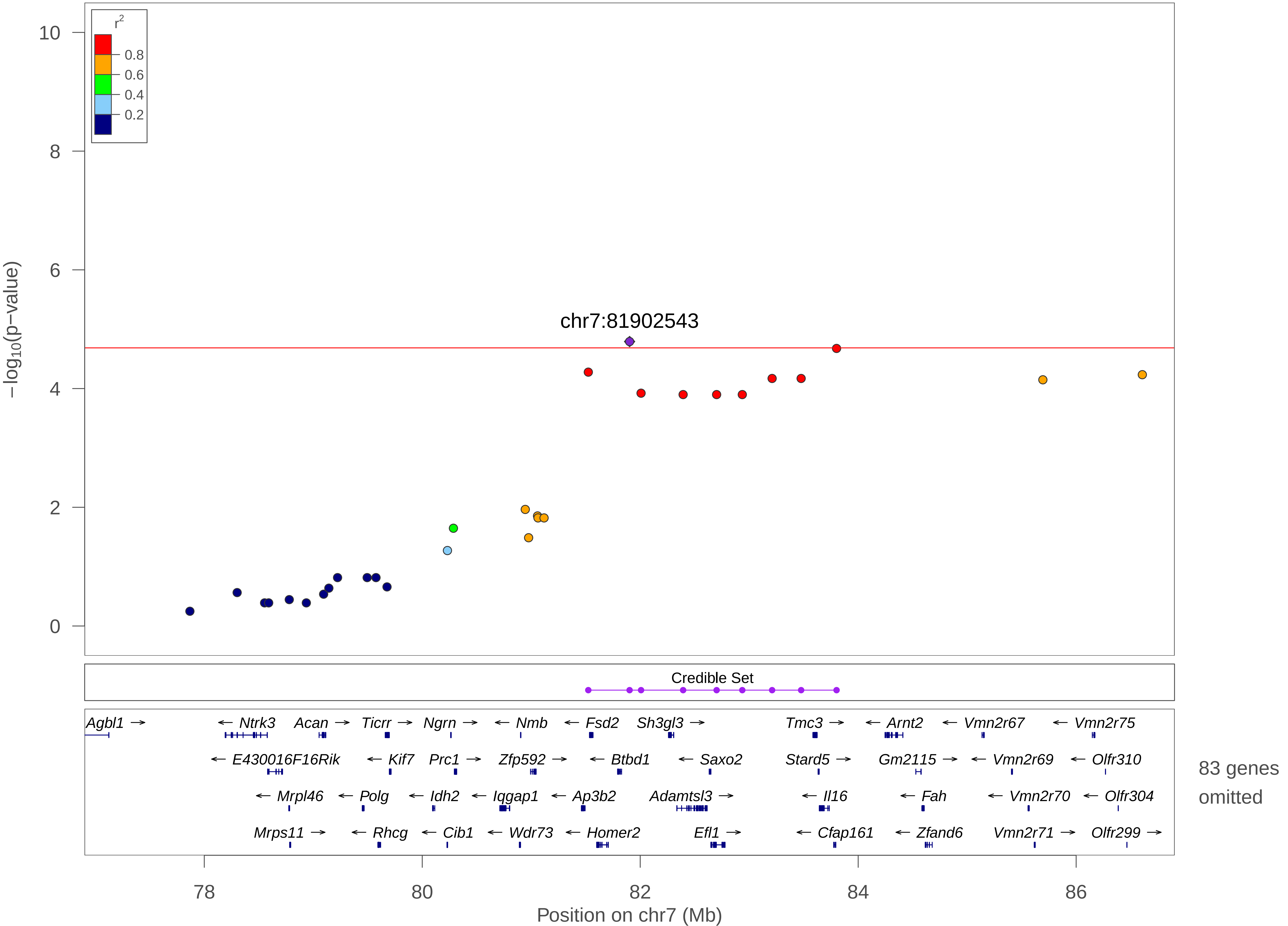

| | |
|---|---|
| date: | Thu Oct 25 16:45:32 2018 |
| build: | F34Array |
| display range: | chr7:76902543−86902543 [76902543−86902543] |
| hilite range: | 0 − 0 [ 0 − 0 ] |
| reference SNP: | chr7:81902543 |
| number of SNPs plotted: | 31 |
| min P−value: | 1.62E−5 [chr7:81902543] |
| max P−value: | 5.64E−1 [chr7:77868097] |
| omitted Genes: | Det1, Mir7−2, Aen |
| omitted Genes: | Isg20, Gm26633, Hapln3 |
| omitted Genes: | Mfge8, Abhd2, Rlbp1 |
| omitted Genes: | Fanci, Mir9−3hg, Mir9−3 |
| omitted Genes: | Plin1, Pex11a, Wdr93 |
| omitted Genes: | Mesp1, Mesp2, Anpep |
| omitted Genes: | Ap3s2, Arpin, Zfp710 |
| omitted Genes: | Gm21057, Sema4b, Gdpgp1 |
| omitted Genes: | Ttll13, Vps33b, Rccd1 |
| omitted Genes: | Unc45a, Hddc3, Man2a2 |
| omitted Genes: | Fes, Furin, Blm |
| omitted Genes: | Crtc3, Gm15880, Zscan2 |
| omitted Genes: | Sec11a, Alpk3, Slc28a1 |
| omitted Genes: | Pde8a, Rps17, Cpeb1 |
| omitted Genes: | BC048679, 2900076A07Rik, Mir1839 |
| omitted Genes: | Whamm, Fam103a1, 3110040N11Rik |
| omitted Genes: | Tm6sf1, Hdgfrp3, Bnc1 |
| omitted Genes: | Gm20744, 4933406J10Rik, Mex3b |
| omitted Genes: | 4933430H16Rik, Mesdc1, Mesdc2 |
| omitted Genes: | Cemip, Abhd17c, 2610206C17Rik |
| omitted Genes: | Olfr291, Olfr290, Vmn2r65 |
| omitted Genes: | Vmn2r66, Vmn2r68, Vmn2r72 |
| omitted Genes: | Vmn2r73, Vmn2r74, Vmn2r76 |
| omitted Genes: | Olfr309, Olfr308, Olfr307 |
| omitted Genes: | Olfr305, Olfr303, Olfr301 |
| omitted Genes: | Olfr298, Olfr297, Olfr295 |
| omitted Genes: | Olfr294, Olfr293, Olfr292 |
| omitted Genes: | Folh1, Vmn2r77 |

Make more plots at http://csg.sph.umich.edu/locuszoom/

Fine−mapping Regions

| snp | chr | pos (Mb) | pp | group | color |
|---|---|---|---|---|---|
| chr7:81523675 | 7 | 81.52367 | 0.10160705 | Credible Set | purple |
| chr7:81902543 | 7 | 81.90254 | 0.31339737 | Credible Set | purple |
| chr7:82007699 | 7 | 82.00770 | 0.04694282 | Credible Set | purple |
| chr7:82393693 | 7 | 82.39369 | 0.04460747 | Credible Set | purple |
| chr7:82701107 | 7 | 82.70111 | 0.04460747 | Credible Set | purple |
| chr7:82936666 | 7 | 82.93667 | 0.04460747 | Credible Set | purple |
| chr7:83210268 | 7 | 83.21027 | 0.08050627 | Credible Set | purple |
| chr7:83476529 | 7 | 83.47653 | 0.08050627 | Credible Set | purple |
| chr7:83801589 | 7 | 83.80159 | 0.24321781 | Credible Set | purple |

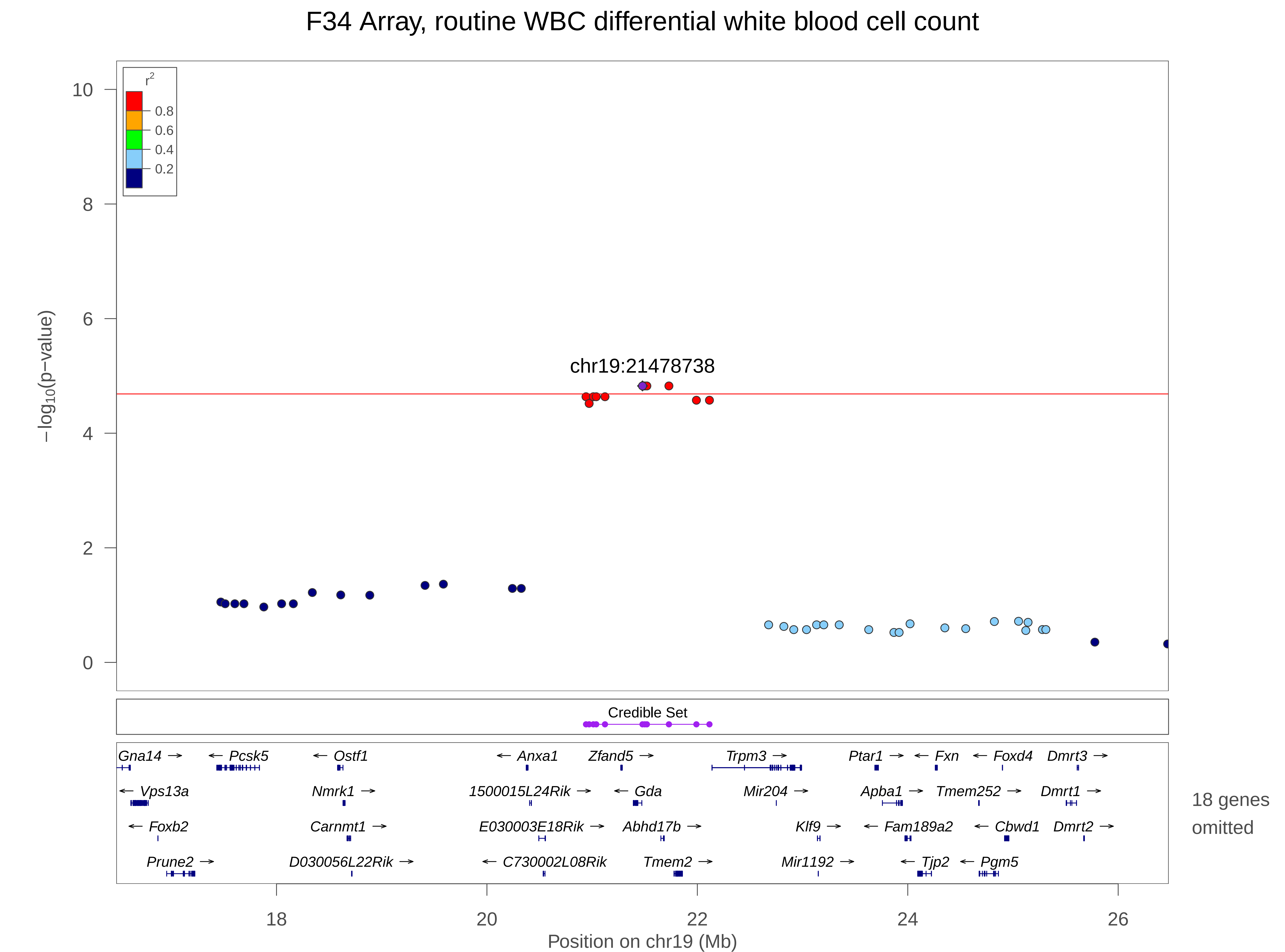

F34 Array, routine WBC differential white blood cell count
r2
0.8
0.6
0.4
0.2
chr19:21478738
−log10(p−value)
Credible Set
Gna14
Pcsk5
Ostf1
Anxa1
Zfand5
Trpm3
Ptar1
Fxn
Foxd4
Dmrt3
Vps13a
Nmrk1
1500015L24Rik
Gda
Mir204
Apba1
Tmem252
Dmrt1
Foxb2
Carnmt1
E030003E18Rik
Abhd17b
Klf9
Fam189a2
Cbwd1
Dmrt2
Prune2
D030056L22Rik
C730002L08Rik
Tmem2
Mir1192
Tjp2
Pgm5
18 genes omitted
Position on chr19 (Mb)

| | |
|---|---|
| date: | Thu Oct 25 16:45:24 2018 |
| build: | F34Array |
| display range: | chr19:16478738−26478738 [16478738−26478738] |
| hilite range: | 0 − 0 [ 0 − 0 ] |
| reference SNP: | chr19:21478738 |
| number of SNPs plotted: | 47 |
| min P−value: | 1.5E−5 [chr19:21478738] |
| max P−value: | 4.76E−1 [chr19:26470094] |
| omitted Genes: | Gcnt1, Rfk, Trpm6 |
| omitted Genes: | Rorb, Aldh1a1, Aldh1a7 |
| omitted Genes: | Tmc1, 4930554I06Rik, 1110059E24Rik |
| omitted Genes: | Smc5, Mamdc2, 1700028P14Rik |
| omitted Genes: | Pip5k1b, Fam122a, Mir3084−1 |
| omitted Genes: | Dock8, Kank1, 2610016A17Rik |

Make more plots at http://csg.sph.umich.edu/locuszoom/

Fine−mapping Regions

| snp | chr | pos (Mb) | pp | group | color |
|---|---|---|---|---|---|
| chr19:20941927 | 19 | 20.94193 | 0.07955138 | Credible Set | purple |
| chr19:20971640 | 19 | 20.97164 | 0.06146381 | Credible Set | purple |
| chr19:21011731 | 19 | 21.01173 | 0.07955138 | Credible Set | purple |
| chr19:21038918 | 19 | 21.03892 | 0.07955138 | Credible Set | purple |
| chr19:21122136 | 19 | 21.12214 | 0.07955138 | Credible Set | purple |
| chr19:21478738 | 19 | 21.47874 | 0.12027742 | Credible Set | purple |
| chr19:21500356 | 19 | 21.50036 | 0.12027742 | Credible Set | purple |
| chr19:21520189 | 19 | 21.52019 | 0.12027742 | Credible Set | purple |
| chr19:21729752 | 19 | 21.72975 | 0.12027742 | Credible Set | purple |
| chr19:21991002 | 19 | 21.99100 | 0.06961049 | Credible Set | purple |
| chr19:22115053 | 19 | 22.11505 | 0.06961049 | Credible Set | purple |

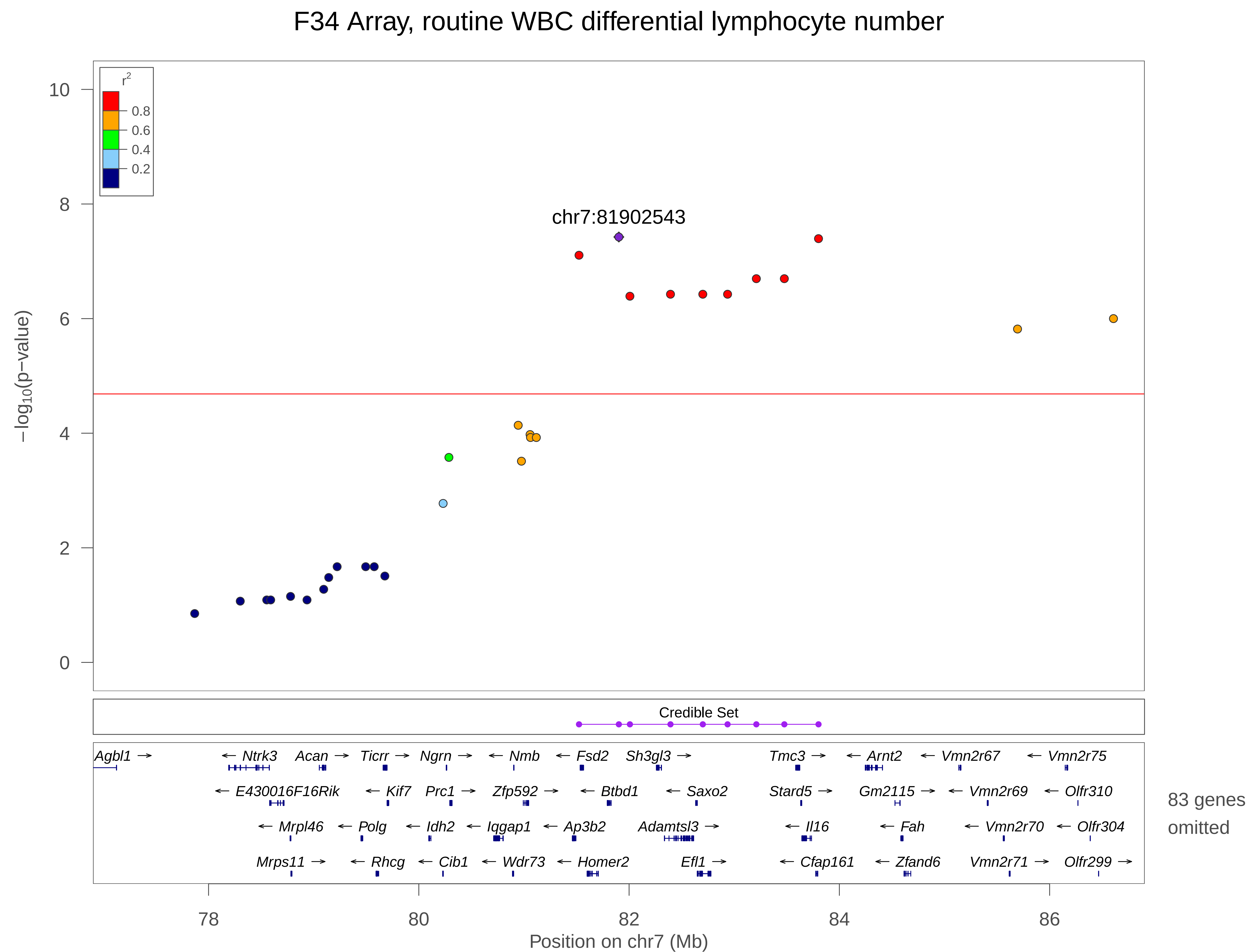
F34 Array, routine WBC differential lymphocyte number
r2
0.8
0.6
0.4
0.2
chr7:81902543
−log10(p−value)
10
8
6
4
2
0
Credible Set
Agbl1
Ntrk3
Acan
Ticrr
Ngrn
Nmb
Fsd2
Sh3gl3
Tmc3
Arnt2
Vmn2r67
Vmn2r75
E430016F16Rik
Kif7
Prc1
Zfp592
Btbd1
Saxo2
Stard5
Gm2115
Vmn2r69
Olfr310
Mrpl46
Polg
Idh2
Iqgap1
Ap3b2
Adamtsl3
Il16
Fah
Vmn2r70
Olfr304
Mrps11
Rhcg
Cib1
Wdr73
Homer2
Efl1
Cfap161
Zfand6
Vmn2r71
Olfr299
83 genes
omitted
78
80
82
84
86
Position on chr7 (Mb)

| | |
|---|---|
| date: | Thu Oct 25 16:45:40 2018 |
| build: | F34Array |
| display range: | chr7:76902543−86902543 [76902543−86902543] |
| hilite range: | 0 − 0 [ 0 − 0 ] |
| reference SNP: | chr7:81902543 |
| number of SNPs plotted: | 31 |
| min P−value: | 3.76E−8 [chr7:81902543] |
| max P−value: | 1.4E−1 [chr7:77868097] |
| omitted Genes: | Det1, Mir7−2, Aen |
| omitted Genes: | Isg20, Gm26633, Hapln3 |
| omitted Genes: | Mfge8, Abhd2, Rlbp1 |
| omitted Genes: | Fanci, Mir9−3hg, Mir9−3 |
| omitted Genes: | Plin1, Pex11a, Wdr93 |
| omitted Genes: | Mesp1, Mesp2, Anpep |
| omitted Genes: | Ap3s2, Arpin, Zfp710 |
| omitted Genes: | Gm21057, Sema4b, Gdpgp1 |
| omitted Genes: | Ttll13, Vps33b, Rccd1 |
| omitted Genes: | Unc45a, Hddc3, Man2a2 |
| omitted Genes: | Fes, Furin, Blm |
| omitted Genes: | Crtc3, Gm15880, Zscan2 |
| omitted Genes: | Sec11a, Alpk3, Slc28a1 |
| omitted Genes: | Pde8a, Rps17, Cpeb1 |
| omitted Genes: | BC048679, 2900076A07Rik, Mir1839 |
| omitted Genes: | Whamm, Fam103a1, 3110040N11Rik |
| omitted Genes: | Tm6sf1, Hdgfrp3, Bnc1 |
| omitted Genes: | Gm20744, 4933406J10Rik, Mex3b |
| omitted Genes: | 4933430H16Rik, Mesdc1, Mesdc2 |
| omitted Genes: | Cemip, Abhd17c, 2610206C17Rik |
| omitted Genes: | Olfr291, Olfr290, Vmn2r65 |
| omitted Genes: | Vmn2r66, Vmn2r68, Vmn2r72 |
| omitted Genes: | Vmn2r73, Vmn2r74, Vmn2r76 |
| omitted Genes: | Olfr309, Olfr308, Olfr307 |
| omitted Genes: | Olfr305, Olfr303, Olfr301 |
| omitted Genes: | Olfr298, Olfr297, Olfr295 |
| omitted Genes: | Olfr294, Olfr293, Olfr292 |
| omitted Genes: | Folh1, Vmn2r77 |

Make more plots at http://csg.sph.umich.edu/locuszoom/

Fine−mapping Regions

| snp | chr | pos (Mb) | pp | group | color |
| --- | --- | --- | --- | --- | --- |
| chr7:81523675 | 7 | 81.52367 | 0.15135341 | Credible Set | purple |
| chr7:81902543 | 7 | 81.90254 | 0.30774554 | Credible Set | purple |
| chr7:82007699 | 7 | 82.00770 | 0.03078917 | Credible Set | purple |
| chr7:82393693 | 7 | 82.39369 | 0.03325479 | Credible Set | purple |
| chr7:82701107 | 7 | 82.70111 | 0.03325479 | Credible Set | purple |
| chr7:82936666 | 7 | 82.93667 | 0.03325479 | Credible Set | purple |
| chr7:83210268 | 7 | 83.21027 | 0.06082235 | Credible Set | purple |
| chr7:83476529 | 7 | 83.47653 | 0.06082235 | Credible Set | purple |
| chr7:83801589 | 7 | 83.80159 | 0.28870281 | Credible Set | purple |

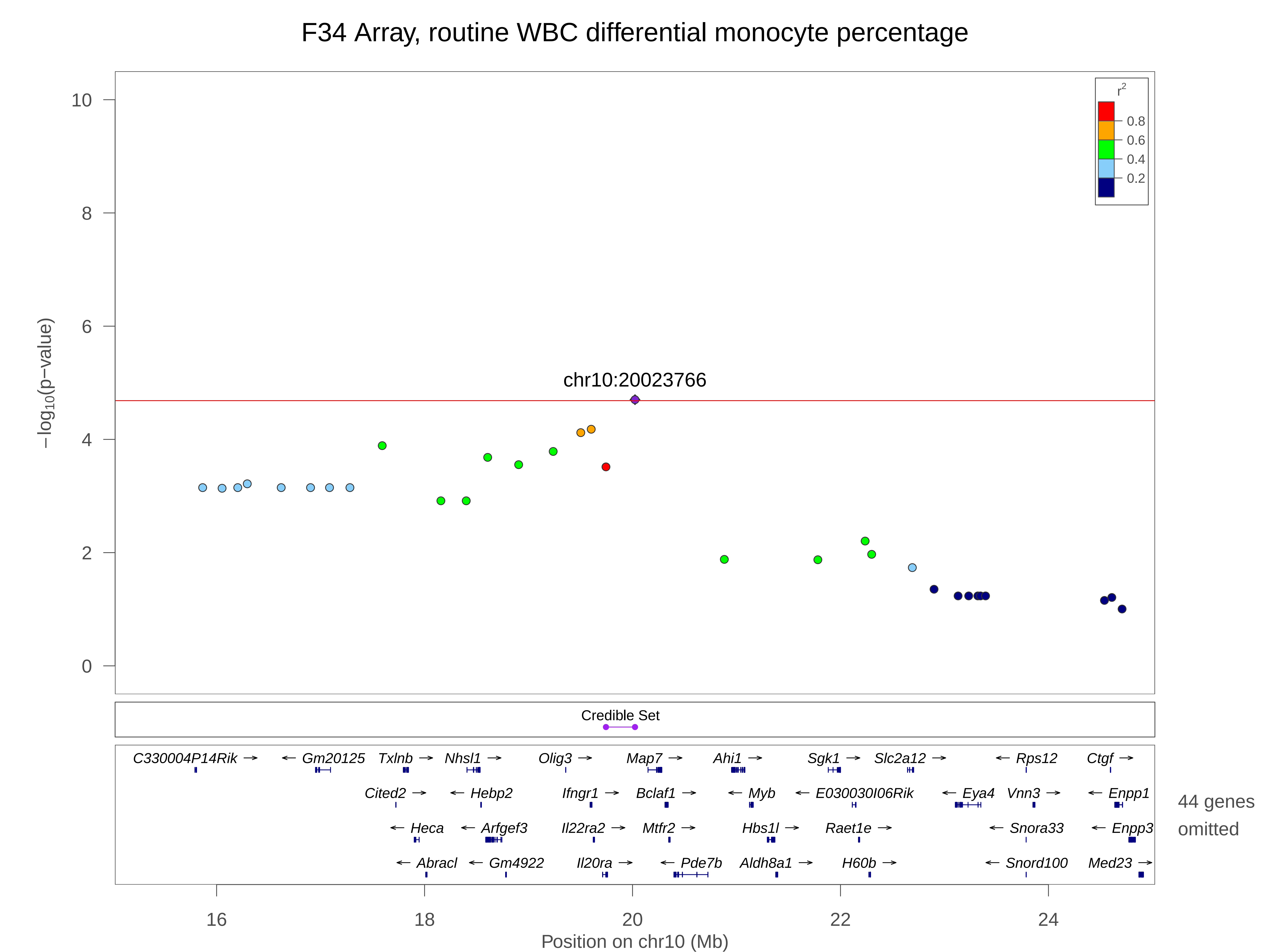
F34 Array, routine WBC differential monocyte percentage
−log10(p−value)
chr10:20023766
r2
0.8
0.6
0.4
0.2
Credible Set
C330004P14Rik →
← Gm20125
Txlnb →
Nhsl1 →
Olig3 →
Map7 →
Ahi1 →
Sgk1 →
Slc2a12 →
← Rps12
Ctgf →
Cited2 →
← Hebp2
Ifngr1 →
Bclaf1 →
← Myb
← E030030I06Rik
← Eya4
Vnn3 →
← Enpp1
← Heca
← Arfgef3
Il22ra2 →
Mtfr2 →
Hbs1l →
Raet1e →
← Snora33
← Enpp3
← Abracl
← Gm4922
Il20ra →
← Pde7b
Aldh8a1 →
H60b →
← Snord100
Med23 →
44 genes omitted
Position on chr10 (Mb)

| | |
|---|---|
| date: | Thu Oct 25 16:45:48 2018 |
| build: | F34Array |
| display range: | chr10:15023766−25023766 [15023766−25023766] |
| hilite range: | 0 − 0 [ 0 − 0 ] |
| reference SNP: | chr10:20023766 |
| number of SNPs plotted: | 33 |
| min P−value: | 1.99E−5 [chr10:20023766] |
| max P−value: | 9.9E−2 [chr10:24708091] |
| omitted Genes: | Reps1, Ect2l, Ccdc28a |
| omitted Genes: | 4930444F02Rik, Perp, Tnfaip3 |
| omitted Genes: | Gm20139, Slc35d3, Pex7 |
| omitted Genes: | Map3k5, 4930405J17Rik, Rpl32l |
| omitted Genes: | Mir6413, 4930455C13Rik, 1700020N01Rik |
| omitted Genes: | Gm5420, 4930444G20Rik, C920009B18Rik |
| omitted Genes: | Raet1d, Gm10825, Tbpl1 |
| omitted Genes: | Mir7663, Tcf21, Slc18b1 |
| omitted Genes: | Vnn1, Taar1, Taar2 |
| omitted Genes: | Taar3, Taar4, Taar5 |
| omitted Genes: | Taar6, Taar7a, Taar7b |
| omitted Genes: | Taar7d, Taar7e, Taar7f |
| omitted Genes: | Taar8a, Taar8b, Taar8c |
| omitted Genes: | Taar9, Stx7, Moxd1 |
| omitted Genes: | Mir6905, Arg1 |

Make more plots at http://csg.sph.umich.edu/locuszoom/

Fine−mapping Regions

| snp | chr | pos (Mb) | pp | group | color |
|---|---|---|---|---|---|
| chr10:19744511 | 10 | 19.74451 | 0.0702341 | Credible Set | purple |
| chr10:20023766 | 10 | 20.02377 | 0.9297659 | Credible Set | purple |

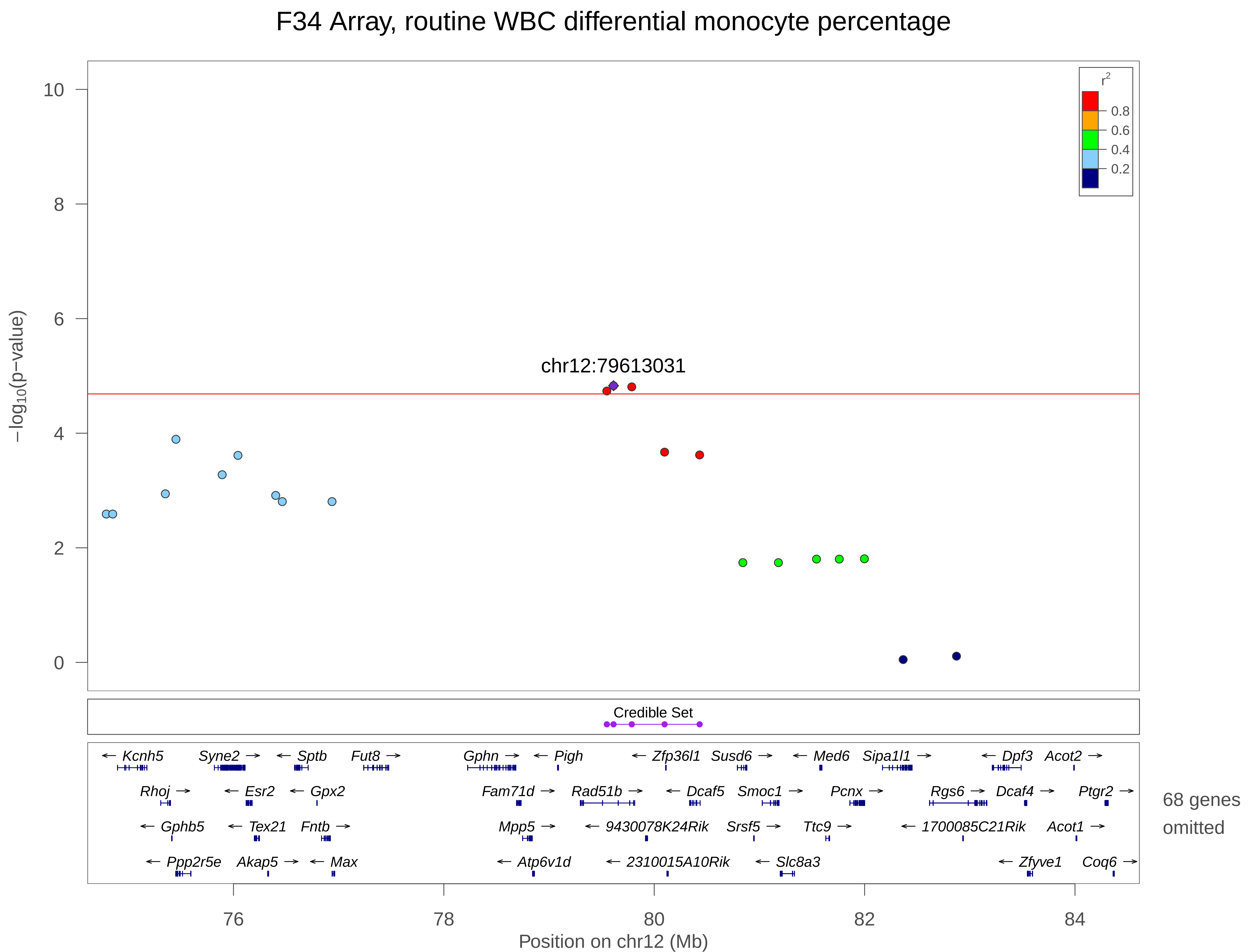
F34 Array, routine WBC differential monocyte percentage
−log10(p−value)
chr12:79613031
r2
0.8
0.6
0.4
0.2
Credible Set
Kcnh5
Syne2
Sptb
Fut8
Gphn
Pigh
Zfp36l1
Susd6
Med6
Sipa1l1
Dpf3
Acot2
Rhoj
Esr2
Gpx2
Fam71d
Rad51b
Dcaf5
Smoc1
Pcnx
Rgs6
Dcaf4
Ptgr2
Gphb5
Tex21
Fntb
Mpp5
9430078K24Rik
Srsf5
Ttc9
1700085C21Rik
Acot1
Ppp2r5e
Akap5
Max
Atp6v1d
2310015A10Rik
Slc8a3
Zfyve1
Coq6
68 genes omitted
Position on chr12 (Mb)

| | |
|---|---|
| date: | Thu Oct 25 16:45:56 2018 |
| build: | F34Array |
| display range: | chr12:74613031−84613031 [74613031−84613031] |
| hilite range: | 0 − 0 [ 0 − 0 ] |
| reference SNP: | chr12:79613031 |
| number of SNPs plotted: | 22 |
| min P−value: | 1.48E−5 [chr12:79613031] |
| max P−value: | 8.93E−1 [chr12:82366650] |
| omitted Genes: | Wdr89, Rplp2−ps1, Sgpp1 |
| omitted Genes: | Mir5101, Mthfd1, Zbtb25 |
| omitted Genes: | Zbtb1, Hspa2, Ppp1r36 |
| omitted Genes: | Gm10451, Mir5135, Plekhg3 |
| omitted Genes: | AI463170, Churc1, Rab15 |
| omitted Genes: | Eif2s1, Plek2, Tmem229b |
| omitted Genes: | Plekhh1, 9230116L04Rik, Arg2 |
| omitted Genes: | Vti1b, Rdh11, Rdh12 |
| omitted Genes: | Zfyve26, Actn1, Mir1843a |
| omitted Genes: | Scarna3b, Exd2, 2310002D06Rik |
| omitted Genes: | Galnt16, Erh, Slc39a9 |
| omitted Genes: | Plekhd1os, Plekhd1, Ccdc177 |
| omitted Genes: | 1700052I22Rik, Gm20337, Slc10a1 |
| omitted Genes: | Mir3067, Cox16, Synj2bp−cox16 |
| omitted Genes: | Gm4787, Adam4, Synj2bp |
| omitted Genes: | Adam21, Map3k9, Rbm25 |
| omitted Genes: | Psen1, Papln, Numb |
| omitted Genes: | 2410016O06Rik, Acot4, Acot3 |
| omitted Genes: | Acot5, Acot6, Dnal1 |
| omitted Genes: | Pnma1, Elmsan1, Zfp410 |
| omitted Genes: | Fam161b, Entpd5, Bbof1 |
| omitted Genes: | Rnf113a2, Aldh6a1, Lin52 |
| omitted Genes: | Vsx2, Abcd4 |

Make more plots at http://csg.sph.umich.edu/locuszoom/

Fine−mapping Regions

| snp | chr | pos (Mb) | pp | group | color |
|---|---|---|---|---|---|
| chr12:79549412 | 12 | 79.54941 | 0.27927771 | Credible Set | purple |
| chr12:79613031 | 12 | 79.61303 | 0.34193524 | Credible Set | purple |
| chr12:79786451 | 12 | 79.78645 | 0.32692496 | Credible Set | purple |
| chr12:80098407 | 12 | 80.09841 | 0.02731771 | Credible Set | purple |
| chr12:80431588 | 12 | 80.43159 | 0.02454438 | Credible Set | purple |

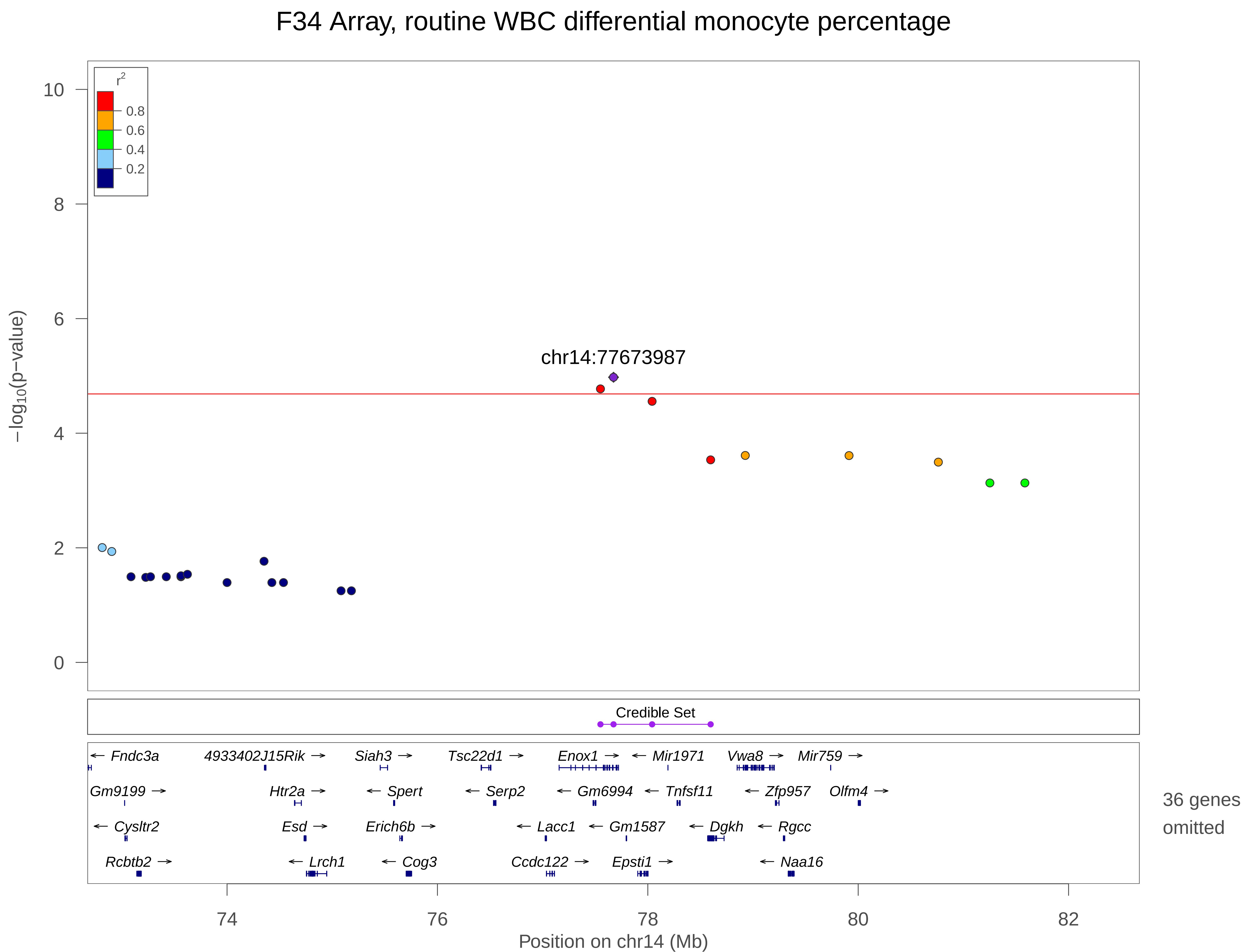
F34 Array, routine WBC differential monocyte percentage
r2
0.8
0.6
0.4
0.2
10
8
6
4
2
0
−log10(p−value)
chr14:77673987
Credible Set
Fndc3a
4933402J15Rik
Siah3
Tsc22d1
Enox1
Mir1971
Vwa8
Mir759
Gm9199
Htr2a
Spert
Serp2
Gm6994
Tnfsf11
Zfp957
Olfm4
Cysltr2
Esd
Erich6b
Lacc1
Gm1587
Dgkh
Rgcc
Rcbtb2
Lrch1
Cog3
Ccdc122
Epsti1
Naa16
36 genes omitted
74
76
78
80
82
Position on chr14 (Mb)

| | |
|---|---|
| date: | Thu Oct 25 16:46:03 2018 |
| build: | F34Array |
| display range: | chr14:72673987−82673987 [72673987−82673987] |
| hilite range: | 0 − 0 [ 0 − 0 ] |
| reference SNP: | chr14:77673987 |
| number of SNPs plotted: | 25 |
| min P−value: | 1.06E−5 [chr14:77673987] |
| max P−value: | 5.62E−2 [chr14:75083773] |
| omitted Genes: | Rb1, Mir687, Lpar6 |
| omitted Genes: | Itm2b, Med4, Nudt15 |
| omitted Genes: | Sucla2, Gm4278, 5031414D18Rik |
| omitted Genes: | Lrrc63, Lcp1, Cpb2 |
| omitted Genes: | Zc3h13, Slc25a30, Gm4285 |
| omitted Genes: | Tpt1, Snora31, Gtf2f2 |
| omitted Genes: | Kctd4, Gpalpp1, Nufip1 |
| omitted Genes: | 4930444M15Rik, 1700108F19Rik, Dnajc15 |
| omitted Genes: | Fam216b, Akap11, Mtrf1 |
| omitted Genes: | Kbtbd7, Kbtbd6, Wbp4 |
| omitted Genes: | Elf1, Sugt1, Lect1 |
| omitted Genes: | 4930452G13Rik, Pcdh8, Gm10845 |

Fine−mapping Regions

| snp | chr | pos (Mb) | pp | group | color |
|---|---|---|---|---|---|
| chr14:77549688 | 14 | 77.54969 | 0.30787002 | Credible Set | purple |
| chr14:77673987 | 14 | 77.67399 | 0.48026350 | Credible Set | purple |
| chr14:78040815 | 14 | 78.04081 | 0.19112811 | Credible Set | purple |
| chr14:78597230 | 14 | 78.59723 | 0.02073838 | Credible Set | purple |

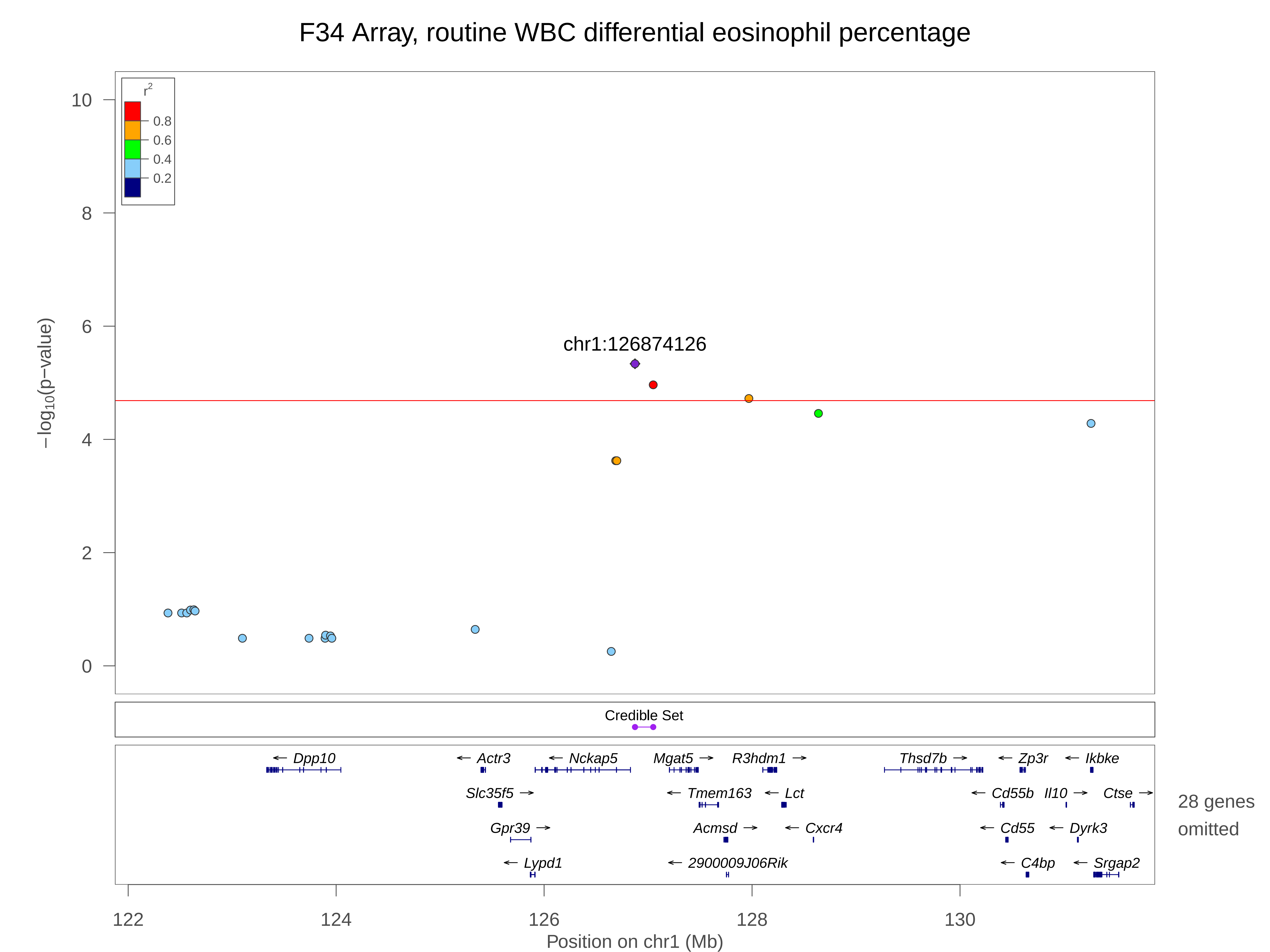

F34 Array, routine WBC differential eosinophil percentage
r2
0.8
0.6
0.4
0.2
10
8
6
4
2
0
−log10(p−value)
chr1:126874126
Credible Set
← Dpp10
← Actr3
← Nckap5
Mgat5 →
R3hdm1 →
Thsd7b →
← Zp3r
← Ikbke
Slc35f5 →
← Tmem163
← Lct
← Cd55b
Il10 →
Ctse →
Gpr39 →
Acmsd →
← Cxcr4
← Cd55
← Dyrk3
← Lypd1
← 2900009J06Rik
← C4bp
← Srgap2
28 genes omitted
122
124
126
128
130
Position on chr1 (Mb)

| | |
|---|---|
| date: | Thu Oct 25 16:46:17 2018 |
| build: | F34Array |
| display range: | chr1:121874126−131874126 [121874126−131874126] |
| hilite range: | 0 − 0 [ 0 − 0 ] |
| reference SNP: | chr1:126874126 |
| number of SNPs plotted: | 22 |
| min P−value: | 4.61E−6 [chr1:126874126] |
| max P−value: | 5.56E−1 [chr1:126645886] |
| omitted Genes: | Ccnt2, Map3k19, Rab3gap1 |
| omitted Genes: | Zranb3, Mir128−1, Ubxn4 |
| omitted Genes: | Mcm6, Dars, C4bp−ps1 |
| omitted Genes: | Pfkfb2, Yod1, AA986860 |
| omitted Genes: | Fcamr, Pigr, Fcmr |
| omitted Genes: | Il24, Il20, Il19 |
| omitted Genes: | Mapkapk2, Eif2d, Rassf5 |
| omitted Genes: | Fam72a, Avpr1b, Rab7b |
| omitted Genes: | Slc26a9, Pm20d1, Slc41a1 |
| omitted Genes: | Rab29 |

Make more plots at http://csg.sph.umich.edu/locuszoom/

Fine−mapping Regions

| snp | chr | pos (Mb) | pp | group | color |
|---|---|---|---|---|---|
| chr1:126874126 | 1 | 126.8741 | 0.694067 | Credible Set | purple |
| chr1:127049404 | 1 | 127.0494 | 0.305933 | Credible Set | purple |

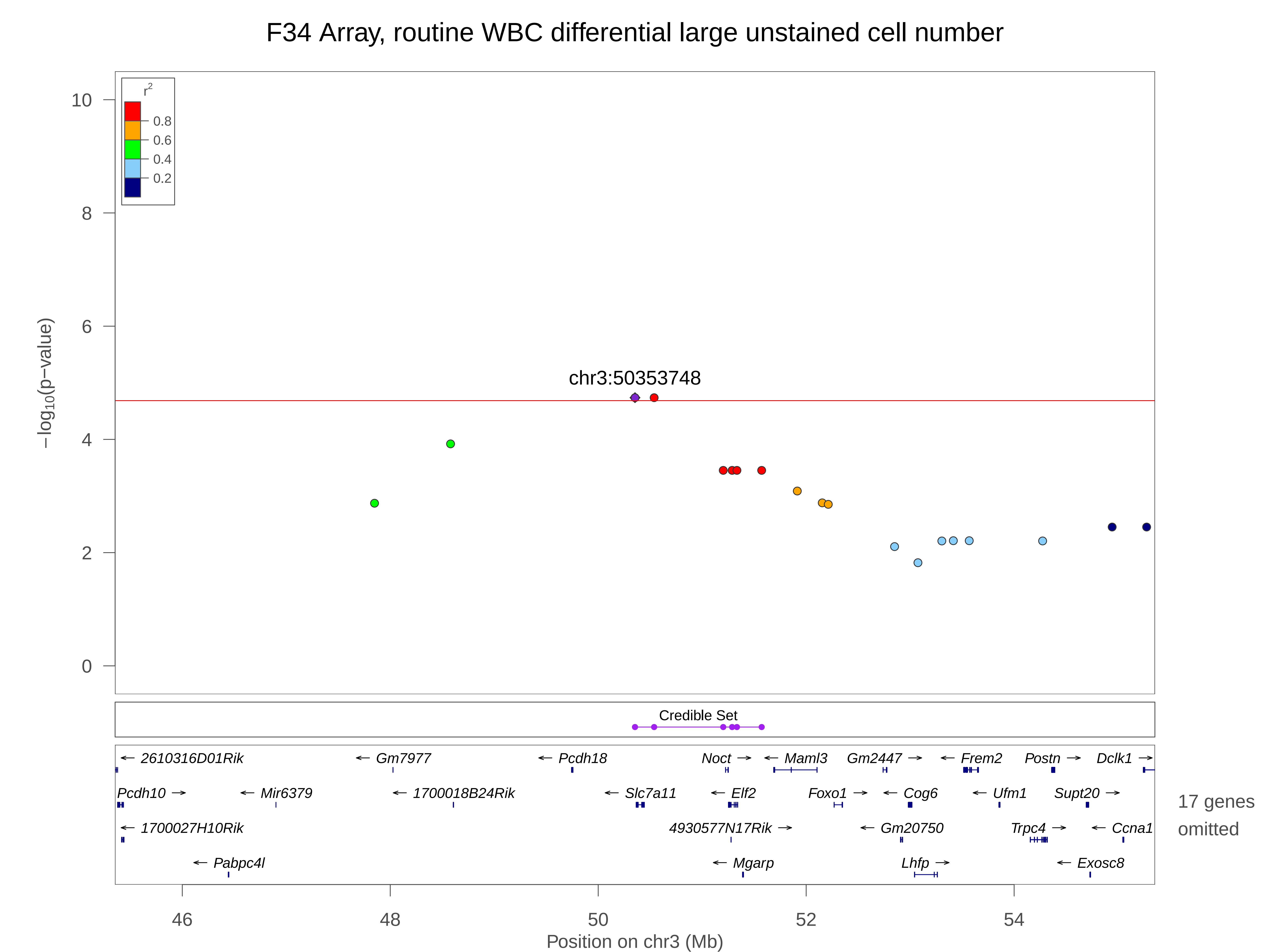
F34 Array, routine WBC differential large unstained cell number
r2
0.8
0.6
0.4
0.2
10
8
6
4
2
0
−log10(p−value)
chr3:50353748
Credible Set
← 2610316D01Rik
← Gm7977
← Pcdh18
Noct →
← Maml3
Gm2447 →
← Frem2
Postn →
Dclk1 →
Pcdh10 →
← Mir6379
← 1700018B24Rik
← Slc7a11
← Elf2
Foxo1 →
← Cog6
← Ufm1
Supt20 →
← 1700027H10Rik
4930577N17Rik →
← Gm20750
Trpc4 →
← Ccna1
← Pabpc4l
← Mgarp
Lhfp →
← Exosc8
17 genes omitted
46
48
50
52
54
Position on chr3 (Mb)

| | |
|---|---|
| date: | Thu Oct 25 16:46:24 2018 |
| build: | F34Array |
| display range: | chr3:45353748−55353748 [45353748−55353748] |
| hilite range: | 0 − 0 [ 0 − 0 ] |
| reference SNP: | chr3:50353748 |
| number of SNPs plotted: | 20 |
| min P−value: | 1.83E−5 [chr3:50353748] |
| max P−value: | 1.51E−2 [chr3:53074908] |
| omitted Genes: | Ndufc1, Naa15, Rab33b |
| omitted Genes: | Setd7, 5031434O11Rik, Mgst2 |
| omitted Genes: | Nhlrc3, Proser1, Stoml3 |
| omitted Genes: | Alg5, Smad9, Rfxap |
| omitted Genes: | Sertm1, 4931419H13Rik, Spg20 |
| omitted Genes: | Ccdc169, Sohlh2 |

Make more plots at http://csg.sph.umich.edu/locuszoom/

Fine−mapping Regions

| snp | chr | pos (Mb) | pp | group | color |
|---|---|---|---|---|---|
| chr3:50353748 | 3 | 50.35375 | 0.44546368 | Credible Set | purple |
| chr3:50537878 | 3 | 50.53788 | 0.44546368 | Credible Set | purple |
| chr3:51202322 | 3 | 51.20232 | 0.02726816 | Credible Set | purple |
| chr3:51287862 | 3 | 51.28786 | 0.02726816 | Credible Set | purple |
| chr3:51333971 | 3 | 51.33397 | 0.02726816 | Credible Set | purple |
| chr3:51571920 | 3 | 51.57192 | 0.02726816 | Credible Set | purple |

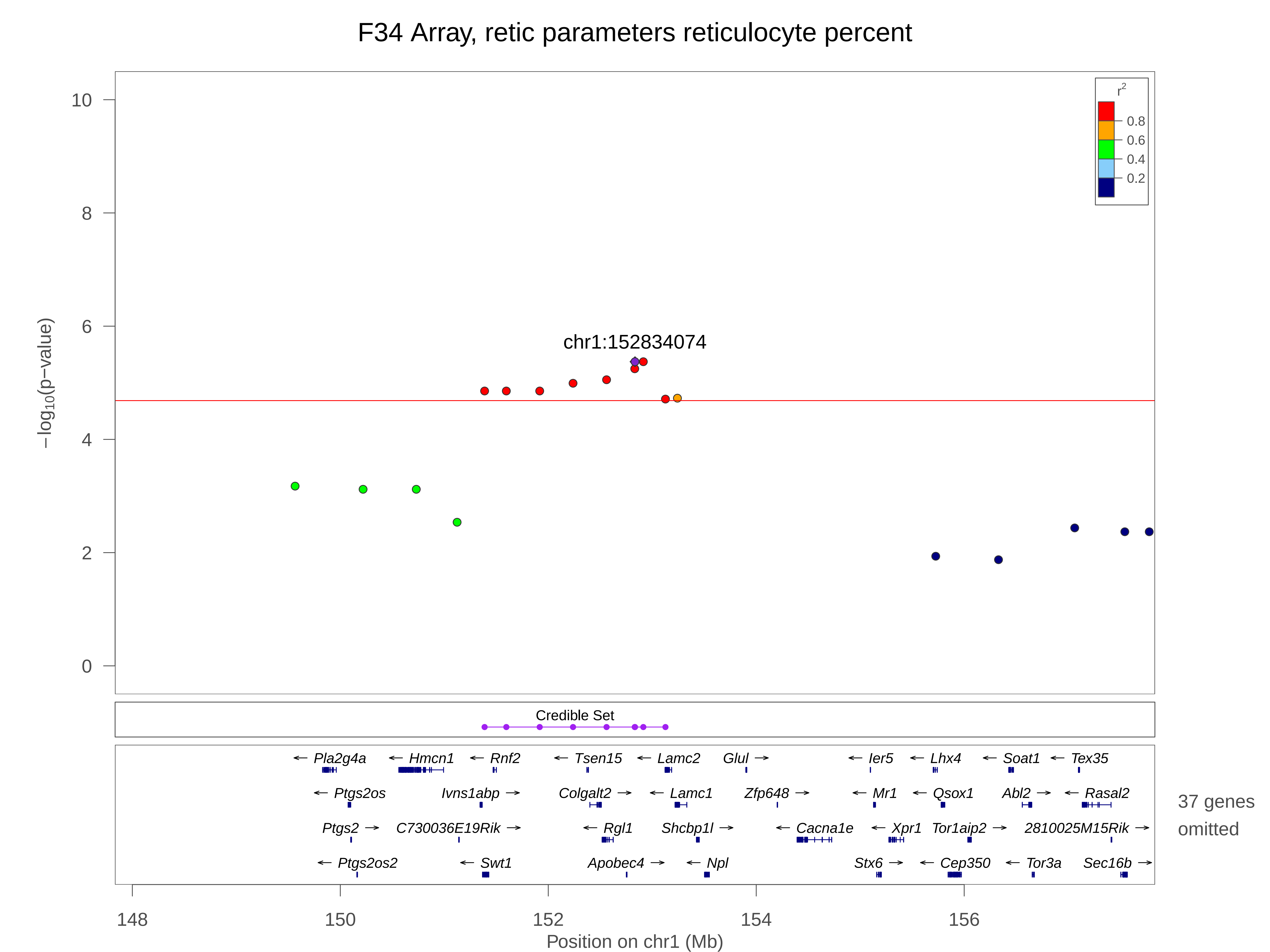

F34 Array, retic parameters reticulocyte percent
−log10(p−value)
10
8
6
4
2
0
r2
0.8
0.6
0.4
0.2
chr1:152834074
Credible Set
← Pla2g4a
← Hmcn1
← Rnf2
← Tsen15
← Lamc2
Glul →
← Ier5
← Lhx4
← Soat1
← Tex35
← Ptgs2os
Ivns1abp →
Colgalt2 →
← Lamc1
Zfp648 →
← Mr1
← Qsox1
Abl2 →
← Rasal2
Ptgs2 →
C730036E19Rik →
← Rgl1
Shcbp1l →
← Cacna1e
← Xpr1
Tor1aip2 →
2810025M15Rik →
← Ptgs2os2
← Swt1
Apobec4 →
← Npl
Stx6 →
← Cep350
← Tor3a
Sec16b →
37 genes omitted
148
150
152
154
156
Position on chr1 (Mb)

| | |
|---|---|
| date: | Thu Oct 25 16:46:31 2018 |
| build: | F34Array |
| display range: | chr1:147834074−157834074 [147834074−157834074] |
| hilite range: | 0 − 0 [ 0 − 0 ] |
| reference SNP: | chr1:152834074 |
| number of SNPs plotted: | 20 |
| min P−value: | 4.24E−6 [chr1:152834074] |
| max P−value: | 1.33E−2 [chr1:156329965] |
| omitted Genes: | Pdc, 2310030A07Rik, BC003331 |
| omitted Genes: | Tpr, Prg4, Trmt1l |
| omitted Genes: | Mir7682, Fam129a, Edem3 |
| omitted Genes: | 1700025G04Rik, Arpc5, Ncf2 |
| omitted Genes: | Smg7, Nmnat2, E330020D12Rik |
| omitted Genes: | Dhx9, Rgs8, C230024C17Rik |
| omitted Genes: | Rgs16, Rnasel, Rgsl1 |
| omitted Genes: | Teddm2, Teddm1b, Teddm1a |
| omitted Genes: | Mir8114, 4930452N14Rik, 4930532M18Rik |
| omitted Genes: | Gm5532, Acbd6, Tor1aip1 |
| omitted Genes: | Fam163a, Tdrd5, Nphs2 |
| omitted Genes: | Fam20b, Ralgps2, Angptl1 |
| omitted Genes: | BC026585 |

Make more plots at http://csg.sph.umich.edu/locuszoom/

Fine−mapping Regions

| snp | chr | pos (Mb) | pp | group | color |
|---|---|---|---|---|---|
| chr1:151387651 | 1 | 151.3877 | 0.06561180 | Credible Set | purple |
| chr1:151596615 | 1 | 151.5966 | 0.06561180 | Credible Set | purple |
| chr1:151917608 | 1 | 151.9176 | 0.06561180 | Credible Set | purple |
| chr1:152238182 | 1 | 152.2382 | 0.08858970 | Credible Set | purple |
| chr1:152560587 | 1 | 152.5606 | 0.10155521 | Credible Set | purple |
| chr1:152831634 | 1 | 152.8316 | 0.15553687 | Credible Set | purple |
| chr1:152834074 | 1 | 152.8341 | 0.20476992 | Credible Set | purple |
| chr1:152914150 | 1 | 152.9142 | 0.20476992 | Credible Set | purple |
| chr1:153126949 | 1 | 153.1269 | 0.04794298 | Credible Set | purple |

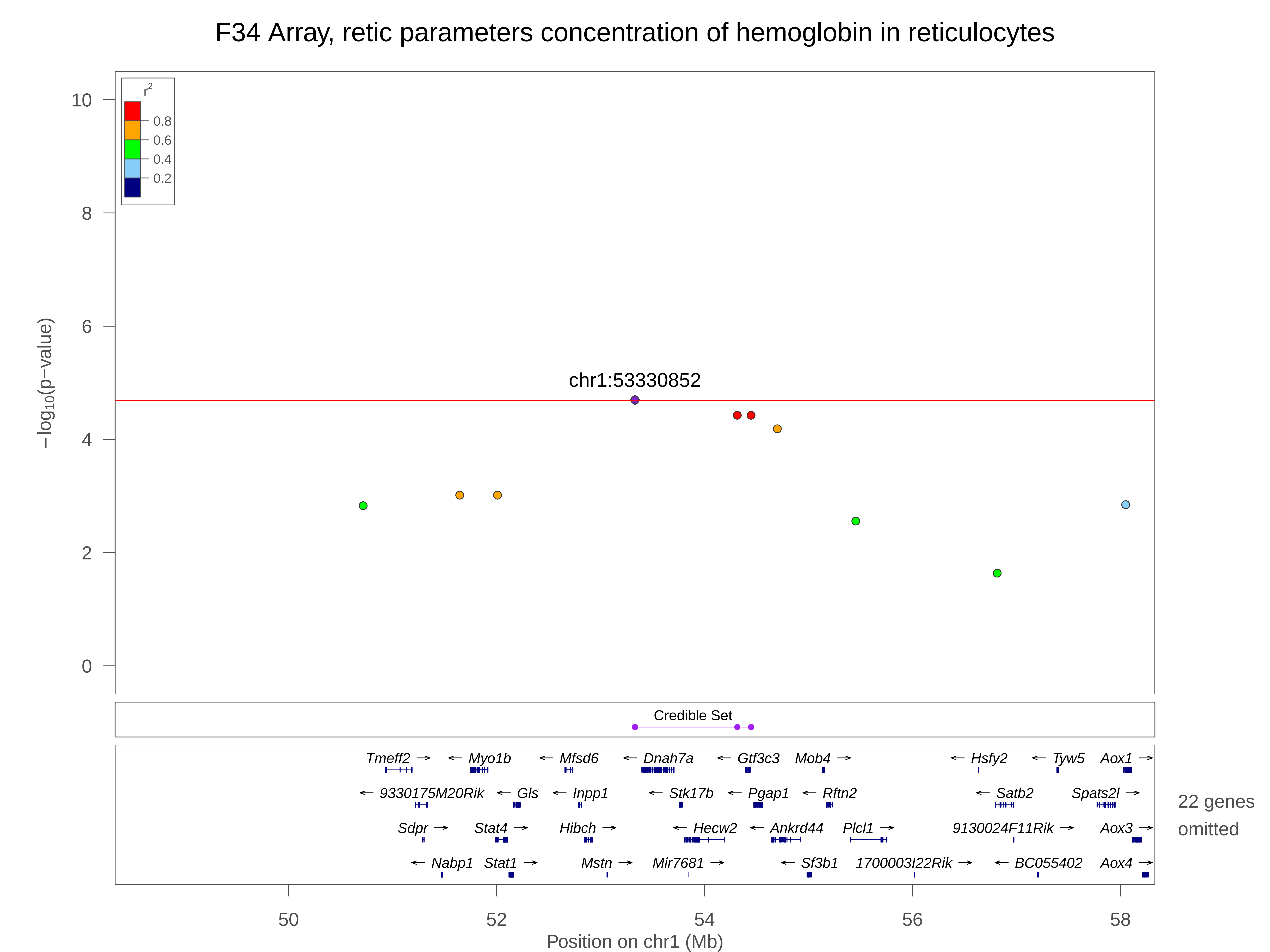
F34 Array, retic parameters concentration of hemoglobin in reticulocytes
r2
0.8
0.6
0.4
0.2
−log10(p−value)
chr1:53330852
Credible Set
Tmeff2
Myo1b
Mfsd6
Dnah7a
Gtf3c3
Mob4
Hsfy2
Tyw5
Aox1
9330175M20Rik
Gls
Inpp1
Stk17b
Pgap1
Rftn2
Satb2
Spats2l
Sdpr
Stat4
Hibch
Hecw2
Ankrd44
Plcl1
9130024F11Rik
Aox3
Nabp1
Stat1
Mstn
Mir7681
Sf3b1
1700003I22Rik
BC055402
Aox4
22 genes omitted
Position on chr1 (Mb)

| | |
|---|---|
| date: | Thu Oct 25 16:46:38 2018 |
| build: | F34Array |
| display range: | chr1:48330852−58330852 [48330852−58330852] |
| hilite range: | 0 − 0 [ 0 − 0 ] |
| reference SNP: | chr1:53330852 |
| number of SNPs plotted: | 11 |
| min P−value: | 2.01E−5 [chr1:53330852] |
| max P−value: | 2.3E−2 [chr1:56814006] |
| omitted Genes: | Nab1, Tmem194b, 1700019D03Rik |
| omitted Genes: | 1700019A02Rik, Pms1, Ormdl1 |
| omitted Genes: | Osgepl1, Asnsd1, Ccdc150 |
| omitted Genes: | Coq10b, Hspd1, Hspe1 |
| omitted Genes: | Mars2, Boll, Gm6644 |
| omitted Genes: | 9130227L01Rik, 4930558J18Rik, 1700066M21Rik |
| omitted Genes: | 9430016H08Rik, Kctd18, Sgol2a |
| omitted Genes: | Aox2 |

Make more plots at http://csg.sph.umich.edu/locuszoom/

Fine−mapping Regions

| snp | chr | pos (Mb) | pp | group | color |
|---|---|---|---|---|---|
| chr1:53330852 | 1 | 53.33085 | 0.4747306 | Credible Set | purple |
| chr1:54314429 | 1 | 54.31443 | 0.2626347 | Credible Set | purple |
| chr1:54447274 | 1 | 54.44727 | 0.2626347 | Credible Set | purple |

F34 Array, retic parameters concentration of hemoglobin in reticulocytes

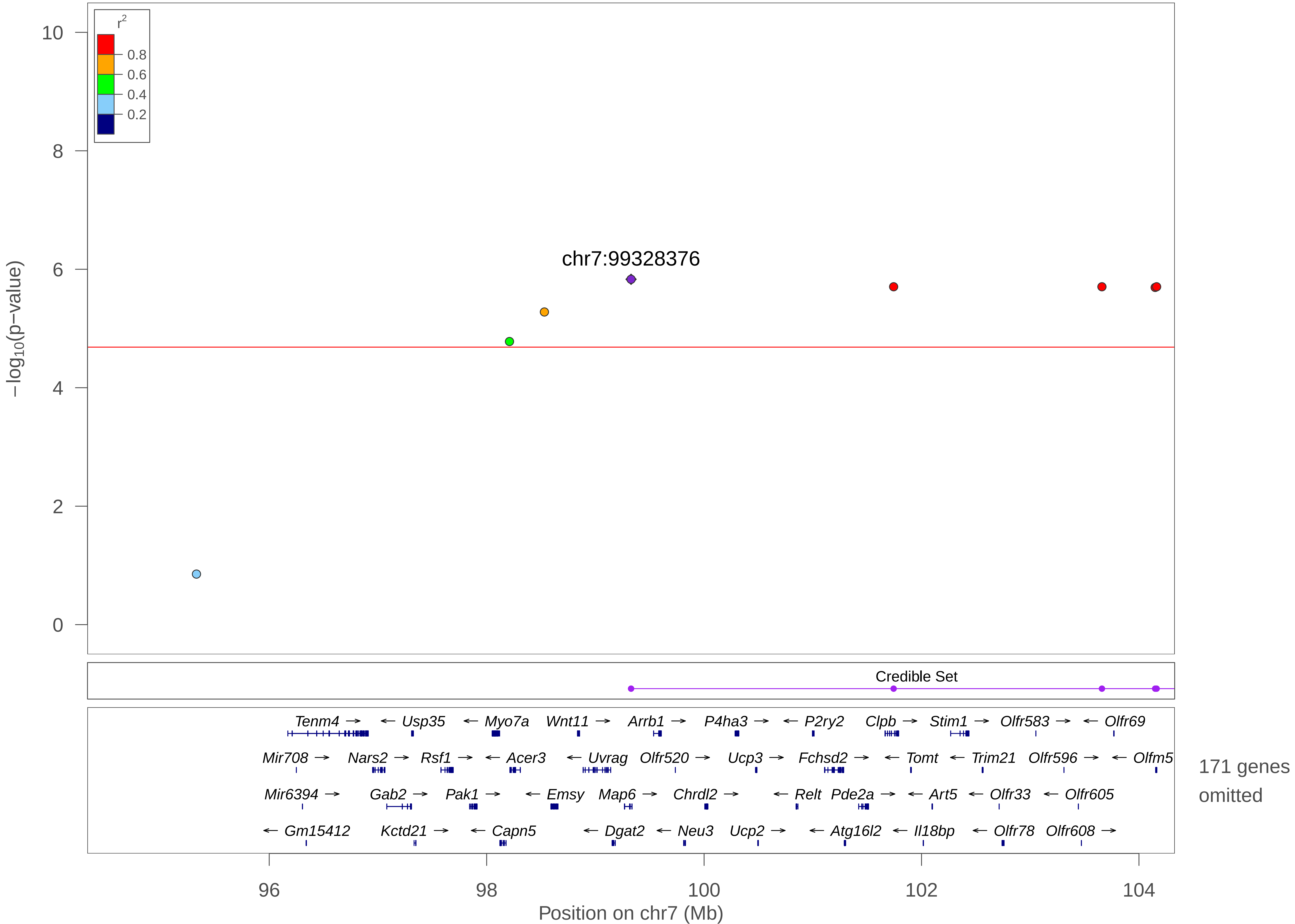

date: Thu Oct 25 16:46:56 2018
build: F34Array
display range: chr7:94328376−104328376 [94328376−104328376]
hilite range: 0 − 0 [ 0 − 0 ]
reference SNP: chr7:99328376
number of SNPs plotted: 9
min P−value: 1.48E−6 [chr7:99328376]
max P−value: 1.4E−1 [chr7:95330810]
omitted Genes: Gm15413, Alg8, Ndufc2
omitted Genes: Thrsp, Kctd14, Ints4
omitted Genes: Aamdc, Rsf1os1, Clns1a
omitted Genes: Aqp11, Gdpd4, Omp
omitted Genes: Gm16938, B3gnt6, Tsku
omitted Genes: Gucy2d, Lrrc32, Prkrir
omitted Genes: Mogat2, Gm26705, Serpinh1
omitted Genes: Gdpd5, Klhl35, Rps3
omitted Genes: Snord15b, Snord15a, Mir326
omitted Genes: Tpbgl, Slco2b1, Olfr521
omitted Genes: Spcs2, Xrra1, Rnf169
omitted Genes: Pold3, Lipt2, Kcne3
omitted Genes: Pgm2l1, Gpx2−ps1, Ppme1
omitted Genes: C2cd3, Dnajb13, Coa4
omitted Genes: Mrpl48, Rab6a, Plekhb1
omitted Genes: Fam168a, Arhgef17, Mir3102
omitted Genes: P2ry6, Stard10, Arap1
omitted Genes: Mir139, Art2a−ps, Art2b
omitted Genes: Phox2a, Inppl1, Folr2
omitted Genes: Folr1, Anapc15, Lamtor1
omitted Genes: Lrrc51, Numa1, Rnf121
omitted Genes: Xndc1, Xntrpc, Trpc2
omitted Genes: Art1, Chrna10, Nup98
omitted Genes: Pgap2, Rhog, Rrm1
omitted Genes: Olfr543, Olfr544, Olfr545
omitted Genes: Olfr547, Olfr549, Olfr550
omitted Genes: Olfr551, Olfr552, Olfr553
omitted Genes: Olfr554, Olfr555, Olfr556
omitted Genes: Trim68, Olfr557, Olfr558
omitted Genes: Olfr559, Olfr560, Olfr561
omitted Genes: Olfr564, Olfr566, Olfr568
Make more plots at http://csg.sph.umich.edu/locuszoom/
omitted Genes: Olfr569, Olfr570, Olfr571

Fine−mapping Regions

| snp | chr | pos (Mb) | pp | group | color |
|---|---|---|---|---|---|
| chr7:99328376 | 7 | 99.32838 | 0.2096164 | Credible Set | purple |
| chr7:101743401 | 7 | 101.74340 | 0.1586110 | Credible Set | purple |
| chr7:103659355 | 7 | 103.65936 | 0.1586110 | Credible Set | purple |
| chr7:104149022 | 7 | 104.14902 | 0.1547811 | Credible Set | purple |
| chr7:104162444 | 7 | 104.16244 | 0.1586110 | Credible Set | purple |
| chr7:104581631 | 7 | 104.58163 | 0.1597694 | Credible Set | purple |

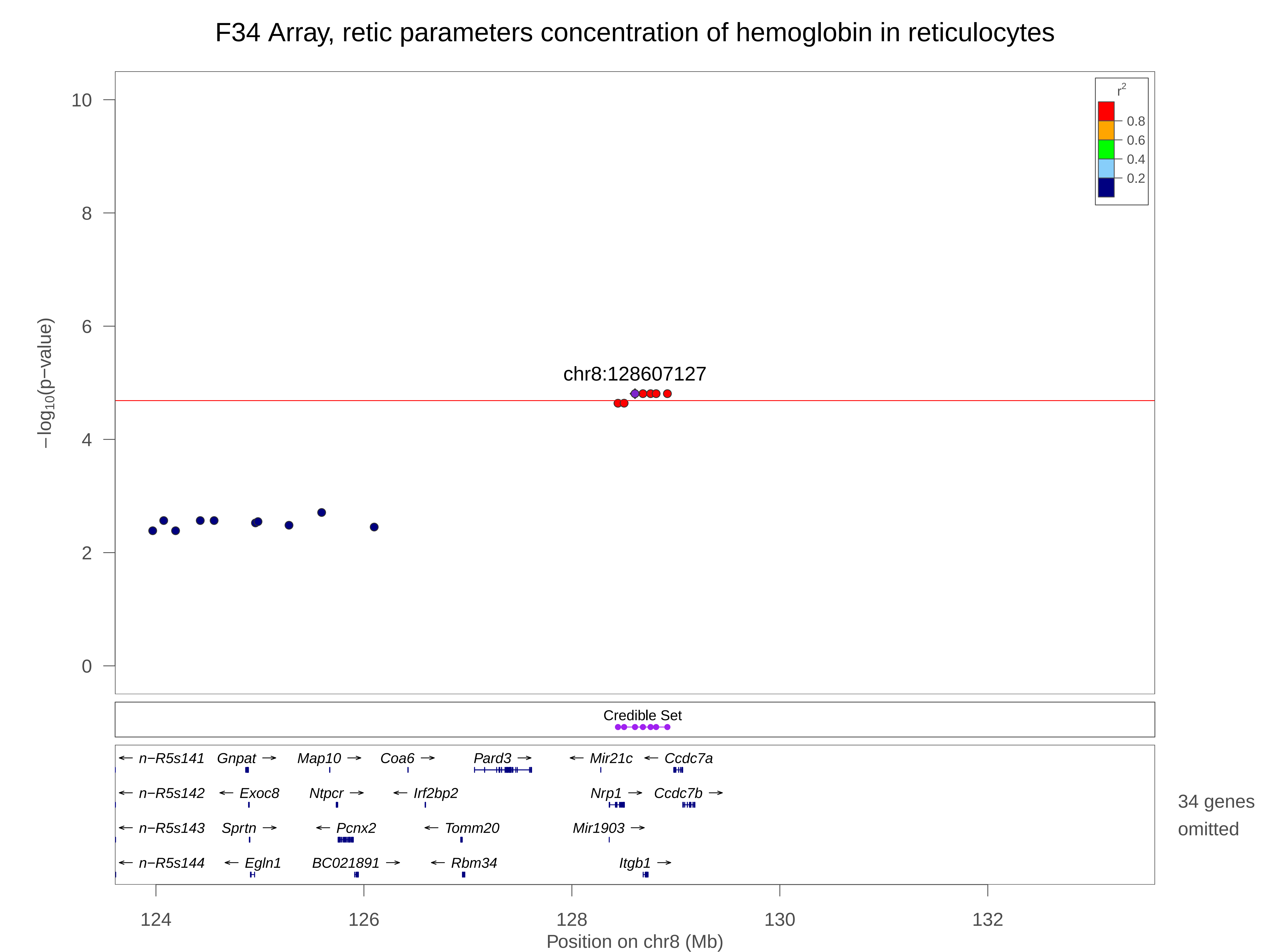

F34 Array, retic parameters concentration of hemoglobin in reticulocytes
r2
0.8
0.6
0.4
0.2
10
8
6
4
2
0
−log10(p−value)
chr8:128607127
Credible Set
← n−R5s141
Gnpat →
Map10 →
Coa6 →
Pard3 →
← Mir21c
← Ccdc7a
← n−R5s142
← Exoc8
Ntpcr →
← Irf2bp2
Nrp1 →
Ccdc7b →
← n−R5s143
Sprtn →
← Pcnx2
← Tomm20
Mir1903 →
← n−R5s144
← Egln1
BC021891 →
← Rbm34
Itgb1 →
34 genes omitted
124
126
128
130
132
Position on chr8 (Mb)

| | |
|---|---|
| date: | Thu Oct 25 16:47:03 2018 |
| build: | F34Array |
| display range: | chr8:123607127−133607127 [123607127−133607127] |
| hilite range: | 0 − 0 [ 0 − 0 ] |
| reference SNP: | chr8:128607127 |
| number of SNPs plotted: | 18 |
| min P−value: | 1.56E−5 [chr8:128607127] |
| max P−value: | 4.1E−3 [chr8:123968866] |
| omitted Genes: | Gm25212, n−R5s146, Gm22291 |
| omitted Genes: | Gm23284, n−R5s149, Rhou |
| omitted Genes: | 6030466F02Rik, Rab4a, Ccsap |
| omitted Genes: | Acta1, Nup133, Abcb10 |
| omitted Genes: | Taf5l, Mir1967, Urb2 |
| omitted Genes: | Galnt2, Pgbd5, Cog2 |
| omitted Genes: | Agt, Capn9, 2310022B05Rik |
| omitted Genes: | Ttc13, Arv1, Fam89a |
| omitted Genes: | Trim67, 2810004N23Rik, Tsnax |
| omitted Genes: | Disc1, Sipa1l2, 4930567H12Rik |
| omitted Genes: | Kcnk1, Slc35f3, Gm17296 |
| omitted Genes: | 2610044O15Rik8 |

Make more plots at http://csg.sph.umich.edu/locuszoom/

Fine−mapping Regions

| snp | chr | pos (Mb) | pp | group | color |
|---|---|---|---|---|---|
| chr8:128443469 | 8 | 128.4435 | 0.1081470 | Credible Set | purple |
| chr8:128502425 | 8 | 128.5024 | 0.1083470 | Credible Set | purple |
| chr8:128607127 | 8 | 128.6071 | 0.1567012 | Credible Set | purple |
| chr8:128683251 | 8 | 128.6833 | 0.1567012 | Credible Set | purple |
| chr8:128757217 | 8 | 128.7572 | 0.1567012 | Credible Set | purple |
| chr8:128810304 | 8 | 128.8103 | 0.1567012 | Credible Set | purple |
| chr8:128918665 | 8 | 128.9187 | 0.1567012 | Credible Set | purple |

F34 Array, retic parameters mean reticulocyte volume

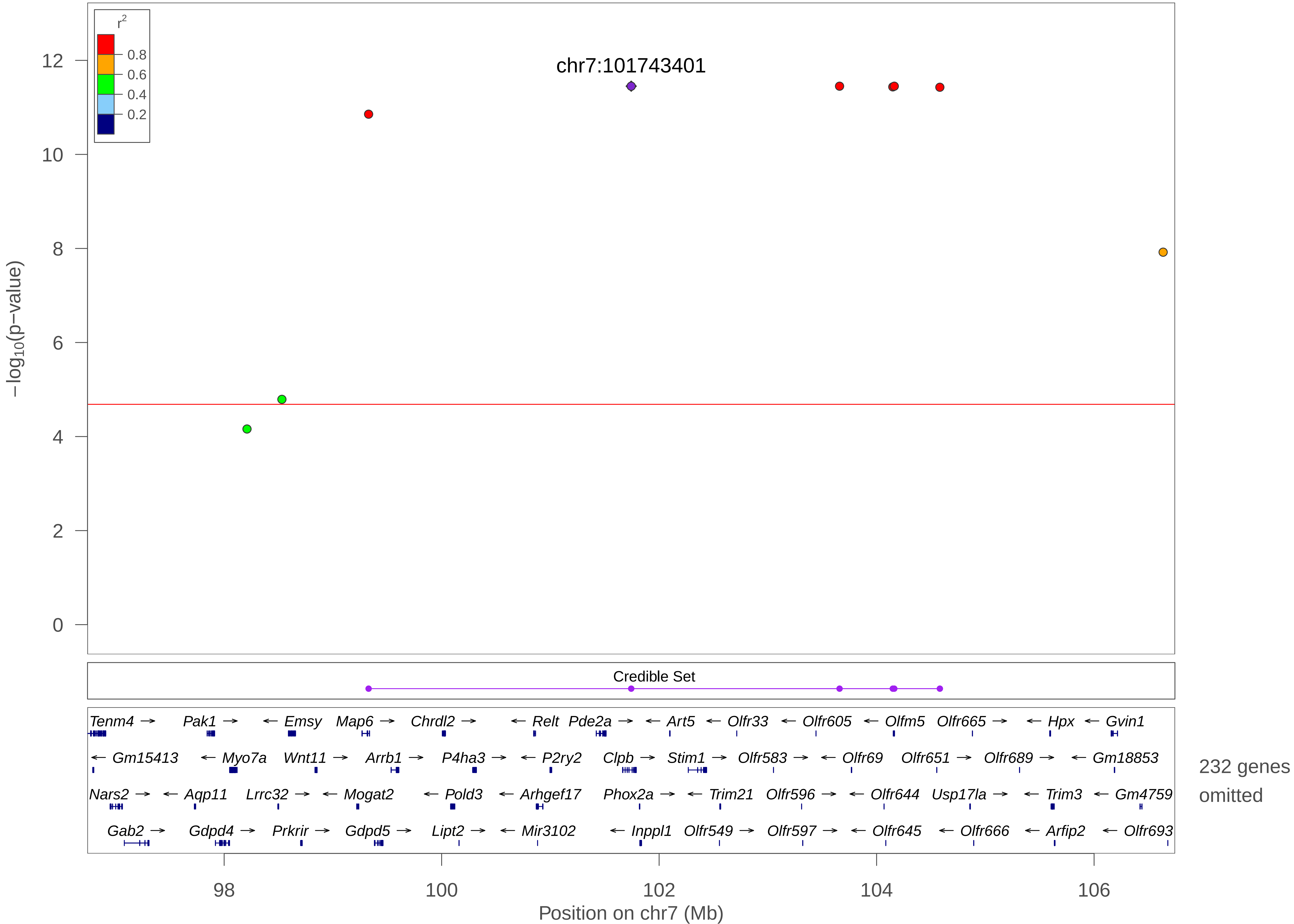

date: Thu Oct 25 16:47:22 2018
build: F34Array
display range: chr7:96743401−106743401 [96743401−106743401]
hilite range: 0 − 0 [ 0 − 0 ]
reference SNP: chr7:101743401
number of SNPs plotted: 10
min P−value: 3.54E−12 [chr7:101743401]
max P−value: 6.9E−5 [chr7:98210049]
omitted Genes: Usp35, Kctd21, Alg8
omitted Genes: Ndufc2, Thrsp, Kctd14
omitted Genes: Ints4, Aamdc, Rsf1
omitted Genes: Rsf1os1, Clns1a, Capn5
omitted Genes: Omp, Gm16938, B3gnt6
omitted Genes: Acer3, Tsku, Gucy2d
omitted Genes: Uvrag, Dgat2, Gm26705
omitted Genes: Serpinh1, Klhl35, Rps3
omitted Genes: Snord15b, Snord15a, Mir326
omitted Genes: Tpbgl, Slco2b1, Olfr520
omitted Genes: Olfr521, Neu3, Spcs2
omitted Genes: Xrra1, Rnf169, Kcne3
omitted Genes: Pgm2l1, Gpx2−ps1, Ppme1
omitted Genes: C2cd3, Ucp3, Ucp2
omitted Genes: Dnajb13, Coa4, Mrpl48
omitted Genes: Rab6a, Plekhb1, Fam168a
omitted Genes: P2ry6, Fchsd2, Atg16l2
omitted Genes: Stard10, Arap1, Mir139
omitted Genes: Art2a−ps, Art2b, Folr2
omitted Genes: Folr1, Anapc15, Tomt
omitted Genes: Lamtor1, Lrrc51, Numa1
omitted Genes: Il18bp, Rnf121, Xndc1
omitted Genes: Xntrpc, Trpc2, Art1
omitted Genes: Chrna10, Nup98, Pgap2
omitted Genes: Rhog, Rrm1, Olfr543
omitted Genes: Olfr544, Olfr545, Olfr547
omitted Genes: Olfr550, Olfr551, Olfr552
omitted Genes: Olfr553, Olfr554, Olfr555
omitted Genes: Olfr556, Trim68, Olfr557
omitted Genes: Olfr558, Olfr559, Olfr78
omitted Genes: Olfr560, Olfr561, Olfr564
omitted Genes: Olfr566, Olfr568, Olfr569

Make more plots at http://csg.sph.umich.edu/locuszoom/

Fine−mapping Regions

| snp | chr | pos (Mb) | pp | group | color |
|---|---|---|---|---|---|
| chr7:99328376 | 7 | 99.32838 | 0.05039541 | Credible Set | purple |
| chr7:101743401 | 7 | 101.74340 | 0.19303482 | Credible Set | purple |
| chr7:103659355 | 7 | 103.65936 | 0.19303482 | Credible Set | purple |
| chr7:104149022 | 7 | 104.14902 | 0.18688221 | Credible Set | purple |
| chr7:104162444 | 7 | 104.16244 | 0.19303482 | Credible Set | purple |
| chr7:104581631 | 7 | 104.58163 | 0.18361792 | Credible Set | purple |

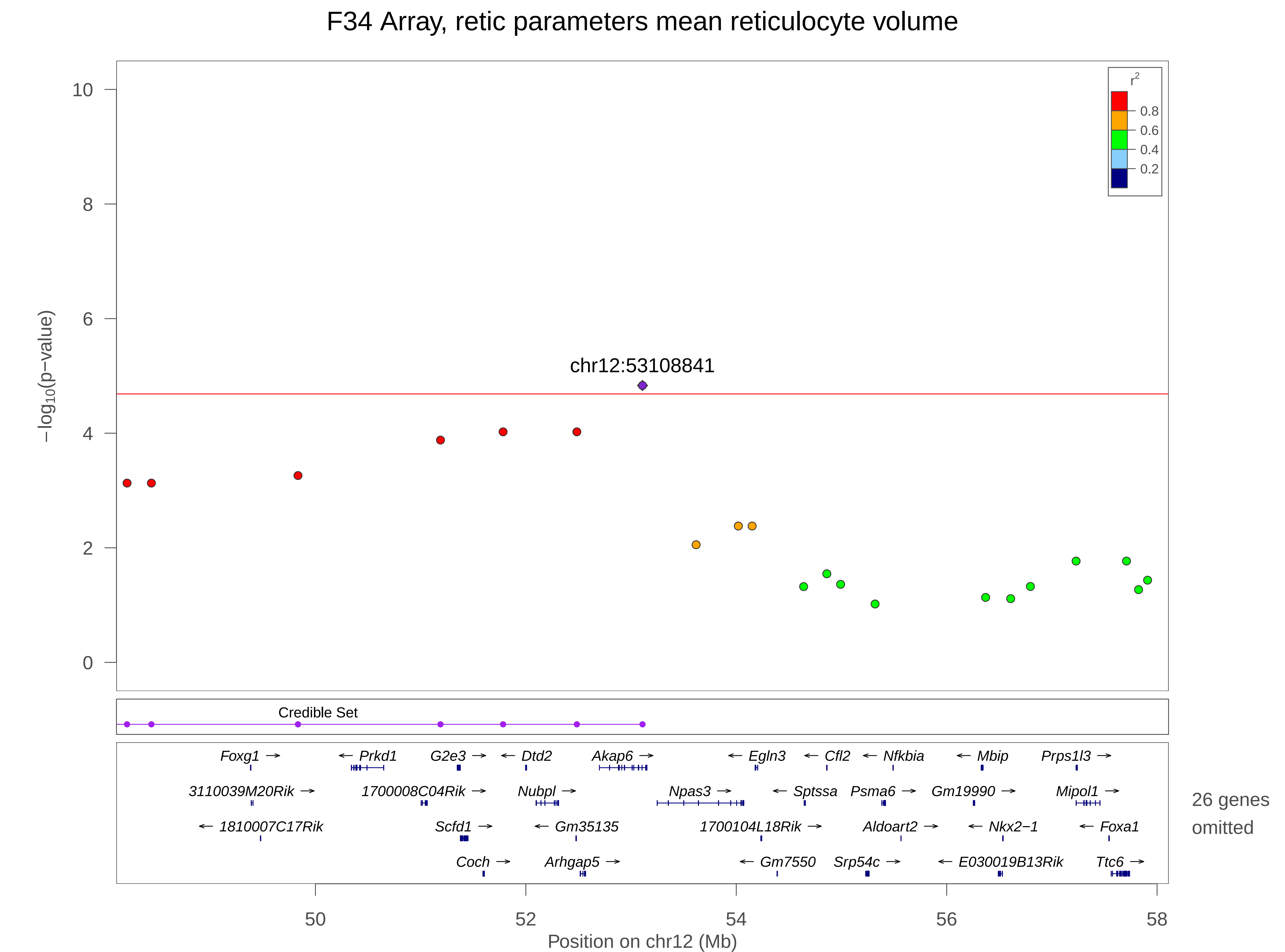
F34 Array, retic parameters mean reticulocyte volume
−log10(p−value)
chr12:53108841
r2
0.8
0.6
0.4
0.2
Credible Set
Foxg1
Prkd1
G2e3
Dtd2
Akap6
Egln3
Cfl2
Nfkbia
Mbip
Prps1l3
3110039M20Rik
1700008C04Rik
Nubpl
Npas3
Sptssa
Psma6
Gm19990
Mipol1
1810007C17Rik
Scfd1
Gm35135
1700104L18Rik
Aldoart2
Nkx2−1
Foxa1
Coch
Arhgap5
Gm7550
Srp54c
E030019B13Rik
Ttc6
26 genes omitted
Position on chr12 (Mb)

| | |
|---|---|
| date: | Thu Oct 25 16:47:11 2018 |
| build: | F34Array |
| display range: | chr12:48108841−58108841 [48108841−58108841] |
| hilite range: | 0 − 0 [ 0 − 0 ] |
| reference SNP: | chr12:53108841 |
| number of SNPs plotted: | 22 |
| min P−value: | 1.47E−5 [chr12:53108841] |
| max P−value: | 9.56E−2 [chr12:55319581] |
| omitted Genes: | Strn3, Ap4s1, Hectd1 |
| omitted Genes: | Heatr5a, Gpr33, 1700031P21Rik |
| omitted Genes: | Mir1892, Eapp, Snx6 |
| omitted Genes: | Baz1a, 2700097O09Rik, Srp54a |
| omitted Genes: | Fam177a, Srp54b, 1700047I17Rik2 |
| omitted Genes: | Ppp2r3c, 1110008L16Rik, Insm2 |
| omitted Genes: | Ralgapa1, Brms1l, C87198 |
| omitted Genes: | Nkx2−9, Pax9, Slc25a21 |
| omitted Genes: | 4921518K17Rik, BC042761 |

Make more plots at http://csg.sph.umich.edu/locuszoom/

Fine−mapping Regions

| snp | chr | pos (Mb) | pp | group | color |
|---|---|---|---|---|---|
| chr12:46941836 | 12 | 46.94184 | 0.01575526 | Credible Set | purple |
| chr12:48209549 | 12 | 48.20955 | 0.01575526 | Credible Set | purple |
| chr12:48441097 | 12 | 48.44110 | 0.01575526 | Credible Set | purple |
| chr12:49834234 | 12 | 49.83423 | 0.02088924 | Credible Set | purple |
| chr12:51188722 | 12 | 51.18872 | 0.07939813 | Credible Set | purple |
| chr12:51783700 | 12 | 51.78370 | 0.10856270 | Credible Set | purple |
| chr12:52484891 | 12 | 52.48489 | 0.10856270 | Credible Set | purple |
| chr12:53108841 | 12 | 53.10884 | 0.63532146 | Credible Set | purple |

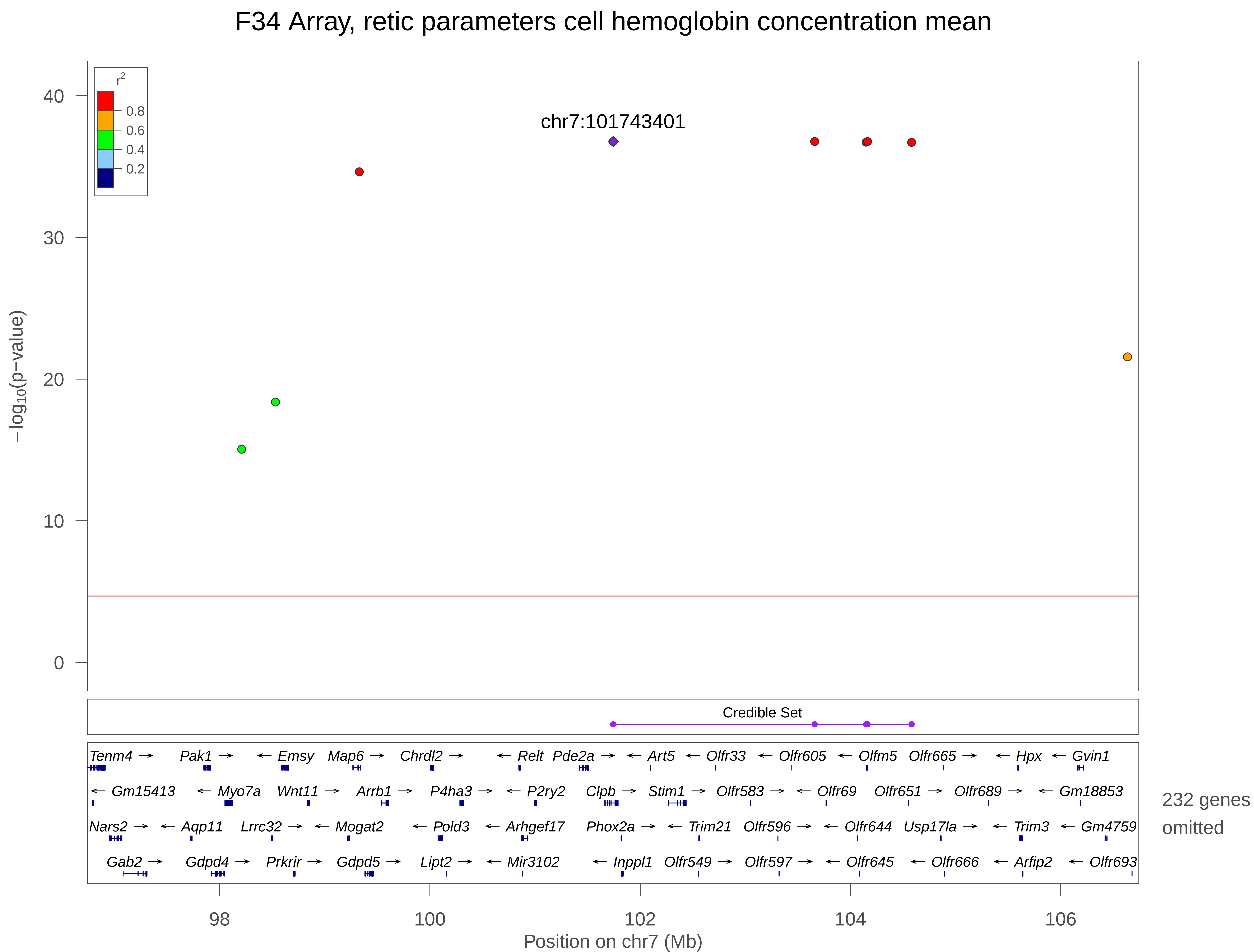
F34 Array, retic parameters cell hemoglobin concentration mean
chr7:101743401
−log10(p−value)
r2
0.8
0.6
0.4
0.2
Credible Set
Tenm4 Pak1 Emsy Map6 Chrdl2 Relt Pde2a Art5 Olfr33 Olfr605 Olfm5 Olfr665 Hpx Gvin1
Gm15413 Myo7a Wnt11 Arrb1 P4ha3 P2ry2 Clpb Stim1 Olfr583 Olfr69 Olfr651 Olfr689 Gm18853
Nars2 Aqp11 Lrrc32 Mogat2 Pold3 Arhgef17 Phox2a Trim21 Olfr596 Olfr644 Usp17la Trim3 Gm4759
Gab2 Gdpd4 Prkrir Gdpd5 Lipt2 Mir3102 Inppl1 Olfr549 Olfr597 Olfr645 Olfr666 Arfip2 Olfr693
232 genes omitted
Position on chr7 (Mb)

date: Thu Oct 25 16:47:43 2018
build: F34Array
display range: chr7:96743401−106743401 [96743401−106743401]
hilite range: 0 − 0 [ 0 − 0 ]
reference SNP: chr7:101743401
number of SNPs plotted: 10
min P−value: 1.69E−37 [chr7:101743401]
max P−value: 8.92E−16 [chr7:98210049]
omitted Genes: Usp35, Kctd21, Alg8
omitted Genes: Ndufc2, Thrsp, Kctd14
omitted Genes: Ints4, Aamdc, Rsf1
omitted Genes: Rsf1os1, Clns1a, Capn5
omitted Genes: Omp, Gm16938, B3gnt6
omitted Genes: Acer3, Tsku, Gucy2d
omitted Genes: Uvrag, Dgat2, Gm26705
omitted Genes: Serpinh1, Klhl35, Rps3
omitted Genes: Snord15b, Snord15a, Mir326
omitted Genes: Tpbgl, Slco2b1, Olfr520
omitted Genes: Olfr521, Neu3, Spcs2
omitted Genes: Xrra1, Rnf169, Kcne3
omitted Genes: Pgm2l1, Gpx2−ps1, Ppme1
omitted Genes: C2cd3, Ucp3, Ucp2
omitted Genes: Dnajb13, Coa4, Mrpl48
omitted Genes: Rab6a, Plekhb1, Fam168a
omitted Genes: P2ry6, Fchsd2, Atg16l2
omitted Genes: Stard10, Arap1, Mir139
omitted Genes: Art2a−ps, Art2b, Folr2
omitted Genes: Folr1, Anapc15, Tomt
omitted Genes: Lamtor1, Lrrc51, Numa1
omitted Genes: Il18bp, Rnf121, Xndc1
omitted Genes: Xntrpc, Trpc2, Art1
omitted Genes: Chrna10, Nup98, Pgap2
omitted Genes: Rhog, Rrm1, Olfr543
omitted Genes: Olfr544, Olfr545, Olfr547
omitted Genes: Olfr550, Olfr551, Olfr552
omitted Genes: Olfr553, Olfr554, Olfr555
omitted Genes: Olfr556, Trim68, Olfr557
omitted Genes: Olfr558, Olfr559, Olfr78
omitted Genes: Olfr560, Olfr561, Olfr564

omitted Genes: Olfr566, Olfr568, Olfr569

Fine−mapping Regions

| snp | chr | pos (Mb) | pp | group | color |
|---|---|---|---|---|---|
| chr7:101743401 | 7 | 101.7434 | 0.2093022 | Credible Set | purple |
| chr7:103659355 | 7 | 103.6594 | 0.2093022 | Credible Set | purple |
| chr7:104149022 | 7 | 104.1490 | 0.1900482 | Credible Set | purple |
| chr7:104162444 | 7 | 104.1624 | 0.2093022 | Credible Set | purple |
| chr7:104581631 | 7 | 104.5816 | 0.1804691 | Credible Set | purple |

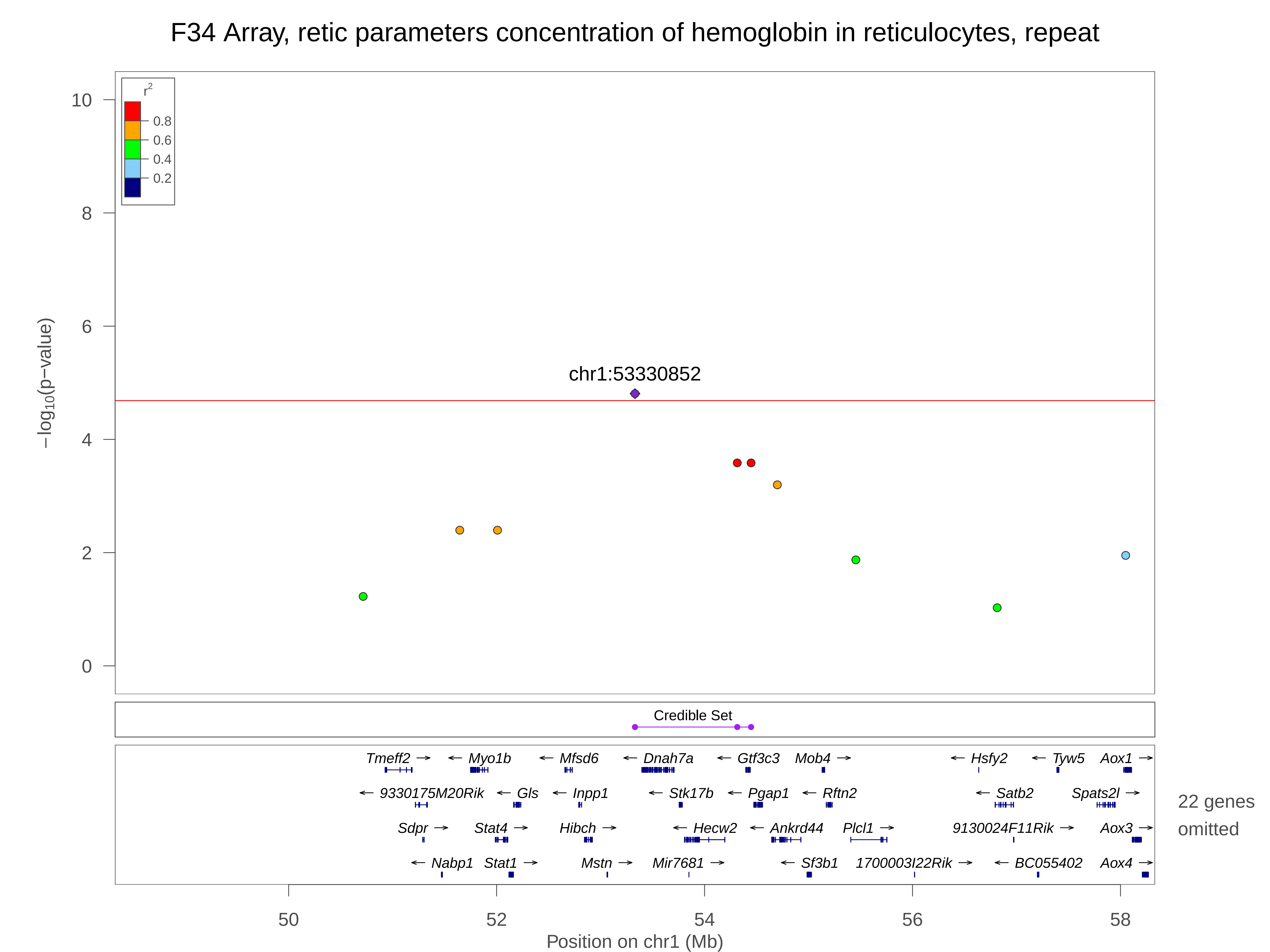
F34 Array, retic parameters concentration of hemoglobin in reticulocytes, repeat
r2
0.8
0.6
0.4
0.2
−log10(p−value)
10
8
6
4
2
0
chr1:53330852
Credible Set
Tmeff2 →
← Myo1b
← Mfsd6
← Dnah7a
← Gtf3c3
Mob4 →
← Hsfy2
← Tyw5
Aox1 →
← 9330175M20Rik
← Gls
← Inpp1
← Stk17b
← Pgap1
← Rftn2
← Satb2
Spats2l →
Sdpr →
Stat4 →
Hibch →
← Hecw2
← Ankrd44
Plcl1 →
9130024F11Rik →
Aox3 →
← Nabp1
Stat1 →
Mstn →
Mir7681 →
← Sf3b1
1700003I22Rik →
← BC055402
Aox4 →
22 genes
omitted
50
52
54
56
58
Position on chr1 (Mb)

| | |
|---|---|
| date: | Thu Oct 25 16:47:59 2018 |
| build: | F34Array |
| display range: | chr1:48330852−58330852 [48330852−58330852] |
| hilite range: | 0 − 0 [ 0 − 0 ] |
| reference SNP: | chr1:53330852 |
| number of SNPs plotted: | 11 |
| min P−value: | 1.55E−5 [chr1:53330852] |
| max P−value: | 9.41E−2 [chr1:56814006] |
| omitted Genes: | Nab1, Tmem194b, 1700019D03Rik |
| omitted Genes: | 1700019A02Rik, Pms1, Ormdl1 |
| omitted Genes: | Osgepl1, Asnsd1, Ccdc150 |
| omitted Genes: | Coq10b, Hspd1, Hspe1 |
| omitted Genes: | Mars2, Boll, Gm6644 |
| omitted Genes: | 9130227L01Rik, 4930558J18Rik, 1700066M21Rik |
| omitted Genes: | 9430016H08Rik, Kctd18, Sgol2a |
| omitted Genes: | Aox2 |

Make more plots at http://csg.sph.umich.edu/locuszoom/

Fine−mapping Regions

| snp | chr | pos (Mb) | pp | group | color |
|---|---|---|---|---|---|
| chr1:53330852 | 1 | 53.33085 | 0.8778534 | Credible Set | purple |
| chr1:54314429 | 1 | 54.31443 | 0.0610733 | Credible Set | purple |
| chr1:54447274 | 1 | 54.44727 | 0.0610733 | Credible Set | purple |

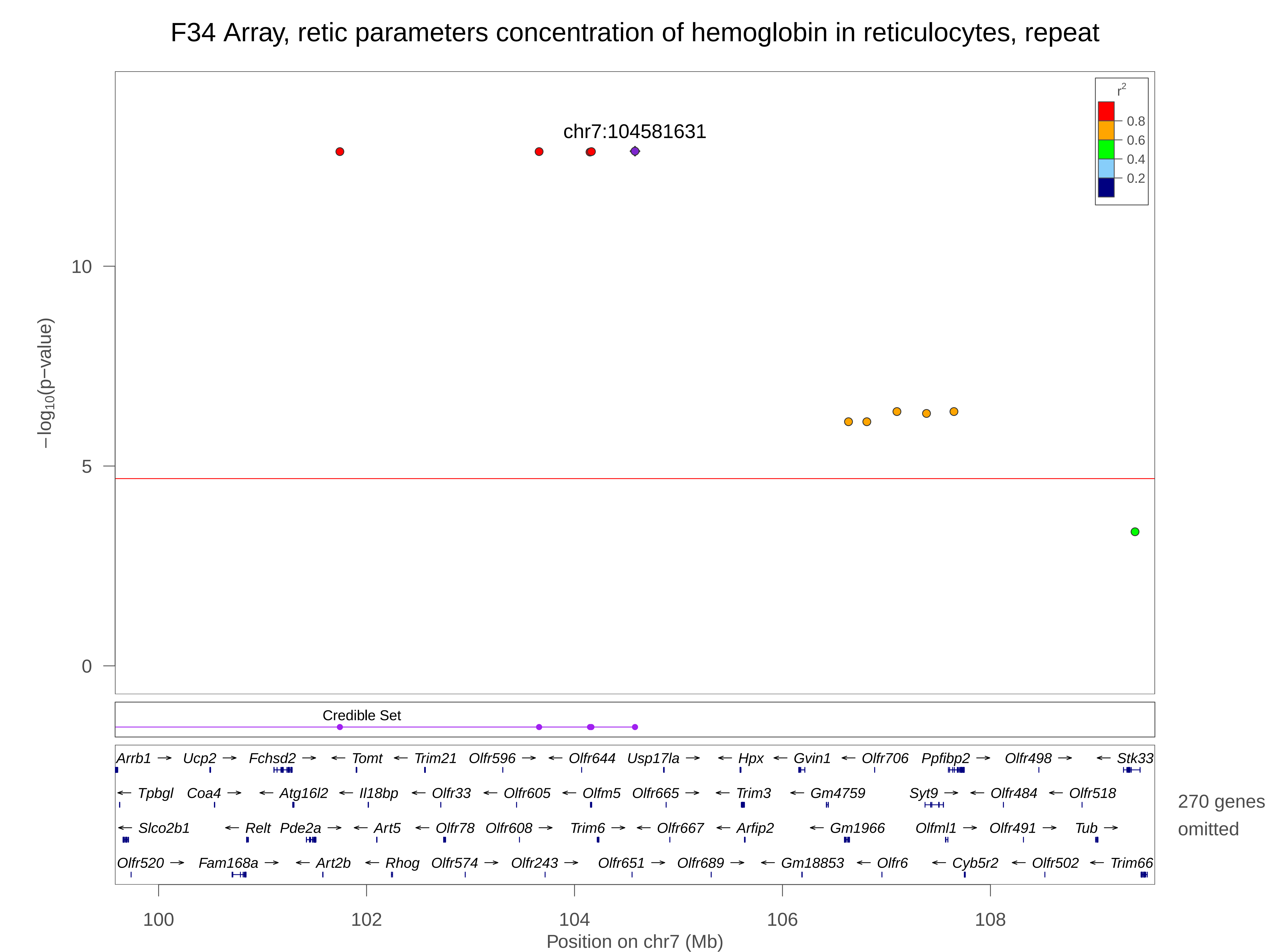
F34 Array, retic parameters concentration of hemoglobin in reticulocytes, repeat
chr7:104581631
r²
0.8
0.6
0.4
0.2
10
5
0
−log10(p−value)
Credible Set
Arrb1 → Ucp2 → Fchsd2 → ← Tomt ← Trim21 Olfr596 → ← Olfr644 Usp17la → ← Hpx ← Gvin1 ← Olfr706 Ppfibp2 → Olfr498 → ← Stk33
← Tpbgl Coa4 → ← Atg16l2 ← Il18bp ← Olfr33 ← Olfr605 ← Olfm5 Olfr665 → ← Trim3 ← Gm4759 Syt9 → ← Olfr484 ← Olfr518
← Slco2b1 ← Relt Pde2a → ← Art5 ← Olfr78 Olfr608 → Trim6 → ← Olfr667 ← Arfip2 ← Gm1966 Olfml1 → Olfr491 → Tub →
Olfr520 → Fam168a → ← Art2b ← Rhog Olfr574 → Olfr243 → Olfr651 → Olfr689 → ← Gm18853 ← Olfr6 ← Cyb5r2 ← Olfr502 ← Trim66
270 genes omitted
100
102
104
106
108
Position on chr7 (Mb)

date: Thu Oct 25 16:48:27 2018
build: F34Array
display range: chr7:99581631−109581631 [99581631−109581631]
hilite range: 0 − 0 [ 0 − 0 ]
reference SNP: chr7:104581631
number of SNPs plotted: 12
min P−value: 1.32E−13 [chr7:104581631]
max P−value: 4.41E−4 [chr7:109390566]
omitted Genes: Olfr521, Neu3, Spcs2
omitted Genes: Xrra1, Rnf169, Chrdl2
omitted Genes: Pold3, Lipt2, Kcne3
omitted Genes: Pgm2l1, Gpx2−ps1, P4ha3
omitted Genes: Ppme1, C2cd3, Ucp3
omitted Genes: Dnajb13, Mrpl48, Rab6a
omitted Genes: Plekhb1, Arhgef17, Mir3102
omitted Genes: P2ry6, P2ry2, Stard10
omitted Genes: Arap1, Mir139, Art2a−ps
omitted Genes: Clpb, Phox2a, Inppl1
omitted Genes: Folr2, Folr1, Anapc15
omitted Genes: Lamtor1, Lrrc51, Numa1
omitted Genes: Rnf121, Xndc1, Xntrpc
omitted Genes: Trpc2, Art1, Chrna10
omitted Genes: Nup98, Pgap2, Stim1
omitted Genes: Rrm1, Olfr543, Olfr544
omitted Genes: Olfr545, Olfr547, Olfr549
omitted Genes: Olfr550, Olfr551, Olfr552
omitted Genes: Olfr553, Olfr554, Olfr555
omitted Genes: Olfr556, Trim68, Olfr557
omitted Genes: Olfr558, Olfr559, Olfr560
omitted Genes: Olfr561, Olfr564, Olfr566
omitted Genes: Olfr568, Olfr569, Olfr570
omitted Genes: Olfr571, Olfr572, Olfr575
omitted Genes: Olfr576, Olfr577, Olfr578
omitted Genes: Olfr582, Olfr583, Olfr584
omitted Genes: Olfr585, Olfr586, Olfr589
omitted Genes: Olfr591, Olfr592, Olfr593
omitted Genes: Olfr594, Usp17ld, Olfr597
omitted Genes: Olfr598, Olfr599, Olfr600
omitted Genes: Olfr601, Olfr603, Usp17lc
Make more plots at http://csg.sph.umich.edu/locuszoom/
omitted Genes: Olfr606, Olfr609, Olfr610

Fine−mapping Regions

| snp | chr | pos (Mb) | pp | group | color |
|---|---|---|---|---|---|
| chr7:99328376 | 7 | 99.32838 | 0.06413313 | Credible Set | purple |
| chr7:101743401 | 7 | 101.74340 | 0.18722938 | Credible Set | purple |
| chr7:103659355 | 7 | 103.65936 | 0.18722938 | Credible Set | purple |
| chr7:104149022 | 7 | 104.14902 | 0.18251656 | Credible Set | purple |
| chr7:104162444 | 7 | 104.16244 | 0.18722938 | Credible Set | purple |
| chr7:104581631 | 7 | 104.58163 | 0.19166215 | Credible Set | purple |

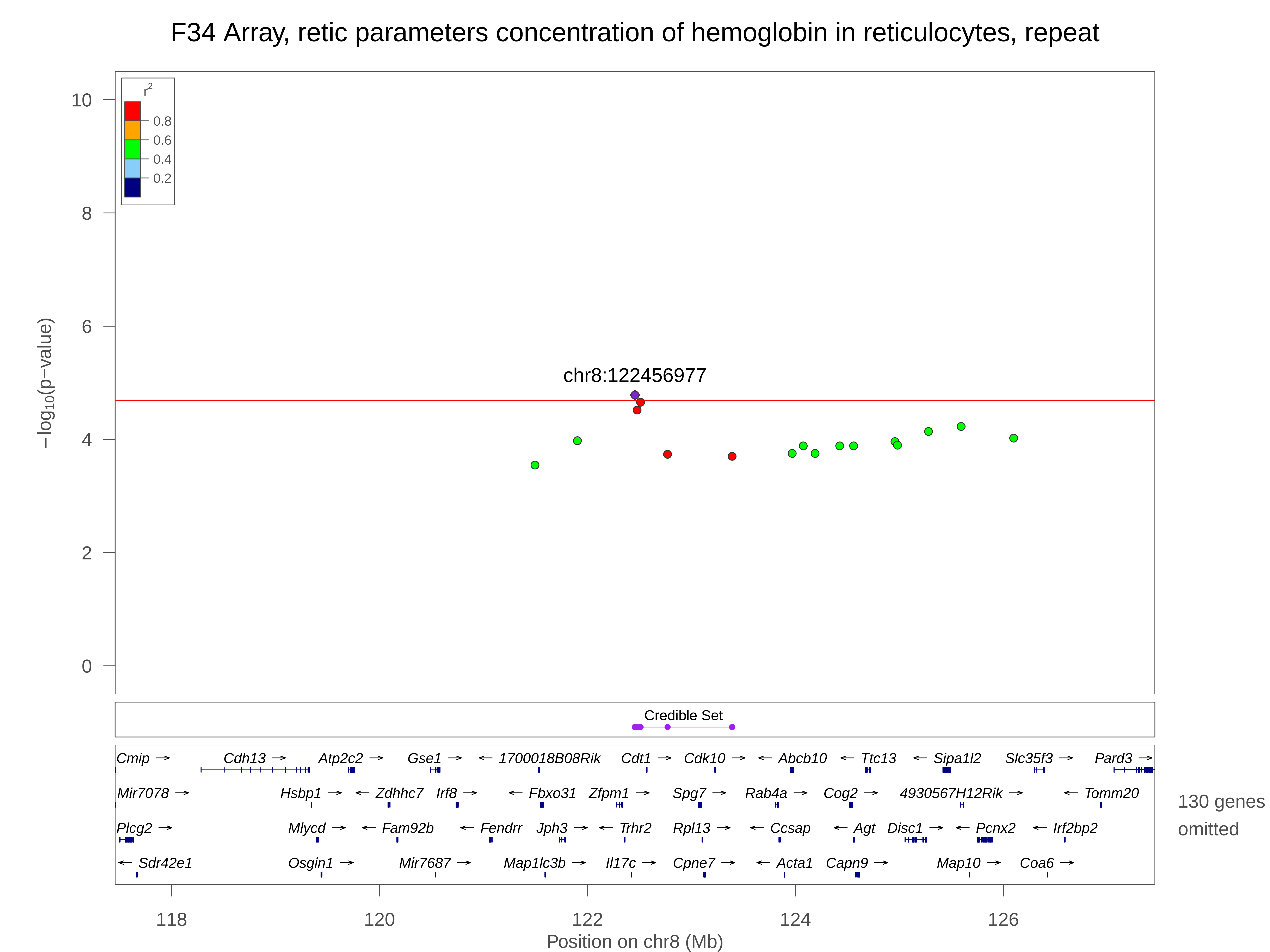

F34 Array, retic parameters concentration of hemoglobin in reticulocytes, repeat
r2
0.8
0.6
0.4
0.2
10
8
6
4
2
0
−log10(p−value)
chr8:122456977
Credible Set
Cmip →
Cdh13 →
Atp2c2 →
Gse1 →
← 1700018B08Rik
Cdt1 →
Cdk10 →
← Abcb10
← Ttc13
← Sipa1l2
Slc35f3 →
Pard3 →
Mir7078 →
Hsbp1 →
← Zdhhc7
Irf8 →
← Fbxo31
Zfpm1 →
Spg7 →
Rab4a →
Cog2 →
4930567H12Rik →
← Tomm20
Plcg2 →
Mlycd →
← Fam92b
← Fendrr
Jph3 →
← Trhr2
Rpl13 →
← Ccsap
← Agt
Disc1 →
← Pcnx2
← Irf2bp2
← Sdr42e1
Osgin1 →
Mir7687 →
Map1lc3b →
Il17c →
Cpne7 →
← Acta1
Capn9 →
Map10 →
Coa6 →
130 genes omitted
118
120
122
124
126
Position on chr8 (Mb)

| | |
|---|---|
| date: | Thu Oct 25 16:48:47 2018 |
| build: | F34Array |
| display range: | chr8:117456977−127456977 [117456977−127456977] |
| hilite range: | 0 − 0 [ 0 − 0 ] |
| reference SNP: | chr8:122456977 |
| number of SNPs plotted: | 18 |
| min P−value: | 1.65E−5 [chr8:122456977] |
| max P−value: | 2.84E−4 [chr8:121495351] |
| omitted Genes: | Gm38416, Hsd17b2, Mphosph6 |
| omitted Genes: | Necab2, Slc38a8, Mbtps1 |
| omitted Genes: | Hsdl1, Dnaaf1, Taf1c |
| omitted Genes: | Adad2, Kcng4, Wfdc1 |
| omitted Genes: | Tldc1, Cotl1, Klhl36 |
| omitted Genes: | Usp10, Crispld2, 6430548M08Rik |
| omitted Genes: | A330074K22Rik, Gins2, Emc8−1190005i06rik |
| omitted Genes: | 1190005I06Rik, Emc8, Cox4i1 |
| omitted Genes: | Foxf1, Mthfsd, Foxc2 |
| omitted Genes: | Foxl1, 1700030M09Rik, Zcchc14 |
| omitted Genes: | Klhdc4, Slc7a5, BC048644 |
| omitted Genes: | Car5a, Banp, Mir7237 |
| omitted Genes: | Zc3h18, Cyba, Mvd |
| omitted Genes: | 9330133O14Rik, Gm20735, Snai3 |
| omitted Genes: | Rnf166, Ctu2, Piezo1 |
| omitted Genes: | Aprt, Galns, Trappc2l |
| omitted Genes: | Pabpn1l, Mir6396, Cbfa2t3 |
| omitted Genes: | Acsf3, Cdh15, Ankrd11 |
| omitted Genes: | 2810013P06Rik, Snord68, Mir7079 |
| omitted Genes: | Mir7080, Sult5a1, Dpep1 |
| omitted Genes: | Chmp1a, Spata33, Spata2l |
| omitted Genes: | 4933417D19Rik, Vps9d1, Zfp276 |
| omitted Genes: | Fanca, Spire2, Tcf25 |
| omitted Genes: | Mc1r, Tubb3, Def8 |
| omitted Genes: | Afg3l1, Dbndd1, Gas8 |
| omitted Genes: | n−R5s100, n−R5s103, n−R5s104 |
| omitted Genes: | n−R5s105, n−R5s108, n−R5s110 |
| omitted Genes: | n−R5s111, n−R5s113, Gm25018 |
| omitted Genes: | n−R5s117, n−R5s121, n−R5s122 |
| omitted Genes: | n−R5s123, n−R5s124, n−R5s128 |
| omitted Genes: | Gm26391, n−R5s133, n−R5s134 |
| omitted Genes: | Rn5s, n−R5s136, n−R5s138 |

Make more plots at http://csg.sph.umich.edu/locuszoom/

Fine−mapping Regions

| snp | chr | pos (Mb) | pp | group | color |
|---|---|---|---|---|---|
| chr8:122456977 | 8 | 122.4570 | 0.39795211 | Credible Set | purple |
| chr8:122477261 | 8 | 122.4773 | 0.22278052 | Credible Set | purple |
| chr8:122510562 | 8 | 122.5106 | 0.30110485 | Credible Set | purple |
| chr8:122770060 | 8 | 122.7701 | 0.04056677 | Credible Set | purple |
| chr8:123390979 | 8 | 123.3910 | 0.03759574 | Credible Set | purple |

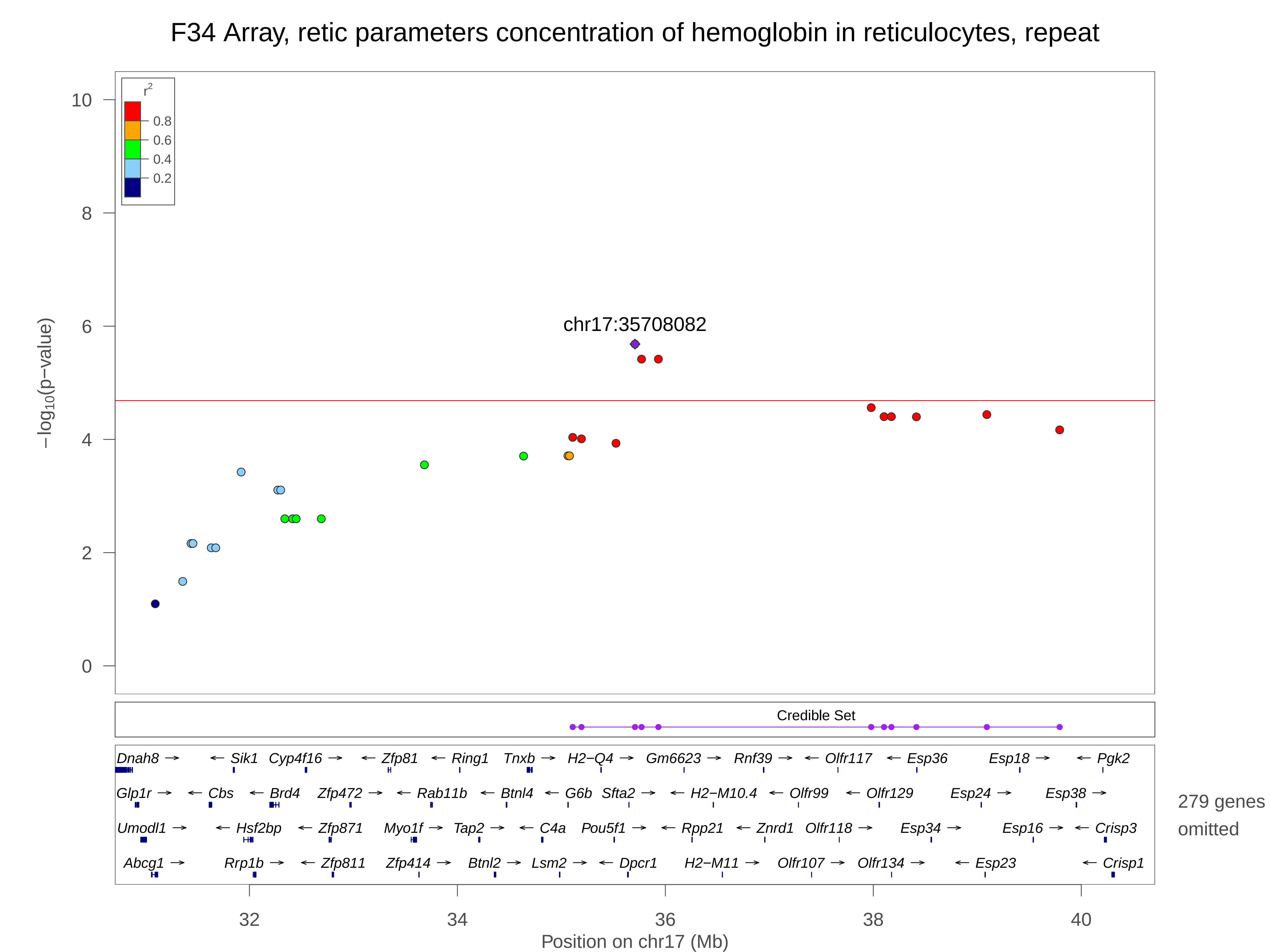

F34 Array, retic parameters concentration of hemoglobin in reticulocytes, repeat
r2
0.8
0.6
0.4
0.2
−log10(p−value)
chr17:35708082
Credible Set
Dnah8 →
← Sik1
Cyp4f16 →
← Zfp81
← Ring1
Tnxb →
H2−Q4 →
Gm6623 →
Rnf39 →
← Olfr117
← Esp36
Esp18 →
← Pgk2
Glp1r →
← Cbs
← Brd4
Zfp472 →
← Rab11b
← Btnl4
← G6b
Sfta2 →
← H2−M10.4
← Olfr99
← Olfr129
Esp24 →
Esp38 →
Umodl1 →
← Hsf2bp
← Zfp871
Myo1f →
Tap2 →
← C4a
Pou5f1 →
← Rpp21
← Znrd1
Olfr118 →
Esp34 →
Esp16 →
← Crisp3
Abcg1 →
Rrp1b →
← Zfp811
Zfp414 →
Btnl2 →
Lsm2 →
← Dpcr1
H2−M11 →
Olfr107 →
Olfr134 →
← Esp23
← Crisp1
279 genes omitted
Position on chr17 (Mb)

| | |
|---|---|
| date: | Thu Oct 25 16:48:13 2018 |
| build: | F34Array |
| display range: | chr17:30708082−40708082 [30708082−40708082] |
| hilite range: | 0 − 0 [ 0 − 0 ] |
| reference SNP: | chr17:35708082 |
| number of SNPs plotted: | 30 |
| min P−value: | 2.07E−6 [chr17:35708082] |
| max P−value: | 8.03E−2 [chr17:31094466] |
| omitted Genes: | Tff3, Tff2, LOC102631757 |
| omitted Genes: | Tff1, Tmprss3, Ubash3a |
| omitted Genes: | Rsph1, Slc37a1, Pde9a |
| omitted Genes: | Wdr4, Ndufv3, Pknox1 |
| omitted Genes: | U2af1, Cryaa, Pdxk−ps |
| omitted Genes: | Notch3, Ephx3, Gm4432 |
| omitted Genes: | Akap8, Akap8l, Wiz |
| omitted Genes: | Rasal3, A530088E08Rik, Pglyrp2 |
| omitted Genes: | Cyp4f39, Cyp4f17, Cyp4f37 |
| omitted Genes: | Cyp4f40, Cyp4f15, Zfp799 |
| omitted Genes: | Zfp870, Cyp4f14, Cyp4f13 |
| omitted Genes: | Cyp4f41−ps, Zfp952, Zfp763 |
| omitted Genes: | 4921501E09Rik, Zfp563, Morc2b |
| omitted Genes: | Olfr55, Olfr239, Gm4461 |
| omitted Genes: | Zfp955a, Olfr63, Zfp955b |
| omitted Genes: | Zfp101, Actl9, Adamts10 |
| omitted Genes: | Pram1, Hnrnpm, March2 |
| omitted Genes: | Angptl4, Kank3, Rps28 |
| omitted Genes: | Ndufa7, Cd320, Kifc1 |
| omitted Genes: | Gm27740, Platr17, BC051226 |
| omitted Genes: | Daxx, Zbtb22, Tapbp |
| omitted Genes: | Rgl2, Pfdn6, Wdr46 |
| omitted Genes: | B3galt4, Rps18, Vps52 |
| omitted Genes: | H2−K2, AA388235, H2−K1 |
| omitted Genes: | Mir219a−1, Mir219c, H2−Ke6 |
| omitted Genes: | Slc39a7, Rxrb, Col11a2 |
| omitted Genes: | BC051537, H2−Oa, Brd2 |
| omitted Genes: | H2−DMa, H2−DMb2, H2−DMb1 |
| omitted Genes: | Psmb9, Tap1, Psmb8 |
| omitted Genes: | H2−Ob, H2−Ab1, H2−Aa |
| omitted Genes: | H2−Eb1, H2−Eb2, Btnl1 |
| omitted Genes: | BC051142, Btnl5−ps, Btnl6 |

Make more plots at http://csg.sph.umich.edu/locuszoom/

Fine−mapping Regions

| snp | chr | pos (Mb) | pp | group | color |
|---|---|---|---|---|---|
| chr17:35109063 | 17 | 35.10906 | 0.01054350 | Credible Set | purple |
| chr17:35193207 | 17 | 35.19321 | 0.00996709 | Credible Set | purple |
| chr17:35708082 | 17 | 35.70808 | 0.39239080 | Credible Set | purple |
| chr17:35771114 | 17 | 35.77111 | 0.21807460 | Credible Set | purple |
| chr17:35932844 | 17 | 35.93284 | 0.21807460 | Credible Set | purple |
| chr17:37979659 | 17 | 37.97966 | 0.03314126 | Credible Set | purple |
| chr17:38103272 | 17 | 38.10327 | 0.02339890 | Credible Set | purple |
| chr17:38174076 | 17 | 38.17408 | 0.02339890 | Credible Set | purple |
| chr17:38414260 | 17 | 38.41426 | 0.02322488 | Credible Set | purple |
| chr17:39091675 | 17 | 39.09168 | 0.02533380 | Credible Set | purple |
| chr17:39792213 | 17 | 39.79221 | 0.01405672 | Credible Set | purple |

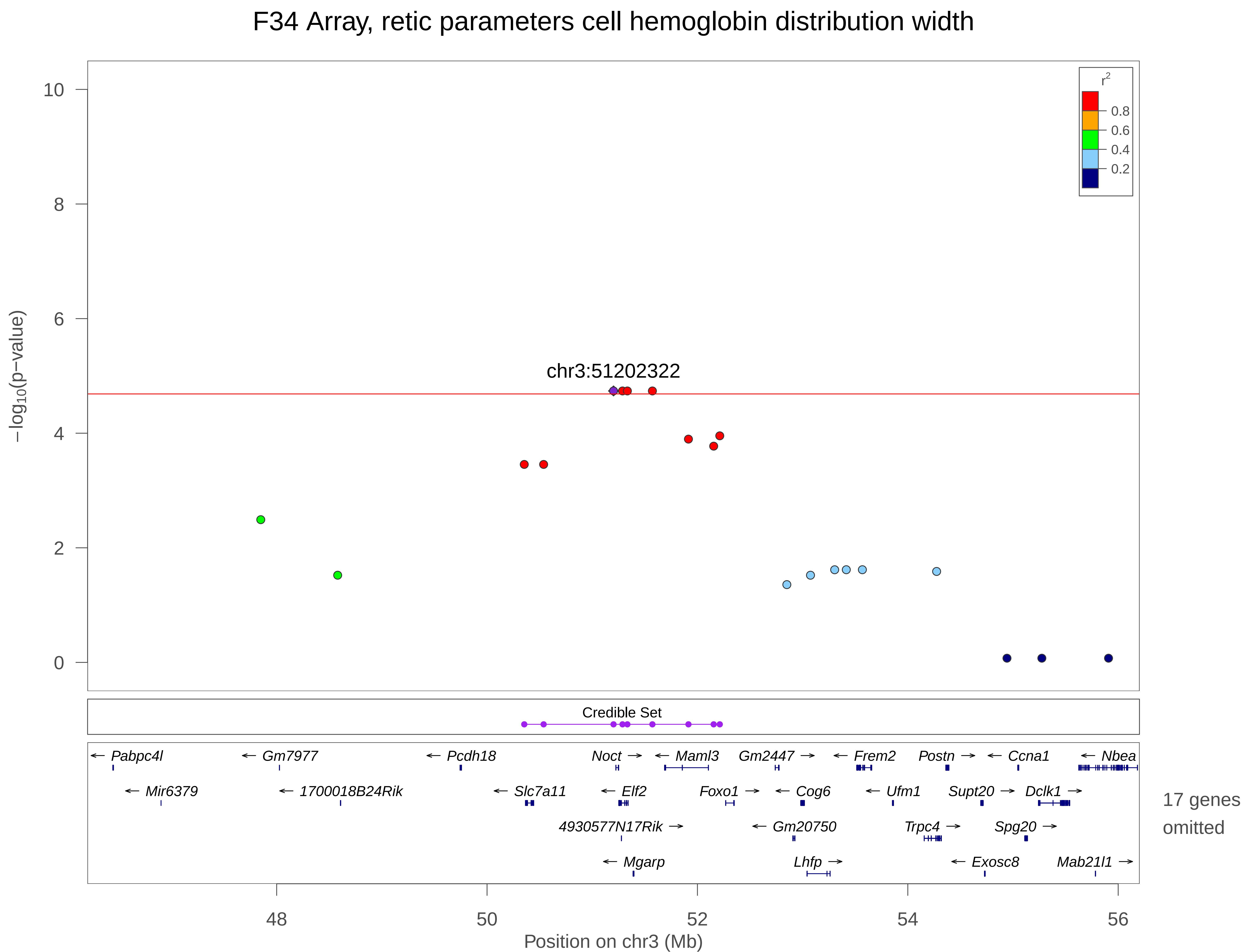

F34 Array, retic parameters cell hemoglobin distribution width
−log10(p−value)
0
2
4
6
8
10
r2
0.8
0.6
0.4
0.2
chr3:51202322
Credible Set
← Pabpc4l
← Gm7977
← Pcdh18
Noct →
← Maml3
Gm2447 →
← Frem2
Postn →
← Ccna1
← Nbea
← Mir6379
← 1700018B24Rik
← Slc7a11
← Elf2
Foxo1 →
← Cog6
← Ufm1
Supt20 →
Dclk1 →
4930577N17Rik →
← Gm20750
Trpc4 →
Spg20 →
← Mgarp
Lhfp →
← Exosc8
Mab21l1 →
17 genes omitted
48
50
52
54
56
Position on chr3 (Mb)

| | |
|---|---|
| date: | Thu Oct 25 16:49:06 2018 |
| build: | F34Array |
| display range: | chr3:46202322−56202322 [46202322−56202322] |
| hilite range: | 0 − 0 [ 0 − 0 ] |
| reference SNP: | chr3:51202322 |
| number of SNPs plotted: | 21 |
| min P−value: | 1.83E−5 [chr3:51202322] |
| max P−value: | 8.44E−1 [chr3:54942736] |
| omitted Genes: | Ndufc1, Naa15, Rab33b |
| omitted Genes: | Setd7, 5031434O11Rik, Mgst2 |
| omitted Genes: | Nhlrc3, Proser1, Stoml3 |
| omitted Genes: | Alg5, Smad9, Rfxap |
| omitted Genes: | Sertm1, 4931419H13Rik, Ccdc169 |
| omitted Genes: | Sohlh2, 4933417G07Rik |

Make more plots at http://csg.sph.umich.edu/locuszoom/

Fine−mapping Regions

| snp | chr | pos (Mb) | pp | group | color |
|---|---|---|---|---|---|
| chr3:50353748 | 3 | 50.35375 | 0.01339687 | Credible Set | purple |
| chr3:50537878 | 3 | 50.53788 | 0.01339687 | Credible Set | purple |
| chr3:51202322 | 3 | 51.20232 | 0.21806426 | Credible Set | purple |
| chr3:51287862 | 3 | 51.28786 | 0.21806426 | Credible Set | purple |
| chr3:51333971 | 3 | 51.33397 | 0.21806426 | Credible Set | purple |
| chr3:51571920 | 3 | 51.57192 | 0.21806426 | Credible Set | purple |
| chr3:51914573 | 3 | 51.91457 | 0.03481626 | Credible Set | purple |
| chr3:52154479 | 3 | 52.15448 | 0.02667820 | Credible Set | purple |
| chr3:52212374 | 3 | 52.21237 | 0.03945476 | Credible Set | purple |

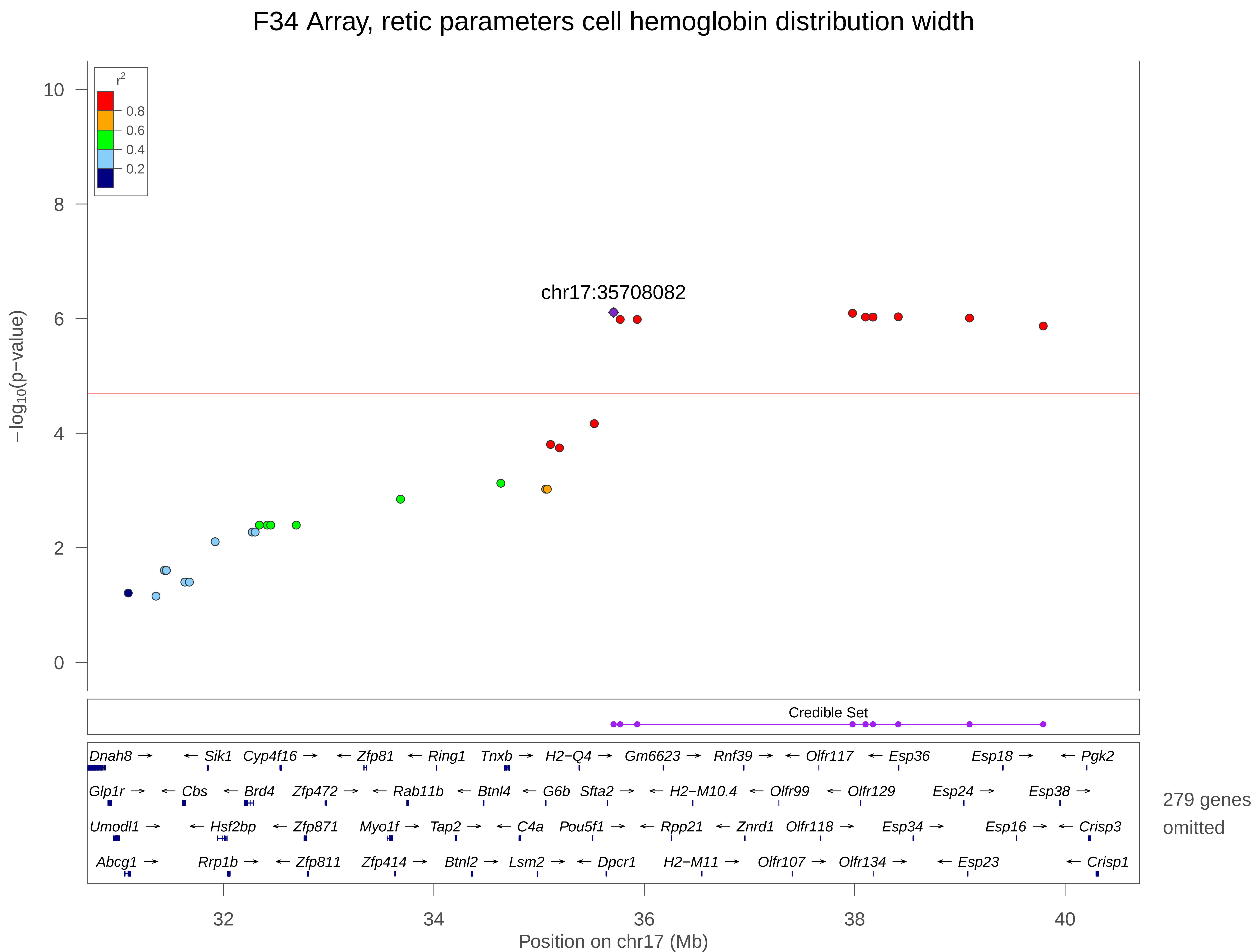

F34 Array, retic parameters cell hemoglobin distribution width
r2
0.8
0.6
0.4
0.2
chr17:35708082
−log10(p−value)
10
8
6
4
2
0
Credible Set
Dnah8 →
← Sik1
Cyp4f16 →
← Zfp81
← Ring1
Tnxb →
H2−Q4 →
Gm6623 →
Rnf39 →
← Olfr117
← Esp36
Esp18 →
← Pgk2
Glp1r →
← Cbs
← Brd4
Zfp472 →
← Rab11b
← Btnl4
← G6b
Sfta2 →
← H2−M10.4
← Olfr99
← Olfr129
Esp24 →
Esp38 →
Umodl1 →
← Hsf2bp
← Zfp871
Myo1f →
Tap2 →
← C4a
Pou5f1 →
← Rpp21
← Znrd1
Olfr118 →
Esp34 →
Esp16 →
← Crisp3
Abcg1 →
Rrp1b →
← Zfp811
Zfp414 →
Btnl2 →
Lsm2 →
← Dpcr1
H2−M11 →
Olfr107 →
Olfr134 →
← Esp23
← Crisp1
279 genes omitted
32
34
36
38
40
Position on chr17 (Mb)

date: Thu Oct 25 16:49:00 2018
build: F34Array
display range: chr17:30708082−40708082 [30708082−40708082]
hilite range: 0 − 0 [ 0 − 0 ]
reference SNP: chr17:35708082
number of SNPs plotted: 30
min P−value: 7.77E−7 [chr17:35708082]
max P−value: 6.97E−2 [chr17:31358358]
omitted Genes: Tff3, Tff2, LOC102631757
omitted Genes: Tff1, Tmprss3, Ubash3a
omitted Genes: Rsph1, Slc37a1, Pde9a
omitted Genes: Wdr4, Ndufv3, Pknox1
omitted Genes: U2af1, Cryaa, Pdxk−ps
omitted Genes: Notch3, Ephx3, Gm4432
omitted Genes: Akap8, Akap8l, Wiz
omitted Genes: Rasal3, A530088E08Rik, Pglyrp2
omitted Genes: Cyp4f39, Cyp4f17, Cyp4f37
omitted Genes: Cyp4f40, Cyp4f15, Zfp799
omitted Genes: Zfp870, Cyp4f14, Cyp4f13
omitted Genes: Cyp4f41−ps, Zfp952, Zfp763
omitted Genes: 4921501E09Rik, Zfp563, Morc2b
omitted Genes: Olfr55, Olfr239, Gm4461
omitted Genes: Zfp955a, Olfr63, Zfp955b
omitted Genes: Zfp101, Actl9, Adamts10
omitted Genes: Pram1, Hnrnpm, March2
omitted Genes: Angptl4, Kank3, Rps28
omitted Genes: Ndufa7, Cd320, Kifc1
omitted Genes: Gm27740, Platr17, BC051226
omitted Genes: Daxx, Zbtb22, Tapbp
omitted Genes: Rgl2, Pfdn6, Wdr46
omitted Genes: B3galt4, Rps18, Vps52
omitted Genes: H2−K2, AA388235, H2−K1
omitted Genes: Mir219a−1, Mir219c, H2−Ke6
omitted Genes: Slc39a7, Rxrb, Col11a2
omitted Genes: BC051537, H2−Oa, Brd2
omitted Genes: H2−DMa, H2−DMb2, H2−DMb1
omitted Genes: Psmb9, Tap1, Psmb8
omitted Genes: H2−Ob, H2−Ab1, H2−Aa
omitted Genes: H2−Eb1, H2−Eb2, Btnl1
Make more plots at http://csg.sph.umich.edu/locuszoom/
omitted Genes: BC051142, Btnl5−ps, Btnl6

Fine−mapping Regions

| snp | chr | pos (Mb) | pp | group | color |
|---|---|---|---|---|---|
| chr17:35708082 | 17 | 35.70808 | 0.13502166 | Credible Set | purple |
| chr17:35771114 | 17 | 35.77111 | 0.10257697 | Credible Set | purple |
| chr17:35932844 | 17 | 35.93284 | 0.10257697 | Credible Set | purple |
| chr17:37979659 | 17 | 37.97966 | 0.13031025 | Credible Set | purple |
| chr17:38103272 | 17 | 38.10327 | 0.11243731 | Credible Set | purple |
| chr17:38174076 | 17 | 38.17408 | 0.11243731 | Credible Set | purple |
| chr17:38414260 | 17 | 38.41426 | 0.11339820 | Credible Set | purple |
| chr17:39091675 | 17 | 39.09168 | 0.10819998 | Credible Set | purple |
| chr17:39792213 | 17 | 39.79221 | 0.07956072 | Credible Set | purple |

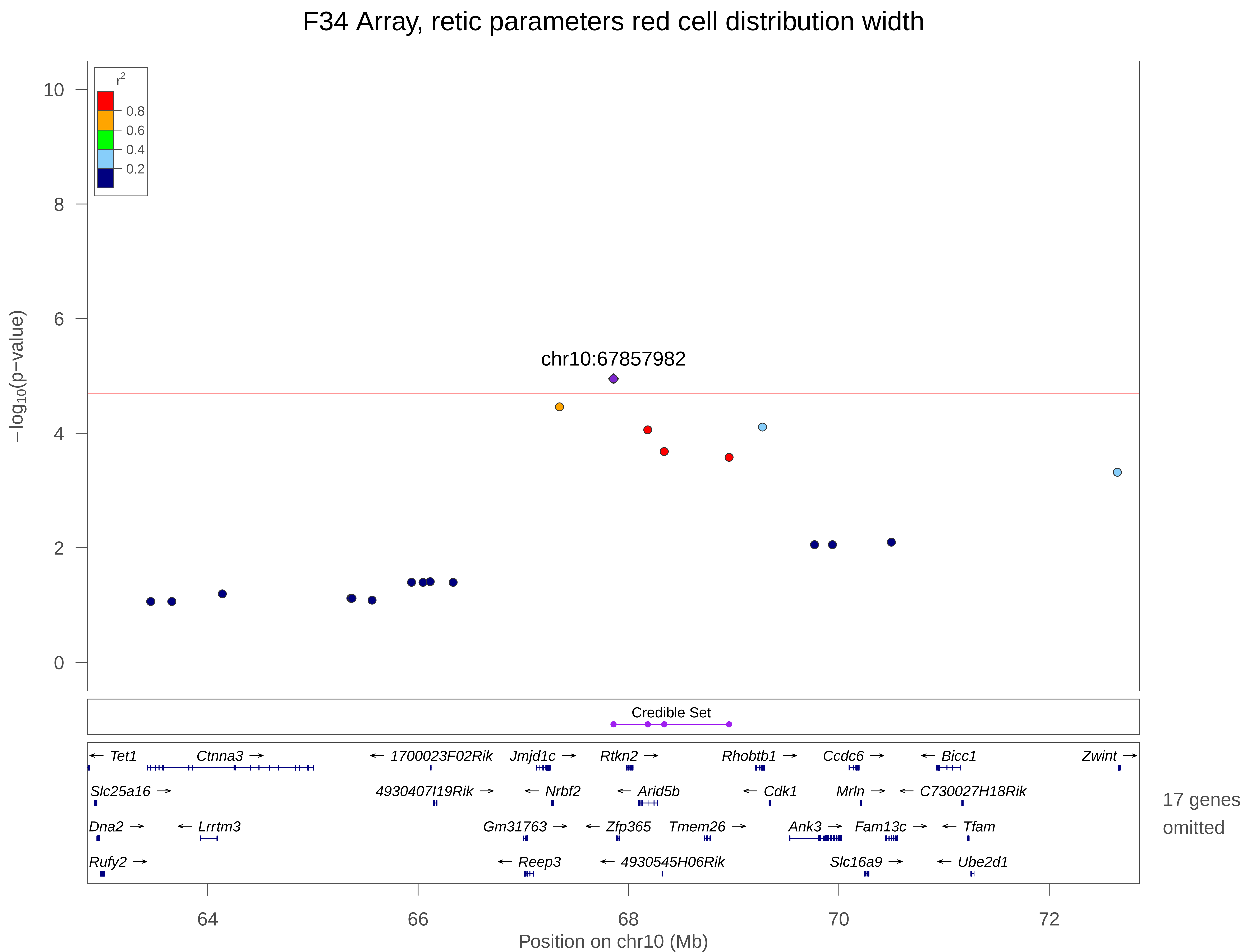

F34 Array, retic parameters red cell distribution width
r2
0.8
0.6
0.4
0.2
10
8
6
4
2
0
−log10(p−value)
chr10:67857982
Credible Set
← Tet1
Ctnna3 →
← 1700023F02Rik
Jmjd1c →
Rtkn2 →
Rhobtb1 →
Ccdc6 →
← Bicc1
Zwint →
Slc25a16 →
4930407I19Rik →
← Nrbf2
← Arid5b
← Cdk1
Mrln →
← C730027H18Rik
Dna2 →
← Lrrtm3
Gm31763 →
← Zfp365
Tmem26 →
Ank3 →
Fam13c →
← Tfam
Rufy2 →
← Reep3
← 4930545H06Rik
Slc16a9 →
← Ube2d1
17 genes omitted
64
66
68
70
72
Position on chr10 (Mb)

| | |
|---|---|
| date: | Thu Oct 25 16:49:14 2018 |
| build: | F34Array |
| display range: | chr10:62857982−72857982 [62857982−72857982] |
| hilite range: | 0 − 0 [ 0 − 0 ] |
| reference SNP: | chr10:67857982 |
| number of SNPs plotted: | 21 |
| min P−value: | 1.12E−5 [chr10:67857982] |
| max P−value: | 8.65E−2 [chr10:63458094] |
| omitted Genes: | Hnrnph3, Pbld2, Pbld1 |
| omitted Genes: | Atoh7, Mypn, Herc4 |
| omitted Genes: | Sirt1, Dnajc12, Egr2 |
| omitted Genes: | Ado, 1700040L02Rik, Phyhipl |
| omitted Genes: | D630013N20Rik, 1700113B09Rik, Cisd1 |
| omitted Genes: | Ipmk, 1700049L16Rik |

Make more plots at http://csg.sph.umich.edu/locuszoom/

Fine−mapping Regions

| snp | chr | pos (Mb) | pp | group | color |
|---|---|---|---|---|---|
| chr10:67857982 | 10 | 67.85798 | 0.79629461 | Credible Set | purple |
| chr10:68183490 | 10 | 68.18349 | 0.11355990 | Credible Set | purple |
| chr10:68340621 | 10 | 68.34062 | 0.04993989 | Credible Set | purple |
| chr10:68956755 | 10 | 68.95676 | 0.04020560 | Credible Set | purple |

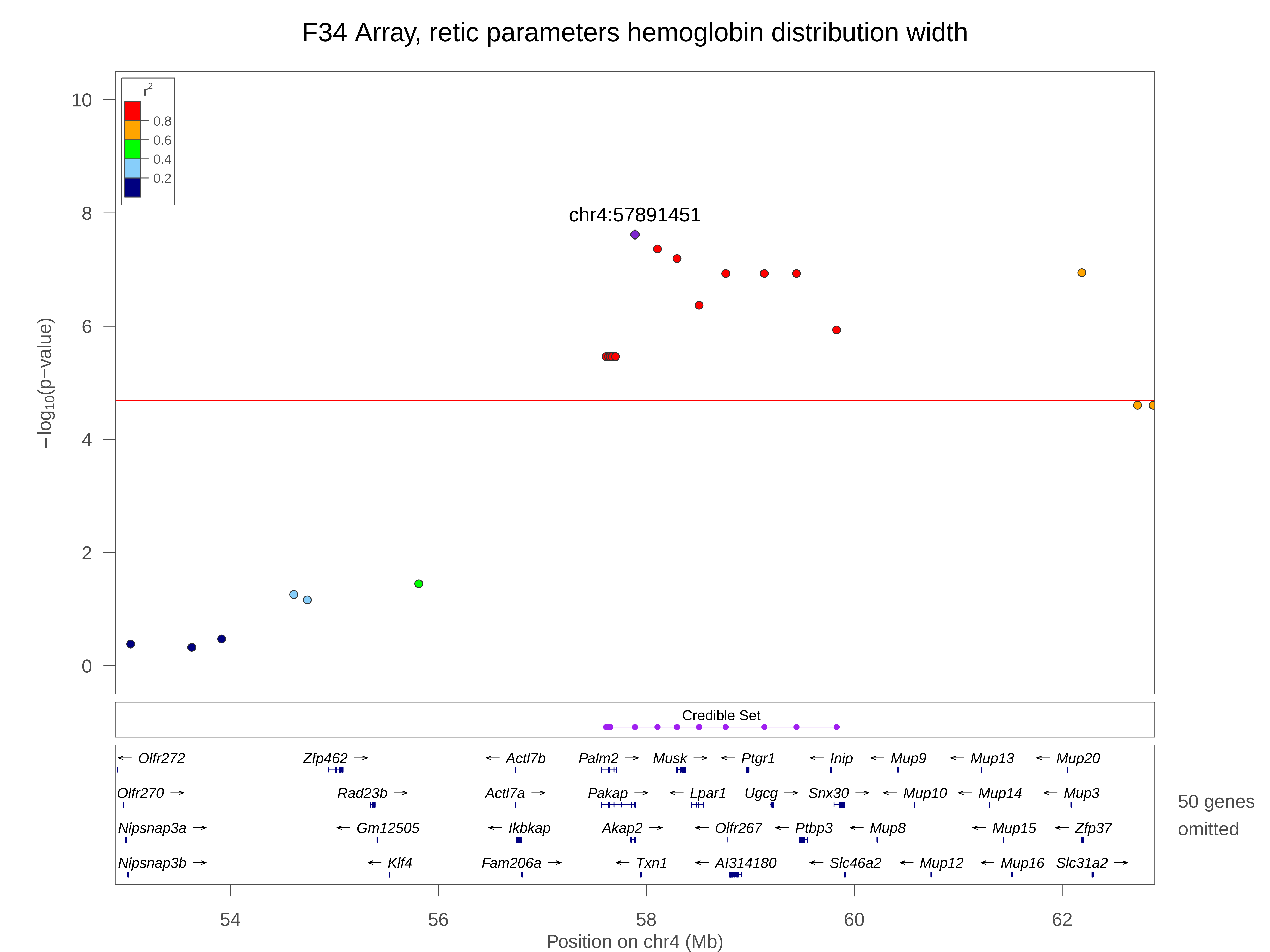

F34 Array, retic parameters hemoglobin distribution width
r2
0.8
0.6
0.4
0.2
chr4:57891451
−log10(p−value)
0
2
4
6
8
10
Credible Set
← Olfr272
Zfp462 →
← Actl7b
Palm2 →
Musk →
← Ptgr1
← Inip
← Mup9
← Mup13
← Mup20
Olfr270 →
Rad23b →
Actl7a →
Pakap →
← Lpar1
Ugcg →
Snx30 →
← Mup10
← Mup14
← Mup3
Nipsnap3a →
← Gm12505
← Ikbkap
Akap2 →
← Olfr267
← Ptbp3
← Mup8
← Mup15
← Zfp37
Nipsnap3b →
← Klf4
Fam206a →
← Txn1
← AI314180
← Slc46a2
← Mup12
← Mup16
Slc31a2 →
50 genes omitted
54
56
58
60
62
Position on chr4 (Mb)

| | |
|---|---|
| date: | Thu Oct 25 16:49:21 2018 |
| build: | F34Array |
| display range: | chr4:52891451−62891451 [52891451−62891451] |
| hilite range: | 0 − 0 [ 0 − 0 ] |
| reference SNP: | chr4:57891451 |
| number of SNPs plotted: | 24 |
| min P−value: | 2.41E−8 [chr4:57891451] |
| max P−value: | 4.69E−1 [chr4:53628462] |
| omitted Genes: | Abca1, 4930522O17Rik, AI427809 |
| omitted Genes: | Slc44a1, Fsd1l, Fktn |
| omitted Genes: | Tal2, Tmem38b, Ctnnal1 |
| omitted Genes: | Tmem245, Mir32, Frrs1l |
| omitted Genes: | Epb41l4b, Gm12530, Ptpn3 |
| omitted Genes: | 1700042G15Rik, D630039A03Rik, Txndc8 |
| omitted Genes: | Svep1, Mir3095, Zkscan16 |
| omitted Genes: | Dnajc25, Gng10, Gm12596 |
| omitted Genes: | Susd1, Hsdl2, E130308A19Rik |
| omitted Genes: | Mup4, Mup6, Mup7 |
| omitted Genes: | Mup2, Mup1, Mup11 |
| omitted Genes: | Mup17, Mup18, Mup19 |
| omitted Genes: | Mup5, Mup21, Fkbp15 |
| omitted Genes: | Slc31a1, Cdc26, Prpf4 |
| omitted Genes: | Rnf183, Wdr31, Bspry |
| omitted Genes: | Hdhd3, Alad, Pole3 |
| omitted Genes: | 4933430I17Rik, Rgs3 |

Make more plots at http://csg.sph.umich.edu/locuszoom/

Fine−mapping Regions

| snp | chr | pos (Mb) | pp | group | color |
|---|---|---|---|---|---|
| chr4:57613710 | 4 | 57.61371 | 0.003029939 | Credible Set | purple |
| chr4:57637755 | 4 | 57.63775 | 0.003029939 | Credible Set | purple |
| chr4:57652730 | 4 | 57.65273 | 0.003029939 | Credible Set | purple |
| chr4:57891451 | 4 | 57.89145 | 0.365446528 | Credible Set | purple |
| chr4:58107891 | 4 | 58.10789 | 0.207204646 | Credible Set | purple |
| chr4:58295311 | 4 | 58.29531 | 0.141999469 | Credible Set | purple |
| chr4:58507849 | 4 | 58.50785 | 0.022635370 | Credible Set | purple |
| chr4:58764292 | 4 | 58.76429 | 0.078645589 | Credible Set | purple |
| chr4:59135384 | 4 | 59.13538 | 0.078645589 | Credible Set | purple |
| chr4:59443996 | 4 | 59.44400 | 0.078645589 | Credible Set | purple |
| chr4:59830858 | 4 | 59.83086 | 0.008597585 | Credible Set | purple |

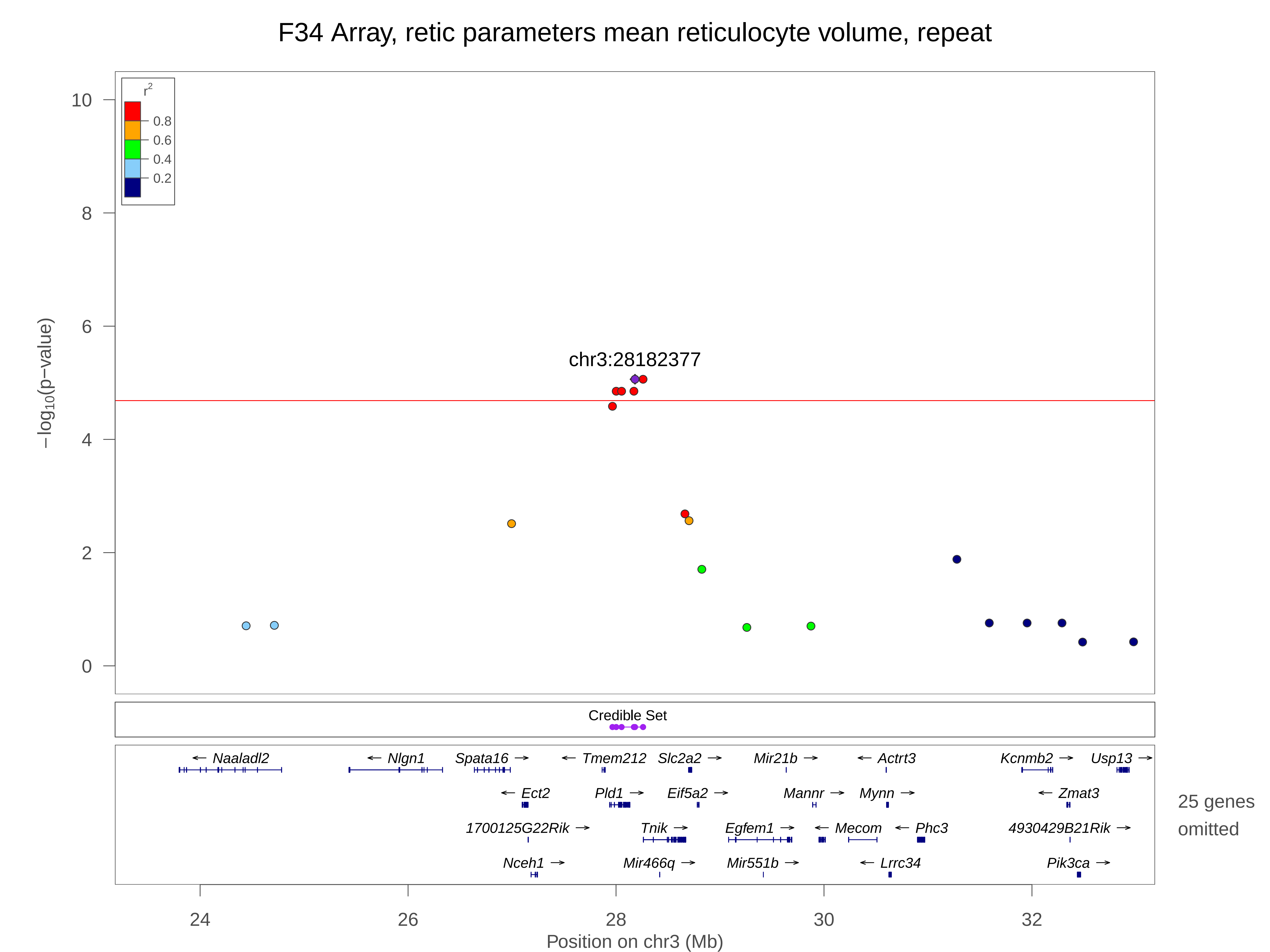
F34 Array, retic parameters mean reticulocyte volume, repeat
r2
0.8
0.6
0.4
0.2
−log10(p−value)
chr3:28182377
Credible Set
Naaladl2
Nlgn1
Spata16
Tmem212
Slc2a2
Mir21b
Actrt3
Kcnmb2
Usp13
Ect2
Pld1
Eif5a2
Mannr
Mynn
Zmat3
1700125G22Rik
Tnik
Egfem1
Mecom
Phc3
4930429B21Rik
Nceh1
Mir466q
Mir551b
Lrrc34
Pik3ca
25 genes omitted
Position on chr3 (Mb)

| | |
|---|---|
| date: | Thu Oct 25 16:49:46 2018 |
| build: | F34Array |
| display range: | chr3:23182377−33182377 [23182377−33182377] |
| hilite range: | 0 − 0 [ 0 − 0 ] |
| reference SNP: | chr3:28182377 |
| number of SNPs plotted: | 21 |
| min P−value: | 8.66E−6 [chr3:28182377] |
| max P−value: | 3.8E−1 [chr3:32486921] |
| omitted Genes: | Tnfsf10, Ghsr, Fndc3b |
| omitted Genes: | Mir3092, Rpl22l1, Gm1527 |
| omitted Genes: | Lrriq4, Lrrc31, Samd7 |
| omitted Genes: | Sec62, Gpr160, Prkci |
| omitted Genes: | Mir7008, Skil, Cldn11 |
| omitted Genes: | Slc7a14, Kcnmb3, Zfp639 |
| omitted Genes: | Mfn1, Gnb4, Actl6a |
| omitted Genes: | Mrpl47, Ndufb5, Pex5l |
| omitted Genes: | 4930419G24Rik |

Make more plots at http://csg.sph.umich.edu/locuszoom/

Fine−mapping Regions

| snp | chr | pos (Mb) | pp | group | color |
|---|---|---|---|---|---|
| chr3:27965717 | 3 | 27.96572 | 0.08266309 | Credible Set | purple |
| chr3:28001696 | 3 | 28.00170 | 0.14814397 | Credible Set | purple |
| chr3:28053395 | 3 | 28.05339 | 0.14814397 | Credible Set | purple |
| chr3:28171809 | 3 | 28.17181 | 0.14814397 | Credible Set | purple |
| chr3:28182377 | 3 | 28.18238 | 0.23576811 | Credible Set | purple |
| chr3:28259887 | 3 | 28.25989 | 0.23576811 | Credible Set | purple |

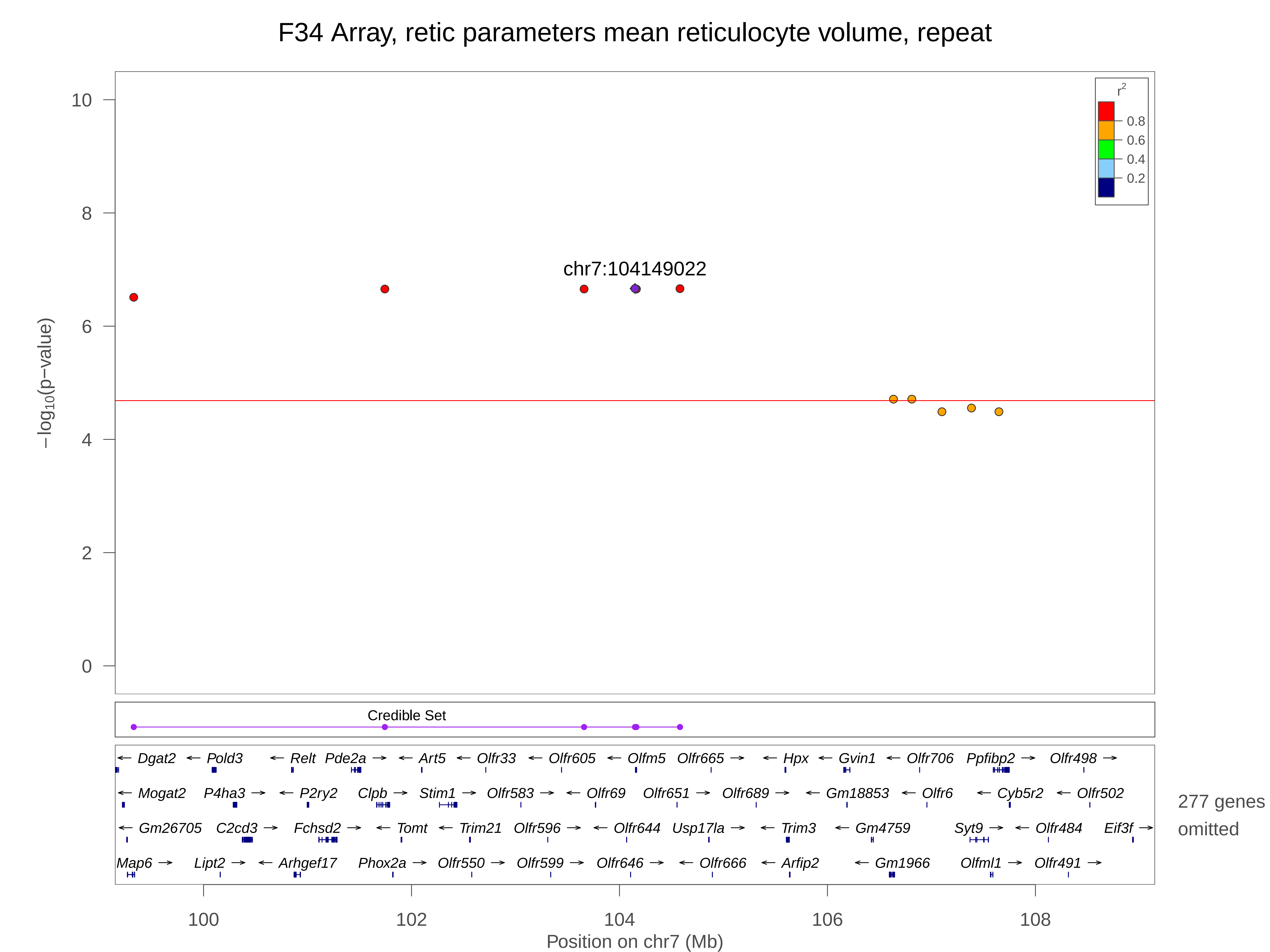

F34 Array, retic parameters mean reticulocyte volume, repeat
chr7:104149022
−log10(p−value)
r2
0.8
0.6
0.4
0.2
Credible Set
Dgat2
Pold3
Relt
Pde2a
Art5
Olfr33
Olfr605
Olfm5
Olfr665
Hpx
Gvin1
Olfr706
Ppfibp2
Olfr498
Mogat2
P4ha3
P2ry2
Clpb
Stim1
Olfr583
Olfr69
Olfr651
Olfr689
Gm18853
Olfr6
Cyb5r2
Olfr502
Gm26705
C2cd3
Fchsd2
Tomt
Trim21
Olfr596
Olfr644
Usp17la
Trim3
Gm4759
Syt9
Olfr484
Eif3f
Map6
Lipt2
Arhgef17
Phox2a
Olfr550
Olfr599
Olfr646
Olfr666
Arfip2
Gm1966
Olfml1
Olfr491
277 genes omitted
Position on chr7 (Mb)

date: Thu Oct 25 16:49:59 2018
build: F34Array
display range: chr7:99149022−109149022 [99149022−109149022]
hilite range: 0 − 0 [ 0 − 0 ]
reference SNP: chr7:104149022
number of SNPs plotted: 12
min P−value: 2.16E−7 [chr7:104149022]
max P−value: 3.24E−5 [chr7:107100725]
omitted Genes: Serpinh1, Gdpd5, Klhl35
omitted Genes: Rps3, Snord15b, Snord15a
omitted Genes: Arrb1, Mir326, Tpbgl
omitted Genes: Slco2b1, Olfr520, Olfr521
omitted Genes: Neu3, Spcs2, Xrra1
omitted Genes: Rnf169, Chrdl2, Kcne3
omitted Genes: Pgm2l1, Gpx2−ps1, Ppme1
omitted Genes: Ucp3, Ucp2, Dnajb13
omitted Genes: Coa4, Mrpl48, Rab6a
omitted Genes: Plekhb1, Fam168a, Mir3102
omitted Genes: P2ry6, Atg16l2, Stard10
omitted Genes: Arap1, Mir139, Art2a−ps
omitted Genes: Art2b, Inppl1, Folr2
omitted Genes: Folr1, Anapc15, Lamtor1
omitted Genes: Lrrc51, Numa1, Il18bp
omitted Genes: Rnf121, Xndc1, Xntrpc
omitted Genes: Trpc2, Art1, Chrna10
omitted Genes: Nup98, Pgap2, Rhog
omitted Genes: Rrm1, Olfr543, Olfr544
omitted Genes: Olfr545, Olfr547, Olfr549
omitted Genes: Olfr551, Olfr552, Olfr553
omitted Genes: Olfr554, Olfr555, Olfr556
omitted Genes: Trim68, Olfr557, Olfr558
omitted Genes: Olfr559, Olfr78, Olfr560
omitted Genes: Olfr561, Olfr564, Olfr566
omitted Genes: Olfr568, Olfr569, Olfr570
omitted Genes: Olfr571, Olfr572, Olfr574
omitted Genes: Olfr575, Olfr576, Olfr577
omitted Genes: Olfr578, Olfr582, Olfr584
omitted Genes: Olfr585, Olfr586, Olfr589
omitted Genes: Olfr591, Olfr592, Olfr593
Make more plots at http://csg.sph.umich.edu/locuszoom/
omitted Genes: Olfr594, Usp17ld, Olfr597

Fine−mapping Regions

| snp | chr | pos (Mb) | pp | group | color |
|---|---|---|---|---|---|
| chr7:99328376 | 7 | 99.32838 | 0.1256159 | Credible Set | purple |
| chr7:101743401 | 7 | 101.74340 | 0.1736837 | Credible Set | purple |
| chr7:103659355 | 7 | 103.65936 | 0.1736837 | Credible Set | purple |
| chr7:104149022 | 7 | 104.14902 | 0.1770891 | Credible Set | purple |
| chr7:104162444 | 7 | 104.16244 | 0.1736837 | Credible Set | purple |
| chr7:104581631 | 7 | 104.58163 | 0.1762440 | Credible Set | purple |

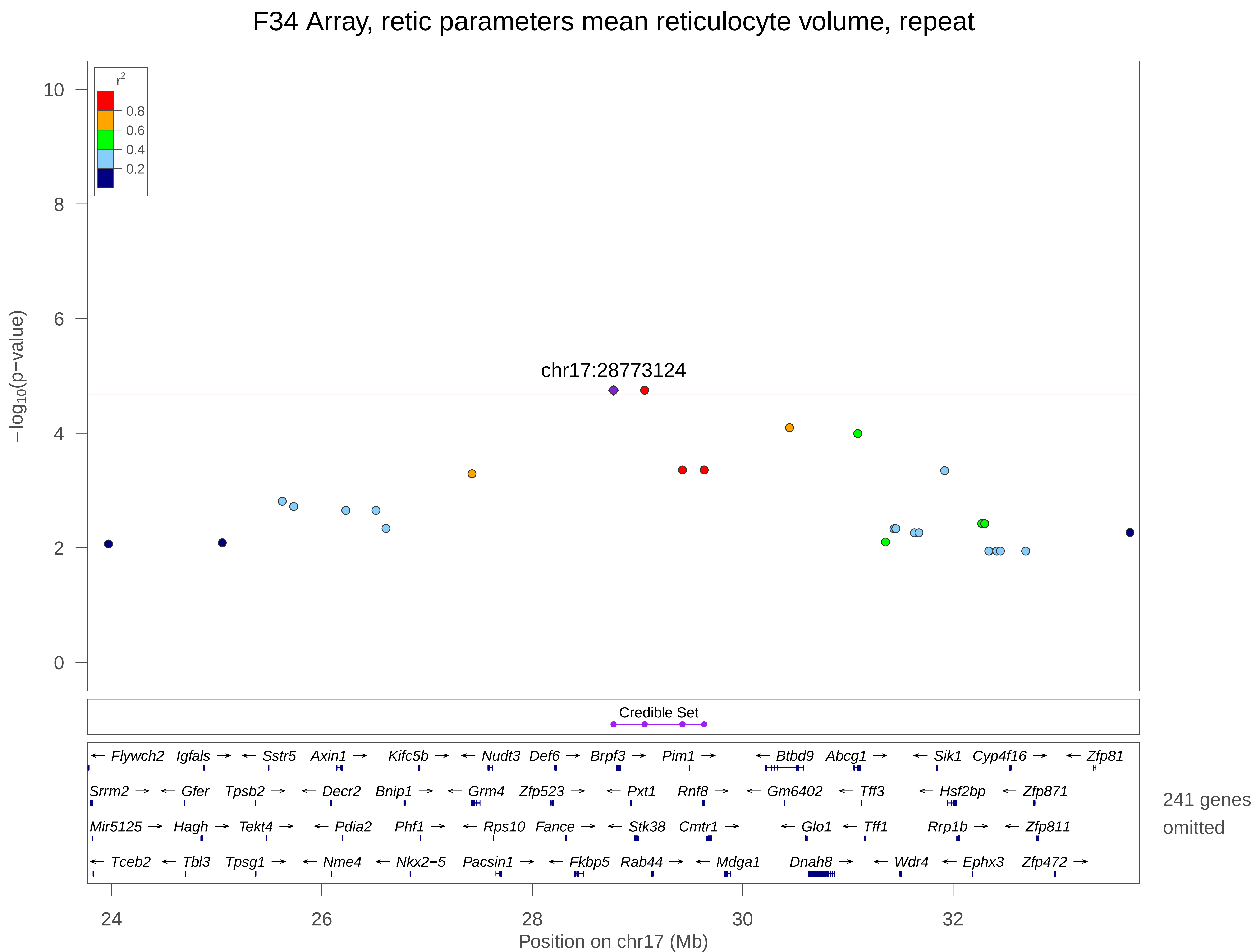

F34 Array, retic parameters mean reticulocyte volume, repeat
r2
0.8
0.6
0.4
0.2
10
8
6
4
2
0
−log10(p−value)
chr17:28773124
Credible Set
← Flywch2 Igfals → ← Sstr5 Axin1 → Kifc5b → ← Nudt3 Def6 → Brpf3 → Pim1 → ← Btbd9 Abcg1 → ← Sik1 Cyp4f16 → ← Zfp81
Srrm2 → ← Gfer Tpsb2 → ← Decr2 Bnip1 → ← Grm4 Zfp523 → ← Pxt1 Rnf8 → ← Gm6402 ← Tff3 ← Hsf2bp ← Zfp871
Mir5125 → Hagh → Tekt4 → ← Pdia2 Phf1 → ← Rps10 Fance → ← Stk38 Cmtr1 → ← Glo1 ← Tff1 Rrp1b → ← Zfp811
← Tceb2 ← Tbl3 Tpsg1 → ← Nme4 ← Nkx2−5 Pacsin1 → ← Fkbp5 Rab44 → ← Mdga1 Dnah8 → ← Wdr4 ← Ephx3 Zfp472 →
241 genes omitted
24
26
28
30
32
Position on chr17 (Mb)

date: Thu Oct 25 16:49:33 2018
build: F34Array
display range: chr17:23773124−33773124 [23773124−33773124]
hilite range: 0 − 0 [ 0 − 0 ]
reference SNP: chr17:28773124
number of SNPs plotted: 28
min P−value: 1.78E−5 [chr17:28773124]
max P−value: 1.14E−2 [chr17:32340563]
omitted Genes: Prss33, Prss41, Prss32
omitted Genes: Prss21, Dcpp1, Dcpp2
omitted Genes: Dcpp3, Sbp, Sbpl
omitted Genes: Prss30, Prss22, Prss27
omitted Genes: Kctd5, Pdpk1, Amdhd2
omitted Genes: Atp6v0c, Tbc1d24, Ntn3
omitted Genes: 1600002H07Rik, Ccnf, Mir5134
omitted Genes: Mir6965, Abca17, Abca3
omitted Genes: D330041H03Rik, Rnps1, Eci1
omitted Genes: Dnase1l2, E4f1, Pgp
omitted Genes: Mlst8, Bricd5, Caskin1
omitted Genes: Traf7, Rab26os, Rab26
omitted Genes: Pkd1, Tsc2, Nthl1
omitted Genes: Slc9a3r2, Npw, Zfp598
omitted Genes: Syngr3, Noxo1, Rnf151
omitted Genes: Snhg9, Snora78, Rps2
omitted Genes: Snora64, Ndufb10, Rpl3l
omitted Genes: Msrb1, Hs3st6, Meiob
omitted Genes: Fahd1, Nubp2, Spsb3
omitted Genes: Eme2, Mrps34, Nme3
omitted Genes: Mapk8ip3, Hn1l, Cramp1l
omitted Genes: Ift140, Tmem204, Telo2
omitted Genes: Ptx4, Clcn7, Ccdc154
omitted Genes: BC003965, Unkl, Gnptg
omitted Genes: Tsr3, Baiap3, Mir3547
omitted Genes: Ube2i, Gm17801, Prss34
omitted Genes: Prss28, Prss29, Tpsab1
omitted Genes: Cacna1h, Sox8, 2810468N07Rik
omitted Genes: Lmf1, Gng13, Chtf18
omitted Genes: Rpusd1, Mslnl, Msln
omitted Genes: Narfl, Haghl, Mir6966
Make more plots at http://csg.sph.umich.edu/locuszoom/
omitted Genes: Ccdc78, Fam173a, Metrn

Fine−mapping Regions

| snp | chr | pos (Mb) | pp | group | color |
|---|---|---|---|---|---|
| chr17:28773124 | 17 | 28.77312 | 0.47684602 | Credible Set | purple |
| chr17:29068384 | 17 | 29.06838 | 0.47684602 | Credible Set | purple |
| chr17:29427987 | 17 | 29.42799 | 0.02315398 | Credible Set | purple |
| chr17:29633989 | 17 | 29.63399 | 0.02315398 | Credible Set | purple |

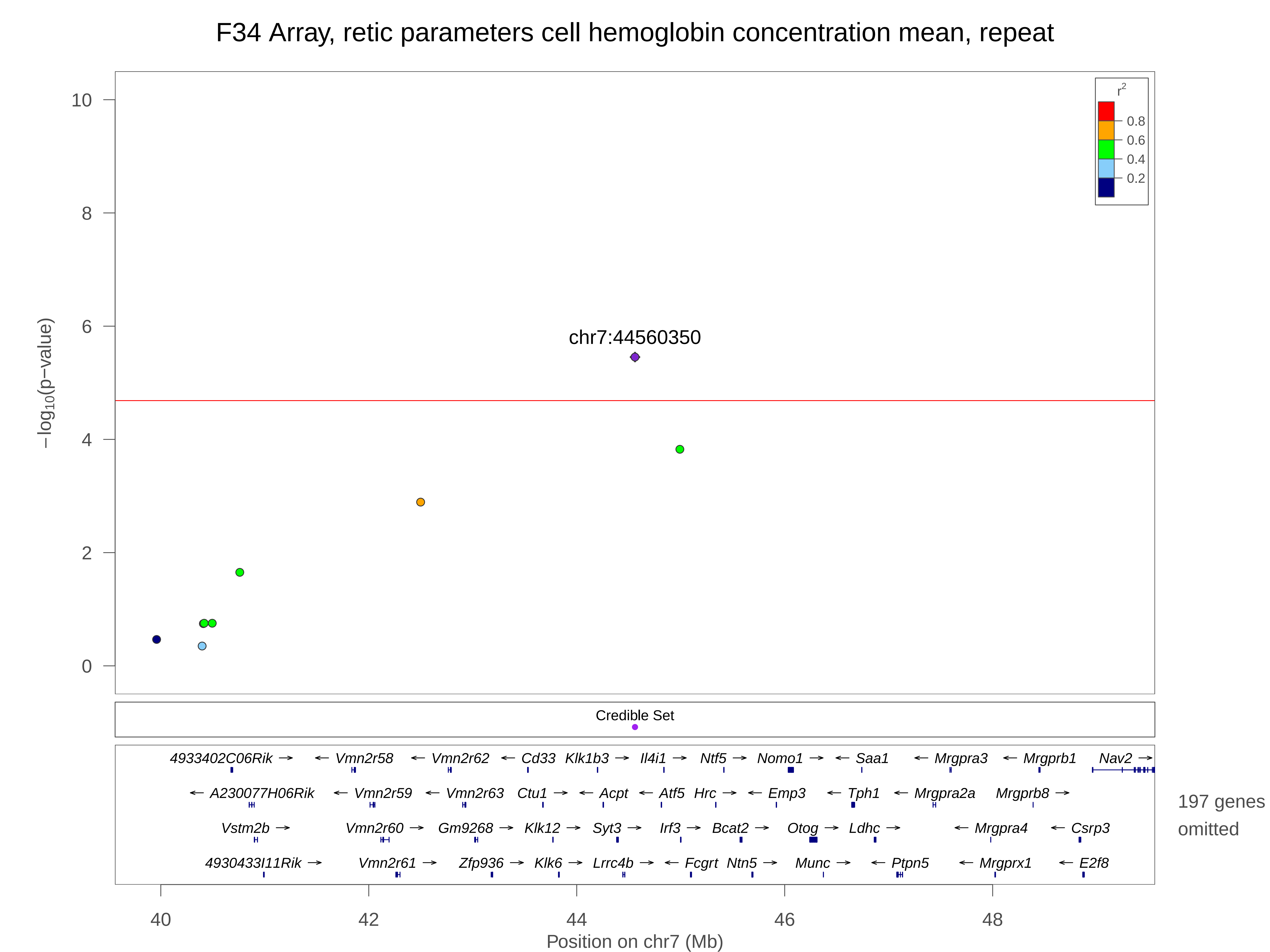

F34 Array, retic parameters cell hemoglobin concentration mean, repeat
−log10(p−value)
chr7:44560350
r2
0.8
0.6
0.4
0.2
Credible Set
4933402C06Rik → ← Vmn2r58 ← Vmn2r62 ← Cd33 Klk1b3 → Il4i1 → Ntf5 → Nomo1 → ← Saa1 ← Mrgpra3 ← Mrgprb1 Nav2 →
← A230077H06Rik ← Vmn2r59 ← Vmn2r63 Ctu1 → ← Acpt ← Atf5 Hrc → ← Emp3 ← Tph1 ← Mrgpra2a Mrgprb8 →
Vstm2b → Vmn2r60 → Gm9268 → Klk12 → Syt3 → Irf3 → Bcat2 → Otog → Ldhc → ← Mrgpra4 ← Csrp3
4930433I11Rik → Vmn2r61 → Zfp936 → Klk6 → Lrrc4b → ← Fcgrt Ntf5 → Munc → ← Ptpn5 ← Mrgprx1 ← E2f8
197 genes omitted
40 42 44 46 48
Position on chr7 (Mb)

date: Thu Oct 25 16:50:26 2018
build: F34Array
display range: chr7:39560350−49560350 [39560350−49560350]
hilite range: 0 − 0 [ 0 − 0 ]
reference SNP: chr7:44560350
number of SNPs plotted: 10
min P−value: 3.51E−6 [chr7:44560350]
max P−value: 4.46E−1 [chr7:40397482]
omitted Genes: Gm4884, 1700110I07Rik, Gm5592
omitted Genes: AI987944, Vmn2r57, AW146154
omitted Genes: Zfp788, Vmn2r−ps54, Vmn2r−ps60
omitted Genes: 4933421I07Rik, Zfp141, 9830147E19Rik
omitted Genes: Gm5595, Gm2381, EU599041
omitted Genes: Zfp715, Siglecf, Siglecg
omitted Genes: Lim2, Nkg7, Cldnd2
omitted Genes: Etfb, Mir7051, Vsig10l
omitted Genes: Iglon5, 4931406B18Rik, Zfp658
omitted Genes: Zfp719, Zfp819, Ceacam18
omitted Genes: Siglece, Klk14, Klk13
omitted Genes: 2310002F09Rik, Klk11, Klk10
omitted Genes: Klk9, Klk8, Klk7
omitted Genes: Klk5, Klk4, Klk15
omitted Genes: Klk1b7−ps, Klk1b8, Klk1b1
omitted Genes: Klk1b9, Klk1b11, Klk1b26
omitted Genes: Klk1b27, Klk1b21, Klk1b22
omitted Genes: Klk1b16, Klk1b24, Klk1b4
omitted Genes: Klk1b5, Klk1, 1700028J19Rik
omitted Genes: 2410002F23Rik, Snord88c, Snord88a
omitted Genes: Clec11a, Shank1, 1700008O03Rik
omitted Genes: Aspdh, Josd2, Mir7052
omitted Genes: 5430431A17Rik, Emc10, Fam71e1
omitted Genes: Mybpc2, Spib, Pold1
omitted Genes: Mir7053, Nr1h2, Napsa
omitted Genes: Kcnc3, Myh14, Izumo2
omitted Genes: Zfp473, Vrk3, Nup62
omitted Genes: Gm21948, Tbc1d17, Akt1s1
omitted Genes: Mir707, Pnkp, Ptov1
omitted Genes: Med25, Fuz, Ap2a1
omitted Genes: Tsks, Cpt1c, Prmt1
omitted Genes: Gm15545, Bcl2l12, Scaf1

Make more plots at http://csg.sph.umich.edu/locuszoom/

Fine−mapping Regions

| snp | chr | pos (Mb) | pp | group | color |
|---|---|---|---|---|---|
| chr7:44560350 | 7 | 44.56035 | 1 | Credible Set | purple |

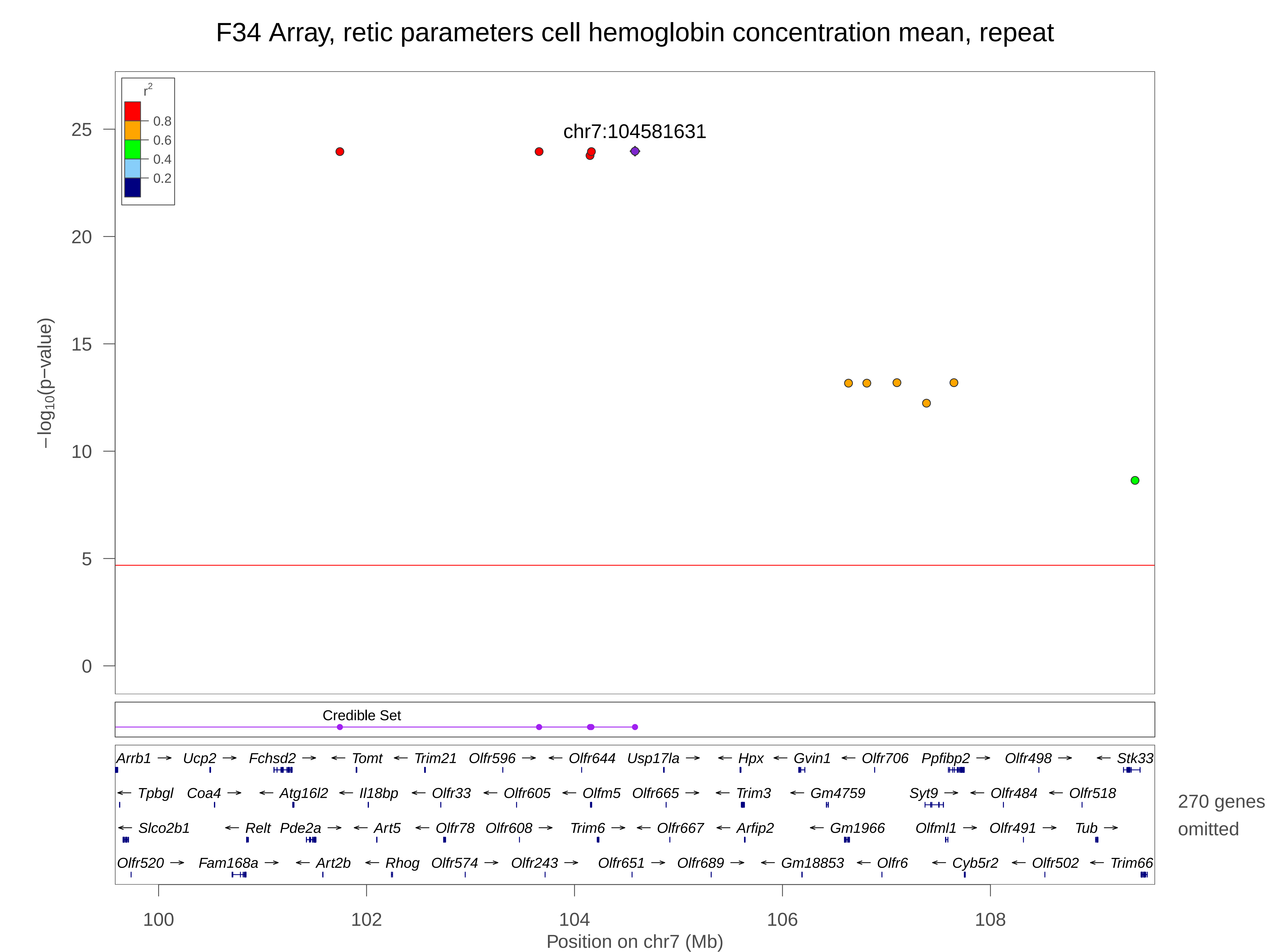
F34 Array, retic parameters cell hemoglobin concentration mean, repeat
r2
0.8
0.6
0.4
0.2
chr7:104581631
−log10(p−value)
25
20
15
10
5
0
Credible Set
Arrb1 → Ucp2 → Fchsd2 → ← Tomt ← Trim21 Olfr596 → ← Olfr644 Usp17la → ← Hpx ← Gvin1 ← Olfr706 Ppfibp2 → Olfr498 → ← Stk33
← Tpbgl Coa4 → ← Atg16l2 ← Il18bp ← Olfr33 ← Olfr605 ← Olfm5 Olfr665 → ← Trim3 ← Gm4759 Syt9 → ← Olfr484 ← Olfr518
← Slco2b1 ← Relt Pde2a → ← Art5 ← Olfr78 Olfr608 → Trim6 → ← Olfr667 ← Arfip2 ← Gm1966 Olfml1 → Olfr491 → Tub →
Olfr520 → Fam168a → ← Art2b ← Rhog Olfr574 → Olfr243 → Olfr651 → Olfr689 → ← Gm18853 ← Olfr6 ← Cyb5r2 ← Olfr502 ← Trim66
270 genes omitted
100
102
104
106
108
Position on chr7 (Mb)

date: Thu Oct 25 16:50:16 2018
build: F34Array
display range: chr7:99581631−109581631 [99581631−109581631]
hilite range: 0 − 0 [ 0 − 0 ]
reference SNP: chr7:104581631
number of SNPs plotted: 12
min P−value: 1.06E−24 [chr7:104581631]
max P−value: 2.29E−9 [chr7:109390566]
omitted Genes: Olfr521, Neu3, Spcs2
omitted Genes: Xrra1, Rnf169, Chrdl2
omitted Genes: Pold3, Lipt2, Kcne3
omitted Genes: Pgm2l1, Gpx2−ps1, P4ha3
omitted Genes: Ppme1, C2cd3, Ucp3
omitted Genes: Dnajb13, Mrpl48, Rab6a
omitted Genes: Plekhb1, Arhgef17, Mir3102
omitted Genes: P2ry6, P2ry2, Stard10
omitted Genes: Arap1, Mir139, Art2a−ps
omitted Genes: Clpb, Phox2a, Inppl1
omitted Genes: Folr2, Folr1, Anapc15
omitted Genes: Lamtor1, Lrrc51, Numa1
omitted Genes: Rnf121, Xndc1, Xntrpc
omitted Genes: Trpc2, Art1, Chrna10
omitted Genes: Nup98, Pgap2, Stim1
omitted Genes: Rrm1, Olfr543, Olfr544
omitted Genes: Olfr545, Olfr547, Olfr549
omitted Genes: Olfr550, Olfr551, Olfr552
omitted Genes: Olfr553, Olfr554, Olfr555
omitted Genes: Olfr556, Trim68, Olfr557
omitted Genes: Olfr558, Olfr559, Olfr560
omitted Genes: Olfr561, Olfr564, Olfr566
omitted Genes: Olfr568, Olfr569, Olfr570
omitted Genes: Olfr571, Olfr572, Olfr575
omitted Genes: Olfr576, Olfr577, Olfr578
omitted Genes: Olfr582, Olfr583, Olfr584
omitted Genes: Olfr585, Olfr586, Olfr589
omitted Genes: Olfr591, Olfr592, Olfr593
omitted Genes: Olfr594, Usp17ld, Olfr597
omitted Genes: Olfr598, Olfr599, Olfr600
omitted Genes: Olfr601, Olfr603, Usp17lc
omitted Genes: Olfr606, Olfr609, Olfr610

Make more plots at http://csg.sph.umich.edu/locuszoom/

Fine−mapping Regions

| snp | chr | pos (Mb) | pp | group | color |
|---|---|---|---|---|---|
| chr7:99328376 | 7 | 99.32838 | 0.1261267 | Credible Set | purple |
| chr7:101743401 | 7 | 101.74340 | 0.1859423 | Credible Set | purple |
| chr7:103659355 | 7 | 103.65936 | 0.1859423 | Credible Set | purple |
| chr7:104149022 | 7 | 104.14902 | 0.1218390 | Credible Set | purple |
| chr7:104162444 | 7 | 104.16244 | 0.1859423 | Credible Set | purple |
| chr7:104581631 | 7 | 104.58163 | 0.1942073 | Credible Set | purple |

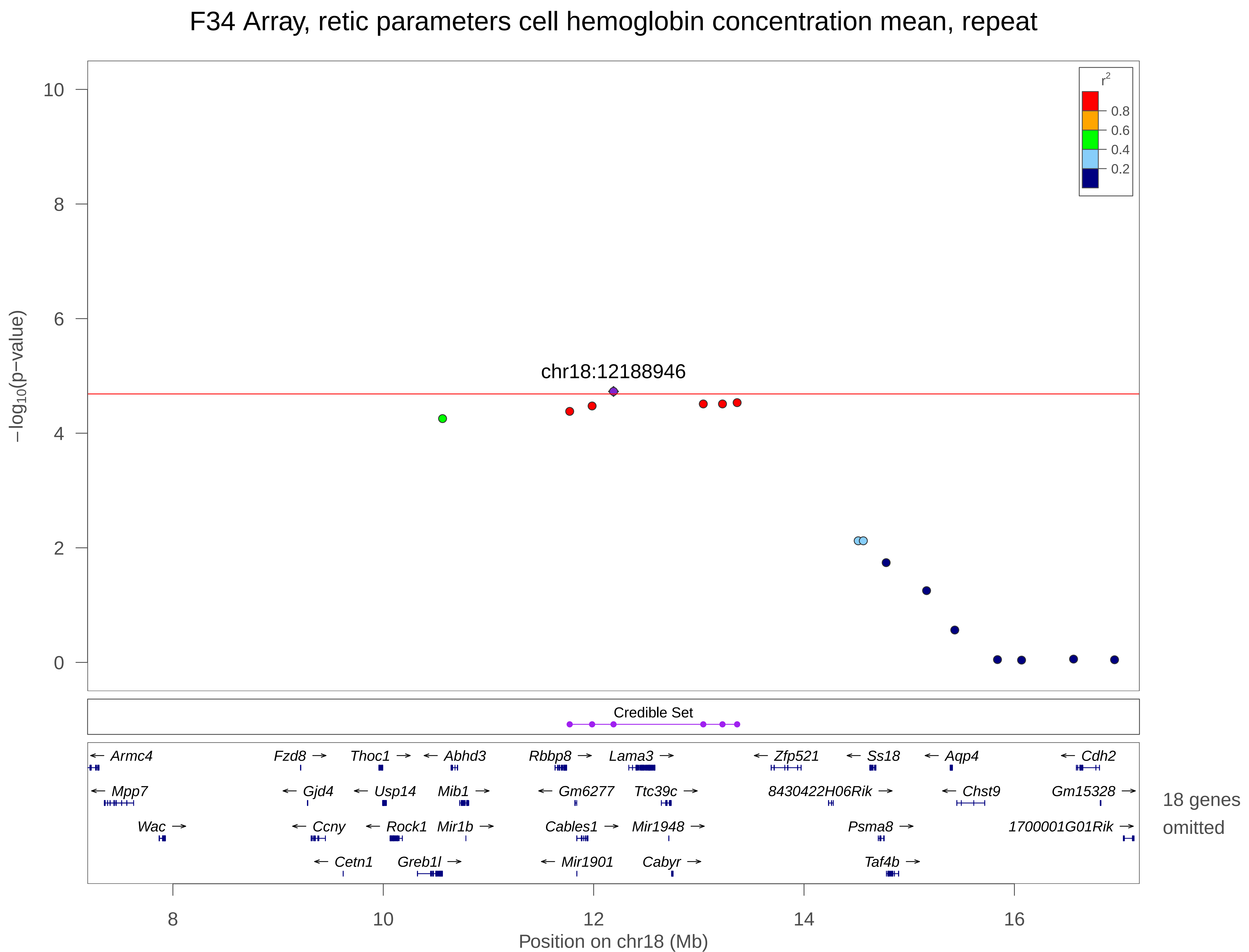
F34 Array, retic parameters cell hemoglobin concentration mean, repeat
r2
0.8
0.6
0.4
0.2
chr18:12188946
−log10(p−value)
10
8
6
4
2
0
Credible Set
Armc4
Fzd8
Thoc1
Abhd3
Rbbp8
Lama3
Zfp521
Ss18
Aqp4
Cdh2
Mpp7
Gjd4
Usp14
Mib1
Gm6277
Ttc39c
8430422H06Rik
Chst9
Gm15328
Wac
Ccny
Rock1
Mir1b
Cables1
Mir1948
Psma8
1700001G01Rik
Cetn1
Greb1l
Mir1901
Cabyr
Taf4b
18 genes omitted
8
10
12
14
16
Position on chr18 (Mb)

| | |
|---|---|
| date: | Thu Oct 25 16:50:04 2018 |
| build: | F34Array |
| display range: | chr18:7188946−17188946 [7188946−17188946] |
| hilite range: | 0 − 0 [ 0 − 0 ] |
| reference SNP: | chr18:12188946 |
| number of SNPs plotted: | 17 |
| min P−value: | 1.87E−5 [chr18:12188946] |
| max P−value: | 9.11E−1 [chr18:16068097] |
| omitted Genes: | Colec12, Esco1, Snrpd1 |
| omitted Genes: | 4930563E18Rik, Mir133a−1, Mir1a−2 |
| omitted Genes: | 1010001N08Rik, Gata6, Tmem241 |
| omitted Genes: | Riok3, 3110002H16Rik, Npc1 |
| omitted Genes: | Ankrd29, Osbpl1a, Impact |
| omitted Genes: | Hrh4, Kctd1, 4933424G05Rik |

Make more plots at http://csg.sph.umich.edu/locuszoom/

Fine−mapping Regions

| snp | chr | pos (Mb) | pp | group | color |
|---|---|---|---|---|---|
| chr18:11772246 | 18 | 11.77225 | 0.1187936 | Credible Set | purple |
| chr18:11985350 | 18 | 11.98535 | 0.1459508 | Credible Set | purple |
| chr18:12188946 | 18 | 12.18895 | 0.2541053 | Credible Set | purple |
| chr18:13042279 | 18 | 13.04228 | 0.1577027 | Credible Set | purple |
| chr18:13224851 | 18 | 13.22485 | 0.1577027 | Credible Set | purple |
| chr18:13364094 | 18 | 13.36409 | 0.1657449 | Credible Set | purple |

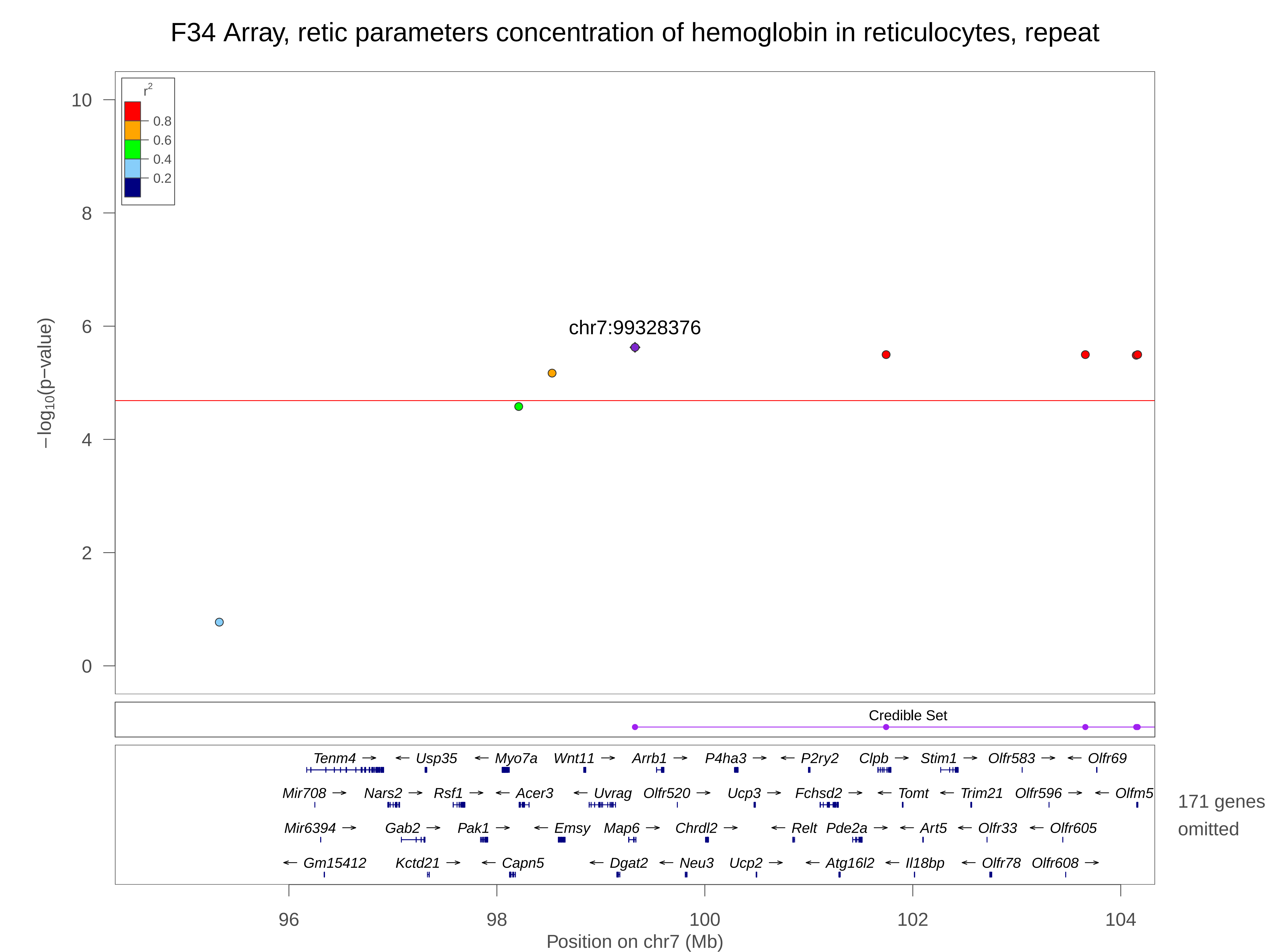

F34 Array, retic parameters concentration of hemoglobin in reticulocytes, repeat
r2
0.8
0.6
0.4
0.2
10
8
6
4
2
0
−log10(p−value)
chr7:99328376
Credible Set
Tenm4 →
← Usp35
← Myo7a
Wnt11 →
Arrb1 →
P4ha3 →
← P2ry2
Clpb →
Stim1 →
Olfr583 →
← Olfr69
Mir708 →
Nars2 →
Rsf1 →
← Acer3
← Uvrag
Olfr520 →
Ucp3 →
Fchsd2 →
← Tomt
← Trim21
Olfr596 →
← Olfm5
Mir6394 →
Gab2 →
Pak1 →
← Emsy
Map6 →
Chrdl2 →
← Relt
Pde2a →
← Art5
← Olfr33
← Olfr605
← Gm15412
Kctd21 →
← Capn5
← Dgat2
← Neu3
Ucp2 →
← Atg16l2
← Il18bp
← Olfr78
Olfr608 →
171 genes omitted
96
98
100
102
104
Position on chr7 (Mb)

date: Thu Oct 25 16:50:45 2018
build: F34Array
display range: chr7:94328376−104328376 [94328376−104328376]
hilite range: 0 − 0 [ 0 − 0 ]
reference SNP: chr7:99328376
number of SNPs plotted: 9
min P−value: 2.37E−6 [chr7:99328376]
max P−value: 1.69E−1 [chr7:95330810]
omitted Genes: Gm15413, Alg8, Ndufc2
omitted Genes: Thrsp, Kctd14, Ints4
omitted Genes: Aamdc, Rsf1os1, Clns1a
omitted Genes: Aqp11, Gdpd4, Omp
omitted Genes: Gm16938, B3gnt6, Tsku
omitted Genes: Gucy2d, Lrrc32, Prkrir
omitted Genes: Mogat2, Gm26705, Serpinh1
omitted Genes: Gdpd5, Klhl35, Rps3
omitted Genes: Snord15b, Snord15a, Mir326
omitted Genes: Tpbgl, Slco2b1, Olfr521
omitted Genes: Spcs2, Xrra1, Rnf169
omitted Genes: Pold3, Lipt2, Kcne3
omitted Genes: Pgm2l1, Gpx2−ps1, Ppme1
omitted Genes: C2cd3, Dnajb13, Coa4
omitted Genes: Mrpl48, Rab6a, Plekhb1
omitted Genes: Fam168a, Arhgef17, Mir3102
omitted Genes: P2ry6, Stard10, Arap1
omitted Genes: Mir139, Art2a−ps, Art2b
omitted Genes: Phox2a, Inppl1, Folr2
omitted Genes: Folr1, Anapc15, Lamtor1
omitted Genes: Lrrc51, Numa1, Rnf121
omitted Genes: Xndc1, Xntrpc, Trpc2
omitted Genes: Art1, Chrna10, Nup98
omitted Genes: Pgap2, Rhog, Rrm1
omitted Genes: Olfr543, Olfr544, Olfr545
omitted Genes: Olfr547, Olfr549, Olfr550
omitted Genes: Olfr551, Olfr552, Olfr553
omitted Genes: Olfr554, Olfr555, Olfr556
omitted Genes: Trim68, Olfr557, Olfr558
omitted Genes: Olfr559, Olfr560, Olfr561
omitted Genes: Olfr564, Olfr566, Olfr568
Make more plots at http://csg.sph.umich.edu/locuszoom/
omitted Genes: Olfr569, Olfr570, Olfr571

Fine−mapping Regions

| snp | chr | pos (Mb) | pp | group | color |
|---|---|---|---|---|---|
| chr7:99328376 | 7 | 99.32838 | 0.2105870 | Credible Set | purple |
| chr7:101743401 | 7 | 101.74340 | 0.1585895 | Credible Set | purple |
| chr7:103659355 | 7 | 103.65936 | 0.1585895 | Credible Set | purple |
| chr7:104149022 | 7 | 104.14902 | 0.1547852 | Credible Set | purple |
| chr7:104162444 | 7 | 104.16244 | 0.1585895 | Credible Set | purple |
| chr7:104581631 | 7 | 104.58163 | 0.1588594 | Credible Set | purple |

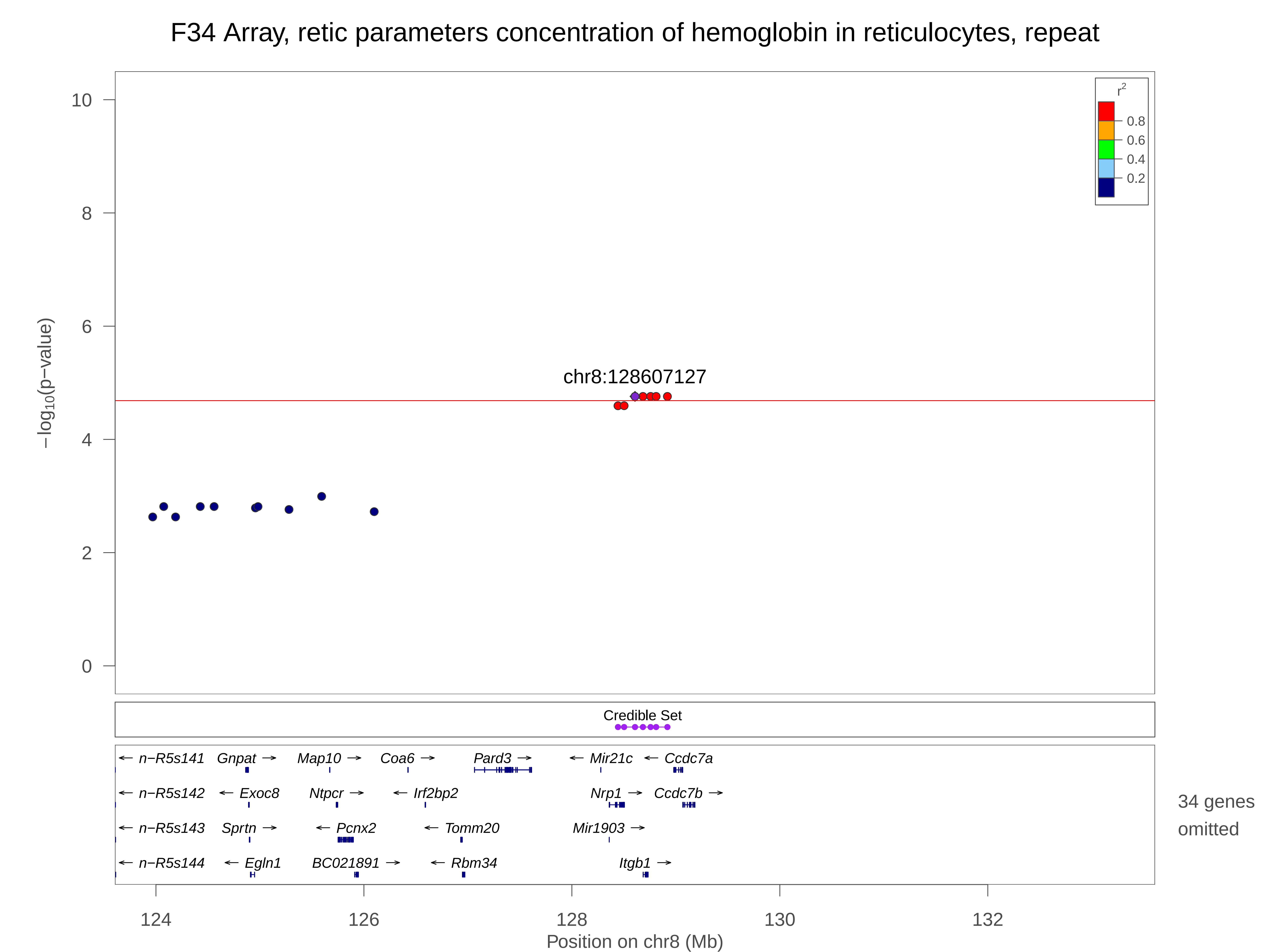
F34 Array, retic parameters concentration of hemoglobin in reticulocytes, repeat
−log10(p−value)
10
8
6
4
2
0
r2
0.8
0.6
0.4
0.2
chr8:128607127
Credible Set
← n−R5s141
Gnpat →
Map10 →
Coa6 →
Pard3 →
← Mir21c
← Ccdc7a
← n−R5s142
← Exoc8
Ntpcr →
← Irf2bp2
Nrp1 →
Ccdc7b →
← n−R5s143
Sprtn →
← Pcnx2
← Tomm20
Mir1903 →
← n−R5s144
← Egln1
BC021891 →
← Rbm34
Itgb1 →
34 genes omitted
124
126
128
130
132
Position on chr8 (Mb)

| | |
|---|---|
| date: | Thu Oct 25 16:50:52 2018 |
| build: | F34Array |
| display range: | chr8:123607127−133607127 [123607127−133607127] |
| hilite range: | 0 − 0 [ 0 − 0 ] |
| reference SNP: | chr8:128607127 |
| number of SNPs plotted: | 18 |
| min P−value: | 1.74E−5 [chr8:128607127] |
| max P−value: | 2.34E−3 [chr8:123968866] |
| omitted Genes: | Gm25212, n−R5s146, Gm22291 |
| omitted Genes: | Gm23284, n−R5s149, Rhou |
| omitted Genes: | 6030466F02Rik, Rab4a, Ccsap |
| omitted Genes: | Acta1, Nup133, Abcb10 |
| omitted Genes: | Taf5l, Mir1967, Urb2 |
| omitted Genes: | Galnt2, Pgbd5, Cog2 |
| omitted Genes: | Agt, Capn9, 2310022B05Rik |
| omitted Genes: | Ttc13, Arv1, Fam89a |
| omitted Genes: | Trim67, 2810004N23Rik, Tsnax |
| omitted Genes: | Disc1, Sipa1l2, 4930567H12Rik |
| omitted Genes: | Kcnk1, Slc35f3, Gm17296 |
| omitted Genes: | 2610044O15Rik8 |

Make more plots at http://csg.sph.umich.edu/locuszoom/

Fine−mapping Regions

| snp | chr | pos (Mb) | pp | group | color |
|---|---|---|---|---|---|
| chr8:128443469 | 8 | 128.4435 | 0.1089254 | Credible Set | purple |
| chr8:128502425 | 8 | 128.5024 | 0.1091172 | Credible Set | purple |
| chr8:128607127 | 8 | 128.6071 | 0.1563915 | Credible Set | purple |
| chr8:128683251 | 8 | 128.6833 | 0.1563915 | Credible Set | purple |
| chr8:128757217 | 8 | 128.7572 | 0.1563915 | Credible Set | purple |
| chr8:128810304 | 8 | 128.8103 | 0.1563915 | Credible Set | purple |
| chr8:128918665 | 8 | 128.9187 | 0.1563915 | Credible Set | purple |

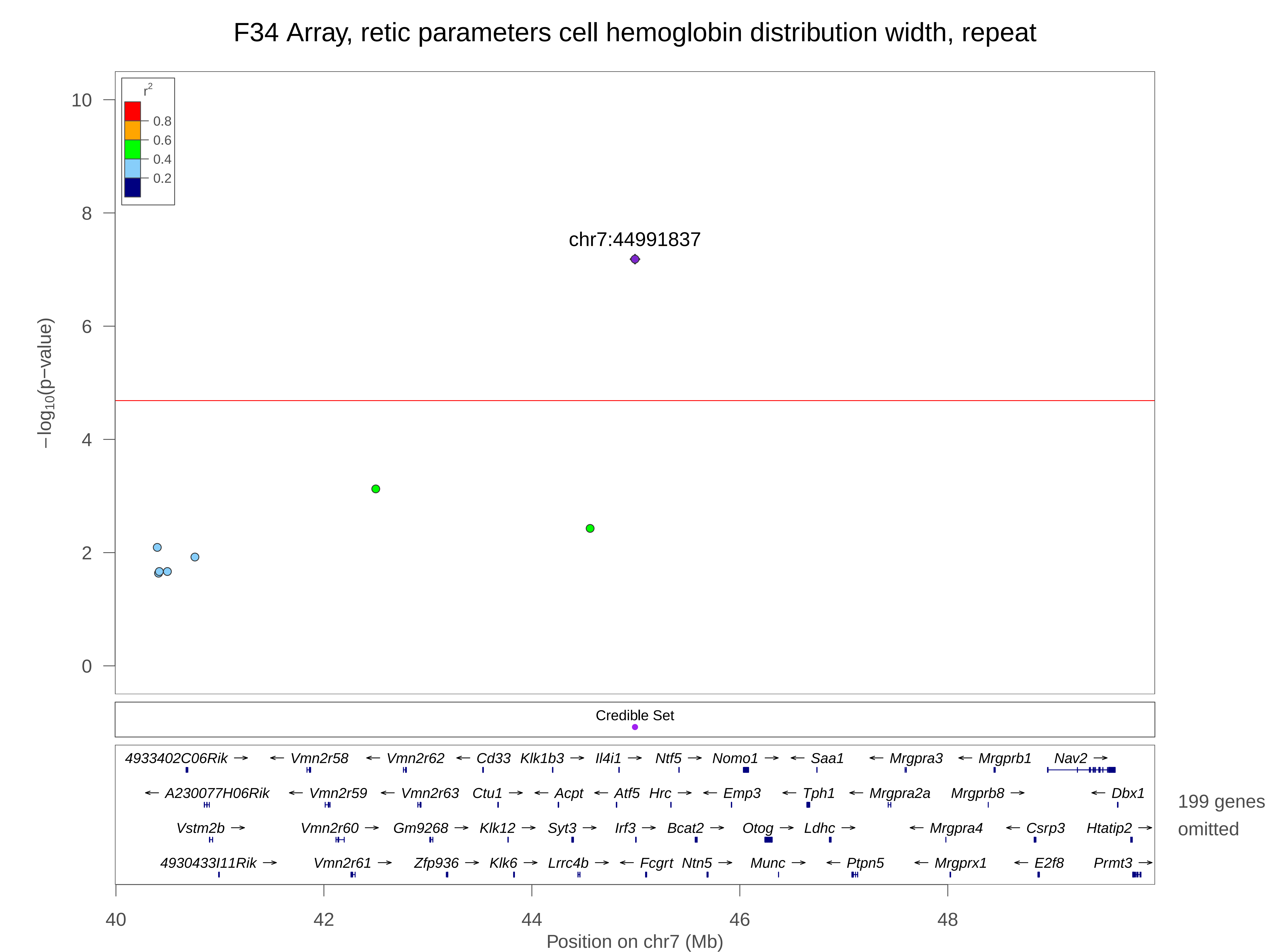
F34 Array, retic parameters cell hemoglobin distribution width, repeat
r2
0.8
0.6
0.4
0.2
10
8
6
4
2
0
−log10(p−value)
chr7:44991837
Credible Set
4933402C06Rik → ← Vmn2r58 ← Vmn2r62 ← Cd33 Klk1b3 → Il4i1 → Ntf5 → Nomo1 → ← Saa1 ← Mrgpra3 ← Mrgprb1 Nav2 →
← A230077H06Rik ← Vmn2r59 ← Vmn2r63 Ctu1 → ← Acpt ← Atf5 Hrc → ← Emp3 ← Tph1 ← Mrgpra2a Mrgprb8 → ← Dbx1
Vstm2b → Vmn2r60 → Gm9268 → Klk12 → Syt3 → Irf3 → Bcat2 → Otog → Ldhc → ← Mrgpra4 ← Csrp3 Htatip2 →
4930433I11Rik → Vmn2r61 → Zfp936 → Klk6 → Lrrc4b → ← Fcgrt Ntn5 → Munc → ← Ptpn5 ← Mrgprx1 ← E2f8 Prmt3 →
199 genes omitted
40
42
44
46
48
Position on chr7 (Mb)

date: Thu Oct 25 16:51:02 2018
build: F34Array
display range: chr7:39991837−49991837 [39991837−49991837]
hilite range: 0 − 0 [ 0 − 0 ]
reference SNP: chr7:44991837
number of SNPs plotted: 9
min P−value: 6.55E−8 [chr7:44991837]
max P−value: 2.31E−2 [chr7:40409511]
omitted Genes: Gm4884, 1700110I07Rik, Gm5592
omitted Genes: AI987944, Vmn2r57, AW146154
omitted Genes: Zfp788, Vmn2r−ps54, Vmn2r−ps60
omitted Genes: 4933421I07Rik, Zfp141, 9830147E19Rik
omitted Genes: Gm5595, Gm2381, EU599041
omitted Genes: Zfp715, Siglecf, Siglecg
omitted Genes: Lim2, Nkg7, Cldnd2
omitted Genes: Etfb, Mir7051, Vsig10l
omitted Genes: Iglon5, 4931406B18Rik, Zfp658
omitted Genes: Zfp719, Zfp819, Ceacam18
omitted Genes: Siglece, Klk14, Klk13
omitted Genes: 2310002F09Rik, Klk11, Klk10
omitted Genes: Klk9, Klk8, Klk7
omitted Genes: Klk5, Klk4, Klk15
omitted Genes: Klk1b7−ps, Klk1b8, Klk1b1
omitted Genes: Klk1b9, Klk1b11, Klk1b26
omitted Genes: Klk1b27, Klk1b21, Klk1b22
omitted Genes: Klk1b16, Klk1b24, Klk1b4
omitted Genes: Klk1b5, Klk1, 1700028J19Rik
omitted Genes: 2410002F23Rik, Snord88c, Snord88a
omitted Genes: Clec11a, Shank1, 1700008O03Rik
omitted Genes: Aspdh, Josd2, Mir7052
omitted Genes: 5430431A17Rik, Emc10, Fam71e1
omitted Genes: Mybpc2, Spib, Pold1
omitted Genes: Mir7053, Nr1h2, Napsa
omitted Genes: Kcnc3, Myh14, Izumo2
omitted Genes: Zfp473, Vrk3, Nup62
omitted Genes: Gm21948, Tbc1d17, Akt1s1
omitted Genes: Mir707, Pnkp, Ptov1
omitted Genes: Med25, Fuz, Ap2a1
omitted Genes: Tsks, Cpt1c, Prmt1
Make more plots at http://csg.sph.umich.edu/locuszoom/
omitted Genes: Gm15545, Bcl2l12, Scaf1

Fine−mapping Regions

| snp | chr | pos (Mb) | pp | group | color |
|---|---|---|---|---|---|
| chr7:44991837 | 7 | 44.99184 | 1 | Credible Set | purple |

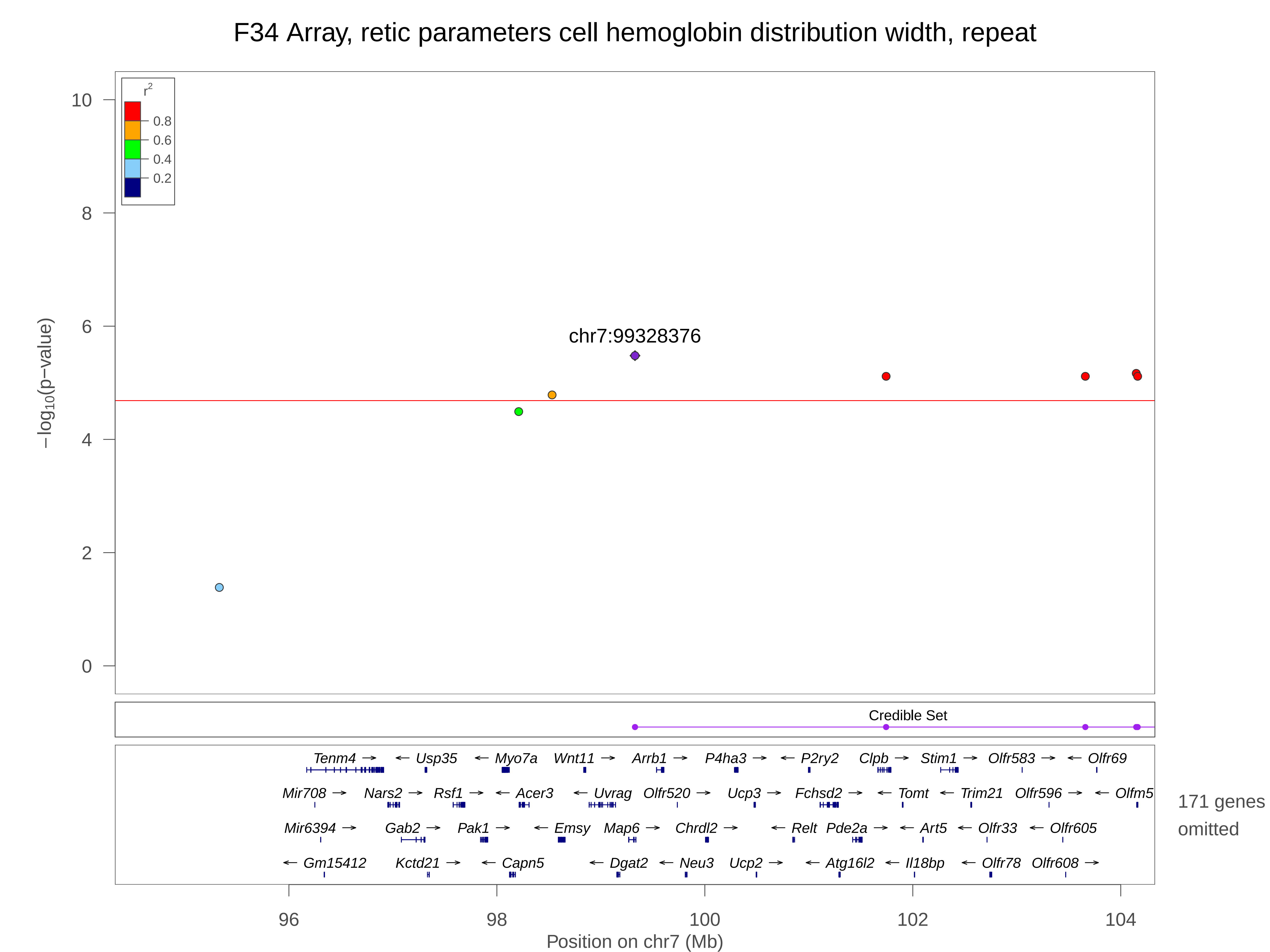

F34 Array, retic parameters cell hemoglobin distribution width, repeat
r2
0.8
0.6
0.4
0.2
10
8
6
4
2
0
−log10(p−value)
chr7:99328376
Credible Set
Tenm4 → ← Usp35 ← Myo7a Wnt11 → Arrb1 → P4ha3 → ← P2ry2 Clpb → Stim1 → Olfr583 → ← Olfr69
Mir708 → Nars2 → Rsf1 → ← Acer3 ← Uvrag Olfr520 → Ucp3 → Fchsd2 → ← Tomt ← Trim21 Olfr596 → ← Olfm5
Mir6394 → Gab2 → Pak1 → ← Emsy Map6 → Chrdl2 → ← Relt Pde2a → ← Art5 ← Olfr33 ← Olfr605
← Gm15412 Kctd21 → ← Capn5 ← Dgat2 ← Neu3 Ucp2 → ← Atg16l2 ← Il18bp ← Olfr78 Olfr608 →
171 genes omitted
96
98
100
102
104
Position on chr7 (Mb)

date: Thu Oct 25 16:51:12 2018
build: F34Array
display range: chr7:94328376−104328376 [94328376−104328376]
hilite range: 0 − 0 [ 0 − 0 ]
reference SNP: chr7:99328376
number of SNPs plotted: 9
min P−value: 3.32E−6 [chr7:99328376]
max P−value: 4.12E−2 [chr7:95330810]
omitted Genes: Gm15413, Alg8, Ndufc2
omitted Genes: Thrsp, Kctd14, Ints4
omitted Genes: Aamdc, Rsf1os1, Clns1a
omitted Genes: Aqp11, Gdpd4, Omp
omitted Genes: Gm16938, B3gnt6, Tsku
omitted Genes: Gucy2d, Lrrc32, Prkrir
omitted Genes: Mogat2, Gm26705, Serpinh1
omitted Genes: Gdpd5, Klhl35, Rps3
omitted Genes: Snord15b, Snord15a, Mir326
omitted Genes: Tpbgl, Slco2b1, Olfr521
omitted Genes: Spcs2, Xrra1, Rnf169
omitted Genes: Pold3, Lipt2, Kcne3
omitted Genes: Pgm2l1, Gpx2−ps1, Ppme1
omitted Genes: C2cd3, Dnajb13, Coa4
omitted Genes: Mrpl48, Rab6a, Plekhb1
omitted Genes: Fam168a, Arhgef17, Mir3102
omitted Genes: P2ry6, Stard10, Arap1
omitted Genes: Mir139, Art2a−ps, Art2b
omitted Genes: Phox2a, Inppl1, Folr2
omitted Genes: Folr1, Anapc15, Lamtor1
omitted Genes: Lrrc51, Numa1, Rnf121
omitted Genes: Xndc1, Xntrpc, Trpc2
omitted Genes: Art1, Chrna10, Nup98
omitted Genes: Pgap2, Rhog, Rrm1
omitted Genes: Olfr543, Olfr544, Olfr545
omitted Genes: Olfr547, Olfr549, Olfr550
omitted Genes: Olfr551, Olfr552, Olfr553
omitted Genes: Olfr554, Olfr555, Olfr556
omitted Genes: Trim68, Olfr557, Olfr558
omitted Genes: Olfr559, Olfr560, Olfr561
omitted Genes: Olfr564, Olfr566, Olfr568
Make more plots at http://csg.sph.umich.edu/locuszoom/
omitted Genes: Olfr569, Olfr570, Olfr571

Fine−mapping Regions

| snp | chr | pos (Mb) | pp | group | color |
| --- | --- | --- | --- | --- | --- |
| chr7:99328376 | 7 | 99.32838 | 0.3050743 | Credible Set | purple |
| chr7:101743401 | 7 | 101.74340 | 0.1364009 | Credible Set | purple |
| chr7:103659355 | 7 | 103.65936 | 0.1364009 | Credible Set | purple |
| chr7:104149022 | 7 | 104.14902 | 0.1531233 | Credible Set | purple |
| chr7:104162444 | 7 | 104.16244 | 0.1364009 | Credible Set | purple |
| chr7:104581631 | 7 | 104.58163 | 0.1325995 | Credible Set | purple |

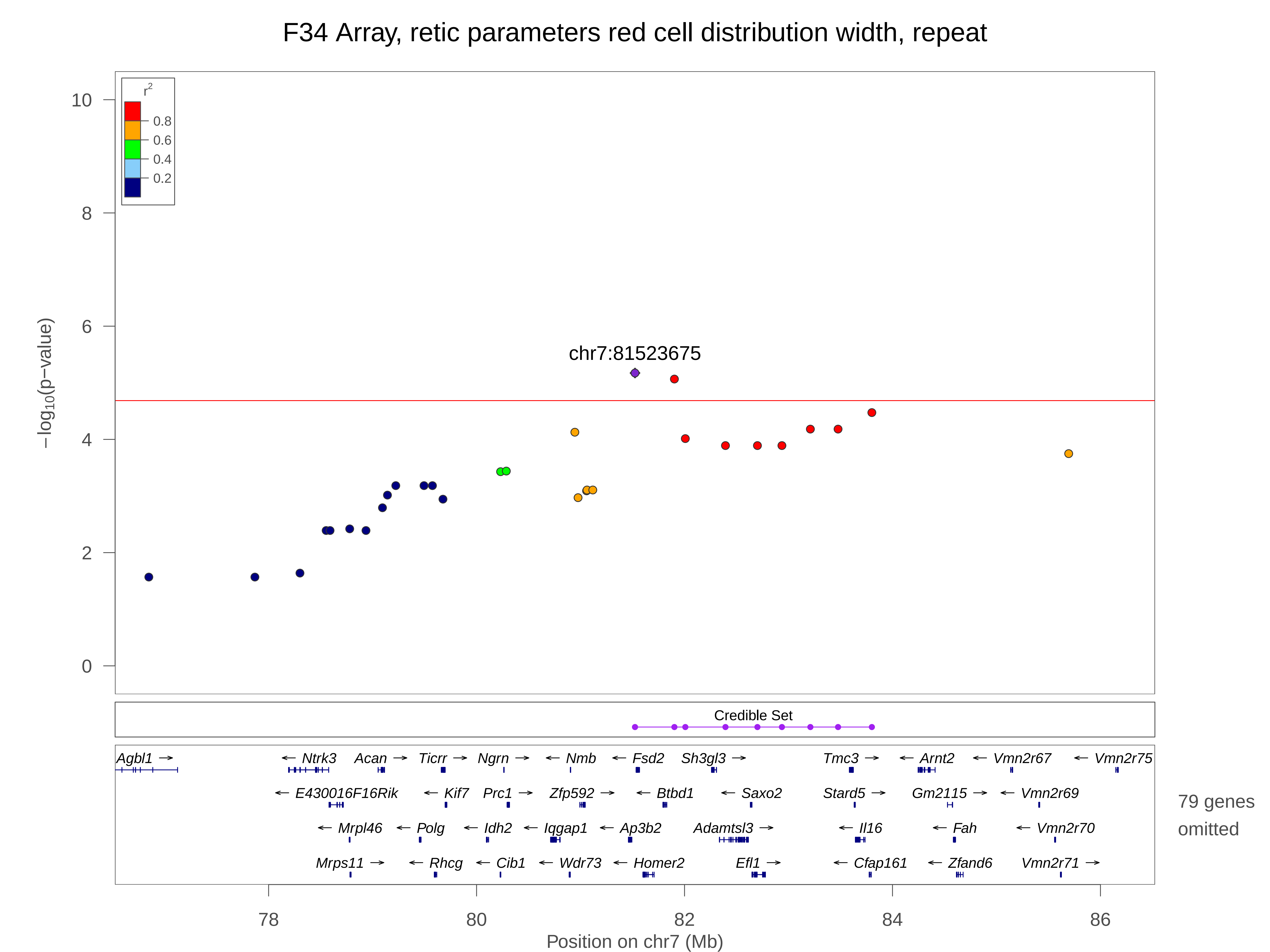

F34 Array, retic parameters red cell distribution width, repeat
r2
0.8
0.6
0.4
0.2
10
8
6
4
2
0
−log10(p−value)
chr7:81523675
Credible Set
Agbl1 →
← Ntrk3
Acan →
Ticrr →
Ngrn →
← Nmb
← Fsd2
Sh3gl3 →
Tmc3 →
← Arnt2
← Vmn2r67
← Vmn2r75
← E430016F16Rik
← Kif7
Prc1 →
Zfp592 →
← Btbd1
← Saxo2
Stard5 →
Gm2115 →
← Vmn2r69
← Mrpl46
← Polg
← Idh2
← Iqgap1
← Ap3b2
Adamtsl3 →
← Il16
← Fah
← Vmn2r70
Mrps11 →
← Rhcg
← Cib1
← Wdr73
← Homer2
Efl1 →
← Cfap161
← Zfand6
Vmn2r71 →
79 genes omitted
78
80
82
84
86
Position on chr7 (Mb)

| | |
|---|---|
| date: | Thu Oct 25 16:51:19 2018 |
| build: | F34Array |
| display range: | chr7:76523675−86523675 [76523675−86523675] |
| hilite range: | 0 − 0 [ 0 − 0 ] |
| reference SNP: | chr7:81523675 |
| number of SNPs plotted: | 31 |
| min P−value: | 6.72E−6 [chr7:81523675] |
| max P−value: | 2.7E−2 [chr7:76848144] |
| omitted Genes: | Det1, Mir7−2, Aen |
| omitted Genes: | Isg20, Gm26633, Hapln3 |
| omitted Genes: | Mfge8, Abhd2, Rlbp1 |
| omitted Genes: | Fanci, Mir9−3hg, Mir9−3 |
| omitted Genes: | Plin1, Pex11a, Wdr93 |
| omitted Genes: | Mesp1, Mesp2, Anpep |
| omitted Genes: | Ap3s2, Arpin, Zfp710 |
| omitted Genes: | Gm21057, Sema4b, Gdpgp1 |
| omitted Genes: | Ttll13, Vps33b, Rccd1 |
| omitted Genes: | Unc45a, Hddc3, Man2a2 |
| omitted Genes: | Fes, Furin, Blm |
| omitted Genes: | Crtc3, Gm15880, Zscan2 |
| omitted Genes: | Sec11a, Alpk3, Slc28a1 |
| omitted Genes: | Pde8a, Rps17, Cpeb1 |
| omitted Genes: | BC048679, 2900076A07Rik, Mir1839 |
| omitted Genes: | Whamm, Fam103a1, 3110040N11Rik |
| omitted Genes: | Tm6sf1, Hdgfrp3, Bnc1 |
| omitted Genes: | Gm20744, 4933406J10Rik, Mex3b |
| omitted Genes: | 4933430H16Rik, Mesdc1, Mesdc2 |
| omitted Genes: | Cemip, Abhd17c, 2610206C17Rik |
| omitted Genes: | Olfr291, Olfr290, Vmn2r65 |
| omitted Genes: | Vmn2r66, Vmn2r68, Vmn2r72 |
| omitted Genes: | Vmn2r73, Vmn2r74, Vmn2r76 |
| omitted Genes: | Olfr310, Olfr309, Olfr308 |
| omitted Genes: | Olfr307, Olfr305, Olfr304 |
| omitted Genes: | Olfr303, Olfr301, Olfr299 |
| omitted Genes: | Olfr298 |

Make more plots at http://csg.sph.umich.edu/locuszoom/

Fine−mapping Regions

| snp | chr | pos (Mb) | pp | group | color |
|---|---|---|---|---|---|
| chr7:81523675 | 7 | 81.52367 | 0.40068110 | Credible Set | purple |
| chr7:81902543 | 7 | 81.90254 | 0.31702067 | Credible Set | purple |
| chr7:82007699 | 7 | 82.00770 | 0.03171553 | Credible Set | purple |
| chr7:82393693 | 7 | 82.39369 | 0.02426518 | Credible Set | purple |
| chr7:82701107 | 7 | 82.70111 | 0.02426518 | Credible Set | purple |
| chr7:82936666 | 7 | 82.93667 | 0.02426518 | Credible Set | purple |
| chr7:83210268 | 7 | 83.21027 | 0.04570385 | Credible Set | purple |
| chr7:83476529 | 7 | 83.47653 | 0.04570385 | Credible Set | purple |
| chr7:83801589 | 7 | 83.80159 | 0.08637946 | Credible Set | purple |

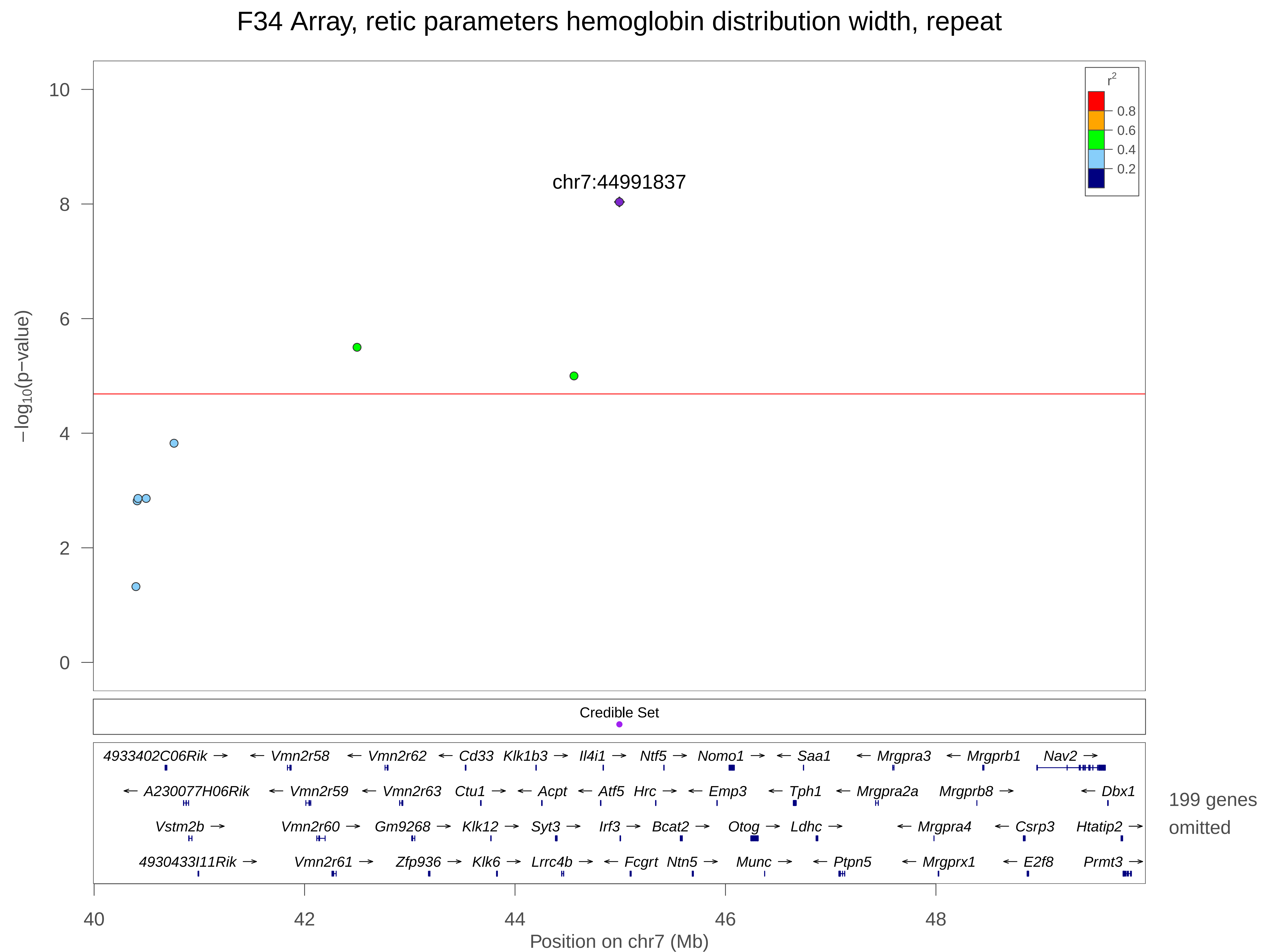
F34 Array, retic parameters hemoglobin distribution width, repeat
chr7:44991837
−log10(p−value)
10
8
6
4
2
0
r2
0.8
0.6
0.4
0.2
Credible Set
4933402C06Rik →
← Vmn2r58
← Vmn2r62
← Cd33
Klk1b3 →
Il4i1 →
Ntf5 →
Nomo1 →
← Saa1
← Mrgpra3
← Mrgprb1
Nav2 →
← A230077H06Rik
← Vmn2r59
← Vmn2r63
Ctu1 →
← Acpt
← Atf5
Hrc →
← Emp3
← Tph1
← Mrgpra2a
Mrgprb8 →
← Dbx1
Vstm2b →
Vmn2r60 →
Gm9268 →
Klk12 →
Syt3 →
Irf3 →
Bcat2 →
Otog →
Ldhc →
← Mrgpra4
← Csrp3
Htatip2 →
4930433I11Rik →
Vmn2r61 →
Zfp936 →
Klk6 →
Lrrc4b →
← Fcgrt
Ntn5 →
Munc →
← Ptpn5
← Mrgprx1
← E2f8
Prmt3 →
199 genes omitted
40
42
44
46
48
Position on chr7 (Mb)

date: Thu Oct 25 16:51:40 2018
build: F34Array
display range: chr7:39991837−49991837 [39991837−49991837]
hilite range: 0 − 0 [ 0 − 0 ]
reference SNP: chr7:44991837
number of SNPs plotted: 9
min P−value: 9.2E−9 [chr7:44991837]
max P−value: 4.75E−2 [chr7:40397482]
omitted Genes: Gm4884, 1700110I07Rik, Gm5592
omitted Genes: AI987944, Vmn2r57, AW146154
omitted Genes: Zfp788, Vmn2r−ps54, Vmn2r−ps60
omitted Genes: 4933421I07Rik, Zfp141, 9830147E19Rik
omitted Genes: Gm5595, Gm2381, EU599041
omitted Genes: Zfp715, Siglecf, Siglecg
omitted Genes: Lim2, Nkg7, Cldnd2
omitted Genes: Etfb, Mir7051, Vsig10l
omitted Genes: Iglon5, 4931406B18Rik, Zfp658
omitted Genes: Zfp719, Zfp819, Ceacam18
omitted Genes: Siglece, Klk14, Klk13
omitted Genes: 2310002F09Rik, Klk11, Klk10
omitted Genes: Klk9, Klk8, Klk7
omitted Genes: Klk5, Klk4, Klk15
omitted Genes: Klk1b7−ps, Klk1b8, Klk1b1
omitted Genes: Klk1b9, Klk1b11, Klk1b26
omitted Genes: Klk1b27, Klk1b21, Klk1b22
omitted Genes: Klk1b16, Klk1b24, Klk1b4
omitted Genes: Klk1b5, Klk1, 1700028J19Rik
omitted Genes: 2410002F23Rik, Snord88c, Snord88a
omitted Genes: Clec11a, Shank1, 1700008O03Rik
omitted Genes: Aspdh, Josd2, Mir7052
omitted Genes: 5430431A17Rik, Emc10, Fam71e1
omitted Genes: Mybpc2, Spib, Pold1
omitted Genes: Mir7053, Nr1h2, Napsa
omitted Genes: Kcnc3, Myh14, Izumo2
omitted Genes: Zfp473, Vrk3, Nup62
omitted Genes: Gm21948, Tbc1d17, Akt1s1
omitted Genes: Mir707, Pnkp, Ptov1
omitted Genes: Med25, Fuz, Ap2a1
omitted Genes: Tsks, Cpt1c, Prmt1
omitted Genes: Gm15545, Bcl2l12, Scaf1

Make more plots at http://csg.sph.umich.edu/locuszoom/

Fine−mapping Regions

| snp | chr | pos (Mb) | pp | group | color |
|---|---|---|---|---|---|
| chr7:44991837 | 7 | 44.99184 | 1 | Credible Set | purple |

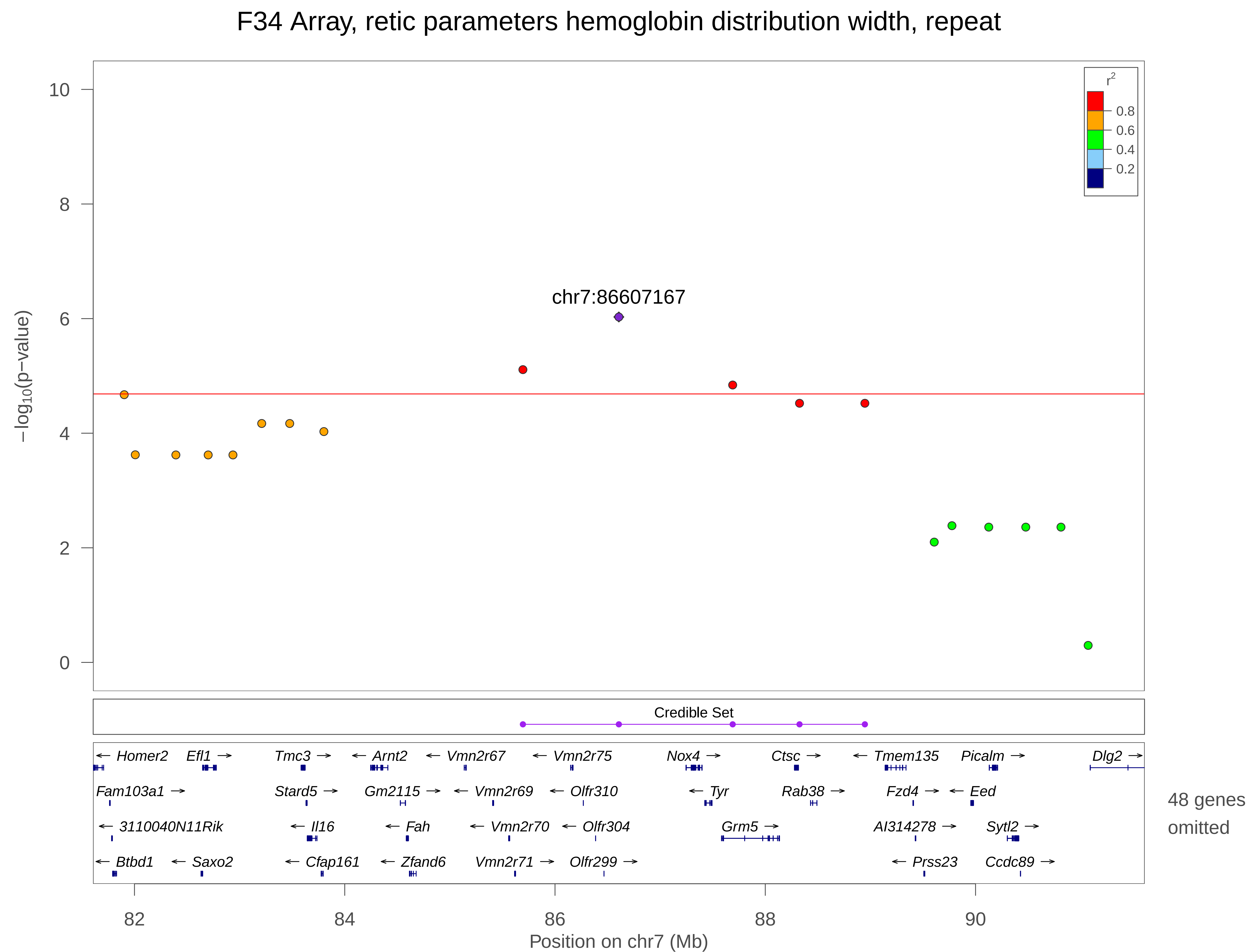
F34 Array, retic parameters hemoglobin distribution width, repeat
chr7:86607167
r2
0.8
0.6
0.4
0.2
−log10(p−value)
Credible Set
Homer2
Efl1
Tmc3
Arnt2
Vmn2r67
Vmn2r75
Nox4
Ctsc
Tmem135
Picalm
Dlg2
Fam103a1
Stard5
Gm2115
Vmn2r69
Olfr310
Tyr
Rab38
Fzd4
Eed
3110040N11Rik
Il16
Fah
Vmn2r70
Olfr304
Grm5
AI314278
Sytl2
Btbd1
Saxo2
Cfap161
Zfand6
Vmn2r71
Olfr299
Prss23
Ccdc89
48 genes omitted
Position on chr7 (Mb)

| | |
|---|---|
| date: | Thu Oct 25 16:51:47 2018 |
| build: | F34Array |
| display range: | chr7:81607167−91607167 [81607167−91607167] |
| hilite range: | 0 − 0 [ 0 − 0 ] |
| reference SNP: | chr7:86607167 |
| number of SNPs plotted: | 20 |
| min P−value: | 9.36E−7 [chr7:86607167] |
| max P−value: | 5.05E−1 [chr7:91070939] |
| omitted Genes: | Tm6sf1, Hdgfrp3, Bnc1 |
| omitted Genes: | Gm20744, Sh3gl3, Adamtsl3 |
| omitted Genes: | 4933406J10Rik, Mex3b, 4933430H16Rik |
| omitted Genes: | Mesdc1, Mesdc2, Cemip |
| omitted Genes: | Abhd17c, 2610206C17Rik, Olfr291 |
| omitted Genes: | Olfr290, Vmn2r65, Vmn2r66 |
| omitted Genes: | Vmn2r68, Vmn2r72, Vmn2r73 |
| omitted Genes: | Vmn2r74, Vmn2r76, Olfr309 |
| omitted Genes: | Olfr308, Olfr307, Olfr305 |
| omitted Genes: | Olfr303, Olfr301, Olfr298 |
| omitted Genes: | Olfr297, Olfr295, Olfr294 |
| omitted Genes: | Olfr293, Olfr292, Folh1 |
| omitted Genes: | Vmn2r77, Vmn2r78, Vmn2r79 |
| omitted Genes: | Me3, Ccdc81, l7Rn6 |
| omitted Genes: | E230029C05Rik, 2310010J17Rik, Ccdc83 |
| omitted Genes: | Crebzf, Tmem126a, Tmem126b |

Make more plots at http://csg.sph.umich.edu/locuszoom/

Fine−mapping Regions

| snp | chr | pos (Mb) | pp | group | color |
|---|---|---|---|---|---|
| chr7:85694603 | 7 | 85.69460 | 0.10284755 | Credible Set | purple |
| chr7:86607167 | 7 | 86.60717 | 0.78343314 | Credible Set | purple |
| chr7:87689885 | 7 | 87.68989 | 0.05701918 | Credible Set | purple |
| chr7:88325650 | 7 | 88.32565 | 0.02835006 | Credible Set | purple |
| chr7:88947234 | 7 | 88.94723 | 0.02835006 | Credible Set | purple |

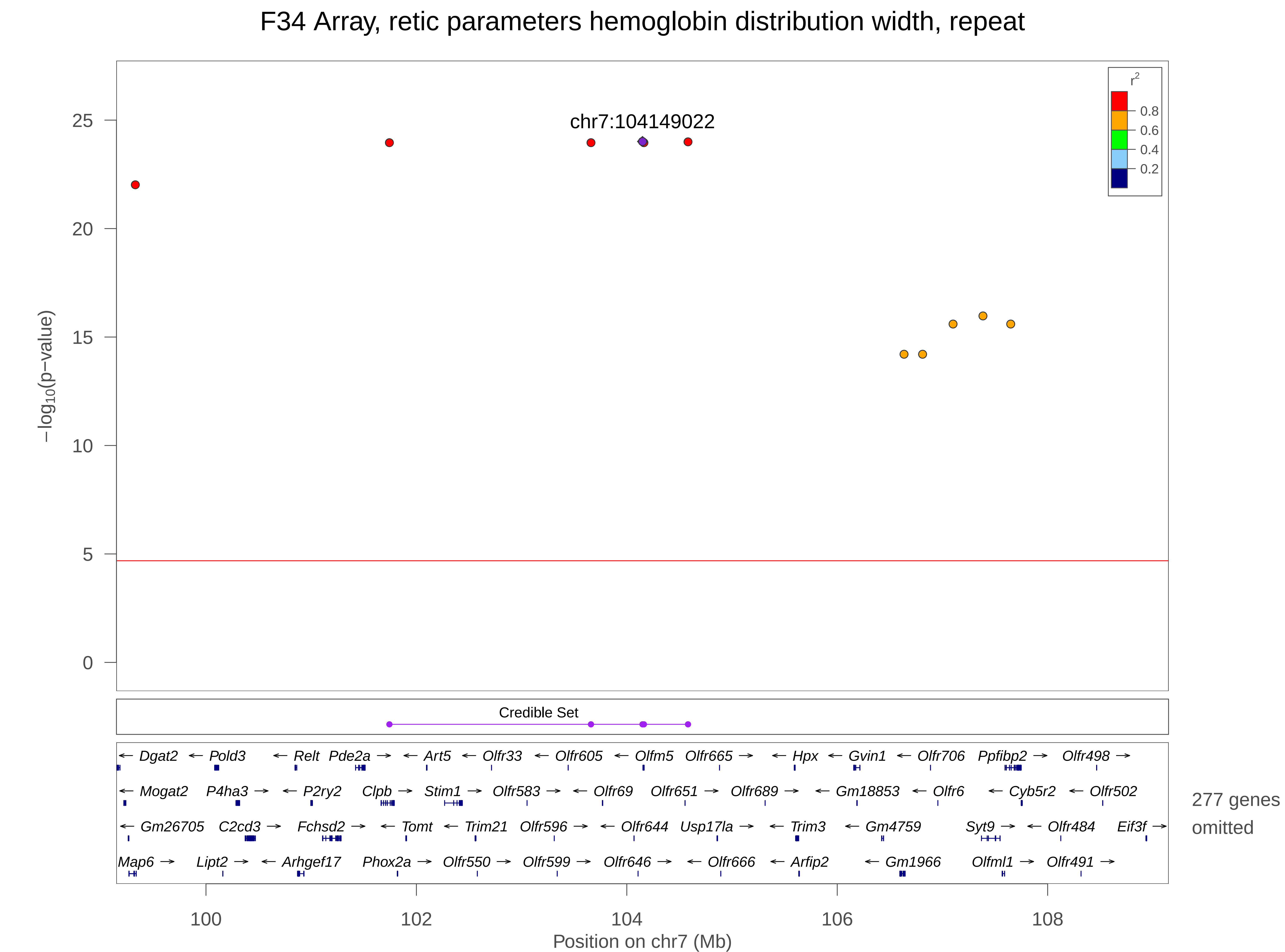
F34 Array, retic parameters hemoglobin distribution width, repeat
chr7:104149022
r2
0.8
0.6
0.4
0.2
−log10(p−value)
25
20
15
10
5
0
Credible Set
← Dgat2 ← Pold3 ← Relt Pde2a → ← Art5 ← Olfr33 ← Olfr605 ← Olfm5 Olfr665 → ← Hpx ← Gvin1 ← Olfr706 Ppfibp2 → Olfr498 →
← Mogat2 P4ha3 → ← P2ry2 Clpb → Stim1 → Olfr583 → ← Olfr69 Olfr651 → Olfr689 → ← Gm18853 ← Olfr6 ← Cyb5r2 ← Olfr502
← Gm26705 C2cd3 → Fchsd2 → ← Tomt ← Trim21 Olfr596 → ← Olfr644 Usp17la → ← Trim3 ← Gm4759 Syt9 → ← Olfr484 Eif3f →
Map6 → Lipt2 → ← Arhgef17 Phox2a → Olfr550 → Olfr599 → Olfr646 → ← Olfr666 ← Arfip2 ← Gm1966 Olfml1 → Olfr491 →
277 genes omitted
100
102
104
106
108
Position on chr7 (Mb)

date: Thu Oct 25 16:51:30 2018
build: F34Array
display range: chr7:99149022−109149022 [99149022−109149022]
hilite range: 0 − 0 [ 0 − 0 ]
reference SNP: chr7:104149022
number of SNPs plotted: 12
min P−value: 9.67E−25 [chr7:104149022]
max P−value: 6.21E−15 [chr7:106634943]
omitted Genes: Serpinh1, Gdpd5, Klhl35
omitted Genes: Rps3, Snord15b, Snord15a
omitted Genes: Arrb1, Mir326, Tpbgl
omitted Genes: Slco2b1, Olfr520, Olfr521
omitted Genes: Neu3, Spcs2, Xrra1
omitted Genes: Rnf169, Chrdl2, Kcne3
omitted Genes: Pgm2l1, Gpx2−ps1, Ppme1
omitted Genes: Ucp3, Ucp2, Dnajb13
omitted Genes: Coa4, Mrpl48, Rab6a
omitted Genes: Plekhb1, Fam168a, Mir3102
omitted Genes: P2ry6, Atg16l2, Stard10
omitted Genes: Arap1, Mir139, Art2a−ps
omitted Genes: Art2b, Inppl1, Folr2
omitted Genes: Folr1, Anapc15, Lamtor1
omitted Genes: Lrrc51, Numa1, Il18bp
omitted Genes: Rnf121, Xndc1, Xntrpc
omitted Genes: Trpc2, Art1, Chrna10
omitted Genes: Nup98, Pgap2, Rhog
omitted Genes: Rrm1, Olfr543, Olfr544
omitted Genes: Olfr545, Olfr547, Olfr549
omitted Genes: Olfr551, Olfr552, Olfr553
omitted Genes: Olfr554, Olfr555, Olfr556
omitted Genes: Trim68, Olfr557, Olfr558
omitted Genes: Olfr559, Olfr78, Olfr560
omitted Genes: Olfr561, Olfr564, Olfr566
omitted Genes: Olfr568, Olfr569, Olfr570
omitted Genes: Olfr571, Olfr572, Olfr574
omitted Genes: Olfr575, Olfr576, Olfr577
omitted Genes: Olfr578, Olfr582, Olfr584
omitted Genes: Olfr585, Olfr586, Olfr589
omitted Genes: Olfr591, Olfr592, Olfr593
Make more plots at http://csg.sph.umich.edu/locuszoom/
omitted Genes: Olfr594, Usp17ld, Olfr597

Fine−mapping Regions

| snp | chr | pos (Mb) | pp | group | color |
|---|---|---|---|---|---|
| chr7:101743401 | 7 | 101.7434 | 0.1909593 | Credible Set | purple |
| chr7:103659355 | 7 | 103.6594 | 0.1909593 | Credible Set | purple |
| chr7:104149022 | 7 | 104.1490 | 0.2159842 | Credible Set | purple |
| chr7:104162444 | 7 | 104.1624 | 0.1909593 | Credible Set | purple |
| chr7:104581631 | 7 | 104.5816 | 0.2088515 | Credible Set | purple |

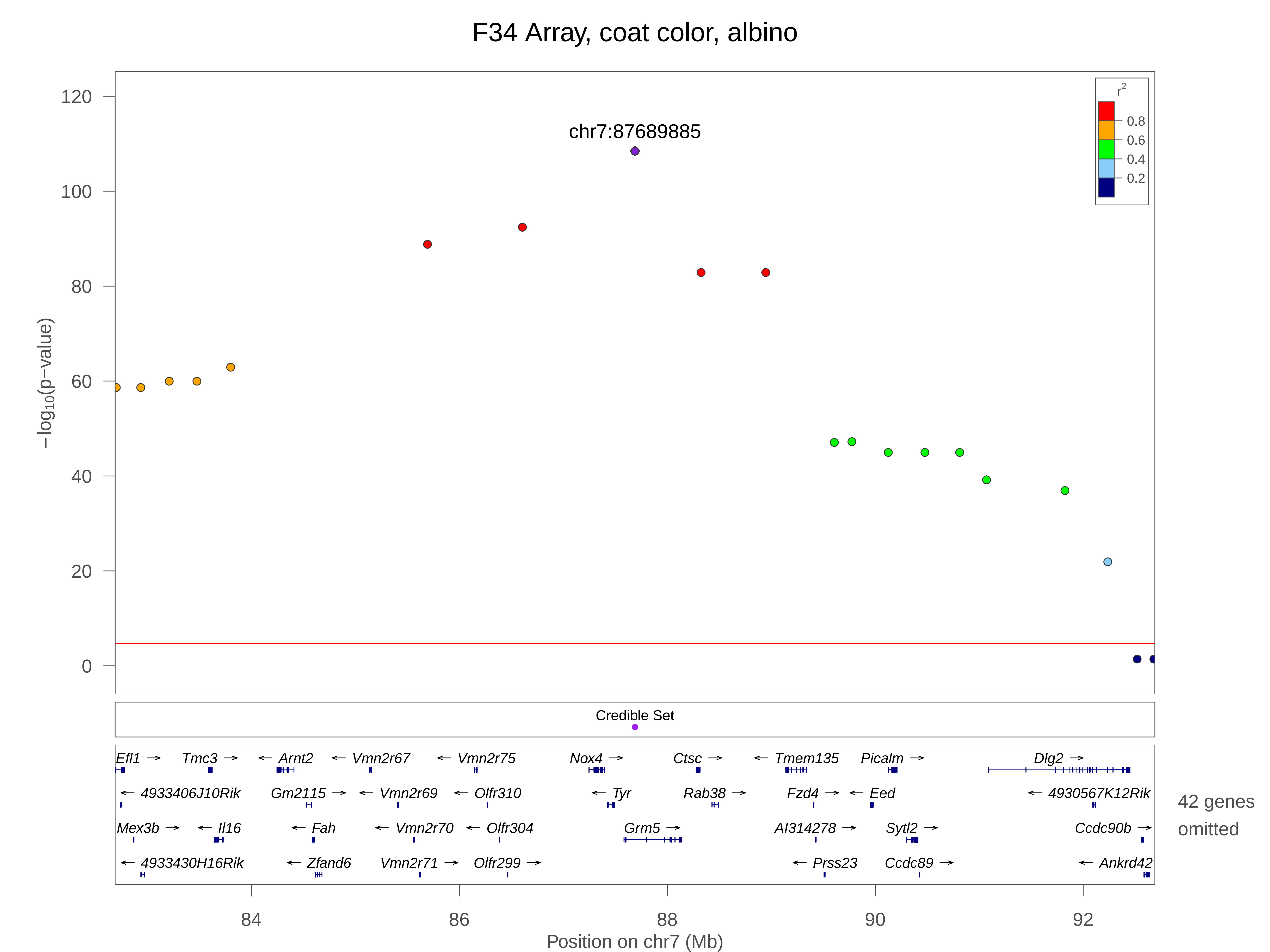

F34 Array, coat color, albino
chr7:87689885
−log10(p−value)
120
100
80
60
40
20
0
r2
0.8
0.6
0.4
0.2
Credible Set
Efl1 → Tmc3 → ← Arnt2 ← Vmn2r67 ← Vmn2r75 Nox4 → Ctsc → ← Tmem135 Picalm → Dlg2 →
← 4933406J10Rik Gm2115 → ← Vmn2r69 ← Olfr310 ← Tyr Rab38 → Fzd4 → ← Eed ← 4930567K12Rik
Mex3b → ← Il16 ← Fah ← Vmn2r70 ← Olfr304 Grm5 → AI314278 → Sytl2 → Ccdc90b →
← 4933430H16Rik ← Zfand6 Vmn2r71 → Olfr299 → ← Prss23 Ccdc89 → ← Ankrd42
42 genes omitted
84
86
88
90
92
Position on chr7 (Mb)

| | |
|---|---|
| date: | Thu Oct 25 16:53:09 2018 |
| build: | F34Array |
| display range: | chr7:82689885−92689885 [82689885−92689885] |
| hilite range: | 0 − 0 [ 0 − 0 ] |
| reference SNP: | chr7:87689885 |
| number of SNPs plotted: | 21 |
| min P−value: | 3.76E−109 [chr7:87689885] |
| max P−value: | 3.73E−2 [chr7:92679853] |
| omitted Genes: | Stard5, Cfap161, Mesdc1 |
| omitted Genes: | Mesdc2, Cemip, Abhd17c |
| omitted Genes: | 2610206C17Rik, Olfr291, Olfr290 |
| omitted Genes: | Vmn2r65, Vmn2r66, Vmn2r68 |
| omitted Genes: | Vmn2r72, Vmn2r73, Vmn2r74 |
| omitted Genes: | Vmn2r76, Olfr309, Olfr308 |
| omitted Genes: | Olfr307, Olfr305, Olfr303 |
| omitted Genes: | Olfr301, Olfr298, Olfr297 |
| omitted Genes: | Olfr295, Olfr294, Olfr293 |
| omitted Genes: | Olfr292, Folh1, Vmn2r77 |
| omitted Genes: | Vmn2r78, Vmn2r79, Me3 |
| omitted Genes: | Ccdc81, l7Rn6, E230029C05Rik |
| omitted Genes: | 2310010J17Rik, Ccdc83, Crebzf |
| omitted Genes: | Tmem126a, Tmem126b, Pcf11 |

Make more plots at http://csg.sph.umich.edu/locuszoom/

Fine−mapping Regions

| snp | chr | pos (Mb) | pp | group | color |
|---|---|---|---|---|---|
| chr7:87689885 | 7 | 87.68989 | 1 | Credible Set | purple |

F34 Array, coat color, agouti

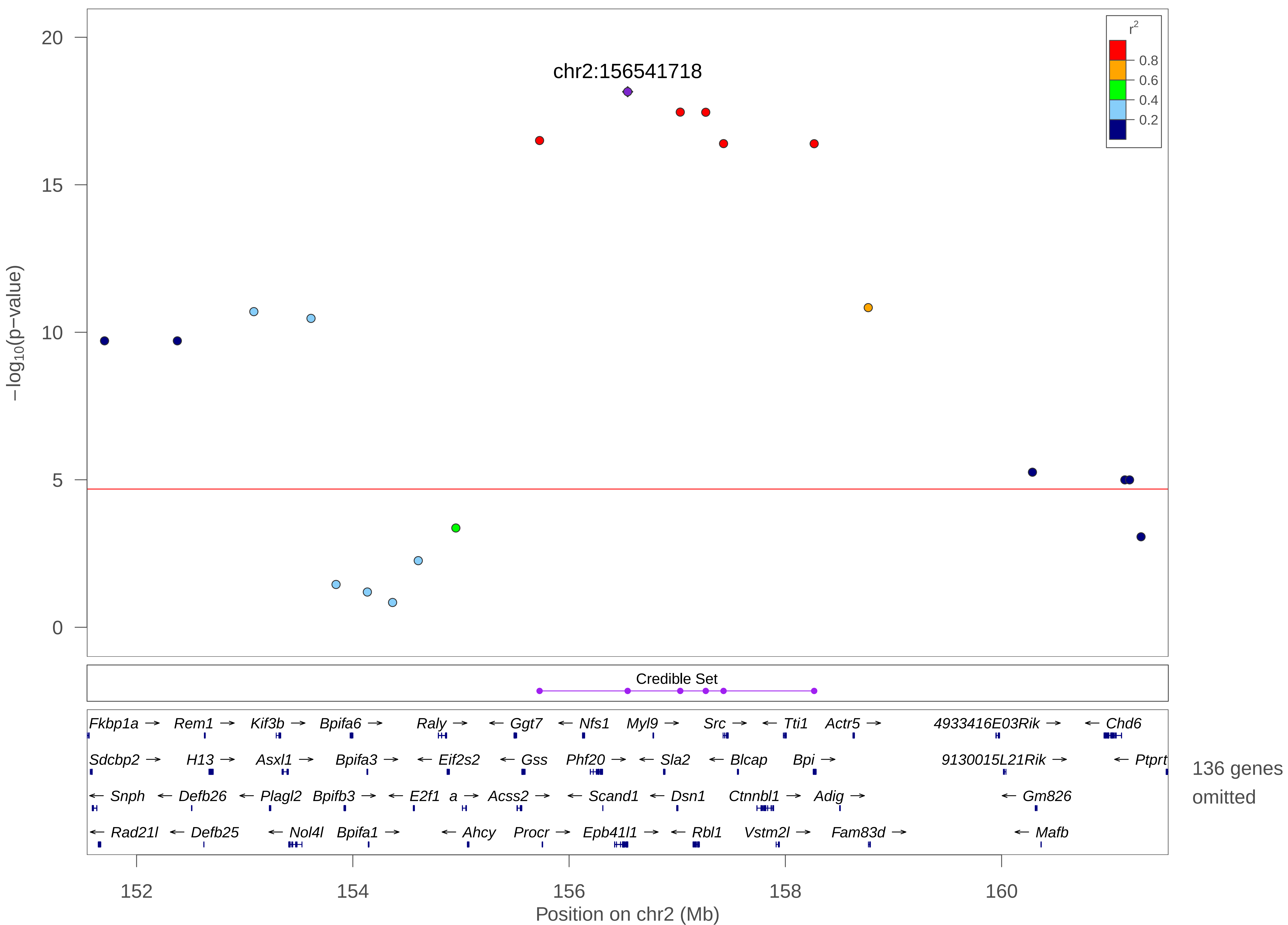

date: Thu Oct 25 16:53:19 2018
build: F34Array
display range: chr2:151541718−161541718 [151541718−161541718]
hilite range: 0 − 0 [ 0 − 0 ]
reference SNP: chr2:156541718
number of SNPs plotted: 21
min P−value: 7.02E−19 [chr2:156541718]
max P−value: 1.44E−1 [chr2:154367620]
omitted Genes: Tmem74b, Psmf1, Rspo4
omitted Genes: Angpt4, Mir1953, Fam110a
omitted Genes: Slc52a3, Scrt2, Srxn1
omitted Genes: Tcf15, Csnk2a1, Tbc1d20
omitted Genes: Rbck1, Trib3, Gm14164
omitted Genes: Nrsn2, Sox12, Zcchc3
omitted Genes: 6820408C15Rik, Defb23, Defb20
omitted Genes: Defb22, Defb28, Defb29
omitted Genes: Defb21, Defb19, Defb45
omitted Genes: Defb36, Mcts2, Id1
omitted Genes: Cox4i2, Bcl2l1, Tpx2
omitted Genes: Mir5622, Mylk2, Foxs1
omitted Genes: Dusp15, Ttll9, Pdrg1
omitted Genes: Xkr7, Ccm2l, Hck
omitted Genes: Tm9sf4, Tspyl3, Pofut1
omitted Genes: 2500004C02Rik, Commd7, Dnmt3b
omitted Genes: Mapre1, Efcab8, Sun5
omitted Genes: Bpifb2, Bpifb6, Bpifb4
omitted Genes: Bpifa2, Bpifa5, Bpifb1
omitted Genes: Bpifb5, Bpifb9a, Bpifb9b
omitted Genes: Cdk5rap1, Snta1, Cbfa2t2
omitted Genes: Necab3, 1700003F12Rik, Actl10
omitted Genes: Pxmp4, Zfp341, Chmp4b
omitted Genes: 2310005A03Rik, Itch, Dynlrb1
omitted Genes: Map1lc3a, Pigu, Mir695
omitted Genes: Trp53inp2, Ncoa6, Acss2os
omitted Genes: Myh7b, Mir499, Trpc4ap
omitted Genes: Edem2, Mmp24, BC029722
omitted Genes: Eif6, Fam83c, Uqcc1
omitted Genes: Gdf5, Cep250, 6430550D23Rik
omitted Genes: Ergic3, Fer1l4, Spag4
Make more plots at http://csg.sph.umich.edu/locuszoom/
omitted Genes: Cpne1, Rbm12, Romo1

Fine−mapping Regions

| snp | chr | pos (Mb) | pp | group | color |
|---|---|---|---|---|---|
| chr2:155727021 | 2 | 155.7270 | 0.01582778 | Credible Set | purple |
| chr2:156541718 | 2 | 156.5417 | 0.67845350 | Credible Set | purple |
| chr2:157028101 | 2 | 157.0281 | 0.14122750 | Credible Set | purple |
| chr2:157263817 | 2 | 157.2638 | 0.13963072 | Credible Set | purple |
| chr2:157428508 | 2 | 157.4285 | 0.01247776 | Credible Set | purple |
| chr2:158266555 | 2 | 158.2666 | 0.01238273 | Credible Set | purple |

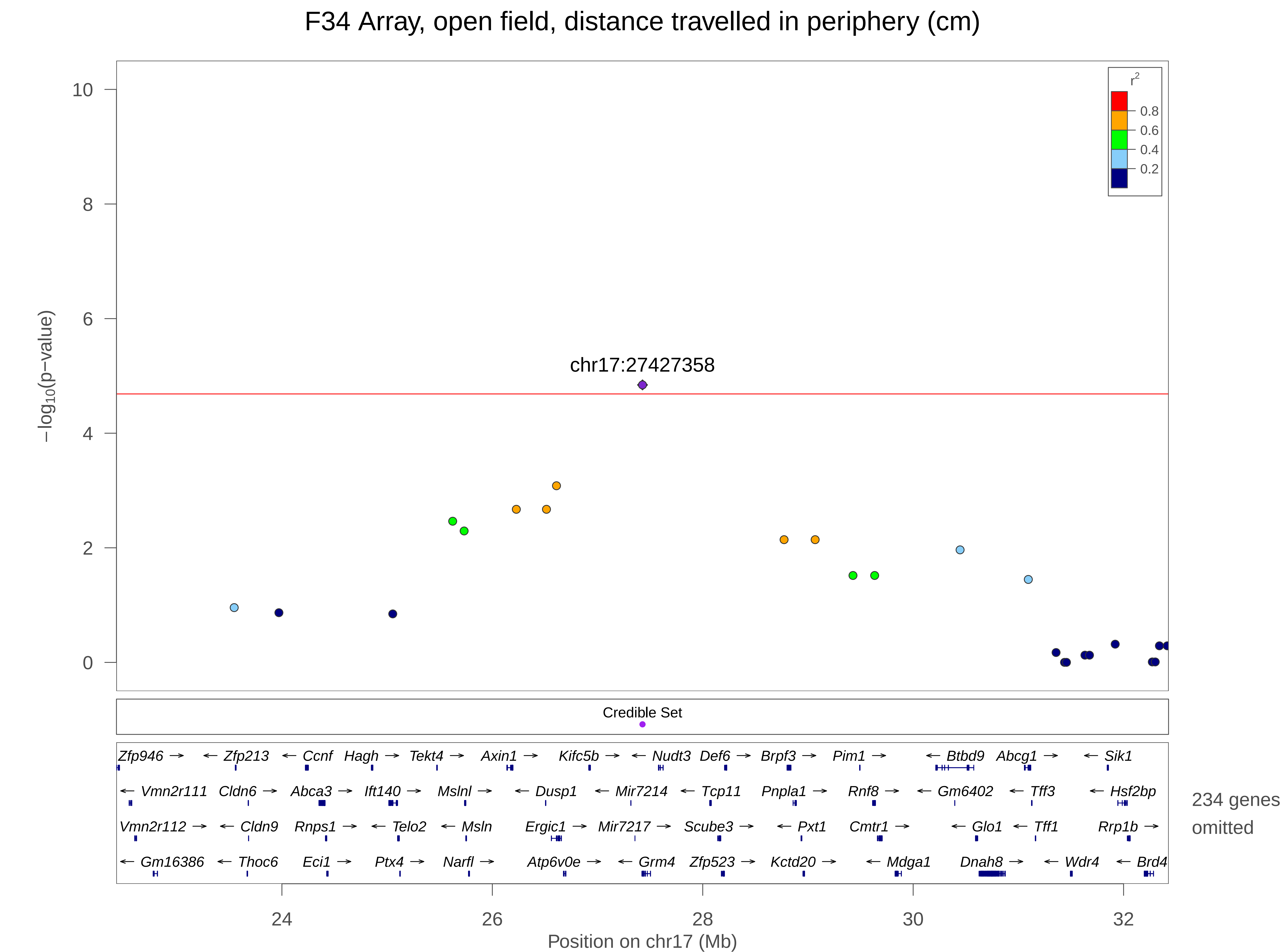

F34 Array, open field, distance travelled in periphery (cm)
−log10(p−value)
chr17:27427358
r2
0.8
0.6
0.4
0.2
Credible Set
Zfp946 → ← Zfp213 ← Ccnf Hagh → Tekt4 → Axin1 → Kifc5b → ← Nudt3 Def6 → Brpf3 → Pim1 → ← Btbd9 Abcg1 → ← Sik1
← Vmn2r111 Cldn6 → Abca3 → Ift140 → Mslnl → ← Dusp1 ← Mir7214 ← Tcp11 Pnpla1 → Rnf8 → ← Gm6402 ← Tff3 ← Hsf2bp
Vmn2r112 → ← Cldn9 Rnps1 → ← Telo2 ← Msln Ergic1 → Mir7217 → Scube3 → ← Pxt1 Cmtr1 → ← Glo1 ← Tff1 Rrp1b →
← Gm16386 ← Thoc6 Eci1 → Ptx4 → Narfl → Atp6v0e → ← Grm4 Zfp523 → Kctd20 → ← Mdga1 Dnah8 → ← Wdr4 ← Brd4
234 genes omitted
Position on chr17 (Mb)

date: Thu Oct 25 16:53:43 2018
build: F34Array
display range: chr17:22427358−32427358 [22427358−32427358]
hilite range: 0 − 0 [ 0 − 0 ]
reference SNP: chr17:27427358
number of SNPs plotted: 26
min P−value: 1.44E−5 [chr17:27427358]
max P−value: 9.97E−1 [chr17:31438721]
omitted Genes: Zfp945, Vmn2r113, Vmn2r−ps129
omitted Genes: Zfp40, Vmn2r114, Vmn2r115
omitted Genes: Vmn2r116, Vmn2r117, 6330415G19Rik
omitted Genes: Zfp13, Zscan10, Mmp25
omitted Genes: Ccdc64b, Hcfc1r1, Tnfrsf12a
omitted Genes: Pkmyt1, Paqr4, Kremen2
omitted Genes: 9530082P21Rik, Flywch1, Flywch2
omitted Genes: Srrm2, Mir5125, Tceb2
omitted Genes: Prss33, Prss41, Prss32
omitted Genes: Prss21, Dcpp1, Dcpp2
omitted Genes: Dcpp3, Sbp, Sbpl
omitted Genes: Prss30, Prss22, Prss27
omitted Genes: Kctd5, Pdpk1, Amdhd2
omitted Genes: Atp6v0c, Tbc1d24, Ntn3
omitted Genes: 1600002H07Rik, Mir5134, Mir6965
omitted Genes: Abca17, D330041H03Rik, Dnase1l2
omitted Genes: E4f1, Pgp, Mlst8
omitted Genes: Bricd5, Caskin1, Traf7
omitted Genes: Rab26os, Rab26, Pkd1
omitted Genes: Tsc2, Nthl1, Slc9a3r2
omitted Genes: Npw, Zfp598, Syngr3
omitted Genes: Gfer, Noxo1, Tbl3
omitted Genes: Rnf151, Snhg9, Snora78
omitted Genes: Rps2, Snora64, Ndufb10
omitted Genes: Rpl3l, Msrb1, Hs3st6
omitted Genes: Meiob, Fahd1, Igfals
omitted Genes: Nubp2, Spsb3, Eme2
omitted Genes: Mrps34, Nme3, Mapk8ip3
omitted Genes: Hn1l, Cramp1l, Tmem204
omitted Genes: Clcn7, Ccdc154, BC003965
omitted Genes: Unkl, Gnptg, Tsr3
Make more plots at http://csg.sph.umich.edu/locuszoom/
omitted Genes: Baiap3, Mir3547, Ube2i

Fine−mapping Regions

| snp | chr | pos (Mb) | pp | group | color |
|---|---|---|---|---|---|
| chr17:27427358 | 17 | 27.42736 | 1 | Credible Set | purple |

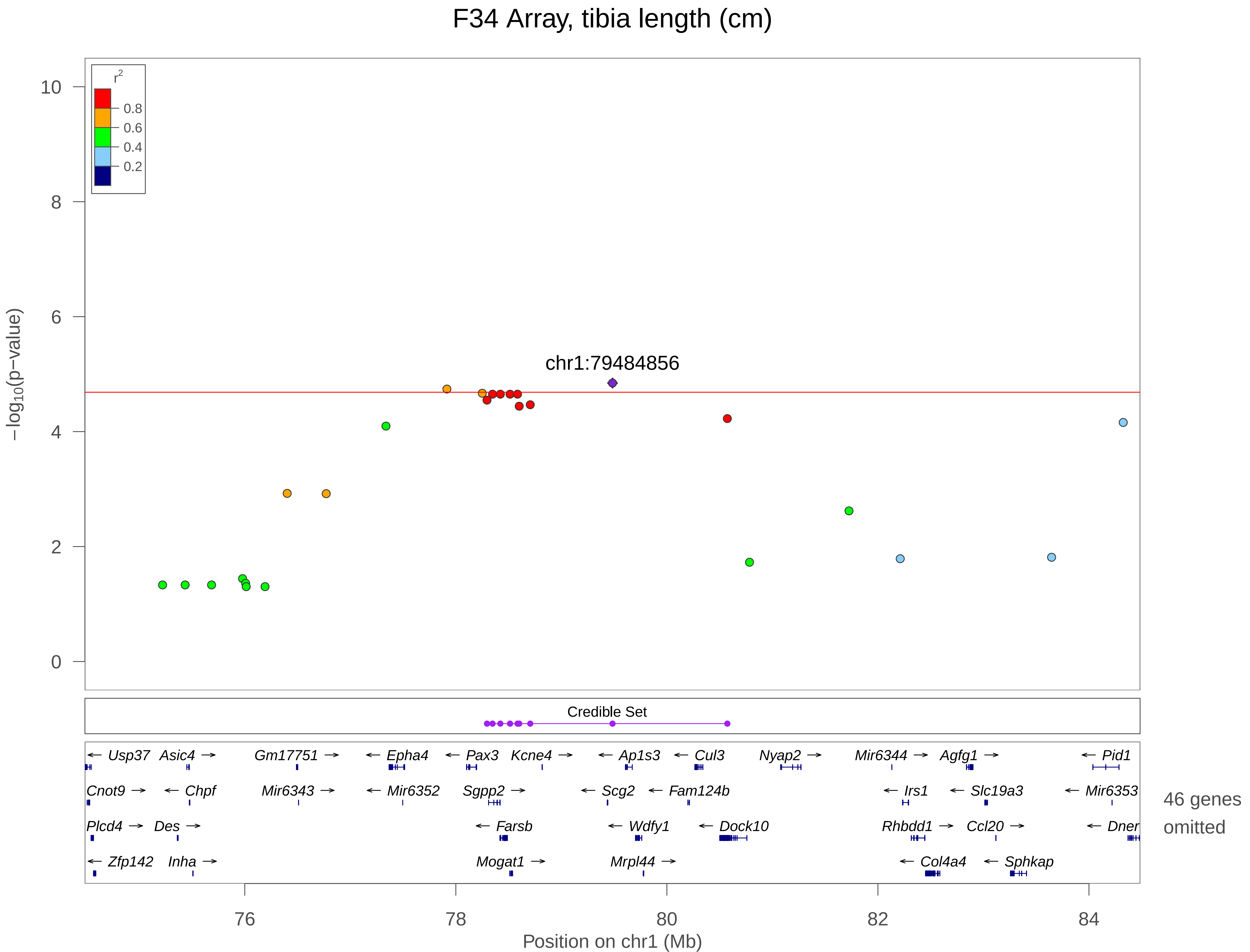

F34 Array, tibia length (cm)
r2
0.8
0.6
0.4
0.2
10
8
6
4
2
0
−log10(p−value)
chr1:79484856
Credible Set
Usp37
Asic4
Gm17751
Epha4
Pax3
Kcne4
Ap1s3
Cul3
Nyap2
Mir6344
Agfg1
Pid1
Cnot9
Chpf
Mir6343
Mir6352
Sgpp2
Scg2
Fam124b
Irs1
Slc19a3
Mir6353
Plcd4
Des
Farsb
Wdfy1
Dock10
Rhbdd1
Ccl20
Dner
Zfp142
Inha
Mogat1
Mrpl44
Col4a4
Sphkap
46 genes omitted
76
78
80
82
84
Position on chr1 (Mb)

| | |
|---|---|
| date: | Thu Oct 25 16:53:49 2018 |
| build: | F34Array |
| display range: | chr1:74484856−84484856 [74484856−84484856] |
| hilite range: | 0 − 0 [ 0 − 0 ] |
| reference SNP: | chr1:79484856 |
| number of SNPs plotted: | 27 |
| min P−value: | 1.43E−5 [chr1:79484856] |
| max P−value: | 4.98E−2 [chr1:76013512] |
| omitted Genes: | Bcs1l, Rnf25, Stk36 |
| omitted Genes: | Ttll4, Cyp27a1, Prkag3 |
| omitted Genes: | Wnt6, Wnt10a, Cdk5r2 |
| omitted Genes: | Fev, Cryba2, Mir375 |
| omitted Genes: | Ccdc108, Ihh, Nhej1 |
| omitted Genes: | Slc23a3, Cnppd1, Fam134a |
| omitted Genes: | Zfand2b, Abcb6, Atg9a |
| omitted Genes: | Ankzf1, Glb1l, Stk16 |
| omitted Genes: | Tuba4a, A630095N17Rik, Dnajb2 |
| omitted Genes: | Ptprn, Resp18, Dnpep |
| omitted Genes: | Gm15179, Speg, Gmppa |
| omitted Genes: | Tmem198, Obsl1, Stk11ip |
| omitted Genes: | Slc4a3, Utp14b, Acsl3 |
| omitted Genes: | Serpine2, 1700016L21Rik, Col4a3 |
| omitted Genes: | Mff, Tm4sf20, Krtap28−13 |
| omitted Genes: | Daw1 |

Make more plots at http://csg.sph.umich.edu/locuszoom/

Fine−mapping Regions

| snp | chr | pos (Mb) | pp | group | color |
|---|---|---|---|---|---|
| chr1:78295279 | 1 | 78.29528 | 0.09930320 | Credible Set | purple |
| chr1:78347824 | 1 | 78.34782 | 0.12464403 | Credible Set | purple |
| chr1:78421886 | 1 | 78.42189 | 0.12464403 | Credible Set | purple |
| chr1:78513810 | 1 | 78.51381 | 0.12464403 | Credible Set | purple |
| chr1:78584391 | 1 | 78.58439 | 0.12464403 | Credible Set | purple |
| chr1:78600600 | 1 | 78.60060 | 0.07883549 | Credible Set | purple |
| chr1:78705181 | 1 | 78.70518 | 0.08339751 | Credible Set | purple |
| chr1:79484856 | 1 | 79.48486 | 0.19063059 | Credible Set | purple |
| chr1:80573002 | 1 | 80.57300 | 0.04925710 | Credible Set | purple |

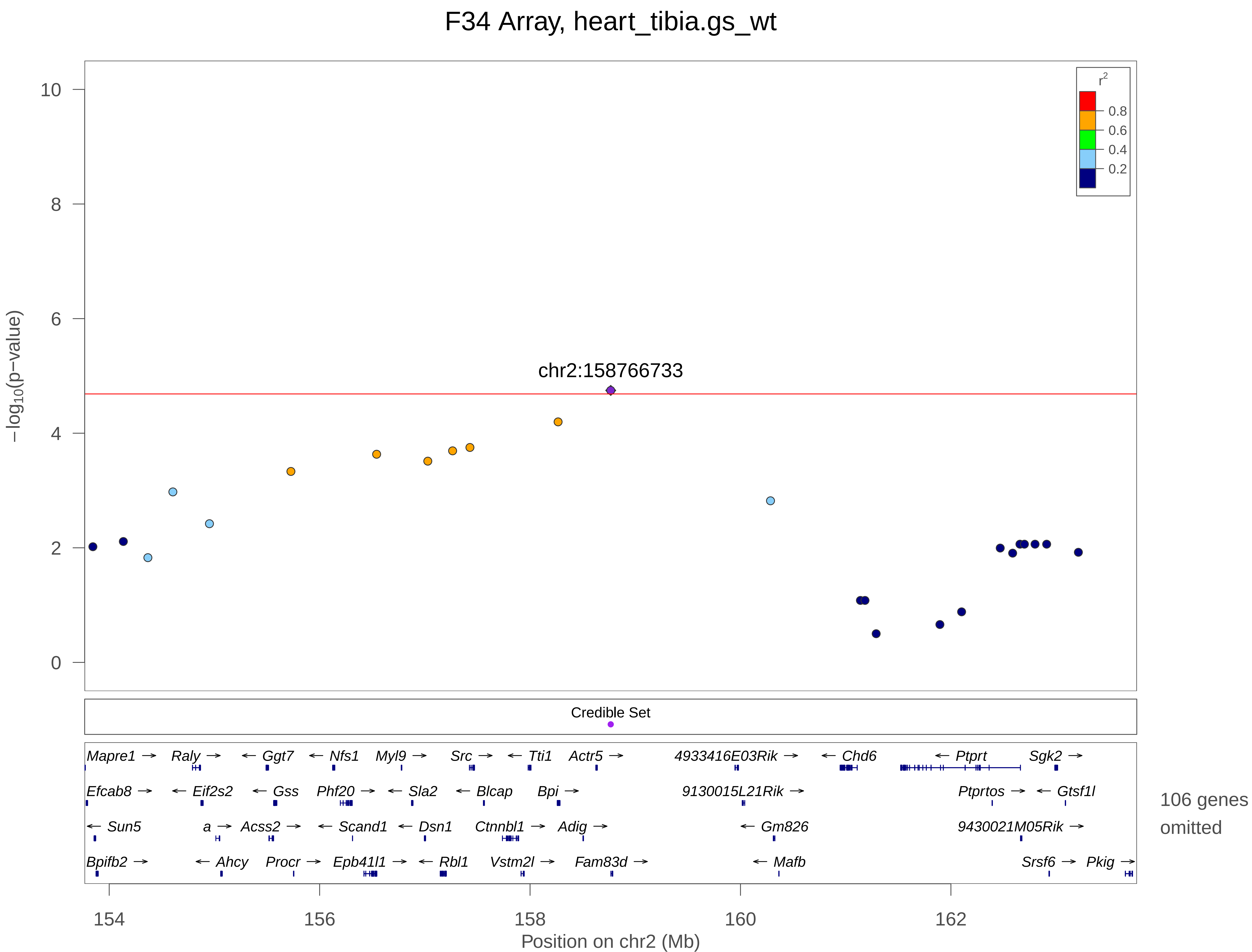

F34 Array, heart_tibia.gs_wt
r2
0.8
0.6
0.4
0.2
10
8
6
4
2
0
−log10(p−value)
chr2:158766733
Credible Set
Mapre1 →
Raly →
← Ggt7
← Nfs1
Myl9 →
Src →
← Tti1
Actr5 →
4933416E03Rik →
← Chd6
← Ptprt
Sgk2 →
Efcab8 →
← Eif2s2
← Gss
Phf20 →
← Sla2
← Blcap
Bpi →
9130015L21Rik →
Ptprtos →
← Gtsf1l
← Sun5
a →
Acss2 →
← Scand1
← Dsn1
Ctnnbl1 →
Adig →
← Gm826
9430021M05Rik →
Bpifb2 →
← Ahcy
Procr →
Epb41l1 →
← Rbl1
Vstm2l →
Fam83d →
← Mafb
Srsf6 →
Pkig →
106 genes omitted
154
156
158
160
162
Position on chr2 (Mb)

| | |
|---|---|
| date: | Thu Oct 25 16:53:57 2018 |
| build: | F34Array |
| display range: | chr2:153766733−163766733 [153766733−163766733] |
| hilite range: | 0 − 0 [ 0 − 0 ] |
| reference SNP: | chr2:158766733 |
| number of SNPs plotted: | 26 |
| min P−value: | 1.79E−5 [chr2:158766733] |
| max P−value: | 3.15E−1 [chr2:161289622] |
| omitted Genes: | Bpifb6, Bpifb3, Bpifb4 |
| omitted Genes: | Bpifa6, Bpifa2, Bpifa3 |
| omitted Genes: | Bpifa1, Bpifa5, Bpifb1 |
| omitted Genes: | Bpifb5, Bpifb9a, Bpifb9b |
| omitted Genes: | Cdk5rap1, Snta1, Cbfa2t2 |
| omitted Genes: | Necab3, 1700003F12Rik, Actl10 |
| omitted Genes: | E2f1, Pxmp4, Zfp341 |
| omitted Genes: | Chmp4b, 2310005A03Rik, Itch |
| omitted Genes: | Dynlrb1, Map1lc3a, Pigu |
| omitted Genes: | Mir695, Trp53inp2, Ncoa6 |
| omitted Genes: | Acss2os, Myh7b, Mir499 |
| omitted Genes: | Trpc4ap, Edem2, Mmp24 |
| omitted Genes: | BC029722, Eif6, Fam83c |
| omitted Genes: | Uqcc1, Gdf5, Cep250 |
| omitted Genes: | 6430550D23Rik, Ergic3, Fer1l4 |
| omitted Genes: | Spag4, Cpne1, Rbm12 |
| omitted Genes: | Romo1, Rbm39, Cnbd2 |
| omitted Genes: | 2900097C17Rik, Aar2, Gm14169 |
| omitted Genes: | Dlgap4, 4930405A21Rik, Tgif2 |
| omitted Genes: | 5430405H02Rik, 1110008F13Rik, Ndrg3 |
| omitted Genes: | Soga1, Tldc2, Samhd1 |
| omitted Genes: | Mroh8, Rpn2, Ghrh |
| omitted Genes: | Manbal, Nnat, Rprd1b |
| omitted Genes: | 2010009K17Rik, Tgm2, D630003M21Rik |
| omitted Genes: | 1700060C20Rik, Mir7685, Lbp |
| omitted Genes: | Snhg17, Snhg11, Ralgapb |
| omitted Genes: | Arhgap40, Gm14204, Slc32a1 |
| omitted Genes: | Mir3474, Ppp1r16b, Dhx35 |
| omitted Genes: | Top1, Plcg1, Zhx3 |
| omitted Genes: | Lpin3, Emilin3, L3mbtl1 |
| omitted Genes: | Ift52, Mybl2, Mir7003 |
| omitted Genes: | Tox2, Jph2, Oser1 |

Make more plots at http://csg.sph.umich.edu/locuszoom/

Fine−mapping Regions

| snp | chr | pos (Mb) | pp | group | color |
|---|---|---|---|---|---|
| chr2:158766733 | 2 | 158.7667 | 1 | Credible Set | purple |

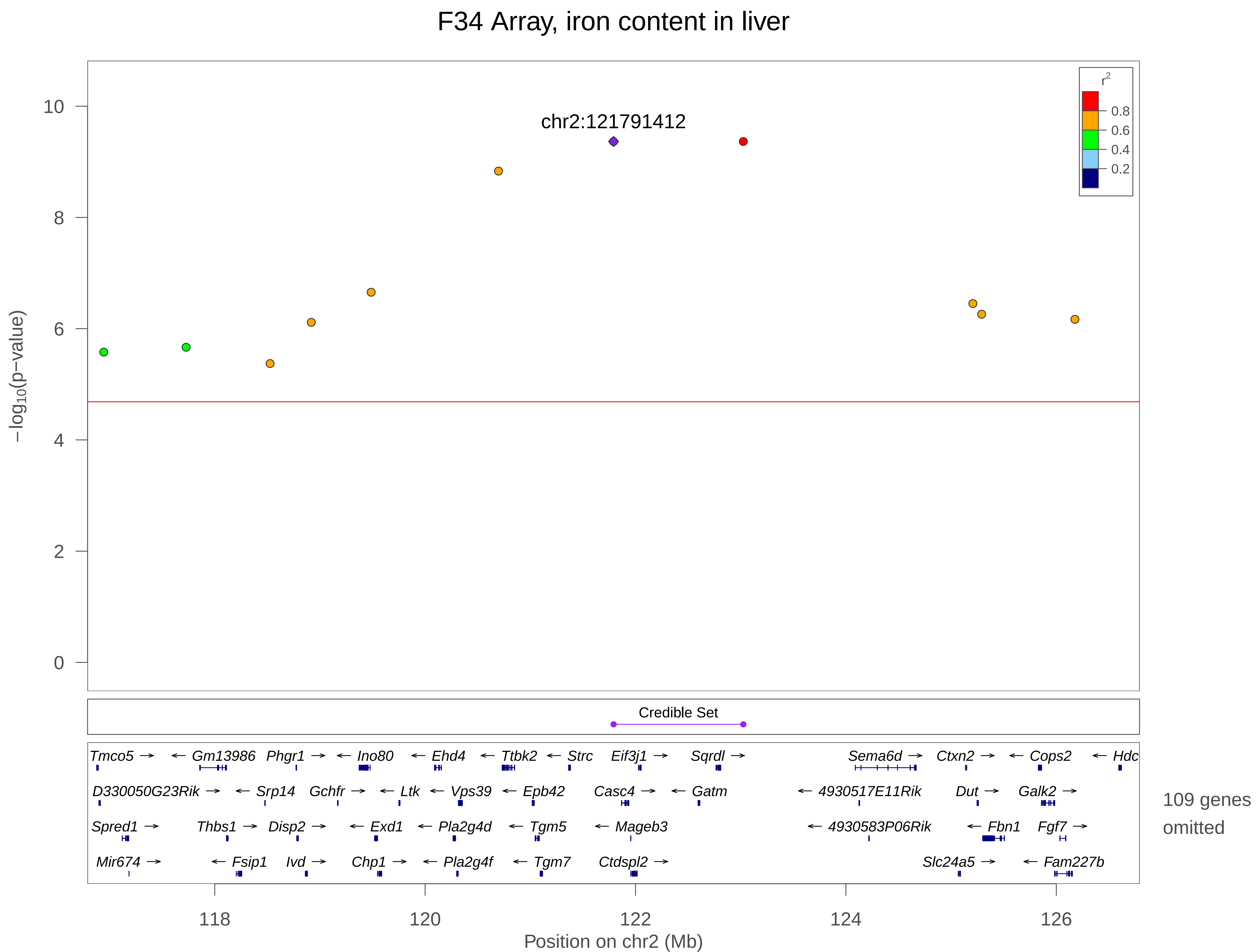

F34 Array, iron content in liver
chr2:121791412
r2
0.8
0.6
0.4
0.2
10
8
6
4
2
0
−log10(p−value)
Credible Set
Tmco5 →
← Gm13986
Phgr1 →
← Ino80
← Ehd4
← Ttbk2
← Strc
Eif3j1 →
Sqrdl →
Sema6d →
Ctxn2 →
← Cops2
← Hdc
D330050G23Rik →
← Srp14
Gchfr →
← Ltk
← Vps39
← Epb42
Casc4 →
← Gatm
← 4930517E11Rik
Dut →
Galk2 →
Spred1 →
Thbs1 →
Disp2 →
← Exd1
← Pla2g4d
← Tgm5
← Mageb3
← 4930583P06Rik
← Fbn1
Fgf7 →
Mir674 →
← Fsip1
Ivd →
Chp1 →
← Pla2g4f
← Tgm7
Ctdspl2 →
Slc24a5 →
← Fam227b
109 genes omitted
118
120
122
124
126
Position on chr2 (Mb)

date: Thu Oct 25 16:54:14 2018
build: F34Array
display range: chr2:116791412−126791412 [116791412−126791412]
hilite range: 0 − 0 [ 0 − 0 ]
reference SNP: chr2:121791412
number of SNPs plotted: 12
min P−value: 4.31E−10 [chr2:121791412]
max P−value: 4.23E−6 [chr2:118526291]
omitted Genes: Fam98b, Rasgrp1, Gm13985
omitted Genes: 4930412B13Rik, Gpr176, 1700054M17Rik
omitted Genes: Eif2ak4, Bmf, Bub1b
omitted Genes: Pak6, Ankrd63, Plcb2
omitted Genes: Inafm2, A430105I19Rik, Knstrn
omitted Genes: Bahd1, Chst14, Ccdc32
omitted Genes: Rpusd2, Casc5, Rad51
omitted Genes: Rmdn3, Dnajc17, Gm14137
omitted Genes: Zfyve19, Ppp1r14d, Spint1
omitted Genes: Rhov, Vps18, Gm14207
omitted Genes: Dll4, Chac1, 1700020I14Rik
omitted Genes: Oip5, Nusap1, Ndufaf1
omitted Genes: Rtf1, Itpka, Rpap1
omitted Genes: Tyro3, Mga, Mapkbp1
omitted Genes: Mir7665, Jmjd7, Gm28042
omitted Genes: Pla2g4b, Pla2g4e, Tmem87a
omitted Genes: Ganc, 4931402G19Rik, Capn3
omitted Genes: Zfp106, Snap23, Lrrc57
omitted Genes: Haus2, Cdan1, AV039307
omitted Genes: Ubr1, Tmem62, Ccndbp1
omitted Genes: Lcmt2, Adal, Zscan29
omitted Genes: Tubgcp4, Trp53bp1, Map1a
omitted Genes: Ppip5k1, Ckmt1, Catsper2
omitted Genes: Pdia3, Ell3, Serf2
omitted Genes: Serinc4, Hypk, Mfap1b
omitted Genes: Mfap1a, Wdr76, Frmd5
omitted Genes: Spg11, Patl2, B2m
omitted Genes: Trim69, Terb2, Sord
omitted Genes: Duox2, Duoxa2, Duoxa1
omitted Genes: Duox1, Shf, Slc28a2
omitted Genes: Bambi−ps1, Gm14085, Spata5l1
omitted Genes: AA467197, Mir147, Slc30a4

Make more plots at http://csg.sph.umich.edu/locuszoom/

Fine−mapping Regions

| snp | chr | pos (Mb) | pp | group | color |
|---|---|---|---|---|---|
| chr2:121791412 | 2 | 121.7914 | 0.5 | Credible Set | purple |
| chr2:123024804 | 2 | 123.0248 | 0.5 | Credible Set | purple |

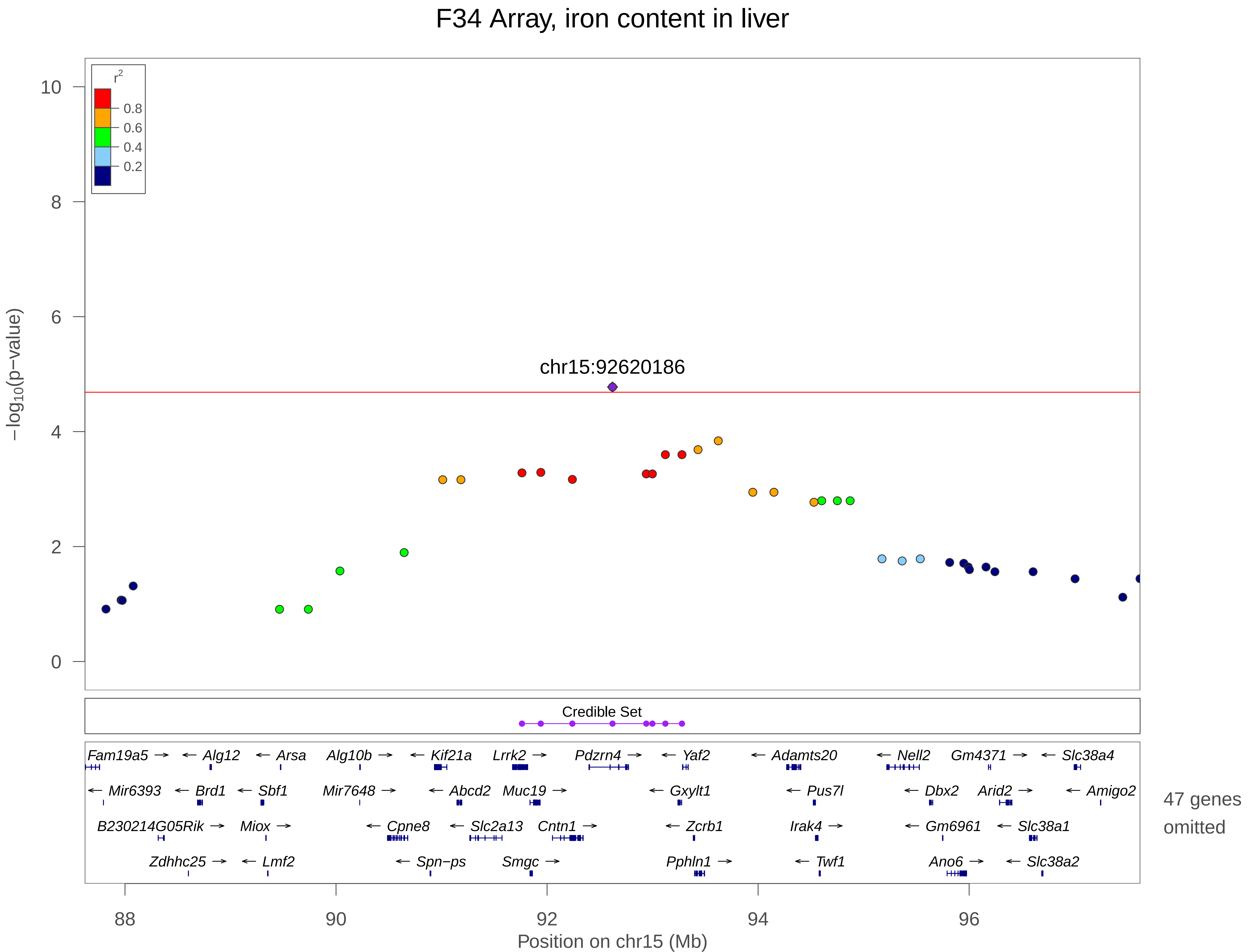

F34 Array, iron content in liver
r2
0.8
0.6
0.4
0.2
10
8
6
4
2
0
−log10(p−value)
chr15:92620186
Credible Set
Fam19a5 →
← Alg12
← Arsa
Alg10b →
← Kif21a
Lrrk2 →
Pdzrn4 →
← Yaf2
← Adamts20
← Nell2
Gm4371 →
← Slc38a4
← Mir6393
← Brd1
← Sbf1
Mir7648 →
← Abcd2
Muc19 →
← Gxylt1
← Pus7l
← Dbx2
Arid2 →
← Amigo2
B230214G05Rik →
Miox →
← Cpne8
← Slc2a13
Cntn1 →
← Zcrb1
Irak4 →
← Gm6961
← Slc38a1
Zdhhc25 →
← Lmf2
← Spn−ps
Smgc →
Pphln1 →
← Twf1
Ano6 →
← Slc38a2
47 genes omitted
88
90
92
94
96
Position on chr15 (Mb)

| | |
|---|---|
| date: | Thu Oct 25 16:54:05 2018 |
| build: | F34Array |
| display range: | chr15:87620186−97620186 [87620186−97620186] |
| hilite range: | 0 − 0 [ 0 − 0 ] |
| reference SNP: | chr15:92620186 |
| number of SNPs plotted: | 40 |
| min P−value: | 1.68E−5 [chr15:92620186] |
| max P−value: | 1.23E−1 [chr15:89464513] |
| omitted Genes: | Zbed4, Creld2, Pim3 |
| omitted Genes: | Ttll8, Mlc1, Mov10l1 |
| omitted Genes: | Panx2, 1810021B22Rik, Trabd |
| omitted Genes: | Selo, Tubgcp6, Hdac10 |
| omitted Genes: | Mapk12, Mapk11, Plxnb2 |
| omitted Genes: | Mir7118, Dennd6b, Mir6958 |
| omitted Genes: | Ppp6r2, Mir6959, Adm2 |
| omitted Genes: | Ncaph2, Sco2, Tymp |
| omitted Genes: | Odf3b, Klhdc7b, Syce3 |
| omitted Genes: | Cpt1b, Gm44502, Chkb |
| omitted Genes: | Mapk8ip2, C230037L18Rik, Shank3 |
| omitted Genes: | Acr, Rabl2, Syt10 |
| omitted Genes: | Prickle1, 9430014N10Rik, D630010B17Rik |
| omitted Genes: | Tmem117, LOC100043315, D030018L15Rik |
| omitted Genes: | 2610037D02Rik, 4833422M21Rik, E330033B04Rik |
| omitted Genes: | Scaf11, Pced1b |

Make more plots at http://csg.sph.umich.edu/locuszoom/

Fine−mapping Regions

| snp | chr | pos (Mb) | pp | group | color |
|---|---|---|---|---|---|
| chr15:91762000 | 15 | 91.76200 | 0.02909231 | Credible Set | purple |
| chr15:91940522 | 15 | 91.94052 | 0.02962923 | Credible Set | purple |
| chr15:92239534 | 15 | 92.23953 | 0.02279485 | Credible Set | purple |
| chr15:92620186 | 15 | 92.62019 | 0.74737959 | Credible Set | purple |
| chr15:92940135 | 15 | 92.94013 | 0.02798625 | Credible Set | purple |
| chr15:92997922 | 15 | 92.99792 | 0.02798625 | Credible Set | purple |
| chr15:93121022 | 15 | 93.12102 | 0.05756576 | Credible Set | purple |
| chr15:93278217 | 15 | 93.27822 | 0.05756576 | Credible Set | purple |

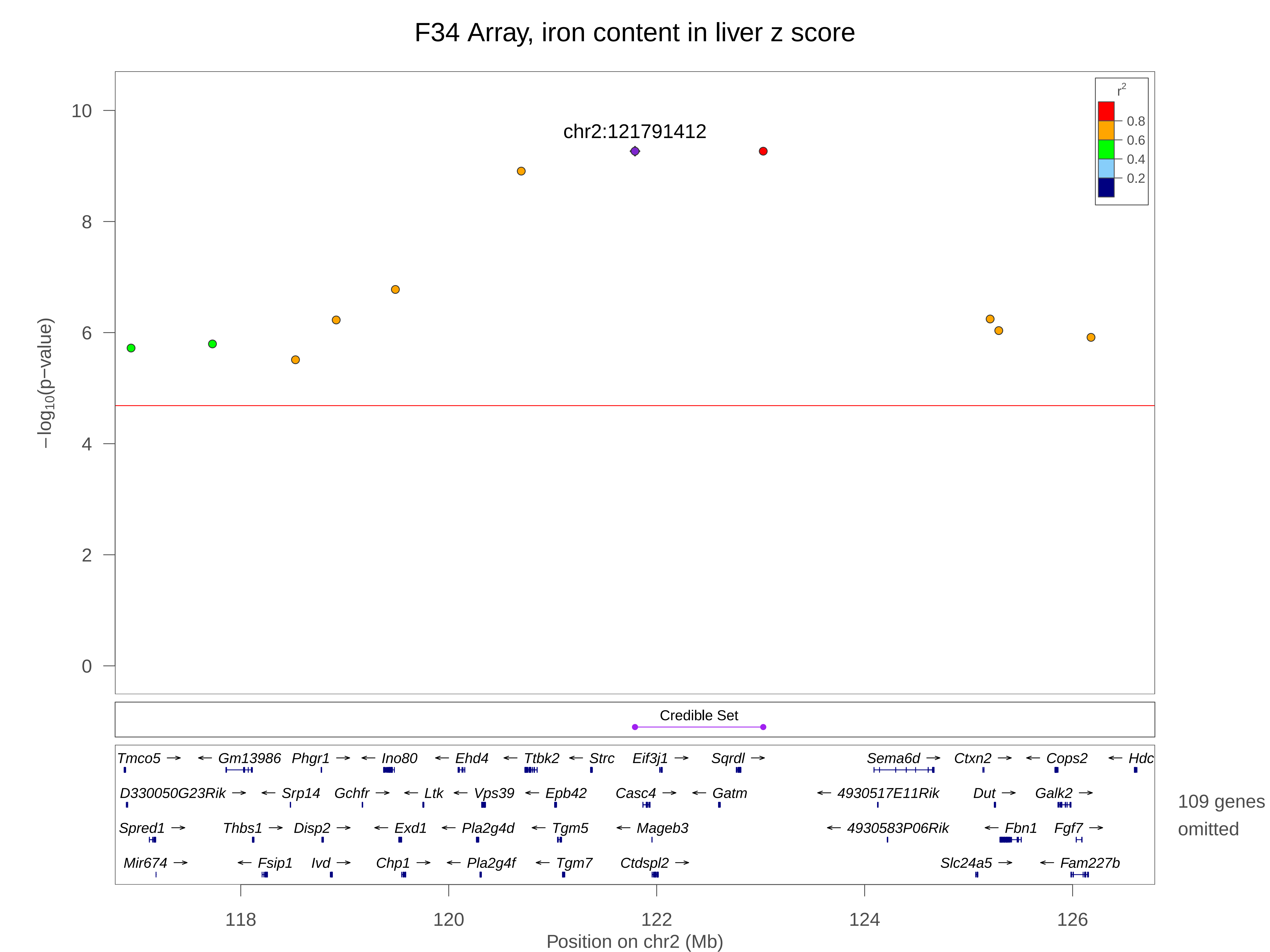
F34 Array, iron content in liver z score
chr2:121791412
r2
0.8
0.6
0.4
0.2
−log10(p−value)
Credible Set
Tmco5 →
← Gm13986
Phgr1 →
← Ino80
← Ehd4
← Ttbk2
← Strc
Eif3j1 →
Sqrdl →
Sema6d →
Ctxn2 →
← Cops2
← Hdc
D330050G23Rik →
← Srp14
Gchfr →
← Ltk
← Vps39
← Epb42
Casc4 →
← Gatm
← 4930517E11Rik
Dut →
Galk2 →
Spred1 →
Thbs1 →
Disp2 →
← Exd1
← Pla2g4d
← Tgm5
← Mageb3
← 4930583P06Rik
← Fbn1
Fgf7 →
Mir674 →
← Fsip1
Ivd →
Chp1 →
← Pla2g4f
← Tgm7
Ctdspl2 →
Slc24a5 →
← Fam227b
109 genes omitted
Position on chr2 (Mb)

date: Thu Oct 25 16:54:32 2018
build: F34Array
display range: chr2:116791412−126791412 [116791412−126791412]
hilite range: 0 − 0 [ 0 − 0 ]
reference SNP: chr2:121791412
number of SNPs plotted: 12
min P−value: 5.41E−10 [chr2:121791412]
max P−value: 3.08E−6 [chr2:118526291]
omitted Genes: Fam98b, Rasgrp1, Gm13985
omitted Genes: 4930412B13Rik, Gpr176, 1700054M17Rik
omitted Genes: Eif2ak4, Bmf, Bub1b
omitted Genes: Pak6, Ankrd63, Plcb2
omitted Genes: Inafm2, A430105I19Rik, Knstrn
omitted Genes: Bahd1, Chst14, Ccdc32
omitted Genes: Rpusd2, Casc5, Rad51
omitted Genes: Rmdn3, Dnajc17, Gm14137
omitted Genes: Zfyve19, Ppp1r14d, Spint1
omitted Genes: Rhov, Vps18, Gm14207
omitted Genes: Dll4, Chac1, 1700020I14Rik
omitted Genes: Oip5, Nusap1, Ndufaf1
omitted Genes: Rtf1, Itpka, Rpap1
omitted Genes: Tyro3, Mga, Mapkbp1
omitted Genes: Mir7665, Jmjd7, Gm28042
omitted Genes: Pla2g4b, Pla2g4e, Tmem87a
omitted Genes: Ganc, 4931402G19Rik, Capn3
omitted Genes: Zfp106, Snap23, Lrrc57
omitted Genes: Haus2, Cdan1, AV039307
omitted Genes: Ubr1, Tmem62, Ccndbp1
omitted Genes: Lcmt2, Adal, Zscan29
omitted Genes: Tubgcp4, Trp53bp1, Map1a
omitted Genes: Ppip5k1, Ckmt1, Catsper2
omitted Genes: Pdia3, Ell3, Serf2
omitted Genes: Serinc4, Hypk, Mfap1b
omitted Genes: Mfap1a, Wdr76, Frmd5
omitted Genes: Spg11, Patl2, B2m
omitted Genes: Trim69, Terb2, Sord
omitted Genes: Duox2, Duoxa2, Duoxa1
omitted Genes: Duox1, Shf, Slc28a2
omitted Genes: Bambi−ps1, Gm14085, Spata5l1
Make more plots at http://csg.sph.umich.edu/locuszoom/
omitted Genes: AA467197, Mir147, Slc30a4

Fine−mapping Regions

| snp | chr | pos (Mb) | pp | group | color |
|---|---|---|---|---|---|
| chr2:121791412 | 2 | 121.7914 | 0.5 | Credible Set | purple |
| chr2:123024804 | 2 | 123.0248 | 0.5 | Credible Set | purple |

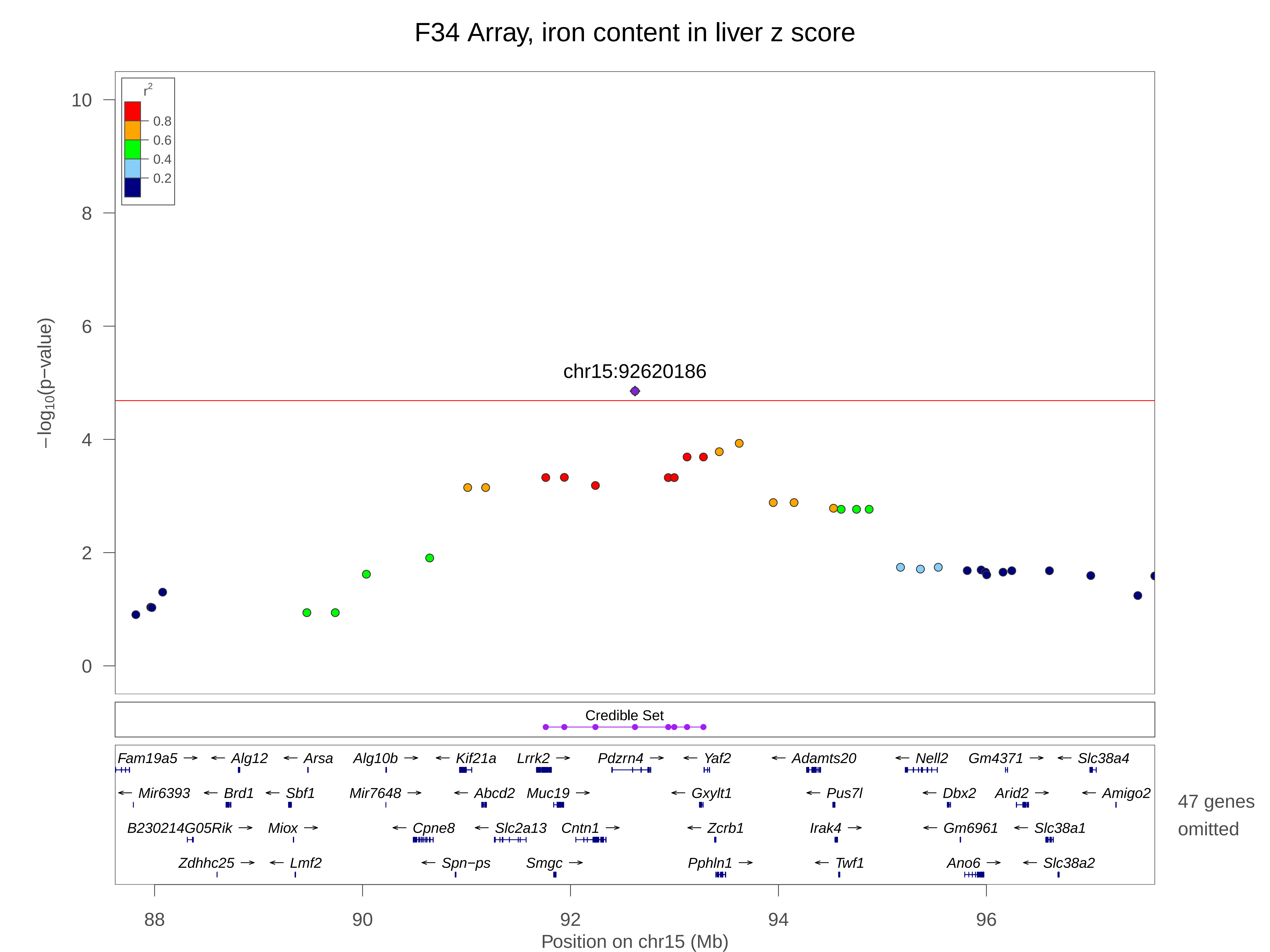
F34 Array, iron content in liver z score
r2
0.8
0.6
0.4
0.2
10
8
6
4
2
0
−log10(p−value)
chr15:92620186
Credible Set
Fam19a5 →
← Alg12
← Arsa
Alg10b →
← Kif21a
Lrrk2 →
Pdzrn4 →
← Yaf2
← Adamts20
← Nell2
Gm4371 →
← Slc38a4
← Mir6393
← Brd1
← Sbf1
Mir7648 →
← Abcd2
Muc19 →
← Gxylt1
← Pus7l
← Dbx2
Arid2 →
← Amigo2
B230214G05Rik →
Miox →
← Cpne8
← Slc2a13
Cntn1 →
← Zcrb1
Irak4 →
← Gm6961
← Slc38a1
Zdhhc25 →
← Lmf2
← Spn−ps
Smgc →
Pphln1 →
← Twf1
Ano6 →
← Slc38a2
47 genes omitted
88
90
92
94
96
Position on chr15 (Mb)

| | |
|---|---|
| date: | Thu Oct 25 16:54:23 2018 |
| build: | F34Array |
| display range: | chr15:87620186−97620186 [87620186−97620186] |
| hilite range: | 0 − 0 [ 0 − 0 ] |
| reference SNP: | chr15:92620186 |
| number of SNPs plotted: | 40 |
| min P−value: | 1.4E−5 [chr15:92620186] |
| max P−value: | 1.25E−1 [chr15:87819815] |
| omitted Genes: | Zbed4, Creld2, Pim3 |
| omitted Genes: | Ttll8, Mlc1, Mov10l1 |
| omitted Genes: | Panx2, 1810021B22Rik, Trabd |
| omitted Genes: | Selo, Tubgcp6, Hdac10 |
| omitted Genes: | Mapk12, Mapk11, Plxnb2 |
| omitted Genes: | Mir7118, Dennd6b, Mir6958 |
| omitted Genes: | Ppp6r2, Mir6959, Adm2 |
| omitted Genes: | Ncaph2, Sco2, Tymp |
| omitted Genes: | Odf3b, Klhdc7b, Syce3 |
| omitted Genes: | Cpt1b, Gm44502, Chkb |
| omitted Genes: | Mapk8ip2, C230037L18Rik, Shank3 |
| omitted Genes: | Acr, Rabl2, Syt10 |
| omitted Genes: | Prickle1, 9430014N10Rik, D630010B17Rik |
| omitted Genes: | Tmem117, LOC100043315, D030018L15Rik |
| omitted Genes: | 2610037D02Rik, 4833422M21Rik, E330033B04Rik |
| omitted Genes: | Scaf11, Pced1b |

Make more plots at http://csg.sph.umich.edu/locuszoom/

Fine−mapping Regions

| snp | chr | pos (Mb) | pp | group | color |
|---|---|---|---|---|---|
| chr15:91762000 | 15 | 91.76200 | 0.02714178 | Credible Set | purple |
| chr15:91940522 | 15 | 91.94052 | 0.02732672 | Credible Set | purple |
| chr15:92239534 | 15 | 92.23953 | 0.02006520 | Credible Set | purple |
| chr15:92620186 | 15 | 92.62019 | 0.75258098 | Credible Set | purple |
| chr15:92940135 | 15 | 92.94013 | 0.02707327 | Credible Set | purple |
| chr15:92997922 | 15 | 92.99792 | 0.02707327 | Credible Set | purple |
| chr15:93121022 | 15 | 93.12102 | 0.05936939 | Credible Set | purple |
| chr15:93278217 | 15 | 93.27822 | 0.05936939 | Credible Set | purple |

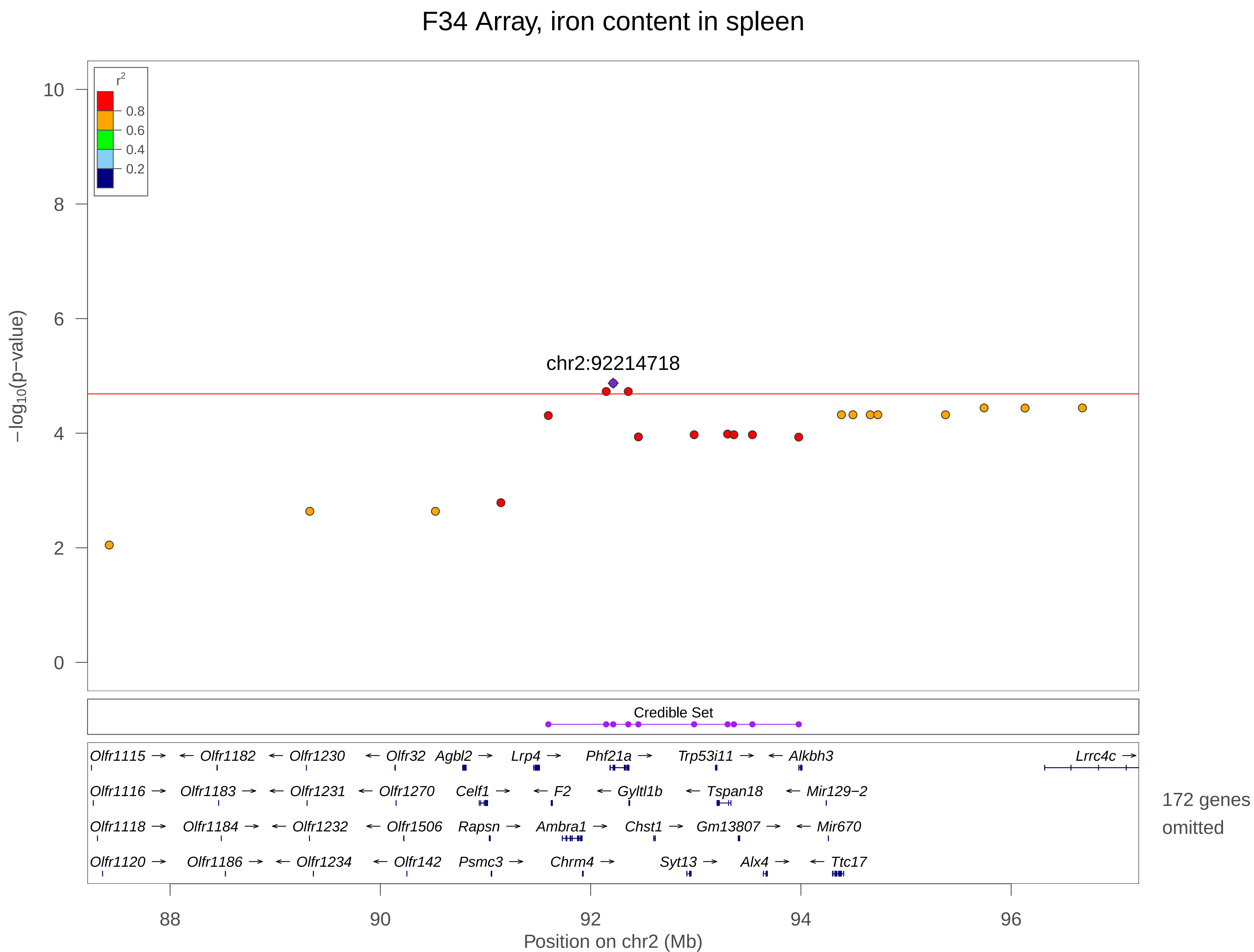
F34 Array, iron content in spleen
r2
0.8
0.6
0.4
0.2
10
8
6
4
2
0
−log10(p−value)
chr2:92214718
Credible Set
Olfr1115 →
← Olfr1182
← Olfr1230
← Olfr32
Agbl2 →
Lrp4 →
Phf21a →
Trp53i11 →
← Alkbh3
Lrrc4c →
Olfr1116 →
Olfr1183 →
← Olfr1231
← Olfr1270
Celf1 →
← F2
← Gyltl1b
← Tspan18
← Mir129−2
Olfr1118 →
Olfr1184 →
← Olfr1232
← Olfr1506
Rapsn →
Ambra1 →
Chst1 →
Gm13807 →
← Mir670
Olfr1120 →
Olfr1186 →
← Olfr1234
← Olfr142
Psmc3 →
Chrm4 →
Syt13 →
Alx4 →
← Ttc17
172 genes omitted
88
90
92
94
96
Position on chr2 (Mb)

date: Thu Oct 25 16:54:50 2018
build: F34Array
display range: chr2:87214718−97214718 [87214718−97214718]
hilite range: 0 − 0 [ 0 − 0 ]
reference SNP: chr2:92214718
number of SNPs plotted: 23
min P−value: 1.34E−5 [chr2:92214718]
max P−value: 8.94E−3 [chr2:87421495]
omitted Genes: Olfr1121, Olfr1122, Olfr1123
omitted Genes: Olfr1124, Olfr1126, Pramel7
omitted Genes: Pramel6, Olfr153, Olfr1128
omitted Genes: Olfr1129, Olfr1130, Olfr1131
omitted Genes: Olfr1132, Olfr1133, Olfr1134
omitted Genes: Olfr1135, Olfr1136, Olfr1137
omitted Genes: Olfr1138, Olfr1140, Olfr1141
omitted Genes: Olfr152, Olfr1143, Olfr1145
omitted Genes: Olfr1148, Olfr1151, Olfr1152
omitted Genes: Olfr1153, Olfr1154, Olfr1155
omitted Genes: Olfr1156, Olfr1157, Olfr74
omitted Genes: Olfr1158, Olfr1160, Olfr1161
omitted Genes: Olfr73, Olfr1162, Olfr1163
omitted Genes: Olfr1164, Olfr1166, Olfr1167
omitted Genes: Olfr1168, Olfr1170, Olfr1173
omitted Genes: Olfr1176, Olfr1178, Olfr1179
omitted Genes: Olfr1180, Olfr1181, Olfr1188
omitted Genes: Olfr1189, Olfr1193, Olfr1195
omitted Genes: Olfr1196, Olfr1197, Olfr1198
omitted Genes: Olfr1199, Olfr1200, Olfr1201
omitted Genes: Olfr1202, Olfr1205, Olfr1204
omitted Genes: Olfr1206, Olfr1208, Olfr1209
omitted Genes: Olfr1211, Olfr1212, Olfr1213
omitted Genes: Olfr1214, Olfr1215, Olfr1216
omitted Genes: Olfr1217, Olfr1218, Olfr1219
omitted Genes: Olfr1220, Olfr1221, Olfr1222
omitted Genes: Olfr1223, Olfr1225, Olfr1226
omitted Genes: Olfr1228, Olfr1229, Olfr1233
omitted Genes: Olfr1238, Olfr1239, Olfr1240
omitted Genes: Olfr1241, Olfr1242, Olfr1243
omitted Genes: Olfr1245, Olfr1246, Olfr1247
Make more plots at http://csg.sph.umich.edu/locuszoom/
omitted Genes: Olfr1248, Olfr1249, Olfr1250

Fine−mapping Regions

| snp | chr | pos (Mb) | pp | group | color |
|---|---|---|---|---|---|
| chr2:91597466 | 2 | 91.59747 | 0.08123578 | Credible Set | purple |
| chr2:92147975 | 2 | 92.14798 | 0.20341544 | Credible Set | purple |
| chr2:92214718 | 2 | 92.21472 | 0.27947380 | Credible Set | purple |
| chr2:92357966 | 2 | 92.35797 | 0.20341544 | Credible Set | purple |
| chr2:92454070 | 2 | 92.45407 | 0.03604329 | Credible Set | purple |
| chr2:92984633 | 2 | 92.98463 | 0.03912820 | Credible Set | purple |
| chr2:93303076 | 2 | 93.30308 | 0.04020689 | Credible Set | purple |
| chr2:93362468 | 2 | 93.36247 | 0.03912820 | Credible Set | purple |
| chr2:93538754 | 2 | 93.53875 | 0.03912820 | Credible Set | purple |
| chr2:93979130 | 2 | 93.97913 | 0.03576123 | Credible Set | purple |

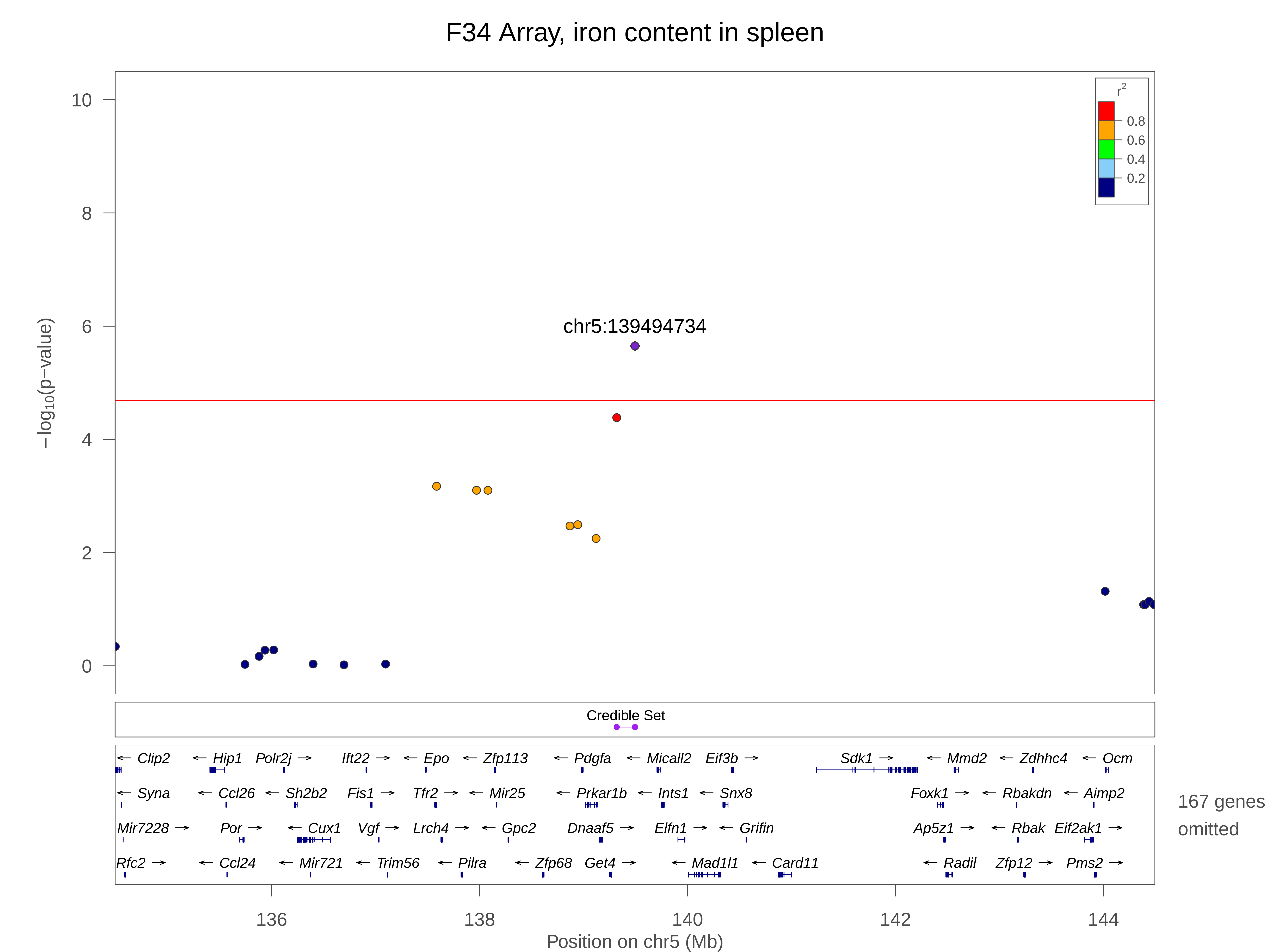
F34 Array, iron content in spleen
chr5:139494734
−log10(p−value)
r2
0.8
0.6
0.4
0.2
Credible Set
← Clip2 ← Hip1 Polr2j → Ift22 → ← Epo ← Zfp113 ← Pdgfa ← Micall2 Eif3b → Sdk1 → ← Mmd2 ← Zdhhc4 ← Ocm
← Syna ← Ccl26 ← Sh2b2 Fis1 → Tfr2 → ← Mir25 ← Prkar1b ← Ints1 ← Snx8 Foxk1 → ← Rbakdn ← Aimp2
Mir7228 → Por → ← Cux1 Vgf → Lrch4 → ← Gpc2 Dnaaf5 → Elfn1 → ← Grifin Ap5z1 → ← Rbak Eif2ak1 →
Rfc2 → ← Ccl24 ← Mir721 ← Trim56 ← Pilra ← Zfp68 Get4 → ← Mad1l1 ← Card11 ← Radil Zfp12 → Pms2 →
167 genes omitted
Position on chr5 (Mb)

date: Thu Oct 25 16:55:01 2018
build: F34Array
display range: chr5:134494734−144494734 [134494734−144494734]
hilite range: 0 − 0 [ 0 − 0 ]
reference SNP: chr5:139494734
number of SNPs plotted: 22
min P−value: 2.24E−6 [chr5:139494734]
max P−value: 9.6E−1 [chr5:136695919]
omitted Genes: Lat2, Eif4h, Limk1
omitted Genes: Eln, Wbscr28, Cldn13
omitted Genes: Wbscr27, Cldn4, Cldn3
omitted Genes: Wbscr25, Abhd11, Abhd11os
omitted Genes: Stx1a, Wbscr22, Dnajc30
omitted Genes: Vps37d, Mlxipl, Tbl2
omitted Genes: Bcl7b, Baz1b, Fzd9
omitted Genes: Fkbp6, Trim50, Nsun5
omitted Genes: Pom121, Mir7033, Rhbdd2
omitted Genes: Tmem120a, Mir7034, Styxl1
omitted Genes: Mdh2, Srrm3, Hspb1
omitted Genes: Ywhag, Ssc4d, Zp3
omitted Genes: Dtx2, Upk3b, Upk3bl
omitted Genes: Rasa4, Mir7035, Lrwd1
omitted Genes: Alkbh4, Orai2, Prkrip1
omitted Genes: Myl10, Col26a1, 4933404O12Rik
omitted Genes: Cldn15, Znhit1, Plod3
omitted Genes: Mir702, Ap1s1, Serpine1
omitted Genes: Ache, Mir8116, Ufsp1
omitted Genes: Srrt, Mir7036, Trip6
omitted Genes: Slc12a9, Ephb4, Zan
omitted Genes: Pop7, Gigyf1, Gnb2
omitted Genes: Mir6418, Actl6b, Mospd3
omitted Genes: Pcolce, Fbxo24, Gm20605
omitted Genes: Sap25, Irs3, Agfg2
omitted Genes: Nyap1, Tsc22d4, Ppp1r35
omitted Genes: Mepce, Zcwpw1, Pilrb1
omitted Genes: Pilrb2, Cyp3a13, Gjc3
omitted Genes: Azgp1, Smok3a, Smok3b
omitted Genes: Smok3c, Zkscan1, Zscan21
omitted Genes: Cops6, Mcm7, Mir93
omitted Genes: Mir106b, Ap4m1, Taf6

Fine−mapping Regions

| snp | chr | pos (Mb) | pp | group | color |
|---|---|---|---|---|---|
| chr5:139318848 | 5 | 139.3188 | 0.05815931 | Credible Set | purple |
| chr5:139494734 | 5 | 139.4947 | 0.94184069 | Credible Set | purple |

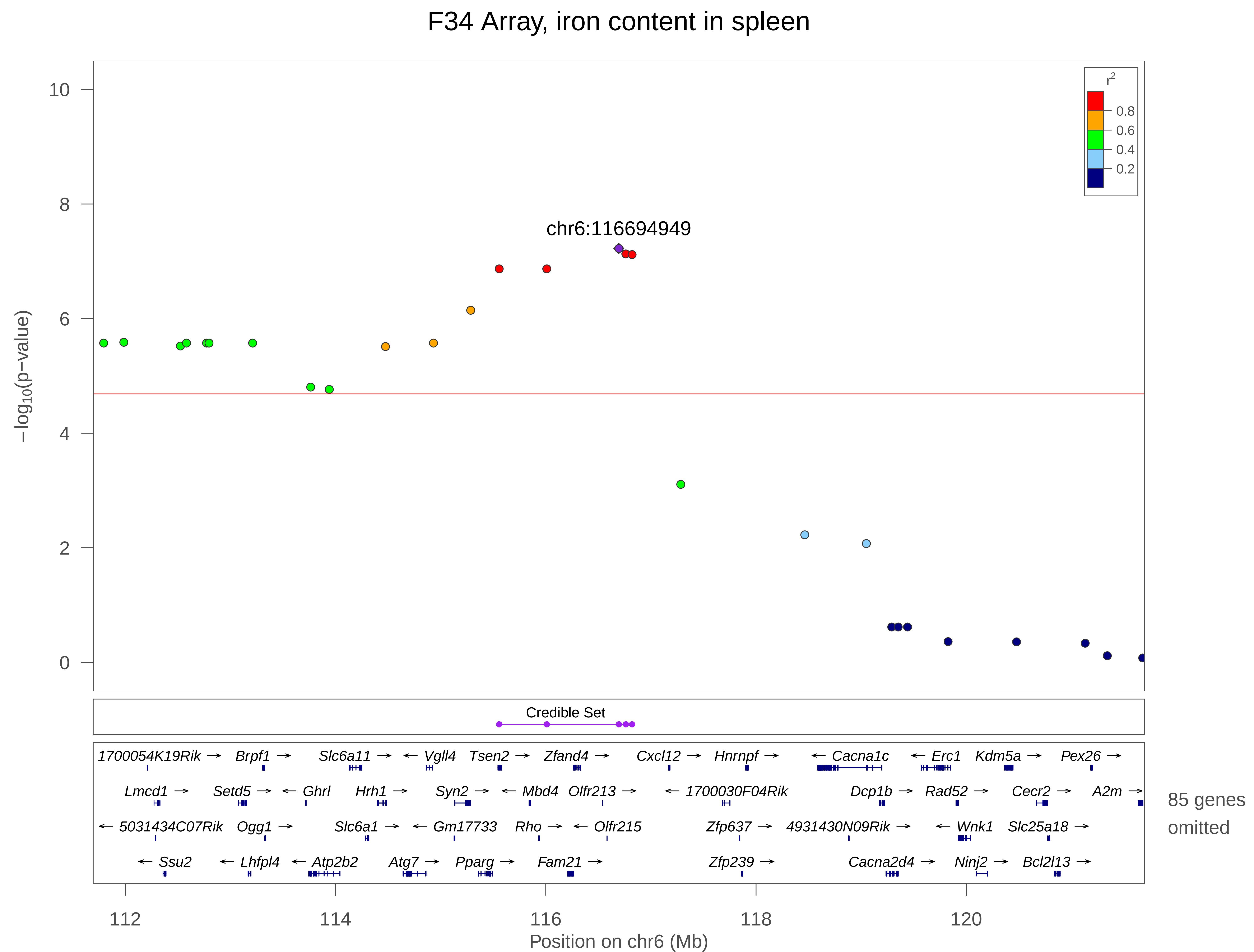
F34 Array, iron content in spleen
r2
0.8
0.6
0.4
0.2
chr6:116694949
Credible Set
1700054K19Rik → Brpf1 → Slc6a11 → ← Vgll4 Tsen2 → Zfand4 → Cxcl12 → Hnrnpf → ← Cacna1c ← Erc1 Kdm5a → Pex26 →
Lmcd1 → Setd5 → ← Ghrl Hrh1 → Syn2 → ← Mbd4 Olfr213 → ← 1700030F04Rik Dcp1b → Rad52 → Cecr2 → A2m →
← 5031434C07Rik Ogg1 → Slc6a1 → ← Gm17733 Rho → ← Olfr215 Zfp637 → 4931430N09Rik → ← Wnk1 Slc25a18 →
← Ssu2 ← Lhfpl4 ← Atp2b2 Atg7 → Pparg → Fam21 → Zfp239 → Cacna2d4 → Ninj2 → Bcl2l13 →
85 genes omitted
−log10(p−value)
Position on chr6 (Mb)
0
2
4
6
8
10
112
114
116
118
120

| | |
|---|---|
| date: | Thu Oct 25 16:55:10 2018 |
| build: | F34Array |
| display range: | chr6:111694949−121694949 [111694949−121694949] |
| hilite range: | 0 − 0 [ 0 − 0 ] |
| reference SNP: | chr6:116694949 |
| number of SNPs plotted: | 29 |
| min P−value: | 5.97E−8 [chr6:116694949] |
| max P−value: | 8.36E−1 [chr6:121677793] |
| omitted Genes: | Cav3, Oxtr, Rad18 |
| omitted Genes: | Srgap3, Thumpd3, Gt(ROSA)26Sor |
| omitted Genes: | Mtmr14, Cpne9, Camk1 |
| omitted Genes: | Tada3, Arpc4, Ttll3 |
| omitted Genes: | Rpusd3, Cidec, Jagn1 |
| omitted Genes: | Il17re, Il17rc, Creld1 |
| omitted Genes: | Prrt3, Emc3, Fancd2 |
| omitted Genes: | Fancd2os, Brk1, Vhl |
| omitted Genes: | Irak2, Tatdn2, Mir7042 |
| omitted Genes: | Sec13, Tamm41, Timp4 |
| omitted Genes: | Mkrn2os, Mkrn2, Raf1 |
| omitted Genes: | Tmem40, Cand2, Mir7660 |
| omitted Genes: | Rpl32, Snora7a, Efcab12 |
| omitted Genes: | Ift122, H1foo, Plxnd1 |
| omitted Genes: | Tmcc1, 9530062K07Rik, March8 |
| omitted Genes: | Alox5, Olfr211, Olfr212 |
| omitted Genes: | Olfr214, Zfp422, Rassf4 |
| omitted Genes: | Mir7043, 8430408G22Rik, Tmem72 |
| omitted Genes: | Fxyd4, Rasgef1a, Mir7044 |
| omitted Genes: | Csgalnact2, Ret, 1700069P05Rik |
| omitted Genes: | Gm30498, Bms1, Zfp248 |
| omitted Genes: | Zfp9, Ankrd26, Lrtm2 |
| omitted Genes: | Adipor2, Wnt5b, Fbxl14 |
| omitted Genes: | 3110021A11Rik, Mir706, B4galnt3 |
| omitted Genes: | Ccdc77, Il17ra, Cecr6 |
| omitted Genes: | Cecr5, 1700072O05Rik, Atp6v1e1 |
| omitted Genes: | Bid, Mical3, Tuba8 |
| omitted Genes: | Usp18, Slc6a13, Slc6a12 |
| omitted Genes: | Iqsec3 |

Make more plots at http://csg.sph.umich.edu/locuszoom/

Fine−mapping Regions

| snp | chr | pos (Mb) | pp | group | color |
|---|---|---|---|---|---|
| chr6:115556641 | 6 | 115.5566 | 0.1292022 | Credible Set | purple |
| chr6:116009500 | 6 | 116.0095 | 0.1292022 | Credible Set | purple |
| chr6:116694949 | 6 | 116.6949 | 0.2854410 | Credible Set | purple |
| chr6:116761437 | 6 | 116.7614 | 0.2310870 | Credible Set | purple |
| chr6:116820969 | 6 | 116.8210 | 0.2250676 | Credible Set | purple |

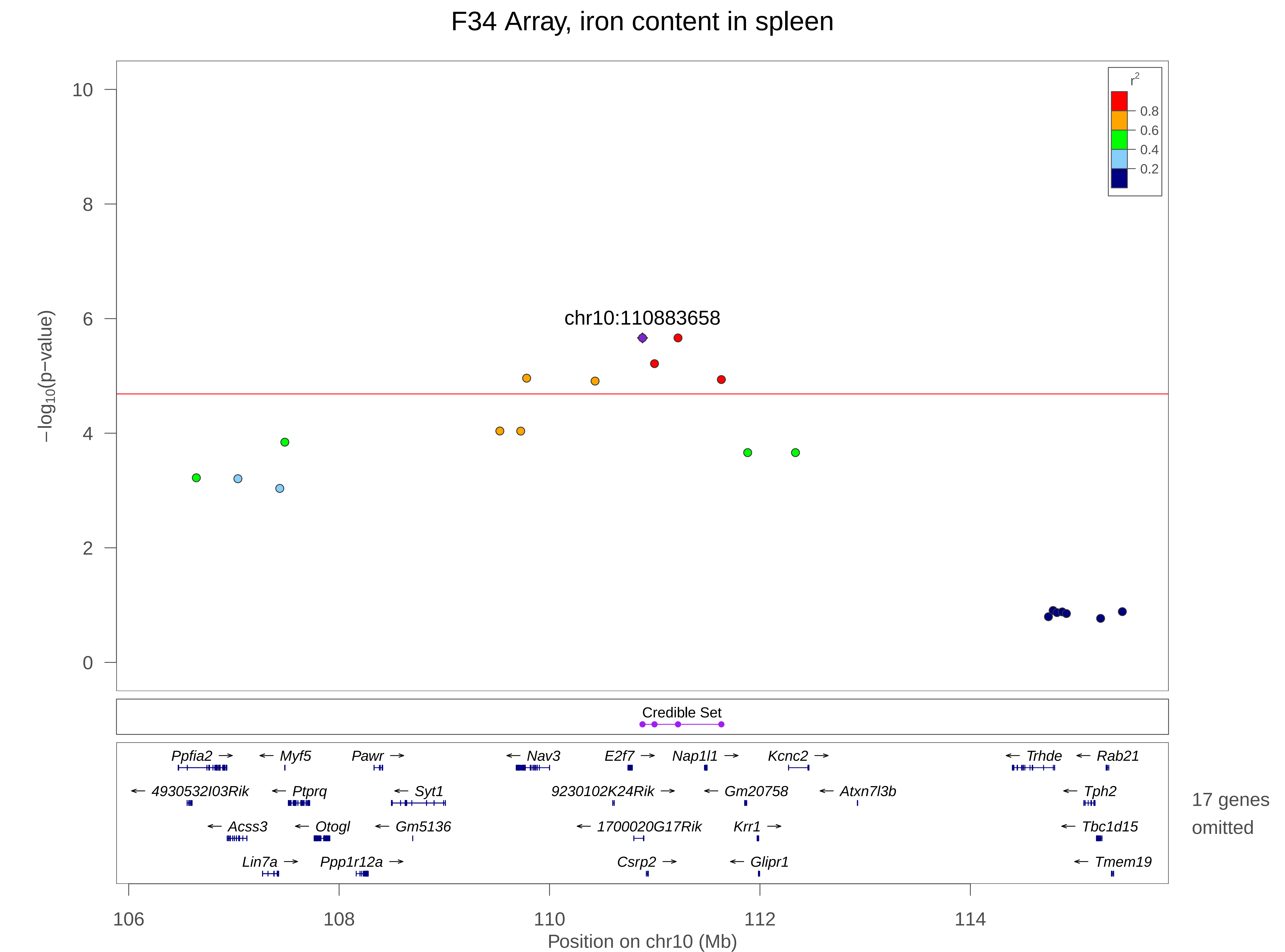

F34 Array, iron content in spleen
r2
0.8
0.6
0.4
0.2
−log10(p−value)
10
8
6
4
2
0
chr10:110883658
Credible Set
Ppfia2 →
← Myf5
Pawr →
← Nav3
E2f7 →
Nap1l1 →
Kcnc2 →
← Trhde
← Rab21
← 4930532I03Rik
← Ptprq
← Syt1
9230102K24Rik →
← Gm20758
← Atxn7l3b
← Tph2
← Acss3
← Otogl
← Gm5136
← 1700020G17Rik
Krr1 →
← Tbc1d15
Lin7a →
Ppp1r12a →
Csrp2 →
← Glipr1
← Tmem19
17 genes omitted
106
108
110
112
114
Position on chr10 (Mb)

| | |
|---|---|
| date: | Thu Oct 25 16:54:40 2018 |
| build: | F34Array |
| display range: | chr10:105883658−115883658 [105883658−115883658] |
| hilite range: | 0 − 0 [ 0 − 0 ] |
| reference SNP: | chr10:110883658 |
| number of SNPs plotted: | 22 |
| min P−value: | 2.17E−6 [chr10:110883658] |
| max P−value: | 1.7E−1 [chr10:115238015] |
| omitted Genes: | Myf6, Gm29685, Zdhhc17 |
| omitted Genes: | Osbpl8, Bbs10, Gm5176 |
| omitted Genes: | Phlda1, 4933440J02Rik, Glipr1l1 |
| omitted Genes: | Glipr1l2, Caps2, 1700010J16Rik |
| omitted Genes: | Thap2, Zfc3h1, Lgr5 |
| omitted Genes: | A930009A15Rik, Tspan8 |

Make more plots at http://csg.sph.umich.edu/locuszoom/

Fine−mapping Regions

| snp | chr | pos (Mb) | pp | group | color |
|---|---|---|---|---|---|
| chr10:110883658 | 10 | 110.8837 | 0.38877202 | Credible Set | purple |
| chr10:110997815 | 10 | 110.9978 | 0.14413964 | Credible Set | purple |
| chr10:111221007 | 10 | 111.2210 | 0.38877202 | Credible Set | purple |
| chr10:111633214 | 10 | 111.6332 | 0.07831632 | Credible Set | purple |

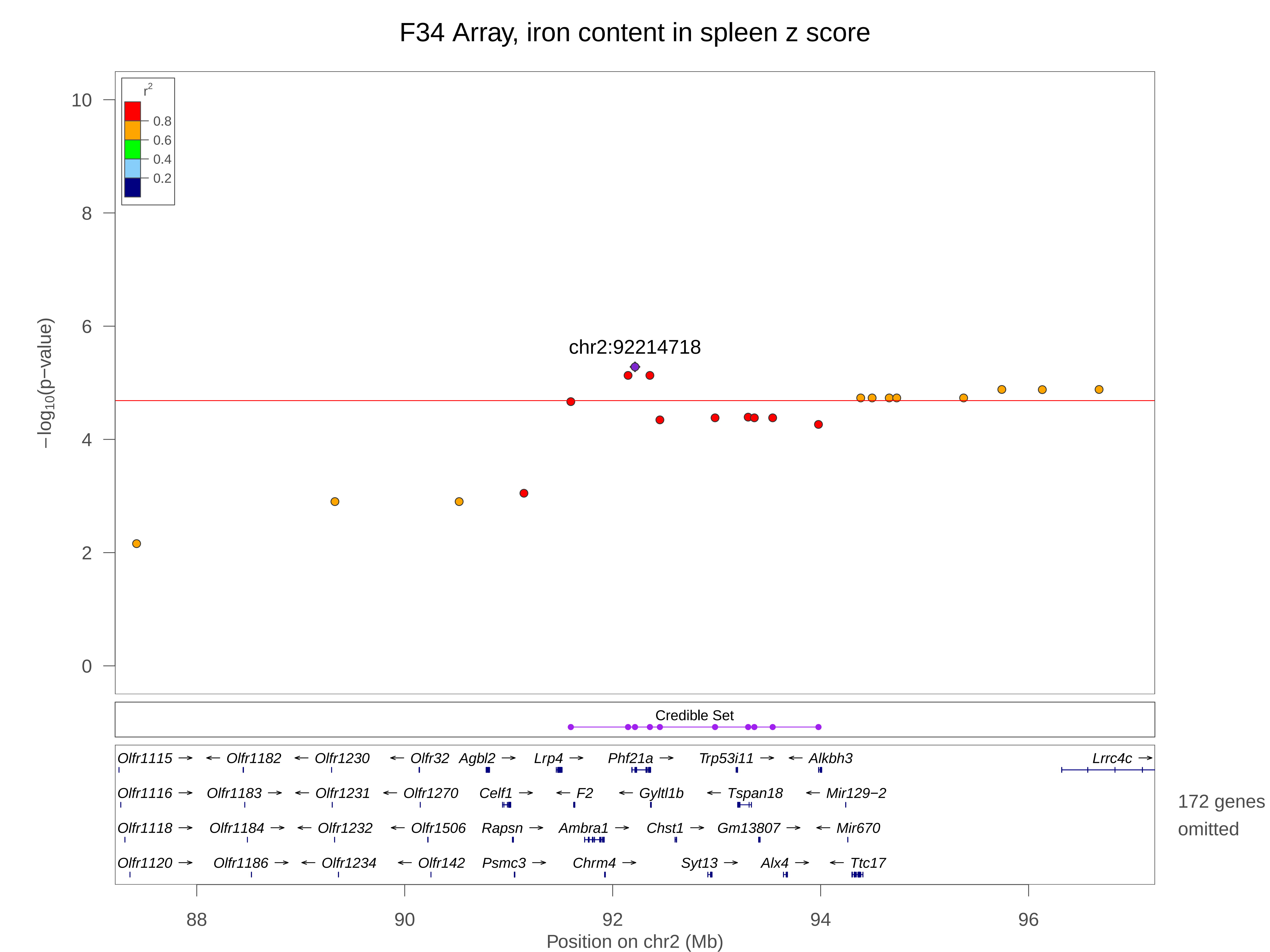

F34 Array, iron content in spleen z score
r2
0.8
0.6
0.4
0.2
10
8
6
4
2
0
−log10(p−value)
chr2:92214718
Credible Set
Olfr1115 → ← Olfr1182 ← Olfr1230 ← Olfr32 Agbl2 → Lrp4 → Phf21a → Trp53i11 → ← Alkbh3 Lrrc4c →
Olfr1116 → Olfr1183 → ← Olfr1231 ← Olfr1270 Celf1 → ← F2 ← Gyltl1b ← Tspan18 ← Mir129−2
Olfr1118 → Olfr1184 → ← Olfr1232 ← Olfr1506 Rapsn → Ambra1 → Chst1 → Gm13807 → ← Mir670
Olfr1120 → Olfr1186 → ← Olfr1234 ← Olfr142 Psmc3 → Chrm4 → Syt13 → Alx4 → ← Ttc17
172 genes omitted
88
90
92
94
96
Position on chr2 (Mb)

| | |
|---|---|
| date: | Thu Oct 25 16:55:26 2018 |
| build: | F34Array |
| display range: | chr2:87214718−97214718 [87214718−97214718] |
| hilite range: | 0 − 0 [ 0 − 0 ] |
| reference SNP: | chr2:92214718 |
| number of SNPs plotted: | 23 |
| min P−value: | 5.23E−6 [chr2:92214718] |
| max P−value: | 6.94E−3 [chr2:87421495] |
| omitted Genes: | Olfr1121, Olfr1122, Olfr1123 |
| omitted Genes: | Olfr1124, Olfr1126, Pramel7 |
| omitted Genes: | Pramel6, Olfr153, Olfr1128 |
| omitted Genes: | Olfr1129, Olfr1130, Olfr1131 |
| omitted Genes: | Olfr1132, Olfr1133, Olfr1134 |
| omitted Genes: | Olfr1135, Olfr1136, Olfr1137 |
| omitted Genes: | Olfr1138, Olfr1140, Olfr1141 |
| omitted Genes: | Olfr152, Olfr1143, Olfr1145 |
| omitted Genes: | Olfr1148, Olfr1151, Olfr1152 |
| omitted Genes: | Olfr1153, Olfr1154, Olfr1155 |
| omitted Genes: | Olfr1156, Olfr1157, Olfr74 |
| omitted Genes: | Olfr1158, Olfr1160, Olfr1161 |
| omitted Genes: | Olfr73, Olfr1162, Olfr1163 |
| omitted Genes: | Olfr1164, Olfr1166, Olfr1167 |
| omitted Genes: | Olfr1168, Olfr1170, Olfr1173 |
| omitted Genes: | Olfr1176, Olfr1178, Olfr1179 |
| omitted Genes: | Olfr1180, Olfr1181, Olfr1188 |
| omitted Genes: | Olfr1189, Olfr1193, Olfr1195 |
| omitted Genes: | Olfr1196, Olfr1197, Olfr1198 |
| omitted Genes: | Olfr1199, Olfr1200, Olfr1201 |
| omitted Genes: | Olfr1202, Olfr1205, Olfr1204 |
| omitted Genes: | Olfr1206, Olfr1208, Olfr1209 |
| omitted Genes: | Olfr1211, Olfr1212, Olfr1213 |
| omitted Genes: | Olfr1214, Olfr1215, Olfr1216 |
| omitted Genes: | Olfr1217, Olfr1218, Olfr1219 |
| omitted Genes: | Olfr1220, Olfr1221, Olfr1222 |
| omitted Genes: | Olfr1223, Olfr1225, Olfr1226 |
| omitted Genes: | Olfr1228, Olfr1229, Olfr1233 |
| omitted Genes: | Olfr1238, Olfr1239, Olfr1240 |
| omitted Genes: | Olfr1241, Olfr1242, Olfr1243 |
| omitted Genes: | Olfr1245, Olfr1246, Olfr1247 |
| omitted Genes: | Olfr1248, Olfr1249, Olfr1250 |

Make more plots at http://csg.sph.umich.edu/locuszoom/

Fine−mapping Regions

| snp | chr | pos (Mb) | pp | group | color |
|---|---|---|---|---|---|
| chr2:91597466 | 2 | 91.59747 | 0.07416765 | Credible Set | purple |
| chr2:92147975 | 2 | 92.14798 | 0.20518626 | Credible Set | purple |
| chr2:92214718 | 2 | 92.21472 | 0.28617800 | Credible Set | purple |
| chr2:92357966 | 2 | 92.35797 | 0.20518626 | Credible Set | purple |
| chr2:92454070 | 2 | 92.45407 | 0.03664601 | Credible Set | purple |
| chr2:92984633 | 2 | 92.98463 | 0.03965422 | Credible Set | purple |
| chr2:93303076 | 2 | 93.30308 | 0.04070338 | Credible Set | purple |
| chr2:93362468 | 2 | 93.36247 | 0.03965422 | Credible Set | purple |
| chr2:93538754 | 2 | 93.53875 | 0.03965422 | Credible Set | purple |
| chr2:93979130 | 2 | 93.97913 | 0.03074496 | Credible Set | purple |

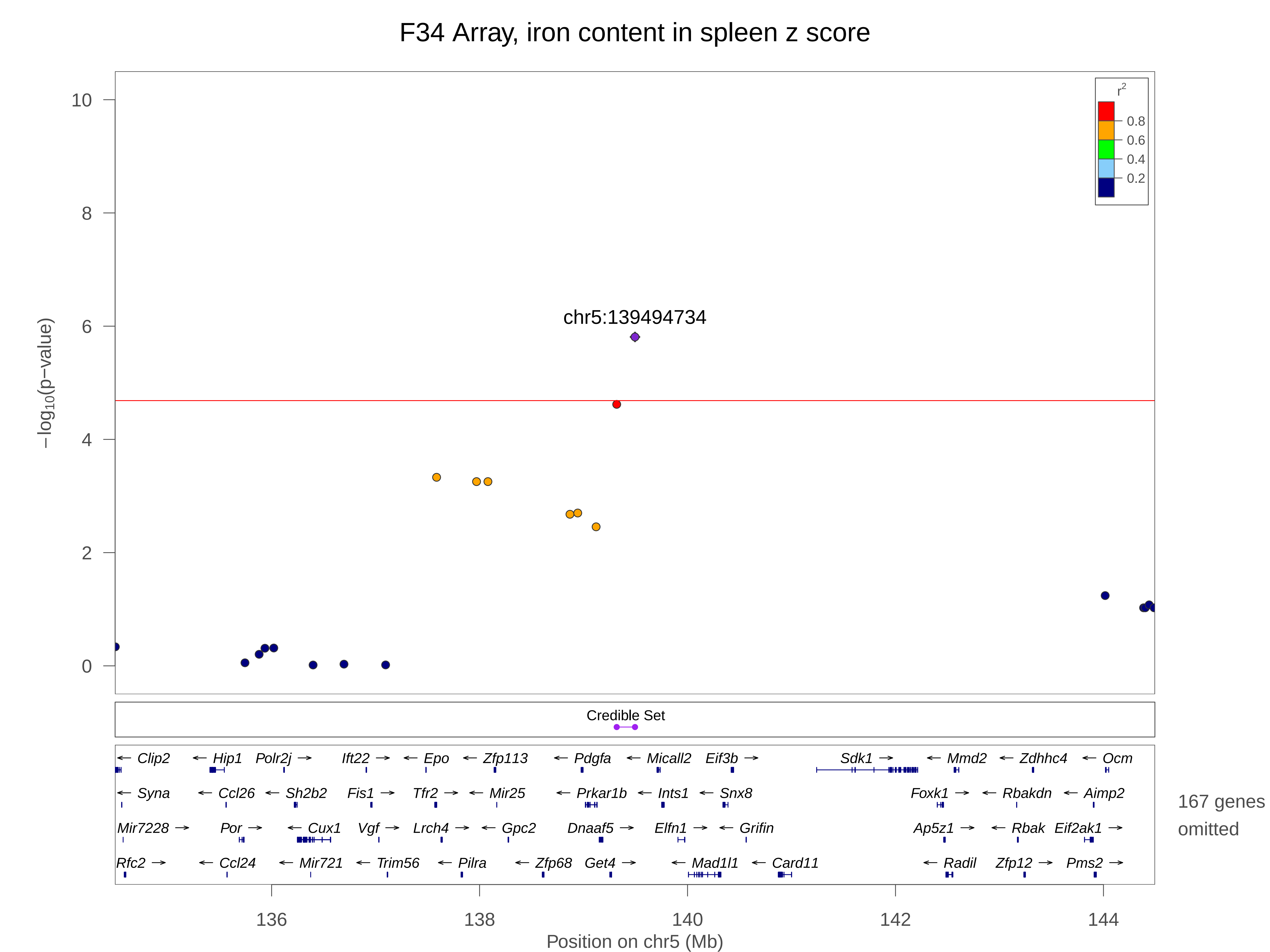

F34 Array, iron content in spleen z score
chr5:139494734
−log10(p−value)
r2
0.8
0.6
0.4
0.2
Credible Set
Clip2
Hip1
Polr2j
Ift22
Epo
Zfp113
Pdgfa
Micall2
Eif3b
Sdk1
Mmd2
Zdhhc4
Ocm
Syna
Ccl26
Sh2b2
Fis1
Tfr2
Mir25
Prkar1b
Ints1
Snx8
Foxk1
Rbakdn
Aimp2
Mir7228
Por
Cux1
Vgf
Lrch4
Gpc2
Dnaaf5
Elfn1
Grifin
Ap5z1
Rbak
Eif2ak1
Rfc2
Ccl24
Mir721
Trim56
Pilra
Zfp68
Get4
Mad1l1
Card11
Radil
Zfp12
Pms2
167 genes omitted
Position on chr5 (Mb)

date: Thu Oct 25 16:55:37 2018
build: F34Array
display range: chr5:134494734−144494734 [134494734−144494734]
hilite range: 0 − 0 [ 0 − 0 ]
reference SNP: chr5:139494734
number of SNPs plotted: 22
min P−value: 1.55E−6 [chr5:139494734]
max P−value: 9.65E−1 [chr5:136398923]
omitted Genes: Lat2, Eif4h, Limk1
omitted Genes: Eln, Wbscr28, Cldn13
omitted Genes: Wbscr27, Cldn4, Cldn3
omitted Genes: Wbscr25, Abhd11, Abhd11os
omitted Genes: Stx1a, Wbscr22, Dnajc30
omitted Genes: Vps37d, Mlxipl, Tbl2
omitted Genes: Bcl7b, Baz1b, Fzd9
omitted Genes: Fkbp6, Trim50, Nsun5
omitted Genes: Pom121, Mir7033, Rhbdd2
omitted Genes: Tmem120a, Mir7034, Styxl1
omitted Genes: Mdh2, Srrm3, Hspb1
omitted Genes: Ywhag, Ssc4d, Zp3
omitted Genes: Dtx2, Upk3b, Upk3bl
omitted Genes: Rasa4, Mir7035, Lrwd1
omitted Genes: Alkbh4, Orai2, Prkrip1
omitted Genes: Myl10, Col26a1, 4933404O12Rik
omitted Genes: Cldn15, Znhit1, Plod3
omitted Genes: Mir702, Ap1s1, Serpine1
omitted Genes: Ache, Mir8116, Ufsp1
omitted Genes: Srrt, Mir7036, Trip6
omitted Genes: Slc12a9, Ephb4, Zan
omitted Genes: Pop7, Gigyf1, Gnb2
omitted Genes: Mir6418, Actl6b, Mospd3
omitted Genes: Pcolce, Fbxo24, Gm20605
omitted Genes: Sap25, Irs3, Agfg2
omitted Genes: Nyap1, Tsc22d4, Ppp1r35
omitted Genes: Mepce, Zcwpw1, Pilrb1
omitted Genes: Pilrb2, Cyp3a13, Gjc3
omitted Genes: Azgp1, Smok3a, Smok3b
omitted Genes: Smok3c, Zkscan1, Zscan21
omitted Genes: Cops6, Mcm7, Mir93
Make more plots at http://csg.sph.umich.edu/locuszoom/
omitted Genes: Mir106b, Ap4m1, Taf6

Fine−mapping Regions

| snp | chr | pos (Mb) | pp | group | color |
|---|---|---|---|---|---|
| chr5:139318848 | 5 | 139.3188 | 0.06774966 | Credible Set | purple |
| chr5:139494734 | 5 | 139.4947 | 0.93225034 | Credible Set | purple |

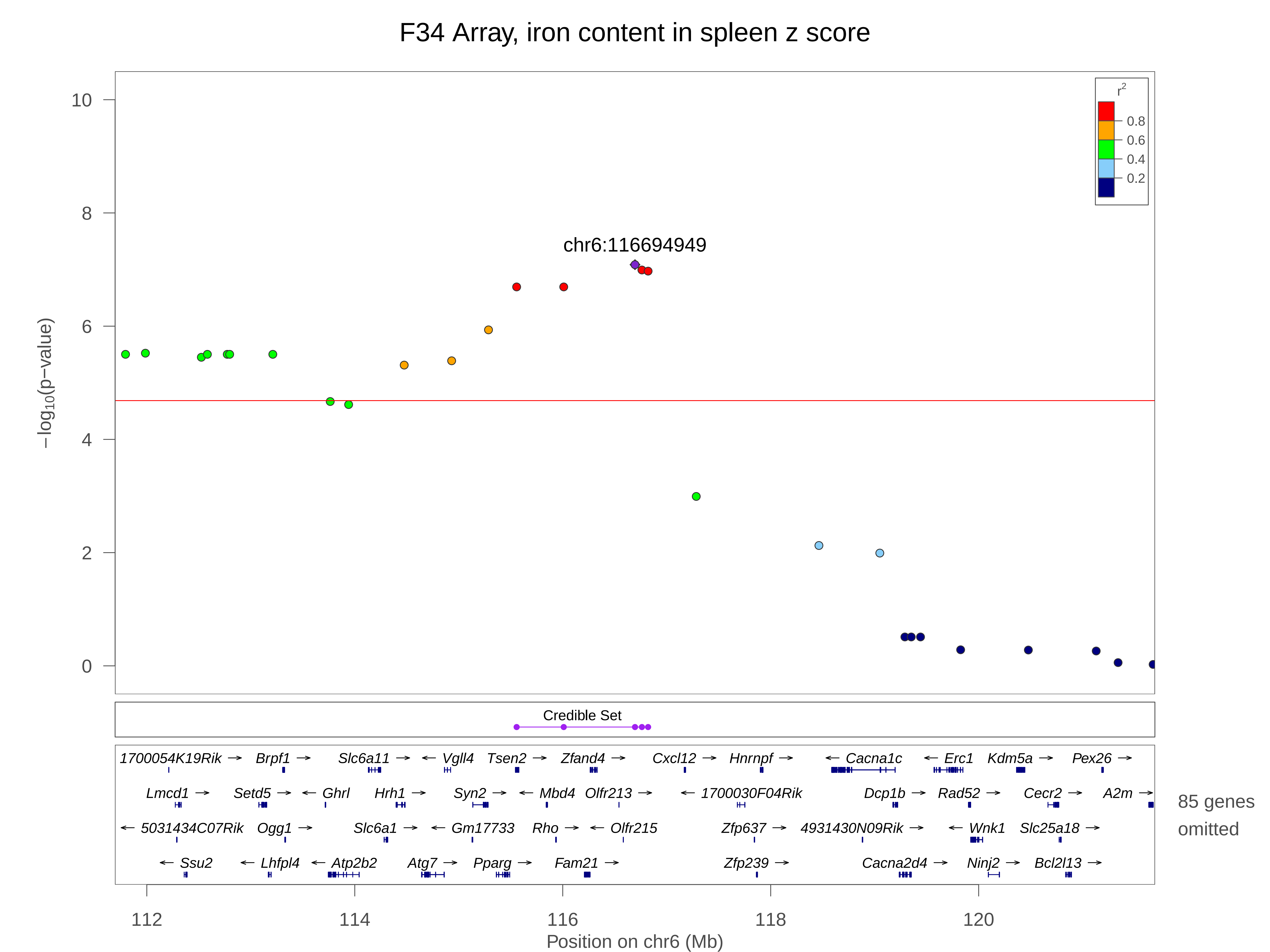

F34 Array, iron content in spleen z score
r2
0.8
0.6
0.4
0.2
chr6:116694949
−log10(p−value)
10
8
6
4
2
0
Credible Set
1700054K19Rik →
Brpf1 →
Slc6a11 →
← Vgll4
Tsen2 →
Zfand4 →
Cxcl12 →
Hnrnpf →
← Cacna1c
← Erc1
Kdm5a →
Pex26 →
Lmcd1 →
Setd5 →
← Ghrl
Hrh1 →
Syn2 →
← Mbd4
Olfr213 →
← 1700030F04Rik
Dcp1b →
Rad52 →
Cecr2 →
A2m →
← 5031434C07Rik
Ogg1 →
Slc6a1 →
← Gm17733
Rho →
← Olfr215
Zfp637 →
4931430N09Rik →
← Wnk1
Slc25a18 →
← Ssu2
← Lhfpl4
← Atp2b2
Atg7 →
Pparg →
Fam21 →
Zfp239 →
Cacna2d4 →
Ninj2 →
Bcl2l13 →
85 genes omitted
112
114
116
118
120
Position on chr6 (Mb)

| | |
|---|---|
| date: | Thu Oct 25 16:55:45 2018 |
| build: | F34Array |
| display range: | chr6:111694949−121694949 [111694949−121694949] |
| hilite range: | 0 − 0 [ 0 − 0 ] |
| reference SNP: | chr6:116694949 |
| number of SNPs plotted: | 29 |
| min P−value: | 8.2E−8 [chr6:116694949] |
| max P−value: | 9.43E−1 [chr6:121677793] |
| omitted Genes: | Cav3, Oxtr, Rad18 |
| omitted Genes: | Srgap3, Thumpd3, Gt(ROSA)26Sor |
| omitted Genes: | Mtmr14, Cpne9, Camk1 |
| omitted Genes: | Tada3, Arpc4, Ttll3 |
| omitted Genes: | Rpusd3, Cidec, Jagn1 |
| omitted Genes: | Il17re, Il17rc, Creld1 |
| omitted Genes: | Prrt3, Emc3, Fancd2 |
| omitted Genes: | Fancd2os, Brk1, Vhl |
| omitted Genes: | Irak2, Tatdn2, Mir7042 |
| omitted Genes: | Sec13, Tamm41, Timp4 |
| omitted Genes: | Mkrn2os, Mkrn2, Raf1 |
| omitted Genes: | Tmem40, Cand2, Mir7660 |
| omitted Genes: | Rpl32, Snora7a, Efcab12 |
| omitted Genes: | Ift122, H1foo, Plxnd1 |
| omitted Genes: | Tmcc1, 9530062K07Rik, March8 |
| omitted Genes: | Alox5, Olfr211, Olfr212 |
| omitted Genes: | Olfr214, Zfp422, Rassf4 |
| omitted Genes: | Mir7043, 8430408G22Rik, Tmem72 |
| omitted Genes: | Fxyd4, Rasgef1a, Mir7044 |
| omitted Genes: | Csgalnact2, Ret, 1700069P05Rik |
| omitted Genes: | Gm30498, Bms1, Zfp248 |
| omitted Genes: | Zfp9, Ankrd26, Lrtm2 |
| omitted Genes: | Adipor2, Wnt5b, Fbxl14 |
| omitted Genes: | 3110021A11Rik, Mir706, B4galnt3 |
| omitted Genes: | Ccdc77, Il17ra, Cecr6 |
| omitted Genes: | Cecr5, 1700072O05Rik, Atp6v1e1 |
| omitted Genes: | Bid, Mical3, Tuba8 |
| omitted Genes: | Usp18, Slc6a13, Slc6a12 |
| omitted Genes: | Iqsec3 |

Make more plots at http://csg.sph.umich.edu/locuszoom/

Fine−mapping Regions

| snp | chr | pos (Mb) | pp | group | color |
|---|---|---|---|---|---|
| chr6:115556641 | 6 | 115.5566 | 0.1218618 | Credible Set | purple |
| chr6:116009500 | 6 | 116.0095 | 0.1218618 | Credible Set | purple |
| chr6:116694949 | 6 | 116.6949 | 0.2923376 | Credible Set | purple |
| chr6:116761437 | 6 | 116.7614 | 0.2372379 | Credible Set | purple |
| chr6:116820969 | 6 | 116.8210 | 0.2267009 | Credible Set | purple |

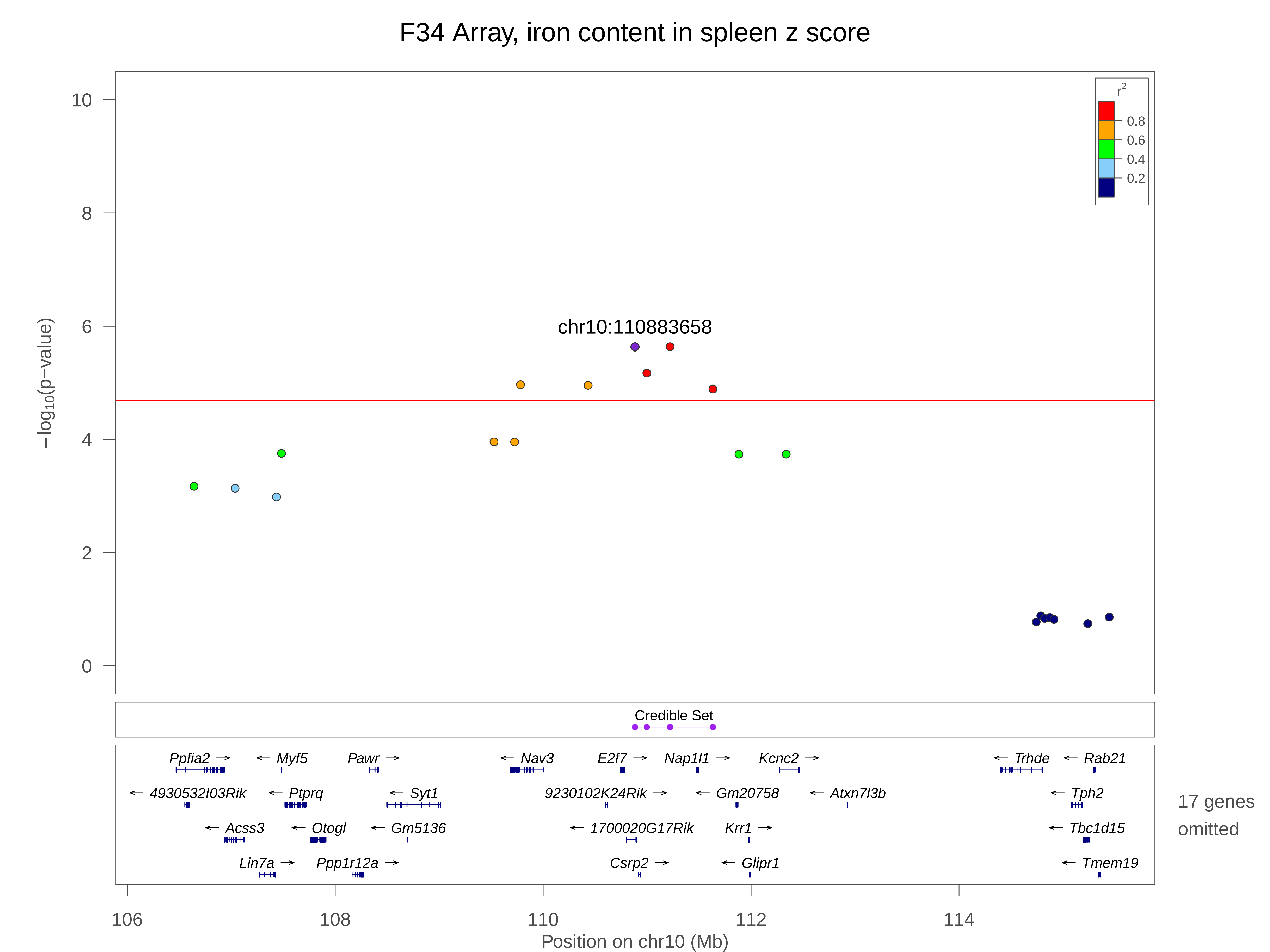

F34 Array, iron content in spleen z score
−log10(p−value)
10
8
6
4
2
0
r2
0.8
0.6
0.4
0.2
chr10:110883658
Credible Set
Ppfia2 →
← Myf5
Pawr →
← Nav3
E2f7 →
Nap1l1 →
Kcnc2 →
← Trhde
← Rab21
← 4930532I03Rik
← Ptprq
← Syt1
9230102K24Rik →
← Gm20758
← Atxn7l3b
← Tph2
← Acss3
← Otogl
← Gm5136
← 1700020G17Rik
Krr1 →
← Tbc1d15
Lin7a →
Ppp1r12a →
Csrp2 →
← Glipr1
← Tmem19
17 genes omitted
106
108
110
112
114
Position on chr10 (Mb)

| | |
|---|---|
| date: | Thu Oct 25 16:55:16 2018 |
| build: | F34Array |
| display range: | chr10:105883658−115883658 [105883658−115883658] |
| hilite range: | 0 − 0 [ 0 − 0 ] |
| reference SNP: | chr10:110883658 |
| number of SNPs plotted: | 22 |
| min P−value: | 2.31E−6 [chr10:110883658] |
| max P−value: | 1.8E−1 [chr10:115238015] |
| omitted Genes: | Myf6, Gm29685, Zdhhc17 |
| omitted Genes: | Osbpl8, Bbs10, Gm5176 |
| omitted Genes: | Phlda1, 4933440J02Rik, Glipr1l1 |
| omitted Genes: | Glipr1l2, Caps2, 1700010J16Rik |
| omitted Genes: | Thap2, Zfc3h1, Lgr5 |
| omitted Genes: | A930009A15Rik, Tspan8 |

Make more plots at http://csg.sph.umich.edu/locuszoom/

Fine−mapping Regions

| snp | chr | pos (Mb) | pp | group | color |
|---|---|---|---|---|---|
| chr10:110883658 | 10 | 110.8837 | 0.39202116 | Credible Set | purple |
| chr10:110997815 | 10 | 110.9978 | 0.14031854 | Credible Set | purple |
| chr10:111221007 | 10 | 111.2210 | 0.39202116 | Credible Set | purple |
| chr10:111633214 | 10 | 111.6332 | 0.07563914 | Credible Set | purple |

F34 Array, locomotor test day 1, total distance travelled in 30min (cm)

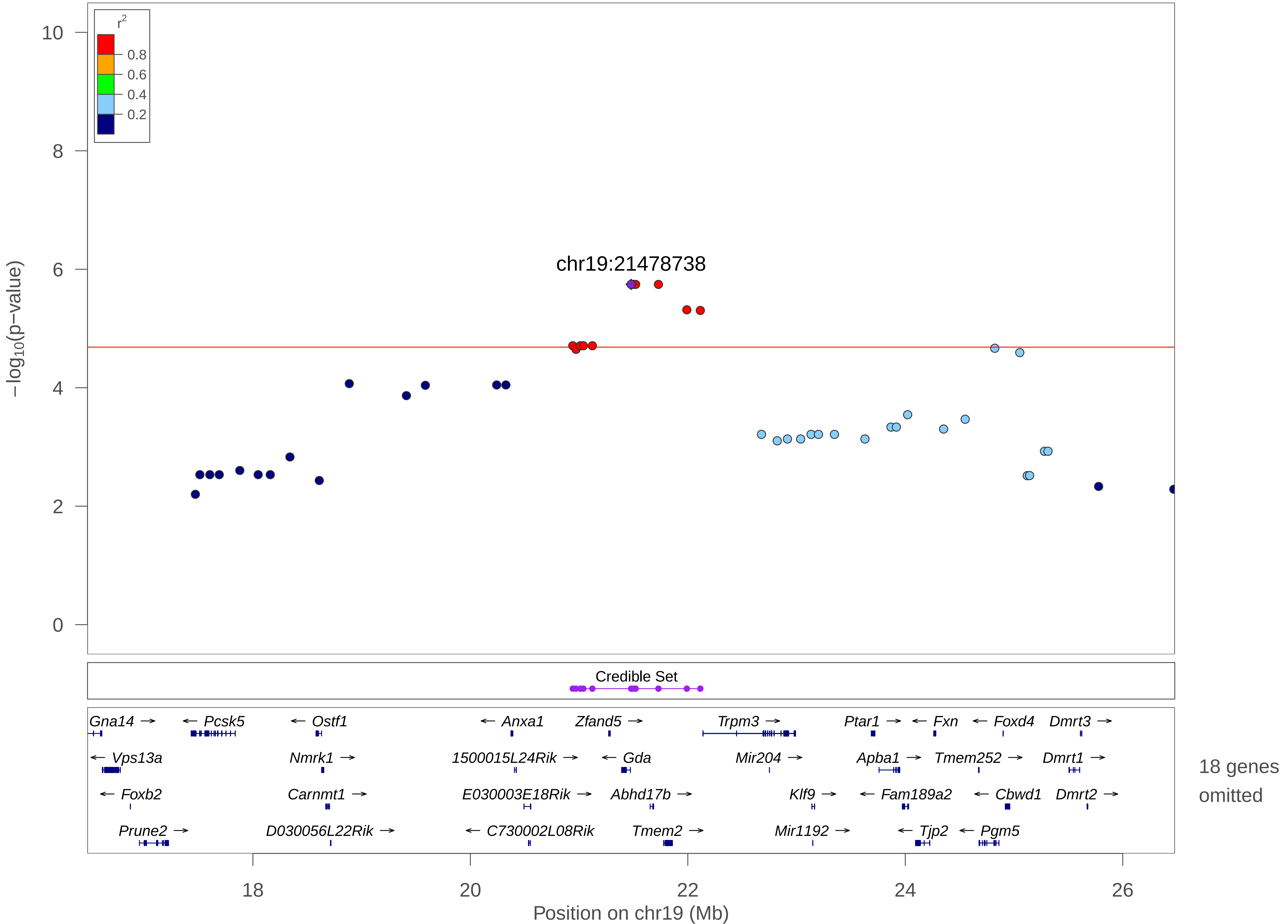

| | |
|---|---|
| date: | Thu Oct 25 16:55:52 2018 |
| build: | F34Array |
| display range: | chr19:16478738−26478738 [16478738−26478738] |
| hilite range: | 0 − 0 [ 0 − 0 ] |
| reference SNP: | chr19:21478738 |
| number of SNPs plotted: | 47 |
| min P−value: | 1.8E−6 [chr19:21478738] |
| max P−value: | 6.3E−3 [chr19:17470922] |
| omitted Genes: | Gcnt1, Rfk, Trpm6 |
| omitted Genes: | Rorb, Aldh1a1, Aldh1a7 |
| omitted Genes: | Tmc1, 4930554I06Rik, 1110059E24Rik |
| omitted Genes: | Smc5, Mamdc2, 1700028P14Rik |
| omitted Genes: | Pip5k1b, Fam122a, Mir3084−1 |
| omitted Genes: | Dock8, Kank1, 2610016A17Rik |

Make more plots at http://csg.sph.umich.edu/locuszoom/

Fine−mapping Regions

| snp | chr | pos (Mb) | pp | group | color |
|---|---|---|---|---|---|
| chr19:20941927 | 19 | 20.94193 | 0.01939458 | Credible Set | purple |
| chr19:20971640 | 19 | 20.97164 | 0.01696307 | Credible Set | purple |
| chr19:21011731 | 19 | 21.01173 | 0.01939458 | Credible Set | purple |
| chr19:21038918 | 19 | 21.03892 | 0.01939458 | Credible Set | purple |
| chr19:21122136 | 19 | 21.12214 | 0.01939458 | Credible Set | purple |
| chr19:21478738 | 19 | 21.47874 | 0.18998540 | Credible Set | purple |
| chr19:21500356 | 19 | 21.50036 | 0.18998540 | Credible Set | purple |
| chr19:21520189 | 19 | 21.52019 | 0.18998540 | Credible Set | purple |
| chr19:21729752 | 19 | 21.72975 | 0.18998540 | Credible Set | purple |
| chr19:21991002 | 19 | 21.99100 | 0.07362660 | Credible Set | purple |
| chr19:22115053 | 19 | 22.11505 | 0.07189041 | Credible Set | purple |

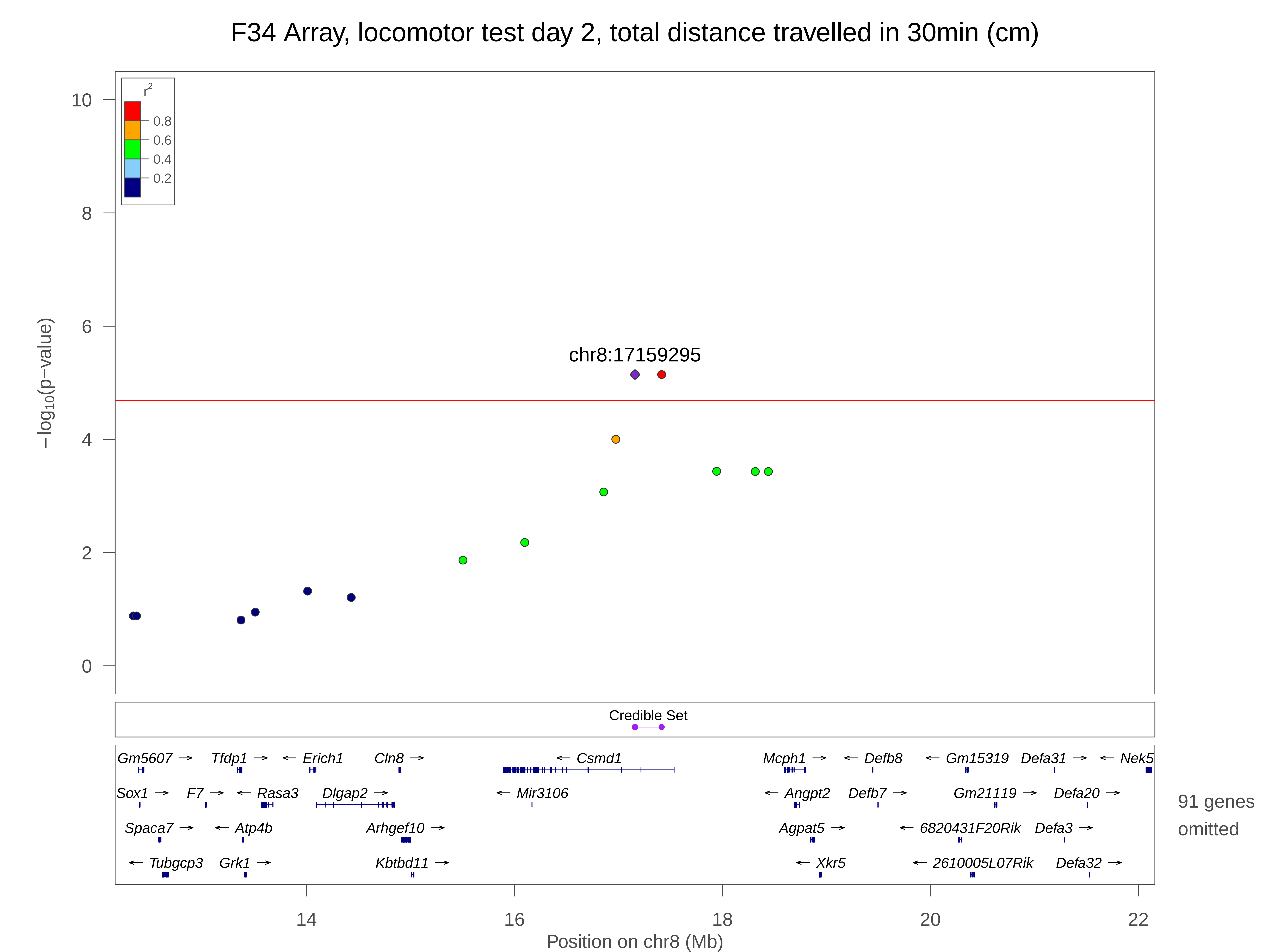
F34 Array, locomotor test day 2, total distance travelled in 30min (cm)
r2
0.8
0.6
0.4
0.2
10
8
6
4
2
0
−log10(p−value)
chr8:17159295
Credible Set
Gm5607 →
Tfdp1 →
← Erich1
Cln8 →
← Csmd1
Mcph1 →
← Defb8
← Gm15319
Defa31 →
← Nek5
Sox1 →
F7 →
← Rasa3
Dlgap2 →
← Mir3106
← Angpt2
Defb7 →
Gm21119 →
Defa20 →
Spaca7 →
← Atp4b
Arhgef10 →
Agpat5 →
← 6820431F20Rik
Defa3 →
← Tubgcp3
Grk1 →
Kbtbd11 →
← Xkr5
← 2610005L07Rik
Defa32 →
91 genes omitted
14
16
18
20
22
Position on chr8 (Mb)

| | |
|---|---|
| date: | Thu Oct 25 16:56:00 2018 |
| build: | F34Array |
| display range: | chr8:12159295−22159295 [12159295−22159295] |
| hilite range: | 0 − 0 [ 0 − 0 ] |
| reference SNP: | chr8:17159295 |
| number of SNPs plotted: | 16 |
| min P−value: | 7.15E−6 [chr8:17159295] |
| max P−value: | 1.55E−1 [chr8:13370321] |
| omitted Genes: | Gm15348, Atp11a, Gm15350 |
| omitted Genes: | Mcf2l, Gm38671, F10 |
| omitted Genes: | Proz, Pcid2, Cul4a |
| omitted Genes: | Mir7065, Lamp1, Grtp1 |
| omitted Genes: | Mir1968, Adprhl1, Dcun1d2 |
| omitted Genes: | Tmco3, Tmem255b, Gas6 |
| omitted Genes: | 1700029H14Rik, 4932443I19Rik, Cdc16 |
| omitted Genes: | Upf3a, AF366264, Champ1 |
| omitted Genes: | Coprs, Fbxo25, Tdrp |
| omitted Genes: | Gm5907, BB014433, Myom2 |
| omitted Genes: | Defb40, Defb37, Defb38 |
| omitted Genes: | Defb39, Defb12, Defb34 |
| omitted Genes: | Spag11b, Spag11a, Defb14 |
| omitted Genes: | Defb4, Defa−ps12, Defb6 |
| omitted Genes: | Defb46, Defb5, Defb3 |
| omitted Genes: | Defa−ps13, 4930467E23Rik, Gm21944 |
| omitted Genes: | Defb33, Gm15056, Gm6040 |
| omitted Genes: | AY761185, Defa21, Defa23 |
| omitted Genes: | Defa35, Defa25, Gm14851 |
| omitted Genes: | Defa30, Defa22, Gm15293 |
| omitted Genes: | Gm15292, Defa5, Defa27 |
| omitted Genes: | Defa29, Defa2, Defa33 |
| omitted Genes: | Defa36, Gm7849, Defa37 |
| omitted Genes: | Defa28, Gm7861, Defa26 |
| omitted Genes: | Defa17, Defa34, Defa−ps1 |
| omitted Genes: | AY761184, Defa24, Defb1 |
| omitted Genes: | Defb50, Defb2, Defb10 |
| omitted Genes: | Defb9, Defb11, Defb15 |
| omitted Genes: | Defb35, Defb13, Fam90a1a |
| omitted Genes: | Ccdc70, Atp7b, Alg11 |
| omitted Genes: | Nek3 |

Make more plots at http://csg.sph.umich.edu/locuszoom/

Fine−mapping Regions

| snp | chr | pos (Mb) | pp | group | color |
|---|---|---|---|---|---|
| chr8:17159295 | 8 | 17.15930 | 0.5 | Credible Set | purple |
| chr8:17415659 | 8 | 17.41566 | 0.5 | Credible Set | purple |

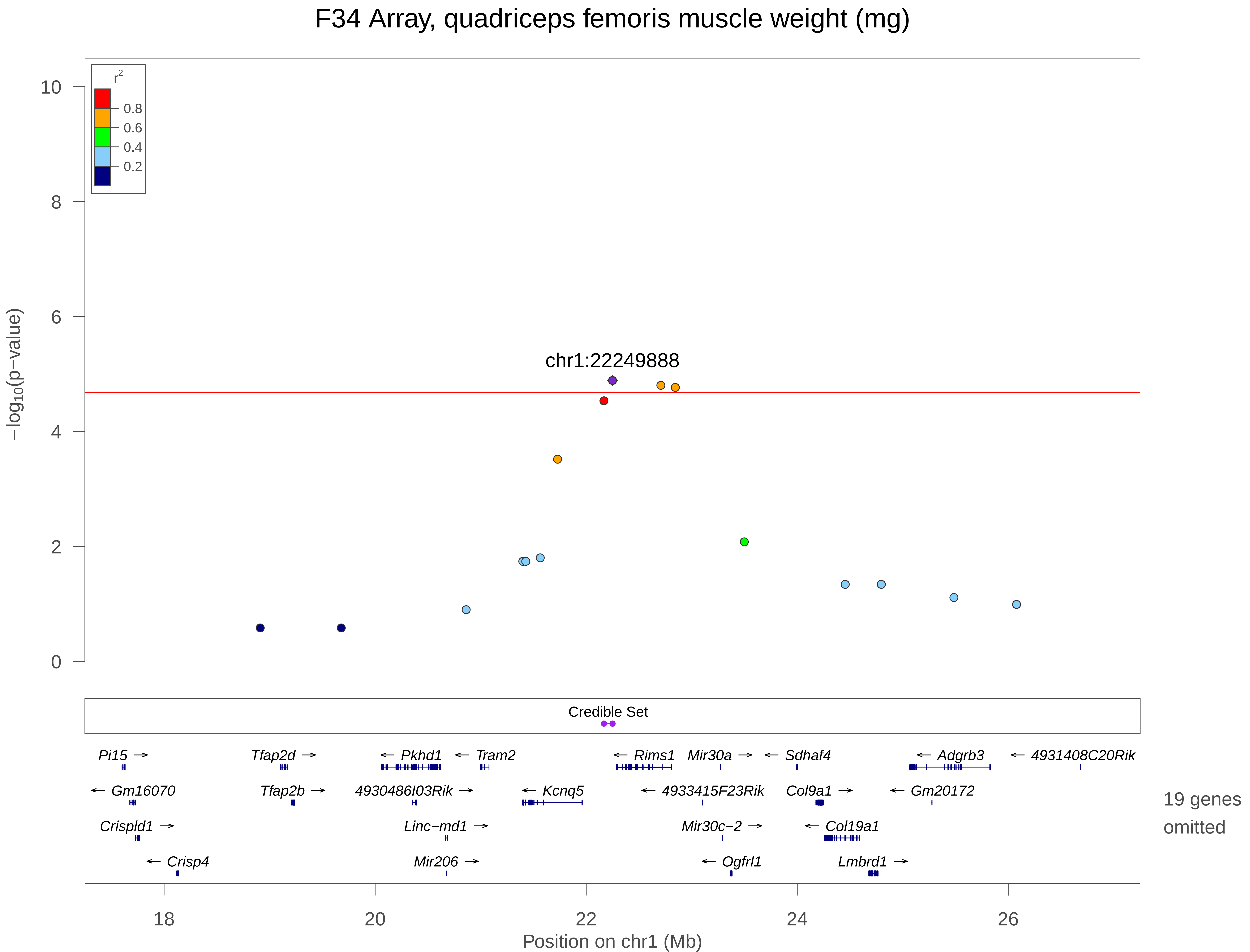
F34 Array, quadriceps femoris muscle weight (mg)
r2
0.8
0.6
0.4
0.2
10
8
6
4
2
0
−log10(p−value)
chr1:22249888
Credible Set
Pi15 →
← Gm16070
Crispld1 →
← Crisp4
Tfap2d →
Tfap2b →
← Pkhd1
4930486I03Rik →
Linc−md1 →
Mir206 →
← Tram2
← Kcnq5
← Rims1
← 4933415F23Rik
Mir30a →
Mir30c−2 →
← Ogfrl1
← Sdhaf4
Col9a1 →
← Col19a1
Lmbrd1 →
← Adgrb3
← Gm20172
← 4931408C20Rik
19 genes omitted
18
20
22
24
26
Position on chr1 (Mb)

| | |
|---|---|
| date: | Thu Oct 25 16:56:13 2018 |
| build: | F34Array |
| display range: | chr1:17249888−27249888 [17249888−27249888] |
| hilite range: | 0 − 0 [ 0 − 0 ] |
| reference SNP: | chr1:22249888 |
| number of SNPs plotted: | 17 |
| min P−value: | 1.29E−5 [chr1:22249888] |
| max P−value: | 2.6E−1 [chr1:18911051] |
| omitted Genes: | Defb44−ps, Defb18, Defb41 |
| omitted Genes: | Mir133b, Il17a, Il17f |
| omitted Genes: | Mcm3, 6720483E21Rik, Paqr8 |
| omitted Genes: | Efhc1, Tmem14a, Gsta3 |
| omitted Genes: | Gm4956, Khdc1a, Khdc1c |
| omitted Genes: | Khdc1b, B3gat2, Smap1 |
| omitted Genes: | Fam135a |

Make more plots at http://csg.sph.umich.edu/locuszoom/

Fine−mapping Regions

| snp | chr | pos (Mb) | pp | group | color |
|---|---|---|---|---|---|
| chr1:22168405 | 1 | 22.16840 | 0.3152276 | Credible Set | purple |
| chr1:22249888 | 1 | 22.24989 | 0.6847724 | Credible Set | purple |

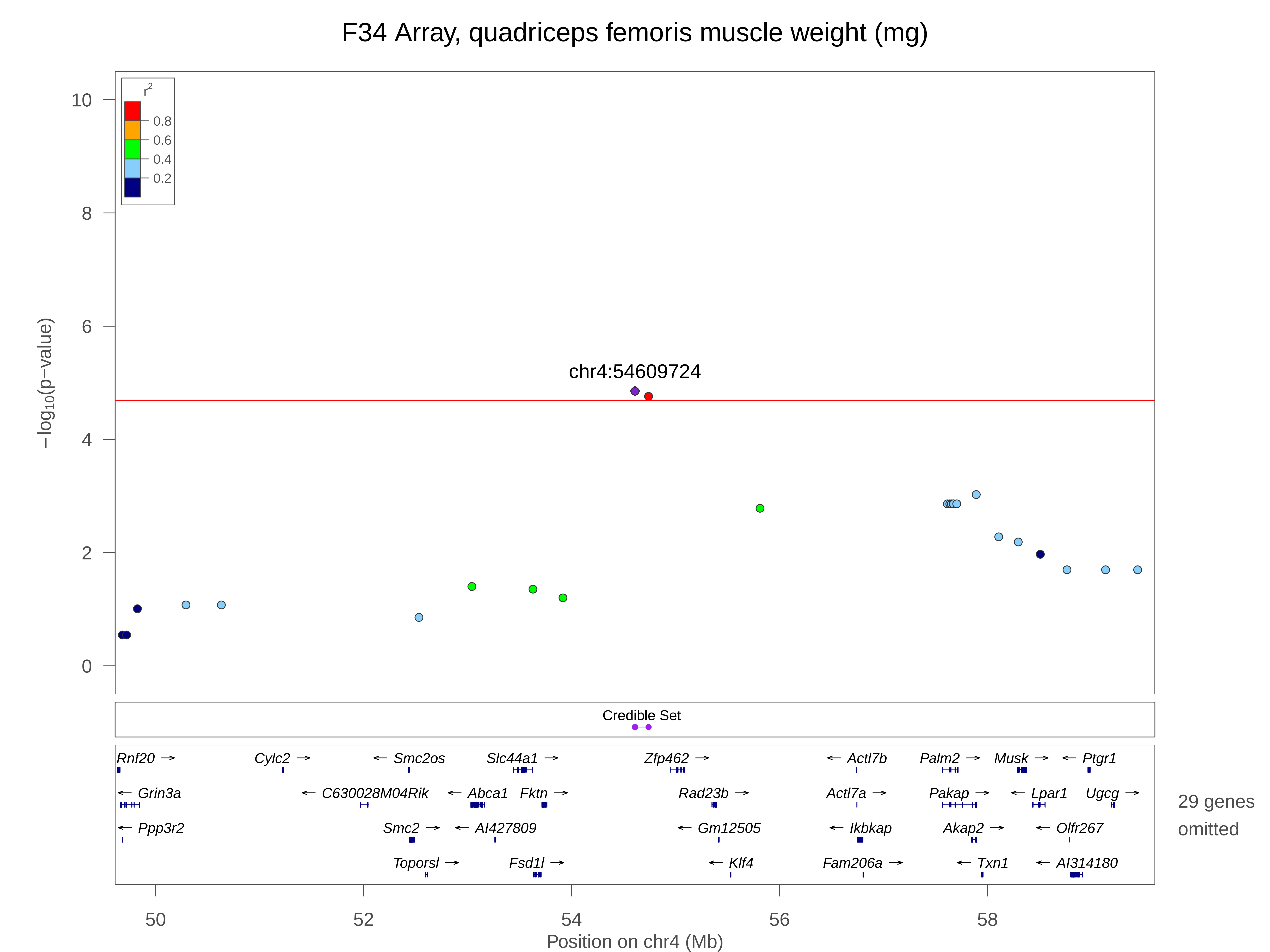
F34 Array, quadriceps femoris muscle weight (mg)
r2
0.8
0.6
0.4
0.2
10
8
6
4
2
0
−log10(p−value)
chr4:54609724
Credible Set
Rnf20 →
Cylc2 →
← Smc2os
Slc44a1 →
Zfp462 →
← Actl7b
Palm2 →
Musk →
← Ptgr1
← Grin3a
← C630028M04Rik
← Abca1
Fktn →
Rad23b →
Actl7a →
Pakap →
← Lpar1
Ugcg →
← Ppp3r2
Smc2 →
← AI427809
← Gm12505
← Ikbkap
Akap2 →
← Olfr267
Toporsl →
Fsd1l →
← Klf4
Fam206a →
← Txn1
← AI314180
29 genes omitted
50
52
54
56
58
Position on chr4 (Mb)

| | |
|---|---|
| date: | Thu Oct 25 16:56:26 2018 |
| build: | F34Array |
| display range: | chr4:49609724−59609724 [49609724−59609724] |
| hilite range: | 0 − 0 [ 0 − 0 ] |
| reference SNP: | chr4:54609724 |
| number of SNPs plotted: | 26 |
| min P−value: | 1.41E−5 [chr4:54609724] |
| max P−value: | 2.85E−1 [chr4:49679066] |
| omitted Genes: | 4930552N02Rik, Olfr275, Olfr273 |
| omitted Genes: | Olfr272, Olfr270, Nipsnap3a |
| omitted Genes: | Nipsnap3b, 4930522O17Rik, Tal2 |
| omitted Genes: | Tmem38b, Ctnnal1, Tmem245 |
| omitted Genes: | Mir32, Frrs1l, Epb41l4b |
| omitted Genes: | Gm12530, Ptpn3, 1700042G15Rik |
| omitted Genes: | D630039A03Rik, Txndc8, Svep1 |
| omitted Genes: | Mir3095, Zkscan16, Dnajc25 |
| omitted Genes: | Gng10, Gm12596, Susd1 |
| omitted Genes: | Ptbp3, Hsdl2 |

Make more plots at http://csg.sph.umich.edu/locuszoom/

Fine−mapping Regions

| snp | chr | pos (Mb) | pp | group | color |
|---|---|---|---|---|---|
| chr4:54609724 | 4 | 54.60972 | 0.5499053 | Credible Set | purple |
| chr4:54740078 | 4 | 54.74008 | 0.4500947 | Credible Set | purple |

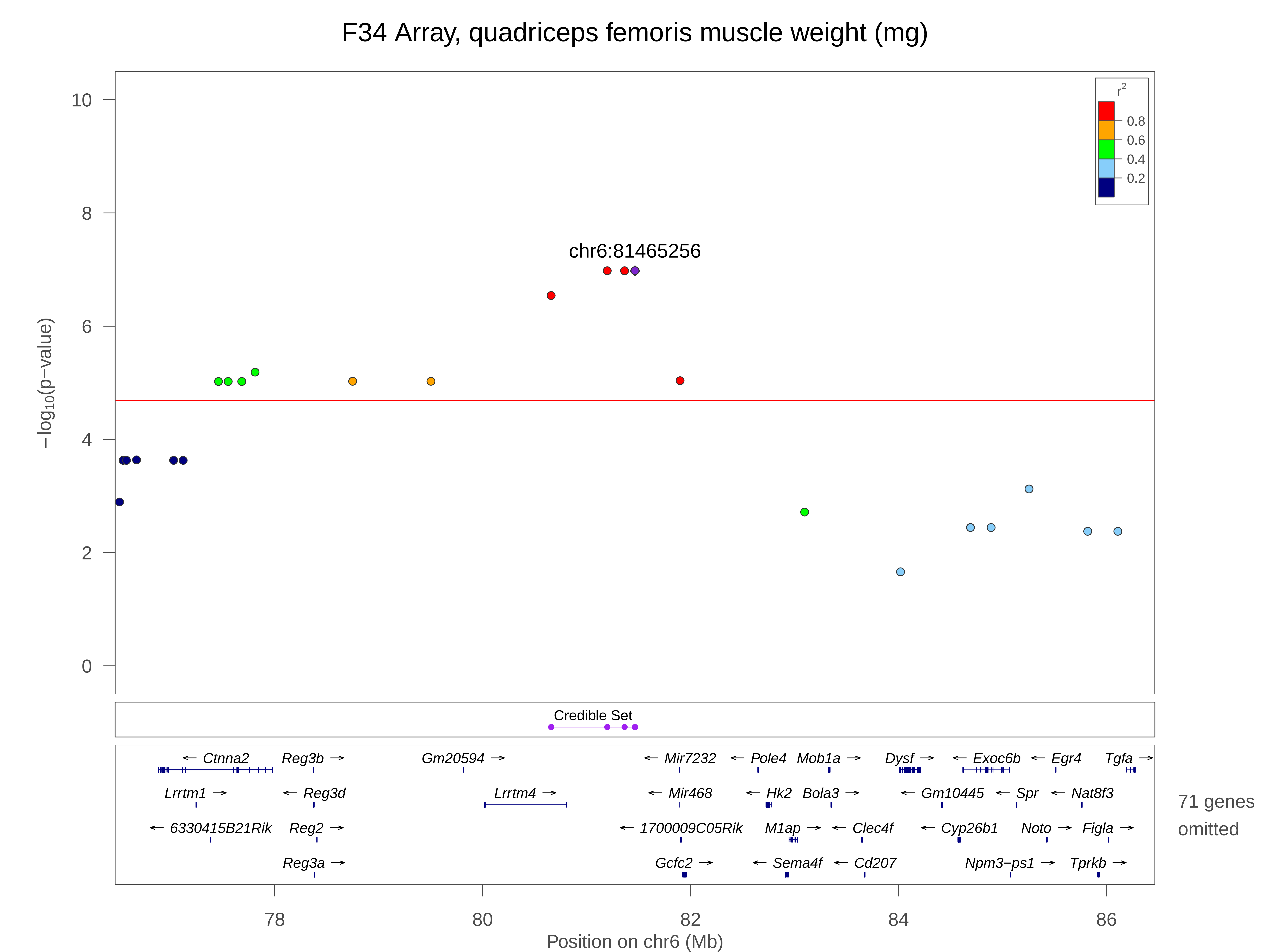

F34 Array, quadriceps femoris muscle weight (mg)
−log10(p−value)
10
8
6
4
2
0
r2
0.8
0.6
0.4
0.2
chr6:81465256
Credible Set
← Ctnna2
Reg3b →
Gm20594 →
← Mir7232
← Pole4
Mob1a →
Dysf →
← Exoc6b
← Egr4
Tgfa →
Lrrtm1 →
← Reg3d
Lrrtm4 →
← Mir468
← Hk2
Bola3 →
← Gm10445
← Spr
← Nat8f3
← 6330415B21Rik
Reg2 →
← 1700009C05Rik
M1ap →
← Clec4f
← Cyp26b1
Noto →
Figla →
Reg3a →
Gcfc2 →
← Sema4f
← Cd207
Npm3−ps1 →
Tprkb →
71 genes omitted
78
80
82
84
86
Position on chr6 (Mb)

| | |
|---|---|
| date: | Thu Oct 25 16:56:34 2018 |
| build: | F34Array |
| display range: | chr6:76465256−86465256 [76465256−86465256] |
| hilite range: | 0 − 0 [ 0 − 0 ] |
| reference SNP: | chr6:81465256 |
| number of SNPs plotted: | 25 |
| min P−value: | 1.05E−7 [chr6:81465256] |
| max P−value: | 2.18E−2 [chr6:84018933] |
| omitted Genes: | Reg1, Reg3g, Gm20362 |
| omitted Genes: | Mrpl19, Eva1a, Tacr1 |
| omitted Genes: | 2310069B03Rik, Dok1, Loxl3 |
| omitted Genes: | Mir7040, Htra2, Aup1 |
| omitted Genes: | Dqx1, Tlx2, Pcgf1 |
| omitted Genes: | Lbx2, Mir3470a, Ccdc142 |
| omitted Genes: | Mrpl53, Mogs, Wbp1 |
| omitted Genes: | Ino80b, Rtkn, Wdr54 |
| omitted Genes: | 1700003E16Rik, Dctn1, Slc4a5 |
| omitted Genes: | Mthfd2, Tet3, Mir6374 |
| omitted Genes: | B230319C09Rik, Dguok, Actg2 |
| omitted Genes: | Stambp, Gm21284, Vax2os |
| omitted Genes: | Vax2, Atp6v1b1, 1700124L16Rik |
| omitted Genes: | Ankrd53, Tex261, Gm7443 |
| omitted Genes: | Nagk, Paip2b, Zfp638 |
| omitted Genes: | Gm5878, Emx1, Sfxn5 |
| omitted Genes: | Rab11fip5, Mir705, Smyd5 |
| omitted Genes: | Pradc1, Cct7, Fbxo41 |
| omitted Genes: | Alms1, Nat8f7, Mir6375 |
| omitted Genes: | Alms1−ps2, Nat8f6, Nat8f5 |
| omitted Genes: | Nat8, Nat8f2, Nat8f4 |
| omitted Genes: | Nat8f1, Dusp11, Add2 |
| omitted Genes: | Fam136a, Snrpg, Pcyox1 |
| omitted Genes: | Tia1, C87436 |

Make more plots at http://csg.sph.umich.edu/locuszoom/

Fine−mapping Regions

| snp | chr | pos (Mb) | pp | group | color |
|---|---|---|---|---|---|
| chr6:80659015 | 6 | 80.65901 | 0.1112528 | Credible Set | purple |
| chr6:81198298 | 6 | 81.19830 | 0.2948104 | Credible Set | purple |
| chr6:81364957 | 6 | 81.36496 | 0.2948104 | Credible Set | purple |
| chr6:81465256 | 6 | 81.46526 | 0.2951438 | Credible Set | purple |

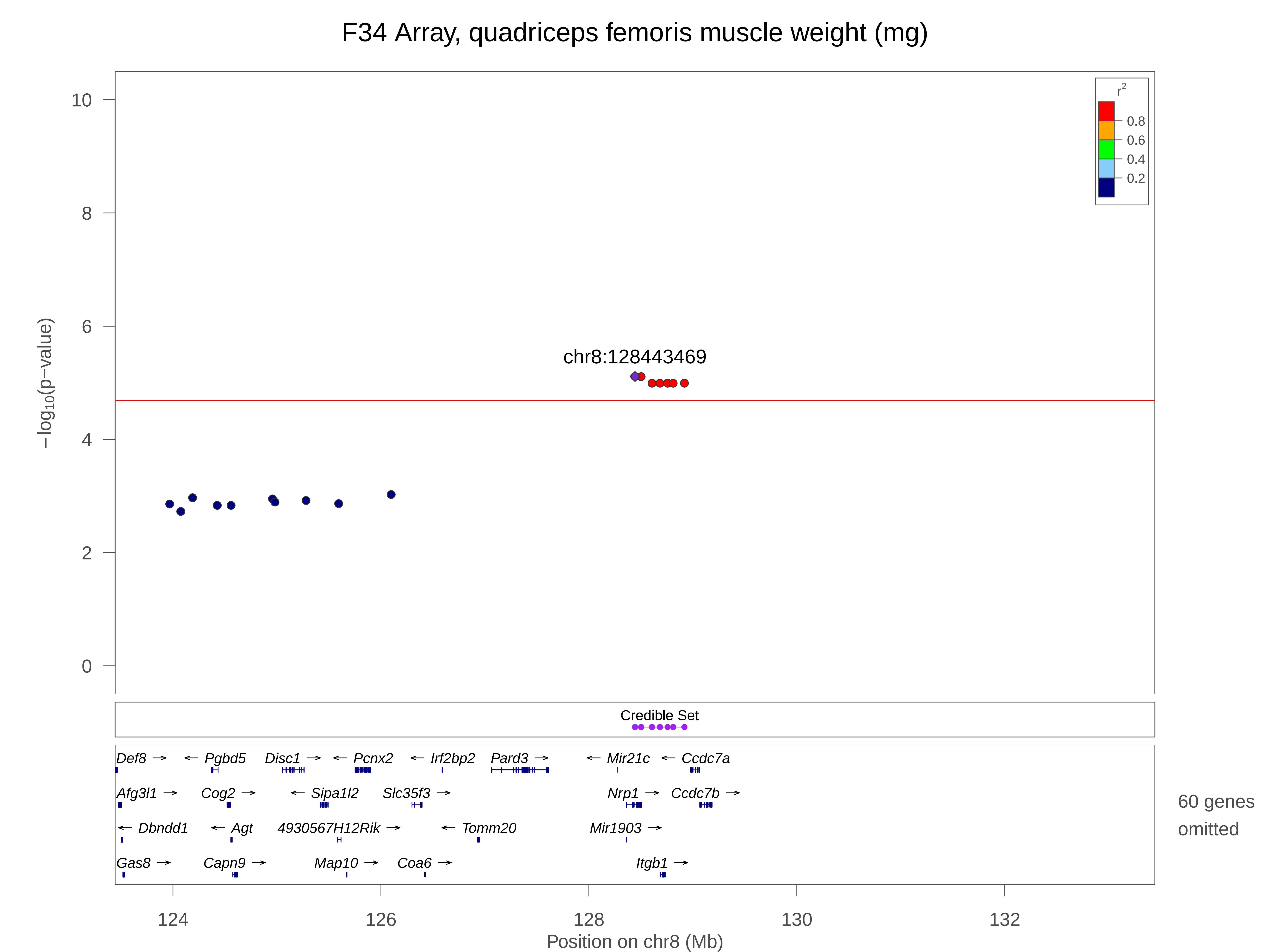

F34 Array, quadriceps femoris muscle weight (mg)
10
8
6
4
2
0
−log10(p−value)
r2
0.8
0.6
0.4
0.2
chr8:128443469
Credible Set
Def8 →
← Pgbd5
Disc1 →
← Pcnx2
← Irf2bp2
Pard3 →
← Mir21c
← Ccdc7a
Afg3l1 →
Cog2 →
← Sipa1l2
Slc35f3 →
Nrp1 →
Ccdc7b →
← Dbndd1
← Agt
4930567H12Rik →
← Tomm20
Mir1903 →
Gas8 →
Capn9 →
Map10 →
Coa6 →
Itgb1 →
60 genes omitted
124
126
128
130
132
Position on chr8 (Mb)

| | |
|---|---|
| date: | Thu Oct 25 16:56:41 2018 |
| build: | F34Array |
| display range: | chr8:123443469−133443469 [123443469−133443469] |
| hilite range: | 0 − 0 [ 0 − 0 ] |
| reference SNP: | chr8:128443469 |
| number of SNPs plotted: | 18 |
| min P−value: | 7.74E−6 [chr8:128443469] |
| max P−value: | 1.87E−3 [chr8:124074818] |
| omitted Genes: | n−R5s100, n−R5s103, n−R5s104 |
| omitted Genes: | n−R5s105, n−R5s108, n−R5s110 |
| omitted Genes: | n−R5s111, n−R5s113, Gm25018 |
| omitted Genes: | n−R5s117, n−R5s121, n−R5s122 |
| omitted Genes: | n−R5s123, n−R5s124, n−R5s128 |
| omitted Genes: | Gm26391, n−R5s133, n−R5s134 |
| omitted Genes: | Rn5s, n−R5s136, n−R5s138 |
| omitted Genes: | n−R5s139, Gm22109, n−R5s141 |
| omitted Genes: | n−R5s142, n−R5s143, n−R5s144 |
| omitted Genes: | Gm25212, n−R5s146, Gm22291 |
| omitted Genes: | Gm23284, n−R5s149, Rhou |
| omitted Genes: | 6030466F02Rik, Rab4a, Ccsap |
| omitted Genes: | Acta1, Nup133, Abcb10 |
| omitted Genes: | Taf5l, Mir1967, Urb2 |
| omitted Genes: | Galnt2, 2310022B05Rik, Ttc13 |
| omitted Genes: | Arv1, Fam89a, Trim67 |
| omitted Genes: | 2810004N23Rik, Gnpat, Exoc8 |
| omitted Genes: | Sprtn, Egln1, Tsnax |
| omitted Genes: | Ntpcr, BC021891, Kcnk1 |
| omitted Genes: | Gm17296, Rbm34, 2610044O15Rik8 |

Make more plots at http://csg.sph.umich.edu/locuszoom/

Fine−mapping Regions

| snp | chr | pos (Mb) | pp | group | color |
|---|---|---|---|---|---|
| chr8:128443469 | 8 | 128.4435 | 0.1709426 | Credible Set | purple |
| chr8:128502425 | 8 | 128.5024 | 0.1701707 | Credible Set | purple |
| chr8:128607127 | 8 | 128.6071 | 0.1317773 | Credible Set | purple |
| chr8:128683251 | 8 | 128.6833 | 0.1317773 | Credible Set | purple |
| chr8:128757217 | 8 | 128.7572 | 0.1317773 | Credible Set | purple |
| chr8:128810304 | 8 | 128.8103 | 0.1317773 | Credible Set | purple |
| chr8:128918665 | 8 | 128.9187 | 0.1317773 | Credible Set | purple |

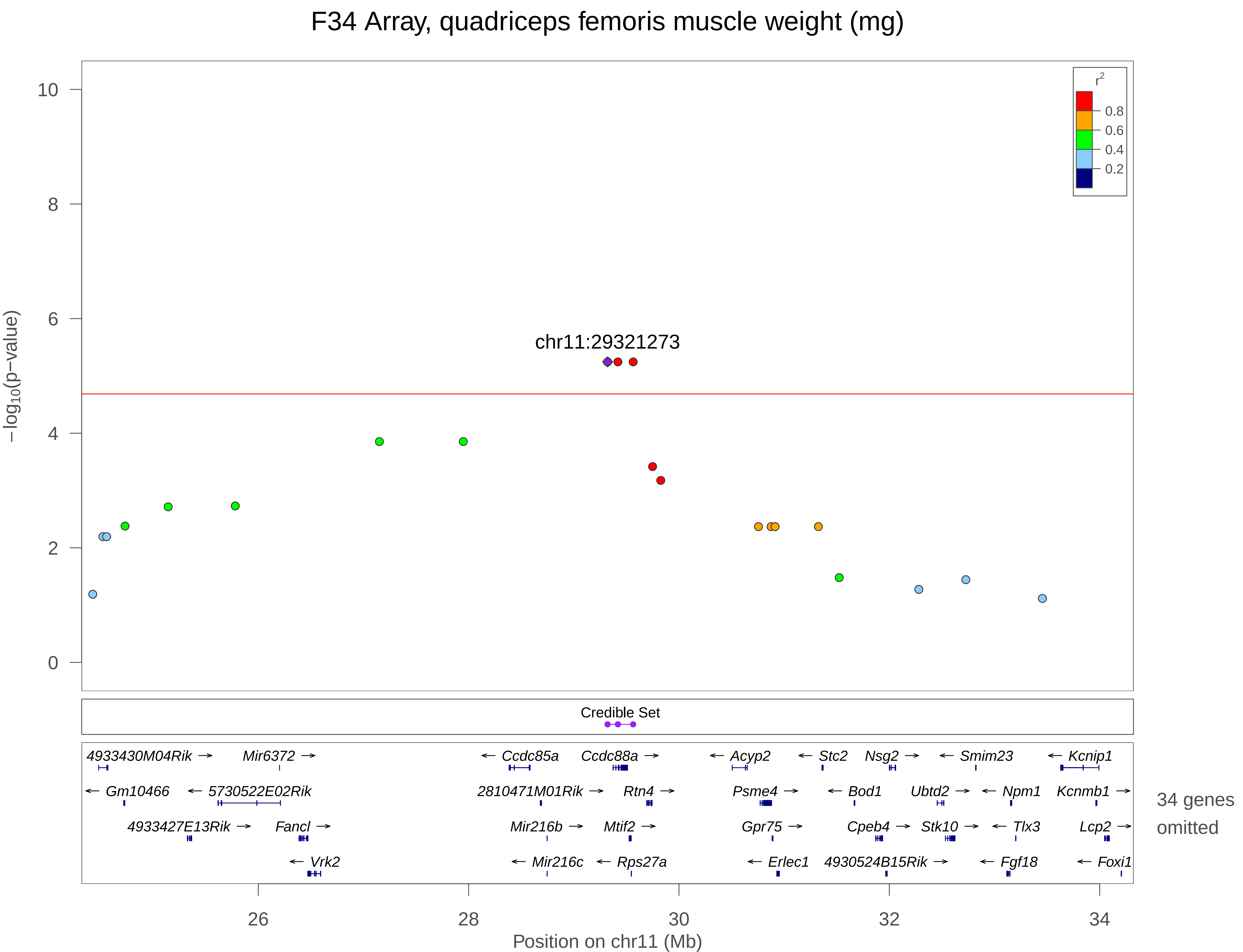
F34 Array, quadriceps femoris muscle weight (mg)
−log10(p−value)
chr11:29321273
r2
0.8
0.6
0.4
0.2
Credible Set
4933430M04Rik → Mir6372 → ← Ccdc85a Ccdc88a → ← Acyp2 ← Stc2 Nsg2 → ← Smim23 ← Kcnip1
← Gm10466 ← 5730522E02Rik 2810471M01Rik → Rtn4 → Psme4 → ← Bod1 Ubtd2 → ← Npm1 Kcnmb1 →
4933427E13Rik → Fancl → Mir216b → Mtif2 → Gpr75 → Cpeb4 → Stk10 → ← Tlx3 Lcp2 →
← Vrk2 ← Mir216c ← Rps27a ← Erlec1 4930524B15Rik → ← Fgf18 ← Foxi1
34 genes omitted
Position on chr11 (Mb)

| | |
|---|---|
| date: | Thu Oct 25 16:56:07 2018 |
| build: | F34Array |
| display range: | chr11:24321273−34321273 [24321273−34321273] |
| hilite range: | 0 − 0 [ 0 − 0 ] |
| reference SNP: | chr11:29321273 |
| number of SNPs plotted: | 22 |
| min P−value: | 5.69E−6 [chr11:29321273] |
| max P−value: | 7.66E−2 [chr11:33455345] |
| omitted Genes: | Gm12070, Mir216a, Mir217 |
| omitted Genes: | Efemp1, Pnpt1, Smek2 |
| omitted Genes: | Cfap36, Prorsd1, Clhc1 |
| omitted Genes: | Eml6, 4931440F15Rik, Sptbn1 |
| omitted Genes: | 4930505A04Rik, Asb3, Chac2 |
| omitted Genes: | D630024D03Rik, Gm12108, Il9r |
| omitted Genes: | Snrnp25, Rhbdf1, Mpg |
| omitted Genes: | Nprl3, Hba−x, Hba−a1 |
| omitted Genes: | Hbq1b, Hba−a2, Hbq1a |
| omitted Genes: | Sh3pxd2b, Efcab9, Fbxw11 |
| omitted Genes: | Ranbp17, Gabrp, Dock2 |
| omitted Genes: | Fam196b |

Make more plots at http://csg.sph.umich.edu/locuszoom/

Fine−mapping Regions

| snp | chr | pos (Mb) | pp | group | color |
|---|---|---|---|---|---|
| chr11:29321273 | 11 | 29.32127 | 0.3300716 | Credible Set | purple |
| chr11:29419287 | 11 | 29.41929 | 0.3300716 | Credible Set | purple |
| chr11:29564549 | 11 | 29.56455 | 0.3300716 | Credible Set | purple |

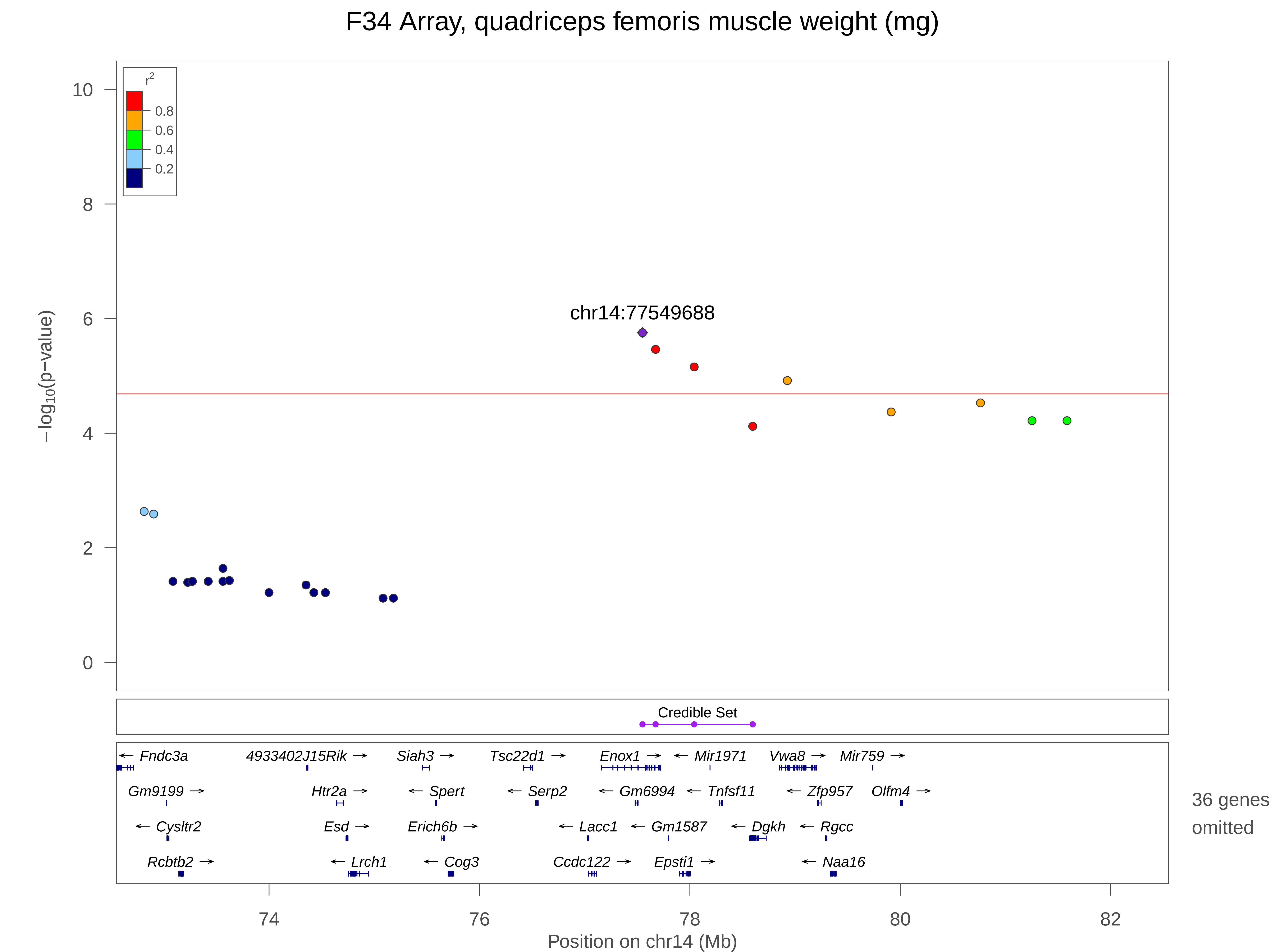
F34 Array, quadriceps femoris muscle weight (mg)
r²
0.8
0.6
0.4
0.2
chr14:77549688
−log10(p−value)
Credible Set
← Fndc3a
4933402J15Rik →
Siah3 →
Tsc22d1 →
Enox1 →
← Mir1971
Vwa8 →
Mir759 →
Gm9199 →
Htr2a →
← Spert
← Serp2
← Gm6994
← Tnfsf11
← Zfp957
Olfm4 →
← Cysltr2
Esd →
Erich6b →
← Lacc1
← Gm1587
← Dgkh
← Rgcc
Rcbtb2 →
← Lrch1
← Cog3
Ccdc122 →
Epsti1 →
← Naa16
36 genes omitted
Position on chr14 (Mb)

| | |
|---|---|
| date: | Thu Oct 25 16:56:20 2018 |
| build: | F34Array |
| display range: | chr14:72549688−82549688 [72549688−82549688] |
| hilite range: | 0 − 0 [ 0 − 0 ] |
| reference SNP: | chr14:77549688 |
| number of SNPs plotted: | 25 |
| min P−value: | 1.76E−6 [chr14:77549688] |
| max P−value: | 7.57E−2 [chr14:75083773] |
| omitted Genes: | Rb1, Mir687, Lpar6 |
| omitted Genes: | Itm2b, Med4, Nudt15 |
| omitted Genes: | Sucla2, Gm4278, 5031414D18Rik |
| omitted Genes: | Lrrc63, Lcp1, Cpb2 |
| omitted Genes: | Zc3h13, Slc25a30, Gm4285 |
| omitted Genes: | Tpt1, Snora31, Gtf2f2 |
| omitted Genes: | Kctd4, Gpalpp1, Nufip1 |
| omitted Genes: | 4930444M15Rik, 1700108F19Rik, Dnajc15 |
| omitted Genes: | Fam216b, Akap11, Mtrf1 |
| omitted Genes: | Kbtbd7, Kbtbd6, Wbp4 |
| omitted Genes: | Elf1, Sugt1, Lect1 |
| omitted Genes: | 4930452G13Rik, Pcdh8, Gm10845 |

Make more plots at http://csg.sph.umich.edu/locuszoom/

Fine−mapping Regions

| snp | chr | pos (Mb) | pp | group | color |
|---|---|---|---|---|---|
| chr14:77549688 | 14 | 77.54969 | 0.54938496 | Credible Set | purple |
| chr14:77673987 | 14 | 77.67399 | 0.28854246 | Credible Set | purple |
| chr14:78040815 | 14 | 78.04081 | 0.14690307 | Credible Set | purple |
| chr14:78597230 | 14 | 78.59723 | 0.01516952 | Credible Set | purple |

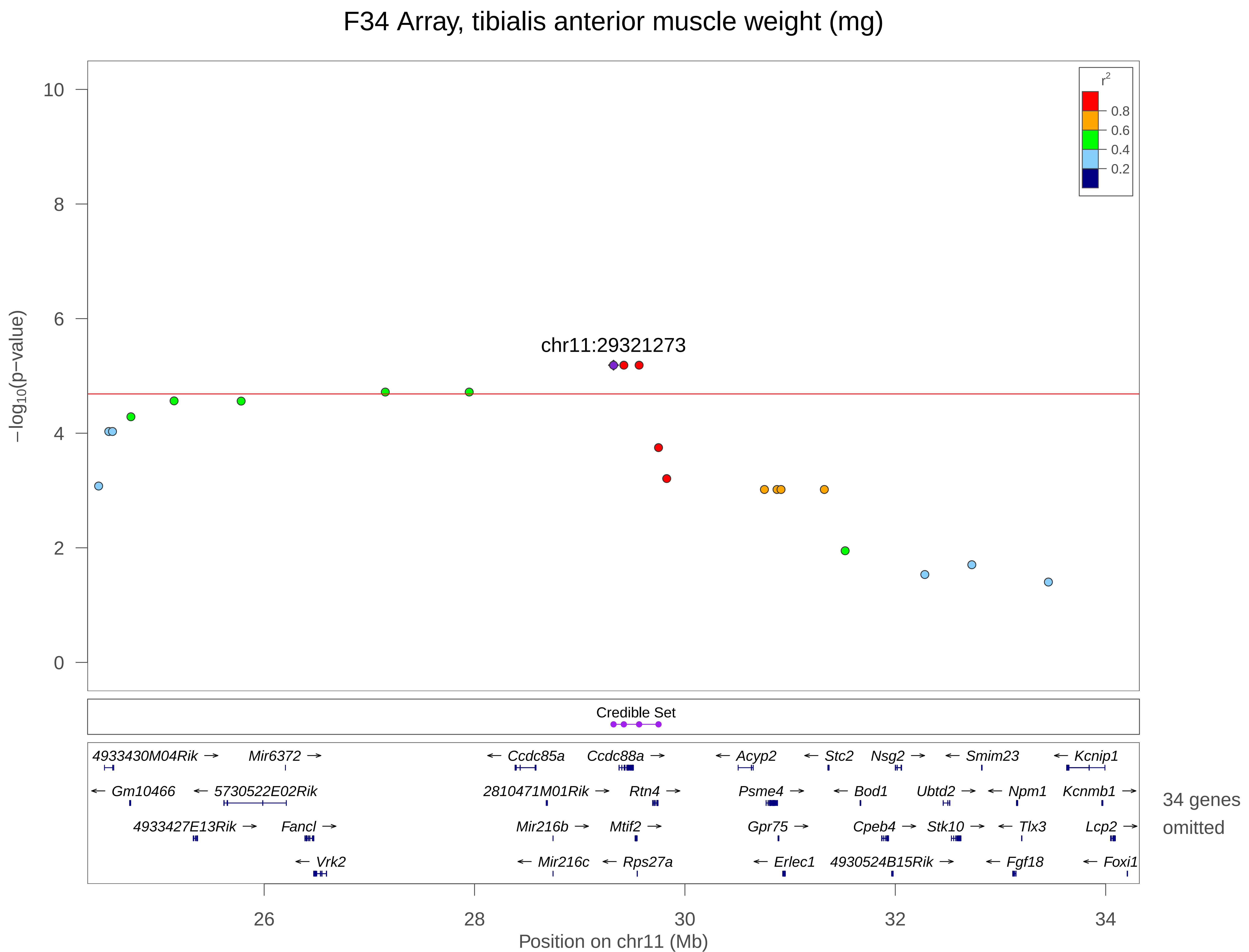

F34 Array, tibialis anterior muscle weight (mg)
−log10(p−value)
10
8
6
4
2
0
chr11:29321273
r2
0.8
0.6
0.4
0.2
Credible Set
4933430M04Rik →
Mir6372 →
← Ccdc85a
Ccdc88a →
← Acyp2
← Stc2
Nsg2 →
← Smim23
← Kcnip1
← Gm10466
← 5730522E02Rik
2810471M01Rik →
Rtn4 →
Psme4 →
← Bod1
Ubtd2 →
← Npm1
Kcnmb1 →
4933427E13Rik →
Fancl →
Mir216b →
Mtif2 →
Gpr75 →
Cpeb4 →
Stk10 →
← Tlx3
Lcp2 →
← Vrk2
← Mir216c
← Rps27a
← Erlec1
4930524B15Rik →
← Fgf18
← Foxi1
34 genes omitted
26
28
30
32
34
Position on chr11 (Mb)

| | |
|---|---|
| date: | Thu Oct 25 16:56:49 2018 |
| build: | F34Array |
| display range: | chr11:24321273−34321273 [24321273−34321273] |
| hilite range: | 0 − 0 [ 0 − 0 ] |
| reference SNP: | chr11:29321273 |
| number of SNPs plotted: | 22 |
| min P−value: | 6.49E−6 [chr11:29321273] |
| max P−value: | 3.95E−2 [chr11:33455345] |
| omitted Genes: | Gm12070, Mir216a, Mir217 |
| omitted Genes: | Efemp1, Pnpt1, Smek2 |
| omitted Genes: | Cfap36, Prorsd1, Clhc1 |
| omitted Genes: | Eml6, 4931440F15Rik, Sptbn1 |
| omitted Genes: | 4930505A04Rik, Asb3, Chac2 |
| omitted Genes: | D630024D03Rik, Gm12108, Il9r |
| omitted Genes: | Snrnp25, Rhbdf1, Mpg |
| omitted Genes: | Nprl3, Hba−x, Hba−a1 |
| omitted Genes: | Hbq1b, Hba−a2, Hbq1a |
| omitted Genes: | Sh3pxd2b, Efcab9, Fbxw11 |
| omitted Genes: | Ranbp17, Gabrp, Dock2 |
| omitted Genes: | Fam196b |

Make more plots at http://csg.sph.umich.edu/locuszoom/

Fine−mapping Regions

| snp | chr | pos (Mb) | pp | group | color |
|---|---|---|---|---|---|
| chr11:29321273 | 11 | 29.32127 | 0.32717292 | Credible Set | purple |
| chr11:29419287 | 11 | 29.41929 | 0.32717292 | Credible Set | purple |
| chr11:29564549 | 11 | 29.56455 | 0.32717292 | Credible Set | purple |
| chr11:29749029 | 11 | 29.74903 | 0.01408268 | Credible Set | purple |

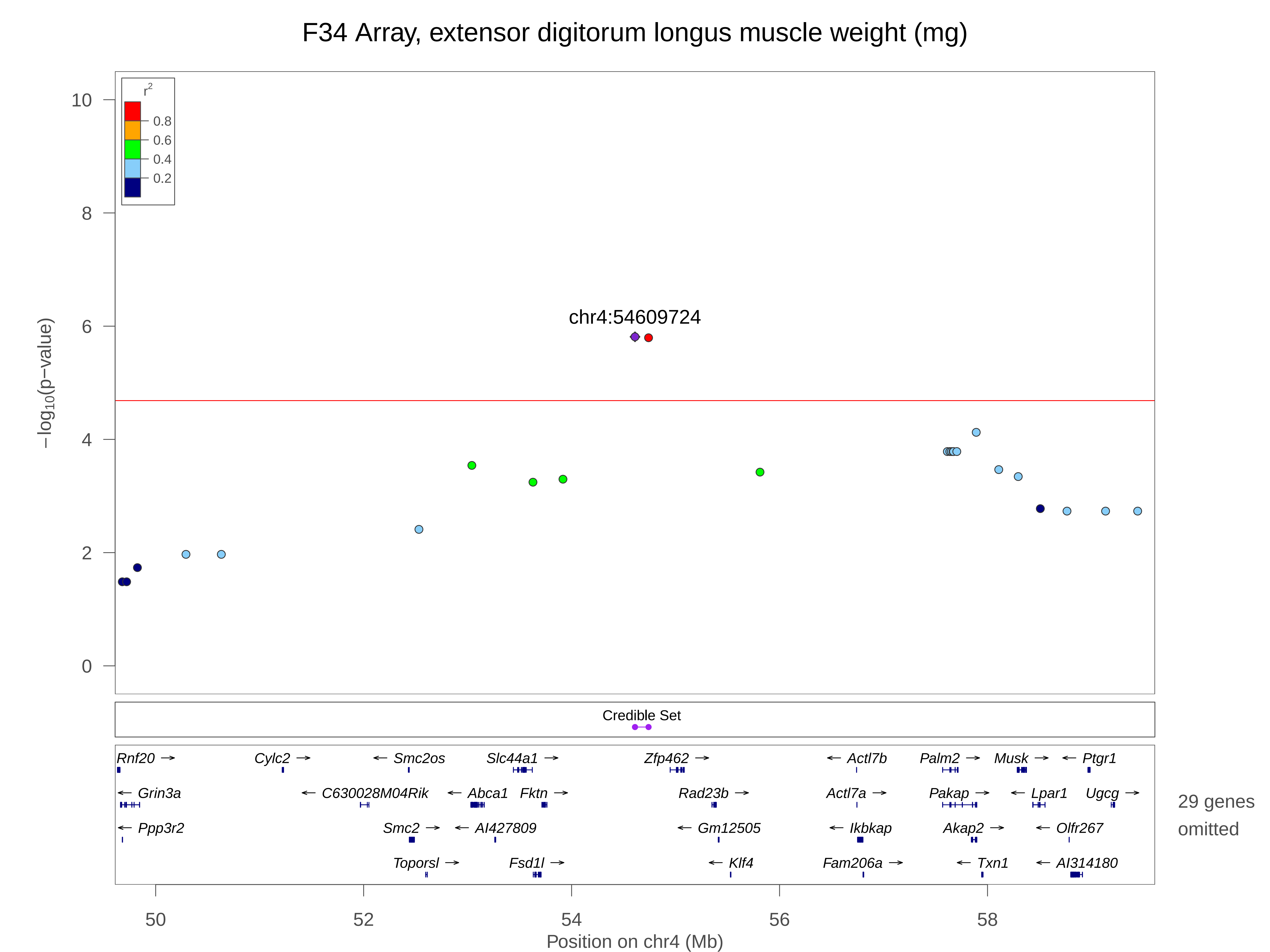
F34 Array, extensor digitorum longus muscle weight (mg)
r2
0.8
0.6
0.4
0.2
−log10(p−value)
10
8
6
4
2
0
chr4:54609724
Credible Set
Rnf20 →
Cylc2 →
← Smc2os
Slc44a1 →
Zfp462 →
← Actl7b
Palm2 →
Musk →
← Ptgr1
← Grin3a
← C630028M04Rik
← Abca1
Fktn →
Rad23b →
Actl7a →
Pakap →
← Lpar1
Ugcg →
← Ppp3r2
Smc2 →
← AI427809
← Gm12505
← Ikbkap
Akap2 →
← Olfr267
Toporsl →
Fsd1l →
← Klf4
Fam206a →
← Txn1
← AI314180
29 genes omitted
50
52
54
56
58
Position on chr4 (Mb)

| | |
|---|---|
| date: | Thu Oct 25 16:56:56 2018 |
| build: | F34Array |
| display range: | chr4:49609724−59609724 [49609724−59609724] |
| hilite range: | 0 − 0 [ 0 − 0 ] |
| reference SNP: | chr4:54609724 |
| number of SNPs plotted: | 26 |
| min P−value: | 1.54E−6 [chr4:54609724] |
| max P−value: | 3.27E−2 [chr4:49679066] |
| omitted Genes: | 4930552N02Rik, Olfr275, Olfr273 |
| omitted Genes: | Olfr272, Olfr270, Nipsnap3a |
| omitted Genes: | Nipsnap3b, 4930522O17Rik, Tal2 |
| omitted Genes: | Tmem38b, Ctnnal1, Tmem245 |
| omitted Genes: | Mir32, Frrs1l, Epb41l4b |
| omitted Genes: | Gm12530, Ptpn3, 1700042G15Rik |
| omitted Genes: | D630039A03Rik, Txndc8, Svep1 |
| omitted Genes: | Mir3095, Zkscan16, Dnajc25 |
| omitted Genes: | Gng10, Gm12596, Susd1 |
| omitted Genes: | Ptbp3, Hsdl2 |

Make more plots at http://csg.sph.umich.edu/locuszoom/

Fine−mapping Regions

| snp | chr | pos (Mb) | pp | group | color |
|---|---|---|---|---|---|
| chr4:54609724 | 4 | 54.60972 | 0.5090756 | Credible Set | purple |
| chr4:54740078 | 4 | 54.74008 | 0.4909244 | Credible Set | purple |

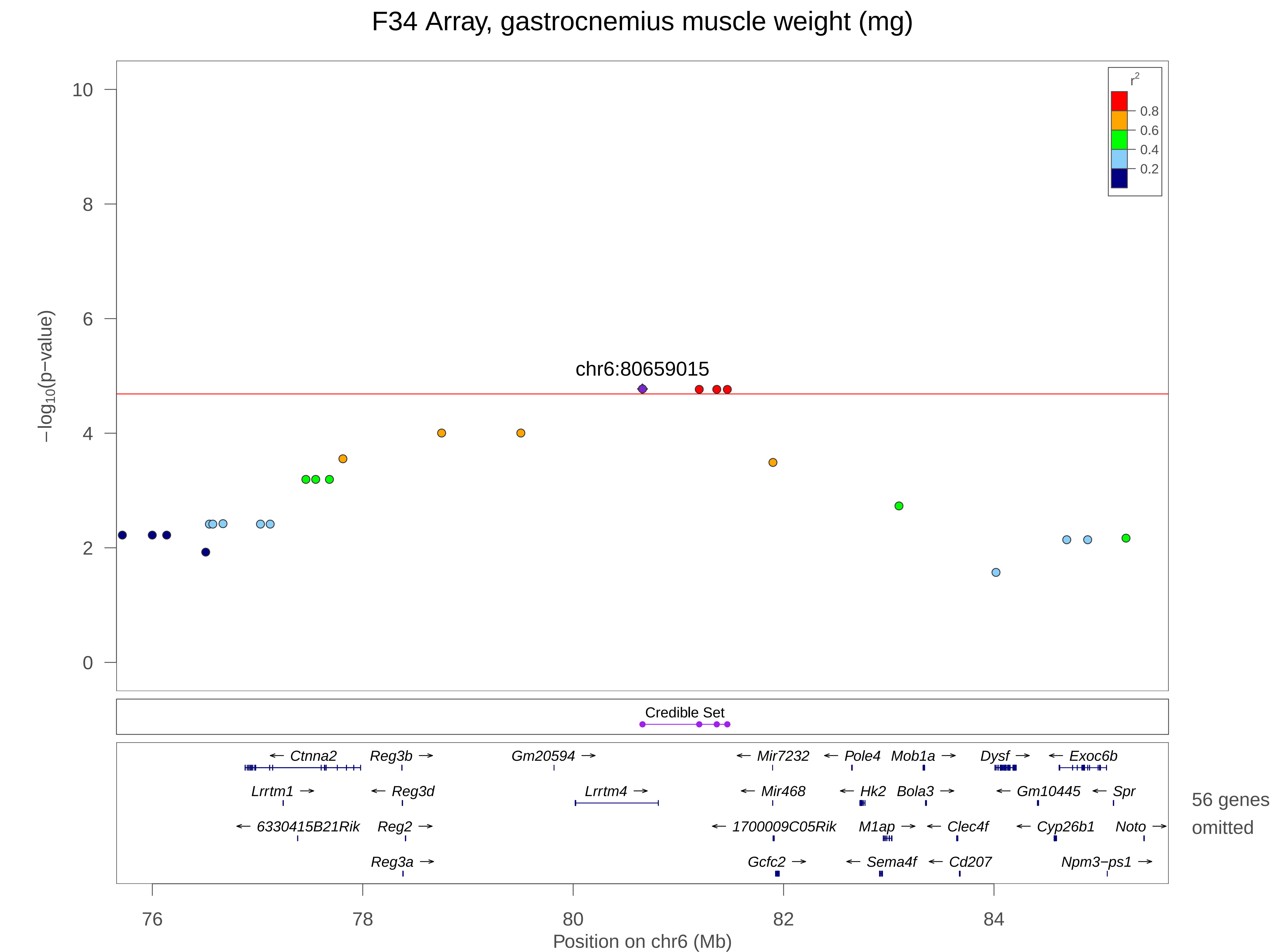
F34 Array, gastrocnemius muscle weight (mg)
r2
0.8
0.6
0.4
0.2
chr6:80659015
−log10(p−value)
10
8
6
4
2
0
Credible Set
← Ctnna2
Reg3b →
Gm20594 →
← Mir7232
← Pole4
Mob1a →
Dysf →
← Exoc6b
Lrrtm1 →
← Reg3d
Lrrtm4 →
← Mir468
← Hk2
Bola3 →
← Gm10445
← Spr
← 6330415B21Rik
Reg2 →
← 1700009C05Rik
M1ap →
← Clec4f
← Cyp26b1
Noto →
Reg3a →
Gcfc2 →
← Sema4f
← Cd207
Npm3−ps1 →
56 genes omitted
76
78
80
82
84
Position on chr6 (Mb)

| | |
|---|---|
| date: | Thu Oct 25 16:57:03 2018 |
| build: | F34Array |
| display range: | chr6:75659015−85659015 [75659015−85659015] |
| hilite range: | 0 − 0 [ 0 − 0 ] |
| reference SNP: | chr6:80659015 |
| number of SNPs plotted: | 26 |
| min P−value: | 1.69E−5 [chr6:80659015] |
| max P−value: | 2.68E−2 [chr6:84018933] |
| omitted Genes: | Reg1, Reg3g, Gm20362 |
| omitted Genes: | Mrpl19, Eva1a, Tacr1 |
| omitted Genes: | 2310069B03Rik, Dok1, Loxl3 |
| omitted Genes: | Mir7040, Htra2, Aup1 |
| omitted Genes: | Dqx1, Tlx2, Pcgf1 |
| omitted Genes: | Lbx2, Mir3470a, Ccdc142 |
| omitted Genes: | Mrpl53, Mogs, Wbp1 |
| omitted Genes: | Ino80b, Rtkn, Wdr54 |
| omitted Genes: | 1700003E16Rik, Dctn1, Slc4a5 |
| omitted Genes: | Mthfd2, Tet3, Mir6374 |
| omitted Genes: | B230319C09Rik, Dguok, Actg2 |
| omitted Genes: | Stambp, Gm21284, Vax2os |
| omitted Genes: | Vax2, Atp6v1b1, 1700124L16Rik |
| omitted Genes: | Ankrd53, Tex261, Gm7443 |
| omitted Genes: | Nagk, Paip2b, Zfp638 |
| omitted Genes: | Gm5878, Emx1, Sfxn5 |
| omitted Genes: | Rab11fip5, Mir705, Smyd5 |
| omitted Genes: | Pradc1, Cct7, Fbxo41 |
| omitted Genes: | Egr4, Alms1 |

Make more plots at http://csg.sph.umich.edu/locuszoom/

Fine−mapping Regions

| snp | chr | pos (Mb) | pp | group | color |
|---|---|---|---|---|---|
| chr6:80659015 | 6 | 80.65901 | 0.2530481 | Credible Set | purple |
| chr6:81198298 | 6 | 81.19830 | 0.2491541 | Credible Set | purple |
| chr6:81364957 | 6 | 81.36496 | 0.2491541 | Credible Set | purple |
| chr6:81465256 | 6 | 81.46526 | 0.2486437 | Credible Set | purple |

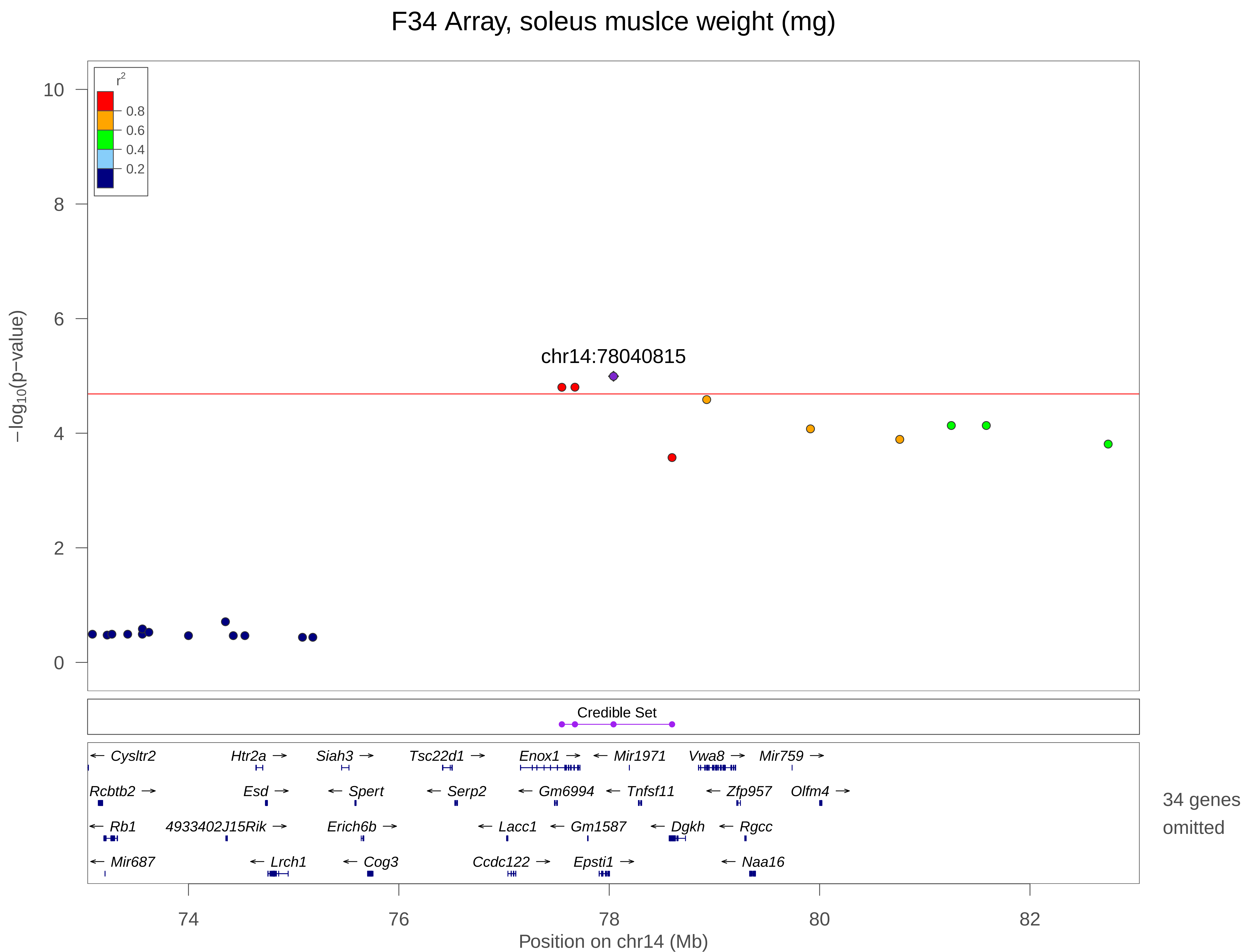
F34 Array, soleus muslce weight (mg)
r2
0.8
0.6
0.4
0.2
chr14:78040815
−log10(p−value)
Credible Set
← Cysltr2
Htr2a →
Siah3 →
Tsc22d1 →
Enox1 →
← Mir1971
Vwa8 →
Mir759 →
Rcbtb2 →
Esd →
← Spert
← Serp2
← Gm6994
← Tnfsf11
← Zfp957
Olfm4 →
← Rb1
4933402J15Rik →
Erich6b →
← Lacc1
← Gm1587
← Dgkh
← Rgcc
← Mir687
← Lrch1
← Cog3
Ccdc122 →
Epsti1 →
← Naa16
34 genes
omitted
Position on chr14 (Mb)

| | |
|---|---|
| date: | Thu Oct 25 16:57:10 2018 |
| build: | F34Array |
| display range: | chr14:73040815−83040815 [73040815−83040815] |
| hilite range: | 0 − 0 [ 0 − 0 ] |
| reference SNP: | chr14:78040815 |
| number of SNPs plotted: | 24 |
| min P−value: | 1.01E−5 [chr14:78040815] |
| max P−value: | 3.64E−1 [chr14:75083773] |
| omitted Genes: | Lpar6, Itm2b, Med4 |
| omitted Genes: | Nudt15, Sucla2, Gm4278 |
| omitted Genes: | 5031414D18Rik, Lrrc63, Lcp1 |
| omitted Genes: | Cpb2, Zc3h13, Slc25a30 |
| omitted Genes: | Gm4285, Tpt1, Snora31 |
| omitted Genes: | Gtf2f2, Kctd4, Gpalpp1 |
| omitted Genes: | Nufip1, 4930444M15Rik, 1700108F19Rik |
| omitted Genes: | Dnajc15, Fam216b, Akap11 |
| omitted Genes: | Mtrf1, Kbtbd7, Kbtbd6 |
| omitted Genes: | Wbp4, Elf1, Sugt1 |
| omitted Genes: | Lect1, 4930452G13Rik, Pcdh8 |
| omitted Genes: | Gm10845 |

Make more plots at http://csg.sph.umich.edu/locuszoom/

Fine−mapping Regions

| snp | chr | pos (Mb) | pp | group | color |
|---|---|---|---|---|---|
| chr14:77549688 | 14 | 77.54969 | 0.2781654 | Credible Set | purple |
| chr14:77673987 | 14 | 77.67399 | 0.2789379 | Credible Set | purple |
| chr14:78040815 | 14 | 78.04081 | 0.4236906 | Credible Set | purple |
| chr14:78597230 | 14 | 78.59723 | 0.0192061 | Credible Set | purple |

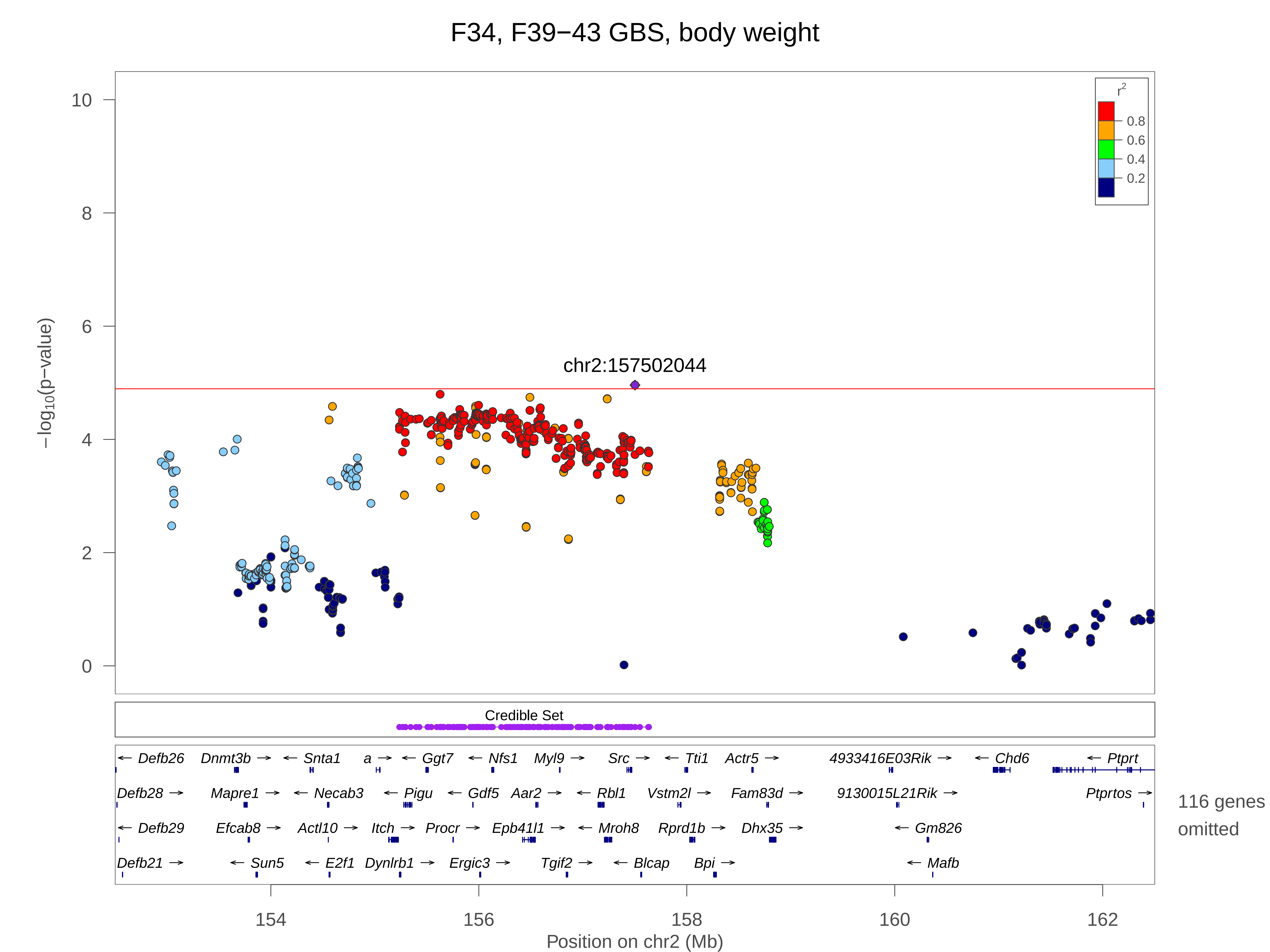
F34, F39−43 GBS, body weight
−log10(p−value)
10
8
6
4
2
0
r2
0.8
0.6
0.4
0.2
chr2:157502044
Credible Set
Defb26
Dnmt3b
Snta1
a
Ggt7
Nfs1
Myl9
Src
Tti1
Actr5
4933416E03Rik
Chd6
Ptprt
Defb28
Mapre1
Necab3
Pigu
Gdf5
Aar2
Rbl1
Vstm2l
Fam83d
9130015L21Rik
Ptprtos
Defb29
Efcab8
Actl10
Itch
Procr
Epb41l1
Mroh8
Rprd1b
Dhx35
Gm826
Defb21
Sun5
E2f1
Dynlrb1
Ergic3
Tgif2
Blcap
Bpi
Mafb
116 genes omitted
154
156
158
160
162
Position on chr2 (Mb)

date: Thu Oct 25 16:57:28 2018
build: F34F3943GBS
display range: chr2:152502044−162502044 [152502044−162502044]
hilite range: 0 − 0 [ 0 − 0 ]
reference SNP: chr2:157502044
number of SNPs plotted: 729
min P−value: 1.1E−5 [chr2:157502044]
max P−value: 9.69E−1 [chr2:161219896]
omitted Genes: Defb19, Defb45, Defb36
omitted Genes: Defb25, Rem1, H13
omitted Genes: Mcts2, Id1, Cox4i2
omitted Genes: Bcl2l1, Tpx2, Mir5622
omitted Genes: Mylk2, Foxs1, Dusp15
omitted Genes: Ttll9, Pdrg1, Xkr7
omitted Genes: Ccm2l, Hck, Tm9sf4
omitted Genes: Tspyl3, Plagl2, Pofut1
omitted Genes: Kif3b, 2500004C02Rik, Asxl1
omitted Genes: Nol4l, Commd7, Bpifb2
omitted Genes: Bpifb6, Bpifb3, Bpifb4
omitted Genes: Bpifa6, Bpifa2, Bpifa3
omitted Genes: Bpifa1, Bpifa5, Bpifb1
omitted Genes: Bpifb5, Bpifb9a, Bpifb9b
omitted Genes: Cdk5rap1, Cbfa2t2, 1700003F12Rik
omitted Genes: Pxmp4, Zfp341, Chmp4b
omitted Genes: Raly, Eif2s2, Ahcy
omitted Genes: 2310005A03Rik, Map1lc3a, Mir695
omitted Genes: Trp53inp2, Ncoa6, Acss2
omitted Genes: Acss2os, Gss, Myh7b
omitted Genes: Mir499, Trpc4ap, Edem2
omitted Genes: Mmp24, BC029722, Eif6
omitted Genes: Fam83c, Uqcc1, Cep250
omitted Genes: 6430550D23Rik, Fer1l4, Spag4
omitted Genes: Cpne1, Rbm12, Romo1
omitted Genes: Rbm39, Phf20, Scand1
omitted Genes: Cnbd2, 2900097C17Rik, Gm14169
omitted Genes: Dlgap4, 4930405A21Rik, 5430405H02Rik
omitted Genes: 1110008F13Rik, Sla2, Ndrg3
omitted Genes: Dsn1, Soga1, Tldc2
omitted Genes: Samhd1, Rpn2, Ghrh
Make more plots at http://csg.sph.umich.edu/locuszoom/
omitted Genes: Manbal, Nnat, Ctnnbl1

Fine−mapping Regions

| snp | chr | pos (Mb) | pp | group | color |
|---|---|---|---|---|---|
| chr2:155236637 | 2 | 155.2366 | 0.003464254 | Credible Set | purple |
| chr2:155236657 | 2 | 155.2367 | 0.003412400 | Credible Set | purple |
| chr2:155236987 | 2 | 155.2370 | 0.003115889 | Credible Set | purple |
| chr2:155237968 | 2 | 155.2380 | 0.005984279 | Credible Set | purple |
| chr2:155265639 | 2 | 155.2656 | 0.004245892 | Credible Set | purple |
| chr2:155266429 | 2 | 155.2664 | 0.001304812 | Credible Set | purple |
| chr2:155266459 | 2 | 155.2665 | 0.004313901 | Credible Set | purple |
| chr2:155291037 | 2 | 155.2910 | 0.002774933 | Credible Set | purple |
| chr2:155291486 | 2 | 155.2915 | 0.005200710 | Credible Set | purple |
| chr2:155294582 | 2 | 155.2946 | 0.004064489 | Credible Set | purple |
| chr2:155294781 | 2 | 155.2948 | 0.001867995 | Credible Set | purple |
| chr2:155294817 | 2 | 155.2948 | 0.001859333 | Credible Set | purple |
| chr2:155344176 | 2 | 155.3442 | 0.004618114 | Credible Set | purple |
| chr2:155396284 | 2 | 155.3963 | 0.004588433 | Credible Set | purple |
| chr2:155427716 | 2 | 155.4277 | 0.004685690 | Credible Set | purple |
| chr2:155506374 | 2 | 155.5064 | 0.003940804 | Credible Set | purple |
| chr2:155514315 | 2 | 155.5143 | 0.004033277 | Credible Set | purple |
| chr2:155542937 | 2 | 155.5429 | 0.004413868 | Credible Set | purple |
| chr2:155543113 | 2 | 155.5431 | 0.002550139 | Credible Set | purple |
| chr2:155543163 | 2 | 155.5432 | 0.002512879 | Credible Set | purple |
| chr2:155596243 | 2 | 155.5962 | 0.003334274 | Credible Set | purple |
| chr2:155596245 | 2 | 155.5962 | 0.003363219 | Credible Set | purple |
| chr2:155596254 | 2 | 155.5963 | 0.003368160 | Credible Set | purple |
| chr2:155626668 | 2 | 155.6267 | 0.004665526 | Credible Set | purple |
| chr2:155626706 | 2 | 155.6267 | 0.004665526 | Credible Set | purple |
| chr2:155627928 | 2 | 155.6279 | 0.012087851 | Credible Set | purple |
| chr2:155630405 | 2 | 155.6304 | 0.004775484 | Credible Set | purple |
| chr2:155630630 | 2 | 155.6306 | 0.004718763 | Credible Set | purple |
| chr2:155630746 | 2 | 155.6307 | 0.004768401 | Credible Set | purple |
| chr2:155631110 | 2 | 155.6311 | 0.004873521 | Credible Set | purple |
| chr2:155632202 | 2 | 155.6322 | 0.004294823 | Credible Set | purple |
| chr2:155632358 | 2 | 155.6324 | 0.004499151 | Credible Set | purple |
| chr2:155632880 | 2 | 155.6329 | 0.004508078 | Credible Set | purple |
| chr2:155633328 | 2 | 155.6333 | 0.004549754 | Credible Set | purple |
| chr2:155640583 | 2 | 155.6406 | 0.003531773 | Credible Set | purple |
| chr2:155640694 | 2 | 155.6407 | 0.005025648 | Credible Set | purple |

Fine−mapping Regions

| snp | chr | pos (Mb) | pp | group | color |
|---|---|---|---|---|---|
| chr2:155640700 | 2 | 155.6407 | 0.005223632 | Credible Set | purple |
| chr2:155641730 | 2 | 155.6417 | 0.004765478 | Credible Set | purple |
| chr2:155647032 | 2 | 155.6470 | 0.004078487 | Credible Set | purple |
| chr2:155647066 | 2 | 155.6471 | 0.003208476 | Credible Set | purple |
| chr2:155659679 | 2 | 155.6597 | 0.004197479 | Credible Set | purple |
| chr2:155659722 | 2 | 155.6597 | 0.004128186 | Credible Set | purple |
| chr2:155664276 | 2 | 155.6643 | 0.004312228 | Credible Set | purple |
| chr2:155703392 | 2 | 155.7034 | 0.001787672 | Credible Set | purple |
| chr2:155703416 | 2 | 155.7034 | 0.001840882 | Credible Set | purple |
| chr2:155703421 | 2 | 155.7034 | 0.001670214 | Credible Set | purple |
| chr2:155718717 | 2 | 155.7187 | 0.004011865 | Credible Set | purple |
| chr2:155718886 | 2 | 155.7189 | 0.003607960 | Credible Set | purple |
| chr2:155718943 | 2 | 155.7189 | 0.003654915 | Credible Set | purple |
| chr2:155748773 | 2 | 155.7488 | 0.004630972 | Credible Set | purple |
| chr2:155754530 | 2 | 155.7545 | 0.004456573 | Credible Set | purple |
| chr2:155754728 | 2 | 155.7547 | 0.004360527 | Credible Set | purple |
| chr2:155765708 | 2 | 155.7657 | 0.004912680 | Credible Set | purple |
| chr2:155787135 | 2 | 155.7871 | 0.004962641 | Credible Set | purple |
| chr2:155787323 | 2 | 155.7873 | 0.004767340 | Credible Set | purple |
| chr2:155804255 | 2 | 155.8043 | 0.002458148 | Credible Set | purple |
| chr2:155804262 | 2 | 155.8043 | 0.002927899 | Credible Set | purple |
| chr2:155804269 | 2 | 155.8043 | 0.002780520 | Credible Set | purple |
| chr2:155813877 | 2 | 155.8139 | 0.005239627 | Credible Set | purple |
| chr2:155816627 | 2 | 155.8166 | 0.006726201 | Credible Set | purple |
| chr2:155818442 | 2 | 155.8184 | 0.005431683 | Credible Set | purple |
| chr2:155819513 | 2 | 155.8195 | 0.005427257 | Credible Set | purple |
| chr2:155819633 | 2 | 155.8196 | 0.003329201 | Credible Set | purple |
| chr2:155819676 | 2 | 155.8197 | 0.003297482 | Credible Set | purple |
| chr2:155827653 | 2 | 155.8277 | 0.004150075 | Credible Set | purple |
| chr2:155827705 | 2 | 155.8277 | 0.003676437 | Credible Set | purple |
| chr2:155828085 | 2 | 155.8281 | 0.005426775 | Credible Set | purple |
| chr2:155828345 | 2 | 155.8283 | 0.005391394 | Credible Set | purple |
| chr2:155828819 | 2 | 155.8288 | 0.005423197 | Credible Set | purple |
| chr2:155830487 | 2 | 155.8305 | 0.005388799 | Credible Set | purple |
| chr2:155830972 | 2 | 155.8310 | 0.004930105 | Credible Set | purple |
| chr2:155843797 | 2 | 155.8438 | 0.005460715 | Credible Set | purple |

Fine−mapping Regions

| snp | chr | pos (Mb) | pp | group | color |
|---|---|---|---|---|---|
| chr2:155843802 | 2 | 155.8438 | 0.005462799 | Credible Set | purple |
| chr2:155845413 | 2 | 155.8454 | 0.005436017 | Credible Set | purple |
| chr2:155859749 | 2 | 155.8597 | 0.005362268 | Credible Set | purple |
| chr2:155860022 | 2 | 155.8600 | 0.004217732 | Credible Set | purple |
| chr2:155917786 | 2 | 155.9178 | 0.003149380 | Credible Set | purple |
| chr2:155917800 | 2 | 155.9178 | 0.003128783 | Credible Set | purple |
| chr2:155934881 | 2 | 155.9349 | 0.003748017 | Credible Set | purple |
| chr2:155935242 | 2 | 155.9352 | 0.003955028 | Credible Set | purple |
| chr2:155941490 | 2 | 155.9415 | 0.004197866 | Credible Set | purple |
| chr2:155941536 | 2 | 155.9415 | 0.004197866 | Credible Set | purple |
| chr2:155962070 | 2 | 155.9621 | 0.004800666 | Credible Set | purple |
| chr2:155969145 | 2 | 155.9691 | 0.005393328 | Credible Set | purple |
| chr2:155969338 | 2 | 155.9693 | 0.004438912 | Credible Set | purple |
| chr2:155973018 | 2 | 155.9730 | 0.005882709 | Credible Set | purple |
| chr2:155973035 | 2 | 155.9730 | 0.005854227 | Credible Set | purple |
| chr2:155973697 | 2 | 155.9737 | 0.005720146 | Credible Set | purple |
| chr2:155974320 | 2 | 155.9743 | 0.005730225 | Credible Set | purple |
| chr2:155974339 | 2 | 155.9743 | 0.005621049 | Credible Set | purple |
| chr2:155974623 | 2 | 155.9746 | 0.005413386 | Credible Set | purple |
| chr2:155985744 | 2 | 155.9857 | 0.005290409 | Credible Set | purple |
| chr2:155986052 | 2 | 155.9861 | 0.005366294 | Credible Set | purple |
| chr2:155988485 | 2 | 155.9885 | 0.005089466 | Credible Set | purple |
| chr2:155989053 | 2 | 155.9891 | 0.005160591 | Credible Set | purple |
| chr2:155991302 | 2 | 155.9913 | 0.005364305 | Credible Set | purple |
| chr2:155991810 | 2 | 155.9918 | 0.005364305 | Credible Set | purple |
| chr2:155992021 | 2 | 155.9920 | 0.005347681 | Credible Set | purple |
| chr2:155997923 | 2 | 155.9979 | 0.007906597 | Credible Set | purple |
| chr2:155998479 | 2 | 155.9985 | 0.005675815 | Credible Set | purple |
| chr2:156011028 | 2 | 156.0110 | 0.005166186 | Credible Set | purple |
| chr2:156011085 | 2 | 156.0111 | 0.005485015 | Credible Set | purple |
| chr2:156011672 | 2 | 156.0117 | 0.005435624 | Credible Set | purple |
| chr2:156039455 | 2 | 156.0395 | 0.005292218 | Credible Set | purple |
| chr2:156040637 | 2 | 156.0406 | 0.005106331 | Credible Set | purple |
| chr2:156040653 | 2 | 156.0407 | 0.005122289 | Credible Set | purple |
| chr2:156041520 | 2 | 156.0415 | 0.005315708 | Credible Set | purple |
| chr2:156041911 | 2 | 156.0419 | 0.004334184 | Credible Set | purple |

Fine−mapping Regions

| snp | chr | pos (Mb) | pp | group | color |
|---|---|---|---|---|---|
| chr2:156041912 | 2 | 156.0419 | 0.004255881 | Credible Set | purple |
| chr2:156043677 | 2 | 156.0437 | 0.005283196 | Credible Set | purple |
| chr2:156071586 | 2 | 156.0716 | 0.005654522 | Credible Set | purple |
| chr2:156073463 | 2 | 156.0735 | 0.003673781 | Credible Set | purple |
| chr2:156077894 | 2 | 156.0779 | 0.004820393 | Credible Set | purple |
| chr2:156078317 | 2 | 156.0783 | 0.004854496 | Credible Set | purple |
| chr2:156078406 | 2 | 156.0784 | 0.005503233 | Credible Set | purple |
| chr2:156078707 | 2 | 156.0787 | 0.004747875 | Credible Set | purple |
| chr2:156078913 | 2 | 156.0789 | 0.004559174 | Credible Set | purple |
| chr2:156082428 | 2 | 156.0824 | 0.004851955 | Credible Set | purple |
| chr2:156082588 | 2 | 156.0826 | 0.004840536 | Credible Set | purple |
| chr2:156082682 | 2 | 156.0827 | 0.005260584 | Credible Set | purple |
| chr2:156111908 | 2 | 156.1119 | 0.005514199 | Credible Set | purple |
| chr2:156111910 | 2 | 156.1119 | 0.005580051 | Credible Set | purple |
| chr2:156124158 | 2 | 156.1242 | 0.005642918 | Credible Set | purple |
| chr2:156124843 | 2 | 156.1248 | 0.005542267 | Credible Set | purple |
| chr2:156133370 | 2 | 156.1334 | 0.006101269 | Credible Set | purple |
| chr2:156133395 | 2 | 156.1334 | 0.006186832 | Credible Set | purple |
| chr2:156133510 | 2 | 156.1335 | 0.004541558 | Credible Set | purple |
| chr2:156215789 | 2 | 156.2158 | 0.004853563 | Credible Set | purple |
| chr2:156259843 | 2 | 156.2598 | 0.002506363 | Credible Set | purple |
| chr2:156267477 | 2 | 156.2675 | 0.004696835 | Credible Set | purple |
| chr2:156269831 | 2 | 156.2698 | 0.004882722 | Credible Set | purple |
| chr2:156282307 | 2 | 156.2823 | 0.004557349 | Credible Set | purple |
| chr2:156302628 | 2 | 156.3026 | 0.005832612 | Credible Set | purple |
| chr2:156302714 | 2 | 156.3027 | 0.003568420 | Credible Set | purple |
| chr2:156302728 | 2 | 156.3027 | 0.003594033 | Credible Set | purple |
| chr2:156302741 | 2 | 156.3027 | 0.004741867 | Credible Set | purple |
| chr2:156303023 | 2 | 156.3030 | 0.004742166 | Credible Set | purple |
| chr2:156305013 | 2 | 156.3050 | 0.002133821 | Credible Set | purple |
| chr2:156323079 | 2 | 156.3231 | 0.004802958 | Credible Set | purple |
| chr2:156343221 | 2 | 156.3432 | 0.004808018 | Credible Set | purple |
| chr2:156343324 | 2 | 156.3433 | 0.003172128 | Credible Set | purple |
| chr2:156368682 | 2 | 156.3687 | 0.003348256 | Credible Set | purple |
| chr2:156368698 | 2 | 156.3687 | 0.004071623 | Credible Set | purple |
| chr2:156378531 | 2 | 156.3785 | 0.002896125 | Credible Set | purple |

Fine−mapping Regions

| snp | chr | pos (Mb) | pp | group | color |
|---|---|---|---|---|---|
| chr2:156387857 | 2 | 156.3879 | 0.002443110 | Credible Set | purple |
| chr2:156401411 | 2 | 156.4014 | 0.002112180 | Credible Set | purple |
| chr2:156407061 | 2 | 156.4071 | 0.001811764 | Credible Set | purple |
| chr2:156407080 | 2 | 156.4071 | 0.001820027 | Credible Set | purple |
| chr2:156407368 | 2 | 156.4074 | 0.002091303 | Credible Set | purple |
| chr2:156417925 | 2 | 156.4179 | 0.002025344 | Credible Set | purple |
| chr2:156419006 | 2 | 156.4190 | 0.002137265 | Credible Set | purple |
| chr2:156423785 | 2 | 156.4238 | 0.002215814 | Credible Set | purple |
| chr2:156449123 | 2 | 156.4491 | 0.001800289 | Credible Set | purple |
| chr2:156449144 | 2 | 156.4491 | 0.001797246 | Credible Set | purple |
| chr2:156453473 | 2 | 156.4535 | 0.001713582 | Credible Set | purple |
| chr2:156453552 | 2 | 156.4536 | 0.001209756 | Credible Set | purple |
| chr2:156453983 | 2 | 156.4540 | 0.001328549 | Credible Set | purple |
| chr2:156455282 | 2 | 156.4553 | 0.001244008 | Credible Set | purple |
| chr2:156455298 | 2 | 156.4553 | 0.001261566 | Credible Set | purple |
| chr2:156465034 | 2 | 156.4650 | 0.002499351 | Credible Set | purple |
| chr2:156465975 | 2 | 156.4660 | 0.003496082 | Credible Set | purple |
| chr2:156468230 | 2 | 156.4682 | 0.003540299 | Credible Set | purple |
| chr2:156474927 | 2 | 156.4749 | 0.002799408 | Credible Set | purple |
| chr2:156483143 | 2 | 156.4831 | 0.002239584 | Credible Set | purple |
| chr2:156488289 | 2 | 156.4883 | 0.002898112 | Credible Set | purple |
| chr2:156491541 | 2 | 156.4915 | 0.006458262 | Credible Set | purple |
| chr2:156491557 | 2 | 156.4916 | 0.006490324 | Credible Set | purple |
| chr2:156521595 | 2 | 156.5216 | 0.001949490 | Credible Set | purple |
| chr2:156521639 | 2 | 156.5216 | 0.001933845 | Credible Set | purple |
| chr2:156530706 | 2 | 156.5307 | 0.001957706 | Credible Set | purple |
| chr2:156532009 | 2 | 156.5320 | 0.003290282 | Credible Set | purple |
| chr2:156532011 | 2 | 156.5320 | 0.003227752 | Credible Set | purple |
| chr2:156532178 | 2 | 156.5322 | 0.002214033 | Credible Set | purple |
| chr2:156567043 | 2 | 156.5670 | 0.003454284 | Credible Set | purple |
| chr2:156567147 | 2 | 156.5671 | 0.004270274 | Credible Set | purple |
| chr2:156567188 | 2 | 156.5672 | 0.004255686 | Credible Set | purple |
| chr2:156567685 | 2 | 156.5677 | 0.003374958 | Credible Set | purple |
| chr2:156576405 | 2 | 156.5764 | 0.003697046 | Credible Set | purple |
| chr2:156576851 | 2 | 156.5769 | 0.003641216 | Credible Set | purple |
| chr2:156577083 | 2 | 156.5771 | 0.003499752 | Credible Set | purple |

Fine−mapping Regions

| snp | chr | pos (Mb) | pp | group | color |
|---|---|---|---|---|---|
| chr2:156585381 | 2 | 156.5854 | 0.0031373417 | Credible Set | purple |
| chr2:156587373 | 2 | 156.5874 | 0.0068286765 | Credible Set | purple |
| chr2:156587398 | 2 | 156.5874 | 0.0066638586 | Credible Set | purple |
| chr2:156592202 | 2 | 156.5922 | 0.0071619124 | Credible Set | purple |
| chr2:156595258 | 2 | 156.5953 | 0.0050107373 | Credible Set | purple |
| chr2:156632710 | 2 | 156.6327 | 0.0035710491 | Credible Set | purple |
| chr2:156632738 | 2 | 156.6327 | 0.0035775720 | Credible Set | purple |
| chr2:156632859 | 2 | 156.6329 | 0.0036849568 | Credible Set | purple |
| chr2:156641079 | 2 | 156.6411 | 0.0037715582 | Credible Set | purple |
| chr2:156641293 | 2 | 156.6413 | 0.0035848273 | Credible Set | purple |
| chr2:156649922 | 2 | 156.6499 | 0.0025984413 | Credible Set | purple |
| chr2:156649929 | 2 | 156.6499 | 0.0026030714 | Credible Set | purple |
| chr2:156650908 | 2 | 156.6509 | 0.0026748524 | Credible Set | purple |
| chr2:156668062 | 2 | 156.6681 | 0.0020853404 | Credible Set | purple |
| chr2:156668618 | 2 | 156.6686 | 0.0022471289 | Credible Set | purple |
| chr2:156704128 | 2 | 156.7041 | 0.0026491077 | Credible Set | purple |
| chr2:156704178 | 2 | 156.7042 | 0.0026521994 | Credible Set | purple |
| chr2:156711128 | 2 | 156.7111 | 0.0029699087 | Credible Set | purple |
| chr2:156743106 | 2 | 156.7431 | 0.0010177636 | Credible Set | purple |
| chr2:156743119 | 2 | 156.7431 | 0.0010261599 | Credible Set | purple |
| chr2:156764741 | 2 | 156.7647 | 0.0015817923 | Credible Set | purple |
| chr2:156764772 | 2 | 156.7648 | 0.0015282847 | Credible Set | purple |
| chr2:156770397 | 2 | 156.7704 | 0.0022390633 | Credible Set | purple |
| chr2:156795214 | 2 | 156.7952 | 0.0021430373 | Credible Set | purple |
| chr2:156795236 | 2 | 156.7952 | 0.0022159256 | Credible Set | purple |
| chr2:156803978 | 2 | 156.8040 | 0.0019834834 | Credible Set | purple |
| chr2:156812752 | 2 | 156.8128 | 0.0032191553 | Credible Set | purple |
| chr2:156825839 | 2 | 156.8258 | 0.0011432370 | Credible Set | purple |
| chr2:156825844 | 2 | 156.8258 | 0.0011455427 | Credible Set | purple |
| chr2:156825876 | 2 | 156.8259 | 0.0011455427 | Credible Set | purple |
| chr2:156845905 | 2 | 156.8459 | 0.0013595092 | Credible Set | purple |
| chr2:156861931 | 2 | 156.8619 | 0.0012955542 | Credible Set | purple |
| chr2:156862185 | 2 | 156.8622 | 0.0007890450 | Credible Set | purple |
| chr2:156862198 | 2 | 156.8622 | 0.0007672412 | Credible Set | purple |
| chr2:156882805 | 2 | 156.8828 | 0.0011521620 | Credible Set | purple |
| chr2:156882992 | 2 | 156.8830 | 0.0008577711 | Credible Set | purple |

Fine−mapping Regions

| snp | chr | pos (Mb) | pp | group | color |
|---|---|---|---|---|---|
| chr2:156883081 | 2 | 156.8831 | 0.0012955563 | Credible Set | purple |
| chr2:156883502 | 2 | 156.8835 | 0.0013118728 | Credible Set | purple |
| chr2:156886515 | 2 | 156.8865 | 0.0012406945 | Credible Set | purple |
| chr2:156886654 | 2 | 156.8867 | 0.0012963735 | Credible Set | purple |
| chr2:156887147 | 2 | 156.8871 | 0.0015148143 | Credible Set | purple |
| chr2:156887169 | 2 | 156.8872 | 0.0015205605 | Credible Set | purple |
| chr2:156948001 | 2 | 156.9480 | 0.0021425069 | Credible Set | purple |
| chr2:156959200 | 2 | 156.9592 | 0.0038667295 | Credible Set | purple |
| chr2:156959224 | 2 | 156.9592 | 0.0039744275 | Credible Set | purple |
| chr2:156959230 | 2 | 156.9592 | 0.0039548773 | Credible Set | purple |
| chr2:156959240 | 2 | 156.9592 | 0.0037438688 | Credible Set | purple |
| chr2:156973206 | 2 | 156.9732 | 0.0017415272 | Credible Set | purple |
| chr2:156973473 | 2 | 156.9735 | 0.0015382514 | Credible Set | purple |
| chr2:157010815 | 2 | 157.0108 | 0.0014694757 | Credible Set | purple |
| chr2:157016739 | 2 | 157.0167 | 0.0014118117 | Credible Set | purple |
| chr2:157017028 | 2 | 157.0170 | 0.0015052076 | Credible Set | purple |
| chr2:157017101 | 2 | 157.0171 | 0.0014015806 | Credible Set | purple |
| chr2:157017479 | 2 | 157.0175 | 0.0016460234 | Credible Set | purple |
| chr2:157017498 | 2 | 157.0175 | 0.0017770082 | Credible Set | purple |
| chr2:157027114 | 2 | 157.0271 | 0.0016627983 | Credible Set | purple |
| chr2:157027115 | 2 | 157.0271 | 0.0016199752 | Credible Set | purple |
| chr2:157027346 | 2 | 157.0273 | 0.0015629572 | Credible Set | purple |
| chr2:157027552 | 2 | 157.0276 | 0.0024364362 | Credible Set | purple |
| chr2:157027594 | 2 | 157.0276 | 0.0014353026 | Credible Set | purple |
| chr2:157033406 | 2 | 157.0334 | 0.0009439426 | Credible Set | purple |
| chr2:157033457 | 2 | 157.0335 | 0.0009444899 | Credible Set | purple |
| chr2:157033569 | 2 | 157.0336 | 0.0010841445 | Credible Set | purple |
| chr2:157033589 | 2 | 157.0336 | 0.0010835272 | Credible Set | purple |
| chr2:157037970 | 2 | 157.0380 | 0.0014006832 | Credible Set | purple |
| chr2:157040364 | 2 | 157.0404 | 0.0011133013 | Credible Set | purple |
| chr2:157040379 | 2 | 157.0404 | 0.0008903294 | Credible Set | purple |
| chr2:157040564 | 2 | 157.0406 | 0.0009705227 | Credible Set | purple |
| chr2:157059250 | 2 | 157.0592 | 0.0010486637 | Credible Set | purple |
| chr2:157059408 | 2 | 157.0594 | 0.0010589980 | Credible Set | purple |
| chr2:157059810 | 2 | 157.0598 | 0.0010928199 | Credible Set | purple |
| chr2:157059893 | 2 | 157.0599 | 0.0009972369 | Credible Set | purple |

Fine−mapping Regions

| snp | chr | pos (Mb) | pp | group | color |
|---|---|---|---|---|---|
| chr2:157063511 | 2 | 157.0635 | 0.0011838957 | Credible Set | purple |
| chr2:157065641 | 2 | 157.0656 | 0.0010353076 | Credible Set | purple |
| chr2:157075626 | 2 | 157.0756 | 0.0011330257 | Credible Set | purple |
| chr2:157075934 | 2 | 157.0759 | 0.0010599377 | Credible Set | purple |
| chr2:157138913 | 2 | 157.1389 | 0.0012723001 | Credible Set | purple |
| chr2:157144291 | 2 | 157.1443 | 0.0013028317 | Credible Set | purple |
| chr2:157145837 | 2 | 157.1458 | 0.0013032310 | Credible Set | purple |
| chr2:157149696 | 2 | 157.1497 | 0.0012348486 | Credible Set | purple |
| chr2:157170168 | 2 | 157.1702 | 0.0012208824 | Credible Set | purple |
| chr2:157234145 | 2 | 157.2341 | 0.0012425076 | Credible Set | purple |
| chr2:157234194 | 2 | 157.2342 | 0.0010569087 | Credible Set | purple |
| chr2:157234584 | 2 | 157.2346 | 0.0011310373 | Credible Set | purple |
| chr2:157234605 | 2 | 157.2346 | 0.0011091951 | Credible Set | purple |
| chr2:157244801 | 2 | 157.2448 | 0.0010773172 | Credible Set | purple |
| chr2:157245205 | 2 | 157.2452 | 0.0010994381 | Credible Set | purple |
| chr2:157245270 | 2 | 157.2453 | 0.0010007732 | Credible Set | purple |
| chr2:157271470 | 2 | 157.2715 | 0.0011382333 | Credible Set | purple |
| chr2:157323455 | 2 | 157.3235 | 0.0007555954 | Credible Set | purple |
| chr2:157323490 | 2 | 157.3235 | 0.0008157944 | Credible Set | purple |
| chr2:157323542 | 2 | 157.3235 | 0.0007564508 | Credible Set | purple |
| chr2:157350851 | 2 | 157.3509 | 0.0008974539 | Credible Set | purple |
| chr2:157356604 | 2 | 157.3566 | 0.0014060960 | Credible Set | purple |
| chr2:157383750 | 2 | 157.3837 | 0.0014465647 | Credible Set | purple |
| chr2:157384030 | 2 | 157.3840 | 0.0023475423 | Credible Set | purple |
| chr2:157384046 | 2 | 157.3840 | 0.0023756954 | Credible Set | purple |
| chr2:157393248 | 2 | 157.3932 | 0.0009728794 | Credible Set | purple |
| chr2:157393326 | 2 | 157.3933 | 0.0008826988 | Credible Set | purple |
| chr2:157393328 | 2 | 157.3933 | 0.0008958535 | Credible Set | purple |
| chr2:157396118 | 2 | 157.3961 | 0.0018808429 | Credible Set | purple |
| chr2:157396325 | 2 | 157.3963 | 0.0011545388 | Credible Set | purple |
| chr2:157396340 | 2 | 157.3963 | 0.0011711049 | Credible Set | purple |
| chr2:157396357 | 2 | 157.3964 | 0.0011652736 | Credible Set | purple |
| chr2:157396513 | 2 | 157.3965 | 0.0020350770 | Credible Set | purple |
| chr2:157401494 | 2 | 157.4015 | 0.0022594395 | Credible Set | purple |
| chr2:157425583 | 2 | 157.4256 | 0.0020445300 | Credible Set | purple |
| chr2:157434588 | 2 | 157.4346 | 0.0018976095 | Credible Set | purple |

Fine−mapping Regions

| snp | chr | pos (Mb) | pp | group | color |
|---|---|---|---|---|---|
| chr2:157439371 | 2 | 157.4394 | 0.001868134 | Credible Set | purple |
| chr2:157455578 | 2 | 157.4556 | 0.001567994 | Credible Set | purple |
| chr2:157458440 | 2 | 157.4584 | 0.001857250 | Credible Set | purple |
| chr2:157464790 | 2 | 157.4648 | 0.002049487 | Credible Set | purple |
| chr2:157464811 | 2 | 157.4648 | 0.002055422 | Credible Set | purple |
| chr2:157464835 | 2 | 157.4648 | 0.002054751 | Credible Set | purple |
| chr2:157464946 | 2 | 157.4649 | 0.002013449 | Credible Set | purple |
| chr2:157465338 | 2 | 157.4653 | 0.002013553 | Credible Set | purple |
| chr2:157502041 | 2 | 157.5020 | 0.001182759 | Credible Set | purple |
| chr2:157502044 | 2 | 157.5020 | 0.017255691 | Credible Set | purple |
| chr2:157549374 | 2 | 157.5494 | 0.001368992 | Credible Set | purple |
| chr2:157630394 | 2 | 157.6304 | 0.001296992 | Credible Set | purple |
| chr2:157630611 | 2 | 157.6306 | 0.001358558 | Credible Set | purple |
| chr2:157635749 | 2 | 157.6357 | 0.001266836 | Credible Set | purple |

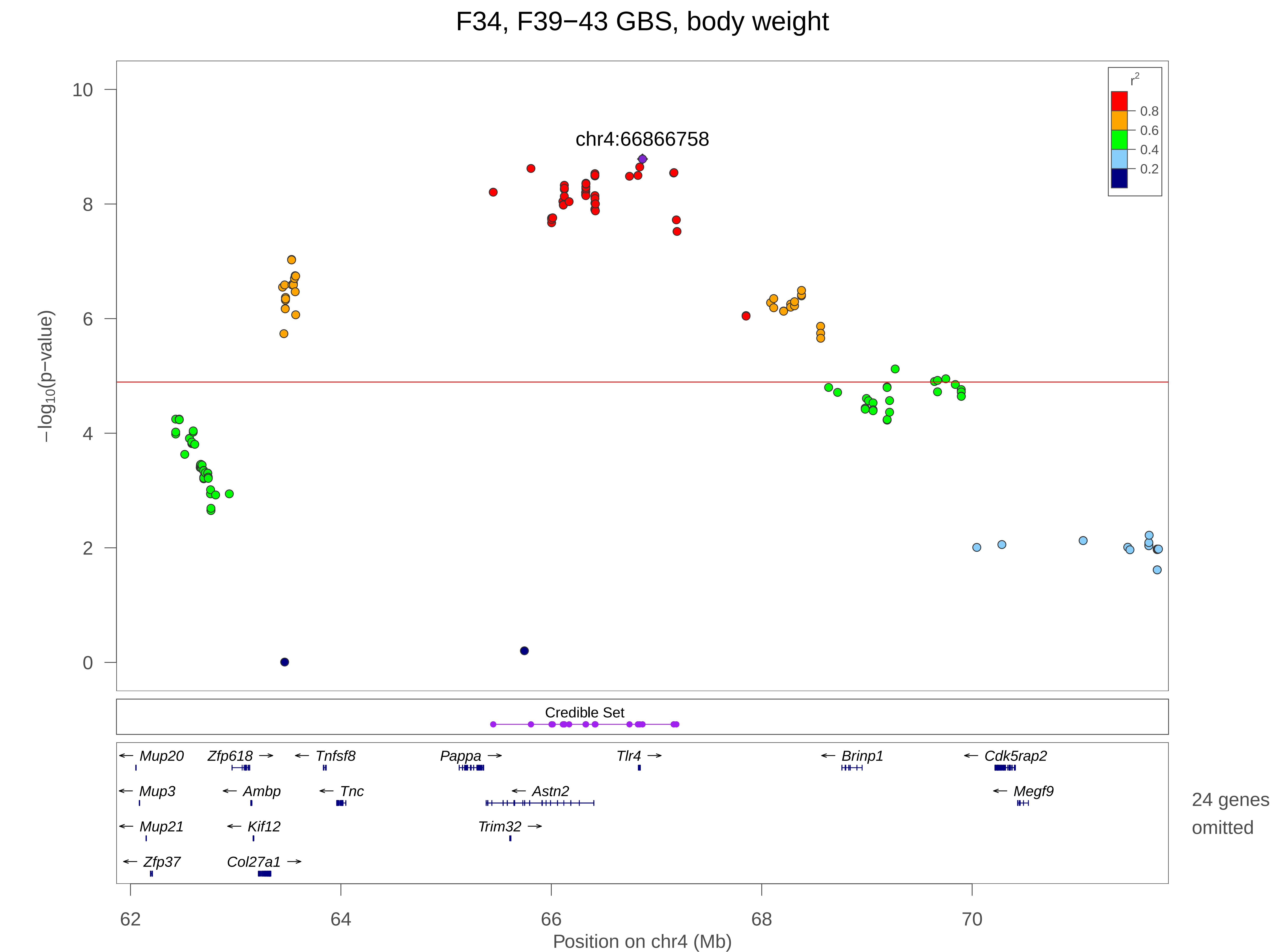

F34, F39−43 GBS, body weight
chr4:66866758
r2
0.8
0.6
0.4
0.2
−log10(p−value)
10
8
6
4
2
0
Credible Set
Mup20
Zfp618
Tnfsf8
Pappa
Tlr4
Brinp1
Cdk5rap2
Mup3
Ambp
Tnc
Astn2
Megf9
Mup21
Kif12
Trim32
Zfp37
Col27a1
24 genes omitted
62
64
66
68
70
Position on chr4 (Mb)

| | |
|---|---|
| date: | Thu Oct 25 16:57:41 2018 |
| build: | F34F3943GBS |
| display range: | chr4:61866758−71866758 [61866758−71866758] |
| hilite range: | 0 − 0 [ 0 − 0 ] |
| reference SNP: | chr4:66866758 |
| number of SNPs plotted: | 176 |
| min P−value: | 1.64E−9 [chr4:66866758] |
| max P−value: | 9.86E−1 [chr4:63465549] |
| omitted Genes: | Slc31a2, Fkbp15, Slc31a1 |
| omitted Genes: | Cdc26, Prpf4, Rnf183 |
| omitted Genes: | Wdr31, Bspry, Hdhd3 |
| omitted Genes: | Alad, Pole3, 4933430I17Rik |
| omitted Genes: | Rgs3, Mir455, Orm1 |
| omitted Genes: | Orm3, Orm2, Akna |
| omitted Genes: | Whrn, Atp6v1g1, 6330416G13Rik |
| omitted Genes: | Gm11213, 1700018C11Rik, Tnfsf15 |

Make more plots at http://csg.sph.umich.edu/locuszoom/

Fine−mapping Regions

| snp | chr | pos (Mb) | pp | group | color |
|---|---|---|---|---|---|
| chr4:65447841 | 4 | 65.44784 | 0.019506224 | Credible Set | purple |
| chr4:65806420 | 4 | 65.80642 | 0.049316694 | Credible Set | purple |
| chr4:66002530 | 4 | 66.00253 | 0.007070884 | Credible Set | purple |
| chr4:66003029 | 4 | 66.00303 | 0.005898172 | Credible Set | purple |
| chr4:66003164 | 4 | 66.00316 | 0.006730268 | Credible Set | purple |
| chr4:66013512 | 4 | 66.01351 | 0.007172830 | Credible Set | purple |
| chr4:66110072 | 4 | 66.11007 | 0.013600870 | Credible Set | purple |
| chr4:66114163 | 4 | 66.11416 | 0.012017017 | Credible Set | purple |
| chr4:66114164 | 4 | 66.11416 | 0.011762290 | Credible Set | purple |
| chr4:66114176 | 4 | 66.11418 | 0.011842761 | Credible Set | purple |
| chr4:66123765 | 4 | 66.12377 | 0.021752254 | Credible Set | purple |
| chr4:66123768 | 4 | 66.12377 | 0.016580614 | Credible Set | purple |
| chr4:66123832 | 4 | 66.12383 | 0.025581610 | Credible Set | purple |
| chr4:66124242 | 4 | 66.12424 | 0.022728535 | Credible Set | purple |
| chr4:66169378 | 4 | 66.16938 | 0.013474615 | Credible Set | purple |
| chr4:66325857 | 4 | 66.32586 | 0.019241218 | Credible Set | purple |
| chr4:66325987 | 4 | 66.32599 | 0.017557542 | Credible Set | purple |
| chr4:66328189 | 4 | 66.32819 | 0.019746700 | Credible Set | purple |
| chr4:66328195 | 4 | 66.32819 | 0.017021073 | Credible Set | purple |
| chr4:66328436 | 4 | 66.32844 | 0.022182463 | Credible Set | purple |
| chr4:66328528 | 4 | 66.32853 | 0.024302960 | Credible Set | purple |
| chr4:66328560 | 4 | 66.32856 | 0.023547948 | Credible Set | purple |
| chr4:66329044 | 4 | 66.32904 | 0.027714988 | Credible Set | purple |
| chr4:66329065 | 4 | 66.32906 | 0.026859513 | Credible Set | purple |
| chr4:66329095 | 4 | 66.32909 | 0.027130315 | Credible Set | purple |
| chr4:66414332 | 4 | 66.41433 | 0.009959476 | Credible Set | purple |
| chr4:66414345 | 4 | 66.41434 | 0.010084430 | Credible Set | purple |
| chr4:66414373 | 4 | 66.41437 | 0.009806950 | Credible Set | purple |
| chr4:66414374 | 4 | 66.41437 | 0.009901791 | Credible Set | purple |
| chr4:66414384 | 4 | 66.41438 | 0.012715423 | Credible Set | purple |
| chr4:66414387 | 4 | 66.41439 | 0.016308798 | Credible Set | purple |
| chr4:66414451 | 4 | 66.41445 | 0.017072865 | Credible Set | purple |
| chr4:66414476 | 4 | 66.41448 | 0.036777925 | Credible Set | purple |
| chr4:66414508 | 4 | 66.41451 | 0.040273642 | Credible Set | purple |
| chr4:66414513 | 4 | 66.41451 | 0.038034578 | Credible Set | purple |
| chr4:66414963 | 4 | 66.41496 | 0.015171452 | Credible Set | purple |

Fine−mapping Regions

| snp | chr | pos (Mb) | pp | group | color |
|---|---|---|---|---|---|
| chr4:66419064 | 4 | 66.41906 | 0.009391583 | Credible Set | purple |
| chr4:66419126 | 4 | 66.41913 | 0.012358438 | Credible Set | purple |
| chr4:66743103 | 4 | 66.74310 | 0.036342646 | Credible Set | purple |
| chr4:66823255 | 4 | 66.82326 | 0.037502254 | Credible Set | purple |
| chr4:66840696 | 4 | 66.84070 | 0.052214273 | Credible Set | purple |
| chr4:66866758 | 4 | 66.86676 | 0.071397622 | Credible Set | purple |
| chr4:67163007 | 4 | 67.16301 | 0.041268735 | Credible Set | purple |
| chr4:67165680 | 4 | 67.16568 | 0.041934791 | Credible Set | purple |
| chr4:67188375 | 4 | 67.18837 | 0.006611697 | Credible Set | purple |

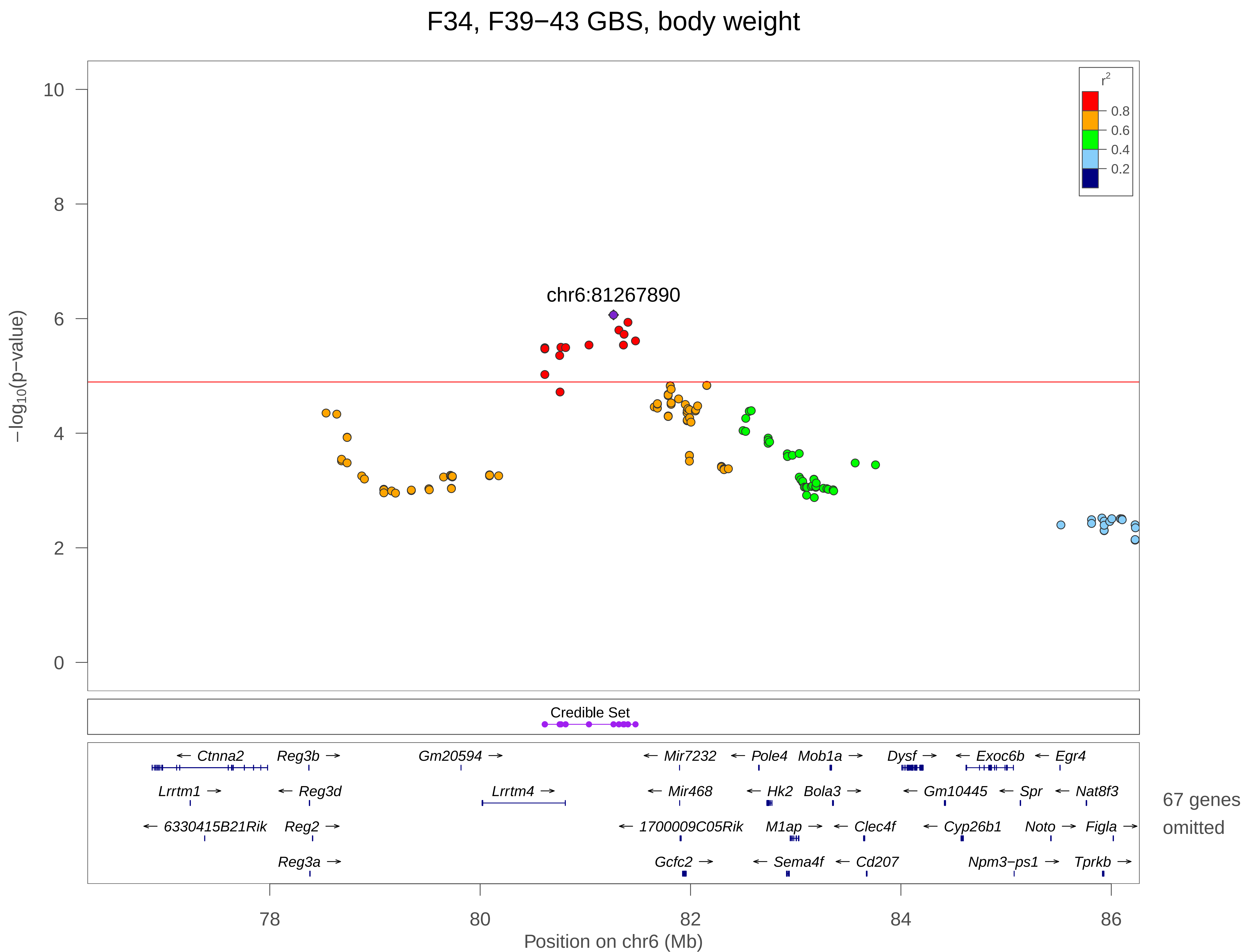

F34, F39−43 GBS, body weight
−log10(p−value)
10
8
6
4
2
0
chr6:81267890
r2
0.8
0.6
0.4
0.2
Credible Set
← Ctnna2
Reg3b →
Gm20594 →
← Mir7232
← Pole4
Mob1a →
Dysf →
← Exoc6b
← Egr4
Lrrtm1 →
← Reg3d
Lrrtm4 →
← Mir468
← Hk2
Bola3 →
← Gm10445
← Spr
← Nat8f3
← 6330415B21Rik
Reg2 →
← 1700009C05Rik
M1ap →
← Clec4f
← Cyp26b1
Noto →
Figla →
Reg3a →
Gcfc2 →
← Sema4f
← Cd207
Npm3−ps1 →
Tprkb →
67 genes omitted
78
80
82
84
86
Position on chr6 (Mb)

| | |
|---|---|
| date: | Thu Oct 25 16:57:51 2018 |
| build: | F34F3943GBS |
| display range: | chr6:76267890−86267890 [76267890−86267890] |
| hilite range: | 0 − 0 [ 0 − 0 ] |
| reference SNP: | chr6:81267890 |
| number of SNPs plotted: | 154 |
| min P−value: | 8.61E−7 [chr6:81267890] |
| max P−value: | 7.33E−3 [chr6:86226371] |
| omitted Genes: | Reg1, Reg3g, Gm20362 |
| omitted Genes: | Mrpl19, Eva1a, Tacr1 |
| omitted Genes: | 2310069B03Rik, Dok1, Loxl3 |
| omitted Genes: | Mir7040, Htra2, Aup1 |
| omitted Genes: | Dqx1, Tlx2, Pcgf1 |
| omitted Genes: | Lbx2, Mir3470a, Ccdc142 |
| omitted Genes: | Mrpl53, Mogs, Wbp1 |
| omitted Genes: | Ino80b, Rtkn, Wdr54 |
| omitted Genes: | 1700003E16Rik, Dctn1, Slc4a5 |
| omitted Genes: | Mthfd2, Tet3, Mir6374 |
| omitted Genes: | B230319C09Rik, Dguok, Actg2 |
| omitted Genes: | Stambp, Gm21284, Vax2os |
| omitted Genes: | Vax2, Atp6v1b1, 1700124L16Rik |
| omitted Genes: | Ankrd53, Tex261, Gm7443 |
| omitted Genes: | Nagk, Paip2b, Zfp638 |
| omitted Genes: | Gm5878, Emx1, Sfxn5 |
| omitted Genes: | Rab11fip5, Mir705, Smyd5 |
| omitted Genes: | Pradc1, Cct7, Fbxo41 |
| omitted Genes: | Alms1, Nat8f7, Mir6375 |
| omitted Genes: | Alms1−ps2, Nat8f6, Nat8f5 |
| omitted Genes: | Nat8, Nat8f2, Nat8f4 |
| omitted Genes: | Nat8f1, Dusp11, Add2 |
| omitted Genes: | Tgfa |

Make more plots at http://csg.sph.umich.edu/locuszoom/

Fine−mapping Regions

| snp | chr | pos (Mb) | pp | group | color |
|---|---|---|---|---|---|
| chr6:80614710 | 6 | 80.61471 | 0.04857390 | Credible Set | purple |
| chr6:80614725 | 6 | 80.61473 | 0.04623828 | Credible Set | purple |
| chr6:80614740 | 6 | 80.61474 | 0.04722318 | Credible Set | purple |
| chr6:80614750 | 6 | 80.61475 | 0.04694999 | Credible Set | purple |
| chr6:80614907 | 6 | 80.61491 | 0.01733174 | Credible Set | purple |
| chr6:80755939 | 6 | 80.75594 | 0.03596034 | Credible Set | purple |
| chr6:80767933 | 6 | 80.76793 | 0.04944593 | Credible Set | purple |
| chr6:80812582 | 6 | 80.81258 | 0.04883514 | Credible Set | purple |
| chr6:81034488 | 6 | 81.03449 | 0.05391742 | Credible Set | purple |
| chr6:81267890 | 6 | 81.26789 | 0.17255878 | Credible Set | purple |
| chr6:81318989 | 6 | 81.31899 | 0.09591536 | Credible Set | purple |
| chr6:81362217 | 6 | 81.36222 | 0.05384616 | Credible Set | purple |
| chr6:81368112 | 6 | 81.36811 | 0.08156347 | Credible Set | purple |
| chr6:81405109 | 6 | 81.40511 | 0.12958790 | Credible Set | purple |
| chr6:81476902 | 6 | 81.47690 | 0.06318092 | Credible Set | purple |

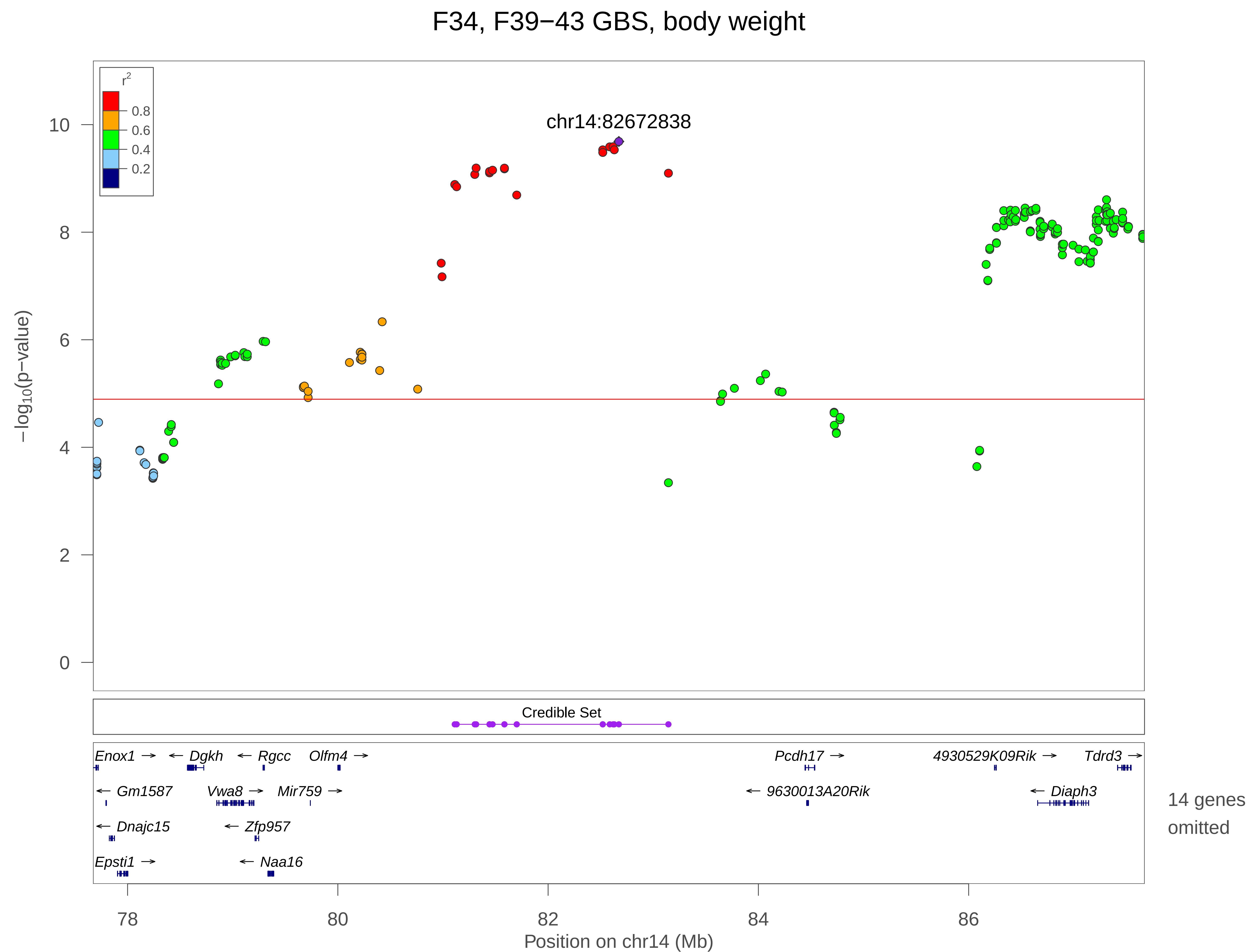

F34, F39−43 GBS, body weight
r2
0.8
0.6
0.4
0.2
chr14:82672838
10
8
6
4
2
0
−log10(p−value)
Credible Set
Enox1 →
← Dgkh
← Rgcc
Olfm4 →
Pcdh17 →
4930529K09Rik →
Tdrd3 →
← Gm1587
Vwa8 →
Mir759 →
← 9630013A20Rik
← Diaph3
← Dnajc15
← Zfp957
Epsti1 →
← Naa16
14 genes omitted
78
80
82
84
86
Position on chr14 (Mb)

| | |
|---|---|
| date: | Thu Oct 25 16:57:18 2018 |
| build: | F34F3943GBS |
| display range: | chr14:77672838−87672838 [77672838−87672838] |
| hilite range: | 0 − 0 [ 0 − 0 ] |
| reference SNP: | chr14:82672838 |
| number of SNPs plotted: | 233 |
| min P−value: | 2.06E−10 [chr14:82672838] |
| max P−value: | 4.55E−4 [chr14:83144370] |
| omitted Genes: | Fam216b, Mir1971, Tnfsf11 |
| omitted Genes: | Akap11, Mtrf1, Kbtbd7 |
| omitted Genes: | Kbtbd6, Wbp4, Elf1 |
| omitted Genes: | Sugt1, Lect1, 4930452G13Rik |
| omitted Genes: | Pcdh8, Gm10845 |

Make more plots at http://csg.sph.umich.edu/locuszoom/

Fine−mapping Regions

| snp | chr | pos (Mb) | pp | group | color |
|---|---|---|---|---|---|
| chr14:81112053 | 14 | 81.11205 | 0.02153824 | Credible Set | purple |
| chr14:81129931 | 14 | 81.12993 | 0.01961019 | Credible Set | purple |
| chr14:81302522 | 14 | 81.30252 | 0.03271591 | Credible Set | purple |
| chr14:81315250 | 14 | 81.31525 | 0.04283704 | Credible Set | purple |
| chr14:81443053 | 14 | 81.44305 | 0.03518459 | Credible Set | purple |
| chr14:81443088 | 14 | 81.44309 | 0.03495130 | Credible Set | purple |
| chr14:81443103 | 14 | 81.44310 | 0.03705697 | Credible Set | purple |
| chr14:81471656 | 14 | 81.47166 | 0.03910768 | Credible Set | purple |
| chr14:81584557 | 14 | 81.58456 | 0.04138683 | Credible Set | purple |
| chr14:81584584 | 14 | 81.58458 | 0.04266022 | Credible Set | purple |
| chr14:81701500 | 14 | 81.70150 | 0.01386607 | Credible Set | purple |
| chr14:82519671 | 14 | 82.51967 | 0.09157956 | Credible Set | purple |
| chr14:82519834 | 14 | 82.51983 | 0.08209422 | Credible Set | purple |
| chr14:82586326 | 14 | 82.58633 | 0.10379728 | Credible Set | purple |
| chr14:82618916 | 14 | 82.61892 | 0.10439366 | Credible Set | purple |
| chr14:82629334 | 14 | 82.62933 | 0.09166977 | Credible Set | purple |
| chr14:82672838 | 14 | 82.67284 | 0.12975387 | Credible Set | purple |
| chr14:83144282 | 14 | 83.14428 | 0.03452068 | Credible Set | purple |

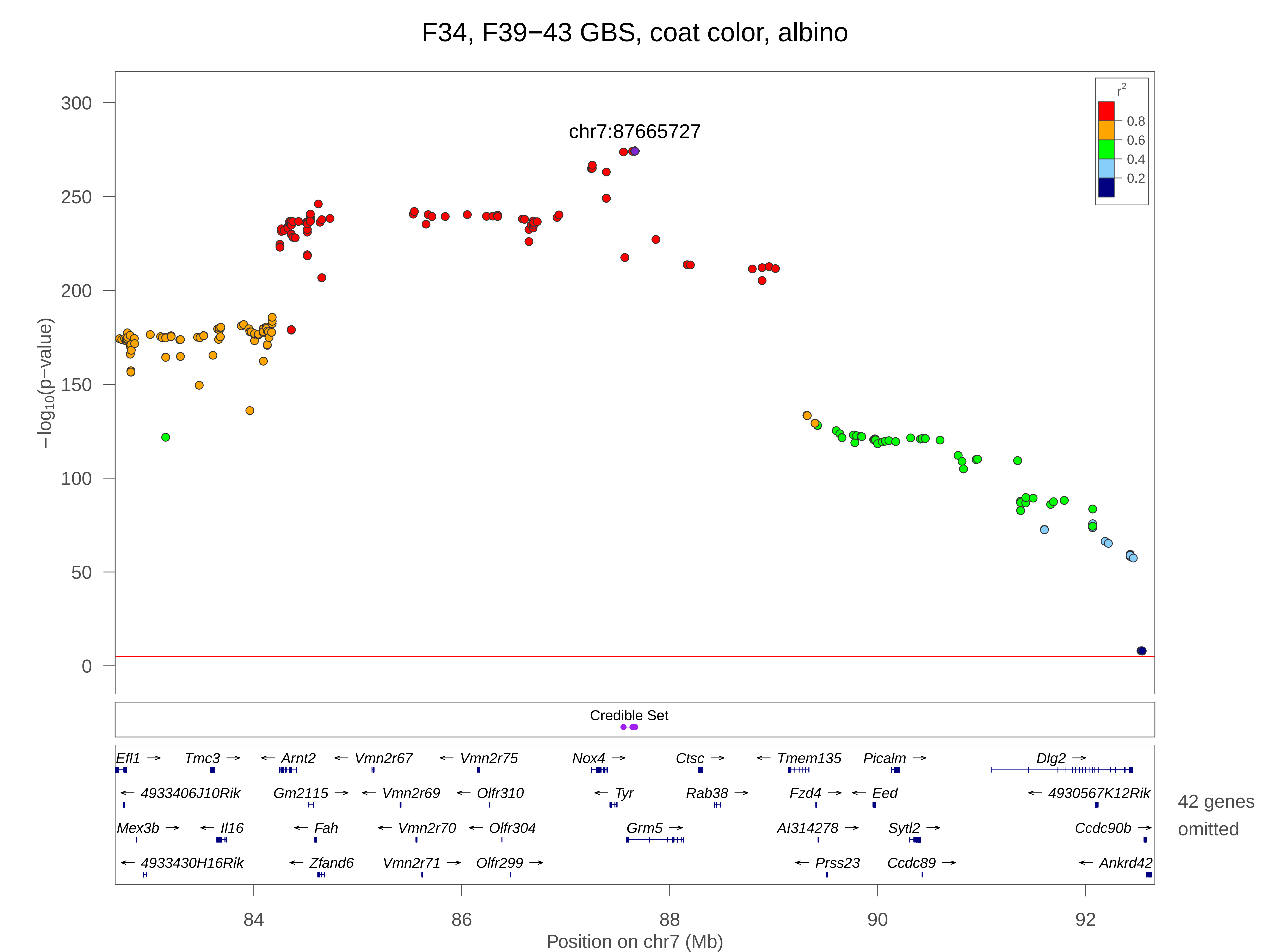

F34, F39−43 GBS, coat color, albino
chr7:87665727
r2
0.8
0.6
0.4
0.2
300
250
200
150
100
50
0
−log10(p−value)
Credible Set
Efl1 →
Tmc3 →
← Arnt2
← Vmn2r67
← Vmn2r75
Nox4 →
Ctsc →
← Tmem135
Picalm →
Dlg2 →
← 4933406J10Rik
Gm2115 →
← Vmn2r69
← Olfr310
← Tyr
Rab38 →
Fzd4 →
← Eed
← 4930567K12Rik
Mex3b →
← Il16
← Fah
← Vmn2r70
← Olfr304
Grm5 →
AI314278 →
Sytl2 →
Ccdc90b →
← 4933430H16Rik
← Zfand6
Vmn2r71 →
Olfr299 →
← Prss23
Ccdc89 →
← Ankrd42
42 genes
omitted
84
86
88
90
92
Position on chr7 (Mb)

| | |
|---|---|
| date: | Thu Oct 25 16:58:20 2018 |
| build: | F34F3943GBS |
| display range: | chr7:82665727−92665727 [82665727−92665727] |
| hilite range: | 0 − 0 [ 0 − 0 ] |
| reference SNP: | chr7:87665727 |
| number of SNPs plotted: | 274 |
| min P−value: | 6.71E−275 [chr7:87665727] |
| max P−value: | 1.33E−8 [chr7:92543164] |
| omitted Genes: | Stard5, Cfap161, Mesdc1 |
| omitted Genes: | Mesdc2, Cemip, Abhd17c |
| omitted Genes: | 2610206C17Rik, Olfr291, Olfr290 |
| omitted Genes: | Vmn2r65, Vmn2r66, Vmn2r68 |
| omitted Genes: | Vmn2r72, Vmn2r73, Vmn2r74 |
| omitted Genes: | Vmn2r76, Olfr309, Olfr308 |
| omitted Genes: | Olfr307, Olfr305, Olfr303 |
| omitted Genes: | Olfr301, Olfr298, Olfr297 |
| omitted Genes: | Olfr295, Olfr294, Olfr293 |
| omitted Genes: | Olfr292, Folh1, Vmn2r77 |
| omitted Genes: | Vmn2r78, Vmn2r79, Me3 |
| omitted Genes: | Ccdc81, I7Rn6, E230029C05Rik |
| omitted Genes: | 2310010J17Rik, Ccdc83, Crebzf |
| omitted Genes: | Tmem126a, Tmem126b, Pcf11 |

Make more plots at http://csg.sph.umich.edu/locuszoom/

Fine−mapping Regions

| snp | chr | pos (Mb) | pp | group | color |
|---|---|---|---|---|---|
| chr7:87554961 | 7 | 87.55496 | 0.1583967 | Credible Set | purple |
| chr7:87642045 | 7 | 87.64204 | 0.3914069 | Credible Set | purple |
| chr7:87665727 | 7 | 87.66573 | 0.4501964 | Credible Set | purple |

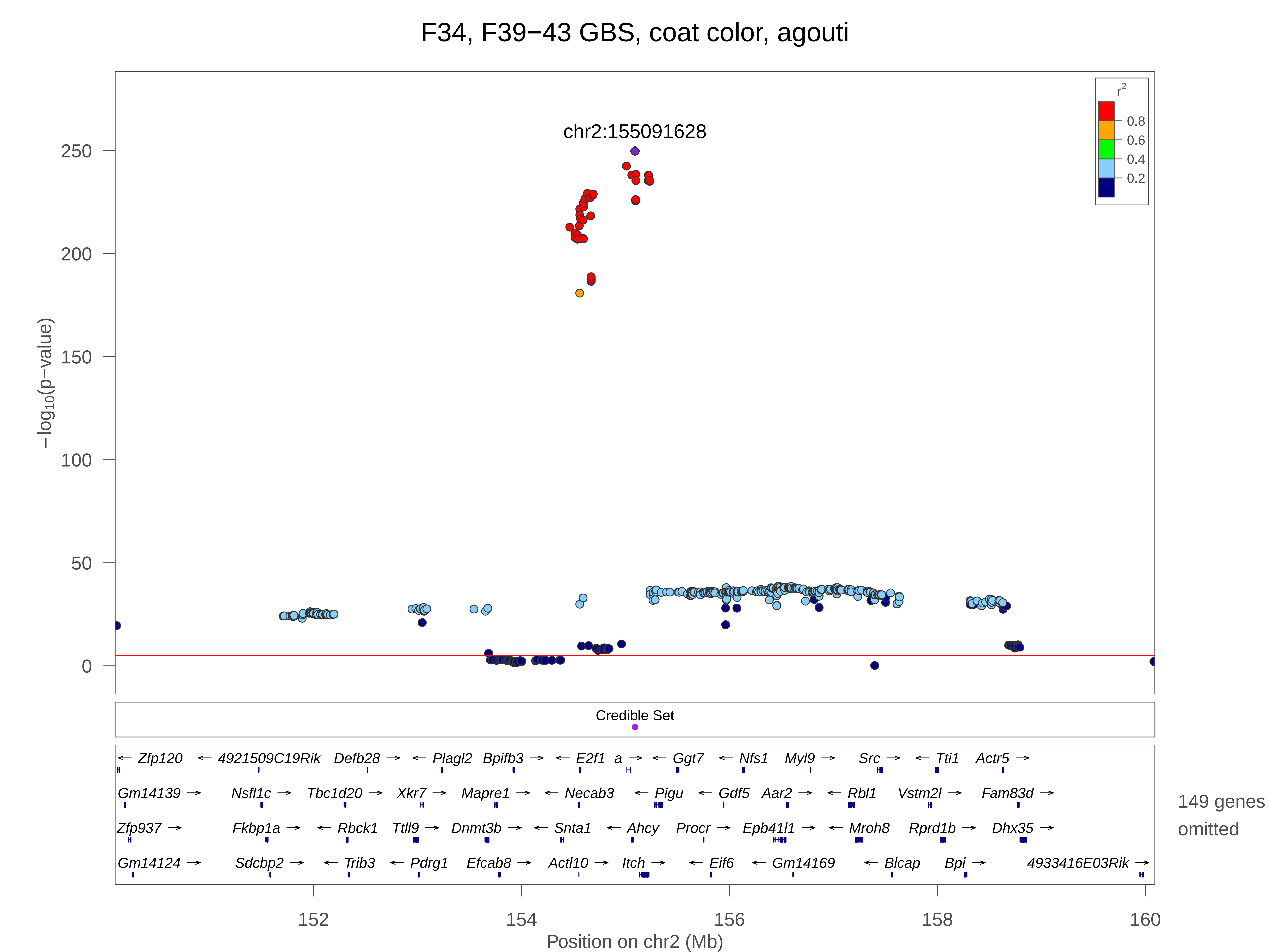
F34, F39−43 GBS, coat color, agouti
chr2:155091628
r2
0.8
0.6
0.4
0.2
250
200
150
100
50
0
−log10(p−value)
Credible Set
← Zfp120
← 4921509C19Rik
Defb28 →
← Plagl2
Bpifb3 →
← E2f1
a →
← Ggt7
← Nfs1
Myl9 →
Src →
← Tti1
Actr5 →
Gm14139 →
Nsfl1c →
Tbc1d20 →
Xkr7 →
Mapre1 →
← Necab3
← Pigu
← Gdf5
Aar2 →
← Rbl1
Vstm2l →
Fam83d →
Zfp937 →
Fkbp1a →
← Rbck1
Ttll9 →
Dnmt3b →
← Snta1
← Ahcy
Procr →
Epb41l1 →
← Mroh8
Rprd1b →
Dhx35 →
Gm14124 →
Sdcbp2 →
← Trib3
← Pdrg1
Efcab8 →
Actl10 →
Itch →
← Eif6
← Gm14169
← Blcap
Bpi →
4933416E03Rik →
149 genes omitted
152
154
156
158
160
Position on chr2 (Mb)

date: Thu Oct 25 16:59:07 2018

build: F34F3943GBS

display range: chr2:150091628−160091628 [150091628−160091628]

hilite range: 0 − 0 [ 0 − 0 ]

reference SNP: chr2:155091628

number of SNPs plotted: 747

min P−value: 1.88E−250 [chr2:155091628]

max P−value: 6.98E−1 [chr2:157396490]

omitted Genes: 3300002I08Rik, Zfp442, Zfp345

omitted Genes: Cst7, Apmap, Acss1

omitted Genes: E130215H24Rik, Vsx1, Entpd6

omitted Genes: Pygb, Abhd12, Gins1

omitted Genes: Ninl, Nanp, Gm14151

omitted Genes: 4930442J19Rik, LOC100125594, Snph

omitted Genes: Rad21l, Tmem74b, Psmf1

omitted Genes: Rspo4, Angpt4, Mir1953

omitted Genes: Fam110a, Slc52a3, Scrt2

omitted Genes: Srxn1, Tcf15, Csnk2a1

omitted Genes: Gm14164, Nrsn2, Sox12

omitted Genes: Zcchc3, 6820408C15Rik, Defb23

omitted Genes: Defb20, Defb22, Defb26

omitted Genes: Defb29, Defb21, Defb19

omitted Genes: Defb45, Defb36, Defb25

omitted Genes: Rem1, H13, Mcts2

omitted Genes: Id1, Cox4i2, Bcl2l1

omitted Genes: Tpx2, Mir5622, Mylk2

omitted Genes: Foxs1, Dusp15, Ccm2l

omitted Genes: Hck, Tm9sf4, Tspyl3

omitted Genes: Pofut1, Kif3b, 2500004C02Rik

omitted Genes: Asxl1, Nol4l, Commd7

omitted Genes: Sun5, Bpifb2, Bpifb6

omitted Genes: Bpifb4, Bpifa6, Bpifa2

omitted Genes: Bpifa3, Bpifa1, Bpifa5

omitted Genes: Bpifb1, Bpifb5, Bpifb9a

omitted Genes: Bpifb9b, Cdk5rap1, Cbfa2t2

omitted Genes: 1700003F12Rik, Pxmp4, Zfp341

omitted Genes: Chmp4b, Raly, Eif2s2

omitted Genes: 2310005A03Rik, Dynlrb1, Map1lc3a

omitted Genes: Mir695, Trp53inp2, Ncoa6

Make more plots at http://csg.sph.umich.edu/locuszoom/

omitted Genes: Acss2, Acss2os, Gss

Fine−mapping Regions

| snp | chr | pos (Mb) | pp | group | color |
|---|---|---|---|---|---|
| chr2:155091628 | 2 | 155.0916 | 0.9999999 | Credible Set | purple |

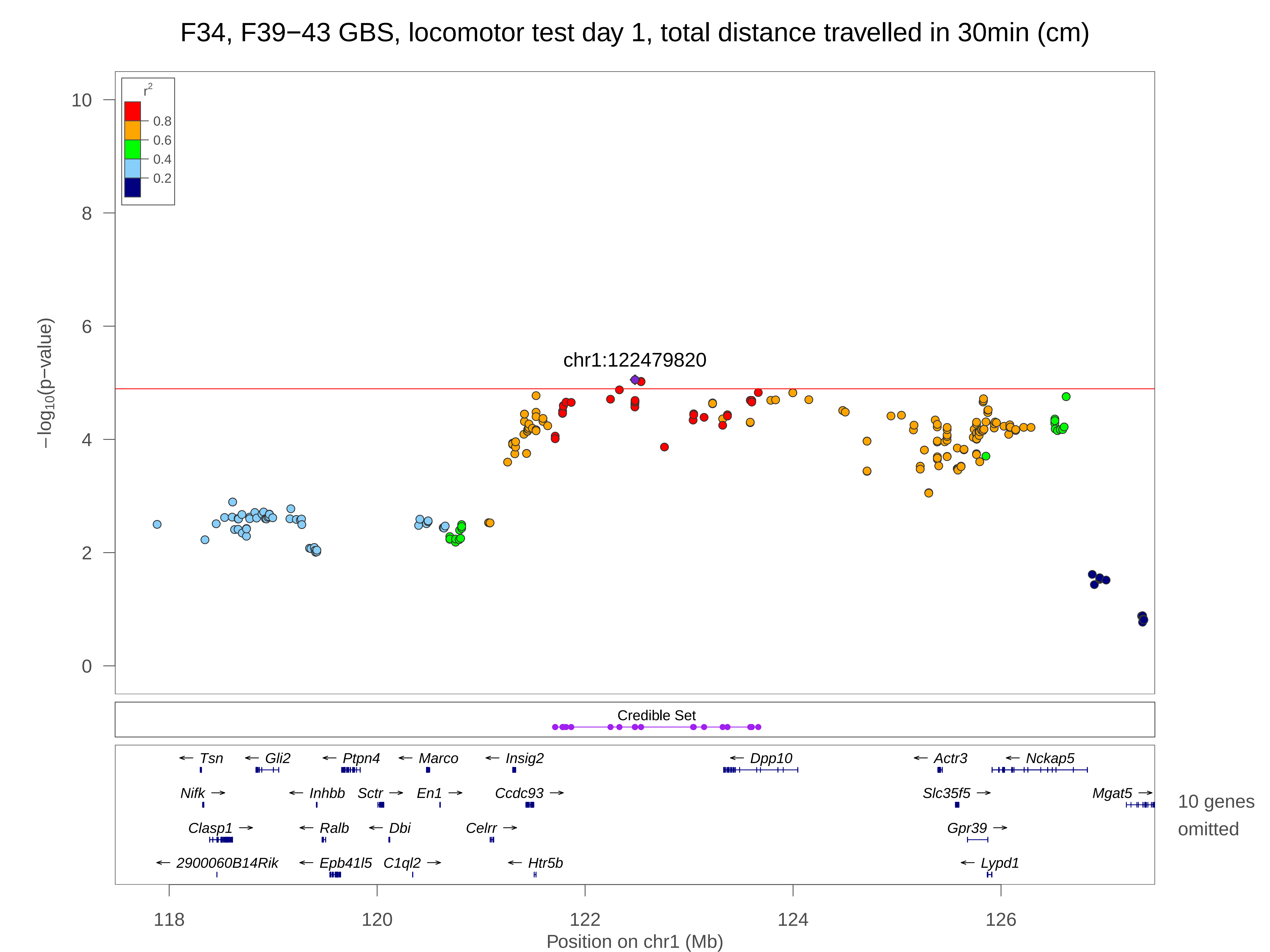
F34, F39−43 GBS, locomotor test day 1, total distance travelled in 30min (cm)
r2
0.8
0.6
0.4
0.2
10
8
6
4
2
0
−log10(p−value)
chr1:122479820
Credible Set
← Tsn
← Gli2
← Ptpn4
← Marco
← Insig2
← Dpp10
← Actr3
← Nckap5
Nifk →
← Inhbb
Sctr →
En1 →
Ccdc93 →
Slc35f5 →
Mgat5 →
Clasp1 →
← Ralb
← Dbi
Celrr →
Gpr39 →
← 2900060B14Rik
← Epb41l5
C1ql2 →
← Htr5b
← Lypd1
10 genes omitted
118
120
122
124
126
Position on chr1 (Mb)

| | |
|---|---|
| date: | Thu Oct 25 16:59:22 2018 |
| build: | F34F3943GBS |
| display range: | chr1:117479820−127479820 [117479820−127479820] |
| hilite range: | 0 − 0 [ 0 − 0 ] |
| reference SNP: | chr1:122479820 |
| number of SNPs plotted: | 268 |
| min P−value: | 8.84E−6 [chr1:122479820] |
| max P−value: | 1.69E−1 [chr1:127361132] |
| omitted Genes: | Tfcp2l1, Mir6346, Tmem185b |
| omitted Genes: | 3830432H09Rik, Tmem177, Cfap221 |
| omitted Genes: | Tmem37, 3110009E18Rik, Steap3 |
| omitted Genes: | Ddx18 |

Make more plots at http://csg.sph.umich.edu/locuszoom/

Fine−mapping Regions

| snp | chr | pos (Mb) | pp | group | color |
|---|---|---|---|---|---|
| chr1:121712291 | 1 | 121.7123 | 0.008285125 | Credible Set | purple |
| chr1:121712326 | 1 | 121.7123 | 0.009119169 | Credible Set | purple |
| chr1:121712327 | 1 | 121.7123 | 0.008217150 | Credible Set | purple |
| chr1:121783108 | 1 | 121.7831 | 0.021818801 | Credible Set | purple |
| chr1:121783777 | 1 | 121.7838 | 0.023997232 | Credible Set | purple |
| chr1:121783797 | 1 | 121.7838 | 0.022021202 | Credible Set | purple |
| chr1:121790517 | 1 | 121.7905 | 0.029622818 | Credible Set | purple |
| chr1:121790555 | 1 | 121.7906 | 0.028975381 | Credible Set | purple |
| chr1:121817004 | 1 | 121.8170 | 0.033807004 | Credible Set | purple |
| chr1:121866576 | 1 | 121.8666 | 0.033392468 | Credible Set | purple |
| chr1:122244126 | 1 | 122.2441 | 0.037839710 | Credible Set | purple |
| chr1:122329939 | 1 | 122.3299 | 0.054349426 | Credible Set | purple |
| chr1:122478661 | 1 | 122.4787 | 0.031348898 | Credible Set | purple |
| chr1:122478665 | 1 | 122.4787 | 0.027951382 | Credible Set | purple |
| chr1:122479820 | 1 | 122.4798 | 0.080446064 | Credible Set | purple |
| chr1:122480027 | 1 | 122.4800 | 0.035461975 | Credible Set | purple |
| chr1:122480078 | 1 | 122.4801 | 0.034018495 | Credible Set | purple |
| chr1:122480093 | 1 | 122.4801 | 0.036042672 | Credible Set | purple |
| chr1:122537959 | 1 | 122.5380 | 0.075581861 | Credible Set | purple |
| chr1:122537965 | 1 | 122.5380 | 0.074489582 | Credible Set | purple |
| chr1:123038644 | 1 | 123.0386 | 0.016858419 | Credible Set | purple |
| chr1:123044323 | 1 | 123.0443 | 0.021494377 | Credible Set | purple |
| chr1:123044326 | 1 | 123.0443 | 0.020646374 | Credible Set | purple |
| chr1:123144261 | 1 | 123.1443 | 0.018770743 | Credible Set | purple |
| chr1:123322958 | 1 | 123.3230 | 0.013793158 | Credible Set | purple |
| chr1:123368825 | 1 | 123.3688 | 0.020709950 | Credible Set | purple |
| chr1:123369170 | 1 | 123.3692 | 0.019699584 | Credible Set | purple |
| chr1:123588436 | 1 | 123.5884 | 0.036196853 | Credible Set | purple |
| chr1:123602768 | 1 | 123.6028 | 0.036249469 | Credible Set | purple |
| chr1:123603027 | 1 | 123.6030 | 0.033855107 | Credible Set | purple |
| chr1:123665450 | 1 | 123.6655 | 0.048953134 | Credible Set | purple |

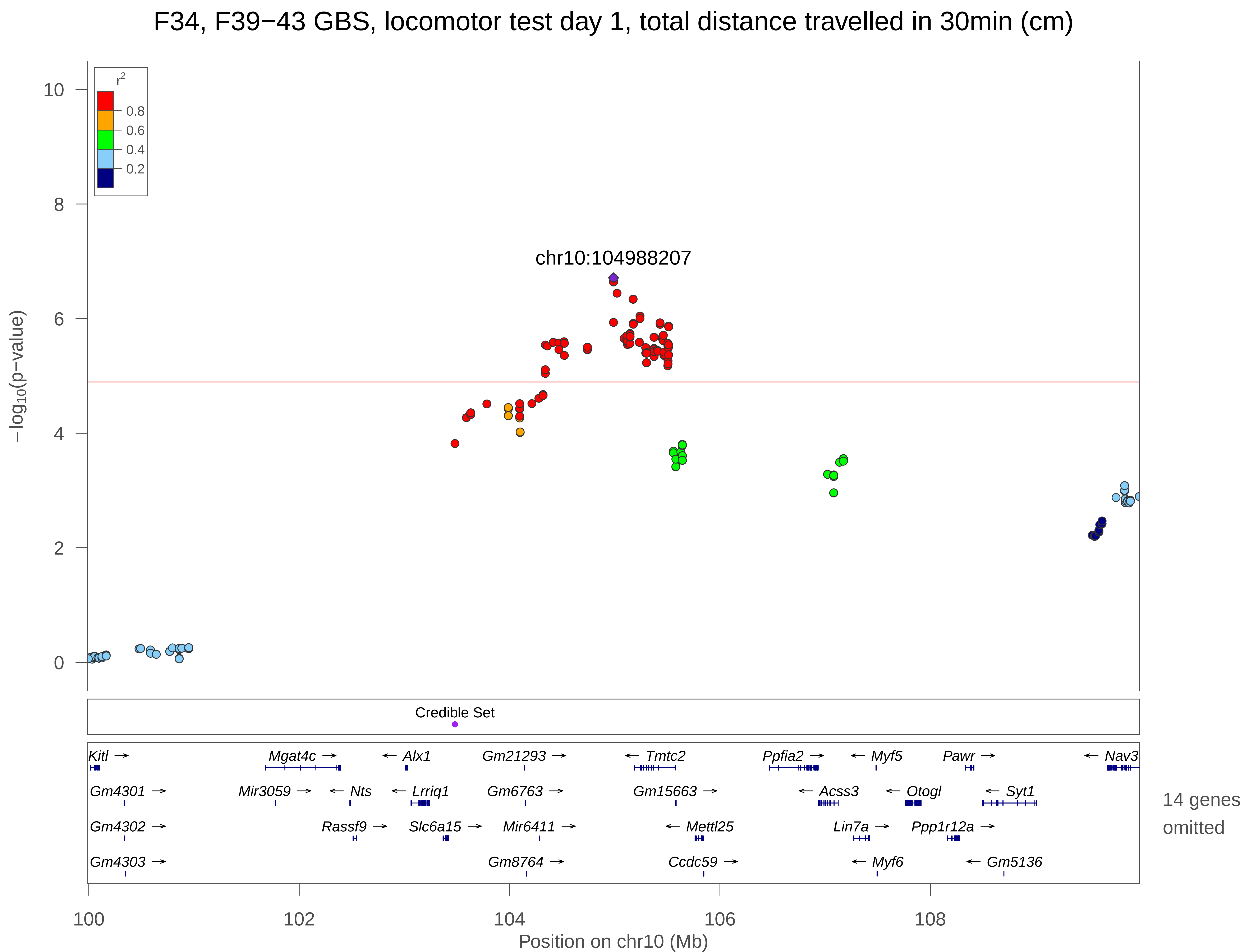
F34, F39−43 GBS, locomotor test day 1, total distance travelled in 30min (cm)
r2
0.8
0.6
0.4
0.2
10
8
6
4
2
0
−log10(p−value)
chr10:104988207
Credible Set
Kitl →
Mgat4c →
← Alx1
Gm21293 →
← Tmtc2
Ppfia2 →
← Myf5
Pawr →
← Nav3
Gm4301 →
Mir3059 →
← Nts
← Lrriq1
Gm6763 →
Gm15663 →
← Acss3
← Otogl
← Syt1
Gm4302 →
Rassf9 →
Slc6a15 →
Mir6411 →
← Mettl25
Lin7a →
Ppp1r12a →
Gm4303 →
Gm8764 →
Ccdc59 →
← Myf6
← Gm5136
14 genes omitted
100
102
104
106
108
Position on chr10 (Mb)

| | |
|---|---|
| date: | Thu Oct 25 16:59:15 2018 |
| build: | F34F3943GBS |
| display range: | chr10:99988207−109988207 [99988207−109988207] |
| hilite range: | 0 − 0 [ 0 − 0 ] |
| reference SNP: | chr10:104988207 |
| number of SNPs plotted: | 216 |
| min P−value: | 1.94E−7 [chr10:104988207] |
| max P−value: | 8.78E−1 [chr10:100031691] |
| omitted Genes: | Gm4305, Gm4307, Gm4312 |
| omitted Genes: | Tmtc3, Cep290, 4930430F08Rik |
| omitted Genes: | 1700017N19Rik, Gm21304, Gm21312 |
| omitted Genes: | Gm20765, Gm4340, 4930532I03Rik |
| omitted Genes: | Ptprq, Gm29685 |

Make more plots at http://csg.sph.umich.edu/locuszoom/

Fine−mapping Regions

| snp | chr | pos (Mb) | pp | group | color |
|---|---|---|---|---|---|
| chr10:103480830 | 10 | 103.4808 | 0 | Credible Set | purple |

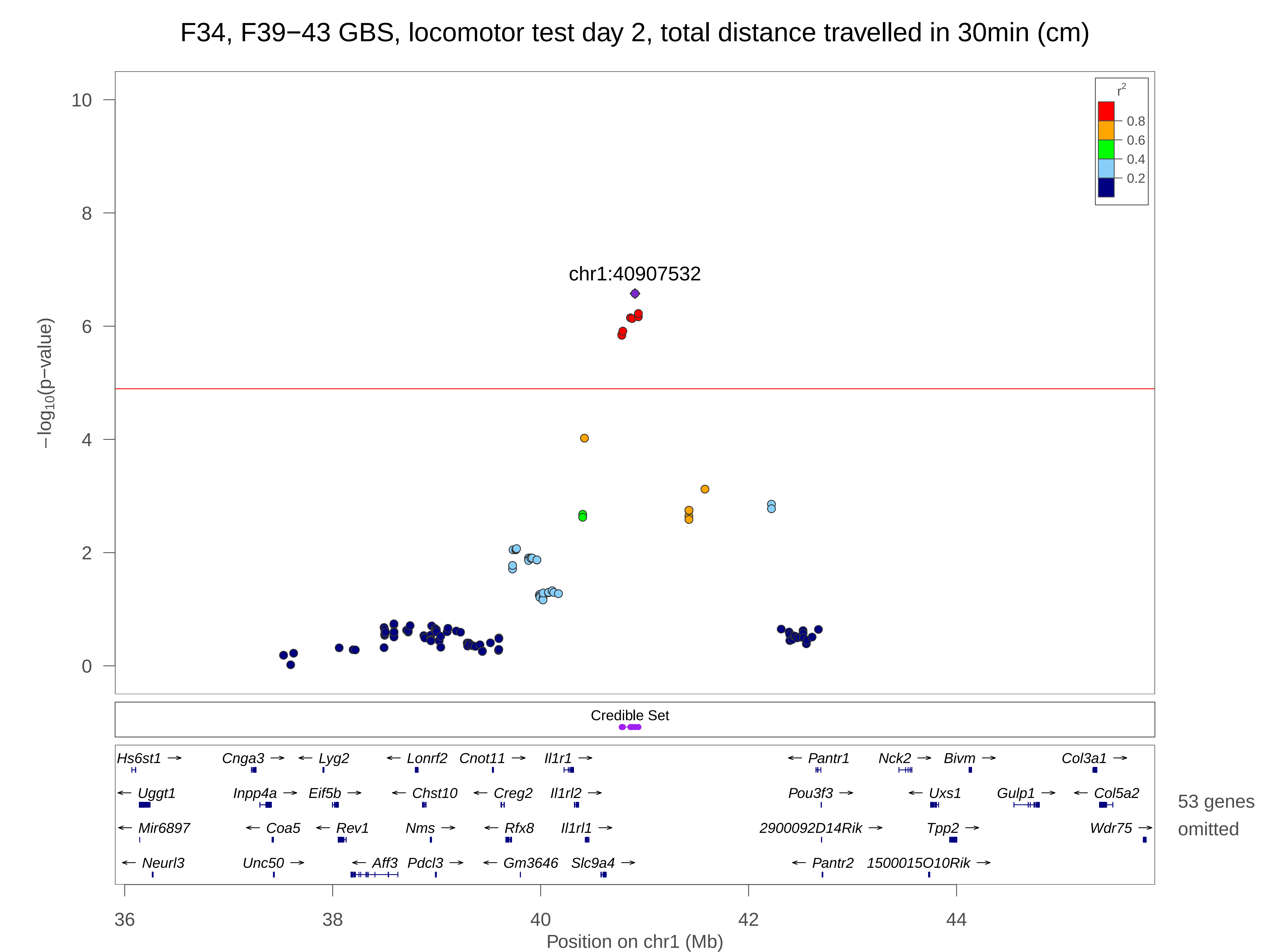
F34, F39−43 GBS, locomotor test day 2, total distance travelled in 30min (cm)
−log10(p−value)
10
8
6
4
2
0
chr1:40907532
r2
0.8
0.6
0.4
0.2
Credible Set
Hs6st1 →
Cnga3 →
← Lyg2
← Lonrf2
Cnot11 →
Il1r1 →
← Pantr1
Nck2 →
Bivm →
Col3a1 →
← Uggt1
Inpp4a →
Eif5b →
← Chst10
← Creg2
Il1rl2 →
Pou3f3 →
← Uxs1
Gulp1 →
← Col5a2
← Mir6897
← Coa5
← Rev1
Nms →
← Rfx8
Il1rl1 →
2900092D14Rik →
Tpp2 →
Wdr75 →
← Neurl3
Unc50 →
← Aff3
Pdcl3 →
← Gm3646
Slc9a4 →
← Pantr2
1500015O10Rik →
53 genes omitted
36
38
40
42
44
Position on chr1 (Mb)

| | |
|---|---|
| date: | Thu Oct 25 16:59:39 2018 |
| build: | F34F3943GBS |
| display range: | chr1:35907532−45907532 [35907532−45907532] |
| hilite range: | 0 − 0 [ 0 − 0 ] |
| reference SNP: | chr1:40907532 |
| number of SNPs plotted: | 157 |
| min P−value: | 2.65E−7 [chr1:40907532] |
| max P−value: | 9.56E−1 [chr1:37596381] |
| omitted Genes: | Arid5a, Kansl3, Mir6898 |
| omitted Genes: | Fer1l5, Lman2l, Cnnm4 |
| omitted Genes: | Cnnm3, Ankrd23, Ankrd39 |
| omitted Genes: | Sema4c, Fam178b, Cox5b |
| omitted Genes: | Actr1b, 4933424G06Rik, Zap70 |
| omitted Genes: | Tmem131, Mgat4a, 4930594C11Rik |
| omitted Genes: | 2010300C02Rik, Tsga10, Lipt1 |
| omitted Genes: | Mitd1, Mrpl30, Lyg1 |
| omitted Genes: | Txndc9, Mir6349, Npas2 |
| omitted Genes: | Rpl31, Tbc1d8, Snord89 |
| omitted Genes: | Rnf149, 1700066B17Rik, Map4k4 |
| omitted Genes: | Gm16894, Il1r2, Il18r1 |
| omitted Genes: | Il18rap, Slc9a2, Mfsd9 |
| omitted Genes: | Tmem182, 4930448I06Rik, Dalir |
| omitted Genes: | Mrps9, Gpr45, Tgfbrap1 |
| omitted Genes: | AI597479, Fhl2, Mettl21c |
| omitted Genes: | Tex30, Kdelc1, Ercc5 |
| omitted Genes: | Mettl21e, 4930521E06Rik |

Make more plots at http://csg.sph.umich.edu/locuszoom/

Fine−mapping Regions

| snp | chr | pos (Mb) | pp | group | color |
|---|---|---|---|---|---|
| chr1:40780351 | 1 | 40.78035 | 0.06182924 | Credible Set | purple |
| chr1:40780677 | 1 | 40.78068 | 0.05996952 | Credible Set | purple |
| chr1:40790450 | 1 | 40.79045 | 0.07076768 | Credible Set | purple |
| chr1:40863930 | 1 | 40.86393 | 0.11903013 | Credible Set | purple |
| chr1:40880208 | 1 | 40.88021 | 0.11549377 | Credible Set | purple |
| chr1:40907532 | 1 | 40.90753 | 0.30844609 | Credible Set | purple |
| chr1:40939551 | 1 | 40.93955 | 0.12347079 | Credible Set | purple |
| chr1:40939963 | 1 | 40.93996 | 0.14099278 | Credible Set | purple |

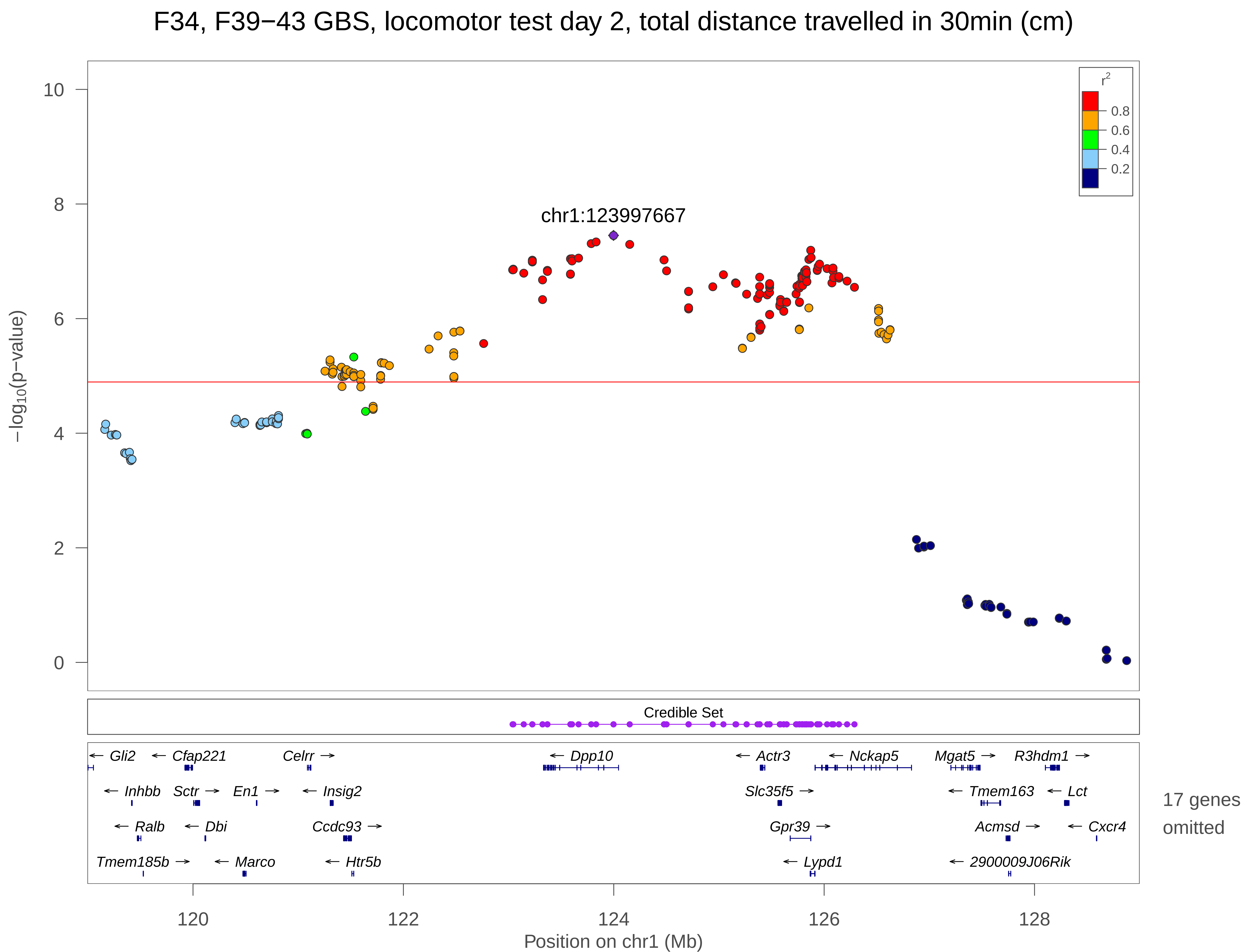
F34, F39−43 GBS, locomotor test day 2, total distance travelled in 30min (cm)
chr1:123997667
r2
0.8
0.6
0.4
0.2
−log10(p−value)
10
8
6
4
2
0
Credible Set
Gli2
Cfap221
Celrr
Dpp10
Actr3
Nckap5
Mgat5
R3hdm1
Inhbb
Sctr
En1
Insig2
Slc35f5
Tmem163
Lct
Ralb
Dbi
Ccdc93
Gpr39
Acmsd
Cxcr4
Tmem185b
Marco
Htr5b
Lypd1
2900009J06Rik
17 genes omitted
120
122
124
126
128
Position on chr1 (Mb)

| | |
|---|---|
| date: | Thu Oct 25 16:59:30 2018 |
| build: | F34F3943GBS |
| display range: | chr1:118997667−128997667 [118997667−128997667] |
| hilite range: | 0 − 0 [ 0 − 0 ] |
| reference SNP: | chr1:123997667 |
| number of SNPs plotted: | 260 |
| min P−value: | 3.53E−8 [chr1:123997667] |
| max P−value: | 9.31E−1 [chr1:128876034] |
| omitted Genes: | 3830432H09Rik, Epb41l5, Ptpn4 |
| omitted Genes: | Tmem177, Tmem37, 3110009E18Rik |
| omitted Genes: | Steap3, C1ql2, Ddx18 |
| omitted Genes: | Ccnt2, Map3k19, Rab3gap1 |
| omitted Genes: | Zranb3, Mir128−1, Ubxn4 |
| omitted Genes: | Mcm6, Dars |

Make more plots at http://csg.sph.umich.edu/locuszoom/

Fine−mapping Regions

| snp | chr | pos (Mb) | pp | group | color |
|---|---|---|---|---|---|
| chr1:123038644 | 1 | 123.0386 | 0.010936661 | Credible Set | purple |
| chr1:123044323 | 1 | 123.0443 | 0.011241536 | Credible Set | purple |
| chr1:123044326 | 1 | 123.0443 | 0.010972144 | Credible Set | purple |
| chr1:123144261 | 1 | 123.1443 | 0.009603843 | Credible Set | purple |
| chr1:123225956 | 1 | 123.2260 | 0.014851880 | Credible Set | purple |
| chr1:123225957 | 1 | 123.2260 | 0.015925704 | Credible Set | purple |
| chr1:123225968 | 1 | 123.2260 | 0.015321794 | Credible Set | purple |
| chr1:123322958 | 1 | 123.3230 | 0.007392939 | Credible Set | purple |
| chr1:123323170 | 1 | 123.3232 | 0.003447493 | Credible Set | purple |
| chr1:123368825 | 1 | 123.3688 | 0.010719158 | Credible Set | purple |
| chr1:123369170 | 1 | 123.3692 | 0.010277294 | Credible Set | purple |
| chr1:123588206 | 1 | 123.5882 | 0.009191190 | Credible Set | purple |
| chr1:123588207 | 1 | 123.5882 | 0.009310789 | Credible Set | purple |
| chr1:123588436 | 1 | 123.5884 | 0.016769146 | Credible Set | purple |
| chr1:123602768 | 1 | 123.6028 | 0.016865161 | Credible Set | purple |
| chr1:123603027 | 1 | 123.6030 | 0.015435306 | Credible Set | purple |
| chr1:123665450 | 1 | 123.6655 | 0.017260549 | Credible Set | purple |
| chr1:123785564 | 1 | 123.7856 | 0.030369351 | Credible Set | purple |
| chr1:123831927 | 1 | 123.8319 | 0.032395972 | Credible Set | purple |
| chr1:123997640 | 1 | 123.9976 | 0.041511594 | Credible Set | purple |
| chr1:123997667 | 1 | 123.9977 | 0.041745092 | Credible Set | purple |
| chr1:124151549 | 1 | 124.1515 | 0.029431303 | Credible Set | purple |
| chr1:124477860 | 1 | 124.4779 | 0.016102671 | Credible Set | purple |
| chr1:124477885 | 1 | 124.4779 | 0.016101238 | Credible Set | purple |
| chr1:124477894 | 1 | 124.4779 | 0.016092643 | Credible Set | purple |
| chr1:124501960 | 1 | 124.5020 | 0.010520156 | Credible Set | purple |
| chr1:124711052 | 1 | 124.7111 | 0.004700363 | Credible Set | purple |
| chr1:124711083 | 1 | 124.7111 | 0.004756787 | Credible Set | purple |
| chr1:124711275 | 1 | 124.7113 | 0.002509492 | Credible Set | purple |
| chr1:124711325 | 1 | 124.7113 | 0.002396614 | Credible Set | purple |
| chr1:124711335 | 1 | 124.7113 | 0.002511671 | Credible Set | purple |
| chr1:124941492 | 1 | 124.9415 | 0.005673875 | Credible Set | purple |
| chr1:125042386 | 1 | 125.0424 | 0.009054356 | Credible Set | purple |
| chr1:125157478 | 1 | 125.1575 | 0.006602808 | Credible Set | purple |
| chr1:125163273 | 1 | 125.1633 | 0.006464504 | Credible Set | purple |
| chr1:125263025 | 1 | 125.2630 | 0.004254125 | Credible Set | purple |

Fine−mapping Regions

| snp | chr | pos (Mb) | pp | group | color |
|---|---|---|---|---|---|
| chr1:125263026 | 1 | 125.2630 | 0.004256381 | Credible Set | purple |
| chr1:125367002 | 1 | 125.3670 | 0.003601180 | Credible Set | purple |
| chr1:125386497 | 1 | 125.3865 | 0.005715202 | Credible Set | purple |
| chr1:125387017 | 1 | 125.3870 | 0.004243755 | Credible Set | purple |
| chr1:125387048 | 1 | 125.3870 | 0.004306048 | Credible Set | purple |
| chr1:125387071 | 1 | 125.3871 | 0.004280330 | Credible Set | purple |
| chr1:125387095 | 1 | 125.3871 | 0.008231250 | Credible Set | purple |
| chr1:125459470 | 1 | 125.4595 | 0.004132014 | Credible Set | purple |
| chr1:125481066 | 1 | 125.4811 | 0.004529628 | Credible Set | purple |
| chr1:125481498 | 1 | 125.4815 | 0.005499793 | Credible Set | purple |
| chr1:125481527 | 1 | 125.4815 | 0.006021720 | Credible Set | purple |
| chr1:125481552 | 1 | 125.4816 | 0.006156479 | Credible Set | purple |
| chr1:125481575 | 1 | 125.4816 | 0.006421286 | Credible Set | purple |
| chr1:125481715 | 1 | 125.4817 | 0.001945740 | Credible Set | purple |
| chr1:125578885 | 1 | 125.5789 | 0.002810203 | Credible Set | purple |
| chr1:125579353 | 1 | 125.5794 | 0.002678265 | Credible Set | purple |
| chr1:125585192 | 1 | 125.5852 | 0.003462203 | Credible Set | purple |
| chr1:125585252 | 1 | 125.5853 | 0.003123502 | Credible Set | purple |
| chr1:125615959 | 1 | 125.6160 | 0.002225522 | Credible Set | purple |
| chr1:125616002 | 1 | 125.6160 | 0.002185989 | Credible Set | purple |
| chr1:125643547 | 1 | 125.6435 | 0.003145129 | Credible Set | purple |
| chr1:125643933 | 1 | 125.6439 | 0.003094798 | Credible Set | purple |
| chr1:125733123 | 1 | 125.7331 | 0.004290058 | Credible Set | purple |
| chr1:125741798 | 1 | 125.7418 | 0.005807763 | Credible Set | purple |
| chr1:125764061 | 1 | 125.7641 | 0.005383415 | Credible Set | purple |
| chr1:125764120 | 1 | 125.7641 | 0.006086535 | Credible Set | purple |
| chr1:125765305 | 1 | 125.7653 | 0.003074121 | Credible Set | purple |
| chr1:125765343 | 1 | 125.7653 | 0.003145375 | Credible Set | purple |
| chr1:125765591 | 1 | 125.7656 | 0.006228165 | Credible Set | purple |
| chr1:125765595 | 1 | 125.7656 | 0.005809238 | Credible Set | purple |
| chr1:125787309 | 1 | 125.7873 | 0.008676441 | Credible Set | purple |
| chr1:125789544 | 1 | 125.7895 | 0.007793146 | Credible Set | purple |
| chr1:125789598 | 1 | 125.7896 | 0.008563652 | Credible Set | purple |
| chr1:125795230 | 1 | 125.7952 | 0.008392435 | Credible Set | purple |
| chr1:125795318 | 1 | 125.7953 | 0.008060562 | Credible Set | purple |
| chr1:125795437 | 1 | 125.7954 | 0.005990406 | Credible Set | purple |

Fine−mapping Regions

| snp | chr | pos (Mb) | pp | group | color |
|---|---|---|---|---|---|
| chr1:125795438 | 1 | 125.7954 | 0.005939926 | Credible Set | purple |
| chr1:125813871 | 1 | 125.8139 | 0.010397190 | Credible Set | purple |
| chr1:125813926 | 1 | 125.8139 | 0.009986284 | Credible Set | purple |
| chr1:125826985 | 1 | 125.8270 | 0.008293128 | Credible Set | purple |
| chr1:125827492 | 1 | 125.8275 | 0.010969665 | Credible Set | purple |
| chr1:125831250 | 1 | 125.8312 | 0.009798995 | Credible Set | purple |
| chr1:125831282 | 1 | 125.8313 | 0.009572848 | Credible Set | purple |
| chr1:125831293 | 1 | 125.8313 | 0.009775255 | Credible Set | purple |
| chr1:125836037 | 1 | 125.8360 | 0.006905259 | Credible Set | purple |
| chr1:125836053 | 1 | 125.8361 | 0.006916447 | Credible Set | purple |
| chr1:125854689 | 1 | 125.8547 | 0.016398789 | Credible Set | purple |
| chr1:125872500 | 1 | 125.8725 | 0.023362569 | Credible Set | purple |
| chr1:125872503 | 1 | 125.8725 | 0.023427212 | Credible Set | purple |
| chr1:125872524 | 1 | 125.8725 | 0.017306554 | Credible Set | purple |
| chr1:125876478 | 1 | 125.8765 | 0.017652913 | Credible Set | purple |
| chr1:125934259 | 1 | 125.9343 | 0.010673487 | Credible Set | purple |
| chr1:125934293 | 1 | 125.9343 | 0.010925334 | Credible Set | purple |
| chr1:125934498 | 1 | 125.9345 | 0.010853985 | Credible Set | purple |
| chr1:125943250 | 1 | 125.9433 | 0.012547516 | Credible Set | purple |
| chr1:125956049 | 1 | 125.9560 | 0.013682406 | Credible Set | purple |
| chr1:126027899 | 1 | 126.0279 | 0.011486449 | Credible Set | purple |
| chr1:126074088 | 1 | 126.0741 | 0.006583365 | Credible Set | purple |
| chr1:126084400 | 1 | 126.0844 | 0.010413670 | Credible Set | purple |
| chr1:126084494 | 1 | 126.0845 | 0.011737507 | Credible Set | purple |
| chr1:126089361 | 1 | 126.0894 | 0.008088780 | Credible Set | purple |
| chr1:126140625 | 1 | 126.1406 | 0.007920435 | Credible Set | purple |
| chr1:126140679 | 1 | 126.1407 | 0.008443348 | Credible Set | purple |
| chr1:126217882 | 1 | 126.2179 | 0.007065695 | Credible Set | purple |
| chr1:126288256 | 1 | 126.2883 | 0.005558527 | Credible Set | purple |

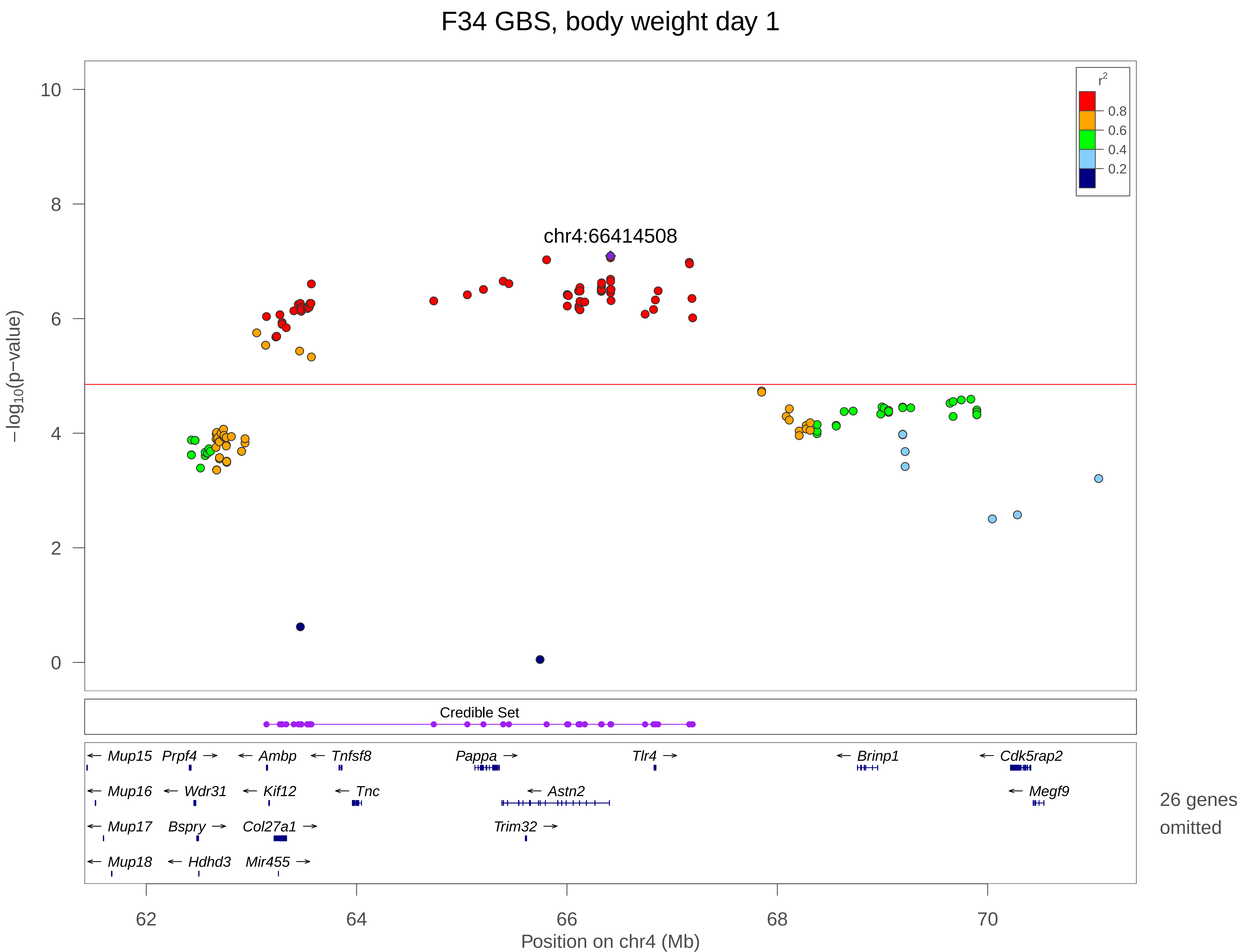

F34 GBS, body weight day 1
chr4:66414508
r2
0.8
0.6
0.4
0.2
−log10(p−value)
10
8
6
4
2
0
Credible Set
Mup15
Prpf4
Ambp
Tnfsf8
Pappa
Tlr4
Brinp1
Cdk5rap2
Mup16
Wdr31
Kif12
Tnc
Astn2
Megf9
Mup17
Bspry
Col27a1
Trim32
Mup18
Hdhd3
Mir455
26 genes omitted
62
64
66
68
70
Position on chr4 (Mb)

| | |
|---|---|
| date: | Thu Oct 25 16:59:56 2018 |
| build: | F34GBS |
| display range: | chr4:61414508−71414508 [61414508−71414508] |
| hilite range: | 0 − 0 [ 0 − 0 ] |
| reference SNP: | chr4:66414508 |
| number of SNPs plotted: | 191 |
| min P−value: | 8.06E−8 [chr4:66414508] |
| max P−value: | 8.91E−1 [chr4:65744347] |
| omitted Genes: | Mup19, Mup5, Mup20 |
| omitted Genes: | Mup3, Mup21, Zfp37 |
| omitted Genes: | Slc31a2, Fkbp15, Slc31a1 |
| omitted Genes: | Cdc26, Rnf183, Alad |
| omitted Genes: | Pole3, 4933430I17Rik, Rgs3 |
| omitted Genes: | Zfp618, Orm1, Orm3 |
| omitted Genes: | Orm2, Akna, Whrn |
| omitted Genes: | Atp6v1g1, 6330416G13Rik, Gm11213 |
| omitted Genes: | 1700018C11Rik, Tnfsf15 |

Make more plots at http://csg.sph.umich.edu/locuszoom/

Fine−mapping Regions

| snp | chr | pos (Mb) | pp | group | color |
|---|---|---|---|---|---|
| chr4:63143387 | 4 | 63.14339 | 0.004920532 | Credible Set | purple |
| chr4:63270812 | 4 | 63.27081 | 0.005286562 | Credible Set | purple |
| chr4:63291699 | 4 | 63.29170 | 0.003919544 | Credible Set | purple |
| chr4:63291735 | 4 | 63.29174 | 0.003895518 | Credible Set | purple |
| chr4:63292002 | 4 | 63.29200 | 0.003641439 | Credible Set | purple |
| chr4:63292021 | 4 | 63.29202 | 0.003657412 | Credible Set | purple |
| chr4:63331035 | 4 | 63.33103 | 0.003198973 | Credible Set | purple |
| chr4:63403594 | 4 | 63.40359 | 0.006146430 | Credible Set | purple |
| chr4:63445899 | 4 | 63.44590 | 0.007880930 | Credible Set | purple |
| chr4:63465528 | 4 | 63.46553 | 0.008176243 | Credible Set | purple |
| chr4:63471028 | 4 | 63.47103 | 0.006807494 | Credible Set | purple |
| chr4:63471086 | 4 | 63.47109 | 0.006701406 | Credible Set | purple |
| chr4:63473150 | 4 | 63.47315 | 0.006064724 | Credible Set | purple |
| chr4:63473181 | 4 | 63.47318 | 0.006026442 | Credible Set | purple |
| chr4:63473231 | 4 | 63.47323 | 0.006770501 | Credible Set | purple |
| chr4:63473937 | 4 | 63.47394 | 0.006987638 | Credible Set | purple |
| chr4:63474009 | 4 | 63.47401 | 0.006476940 | Credible Set | purple |
| chr4:63531405 | 4 | 63.53140 | 0.006799207 | Credible Set | purple |
| chr4:63531436 | 4 | 63.53144 | 0.006706997 | Credible Set | purple |
| chr4:63535096 | 4 | 63.53510 | 0.007191464 | Credible Set | purple |
| chr4:63548238 | 4 | 63.54824 | 0.007458420 | Credible Set | purple |
| chr4:63548572 | 4 | 63.54857 | 0.007001992 | Credible Set | purple |
| chr4:63558825 | 4 | 63.55882 | 0.008222442 | Credible Set | purple |
| chr4:63565618 | 4 | 63.56562 | 0.008176243 | Credible Set | purple |
| chr4:63565712 | 4 | 63.56571 | 0.008176243 | Credible Set | purple |
| chr4:63570198 | 4 | 63.57020 | 0.017426428 | Credible Set | purple |
| chr4:64733521 | 4 | 64.73352 | 0.009019328 | Credible Set | purple |
| chr4:65052703 | 4 | 65.05270 | 0.011424417 | Credible Set | purple |
| chr4:65205920 | 4 | 65.20592 | 0.014056399 | Credible Set | purple |
| chr4:65393970 | 4 | 65.39397 | 0.019387756 | Credible Set | purple |
| chr4:65447841 | 4 | 65.44784 | 0.017604254 | Credible Set | purple |
| chr4:65806420 | 4 | 65.80642 | 0.044483904 | Credible Set | purple |
| chr4:66002530 | 4 | 66.00253 | 0.011555098 | Credible Set | purple |
| chr4:66003029 | 4 | 66.00303 | 0.007394839 | Credible Set | purple |
| chr4:66003164 | 4 | 66.00316 | 0.011297129 | Credible Set | purple |
| chr4:66013512 | 4 | 66.01351 | 0.011044254 | Credible Set | purple |

Fine−mapping Regions

| snp | chr | pos (Mb) | pp | group | color |
|---|---|---|---|---|---|
| chr4:66110072 | 4 | 66.11007 | 0.013212622 | Credible Set | purple |
| chr4:66114163 | 4 | 66.11416 | 0.007359190 | Credible Set | purple |
| chr4:66114164 | 4 | 66.11416 | 0.006808908 | Credible Set | purple |
| chr4:66114176 | 4 | 66.11418 | 0.006844079 | Credible Set | purple |
| chr4:66123765 | 4 | 66.12377 | 0.008867952 | Credible Set | purple |
| chr4:66123768 | 4 | 66.12377 | 0.006380854 | Credible Set | purple |
| chr4:66123832 | 4 | 66.12383 | 0.015171792 | Credible Set | purple |
| chr4:66124242 | 4 | 66.12424 | 0.013213448 | Credible Set | purple |
| chr4:66169378 | 4 | 66.16938 | 0.008650737 | Credible Set | purple |
| chr4:66325857 | 4 | 66.32586 | 0.014138122 | Credible Set | purple |
| chr4:66325987 | 4 | 66.32599 | 0.013078820 | Credible Set | purple |
| chr4:66328189 | 4 | 66.32819 | 0.013209608 | Credible Set | purple |
| chr4:66328195 | 4 | 66.32819 | 0.013217499 | Credible Set | purple |
| chr4:66328436 | 4 | 66.32844 | 0.013823825 | Credible Set | purple |
| chr4:66328528 | 4 | 66.32853 | 0.015206020 | Credible Set | purple |
| chr4:66328560 | 4 | 66.32856 | 0.013843567 | Credible Set | purple |
| chr4:66329044 | 4 | 66.32904 | 0.017044709 | Credible Set | purple |
| chr4:66329065 | 4 | 66.32906 | 0.018199854 | Credible Set | purple |
| chr4:66329095 | 4 | 66.32909 | 0.018061779 | Credible Set | purple |
| chr4:66414332 | 4 | 66.41433 | 0.012736841 | Credible Set | purple |
| chr4:66414345 | 4 | 66.41434 | 0.012978584 | Credible Set | purple |
| chr4:66414373 | 4 | 66.41437 | 0.012452192 | Credible Set | purple |
| chr4:66414374 | 4 | 66.41437 | 0.012460381 | Credible Set | purple |
| chr4:66414384 | 4 | 66.41438 | 0.014019326 | Credible Set | purple |
| chr4:66414387 | 4 | 66.41439 | 0.019585308 | Credible Set | purple |
| chr4:66414451 | 4 | 66.41445 | 0.020897239 | Credible Set | purple |
| chr4:66414476 | 4 | 66.41448 | 0.048901218 | Credible Set | purple |
| chr4:66414508 | 4 | 66.41451 | 0.051683313 | Credible Set | purple |
| chr4:66414513 | 4 | 66.41451 | 0.048088101 | Credible Set | purple |
| chr4:66414963 | 4 | 66.41496 | 0.019242291 | Credible Set | purple |
| chr4:66419064 | 4 | 66.41906 | 0.009116121 | Credible Set | purple |
| chr4:66419126 | 4 | 66.41913 | 0.014049268 | Credible Set | purple |
| chr4:66743103 | 4 | 66.74310 | 0.005389529 | Credible Set | purple |
| chr4:66823255 | 4 | 66.82326 | 0.006460812 | Credible Set | purple |
| chr4:66840696 | 4 | 66.84070 | 0.009357500 | Credible Set | purple |
| chr4:66866758 | 4 | 66.86676 | 0.013333133 | Credible Set | purple |

Fine−mapping Regions

| snp | chr | pos (Mb) | pp | group | color |
|---|---|---|---|---|---|
| chr4:67163007 | 4 | 67.16301 | 0.040364868 | Credible Set | purple |
| chr4:67165680 | 4 | 67.16568 | 0.037908162 | Credible Set | purple |
| chr4:67188375 | 4 | 67.18837 | 0.009903913 | Credible Set | purple |
| chr4:67194875 | 4 | 67.19487 | 0.004679250 | Credible Set | purple |

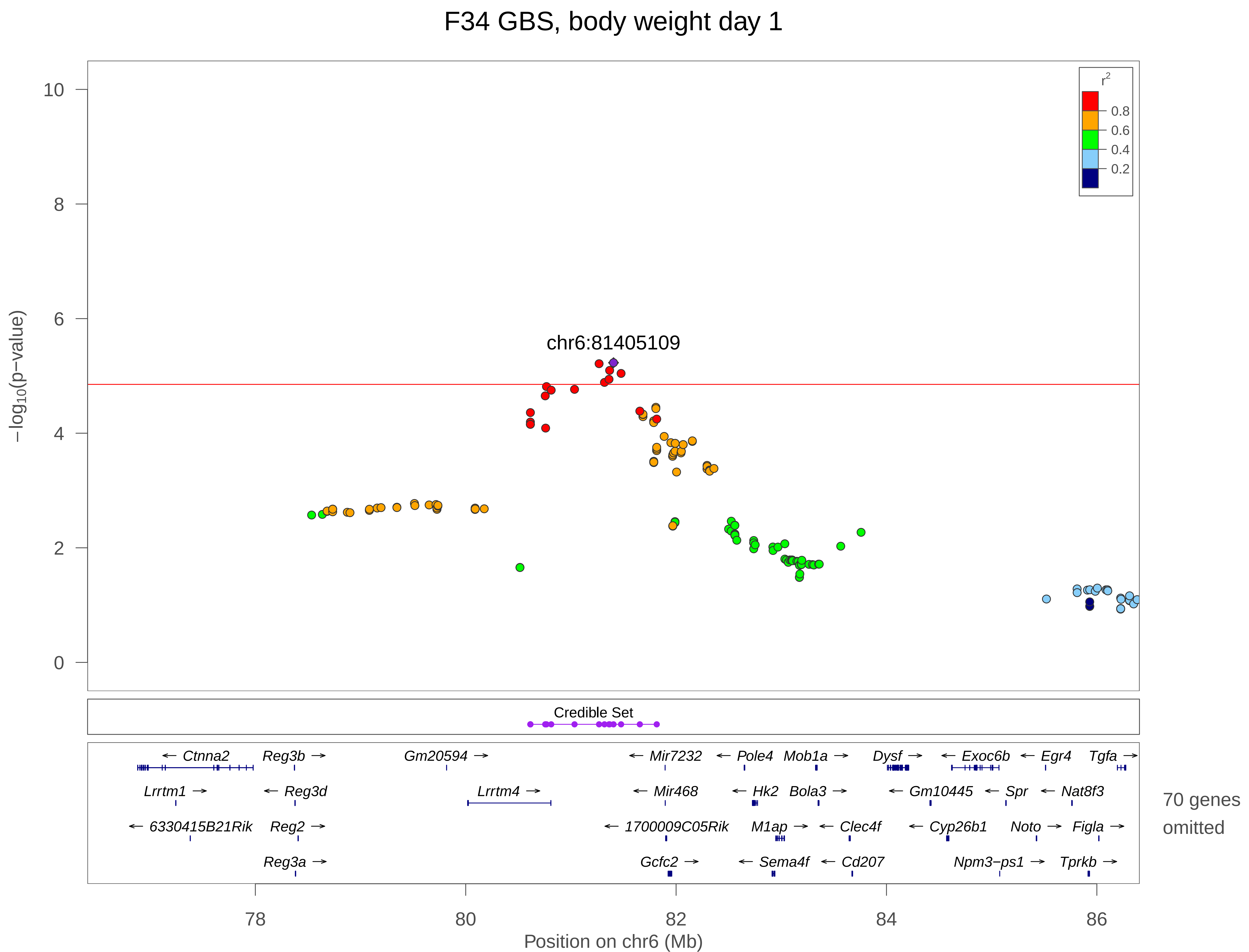
F34 GBS, body weight day 1
chr6:81405109
−log10(p−value)
r2
0.8
0.6
0.4
0.2
Credible Set
← Ctnna2
Reg3b →
Gm20594 →
← Mir7232
← Pole4
Mob1a →
Dysf →
← Exoc6b
← Egr4
Tgfa →
Lrrtm1 →
← Reg3d
Lrrtm4 →
← Mir468
← Hk2
Bola3 →
← Gm10445
← Spr
← Nat8f3
← 6330415B21Rik
Reg2 →
← 1700009C05Rik
M1ap →
← Clec4f
← Cyp26b1
Noto →
Figla →
Reg3a →
Gcfc2 →
← Sema4f
← Cd207
Npm3−ps1 →
Tprkb →
70 genes omitted
Position on chr6 (Mb)

| | |
|---|---|
| date: | Thu Oct 25 17:00:07 2018 |
| build: | F34GBS |
| display range: | chr6:76405109−86405109 [76405109−86405109] |
| hilite range: | 0 − 0 [ 0 − 0 ] |
| reference SNP: | chr6:81405109 |
| number of SNPs plotted: | 172 |
| min P−value: | 5.89E−6 [chr6:81405109] |
| max P−value: | 1.17E−1 [chr6:86226324] |
| omitted Genes: | Reg1, Reg3g, Gm20362 |
| omitted Genes: | Mrpl19, Eva1a, Tacr1 |
| omitted Genes: | 2310069B03Rik, Dok1, Loxl3 |
| omitted Genes: | Mir7040, Htra2, Aup1 |
| omitted Genes: | Dqx1, Tlx2, Pcgf1 |
| omitted Genes: | Lbx2, Mir3470a, Ccdc142 |
| omitted Genes: | Mrpl53, Mogs, Wbp1 |
| omitted Genes: | Ino80b, Rtkn, Wdr54 |
| omitted Genes: | 1700003E16Rik, Dctn1, Slc4a5 |
| omitted Genes: | Mthfd2, Tet3, Mir6374 |
| omitted Genes: | B230319C09Rik, Dguok, Actg2 |
| omitted Genes: | Stambp, Gm21284, Vax2os |
| omitted Genes: | Vax2, Atp6v1b1, 1700124L16Rik |
| omitted Genes: | Ankrd53, Tex261, Gm7443 |
| omitted Genes: | Nagk, Paip2b, Zfp638 |
| omitted Genes: | Gm5878, Emx1, Sfxn5 |
| omitted Genes: | Rab11fip5, Mir705, Smyd5 |
| omitted Genes: | Pradc1, Cct7, Fbxo41 |
| omitted Genes: | Alms1, Nat8f7, Mir6375 |
| omitted Genes: | Alms1−ps2, Nat8f6, Nat8f5 |
| omitted Genes: | Nat8, Nat8f2, Nat8f4 |
| omitted Genes: | Nat8f1, Dusp11, Add2 |
| omitted Genes: | Fam136a, Snrpg, Pcyox1 |
| omitted Genes: | Tia1 |

Make more plots at http://csg.sph.umich.edu/locuszoom/

Fine−mapping Regions

| snp | chr | pos (Mb) | pp | group | color |
|---|---|---|---|---|---|
| chr6:80614710 | 6 | 80.61471 | 0.01554354 | Credible Set | purple |
| chr6:80614725 | 6 | 80.61473 | 0.01423141 | Credible Set | purple |
| chr6:80614740 | 6 | 80.61474 | 0.01429243 | Credible Set | purple |
| chr6:80614750 | 6 | 80.61475 | 0.01423141 | Credible Set | purple |
| chr6:80614907 | 6 | 80.61491 | 0.02227719 | Credible Set | purple |
| chr6:80755939 | 6 | 80.75594 | 0.04218344 | Credible Set | purple |
| chr6:80759589 | 6 | 80.75959 | 0.01234583 | Credible Set | purple |
| chr6:80767933 | 6 | 80.76793 | 0.06018603 | Credible Set | purple |
| chr6:80812582 | 6 | 80.81258 | 0.05245386 | Credible Set | purple |
| chr6:81034488 | 6 | 81.03449 | 0.05411055 | Credible Set | purple |
| chr6:81267890 | 6 | 81.26789 | 0.14493198 | Credible Set | purple |
| chr6:81318989 | 6 | 81.31899 | 0.07060812 | Credible Set | purple |
| chr6:81362217 | 6 | 81.36222 | 0.07964570 | Credible Set | purple |
| chr6:81368112 | 6 | 81.36811 | 0.11190384 | Credible Set | purple |
| chr6:81405109 | 6 | 81.40511 | 0.15035759 | Credible Set | purple |
| chr6:81476902 | 6 | 81.47690 | 0.09971273 | Credible Set | purple |
| chr6:81655195 | 6 | 81.65520 | 0.02357117 | Credible Set | purple |
| chr6:81815817 | 6 | 81.81582 | 0.01741315 | Credible Set | purple |

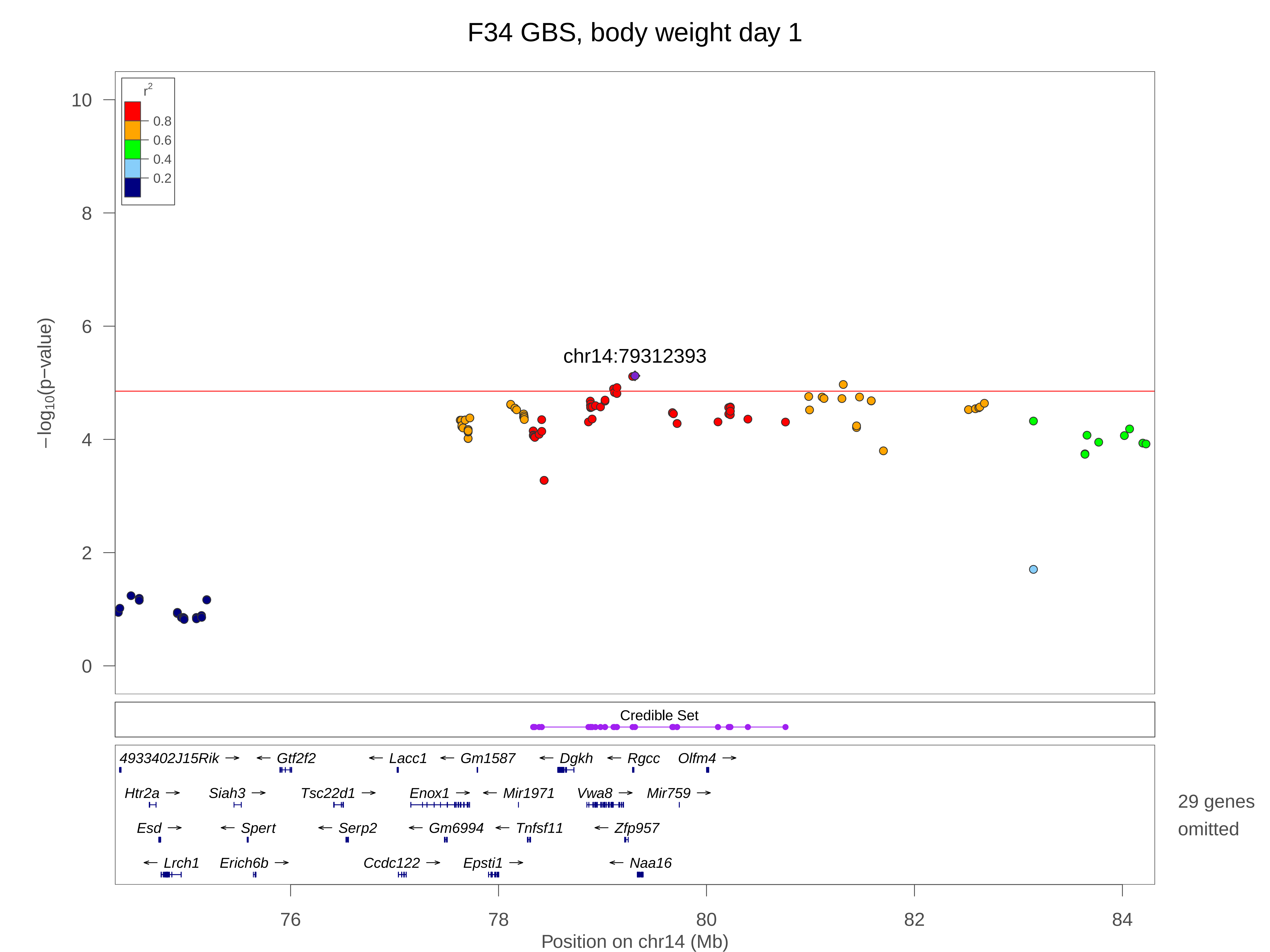

F34 GBS, body weight day 1
r²
0.8
0.6
0.4
0.2
10
8
6
4
2
0
−log10(p−value)
chr14:79312393
Credible Set
4933402J15Rik →
← Gtf2f2
← Lacc1
← Gm1587
← Dgkh
← Rgcc
Olfm4 →
Htr2a →
Siah3 →
Tsc22d1 →
Enox1 →
← Mir1971
Vwa8 →
Mir759 →
Esd →
← Spert
← Serp2
← Gm6994
← Tnfsf11
← Zfp957
← Lrch1
Erich6b →
Ccdc122 →
Epsti1 →
← Naa16
29 genes omitted
76
78
80
82
84
Position on chr14 (Mb)

| | |
|---|---|
| date: | Thu Oct 25 16:59:47 2018 |
| build: | F34GBS |
| display range: | chr14:74312393−84312393 [74312393−84312393] |
| hilite range: | 0 − 0 [ 0 − 0 ] |
| reference SNP: | chr14:79312393 |
| number of SNPs plotted: | 127 |
| min P−value: | 7.53E−6 [chr14:79312393] |
| max P−value: | 1.52E−1 [chr14:74975984] |
| omitted Genes: | Gm4278, 5031414D18Rik, Lrrc63 |
| omitted Genes: | Lcp1, Cpb2, Zc3h13 |
| omitted Genes: | Cog3, Slc25a30, Gm4285 |
| omitted Genes: | Tpt1, Snora31, Kctd4 |
| omitted Genes: | Gpalpp1, Nufip1, 4930444M15Rik |
| omitted Genes: | 1700108F19Rik, Dnajc15, Fam216b |
| omitted Genes: | Akap11, Mtrf1, Kbtbd7 |
| omitted Genes: | Kbtbd6, Wbp4, Elf1 |
| omitted Genes: | Sugt1, Lect1, 4930452G13Rik |
| omitted Genes: | Pcdh8, Gm10845 |

Make more plots at http://csg.sph.umich.edu/locuszoom/

Fine−mapping Regions

| snp | chr | pos (Mb) | pp | group | color |
|---|---|---|---|---|---|
| chr14:78333693 | 14 | 78.33369 | 0.009596012 | Credible Set | purple |
| chr14:78333727 | 14 | 78.33373 | 0.009605429 | Credible Set | purple |
| chr14:78334235 | 14 | 78.33424 | 0.008176264 | Credible Set | purple |
| chr14:78334352 | 14 | 78.33435 | 0.008002390 | Credible Set | purple |
| chr14:78342935 | 14 | 78.34293 | 0.007662936 | Credible Set | purple |
| chr14:78349294 | 14 | 78.34929 | 0.007479411 | Credible Set | purple |
| chr14:78390714 | 14 | 78.39071 | 0.008413361 | Credible Set | purple |
| chr14:78415035 | 14 | 78.41504 | 0.014815595 | Credible Set | purple |
| chr14:78415900 | 14 | 78.41590 | 0.009447704 | Credible Set | purple |
| chr14:78864748 | 14 | 78.86475 | 0.013558704 | Credible Set | purple |
| chr14:78882395 | 14 | 78.88240 | 0.030401999 | Credible Set | purple |
| chr14:78884278 | 14 | 78.88428 | 0.025644109 | Credible Set | purple |
| chr14:78885403 | 14 | 78.88540 | 0.023519885 | Credible Set | purple |
| chr14:78885542 | 14 | 78.88554 | 0.027055887 | Credible Set | purple |
| chr14:78885909 | 14 | 78.88591 | 0.024172435 | Credible Set | purple |
| chr14:78900265 | 14 | 78.90027 | 0.015186616 | Credible Set | purple |
| chr14:78900272 | 14 | 78.90027 | 0.024273387 | Credible Set | purple |
| chr14:78932384 | 14 | 78.93238 | 0.025452236 | Credible Set | purple |
| chr14:78980736 | 14 | 78.98074 | 0.024189889 | Credible Set | purple |
| chr14:79024151 | 14 | 79.02415 | 0.030356851 | Credible Set | purple |
| chr14:79024207 | 14 | 79.02421 | 0.031572210 | Credible Set | purple |
| chr14:79105928 | 14 | 79.10593 | 0.048647173 | Credible Set | purple |
| chr14:79115207 | 14 | 79.11521 | 0.042428195 | Credible Set | purple |
| chr14:79138261 | 14 | 79.13826 | 0.040676940 | Credible Set | purple |
| chr14:79138486 | 14 | 79.13849 | 0.051279397 | Credible Set | purple |
| chr14:79289663 | 14 | 79.28966 | 0.078969096 | Credible Set | purple |
| chr14:79312393 | 14 | 79.31239 | 0.080912275 | Credible Set | purple |
| chr14:79672373 | 14 | 79.67237 | 0.019294935 | Credible Set | purple |
| chr14:79672387 | 14 | 79.67239 | 0.019192702 | Credible Set | purple |
| chr14:79672589 | 14 | 79.67259 | 0.019449424 | Credible Set | purple |
| chr14:79682220 | 14 | 79.68222 | 0.018595042 | Credible Set | purple |
| chr14:79717330 | 14 | 79.71733 | 0.012796695 | Credible Set | purple |
| chr14:79717420 | 14 | 79.71742 | 0.012748553 | Credible Set | purple |
| chr14:80110393 | 14 | 80.11039 | 0.013594815 | Credible Set | purple |
| chr14:80213369 | 14 | 80.21337 | 0.023644892 | Credible Set | purple |
| chr14:80214058 | 14 | 80.21406 | 0.018461552 | Credible Set | purple |

Fine−mapping Regions

| snp | chr | pos (Mb) | pp | group | color |
|---|---|---|---|---|---|
| chr14:80228969 | 14 | 80.22897 | 0.01790472 | Credible Set | purple |
| chr14:80229277 | 14 | 80.22928 | 0.02419785 | Credible Set | purple |
| chr14:80229281 | 14 | 80.22928 | 0.02364378 | Credible Set | purple |
| chr14:80229399 | 14 | 80.22940 | 0.02054354 | Credible Set | purple |
| chr14:80397940 | 14 | 80.39794 | 0.01511391 | Credible Set | purple |
| chr14:80759627 | 14 | 80.75963 | 0.01351342 | Credible Set | purple |

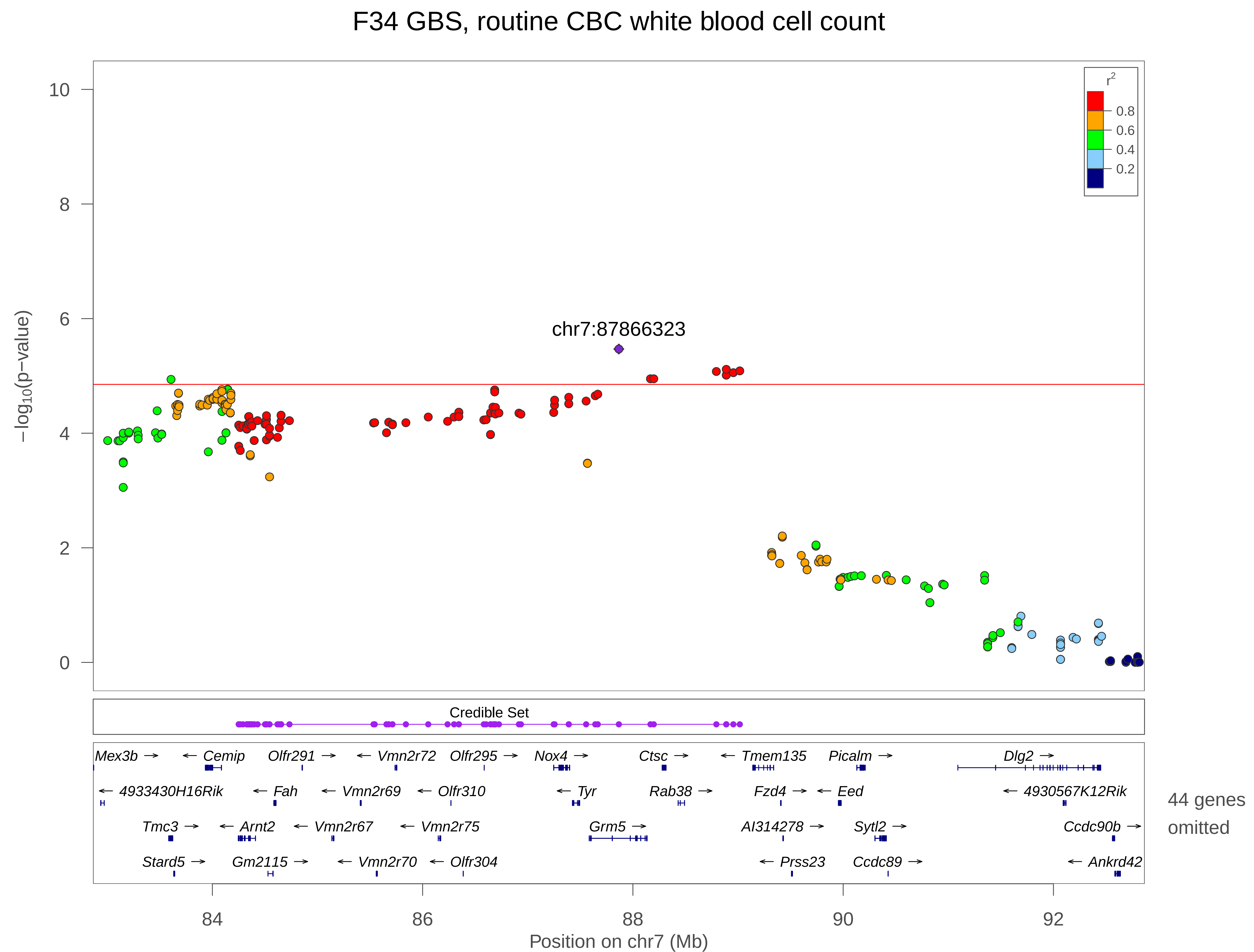
F34 GBS, routine CBC white blood cell count
chr7:87866323
−log10(p−value)
r2
0.8
0.6
0.4
0.2
Credible Set
Mex3b
Cemip
Olfr291
Vmn2r72
Olfr295
Nox4
Ctsc
Tmem135
Picalm
Dlg2
4933430H16Rik
Fah
Vmn2r69
Olfr310
Tyr
Rab38
Fzd4
Eed
4930567K12Rik
Tmc3
Arnt2
Vmn2r67
Vmn2r75
Grm5
AI314278
Sytl2
Ccdc90b
Stard5
Gm2115
Vmn2r70
Olfr304
Prss23
Ccdc89
Ankrd42
44 genes omitted
Position on chr7 (Mb)

| | |
|---|---|
| date: | Thu Oct 25 17:00:43 2018 |
| build: | F34GBS |
| display range: | chr7:82866323−92866323 [82866323−92866323] |
| hilite range: | 0 − 0 [ 0 − 0 ] |
| reference SNP: | chr7:87866323 |
| number of SNPs plotted: | 272 |
| min P−value: | 3.4E−6 [chr7:87866323] |
| max P−value: | 10E−1 [chr7:92800528] |
| omitted Genes: | Il16, Cfap161, Mesdc1 |
| omitted Genes: | Mesdc2, Abhd17c, Zfand6 |
| omitted Genes: | 2610206C17Rik, Olfr290, Vmn2r65 |
| omitted Genes: | Vmn2r66, Vmn2r68, Vmn2r71 |
| omitted Genes: | Vmn2r73, Vmn2r74, Vmn2r76 |
| omitted Genes: | Olfr309, Olfr308, Olfr307 |
| omitted Genes: | Olfr305, Olfr303, Olfr301 |
| omitted Genes: | Olfr299, Olfr298, Olfr297 |
| omitted Genes: | Olfr294, Olfr293, Olfr292 |
| omitted Genes: | Folh1, Vmn2r77, Vmn2r78 |
| omitted Genes: | Vmn2r79, Me3, Ccdc81 |
| omitted Genes: | l7Rn6, E230029C05Rik, 2310010J17Rik |
| omitted Genes: | Ccdc83, Crebzf, Tmem126a |
| omitted Genes: | Tmem126b, Pcf11, 4632427E13Rik |
| omitted Genes: | Rab30, Ddias |

Make more plots at http://csg.sph.umich.edu/locuszoom/

Fine−mapping Regions

| snp | chr | pos (Mb) | pp | group | color |
|---|---|---|---|---|---|
| chr7:84250985 | 7 | 84.25099 | 0.005501709 | Credible Set | purple |
| chr7:84251011 | 7 | 84.25101 | 0.005475841 | Credible Set | purple |
| chr7:84265685 | 7 | 84.26569 | 0.005016557 | Credible Set | purple |
| chr7:84291540 | 7 | 84.29154 | 0.005329965 | Credible Set | purple |
| chr7:84327288 | 7 | 84.32729 | 0.005235804 | Credible Set | purple |
| chr7:84327808 | 7 | 84.32781 | 0.004717339 | Credible Set | purple |
| chr7:84327850 | 7 | 84.32785 | 0.004793861 | Credible Set | purple |
| chr7:84338595 | 7 | 84.33859 | 0.005749274 | Credible Set | purple |
| chr7:84343118 | 7 | 84.34312 | 0.005686161 | Credible Set | purple |
| chr7:84344805 | 7 | 84.34480 | 0.007572429 | Credible Set | purple |
| chr7:84347423 | 7 | 84.34742 | 0.007649440 | Credible Set | purple |
| chr7:84360038 | 7 | 84.36004 | 0.006032025 | Credible Set | purple |
| chr7:84360626 | 7 | 84.36063 | 0.005934546 | Credible Set | purple |
| chr7:84360642 | 7 | 84.36064 | 0.005934546 | Credible Set | purple |
| chr7:84360703 | 7 | 84.36070 | 0.005934546 | Credible Set | purple |
| chr7:84360714 | 7 | 84.36071 | 0.005934546 | Credible Set | purple |
| chr7:84360716 | 7 | 84.36072 | 0.005934546 | Credible Set | purple |
| chr7:84375526 | 7 | 84.37553 | 0.006240815 | Credible Set | purple |
| chr7:84375577 | 7 | 84.37558 | 0.005305038 | Credible Set | purple |
| chr7:84375642 | 7 | 84.37564 | 0.005305038 | Credible Set | purple |
| chr7:84397575 | 7 | 84.39758 | 0.003067353 | Credible Set | purple |
| chr7:84397597 | 7 | 84.39760 | 0.003067353 | Credible Set | purple |
| chr7:84430102 | 7 | 84.43010 | 0.006523611 | Credible Set | purple |
| chr7:84501082 | 7 | 84.50108 | 0.005692555 | Credible Set | purple |
| chr7:84514082 | 7 | 84.51408 | 0.006742649 | Credible Set | purple |
| chr7:84514444 | 7 | 84.51444 | 0.006831071 | Credible Set | purple |
| chr7:84514505 | 7 | 84.51450 | 0.007872161 | Credible Set | purple |
| chr7:84514564 | 7 | 84.51456 | 0.003141971 | Credible Set | purple |
| chr7:84514575 | 7 | 84.51457 | 0.003159374 | Credible Set | purple |
| chr7:84514734 | 7 | 84.51473 | 0.005690317 | Credible Set | purple |
| chr7:84543346 | 7 | 84.54335 | 0.003607470 | Credible Set | purple |
| chr7:84543355 | 7 | 84.54336 | 0.003751895 | Credible Set | purple |
| chr7:84543782 | 7 | 84.54378 | 0.004868379 | Credible Set | purple |
| chr7:84619632 | 7 | 84.61963 | 0.003458170 | Credible Set | purple |
| chr7:84636759 | 7 | 84.63676 | 0.004954218 | Credible Set | purple |
| chr7:84653095 | 7 | 84.65309 | 0.006367397 | Credible Set | purple |

Fine−mapping Regions

| snp | chr | pos (Mb) | pp | group | color |
|---|---|---|---|---|---|
| chr7:84653660 | 7 | 84.65366 | 0.008037400 | Credible Set | purple |
| chr7:84653702 | 7 | 84.65370 | 0.007946669 | Credible Set | purple |
| chr7:84653707 | 7 | 84.65371 | 0.007985099 | Credible Set | purple |
| chr7:84733186 | 7 | 84.73319 | 0.006513920 | Credible Set | purple |
| chr7:85533207 | 7 | 85.53321 | 0.005973992 | Credible Set | purple |
| chr7:85543843 | 7 | 85.54384 | 0.006046224 | Credible Set | purple |
| chr7:85656243 | 7 | 85.65624 | 0.004116651 | Credible Set | purple |
| chr7:85677494 | 7 | 85.67749 | 0.006148412 | Credible Set | purple |
| chr7:85712663 | 7 | 85.71266 | 0.005744222 | Credible Set | purple |
| chr7:85712975 | 7 | 85.71298 | 0.005566094 | Credible Set | purple |
| chr7:85840665 | 7 | 85.84067 | 0.006027668 | Credible Set | purple |
| chr7:86053386 | 7 | 86.05339 | 0.007480299 | Credible Set | purple |
| chr7:86237374 | 7 | 86.23737 | 0.006361786 | Credible Set | purple |
| chr7:86298979 | 7 | 86.29898 | 0.007477890 | Credible Set | purple |
| chr7:86344420 | 7 | 86.34442 | 0.008818260 | Credible Set | purple |
| chr7:86344437 | 7 | 86.34444 | 0.009002727 | Credible Set | purple |
| chr7:86344488 | 7 | 86.34449 | 0.007594144 | Credible Set | purple |
| chr7:86582139 | 7 | 86.58214 | 0.006693461 | Credible Set | purple |
| chr7:86603770 | 7 | 86.60377 | 0.006766168 | Credible Set | purple |
| chr7:86645719 | 7 | 86.64572 | 0.003824593 | Credible Set | purple |
| chr7:86645792 | 7 | 86.64579 | 0.003862274 | Credible Set | purple |
| chr7:86645947 | 7 | 86.64595 | 0.008697316 | Credible Set | purple |
| chr7:86645990 | 7 | 86.64599 | 0.008776408 | Credible Set | purple |
| chr7:86668998 | 7 | 86.66900 | 0.010918322 | Credible Set | purple |
| chr7:86685426 | 7 | 86.68543 | 0.021104555 | Credible Set | purple |
| chr7:86685507 | 7 | 86.68551 | 0.019524378 | Credible Set | purple |
| chr7:86685648 | 7 | 86.68565 | 0.008808600 | Credible Set | purple |
| chr7:86692214 | 7 | 86.69221 | 0.010800522 | Credible Set | purple |
| chr7:86692491 | 7 | 86.69249 | 0.008386308 | Credible Set | purple |
| chr7:86725518 | 7 | 86.72552 | 0.008731934 | Credible Set | purple |
| chr7:86916161 | 7 | 86.91616 | 0.008683614 | Credible Set | purple |
| chr7:86934710 | 7 | 86.93471 | 0.008346359 | Credible Set | purple |
| chr7:87246786 | 7 | 87.24679 | 0.008888467 | Credible Set | purple |
| chr7:87254690 | 7 | 87.25469 | 0.011770641 | Credible Set | purple |
| chr7:87255156 | 7 | 87.25516 | 0.014260261 | Credible Set | purple |
| chr7:87389952 | 7 | 87.38995 | 0.015900164 | Credible Set | purple |

Fine−mapping Regions

| snp | chr | pos (Mb) | pp | group | color |
|---|---|---|---|---|---|
| chr7:87389989 | 7 | 87.38999 | 0.01238772 | Credible Set | purple |
| chr7:87390014 | 7 | 87.39001 | 0.01246453 | Credible Set | purple |
| chr7:87554961 | 7 | 87.55496 | 0.01375690 | Credible Set | purple |
| chr7:87642045 | 7 | 87.64204 | 0.01685574 | Credible Set | purple |
| chr7:87665727 | 7 | 87.66573 | 0.01790927 | Credible Set | purple |
| chr7:87866323 | 7 | 87.86632 | 0.10136765 | Credible Set | purple |
| chr7:88167216 | 7 | 88.16722 | 0.03228100 | Credible Set | purple |
| chr7:88198240 | 7 | 88.19824 | 0.03228262 | Credible Set | purple |
| chr7:88793653 | 7 | 88.79365 | 0.04273059 | Credible Set | purple |
| chr7:88888100 | 7 | 88.88810 | 0.03716292 | Credible Set | purple |
| chr7:88888124 | 7 | 88.88812 | 0.03716292 | Credible Set | purple |
| chr7:88888200 | 7 | 88.88820 | 0.04640861 | Credible Set | purple |
| chr7:88955188 | 7 | 88.95519 | 0.04081574 | Credible Set | purple |
| chr7:89016577 | 7 | 89.01658 | 0.04373922 | Credible Set | purple |

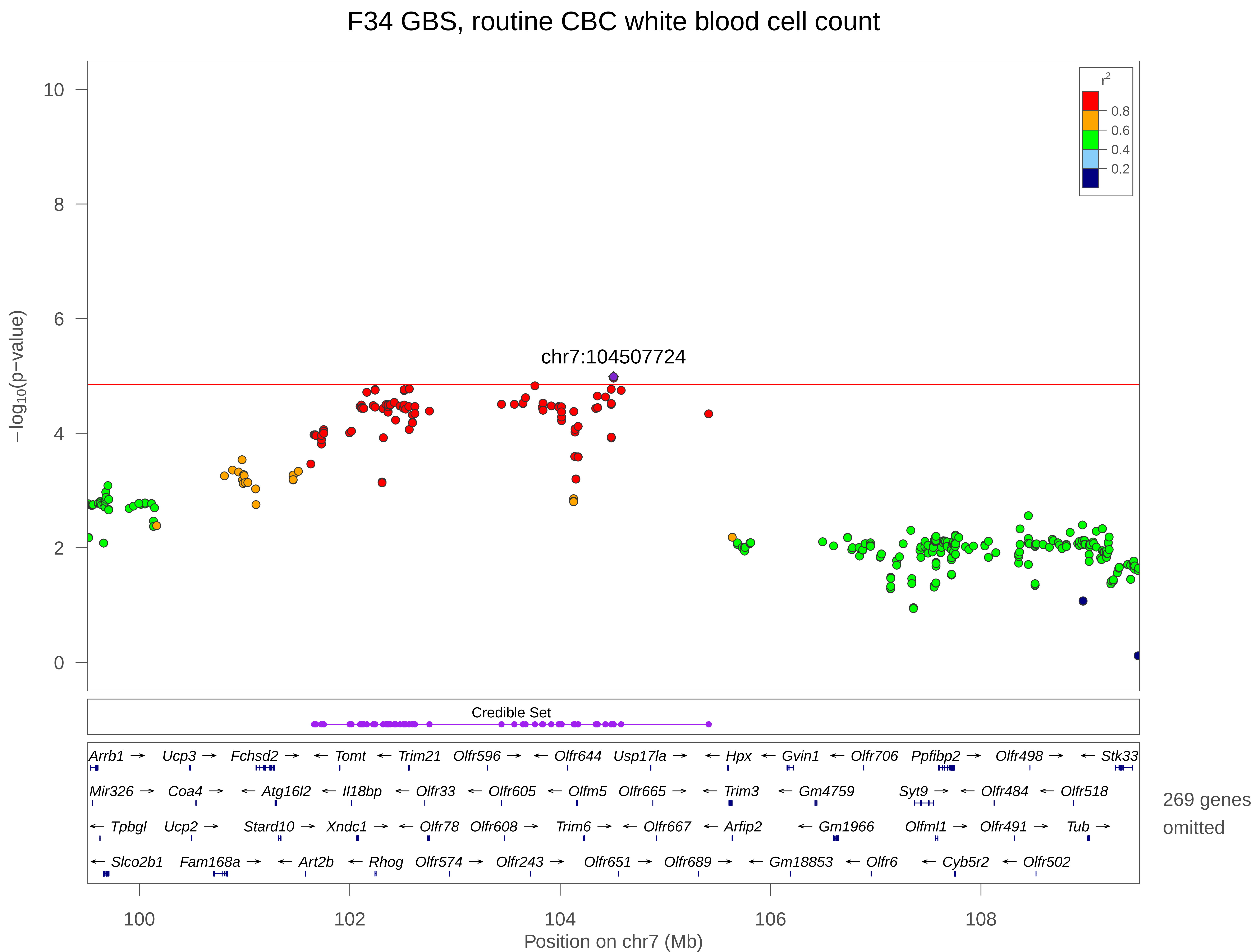
F34 GBS, routine CBC white blood cell count
r2
0.8
0.6
0.4
0.2
chr7:104507724
−log10(p−value)
Credible Set
Arrb1 →
Ucp3 →
Fchsd2 →
← Tomt
← Trim21
Olfr596 →
← Olfr644
Usp17la →
← Hpx
← Gvin1
← Olfr706
Ppfibp2 →
Olfr498 →
← Stk33
Mir326 →
Coa4 →
← Atg16l2
← Il18bp
← Olfr33
← Olfr605
← Olfm5
Olfr665 →
← Trim3
← Gm4759
Syt9 →
← Olfr484
← Olfr518
← Tpbgl
Ucp2 →
Stard10 →
Xndc1 →
← Olfr78
Olfr608 →
Trim6 →
← Olfr667
← Arfip2
← Gm1966
Olfml1 →
Olfr491 →
Tub →
← Slco2b1
Fam168a →
← Art2b
← Rhog
Olfr574 →
Olfr243 →
Olfr651 →
Olfr689 →
← Gm18853
← Olfr6
← Cyb5r2
← Olfr502
269 genes omitted
Position on chr7 (Mb)

date: Thu Oct 25 17:00:32 2018
build: F34GBS
display range: chr7:99507724−109507724 [99507724−109507724]
hilite range: 0 − 0 [ 0 − 0 ]
reference SNP: chr7:104507724
number of SNPs plotted: 453
min P−value: 1.03E−5 [chr7:104507724]
max P−value: 7.67E−1 [chr7:109493860]
omitted Genes: Olfr520, Olfr521, Neu3
omitted Genes: Spcs2, Xrra1, Rnf169
omitted Genes: Chrdl2, Pold3, Lipt2
omitted Genes: Kcne3, Pgm2l1, Gpx2−ps1
omitted Genes: P4ha3, Ppme1, C2cd3
omitted Genes: Dnajb13, Mrpl48, Rab6a
omitted Genes: Plekhb1, Relt, Arhgef17
omitted Genes: Mir3102, P2ry6, P2ry2
omitted Genes: Arap1, Pde2a, Mir139
omitted Genes: Art2a−ps, Clpb, Phox2a
omitted Genes: Inppl1, Folr2, Folr1
omitted Genes: Anapc15, Lamtor1, Lrrc51
omitted Genes: Numa1, Rnf121, Xntrpc
omitted Genes: Trpc2, Art5, Art1
omitted Genes: Chrna10, Nup98, Pgap2
omitted Genes: Stim1, Rrm1, Olfr543
omitted Genes: Olfr544, Olfr545, Olfr547
omitted Genes: Olfr549, Olfr550, Olfr551
omitted Genes: Olfr552, Olfr553, Olfr554
omitted Genes: Olfr555, Olfr556, Trim68
omitted Genes: Olfr557, Olfr558, Olfr559
omitted Genes: Olfr560, Olfr561, Olfr564
omitted Genes: Olfr566, Olfr568, Olfr569
omitted Genes: Olfr570, Olfr571, Olfr572
omitted Genes: Olfr575, Olfr576, Olfr577
omitted Genes: Olfr578, Olfr582, Olfr583
omitted Genes: Olfr584, Olfr585, Olfr586
omitted Genes: Olfr589, Olfr591, Olfr592
omitted Genes: Olfr593, Olfr594, Usp17ld
omitted Genes: Olfr597, Olfr598, Olfr599
omitted Genes: Olfr600, Olfr601, Olfr603
omitted Genes: Usp17lc, Olfr606, Olfr609

Make more plots at http://csg.sph.umich.edu/locuszoom/

Fine−mapping Regions

| snp | chr | pos (Mb) | pp | group | color |
|---|---|---|---|---|---|
| chr7:101660383 | 7 | 101.6604 | 0.003134497 | Credible Set | purple |
| chr7:101669186 | 7 | 101.6692 | 0.003119559 | Credible Set | purple |
| chr7:101673483 | 7 | 101.6735 | 0.003137235 | Credible Set | purple |
| chr7:101680140 | 7 | 101.6801 | 0.003034775 | Credible Set | purple |
| chr7:101731061 | 7 | 101.7311 | 0.003066289 | Credible Set | purple |
| chr7:101731110 | 7 | 101.7311 | 0.002205150 | Credible Set | purple |
| chr7:101731130 | 7 | 101.7311 | 0.002601388 | Credible Set | purple |
| chr7:101731658 | 7 | 101.7317 | 0.003051814 | Credible Set | purple |
| chr7:101751790 | 7 | 101.7518 | 0.003815374 | Credible Set | purple |
| chr7:101752580 | 7 | 101.7526 | 0.003585198 | Credible Set | purple |
| chr7:101752971 | 7 | 101.7530 | 0.003329312 | Credible Set | purple |
| chr7:101998679 | 7 | 101.9987 | 0.003384327 | Credible Set | purple |
| chr7:102016307 | 7 | 102.0163 | 0.003595272 | Credible Set | purple |
| chr7:102098054 | 7 | 102.0981 | 0.009278800 | Credible Set | purple |
| chr7:102110109 | 7 | 102.1101 | 0.009760998 | Credible Set | purple |
| chr7:102110501 | 7 | 102.1105 | 0.008607181 | Credible Set | purple |
| chr7:102116318 | 7 | 102.1163 | 0.008893541 | Credible Set | purple |
| chr7:102116430 | 7 | 102.1164 | 0.008893541 | Credible Set | purple |
| chr7:102116478 | 7 | 102.1165 | 0.008598750 | Credible Set | purple |
| chr7:102132562 | 7 | 102.1326 | 0.008566080 | Credible Set | purple |
| chr7:102162058 | 7 | 102.1621 | 0.015849017 | Credible Set | purple |
| chr7:102223396 | 7 | 102.2234 | 0.009521621 | Credible Set | purple |
| chr7:102240695 | 7 | 102.2407 | 0.008983457 | Credible Set | purple |
| chr7:102241191 | 7 | 102.2412 | 0.017604614 | Credible Set | purple |
| chr7:102241211 | 7 | 102.2412 | 0.017218708 | Credible Set | purple |
| chr7:102241212 | 7 | 102.2412 | 0.017212173 | Credible Set | purple |
| chr7:102317213 | 7 | 102.3172 | 0.008417846 | Credible Set | purple |
| chr7:102317246 | 7 | 102.3172 | 0.008523494 | Credible Set | purple |
| chr7:102319586 | 7 | 102.3196 | 0.002807123 | Credible Set | purple |
| chr7:102344970 | 7 | 102.3450 | 0.009893181 | Credible Set | purple |
| chr7:102363897 | 7 | 102.3639 | 0.009878185 | Credible Set | purple |
| chr7:102364204 | 7 | 102.3642 | 0.007674797 | Credible Set | purple |
| chr7:102364223 | 7 | 102.3642 | 0.007409394 | Credible Set | purple |
| chr7:102364315 | 7 | 102.3643 | 0.009087320 | Credible Set | purple |
| chr7:102364326 | 7 | 102.3643 | 0.009087320 | Credible Set | purple |
| chr7:102364411 | 7 | 102.3644 | 0.009740598 | Credible Set | purple |

Fine−mapping Regions

| snp | chr | pos (Mb) | pp | group | color |
|---|---|---|---|---|---|
| chr7:102385520 | 7 | 102.3855 | 0.009649962 | Credible Set | purple |
| chr7:102385536 | 7 | 102.3855 | 0.009838476 | Credible Set | purple |
| chr7:102423165 | 7 | 102.4232 | 0.010766295 | Credible Set | purple |
| chr7:102435692 | 7 | 102.4357 | 0.005481659 | Credible Set | purple |
| chr7:102435720 | 7 | 102.4357 | 0.005481659 | Credible Set | purple |
| chr7:102478469 | 7 | 102.4785 | 0.009373951 | Credible Set | purple |
| chr7:102510475 | 7 | 102.5105 | 0.008829658 | Credible Set | purple |
| chr7:102510607 | 7 | 102.5106 | 0.008615186 | Credible Set | purple |
| chr7:102516878 | 7 | 102.5169 | 0.009752256 | Credible Set | purple |
| chr7:102517238 | 7 | 102.5172 | 0.016944462 | Credible Set | purple |
| chr7:102517241 | 7 | 102.5172 | 0.016914451 | Credible Set | purple |
| chr7:102517247 | 7 | 102.5172 | 0.017488843 | Credible Set | purple |
| chr7:102531652 | 7 | 102.5317 | 0.008330102 | Credible Set | purple |
| chr7:102561708 | 7 | 102.5617 | 0.009182845 | Credible Set | purple |
| chr7:102565262 | 7 | 102.5653 | 0.018126622 | Credible Set | purple |
| chr7:102565292 | 7 | 102.5653 | 0.017976917 | Credible Set | purple |
| chr7:102565307 | 7 | 102.5653 | 0.018158430 | Credible Set | purple |
| chr7:102565323 | 7 | 102.5653 | 0.018008463 | Credible Set | purple |
| chr7:102565329 | 7 | 102.5653 | 0.018008463 | Credible Set | purple |
| chr7:102565410 | 7 | 102.5654 | 0.003824409 | Credible Set | purple |
| chr7:102596442 | 7 | 102.5964 | 0.006676858 | Credible Set | purple |
| chr7:102596578 | 7 | 102.5966 | 0.005007568 | Credible Set | purple |
| chr7:102596636 | 7 | 102.5966 | 0.004930290 | Credible Set | purple |
| chr7:102619418 | 7 | 102.6194 | 0.009189028 | Credible Set | purple |
| chr7:102619431 | 7 | 102.6194 | 0.009164687 | Credible Set | purple |
| chr7:102619466 | 7 | 102.6195 | 0.007013656 | Credible Set | purple |
| chr7:102757499 | 7 | 102.7575 | 0.007726660 | Credible Set | purple |
| chr7:103442570 | 7 | 103.4426 | 0.010005887 | Credible Set | purple |
| chr7:103564836 | 7 | 103.5648 | 0.009997328 | Credible Set | purple |
| chr7:103646439 | 7 | 103.6464 | 0.010250965 | Credible Set | purple |
| chr7:103646709 | 7 | 103.6467 | 0.010462968 | Credible Set | purple |
| chr7:103671033 | 7 | 103.6710 | 0.012894201 | Credible Set | purple |
| chr7:103760301 | 7 | 103.7603 | 0.020276455 | Credible Set | purple |
| chr7:103829200 | 7 | 103.8292 | 0.008920566 | Credible Set | purple |
| chr7:103837099 | 7 | 103.8371 | 0.010470456 | Credible Set | purple |
| chr7:103837213 | 7 | 103.8372 | 0.007981410 | Credible Set | purple |

Fine−mapping Regions

| snp | chr | pos (Mb) | pp | group | color |
|---|---|---|---|---|---|
| chr7:103915503 | 7 | 103.9155 | 0.009415993 | Credible Set | purple |
| chr7:103986792 | 7 | 103.9868 | 0.009194444 | Credible Set | purple |
| chr7:103986795 | 7 | 103.9868 | 0.009194444 | Credible Set | purple |
| chr7:103986857 | 7 | 103.9869 | 0.008944166 | Credible Set | purple |
| chr7:103986860 | 7 | 103.9869 | 0.008944166 | Credible Set | purple |
| chr7:104012808 | 7 | 104.0128 | 0.009113036 | Credible Set | purple |
| chr7:104013370 | 7 | 104.0134 | 0.005364078 | Credible Set | purple |
| chr7:104013395 | 7 | 104.0134 | 0.006178234 | Credible Set | purple |
| chr7:104013467 | 7 | 104.0135 | 0.007492059 | Credible Set | purple |
| chr7:104129734 | 7 | 104.1297 | 0.007593289 | Credible Set | purple |
| chr7:104129736 | 7 | 104.1297 | 0.007593289 | Credible Set | purple |
| chr7:104138934 | 7 | 104.1389 | 0.001377872 | Credible Set | purple |
| chr7:104138936 | 7 | 104.1389 | 0.001381271 | Credible Set | purple |
| chr7:104138968 | 7 | 104.1390 | 0.001381271 | Credible Set | purple |
| chr7:104141343 | 7 | 104.1413 | 0.003480258 | Credible Set | purple |
| chr7:104141705 | 7 | 104.1417 | 0.003919792 | Credible Set | purple |
| chr7:104141768 | 7 | 104.1418 | 0.003857797 | Credible Set | purple |
| chr7:104170518 | 7 | 104.1705 | 0.001363535 | Credible Set | purple |
| chr7:104170588 | 7 | 104.1706 | 0.004306815 | Credible Set | purple |
| chr7:104170594 | 7 | 104.1706 | 0.004293377 | Credible Set | purple |
| chr7:104170618 | 7 | 104.1706 | 0.004315962 | Credible Set | purple |
| chr7:104170662 | 7 | 104.1707 | 0.004315962 | Credible Set | purple |
| chr7:104338784 | 7 | 104.3388 | 0.008594047 | Credible Set | purple |
| chr7:104338810 | 7 | 104.3388 | 0.008541118 | Credible Set | purple |
| chr7:104353491 | 7 | 104.3535 | 0.013734151 | Credible Set | purple |
| chr7:104356230 | 7 | 104.3562 | 0.008810938 | Credible Set | purple |
| chr7:104430205 | 7 | 104.4302 | 0.013252966 | Credible Set | purple |
| chr7:104485236 | 7 | 104.4852 | 0.017776242 | Credible Set | purple |
| chr7:104485721 | 7 | 104.4857 | 0.002784109 | Credible Set | purple |
| chr7:104485727 | 7 | 104.4857 | 0.002878544 | Credible Set | purple |
| chr7:104486022 | 7 | 104.4860 | 0.009960934 | Credible Set | purple |
| chr7:104486041 | 7 | 104.4860 | 0.010332955 | Credible Set | purple |
| chr7:104507724 | 7 | 104.5077 | 0.028757333 | Credible Set | purple |
| chr7:104508127 | 7 | 104.5081 | 0.027279761 | Credible Set | purple |
| chr7:104508173 | 7 | 104.5082 | 0.027266602 | Credible Set | purple |
| chr7:104580091 | 7 | 104.5801 | 0.017066721 | Credible Set | purple |

Fine−mapping Regions

| snp | chr | pos (Mb) | pp | group | color |
|---|---|---|---|---|---|
| chr7:105411752 | 7 | 105.4118 | 0.006951717 | Credible Set | purple |

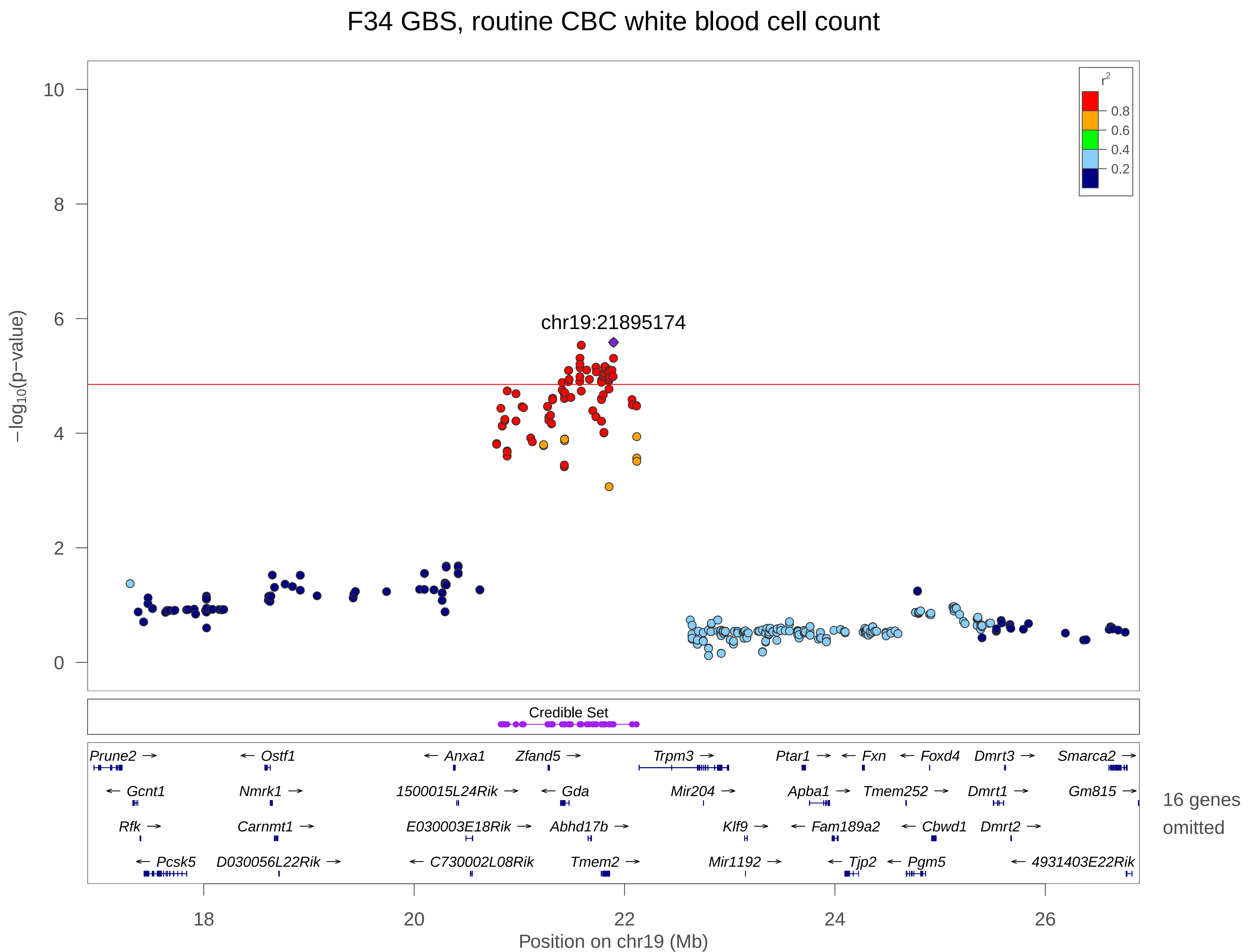
F34 GBS, routine CBC white blood cell count
−log10(p−value)
chr19:21895174
r2
0.8
0.6
0.4
0.2
Credible Set
Prune2 →
← Ostf1
← Anxa1
Zfand5 →
Trpm3 →
Ptar1 →
← Fxn
← Foxd4
Dmrt3 →
Smarca2 →
← Gcnt1
Nmrk1 →
1500015L24Rik →
← Gda
Mir204 →
Apba1 →
Tmem252 →
Dmrt1 →
Gm815 →
Rfk →
Carnmt1 →
E030003E18Rik →
Abhd17b →
Klf9 →
← Fam189a2
← Cbwd1
Dmrt2 →
← Pcsk5
D030056L22Rik →
← C730002L08Rik
Tmem2 →
Mir1192 →
← Tjp2
← Pgm5
← 4931403E22Rik
16 genes omitted
Position on chr19 (Mb)

| | |
|---|---|
| date: | Thu Oct 25 17:00:16 2018 |
| build: | F34GBS |
| display range: | chr19:16895174−26895174 [16895174−26895174] |
| hilite range: | 0 − 0 [ 0 − 0 ] |
| reference SNP: | chr19:21895174 |
| number of SNPs plotted: | 470 |
| min P−value: | 2.59E−6 [chr19:21895174] |
| max P−value: | 7.62E−1 [chr19:22798742] |
| omitted Genes: | Trpm6, Rorb, Aldh1a1 |
| omitted Genes: | Aldh1a7, Tmc1, 4930554I06Rik |
| omitted Genes: | 1110059E24Rik, Smc5, Mamdc2 |
| omitted Genes: | 1700028P14Rik, Pip5k1b, Fam122a |
| omitted Genes: | Mir3084−1, Dock8, Kank1 |
| omitted Genes: | 2610016A17Rik |

Make more plots at http://csg.sph.umich.edu/locuszoom/

Fine−mapping Regions

| snp | chr | pos (Mb) | pp | group | color |
|---|---|---|---|---|---|
| chr19:20824062 | 19 | 20.82406 | 0.003479946 | Credible Set | purple |
| chr19:20837340 | 19 | 20.83734 | 0.001766335 | Credible Set | purple |
| chr19:20837342 | 19 | 20.83734 | 0.001792071 | Credible Set | purple |
| chr19:20861811 | 19 | 20.86181 | 0.002176357 | Credible Set | purple |
| chr19:20861821 | 19 | 20.86182 | 0.002290864 | Credible Set | purple |
| chr19:20884530 | 19 | 20.88453 | 0.006761317 | Credible Set | purple |
| chr19:20967742 | 19 | 20.96774 | 0.002144878 | Credible Set | purple |
| chr19:20967772 | 19 | 20.96777 | 0.002147991 | Credible Set | purple |
| chr19:20967840 | 19 | 20.96784 | 0.006083086 | Credible Set | purple |
| chr19:20967865 | 19 | 20.96786 | 0.006093708 | Credible Set | purple |
| chr19:21025668 | 19 | 21.02567 | 0.003706715 | Credible Set | purple |
| chr19:21039744 | 19 | 21.03974 | 0.003568918 | Credible Set | purple |
| chr19:21268525 | 19 | 21.26853 | 0.003719266 | Credible Set | purple |
| chr19:21268527 | 19 | 21.26853 | 0.003726990 | Credible Set | purple |
| chr19:21268550 | 19 | 21.26855 | 0.003758563 | Credible Set | purple |
| chr19:21280282 | 19 | 21.28028 | 0.002478884 | Credible Set | purple |
| chr19:21280286 | 19 | 21.28029 | 0.002210847 | Credible Set | purple |
| chr19:21296098 | 19 | 21.29610 | 0.002672970 | Credible Set | purple |
| chr19:21305518 | 19 | 21.30552 | 0.001923346 | Credible Set | purple |
| chr19:21305568 | 19 | 21.30557 | 0.001952204 | Credible Set | purple |
| chr19:21316234 | 19 | 21.31623 | 0.005121764 | Credible Set | purple |
| chr19:21316268 | 19 | 21.31627 | 0.004811437 | Credible Set | purple |
| chr19:21316305 | 19 | 21.31630 | 0.004823994 | Credible Set | purple |
| chr19:21407108 | 19 | 21.40711 | 0.006981278 | Credible Set | purple |
| chr19:21407182 | 19 | 21.40718 | 0.009324381 | Credible Set | purple |
| chr19:21428151 | 19 | 21.42815 | 0.005962054 | Credible Set | purple |
| chr19:21428338 | 19 | 21.42834 | 0.005063110 | Credible Set | purple |
| chr19:21428343 | 19 | 21.42834 | 0.005063110 | Credible Set | purple |
| chr19:21430361 | 19 | 21.43036 | 0.006292518 | Credible Set | purple |
| chr19:21430412 | 19 | 21.43041 | 0.006353994 | Credible Set | purple |
| chr19:21467882 | 19 | 21.46788 | 0.009646199 | Credible Set | purple |
| chr19:21467967 | 19 | 21.46797 | 0.014776152 | Credible Set | purple |
| chr19:21467982 | 19 | 21.46798 | 0.014850830 | Credible Set | purple |
| chr19:21467985 | 19 | 21.46798 | 0.014850830 | Credible Set | purple |
| chr19:21467987 | 19 | 21.46799 | 0.014813541 | Credible Set | purple |
| chr19:21472509 | 19 | 21.47251 | 0.010549821 | Credible Set | purple |

Fine−mapping Regions

| snp | chr | pos (Mb) | pp | group | color |
|---|---|---|---|---|---|
| chr19:21472526 | 19 | 21.47253 | 0.010517116 | Credible Set | purple |
| chr19:21489550 | 19 | 21.48955 | 0.005257509 | Credible Set | purple |
| chr19:21489579 | 19 | 21.48958 | 0.005292122 | Credible Set | purple |
| chr19:21574198 | 19 | 21.57420 | 0.009713901 | Credible Set | purple |
| chr19:21574202 | 19 | 21.57420 | 0.009748026 | Credible Set | purple |
| chr19:21575782 | 19 | 21.57578 | 0.011679622 | Credible Set | purple |
| chr19:21576364 | 19 | 21.57636 | 0.023787792 | Credible Set | purple |
| chr19:21576365 | 19 | 21.57636 | 0.016462767 | Credible Set | purple |
| chr19:21576374 | 19 | 21.57637 | 0.018862607 | Credible Set | purple |
| chr19:21588462 | 19 | 21.58846 | 0.039243530 | Credible Set | purple |
| chr19:21588470 | 19 | 21.58847 | 0.039191513 | Credible Set | purple |
| chr19:21588527 | 19 | 21.58853 | 0.038982712 | Credible Set | purple |
| chr19:21588528 | 19 | 21.58853 | 0.039275032 | Credible Set | purple |
| chr19:21588608 | 19 | 21.58861 | 0.006724028 | Credible Set | purple |
| chr19:21640403 | 19 | 21.64040 | 0.015070885 | Credible Set | purple |
| chr19:21666239 | 19 | 21.66624 | 0.010518703 | Credible Set | purple |
| chr19:21697884 | 19 | 21.69788 | 0.003177776 | Credible Set | purple |
| chr19:21727318 | 19 | 21.72732 | 0.002543642 | Credible Set | purple |
| chr19:21727391 | 19 | 21.72739 | 0.002520819 | Credible Set | purple |
| chr19:21727561 | 19 | 21.72756 | 0.016824376 | Credible Set | purple |
| chr19:21731633 | 19 | 21.73163 | 0.014019582 | Credible Set | purple |
| chr19:21780788 | 19 | 21.78079 | 0.010236000 | Credible Set | purple |
| chr19:21780850 | 19 | 21.78085 | 0.009426365 | Credible Set | purple |
| chr19:21780922 | 19 | 21.78092 | 0.002127112 | Credible Set | purple |
| chr19:21780928 | 19 | 21.78093 | 0.002124246 | Credible Set | purple |
| chr19:21780934 | 19 | 21.78093 | 0.002134882 | Credible Set | purple |
| chr19:21780935 | 19 | 21.78093 | 0.002134882 | Credible Set | purple |
| chr19:21781005 | 19 | 21.78101 | 0.005043833 | Credible Set | purple |
| chr19:21781019 | 19 | 21.78102 | 0.004855973 | Credible Set | purple |
| chr19:21798450 | 19 | 21.79845 | 0.011913874 | Credible Set | purple |
| chr19:21798541 | 19 | 21.79854 | 0.005838403 | Credible Set | purple |
| chr19:21803806 | 19 | 21.80381 | 0.001393950 | Credible Set | purple |
| chr19:21803819 | 19 | 21.80382 | 0.001362329 | Credible Set | purple |
| chr19:21803853 | 19 | 21.80385 | 0.001388858 | Credible Set | purple |
| chr19:21803859 | 19 | 21.80386 | 0.001390581 | Credible Set | purple |
| chr19:21804356 | 19 | 21.80436 | 0.012053895 | Credible Set | purple |

Fine−mapping Regions

| snp | chr | pos (Mb) | pp | group | color |
|---|---|---|---|---|---|
| chr19:21804420 | 19 | 21.80442 | 0.013442321 | Credible Set | purple |
| chr19:21804582 | 19 | 21.80458 | 0.013518135 | Credible Set | purple |
| chr19:21812258 | 19 | 21.81226 | 0.015579607 | Credible Set | purple |
| chr19:21812298 | 19 | 21.81230 | 0.015638761 | Credible Set | purple |
| chr19:21812594 | 19 | 21.81259 | 0.017218328 | Credible Set | purple |
| chr19:21814790 | 19 | 21.81479 | 0.017124033 | Credible Set | purple |
| chr19:21814806 | 19 | 21.81481 | 0.017236345 | Credible Set | purple |
| chr19:21851795 | 19 | 21.85179 | 0.009981769 | Credible Set | purple |
| chr19:21852236 | 19 | 21.85224 | 0.011199702 | Credible Set | purple |
| chr19:21852273 | 19 | 21.85227 | 0.010492163 | Credible Set | purple |
| chr19:21852296 | 19 | 21.85230 | 0.007290938 | Credible Set | purple |
| chr19:21857059 | 19 | 21.85706 | 0.011777903 | Credible Set | purple |
| chr19:21857128 | 19 | 21.85713 | 0.011292658 | Credible Set | purple |
| chr19:21857704 | 19 | 21.85770 | 0.015183882 | Credible Set | purple |
| chr19:21857707 | 19 | 21.85771 | 0.014478042 | Credible Set | purple |
| chr19:21880097 | 19 | 21.88010 | 0.014906049 | Credible Set | purple |
| chr19:21889764 | 19 | 21.88976 | 0.011703019 | Credible Set | purple |
| chr19:21889768 | 19 | 21.88977 | 0.011733833 | Credible Set | purple |
| chr19:21889777 | 19 | 21.88978 | 0.011703019 | Credible Set | purple |
| chr19:21889831 | 19 | 21.88983 | 0.011723340 | Credible Set | purple |
| chr19:21889832 | 19 | 21.88983 | 0.011730755 | Credible Set | purple |
| chr19:21889838 | 19 | 21.88984 | 0.011882834 | Credible Set | purple |
| chr19:21895163 | 19 | 21.89516 | 0.023703299 | Credible Set | purple |
| chr19:21895174 | 19 | 21.89517 | 0.043781690 | Credible Set | purple |
| chr19:22070939 | 19 | 22.07094 | 0.004872101 | Credible Set | purple |
| chr19:22073618 | 19 | 22.07362 | 0.003957677 | Credible Set | purple |
| chr19:22113626 | 19 | 22.11363 | 0.003856117 | Credible Set | purple |
| chr19:22113628 | 19 | 22.11363 | 0.003806433 | Credible Set | purple |
| chr19:22113687 | 19 | 22.11369 | 0.003804155 | Credible Set | purple |

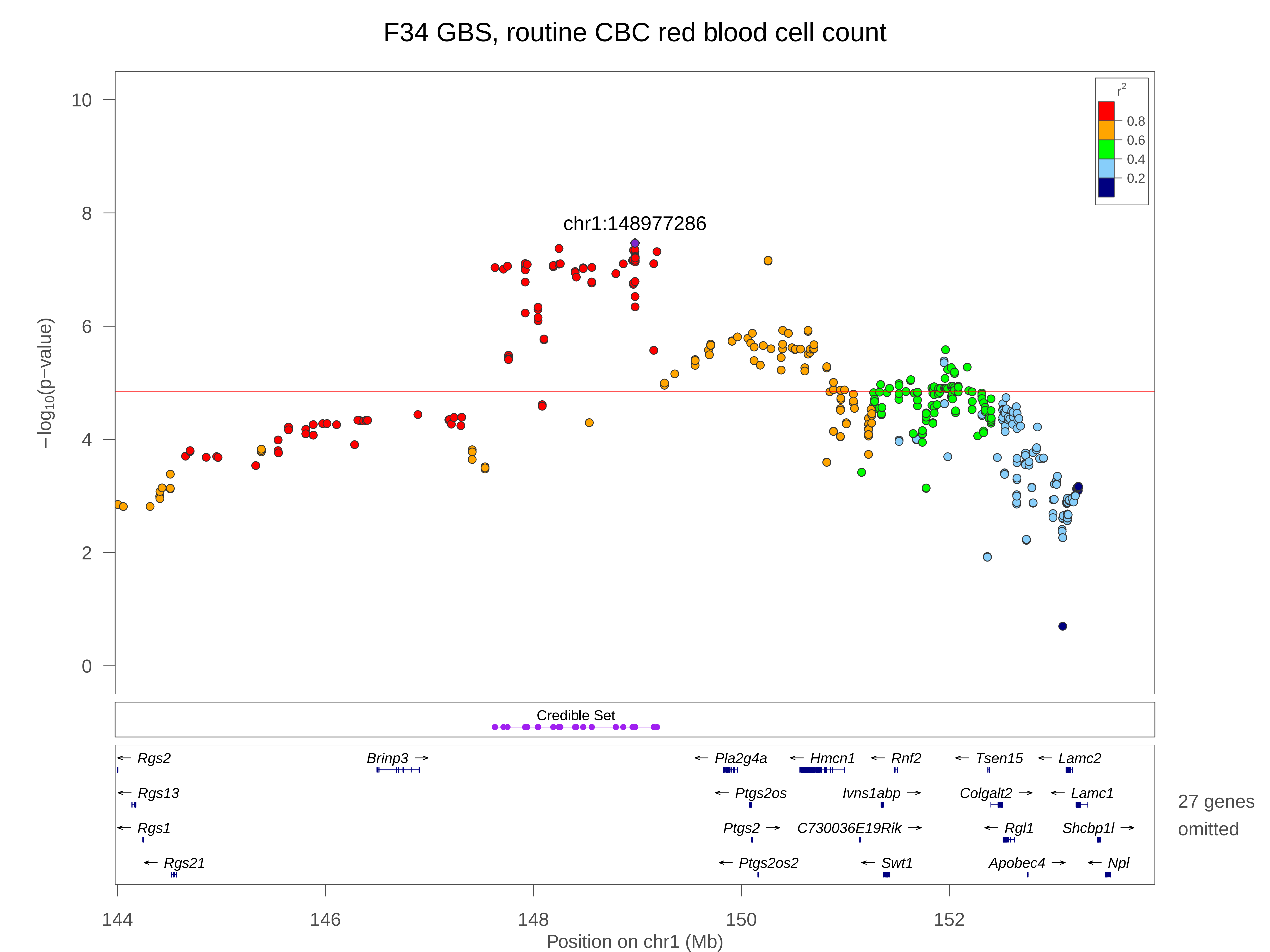

F34 GBS, routine CBC red blood cell count
r2
0.8
0.6
0.4
0.2
chr1:148977286
10
8
6
4
2
0
−log10(p−value)
Credible Set
Rgs2
Rgs13
Rgs1
Rgs21
Brinp3
Pla2g4a
Ptgs2os
Ptgs2
Ptgs2os2
Hmcn1
C730036E19Rik
Ivns1abp
Rnf2
Swt1
Tsen15
Colgalt2
Rgl1
Apobec4
Lamc2
Lamc1
Shcbp1l
Npl
27 genes omitted
144
146
148
150
152
Position on chr1 (Mb)

| | |
|---|---|
| date: | Thu Oct 25 17:03:27 2018 |
| build: | F34GBS |
| display range: | chr1:143977286−153977286 [143977286−153977286] |
| hilite range: | 0 − 0 [ 0 − 0 ] |
| reference SNP: | chr1:148977286 |
| number of SNPs plotted: | 507 |
| min P−value: | 3.42E−8 [chr1:148977286] |
| max P−value: | 2E−1 [chr1:153090942] |
| omitted Genes: | Rgs18, Pdc, 2310030A07Rik |
| omitted Genes: | BC003331, Tpr, Prg4 |
| omitted Genes: | Trmt1l, Mir7682, Fam129a |
| omitted Genes: | Edem3, 1700025G04Rik, Arpc5 |
| omitted Genes: | Ncf2, Smg7, Nmnat2 |
| omitted Genes: | E330020D12Rik, Dhx9, Rgs8 |
| omitted Genes: | C230024C17Rik, Rgs16, Rnasel |
| omitted Genes: | Rgsl1, Teddm2, Teddm1b |
| omitted Genes: | Teddm1a, Mir8114, Glul |

Make more plots at http://csg.sph.umich.edu/locuszoom/

Fine−mapping Regions

| snp | chr | pos (Mb) | pp | group | color |
|---|---|---|---|---|---|
| chr1:147630646 | 1 | 147.6306 | 0.016770111 | Credible Set | purple |
| chr1:147712367 | 1 | 147.7124 | 0.015760618 | Credible Set | purple |
| chr1:147751084 | 1 | 147.7511 | 0.017691123 | Credible Set | purple |
| chr1:147921174 | 1 | 147.9212 | 0.009486611 | Credible Set | purple |
| chr1:147921280 | 1 | 147.9213 | 0.002813986 | Credible Set | purple |
| chr1:147921379 | 1 | 147.9214 | 0.018112685 | Credible Set | purple |
| chr1:147921380 | 1 | 147.9214 | 0.018157020 | Credible Set | purple |
| chr1:147921465 | 1 | 147.9215 | 0.019651103 | Credible Set | purple |
| chr1:147921979 | 1 | 147.9220 | 0.015252863 | Credible Set | purple |
| chr1:147940897 | 1 | 147.9409 | 0.019030575 | Credible Set | purple |
| chr1:148044707 | 1 | 148.0447 | 0.003235533 | Credible Set | purple |
| chr1:148044765 | 1 | 148.0448 | 0.003542318 | Credible Set | purple |
| chr1:148045260 | 1 | 148.0453 | 0.002364531 | Credible Set | purple |
| chr1:148191398 | 1 | 148.1914 | 0.017338943 | Credible Set | purple |
| chr1:148191461 | 1 | 148.1915 | 0.018222125 | Credible Set | purple |
| chr1:148246268 | 1 | 148.2463 | 0.035525322 | Credible Set | purple |
| chr1:148246270 | 1 | 148.2463 | 0.035553321 | Credible Set | purple |
| chr1:148246308 | 1 | 148.2463 | 0.035553321 | Credible Set | purple |
| chr1:148246370 | 1 | 148.2464 | 0.019228793 | Credible Set | purple |
| chr1:148259002 | 1 | 148.2590 | 0.019519378 | Credible Set | purple |
| chr1:148401594 | 1 | 148.4016 | 0.014359338 | Credible Set | purple |
| chr1:148401658 | 1 | 148.4017 | 0.013724217 | Credible Set | purple |
| chr1:148412829 | 1 | 148.4128 | 0.011526073 | Credible Set | purple |
| chr1:148479166 | 1 | 148.4792 | 0.016637040 | Credible Set | purple |
| chr1:148479488 | 1 | 148.4795 | 0.016101327 | Credible Set | purple |
| chr1:148561626 | 1 | 148.5616 | 0.016884578 | Credible Set | purple |
| chr1:148561643 | 1 | 148.5616 | 0.009132927 | Credible Set | purple |
| chr1:148561679 | 1 | 148.5617 | 0.009533432 | Credible Set | purple |
| chr1:148793161 | 1 | 148.7932 | 0.013149282 | Credible Set | purple |
| chr1:148793216 | 1 | 148.7932 | 0.013271295 | Credible Set | purple |
| chr1:148864191 | 1 | 148.8642 | 0.019448661 | Credible Set | purple |
| chr1:148954162 | 1 | 148.9542 | 0.022275747 | Credible Set | purple |
| chr1:148954168 | 1 | 148.9542 | 0.022275747 | Credible Set | purple |
| chr1:148954207 | 1 | 148.9542 | 0.022275747 | Credible Set | purple |
| chr1:148961834 | 1 | 148.9618 | 0.008683184 | Credible Set | purple |
| chr1:148961852 | 1 | 148.9619 | 0.009084529 | Credible Set | purple |

Fine−mapping Regions

| snp | chr | pos (Mb) | pp | group | color |
|---|---|---|---|---|---|
| chr1:148962391 | 1 | 148.9624 | 0.022799495 | Credible Set | purple |
| chr1:148962448 | 1 | 148.9624 | 0.022489334 | Credible Set | purple |
| chr1:148962557 | 1 | 148.9626 | 0.033096424 | Credible Set | purple |
| chr1:148977255 | 1 | 148.9773 | 0.031479271 | Credible Set | purple |
| chr1:148977279 | 1 | 148.9773 | 0.033707946 | Credible Set | purple |
| chr1:148977286 | 1 | 148.9773 | 0.043873858 | Credible Set | purple |
| chr1:148977945 | 1 | 148.9779 | 0.009703785 | Credible Set | purple |
| chr1:148977976 | 1 | 148.9780 | 0.009699202 | Credible Set | purple |
| chr1:148977977 | 1 | 148.9780 | 0.003578461 | Credible Set | purple |
| chr1:148977984 | 1 | 148.9780 | 0.005377246 | Credible Set | purple |
| chr1:148978648 | 1 | 148.9786 | 0.025925582 | Credible Set | purple |
| chr1:148978689 | 1 | 148.9787 | 0.025159514 | Credible Set | purple |
| chr1:148978796 | 1 | 148.9788 | 0.024447543 | Credible Set | purple |
| chr1:148978842 | 1 | 148.9788 | 0.020973256 | Credible Set | purple |
| chr1:148978923 | 1 | 148.9789 | 0.022403352 | Credible Set | purple |
| chr1:148979220 | 1 | 148.9792 | 0.024483940 | Credible Set | purple |
| chr1:149157510 | 1 | 149.1575 | 0.019554960 | Credible Set | purple |
| chr1:149188240 | 1 | 149.1882 | 0.031394233 | Credible Set | purple |

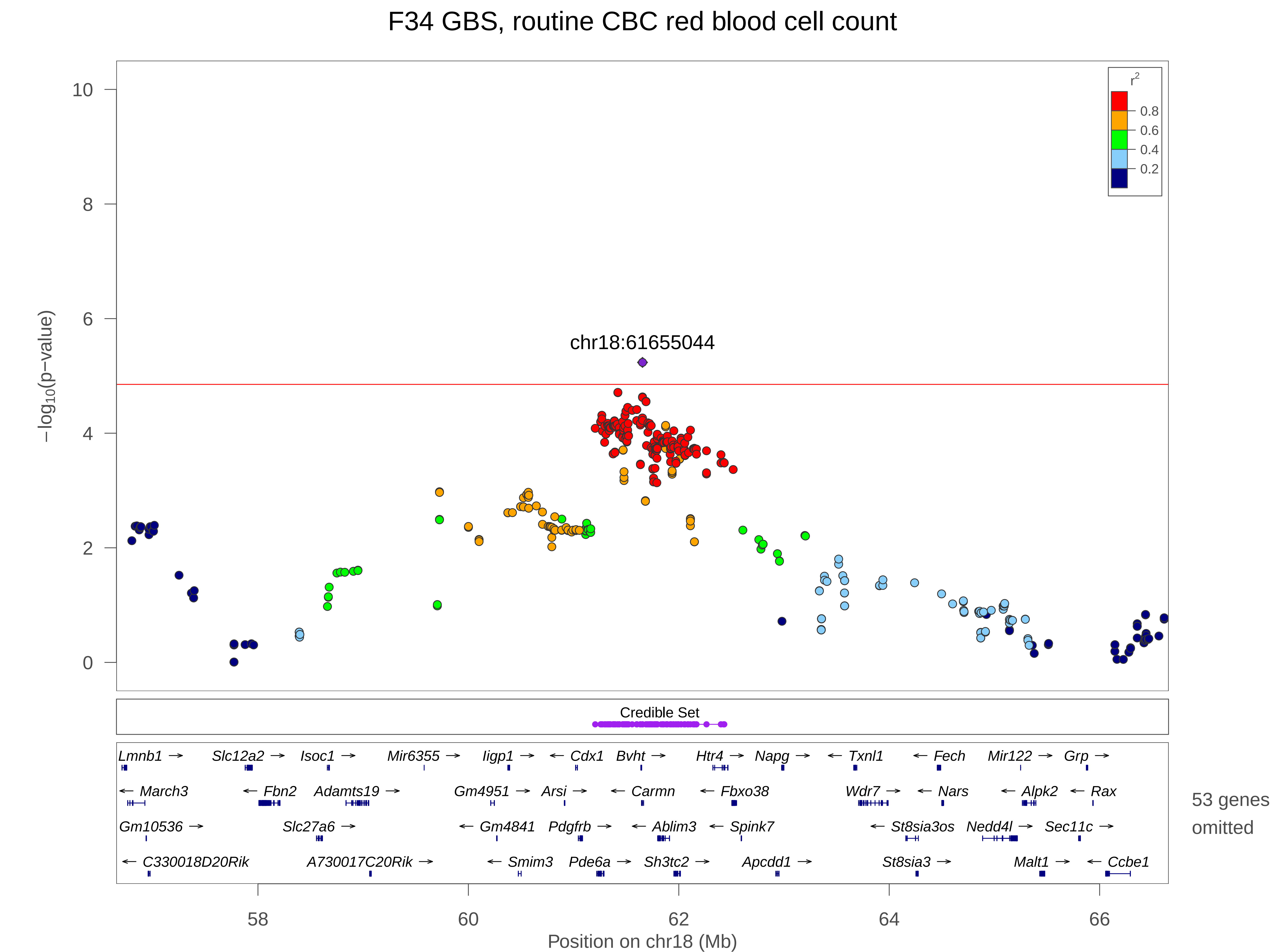
F34 GBS, routine CBC red blood cell count
−log10(p−value)
10
8
6
4
2
0
r2
0.8
0.6
0.4
0.2
chr18:61655044
Credible Set
Lmnb1 →
Slc12a2 →
Isoc1 →
Mir6355 →
Iigp1 →
← Cdx1
Bvht →
Htr4 →
Napg →
← Txnl1
← Fech
Mir122 →
Grp →
← March3
← Fbn2
Adamts19 →
Gm4951 →
Arsi →
← Carmn
← Fbxo38
Wdr7 →
← Nars
← Alpk2
← Rax
Gm10536 →
Slc27a6 →
← Gm4841
Pdgfrb →
← Ablim3
← Spink7
← St8sia3os
Nedd4l →
Sec11c →
← C330018D20Rik
A730017C20Rik →
← Smim3
Pde6a →
Sh3tc2 →
Apcdd1 →
St8sia3 →
Malt1 →
← Ccbe1
53 genes omitted
58
60
62
64
66
Position on chr18 (Mb)

| | |
|---|---|
| date: | Thu Oct 25 17:03:38 2018 |
| build: | F34GBS |
| display range: | chr18:56655044−66655044 [56655044−66655044] |
| hilite range: | 0 − 0 [ 0 − 0 ] |
| reference SNP: | chr18:61655044 |
| number of SNPs plotted: | 476 |
| min P−value: | 5.81E−6 [chr18:61655044] |
| max P−value: | 9.88E−1 [chr18:57772170] |
| omitted Genes: | Megf10, Prrc1, Ctxn3 |
| omitted Genes: | 1700066O22Rik, 1700011I03Rik, 4930511M06Rik |
| omitted Genes: | 2210409D07Rik, Chsy3, F830016B08Rik |
| omitted Genes: | Dctn4, Rbm22, Myoz3 |
| omitted Genes: | Synpo, G630071F17Rik, Ndst1 |
| omitted Genes: | Rps14, Cd74, Mir5107 |
| omitted Genes: | Tcof1, Camk2a, Mir7220 |
| omitted Genes: | Mir6982, Slc6a7, Csf1r |
| omitted Genes: | Mir6983, Hmgxb3, Slc26a2 |
| omitted Genes: | Ppargc1b, Mir378a, A530050N04Rik |
| omitted Genes: | Arhgef37, Csnk1a1, Mir145a |
| omitted Genes: | Mir143, Il17b, Pcyox1l |
| omitted Genes: | Grpel2, 1500015A07Rik, Afap1l1 |
| omitted Genes: | Adrb2, Spink10, Spink13 |
| omitted Genes: | 2700046A07Rik, Piezo2, Onecut2 |
| omitted Genes: | Atp8b1, Zfp532, Oacyl |
| omitted Genes: | Cplx4, Lman1, Mir694 |
| omitted Genes: | Pmaip1, Gm9926 |

Make more plots at http://csg.sph.umich.edu/locuszoom/

Fine−mapping Regions

| snp | chr | pos (Mb) | pp | group | color |
|---|---|---|---|---|---|
| chr18:61205705 | 18 | 61.20571 | 0.005448370 | Credible Set | purple |
| chr18:61257201 | 18 | 61.25720 | 0.006945231 | Credible Set | purple |
| chr18:61257377 | 18 | 61.25738 | 0.006966032 | Credible Set | purple |
| chr18:61268578 | 18 | 61.26858 | 0.008962725 | Credible Set | purple |
| chr18:61268584 | 18 | 61.26858 | 0.007777763 | Credible Set | purple |
| chr18:61275165 | 18 | 61.27517 | 0.004888103 | Credible Set | purple |
| chr18:61294899 | 18 | 61.29490 | 0.003202445 | Credible Set | purple |
| chr18:61295744 | 18 | 61.29574 | 0.005984639 | Credible Set | purple |
| chr18:61307248 | 18 | 61.30725 | 0.004362501 | Credible Set | purple |
| chr18:61307278 | 18 | 61.30728 | 0.004369899 | Credible Set | purple |
| chr18:61322420 | 18 | 61.32242 | 0.006295605 | Credible Set | purple |
| chr18:61325885 | 18 | 61.32588 | 0.005969946 | Credible Set | purple |
| chr18:61325983 | 18 | 61.32598 | 0.006591703 | Credible Set | purple |
| chr18:61326493 | 18 | 61.32649 | 0.006001677 | Credible Set | purple |
| chr18:61326662 | 18 | 61.32666 | 0.005943314 | Credible Set | purple |
| chr18:61326689 | 18 | 61.32669 | 0.005943314 | Credible Set | purple |
| chr18:61336840 | 18 | 61.33684 | 0.005959787 | Credible Set | purple |
| chr18:61340720 | 18 | 61.34072 | 0.004978988 | Credible Set | purple |
| chr18:61340729 | 18 | 61.34073 | 0.004985735 | Credible Set | purple |
| chr18:61344728 | 18 | 61.34473 | 0.005558116 | Credible Set | purple |
| chr18:61351043 | 18 | 61.35104 | 0.005735564 | Credible Set | purple |
| chr18:61351491 | 18 | 61.35149 | 0.005563928 | Credible Set | purple |
| chr18:61376042 | 18 | 61.37604 | 0.002083723 | Credible Set | purple |
| chr18:61376045 | 18 | 61.37604 | 0.002067396 | Credible Set | purple |
| chr18:61376483 | 18 | 61.37648 | 0.006195200 | Credible Set | purple |
| chr18:61376688 | 18 | 61.37669 | 0.005969946 | Credible Set | purple |
| chr18:61376841 | 18 | 61.37684 | 0.005969946 | Credible Set | purple |
| chr18:61376921 | 18 | 61.37692 | 0.005969946 | Credible Set | purple |
| chr18:61382329 | 18 | 61.38233 | 0.006741293 | Credible Set | purple |
| chr18:61382496 | 18 | 61.38250 | 0.005969946 | Credible Set | purple |
| chr18:61386134 | 18 | 61.38613 | 0.005697101 | Credible Set | purple |
| chr18:61386274 | 18 | 61.38627 | 0.005969946 | Credible Set | purple |
| chr18:61388160 | 18 | 61.38816 | 0.006259177 | Credible Set | purple |
| chr18:61388169 | 18 | 61.38817 | 0.007260709 | Credible Set | purple |
| chr18:61388197 | 18 | 61.38820 | 0.007148277 | Credible Set | purple |
| chr18:61389384 | 18 | 61.38938 | 0.005988841 | Credible Set | purple |

Fine−mapping Regions

| snp | chr | pos (Mb) | pp | group | color |
|---|---|---|---|---|---|
| chr18:61392736 | 18 | 61.39274 | 0.002216639 | Credible Set | purple |
| chr18:61392737 | 18 | 61.39274 | 0.002216639 | Credible Set | purple |
| chr18:61392782 | 18 | 61.39278 | 0.002216639 | Credible Set | purple |
| chr18:61392828 | 18 | 61.39283 | 0.002171816 | Credible Set | purple |
| chr18:61392844 | 18 | 61.39284 | 0.002171816 | Credible Set | purple |
| chr18:61400213 | 18 | 61.40021 | 0.005969946 | Credible Set | purple |
| chr18:61415978 | 18 | 61.41598 | 0.006348102 | Credible Set | purple |
| chr18:61421022 | 18 | 61.42102 | 0.021218291 | Credible Set | purple |
| chr18:61421081 | 18 | 61.42108 | 0.021462381 | Credible Set | purple |
| chr18:61434374 | 18 | 61.43437 | 0.005567321 | Credible Set | purple |
| chr18:61435019 | 18 | 61.43502 | 0.004474155 | Credible Set | purple |
| chr18:61435052 | 18 | 61.43505 | 0.004336106 | Credible Set | purple |
| chr18:61466355 | 18 | 61.46636 | 0.007011180 | Credible Set | purple |
| chr18:61466435 | 18 | 61.46643 | 0.004066953 | Credible Set | purple |
| chr18:61466477 | 18 | 61.46648 | 0.004009173 | Credible Set | purple |
| chr18:61466745 | 18 | 61.46675 | 0.003794946 | Credible Set | purple |
| chr18:61475793 | 18 | 61.47579 | 0.005003903 | Credible Set | purple |
| chr18:61475799 | 18 | 61.47580 | 0.005003903 | Credible Set | purple |
| chr18:61475844 | 18 | 61.47584 | 0.005582395 | Credible Set | purple |
| chr18:61488563 | 18 | 61.48856 | 0.003606624 | Credible Set | purple |
| chr18:61488568 | 18 | 61.48857 | 0.003606624 | Credible Set | purple |
| chr18:61488569 | 18 | 61.48857 | 0.003606624 | Credible Set | purple |
| chr18:61488585 | 18 | 61.48859 | 0.005969946 | Credible Set | purple |
| chr18:61488640 | 18 | 61.48864 | 0.008877543 | Credible Set | purple |
| chr18:61488644 | 18 | 61.48864 | 0.008877543 | Credible Set | purple |
| chr18:61497425 | 18 | 61.49742 | 0.010461353 | Credible Set | purple |
| chr18:61502586 | 18 | 61.50259 | 0.004116200 | Credible Set | purple |
| chr18:61505684 | 18 | 61.50568 | 0.003667400 | Credible Set | purple |
| chr18:61505787 | 18 | 61.50579 | 0.003694007 | Credible Set | purple |
| chr18:61506314 | 18 | 61.50631 | 0.004005741 | Credible Set | purple |
| chr18:61506364 | 18 | 61.50636 | 0.003249537 | Credible Set | purple |
| chr18:61506393 | 18 | 61.50639 | 0.003338023 | Credible Set | purple |
| chr18:61513675 | 18 | 61.51367 | 0.005053807 | Credible Set | purple |
| chr18:61513686 | 18 | 61.51369 | 0.005121434 | Credible Set | purple |
| chr18:61514087 | 18 | 61.51409 | 0.012051866 | Credible Set | purple |
| chr18:61514093 | 18 | 61.51409 | 0.012088210 | Credible Set | purple |

Fine−mapping Regions

| snp | chr | pos (Mb) | pp | group | color |
|---|---|---|---|---|---|
| chr18:61518663 | 18 | 61.51866 | 0.006551351 | Credible Set | purple |
| chr18:61522253 | 18 | 61.52225 | 0.004134208 | Credible Set | purple |
| chr18:61522261 | 18 | 61.52226 | 0.004081616 | Credible Set | purple |
| chr18:61556863 | 18 | 61.55686 | 0.010791463 | Credible Set | purple |
| chr18:61600372 | 18 | 61.60037 | 0.007326827 | Credible Set | purple |
| chr18:61600381 | 18 | 61.60038 | 0.007297248 | Credible Set | purple |
| chr18:61600401 | 18 | 61.60040 | 0.007302610 | Credible Set | purple |
| chr18:61600428 | 18 | 61.60043 | 0.007315662 | Credible Set | purple |
| chr18:61600547 | 18 | 61.60055 | 0.011113437 | Credible Set | purple |
| chr18:61600608 | 18 | 61.60061 | 0.011132748 | Credible Set | purple |
| chr18:61634745 | 18 | 61.63475 | 0.006181847 | Credible Set | purple |
| chr18:61634869 | 18 | 61.63487 | 0.006592431 | Credible Set | purple |
| chr18:61635121 | 18 | 61.63512 | 0.001412198 | Credible Set | purple |
| chr18:61635169 | 18 | 61.63517 | 0.001379070 | Credible Set | purple |
| chr18:61653762 | 18 | 61.65376 | 0.008072224 | Credible Set | purple |
| chr18:61654922 | 18 | 61.65492 | 0.008071189 | Credible Set | purple |
| chr18:61654978 | 18 | 61.65498 | 0.018033908 | Credible Set | purple |
| chr18:61655005 | 18 | 61.65501 | 0.017658601 | Credible Set | purple |
| chr18:61655044 | 18 | 61.65504 | 0.067766511 | Credible Set | purple |
| chr18:61655445 | 18 | 61.65545 | 0.007365164 | Credible Set | purple |
| chr18:61688361 | 18 | 61.68836 | 0.015176045 | Credible Set | purple |
| chr18:61688363 | 18 | 61.68836 | 0.015252225 | Credible Set | purple |
| chr18:61688400 | 18 | 61.68840 | 0.015014264 | Credible Set | purple |
| chr18:61693349 | 18 | 61.69335 | 0.002842604 | Credible Set | purple |
| chr18:61706715 | 18 | 61.70672 | 0.004681497 | Credible Set | purple |
| chr18:61707062 | 18 | 61.70706 | 0.006397523 | Credible Set | purple |
| chr18:61707121 | 18 | 61.70712 | 0.006710377 | Credible Set | purple |
| chr18:61722238 | 18 | 61.72224 | 0.006544158 | Credible Set | purple |
| chr18:61722262 | 18 | 61.72226 | 0.005921844 | Credible Set | purple |
| chr18:61726167 | 18 | 61.72617 | 0.005999756 | Credible Set | purple |
| chr18:61726194 | 18 | 61.72619 | 0.006024756 | Credible Set | purple |
| chr18:61726206 | 18 | 61.72621 | 0.005999756 | Credible Set | purple |
| chr18:61726239 | 18 | 61.72624 | 0.005859850 | Credible Set | purple |
| chr18:61737597 | 18 | 61.73760 | 0.002605931 | Credible Set | purple |
| chr18:61737612 | 18 | 61.73761 | 0.002596645 | Credible Set | purple |
| chr18:61737626 | 18 | 61.73763 | 0.002587908 | Credible Set | purple |

Fine−mapping Regions

| snp | chr | pos (Mb) | pp | group | color |
|---|---|---|---|---|---|
| chr18:61737739 | 18 | 61.73774 | 0.006042115 | Credible Set | purple |
| chr18:61752870 | 18 | 61.75287 | 0.002491824 | Credible Set | purple |
| chr18:61752916 | 18 | 61.75292 | 0.002042854 | Credible Set | purple |
| chr18:61766474 | 18 | 61.76647 | 0.003275071 | Credible Set | purple |
| chr18:61773121 | 18 | 61.77312 | 0.002695951 | Credible Set | purple |
| chr18:61773529 | 18 | 61.77353 | 0.002488509 | Credible Set | purple |
| chr18:61773667 | 18 | 61.77367 | 0.001992165 | Credible Set | purple |
| chr18:61773688 | 18 | 61.77369 | 0.001957495 | Credible Set | purple |
| chr18:61773748 | 18 | 61.77375 | 0.001957495 | Credible Set | purple |
| chr18:61774150 | 18 | 61.77415 | 0.002437604 | Credible Set | purple |
| chr18:61774277 | 18 | 61.77428 | 0.002469121 | Credible Set | purple |
| chr18:61785401 | 18 | 61.78540 | 0.002463153 | Credible Set | purple |
| chr18:61785450 | 18 | 61.78545 | 0.002451525 | Credible Set | purple |
| chr18:61785465 | 18 | 61.78547 | 0.002445143 | Credible Set | purple |
| chr18:61786082 | 18 | 61.78608 | 0.002443676 | Credible Set | purple |
| chr18:61786403 | 18 | 61.78640 | 0.002551589 | Credible Set | purple |
| chr18:61790408 | 18 | 61.79041 | 0.002564550 | Credible Set | purple |
| chr18:61790488 | 18 | 61.79049 | 0.002364411 | Credible Set | purple |
| chr18:61791422 | 18 | 61.79142 | 0.002508820 | Credible Set | purple |
| chr18:61792239 | 18 | 61.79224 | 0.001755982 | Credible Set | purple |
| chr18:61792326 | 18 | 61.79233 | 0.002617102 | Credible Set | purple |
| chr18:61795073 | 18 | 61.79507 | 0.003692253 | Credible Set | purple |
| chr18:61795217 | 18 | 61.79522 | 0.002538919 | Credible Set | purple |
| chr18:61795218 | 18 | 61.79522 | 0.002538919 | Credible Set | purple |
| chr18:61795225 | 18 | 61.79523 | 0.002538919 | Credible Set | purple |
| chr18:61795296 | 18 | 61.79530 | 0.004117110 | Credible Set | purple |
| chr18:61795298 | 18 | 61.79530 | 0.004350738 | Credible Set | purple |
| chr18:61832086 | 18 | 61.83209 | 0.003770205 | Credible Set | purple |
| chr18:61847573 | 18 | 61.84757 | 0.003388558 | Credible Set | purple |
| chr18:61847582 | 18 | 61.84758 | 0.003248936 | Credible Set | purple |
| chr18:61847690 | 18 | 61.84769 | 0.003151108 | Credible Set | purple |
| chr18:61858191 | 18 | 61.85819 | 0.003184572 | Credible Set | purple |
| chr18:61882775 | 18 | 61.88278 | 0.003208489 | Credible Set | purple |
| chr18:61882784 | 18 | 61.88278 | 0.003204335 | Credible Set | purple |
| chr18:61882815 | 18 | 61.88282 | 0.003204335 | Credible Set | purple |
| chr18:61882830 | 18 | 61.88283 | 0.003204335 | Credible Set | purple |

Fine−mapping Regions

| snp | chr | pos (Mb) | pp | group | color |
|---|---|---|---|---|---|
| chr18:61890944 | 18 | 61.89094 | 0.004054650 | Credible Set | purple |
| chr18:61891389 | 18 | 61.89139 | 0.003143261 | Credible Set | purple |
| chr18:61891400 | 18 | 61.89140 | 0.003273676 | Credible Set | purple |
| chr18:61917922 | 18 | 61.91792 | 0.002036727 | Credible Set | purple |
| chr18:61921624 | 18 | 61.92162 | 0.002492555 | Credible Set | purple |
| chr18:61921661 | 18 | 61.92166 | 0.002604104 | Credible Set | purple |
| chr18:61921694 | 18 | 61.92169 | 0.002562519 | Credible Set | purple |
| chr18:61922273 | 18 | 61.92227 | 0.001533529 | Credible Set | purple |
| chr18:61927547 | 18 | 61.92755 | 0.002698341 | Credible Set | purple |
| chr18:61936945 | 18 | 61.93695 | 0.003369307 | Credible Set | purple |
| chr18:61949121 | 18 | 61.94912 | 0.002893394 | Credible Set | purple |
| chr18:61952249 | 18 | 61.95225 | 0.002664253 | Credible Set | purple |
| chr18:61952524 | 18 | 61.95252 | 0.002634229 | Credible Set | purple |
| chr18:61952649 | 18 | 61.95265 | 0.004964955 | Credible Set | purple |
| chr18:61971767 | 18 | 61.97177 | 0.001571504 | Credible Set | purple |
| chr18:61973442 | 18 | 61.97344 | 0.001457635 | Credible Set | purple |
| chr18:61973478 | 18 | 61.97348 | 0.001453488 | Credible Set | purple |
| chr18:61991352 | 18 | 61.99135 | 0.002582605 | Credible Set | purple |
| chr18:61991384 | 18 | 61.99138 | 0.002590106 | Credible Set | purple |
| chr18:61992210 | 18 | 61.99221 | 0.002947759 | Credible Set | purple |
| chr18:61992519 | 18 | 61.99252 | 0.002740291 | Credible Set | purple |
| chr18:61999716 | 18 | 61.99972 | 0.002319199 | Credible Set | purple |
| chr18:62021643 | 18 | 62.02164 | 0.003763327 | Credible Set | purple |
| chr18:62021683 | 18 | 62.02168 | 0.003719783 | Credible Set | purple |
| chr18:62022099 | 18 | 62.02210 | 0.003465092 | Credible Set | purple |
| chr18:62051825 | 18 | 62.05183 | 0.002490355 | Credible Set | purple |
| chr18:62051845 | 18 | 62.05185 | 0.002321060 | Credible Set | purple |
| chr18:62055840 | 18 | 62.05584 | 0.003121373 | Credible Set | purple |
| chr18:62055876 | 18 | 62.05588 | 0.003121373 | Credible Set | purple |
| chr18:62055905 | 18 | 62.05591 | 0.003106380 | Credible Set | purple |
| chr18:62059126 | 18 | 62.05913 | 0.001941320 | Credible Set | purple |
| chr18:62059178 | 18 | 62.05918 | 0.001973137 | Credible Set | purple |
| chr18:62085794 | 18 | 62.08579 | 0.003906365 | Credible Set | purple |
| chr18:62085799 | 18 | 62.08580 | 0.003906365 | Credible Set | purple |
| chr18:62085840 | 18 | 62.08584 | 0.003906365 | Credible Set | purple |
| chr18:62092239 | 18 | 62.09224 | 0.002169119 | Credible Set | purple |

Fine−mapping Regions

| snp | chr | pos (Mb) | pp | group | color |
|---|---|---|---|---|---|
| chr18:62110682 | 18 | 62.11068 | 0.005086505 | Credible Set | purple |
| chr18:62137890 | 18 | 62.13789 | 0.002512666 | Credible Set | purple |
| chr18:62137914 | 18 | 62.13791 | 0.002512666 | Credible Set | purple |
| chr18:62147016 | 18 | 62.14702 | 0.002558800 | Credible Set | purple |
| chr18:62148467 | 18 | 62.14847 | 0.002515514 | Credible Set | purple |
| chr18:62151389 | 18 | 62.15139 | 0.002438614 | Credible Set | purple |
| chr18:62167241 | 18 | 62.16724 | 0.002489034 | Credible Set | purple |
| chr18:62167915 | 18 | 62.16792 | 0.002047695 | Credible Set | purple |
| chr18:62263390 | 18 | 62.26339 | 0.002328210 | Credible Set | purple |
| chr18:62400951 | 18 | 62.40095 | 0.002008048 | Credible Set | purple |
| chr18:62401009 | 18 | 62.40101 | 0.001472867 | Credible Set | purple |
| chr18:62425156 | 18 | 62.42516 | 0.001508041 | Credible Set | purple |
| chr18:62431611 | 18 | 62.43161 | 0.001468718 | Credible Set | purple |
| chr18:62431828 | 18 | 62.43183 | 0.001476917 | Credible Set | purple |

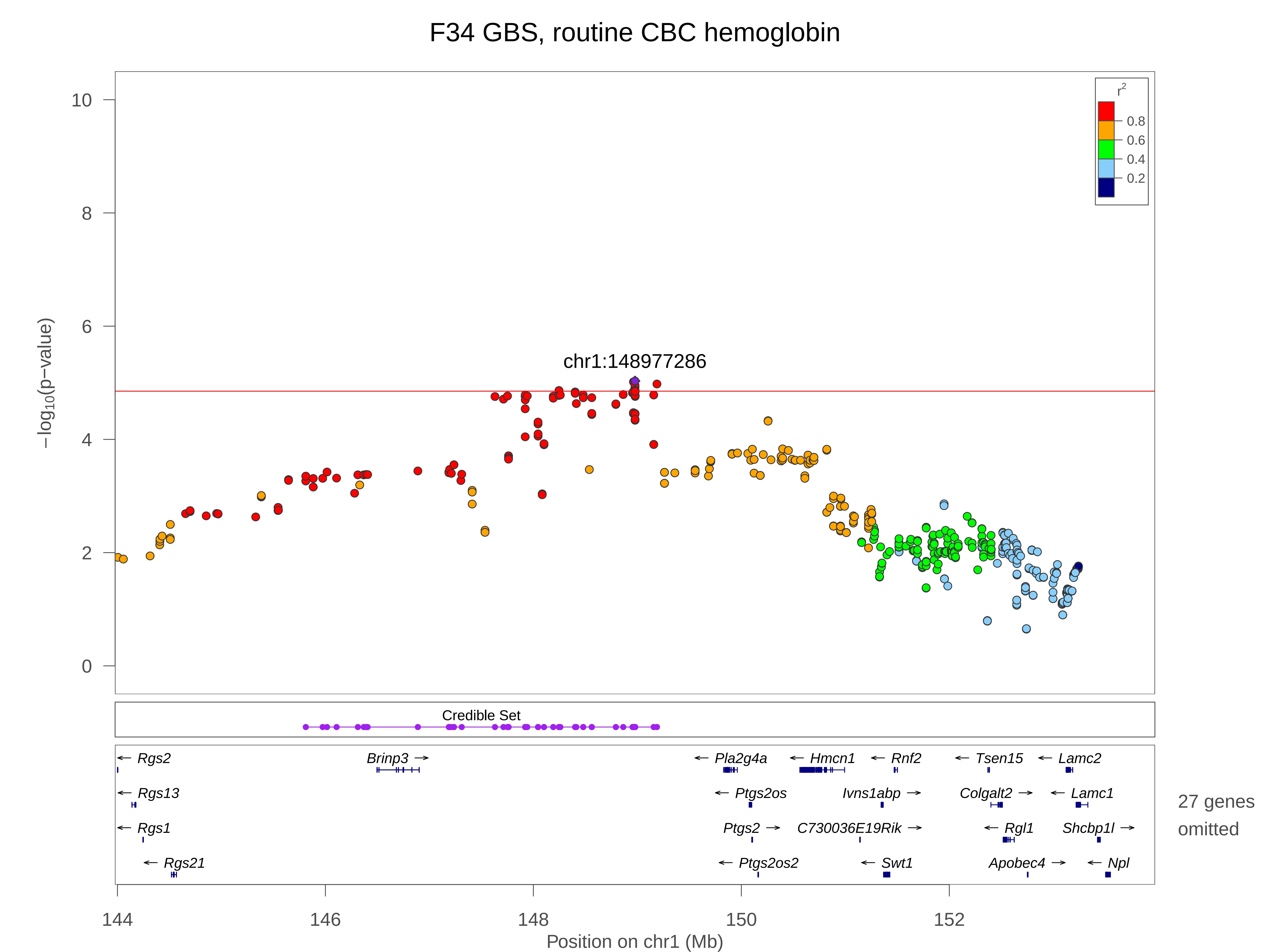

F34 GBS, routine CBC hemoglobin
chr1:148977286
−log10(p−value)
0
2
4
6
8
10
r2
0.8
0.6
0.4
0.2
Credible Set
← Rgs2
← Rgs13
← Rgs1
← Rgs21
Brinp3 →
← Pla2g4a
← Ptgs2os
Ptgs2 →
← Ptgs2os2
← Hmcn1
Ivns1abp →
C730036E19Rik →
← Swt1
← Rnf2
← Tsen15
Colgalt2 →
← Rgl1
Apobec4 →
← Lamc2
← Lamc1
Shcbp1l →
← Npl
27 genes omitted
144
146
148
150
152
Position on chr1 (Mb)

| | |
|---|---|
| date: | Thu Oct 25 17:05:22 2018 |
| build: | F34GBS |
| display range: | chr1:143977286−153977286 [143977286−153977286] |
| hilite range: | 0 − 0 [ 0 − 0 ] |
| reference SNP: | chr1:148977286 |
| number of SNPs plotted: | 507 |
| min P−value: | 9.3E−6 [chr1:148977286] |
| max P−value: | 2.24E−1 [chr1:152741578] |
| omitted Genes: | Rgs18, Pdc, 2310030A07Rik |
| omitted Genes: | BC003331, Tpr, Prg4 |
| omitted Genes: | Trmt1l, Mir7682, Fam129a |
| omitted Genes: | Edem3, 1700025G04Rik, Arpc5 |
| omitted Genes: | Ncf2, Smg7, Nmnat2 |
| omitted Genes: | E330020D12Rik, Dhx9, Rgs8 |
| omitted Genes: | C230024C17Rik, Rgs16, Rnasel |
| omitted Genes: | Rgsl1, Teddm2, Teddm1b |
| omitted Genes: | Teddm1a, Mir8114, Glul |

Make more plots at http://csg.sph.umich.edu/locuszoom/

Fine−mapping Regions

| snp | chr | pos (Mb) | pp | group | color |
|---|---|---|---|---|---|
| chr1:145811289 | 1 | 145.8113 | 0.0008577369 | Credible Set | purple |
| chr1:145973832 | 1 | 145.9738 | 0.0007888857 | Credible Set | purple |
| chr1:146015408 | 1 | 146.0154 | 0.0010040533 | Credible Set | purple |
| chr1:146107352 | 1 | 146.1074 | 0.0007967460 | Credible Set | purple |
| chr1:146312032 | 1 | 146.3120 | 0.0009021361 | Credible Set | purple |
| chr1:146368627 | 1 | 146.3686 | 0.0008993730 | Credible Set | purple |
| chr1:146391182 | 1 | 146.3912 | 0.0009119973 | Credible Set | purple |
| chr1:146391258 | 1 | 146.3913 | 0.0009119973 | Credible Set | purple |
| chr1:146405751 | 1 | 146.4058 | 0.0009091695 | Credible Set | purple |
| chr1:146889105 | 1 | 146.8891 | 0.0010454772 | Credible Set | purple |
| chr1:147187489 | 1 | 147.1875 | 0.0010033201 | Credible Set | purple |
| chr1:147187492 | 1 | 147.1875 | 0.0009753461 | Credible Set | purple |
| chr1:147194857 | 1 | 147.1949 | 0.0011050430 | Credible Set | purple |
| chr1:147211205 | 1 | 147.2112 | 0.0009529823 | Credible Set | purple |
| chr1:147236178 | 1 | 147.2362 | 0.0013229254 | Credible Set | purple |
| chr1:147236210 | 1 | 147.2362 | 0.0013215781 | Credible Set | purple |
| chr1:147310927 | 1 | 147.3109 | 0.0009236106 | Credible Set | purple |
| chr1:147630646 | 1 | 147.6306 | 0.0182459695 | Credible Set | purple |
| chr1:147712367 | 1 | 147.7124 | 0.0165009220 | Credible Set | purple |
| chr1:147751084 | 1 | 147.7511 | 0.0186180085 | Credible Set | purple |
| chr1:147760964 | 1 | 147.7610 | 0.0018628660 | Credible Set | purple |
| chr1:147760989 | 1 | 147.7610 | 0.0017353471 | Credible Set | purple |
| chr1:147761018 | 1 | 147.7610 | 0.0017136111 | Credible Set | purple |
| chr1:147761024 | 1 | 147.7610 | 0.0016379357 | Credible Set | purple |
| chr1:147921174 | 1 | 147.9212 | 0.0113437693 | Credible Set | purple |
| chr1:147921280 | 1 | 147.9213 | 0.0038550808 | Credible Set | purple |
| chr1:147921379 | 1 | 147.9214 | 0.0184018429 | Credible Set | purple |
| chr1:147921380 | 1 | 147.9214 | 0.0184305278 | Credible Set | purple |
| chr1:147921465 | 1 | 147.9215 | 0.0193593349 | Credible Set | purple |
| chr1:147921979 | 1 | 147.9220 | 0.0160262666 | Credible Set | purple |
| chr1:147940897 | 1 | 147.9409 | 0.0186561195 | Credible Set | purple |
| chr1:148044707 | 1 | 148.0447 | 0.0063120964 | Credible Set | purple |
| chr1:148044765 | 1 | 148.0448 | 0.0067958992 | Credible Set | purple |
| chr1:148045257 | 1 | 148.0453 | 0.0039575226 | Credible Set | purple |
| chr1:148045260 | 1 | 148.0453 | 0.0043140694 | Credible Set | purple |
| chr1:148102161 | 1 | 148.1022 | 0.0028430468 | Credible Set | purple |

Fine−mapping Regions

| snp | chr | pos (Mb) | pp | group | color |
|---|---|---|---|---|---|
| chr1:148102208 | 1 | 148.1022 | 0.002957991 | Credible Set | purple |
| chr1:148191398 | 1 | 148.1914 | 0.018498438 | Credible Set | purple |
| chr1:148191461 | 1 | 148.1915 | 0.017098490 | Credible Set | purple |
| chr1:148246268 | 1 | 148.2463 | 0.023058953 | Credible Set | purple |
| chr1:148246270 | 1 | 148.2463 | 0.022938019 | Credible Set | purple |
| chr1:148246308 | 1 | 148.2463 | 0.022938019 | Credible Set | purple |
| chr1:148246370 | 1 | 148.2464 | 0.019034858 | Credible Set | purple |
| chr1:148259002 | 1 | 148.2590 | 0.019304910 | Credible Set | purple |
| chr1:148401594 | 1 | 148.4016 | 0.021791085 | Credible Set | purple |
| chr1:148401658 | 1 | 148.4017 | 0.020635979 | Credible Set | purple |
| chr1:148412829 | 1 | 148.4128 | 0.013877302 | Credible Set | purple |
| chr1:148479166 | 1 | 148.4792 | 0.019399154 | Credible Set | purple |
| chr1:148479488 | 1 | 148.4795 | 0.017410552 | Credible Set | purple |
| chr1:148561626 | 1 | 148.5616 | 0.017452817 | Credible Set | purple |
| chr1:148561643 | 1 | 148.5616 | 0.009100144 | Credible Set | purple |
| chr1:148561679 | 1 | 148.5617 | 0.009496714 | Credible Set | purple |
| chr1:148793161 | 1 | 148.7932 | 0.013416499 | Credible Set | purple |
| chr1:148793216 | 1 | 148.7932 | 0.013720055 | Credible Set | purple |
| chr1:148864191 | 1 | 148.8642 | 0.019700105 | Credible Set | purple |
| chr1:148954162 | 1 | 148.9542 | 0.020982278 | Credible Set | purple |
| chr1:148954168 | 1 | 148.9542 | 0.020982278 | Credible Set | purple |
| chr1:148954207 | 1 | 148.9542 | 0.020982278 | Credible Set | purple |
| chr1:148961834 | 1 | 148.9618 | 0.009444397 | Credible Set | purple |
| chr1:148961852 | 1 | 148.9619 | 0.009767569 | Credible Set | purple |
| chr1:148962391 | 1 | 148.9624 | 0.022250022 | Credible Set | purple |
| chr1:148962448 | 1 | 148.9624 | 0.021115456 | Credible Set | purple |
| chr1:148962557 | 1 | 148.9626 | 0.032215213 | Credible Set | purple |
| chr1:148977255 | 1 | 148.9773 | 0.026336507 | Credible Set | purple |
| chr1:148977279 | 1 | 148.9773 | 0.028310124 | Credible Set | purple |
| chr1:148977286 | 1 | 148.9773 | 0.033309888 | Credible Set | purple |
| chr1:148977945 | 1 | 148.9779 | 0.009454303 | Credible Set | purple |
| chr1:148977976 | 1 | 148.9780 | 0.009250681 | Credible Set | purple |
| chr1:148977977 | 1 | 148.9780 | 0.007291518 | Credible Set | purple |
| chr1:148977984 | 1 | 148.9780 | 0.007549454 | Credible Set | purple |
| chr1:148978648 | 1 | 148.9786 | 0.023116484 | Credible Set | purple |
| chr1:148978689 | 1 | 148.9787 | 0.022578943 | Credible Set | purple |

Fine−mapping Regions

| snp | chr | pos (Mb) | pp | group | color |
|---|---|---|---|---|---|
| chr1:148978796 | 1 | 148.9788 | 0.022166155 | Credible Set | purple |
| chr1:148978842 | 1 | 148.9788 | 0.018304987 | Credible Set | purple |
| chr1:148978923 | 1 | 148.9789 | 0.019015093 | Credible Set | purple |
| chr1:148979220 | 1 | 148.9792 | 0.022250380 | Credible Set | purple |
| chr1:149157510 | 1 | 149.1575 | 0.019350960 | Credible Set | purple |
| chr1:149157713 | 1 | 149.1577 | 0.002885141 | Credible Set | purple |
| chr1:149157735 | 1 | 149.1577 | 0.002866823 | Credible Set | purple |
| chr1:149188240 | 1 | 149.1882 | 0.029664902 | Credible Set | purple |

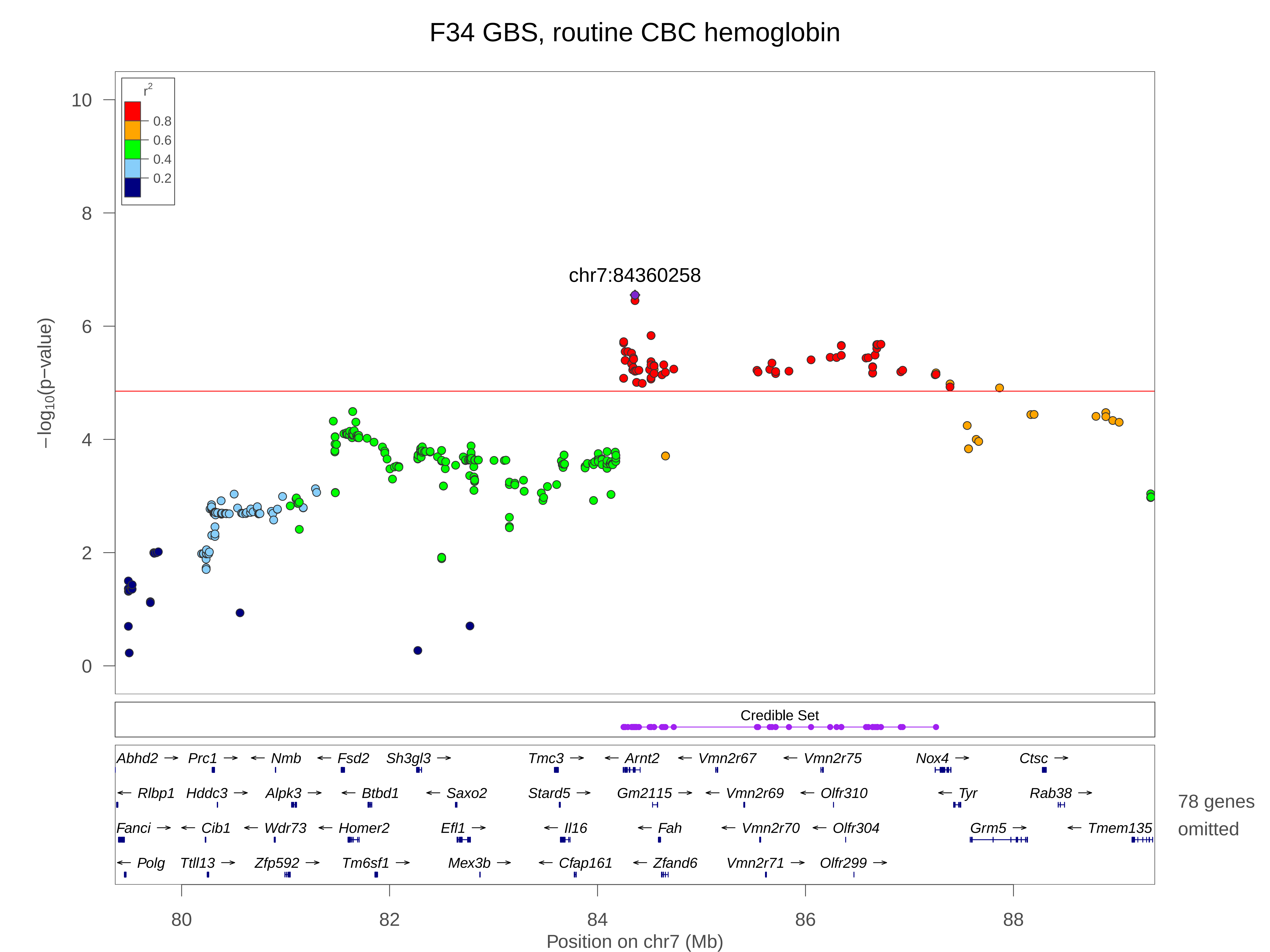
F34 GBS, routine CBC hemoglobin
r2
0.8
0.6
0.4
0.2
chr7:84360258
−log10(p−value)
10
8
6
4
2
0
Credible Set
Abhd2 → Prc1 → ← Nmb ← Fsd2 Sh3gl3 → Tmc3 → ← Arnt2 ← Vmn2r67 ← Vmn2r75 Nox4 → Ctsc →
← Rlbp1 Hddc3 → Alpk3 → ← Btbd1 ← Saxo2 Stard5 → Gm2115 → ← Vmn2r69 ← Olfr310 ← Tyr Rab38 →
Fanci → ← Cib1 ← Wdr73 ← Homer2 Efl1 → ← Il16 ← Fah ← Vmn2r70 ← Olfr304 Grm5 → ← Tmem135
← Polg Ttll13 → Zfp592 → Tm6sf1 → Mex3b → ← Cfap161 ← Zfand6 Vmn2r71 → Olfr299 →
78 genes omitted
80
82
84
86
88
Position on chr7 (Mb)

| | |
|---|---|
| date: | Thu Oct 25 17:05:33 2018 |
| build: | F34GBS |
| display range: | chr7:79360258−89360258 [79360258−89360258] |
| hilite range: | 0 − 0 [ 0 − 0 ] |
| reference SNP: | chr7:84360258 |
| number of SNPs plotted: | 391 |
| min P−value: | 2.81E−7 [chr7:84360258] |
| max P−value: | 5.91E−1 [chr7:79496275] |
| omitted Genes: | Mir9−3hg, Mir9−3, Rhcg |
| omitted Genes: | Ticrr, Kif7, Plin1 |
| omitted Genes: | Pex11a, Wdr93, Mesp1 |
| omitted Genes: | Mesp2, Anpep, Ap3s2 |
| omitted Genes: | Arpin, Zfp710, Gm21057 |
| omitted Genes: | Idh2, Sema4b, Gdpgp1 |
| omitted Genes: | Ngrn, Vps33b, Rccd1 |
| omitted Genes: | Unc45a, Man2a2, Fes |
| omitted Genes: | Furin, Blm, Crtc3 |
| omitted Genes: | Gm15880, Iqgap1, Zscan2 |
| omitted Genes: | Sec11a, Slc28a1, Pde8a |
| omitted Genes: | Rps17, Cpeb1, Ap3b2 |
| omitted Genes: | BC048679, 2900076A07Rik, Mir1839 |
| omitted Genes: | Whamm, Fam103a1, 3110040N11Rik |
| omitted Genes: | Hdgfrp3, Bnc1, Gm20744 |
| omitted Genes: | Adamtsl3, 4933406J10Rik, 4933430H16Rik |
| omitted Genes: | Mesdc1, Mesdc2, Cemip |
| omitted Genes: | Abhd17c, 2610206C17Rik, Olfr291 |
| omitted Genes: | Olfr290, Vmn2r65, Vmn2r66 |
| omitted Genes: | Vmn2r68, Vmn2r72, Vmn2r73 |
| omitted Genes: | Vmn2r74, Vmn2r76, Olfr309 |
| omitted Genes: | Olfr308, Olfr307, Olfr305 |
| omitted Genes: | Olfr303, Olfr301, Olfr298 |
| omitted Genes: | Olfr297, Olfr295, Olfr294 |
| omitted Genes: | Olfr293, Olfr292, Folh1 |
| omitted Genes: | Vmn2r77, Vmn2r78, Vmn2r79 |

Make more plots at http://csg.sph.umich.edu/locuszoom/

Fine−mapping Regions

| snp | chr | pos (Mb) | pp | group | color |
|---|---|---|---|---|---|
| chr7:84250985 | 7 | 84.25099 | 0.004369210 | Credible Set | purple |
| chr7:84251011 | 7 | 84.25101 | 0.004389647 | Credible Set | purple |
| chr7:84251159 | 7 | 84.25116 | 0.017322996 | Credible Set | purple |
| chr7:84251209 | 7 | 84.25121 | 0.018238578 | Credible Set | purple |
| chr7:84265410 | 7 | 84.26541 | 0.008791376 | Credible Set | purple |
| chr7:84265685 | 7 | 84.26569 | 0.012380497 | Credible Set | purple |
| chr7:84291540 | 7 | 84.29154 | 0.012397689 | Credible Set | purple |
| chr7:84327288 | 7 | 84.32729 | 0.011673098 | Credible Set | purple |
| chr7:84327808 | 7 | 84.32781 | 0.007835286 | Credible Set | purple |
| chr7:84327850 | 7 | 84.32785 | 0.007649291 | Credible Set | purple |
| chr7:84338595 | 7 | 84.33859 | 0.006078888 | Credible Set | purple |
| chr7:84343118 | 7 | 84.34312 | 0.006637969 | Credible Set | purple |
| chr7:84344805 | 7 | 84.34480 | 0.009692791 | Credible Set | purple |
| chr7:84347423 | 7 | 84.34742 | 0.009211168 | Credible Set | purple |
| chr7:84360038 | 7 | 84.36004 | 0.005835539 | Credible Set | purple |
| chr7:84360224 | 7 | 84.36022 | 0.111211961 | Credible Set | purple |
| chr7:84360247 | 7 | 84.36025 | 0.112493837 | Credible Set | purple |
| chr7:84360258 | 7 | 84.36026 | 0.113748596 | Credible Set | purple |
| chr7:84360279 | 7 | 84.36028 | 0.091143796 | Credible Set | purple |
| chr7:84360626 | 7 | 84.36063 | 0.005773288 | Credible Set | purple |
| chr7:84360642 | 7 | 84.36064 | 0.005773288 | Credible Set | purple |
| chr7:84360703 | 7 | 84.36070 | 0.005773288 | Credible Set | purple |
| chr7:84360714 | 7 | 84.36071 | 0.005773288 | Credible Set | purple |
| chr7:84360716 | 7 | 84.36072 | 0.005773288 | Credible Set | purple |
| chr7:84375526 | 7 | 84.37553 | 0.005862583 | Credible Set | purple |
| chr7:84375577 | 7 | 84.37558 | 0.003735568 | Credible Set | purple |
| chr7:84375642 | 7 | 84.37564 | 0.003735568 | Credible Set | purple |
| chr7:84397575 | 7 | 84.39758 | 0.005982792 | Credible Set | purple |
| chr7:84397597 | 7 | 84.39760 | 0.005982792 | Credible Set | purple |
| chr7:84501082 | 7 | 84.50108 | 0.006122093 | Credible Set | purple |
| chr7:84514082 | 7 | 84.51408 | 0.004245872 | Credible Set | purple |
| chr7:84514444 | 7 | 84.51444 | 0.008375030 | Credible Set | purple |
| chr7:84514505 | 7 | 84.51450 | 0.007391224 | Credible Set | purple |
| chr7:84514564 | 7 | 84.51456 | 0.023163768 | Credible Set | purple |
| chr7:84514575 | 7 | 84.51457 | 0.023206018 | Credible Set | purple |
| chr7:84514734 | 7 | 84.51473 | 0.004472572 | Credible Set | purple |

Fine−mapping Regions

| snp | chr | pos (Mb) | pp | group | color |
|---|---|---|---|---|---|
| chr7:84543346 | 7 | 84.54335 | 0.007165240 | Credible Set | purple |
| chr7:84543355 | 7 | 84.54336 | 0.007023433 | Credible Set | purple |
| chr7:84543782 | 7 | 84.54378 | 0.005281014 | Credible Set | purple |
| chr7:84619632 | 7 | 84.61963 | 0.005012820 | Credible Set | purple |
| chr7:84636759 | 7 | 84.63676 | 0.007415041 | Credible Set | purple |
| chr7:84653095 | 7 | 84.65309 | 0.005521391 | Credible Set | purple |
| chr7:84733186 | 7 | 84.73319 | 0.006266030 | Credible Set | purple |
| chr7:85533207 | 7 | 85.53321 | 0.005977768 | Credible Set | purple |
| chr7:85543843 | 7 | 85.54384 | 0.005573826 | Credible Set | purple |
| chr7:85656243 | 7 | 85.65624 | 0.006198745 | Credible Set | purple |
| chr7:85677494 | 7 | 85.67749 | 0.007926949 | Credible Set | purple |
| chr7:85712663 | 7 | 85.71266 | 0.005255469 | Credible Set | purple |
| chr7:85712975 | 7 | 85.71298 | 0.005700249 | Credible Set | purple |
| chr7:85840665 | 7 | 85.84067 | 0.005779641 | Credible Set | purple |
| chr7:86053386 | 7 | 86.05339 | 0.008986956 | Credible Set | purple |
| chr7:86237374 | 7 | 86.23737 | 0.009910253 | Credible Set | purple |
| chr7:86298979 | 7 | 86.29898 | 0.009829927 | Credible Set | purple |
| chr7:86344420 | 7 | 86.34442 | 0.015774234 | Credible Set | purple |
| chr7:86344437 | 7 | 86.34444 | 0.015571005 | Credible Set | purple |
| chr7:86344488 | 7 | 86.34449 | 0.010698252 | Credible Set | purple |
| chr7:86582139 | 7 | 86.58214 | 0.009631569 | Credible Set | purple |
| chr7:86603770 | 7 | 86.60377 | 0.009763695 | Credible Set | purple |
| chr7:86645719 | 7 | 86.64572 | 0.005342458 | Credible Set | purple |
| chr7:86645792 | 7 | 86.64579 | 0.005397533 | Credible Set | purple |
| chr7:86645947 | 7 | 86.64595 | 0.006879130 | Credible Set | purple |
| chr7:86645990 | 7 | 86.64599 | 0.006780100 | Credible Set | purple |
| chr7:86668998 | 7 | 86.66900 | 0.010806677 | Credible Set | purple |
| chr7:86685648 | 7 | 86.68565 | 0.013988330 | Credible Set | purple |
| chr7:86692214 | 7 | 86.69221 | 0.015974588 | Credible Set | purple |
| chr7:86692491 | 7 | 86.69249 | 0.016372079 | Credible Set | purple |
| chr7:86725518 | 7 | 86.72552 | 0.016501220 | Credible Set | purple |
| chr7:86916161 | 7 | 86.91616 | 0.005627650 | Credible Set | purple |
| chr7:86934710 | 7 | 86.93471 | 0.005992317 | Credible Set | purple |
| chr7:87255156 | 7 | 87.25516 | 0.005102177 | Credible Set | purple |

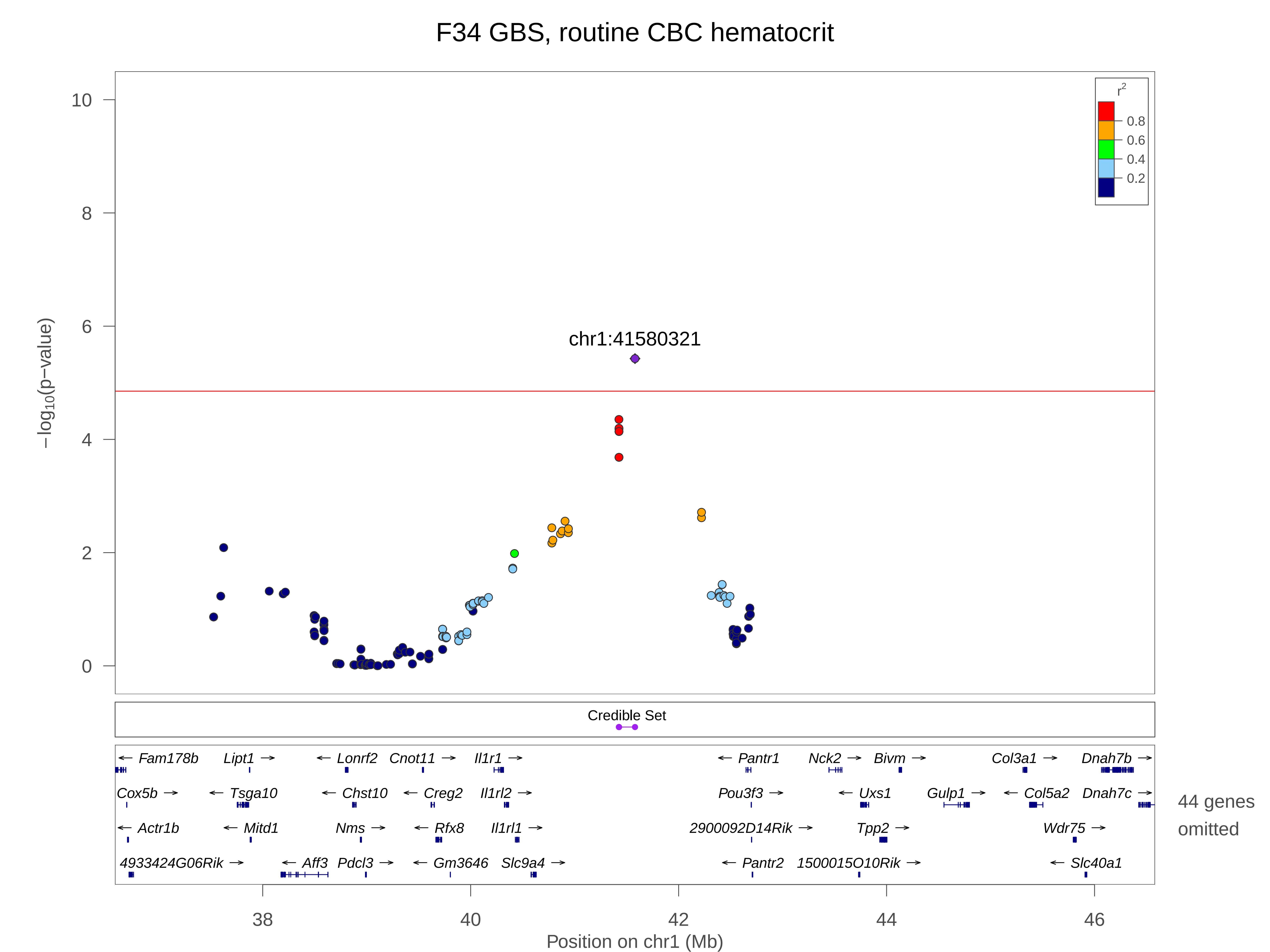
F34 GBS, routine CBC hematocrit
−log10(p−value)
10
8
6
4
2
0
chr1:41580321
r2
0.8
0.6
0.4
0.2
Credible Set
← Fam178b
Lipt1 →
← Lonrf2
Cnot11 →
Il1r1 →
← Pantr1
Nck2 →
Bivm →
Col3a1 →
Dnah7b →
Cox5b →
← Tsga10
← Chst10
← Creg2
Il1rl2 →
Pou3f3 →
← Uxs1
Gulp1 →
← Col5a2
Dnah7c →
← Actr1b
← Mitd1
Nms →
← Rfx8
Il1rl1 →
2900092D14Rik →
Tpp2 →
Wdr75 →
4933424G06Rik →
← Aff3
Pdcl3 →
← Gm3646
Slc9a4 →
← Pantr2
1500015O10Rik →
← Slc40a1
44 genes omitted
38
40
42
44
46
Position on chr1 (Mb)

| | |
|---|---|
| date: | Thu Oct 25 17:10:49 2018 |
| build: | F34GBS |
| display range: | chr1:36580321−46580321 [36580321−46580321] |
| hilite range: | 0 − 0 [ 0 − 0 ] |
| reference SNP: | chr1:41580321 |
| number of SNPs plotted: | 187 |
| min P−value: | 3.74E−6 [chr1:41580321] |
| max P−value: | 9.96E−1 [chr1:39101479] |
| omitted Genes: | Zap70, Tmem131, Cnga3 |
| omitted Genes: | Inpp4a, Coa5, Unc50 |
| omitted Genes: | Mgat4a, 4930594C11Rik, 2010300C02Rik |
| omitted Genes: | Mrpl30, Lyg2, Lyg1 |
| omitted Genes: | Txndc9, Eif5b, Rev1 |
| omitted Genes: | Mir6349, Npas2, Rpl31 |
| omitted Genes: | Tbc1d8, Snord89, Rnf149 |
| omitted Genes: | 1700066B17Rik, Map4k4, Gm16894 |
| omitted Genes: | Il1r2, Il18r1, Il18rap |
| omitted Genes: | Slc9a2, Mfsd9, Tmem182 |
| omitted Genes: | 4930448I06Rik, Dalir, Mrps9 |
| omitted Genes: | Gpr45, Tgfbrap1, AI597479 |
| omitted Genes: | Fhl2, Mettl21c, Tex30 |
| omitted Genes: | Kdelc1, Ercc5, Mettl21e |
| omitted Genes: | 4930521E06Rik, Stk−ps2 |

Make more plots at http://csg.sph.umich.edu/locuszoom/

Fine−mapping Regions

| snp | chr | pos (Mb) | pp | group | color |
|---|---|---|---|---|---|
| chr1:41425834 | 1 | 41.42583 | 0.06690913 | Credible Set | purple |
| chr1:41426242 | 1 | 41.42624 | 0.04624021 | Credible Set | purple |
| chr1:41426343 | 1 | 41.42634 | 0.01563230 | Credible Set | purple |
| chr1:41426398 | 1 | 41.42640 | 0.01563230 | Credible Set | purple |
| chr1:41426422 | 1 | 41.42642 | 0.01563230 | Credible Set | purple |
| chr1:41426449 | 1 | 41.42645 | 0.04767386 | Credible Set | purple |
| chr1:41426503 | 1 | 41.42650 | 0.04195553 | Credible Set | purple |
| chr1:41426526 | 1 | 41.42653 | 0.04211278 | Credible Set | purple |
| chr1:41580321 | 1 | 41.58032 | 0.70821160 | Credible Set | purple |

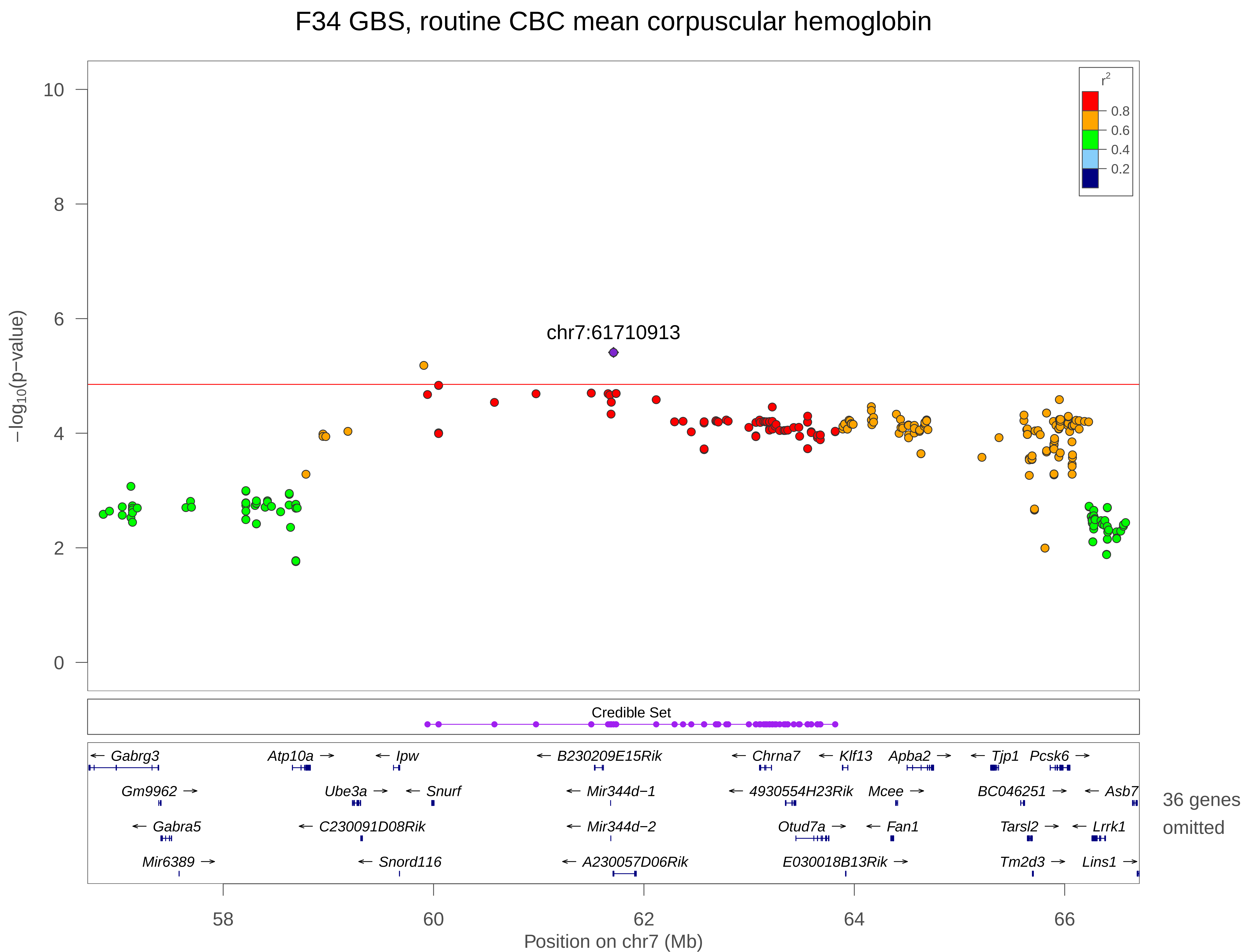

F34 GBS, routine CBC mean corpuscular hemoglobin
−log10(p−value)
chr7:61710913
r2
0.8
0.6
0.4
0.2
Credible Set
← Gabrg3
Atp10a →
← Ipw
← B230209E15Rik
← Chrna7
← Klf13
Apba2 →
← Tjp1
Pcsk6 →
Gm9962 →
Ube3a →
← Snurf
← Mir344d−1
← 4930554H23Rik
Mcee →
BC046251 →
← Asb7
← Gabra5
← C230091D08Rik
← Mir344d−2
Otud7a →
← Fan1
Tarsl2 →
← Lrrk1
Mir6389 →
← Snord116
← A230057D06Rik
E030018B13Rik →
Tm2d3 →
Lins1 →
36 genes omitted
Position on chr7 (Mb)

| | |
|---|---|
| date: | Thu Oct 25 17:11:36 2018 |
| build: | F34GBS |
| display range: | chr7:56710913−66710913 [56710913−66710913] |
| hilite range: | 0 − 0 [ 0 − 0 ] |
| reference SNP: | chr7:61710913 |
| number of SNPs plotted: | 336 |
| min P−value: | 3.89E−6 [chr7:61710913] |
| max P−value: | 1.74E−2 [chr7:58690611] |
| omitted Genes: | Gabrb3, Snord116l2, Snord116l1 |
| omitted Genes: | Snhg14, Snord64, Snrpn |
| omitted Genes: | Gm7367, Mir344d−3, Mir344e |
| omitted Genes: | Mir344h−1, Mir344h−2, Mir344b |
| omitted Genes: | Mir344c, Mir344, Mir344−2 |
| omitted Genes: | A330076H08Rik, Mir344g, Mir344f |
| omitted Genes: | Mir344i, Ndn, Magel2 |
| omitted Genes: | Mkrn3, Peg12, Trpm1 |
| omitted Genes: | Mir211, Mir6416, Mtmr10 |
| omitted Genes: | Mphosph10, Fam189a1, Ndnl2 |
| omitted Genes: | 1810008I18Rik, Snrpa1, Vimp |
| omitted Genes: | Chsy1, Mir7057, Aldh1a3 |

Make more plots at http://csg.sph.umich.edu/locuszoom/

Fine−mapping Regions

| snp | chr | pos (Mb) | pp | group | color |
|---|---|---|---|---|---|
| chr7:59942572 | 7 | 59.94257 | 0.022251592 | Credible Set | purple |
| chr7:60047736 | 7 | 60.04774 | 0.005072398 | Credible Set | purple |
| chr7:60047743 | 7 | 60.04774 | 0.005167092 | Credible Set | purple |
| chr7:60047769 | 7 | 60.04777 | 0.005094818 | Credible Set | purple |
| chr7:60047773 | 7 | 60.04777 | 0.005027397 | Credible Set | purple |
| chr7:60047999 | 7 | 60.04800 | 0.031474306 | Credible Set | purple |
| chr7:60048040 | 7 | 60.04804 | 0.031730538 | Credible Set | purple |
| chr7:60578939 | 7 | 60.57894 | 0.016483308 | Credible Set | purple |
| chr7:60974140 | 7 | 60.97414 | 0.022912960 | Credible Set | purple |
| chr7:61499181 | 7 | 61.49918 | 0.023608267 | Credible Set | purple |
| chr7:61499595 | 7 | 61.49959 | 0.023454797 | Credible Set | purple |
| chr7:61659201 | 7 | 61.65920 | 0.023000487 | Credible Set | purple |
| chr7:61675494 | 7 | 61.67549 | 0.021831864 | Credible Set | purple |
| chr7:61687345 | 7 | 61.68735 | 0.010537402 | Credible Set | purple |
| chr7:61689644 | 7 | 61.68964 | 0.016600154 | Credible Set | purple |
| chr7:61689650 | 7 | 61.68965 | 0.016600154 | Credible Set | purple |
| chr7:61710907 | 7 | 61.71091 | 0.111123981 | Credible Set | purple |
| chr7:61710913 | 7 | 61.71091 | 0.112099696 | Credible Set | purple |
| chr7:61733905 | 7 | 61.73391 | 0.023124976 | Credible Set | purple |
| chr7:61736166 | 7 | 61.73617 | 0.023167659 | Credible Set | purple |
| chr7:62116766 | 7 | 62.11677 | 0.018268790 | Credible Set | purple |
| chr7:62291228 | 7 | 62.29123 | 0.007861675 | Credible Set | purple |
| chr7:62291477 | 7 | 62.29148 | 0.007861675 | Credible Set | purple |
| chr7:62371914 | 7 | 62.37191 | 0.008020960 | Credible Set | purple |
| chr7:62450245 | 7 | 62.45025 | 0.005363873 | Credible Set | purple |
| chr7:62571648 | 7 | 62.57165 | 0.007506324 | Credible Set | purple |
| chr7:62572320 | 7 | 62.57232 | 0.007872096 | Credible Set | purple |
| chr7:62572352 | 7 | 62.57235 | 0.007872096 | Credible Set | purple |
| chr7:62572353 | 7 | 62.57235 | 0.007872096 | Credible Set | purple |
| chr7:62684003 | 7 | 62.68400 | 0.008042015 | Credible Set | purple |
| chr7:62684079 | 7 | 62.68408 | 0.008154924 | Credible Set | purple |
| chr7:62684449 | 7 | 62.68445 | 0.008203114 | Credible Set | purple |
| chr7:62696801 | 7 | 62.69680 | 0.007999415 | Credible Set | purple |
| chr7:62706977 | 7 | 62.70698 | 0.007787941 | Credible Set | purple |
| chr7:62782840 | 7 | 62.78284 | 0.008419134 | Credible Set | purple |
| chr7:62800795 | 7 | 62.80080 | 0.008053060 | Credible Set | purple |

Fine−mapping Regions

| snp | chr | pos (Mb) | pp | group | color |
|---|---|---|---|---|---|
| chr7:62997563 | 7 | 62.99756 | 0.006358640 | Credible Set | purple |
| chr7:63064073 | 7 | 63.06407 | 0.004623203 | Credible Set | purple |
| chr7:63064081 | 7 | 63.06408 | 0.004478329 | Credible Set | purple |
| chr7:63064100 | 7 | 63.06410 | 0.004497934 | Credible Set | purple |
| chr7:63064751 | 7 | 63.06475 | 0.007679605 | Credible Set | purple |
| chr7:63099443 | 7 | 63.09944 | 0.008398832 | Credible Set | purple |
| chr7:63105975 | 7 | 63.10598 | 0.007676573 | Credible Set | purple |
| chr7:63139467 | 7 | 63.13947 | 0.007945851 | Credible Set | purple |
| chr7:63153842 | 7 | 63.15384 | 0.007872096 | Credible Set | purple |
| chr7:63167968 | 7 | 63.16797 | 0.007872096 | Credible Set | purple |
| chr7:63193963 | 7 | 63.19396 | 0.007759763 | Credible Set | purple |
| chr7:63194039 | 7 | 63.19404 | 0.006037196 | Credible Set | purple |
| chr7:63194056 | 7 | 63.19406 | 0.005732830 | Credible Set | purple |
| chr7:63194259 | 7 | 63.19426 | 0.007935149 | Credible Set | purple |
| chr7:63194342 | 7 | 63.19434 | 0.007872096 | Credible Set | purple |
| chr7:63194487 | 7 | 63.19449 | 0.007886203 | Credible Set | purple |
| chr7:63194494 | 7 | 63.19449 | 0.007886203 | Credible Set | purple |
| chr7:63220109 | 7 | 63.22011 | 0.013813942 | Credible Set | purple |
| chr7:63221022 | 7 | 63.22102 | 0.008002939 | Credible Set | purple |
| chr7:63223436 | 7 | 63.22344 | 0.005961252 | Credible Set | purple |
| chr7:63246570 | 7 | 63.24657 | 0.006683077 | Credible Set | purple |
| chr7:63256470 | 7 | 63.25647 | 0.007116144 | Credible Set | purple |
| chr7:63291026 | 7 | 63.29103 | 0.005650557 | Credible Set | purple |
| chr7:63332548 | 7 | 63.33255 | 0.005642547 | Credible Set | purple |
| chr7:63345554 | 7 | 63.34555 | 0.005661611 | Credible Set | purple |
| chr7:63366897 | 7 | 63.36690 | 0.005729901 | Credible Set | purple |
| chr7:63367321 | 7 | 63.36732 | 0.005729901 | Credible Set | purple |
| chr7:63424326 | 7 | 63.42433 | 0.006328016 | Credible Set | purple |
| chr7:63472965 | 7 | 63.47297 | 0.006374090 | Credible Set | purple |
| chr7:63480226 | 7 | 63.48023 | 0.004537845 | Credible Set | purple |
| chr7:63555396 | 7 | 63.55540 | 0.007725729 | Credible Set | purple |
| chr7:63555456 | 7 | 63.55546 | 0.007799817 | Credible Set | purple |
| chr7:63556071 | 7 | 63.55607 | 0.007792293 | Credible Set | purple |
| chr7:63556106 | 7 | 63.55611 | 0.007792293 | Credible Set | purple |
| chr7:63556195 | 7 | 63.55620 | 0.002840500 | Credible Set | purple |
| chr7:63556256 | 7 | 63.55626 | 0.002843255 | Credible Set | purple |

Fine−mapping Regions

| snp | chr | pos (Mb) | pp | group | color |
|---|---|---|---|---|---|
| chr7:63556415 | 7 | 63.55642 | 0.009769688 | Credible Set | purple |
| chr7:63590413 | 7 | 63.59041 | 0.005388610 | Credible Set | purple |
| chr7:63590941 | 7 | 63.59094 | 0.005365126 | Credible Set | purple |
| chr7:63591000 | 7 | 63.59100 | 0.005214284 | Credible Set | purple |
| chr7:63649222 | 7 | 63.64922 | 0.004414043 | Credible Set | purple |
| chr7:63649227 | 7 | 63.64923 | 0.004306175 | Credible Set | purple |
| chr7:63650750 | 7 | 63.65075 | 0.004683786 | Credible Set | purple |
| chr7:63676214 | 7 | 63.67621 | 0.004782311 | Credible Set | purple |
| chr7:63676590 | 7 | 63.67659 | 0.003980815 | Credible Set | purple |
| chr7:63677624 | 7 | 63.67762 | 0.004767654 | Credible Set | purple |
| chr7:63818257 | 7 | 63.81826 | 0.005387385 | Credible Set | purple |
| chr7:63818340 | 7 | 63.81834 | 0.005490960 | Credible Set | purple |

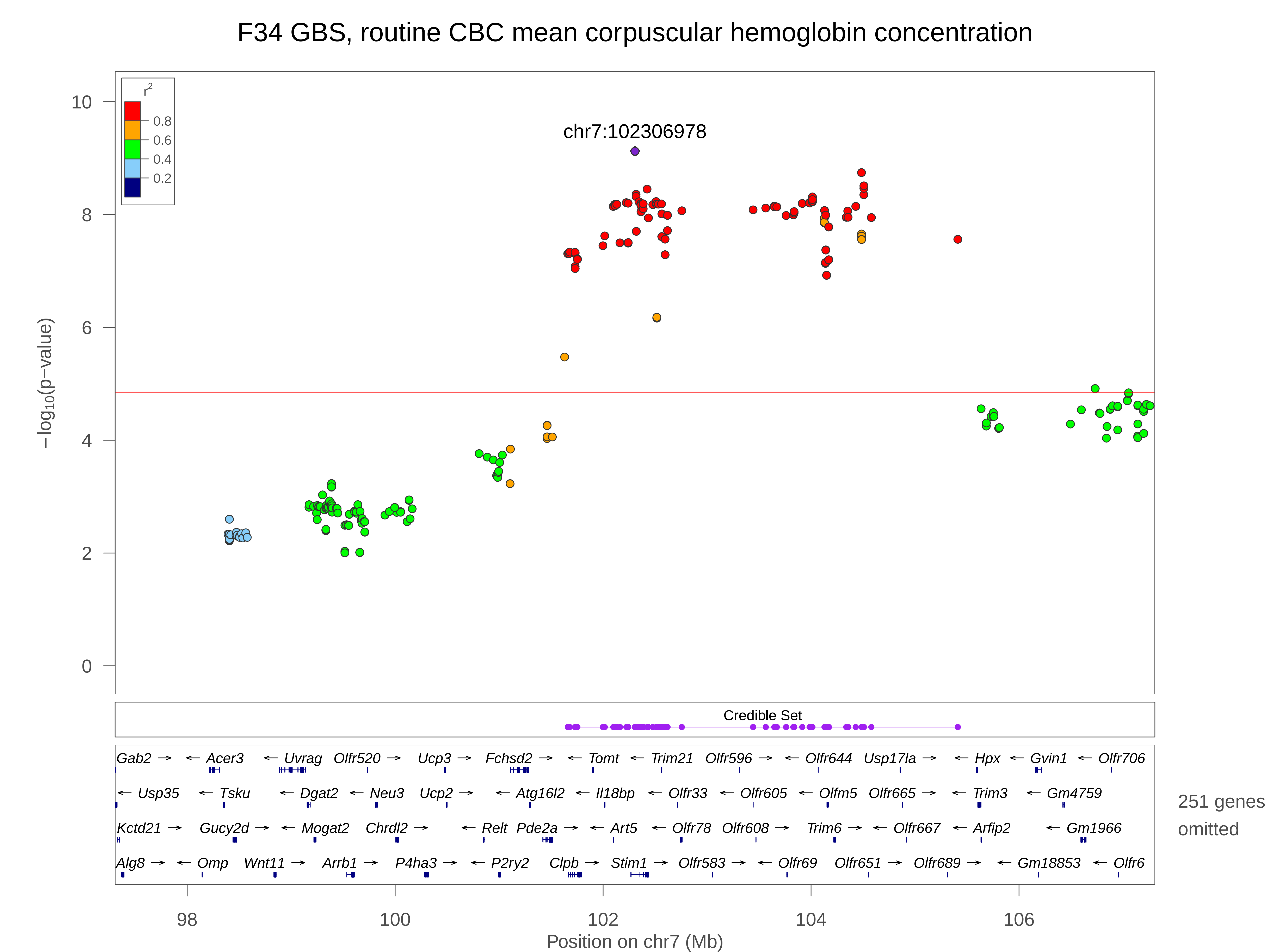
F34 GBS, routine CBC mean corpuscular hemoglobin concentration
chr7:102306978
r2
0.8
0.6
0.4
0.2
−log10(p−value)
Credible Set
Gab2 → ← Acer3 ← Uvrag Olfr520 → Ucp3 → Fchsd2 → ← Tomt ← Trim21 Olfr596 → ← Olfr644 Usp17la → ← Hpx ← Gvin1 ← Olfr706
← Usp35 ← Tsku ← Dgat2 ← Neu3 Ucp2 → ← Atg16l2 ← Il18bp ← Olfr33 ← Olfr605 ← Olfm5 Olfr665 → ← Trim3 ← Gm4759
Kctd21 → Gucy2d → ← Mogat2 Chrdl2 → ← Relt Pde2a → ← Art5 ← Olfr78 Olfr608 → Trim6 → ← Olfr667 ← Arfip2 ← Gm1966
Alg8 → ← Omp Wnt11 → Arrb1 → P4ha3 → ← P2ry2 Clpb → Stim1 → Olfr583 → ← Olfr69 Olfr651 → Olfr689 → ← Gm18853 ← Olfr6
251 genes omitted
Position on chr7 (Mb)

date: Thu Oct 25 17:11:51 2018
build: F34GBS
display range: chr7:97306978−107306978 [97306978−107306978]
hilite range: 0 − 0 [ 0 − 0 ]
reference SNP: chr7:102306978
number of SNPs plotted: 301
min P−value: 7.53E−10 [chr7:102306978]
max P−value: 9.94E−3 [chr7:99516884]
omitted Genes: Ndufc2, Thrsp, Kctd14
omitted Genes: Ints4, Aamdc, Rsf1
omitted Genes: Rsf1os1, Clns1a, Aqp11
omitted Genes: Pak1, Gdpd4, Myo7a
omitted Genes: Capn5, Gm16938, B3gnt6
omitted Genes: Lrrc32, Emsy, Prkrir
omitted Genes: Gm26705, Map6, Serpinh1
omitted Genes: Gdpd5, Klhl35, Rps3
omitted Genes: Snord15b, Snord15a, Mir326
omitted Genes: Tpbgl, Slco2b1, Olfr521
omitted Genes: Spcs2, Xrra1, Rnf169
omitted Genes: Pold3, Lipt2, Kcne3
omitted Genes: Pgm2l1, Gpx2−ps1, Ppme1
omitted Genes: C2cd3, Dnajb13, Coa4
omitted Genes: Mrpl48, Rab6a, Plekhb1
omitted Genes: Fam168a, Arhgef17, Mir3102
omitted Genes: P2ry6, Stard10, Arap1
omitted Genes: Mir139, Art2a−ps, Art2b
omitted Genes: Phox2a, Inppl1, Folr2
omitted Genes: Folr1, Anapc15, Lamtor1
omitted Genes: Lrrc51, Numa1, Rnf121
omitted Genes: Xndc1, Xntrpc, Trpc2
omitted Genes: Art1, Chrna10, Nup98
omitted Genes: Pgap2, Rhog, Rrm1
omitted Genes: Olfr543, Olfr544, Olfr545
omitted Genes: Olfr547, Olfr549, Olfr550
omitted Genes: Olfr551, Olfr552, Olfr553
omitted Genes: Olfr554, Olfr555, Olfr556
omitted Genes: Trim68, Olfr557, Olfr558
omitted Genes: Olfr559, Olfr560, Olfr561
omitted Genes: Olfr564, Olfr566, Olfr568
Make more plots at http://csg.sph.umich.edu/locuszoom/
omitted Genes: Olfr569, Olfr570, Olfr571

Fine−mapping Regions

| snp | chr | pos (Mb) | pp | group | color |
|---|---|---|---|---|---|
| chr7:101660383 | 7 | 101.6604 | 0.001576280 | Credible Set | purple |
| chr7:101669186 | 7 | 101.6692 | 0.001627721 | Credible Set | purple |
| chr7:101673483 | 7 | 101.6735 | 0.001581352 | Credible Set | purple |
| chr7:101680140 | 7 | 101.6801 | 0.001682982 | Credible Set | purple |
| chr7:101731061 | 7 | 101.7311 | 0.001596039 | Credible Set | purple |
| chr7:101731110 | 7 | 101.7311 | 0.001660867 | Credible Set | purple |
| chr7:101751790 | 7 | 101.7518 | 0.001282302 | Credible Set | purple |
| chr7:101752580 | 7 | 101.7526 | 0.001291531 | Credible Set | purple |
| chr7:101752971 | 7 | 101.7530 | 0.001254355 | Credible Set | purple |
| chr7:101998679 | 7 | 101.9987 | 0.002156079 | Credible Set | purple |
| chr7:102016307 | 7 | 102.0163 | 0.003192659 | Credible Set | purple |
| chr7:102098054 | 7 | 102.0981 | 0.010253204 | Credible Set | purple |
| chr7:102110109 | 7 | 102.1101 | 0.011040329 | Credible Set | purple |
| chr7:102110501 | 7 | 102.1105 | 0.011050674 | Credible Set | purple |
| chr7:102116318 | 7 | 102.1163 | 0.010660109 | Credible Set | purple |
| chr7:102116430 | 7 | 102.1164 | 0.010660109 | Credible Set | purple |
| chr7:102116478 | 7 | 102.1165 | 0.010555237 | Credible Set | purple |
| chr7:102132562 | 7 | 102.1326 | 0.011196897 | Credible Set | purple |
| chr7:102162058 | 7 | 102.1621 | 0.002421824 | Credible Set | purple |
| chr7:102223396 | 7 | 102.2234 | 0.011906611 | Credible Set | purple |
| chr7:102240695 | 7 | 102.2407 | 0.011670182 | Credible Set | purple |
| chr7:102241191 | 7 | 102.2412 | 0.002444228 | Credible Set | purple |
| chr7:102241211 | 7 | 102.2412 | 0.002396435 | Credible Set | purple |
| chr7:102241212 | 7 | 102.2412 | 0.002438744 | Credible Set | purple |
| chr7:102306968 | 7 | 102.3070 | 0.090949863 | Credible Set | purple |
| chr7:102306978 | 7 | 102.3070 | 0.092233491 | Credible Set | purple |
| chr7:102317213 | 7 | 102.3172 | 0.016661476 | Credible Set | purple |
| chr7:102317246 | 7 | 102.3172 | 0.015248052 | Credible Set | purple |
| chr7:102319586 | 7 | 102.3196 | 0.003814712 | Credible Set | purple |
| chr7:102344970 | 7 | 102.3450 | 0.012334078 | Credible Set | purple |
| chr7:102363897 | 7 | 102.3639 | 0.011227081 | Credible Set | purple |
| chr7:102364204 | 7 | 102.3642 | 0.010507328 | Credible Set | purple |
| chr7:102364223 | 7 | 102.3642 | 0.011395775 | Credible Set | purple |
| chr7:102364315 | 7 | 102.3643 | 0.010787722 | Credible Set | purple |
| chr7:102364326 | 7 | 102.3643 | 0.010787722 | Credible Set | purple |
| chr7:102364411 | 7 | 102.3644 | 0.008270994 | Credible Set | purple |

Fine−mapping Regions

| snp | chr | pos (Mb) | pp | group | color |
|---|---|---|---|---|---|
| chr7:102385520 | 7 | 102.3855 | 0.009325765 | Credible Set | purple |
| chr7:102385536 | 7 | 102.3855 | 0.011386844 | Credible Set | purple |
| chr7:102423165 | 7 | 102.4232 | 0.020416283 | Credible Set | purple |
| chr7:102435692 | 7 | 102.4357 | 0.006496564 | Credible Set | purple |
| chr7:102435720 | 7 | 102.4357 | 0.006496564 | Credible Set | purple |
| chr7:102478469 | 7 | 102.4785 | 0.011012408 | Credible Set | purple |
| chr7:102510475 | 7 | 102.5105 | 0.011340650 | Credible Set | purple |
| chr7:102510607 | 7 | 102.5106 | 0.012432072 | Credible Set | purple |
| chr7:102516878 | 7 | 102.5169 | 0.011305657 | Credible Set | purple |
| chr7:102531652 | 7 | 102.5317 | 0.011241254 | Credible Set | purple |
| chr7:102561708 | 7 | 102.5617 | 0.011341837 | Credible Set | purple |
| chr7:102565262 | 7 | 102.5653 | 0.003088087 | Credible Set | purple |
| chr7:102565292 | 7 | 102.5653 | 0.003093001 | Credible Set | purple |
| chr7:102565307 | 7 | 102.5653 | 0.003081350 | Credible Set | purple |
| chr7:102565323 | 7 | 102.5653 | 0.003086252 | Credible Set | purple |
| chr7:102565329 | 7 | 102.5653 | 0.003086252 | Credible Set | purple |
| chr7:102565410 | 7 | 102.5654 | 0.007625065 | Credible Set | purple |
| chr7:102596442 | 7 | 102.5964 | 0.002803786 | Credible Set | purple |
| chr7:102596578 | 7 | 102.5966 | 0.001512829 | Credible Set | purple |
| chr7:102596636 | 7 | 102.5966 | 0.001510423 | Credible Set | purple |
| chr7:102619418 | 7 | 102.6194 | 0.007215964 | Credible Set | purple |
| chr7:102619431 | 7 | 102.6194 | 0.007203272 | Credible Set | purple |
| chr7:102619466 | 7 | 102.6195 | 0.003940197 | Credible Set | purple |
| chr7:102757499 | 7 | 102.7575 | 0.008632259 | Credible Set | purple |
| chr7:103442570 | 7 | 103.4426 | 0.008952930 | Credible Set | purple |
| chr7:103564836 | 7 | 103.5648 | 0.009614334 | Credible Set | purple |
| chr7:103646439 | 7 | 103.6464 | 0.010419108 | Credible Set | purple |
| chr7:103646709 | 7 | 103.6467 | 0.010119398 | Credible Set | purple |
| chr7:103671033 | 7 | 103.6710 | 0.010059662 | Credible Set | purple |
| chr7:103760301 | 7 | 103.7603 | 0.007158021 | Credible Set | purple |
| chr7:103829200 | 7 | 103.8292 | 0.007327551 | Credible Set | purple |
| chr7:103837099 | 7 | 103.8371 | 0.007894645 | Credible Set | purple |
| chr7:103837213 | 7 | 103.8372 | 0.008350379 | Credible Set | purple |
| chr7:103915503 | 7 | 103.9155 | 0.011543428 | Credible Set | purple |
| chr7:103986792 | 7 | 103.9868 | 0.011984438 | Credible Set | purple |
| chr7:103986795 | 7 | 103.9868 | 0.011984438 | Credible Set | purple |

Fine−mapping Regions

| snp | chr | pos (Mb) | pp | group | color |
|---|---|---|---|---|---|
| chr7:103986857 | 7 | 103.9869 | 0.011749707 | Credible Set | purple |
| chr7:103986860 | 7 | 103.9869 | 0.011749707 | Credible Set | purple |
| chr7:104012808 | 7 | 104.0128 | 0.012334102 | Credible Set | purple |
| chr7:104013370 | 7 | 104.0134 | 0.012542197 | Credible Set | purple |
| chr7:104013395 | 7 | 104.0134 | 0.014993762 | Credible Set | purple |
| chr7:104013467 | 7 | 104.0135 | 0.013443169 | Credible Set | purple |
| chr7:104129734 | 7 | 104.1297 | 0.008725313 | Credible Set | purple |
| chr7:104129736 | 7 | 104.1297 | 0.008725313 | Credible Set | purple |
| chr7:104141343 | 7 | 104.1413 | 0.007256169 | Credible Set | purple |
| chr7:104141705 | 7 | 104.1417 | 0.001834794 | Credible Set | purple |
| chr7:104141768 | 7 | 104.1418 | 0.001817733 | Credible Set | purple |
| chr7:104170479 | 7 | 104.1705 | 0.001239540 | Credible Set | purple |
| chr7:104170518 | 7 | 104.1705 | 0.001234318 | Credible Set | purple |
| chr7:104170543 | 7 | 104.1705 | 0.001236844 | Credible Set | purple |
| chr7:104170588 | 7 | 104.1706 | 0.004537395 | Credible Set | purple |
| chr7:104170594 | 7 | 104.1706 | 0.004523155 | Credible Set | purple |
| chr7:104170618 | 7 | 104.1706 | 0.004549326 | Credible Set | purple |
| chr7:104170662 | 7 | 104.1707 | 0.004549326 | Credible Set | purple |
| chr7:104338784 | 7 | 104.3388 | 0.006676217 | Credible Set | purple |
| chr7:104338810 | 7 | 104.3388 | 0.006689773 | Credible Set | purple |
| chr7:104353491 | 7 | 104.3535 | 0.008546463 | Credible Set | purple |
| chr7:104356230 | 7 | 104.3562 | 0.006682156 | Credible Set | purple |
| chr7:104430205 | 7 | 104.4302 | 0.010279656 | Credible Set | purple |
| chr7:104485236 | 7 | 104.4852 | 0.039314262 | Credible Set | purple |
| chr7:104507724 | 7 | 104.5077 | 0.016280612 | Credible Set | purple |
| chr7:104508127 | 7 | 104.5081 | 0.021038879 | Credible Set | purple |
| chr7:104508173 | 7 | 104.5082 | 0.023240353 | Credible Set | purple |
| chr7:104580091 | 7 | 104.5801 | 0.006597281 | Credible Set | purple |
| chr7:105411752 | 7 | 105.4118 | 0.002787939 | Credible Set | purple |

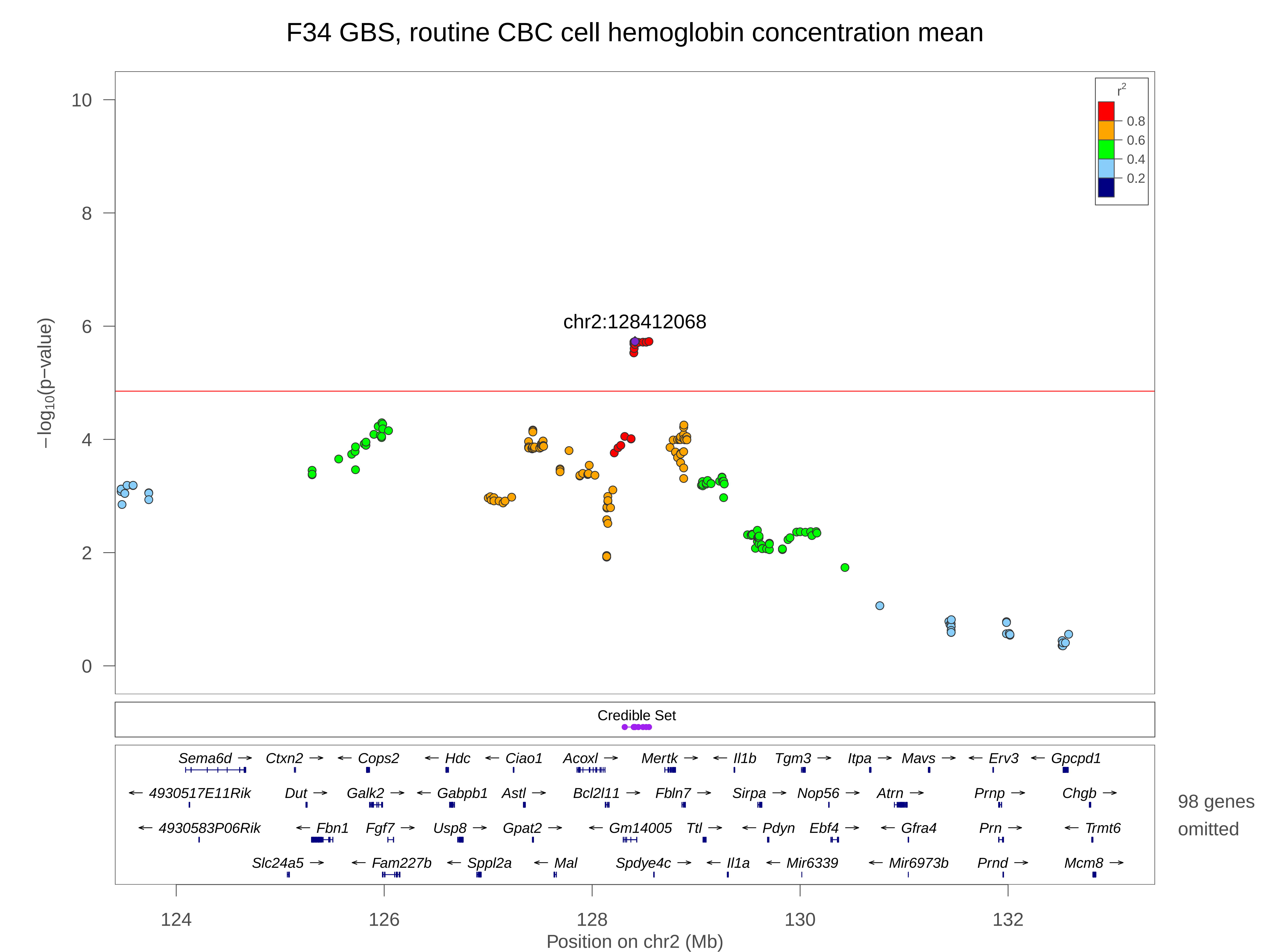

F34 GBS, routine CBC cell hemoglobin concentration mean
−log10(p−value)
10
8
6
4
2
0
chr2:128412068
r2
0.8
0.6
0.4
0.2
Credible Set
Sema6d → Ctxn2 → ← Cops2 ← Hdc ← Ciao1 Acoxl → Mertk → ← Il1b Tgm3 → Itpa → Mavs → ← Erv3 ← Gpcpd1
← 4930517E11Rik Dut → Galk2 → ← Gabpb1 Astl → Bcl2l11 → Fbln7 → Sirpa → Nop56 → Atrn → Prnp → Chgb →
← 4930583P06Rik ← Fbn1 Fgf7 → Usp8 → Gpat2 → ← Gm14005 Ttl → ← Pdyn Ebf4 → ← Gfra4 Prn → ← Trmt6
Slc24a5 → ← Fam227b ← Sppl2a ← Mal Spdye4c → ← Il1a ← Mir6339 ← Mir6973b Prnd → Mcm8 →
98 genes omitted
124
126
128
130
132
Position on chr2 (Mb)

date: Thu Oct 25 17:12:03 2018
build: F34GBS
display range: chr2:123412068−133412068 [123412068−133412068]
hilite range: 0 − 0 [ 0 − 0 ]
reference SNP: chr2:128412068
number of SNPs plotted: 238
min P−value: 1.86E−6 [chr2:128412068]
max P−value: 4.43E−1 [chr2:132526704]
omitted Genes: Myef2, Slc12a1, Cep152
omitted Genes: Shc4, Eid1, Secisbp2l
omitted Genes: Dtwd1, Atp8b4, Slc27a2
omitted Genes: Usp50, Trpm7, Mir3473g
omitted Genes: Ap4e1, Blvra, Ncaph
omitted Genes: Itpripl1, 1810024B03Rik, Snrnp200
omitted Genes: Tmem127, Stard7, Dusp2
omitted Genes: Adra2b, Fahd2a, Kcnip3
omitted Genes: Prom2, Zfp661, Mrps5
omitted Genes: Mall, Nphp1, 1500011K16Rik
omitted Genes: Bub1, Anapc1, Tmem87b
omitted Genes: Zc3h8, Zc3h6, Polr1b
omitted Genes: Chchd5, AI847159, Slc20a1
omitted Genes: A730036I17Rik, Ckap2l, Gm14023
omitted Genes: Il1bos, F830045P16Rik, 4932416H05Rik
omitted Genes: Stk35, AU015228, Tgm6
omitted Genes: Snrpb, Tmc2, Snord110
omitted Genes: Snord57, Idh3b, Cpxm1
omitted Genes: 1700020A23Rik, Tmem239, Pced1a
omitted Genes: Vps16, Ptpra, 4930473A02Rik
omitted Genes: Mrps26, Oxt, Avp
omitted Genes: Ubox5, Fastkd5, Lzts3
omitted Genes: Ddrgk1, Slc4a11, 4930402H24Rik
omitted Genes: Gm14057, A730017L22Rik, Adam33
omitted Genes: Siglec1, Hspa12b, 1700037H04Rik
omitted Genes: Spef1, Cenpb, Cdc25b
omitted Genes: Ap5s1, Pank2, Mir103−2
omitted Genes: Rnf24, Smox, Adra1d
omitted Genes: 5330413P13Rik, Rassf2, Slc23a2
omitted Genes: Tmem230, Pcna, Cds2
omitted Genes: Prokr2, 4921508D12Rik, 1700026D11Rik
Make more plots at http://csg.sph.umich.edu/locuszoom/
omitted Genes: AU019990, 1110034G24Rik, Crls1

Fine−mapping Regions

| snp | chr | pos (Mb) | pp | group | color |
|---|---|---|---|---|---|
| chr2:128313171 | 2 | 128.3132 | 0.002552786 | Credible Set | purple |
| chr2:128400308 | 2 | 128.4003 | 0.065190969 | Credible Set | purple |
| chr2:128400919 | 2 | 128.4009 | 0.064934418 | Credible Set | purple |
| chr2:128401561 | 2 | 128.4016 | 0.092386329 | Credible Set | purple |
| chr2:128401595 | 2 | 128.4016 | 0.099703027 | Credible Set | purple |
| chr2:128403982 | 2 | 128.4040 | 0.077076500 | Credible Set | purple |
| chr2:128412068 | 2 | 128.4121 | 0.101536886 | Credible Set | purple |
| chr2:128412554 | 2 | 128.4126 | 0.090071239 | Credible Set | purple |
| chr2:128444910 | 2 | 128.4449 | 0.098108275 | Credible Set | purple |
| chr2:128488809 | 2 | 128.4888 | 0.098785461 | Credible Set | purple |
| chr2:128519692 | 2 | 128.5197 | 0.098656916 | Credible Set | purple |
| chr2:128546244 | 2 | 128.5462 | 0.101523739 | Credible Set | purple |

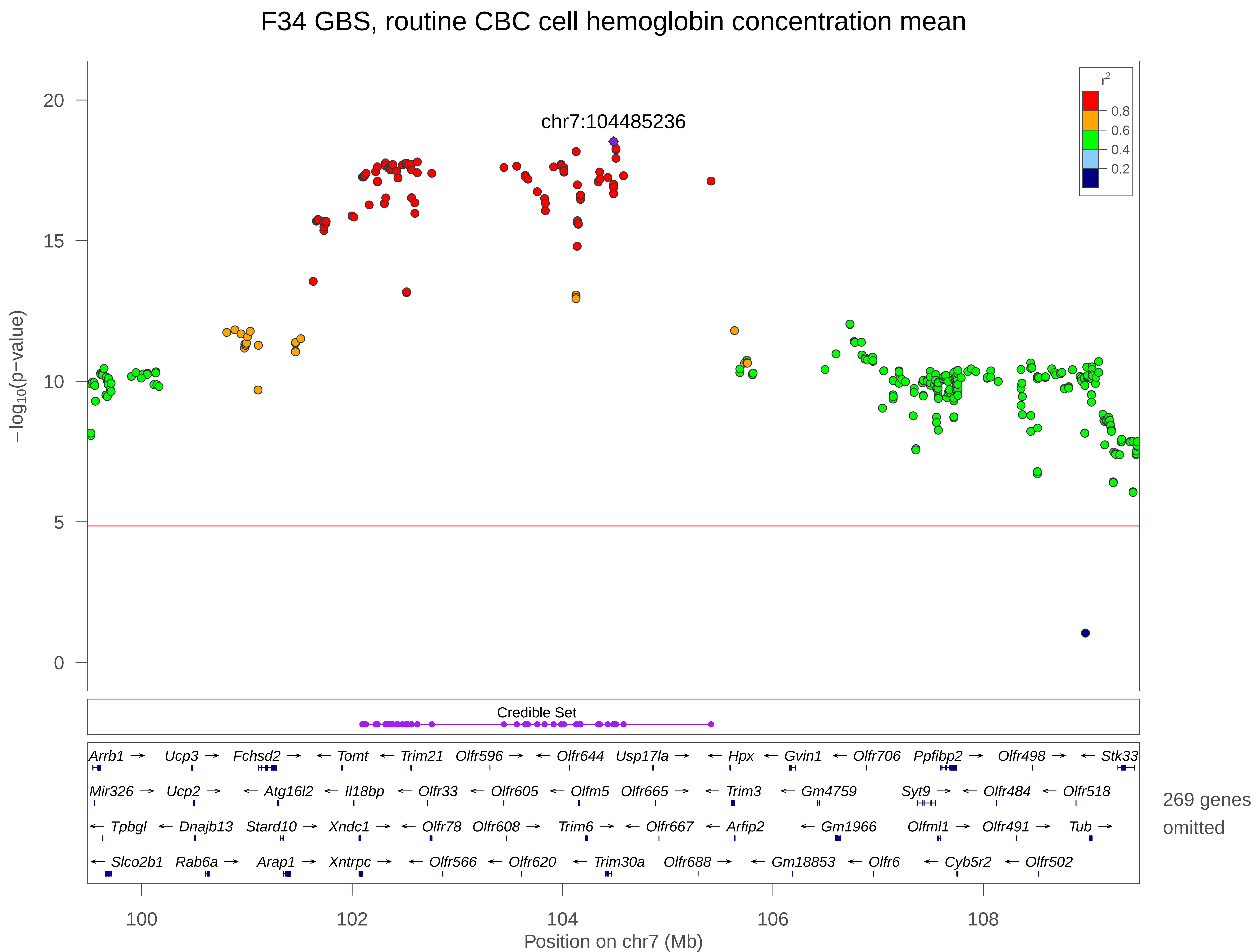

F34 GBS, routine CBC cell hemoglobin concentration mean
chr7:104485236
r2
0.8
0.6
0.4
0.2
−log10(p−value)
20
15
10
5
0
Credible Set
Arrb1 →
Ucp3 →
Fchsd2 →
← Tomt
← Trim21
Olfr596 →
← Olfr644
Usp17la →
← Hpx
← Gvin1
← Olfr706
Ppfibp2 →
Olfr498 →
← Stk33
Mir326 →
Ucp2 →
← Atg16l2
← Il18bp
← Olfr33
← Olfr605
← Olfm5
Olfr665 →
← Trim3
← Gm4759
Syt9 →
← Olfr484
← Olfr518
← Tpbgl
← Dnajb13
Stard10 →
Xndc1 →
← Olfr78
Olfr608 →
Trim6 →
← Olfr667
← Arfip2
← Gm1966
Olfml1 →
Olfr491 →
Tub →
← Slco2b1
Rab6a →
Arap1 →
Xntrpc →
← Olfr566
← Olfr620
← Trim30a
Olfr688 →
← Gm18853
← Olfr6
← Cyb5r2
← Olfr502
269 genes omitted
100
102
104
106
108
Position on chr7 (Mb)

date: Thu Oct 25 17:12:15 2018
build: F34GBS
display range: chr7:99485236−109485236 [99485236−109485236]
hilite range: 0 − 0 [ 0 − 0 ]
reference SNP: chr7:104485236
number of SNPs plotted: 448
min P−value: 2.98E−19 [chr7:104485236]
max P−value: 9.1E−2 [chr7:108970724]
omitted Genes: Olfr520, Olfr521, Neu3
omitted Genes: Spcs2, Xrra1, Rnf169
omitted Genes: Chrdl2, Pold3, Lipt2
omitted Genes: Kcne3, Pgm2l1, Gpx2−ps1
omitted Genes: P4ha3, Ppme1, C2cd3
omitted Genes: Coa4, Mrpl48, Plekhb1
omitted Genes: Fam168a, Relt, Arhgef17
omitted Genes: Mir3102, P2ry6, P2ry2
omitted Genes: Pde2a, Mir139, Art2a−ps
omitted Genes: Art2b, Clpb, Phox2a
omitted Genes: Inppl1, Folr2, Folr1
omitted Genes: Anapc15, Lamtor1, Lrrc51
omitted Genes: Numa1, Rnf121, Trpc2
omitted Genes: Art5, Art1, Chrna10
omitted Genes: Nup98, Pgap2, Rhog
omitted Genes: Stim1, Rrm1, Olfr543
omitted Genes: Olfr544, Olfr545, Olfr547
omitted Genes: Olfr549, Olfr550, Olfr551
omitted Genes: Olfr552, Olfr553, Olfr554
omitted Genes: Olfr555, Olfr556, Trim68
omitted Genes: Olfr557, Olfr558, Olfr559
omitted Genes: Olfr560, Olfr561, Olfr564
omitted Genes: Olfr568, Olfr569, Olfr570
omitted Genes: Olfr571, Olfr572, Olfr574
omitted Genes: Olfr575, Olfr576, Olfr577
omitted Genes: Olfr578, Olfr582, Olfr583
omitted Genes: Olfr584, Olfr585, Olfr586
omitted Genes: Olfr589, Olfr591, Olfr592
omitted Genes: Olfr593, Olfr594, Usp17ld
omitted Genes: Olfr597, Olfr598, Olfr599
omitted Genes: Olfr600, Olfr601, Olfr603
omitted Genes: Usp17lc, Olfr606, Olfr609

Make more plots at http://csg.sph.umich.edu/locuszoom/

### Fine−mapping Regions

| snp | chr | pos (Mb) | pp | group | color |
|---|---|---|---|---|---|
| chr7:102098054 | 7 | 102.0981 | 0.005972896 | Credible Set | purple |
| chr7:102110109 | 7 | 102.1101 | 0.006094195 | Credible Set | purple |
| chr7:102110501 | 7 | 102.1105 | 0.005982939 | Credible Set | purple |
| chr7:102116318 | 7 | 102.1163 | 0.006449616 | Credible Set | purple |
| chr7:102116430 | 7 | 102.1164 | 0.006449616 | Credible Set | purple |
| chr7:102116478 | 7 | 102.1165 | 0.006918295 | Credible Set | purple |
| chr7:102132562 | 7 | 102.1326 | 0.008098267 | Credible Set | purple |
| chr7:102223396 | 7 | 102.2234 | 0.009250125 | Credible Set | purple |
| chr7:102240695 | 7 | 102.2407 | 0.013700692 | Credible Set | purple |
| chr7:102241191 | 7 | 102.2412 | 0.004215671 | Credible Set | purple |
| chr7:102241211 | 7 | 102.2412 | 0.004050529 | Credible Set | purple |
| chr7:102241212 | 7 | 102.2412 | 0.004208170 | Credible Set | purple |
| chr7:102317213 | 7 | 102.3172 | 0.018798761 | Credible Set | purple |
| chr7:102317246 | 7 | 102.3172 | 0.014858524 | Credible Set | purple |
| chr7:102319586 | 7 | 102.3196 | 0.001107848 | Credible Set | purple |
| chr7:102344970 | 7 | 102.3450 | 0.012218346 | Credible Set | purple |
| chr7:102363897 | 7 | 102.3639 | 0.011533835 | Credible Set | purple |
| chr7:102364204 | 7 | 102.3642 | 0.011806281 | Credible Set | purple |
| chr7:102364223 | 7 | 102.3642 | 0.013118590 | Credible Set | purple |
| chr7:102364315 | 7 | 102.3643 | 0.012160089 | Credible Set | purple |
| chr7:102364326 | 7 | 102.3643 | 0.012160089 | Credible Set | purple |
| chr7:102364411 | 7 | 102.3644 | 0.010644735 | Credible Set | purple |
| chr7:102385520 | 7 | 102.3855 | 0.010879315 | Credible Set | purple |
| chr7:102385536 | 7 | 102.3855 | 0.016478430 | Credible Set | purple |
| chr7:102423165 | 7 | 102.4232 | 0.009460741 | Credible Set | purple |
| chr7:102435692 | 7 | 102.4357 | 0.005546541 | Credible Set | purple |
| chr7:102435720 | 7 | 102.4357 | 0.005546541 | Credible Set | purple |
| chr7:102478469 | 7 | 102.4785 | 0.016049128 | Credible Set | purple |
| chr7:102510475 | 7 | 102.5105 | 0.017343491 | Credible Set | purple |
| chr7:102510607 | 7 | 102.5106 | 0.018257457 | Credible Set | purple |
| chr7:102516878 | 7 | 102.5169 | 0.018218312 | Credible Set | purple |
| chr7:102531652 | 7 | 102.5317 | 0.016054242 | Credible Set | purple |
| chr7:102561708 | 7 | 102.5617 | 0.016638150 | Credible Set | purple |
| chr7:102565262 | 7 | 102.5653 | 0.001122496 | Credible Set | purple |
| chr7:102565292 | 7 | 102.5653 | 0.001113980 | Credible Set | purple |
| chr7:102565307 | 7 | 102.5653 | 0.001111050 | Credible Set | purple |

Fine−mapping Regions

| snp | chr | pos (Mb) | pp | group | color |
|---|---|---|---|---|---|
| chr7:102565323 | 7 | 102.5653 | 0.0011026237 | Credible Set | purple |
| chr7:102565329 | 7 | 102.5653 | 0.0011026237 | Credible Set | purple |
| chr7:102565410 | 7 | 102.5654 | 0.0105738196 | Credible Set | purple |
| chr7:102619418 | 7 | 102.6194 | 0.0202781359 | Credible Set | purple |
| chr7:102619431 | 7 | 102.6194 | 0.0202198054 | Credible Set | purple |
| chr7:102619466 | 7 | 102.6195 | 0.0084858925 | Credible Set | purple |
| chr7:102757499 | 7 | 102.7575 | 0.0081127224 | Credible Set | purple |
| chr7:103442570 | 7 | 103.4426 | 0.0129377802 | Credible Set | purple |
| chr7:103564836 | 7 | 103.5648 | 0.0143513009 | Credible Set | purple |
| chr7:103646439 | 7 | 103.6464 | 0.0068125725 | Credible Set | purple |
| chr7:103646709 | 7 | 103.6467 | 0.0060312839 | Credible Set | purple |
| chr7:103671033 | 7 | 103.6710 | 0.0050477275 | Credible Set | purple |
| chr7:103760301 | 7 | 103.7603 | 0.0018296339 | Credible Set | purple |
| chr7:103829200 | 7 | 103.8292 | 0.0010534099 | Credible Set | purple |
| chr7:103915503 | 7 | 103.9155 | 0.0135794009 | Credible Set | purple |
| chr7:103986792 | 7 | 103.9868 | 0.0166677107 | Credible Set | purple |
| chr7:103986795 | 7 | 103.9868 | 0.0166677107 | Credible Set | purple |
| chr7:103986857 | 7 | 103.9869 | 0.0152014253 | Credible Set | purple |
| chr7:103986860 | 7 | 103.9869 | 0.0152014253 | Credible Set | purple |
| chr7:104012808 | 7 | 104.0128 | 0.0126737350 | Credible Set | purple |
| chr7:104013370 | 7 | 104.0134 | 0.0101781509 | Credible Set | purple |
| chr7:104013395 | 7 | 104.0134 | 0.0088625137 | Credible Set | purple |
| chr7:104013467 | 7 | 104.0135 | 0.0096893227 | Credible Set | purple |
| chr7:104129734 | 7 | 104.1297 | 0.0467498262 | Credible Set | purple |
| chr7:104129736 | 7 | 104.1297 | 0.0467498262 | Credible Set | purple |
| chr7:104141343 | 7 | 104.1413 | 0.0031826073 | Credible Set | purple |
| chr7:104170479 | 7 | 104.1705 | 0.0010029286 | Credible Set | purple |
| chr7:104170506 | 7 | 104.1705 | 0.0010005741 | Credible Set | purple |
| chr7:104170522 | 7 | 104.1705 | 0.0009976148 | Credible Set | purple |
| chr7:104170588 | 7 | 104.1706 | 0.0013779504 | Credible Set | purple |
| chr7:104170594 | 7 | 104.1706 | 0.0013657787 | Credible Set | purple |
| chr7:104170618 | 7 | 104.1706 | 0.0013900693 | Credible Set | purple |
| chr7:104170662 | 7 | 104.1707 | 0.0013900693 | Credible Set | purple |
| chr7:104338784 | 7 | 104.3388 | 0.0040315581 | Credible Set | purple |
| chr7:104338810 | 7 | 104.3388 | 0.0040597580 | Credible Set | purple |
| chr7:104353491 | 7 | 104.3535 | 0.0089776067 | Credible Set | purple |

Fine−mapping Regions

| snp | chr | pos (Mb) | pp | group | color |
|---|---|---|---|---|---|
| chr7:104356230 | 7 | 104.3562 | 0.005120499 | Credible Set | purple |
| chr7:104430205 | 7 | 104.4302 | 0.005773948 | Credible Set | purple |
| chr7:104485236 | 7 | 104.4852 | 0.105711391 | Credible Set | purple |
| chr7:104485721 | 7 | 104.4857 | 0.003369688 | Credible Set | purple |
| chr7:104485727 | 7 | 104.4857 | 0.002568336 | Credible Set | purple |
| chr7:104486022 | 7 | 104.4860 | 0.001532997 | Credible Set | purple |
| chr7:104486041 | 7 | 104.4860 | 0.001573481 | Credible Set | purple |
| chr7:104507724 | 7 | 104.5077 | 0.027210172 | Credible Set | purple |
| chr7:104508127 | 7 | 104.5081 | 0.053826633 | Credible Set | purple |
| chr7:104508173 | 7 | 104.5082 | 0.060422735 | Credible Set | purple |
| chr7:104580091 | 7 | 104.5801 | 0.006650589 | Credible Set | purple |
| chr7:105411752 | 7 | 105.4118 | 0.004324642 | Credible Set | purple |

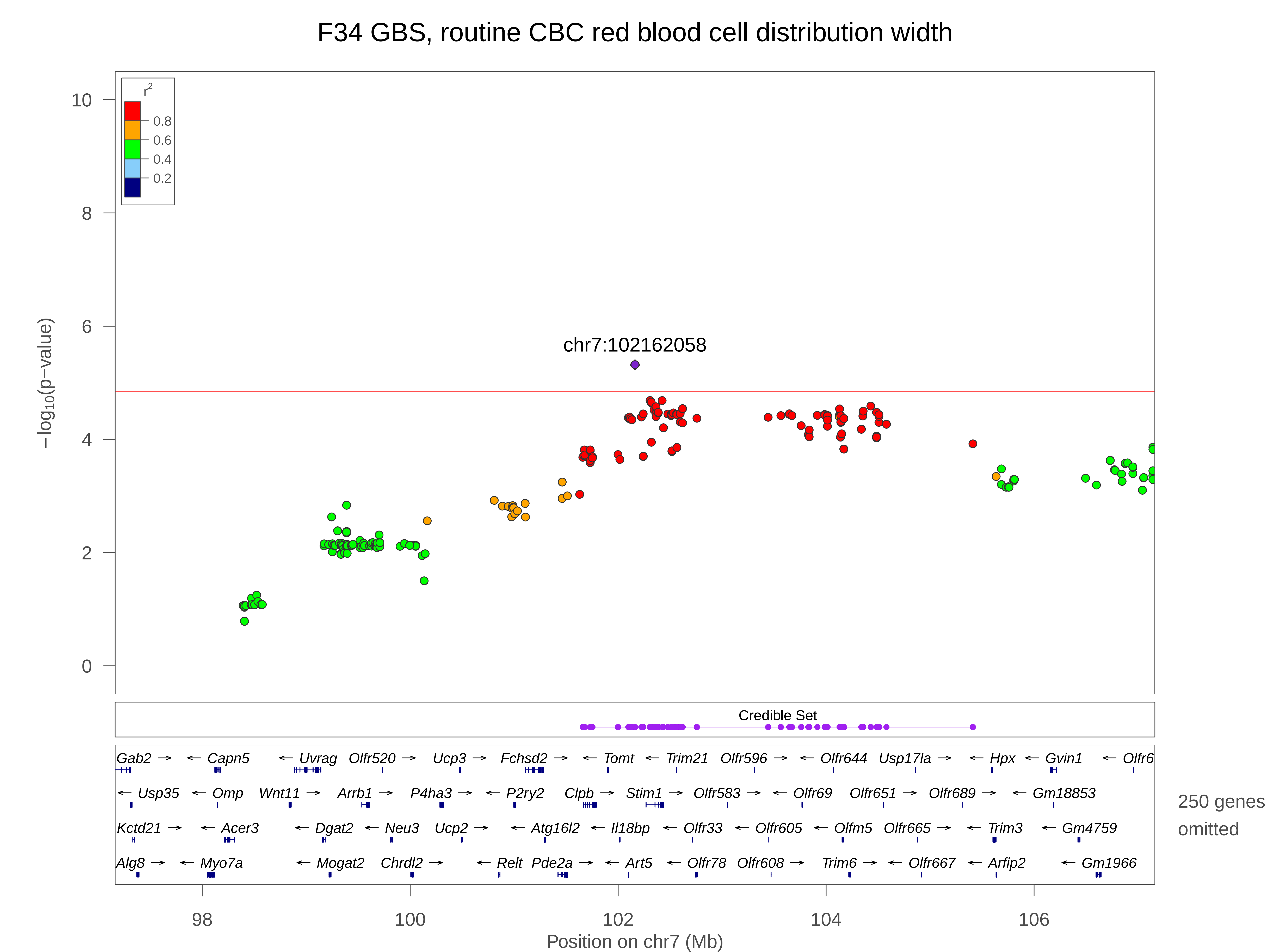
F34 GBS, routine CBC red blood cell distribution width
r2
0.8
0.6
0.4
0.2
10
8
6
4
2
0
−log10(p−value)
chr7:102162058
Credible Set
Gab2 → ← Capn5 ← Uvrag Olfr520 → Ucp3 → Fchsd2 → ← Tomt ← Trim21 Olfr596 → ← Olfr644 Usp17la → ← Hpx ← Gvin1 ← Olfr6
← Usp35 ← Omp Wnt11 → Arrb1 → P4ha3 → ← P2ry2 Clpb → Stim1 → Olfr583 → ← Olfr69 Olfr651 → Olfr689 → ← Gm18853
Kctd21 → ← Acer3 ← Dgat2 ← Neu3 Ucp2 → ← Atg16l2 ← Il18bp ← Olfr33 ← Olfr605 ← Olfm5 Olfr665 → ← Trim3 ← Gm4759
Alg8 → ← Myo7a ← Mogat2 Chrdl2 → ← Relt Pde2a → ← Art5 ← Olfr78 Olfr608 → Trim6 → ← Olfr667 ← Arfip2 ← Gm1966
250 genes omitted
98
100
102
104
106
Position on chr7 (Mb)

date: Thu Oct 25 17:12:39 2018

build: F34GBS

display range: chr7:97162058−107162058 [97162058−107162058]

hilite range: 0 − 0 [ 0 − 0 ]

reference SNP: chr7:102162058

number of SNPs plotted: 295

min P−value: 4.78E−6 [chr7:102162058]

max P−value: 1.64E−1 [chr7:98406311]

omitted Genes: Ndufc2, Thrsp, Kctd14

omitted Genes: Ints4, Aamdc, Rsf1

omitted Genes: Rsf1os1, Clns1a, Aqp11

omitted Genes: Pak1, Gdpd4, Gm16938

omitted Genes: B3gnt6, Tsku, Gucy2d

omitted Genes: Lrrc32, Emsy, Prkrir

omitted Genes: Gm26705, Map6, Serpinh1

omitted Genes: Gdpd5, Klhl35, Rps3

omitted Genes: Snord15b, Snord15a, Mir326

omitted Genes: Tpbgl, Slco2b1, Olfr521

omitted Genes: Spcs2, Xrra1, Rnf169

omitted Genes: Pold3, Lipt2, Kcne3

omitted Genes: Pgm2l1, Gpx2−ps1, Ppme1

omitted Genes: C2cd3, Dnajb13, Coa4

omitted Genes: Mrpl48, Rab6a, Plekhb1

omitted Genes: Fam168a, Arhgef17, Mir3102

omitted Genes: P2ry6, Stard10, Arap1

omitted Genes: Mir139, Art2a−ps, Art2b

omitted Genes: Phox2a, Inppl1, Folr2

omitted Genes: Folr1, Anapc15, Lamtor1

omitted Genes: Lrrc51, Numa1, Rnf121

omitted Genes: Xndc1, Xntrpc, Trpc2

omitted Genes: Art1, Chrna10, Nup98

omitted Genes: Pgap2, Rhog, Rrm1

omitted Genes: Olfr543, Olfr544, Olfr545

omitted Genes: Olfr547, Olfr549, Olfr550

omitted Genes: Olfr551, Olfr552, Olfr553

omitted Genes: Olfr554, Olfr555, Olfr556

omitted Genes: Trim68, Olfr557, Olfr558

omitted Genes: Olfr559, Olfr560, Olfr561

omitted Genes: Olfr564, Olfr566, Olfr568

Make more plots at http://csg.sph.umich.edu/locuszoom/

omitted Genes: Olfr569, Olfr570, Olfr571

Fine−mapping Regions

| snp | chr | pos (Mb) | pp | group | color |
|---|---|---|---|---|---|
| chr7:101660383 | 7 | 101.6604 | 0.002089386 | Credible Set | purple |
| chr7:101669186 | 7 | 101.6692 | 0.002230006 | Credible Set | purple |
| chr7:101673483 | 7 | 101.6735 | 0.002772393 | Credible Set | purple |
| chr7:101680140 | 7 | 101.6801 | 0.002283530 | Credible Set | purple |
| chr7:101731061 | 7 | 101.7311 | 0.002659947 | Credible Set | purple |
| chr7:101731110 | 7 | 101.7311 | 0.002775208 | Credible Set | purple |
| chr7:101751790 | 7 | 101.7518 | 0.002124253 | Credible Set | purple |
| chr7:101752580 | 7 | 101.7526 | 0.002142424 | Credible Set | purple |
| chr7:101998679 | 7 | 101.9987 | 0.002318842 | Credible Set | purple |
| chr7:102098054 | 7 | 102.0981 | 0.009548508 | Credible Set | purple |
| chr7:102110109 | 7 | 102.1101 | 0.009503956 | Credible Set | purple |
| chr7:102110501 | 7 | 102.1105 | 0.009837912 | Credible Set | purple |
| chr7:102116318 | 7 | 102.1163 | 0.009043904 | Credible Set | purple |
| chr7:102116430 | 7 | 102.1164 | 0.009043904 | Credible Set | purple |
| chr7:102116478 | 7 | 102.1165 | 0.009176579 | Credible Set | purple |
| chr7:102132562 | 7 | 102.1326 | 0.008802086 | Credible Set | purple |
| chr7:102162058 | 7 | 102.1621 | 0.074805787 | Credible Set | purple |
| chr7:102223396 | 7 | 102.2234 | 0.009811606 | Credible Set | purple |
| chr7:102240695 | 7 | 102.2407 | 0.011072285 | Credible Set | purple |
| chr7:102241191 | 7 | 102.2412 | 0.002179105 | Credible Set | purple |
| chr7:102241211 | 7 | 102.2412 | 0.002151102 | Credible Set | purple |
| chr7:102241212 | 7 | 102.2412 | 0.002179322 | Credible Set | purple |
| chr7:102306968 | 7 | 102.3070 | 0.018536215 | Credible Set | purple |
| chr7:102306978 | 7 | 102.3070 | 0.018538116 | Credible Set | purple |
| chr7:102317213 | 7 | 102.3172 | 0.017128207 | Credible Set | purple |
| chr7:102317246 | 7 | 102.3172 | 0.017508287 | Credible Set | purple |
| chr7:102319586 | 7 | 102.3196 | 0.003715643 | Credible Set | purple |
| chr7:102344970 | 7 | 102.3450 | 0.012892797 | Credible Set | purple |
| chr7:102363897 | 7 | 102.3639 | 0.010794577 | Credible Set | purple |
| chr7:102364204 | 7 | 102.3642 | 0.012873946 | Credible Set | purple |
| chr7:102364223 | 7 | 102.3642 | 0.014588118 | Credible Set | purple |
| chr7:102364315 | 7 | 102.3643 | 0.011194140 | Credible Set | purple |
| chr7:102364326 | 7 | 102.3643 | 0.011194140 | Credible Set | purple |
| chr7:102364411 | 7 | 102.3644 | 0.009946970 | Credible Set | purple |
| chr7:102385520 | 7 | 102.3855 | 0.011458601 | Credible Set | purple |
| chr7:102385536 | 7 | 102.3855 | 0.011787086 | Credible Set | purple |

Fine−mapping Regions

| snp | chr | pos (Mb) | pp | group | color |
|---|---|---|---|---|---|
| chr7:102423165 | 7 | 102.4232 | 0.018541508 | Credible Set | purple |
| chr7:102435692 | 7 | 102.4357 | 0.006490996 | Credible Set | purple |
| chr7:102435720 | 7 | 102.4357 | 0.006490996 | Credible Set | purple |
| chr7:102478469 | 7 | 102.4785 | 0.011022684 | Credible Set | purple |
| chr7:102510475 | 7 | 102.5105 | 0.010378824 | Credible Set | purple |
| chr7:102510607 | 7 | 102.5106 | 0.010981256 | Credible Set | purple |
| chr7:102516878 | 7 | 102.5169 | 0.010650670 | Credible Set | purple |
| chr7:102517238 | 7 | 102.5172 | 0.002603268 | Credible Set | purple |
| chr7:102517241 | 7 | 102.5172 | 0.002601938 | Credible Set | purple |
| chr7:102517247 | 7 | 102.5172 | 0.002675521 | Credible Set | purple |
| chr7:102531652 | 7 | 102.5317 | 0.011523240 | Credible Set | purple |
| chr7:102561708 | 7 | 102.5617 | 0.011019190 | Credible Set | purple |
| chr7:102565262 | 7 | 102.5653 | 0.003022905 | Credible Set | purple |
| chr7:102565292 | 7 | 102.5653 | 0.003041713 | Credible Set | purple |
| chr7:102565307 | 7 | 102.5653 | 0.003012704 | Credible Set | purple |
| chr7:102565323 | 7 | 102.5653 | 0.003031443 | Credible Set | purple |
| chr7:102565329 | 7 | 102.5653 | 0.003031443 | Credible Set | purple |
| chr7:102565410 | 7 | 102.5654 | 0.010811193 | Credible Set | purple |
| chr7:102596442 | 7 | 102.5964 | 0.011150038 | Credible Set | purple |
| chr7:102596578 | 7 | 102.5966 | 0.008116217 | Credible Set | purple |
| chr7:102596636 | 7 | 102.5966 | 0.008119380 | Credible Set | purple |
| chr7:102619418 | 7 | 102.6194 | 0.013546316 | Credible Set | purple |
| chr7:102619431 | 7 | 102.6194 | 0.013583107 | Credible Set | purple |
| chr7:102619466 | 7 | 102.6195 | 0.007825871 | Credible Set | purple |
| chr7:102757499 | 7 | 102.7575 | 0.009370125 | Credible Set | purple |
| chr7:103442570 | 7 | 103.4426 | 0.009727824 | Credible Set | purple |
| chr7:103564836 | 7 | 103.5648 | 0.010363188 | Credible Set | purple |
| chr7:103646439 | 7 | 103.6464 | 0.010875360 | Credible Set | purple |
| chr7:103646709 | 7 | 103.6467 | 0.011132242 | Credible Set | purple |
| chr7:103671033 | 7 | 103.6710 | 0.010389433 | Credible Set | purple |
| chr7:103760301 | 7 | 103.7603 | 0.007035344 | Credible Set | purple |
| chr7:103829200 | 7 | 103.8292 | 0.004976044 | Credible Set | purple |
| chr7:103837099 | 7 | 103.8371 | 0.004571138 | Credible Set | purple |
| chr7:103837213 | 7 | 103.8372 | 0.005961047 | Credible Set | purple |
| chr7:103915503 | 7 | 103.9155 | 0.010463737 | Credible Set | purple |
| chr7:103986792 | 7 | 103.9868 | 0.010812874 | Credible Set | purple |

Fine−mapping Regions

| snp | chr | pos (Mb) | pp | group | color |
|---|---|---|---|---|---|
| chr7:103986795 | 7 | 103.9868 | 0.010812874 | Credible Set | purple |
| chr7:103986857 | 7 | 103.9869 | 0.010624554 | Credible Set | purple |
| chr7:103986860 | 7 | 103.9869 | 0.010624554 | Credible Set | purple |
| chr7:104012808 | 7 | 104.0128 | 0.006875938 | Credible Set | purple |
| chr7:104013370 | 7 | 104.0134 | 0.010436535 | Credible Set | purple |
| chr7:104013395 | 7 | 104.0134 | 0.010211190 | Credible Set | purple |
| chr7:104013467 | 7 | 104.0135 | 0.008727574 | Credible Set | purple |
| chr7:104129734 | 7 | 104.1297 | 0.013437577 | Credible Set | purple |
| chr7:104129736 | 7 | 104.1297 | 0.013437577 | Credible Set | purple |
| chr7:104138934 | 7 | 104.1389 | 0.004518107 | Credible Set | purple |
| chr7:104138936 | 7 | 104.1389 | 0.004528438 | Credible Set | purple |
| chr7:104138968 | 7 | 104.1390 | 0.004528438 | Credible Set | purple |
| chr7:104141343 | 7 | 104.1413 | 0.010287885 | Credible Set | purple |
| chr7:104141705 | 7 | 104.1417 | 0.007971042 | Credible Set | purple |
| chr7:104141768 | 7 | 104.1418 | 0.008141437 | Credible Set | purple |
| chr7:104149893 | 7 | 104.1499 | 0.005108275 | Credible Set | purple |
| chr7:104149911 | 7 | 104.1499 | 0.005143698 | Credible Set | purple |
| chr7:104170479 | 7 | 104.1705 | 0.009227558 | Credible Set | purple |
| chr7:104170506 | 7 | 104.1705 | 0.009263429 | Credible Set | purple |
| chr7:104170518 | 7 | 104.1705 | 0.009227143 | Credible Set | purple |
| chr7:104170522 | 7 | 104.1705 | 0.009254306 | Credible Set | purple |
| chr7:104170543 | 7 | 104.1705 | 0.009248551 | Credible Set | purple |
| chr7:104170588 | 7 | 104.1706 | 0.002873580 | Credible Set | purple |
| chr7:104170594 | 7 | 104.1706 | 0.002857667 | Credible Set | purple |
| chr7:104170618 | 7 | 104.1706 | 0.002863148 | Credible Set | purple |
| chr7:104170662 | 7 | 104.1707 | 0.002863148 | Credible Set | purple |
| chr7:104338784 | 7 | 104.3388 | 0.006147177 | Credible Set | purple |
| chr7:104338810 | 7 | 104.3388 | 0.006117691 | Credible Set | purple |
| chr7:104353491 | 7 | 104.3535 | 0.010200340 | Credible Set | purple |
| chr7:104356230 | 7 | 104.3562 | 0.012345949 | Credible Set | purple |
| chr7:104430205 | 7 | 104.4302 | 0.014991492 | Credible Set | purple |
| chr7:104485236 | 7 | 104.4852 | 0.011749030 | Credible Set | purple |
| chr7:104485721 | 7 | 104.4857 | 0.004407447 | Credible Set | purple |
| chr7:104485727 | 7 | 104.4857 | 0.004419985 | Credible Set | purple |
| chr7:104486022 | 7 | 104.4860 | 0.004691478 | Credible Set | purple |
| chr7:104486041 | 7 | 104.4860 | 0.004526093 | Credible Set | purple |

Fine−mapping Regions

| snp | chr | pos (Mb) | pp | group | color |
|---|---|---|---|---|---|
| chr7:104507724 | 7 | 104.5077 | 0.007987074 | Credible Set | purple |
| chr7:104508127 | 7 | 104.5081 | 0.010042658 | Credible Set | purple |
| chr7:104508173 | 7 | 104.5082 | 0.010682151 | Credible Set | purple |
| chr7:104580091 | 7 | 104.5801 | 0.007416009 | Credible Set | purple |
| chr7:105411752 | 7 | 105.4118 | 0.003497803 | Credible Set | purple |

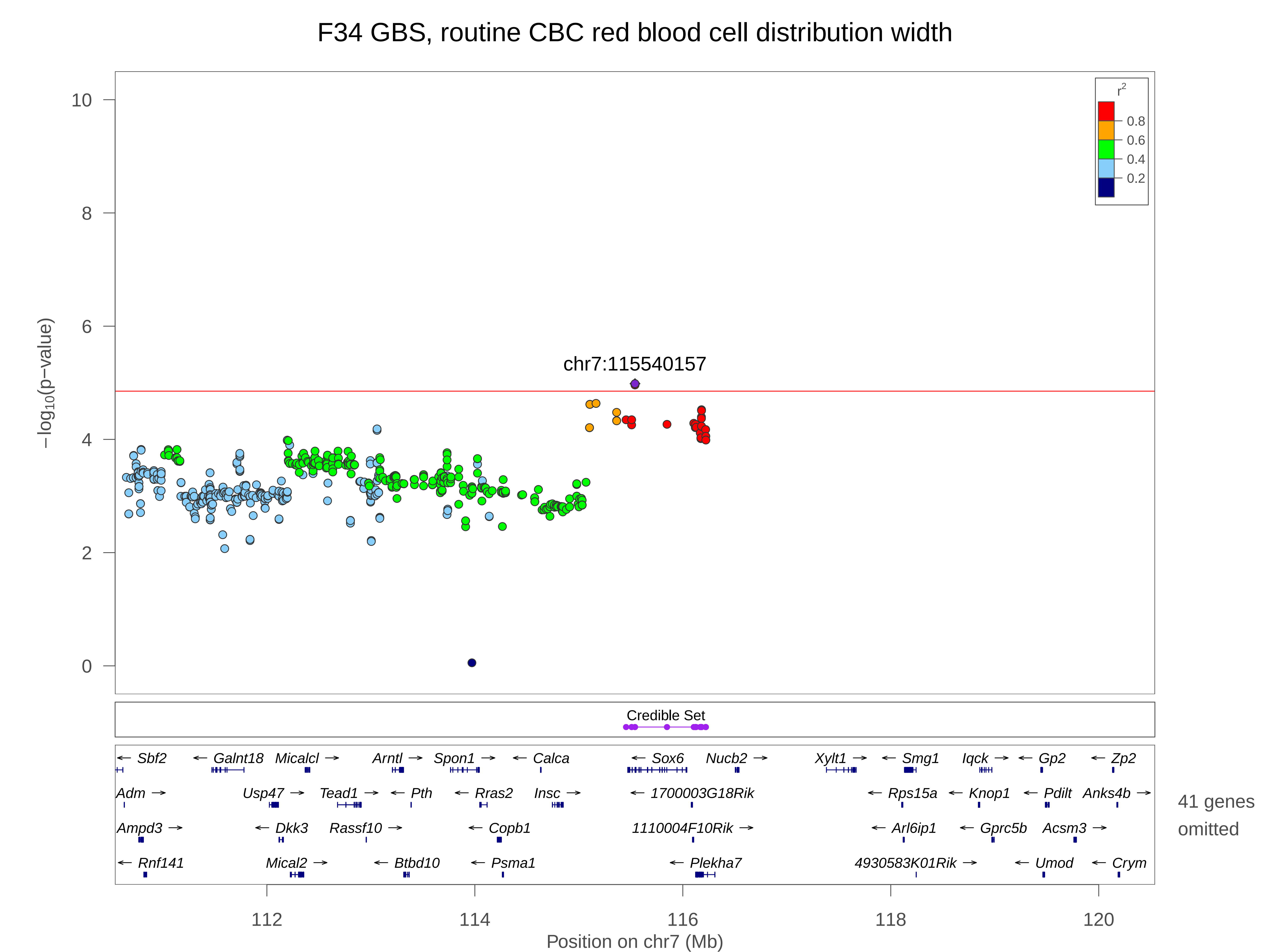

F34 GBS, routine CBC red blood cell distribution width
−log10(p−value)
10
8
6
4
2
0
r2
0.8
0.6
0.4
0.2
chr7:115540157
Credible Set
← Sbf2
← Galnt18
Micalcl →
Arntl →
Spon1 →
← Calca
← Sox6
Nucb2 →
Xylt1 →
← Smg1
Iqck →
← Gp2
← Zp2
Adm →
Usp47 →
Tead1 →
← Pth
← Rras2
Insc →
← 1700003G18Rik
← Rps15a
← Knop1
← Pdilt
Anks4b →
Ampd3 →
← Dkk3
Rassf10 →
← Copb1
1110004F10Rik →
← Arl6ip1
← Gprc5b
Acsm3 →
← Rnf141
Mical2 →
← Btbd10
← Psma1
← Plekha7
4930583K01Rik →
← Umod
← Crym
41 genes omitted
112
114
116
118
120
Position on chr7 (Mb)

| | |
|---|---|
| date: | Thu Oct 25 17:12:50 2018 |
| build: | F34GBS |
| display range: | chr7:110540157−120540157 [110540157−120540157] |
| hilite range: | 0 − 0 [ 0 − 0 ] |
| reference SNP: | chr7:115540157 |
| number of SNPs plotted: | 670 |
| min P−value: | 1.04E−5 [chr7:115540157] |
| max P−value: | 8.83E−1 [chr7:113971813] |
| omitted Genes: | Lyve1, Mrvi1, Gm16336 |
| omitted Genes: | Ctr9, Eif4g2, 1700012D14Rik |
| omitted Genes: | Parva, 4930543E12Rik, Far1 |
| omitted Genes: | Far1os, 4933406I18Rik, Pde3b |
| omitted Genes: | Cyp2r1, Calcb, A730082K24Rik |
| omitted Genes: | Rps13, Snord14a, Pik3c2a |
| omitted Genes: | Syt17, Itpripl2, Coq7 |
| omitted Genes: | Tmc7, Tmc5, Gde1 |
| omitted Genes: | Ccp110, 9030624J02Rik, Gpr139 |
| omitted Genes: | Acsm5, Acsm2, Acsm1 |
| omitted Genes: | Acsm4, Thumpd1, Eri2 |
| omitted Genes: | 2610020H08Rik, Dcun1d3, Lyrm1 |
| omitted Genes: | Tmem159, 4930505K13Rik, Abca14 |
| omitted Genes: | Abca15, Abca16 |

Make more plots at http://csg.sph.umich.edu/locuszoom/

Fine−mapping Regions

| snp | chr | pos (Mb) | pp | group | color |
|---|---|---|---|---|---|
| chr7:115454415 | 7 | 115.4544 | 0.03400282 | Credible Set | purple |
| chr7:115507677 | 7 | 115.5077 | 0.02790172 | Credible Set | purple |
| chr7:115508170 | 7 | 115.5082 | 0.03407172 | Credible Set | purple |
| chr7:115540157 | 7 | 115.5402 | 0.13779863 | Credible Set | purple |
| chr7:115540180 | 7 | 115.5402 | 0.13091693 | Credible Set | purple |
| chr7:115847912 | 7 | 115.8479 | 0.02857463 | Credible Set | purple |
| chr7:116105450 | 7 | 116.1055 | 0.02973166 | Credible Set | purple |
| chr7:116120266 | 7 | 116.1203 | 0.02517664 | Credible Set | purple |
| chr7:116121716 | 7 | 116.1217 | 0.02862464 | Credible Set | purple |
| chr7:116128732 | 7 | 116.1287 | 0.02548667 | Credible Set | purple |
| chr7:116167789 | 7 | 116.1678 | 0.02048942 | Credible Set | purple |
| chr7:116167891 | 7 | 116.1679 | 0.02057105 | Credible Set | purple |
| chr7:116173642 | 7 | 116.1736 | 0.01646756 | Credible Set | purple |
| chr7:116173781 | 7 | 116.1738 | 0.01699213 | Credible Set | purple |
| chr7:116179005 | 7 | 116.1790 | 0.03788025 | Credible Set | purple |
| chr7:116179028 | 7 | 116.1790 | 0.03574226 | Credible Set | purple |
| chr7:116179466 | 7 | 116.1795 | 0.04923355 | Credible Set | purple |
| chr7:116179541 | 7 | 116.1795 | 0.02626363 | Credible Set | purple |
| chr7:116179576 | 7 | 116.1796 | 0.02640221 | Credible Set | purple |
| chr7:116179586 | 7 | 116.1796 | 0.02650116 | Credible Set | purple |
| chr7:116179595 | 7 | 116.1796 | 0.02657195 | Credible Set | purple |
| chr7:116180032 | 7 | 116.1800 | 0.05017761 | Credible Set | purple |
| chr7:116180072 | 7 | 116.1801 | 0.04847339 | Credible Set | purple |
| chr7:116219326 | 7 | 116.2193 | 0.02336931 | Credible Set | purple |
| chr7:116219327 | 7 | 116.2193 | 0.02334005 | Credible Set | purple |
| chr7:116221282 | 7 | 116.2213 | 0.01802601 | Credible Set | purple |
| chr7:116221389 | 7 | 116.2214 | 0.01562628 | Credible Set | purple |
| chr7:116221407 | 7 | 116.2214 | 0.01558612 | Credible Set | purple |

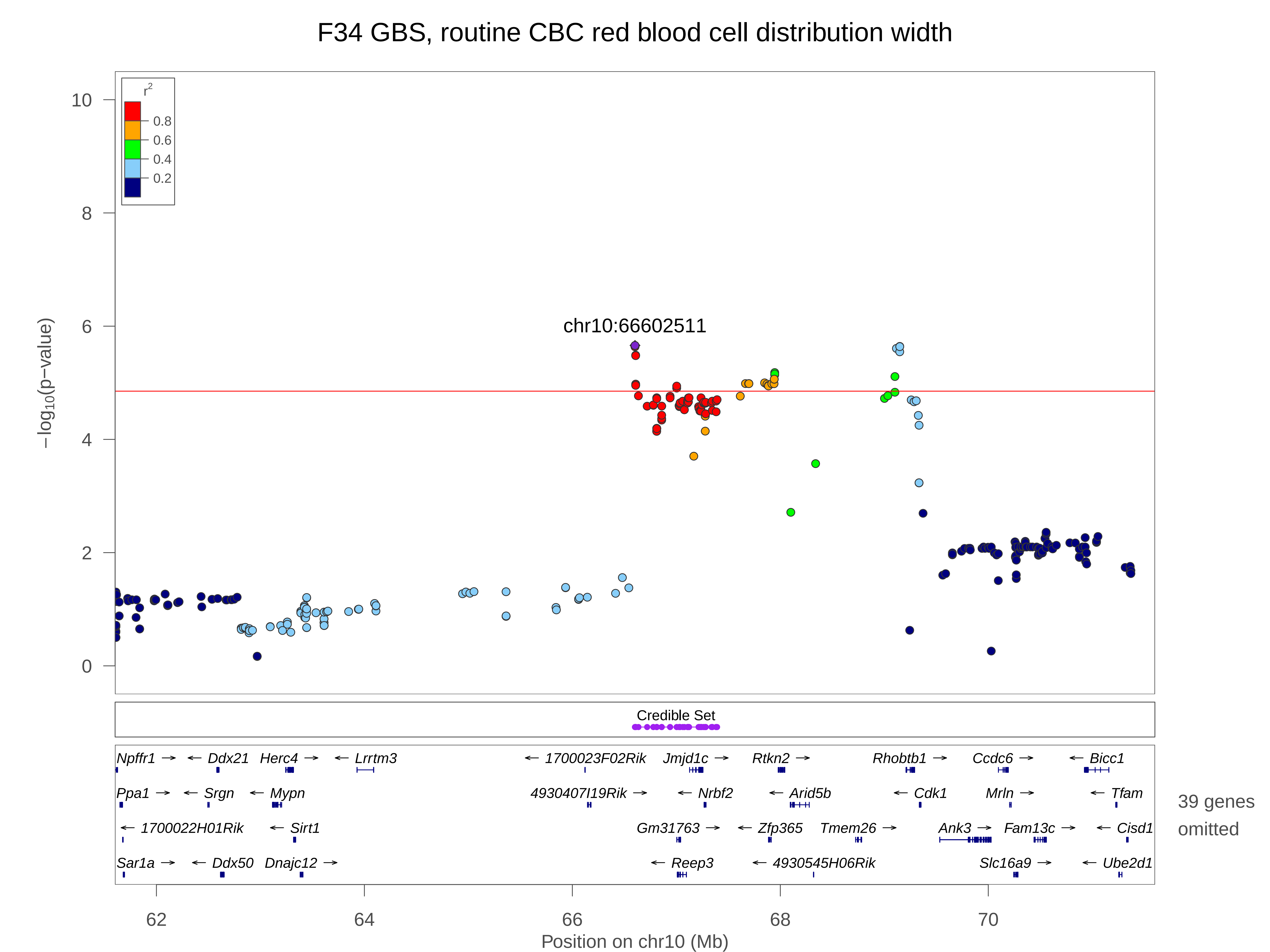

F34 GBS, routine CBC red blood cell distribution width
chr10:66602511
r2
0.8
0.6
0.4
0.2
−log10(p−value)
Credible Set
Npffr1 →
← Ddx21
Herc4 →
← Lrrtm3
← 1700023F02Rik
Jmjd1c →
Rtkn2 →
Rhobtb1 →
Ccdc6 →
← Bicc1
Ppa1 →
← Srgn
← Mypn
4930407I19Rik →
← Nrbf2
← Arid5b
← Cdk1
Mrln →
← Tfam
← 1700022H01Rik
← Sirt1
Gm31763 →
← Zfp365
Tmem26 →
Ank3 →
Fam13c →
← Cisd1
Sar1a →
← Ddx50
Dnajc12 →
← Reep3
← 4930545H06Rik
Slc16a9 →
← Ube2d1
39 genes omitted
Position on chr10 (Mb)

| | |
|---|---|
| date: | Thu Oct 25 17:12:25 2018 |
| build: | F34GBS |
| display range: | chr10:61602511−71602511 [61602511−71602511] |
| hilite range: | 0 − 0 [ 0 − 0 ] |
| reference SNP: | chr10:66602511 |
| number of SNPs plotted: | 346 |
| min P−value: | 2.19E−6 [chr10:66602511] |
| max P−value: | 6.82E−1 [chr10:62969101] |
| omitted Genes: | Tysnd1, Aifm2, H2afy2 |
| omitted Genes: | Mir5108, Col13a1, Gm5424 |
| omitted Genes: | 2010107G23Rik, Neurog3, Tspan15 |
| omitted Genes: | Mir7662, Tacr2, Hk1 |
| omitted Genes: | Hk1os, Hkdc1, Supv3l1 |
| omitted Genes: | 4930507D05Rik, Vps26a, Kif1bp |
| omitted Genes: | Stox1, Mir7215, Ccar1 |
| omitted Genes: | Snord98, Tet1, Slc25a16 |
| omitted Genes: | Dna2, Rufy2, Hnrnph3 |
| omitted Genes: | Pbld2, Pbld1, Atoh7 |
| omitted Genes: | Ctnna3, Egr2, Ado |
| omitted Genes: | 1700040L02Rik, Phyhipl, D630013N20Rik |
| omitted Genes: | C730027H18Rik, 1700113B09Rik, Ipmk |

Make more plots at http://csg.sph.umich.edu/locuszoom/

Fine−mapping Regions

| snp | chr | pos (Mb) | pp | group | color |
|---|---|---|---|---|---|
| chr10:66602509 | 10 | 66.60251 | 0.113276210 | Credible Set | purple |
| chr10:66602511 | 10 | 66.60251 | 0.113276210 | Credible Set | purple |
| chr10:66608856 | 10 | 66.60886 | 0.076752560 | Credible Set | purple |
| chr10:66608885 | 10 | 66.60889 | 0.076752560 | Credible Set | purple |
| chr10:66609052 | 10 | 66.60905 | 0.022131543 | Credible Set | purple |
| chr10:66609080 | 10 | 66.60908 | 0.022131543 | Credible Set | purple |
| chr10:66609124 | 10 | 66.60912 | 0.022131543 | Credible Set | purple |
| chr10:66635369 | 10 | 66.63537 | 0.014610950 | Credible Set | purple |
| chr10:66719524 | 10 | 66.71952 | 0.009750032 | Credible Set | purple |
| chr10:66778997 | 10 | 66.77900 | 0.010120631 | Credible Set | purple |
| chr10:66779040 | 10 | 66.77904 | 0.010120631 | Credible Set | purple |
| chr10:66811497 | 10 | 66.81150 | 0.013836482 | Credible Set | purple |
| chr10:66811529 | 10 | 66.81153 | 0.013141887 | Credible Set | purple |
| chr10:66811581 | 10 | 66.81158 | 0.004147612 | Credible Set | purple |
| chr10:66811611 | 10 | 66.81161 | 0.004147612 | Credible Set | purple |
| chr10:66811619 | 10 | 66.81162 | 0.004147612 | Credible Set | purple |
| chr10:66811684 | 10 | 66.81168 | 0.004147612 | Credible Set | purple |
| chr10:66811694 | 10 | 66.81169 | 0.004147612 | Credible Set | purple |
| chr10:66811705 | 10 | 66.81171 | 0.004087190 | Credible Set | purple |
| chr10:66859150 | 10 | 66.85915 | 0.005671287 | Credible Set | purple |
| chr10:66859163 | 10 | 66.85916 | 0.006045913 | Credible Set | purple |
| chr10:66859201 | 10 | 66.85920 | 0.006972709 | Credible Set | purple |
| chr10:66859671 | 10 | 66.85967 | 0.009750032 | Credible Set | purple |
| chr10:66940094 | 10 | 66.94009 | 0.014610950 | Credible Set | purple |
| chr10:66940121 | 10 | 66.94012 | 0.013141887 | Credible Set | purple |
| chr10:67003583 | 10 | 67.00358 | 0.020367602 | Credible Set | purple |
| chr10:67003588 | 10 | 67.00359 | 0.022131543 | Credible Set | purple |
| chr10:67026213 | 10 | 67.02621 | 0.010120631 | Credible Set | purple |
| chr10:67029219 | 10 | 67.02922 | 0.009750032 | Credible Set | purple |
| chr10:67029237 | 10 | 67.02924 | 0.009750032 | Credible Set | purple |
| chr10:67034406 | 10 | 67.03441 | 0.010521406 | Credible Set | purple |
| chr10:67036263 | 10 | 67.03626 | 0.010956238 | Credible Set | purple |
| chr10:67061736 | 10 | 67.06174 | 0.011947227 | Credible Set | purple |
| chr10:67077389 | 10 | 67.07739 | 0.008509800 | Credible Set | purple |
| chr10:67107858 | 10 | 67.10786 | 0.011429699 | Credible Set | purple |
| chr10:67107945 | 10 | 67.10795 | 0.010956238 | Credible Set | purple |

Fine−mapping Regions

| snp | chr | pos (Mb) | pp | group | color |
|---|---|---|---|---|---|
| chr10:67108562 | 10 | 67.10856 | 0.011947227 | Credible Set | purple |
| chr10:67115847 | 10 | 67.11585 | 0.011429699 | Credible Set | purple |
| chr10:67116164 | 10 | 67.11616 | 0.013836482 | Credible Set | purple |
| chr10:67121830 | 10 | 67.12183 | 0.013836482 | Credible Set | purple |
| chr10:67121898 | 10 | 67.12190 | 0.013836482 | Credible Set | purple |
| chr10:67214197 | 10 | 67.21420 | 0.009750032 | Credible Set | purple |
| chr10:67217456 | 10 | 67.21746 | 0.009086602 | Credible Set | purple |
| chr10:67217505 | 10 | 67.21751 | 0.009086602 | Credible Set | purple |
| chr10:67226747 | 10 | 67.22675 | 0.008248714 | Credible Set | purple |
| chr10:67237234 | 10 | 67.23723 | 0.008509800 | Credible Set | purple |
| chr10:67237236 | 10 | 67.23724 | 0.008003592 | Credible Set | purple |
| chr10:67237389 | 10 | 67.23739 | 0.013836482 | Credible Set | purple |
| chr10:67246731 | 10 | 67.24673 | 0.010120631 | Credible Set | purple |
| chr10:67264982 | 10 | 67.26498 | 0.010956238 | Credible Set | purple |
| chr10:67265380 | 10 | 67.26538 | 0.011429699 | Credible Set | purple |
| chr10:67281545 | 10 | 67.28154 | 0.010956238 | Credible Set | purple |
| chr10:67281940 | 10 | 67.28194 | 0.011429699 | Credible Set | purple |
| chr10:67282089 | 10 | 67.28209 | 0.007350506 | Credible Set | purple |
| chr10:67338317 | 10 | 67.33832 | 0.010956238 | Credible Set | purple |
| chr10:67345731 | 10 | 67.34573 | 0.011947227 | Credible Set | purple |
| chr10:67346184 | 10 | 67.34618 | 0.008248714 | Credible Set | purple |
| chr10:67382855 | 10 | 67.38286 | 0.007773003 | Credible Set | purple |
| chr10:67383022 | 10 | 67.38302 | 0.011947227 | Credible Set | purple |
| chr10:67391783 | 10 | 67.39178 | 0.012515334 | Credible Set | purple |

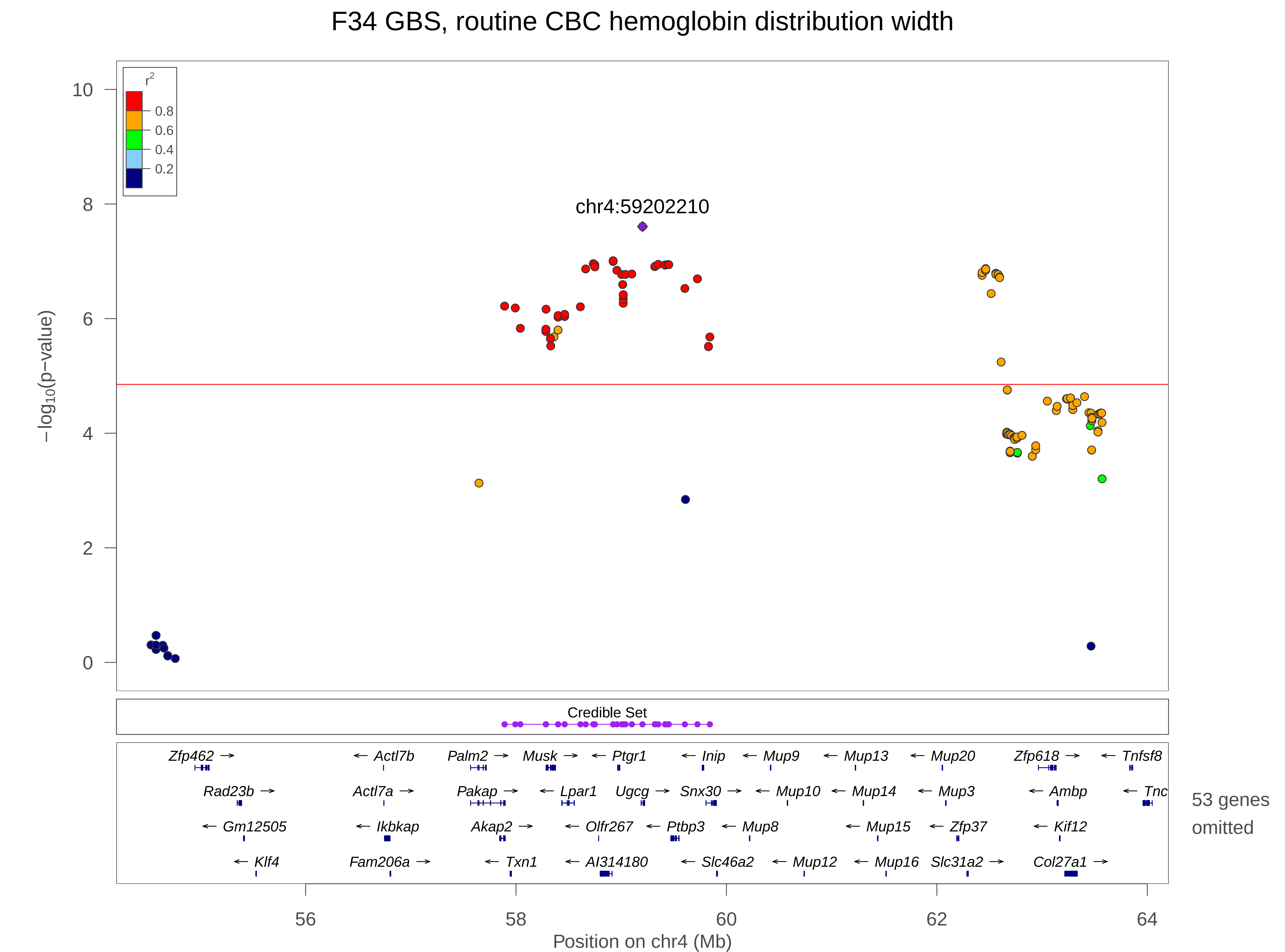
F34 GBS, routine CBC hemoglobin distribution width
r2
0.8
0.6
0.4
0.2
chr4:59202210
−log10(p−value)
Credible Set
Zfp462
Rad23b
Gm12505
Klf4
Actl7b
Actl7a
Ikbkap
Fam206a
Palm2
Pakap
Akap2
Txn1
Musk
Lpar1
Olfr267
AI314180
Ptgr1
Ugcg
Ptbp3
Inip
Snx30
Slc46a2
Mup9
Mup10
Mup8
Mup12
Mup13
Mup14
Mup15
Mup16
Mup20
Mup3
Zfp37
Slc31a2
Zfp618
Ambp
Kif12
Col27a1
Tnfsf8
Tnc
53 genes omitted
Position on chr4 (Mb)

| | |
|---|---|
| date: | Thu Oct 25 17:01:02 2018 |
| build: | F34GBS |
| display range: | chr4:54202210−64202210 [54202210−64202210] |
| hilite range: | 0 − 0 [ 0 − 0 ] |
| reference SNP: | chr4:59202210 |
| number of SNPs plotted: | 157 |
| min P−value: | 2.47E−8 [chr4:59202210] |
| max P−value: | 8.56E−1 [chr4:54761468] |
| omitted Genes: | Ctnnal1, Tmem245, Mir32 |
| omitted Genes: | Frrs1l, Epb41l4b, Gm12530 |
| omitted Genes: | Ptpn3, 1700042G15Rik, D630039A03Rik |
| omitted Genes: | Txndc8, Svep1, Mir3095 |
| omitted Genes: | Zkscan16, Dnajc25, Gng10 |
| omitted Genes: | Gm12596, Susd1, Hsdl2 |
| omitted Genes: | E130308A19Rik, Mup4, Mup6 |
| omitted Genes: | Mup7, Mup2, Mup1 |
| omitted Genes: | Mup11, Mup17, Mup18 |
| omitted Genes: | Mup19, Mup5, Mup21 |
| omitted Genes: | Fkbp15, Slc31a1, Cdc26 |
| omitted Genes: | Prpf4, Rnf183, Wdr31 |
| omitted Genes: | Bspry, Hdhd3, Alad |
| omitted Genes: | Pole3, 4933430I17Rik, Rgs3 |
| omitted Genes: | Mir455, Orm1, Orm3 |
| omitted Genes: | Orm2, Akna, Whrn |
| omitted Genes: | Atp6v1g1, 6330416G13Rik, Gm11213 |
| omitted Genes: | 1700018C11Rik, Tnfsf15 |

Make more plots at http://csg.sph.umich.edu/locuszoom/

Fine−mapping Regions

| snp | chr | pos (Mb) | pp | group | color |
|---|---|---|---|---|---|
| chr4:57891427 | 4 | 57.89143 | 0.007748913 | Credible Set | purple |
| chr4:57891451 | 4 | 57.89145 | 0.007748913 | Credible Set | purple |
| chr4:57993173 | 4 | 57.99317 | 0.007207393 | Credible Set | purple |
| chr4:58041091 | 4 | 58.04109 | 0.003285135 | Credible Set | purple |
| chr4:58283671 | 4 | 58.28367 | 0.002906190 | Credible Set | purple |
| chr4:58283721 | 4 | 58.28372 | 0.003173812 | Credible Set | purple |
| chr4:58285477 | 4 | 58.28548 | 0.006881396 | Credible Set | purple |
| chr4:58399947 | 4 | 58.39995 | 0.005058829 | Credible Set | purple |
| chr4:58400011 | 4 | 58.40001 | 0.005378042 | Credible Set | purple |
| chr4:58461854 | 4 | 58.46185 | 0.005181592 | Credible Set | purple |
| chr4:58462401 | 4 | 58.46240 | 0.005633158 | Credible Set | purple |
| chr4:58611965 | 4 | 58.61196 | 0.007548703 | Credible Set | purple |
| chr4:58662154 | 4 | 58.66215 | 0.032809373 | Credible Set | purple |
| chr4:58735042 | 4 | 58.73504 | 0.040294408 | Credible Set | purple |
| chr4:58748774 | 4 | 58.74877 | 0.038686690 | Credible Set | purple |
| chr4:58749245 | 4 | 58.74925 | 0.035507815 | Credible Set | purple |
| chr4:58923167 | 4 | 58.92317 | 0.043927718 | Credible Set | purple |
| chr4:58923190 | 4 | 58.92319 | 0.045160814 | Credible Set | purple |
| chr4:58958681 | 4 | 58.95868 | 0.031124327 | Credible Set | purple |
| chr4:59004314 | 4 | 59.00431 | 0.026373074 | Credible Set | purple |
| chr4:59013917 | 4 | 59.01392 | 0.017881488 | Credible Set | purple |
| chr4:59019003 | 4 | 59.01900 | 0.008656517 | Credible Set | purple |
| chr4:59019032 | 4 | 59.01903 | 0.010075153 | Credible Set | purple |
| chr4:59019072 | 4 | 59.01907 | 0.011759463 | Credible Set | purple |
| chr4:59019113 | 4 | 59.01911 | 0.012105110 | Credible Set | purple |
| chr4:59040591 | 4 | 59.04059 | 0.026499791 | Credible Set | purple |
| chr4:59100640 | 4 | 59.10064 | 0.026917255 | Credible Set | purple |
| chr4:59202210 | 4 | 59.20221 | 0.171013248 | Credible Set | purple |
| chr4:59319020 | 4 | 59.31902 | 0.035880347 | Credible Set | purple |
| chr4:59324851 | 4 | 59.32485 | 0.036725464 | Credible Set | purple |
| chr4:59352342 | 4 | 59.35234 | 0.039231485 | Credible Set | purple |
| chr4:59417447 | 4 | 59.41745 | 0.038287627 | Credible Set | purple |
| chr4:59417451 | 4 | 59.41745 | 0.038167661 | Credible Set | purple |
| chr4:59417505 | 4 | 59.41750 | 0.038047320 | Credible Set | purple |
| chr4:59444538 | 4 | 59.44454 | 0.038964344 | Credible Set | purple |
| chr4:59451945 | 4 | 59.45195 | 0.038747805 | Credible Set | purple |

Fine−mapping Regions

| snp | chr | pos (Mb) | pp | group | color |
|---|---|---|---|---|---|
| chr4:59604913 | 4 | 59.60491 | 0.015387839 | Credible Set | purple |
| chr4:59724135 | 4 | 59.72413 | 0.022336546 | Credible Set | purple |
| chr4:59842519 | 4 | 59.84252 | 0.002352616 | Credible Set | purple |

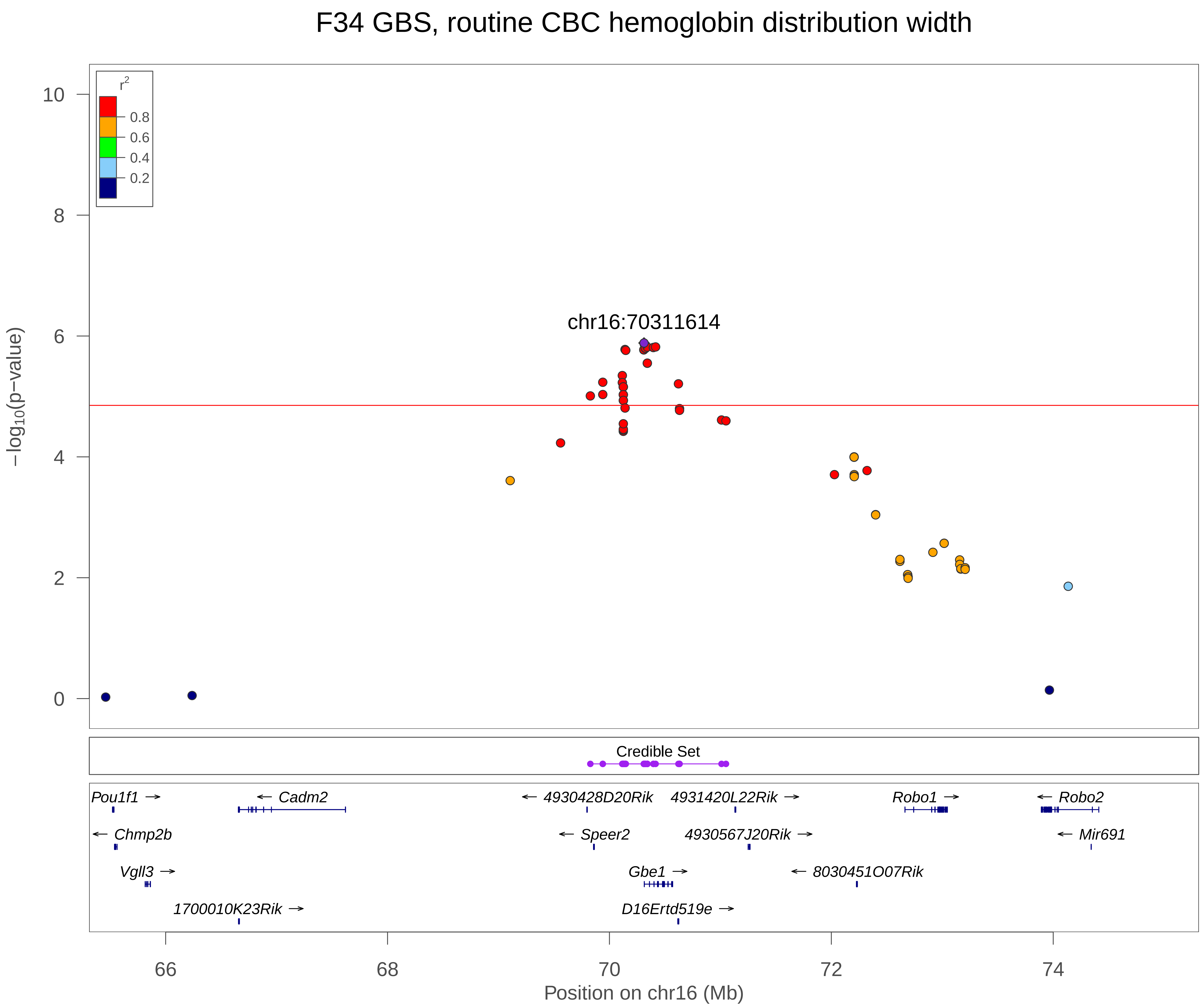
F34 GBS, routine CBC hemoglobin distribution width
r2
0.8
0.6
0.4
0.2
10
8
6
4
2
0
−log10(p−value)
chr16:70311614
Credible Set
Pou1f1 →
← Cadm2
← 4930428D20Rik
4931420L22Rik →
Robo1 →
← Robo2
← Chmp2b
← Speer2
4930567J20Rik →
← Mir691
Vgll3 →
Gbe1 →
← 8030451O07Rik
1700010K23Rik →
D16Ertd519e →
66
68
70
72
74
Position on chr16 (Mb)

| | |
|---|---|
| date: | Thu Oct 25 17:00:53 2018 |
| build: | F34GBS |
| display range: | chr16:65311614−75311614 [65311614−75311614] |
| hilite range: | 0 − 0 [ 0 − 0 ] |
| reference SNP: | chr16:70311614 |
| number of SNPs plotted: | 80 |
| min P−value: | 1.3E−6 [chr16:70311614] |
| max P−value: | 9.45E−1 [chr16:65459428] |

Make more plots at http://csg.sph.umich.edu/locuszoom/

Fine−mapping Regions

| snp | chr | pos (Mb) | pp | group | color |
|---|---|---|---|---|---|
| chr16:69827834 | 16 | 69.82783 | 0.008250572 | Credible Set | purple |
| chr16:69939841 | 16 | 69.93984 | 0.013575024 | Credible Set | purple |
| chr16:69939906 | 16 | 69.93991 | 0.008675613 | Credible Set | purple |
| chr16:70116129 | 16 | 70.11613 | 0.017336091 | Credible Set | purple |
| chr16:70116236 | 16 | 70.11624 | 0.013409284 | Credible Set | purple |
| chr16:70124826 | 16 | 70.12483 | 0.011452122 | Credible Set | purple |
| chr16:70124828 | 16 | 70.12483 | 0.011452122 | Credible Set | purple |
| chr16:70124838 | 16 | 70.12484 | 0.011452122 | Credible Set | purple |
| chr16:70124859 | 16 | 70.12486 | 0.011452122 | Credible Set | purple |
| chr16:70124867 | 16 | 70.12487 | 0.011452122 | Credible Set | purple |
| chr16:70124868 | 16 | 70.12487 | 0.011452122 | Credible Set | purple |
| chr16:70124887 | 16 | 70.12489 | 0.011452122 | Credible Set | purple |
| chr16:70124959 | 16 | 70.12496 | 0.008623924 | Credible Set | purple |
| chr16:70124962 | 16 | 70.12496 | 0.008623924 | Credible Set | purple |
| chr16:70124987 | 16 | 70.12499 | 0.008623924 | Credible Set | purple |
| chr16:70125117 | 16 | 70.12512 | 0.003001133 | Credible Set | purple |
| chr16:70125367 | 16 | 70.12537 | 0.006971344 | Credible Set | purple |
| chr16:70125371 | 16 | 70.12537 | 0.006996982 | Credible Set | purple |
| chr16:70125418 | 16 | 70.12542 | 0.007010792 | Credible Set | purple |
| chr16:70140593 | 16 | 70.14059 | 0.044975696 | Credible Set | purple |
| chr16:70140761 | 16 | 70.14076 | 0.005327267 | Credible Set | purple |
| chr16:70140804 | 16 | 70.14080 | 0.005308280 | Credible Set | purple |
| chr16:70146841 | 16 | 70.14684 | 0.043580539 | Credible Set | purple |
| chr16:70146890 | 16 | 70.14689 | 0.043580539 | Credible Set | purple |
| chr16:70309612 | 16 | 70.30961 | 0.044244068 | Credible Set | purple |
| chr16:70309921 | 16 | 70.30992 | 0.044612838 | Credible Set | purple |
| chr16:70311614 | 16 | 70.31161 | 0.057019078 | Credible Set | purple |
| chr16:70314976 | 16 | 70.31498 | 0.054154550 | Credible Set | purple |
| chr16:70323729 | 16 | 70.32373 | 0.050477142 | Credible Set | purple |
| chr16:70324316 | 16 | 70.32432 | 0.046671903 | Credible Set | purple |
| chr16:70324391 | 16 | 70.32439 | 0.047358827 | Credible Set | purple |
| chr16:70341194 | 16 | 70.34119 | 0.049141270 | Credible Set | purple |
| chr16:70341347 | 16 | 70.34135 | 0.027153292 | Credible Set | purple |
| chr16:70341354 | 16 | 70.34135 | 0.027284663 | Credible Set | purple |
| chr16:70341358 | 16 | 70.34136 | 0.027246633 | Credible Set | purple |
| chr16:70395502 | 16 | 70.39550 | 0.048288713 | Credible Set | purple |

Fine−mapping Regions

| snp | chr | pos (Mb) | pp | group | color |
|---|---|---|---|---|---|
| chr16:70396913 | 16 | 70.39691 | 0.048411322 | Credible Set | purple |
| chr16:70416028 | 16 | 70.41603 | 0.049357532 | Credible Set | purple |
| chr16:70621955 | 16 | 70.62195 | 0.012817652 | Credible Set | purple |
| chr16:70631684 | 16 | 70.63168 | 0.005194439 | Credible Set | purple |
| chr16:70631718 | 16 | 70.63172 | 0.005201000 | Credible Set | purple |
| chr16:70631722 | 16 | 70.63172 | 0.004885828 | Credible Set | purple |
| chr16:71010303 | 16 | 71.01030 | 0.003440319 | Credible Set | purple |
| chr16:71050225 | 16 | 71.05022 | 0.003338409 | Credible Set | purple |

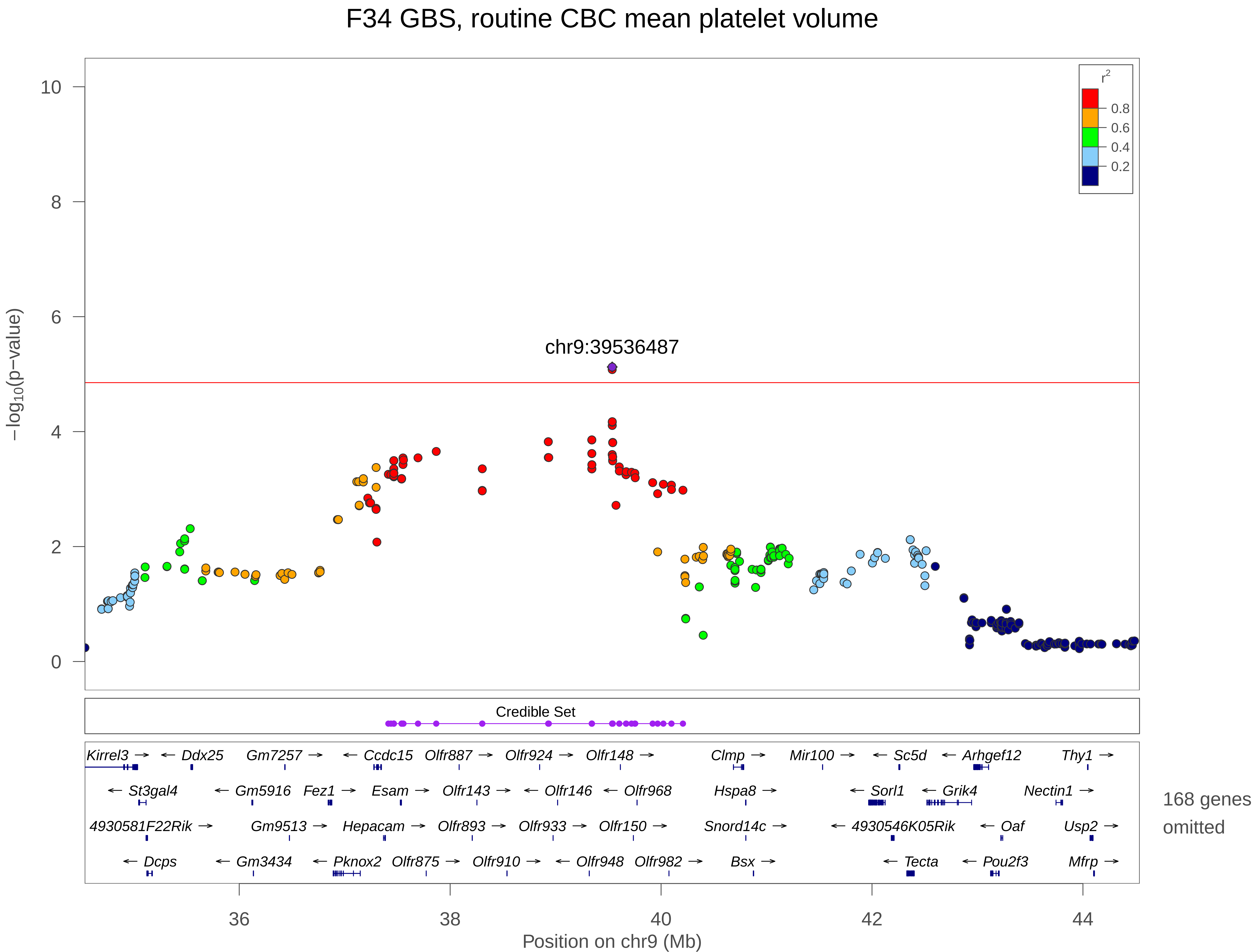
F34 GBS, routine CBC mean platelet volume
−log10(p−value)
chr9:39536487
r2
0.8
0.6
0.4
0.2
Credible Set
Kirrel3 →
← Ddx25
Gm7257 →
← Ccdc15
Olfr887 →
Olfr924 →
Olfr148 →
Clmp →
Mir100 →
← Sc5d
← Arhgef12
Thy1 →
← St3gal4
← Gm5916
Fez1 →
Esam →
Olfr143 →
← Olfr146
← Olfr968
Hspa8 →
← Sorl1
← Grik4
Nectin1 →
4930581F22Rik →
Gm9513 →
Hepacam →
Olfr893 →
Olfr933 →
Olfr150 →
Snord14c →
← 4930546K05Rik
← Oaf
Usp2 →
← Dcps
← Gm3434
← Pknox2
Olfr875 →
Olfr910 →
← Olfr948
Olfr982 →
Bsx →
← Tecta
← Pou2f3
Mfrp →
168 genes omitted
Position on chr9 (Mb)

date: Thu Oct 25 17:01:26 2018
build: F34GBS
display range: chr9:34536487−44536487 [34536487−44536487]
hilite range: 0 − 0 [ 0 − 0 ]
reference SNP: chr9:39536487
number of SNPs plotted: 396
min P−value: 7.51E−6 [chr9:39536487]
max P−value: 5.98E−1 [chr9:43965763]
omitted Genes: Tirap, Foxred1, Srpr
omitted Genes: Fam118b, Rpusd4, 4933422A05Rik
omitted Genes: Cdon, Pus3, Hyls1
omitted Genes: Pate4, Pate3, Pate2
omitted Genes: Pate1, Gm17677, Gm17727
omitted Genes: D730048I06Rik, 9230110F15Rik, Gm3428
omitted Genes: Gm5615, Gm17689, A630095E13Rik
omitted Genes: 1700027I24Rik, Acrv1, Chek1
omitted Genes: Stt3a, Ei24, Tmem218
omitted Genes: Slc37a2, Robo4, Robo3
omitted Genes: Msantd2, Vsig2, Nrgn
omitted Genes: Spa17, Siae, Tbrg1
omitted Genes: Panx3, Olfr160, Olfr151
omitted Genes: Olfr874, Olfr876, Olfr877
omitted Genes: Olfr145, Olfr878, Olfr881
omitted Genes: Olfr883, Olfr884, Olfr885
omitted Genes: Olfr888, Olfr889, Olfr890
omitted Genes: Olfr891, Olfr894, Olfr895
omitted Genes: Olfr25, Olfr898, Olfr250
omitted Genes: Olfr251, Olfr147, Olfr901
omitted Genes: Olfr902, Olfr904, Olfr905
omitted Genes: Olfr906, Olfr907, Olfr908
omitted Genes: Olfr911−ps1, Olfr912, Olfr913
omitted Genes: Olfr914, Olfr915, Olfr916
omitted Genes: Olfr917, Olfr918, Olfr919
omitted Genes: Vwa5a, Olfr920, Olfr921
omitted Genes: Olfr922, Olfr923, Olfr26
omitted Genes: Olfr926, Olfr930, Olfr934
omitted Genes: Olfr935, Olfr936, Olfr937
omitted Genes: Olfr938, Olfr27, Olfr943
omitted Genes: Olfr944, Olfr1537, Olfr945
Make more plots at http://csg.sph.umich.edu/locuszoom/
omitted Genes: Olfr947−ps1, Olfr951, Olfr952

Fine−mapping Regions

| snp | chr | pos (Mb) | pp | group | color |
|---|---|---|---|---|---|
| chr9:37414290 | 9 | 37.41429 | 0.003422972 | Credible Set | purple |
| chr9:37438921 | 9 | 37.43892 | 0.003366014 | Credible Set | purple |
| chr9:37465579 | 9 | 37.46558 | 0.004211718 | Credible Set | purple |
| chr9:37465802 | 9 | 37.46580 | 0.005689811 | Credible Set | purple |
| chr9:37465804 | 9 | 37.46580 | 0.005712100 | Credible Set | purple |
| chr9:37465813 | 9 | 37.46581 | 0.005720340 | Credible Set | purple |
| chr9:37465903 | 9 | 37.46590 | 0.003143168 | Credible Set | purple |
| chr9:37465905 | 9 | 37.46590 | 0.003119966 | Credible Set | purple |
| chr9:37465916 | 9 | 37.46592 | 0.003139806 | Credible Set | purple |
| chr9:37465963 | 9 | 37.46596 | 0.003217177 | Credible Set | purple |
| chr9:37466086 | 9 | 37.46609 | 0.003598142 | Credible Set | purple |
| chr9:37466137 | 9 | 37.46614 | 0.003550052 | Credible Set | purple |
| chr9:37539552 | 9 | 37.53955 | 0.002926010 | Credible Set | purple |
| chr9:37539568 | 9 | 37.53957 | 0.002869104 | Credible Set | purple |
| chr9:37539584 | 9 | 37.53958 | 0.002923584 | Credible Set | purple |
| chr9:37539604 | 9 | 37.53960 | 0.002902147 | Credible Set | purple |
| chr9:37552798 | 9 | 37.55280 | 0.004936077 | Credible Set | purple |
| chr9:37553112 | 9 | 37.55311 | 0.006324244 | Credible Set | purple |
| chr9:37557424 | 9 | 37.55742 | 0.005854382 | Credible Set | purple |
| chr9:37695097 | 9 | 37.69510 | 0.006348683 | Credible Set | purple |
| chr9:37868086 | 9 | 37.86809 | 0.008063101 | Credible Set | purple |
| chr9:38304315 | 9 | 38.30432 | 0.001881520 | Credible Set | purple |
| chr9:38304328 | 9 | 38.30433 | 0.001847651 | Credible Set | purple |
| chr9:38304643 | 9 | 38.30464 | 0.004207367 | Credible Set | purple |
| chr9:38930488 | 9 | 38.93049 | 0.011644150 | Credible Set | purple |
| chr9:38930537 | 9 | 38.93054 | 0.011611415 | Credible Set | purple |
| chr9:38930589 | 9 | 38.93059 | 0.006431069 | Credible Set | purple |
| chr9:38935078 | 9 | 38.93508 | 0.006400153 | Credible Set | purple |
| chr9:39343278 | 9 | 39.34328 | 0.012427759 | Credible Set | purple |
| chr9:39343281 | 9 | 39.34328 | 0.012427759 | Credible Set | purple |
| chr9:39343318 | 9 | 39.34332 | 0.012427759 | Credible Set | purple |
| chr9:39343350 | 9 | 39.34335 | 0.012486147 | Credible Set | purple |
| chr9:39343392 | 9 | 39.34339 | 0.007439149 | Credible Set | purple |
| chr9:39343432 | 9 | 39.34343 | 0.007463132 | Credible Set | purple |
| chr9:39343515 | 9 | 39.34351 | 0.004209055 | Credible Set | purple |
| chr9:39343529 | 9 | 39.34353 | 0.004194884 | Credible Set | purple |

Fine−mapping Regions

| snp | chr | pos (Mb) | pp | group | color |
|---|---|---|---|---|---|
| chr9:39343543 | 9 | 39.34354 | 0.004194884 | Credible Set | purple |
| chr9:39343573 | 9 | 39.34357 | 0.004194884 | Credible Set | purple |
| chr9:39343574 | 9 | 39.34357 | 0.004194884 | Credible Set | purple |
| chr9:39343835 | 9 | 39.34383 | 0.004907975 | Credible Set | purple |
| chr9:39343836 | 9 | 39.34384 | 0.004888817 | Credible Set | purple |
| chr9:39536486 | 9 | 39.53649 | 0.198249432 | Credible Set | purple |
| chr9:39536487 | 9 | 39.53649 | 0.199624197 | Credible Set | purple |
| chr9:39536502 | 9 | 39.53650 | 0.180698952 | Credible Set | purple |
| chr9:39536802 | 9 | 39.53680 | 0.007177750 | Credible Set | purple |
| chr9:39536806 | 9 | 39.53681 | 0.007167807 | Credible Set | purple |
| chr9:39537143 | 9 | 39.53714 | 0.021501081 | Credible Set | purple |
| chr9:39537177 | 9 | 39.53718 | 0.024301392 | Credible Set | purple |
| chr9:39537178 | 9 | 39.53718 | 0.024833881 | Credible Set | purple |
| chr9:39540755 | 9 | 39.54075 | 0.005834370 | Credible Set | purple |
| chr9:39540852 | 9 | 39.54085 | 0.005659179 | Credible Set | purple |
| chr9:39540971 | 9 | 39.54097 | 0.006569202 | Credible Set | purple |
| chr9:39540972 | 9 | 39.54097 | 0.006574866 | Credible Set | purple |
| chr9:39541231 | 9 | 39.54123 | 0.011349162 | Credible Set | purple |
| chr9:39541239 | 9 | 39.54124 | 0.011284875 | Credible Set | purple |
| chr9:39541258 | 9 | 39.54126 | 0.011218388 | Credible Set | purple |
| chr9:39603580 | 9 | 39.60358 | 0.004517231 | Credible Set | purple |
| chr9:39604144 | 9 | 39.60414 | 0.003867247 | Credible Set | purple |
| chr9:39667289 | 9 | 39.66729 | 0.003367749 | Credible Set | purple |
| chr9:39669535 | 9 | 39.66954 | 0.003774157 | Credible Set | purple |
| chr9:39719870 | 9 | 39.71987 | 0.003698203 | Credible Set | purple |
| chr9:39751082 | 9 | 39.75108 | 0.003555469 | Credible Set | purple |
| chr9:39754805 | 9 | 39.75480 | 0.003010232 | Credible Set | purple |
| chr9:39920209 | 9 | 39.92021 | 0.002512328 | Credible Set | purple |
| chr9:39967389 | 9 | 39.96739 | 0.001665887 | Credible Set | purple |
| chr9:40021684 | 9 | 40.02168 | 0.002365859 | Credible Set | purple |
| chr9:40098115 | 9 | 40.09811 | 0.002289875 | Credible Set | purple |
| chr9:40098581 | 9 | 40.09858 | 0.001942722 | Credible Set | purple |
| chr9:40207245 | 9 | 40.20725 | 0.001895759 | Credible Set | purple |

### F34 GBS, routine CBC mean platelet volume

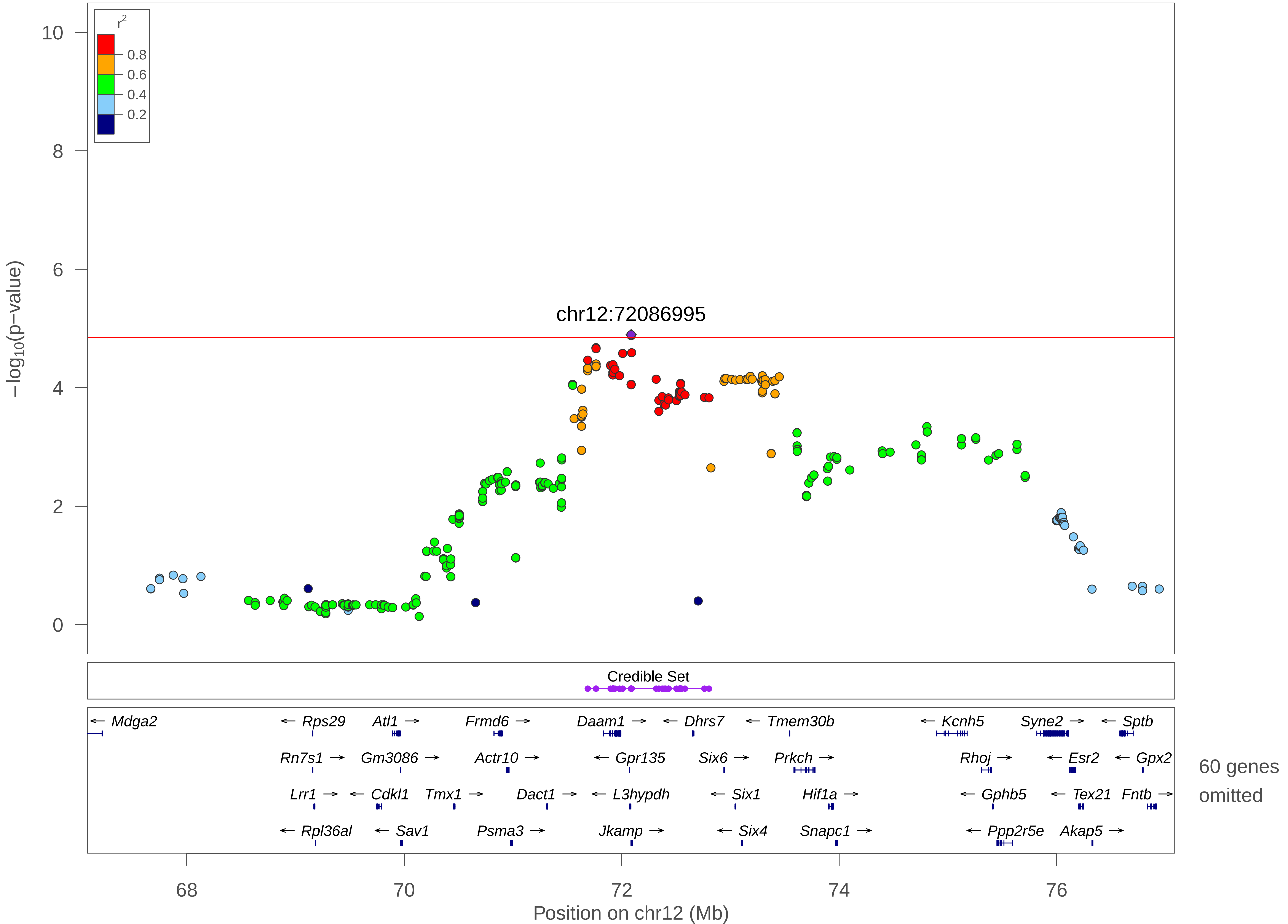

| | |
|---|---|
| date: | Thu Oct 25 17:01:13 2018 |
| build: | F34GBS |
| display range: | chr12:67086995−77086995 [67086995−77086995] |
| hilite range: | 0 − 0 [ 0 − 0 ] |
| reference SNP: | chr12:72086995 |
| number of SNPs plotted: | 340 |
| min P−value: | 1.27E−5 [chr12:72086995] |
| max P−value: | 7.27E−1 [chr12:70137276] |
| omitted Genes: | Mgat2, Dnaaf2, 9330151L19Rik |
| omitted Genes: | Pole2, Klhdc1, Klhdc2 |
| omitted Genes: | Nemf, Rn7s2, Arf6 |
| omitted Genes: | 5830428M24Rik, Vcpkmt, Sos2 |
| omitted Genes: | L2hgdh, Atp5s, Mir681 |
| omitted Genes: | 4930512B01Rik, Map4k5, 4931403G20Rik |
| omitted Genes: | Nin, Abhd12b, Pygl |
| omitted Genes: | F730035M05Rik, Trim9, 3110056K07Rik |
| omitted Genes: | Arid4a, Tomm20l, Timm9 |
| omitted Genes: | 2700049A03Rik, 1700083H02Rik, 4930404H11Rik |
| omitted Genes: | Ccdc175, Rtn1, Lrrc9 |
| omitted Genes: | Pcnx4, Ppm1a, 4930447C04Rik |
| omitted Genes: | Mnat1, Trmt5, Slc38a6 |
| omitted Genes: | D830013O20Rik, 2210039B01Rik, Syt16 |
| omitted Genes: | 1700086L19Rik, Dbpht2, Wdr89 |
| omitted Genes: | Rplp2−ps1, Sgpp1, Mir5101 |
| omitted Genes: | Mthfd1, Zbtb25, Zbtb1 |
| omitted Genes: | Hspa2, Ppp1r36, Gm10451 |
| omitted Genes: | Mir5135, Plekhg3, AI463170 |
| omitted Genes: | Churc1, Rab15, Max |

Make more plots at http://csg.sph.umich.edu/locuszoom/

Fine−mapping Regions

| snp | chr | pos (Mb) | pp | group | color |
|---|---|---|---|---|---|
| chr12:71688023 | 12 | 71.68802 | 0.035804279 | Credible Set | purple |
| chr12:71763688 | 12 | 71.76369 | 0.056816257 | Credible Set | purple |
| chr12:71764110 | 12 | 71.76411 | 0.054725343 | Credible Set | purple |
| chr12:71898279 | 12 | 71.89828 | 0.029427655 | Credible Set | purple |
| chr12:71918314 | 12 | 71.91831 | 0.020766950 | Credible Set | purple |
| chr12:71919408 | 12 | 71.91941 | 0.022845369 | Credible Set | purple |
| chr12:71919477 | 12 | 71.91948 | 0.030259950 | Credible Set | purple |
| chr12:71919515 | 12 | 71.91952 | 0.030291080 | Credible Set | purple |
| chr12:71919520 | 12 | 71.91952 | 0.030291080 | Credible Set | purple |
| chr12:71923446 | 12 | 71.92345 | 0.022812867 | Credible Set | purple |
| chr12:71937615 | 12 | 71.93761 | 0.025669447 | Credible Set | purple |
| chr12:71980528 | 12 | 71.98053 | 0.020187761 | Credible Set | purple |
| chr12:72009199 | 12 | 72.00920 | 0.045918376 | Credible Set | purple |
| chr12:72086662 | 12 | 72.08666 | 0.014723900 | Credible Set | purple |
| chr12:72086706 | 12 | 72.08671 | 0.014634966 | Credible Set | purple |
| chr12:72086724 | 12 | 72.08672 | 0.014459619 | Credible Set | purple |
| chr12:72086792 | 12 | 72.08679 | 0.088935759 | Credible Set | purple |
| chr12:72086995 | 12 | 72.08700 | 0.091752314 | Credible Set | purple |
| chr12:72087027 | 12 | 72.08703 | 0.091238103 | Credible Set | purple |
| chr12:72091767 | 12 | 72.09177 | 0.047107639 | Credible Set | purple |
| chr12:72316934 | 12 | 72.31693 | 0.017748789 | Credible Set | purple |
| chr12:72342228 | 12 | 72.34223 | 0.008174200 | Credible Set | purple |
| chr12:72372220 | 12 | 72.37222 | 0.009361947 | Credible Set | purple |
| chr12:72389901 | 12 | 72.38990 | 0.007115213 | Credible Set | purple |
| chr12:72404532 | 12 | 72.40453 | 0.006890577 | Credible Set | purple |
| chr12:72430018 | 12 | 72.43002 | 0.008940829 | Credible Set | purple |
| chr12:72432423 | 12 | 72.43242 | 0.008317797 | Credible Set | purple |
| chr12:72503712 | 12 | 72.50371 | 0.008093364 | Credible Set | purple |
| chr12:72530507 | 12 | 72.53051 | 0.011173208 | Credible Set | purple |
| chr12:72531030 | 12 | 72.53103 | 0.009498292 | Credible Set | purple |
| chr12:72531036 | 12 | 72.53104 | 0.009612957 | Credible Set | purple |
| chr12:72534336 | 12 | 72.53434 | 0.010714374 | Credible Set | purple |
| chr12:72542646 | 12 | 72.54265 | 0.011189095 | Credible Set | purple |
| chr12:72542706 | 12 | 72.54271 | 0.009769147 | Credible Set | purple |
| chr12:72543184 | 12 | 72.54318 | 0.015260124 | Credible Set | purple |
| chr12:72543205 | 12 | 72.54321 | 0.014846908 | Credible Set | purple |

Fine−mapping Regions

| snp | chr | pos (Mb) | pp | group | color |
|---|---|---|---|---|---|
| chr12:72551877 | 12 | 72.55188 | 0.011070755 | Credible Set | purple |
| chr12:72582082 | 12 | 72.58208 | 0.009998311 | Credible Set | purple |
| chr12:72761155 | 12 | 72.76116 | 0.009115619 | Credible Set | purple |
| chr12:72803219 | 12 | 72.80322 | 0.008967203 | Credible Set | purple |

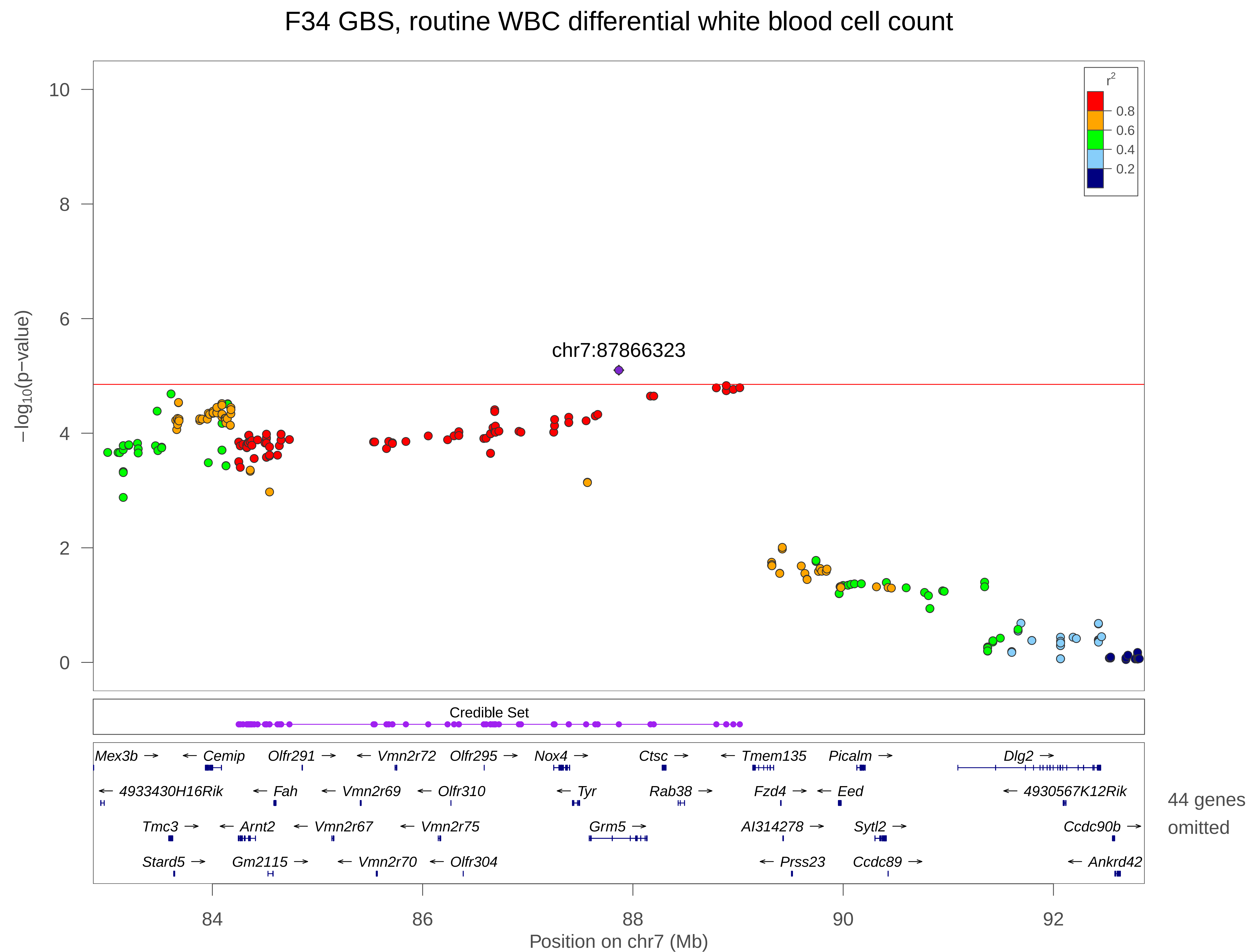
F34 GBS, routine WBC differential white blood cell count
chr7:87866323
r2
0.8
0.6
0.4
0.2
−log10(p−value)
Credible Set
Mex3b
Cemip
Olfr291
Vmn2r72
Olfr295
Nox4
Ctsc
Tmem135
Picalm
Dlg2
4933430H16Rik
Fah
Vmn2r69
Olfr310
Tyr
Rab38
Fzd4
Eed
4930567K12Rik
Tmc3
Arnt2
Vmn2r67
Vmn2r75
Grm5
AI314278
Sytl2
Ccdc90b
Stard5
Gm2115
Vmn2r70
Olfr304
Prss23
Ccdc89
Ankrd42
44 genes omitted
Position on chr7 (Mb)

| | |
|---|---|
| date: | Thu Oct 25 17:02:03 2018 |
| build: | F34GBS |
| display range: | chr7:82866323−92866323 [82866323−92866323] |
| hilite range: | 0 − 0 [ 0 − 0 ] |
| reference SNP: | chr7:87866323 |
| number of SNPs plotted: | 272 |
| min P−value: | 7.94E−6 [chr7:87866323] |
| max P−value: | 8.93E−1 [chr7:92689613] |
| omitted Genes: | Il16, Cfap161, Mesdc1 |
| omitted Genes: | Mesdc2, Abhd17c, Zfand6 |
| omitted Genes: | 2610206C17Rik, Olfr290, Vmn2r65 |
| omitted Genes: | Vmn2r66, Vmn2r68, Vmn2r71 |
| omitted Genes: | Vmn2r73, Vmn2r74, Vmn2r76 |
| omitted Genes: | Olfr309, Olfr308, Olfr307 |
| omitted Genes: | Olfr305, Olfr303, Olfr301 |
| omitted Genes: | Olfr299, Olfr298, Olfr297 |
| omitted Genes: | Olfr294, Olfr293, Olfr292 |
| omitted Genes: | Folh1, Vmn2r77, Vmn2r78 |
| omitted Genes: | Vmn2r79, Me3, Ccdc81 |
| omitted Genes: | l7Rn6, E230029C05Rik, 2310010J17Rik |
| omitted Genes: | Ccdc83, Crebzf, Tmem126a |
| omitted Genes: | Tmem126b, Pcf11, 4632427E13Rik |
| omitted Genes: | Rab30, Ddias |

Make more plots at http://csg.sph.umich.edu/locuszoom/

Fine−mapping Regions

| snp | chr | pos (Mb) | pp | group | color |
|---|---|---|---|---|---|
| chr7:84250985 | 7 | 84.25099 | 0.005842461 | Credible Set | purple |
| chr7:84251011 | 7 | 84.25101 | 0.005822474 | Credible Set | purple |
| chr7:84265685 | 7 | 84.26569 | 0.005053687 | Credible Set | purple |
| chr7:84291540 | 7 | 84.29154 | 0.005344684 | Credible Set | purple |
| chr7:84327288 | 7 | 84.32729 | 0.005250490 | Credible Set | purple |
| chr7:84327808 | 7 | 84.32781 | 0.004707908 | Credible Set | purple |
| chr7:84327850 | 7 | 84.32785 | 0.004781219 | Credible Set | purple |
| chr7:84338595 | 7 | 84.33859 | 0.005694824 | Credible Set | purple |
| chr7:84343118 | 7 | 84.34312 | 0.005632570 | Credible Set | purple |
| chr7:84344805 | 7 | 84.34480 | 0.007499805 | Credible Set | purple |
| chr7:84347423 | 7 | 84.34742 | 0.007578390 | Credible Set | purple |
| chr7:84360038 | 7 | 84.36004 | 0.005936887 | Credible Set | purple |
| chr7:84360626 | 7 | 84.36063 | 0.005872583 | Credible Set | purple |
| chr7:84360642 | 7 | 84.36064 | 0.005872583 | Credible Set | purple |
| chr7:84360703 | 7 | 84.36070 | 0.005872583 | Credible Set | purple |
| chr7:84360714 | 7 | 84.36071 | 0.005872583 | Credible Set | purple |
| chr7:84360716 | 7 | 84.36072 | 0.005872583 | Credible Set | purple |
| chr7:84375526 | 7 | 84.37553 | 0.006162539 | Credible Set | purple |
| chr7:84375577 | 7 | 84.37558 | 0.005161287 | Credible Set | purple |
| chr7:84375642 | 7 | 84.37564 | 0.005161287 | Credible Set | purple |
| chr7:84397575 | 7 | 84.39758 | 0.003135446 | Credible Set | purple |
| chr7:84397597 | 7 | 84.39760 | 0.003135446 | Credible Set | purple |
| chr7:84430102 | 7 | 84.43010 | 0.006337133 | Credible Set | purple |
| chr7:84501082 | 7 | 84.50108 | 0.005628534 | Credible Set | purple |
| chr7:84514082 | 7 | 84.51408 | 0.006610877 | Credible Set | purple |
| chr7:84514444 | 7 | 84.51444 | 0.006903711 | Credible Set | purple |
| chr7:84514505 | 7 | 84.51450 | 0.007875050 | Credible Set | purple |
| chr7:84514564 | 7 | 84.51456 | 0.003276428 | Credible Set | purple |
| chr7:84514575 | 7 | 84.51457 | 0.003296762 | Credible Set | purple |
| chr7:84514734 | 7 | 84.51473 | 0.005598543 | Credible Set | purple |
| chr7:84543346 | 7 | 84.54335 | 0.003440904 | Credible Set | purple |
| chr7:84543355 | 7 | 84.54336 | 0.003574188 | Credible Set | purple |
| chr7:84543782 | 7 | 84.54378 | 0.004863619 | Credible Set | purple |
| chr7:84619632 | 7 | 84.61963 | 0.003553096 | Credible Set | purple |
| chr7:84636759 | 7 | 84.63676 | 0.005073750 | Credible Set | purple |
| chr7:84653095 | 7 | 84.65309 | 0.006302011 | Credible Set | purple |

Fine−mapping Regions

| snp | chr | pos (Mb) | pp | group | color |
|---|---|---|---|---|---|
| chr7:84653660 | 7 | 84.65366 | 0.007889859 | Credible Set | purple |
| chr7:84653702 | 7 | 84.65370 | 0.007805421 | Credible Set | purple |
| chr7:84653707 | 7 | 84.65371 | 0.007842187 | Credible Set | purple |
| chr7:84733186 | 7 | 84.73319 | 0.006423158 | Credible Set | purple |
| chr7:85533207 | 7 | 85.53321 | 0.005864151 | Credible Set | purple |
| chr7:85543843 | 7 | 85.54384 | 0.005864696 | Credible Set | purple |
| chr7:85656243 | 7 | 85.65624 | 0.004561726 | Credible Set | purple |
| chr7:85677494 | 7 | 85.67749 | 0.005984206 | Credible Set | purple |
| chr7:85712663 | 7 | 85.71266 | 0.005681281 | Credible Set | purple |
| chr7:85712975 | 7 | 85.71298 | 0.005519092 | Credible Set | purple |
| chr7:85840665 | 7 | 85.84067 | 0.005963647 | Credible Set | purple |
| chr7:86053386 | 7 | 86.05339 | 0.007364597 | Credible Set | purple |
| chr7:86237374 | 7 | 86.23737 | 0.006370283 | Credible Set | purple |
| chr7:86298979 | 7 | 86.29898 | 0.007388638 | Credible Set | purple |
| chr7:86344420 | 7 | 86.34442 | 0.008471295 | Credible Set | purple |
| chr7:86344437 | 7 | 86.34444 | 0.008652510 | Credible Set | purple |
| chr7:86344488 | 7 | 86.34449 | 0.007467535 | Credible Set | purple |
| chr7:86582139 | 7 | 86.58214 | 0.006677627 | Credible Set | purple |
| chr7:86603770 | 7 | 86.60377 | 0.006743165 | Credible Set | purple |
| chr7:86645719 | 7 | 86.64572 | 0.003796364 | Credible Set | purple |
| chr7:86645792 | 7 | 86.64579 | 0.003832019 | Credible Set | purple |
| chr7:86645947 | 7 | 86.64595 | 0.007970007 | Credible Set | purple |
| chr7:86645990 | 7 | 86.64599 | 0.008014571 | Credible Set | purple |
| chr7:86668998 | 7 | 86.66900 | 0.010065420 | Credible Set | purple |
| chr7:86685426 | 7 | 86.68543 | 0.019941447 | Credible Set | purple |
| chr7:86685507 | 7 | 86.68551 | 0.018570266 | Credible Set | purple |
| chr7:86685648 | 7 | 86.68565 | 0.008771573 | Credible Set | purple |
| chr7:86692214 | 7 | 86.69221 | 0.010704621 | Credible Set | purple |
| chr7:86692491 | 7 | 86.69249 | 0.008450428 | Credible Set | purple |
| chr7:86725518 | 7 | 86.72552 | 0.008818564 | Credible Set | purple |
| chr7:86916161 | 7 | 86.91616 | 0.008757632 | Credible Set | purple |
| chr7:86934710 | 7 | 86.93471 | 0.008484704 | Credible Set | purple |
| chr7:87246786 | 7 | 87.24679 | 0.008476837 | Credible Set | purple |
| chr7:87254690 | 7 | 87.25469 | 0.010871267 | Credible Set | purple |
| chr7:87255156 | 7 | 87.25516 | 0.013773680 | Credible Set | purple |
| chr7:87389952 | 7 | 87.38995 | 0.014965327 | Credible Set | purple |

Fine−mapping Regions

| snp | chr | pos (Mb) | pp | group | color |
|---|---|---|---|---|---|
| chr7:87389989 | 7 | 87.38999 | 0.01219040 | Credible Set | purple |
| chr7:87390014 | 7 | 87.39001 | 0.01226061 | Credible Set | purple |
| chr7:87554961 | 7 | 87.55496 | 0.01308847 | Credible Set | purple |
| chr7:87642045 | 7 | 87.64204 | 0.01571216 | Credible Set | purple |
| chr7:87665727 | 7 | 87.66573 | 0.01666577 | Credible Set | purple |
| chr7:87866323 | 7 | 87.86632 | 0.09063412 | Credible Set | purple |
| chr7:88167216 | 7 | 88.16722 | 0.03354206 | Credible Set | purple |
| chr7:88198240 | 7 | 88.19824 | 0.03355873 | Credible Set | purple |
| chr7:88793653 | 7 | 88.79365 | 0.04597496 | Credible Set | purple |
| chr7:88888100 | 7 | 88.88810 | 0.04127148 | Credible Set | purple |
| chr7:88888124 | 7 | 88.88812 | 0.04127148 | Credible Set | purple |
| chr7:88888200 | 7 | 88.88820 | 0.05013049 | Credible Set | purple |
| chr7:88955188 | 7 | 88.95519 | 0.04334955 | Credible Set | purple |
| chr7:89016577 | 7 | 89.01658 | 0.04618414 | Credible Set | purple |

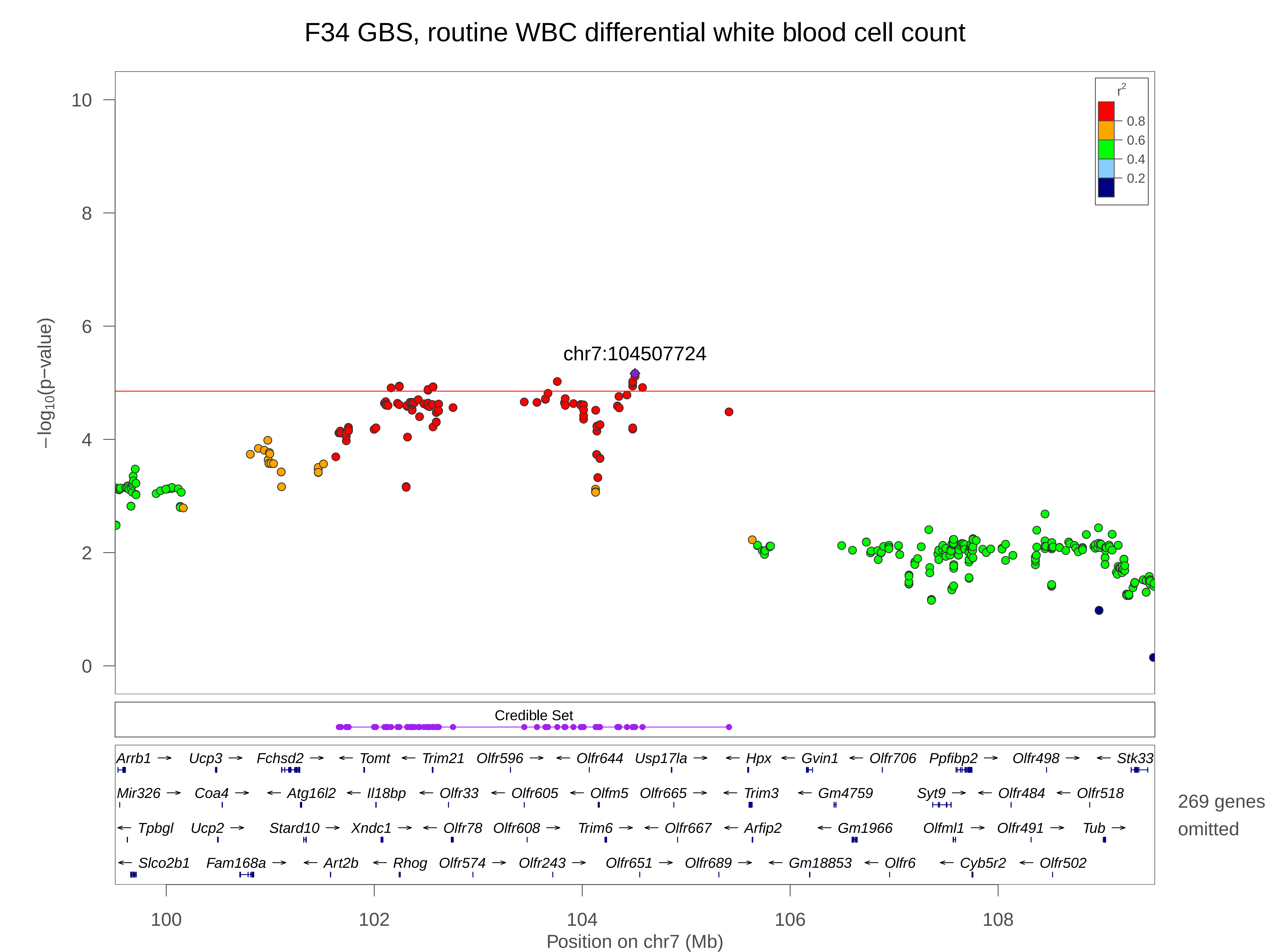

F34 GBS, routine WBC differential white blood cell count
r2
0.8
0.6
0.4
0.2
chr7:104507724
−log10(p−value)
10
8
6
4
2
0
Credible Set
Arrb1 → Ucp3 → Fchsd2 → ← Tomt ← Trim21 Olfr596 → ← Olfr644 Usp17la → ← Hpx ← Gvin1 ← Olfr706 Ppfibp2 → Olfr498 → ← Stk33
Mir326 → Coa4 → ← Atg16l2 ← Il18bp ← Olfr33 ← Olfr605 ← Olfm5 Olfr665 → ← Trim3 ← Gm4759 Syt9 → ← Olfr484 ← Olfr518
← Tpbgl Ucp2 → Stard10 → Xndc1 → ← Olfr78 Olfr608 → Trim6 → ← Olfr667 ← Arfip2 ← Gm1966 Olfml1 → Olfr491 → Tub →
← Slco2b1 Fam168a → ← Art2b ← Rhog Olfr574 → Olfr243 → Olfr651 → Olfr689 → ← Gm18853 ← Olfr6 ← Cyb5r2 ← Olfr502
269 genes omitted
100
102
104
106
108
Position on chr7 (Mb)

date: Thu Oct 25 17:01:51 2018
build: F34GBS
display range: chr7:99507724−109507724 [99507724−109507724]
hilite range: 0 − 0 [ 0 − 0 ]
reference SNP: chr7:104507724
number of SNPs plotted: 453
min P−value: 6.84E−6 [chr7:104507724]
max P−value: 7.11E−1 [chr7:109493860]
omitted Genes: Olfr520, Olfr521, Neu3
omitted Genes: Spcs2, Xrra1, Rnf169
omitted Genes: Chrdl2, Pold3, Lipt2
omitted Genes: Kcne3, Pgm2l1, Gpx2−ps1
omitted Genes: P4ha3, Ppme1, C2cd3
omitted Genes: Dnajb13, Mrpl48, Rab6a
omitted Genes: Plekhb1, Relt, Arhgef17
omitted Genes: Mir3102, P2ry6, P2ry2
omitted Genes: Arap1, Pde2a, Mir139
omitted Genes: Art2a−ps, Clpb, Phox2a
omitted Genes: Inppl1, Folr2, Folr1
omitted Genes: Anapc15, Lamtor1, Lrrc51
omitted Genes: Numa1, Rnf121, Xntrpc
omitted Genes: Trpc2, Art5, Art1
omitted Genes: Chrna10, Nup98, Pgap2
omitted Genes: Stim1, Rrm1, Olfr543
omitted Genes: Olfr544, Olfr545, Olfr547
omitted Genes: Olfr549, Olfr550, Olfr551
omitted Genes: Olfr552, Olfr553, Olfr554
omitted Genes: Olfr555, Olfr556, Trim68
omitted Genes: Olfr557, Olfr558, Olfr559
omitted Genes: Olfr560, Olfr561, Olfr564
omitted Genes: Olfr566, Olfr568, Olfr569
omitted Genes: Olfr570, Olfr571, Olfr572
omitted Genes: Olfr575, Olfr576, Olfr577
omitted Genes: Olfr578, Olfr582, Olfr583
omitted Genes: Olfr584, Olfr585, Olfr586
omitted Genes: Olfr589, Olfr591, Olfr592
omitted Genes: Olfr593, Olfr594, Usp17ld
omitted Genes: Olfr597, Olfr598, Olfr599
omitted Genes: Olfr600, Olfr601, Olfr603
omitted Genes: Usp17le, Olfr606, Olfr609

Make more plots at http://csg.sph.umich.edu/locuszoom/

Fine−mapping Regions

| snp | chr | pos (Mb) | pp | group | color |
|---|---|---|---|---|---|
| chr7:101660383 | 7 | 101.6604 | 0.002947822 | Credible Set | purple |
| chr7:101669186 | 7 | 101.6692 | 0.002986122 | Credible Set | purple |
| chr7:101673483 | 7 | 101.6735 | 0.003165845 | Credible Set | purple |
| chr7:101680140 | 7 | 101.6801 | 0.002925323 | Credible Set | purple |
| chr7:101731061 | 7 | 101.7311 | 0.002990668 | Credible Set | purple |
| chr7:101731110 | 7 | 101.7311 | 0.002174403 | Credible Set | purple |
| chr7:101731130 | 7 | 101.7311 | 0.002611160 | Credible Set | purple |
| chr7:101731658 | 7 | 101.7317 | 0.002882023 | Credible Set | purple |
| chr7:101751790 | 7 | 101.7518 | 0.003659142 | Credible Set | purple |
| chr7:101752580 | 7 | 101.7526 | 0.003433699 | Credible Set | purple |
| chr7:101752971 | 7 | 101.7530 | 0.003192238 | Credible Set | purple |
| chr7:101998679 | 7 | 101.9987 | 0.003380757 | Credible Set | purple |
| chr7:102016307 | 7 | 102.0163 | 0.003587087 | Credible Set | purple |
| chr7:102098054 | 7 | 102.0981 | 0.009274583 | Credible Set | purple |
| chr7:102110109 | 7 | 102.1101 | 0.009854129 | Credible Set | purple |
| chr7:102110501 | 7 | 102.1105 | 0.008638142 | Credible Set | purple |
| chr7:102116318 | 7 | 102.1163 | 0.008859280 | Credible Set | purple |
| chr7:102116430 | 7 | 102.1164 | 0.008859280 | Credible Set | purple |
| chr7:102116478 | 7 | 102.1165 | 0.008555806 | Credible Set | purple |
| chr7:102132562 | 7 | 102.1326 | 0.008467642 | Credible Set | purple |
| chr7:102162058 | 7 | 102.1621 | 0.016793280 | Credible Set | purple |
| chr7:102223396 | 7 | 102.2234 | 0.009244404 | Credible Set | purple |
| chr7:102240695 | 7 | 102.2407 | 0.008762541 | Credible Set | purple |
| chr7:102241191 | 7 | 102.2412 | 0.018032301 | Credible Set | purple |
| chr7:102241211 | 7 | 102.2412 | 0.017605212 | Credible Set | purple |
| chr7:102241212 | 7 | 102.2412 | 0.017636586 | Credible Set | purple |
| chr7:102317213 | 7 | 102.3172 | 0.008197054 | Credible Set | purple |
| chr7:102317246 | 7 | 102.3172 | 0.008540130 | Credible Set | purple |
| chr7:102319586 | 7 | 102.3196 | 0.002515012 | Credible Set | purple |
| chr7:102344970 | 7 | 102.3450 | 0.009606139 | Credible Set | purple |
| chr7:102363897 | 7 | 102.3639 | 0.009620264 | Credible Set | purple |
| chr7:102364204 | 7 | 102.3642 | 0.007336519 | Credible Set | purple |
| chr7:102364223 | 7 | 102.3642 | 0.007054151 | Credible Set | purple |
| chr7:102364315 | 7 | 102.3643 | 0.008821599 | Credible Set | purple |
| chr7:102364326 | 7 | 102.3643 | 0.008821599 | Credible Set | purple |
| chr7:102364411 | 7 | 102.3644 | 0.009415387 | Credible Set | purple |

Fine−mapping Regions

| snp | chr | pos (Mb) | pp | group | color |
|---|---|---|---|---|---|
| chr7:102385520 | 7 | 102.3855 | 0.009422644 | Credible Set | purple |
| chr7:102385536 | 7 | 102.3855 | 0.009588643 | Credible Set | purple |
| chr7:102423165 | 7 | 102.4232 | 0.010639022 | Credible Set | purple |
| chr7:102435692 | 7 | 102.4357 | 0.005507396 | Credible Set | purple |
| chr7:102435720 | 7 | 102.4357 | 0.005507396 | Credible Set | purple |
| chr7:102478469 | 7 | 102.4785 | 0.009058694 | Credible Set | purple |
| chr7:102510475 | 7 | 102.5105 | 0.008575601 | Credible Set | purple |
| chr7:102510607 | 7 | 102.5106 | 0.008430344 | Credible Set | purple |
| chr7:102516878 | 7 | 102.5169 | 0.009321505 | Credible Set | purple |
| chr7:102517238 | 7 | 102.5172 | 0.015357203 | Credible Set | purple |
| chr7:102517241 | 7 | 102.5172 | 0.015341995 | Credible Set | purple |
| chr7:102517247 | 7 | 102.5172 | 0.015863699 | Credible Set | purple |
| chr7:102531652 | 7 | 102.5317 | 0.008086830 | Credible Set | purple |
| chr7:102561708 | 7 | 102.5617 | 0.008830154 | Credible Set | purple |
| chr7:102565262 | 7 | 102.5653 | 0.017490592 | Credible Set | purple |
| chr7:102565292 | 7 | 102.5653 | 0.017347028 | Credible Set | purple |
| chr7:102565307 | 7 | 102.5653 | 0.017521476 | Credible Set | purple |
| chr7:102565323 | 7 | 102.5653 | 0.017377646 | Credible Set | purple |
| chr7:102565329 | 7 | 102.5653 | 0.017377646 | Credible Set | purple |
| chr7:102565410 | 7 | 102.5654 | 0.003709602 | Credible Set | purple |
| chr7:102596442 | 7 | 102.5964 | 0.006464016 | Credible Set | purple |
| chr7:102596578 | 7 | 102.5966 | 0.004511411 | Credible Set | purple |
| chr7:102596636 | 7 | 102.5966 | 0.004443758 | Credible Set | purple |
| chr7:102619418 | 7 | 102.6194 | 0.008982864 | Credible Set | purple |
| chr7:102619431 | 7 | 102.6194 | 0.008957417 | Credible Set | purple |
| chr7:102619466 | 7 | 102.6195 | 0.006884649 | Credible Set | purple |
| chr7:102757499 | 7 | 102.7575 | 0.007829380 | Credible Set | purple |
| chr7:103442570 | 7 | 103.4426 | 0.009741931 | Credible Set | purple |
| chr7:103564836 | 7 | 103.5648 | 0.009571513 | Credible Set | purple |
| chr7:103646439 | 7 | 103.6464 | 0.010767138 | Credible Set | purple |
| chr7:103646709 | 7 | 103.6467 | 0.011039266 | Credible Set | purple |
| chr7:103671033 | 7 | 103.6710 | 0.013611164 | Credible Set | purple |
| chr7:103760301 | 7 | 103.7603 | 0.021492054 | Credible Set | purple |
| chr7:103829200 | 7 | 103.8292 | 0.009423901 | Credible Set | purple |
| chr7:103837099 | 7 | 103.8371 | 0.011088190 | Credible Set | purple |
| chr7:103837213 | 7 | 103.8372 | 0.008502119 | Credible Set | purple |

Fine−mapping Regions

| snp | chr | pos (Mb) | pp | group | color |
|---|---|---|---|---|---|
| chr7:103915503 | 7 | 103.9155 | 0.009147836 | Credible Set | purple |
| chr7:103986792 | 7 | 103.9868 | 0.008826500 | Credible Set | purple |
| chr7:103986795 | 7 | 103.9868 | 0.008826500 | Credible Set | purple |
| chr7:103986857 | 7 | 103.9869 | 0.008548117 | Credible Set | purple |
| chr7:103986860 | 7 | 103.9869 | 0.008548117 | Credible Set | purple |
| chr7:104012808 | 7 | 104.0128 | 0.008714889 | Credible Set | purple |
| chr7:104013370 | 7 | 104.0134 | 0.004993329 | Credible Set | purple |
| chr7:104013395 | 7 | 104.0134 | 0.005737745 | Credible Set | purple |
| chr7:104013467 | 7 | 104.0135 | 0.007168033 | Credible Set | purple |
| chr7:104129734 | 7 | 104.1297 | 0.007066361 | Credible Set | purple |
| chr7:104129736 | 7 | 104.1297 | 0.007066361 | Credible Set | purple |
| chr7:104138936 | 7 | 104.1389 | 0.001289798 | Credible Set | purple |
| chr7:104138968 | 7 | 104.1390 | 0.001289798 | Credible Set | purple |
| chr7:104141343 | 7 | 104.1413 | 0.003154168 | Credible Set | purple |
| chr7:104141705 | 7 | 104.1417 | 0.003837844 | Credible Set | purple |
| chr7:104141768 | 7 | 104.1418 | 0.003756126 | Credible Set | purple |
| chr7:104170588 | 7 | 104.1706 | 0.004039617 | Credible Set | purple |
| chr7:104170594 | 7 | 104.1706 | 0.004024903 | Credible Set | purple |
| chr7:104170618 | 7 | 104.1706 | 0.004047853 | Credible Set | purple |
| chr7:104170662 | 7 | 104.1707 | 0.004047853 | Credible Set | purple |
| chr7:104338784 | 7 | 104.3388 | 0.008341737 | Credible Set | purple |
| chr7:104338810 | 7 | 104.3388 | 0.008275938 | Credible Set | purple |
| chr7:104353491 | 7 | 104.3535 | 0.012048517 | Credible Set | purple |
| chr7:104356230 | 7 | 104.3562 | 0.007725665 | Credible Set | purple |
| chr7:104430205 | 7 | 104.4302 | 0.012740275 | Credible Set | purple |
| chr7:104485236 | 7 | 104.4852 | 0.018029211 | Credible Set | purple |
| chr7:104485721 | 7 | 104.4857 | 0.003407890 | Credible Set | purple |
| chr7:104485727 | 7 | 104.4857 | 0.003581521 | Credible Set | purple |
| chr7:104486022 | 7 | 104.4860 | 0.020443497 | Credible Set | purple |
| chr7:104486041 | 7 | 104.4860 | 0.021482585 | Credible Set | purple |
| chr7:104507724 | 7 | 104.5077 | 0.029421529 | Credible Set | purple |
| chr7:104508127 | 7 | 104.5081 | 0.026461520 | Credible Set | purple |
| chr7:104508173 | 7 | 104.5082 | 0.026290096 | Credible Set | purple |
| chr7:104580091 | 7 | 104.5801 | 0.017032766 | Credible Set | purple |
| chr7:105411752 | 7 | 105.4118 | 0.006632638 | Credible Set | purple |

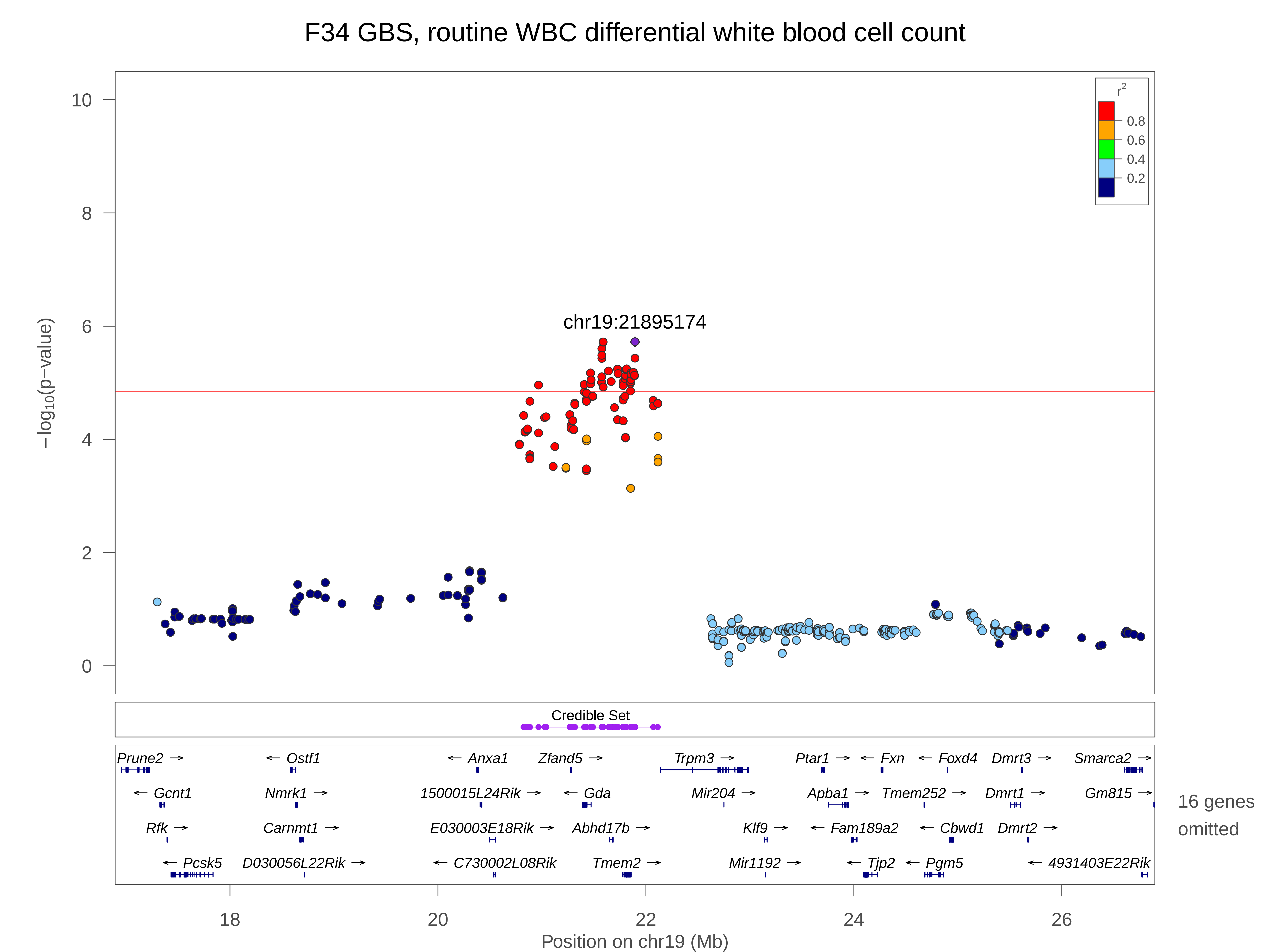
F34 GBS, routine WBC differential white blood cell count
−log10(p−value)
chr19:21895174
r2
0.8
0.6
0.4
0.2
Credible Set
Prune2 → ← Ostf1 ← Anxa1 Zfand5 → Trpm3 → Ptar1 → ← Fxn ← Foxd4 Dmrt3 → Smarca2 →
← Gcnt1 Nmrk1 → 1500015L24Rik → ← Gda Mir204 → Apba1 → Tmem252 → Dmrt1 → Gm815 →
Rfk → Carnmt1 → E030003E18Rik → Abhd17b → Klf9 → ← Fam189a2 ← Cbwd1 Dmrt2 →
← Pcsk5 D030056L22Rik → ← C730002L08Rik Tmem2 → Mir1192 → ← Tjp2 ← Pgm5 ← 4931403E22Rik
16 genes omitted
Position on chr19 (Mb)

| | |
|---|---|
| date: | Thu Oct 25 17:01:36 2018 |
| build: | F34GBS |
| display range: | chr19:16895174−26895174 [16895174−26895174] |
| hilite range: | 0 − 0 [ 0 − 0 ] |
| reference SNP: | chr19:21895174 |
| number of SNPs plotted: | 470 |
| min P−value: | 1.88E−6 [chr19:21895174] |
| max P−value: | 8.76E−1 [chr19:22798742] |
| omitted Genes: | Trpm6, Rorb, Aldh1a1 |
| omitted Genes: | Aldh1a7, Tmc1, 4930554I06Rik |
| omitted Genes: | 1110059E24Rik, Smc5, Mamdc2 |
| omitted Genes: | 1700028P14Rik, Pip5k1b, Fam122a |
| omitted Genes: | Mir3084−1, Dock8, Kank1 |
| omitted Genes: | 2610016A17Rik |

Make more plots at http://csg.sph.umich.edu/locuszoom/

Fine−mapping Regions

| snp | chr | pos (Mb) | pp | group | color |
|---|---|---|---|---|---|
| chr19:20824062 | 19 | 20.82406 | 0.002585794 | Credible Set | purple |
| chr19:20837340 | 19 | 20.83734 | 0.001363790 | Credible Set | purple |
| chr19:20837342 | 19 | 20.83734 | 0.001382413 | Credible Set | purple |
| chr19:20861811 | 19 | 20.86181 | 0.001489729 | Credible Set | purple |
| chr19:20861821 | 19 | 20.86182 | 0.001545357 | Credible Set | purple |
| chr19:20884530 | 19 | 20.88453 | 0.004479106 | Credible Set | purple |
| chr19:20967742 | 19 | 20.96774 | 0.001319861 | Credible Set | purple |
| chr19:20967772 | 19 | 20.96777 | 0.001322070 | Credible Set | purple |
| chr19:20967840 | 19 | 20.96784 | 0.008395932 | Credible Set | purple |
| chr19:20967865 | 19 | 20.96786 | 0.008409044 | Credible Set | purple |
| chr19:21025668 | 19 | 21.02567 | 0.002384590 | Credible Set | purple |
| chr19:21039744 | 19 | 21.03974 | 0.002462130 | Credible Set | purple |
| chr19:21268525 | 19 | 21.26853 | 0.002643325 | Credible Set | purple |
| chr19:21268527 | 19 | 21.26853 | 0.002651431 | Credible Set | purple |
| chr19:21268550 | 19 | 21.26855 | 0.002674964 | Credible Set | purple |
| chr19:21280282 | 19 | 21.28028 | 0.001746321 | Credible Set | purple |
| chr19:21280286 | 19 | 21.28029 | 0.001572745 | Credible Set | purple |
| chr19:21296098 | 19 | 21.29610 | 0.002126306 | Credible Set | purple |
| chr19:21305518 | 19 | 21.30552 | 0.001490133 | Credible Set | purple |
| chr19:21305568 | 19 | 21.30557 | 0.001515579 | Credible Set | purple |
| chr19:21316234 | 19 | 21.31623 | 0.004175380 | Credible Set | purple |
| chr19:21316268 | 19 | 21.31627 | 0.003944184 | Credible Set | purple |
| chr19:21316305 | 19 | 21.31630 | 0.003962344 | Credible Set | purple |
| chr19:21407108 | 19 | 21.40711 | 0.006454658 | Credible Set | purple |
| chr19:21407182 | 19 | 21.40718 | 0.008588335 | Credible Set | purple |
| chr19:21428151 | 19 | 21.42815 | 0.004829997 | Credible Set | purple |
| chr19:21428338 | 19 | 21.42834 | 0.004467193 | Credible Set | purple |
| chr19:21428343 | 19 | 21.42834 | 0.004467193 | Credible Set | purple |
| chr19:21430361 | 19 | 21.43036 | 0.006153998 | Credible Set | purple |
| chr19:21430412 | 19 | 21.43041 | 0.006138345 | Credible Set | purple |
| chr19:21467882 | 19 | 21.46788 | 0.008793568 | Credible Set | purple |
| chr19:21467967 | 19 | 21.46797 | 0.013428357 | Credible Set | purple |
| chr19:21467982 | 19 | 21.46798 | 0.013506313 | Credible Set | purple |
| chr19:21467985 | 19 | 21.46798 | 0.013506313 | Credible Set | purple |
| chr19:21467987 | 19 | 21.46799 | 0.013467367 | Credible Set | purple |
| chr19:21472509 | 19 | 21.47251 | 0.010292809 | Credible Set | purple |

Fine−mapping Regions

| snp | chr | pos (Mb) | pp | group | color |
|---|---|---|---|---|---|
| chr19:21472526 | 19 | 21.47253 | 0.010257024 | Credible Set | purple |
| chr19:21489550 | 19 | 21.48955 | 0.005412916 | Credible Set | purple |
| chr19:21489579 | 19 | 21.48958 | 0.005467641 | Credible Set | purple |
| chr19:21574198 | 19 | 21.57420 | 0.009320179 | Credible Set | purple |
| chr19:21574202 | 19 | 21.57420 | 0.009346112 | Credible Set | purple |
| chr19:21575782 | 19 | 21.57578 | 0.011615154 | Credible Set | purple |
| chr19:21576364 | 19 | 21.57636 | 0.034714558 | Credible Set | purple |
| chr19:21576365 | 19 | 21.57636 | 0.023734065 | Credible Set | purple |
| chr19:21576374 | 19 | 21.57637 | 0.026780454 | Credible Set | purple |
| chr19:21588462 | 19 | 21.58846 | 0.045063501 | Credible Set | purple |
| chr19:21588470 | 19 | 21.58847 | 0.045005342 | Credible Set | purple |
| chr19:21588527 | 19 | 21.58853 | 0.044776888 | Credible Set | purple |
| chr19:21588528 | 19 | 21.58853 | 0.045107340 | Credible Set | purple |
| chr19:21588608 | 19 | 21.58861 | 0.007800315 | Credible Set | purple |
| chr19:21640403 | 19 | 21.64040 | 0.014535903 | Credible Set | purple |
| chr19:21666239 | 19 | 21.66624 | 0.009647457 | Credible Set | purple |
| chr19:21697884 | 19 | 21.69788 | 0.003516256 | Credible Set | purple |
| chr19:21727318 | 19 | 21.72732 | 0.002211662 | Credible Set | purple |
| chr19:21727391 | 19 | 21.72739 | 0.002191642 | Credible Set | purple |
| chr19:21727561 | 19 | 21.72756 | 0.015647139 | Credible Set | purple |
| chr19:21731633 | 19 | 21.73163 | 0.013131097 | Credible Set | purple |
| chr19:21780788 | 19 | 21.78079 | 0.009483067 | Credible Set | purple |
| chr19:21780850 | 19 | 21.78085 | 0.008262784 | Credible Set | purple |
| chr19:21780922 | 19 | 21.78092 | 0.002102234 | Credible Set | purple |
| chr19:21780928 | 19 | 21.78093 | 0.002099716 | Credible Set | purple |
| chr19:21780934 | 19 | 21.78093 | 0.002111114 | Credible Set | purple |
| chr19:21780935 | 19 | 21.78093 | 0.002111114 | Credible Set | purple |
| chr19:21781005 | 19 | 21.78101 | 0.004950729 | Credible Set | purple |
| chr19:21781019 | 19 | 21.78102 | 0.004702405 | Credible Set | purple |
| chr19:21798450 | 19 | 21.79845 | 0.010984485 | Credible Set | purple |
| chr19:21798541 | 19 | 21.79854 | 0.005465049 | Credible Set | purple |
| chr19:21803806 | 19 | 21.80381 | 0.001112543 | Credible Set | purple |
| chr19:21804356 | 19 | 21.80436 | 0.010674368 | Credible Set | purple |
| chr19:21804420 | 19 | 21.80442 | 0.012092035 | Credible Set | purple |
| chr19:21804582 | 19 | 21.80458 | 0.012166277 | Credible Set | purple |
| chr19:21812258 | 19 | 21.81226 | 0.015158100 | Credible Set | purple |

Fine−mapping Regions

| snp | chr | pos (Mb) | pp | group | color |
|---|---|---|---|---|---|
| chr19:21812298 | 19 | 21.81230 | 0.015237991 | Credible Set | purple |
| chr19:21812594 | 19 | 21.81259 | 0.015614975 | Credible Set | purple |
| chr19:21814790 | 19 | 21.81479 | 0.015708525 | Credible Set | purple |
| chr19:21814806 | 19 | 21.81481 | 0.015794260 | Credible Set | purple |
| chr19:21851795 | 19 | 21.85179 | 0.008952670 | Credible Set | purple |
| chr19:21852236 | 19 | 21.85224 | 0.010187775 | Credible Set | purple |
| chr19:21852273 | 19 | 21.85227 | 0.009546923 | Credible Set | purple |
| chr19:21852296 | 19 | 21.85230 | 0.006664577 | Credible Set | purple |
| chr19:21857059 | 19 | 21.85706 | 0.010695036 | Credible Set | purple |
| chr19:21857128 | 19 | 21.85713 | 0.010316721 | Credible Set | purple |
| chr19:21857704 | 19 | 21.85770 | 0.013519818 | Credible Set | purple |
| chr19:21857707 | 19 | 21.85771 | 0.012844288 | Credible Set | purple |
| chr19:21880097 | 19 | 21.88010 | 0.013786204 | Credible Set | purple |
| chr19:21889764 | 19 | 21.88976 | 0.012094988 | Credible Set | purple |
| chr19:21889768 | 19 | 21.88977 | 0.012127490 | Credible Set | purple |
| chr19:21889777 | 19 | 21.88978 | 0.012094988 | Credible Set | purple |
| chr19:21889831 | 19 | 21.88983 | 0.012121119 | Credible Set | purple |
| chr19:21889832 | 19 | 21.88983 | 0.012134329 | Credible Set | purple |
| chr19:21889838 | 19 | 21.88984 | 0.012277854 | Credible Set | purple |
| chr19:21895163 | 19 | 21.89516 | 0.023993456 | Credible Set | purple |
| chr19:21895174 | 19 | 21.89517 | 0.045457301 | Credible Set | purple |
| chr19:22070939 | 19 | 22.07094 | 0.004641355 | Credible Set | purple |
| chr19:22073618 | 19 | 22.07362 | 0.003737414 | Credible Set | purple |
| chr19:22113626 | 19 | 22.11363 | 0.004175365 | Credible Set | purple |
| chr19:22113628 | 19 | 22.11363 | 0.004116085 | Credible Set | purple |
| chr19:22113687 | 19 | 22.11369 | 0.004108361 | Credible Set | purple |

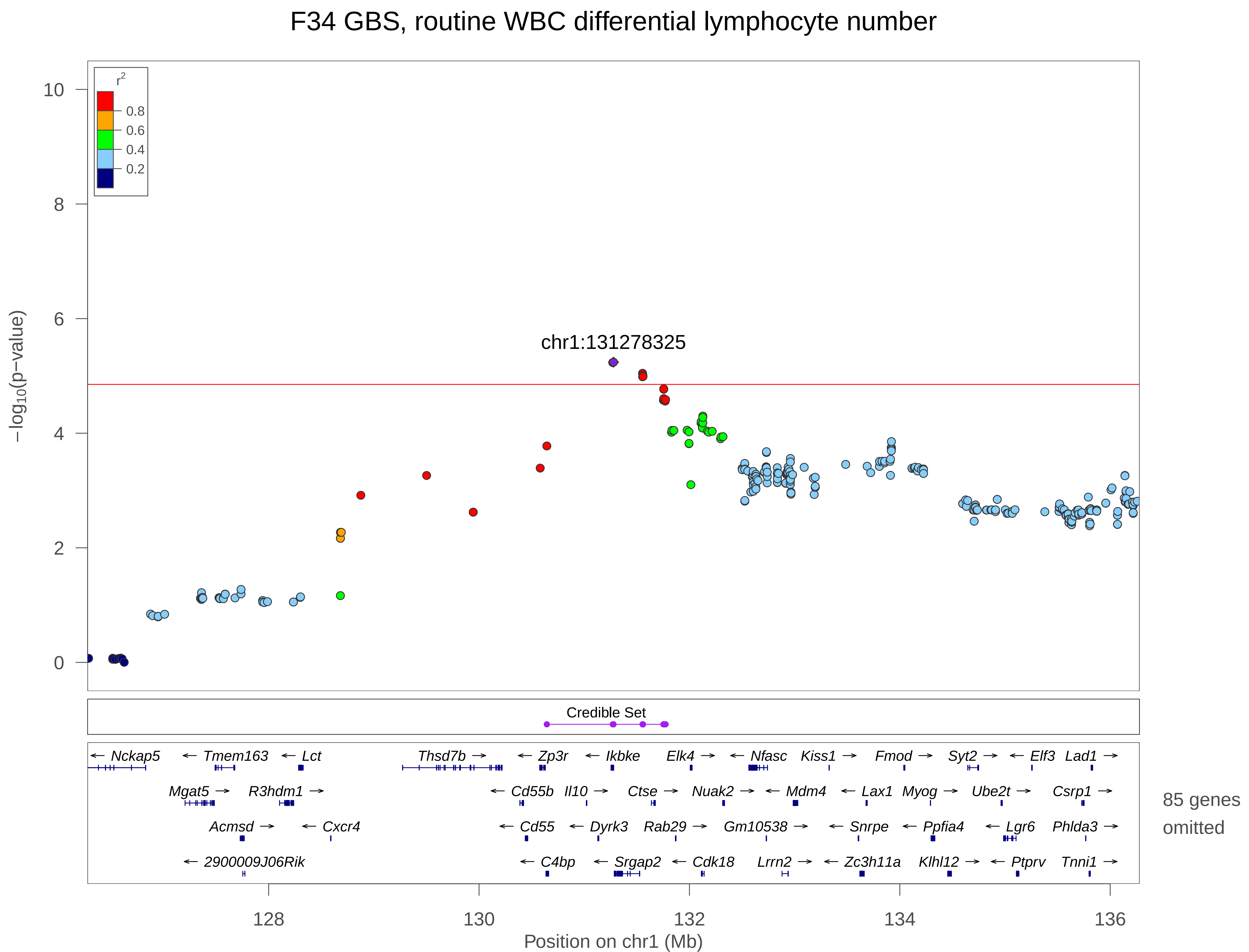
F34 GBS, routine WBC differential lymphocyte number
r2
0.8
0.6
0.4
0.2
chr1:131278325
−log10(p−value)
0
2
4
6
8
10
Credible Set
← Nckap5
← Tmem163
← Lct
Thsd7b →
← Zp3r
← Ikbke
Elk4 →
← Nfasc
Kiss1 →
Fmod →
Syt2 →
← Elf3
Lad1 →
Mgat5 →
R3hdm1 →
← Cd55b
Il10 →
Ctse →
Nuak2 →
← Mdm4
← Lax1
Myog →
Ube2t →
Csrp1 →
Acmsd →
← Cxcr4
← Cd55
← Dyrk3
Rab29 →
Gm10538 →
← Snrpe
← Ppfia4
← Lgr6
Phlda3 →
← 2900009J06Rik
← C4bp
← Srgap2
← Cdk18
Lrrn2 →
← Zc3h11a
Klhl12 →
← Ptprv
Tnni1 →
85 genes omitted
128
130
132
134
136
Position on chr1 (Mb)

| | |
|---|---|
| date: | Thu Oct 25 17:02:15 2018 |
| build: | F34GBS |
| display range: | chr1:126278325−136278325 [126278325−136278325] |
| hilite range: | 0 − 0 [ 0 − 0 ] |
| reference SNP: | chr1:131278325 |
| number of SNPs plotted: | 331 |
| min P−value: | 5.76E−6 [chr1:131278325] |
| max P−value: | 9.95E−1 [chr1:126626341] |
| omitted Genes: | Ccnt2, Map3k19, Rab3gap1 |
| omitted Genes: | Zranb3, Mir128−1, Ubxn4 |
| omitted Genes: | Mcm6, Dars, C4bp−ps1 |
| omitted Genes: | Pfkfb2, Yod1, AA986860 |
| omitted Genes: | Fcamr, Pigr, Fcmr |
| omitted Genes: | Il24, Il20, Il19 |
| omitted Genes: | Mapkapk2, Eif2d, Rassf5 |
| omitted Genes: | Fam72a, Avpr1b, Rab7b |
| omitted Genes: | Slc26a9, Pm20d1, Slc41a1 |
| omitted Genes: | Nucks1, Slc45a3, Mfsd4a |
| omitted Genes: | Lemd1, Mir135b, Klhdc8a |
| omitted Genes: | Tmcc2, Dstyk, Rbbp5 |
| omitted Genes: | Tmem81, Cntn2, Pik3c2b |
| omitted Genes: | Ppp1r15b, Plekha6, Gm19461 |
| omitted Genes: | Golt1a, Ren1, Etnk2 |
| omitted Genes: | Sox13, Zbed6, Atp2b4 |
| omitted Genes: | Mir6903, Optc, Prelp |
| omitted Genes: | Btg2, Chit1, Chil1 |
| omitted Genes: | Mybph, Adora1, Tmem183a |
| omitted Genes: | 4933406M09Rik, Cyb5r1, Adipor1 |
| omitted Genes: | Rabif, Mgat4e, Kdm5b |
| omitted Genes: | Ppp1r12b, Ptpn7, Arl8a |
| omitted Genes: | Gpr37l1, Rnpep, Timm17a |
| omitted Genes: | Lmod1, Shisa4, Ipo9 |
| omitted Genes: | Nav1, Mir1231, Tnnt2 |
| omitted Genes: | Pkp1, Igfn1, Tmem9 |
| omitted Genes: | Ascl5, Cacna1s, Gm15850 |
| omitted Genes: | Kif21b, 5730559C18Rik, Gpr25 |
| omitted Genes: | Camsap2 |

Make more plots at http://csg.sph.umich.edu/locuszoom/

Fine−mapping Regions

| snp | chr | pos (Mb) | pp | group | color |
|---|---|---|---|---|---|
| chr1:130644276 | 1 | 130.6443 | 0.005171124 | Credible Set | purple |
| chr1:131271228 | 1 | 131.2712 | 0.124800595 | Credible Set | purple |
| chr1:131278325 | 1 | 131.2783 | 0.126561546 | Credible Set | purple |
| chr1:131554333 | 1 | 131.5543 | 0.082666046 | Credible Set | purple |
| chr1:131554917 | 1 | 131.5549 | 0.072325200 | Credible Set | purple |
| chr1:131557434 | 1 | 131.5574 | 0.077571556 | Credible Set | purple |
| chr1:131557903 | 1 | 131.5579 | 0.073665721 | Credible Set | purple |
| chr1:131558017 | 1 | 131.5580 | 0.072914337 | Credible Set | purple |
| chr1:131558379 | 1 | 131.5584 | 0.073069638 | Credible Set | purple |
| chr1:131752804 | 1 | 131.7528 | 0.029456788 | Credible Set | purple |
| chr1:131755038 | 1 | 131.7550 | 0.030210485 | Credible Set | purple |
| chr1:131755217 | 1 | 131.7552 | 0.031330051 | Credible Set | purple |
| chr1:131755634 | 1 | 131.7556 | 0.044790792 | Credible Set | purple |
| chr1:131755661 | 1 | 131.7557 | 0.045572795 | Credible Set | purple |
| chr1:131755687 | 1 | 131.7557 | 0.045290624 | Credible Set | purple |
| chr1:131769555 | 1 | 131.7696 | 0.028472515 | Credible Set | purple |
| chr1:131773688 | 1 | 131.7737 | 0.030055953 | Credible Set | purple |

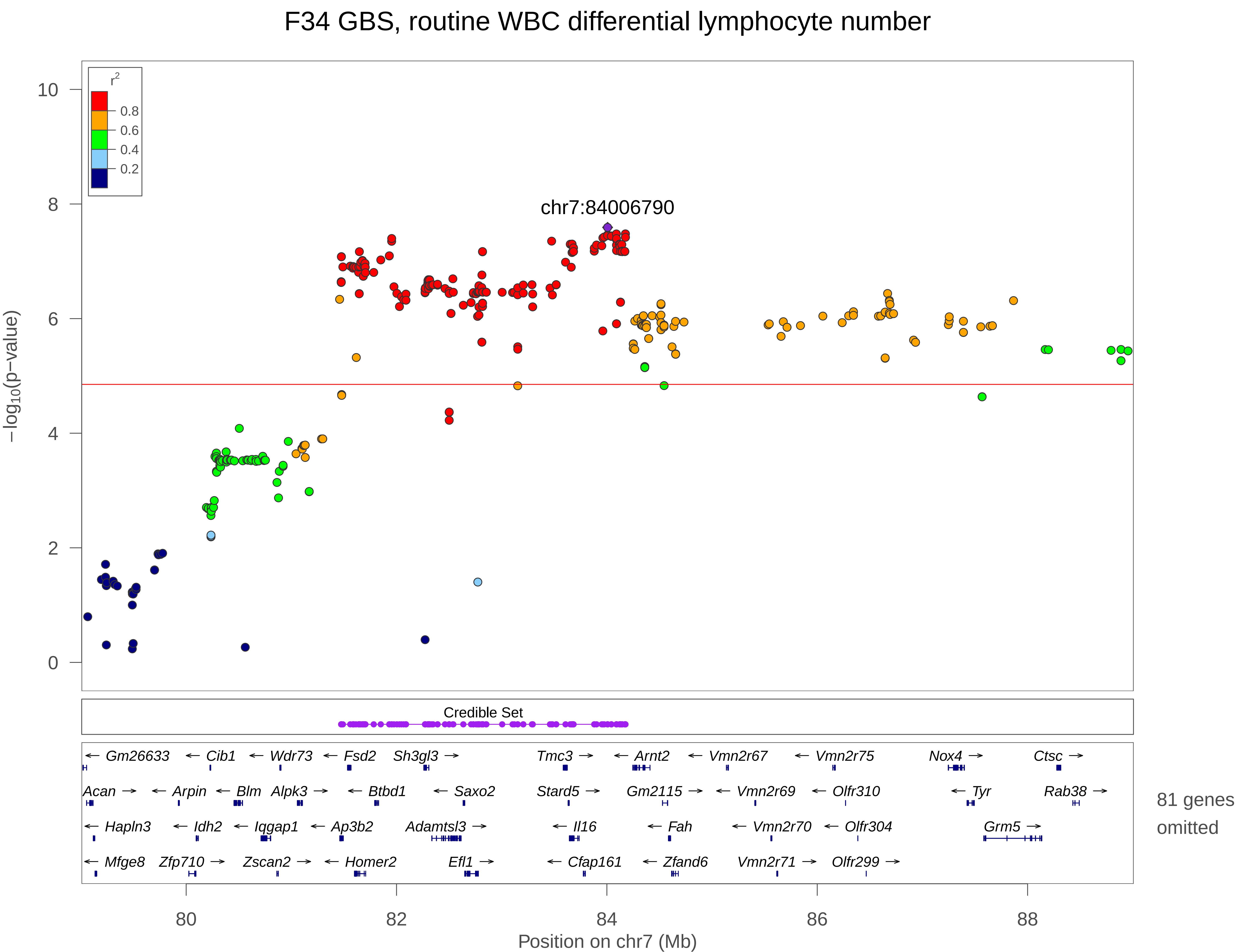

F34 GBS, routine WBC differential lymphocyte number
r2
0.8
0.6
0.4
0.2
chr7:84006790
−log10(p−value)
10
8
6
4
2
0
Credible Set
← Gm26633
← Cib1
← Wdr73
← Fsd2
Sh3gl3 →
Tmc3 →
← Arnt2
← Vmn2r67
← Vmn2r75
Nox4 →
Ctsc →
Acan →
← Arpin
← Blm
Alpk3 →
← Btbd1
← Saxo2
Stard5 →
Gm2115 →
← Vmn2r69
← Olfr310
← Tyr
Rab38 →
← Hapln3
← Idh2
← Iqgap1
← Ap3b2
Adamtsl3 →
← Il16
← Fah
← Vmn2r70
← Olfr304
Grm5 →
← Mfge8
Zfp710 →
Zscan2 →
← Homer2
Efl1 →
← Cfap161
← Zfand6
Vmn2r71 →
Olfr299 →
81 genes omitted
80
82
84
86
88
Position on chr7 (Mb)

| | |
|---|---|
| date: | Thu Oct 25 17:02:46 2018 |
| build: | F34GBS |
| display range: | chr7:79006790−89006790 [79006790−89006790] |
| hilite range: | 0 − 0 [ 0 − 0 ] |
| reference SNP: | chr7:84006790 |
| number of SNPs plotted: | 397 |
| min P−value: | 2.56E−8 [chr7:84006790] |
| max P−value: | 5.8E−1 [chr7:79487919] |
| omitted Genes: | Abhd2, Rlbp1, Fanci |
| omitted Genes: | Polg, Mir9−3hg, Mir9−3 |
| omitted Genes: | Rhcg, Ticrr, Kif7 |
| omitted Genes: | Plin1, Pex11a, Wdr93 |
| omitted Genes: | Mesp1, Mesp2, Anpep |
| omitted Genes: | Ap3s2, Gm21057, Sema4b |
| omitted Genes: | Gdpgp1, Ttll13, Ngrn |
| omitted Genes: | Vps33b, Prc1, Rccd1 |
| omitted Genes: | Unc45a, Hddc3, Man2a2 |
| omitted Genes: | Fes, Furin, Crtc3 |
| omitted Genes: | Gm15880, Nmb, Sec11a |
| omitted Genes: | Zfp592, Slc28a1, Pde8a |
| omitted Genes: | Rps17, Cpeb1, BC048679 |
| omitted Genes: | 2900076A07Rik, Mir1839, Whamm |
| omitted Genes: | Fam103a1, 3110040N11Rik, Tm6sf1 |
| omitted Genes: | Hdgfrp3, Bnc1, Gm20744 |
| omitted Genes: | 4933406J10Rik, Mex3b, 4933430H16Rik |
| omitted Genes: | Mesdc1, Mesdc2, Cemip |
| omitted Genes: | Abhd17c, 2610206C17Rik, Olfr291 |
| omitted Genes: | Olfr290, Vmn2r65, Vmn2r66 |
| omitted Genes: | Vmn2r68, Vmn2r72, Vmn2r73 |
| omitted Genes: | Vmn2r74, Vmn2r76, Olfr309 |
| omitted Genes: | Olfr308, Olfr307, Olfr305 |
| omitted Genes: | Olfr303, Olfr301, Olfr298 |
| omitted Genes: | Olfr297, Olfr295, Olfr294 |
| omitted Genes: | Olfr293, Olfr292, Folh1 |
| omitted Genes: | Vmn2r77, Vmn2r78, Vmn2r79 |

Make more plots at http://csg.sph.umich.edu/locuszoom/

Fine−mapping Regions

| snp | chr | pos (Mb) | pp | group | color |
|---|---|---|---|---|---|
| chr7:81473821 | 7 | 81.47382 | 0.003026101 | Credible Set | purple |
| chr7:81473841 | 7 | 81.47384 | 0.003122681 | Credible Set | purple |
| chr7:81475123 | 7 | 81.47512 | 0.003052756 | Credible Set | purple |
| chr7:81475858 | 7 | 81.47586 | 0.008274026 | Credible Set | purple |
| chr7:81475869 | 7 | 81.47587 | 0.008217810 | Credible Set | purple |
| chr7:81489705 | 7 | 81.48971 | 0.005538830 | Credible Set | purple |
| chr7:81560306 | 7 | 81.56031 | 0.005716971 | Credible Set | purple |
| chr7:81584607 | 7 | 81.58461 | 0.005287670 | Credible Set | purple |
| chr7:81587028 | 7 | 81.58703 | 0.005626033 | Credible Set | purple |
| chr7:81587096 | 7 | 81.58710 | 0.005586258 | Credible Set | purple |
| chr7:81595380 | 7 | 81.59538 | 0.005512988 | Credible Set | purple |
| chr7:81614445 | 7 | 81.61445 | 0.005449247 | Credible Set | purple |
| chr7:81639723 | 7 | 81.63972 | 0.004486953 | Credible Set | purple |
| chr7:81641021 | 7 | 81.64102 | 0.005527387 | Credible Set | purple |
| chr7:81641166 | 7 | 81.64117 | 0.005560081 | Credible Set | purple |
| chr7:81645454 | 7 | 81.64545 | 0.001969161 | Credible Set | purple |
| chr7:81645464 | 7 | 81.64546 | 0.001950338 | Credible Set | purple |
| chr7:81646679 | 7 | 81.64668 | 0.010038701 | Credible Set | purple |
| chr7:81653902 | 7 | 81.65390 | 0.005661306 | Credible Set | purple |
| chr7:81660191 | 7 | 81.66019 | 0.006627368 | Credible Set | purple |
| chr7:81675727 | 7 | 81.67573 | 0.007133144 | Credible Set | purple |
| chr7:81675758 | 7 | 81.67576 | 0.007000544 | Credible Set | purple |
| chr7:81683805 | 7 | 81.68381 | 0.003858310 | Credible Set | purple |
| chr7:81683816 | 7 | 81.68382 | 0.003852926 | Credible Set | purple |
| chr7:81691936 | 7 | 81.69194 | 0.005884701 | Credible Set | purple |
| chr7:81691946 | 7 | 81.69195 | 0.005610961 | Credible Set | purple |
| chr7:81700477 | 7 | 81.70048 | 0.006368899 | Credible Set | purple |
| chr7:81702582 | 7 | 81.70258 | 0.005470962 | Credible Set | purple |
| chr7:81704380 | 7 | 81.70438 | 0.004426288 | Credible Set | purple |
| chr7:81783181 | 7 | 81.78318 | 0.004471243 | Credible Set | purple |
| chr7:81850247 | 7 | 81.85025 | 0.007272302 | Credible Set | purple |
| chr7:81931360 | 7 | 81.93136 | 0.008491558 | Credible Set | purple |
| chr7:81931538 | 7 | 81.93154 | 0.008543284 | Credible Set | purple |
| chr7:81953706 | 7 | 81.95371 | 0.015024622 | Credible Set | purple |
| chr7:81954104 | 7 | 81.95410 | 0.016755415 | Credible Set | purple |
| chr7:81975588 | 7 | 81.97559 | 0.002564800 | Credible Set | purple |

Fine−mapping Regions

| snp | chr | pos (Mb) | pp | group | color |
|---|---|---|---|---|---|
| chr7:82003279 | 7 | 82.00328 | 0.001981678 | Credible Set | purple |
| chr7:82027931 | 7 | 82.02793 | 0.001189539 | Credible Set | purple |
| chr7:82047190 | 7 | 82.04719 | 0.001740457 | Credible Set | purple |
| chr7:82068213 | 7 | 82.06821 | 0.001548150 | Credible Set | purple |
| chr7:82089480 | 7 | 82.08948 | 0.001930510 | Credible Set | purple |
| chr7:82089724 | 7 | 82.08972 | 0.001520962 | Credible Set | purple |
| chr7:82271335 | 7 | 82.27133 | 0.002321685 | Credible Set | purple |
| chr7:82271484 | 7 | 82.27148 | 0.002030760 | Credible Set | purple |
| chr7:82271646 | 7 | 82.27165 | 0.002074878 | Credible Set | purple |
| chr7:82271673 | 7 | 82.27167 | 0.002076964 | Credible Set | purple |
| chr7:82274291 | 7 | 82.27429 | 0.002450207 | Credible Set | purple |
| chr7:82274554 | 7 | 82.27455 | 0.002412916 | Credible Set | purple |
| chr7:82297457 | 7 | 82.29746 | 0.002914205 | Credible Set | purple |
| chr7:82299107 | 7 | 82.29911 | 0.003198637 | Credible Set | purple |
| chr7:82299132 | 7 | 82.29913 | 0.003278590 | Credible Set | purple |
| chr7:82299530 | 7 | 82.29953 | 0.002786006 | Credible Set | purple |
| chr7:82303408 | 7 | 82.30341 | 0.003303529 | Credible Set | purple |
| chr7:82303412 | 7 | 82.30341 | 0.003374319 | Credible Set | purple |
| chr7:82303615 | 7 | 82.30361 | 0.002392926 | Credible Set | purple |
| chr7:82303634 | 7 | 82.30363 | 0.002376889 | Credible Set | purple |
| chr7:82313033 | 7 | 82.31303 | 0.002812280 | Credible Set | purple |
| chr7:82313147 | 7 | 82.31315 | 0.002999856 | Credible Set | purple |
| chr7:82313292 | 7 | 82.31329 | 0.002812280 | Credible Set | purple |
| chr7:82313462 | 7 | 82.31346 | 0.002829550 | Credible Set | purple |
| chr7:82314291 | 7 | 82.31429 | 0.003366730 | Credible Set | purple |
| chr7:82314356 | 7 | 82.31436 | 0.003320052 | Credible Set | purple |
| chr7:82314779 | 7 | 82.31478 | 0.002616731 | Credible Set | purple |
| chr7:82314816 | 7 | 82.31482 | 0.002683564 | Credible Set | purple |
| chr7:82335262 | 7 | 82.33526 | 0.002733645 | Credible Set | purple |
| chr7:82347625 | 7 | 82.34762 | 0.002764231 | Credible Set | purple |
| chr7:82390271 | 7 | 82.39027 | 0.002728454 | Credible Set | purple |
| chr7:82390280 | 7 | 82.39028 | 0.002829724 | Credible Set | purple |
| chr7:82460970 | 7 | 82.46097 | 0.002394497 | Credible Set | purple |
| chr7:82500110 | 7 | 82.50011 | 0.002010226 | Credible Set | purple |
| chr7:82500157 | 7 | 82.50016 | 0.001994627 | Credible Set | purple |
| chr7:82500479 | 7 | 82.50048 | 0.002167026 | Credible Set | purple |

Fine−mapping Regions

| snp | chr | pos (Mb) | pp | group | color |
|---|---|---|---|---|---|
| chr7:82501151 | 7 | 82.50115 | 0.001973573 | Credible Set | purple |
| chr7:82535591 | 7 | 82.53559 | 0.003493037 | Credible Set | purple |
| chr7:82539877 | 7 | 82.53988 | 0.002072976 | Credible Set | purple |
| chr7:82635047 | 7 | 82.63505 | 0.001252680 | Credible Set | purple |
| chr7:82708811 | 7 | 82.70881 | 0.001388179 | Credible Set | purple |
| chr7:82730396 | 7 | 82.73040 | 0.001964861 | Credible Set | purple |
| chr7:82730540 | 7 | 82.73054 | 0.002049199 | Credible Set | purple |
| chr7:82756984 | 7 | 82.75698 | 0.001951843 | Credible Set | purple |
| chr7:82770538 | 7 | 82.77054 | 0.002082988 | Credible Set | purple |
| chr7:82773085 | 7 | 82.77308 | 0.002101881 | Credible Set | purple |
| chr7:82782979 | 7 | 82.78298 | 0.002082988 | Credible Set | purple |
| chr7:82783483 | 7 | 82.78348 | 0.002626973 | Credible Set | purple |
| chr7:82783548 | 7 | 82.78355 | 0.002667989 | Credible Set | purple |
| chr7:82787558 | 7 | 82.78756 | 0.002145889 | Credible Set | purple |
| chr7:82810186 | 7 | 82.81019 | 0.002491472 | Credible Set | purple |
| chr7:82811761 | 7 | 82.81176 | 0.004050773 | Credible Set | purple |
| chr7:82817547 | 7 | 82.81755 | 0.009973121 | Credible Set | purple |
| chr7:82817550 | 7 | 82.81755 | 0.010080208 | Credible Set | purple |
| chr7:82817963 | 7 | 82.81796 | 0.001361527 | Credible Set | purple |
| chr7:82817972 | 7 | 82.81797 | 0.001191165 | Credible Set | purple |
| chr7:82817975 | 7 | 82.81798 | 0.001348065 | Credible Set | purple |
| chr7:82818115 | 7 | 82.81812 | 0.002079740 | Credible Set | purple |
| chr7:82820779 | 7 | 82.82078 | 0.002082988 | Credible Set | purple |
| chr7:82820783 | 7 | 82.82078 | 0.002082988 | Credible Set | purple |
| chr7:82820806 | 7 | 82.82081 | 0.002082988 | Credible Set | purple |
| chr7:82850796 | 7 | 82.85080 | 0.002082988 | Credible Set | purple |
| chr7:82854160 | 7 | 82.85416 | 0.002073230 | Credible Set | purple |
| chr7:83004458 | 7 | 83.00446 | 0.002069762 | Credible Set | purple |
| chr7:83103367 | 7 | 83.10337 | 0.002049895 | Credible Set | purple |
| chr7:83118843 | 7 | 83.11884 | 0.002069472 | Credible Set | purple |
| chr7:83151951 | 7 | 83.15195 | 0.001876320 | Credible Set | purple |
| chr7:83152293 | 7 | 83.15229 | 0.002464111 | Credible Set | purple |
| chr7:83204993 | 7 | 83.20499 | 0.002006845 | Credible Set | purple |
| chr7:83205102 | 7 | 83.20510 | 0.002745910 | Credible Set | purple |
| chr7:83288039 | 7 | 83.28804 | 0.002776773 | Credible Set | purple |
| chr7:83294270 | 7 | 83.29427 | 0.001176128 | Credible Set | purple |

Fine−mapping Regions

| snp | chr | pos (Mb) | pp | group | color |
|---|---|---|---|---|---|
| chr7:83294302 | 7 | 83.29430 | 0.001176128 | Credible Set | purple |
| chr7:83294391 | 7 | 83.29439 | 0.001934993 | Credible Set | purple |
| chr7:83459164 | 7 | 83.45916 | 0.002437130 | Credible Set | purple |
| chr7:83474799 | 7 | 83.47480 | 0.015115469 | Credible Set | purple |
| chr7:83481871 | 7 | 83.48187 | 0.001864795 | Credible Set | purple |
| chr7:83518242 | 7 | 83.51824 | 0.002756305 | Credible Set | purple |
| chr7:83518263 | 7 | 83.51826 | 0.002787826 | Credible Set | purple |
| chr7:83606909 | 7 | 83.60691 | 0.006664566 | Credible Set | purple |
| chr7:83651842 | 7 | 83.65184 | 0.013406210 | Credible Set | purple |
| chr7:83661158 | 7 | 83.66116 | 0.005479029 | Credible Set | purple |
| chr7:83668857 | 7 | 83.66886 | 0.012777413 | Credible Set | purple |
| chr7:83668910 | 7 | 83.66891 | 0.009708079 | Credible Set | purple |
| chr7:83669157 | 7 | 83.66916 | 0.013482005 | Credible Set | purple |
| chr7:83677943 | 7 | 83.67794 | 0.010030132 | Credible Set | purple |
| chr7:83677967 | 7 | 83.67797 | 0.010198684 | Credible Set | purple |
| chr7:83682817 | 7 | 83.68282 | 0.011631887 | Credible Set | purple |
| chr7:83682921 | 7 | 83.68292 | 0.010079954 | Credible Set | purple |
| chr7:83879999 | 7 | 83.88000 | 0.010229013 | Credible Set | purple |
| chr7:83880145 | 7 | 83.88014 | 0.011484956 | Credible Set | purple |
| chr7:83901893 | 7 | 83.90189 | 0.013011030 | Credible Set | purple |
| chr7:83951649 | 7 | 83.95165 | 0.012608519 | Credible Set | purple |
| chr7:83961926 | 7 | 83.96193 | 0.017167467 | Credible Set | purple |
| chr7:83974098 | 7 | 83.97410 | 0.017752599 | Credible Set | purple |
| chr7:84006464 | 7 | 84.00646 | 0.019292341 | Credible Set | purple |
| chr7:84006790 | 7 | 84.00679 | 0.025817660 | Credible Set | purple |
| chr7:84006899 | 7 | 84.00690 | 0.019285117 | Credible Set | purple |
| chr7:84007186 | 7 | 84.00719 | 0.018768817 | Credible Set | purple |
| chr7:84042795 | 7 | 84.04279 | 0.018504064 | Credible Set | purple |
| chr7:84042808 | 7 | 84.04281 | 0.018167839 | Credible Set | purple |
| chr7:84043291 | 7 | 84.04329 | 0.018286351 | Credible Set | purple |
| chr7:84090415 | 7 | 84.09041 | 0.020030694 | Credible Set | purple |
| chr7:84091016 | 7 | 84.09102 | 0.016138703 | Credible Set | purple |
| chr7:84091395 | 7 | 84.09140 | 0.012946410 | Credible Set | purple |
| chr7:84091408 | 7 | 84.09141 | 0.016709005 | Credible Set | purple |
| chr7:84091894 | 7 | 84.09189 | 0.010483652 | Credible Set | purple |
| chr7:84119855 | 7 | 84.11986 | 0.013429388 | Credible Set | purple |

Fine−mapping Regions

| snp | chr | pos (Mb) | pp | group | color |
|---|---|---|---|---|---|
| chr7:84124190 | 7 | 84.12419 | 0.013337639 | Credible Set | purple |
| chr7:84124515 | 7 | 84.12452 | 0.012123053 | Credible Set | purple |
| chr7:84129412 | 7 | 84.12941 | 0.001416950 | Credible Set | purple |
| chr7:84129418 | 7 | 84.12942 | 0.001402105 | Credible Set | purple |
| chr7:84129648 | 7 | 84.12965 | 0.010095752 | Credible Set | purple |
| chr7:84142029 | 7 | 84.14203 | 0.013222649 | Credible Set | purple |
| chr7:84146037 | 7 | 84.14604 | 0.010252076 | Credible Set | purple |
| chr7:84146051 | 7 | 84.14605 | 0.010032074 | Credible Set | purple |
| chr7:84170763 | 7 | 84.17076 | 0.010055338 | Credible Set | purple |
| chr7:84170827 | 7 | 84.17083 | 0.010151340 | Credible Set | purple |
| chr7:84175856 | 7 | 84.17586 | 0.017693356 | Credible Set | purple |
| chr7:84176468 | 7 | 84.17647 | 0.020084459 | Credible Set | purple |
| chr7:84176482 | 7 | 84.17648 | 0.017566262 | Credible Set | purple |

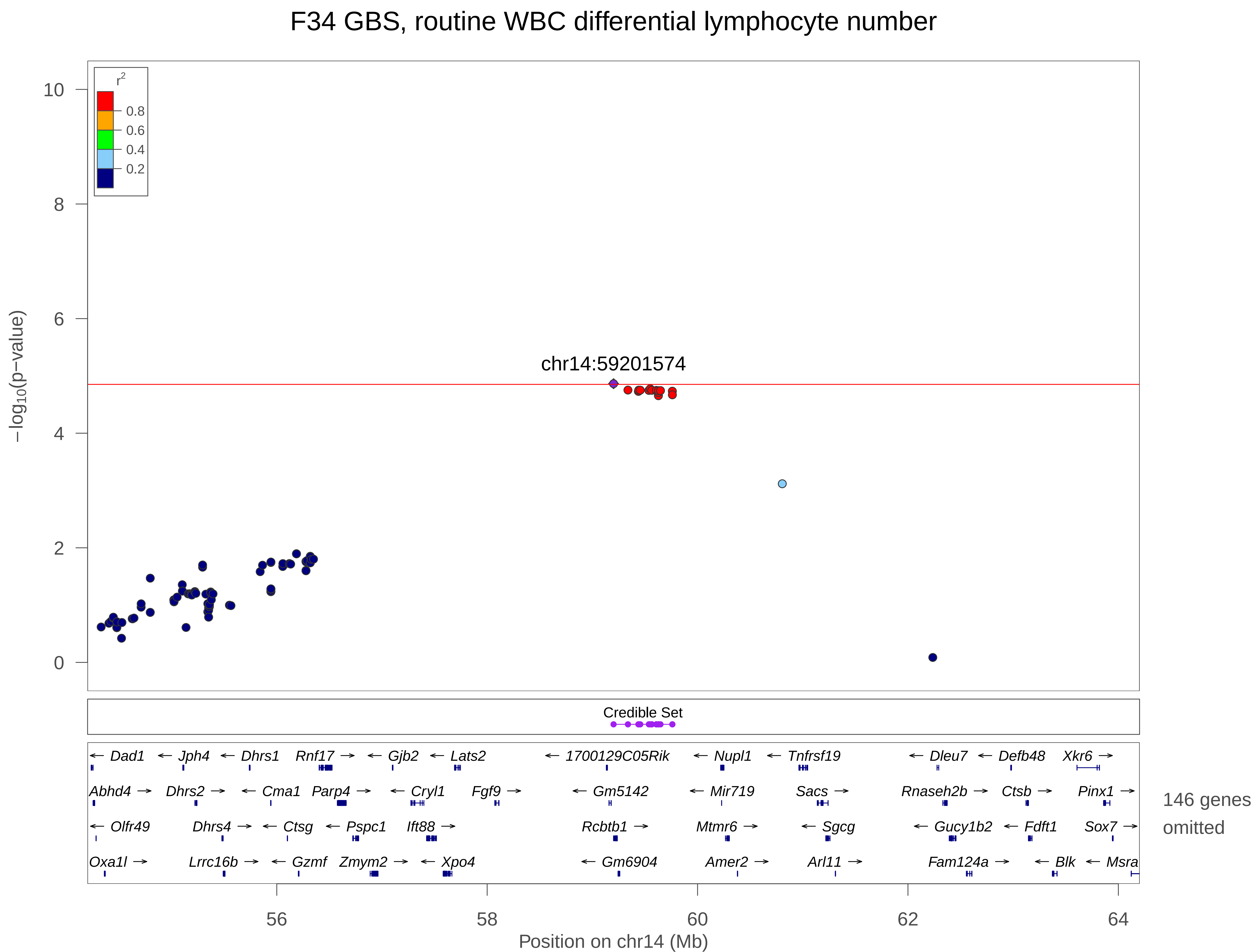

F34 GBS, routine WBC differential lymphocyte number
r2
0.8
0.6
0.4
0.2
10
8
6
4
2
0
−log10(p−value)
chr14:59201574
Credible Set
← Dad1
← Jph4
← Dhrs1
Rnf17 →
← Gjb2
← Lats2
← 1700129C05Rik
← Nupl1
← Tnfrsf19
← Dleu7
← Defb48
Xkr6 →
Abhd4 →
Dhrs2 →
← Cma1
Parp4 →
← Cryl1
Fgf9 →
← Gm5142
← Mir719
Sacs →
Rnaseh2b →
Ctsb →
Pinx1 →
← Olfr49
Dhrs4 →
← Ctsg
← Pspc1
Ift88 →
Rcbtb1 →
Mtmr6 →
← Sgcg
← Gucy1b2
← Fdft1
Sox7 →
Oxa1l →
Lrrc16b →
← Gzmf
Zmym2 →
← Xpo4
← Gm6904
Amer2 →
Arl11 →
Fam124a →
← Blk
← Msra
146 genes omitted
56
58
60
62
64
Position on chr14 (Mb)

| | |
|---|---|
| date: | Thu Oct 25 17:02:27 2018 |
| build: | F34GBS |
| display range: | chr14:54201574−64201574 [54201574−64201574] |
| hilite range: | 0 − 0 [ 0 − 0 ] |
| reference SNP: | chr14:59201574 |
| number of SNPs plotted: | 118 |
| min P−value: | 1.37E−5 [chr14:59201574] |
| max P−value: | 8.19E−1 [chr14:62236131] |
| omitted Genes: | Slc7a7, Mrpl52, Mmp14 |
| omitted Genes: | Lrp10, Rem2, Prmt5 |
| omitted Genes: | Haus4, Ajuba, 4931414P19Rik |
| omitted Genes: | Psmb5, Mir686, Psmb11 |
| omitted Genes: | Cdh24, Acin1, Mir6948 |
| omitted Genes: | 4930579G18Rik, 1700123O20Rik, Cebpe |
| omitted Genes: | Slc7a8, Homez, Ppp1r3e |
| omitted Genes: | Bcl2l2, Pabpn1, Slc22a17 |
| omitted Genes: | Efs, Il25, Cmtm5 |
| omitted Genes: | Myh6, Mir208a, Mhrt |
| omitted Genes: | Myh7, Mir208b, Ngdn |
| omitted Genes: | Gm10364, Zfhx2, Zfhx2os |
| omitted Genes: | Thtpa, Ap1g2, Cpne6 |
| omitted Genes: | Nrl, Pck2, Dcaf11 |
| omitted Genes: | Fitm1, Psme1, Emc9 |
| omitted Genes: | Psme2, Rnf31, Irf9 |
| omitted Genes: | Rec8, Ipo4, Tm9sf1 |
| omitted Genes: | Tssk4, Mdp1, Nedd8 |
| omitted Genes: | Gmpr2, Tinf2, Tgm1 |
| omitted Genes: | Rabggta, Nop9, Cideb |
| omitted Genes: | Ltb4r2, Ltb4r1, Adcy4 |
| omitted Genes: | Ripk3, Nfatc4, Nynrin |
| omitted Genes: | Cbln3, Khnyn, Sdr39u1 |
| omitted Genes: | Mcpt−ps1, Cma2, Gm5801 |
| omitted Genes: | Mcpt1, Mcpt9, Mcpt2 |
| omitted Genes: | Mcpt4, Mcpt8, Mir6949 |
| omitted Genes: | Gzme, Gzmd, Gzmg |
| omitted Genes: | Gzmn, Gzmc, Gzmb |
| omitted Genes: | Atp12a, Cenpj, Mphosph8 |
| omitted Genes: | Gm16973, Zmym5, B020004C17Rik |
| omitted Genes: | Gja3, Gjb6, Il17d |
| omitted Genes: | Eef1akmt1, Sap18, Mir3077 |

Make more plots at http://csg.sph.umich.edu/locuszoom/

Fine−mapping Regions

| snp | chr | pos (Mb) | pp | group | color |
|---|---|---|---|---|---|
| chr14:59201574 | 14 | 59.20157 | 0.07261897 | Credible Set | purple |
| chr14:59338108 | 14 | 59.33811 | 0.05689636 | Credible Set | purple |
| chr14:59437591 | 14 | 59.43759 | 0.05409787 | Credible Set | purple |
| chr14:59440712 | 14 | 59.44071 | 0.05670075 | Credible Set | purple |
| chr14:59455253 | 14 | 59.45525 | 0.05663948 | Credible Set | purple |
| chr14:59537225 | 14 | 59.53722 | 0.05615352 | Credible Set | purple |
| chr14:59549690 | 14 | 59.54969 | 0.05971186 | Credible Set | purple |
| chr14:59564460 | 14 | 59.56446 | 0.05645541 | Credible Set | purple |
| chr14:59565568 | 14 | 59.56557 | 0.05645541 | Credible Set | purple |
| chr14:59608014 | 14 | 59.60801 | 0.05611562 | Credible Set | purple |
| chr14:59608030 | 14 | 59.60803 | 0.05611562 | Credible Set | purple |
| chr14:59625852 | 14 | 59.62585 | 0.05098621 | Credible Set | purple |
| chr14:59627887 | 14 | 59.62789 | 0.04583768 | Credible Set | purple |
| chr14:59628306 | 14 | 59.62831 | 0.05192754 | Credible Set | purple |
| chr14:59628310 | 14 | 59.62831 | 0.05563060 | Credible Set | purple |
| chr14:59647532 | 14 | 59.64753 | 0.05574007 | Credible Set | purple |
| chr14:59760158 | 14 | 59.76016 | 0.05448814 | Credible Set | purple |
| chr14:59760629 | 14 | 59.76063 | 0.04742890 | Credible Set | purple |

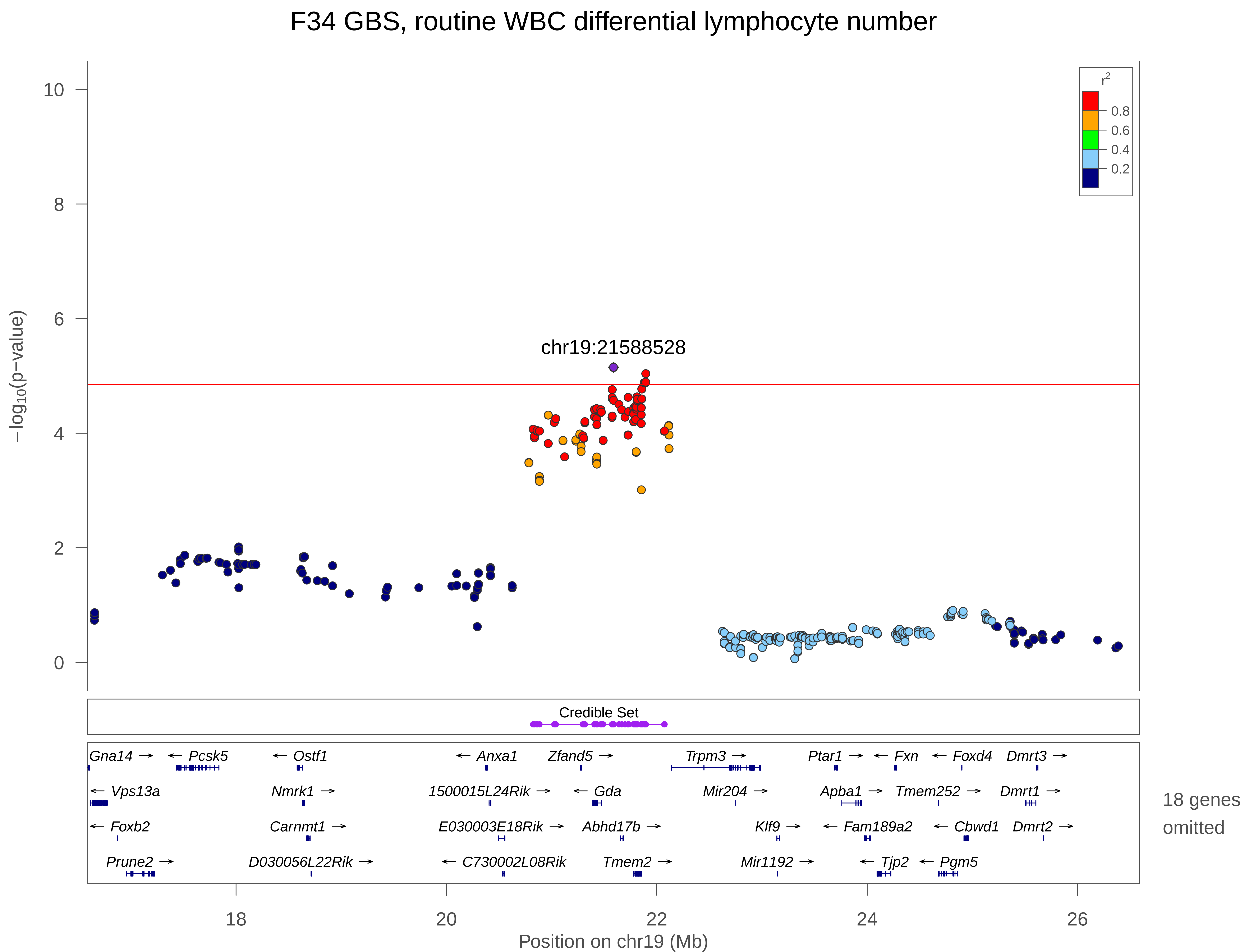
F34 GBS, routine WBC differential lymphocyte number
−log10(p−value)
0
2
4
6
8
10
chr19:21588528
r2
0.8
0.6
0.4
0.2
Credible Set
Gna14 →
← Pcsk5
← Ostf1
← Anxa1
Zfand5 →
Trpm3 →
Ptar1 →
← Fxn
← Foxd4
Dmrt3 →
← Vps13a
Nmrk1 →
1500015L24Rik →
← Gda
Mir204 →
Apba1 →
Tmem252 →
Dmrt1 →
← Foxb2
Carnmt1 →
E030003E18Rik →
Abhd17b →
Klf9 →
← Fam189a2
← Cbwd1
Dmrt2 →
Prune2 →
D030056L22Rik →
← C730002L08Rik
Tmem2 →
Mir1192 →
← Tjp2
← Pgm5
18 genes omitted
18
20
22
24
26
Position on chr19 (Mb)

| | |
|---|---|
| date: | Thu Oct 25 17:02:35 2018 |
| build: | F34GBS |
| display range: | chr19:16588528−26588528 [16588528−26588528] |
| hilite range: | 0 − 0 [ 0 − 0 ] |
| reference SNP: | chr19:21588528 |
| number of SNPs plotted: | 468 |
| min P−value: | 7.07E−6 [chr19:21588528] |
| max P−value: | 8.66E−1 [chr19:23311843] |
| omitted Genes: | Gcnt1, Rfk, Trpm6 |
| omitted Genes: | Rorb, Aldh1a1, Aldh1a7 |
| omitted Genes: | Tmc1, 4930554I06Rik, 1110059E24Rik |
| omitted Genes: | Smc5, Mamdc2, 1700028P14Rik |
| omitted Genes: | Pip5k1b, Fam122a, Mir3084−1 |
| omitted Genes: | Dock8, Kank1, 2610016A17Rik |

Make more plots at http://csg.sph.umich.edu/locuszoom/

Fine−mapping Regions

| snp | chr | pos (Mb) | pp | group | color |
|---|---|---|---|---|---|
| chr19:20824062 | 19 | 20.82406 | 0.004208088 | Credible Set | purple |
| chr19:20837340 | 19 | 20.83734 | 0.003016468 | Credible Set | purple |
| chr19:20837342 | 19 | 20.83734 | 0.003234779 | Credible Set | purple |
| chr19:20861811 | 19 | 20.86181 | 0.003891778 | Credible Set | purple |
| chr19:20861821 | 19 | 20.86182 | 0.003985128 | Credible Set | purple |
| chr19:20884530 | 19 | 20.88453 | 0.003909079 | Credible Set | purple |
| chr19:21025668 | 19 | 21.02567 | 0.005414121 | Credible Set | purple |
| chr19:21039744 | 19 | 21.03974 | 0.006259725 | Credible Set | purple |
| chr19:21296098 | 19 | 21.29610 | 0.003285971 | Credible Set | purple |
| chr19:21305518 | 19 | 21.30552 | 0.002970110 | Credible Set | purple |
| chr19:21305568 | 19 | 21.30557 | 0.003007889 | Credible Set | purple |
| chr19:21316234 | 19 | 21.31623 | 0.005335891 | Credible Set | purple |
| chr19:21316268 | 19 | 21.31627 | 0.005460410 | Credible Set | purple |
| chr19:21316305 | 19 | 21.31630 | 0.005592177 | Credible Set | purple |
| chr19:21407108 | 19 | 21.40711 | 0.006735416 | Credible Set | purple |
| chr19:21407182 | 19 | 21.40718 | 0.008862111 | Credible Set | purple |
| chr19:21428151 | 19 | 21.42815 | 0.006314288 | Credible Set | purple |
| chr19:21428338 | 19 | 21.42834 | 0.009176532 | Credible Set | purple |
| chr19:21428343 | 19 | 21.42834 | 0.009176532 | Credible Set | purple |
| chr19:21430361 | 19 | 21.43036 | 0.004950147 | Credible Set | purple |
| chr19:21430412 | 19 | 21.43041 | 0.005018073 | Credible Set | purple |
| chr19:21467882 | 19 | 21.46788 | 0.007875532 | Credible Set | purple |
| chr19:21467967 | 19 | 21.46797 | 0.008801371 | Credible Set | purple |
| chr19:21467982 | 19 | 21.46798 | 0.008818367 | Credible Set | purple |
| chr19:21467985 | 19 | 21.46798 | 0.008818367 | Credible Set | purple |
| chr19:21467987 | 19 | 21.46799 | 0.008809916 | Credible Set | purple |
| chr19:21472509 | 19 | 21.47251 | 0.008053099 | Credible Set | purple |
| chr19:21472526 | 19 | 21.47253 | 0.008032139 | Credible Set | purple |
| chr19:21489579 | 19 | 21.48958 | 0.002760087 | Credible Set | purple |
| chr19:21574198 | 19 | 21.57420 | 0.006543297 | Credible Set | purple |
| chr19:21574202 | 19 | 21.57420 | 0.006568182 | Credible Set | purple |
| chr19:21575782 | 19 | 21.57578 | 0.006917508 | Credible Set | purple |
| chr19:21576364 | 19 | 21.57636 | 0.018961711 | Credible Set | purple |
| chr19:21576365 | 19 | 21.57636 | 0.013504963 | Credible Set | purple |
| chr19:21576374 | 19 | 21.57637 | 0.014026219 | Credible Set | purple |
| chr19:21588462 | 19 | 21.58846 | 0.044572147 | Credible Set | purple |

Fine−mapping Regions

| snp | chr | pos (Mb) | pp | group | color |
|---|---|---|---|---|---|
| chr19:21588470 | 19 | 21.58847 | 0.044496810 | Credible Set | purple |
| chr19:21588527 | 19 | 21.58853 | 0.044205261 | Credible Set | purple |
| chr19:21588528 | 19 | 21.58853 | 0.044692835 | Credible Set | purple |
| chr19:21588608 | 19 | 21.58861 | 0.012585851 | Credible Set | purple |
| chr19:21640403 | 19 | 21.64040 | 0.010808190 | Credible Set | purple |
| chr19:21666239 | 19 | 21.66624 | 0.008798367 | Credible Set | purple |
| chr19:21697884 | 19 | 21.69788 | 0.006619425 | Credible Set | purple |
| chr19:21727318 | 19 | 21.72732 | 0.003379409 | Credible Set | purple |
| chr19:21727391 | 19 | 21.72739 | 0.003347427 | Credible Set | purple |
| chr19:21727561 | 19 | 21.72756 | 0.014152365 | Credible Set | purple |
| chr19:21731633 | 19 | 21.73163 | 0.008213531 | Credible Set | purple |
| chr19:21780788 | 19 | 21.78079 | 0.009395344 | Credible Set | purple |
| chr19:21780850 | 19 | 21.78085 | 0.009250484 | Credible Set | purple |
| chr19:21780922 | 19 | 21.78092 | 0.005559132 | Credible Set | purple |
| chr19:21780928 | 19 | 21.78093 | 0.005559389 | Credible Set | purple |
| chr19:21780934 | 19 | 21.78093 | 0.005592690 | Credible Set | purple |
| chr19:21780935 | 19 | 21.78093 | 0.005592690 | Credible Set | purple |
| chr19:21781005 | 19 | 21.78101 | 0.007931772 | Credible Set | purple |
| chr19:21781019 | 19 | 21.78102 | 0.007433931 | Credible Set | purple |
| chr19:21798450 | 19 | 21.79845 | 0.010138868 | Credible Set | purple |
| chr19:21798541 | 19 | 21.79854 | 0.006052896 | Credible Set | purple |
| chr19:21804356 | 19 | 21.80436 | 0.009010334 | Credible Set | purple |
| chr19:21804420 | 19 | 21.80442 | 0.009692442 | Credible Set | purple |
| chr19:21804582 | 19 | 21.80458 | 0.009749274 | Credible Set | purple |
| chr19:21812258 | 19 | 21.81226 | 0.014244974 | Credible Set | purple |
| chr19:21812298 | 19 | 21.81230 | 0.014392275 | Credible Set | purple |
| chr19:21812594 | 19 | 21.81259 | 0.012409000 | Credible Set | purple |
| chr19:21814790 | 19 | 21.81479 | 0.013046734 | Credible Set | purple |
| chr19:21814806 | 19 | 21.81481 | 0.013079562 | Credible Set | purple |
| chr19:21851795 | 19 | 21.85179 | 0.007261849 | Credible Set | purple |
| chr19:21852236 | 19 | 21.85224 | 0.009561590 | Credible Set | purple |
| chr19:21852273 | 19 | 21.85227 | 0.009436247 | Credible Set | purple |
| chr19:21852296 | 19 | 21.85230 | 0.005195750 | Credible Set | purple |
| chr19:21857059 | 19 | 21.85706 | 0.013266207 | Credible Set | purple |
| chr19:21857128 | 19 | 21.85713 | 0.013288766 | Credible Set | purple |
| chr19:21857704 | 19 | 21.85770 | 0.019681719 | Credible Set | purple |

Fine−mapping Regions

| snp | chr | pos (Mb) | pp | group | color |
|---|---|---|---|---|---|
| chr19:21857707 | 19 | 21.85771 | 0.019373102 | Credible Set | purple |
| chr19:21880097 | 19 | 21.88010 | 0.024721873 | Credible Set | purple |
| chr19:21889764 | 19 | 21.88976 | 0.024668792 | Credible Set | purple |
| chr19:21889768 | 19 | 21.88977 | 0.024732248 | Credible Set | purple |
| chr19:21889777 | 19 | 21.88978 | 0.024668792 | Credible Set | purple |
| chr19:21889831 | 19 | 21.88983 | 0.024694632 | Credible Set | purple |
| chr19:21889832 | 19 | 21.88983 | 0.024767928 | Credible Set | purple |
| chr19:21889838 | 19 | 21.88984 | 0.025160262 | Credible Set | purple |
| chr19:21895163 | 19 | 21.89516 | 0.025304821 | Credible Set | purple |
| chr19:21895174 | 19 | 21.89517 | 0.035029357 | Credible Set | purple |
| chr19:22070939 | 19 | 22.07094 | 0.003881712 | Credible Set | purple |
| chr19:22073618 | 19 | 22.07362 | 0.003931898 | Credible Set | purple |

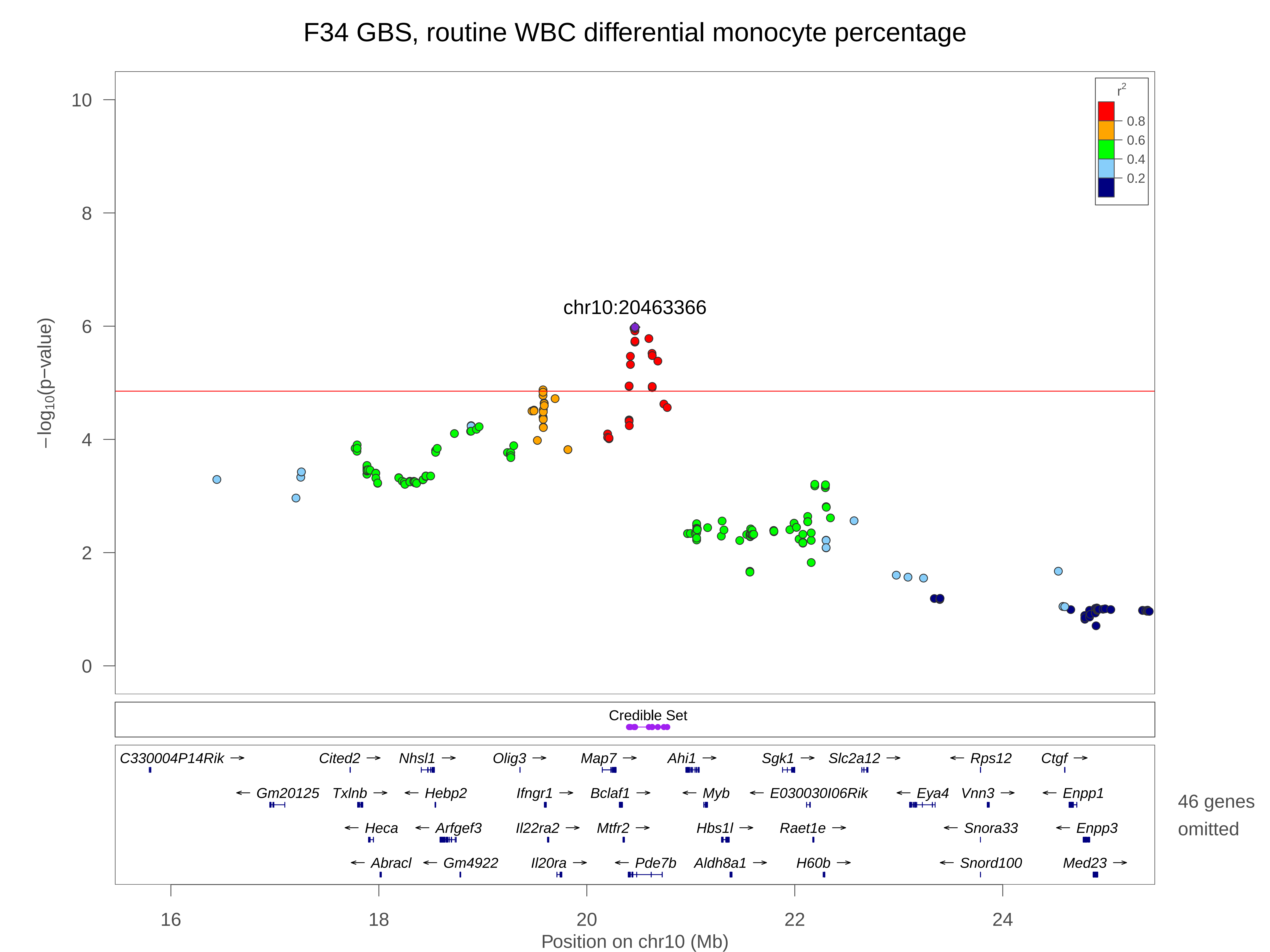
F34 GBS, routine WBC differential monocyte percentage
chr10:20463366
r2
0.8
0.6
0.4
0.2
−log10(p−value)
10
8
6
4
2
0
Credible Set
C330004P14Rik →
Cited2 →
Nhsl1 →
Olig3 →
Map7 →
Ahi1 →
Sgk1 →
Slc2a12 →
← Rps12
Ctgf →
← Gm20125
Txlnb →
← Hebp2
Ifngr1 →
Bclaf1 →
← Myb
← E030030I06Rik
← Eya4
Vnn3 →
← Enpp1
← Heca
← Arfgef3
Il22ra2 →
Mtfr2 →
Hbs1l →
Raet1e →
← Snora33
← Enpp3
← Abracl
← Gm4922
Il20ra →
← Pde7b
Aldh8a1 →
H60b →
← Snord100
Med23 →
46 genes omitted
16
18
20
22
24
Position on chr10 (Mb)

| | |
|---|---|
| date: | Thu Oct 25 17:02:59 2018 |
| build: | F34GBS |
| display range: | chr10:15463366−25463366 [15463366−25463366] |
| hilite range: | 0 − 0 [ 0 − 0 ] |
| reference SNP: | chr10:20463366 |
| number of SNPs plotted: | 243 |
| min P−value: | 1.04E−6 [chr10:20463366] |
| max P−value: | 1.96E−1 [chr10:24897369] |
| omitted Genes: | Reps1, Ect2l, Ccdc28a |
| omitted Genes: | 4930444F02Rik, Perp, Tnfaip3 |
| omitted Genes: | Gm20139, Slc35d3, Pex7 |
| omitted Genes: | Map3k5, 4930405J17Rik, Rpl32l |
| omitted Genes: | Mir6413, 4930455C13Rik, 1700020N01Rik |
| omitted Genes: | Gm5420, 4930444G20Rik, C920009B18Rik |
| omitted Genes: | Raet1d, Gm10825, Tbpl1 |
| omitted Genes: | Mir7663, Tcf21, Slc18b1 |
| omitted Genes: | Vnn1, Taar1, Taar2 |
| omitted Genes: | Taar3, Taar4, Taar5 |
| omitted Genes: | Taar6, Taar7a, Taar7b |
| omitted Genes: | Taar7d, Taar7e, Taar7f |
| omitted Genes: | Taar8a, Taar8b, Taar8c |
| omitted Genes: | Taar9, Stx7, Moxd1 |
| omitted Genes: | Mir6905, Arg1, Akap7 |
| omitted Genes: | Epb41l2 |

Make more plots at http://csg.sph.umich.edu/locuszoom/

Fine−mapping Regions

| snp | chr | pos (Mb) | pp | group | color |
| --- | --- | --- | --- | --- | --- |
| chr10:20406341 | 10 | 20.40634 | 0.001638727 | Credible Set | purple |
| chr10:20406586 | 10 | 20.40659 | 0.006263026 | Credible Set | purple |
| chr10:20406630 | 10 | 20.40663 | 0.005763847 | Credible Set | purple |
| chr10:20406656 | 10 | 20.40666 | 0.006263026 | Credible Set | purple |
| chr10:20419014 | 10 | 20.41901 | 0.021720278 | Credible Set | purple |
| chr10:20419458 | 10 | 20.41946 | 0.013311268 | Credible Set | purple |
| chr10:20453644 | 10 | 20.45364 | 0.062429059 | Credible Set | purple |
| chr10:20462020 | 10 | 20.46202 | 0.062429059 | Credible Set | purple |
| chr10:20462039 | 10 | 20.46204 | 0.062429059 | Credible Set | purple |
| chr10:20462485 | 10 | 20.46249 | 0.032056139 | Credible Set | purple |
| chr10:20462491 | 10 | 20.46249 | 0.032056139 | Credible Set | purple |
| chr10:20462498 | 10 | 20.46250 | 0.032056139 | Credible Set | purple |
| chr10:20462516 | 10 | 20.46252 | 0.032056139 | Credible Set | purple |
| chr10:20462589 | 10 | 20.46259 | 0.062429059 | Credible Set | purple |
| chr10:20462616 | 10 | 20.46262 | 0.062429059 | Credible Set | purple |
| chr10:20462619 | 10 | 20.46262 | 0.062429059 | Credible Set | purple |
| chr10:20462638 | 10 | 20.46264 | 0.062429059 | Credible Set | purple |
| chr10:20462706 | 10 | 20.46271 | 0.062429059 | Credible Set | purple |
| chr10:20463366 | 10 | 20.46337 | 0.062429059 | Credible Set | purple |
| chr10:20463388 | 10 | 20.46339 | 0.062429059 | Credible Set | purple |
| chr10:20463396 | 10 | 20.46340 | 0.062429059 | Credible Set | purple |
| chr10:20596835 | 10 | 20.59683 | 0.032056139 | Credible Set | purple |
| chr10:20627018 | 10 | 20.62702 | 0.021720278 | Credible Set | purple |
| chr10:20628820 | 10 | 20.62882 | 0.005763847 | Credible Set | purple |
| chr10:20628826 | 10 | 20.62883 | 0.005763847 | Credible Set | purple |
| chr10:20628827 | 10 | 20.62883 | 0.005763847 | Credible Set | purple |
| chr10:20628851 | 10 | 20.62885 | 0.005763847 | Credible Set | purple |
| chr10:20629308 | 10 | 20.62931 | 0.021720278 | Credible Set | purple |
| chr10:20683025 | 10 | 20.68303 | 0.016484013 | Credible Set | purple |
| chr10:20740878 | 10 | 20.74088 | 0.002977462 | Credible Set | purple |
| chr10:20773316 | 10 | 20.77332 | 0.002661898 | Credible Set | purple |

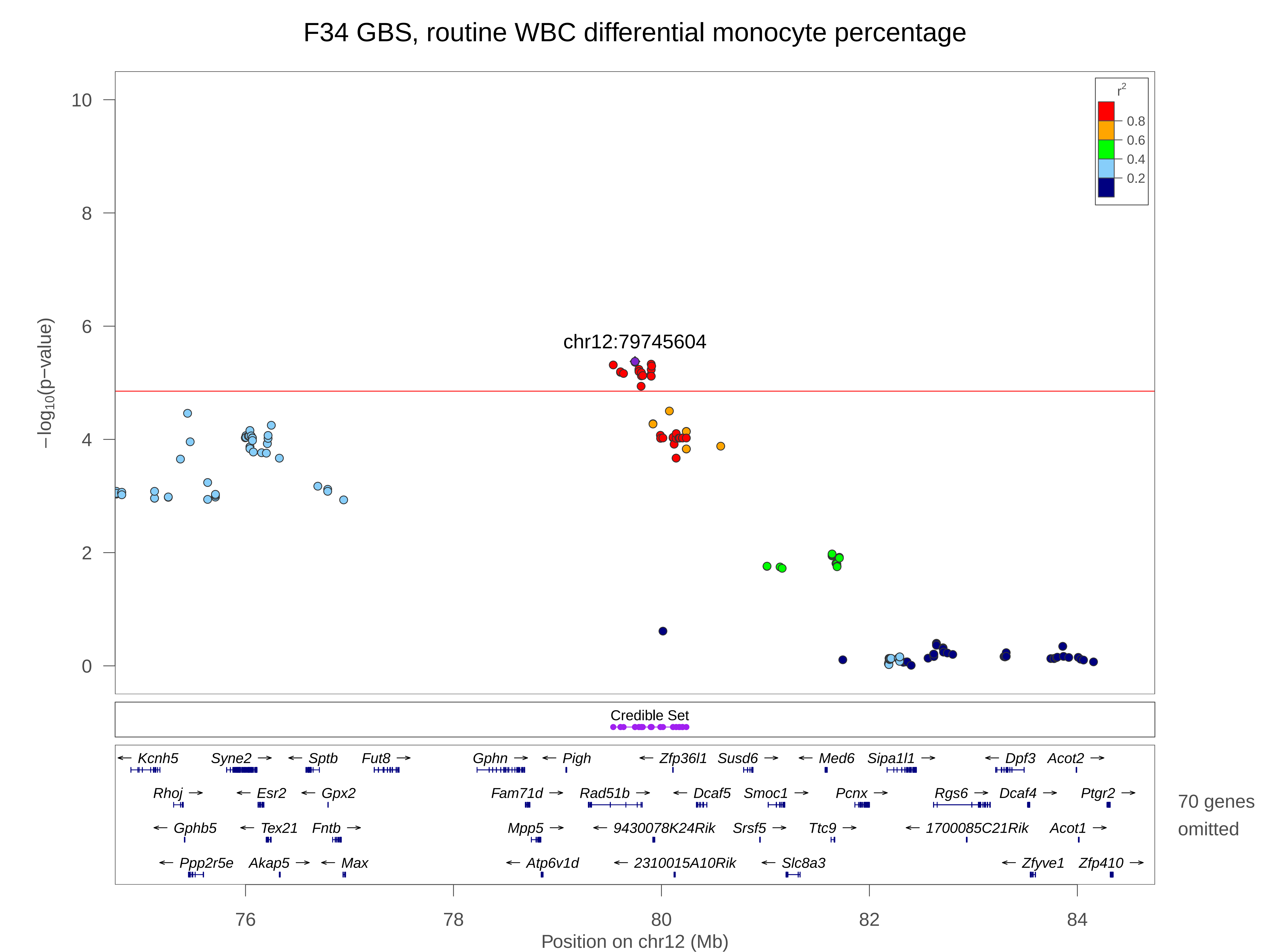
F34 GBS, routine WBC differential monocyte percentage
−log10(p−value)
chr12:79745604
r2
0.8
0.6
0.4
0.2
Credible Set
Kcnh5
Syne2
Sptb
Fut8
Gphn
Pigh
Zfp36l1
Susd6
Med6
Sipa1l1
Dpf3
Acot2
Rhoj
Esr2
Gpx2
Fam71d
Rad51b
Dcaf5
Smoc1
Pcnx
Rgs6
Dcaf4
Ptgr2
Gphb5
Tex21
Fntb
Mpp5
9430078K24Rik
Srsf5
Ttc9
1700085C21Rik
Acot1
Ppp2r5e
Akap5
Max
Atp6v1d
2310015A10Rik
Slc8a3
Zfyve1
Zfp410
70 genes omitted
Position on chr12 (Mb)

| | |
|---|---|
| date: | Thu Oct 25 17:03:09 2018 |
| build: | F34GBS |
| display range: | chr12:74745604−84745604 [74745604−84745604] |
| hilite range: | 0 − 0 [ 0 − 0 ] |
| reference SNP: | chr12:79745604 |
| number of SNPs plotted: | 197 |
| min P−value: | 4.2E−6 [chr12:79745604] |
| max P−value: | 9.78E−1 [chr12:82401670] |
| omitted Genes: | Wdr89, Rplp2−ps1, Sgpp1 |
| omitted Genes: | Mir5101, Mthfd1, Zbtb25 |
| omitted Genes: | Zbtb1, Hspa2, Ppp1r36 |
| omitted Genes: | Gm10451, Mir5135, Plekhg3 |
| omitted Genes: | AI463170, Churc1, Rab15 |
| omitted Genes: | Eif2s1, Plek2, Tmem229b |
| omitted Genes: | Plekhh1, 9230116L04Rik, Arg2 |
| omitted Genes: | Vti1b, Rdh11, Rdh12 |
| omitted Genes: | Zfyve26, Actn1, Mir1843a |
| omitted Genes: | Scarna3b, Exd2, 2310002D06Rik |
| omitted Genes: | Galnt16, Erh, Slc39a9 |
| omitted Genes: | Plekhd1os, Plekhd1, Ccdc177 |
| omitted Genes: | 1700052I22Rik, Gm20337, Slc10a1 |
| omitted Genes: | Mir3067, Cox16, Synj2bp−cox16 |
| omitted Genes: | Gm4787, Adam4, Synj2bp |
| omitted Genes: | Adam21, Map3k9, Rbm25 |
| omitted Genes: | Psen1, Papln, Numb |
| omitted Genes: | 2410016O06Rik, Acot4, Acot3 |
| omitted Genes: | Acot5, Acot6, Dnal1 |
| omitted Genes: | Pnma1, Elmsan1, Fam161b |
| omitted Genes: | Coq6, Entpd5, Bbof1 |
| omitted Genes: | Rnf113a2, Aldh6a1, Lin52 |
| omitted Genes: | Vsx2, Abcd4, Vrtn |
| omitted Genes: | Syndig1l |

Make more plots at http://csg.sph.umich.edu/locuszoom/

Fine−mapping Regions

| snp | chr | pos (Mb) | pp | group | color |
|---|---|---|---|---|---|
| chr12:79536445 | 12 | 79.53645 | 0.055357895 | Credible Set | purple |
| chr12:79606302 | 12 | 79.60630 | 0.041507626 | Credible Set | purple |
| chr12:79606341 | 12 | 79.60634 | 0.042276041 | Credible Set | purple |
| chr12:79606519 | 12 | 79.60652 | 0.042431669 | Credible Set | purple |
| chr12:79635483 | 12 | 79.63548 | 0.039699130 | Credible Set | purple |
| chr12:79745599 | 12 | 79.74560 | 0.063241943 | Credible Set | purple |
| chr12:79745602 | 12 | 79.74560 | 0.061511193 | Credible Set | purple |
| chr12:79745604 | 12 | 79.74560 | 0.063375677 | Credible Set | purple |
| chr12:79783055 | 12 | 79.78306 | 0.044553084 | Credible Set | purple |
| chr12:79783259 | 12 | 79.78326 | 0.046672301 | Credible Set | purple |
| chr12:79783633 | 12 | 79.78363 | 0.042733209 | Credible Set | purple |
| chr12:79804344 | 12 | 79.80434 | 0.024183296 | Credible Set | purple |
| chr12:79807304 | 12 | 79.80730 | 0.036115502 | Credible Set | purple |
| chr12:79807400 | 12 | 79.80740 | 0.040986996 | Credible Set | purple |
| chr12:79819960 | 12 | 79.81996 | 0.036553407 | Credible Set | purple |
| chr12:79896148 | 12 | 79.89615 | 0.037212111 | Credible Set | purple |
| chr12:79901257 | 12 | 79.90126 | 0.057122572 | Credible Set | purple |
| chr12:79901553 | 12 | 79.90155 | 0.046158253 | Credible Set | purple |
| chr12:79901695 | 12 | 79.90170 | 0.035686737 | Credible Set | purple |
| chr12:79901736 | 12 | 79.90174 | 0.035686737 | Credible Set | purple |
| chr12:79905916 | 12 | 79.90592 | 0.052976006 | Credible Set | purple |
| chr12:79989743 | 12 | 79.98974 | 0.003638209 | Credible Set | purple |
| chr12:79991638 | 12 | 79.99164 | 0.003233201 | Credible Set | purple |
| chr12:80014258 | 12 | 80.01426 | 0.003275603 | Credible Set | purple |
| chr12:80111333 | 12 | 80.11133 | 0.003289963 | Credible Set | purple |
| chr12:80111773 | 12 | 80.11177 | 0.003369705 | Credible Set | purple |
| chr12:80141156 | 12 | 80.14116 | 0.003240498 | Credible Set | purple |
| chr12:80141872 | 12 | 80.14187 | 0.003885293 | Credible Set | purple |
| chr12:80167644 | 12 | 80.16764 | 0.003260801 | Credible Set | purple |
| chr12:80176656 | 12 | 80.17666 | 0.003264987 | Credible Set | purple |
| chr12:80200997 | 12 | 80.20100 | 0.003275603 | Credible Set | purple |
| chr12:80201015 | 12 | 80.20101 | 0.003275603 | Credible Set | purple |
| chr12:80203854 | 12 | 80.20385 | 0.003275603 | Credible Set | purple |
| chr12:80203855 | 12 | 80.20386 | 0.003275603 | Credible Set | purple |
| chr12:80239462 | 12 | 80.23946 | 0.003274306 | Credible Set | purple |

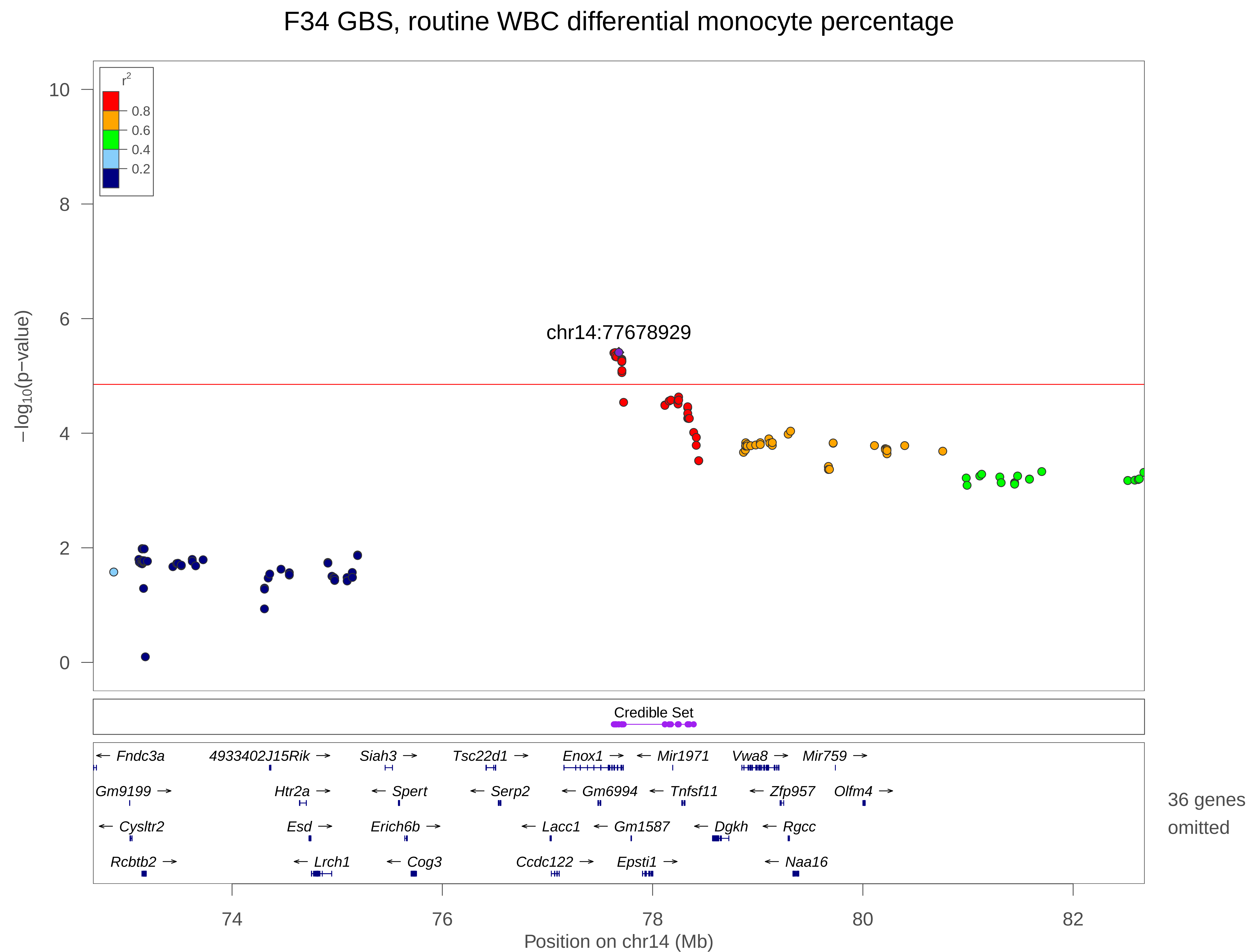

F34 GBS, routine WBC differential monocyte percentage
r2
0.8
0.6
0.4
0.2
10
8
6
4
2
0
−log10(p−value)
chr14:77678929
Credible Set
Fndc3a
4933402J15Rik
Siah3
Tsc22d1
Enox1
Mir1971
Vwa8
Mir759
Gm9199
Htr2a
Spert
Serp2
Gm6994
Tnfsf11
Zfp957
Olfm4
Cysltr2
Esd
Erich6b
Lacc1
Gm1587
Dgkh
Rgcc
Rcbtb2
Lrch1
Cog3
Ccdc122
Epsti1
Naa16
36 genes omitted
74
76
78
80
82
Position on chr14 (Mb)

| | |
|---|---|
| date: | Thu Oct 25 17:03:18 2018 |
| build: | F34GBS |
| display range: | chr14:72678929−82678929 [72678929−82678929] |
| hilite range: | 0 − 0 [ 0 − 0 ] |
| reference SNP: | chr14:77678929 |
| number of SNPs plotted: | 153 |
| min P−value: | 3.88E−6 [chr14:77678929] |
| max P−value: | 7.99E−1 [chr14:73175203] |
| omitted Genes: | Rb1, Mir687, Lpar6 |
| omitted Genes: | Itm2b, Med4, Nudt15 |
| omitted Genes: | Sucla2, Gm4278, 5031414D18Rik |
| omitted Genes: | Lrrc63, Lcp1, Cpb2 |
| omitted Genes: | Zc3h13, Slc25a30, Gm4285 |
| omitted Genes: | Tpt1, Snora31, Gtf2f2 |
| omitted Genes: | Kctd4, Gpalpp1, Nufip1 |
| omitted Genes: | 4930444M15Rik, 1700108F19Rik, Dnajc15 |
| omitted Genes: | Fam216b, Akap11, Mtrf1 |
| omitted Genes: | Kbtbd7, Kbtbd6, Wbp4 |
| omitted Genes: | Elf1, Sugt1, Lect1 |
| omitted Genes: | 4930452G13Rik, Pcdh8, Gm10845 |

Make more plots at http://csg.sph.umich.edu/locuszoom/

Fine−mapping Regions

| snp | chr | pos (Mb) | pp | group | color |
|---|---|---|---|---|---|
| chr14:77631724 | 14 | 77.63172 | 0.060931650 | Credible Set | purple |
| chr14:77642597 | 14 | 77.64260 | 0.061706763 | Credible Set | purple |
| chr14:77642802 | 14 | 77.64280 | 0.061706763 | Credible Set | purple |
| chr14:77647283 | 14 | 77.64728 | 0.052703501 | Credible Set | purple |
| chr14:77647687 | 14 | 77.64769 | 0.054714719 | Credible Set | purple |
| chr14:77658356 | 14 | 77.65836 | 0.052057048 | Credible Set | purple |
| chr14:77678847 | 14 | 77.67885 | 0.062164900 | Credible Set | purple |
| chr14:77678929 | 14 | 77.67893 | 0.062313732 | Credible Set | purple |
| chr14:77707033 | 14 | 77.70703 | 0.047953004 | Credible Set | purple |
| chr14:77707351 | 14 | 77.70735 | 0.047078685 | Credible Set | purple |
| chr14:77707392 | 14 | 77.70739 | 0.045308725 | Credible Set | purple |
| chr14:77707431 | 14 | 77.70743 | 0.044366826 | Credible Set | purple |
| chr14:77707491 | 14 | 77.70749 | 0.043387652 | Credible Set | purple |
| chr14:77707493 | 14 | 77.70749 | 0.044179531 | Credible Set | purple |
| chr14:77707780 | 14 | 77.70778 | 0.030242221 | Credible Set | purple |
| chr14:77707832 | 14 | 77.70783 | 0.028586543 | Credible Set | purple |
| chr14:77708407 | 14 | 77.70841 | 0.030906687 | Credible Set | purple |
| chr14:77724180 | 14 | 77.72418 | 0.009154335 | Credible Set | purple |
| chr14:78116598 | 14 | 78.11660 | 0.008271824 | Credible Set | purple |
| chr14:78116913 | 14 | 78.11691 | 0.008135150 | Credible Set | purple |
| chr14:78157408 | 14 | 78.15741 | 0.009649219 | Credible Set | purple |
| chr14:78173886 | 14 | 78.17389 | 0.009978527 | Credible Set | purple |
| chr14:78240912 | 14 | 78.24091 | 0.009608270 | Credible Set | purple |
| chr14:78240962 | 14 | 78.24096 | 0.009383688 | Credible Set | purple |
| chr14:78241501 | 14 | 78.24150 | 0.008658576 | Credible Set | purple |
| chr14:78241810 | 14 | 78.24181 | 0.008582225 | Credible Set | purple |
| chr14:78245017 | 14 | 78.24502 | 0.009894615 | Credible Set | purple |
| chr14:78245066 | 14 | 78.24507 | 0.010025453 | Credible Set | purple |
| chr14:78247459 | 14 | 78.24746 | 0.011247210 | Credible Set | purple |
| chr14:78247805 | 14 | 78.24780 | 0.009955304 | Credible Set | purple |
| chr14:78333693 | 14 | 78.33369 | 0.007444743 | Credible Set | purple |
| chr14:78333727 | 14 | 78.33373 | 0.007735171 | Credible Set | purple |
| chr14:78334235 | 14 | 78.33424 | 0.004976995 | Credible Set | purple |
| chr14:78334352 | 14 | 78.33435 | 0.005985134 | Credible Set | purple |
| chr14:78342935 | 14 | 78.34293 | 0.004937596 | Credible Set | purple |
| chr14:78349294 | 14 | 78.34929 | 0.004956184 | Credible Set | purple |

Fine−mapping Regions

| snp | chr | pos (Mb) | pp | group | color |
|---|---|---|---|---|---|
| chr14:78390714 | 14 | 78.39071 | 0.002912681 | Credible Set | purple |

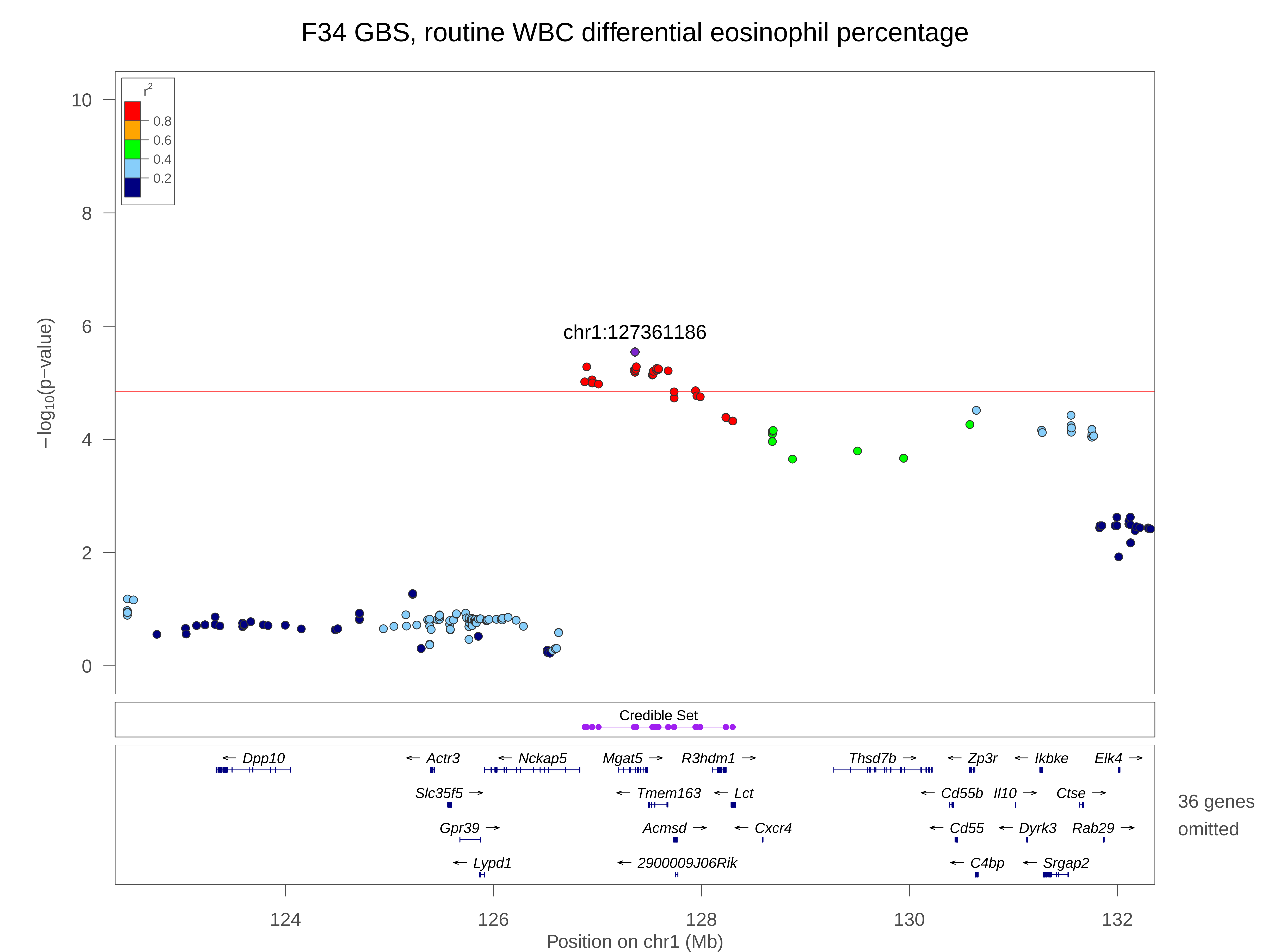
F34 GBS, routine WBC differential eosinophil percentage
r2
0.8
0.6
0.4
0.2
10
8
6
4
2
0
−log10(p−value)
chr1:127361186
Credible Set
← Dpp10
← Actr3
← Nckap5
Mgat5 →
R3hdm1 →
Thsd7b →
← Zp3r
← Ikbke
Elk4 →
Slc35f5 →
← Tmem163
← Lct
← Cd55b
Il10 →
Ctse →
Gpr39 →
Acmsd →
← Cxcr4
← Cd55
← Dyrk3
Rab29 →
← Lypd1
← 2900009J06Rik
← C4bp
← Srgap2
36 genes omitted
124
126
128
130
132
Position on chr1 (Mb)

| | |
|---|---|
| date: | Thu Oct 25 17:03:50 2018 |
| build: | F34GBS |
| display range: | chr1:122361186−132361186 [122361186−132361186] |
| hilite range: | 0 − 0 [ 0 − 0 ] |
| reference SNP: | chr1:127361186 |
| number of SNPs plotted: | 227 |
| min P−value: | 2.85E−6 [chr1:127361186] |
| max P−value: | 5.97E−1 [chr1:126542829] |
| omitted Genes: | Ccnt2, Map3k19, Rab3gap1 |
| omitted Genes: | Zranb3, Mir128−1, Ubxn4 |
| omitted Genes: | Mcm6, Dars, C4bp−ps1 |
| omitted Genes: | Pfkfb2, Yod1, AA986860 |
| omitted Genes: | Fcamr, Pigr, Fcmr |
| omitted Genes: | Il24, Il20, Il19 |
| omitted Genes: | Mapkapk2, Eif2d, Rassf5 |
| omitted Genes: | Fam72a, Avpr1b, Rab7b |
| omitted Genes: | Slc26a9, Pm20d1, Slc41a1 |
| omitted Genes: | Nucks1, Slc45a3, Mfsd4a |
| omitted Genes: | Cdk18, Lemd1, Mir135b |
| omitted Genes: | Klhdc8a, Nuak2, Tmcc2 |

Make more plots at http://csg.sph.umich.edu/locuszoom/

Fine−mapping Regions

| snp | chr | pos (Mb) | pp | group | color |
|---|---|---|---|---|---|
| chr1:126877472 | 1 | 126.8775 | 0.024408867 | Credible Set | purple |
| chr1:126897506 | 1 | 126.8975 | 0.043620608 | Credible Set | purple |
| chr1:126948531 | 1 | 126.9485 | 0.026317821 | Credible Set | purple |
| chr1:126948932 | 1 | 126.9489 | 0.023233275 | Credible Set | purple |
| chr1:126948973 | 1 | 126.9490 | 0.023366825 | Credible Set | purple |
| chr1:127010566 | 1 | 127.0106 | 0.022312139 | Credible Set | purple |
| chr1:127351820 | 1 | 127.3518 | 0.038332922 | Credible Set | purple |
| chr1:127360988 | 1 | 127.3610 | 0.035420933 | Credible Set | purple |
| chr1:127360991 | 1 | 127.3610 | 0.035420933 | Credible Set | purple |
| chr1:127361132 | 1 | 127.3611 | 0.037356199 | Credible Set | purple |
| chr1:127361186 | 1 | 127.3612 | 0.078016154 | Credible Set | purple |
| chr1:127366123 | 1 | 127.3661 | 0.039908366 | Credible Set | purple |
| chr1:127367650 | 1 | 127.3676 | 0.041023204 | Credible Set | purple |
| chr1:127370999 | 1 | 127.3710 | 0.039818213 | Credible Set | purple |
| chr1:127373912 | 1 | 127.3739 | 0.043808802 | Credible Set | purple |
| chr1:127528010 | 1 | 127.5280 | 0.031706054 | Credible Set | purple |
| chr1:127534149 | 1 | 127.5341 | 0.035372315 | Credible Set | purple |
| chr1:127534857 | 1 | 127.5349 | 0.032409527 | Credible Set | purple |
| chr1:127537075 | 1 | 127.5371 | 0.036834344 | Credible Set | purple |
| chr1:127569392 | 1 | 127.5694 | 0.038263434 | Credible Set | purple |
| chr1:127569489 | 1 | 127.5695 | 0.041012765 | Credible Set | purple |
| chr1:127586872 | 1 | 127.5869 | 0.039431793 | Credible Set | purple |
| chr1:127586881 | 1 | 127.5869 | 0.040471969 | Credible Set | purple |
| chr1:127679838 | 1 | 127.6798 | 0.037394863 | Credible Set | purple |
| chr1:127737207 | 1 | 127.7372 | 0.013012840 | Credible Set | purple |
| chr1:127737208 | 1 | 127.7372 | 0.016523559 | Credible Set | purple |
| chr1:127942547 | 1 | 127.9425 | 0.017181009 | Credible Set | purple |
| chr1:127943084 | 1 | 127.9431 | 0.017276226 | Credible Set | purple |
| chr1:127956570 | 1 | 127.9566 | 0.014120763 | Credible Set | purple |
| chr1:127988443 | 1 | 127.9884 | 0.013632031 | Credible Set | purple |
| chr1:128235181 | 1 | 128.2352 | 0.006193451 | Credible Set | purple |
| chr1:128235351 | 1 | 128.2354 | 0.006090698 | Credible Set | purple |
| chr1:128300314 | 1 | 128.3003 | 0.005360983 | Credible Set | purple |

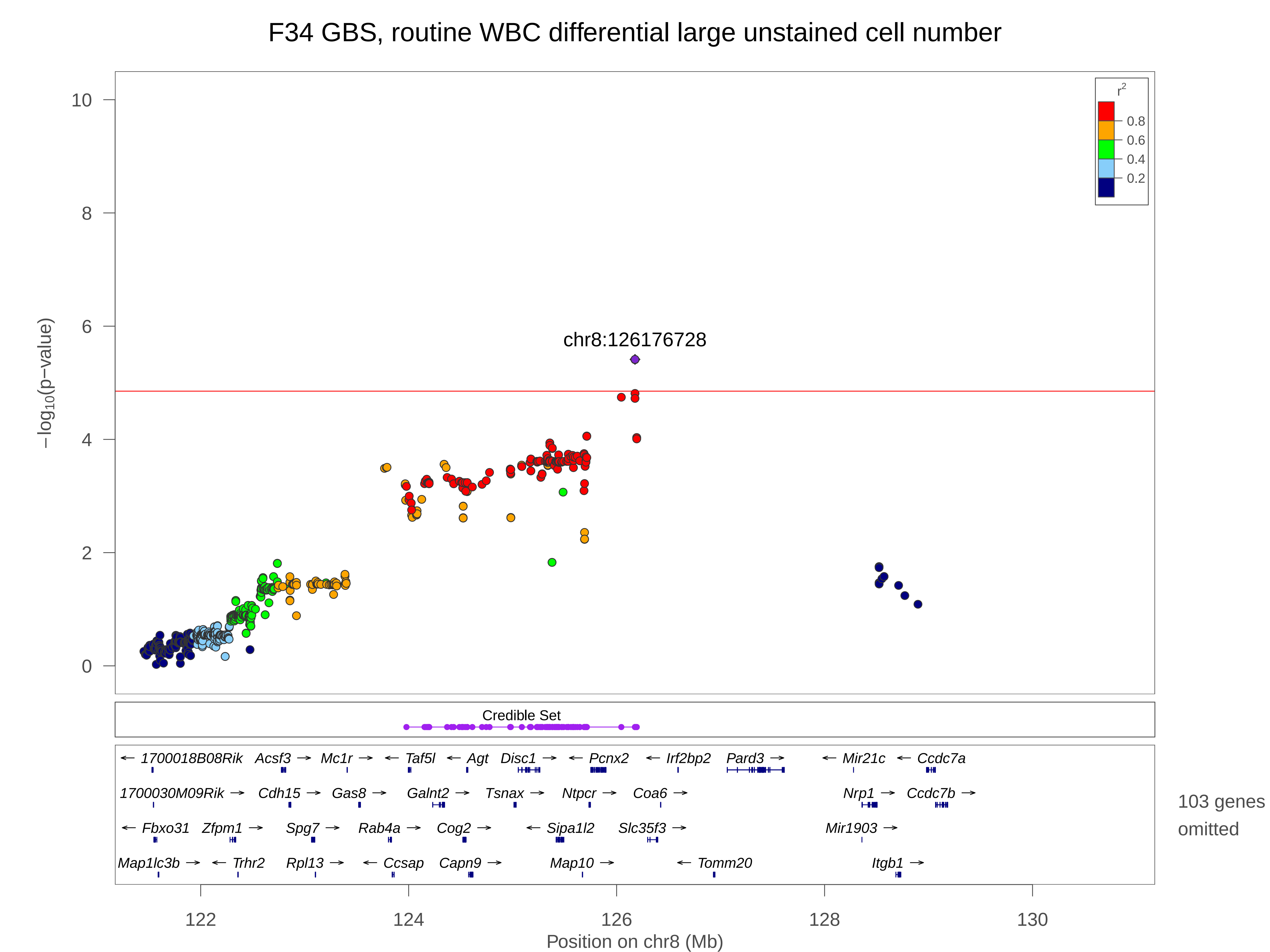
F34 GBS, routine WBC differential large unstained cell number
r2
0.8
0.6
0.4
0.2
chr8:126176728
−log10(p−value)
10
8
6
4
2
0
Credible Set
← 1700018B08Rik
Acsf3 →
Mc1r →
← Taf5l
← Agt
Disc1 →
← Pcnx2
← Irf2bp2
Pard3 →
← Mir21c
← Ccdc7a
1700030M09Rik →
Cdh15 →
Gas8 →
Galnt2 →
Tsnax →
Ntpcr →
Coa6 →
Nrp1 →
Ccdc7b →
← Fbxo31
Zfpm1 →
Spg7 →
Rab4a →
Cog2 →
← Sipa1l2
Slc35f3 →
Mir1903 →
Map1lc3b →
← Trhr2
Rpl13 →
← Ccsap
Capn9 →
Map10 →
← Tomm20
Itgb1 →
103 genes omitted
122
124
126
128
130
Position on chr8 (Mb)

date: Thu Oct 25 17:04:01 2018
build: F34GBS
display range: chr8:121176728−131176728 [121176728−131176728]
hilite range: 0 − 0 [ 0 − 0 ]
reference SNP: chr8:126176728
number of SNPs plotted: 850
min P−value: 3.86E−6 [chr8:126176728]
max P−value: 9.45E−1 [chr8:121573290]
omitted Genes: Zcchc14, Jph3, Klhdc4
omitted Genes: Slc7a5, BC048644, Car5a
omitted Genes: Banp, Mir7237, Zc3h18
omitted Genes: Il17c, Cyba, Mvd
omitted Genes: 9330133O14Rik, Gm20735, Snai3
omitted Genes: Rnf166, Ctu2, Piezo1
omitted Genes: Cdt1, Aprt, Galns
omitted Genes: Trappc2l, Pabpn1l, Mir6396
omitted Genes: Cbfa2t3, Ankrd11, 2810013P06Rik
omitted Genes: Snord68, Mir7079, Cpne7
omitted Genes: Mir7080, Sult5a1, Dpep1
omitted Genes: Chmp1a, Spata33, Cdk10
omitted Genes: Spata2l, 4933417D19Rik, Vps9d1
omitted Genes: Zfp276, Fanca, Spire2
omitted Genes: Tcf25, Tubb3, Def8
omitted Genes: Afg3l1, Dbndd1, n−R5s100
omitted Genes: n−R5s103, n−R5s104, n−R5s105
omitted Genes: n−R5s108, n−R5s110, n−R5s111
omitted Genes: n−R5s113, Gm25018, n−R5s117
omitted Genes: n−R5s121, n−R5s122, n−R5s123
omitted Genes: n−R5s124, n−R5s128, Gm26391
omitted Genes: n−R5s133, n−R5s134, Rn5s
omitted Genes: n−R5s136, n−R5s138, n−R5s139
omitted Genes: Gm22109, n−R5s141, n−R5s142
omitted Genes: n−R5s143, n−R5s144, Gm25212
omitted Genes: n−R5s146, Gm22291, Gm23284
omitted Genes: n−R5s149, Rhou, 6030466F02Rik
omitted Genes: Acta1, Nup133, Abcb10
omitted Genes: Mir1967, Urb2, Pgbd5
omitted Genes: 2310022B05Rik, Ttc13, Arv1
omitted Genes: Fam89a, Trim67, 2810004N23Rik
Make more plots at http://csg.sph.umich.edu/locuszoom/
omitted Genes: Gnpat, Exoc8, Sprtn

Fine−mapping Regions

| snp | chr | pos (Mb) | pp | group | color |
|---|---|---|---|---|---|
| chr8:123978600 | 8 | 123.9786 | 0.001528413 | Credible Set | purple |
| chr8:124152632 | 8 | 124.1526 | 0.001694062 | Credible Set | purple |
| chr8:124159766 | 8 | 124.1598 | 0.001867781 | Credible Set | purple |
| chr8:124173734 | 8 | 124.1737 | 0.002006165 | Credible Set | purple |
| chr8:124180762 | 8 | 124.1808 | 0.001746288 | Credible Set | purple |
| chr8:124189418 | 8 | 124.1894 | 0.001759202 | Credible Set | purple |
| chr8:124196154 | 8 | 124.1962 | 0.001685875 | Credible Set | purple |
| chr8:124196376 | 8 | 124.1964 | 0.001796655 | Credible Set | purple |
| chr8:124197071 | 8 | 124.1971 | 0.001759825 | Credible Set | purple |
| chr8:124197277 | 8 | 124.1973 | 0.001700089 | Credible Set | purple |
| chr8:124370035 | 8 | 124.3700 | 0.002142741 | Credible Set | purple |
| chr8:124412466 | 8 | 124.4125 | 0.002019464 | Credible Set | purple |
| chr8:124433098 | 8 | 124.4331 | 0.001685289 | Credible Set | purple |
| chr8:124487999 | 8 | 124.4880 | 0.001859850 | Credible Set | purple |
| chr8:124510054 | 8 | 124.5101 | 0.001757004 | Credible Set | purple |
| chr8:124520576 | 8 | 124.5206 | 0.001755578 | Credible Set | purple |
| chr8:124520641 | 8 | 124.5206 | 0.001765541 | Credible Set | purple |
| chr8:124543418 | 8 | 124.5434 | 0.001763569 | Credible Set | purple |
| chr8:124563037 | 8 | 124.5630 | 0.001766275 | Credible Set | purple |
| chr8:124563111 | 8 | 124.5631 | 0.001763569 | Credible Set | purple |
| chr8:124563221 | 8 | 124.5632 | 0.001763569 | Credible Set | purple |
| chr8:124612523 | 8 | 124.6125 | 0.001492897 | Credible Set | purple |
| chr8:124705504 | 8 | 124.7055 | 0.001650005 | Credible Set | purple |
| chr8:124746311 | 8 | 124.7463 | 0.001893122 | Credible Set | purple |
| chr8:124777638 | 8 | 124.7776 | 0.002595270 | Credible Set | purple |
| chr8:124977284 | 8 | 124.9773 | 0.002787824 | Credible Set | purple |
| chr8:124977290 | 8 | 124.9773 | 0.002837849 | Credible Set | purple |
| chr8:124977353 | 8 | 124.9774 | 0.002955531 | Credible Set | purple |
| chr8:124979975 | 8 | 124.9800 | 0.002529348 | Credible Set | purple |
| chr8:124982183 | 8 | 124.9822 | 0.002892122 | Credible Set | purple |
| chr8:125088462 | 8 | 125.0885 | 0.003243641 | Credible Set | purple |
| chr8:125166834 | 8 | 125.1668 | 0.003794082 | Credible Set | purple |
| chr8:125175162 | 8 | 125.1752 | 0.002742517 | Credible Set | purple |
| chr8:125175172 | 8 | 125.1752 | 0.002755058 | Credible Set | purple |
| chr8:125175263 | 8 | 125.1753 | 0.004298180 | Credible Set | purple |
| chr8:125175309 | 8 | 125.1753 | 0.004334181 | Credible Set | purple |

Fine−mapping Regions

| snp | chr | pos (Mb) | pp | group | color |
|---|---|---|---|---|---|
| chr8:125175313 | 8 | 125.1753 | 0.004323931 | Credible Set | purple |
| chr8:125232862 | 8 | 125.2329 | 0.003950738 | Credible Set | purple |
| chr8:125232967 | 8 | 125.2330 | 0.003950738 | Credible Set | purple |
| chr8:125236805 | 8 | 125.2368 | 0.003835431 | Credible Set | purple |
| chr8:125244581 | 8 | 125.2446 | 0.003950738 | Credible Set | purple |
| chr8:125261827 | 8 | 125.2618 | 0.004009503 | Credible Set | purple |
| chr8:125271155 | 8 | 125.2712 | 0.002152626 | Credible Set | purple |
| chr8:125283686 | 8 | 125.2837 | 0.002462780 | Credible Set | purple |
| chr8:125283700 | 8 | 125.2837 | 0.002427817 | Credible Set | purple |
| chr8:125313378 | 8 | 125.3134 | 0.003960228 | Credible Set | purple |
| chr8:125327590 | 8 | 125.3276 | 0.003962498 | Credible Set | purple |
| chr8:125327926 | 8 | 125.3279 | 0.003932665 | Credible Set | purple |
| chr8:125328592 | 8 | 125.3286 | 0.005002569 | Credible Set | purple |
| chr8:125337596 | 8 | 125.3376 | 0.003956013 | Credible Set | purple |
| chr8:125339832 | 8 | 125.3398 | 0.004043053 | Credible Set | purple |
| chr8:125339948 | 8 | 125.3399 | 0.004403995 | Credible Set | purple |
| chr8:125352885 | 8 | 125.3529 | 0.003950738 | Credible Set | purple |
| chr8:125357050 | 8 | 125.3571 | 0.004153951 | Credible Set | purple |
| chr8:125357070 | 8 | 125.3571 | 0.003992568 | Credible Set | purple |
| chr8:125357611 | 8 | 125.3576 | 0.007998043 | Credible Set | purple |
| chr8:125357613 | 8 | 125.3576 | 0.008011698 | Credible Set | purple |
| chr8:125357664 | 8 | 125.3577 | 0.007290025 | Credible Set | purple |
| chr8:125358000 | 8 | 125.3580 | 0.003948422 | Credible Set | purple |
| chr8:125381350 | 8 | 125.3813 | 0.003950738 | Credible Set | purple |
| chr8:125381413 | 8 | 125.3814 | 0.003983447 | Credible Set | purple |
| chr8:125381419 | 8 | 125.3814 | 0.003983447 | Credible Set | purple |
| chr8:125381501 | 8 | 125.3815 | 0.006456504 | Credible Set | purple |
| chr8:125381543 | 8 | 125.3815 | 0.006588992 | Credible Set | purple |
| chr8:125402101 | 8 | 125.4021 | 0.003438611 | Credible Set | purple |
| chr8:125402104 | 8 | 125.4021 | 0.003367583 | Credible Set | purple |
| chr8:125413718 | 8 | 125.4137 | 0.003950738 | Credible Set | purple |
| chr8:125418482 | 8 | 125.4185 | 0.003950738 | Credible Set | purple |
| chr8:125418672 | 8 | 125.4187 | 0.003950738 | Credible Set | purple |
| chr8:125418880 | 8 | 125.4189 | 0.003950738 | Credible Set | purple |
| chr8:125424267 | 8 | 125.4243 | 0.003892193 | Credible Set | purple |
| chr8:125431865 | 8 | 125.4319 | 0.002928213 | Credible Set | purple |

Fine−mapping Regions

| snp | chr | pos (Mb) | pp | group | color |
|---|---|---|---|---|---|
| chr8:125434348 | 8 | 125.4343 | 0.004035697 | Credible Set | purple |
| chr8:125434876 | 8 | 125.4349 | 0.003964326 | Credible Set | purple |
| chr8:125435327 | 8 | 125.4353 | 0.003969090 | Credible Set | purple |
| chr8:125435876 | 8 | 125.4359 | 0.003944932 | Credible Set | purple |
| chr8:125442038 | 8 | 125.4420 | 0.005071274 | Credible Set | purple |
| chr8:125442845 | 8 | 125.4428 | 0.003941202 | Credible Set | purple |
| chr8:125445748 | 8 | 125.4457 | 0.003789302 | Credible Set | purple |
| chr8:125447109 | 8 | 125.4471 | 0.003928053 | Credible Set | purple |
| chr8:125470813 | 8 | 125.4708 | 0.003830442 | Credible Set | purple |
| chr8:125475610 | 8 | 125.4756 | 0.003947334 | Credible Set | purple |
| chr8:125490914 | 8 | 125.4909 | 0.003947334 | Credible Set | purple |
| chr8:125519869 | 8 | 125.5199 | 0.003947334 | Credible Set | purple |
| chr8:125532800 | 8 | 125.5328 | 0.003950738 | Credible Set | purple |
| chr8:125532879 | 8 | 125.5329 | 0.003950738 | Credible Set | purple |
| chr8:125535705 | 8 | 125.5357 | 0.005200027 | Credible Set | purple |
| chr8:125535715 | 8 | 125.5357 | 0.005188449 | Credible Set | purple |
| chr8:125536481 | 8 | 125.5365 | 0.004393031 | Credible Set | purple |
| chr8:125560754 | 8 | 125.5608 | 0.004745899 | Credible Set | purple |
| chr8:125576647 | 8 | 125.5766 | 0.004973082 | Credible Set | purple |
| chr8:125584598 | 8 | 125.5846 | 0.003112353 | Credible Set | purple |
| chr8:125584609 | 8 | 125.5846 | 0.004086251 | Credible Set | purple |
| chr8:125585107 | 8 | 125.5851 | 0.004664570 | Credible Set | purple |
| chr8:125602298 | 8 | 125.6023 | 0.004606692 | Credible Set | purple |
| chr8:125621974 | 8 | 125.6220 | 0.004828224 | Credible Set | purple |
| chr8:125645785 | 8 | 125.6458 | 0.004040086 | Credible Set | purple |
| chr8:125645803 | 8 | 125.6458 | 0.004040086 | Credible Set | purple |
| chr8:125686823 | 8 | 125.6868 | 0.005324217 | Credible Set | purple |
| chr8:125689213 | 8 | 125.6892 | 0.005092299 | Credible Set | purple |
| chr8:125689398 | 8 | 125.6894 | 0.005096822 | Credible Set | purple |
| chr8:125691437 | 8 | 125.6914 | 0.001702796 | Credible Set | purple |
| chr8:125697075 | 8 | 125.6971 | 0.003270429 | Credible Set | purple |
| chr8:125705281 | 8 | 125.7053 | 0.004040400 | Credible Set | purple |
| chr8:125705285 | 8 | 125.7053 | 0.003828675 | Credible Set | purple |
| chr8:125705796 | 8 | 125.7058 | 0.004714132 | Credible Set | purple |
| chr8:125712995 | 8 | 125.7130 | 0.010388986 | Credible Set | purple |
| chr8:125712999 | 8 | 125.7130 | 0.010446069 | Credible Set | purple |

Fine−mapping Regions

| snp | chr | pos (Mb) | pp | group | color |
|---|---|---|---|---|---|
| chr8:125713002 | 8 | 125.7130 | 0.010312410 | Credible Set | purple |
| chr8:125713148 | 8 | 125.7131 | 0.004560446 | Credible Set | purple |
| chr8:126045228 | 8 | 126.0452 | 0.046640253 | Credible Set | purple |
| chr8:126176719 | 8 | 126.1767 | 0.200755065 | Credible Set | purple |
| chr8:126176728 | 8 | 126.1767 | 0.202885381 | Credible Set | purple |
| chr8:126176792 | 8 | 126.1768 | 0.054057454 | Credible Set | purple |
| chr8:126176848 | 8 | 126.1768 | 0.044585097 | Credible Set | purple |
| chr8:126193117 | 8 | 126.1931 | 0.009681698 | Credible Set | purple |
| chr8:126193126 | 8 | 126.1931 | 0.009858927 | Credible Set | purple |
| chr8:126193186 | 8 | 126.1932 | 0.009338563 | Credible Set | purple |

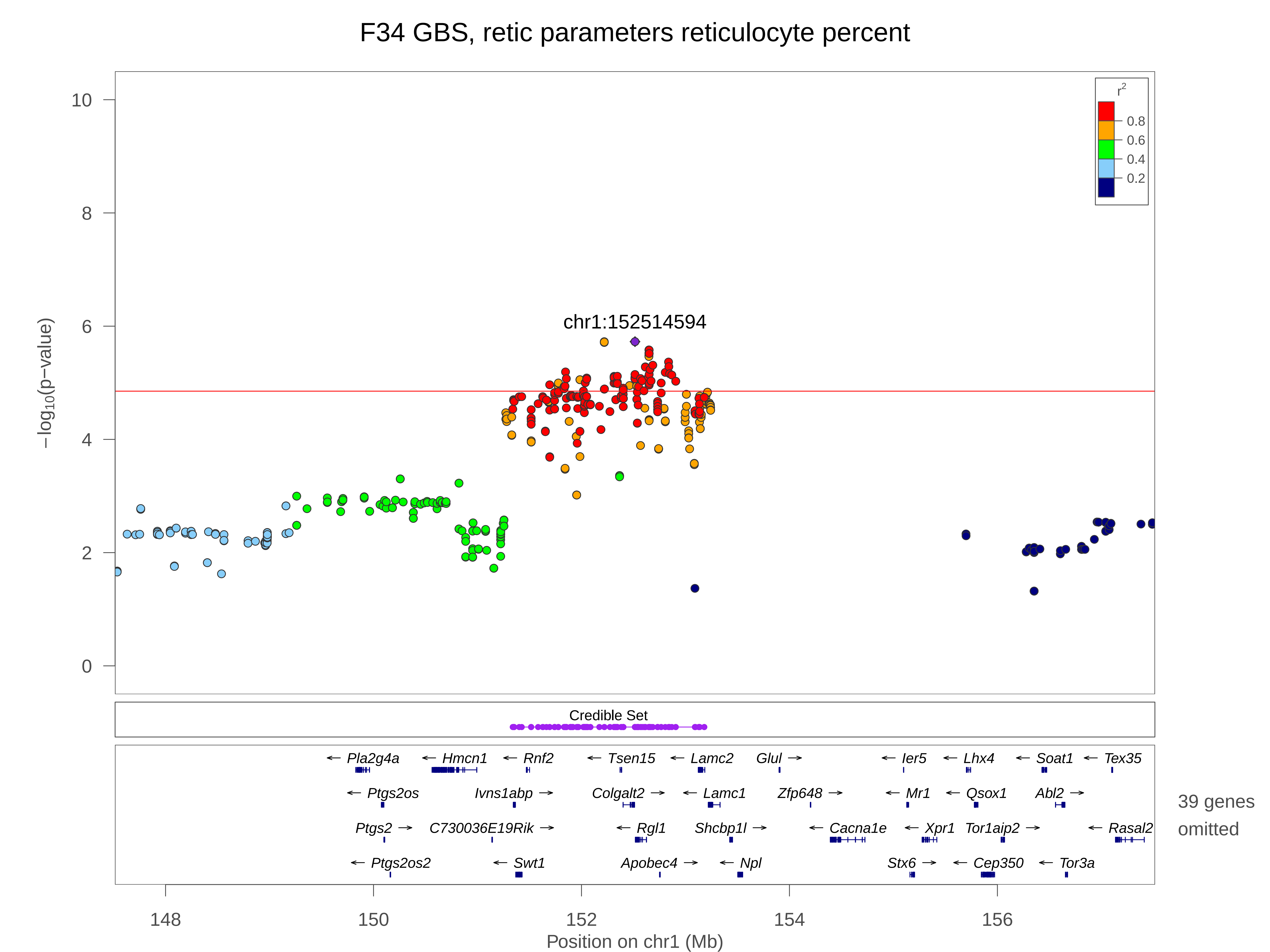
F34 GBS, retic parameters reticulocyte percent
−log10(p−value)
chr1:152514594
r2
0.8
0.6
0.4
0.2
Credible Set
Pla2g4a
Hmcn1
Rnf2
Tsen15
Lamc2
Glul
Ier5
Lhx4
Soat1
Tex35
Ptgs2os
Ivns1abp
Colgalt2
Lamc1
Zfp648
Mr1
Qsox1
Abl2
Ptgs2
C730036E19Rik
Rgl1
Shcbp1l
Cacna1e
Xpr1
Tor1aip2
Rasal2
Ptgs2os2
Swt1
Apobec4
Npl
Stx6
Cep350
Tor3a
39 genes omitted
Position on chr1 (Mb)

| | |
|---|---|
| date: | Thu Oct 25 17:04:13 2018 |
| build: | F34GBS |
| display range: | chr1:147514594−157514594 [147514594−157514594] |
| hilite range: | 0 − 0 [ 0 − 0 ] |
| reference SNP: | chr1:152514594 |
| number of SNPs plotted: | 499 |
| min P−value: | 1.88E−6 [chr1:152514594] |
| max P−value: | 4.8E−2 [chr1:156353025] |
| omitted Genes: | Pdc, 2310030A07Rik, BC003331 |
| omitted Genes: | Tpr, Prg4, Trmt1l |
| omitted Genes: | Mir7682, Fam129a, Edem3 |
| omitted Genes: | 1700025G04Rik, Arpc5, Ncf2 |
| omitted Genes: | Smg7, Nmnat2, E330020D12Rik |
| omitted Genes: | Dhx9, Rgs8, C230024C17Rik |
| omitted Genes: | Rgs16, Rnasel, Rgsl1 |
| omitted Genes: | Teddm2, Teddm1b, Teddm1a |
| omitted Genes: | Mir8114, 4930452N14Rik, 4930532M18Rik |
| omitted Genes: | Gm5532, Acbd6, Tor1aip1 |
| omitted Genes: | Fam163a, Tdrd5, Nphs2 |
| omitted Genes: | Fam20b, Ralgps2, Angptl1 |
| omitted Genes: | 2810025M15Rik, BC026585, Sec16b |

Make more plots at http://csg.sph.umich.edu/locuszoom/

Fine−mapping Regions

| snp | chr | pos (Mb) | pp | group | color |
|---|---|---|---|---|---|
| chr1:151339098 | 1 | 151.3391 | 0.002896438 | Credible Set | purple |
| chr1:151346919 | 1 | 151.3469 | 0.004173767 | Credible Set | purple |
| chr1:151346950 | 1 | 151.3469 | 0.004050923 | Credible Set | purple |
| chr1:151351933 | 1 | 151.3519 | 0.003903870 | Credible Set | purple |
| chr1:151352397 | 1 | 151.3524 | 0.003897974 | Credible Set | purple |
| chr1:151400733 | 1 | 151.4007 | 0.004666141 | Credible Set | purple |
| chr1:151424994 | 1 | 151.4250 | 0.004691589 | Credible Set | purple |
| chr1:151515546 | 1 | 151.5155 | 0.001985201 | Credible Set | purple |
| chr1:151515567 | 1 | 151.5156 | 0.002067244 | Credible Set | purple |
| chr1:151516022 | 1 | 151.5160 | 0.001823672 | Credible Set | purple |
| chr1:151516157 | 1 | 151.5162 | 0.002830805 | Credible Set | purple |
| chr1:151582944 | 1 | 151.5829 | 0.003570811 | Credible Set | purple |
| chr1:151626734 | 1 | 151.6267 | 0.004691589 | Credible Set | purple |
| chr1:151630091 | 1 | 151.6301 | 0.004466706 | Credible Set | purple |
| chr1:151663201 | 1 | 151.6632 | 0.004119639 | Credible Set | purple |
| chr1:151693985 | 1 | 151.6940 | 0.002783331 | Credible Set | purple |
| chr1:151694036 | 1 | 151.6940 | 0.007437689 | Credible Set | purple |
| chr1:151741376 | 1 | 151.7414 | 0.004027916 | Credible Set | purple |
| chr1:151741404 | 1 | 151.7414 | 0.002907849 | Credible Set | purple |
| chr1:151741409 | 1 | 151.7414 | 0.002907849 | Credible Set | purple |
| chr1:151741410 | 1 | 151.7414 | 0.002902968 | Credible Set | purple |
| chr1:151741413 | 1 | 151.7414 | 0.002907849 | Credible Set | purple |
| chr1:151741477 | 1 | 151.7415 | 0.005100188 | Credible Set | purple |
| chr1:151741478 | 1 | 151.7415 | 0.005491751 | Credible Set | purple |
| chr1:151777225 | 1 | 151.7772 | 0.005626348 | Credible Set | purple |
| chr1:151777722 | 1 | 151.7777 | 0.005341084 | Credible Set | purple |
| chr1:151777765 | 1 | 151.7778 | 0.005657913 | Credible Set | purple |
| chr1:151832714 | 1 | 151.8327 | 0.006949675 | Credible Set | purple |
| chr1:151832824 | 1 | 151.8328 | 0.006358324 | Credible Set | purple |
| chr1:151841655 | 1 | 151.8417 | 0.007041982 | Credible Set | purple |
| chr1:151846455 | 1 | 151.8465 | 0.012261176 | Credible Set | purple |
| chr1:151854124 | 1 | 151.8541 | 0.009435657 | Credible Set | purple |
| chr1:151854136 | 1 | 151.8541 | 0.003033152 | Credible Set | purple |
| chr1:151854478 | 1 | 151.8545 | 0.004411419 | Credible Set | purple |
| chr1:151854533 | 1 | 151.8545 | 0.004394856 | Credible Set | purple |
| chr1:151893115 | 1 | 151.8931 | 0.004691589 | Credible Set | purple |

Fine−mapping Regions

| snp | chr | pos (Mb) | pp | group | color |
|---|---|---|---|---|---|
| chr1:151893154 | 1 | 151.8932 | 0.004691589 | Credible Set | purple |
| chr1:151893169 | 1 | 151.8932 | 0.004876621 | Credible Set | purple |
| chr1:151896769 | 1 | 151.8968 | 0.004891800 | Credible Set | purple |
| chr1:151907336 | 1 | 151.9073 | 0.004865930 | Credible Set | purple |
| chr1:151916998 | 1 | 151.9170 | 0.004691589 | Credible Set | purple |
| chr1:151917004 | 1 | 151.9170 | 0.004691589 | Credible Set | purple |
| chr1:151953861 | 1 | 151.9539 | 0.004720731 | Credible Set | purple |
| chr1:151963849 | 1 | 151.9638 | 0.002965971 | Credible Set | purple |
| chr1:151964084 | 1 | 151.9641 | 0.004551512 | Credible Set | purple |
| chr1:151971418 | 1 | 151.9714 | 0.004691589 | Credible Set | purple |
| chr1:152018151 | 1 | 152.0182 | 0.004898524 | Credible Set | purple |
| chr1:152018723 | 1 | 152.0187 | 0.005853185 | Credible Set | purple |
| chr1:152019427 | 1 | 152.0194 | 0.005074529 | Credible Set | purple |
| chr1:152027497 | 1 | 152.0275 | 0.002520202 | Credible Set | purple |
| chr1:152027513 | 1 | 152.0275 | 0.003226878 | Credible Set | purple |
| chr1:152027772 | 1 | 152.0278 | 0.003402495 | Credible Set | purple |
| chr1:152030727 | 1 | 152.0307 | 0.004691589 | Credible Set | purple |
| chr1:152030740 | 1 | 152.0307 | 0.004691589 | Credible Set | purple |
| chr1:152030750 | 1 | 152.0308 | 0.004691589 | Credible Set | purple |
| chr1:152030755 | 1 | 152.0308 | 0.004691589 | Credible Set | purple |
| chr1:152030782 | 1 | 152.0308 | 0.003791684 | Credible Set | purple |
| chr1:152035922 | 1 | 152.0359 | 0.008054665 | Credible Set | purple |
| chr1:152035925 | 1 | 152.0359 | 0.008056079 | Credible Set | purple |
| chr1:152047326 | 1 | 152.0473 | 0.004699510 | Credible Set | purple |
| chr1:152049281 | 1 | 152.0493 | 0.004747493 | Credible Set | purple |
| chr1:152049576 | 1 | 152.0496 | 0.004709191 | Credible Set | purple |
| chr1:152050080 | 1 | 152.0501 | 0.009673176 | Credible Set | purple |
| chr1:152050132 | 1 | 152.0501 | 0.009271428 | Credible Set | purple |
| chr1:152060153 | 1 | 152.0602 | 0.003487767 | Credible Set | purple |
| chr1:152060186 | 1 | 152.0602 | 0.003468602 | Credible Set | purple |
| chr1:152085317 | 1 | 152.0853 | 0.003414125 | Credible Set | purple |
| chr1:152085323 | 1 | 152.0853 | 0.003507709 | Credible Set | purple |
| chr1:152085887 | 1 | 152.0859 | 0.003446032 | Credible Set | purple |
| chr1:152172042 | 1 | 152.1720 | 0.003232799 | Credible Set | purple |
| chr1:152219150 | 1 | 152.2192 | 0.006258417 | Credible Set | purple |
| chr1:152221047 | 1 | 152.2210 | 0.006330232 | Credible Set | purple |

Fine−mapping Regions

| snp | chr | pos (Mb) | pp | group | color |
|---|---|---|---|---|---|
| chr1:152273304 | 1 | 152.2733 | 0.002635169 | Credible Set | purple |
| chr1:152312984 | 1 | 152.3130 | 0.007935417 | Credible Set | purple |
| chr1:152313303 | 1 | 152.3133 | 0.010321796 | Credible Set | purple |
| chr1:152313315 | 1 | 152.3133 | 0.009863235 | Credible Set | purple |
| chr1:152313427 | 1 | 152.3134 | 0.010165857 | Credible Set | purple |
| chr1:152313450 | 1 | 152.3134 | 0.009950365 | Credible Set | purple |
| chr1:152313626 | 1 | 152.3136 | 0.009508917 | Credible Set | purple |
| chr1:152313656 | 1 | 152.3137 | 0.009863648 | Credible Set | purple |
| chr1:152329709 | 1 | 152.3297 | 0.004194085 | Credible Set | purple |
| chr1:152329711 | 1 | 152.3297 | 0.004150873 | Credible Set | purple |
| chr1:152330012 | 1 | 152.3300 | 0.007932320 | Credible Set | purple |
| chr1:152344680 | 1 | 152.3447 | 0.008269548 | Credible Set | purple |
| chr1:152344704 | 1 | 152.3447 | 0.010358700 | Credible Set | purple |
| chr1:152345178 | 1 | 152.3452 | 0.007767833 | Credible Set | purple |
| chr1:152382093 | 1 | 152.3821 | 0.004450088 | Credible Set | purple |
| chr1:152382197 | 1 | 152.3822 | 0.004841214 | Credible Set | purple |
| chr1:152400891 | 1 | 152.4009 | 0.005853289 | Credible Set | purple |
| chr1:152401044 | 1 | 152.4010 | 0.005123119 | Credible Set | purple |
| chr1:152401140 | 1 | 152.4011 | 0.006527809 | Credible Set | purple |
| chr1:152401179 | 1 | 152.4012 | 0.006118288 | Credible Set | purple |
| chr1:152401759 | 1 | 152.4018 | 0.003179731 | Credible Set | purple |
| chr1:152402888 | 1 | 152.4029 | 0.004367868 | Credible Set | purple |
| chr1:152402907 | 1 | 152.4029 | 0.004367868 | Credible Set | purple |
| chr1:152402941 | 1 | 152.4029 | 0.004367868 | Credible Set | purple |
| chr1:152513693 | 1 | 152.5137 | 0.009901076 | Credible Set | purple |
| chr1:152513781 | 1 | 152.5138 | 0.009373920 | Credible Set | purple |
| chr1:152514271 | 1 | 152.5143 | 0.009403894 | Credible Set | purple |
| chr1:152514375 | 1 | 152.5144 | 0.011093062 | Credible Set | purple |
| chr1:152514594 | 1 | 152.5146 | 0.039929574 | Credible Set | purple |
| chr1:152514647 | 1 | 152.5146 | 0.039832481 | Credible Set | purple |
| chr1:152531770 | 1 | 152.5318 | 0.004235411 | Credible Set | purple |
| chr1:152536828 | 1 | 152.5368 | 0.001677710 | Credible Set | purple |
| chr1:152536849 | 1 | 152.5368 | 0.001694258 | Credible Set | purple |
| chr1:152536958 | 1 | 152.5370 | 0.005604155 | Credible Set | purple |
| chr1:152545294 | 1 | 152.5453 | 0.003390846 | Credible Set | purple |
| chr1:152545312 | 1 | 152.5453 | 0.003390846 | Credible Set | purple |

Fine−mapping Regions

| snp | chr | pos (Mb) | pp | group | color |
|---|---|---|---|---|---|
| chr1:152548725 | 1 | 152.5487 | 0.006900719 | Credible Set | purple |
| chr1:152568046 | 1 | 152.5680 | 0.009478547 | Credible Set | purple |
| chr1:152582751 | 1 | 152.5828 | 0.008722077 | Credible Set | purple |
| chr1:152600723 | 1 | 152.6007 | 0.005880718 | Credible Set | purple |
| chr1:152614121 | 1 | 152.6141 | 0.014948522 | Credible Set | purple |
| chr1:152614133 | 1 | 152.6141 | 0.014797893 | Credible Set | purple |
| chr1:152644575 | 1 | 152.6446 | 0.010114633 | Credible Set | purple |
| chr1:152650776 | 1 | 152.6508 | 0.007548897 | Credible Set | purple |
| chr1:152650892 | 1 | 152.6509 | 0.007345767 | Credible Set | purple |
| chr1:152650962 | 1 | 152.6510 | 0.011127781 | Credible Set | purple |
| chr1:152650976 | 1 | 152.6510 | 0.028765589 | Credible Set | purple |
| chr1:152651016 | 1 | 152.6510 | 0.025133576 | Credible Set | purple |
| chr1:152651777 | 1 | 152.6518 | 0.007821710 | Credible Set | purple |
| chr1:152659220 | 1 | 152.6592 | 0.013518611 | Credible Set | purple |
| chr1:152669121 | 1 | 152.6691 | 0.008623833 | Credible Set | purple |
| chr1:152685949 | 1 | 152.6859 | 0.015942450 | Credible Set | purple |
| chr1:152731898 | 1 | 152.7319 | 0.003892579 | Credible Set | purple |
| chr1:152731899 | 1 | 152.7319 | 0.003604457 | Credible Set | purple |
| chr1:152731910 | 1 | 152.7319 | 0.003205477 | Credible Set | purple |
| chr1:152732468 | 1 | 152.7325 | 0.002820924 | Credible Set | purple |
| chr1:152732477 | 1 | 152.7325 | 0.002848003 | Credible Set | purple |
| chr1:152732522 | 1 | 152.7325 | 0.002607363 | Credible Set | purple |
| chr1:152766369 | 1 | 152.7664 | 0.007971496 | Credible Set | purple |
| chr1:152766418 | 1 | 152.7664 | 0.005413035 | Credible Set | purple |
| chr1:152805855 | 1 | 152.8059 | 0.012036101 | Credible Set | purple |
| chr1:152837203 | 1 | 152.8372 | 0.018034009 | Credible Set | purple |
| chr1:152840027 | 1 | 152.8400 | 0.015050244 | Credible Set | purple |
| chr1:152847662 | 1 | 152.8477 | 0.011433616 | Credible Set | purple |
| chr1:152868708 | 1 | 152.8687 | 0.010864872 | Credible Set | purple |
| chr1:152906391 | 1 | 152.9064 | 0.008534767 | Credible Set | purple |
| chr1:152906398 | 1 | 152.9064 | 0.008530725 | Credible Set | purple |
| chr1:153091176 | 1 | 153.0912 | 0.002688364 | Credible Set | purple |
| chr1:153091194 | 1 | 153.0912 | 0.002722811 | Credible Set | purple |
| chr1:153096224 | 1 | 153.0962 | 0.002384013 | Credible Set | purple |
| chr1:153128621 | 1 | 153.1286 | 0.002474947 | Credible Set | purple |
| chr1:153129186 | 1 | 153.1292 | 0.002387507 | Credible Set | purple |

Fine−mapping Regions

| snp | chr | pos (Mb) | pp | group | color |
|---|---|---|---|---|---|
| chr1:153129276 | 1 | 153.1293 | 0.004400066 | Credible Set | purple |
| chr1:153134434 | 1 | 153.1344 | 0.002371447 | Credible Set | purple |
| chr1:153134521 | 1 | 153.1345 | 0.003527590 | Credible Set | purple |
| chr1:153136723 | 1 | 153.1367 | 0.002653707 | Credible Set | purple |
| chr1:153180646 | 1 | 153.1806 | 0.004571288 | Credible Set | purple |
| chr1:153180739 | 1 | 153.1807 | 0.004551653 | Credible Set | purple |

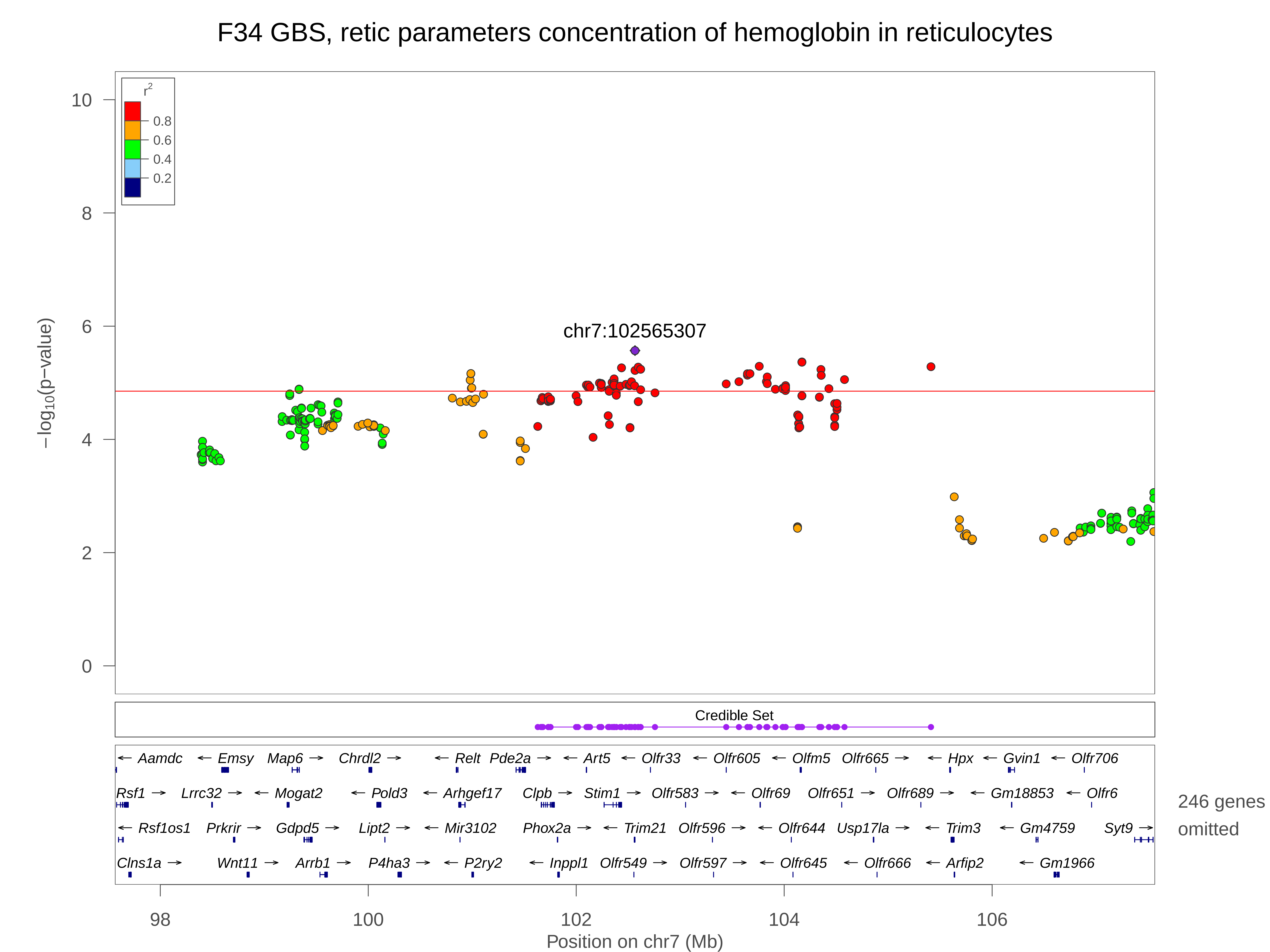

F34 GBS, retic parameters concentration of hemoglobin in reticulocytes
r2
0.8
0.6
0.4
0.2
−log10(p−value)
chr7:102565307
Credible Set
← Aamdc ← Emsy Map6 → Chrdl2 → ← Relt Pde2a → ← Art5 ← Olfr33 ← Olfr605 ← Olfm5 Olfr665 → ← Hpx ← Gvin1 ← Olfr706
Rsf1 → Lrrc32 → ← Mogat2 ← Pold3 ← Arhgef17 Clpb → Stim1 → Olfr583 → ← Olfr69 Olfr651 → Olfr689 → ← Gm18853 ← Olfr6
← Rsf1os1 Prkrir → Gdpd5 → Lipt2 → ← Mir3102 Phox2a → ← Trim21 Olfr596 → ← Olfr644 Usp17la → ← Trim3 ← Gm4759 Syt9 →
Clns1a → Wnt11 → Arrb1 → P4ha3 → ← P2ry2 ← Inppl1 Olfr549 → Olfr597 → ← Olfr645 ← Olfr666 ← Arfip2 ← Gm1966
246 genes omitted
Position on chr7 (Mb)

date: Thu Oct 25 17:04:37 2018
build: F34GBS
display range: chr7:97565307−107565307 [97565307−107565307]
hilite range: 0 − 0 [ 0 − 0 ]
reference SNP: chr7:102565307
number of SNPs plotted: 328
min P−value: 2.7E−6 [chr7:102565307]
max P−value: 6.35E−3 [chr7:107333153]
omitted Genes: Aqp11, Pak1, Gdpd4
omitted Genes: Myo7a, Capn5, Omp
omitted Genes: Gm16938, B3gnt6, Acer3
omitted Genes: Tsku, Gucy2d, Uvrag
omitted Genes: Dgat2, Gm26705, Serpinh1
omitted Genes: Klhl35, Rps3, Snord15b
omitted Genes: Snord15a, Mir326, Tpbgl
omitted Genes: Slco2b1, Olfr520, Olfr521
omitted Genes: Neu3, Spcs2, Xrra1
omitted Genes: Rnf169, Kcne3, Pgm2l1
omitted Genes: Gpx2−ps1, Ppme1, C2cd3
omitted Genes: Ucp3, Ucp2, Dnajb13
omitted Genes: Coa4, Mrpl48, Rab6a
omitted Genes: Plekhb1, Fam168a, P2ry6
omitted Genes: Fchsd2, Atg16l2, Stard10
omitted Genes: Arap1, Mir139, Art2a−ps
omitted Genes: Art2b, Folr2, Folr1
omitted Genes: Anapc15, Tomt, Lamtor1
omitted Genes: Lrrc51, Numa1, Il18bp
omitted Genes: Rnf121, Xndc1, Xntrpc
omitted Genes: Trpc2, Art1, Chrna10
omitted Genes: Nup98, Pgap2, Rhog
omitted Genes: Rrm1, Olfr543, Olfr544
omitted Genes: Olfr545, Olfr547, Olfr550
omitted Genes: Olfr551, Olfr552, Olfr553
omitted Genes: Olfr554, Olfr555, Olfr556
omitted Genes: Trim68, Olfr557, Olfr558
omitted Genes: Olfr559, Olfr78, Olfr560
omitted Genes: Olfr561, Olfr564, Olfr566
omitted Genes: Olfr568, Olfr569, Olfr570
omitted Genes: Olfr571, Olfr572, Olfr574
Make more plots at http://csg.sph.umich.edu/locuszoom/
omitted Genes: Olfr575, Olfr576, Olfr577

Fine−mapping Regions

| snp | chr | pos (Mb) | pp | group | color |
|---|---|---|---|---|---|
| chr7:101630176 | 7 | 101.6302 | 0.001666218 | Credible Set | purple |
| chr7:101660383 | 7 | 101.6604 | 0.004478869 | Credible Set | purple |
| chr7:101669186 | 7 | 101.6692 | 0.004685483 | Credible Set | purple |
| chr7:101673483 | 7 | 101.6735 | 0.005106436 | Credible Set | purple |
| chr7:101680140 | 7 | 101.6801 | 0.004873251 | Credible Set | purple |
| chr7:101731061 | 7 | 101.7311 | 0.004366629 | Credible Set | purple |
| chr7:101731110 | 7 | 101.7311 | 0.004834470 | Credible Set | purple |
| chr7:101731130 | 7 | 101.7311 | 0.005250993 | Credible Set | purple |
| chr7:101731658 | 7 | 101.7317 | 0.004500023 | Credible Set | purple |
| chr7:101751790 | 7 | 101.7518 | 0.004464519 | Credible Set | purple |
| chr7:101752580 | 7 | 101.7526 | 0.004669091 | Credible Set | purple |
| chr7:101752971 | 7 | 101.7530 | 0.004775575 | Credible Set | purple |
| chr7:101998679 | 7 | 101.9987 | 0.005446458 | Credible Set | purple |
| chr7:102016307 | 7 | 102.0163 | 0.004346337 | Credible Set | purple |
| chr7:102098054 | 7 | 102.0981 | 0.008271975 | Credible Set | purple |
| chr7:102110109 | 7 | 102.1101 | 0.007713600 | Credible Set | purple |
| chr7:102110501 | 7 | 102.1105 | 0.007717741 | Credible Set | purple |
| chr7:102116318 | 7 | 102.1163 | 0.008094142 | Credible Set | purple |
| chr7:102116430 | 7 | 102.1164 | 0.008094142 | Credible Set | purple |
| chr7:102116478 | 7 | 102.1165 | 0.008281684 | Credible Set | purple |
| chr7:102132562 | 7 | 102.1326 | 0.007649262 | Credible Set | purple |
| chr7:102223396 | 7 | 102.2234 | 0.008932092 | Credible Set | purple |
| chr7:102240695 | 7 | 102.2407 | 0.007499838 | Credible Set | purple |
| chr7:102241191 | 7 | 102.2412 | 0.008790987 | Credible Set | purple |
| chr7:102241211 | 7 | 102.2412 | 0.008585367 | Credible Set | purple |
| chr7:102241212 | 7 | 102.2412 | 0.008450446 | Credible Set | purple |
| chr7:102306968 | 7 | 102.3070 | 0.002505706 | Credible Set | purple |
| chr7:102306978 | 7 | 102.3070 | 0.002523243 | Credible Set | purple |
| chr7:102317213 | 7 | 102.3172 | 0.006912562 | Credible Set | purple |
| chr7:102317246 | 7 | 102.3172 | 0.006473017 | Credible Set | purple |
| chr7:102319586 | 7 | 102.3196 | 0.001794277 | Credible Set | purple |
| chr7:102344970 | 7 | 102.3450 | 0.009168286 | Credible Set | purple |
| chr7:102363897 | 7 | 102.3639 | 0.008006639 | Credible Set | purple |
| chr7:102364204 | 7 | 102.3642 | 0.009810287 | Credible Set | purple |
| chr7:102364223 | 7 | 102.3642 | 0.010503286 | Credible Set | purple |
| chr7:102364315 | 7 | 102.3643 | 0.008789794 | Credible Set | purple |

Fine−mapping Regions

| snp | chr | pos (Mb) | pp | group | color |
|---|---|---|---|---|---|
| chr7:102364326 | 7 | 102.3643 | 0.008789794 | Credible Set | purple |
| chr7:102364411 | 7 | 102.3644 | 0.008263564 | Credible Set | purple |
| chr7:102385520 | 7 | 102.3855 | 0.006047459 | Credible Set | purple |
| chr7:102385536 | 7 | 102.3855 | 0.005554218 | Credible Set | purple |
| chr7:102423165 | 7 | 102.4232 | 0.007919867 | Credible Set | purple |
| chr7:102435692 | 7 | 102.4357 | 0.016139724 | Credible Set | purple |
| chr7:102435720 | 7 | 102.4357 | 0.016139724 | Credible Set | purple |
| chr7:102478469 | 7 | 102.4785 | 0.008488158 | Credible Set | purple |
| chr7:102510475 | 7 | 102.5105 | 0.008124566 | Credible Set | purple |
| chr7:102510607 | 7 | 102.5106 | 0.008083571 | Credible Set | purple |
| chr7:102516878 | 7 | 102.5169 | 0.008314005 | Credible Set | purple |
| chr7:102531652 | 7 | 102.5317 | 0.009387117 | Credible Set | purple |
| chr7:102561708 | 7 | 102.5617 | 0.008035262 | Credible Set | purple |
| chr7:102565262 | 7 | 102.5653 | 0.031563442 | Credible Set | purple |
| chr7:102565292 | 7 | 102.5653 | 0.031472204 | Credible Set | purple |
| chr7:102565307 | 7 | 102.5653 | 0.031602621 | Credible Set | purple |
| chr7:102565323 | 7 | 102.5653 | 0.031511275 | Credible Set | purple |
| chr7:102565329 | 7 | 102.5653 | 0.031511275 | Credible Set | purple |
| chr7:102565410 | 7 | 102.5654 | 0.014576406 | Credible Set | purple |
| chr7:102596442 | 7 | 102.5964 | 0.016595746 | Credible Set | purple |
| chr7:102596578 | 7 | 102.5966 | 0.004329223 | Credible Set | purple |
| chr7:102596636 | 7 | 102.5966 | 0.004357565 | Credible Set | purple |
| chr7:102619418 | 7 | 102.6194 | 0.015290632 | Credible Set | purple |
| chr7:102619431 | 7 | 102.6194 | 0.015253688 | Credible Set | purple |
| chr7:102619466 | 7 | 102.6195 | 0.006888519 | Credible Set | purple |
| chr7:102757499 | 7 | 102.7575 | 0.006102586 | Credible Set | purple |
| chr7:103442570 | 7 | 103.4426 | 0.008636898 | Credible Set | purple |
| chr7:103564836 | 7 | 103.5648 | 0.009421142 | Credible Set | purple |
| chr7:103646439 | 7 | 103.6464 | 0.012111876 | Credible Set | purple |
| chr7:103646709 | 7 | 103.6467 | 0.012830230 | Credible Set | purple |
| chr7:103671033 | 7 | 103.6710 | 0.012844440 | Credible Set | purple |
| chr7:103760301 | 7 | 103.7603 | 0.017133624 | Credible Set | purple |
| chr7:103829200 | 7 | 103.8292 | 0.009447507 | Credible Set | purple |
| chr7:103837099 | 7 | 103.8371 | 0.011322627 | Credible Set | purple |
| chr7:103837213 | 7 | 103.8372 | 0.008715849 | Credible Set | purple |
| chr7:103915503 | 7 | 103.9155 | 0.007023448 | Credible Set | purple |

Fine−mapping Regions

| snp | chr | pos (Mb) | pp | group | color |
|---|---|---|---|---|---|
| chr7:103986792 | 7 | 103.9868 | 0.007285178 | Credible Set | purple |
| chr7:103986795 | 7 | 103.9868 | 0.007285178 | Credible Set | purple |
| chr7:103986857 | 7 | 103.9869 | 0.007129979 | Credible Set | purple |
| chr7:103986860 | 7 | 103.9869 | 0.007129979 | Credible Set | purple |
| chr7:104012808 | 7 | 104.0128 | 0.006678759 | Credible Set | purple |
| chr7:104013370 | 7 | 104.0134 | 0.007235938 | Credible Set | purple |
| chr7:104013395 | 7 | 104.0134 | 0.008094198 | Credible Set | purple |
| chr7:104013467 | 7 | 104.0135 | 0.007587979 | Credible Set | purple |
| chr7:104129734 | 7 | 104.1297 | 0.002594766 | Credible Set | purple |
| chr7:104129736 | 7 | 104.1297 | 0.002594766 | Credible Set | purple |
| chr7:104138934 | 7 | 104.1389 | 0.001858189 | Credible Set | purple |
| chr7:104138936 | 7 | 104.1389 | 0.001865475 | Credible Set | purple |
| chr7:104138968 | 7 | 104.1390 | 0.001865475 | Credible Set | purple |
| chr7:104141705 | 7 | 104.1417 | 0.002378657 | Credible Set | purple |
| chr7:104141768 | 7 | 104.1418 | 0.002450029 | Credible Set | purple |
| chr7:104149893 | 7 | 104.1499 | 0.001648049 | Credible Set | purple |
| chr7:104170479 | 7 | 104.1705 | 0.020160451 | Credible Set | purple |
| chr7:104170506 | 7 | 104.1705 | 0.020225329 | Credible Set | purple |
| chr7:104170518 | 7 | 104.1705 | 0.020295584 | Credible Set | purple |
| chr7:104170522 | 7 | 104.1705 | 0.020184594 | Credible Set | purple |
| chr7:104170543 | 7 | 104.1705 | 0.020080050 | Credible Set | purple |
| chr7:104170588 | 7 | 104.1706 | 0.005423359 | Credible Set | purple |
| chr7:104170594 | 7 | 104.1706 | 0.005434988 | Credible Set | purple |
| chr7:104170618 | 7 | 104.1706 | 0.005458156 | Credible Set | purple |
| chr7:104170662 | 7 | 104.1707 | 0.005458156 | Credible Set | purple |
| chr7:104338784 | 7 | 104.3388 | 0.005132344 | Credible Set | purple |
| chr7:104338810 | 7 | 104.3388 | 0.005188055 | Credible Set | purple |
| chr7:104353491 | 7 | 104.3535 | 0.015143062 | Credible Set | purple |
| chr7:104356230 | 7 | 104.3562 | 0.012025411 | Credible Set | purple |
| chr7:104430205 | 7 | 104.4302 | 0.007189441 | Credible Set | purple |
| chr7:104485236 | 7 | 104.4852 | 0.004012765 | Credible Set | purple |
| chr7:104485721 | 7 | 104.4857 | 0.001742260 | Credible Set | purple |
| chr7:104485727 | 7 | 104.4857 | 0.001658591 | Credible Set | purple |
| chr7:104486022 | 7 | 104.4860 | 0.002426149 | Credible Set | purple |
| chr7:104486041 | 7 | 104.4860 | 0.002328465 | Credible Set | purple |
| chr7:104507724 | 7 | 104.5077 | 0.003180216 | Credible Set | purple |

Fine−mapping Regions

| snp | chr | pos (Mb) | pp | group | color |
|---|---|---|---|---|---|
| chr7:104508127 | 7 | 104.5081 | 0.003511477 | Credible Set | purple |
| chr7:104508173 | 7 | 104.5082 | 0.004054181 | Credible Set | purple |
| chr7:104580091 | 7 | 104.5801 | 0.010219445 | Credible Set | purple |
| chr7:105411752 | 7 | 105.4118 | 0.016860666 | Credible Set | purple |

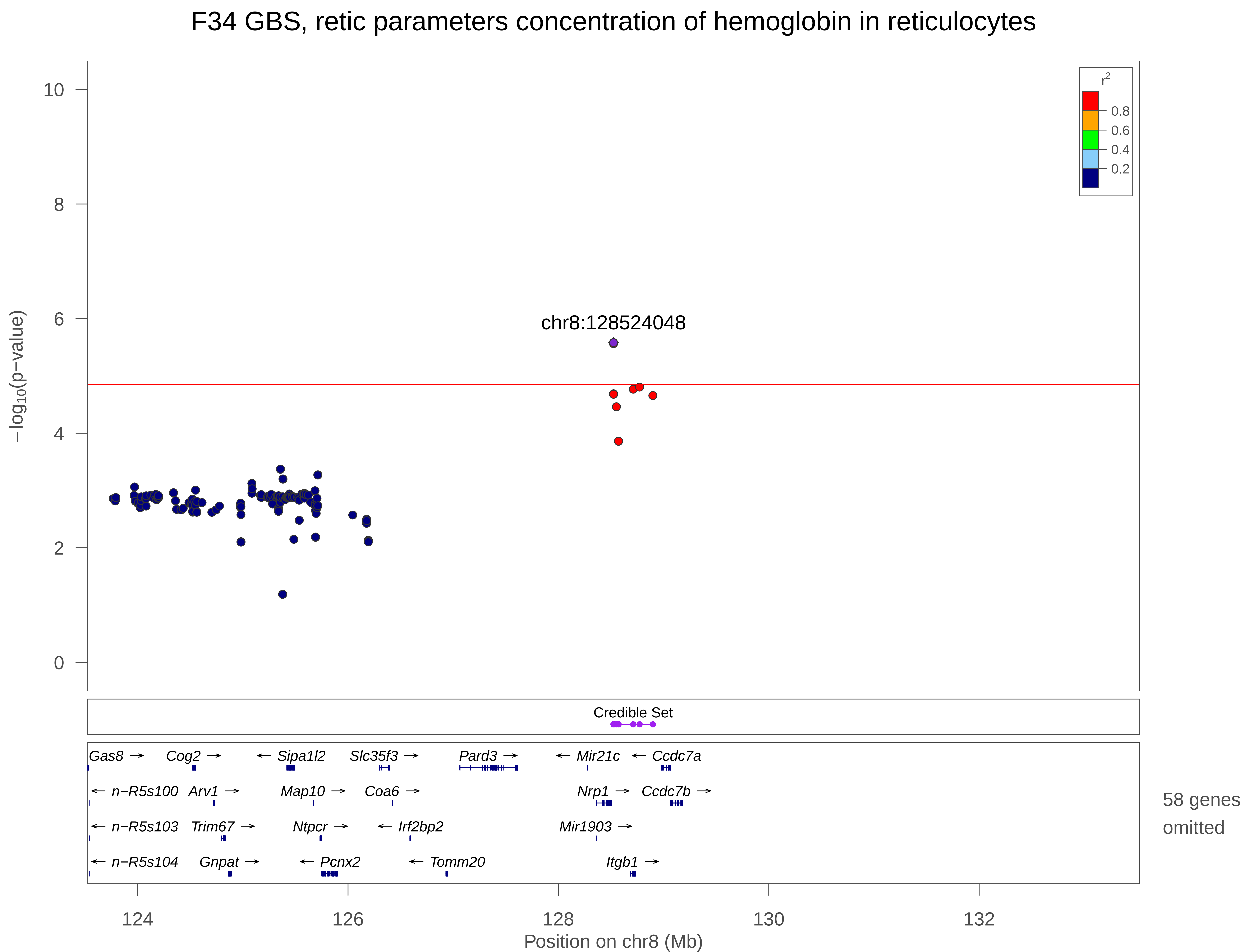

F34 GBS, retic parameters concentration of hemoglobin in reticulocytes
−log10(p−value)
10
8
6
4
2
0
chr8:128524048
r2
0.8
0.6
0.4
0.2
Credible Set
Gas8
Cog2
Sipa1l2
Slc35f3
Pard3
Mir21c
Ccdc7a
n−R5s100
Arv1
Map10
Coa6
Nrp1
Ccdc7b
n−R5s103
Trim67
Ntpcr
Irf2bp2
Mir1903
n−R5s104
Gnpat
Pcnx2
Tomm20
Itgb1
58 genes omitted
124
126
128
130
132
Position on chr8 (Mb)

| | |
|---|---|
| date: | Thu Oct 25 17:04:47 2018 |
| build: | F34GBS |
| display range: | chr8:123524048−133524048 [123524048−133524048] |
| hilite range: | 0 − 0 [ 0 − 0 ] |
| reference SNP: | chr8:128524048 |
| number of SNPs plotted: | 191 |
| min P−value: | 2.62E−6 [chr8:128524048] |
| max P−value: | 6.49E−2 [chr8:125378859] |
| omitted Genes: | n−R5s105, n−R5s108, n−R5s110 |
| omitted Genes: | n−R5s111, n−R5s113, Gm25018 |
| omitted Genes: | n−R5s117, n−R5s121, n−R5s122 |
| omitted Genes: | n−R5s123, n−R5s124, n−R5s128 |
| omitted Genes: | Gm26391, n−R5s133, n−R5s134 |
| omitted Genes: | Rn5s, n−R5s136, n−R5s138 |
| omitted Genes: | n−R5s139, Gm22109, n−R5s141 |
| omitted Genes: | n−R5s142, n−R5s143, n−R5s144 |
| omitted Genes: | Gm25212, n−R5s146, Gm22291 |
| omitted Genes: | Gm23284, n−R5s149, Rhou |
| omitted Genes: | 6030466F02Rik, Rab4a, Ccsap |
| omitted Genes: | Acta1, Nup133, Abcb10 |
| omitted Genes: | Taf5l, Mir1967, Urb2 |
| omitted Genes: | Galnt2, Pgbd5, Agt |
| omitted Genes: | Capn9, 2310022B05Rik, Ttc13 |
| omitted Genes: | Fam89a, 2810004N23Rik, Exoc8 |
| omitted Genes: | Sprtn, Egln1, Tsnax |
| omitted Genes: | Disc1, 4930567H12Rik, BC021891 |
| omitted Genes: | Kcnk1, Gm17296, Rbm34 |
| omitted Genes: | 2610044O15Rik8 |

Make more plots at http://csg.sph.umich.edu/locuszoom/

Fine−mapping Regions

| snp | chr | pos (Mb) | pp | group | color |
|---|---|---|---|---|---|
| chr8:128523972 | 8 | 128.5240 | 0.033717596 | Credible Set | purple |
| chr8:128524047 | 8 | 128.5240 | 0.238995812 | Credible Set | purple |
| chr8:128524048 | 8 | 128.5240 | 0.241872420 | Credible Set | purple |
| chr8:128524081 | 8 | 128.5241 | 0.232170029 | Credible Set | purple |
| chr8:128524424 | 8 | 128.5244 | 0.033841364 | Credible Set | purple |
| chr8:128524425 | 8 | 128.5244 | 0.033257818 | Credible Set | purple |
| chr8:128524441 | 8 | 128.5244 | 0.033192537 | Credible Set | purple |
| chr8:128551483 | 8 | 128.5515 | 0.020601125 | Credible Set | purple |
| chr8:128571869 | 8 | 128.5719 | 0.005561569 | Credible Set | purple |
| chr8:128571887 | 8 | 128.5719 | 0.005561968 | Credible Set | purple |
| chr8:128711717 | 8 | 128.7117 | 0.040376936 | Credible Set | purple |
| chr8:128771719 | 8 | 128.7717 | 0.043695102 | Credible Set | purple |
| chr8:128898075 | 8 | 128.8981 | 0.031594156 | Credible Set | purple |

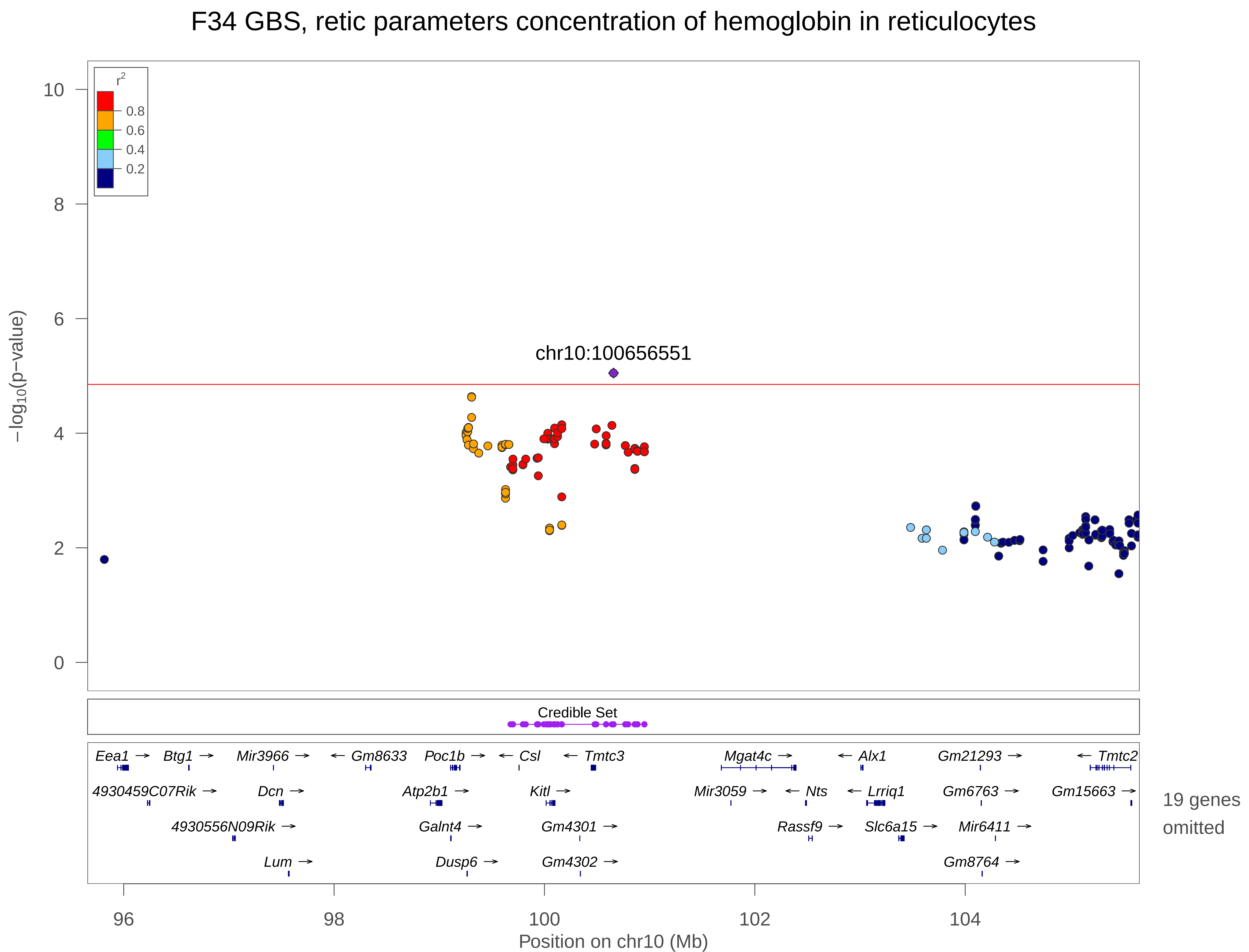

F34 GBS, retic parameters concentration of hemoglobin in reticulocytes
r2
0.8
0.6
0.4
0.2
chr10:100656551
−log10(p−value)
10
8
6
4
2
0
Credible Set
Eea1
Btg1
Mir3966
Gm8633
Poc1b
Csl
Tmtc3
Mgat4c
Alx1
Gm21293
Tmtc2
4930459C07Rik
Dcn
Atp2b1
Kitl
Mir3059
Nts
Lrriq1
Gm6763
Gm15663
4930556N09Rik
Galnt4
Gm4301
Rassf9
Slc6a15
Mir6411
Lum
Dusp6
Gm4302
Gm8764
19 genes omitted
96
98
100
102
104
Position on chr10 (Mb)

| | |
|---|---|
| date: | Thu Oct 25 17:04:25 2018 |
| build: | F34GBS |
| display range: | chr10:95656551−105656551 [95656551−105656551] |
| hilite range: | 0 − 0 [ 0 − 0 ] |
| reference SNP: | chr10:100656551 |
| number of SNPs plotted: | 225 |
| min P−value: | 8.91E−6 [chr10:100656551] |
| max P−value: | 2.83E−2 [chr10:105460999] |
| omitted Genes: | 4930473O22Rik, Kera, Epyc |
| omitted Genes: | Gm10754, Ccer1, 4930525C09Rik |
| omitted Genes: | Gm20110, Gm18409, Gm4303 |
| omitted Genes: | Gm4305, Gm4307, Gm4312 |
| omitted Genes: | Cep290, 4930430F08Rik, 1700017N19Rik |
| omitted Genes: | Gm21304, Gm21312, Gm20765 |
| omitted Genes: | Gm4340 |

Make more plots at http://csg.sph.umich.edu/locuszoom/

Fine−mapping Regions

| snp | chr | pos (Mb) | pp | group | color |
|---|---|---|---|---|---|
| chr10:99678204 | 10 | 99.67820 | 0.003695785 | Credible Set | purple |
| chr10:99699271 | 10 | 99.69927 | 0.004004593 | Credible Set | purple |
| chr10:99699681 | 10 | 99.69968 | 0.003455693 | Credible Set | purple |
| chr10:99700646 | 10 | 99.70065 | 0.004977286 | Credible Set | purple |
| chr10:99795502 | 10 | 99.79550 | 0.004015112 | Credible Set | purple |
| chr10:99795653 | 10 | 99.79565 | 0.004090384 | Credible Set | purple |
| chr10:99821621 | 10 | 99.82162 | 0.004993822 | Credible Set | purple |
| chr10:99929872 | 10 | 99.92987 | 0.005147946 | Credible Set | purple |
| chr10:99930010 | 10 | 99.93001 | 0.005147946 | Credible Set | purple |
| chr10:99941475 | 10 | 99.94147 | 0.005237917 | Credible Set | purple |
| chr10:99994442 | 10 | 99.99444 | 0.010728454 | Credible Set | purple |
| chr10:100020266 | 10 | 100.02027 | 0.010728454 | Credible Set | purple |
| chr10:100031418 | 10 | 100.03142 | 0.010569132 | Credible Set | purple |
| chr10:100031691 | 10 | 100.03169 | 0.013242003 | Credible Set | purple |
| chr10:100048262 | 10 | 100.04826 | 0.010728454 | Credible Set | purple |
| chr10:100048350 | 10 | 100.04835 | 0.010728454 | Credible Set | purple |
| chr10:100048362 | 10 | 100.04836 | 0.010728454 | Credible Set | purple |
| chr10:100055412 | 10 | 100.05541 | 0.010728454 | Credible Set | purple |
| chr10:100055418 | 10 | 100.05542 | 0.010728454 | Credible Set | purple |
| chr10:100086663 | 10 | 100.08666 | 0.010728454 | Credible Set | purple |
| chr10:100096296 | 10 | 100.09630 | 0.008868477 | Credible Set | purple |
| chr10:100096708 | 10 | 100.09671 | 0.010728454 | Credible Set | purple |
| chr10:100096785 | 10 | 100.09678 | 0.016158474 | Credible Set | purple |
| chr10:100096807 | 10 | 100.09681 | 0.016158474 | Credible Set | purple |
| chr10:100124479 | 10 | 100.12448 | 0.011605958 | Credible Set | purple |
| chr10:100125049 | 10 | 100.12505 | 0.013368238 | Credible Set | purple |
| chr10:100164149 | 10 | 100.16415 | 0.018302663 | Credible Set | purple |
| chr10:100164180 | 10 | 100.16418 | 0.018302663 | Credible Set | purple |
| chr10:100164914 | 10 | 100.16491 | 0.015790220 | Credible Set | purple |
| chr10:100477167 | 10 | 100.47717 | 0.008760724 | Credible Set | purple |
| chr10:100492319 | 10 | 100.49232 | 0.015612511 | Credible Set | purple |
| chr10:100585564 | 10 | 100.58556 | 0.008502828 | Credible Set | purple |
| chr10:100585784 | 10 | 100.58578 | 0.009035353 | Credible Set | purple |
| chr10:100585854 | 10 | 100.58585 | 0.008814260 | Credible Set | purple |
| chr10:100586507 | 10 | 100.58651 | 0.012102948 | Credible Set | purple |
| chr10:100641295 | 10 | 100.64129 | 0.017827993 | Credible Set | purple |

Fine−mapping Regions

| snp | chr | pos (Mb) | pp | group | color |
|---|---|---|---|---|---|
| chr10:100656540 | 10 | 100.6565 | 0.130504087 | Credible Set | purple |
| chr10:100656551 | 10 | 100.6566 | 0.130504087 | Credible Set | purple |
| chr10:100656588 | 10 | 100.6566 | 0.130504087 | Credible Set | purple |
| chr10:100656608 | 10 | 100.6566 | 0.130504087 | Credible Set | purple |
| chr10:100768688 | 10 | 100.7687 | 0.008260169 | Credible Set | purple |
| chr10:100768699 | 10 | 100.7687 | 0.008260169 | Credible Set | purple |
| chr10:100768710 | 10 | 100.7687 | 0.008260169 | Credible Set | purple |
| chr10:100768727 | 10 | 100.7687 | 0.008260169 | Credible Set | purple |
| chr10:100795827 | 10 | 100.7958 | 0.006440819 | Credible Set | purple |
| chr10:100858643 | 10 | 100.8586 | 0.007233931 | Credible Set | purple |
| chr10:100858698 | 10 | 100.8587 | 0.007455446 | Credible Set | purple |
| chr10:100858701 | 10 | 100.8587 | 0.007380061 | Credible Set | purple |
| chr10:100858992 | 10 | 100.8590 | 0.003502672 | Credible Set | purple |
| chr10:100859031 | 10 | 100.8590 | 0.003372882 | Credible Set | purple |
| chr10:100859032 | 10 | 100.8590 | 0.003425095 | Credible Set | purple |
| chr10:100884062 | 10 | 100.8841 | 0.006861303 | Credible Set | purple |
| chr10:100884636 | 10 | 100.8846 | 0.006674317 | Credible Set | purple |
| chr10:100949966 | 10 | 100.9500 | 0.007943553 | Credible Set | purple |
| chr10:100950077 | 10 | 100.9501 | 0.006555414 | Credible Set | purple |
| chr10:100950118 | 10 | 100.9501 | 0.006555414 | Credible Set | purple |

F34 GBS, retic parameters mean reticulocyte volume

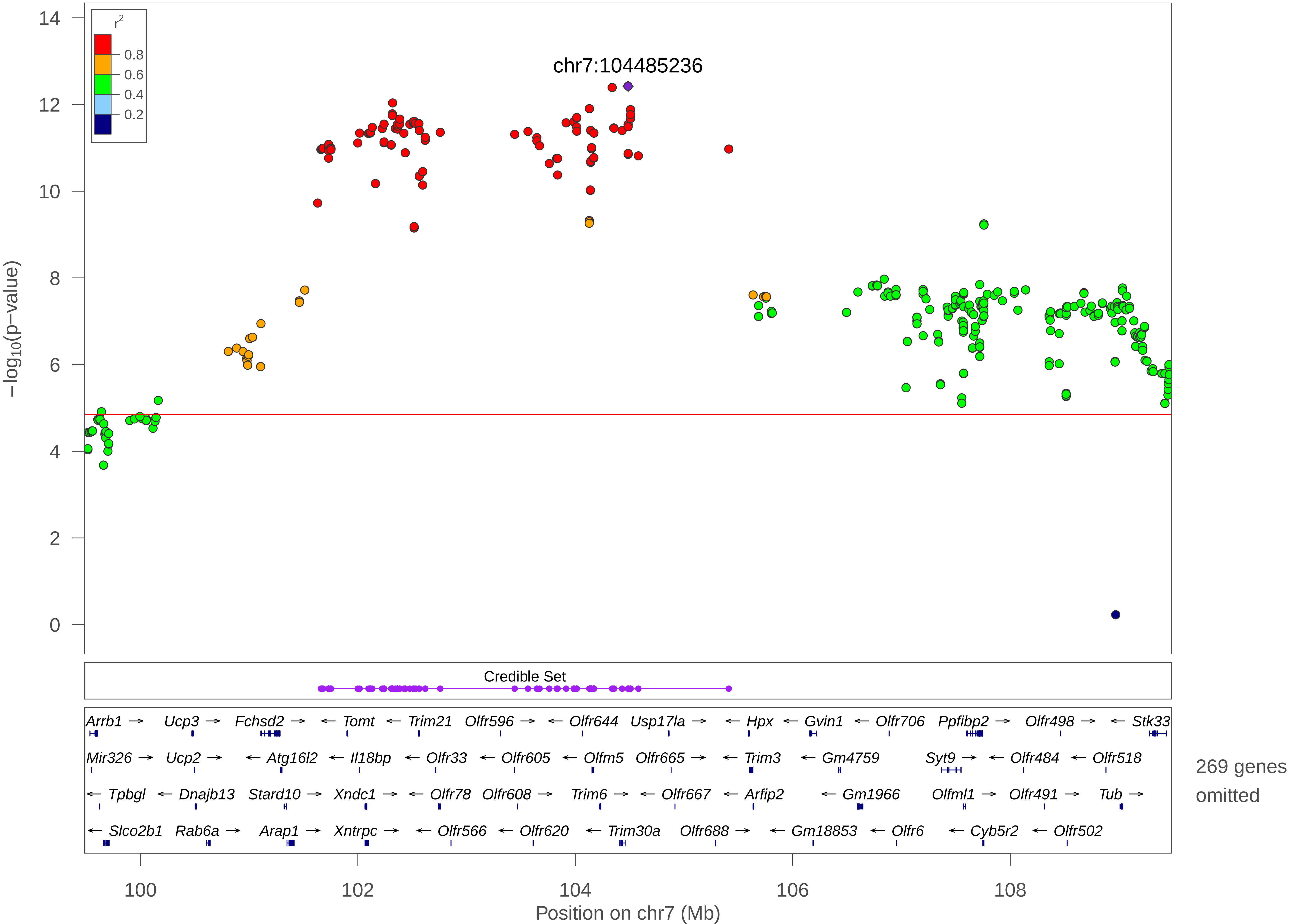

date: Thu Oct 25 17:05:11 2018
build: F34GBS
display range: chr7:99485236−109485236 [99485236−109485236]
hilite range: 0 − 0 [ 0 − 0 ]
reference SNP: chr7:104485236
number of SNPs plotted: 448
min P−value: 3.78E−13 [chr7:104485236]
max P−value: 5.92E−1 [chr7:108970724]
omitted Genes: Olfr520, Olfr521, Neu3
omitted Genes: Spcs2, Xrra1, Rnf169
omitted Genes: Chrdl2, Pold3, Lipt2
omitted Genes: Kcne3, Pgm2l1, Gpx2−ps1
omitted Genes: P4ha3, Ppme1, C2cd3
omitted Genes: Coa4, Mrpl48, Plekhb1
omitted Genes: Fam168a, Relt, Arhgef17
omitted Genes: Mir3102, P2ry6, P2ry2
omitted Genes: Pde2a, Mir139, Art2a−ps
omitted Genes: Art2b, Clpb, Phox2a
omitted Genes: Inppl1, Folr2, Folr1
omitted Genes: Anapc15, Lamtor1, Lrrc51
omitted Genes: Numa1, Rnf121, Trpc2
omitted Genes: Art5, Art1, Chrna10
omitted Genes: Nup98, Pgap2, Rhog
omitted Genes: Stim1, Rrm1, Olfr543
omitted Genes: Olfr544, Olfr545, Olfr547
omitted Genes: Olfr549, Olfr550, Olfr551
omitted Genes: Olfr552, Olfr553, Olfr554
omitted Genes: Olfr555, Olfr556, Trim68
omitted Genes: Olfr557, Olfr558, Olfr559
omitted Genes: Olfr560, Olfr561, Olfr564
omitted Genes: Olfr568, Olfr569, Olfr570
omitted Genes: Olfr571, Olfr572, Olfr574
omitted Genes: Olfr575, Olfr576, Olfr577
omitted Genes: Olfr578, Olfr582, Olfr583
omitted Genes: Olfr584, Olfr585, Olfr586
omitted Genes: Olfr589, Olfr591, Olfr592
omitted Genes: Olfr593, Olfr594, Usp17ld
omitted Genes: Olfr597, Olfr598, Olfr599
omitted Genes: Olfr600, Olfr601, Olfr603
omitted Genes: Usp17lc, Olfr606, Olfr609

Fine−mapping Regions

| snp | chr | pos (Mb) | pp | group | color |
|---|---|---|---|---|---|
| chr7:101660383 | 7 | 101.6604 | 0.002922584 | Credible Set | purple |
| chr7:101669186 | 7 | 101.6692 | 0.002980118 | Credible Set | purple |
| chr7:101673483 | 7 | 101.6735 | 0.003100272 | Credible Set | purple |
| chr7:101680140 | 7 | 101.6801 | 0.003089270 | Credible Set | purple |
| chr7:101731061 | 7 | 101.7311 | 0.003264047 | Credible Set | purple |
| chr7:101731110 | 7 | 101.7311 | 0.002720615 | Credible Set | purple |
| chr7:101731130 | 7 | 101.7311 | 0.001858530 | Credible Set | purple |
| chr7:101731658 | 7 | 101.7317 | 0.003812740 | Credible Set | purple |
| chr7:101751790 | 7 | 101.7518 | 0.003150088 | Credible Set | purple |
| chr7:101752580 | 7 | 101.7526 | 0.003097184 | Credible Set | purple |
| chr7:101752971 | 7 | 101.7530 | 0.002881860 | Credible Set | purple |
| chr7:101998679 | 7 | 101.9987 | 0.004105885 | Credible Set | purple |
| chr7:102016307 | 7 | 102.0163 | 0.006874705 | Credible Set | purple |
| chr7:102098054 | 7 | 102.0981 | 0.006735745 | Credible Set | purple |
| chr7:102110109 | 7 | 102.1101 | 0.006940155 | Credible Set | purple |
| chr7:102110501 | 7 | 102.1105 | 0.007050173 | Credible Set | purple |
| chr7:102116318 | 7 | 102.1163 | 0.006921301 | Credible Set | purple |
| chr7:102116430 | 7 | 102.1164 | 0.006921301 | Credible Set | purple |
| chr7:102116478 | 7 | 102.1165 | 0.007172341 | Credible Set | purple |
| chr7:102132562 | 7 | 102.1326 | 0.009237227 | Credible Set | purple |
| chr7:102223396 | 7 | 102.2234 | 0.008678256 | Credible Set | purple |
| chr7:102240695 | 7 | 102.2407 | 0.010954729 | Credible Set | purple |
| chr7:102241191 | 7 | 102.2412 | 0.004136877 | Credible Set | purple |
| chr7:102241211 | 7 | 102.2412 | 0.004235478 | Credible Set | purple |
| chr7:102241212 | 7 | 102.2412 | 0.004323937 | Credible Set | purple |
| chr7:102306968 | 7 | 102.3070 | 0.003675950 | Credible Set | purple |
| chr7:102306978 | 7 | 102.3070 | 0.003746813 | Credible Set | purple |
| chr7:102317213 | 7 | 102.3172 | 0.018744530 | Credible Set | purple |
| chr7:102317246 | 7 | 102.3172 | 0.017143486 | Credible Set | purple |
| chr7:102319586 | 7 | 102.3196 | 0.033053856 | Credible Set | purple |
| chr7:102344970 | 7 | 102.3450 | 0.008763101 | Credible Set | purple |
| chr7:102363897 | 7 | 102.3639 | 0.010237186 | Credible Set | purple |
| chr7:102364204 | 7 | 102.3642 | 0.009390003 | Credible Set | purple |
| chr7:102364223 | 7 | 102.3642 | 0.008475725 | Credible Set | purple |
| chr7:102364315 | 7 | 102.3643 | 0.009876047 | Credible Set | purple |
| chr7:102364326 | 7 | 102.3643 | 0.009876047 | Credible Set | purple |

Fine−mapping Regions

| snp | chr | pos (Mb) | pp | group | color |
|---|---|---|---|---|---|
| chr7:102364411 | 7 | 102.3644 | 0.010896742 | Credible Set | purple |
| chr7:102385520 | 7 | 102.3855 | 0.011063609 | Credible Set | purple |
| chr7:102385536 | 7 | 102.3855 | 0.014229578 | Credible Set | purple |
| chr7:102423165 | 7 | 102.4232 | 0.006809155 | Credible Set | purple |
| chr7:102435692 | 7 | 102.4357 | 0.002460061 | Credible Set | purple |
| chr7:102435720 | 7 | 102.4357 | 0.002460061 | Credible Set | purple |
| chr7:102478469 | 7 | 102.4785 | 0.010867907 | Credible Set | purple |
| chr7:102510475 | 7 | 102.5105 | 0.011608096 | Credible Set | purple |
| chr7:102510607 | 7 | 102.5106 | 0.012399442 | Credible Set | purple |
| chr7:102516878 | 7 | 102.5169 | 0.012715001 | Credible Set | purple |
| chr7:102531652 | 7 | 102.5317 | 0.011438024 | Credible Set | purple |
| chr7:102561708 | 7 | 102.5617 | 0.011251725 | Credible Set | purple |
| chr7:102565410 | 7 | 102.5654 | 0.007851336 | Credible Set | purple |
| chr7:102619418 | 7 | 102.6194 | 0.004724310 | Credible Set | purple |
| chr7:102619431 | 7 | 102.6194 | 0.004725741 | Credible Set | purple |
| chr7:102619466 | 7 | 102.6195 | 0.005514189 | Credible Set | purple |
| chr7:102757499 | 7 | 102.7575 | 0.007152918 | Credible Set | purple |
| chr7:103442570 | 7 | 103.4426 | 0.006445316 | Credible Set | purple |
| chr7:103564836 | 7 | 103.5648 | 0.007471790 | Credible Set | purple |
| chr7:103646439 | 7 | 103.6464 | 0.005434970 | Credible Set | purple |
| chr7:103646709 | 7 | 103.6467 | 0.004594335 | Credible Set | purple |
| chr7:103671033 | 7 | 103.6710 | 0.003535162 | Credible Set | purple |
| chr7:103760301 | 7 | 103.7603 | 0.001406962 | Credible Set | purple |
| chr7:103829200 | 7 | 103.8292 | 0.001838384 | Credible Set | purple |
| chr7:103837213 | 7 | 103.8372 | 0.001835699 | Credible Set | purple |
| chr7:103915503 | 7 | 103.9155 | 0.011708764 | Credible Set | purple |
| chr7:103986792 | 7 | 103.9868 | 0.012442039 | Credible Set | purple |
| chr7:103986795 | 7 | 103.9868 | 0.012442039 | Credible Set | purple |
| chr7:103986857 | 7 | 103.9869 | 0.012419049 | Credible Set | purple |
| chr7:103986860 | 7 | 103.9869 | 0.012419049 | Credible Set | purple |
| chr7:104012808 | 7 | 104.0128 | 0.015074030 | Credible Set | purple |
| chr7:104013370 | 7 | 104.0134 | 0.009389331 | Credible Set | purple |
| chr7:104013395 | 7 | 104.0134 | 0.007585969 | Credible Set | purple |
| chr7:104013467 | 7 | 104.0135 | 0.015455171 | Credible Set | purple |
| chr7:104129734 | 7 | 104.1297 | 0.024487194 | Credible Set | purple |
| chr7:104129736 | 7 | 104.1297 | 0.024487194 | Credible Set | purple |

Fine−mapping Regions

| snp | chr | pos (Mb) | pp | group | color |
|---|---|---|---|---|---|
| chr7:104141343 | 7 | 104.1413 | 0.007884958 | Credible Set | purple |
| chr7:104141705 | 7 | 104.1417 | 0.001493692 | Credible Set | purple |
| chr7:104141768 | 7 | 104.1418 | 0.001575630 | Credible Set | purple |
| chr7:104149893 | 7 | 104.1499 | 0.003036090 | Credible Set | purple |
| chr7:104149911 | 7 | 104.1499 | 0.003218082 | Credible Set | purple |
| chr7:104170479 | 7 | 104.1705 | 0.001907269 | Credible Set | purple |
| chr7:104170506 | 7 | 104.1705 | 0.001894180 | Credible Set | purple |
| chr7:104170518 | 7 | 104.1705 | 0.001879994 | Credible Set | purple |
| chr7:104170522 | 7 | 104.1705 | 0.001890011 | Credible Set | purple |
| chr7:104170543 | 7 | 104.1705 | 0.001910915 | Credible Set | purple |
| chr7:104170588 | 7 | 104.1706 | 0.006882723 | Credible Set | purple |
| chr7:104170594 | 7 | 104.1706 | 0.006857523 | Credible Set | purple |
| chr7:104170618 | 7 | 104.1706 | 0.006874262 | Credible Set | purple |
| chr7:104170662 | 7 | 104.1707 | 0.006874262 | Credible Set | purple |
| chr7:104338784 | 7 | 104.3388 | 0.073962290 | Credible Set | purple |
| chr7:104338810 | 7 | 104.3388 | 0.073651039 | Credible Set | purple |
| chr7:104353491 | 7 | 104.3535 | 0.008981800 | Credible Set | purple |
| chr7:104356230 | 7 | 104.3562 | 0.008816571 | Credible Set | purple |
| chr7:104430205 | 7 | 104.4302 | 0.007838657 | Credible Set | purple |
| chr7:104485236 | 7 | 104.4852 | 0.079147235 | Credible Set | purple |
| chr7:104485721 | 7 | 104.4857 | 0.010814539 | Credible Set | purple |
| chr7:104485727 | 7 | 104.4857 | 0.009539969 | Credible Set | purple |
| chr7:104486022 | 7 | 104.4860 | 0.002262679 | Credible Set | purple |
| chr7:104486041 | 7 | 104.4860 | 0.002383354 | Credible Set | purple |
| chr7:104507724 | 7 | 104.5077 | 0.014686410 | Credible Set | purple |
| chr7:104508127 | 7 | 104.5081 | 0.023288637 | Credible Set | purple |
| chr7:104508173 | 7 | 104.5082 | 0.017930859 | Credible Set | purple |
| chr7:104580091 | 7 | 104.5801 | 0.002099295 | Credible Set | purple |
| chr7:105411752 | 7 | 105.4118 | 0.002993724 | Credible Set | purple |

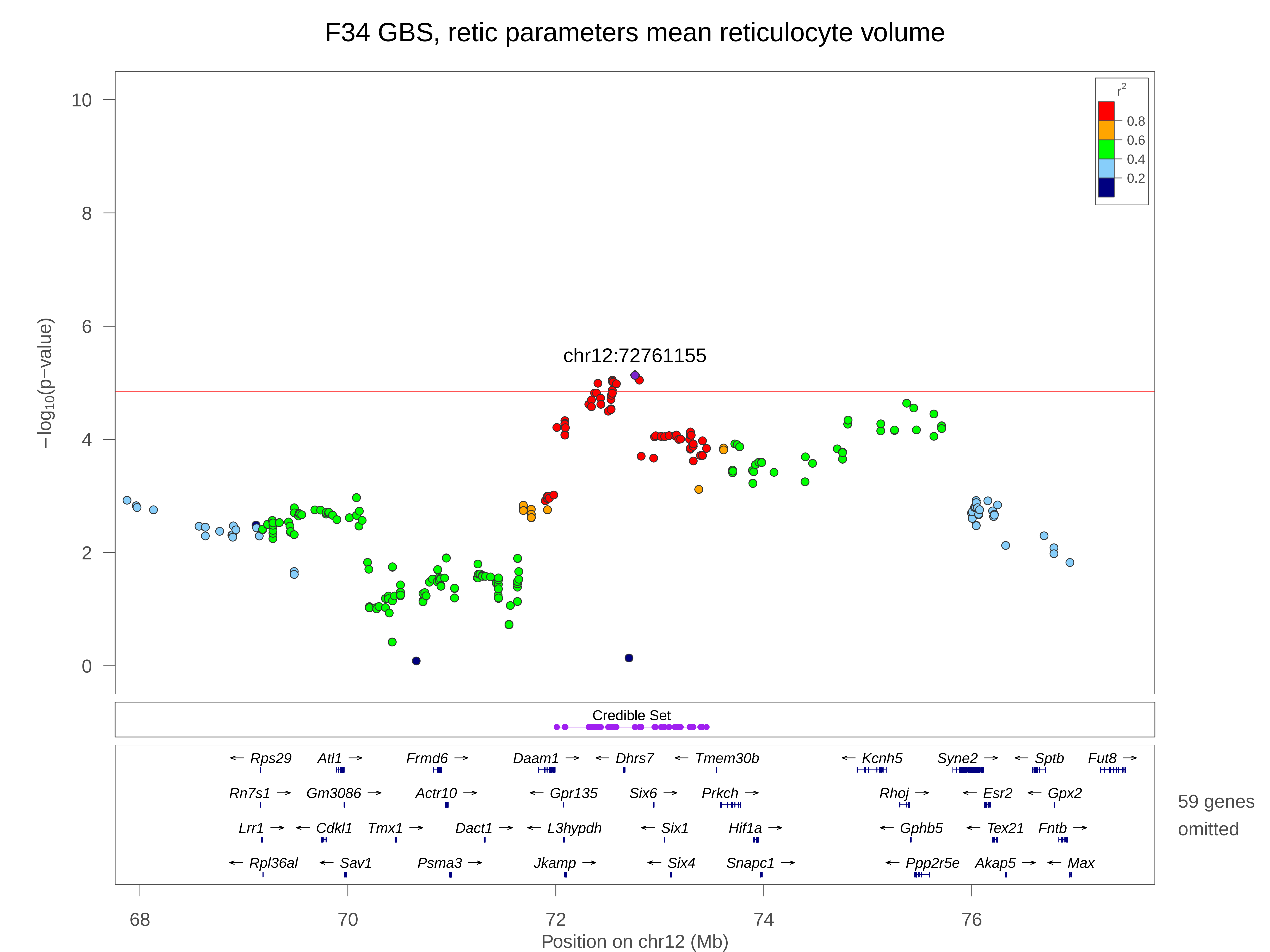
F34 GBS, retic parameters mean reticulocyte volume
−log10(p−value)
0
2
4
6
8
10
r2
0.8
0.6
0.4
0.2
chr12:72761155
Credible Set
Rps29
Atl1
Frmd6
Daam1
Dhrs7
Tmem30b
Kcnh5
Syne2
Sptb
Fut8
Rn7s1
Gm3086
Actr10
Gpr135
Six6
Prkch
Rhoj
Esr2
Gpx2
Lrr1
Cdkl1
Tmx1
Dact1
L3hypdh
Six1
Hif1a
Gphb5
Tex21
Fntb
Rpl36al
Sav1
Psma3
Jkamp
Six4
Snapc1
Ppp2r5e
Akap5
Max
59 genes omitted
68
70
72
74
76
Position on chr12 (Mb)

| | |
|---|---|
| date: | Thu Oct 25 17:04:57 2018 |
| build: | F34GBS |
| display range: | chr12:67761155−77761155 [67761155−77761155] |
| hilite range: | 0 − 0 [ 0 − 0 ] |
| reference SNP: | chr12:72761155 |
| number of SNPs plotted: | 336 |
| min P−value: | 7.38E−6 [chr12:72761155] |
| max P−value: | 8.19E−1 [chr12:70656926] |
| omitted Genes: | Mgat2, Dnaaf2, 9330151L19Rik |
| omitted Genes: | Pole2, Klhdc1, Klhdc2 |
| omitted Genes: | Nemf, Rn7s2, Arf6 |
| omitted Genes: | 5830428M24Rik, Vcpkmt, Sos2 |
| omitted Genes: | L2hgdh, Atp5s, Mir681 |
| omitted Genes: | 4930512B01Rik, Map4k5, 4931403G20Rik |
| omitted Genes: | Nin, Abhd12b, Pygl |
| omitted Genes: | F730035M05Rik, Trim9, 3110056K07Rik |
| omitted Genes: | Arid4a, Tomm20l, Timm9 |
| omitted Genes: | 2700049A03Rik, 1700083H02Rik, 4930404H11Rik |
| omitted Genes: | Ccdc175, Rtn1, Lrrc9 |
| omitted Genes: | Pcnx4, Ppm1a, 4930447C04Rik |
| omitted Genes: | Mnat1, Trmt5, Slc38a6 |
| omitted Genes: | D830013O20Rik, 2210039B01Rik, Syt16 |
| omitted Genes: | 1700086L19Rik, Dbpht2, Wdr89 |
| omitted Genes: | Rplp2−ps1, Sgpp1, Mir5101 |
| omitted Genes: | Mthfd1, Zbtb25, Zbtb1 |
| omitted Genes: | Hspa2, Ppp1r36, Gm10451 |
| omitted Genes: | Mir5135, Plekhg3, AI463170 |
| omitted Genes: | Churc1, Rab15 |

Make more plots at http://csg.sph.umich.edu/locuszoom/

Fine−mapping Regions

| snp | chr | pos (Mb) | pp | group | color |
|---|---|---|---|---|---|
| chr12:72009199 | 12 | 72.00920 | 0.009313544 | Credible Set | purple |
| chr12:72086662 | 12 | 72.08666 | 0.012109109 | Credible Set | purple |
| chr12:72086706 | 12 | 72.08671 | 0.010931224 | Credible Set | purple |
| chr12:72086724 | 12 | 72.08672 | 0.010616944 | Credible Set | purple |
| chr12:72086792 | 12 | 72.08679 | 0.007133110 | Credible Set | purple |
| chr12:72086995 | 12 | 72.08700 | 0.006955550 | Credible Set | purple |
| chr12:72087027 | 12 | 72.08703 | 0.006936629 | Credible Set | purple |
| chr12:72091767 | 12 | 72.09177 | 0.009223625 | Credible Set | purple |
| chr12:72316934 | 12 | 72.31693 | 0.022792517 | Credible Set | purple |
| chr12:72342139 | 12 | 72.34214 | 0.026925751 | Credible Set | purple |
| chr12:72342228 | 12 | 72.34223 | 0.020744439 | Credible Set | purple |
| chr12:72372220 | 12 | 72.37222 | 0.035303028 | Credible Set | purple |
| chr12:72389901 | 12 | 72.38990 | 0.035282384 | Credible Set | purple |
| chr12:72404532 | 12 | 72.40453 | 0.051569904 | Credible Set | purple |
| chr12:72430018 | 12 | 72.43002 | 0.028832638 | Credible Set | purple |
| chr12:72432423 | 12 | 72.43242 | 0.022760314 | Credible Set | purple |
| chr12:72503712 | 12 | 72.50371 | 0.017448019 | Credible Set | purple |
| chr12:72530507 | 12 | 72.53051 | 0.027468994 | Credible Set | purple |
| chr12:72531030 | 12 | 72.53103 | 0.019107273 | Credible Set | purple |
| chr12:72531036 | 12 | 72.53104 | 0.018537023 | Credible Set | purple |
| chr12:72534336 | 12 | 72.53434 | 0.032352122 | Credible Set | purple |
| chr12:72542646 | 12 | 72.54265 | 0.039247508 | Credible Set | purple |
| chr12:72542706 | 12 | 72.54271 | 0.034964712 | Credible Set | purple |
| chr12:72543184 | 12 | 72.54318 | 0.058214784 | Credible Set | purple |
| chr12:72543205 | 12 | 72.54321 | 0.055352205 | Credible Set | purple |
| chr12:72551877 | 12 | 72.55188 | 0.053745692 | Credible Set | purple |
| chr12:72582082 | 12 | 72.58208 | 0.050431926 | Credible Set | purple |
| chr12:72761155 | 12 | 72.76116 | 0.070149570 | Credible Set | purple |
| chr12:72803219 | 12 | 72.80322 | 0.058077048 | Credible Set | purple |
| chr12:72820664 | 12 | 72.82066 | 0.003091247 | Credible Set | purple |
| chr12:72948661 | 12 | 72.94866 | 0.006497966 | Credible Set | purple |
| chr12:72960023 | 12 | 72.96002 | 0.006766562 | Credible Set | purple |
| chr12:73012360 | 12 | 73.01236 | 0.006588722 | Credible Set | purple |
| chr12:73046310 | 12 | 73.04631 | 0.006532534 | Credible Set | purple |
| chr12:73088646 | 12 | 73.08865 | 0.006792230 | Credible Set | purple |
| chr12:73144833 | 12 | 73.14483 | 0.006777596 | Credible Set | purple |

Fine−mapping Regions

| snp | chr | pos (Mb) | pp | group | color |
|---|---|---|---|---|---|
| chr12:73160141 | 12 | 73.16014 | 0.006973151 | Credible Set | purple |
| chr12:73182054 | 12 | 73.18205 | 0.005897369 | Credible Set | purple |
| chr12:73200528 | 12 | 73.20053 | 0.005957969 | Credible Set | purple |
| chr12:73287081 | 12 | 73.28708 | 0.005871430 | Credible Set | purple |
| chr12:73293245 | 12 | 73.29324 | 0.004042649 | Credible Set | purple |
| chr12:73293285 | 12 | 73.29328 | 0.004179006 | Credible Set | purple |
| chr12:73293811 | 12 | 73.29381 | 0.007219253 | Credible Set | purple |
| chr12:73293812 | 12 | 73.29381 | 0.006934058 | Credible Set | purple |
| chr12:73294375 | 12 | 73.29438 | 0.007853504 | Credible Set | purple |
| chr12:73301700 | 12 | 73.30170 | 0.006913792 | Credible Set | purple |
| chr12:73320679 | 12 | 73.32068 | 0.004514037 | Credible Set | purple |
| chr12:73320695 | 12 | 73.32070 | 0.004514037 | Credible Set | purple |
| chr12:73320730 | 12 | 73.32073 | 0.004941685 | Credible Set | purple |
| chr12:73390527 | 12 | 73.39053 | 0.003172300 | Credible Set | purple |
| chr12:73409987 | 12 | 73.40999 | 0.003174178 | Credible Set | purple |
| chr12:73409994 | 12 | 73.40999 | 0.003175461 | Credible Set | purple |
| chr12:73410037 | 12 | 73.41004 | 0.005585985 | Credible Set | purple |
| chr12:73449146 | 12 | 73.44915 | 0.004169872 | Credible Set | purple |

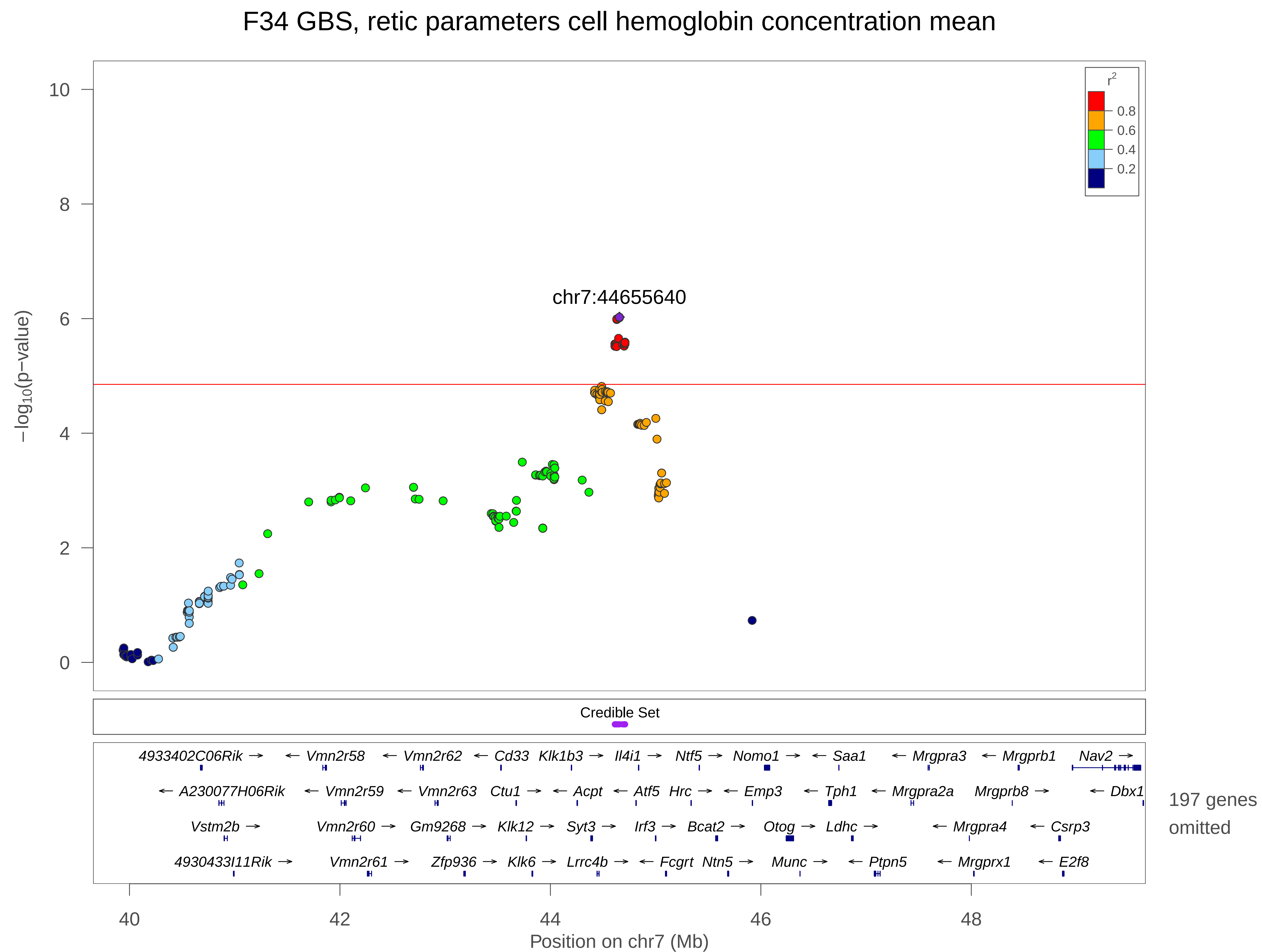
F34 GBS, retic parameters cell hemoglobin concentration mean
chr7:44655640
−log10(p−value)
r2
0.8
0.6
0.4
0.2
Credible Set
4933402C06Rik → ← Vmn2r58 ← Vmn2r62 ← Cd33 Klk1b3 → Il4i1 → Ntf5 → Nomo1 → ← Saa1 ← Mrgpra3 ← Mrgprb1 Nav2 →
← A230077H06Rik ← Vmn2r59 ← Vmn2r63 Ctu1 → ← Acpt ← Atf5 Hrc → ← Emp3 ← Tph1 ← Mrgpra2a Mrgprb8 → ← Dbx1
Vstm2b → Vmn2r60 → Gm9268 → Klk12 → Syt3 → Irf3 → Bcat2 → Otog → Ldhc → ← Mrgpra4 ← Csrp3
4930433I11Rik → Vmn2r61 → Zfp936 → Klk6 → Lrrc4b → ← Fcgrt Ntn5 → Munc → ← Ptpn5 ← Mrgprx1 ← E2f8
197 genes omitted
Position on chr7 (Mb)

date: Thu Oct 25 17:06:00 2018
build: F34GBS
display range: chr7:39655640−49655640 [39655640−49655640]
hilite range: 0 − 0 [ 0 − 0 ]
reference SNP: chr7:44655640
number of SNPs plotted: 247
min P−value: 9.42E−7 [chr7:44655640]
max P−value: 9.76E−1 [chr7:40177351]
omitted Genes: Gm4884, 1700110I07Rik, Gm5592
omitted Genes: AI987944, Vmn2r57, AW146154
omitted Genes: Zfp788, Vmn2r−ps54, Vmn2r−ps60
omitted Genes: 4933421I07Rik, Zfp141, 9830147E19Rik
omitted Genes: Gm5595, Gm2381, EU599041
omitted Genes: Zfp715, Siglecf, Siglecg
omitted Genes: Lim2, Nkg7, Cldnd2
omitted Genes: Etfb, Mir7051, Vsig10l
omitted Genes: Iglon5, 4931406B18Rik, Zfp658
omitted Genes: Zfp719, Zfp819, Ceacam18
omitted Genes: Siglece, Klk14, Klk13
omitted Genes: 2310002F09Rik, Klk11, Klk10
omitted Genes: Klk9, Klk8, Klk7
omitted Genes: Klk5, Klk4, Klk15
omitted Genes: Klk1b7−ps, Klk1b8, Klk1b1
omitted Genes: Klk1b9, Klk1b11, Klk1b26
omitted Genes: Klk1b27, Klk1b21, Klk1b22
omitted Genes: Klk1b16, Klk1b24, Klk1b4
omitted Genes: Klk1b5, Klk1, 1700028J19Rik
omitted Genes: 2410002F23Rik, Snord88c, Snord88a
omitted Genes: Clec11a, Shank1, 1700008O03Rik
omitted Genes: Aspdh, Josd2, Mir7052
omitted Genes: 5430431A17Rik, Emc10, Fam71e1
omitted Genes: Mybpc2, Spib, Pold1
omitted Genes: Mir7053, Nr1h2, Napsa
omitted Genes: Kcnc3, Myh14, Izumo2
omitted Genes: Zfp473, Vrk3, Nup62
omitted Genes: Gm21948, Tbc1d17, Akt1s1
omitted Genes: Mir707, Pnkp, Ptov1
omitted Genes: Med25, Fuz, Ap2a1
omitted Genes: Tsks, Cpt1c, Prmt1
Make more plots at http://csg.sph.umich.edu/locuszoom/
omitted Genes: Gm15545, Bcl2l12, Scaf1

Fine−mapping Regions

| snp | chr | pos (Mb) | pp | group | color |
|---|---|---|---|---|---|
| chr7:44612740 | 7 | 44.61274 | 0.04019643 | Credible Set | purple |
| chr7:44613080 | 7 | 44.61308 | 0.03673297 | Credible Set | purple |
| chr7:44623189 | 7 | 44.62319 | 0.04042732 | Credible Set | purple |
| chr7:44625757 | 7 | 44.62576 | 0.03771995 | Credible Set | purple |
| chr7:44628444 | 7 | 44.62844 | 0.03622587 | Credible Set | purple |
| chr7:44630865 | 7 | 44.63087 | 0.10304051 | Credible Set | purple |
| chr7:44630890 | 7 | 44.63089 | 0.10554906 | Credible Set | purple |
| chr7:44647914 | 7 | 44.64791 | 0.04987531 | Credible Set | purple |
| chr7:44655619 | 7 | 44.65562 | 0.11031480 | Credible Set | purple |
| chr7:44655640 | 7 | 44.65564 | 0.11308659 | Credible Set | purple |
| chr7:44685749 | 7 | 44.68575 | 0.03943631 | Credible Set | purple |
| chr7:44700610 | 7 | 44.70061 | 0.03691384 | Credible Set | purple |
| chr7:44700692 | 7 | 44.70069 | 0.04008230 | Credible Set | purple |
| chr7:44708823 | 7 | 44.70882 | 0.04008230 | Credible Set | purple |
| chr7:44708981 | 7 | 44.70898 | 0.04271153 | Credible Set | purple |
| chr7:44708990 | 7 | 44.70899 | 0.04311534 | Credible Set | purple |
| chr7:44708997 | 7 | 44.70900 | 0.04224478 | Credible Set | purple |
| chr7:44709016 | 7 | 44.70902 | 0.04224478 | Credible Set | purple |

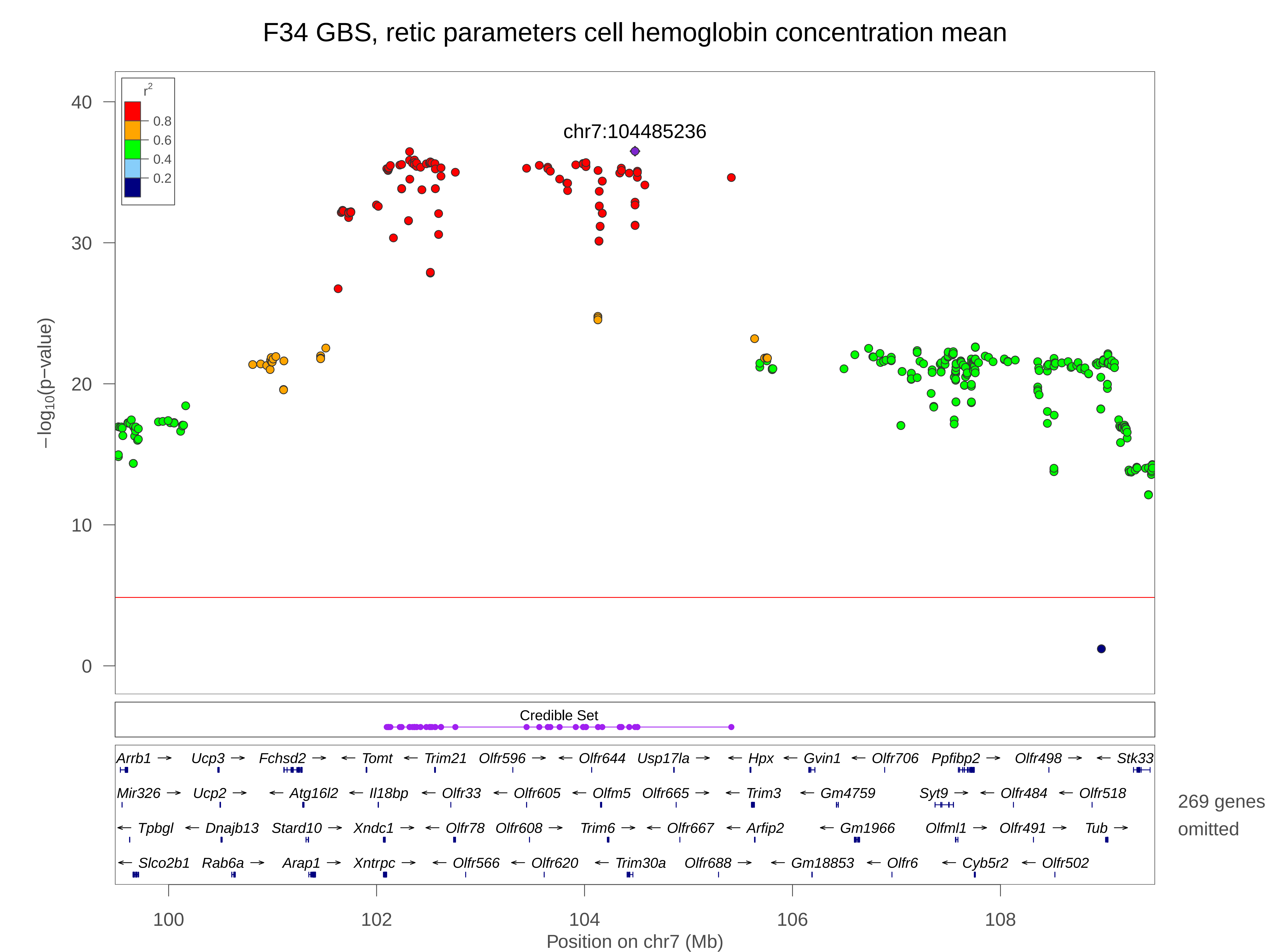

F34 GBS, retic parameters cell hemoglobin concentration mean
r2
0.8
0.6
0.4
0.2
chr7:104485236
40
30
20
10
0
−log10(p−value)
Credible Set
Arrb1 → Ucp3 → Fchsd2 → ← Tomt ← Trim21 Olfr596 → ← Olfr644 Usp17la → ← Hpx ← Gvin1 ← Olfr706 Ppfibp2 → Olfr498 → ← Stk33
Mir326 → Ucp2 → ← Atg16l2 ← Il18bp ← Olfr33 ← Olfr605 ← Olfm5 Olfr665 → ← Trim3 ← Gm4759 Syt9 → ← Olfr484 ← Olfr518
← Tpbgl ← Dnajb13 Stard10 → Xndc1 → ← Olfr78 Olfr608 → Trim6 → ← Olfr667 ← Arfip2 ← Gm1966 Olfml1 → Olfr491 → Tub →
← Slco2b1 Rab6a → Arap1 → Xntrpc → ← Olfr566 ← Olfr620 ← Trim30a Olfr688 → ← Gm18853 ← Olfr6 ← Cyb5r2 ← Olfr502
269 genes omitted
100
102
104
106
108
Position on chr7 (Mb)

date: Thu Oct 25 17:05:47 2018
build: F34GBS
display range: chr7:99485236−109485236 [99485236−109485236]
hilite range: 0 − 0 [ 0 − 0 ]
reference SNP: chr7:104485236
number of SNPs plotted: 448
min P−value: 3.19E−37 [chr7:104485236]
max P−value: 6.26E−2 [chr7:108970724]
omitted Genes: Olfr520, Olfr521, Neu3
omitted Genes: Spcs2, Xrra1, Rnf169
omitted Genes: Chrdl2, Pold3, Lipt2
omitted Genes: Kcne3, Pgm2l1, Gpx2−ps1
omitted Genes: P4ha3, Ppme1, C2cd3
omitted Genes: Coa4, Mrpl48, Plekhb1
omitted Genes: Fam168a, Relt, Arhgef17
omitted Genes: Mir3102, P2ry6, P2ry2
omitted Genes: Pde2a, Mir139, Art2a−ps
omitted Genes: Art2b, Clpb, Phox2a
omitted Genes: Inppl1, Folr2, Folr1
omitted Genes: Anapc15, Lamtor1, Lrrc51
omitted Genes: Numa1, Rnf121, Trpc2
omitted Genes: Art5, Art1, Chrna10
omitted Genes: Nup98, Pgap2, Rhog
omitted Genes: Stim1, Rrm1, Olfr543
omitted Genes: Olfr544, Olfr545, Olfr547
omitted Genes: Olfr549, Olfr550, Olfr551
omitted Genes: Olfr552, Olfr553, Olfr554
omitted Genes: Olfr555, Olfr556, Trim68
omitted Genes: Olfr557, Olfr558, Olfr559
omitted Genes: Olfr560, Olfr561, Olfr564
omitted Genes: Olfr568, Olfr569, Olfr570
omitted Genes: Olfr571, Olfr572, Olfr574
omitted Genes: Olfr575, Olfr576, Olfr577
omitted Genes: Olfr578, Olfr582, Olfr583
omitted Genes: Olfr584, Olfr585, Olfr586
omitted Genes: Olfr589, Olfr591, Olfr592
omitted Genes: Olfr593, Olfr594, Usp17ld
omitted Genes: Olfr597, Olfr598, Olfr599
omitted Genes: Olfr600, Olfr601, Olfr603
omitted Genes: Usp17le, Olfr606, Olfr609

Make more plots at http://csg.sph.umich.edu/locuszoom/

Fine−mapping Regions

| snp | chr | pos (Mb) | pp | group | color |
|---|---|---|---|---|---|
| chr7:102098054 | 7 | 102.0981 | 0.008086265 | Credible Set | purple |
| chr7:102110109 | 7 | 102.1101 | 0.006000914 | Credible Set | purple |
| chr7:102110501 | 7 | 102.1105 | 0.006483575 | Credible Set | purple |
| chr7:102116318 | 7 | 102.1163 | 0.007997225 | Credible Set | purple |
| chr7:102116430 | 7 | 102.1164 | 0.007997225 | Credible Set | purple |
| chr7:102116478 | 7 | 102.1165 | 0.009340079 | Credible Set | purple |
| chr7:102132562 | 7 | 102.1326 | 0.013561780 | Credible Set | purple |
| chr7:102223396 | 7 | 102.2234 | 0.014473276 | Credible Set | purple |
| chr7:102240695 | 7 | 102.2407 | 0.016062135 | Credible Set | purple |
| chr7:102317213 | 7 | 102.3172 | 0.031915585 | Credible Set | purple |
| chr7:102317246 | 7 | 102.3172 | 0.128817713 | Credible Set | purple |
| chr7:102319586 | 7 | 102.3196 | 0.001470396 | Credible Set | purple |
| chr7:102344970 | 7 | 102.3450 | 0.019659537 | Credible Set | purple |
| chr7:102363897 | 7 | 102.3639 | 0.015557967 | Credible Set | purple |
| chr7:102364204 | 7 | 102.3642 | 0.027955625 | Credible Set | purple |
| chr7:102364223 | 7 | 102.3642 | 0.032978368 | Credible Set | purple |
| chr7:102364315 | 7 | 102.3643 | 0.021621776 | Credible Set | purple |
| chr7:102364326 | 7 | 102.3643 | 0.021621776 | Credible Set | purple |
| chr7:102364411 | 7 | 102.3644 | 0.015597870 | Credible Set | purple |
| chr7:102385520 | 7 | 102.3855 | 0.011607758 | Credible Set | purple |
| chr7:102385536 | 7 | 102.3855 | 0.019966710 | Credible Set | purple |
| chr7:102423165 | 7 | 102.4232 | 0.010141303 | Credible Set | purple |
| chr7:102478469 | 7 | 102.4785 | 0.017703605 | Credible Set | purple |
| chr7:102510475 | 7 | 102.5105 | 0.019860576 | Credible Set | purple |
| chr7:102510607 | 7 | 102.5106 | 0.021177480 | Credible Set | purple |
| chr7:102516878 | 7 | 102.5169 | 0.024302819 | Credible Set | purple |
| chr7:102531652 | 7 | 102.5317 | 0.021019403 | Credible Set | purple |
| chr7:102561708 | 7 | 102.5617 | 0.018269972 | Credible Set | purple |
| chr7:102565410 | 7 | 102.5654 | 0.007652250 | Credible Set | purple |
| chr7:102619418 | 7 | 102.6194 | 0.009260030 | Credible Set | purple |
| chr7:102619431 | 7 | 102.6194 | 0.009201823 | Credible Set | purple |
| chr7:102619466 | 7 | 102.6195 | 0.002380883 | Credible Set | purple |
| chr7:102757499 | 7 | 102.7575 | 0.004519018 | Credible Set | purple |
| chr7:103442570 | 7 | 103.4426 | 0.008593713 | Credible Set | purple |
| chr7:103564836 | 7 | 103.5648 | 0.013800293 | Credible Set | purple |
| chr7:103646439 | 7 | 103.6464 | 0.010268750 | Credible Set | purple |

Fine−mapping Regions

| snp | chr | pos (Mb) | pp | group | color |
|---|---|---|---|---|---|
| chr7:103646709 | 7 | 103.6467 | 0.007973212 | Credible Set | purple |
| chr7:103671033 | 7 | 103.6710 | 0.005323470 | Credible Set | purple |
| chr7:103760301 | 7 | 103.7603 | 0.001495665 | Credible Set | purple |
| chr7:103915503 | 7 | 103.9155 | 0.015003463 | Credible Set | purple |
| chr7:103986792 | 7 | 103.9868 | 0.018843635 | Credible Set | purple |
| chr7:103986795 | 7 | 103.9868 | 0.018843635 | Credible Set | purple |
| chr7:103986857 | 7 | 103.9869 | 0.017031546 | Credible Set | purple |
| chr7:103986860 | 7 | 103.9869 | 0.017031546 | Credible Set | purple |
| chr7:104012808 | 7 | 104.0128 | 0.016708489 | Credible Set | purple |
| chr7:104013370 | 7 | 104.0134 | 0.011923804 | Credible Set | purple |
| chr7:104013395 | 7 | 104.0134 | 0.011183186 | Credible Set | purple |
| chr7:104013467 | 7 | 104.0135 | 0.021486394 | Credible Set | purple |
| chr7:104129734 | 7 | 104.1297 | 0.006038278 | Credible Set | purple |
| chr7:104129736 | 7 | 104.1297 | 0.006038278 | Credible Set | purple |
| chr7:104170618 | 7 | 104.1706 | 0.001081113 | Credible Set | purple |
| chr7:104338784 | 7 | 104.3388 | 0.004006456 | Credible Set | purple |
| chr7:104338810 | 7 | 104.3388 | 0.003972653 | Credible Set | purple |
| chr7:104353491 | 7 | 104.3535 | 0.008827808 | Credible Set | purple |
| chr7:104356230 | 7 | 104.3562 | 0.006046661 | Credible Set | purple |
| chr7:104430205 | 7 | 104.4302 | 0.003941042 | Credible Set | purple |
| chr7:104485236 | 7 | 104.4852 | 0.137614778 | Credible Set | purple |
| chr7:104507724 | 7 | 104.5077 | 0.001975080 | Credible Set | purple |
| chr7:104508127 | 7 | 104.5081 | 0.005362379 | Credible Set | purple |
| chr7:104508173 | 7 | 104.5082 | 0.004347901 | Credible Set | purple |
| chr7:105411752 | 7 | 105.4118 | 0.001912813 | Credible Set | purple |

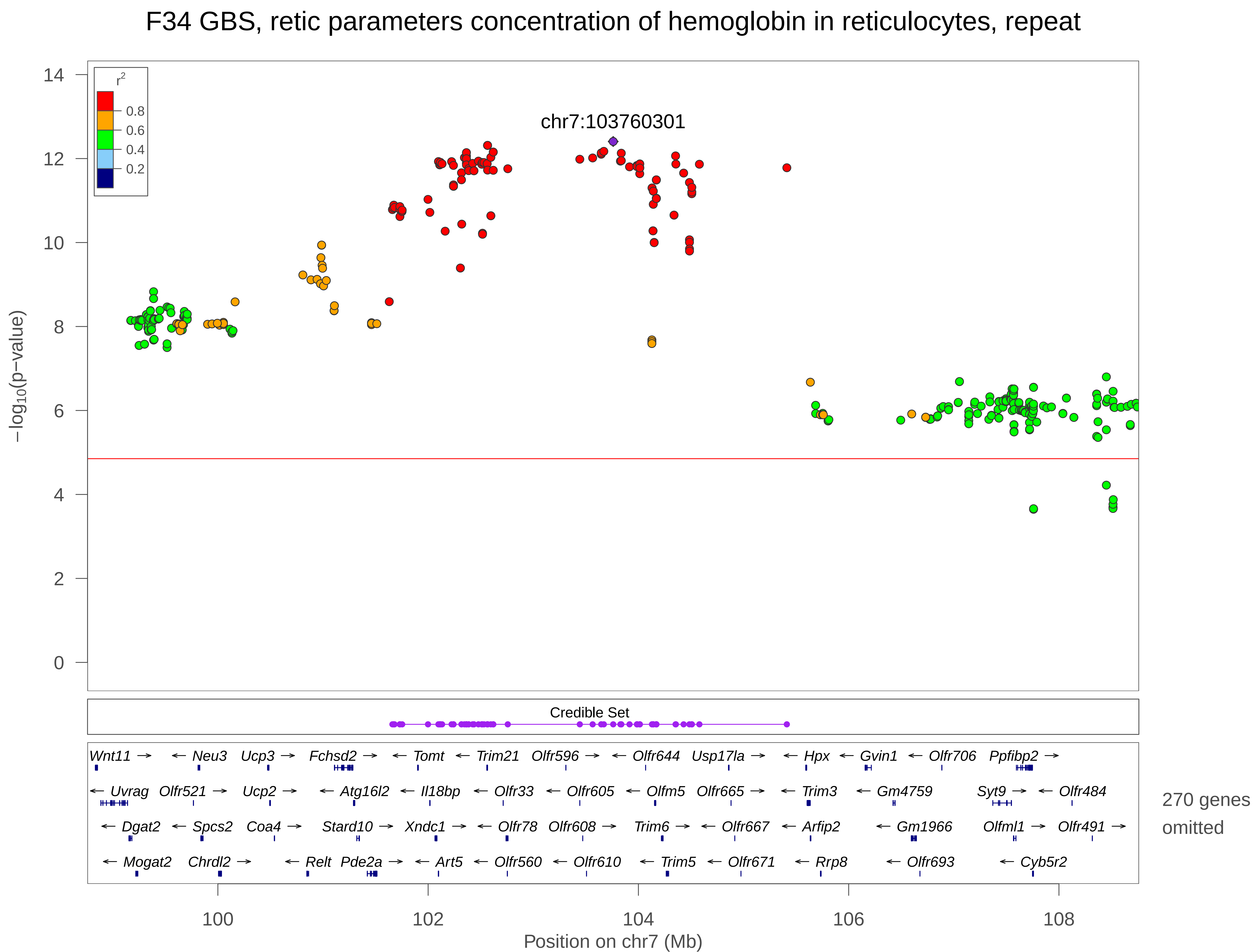

F34 GBS, retic parameters concentration of hemoglobin in reticulocytes, repeat
chr7:103760301
r2
0.8
0.6
0.4
0.2
−log10(p−value)
14
12
10
8
6
4
2
0
Credible Set
Wnt11 → ← Neu3 Ucp3 → Fchsd2 → ← Tomt ← Trim21 Olfr596 → ← Olfr644 Usp17la → ← Hpx ← Gvin1 ← Olfr706 Ppfibp2 →
← Uvrag Olfr521 → Ucp2 → ← Atg16l2 ← Il18bp ← Olfr33 ← Olfr605 ← Olfm5 Olfr665 → ← Trim3 ← Gm4759 Syt9 → ← Olfr484
← Dgat2 ← Spcs2 Coa4 → Stard10 → Xndc1 → ← Olfr78 Olfr608 → Trim6 → ← Olfr667 ← Arfip2 ← Gm1966 Olfml1 → Olfr491 →
← Mogat2 Chrdl2 → ← Relt Pde2a → ← Art5 ← Olfr560 ← Olfr610 ← Trim5 ← Olfr671 ← Rrp8 ← Olfr693 ← Cyb5r2
270 genes omitted
100
102
104
106
108
Position on chr7 (Mb)

date: Thu Oct 25 17:06:40 2018
build: F34GBS
display range: chr7:98760301−108760301 [98760301−108760301]
hilite range: 0 − 0 [ 0 − 0 ]
reference SNP: chr7:103760301
number of SNPs plotted: 417
min P−value: 3.92E−13 [chr7:103760301]
max P−value: 2.25E−4 [chr7:107758350]
omitted Genes: Gm26705, Map6, Serpinh1
omitted Genes: Gdpd5, Klhl35, Rps3
omitted Genes: Snord15b, Snord15a, Arrb1
omitted Genes: Mir326, Tpbgl, Slco2b1
omitted Genes: Olfr520, Xrra1, Rnf169
omitted Genes: Pold3, Lipt2, Kcne3
omitted Genes: Pgm2l1, Gpx2−ps1, P4ha3
omitted Genes: Ppme1, C2cd3, Dnajb13
omitted Genes: Mrpl48, Rab6a, Plekhb1
omitted Genes: Fam168a, Arhgef17, Mir3102
omitted Genes: P2ry6, P2ry2, Arap1
omitted Genes: Mir139, Art2a−ps, Art2b
omitted Genes: Clpb, Phox2a, Inppl1
omitted Genes: Folr2, Folr1, Anapc15
omitted Genes: Lamtor1, Lrrc51, Numa1
omitted Genes: Rnf121, Xntrpc, Trpc2
omitted Genes: Art1, Chrna10, Nup98
omitted Genes: Pgap2, Rhog, Stim1
omitted Genes: Rrm1, Olfr543, Olfr544
omitted Genes: Olfr545, Olfr547, Olfr549
omitted Genes: Olfr550, Olfr551, Olfr552
omitted Genes: Olfr553, Olfr554, Olfr555
omitted Genes: Olfr556, Trim68, Olfr557
omitted Genes: Olfr558, Olfr559, Olfr561
omitted Genes: Olfr564, Olfr566, Olfr568
omitted Genes: Olfr569, Olfr570, Olfr571
omitted Genes: Olfr572, Olfr574, Olfr575
omitted Genes: Olfr576, Olfr577, Olfr578
omitted Genes: Olfr582, Olfr583, Olfr584
omitted Genes: Olfr585, Olfr586, Olfr589
omitted Genes: Olfr591, Olfr592, Olfr593
omitted Genes: Olfr594, Usp17ld, Olfr597

Make more plots at http://csg.sph.umich.edu/locuszoom/

Fine−mapping Regions

| snp | chr | pos (Mb) | pp | group | color |
|---|---|---|---|---|---|
| chr7:101660383 | 7 | 101.6604 | 0.0009929434 | Credible Set | purple |
| chr7:101669186 | 7 | 101.6692 | 0.0010713666 | Credible Set | purple |
| chr7:101673483 | 7 | 101.6735 | 0.0012617061 | Credible Set | purple |
| chr7:101680140 | 7 | 101.6801 | 0.0011142265 | Credible Set | purple |
| chr7:101731061 | 7 | 101.7311 | 0.0010041356 | Credible Set | purple |
| chr7:101731110 | 7 | 101.7311 | 0.0010118919 | Credible Set | purple |
| chr7:101731130 | 7 | 101.7311 | 0.0011696765 | Credible Set | purple |
| chr7:101752971 | 7 | 101.7530 | 0.0009629975 | Credible Set | purple |
| chr7:101998679 | 7 | 101.9987 | 0.0017175575 | Credible Set | purple |
| chr7:102098054 | 7 | 102.0981 | 0.0131280765 | Credible Set | purple |
| chr7:102110109 | 7 | 102.1101 | 0.0109895379 | Credible Set | purple |
| chr7:102110501 | 7 | 102.1105 | 0.0112663096 | Credible Set | purple |
| chr7:102116318 | 7 | 102.1163 | 0.0120776507 | Credible Set | purple |
| chr7:102116430 | 7 | 102.1164 | 0.0120776507 | Credible Set | purple |
| chr7:102116478 | 7 | 102.1165 | 0.0125998291 | Credible Set | purple |
| chr7:102132562 | 7 | 102.1326 | 0.0116375091 | Credible Set | purple |
| chr7:102223396 | 7 | 102.2234 | 0.0130936504 | Credible Set | purple |
| chr7:102240695 | 7 | 102.2407 | 0.0106787116 | Credible Set | purple |
| chr7:102241191 | 7 | 102.2412 | 0.0037464787 | Credible Set | purple |
| chr7:102241211 | 7 | 102.2412 | 0.0034665230 | Credible Set | purple |
| chr7:102241212 | 7 | 102.2412 | 0.0034817014 | Credible Set | purple |
| chr7:102317213 | 7 | 102.3172 | 0.0071701726 | Credible Set | purple |
| chr7:102317246 | 7 | 102.3172 | 0.0049213778 | Credible Set | purple |
| chr7:102344970 | 7 | 102.3450 | 0.0161887131 | Credible Set | purple |
| chr7:102363897 | 7 | 102.3639 | 0.0120464049 | Credible Set | purple |
| chr7:102364204 | 7 | 102.3642 | 0.0177104280 | Credible Set | purple |
| chr7:102364223 | 7 | 102.3642 | 0.0212434342 | Credible Set | purple |
| chr7:102364315 | 7 | 102.3643 | 0.0152663070 | Credible Set | purple |
| chr7:102364326 | 7 | 102.3643 | 0.0152663070 | Credible Set | purple |
| chr7:102364411 | 7 | 102.3644 | 0.0109717078 | Credible Set | purple |
| chr7:102385520 | 7 | 102.3855 | 0.0089801886 | Credible Set | purple |
| chr7:102385536 | 7 | 102.3855 | 0.0080702215 | Credible Set | purple |
| chr7:102423165 | 7 | 102.4232 | 0.0119005748 | Credible Set | purple |
| chr7:102435692 | 7 | 102.4357 | 0.0080011307 | Credible Set | purple |
| chr7:102435720 | 7 | 102.4357 | 0.0080011307 | Credible Set | purple |
| chr7:102478469 | 7 | 102.4785 | 0.0134630192 | Credible Set | purple |

Fine−mapping Regions

| snp | chr | pos (Mb) | pp | group | color |
|---|---|---|---|---|---|
| chr7:102510475 | 7 | 102.5105 | 0.011760647 | Credible Set | purple |
| chr7:102510607 | 7 | 102.5106 | 0.011380137 | Credible Set | purple |
| chr7:102516878 | 7 | 102.5169 | 0.012505242 | Credible Set | purple |
| chr7:102531652 | 7 | 102.5317 | 0.012456754 | Credible Set | purple |
| chr7:102561708 | 7 | 102.5617 | 0.011694669 | Credible Set | purple |
| chr7:102565262 | 7 | 102.5653 | 0.031327334 | Credible Set | purple |
| chr7:102565292 | 7 | 102.5653 | 0.031346426 | Credible Set | purple |
| chr7:102565307 | 7 | 102.5653 | 0.031369672 | Credible Set | purple |
| chr7:102565323 | 7 | 102.5653 | 0.031388803 | Credible Set | purple |
| chr7:102565329 | 7 | 102.5653 | 0.031388803 | Credible Set | purple |
| chr7:102565410 | 7 | 102.5654 | 0.008295208 | Credible Set | purple |
| chr7:102596442 | 7 | 102.5964 | 0.016460981 | Credible Set | purple |
| chr7:102619418 | 7 | 102.6194 | 0.021930790 | Credible Set | purple |
| chr7:102619431 | 7 | 102.6194 | 0.021874623 | Credible Set | purple |
| chr7:102619466 | 7 | 102.6195 | 0.008212079 | Credible Set | purple |
| chr7:102757499 | 7 | 102.7575 | 0.008925640 | Credible Set | purple |
| chr7:103442570 | 7 | 103.4426 | 0.014881387 | Credible Set | purple |
| chr7:103564836 | 7 | 103.5648 | 0.015953655 | Credible Set | purple |
| chr7:103646439 | 7 | 103.6464 | 0.019486914 | Credible Set | purple |
| chr7:103646709 | 7 | 103.6467 | 0.020896343 | Credible Set | purple |
| chr7:103671033 | 7 | 103.6710 | 0.022759804 | Credible Set | purple |
| chr7:103760301 | 7 | 103.7603 | 0.038647383 | Credible Set | purple |
| chr7:103829200 | 7 | 103.8292 | 0.013434031 | Credible Set | purple |
| chr7:103837099 | 7 | 103.8371 | 0.020615481 | Credible Set | purple |
| chr7:103837213 | 7 | 103.8372 | 0.013790586 | Credible Set | purple |
| chr7:103915503 | 7 | 103.9155 | 0.009890892 | Credible Set | purple |
| chr7:103986792 | 7 | 103.9868 | 0.010600819 | Credible Set | purple |
| chr7:103986795 | 7 | 103.9868 | 0.010600819 | Credible Set | purple |
| chr7:103986857 | 7 | 103.9869 | 0.010024565 | Credible Set | purple |
| chr7:103986860 | 7 | 103.9869 | 0.010024565 | Credible Set | purple |
| chr7:104012808 | 7 | 104.0128 | 0.006829287 | Credible Set | purple |
| chr7:104013370 | 7 | 104.0134 | 0.009540389 | Credible Set | purple |
| chr7:104013395 | 7 | 104.0134 | 0.011583584 | Credible Set | purple |
| chr7:104013467 | 7 | 104.0135 | 0.009304569 | Credible Set | purple |
| chr7:104129734 | 7 | 104.1297 | 0.003174973 | Credible Set | purple |
| chr7:104129736 | 7 | 104.1297 | 0.003174973 | Credible Set | purple |

Fine−mapping Regions

| snp | chr | pos (Mb) | pp | group | color |
|---|---|---|---|---|---|
| chr7:104141343 | 7 | 104.1413 | 0.001317245 | Credible Set | purple |
| chr7:104141705 | 7 | 104.1417 | 0.002670671 | Credible Set | purple |
| chr7:104141768 | 7 | 104.1418 | 0.002679300 | Credible Set | purple |
| chr7:104170479 | 7 | 104.1705 | 0.001806270 | Credible Set | purple |
| chr7:104170506 | 7 | 104.1705 | 0.001812380 | Credible Set | purple |
| chr7:104170518 | 7 | 104.1705 | 0.001828490 | Credible Set | purple |
| chr7:104170522 | 7 | 104.1705 | 0.001814914 | Credible Set | purple |
| chr7:104170543 | 7 | 104.1705 | 0.001793815 | Credible Set | purple |
| chr7:104170588 | 7 | 104.1706 | 0.004880965 | Credible Set | purple |
| chr7:104170594 | 7 | 104.1706 | 0.004862902 | Credible Set | purple |
| chr7:104170618 | 7 | 104.1706 | 0.004913876 | Credible Set | purple |
| chr7:104170662 | 7 | 104.1707 | 0.004913876 | Credible Set | purple |
| chr7:104353491 | 7 | 104.3535 | 0.017811187 | Credible Set | purple |
| chr7:104356230 | 7 | 104.3562 | 0.011465954 | Credible Set | purple |
| chr7:104430205 | 7 | 104.4302 | 0.007067539 | Credible Set | purple |
| chr7:104485236 | 7 | 104.4852 | 0.004261719 | Credible Set | purple |
| chr7:104507724 | 7 | 104.5077 | 0.002328720 | Credible Set | purple |
| chr7:104508127 | 7 | 104.5081 | 0.002551765 | Credible Set | purple |
| chr7:104508173 | 7 | 104.5082 | 0.003282108 | Credible Set | purple |
| chr7:104580091 | 7 | 104.5801 | 0.011377779 | Credible Set | purple |
| chr7:105411752 | 7 | 105.4118 | 0.009416495 | Credible Set | purple |

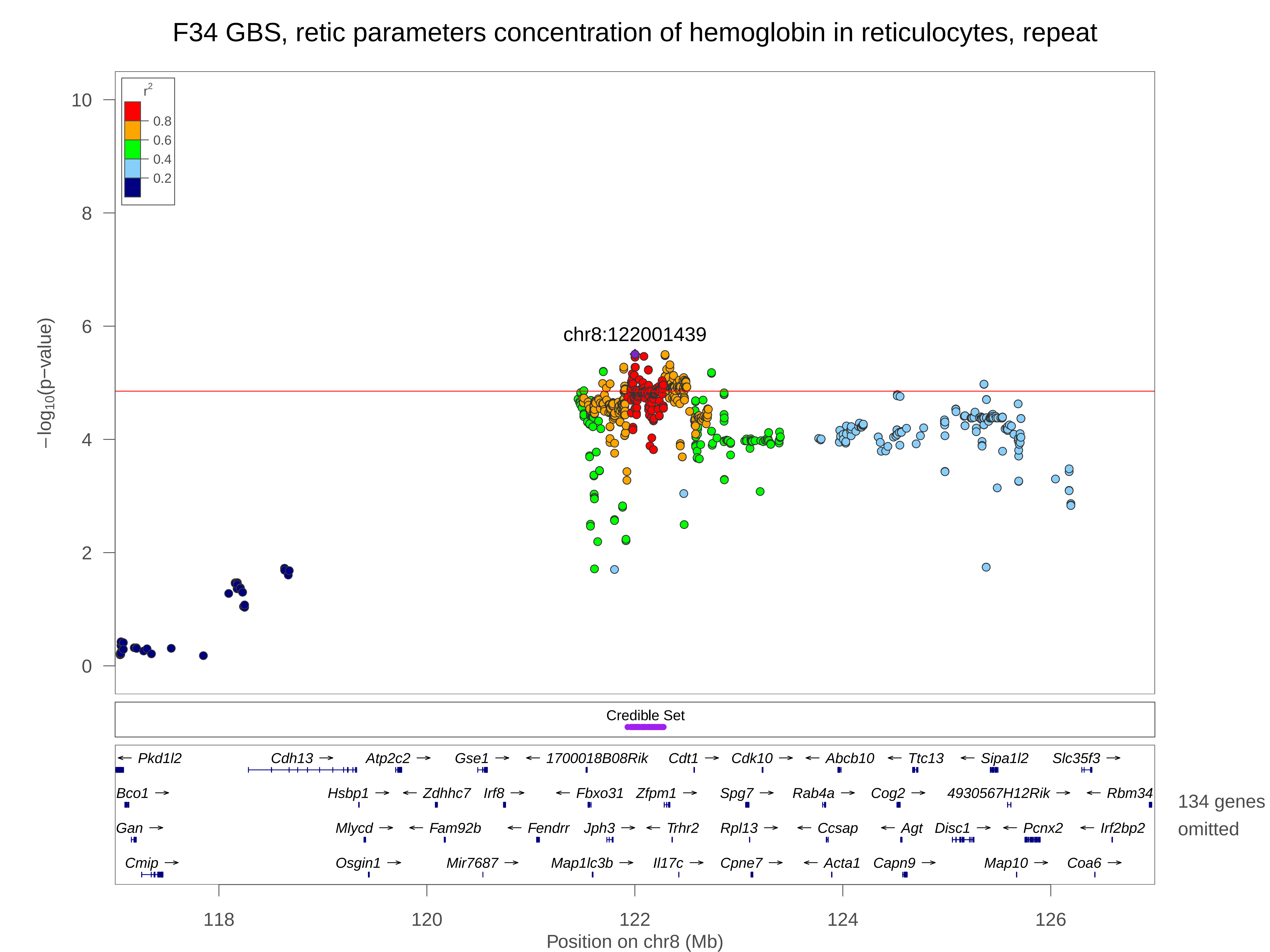
F34 GBS, retic parameters concentration of hemoglobin in reticulocytes, repeat
r2
0.8
0.6
0.4
0.2
10
8
6
4
2
0
−log10(p−value)
chr8:122001439
Credible Set
Pkd1l2
Cdh13
Atp2c2
Gse1
1700018B08Rik
Cdt1
Cdk10
Abcb10
Ttc13
Sipa1l2
Slc35f3
Bco1
Hsbp1
Zdhhc7
Irf8
Fbxo31
Zfpm1
Spg7
Rab4a
Cog2
4930567H12Rik
Rbm34
Gan
Mlycd
Fam92b
Fendrr
Jph3
Trhr2
Rpl13
Ccsap
Agt
Disc1
Pcnx2
Irf2bp2
Cmip
Osgin1
Mir7687
Map1lc3b
Il17c
Cpne7
Acta1
Capn9
Map10
Coa6
134 genes omitted
118
120
122
124
126
Position on chr8 (Mb)

date: Thu Oct 25 17:06:54 2018
build: F34GBS
display range: chr8:117001439−127001439 [117001439−127001439]
hilite range: 0 − 0 [ 0 − 0 ]
reference SNP: chr8:122001439
number of SNPs plotted: 886
min P−value: 3.12E−6 [chr8:122001439]
max P−value: 6.6E−1 [chr8:117850640]
omitted Genes: Mir7077, Mir7078, Plcg2
omitted Genes: Sdr42e1, Gm38416, Hsd17b2
omitted Genes: Mphosph6, Necab2, Slc38a8
omitted Genes: Mbtps1, Hsdl1, Dnaaf1
omitted Genes: Taf1c, Adad2, Kcng4
omitted Genes: Wfdc1, Tldc1, Cotl1
omitted Genes: Klhl36, Usp10, Crispld2
omitted Genes: 6430548M08Rik, A330074K22Rik, Gins2
omitted Genes: Emc8−1190005i06rik, 1190005I06Rik, Emc8
omitted Genes: Cox4i1, Foxf1, Mthfsd
omitted Genes: Foxc2, Foxl1, 1700030M09Rik
omitted Genes: Zcchc14, Klhdc4, Slc7a5
omitted Genes: BC048644, Car5a, Banp
omitted Genes: Mir7237, Zc3h18, Cyba
omitted Genes: Mvd, 9330133O14Rik, Gm20735
omitted Genes: Snai3, Rnf166, Ctu2
omitted Genes: Piezo1, Aprt, Galns
omitted Genes: Trappc2l, Pabpn1l, Mir6396
omitted Genes: Cbfa2t3, Acsf3, Cdh15
omitted Genes: Ankrd11, 2810013P06Rik, Snord68
omitted Genes: Mir7079, Mir7080, Sult5a1
omitted Genes: Dpep1, Chmp1a, Spata33
omitted Genes: Spata2l, 4933417D19Rik, Vps9d1
omitted Genes: Zfp276, Fanca, Spire2
omitted Genes: Tcf25, Mc1r, Tubb3
omitted Genes: Def8, Afg3l1, Dbndd1
omitted Genes: Gas8, n−R5s100, n−R5s103
omitted Genes: n−R5s104, n−R5s105, n−R5s108
omitted Genes: n−R5s110, n−R5s111, n−R5s113
omitted Genes: Gm25018, n−R5s117, n−R5s121
omitted Genes: n−R5s122, n−R5s123, n−R5s124
Make more plots at http://csg.sph.umich.edu/locuszoom/
omitted Genes: n−R5s128, Gm26391, n−R5s133

Fine−mapping Regions

| snp | chr | pos (Mb) | pp | group | color |
|---|---|---|---|---|---|
| chr8:121930880 | 8 | 121.9309 | 0.004949384 | Credible Set | purple |
| chr8:121930977 | 8 | 121.9310 | 0.004884971 | Credible Set | purple |
| chr8:121960554 | 8 | 121.9606 | 0.006118467 | Credible Set | purple |
| chr8:121961919 | 8 | 121.9619 | 0.006741756 | Credible Set | purple |
| chr8:121966290 | 8 | 121.9663 | 0.007481209 | Credible Set | purple |
| chr8:121966311 | 8 | 121.9663 | 0.002658600 | Credible Set | purple |
| chr8:121966312 | 8 | 121.9663 | 0.002684870 | Credible Set | purple |
| chr8:121966322 | 8 | 121.9663 | 0.002636304 | Credible Set | purple |
| chr8:121966639 | 8 | 121.9666 | 0.004326518 | Credible Set | purple |
| chr8:121966705 | 8 | 121.9667 | 0.008656549 | Credible Set | purple |
| chr8:121966756 | 8 | 121.9668 | 0.008911267 | Credible Set | purple |
| chr8:121966990 | 8 | 121.9670 | 0.007370962 | Credible Set | purple |
| chr8:121966991 | 8 | 121.9670 | 0.006903072 | Credible Set | purple |
| chr8:121966992 | 8 | 121.9670 | 0.006856297 | Credible Set | purple |
| chr8:121974244 | 8 | 121.9742 | 0.008208380 | Credible Set | purple |
| chr8:121974249 | 8 | 121.9742 | 0.008226306 | Credible Set | purple |
| chr8:121974305 | 8 | 121.9743 | 0.008161931 | Credible Set | purple |
| chr8:121974325 | 8 | 121.9743 | 0.008179746 | Credible Set | purple |
| chr8:121976235 | 8 | 121.9762 | 0.010063614 | Credible Set | purple |
| chr8:121976517 | 8 | 121.9765 | 0.012085593 | Credible Set | purple |
| chr8:121976521 | 8 | 121.9765 | 0.012129693 | Credible Set | purple |
| chr8:121981546 | 8 | 121.9815 | 0.008211202 | Credible Set | purple |
| chr8:121995055 | 8 | 121.9951 | 0.004449220 | Credible Set | purple |
| chr8:121995078 | 8 | 121.9951 | 0.004578666 | Credible Set | purple |
| chr8:121995079 | 8 | 121.9951 | 0.004883014 | Credible Set | purple |
| chr8:121995687 | 8 | 121.9957 | 0.011467249 | Credible Set | purple |
| chr8:122001362 | 8 | 122.0014 | 0.015662249 | Credible Set | purple |
| chr8:122001439 | 8 | 122.0014 | 0.025989359 | Credible Set | purple |
| chr8:122001442 | 8 | 122.0014 | 0.023059665 | Credible Set | purple |
| chr8:122001818 | 8 | 122.0018 | 0.005741075 | Credible Set | purple |
| chr8:122001981 | 8 | 122.0020 | 0.006054676 | Credible Set | purple |
| chr8:122002025 | 8 | 122.0020 | 0.003179842 | Credible Set | purple |
| chr8:122002067 | 8 | 122.0021 | 0.003179842 | Credible Set | purple |
| chr8:122002101 | 8 | 122.0021 | 0.003179842 | Credible Set | purple |
| chr8:122002127 | 8 | 122.0021 | 0.006030626 | Credible Set | purple |
| chr8:122002218 | 8 | 122.0022 | 0.006030626 | Credible Set | purple |

Fine−mapping Regions

| snp | chr | pos (Mb) | pp | group | color |
|---|---|---|---|---|---|
| chr8:122003523 | 8 | 122.0035 | 0.006279445 | Credible Set | purple |
| chr8:122003541 | 8 | 122.0035 | 0.006449517 | Credible Set | purple |
| chr8:122003853 | 8 | 122.0039 | 0.005918232 | Credible Set | purple |
| chr8:122003945 | 8 | 122.0039 | 0.006484190 | Credible Set | purple |
| chr8:122005311 | 8 | 122.0053 | 0.005339823 | Credible Set | purple |
| chr8:122005336 | 8 | 122.0053 | 0.005339823 | Credible Set | purple |
| chr8:122005348 | 8 | 122.0053 | 0.015674070 | Credible Set | purple |
| chr8:122010526 | 8 | 122.0105 | 0.006030626 | Credible Set | purple |
| chr8:122015785 | 8 | 122.0158 | 0.005369289 | Credible Set | purple |
| chr8:122015943 | 8 | 122.0159 | 0.002622184 | Credible Set | purple |
| chr8:122015966 | 8 | 122.0160 | 0.002472665 | Credible Set | purple |
| chr8:122018214 | 8 | 122.0182 | 0.005907470 | Credible Set | purple |
| chr8:122018241 | 8 | 122.0182 | 0.005545572 | Credible Set | purple |
| chr8:122018758 | 8 | 122.0188 | 0.003254292 | Credible Set | purple |
| chr8:122020302 | 8 | 122.0203 | 0.005191434 | Credible Set | purple |
| chr8:122023460 | 8 | 122.0235 | 0.006425334 | Credible Set | purple |
| chr8:122024035 | 8 | 122.0240 | 0.004310108 | Credible Set | purple |
| chr8:122026904 | 8 | 122.0269 | 0.005462509 | Credible Set | purple |
| chr8:122031170 | 8 | 122.0312 | 0.004907366 | Credible Set | purple |
| chr8:122032377 | 8 | 122.0324 | 0.005793253 | Credible Set | purple |
| chr8:122032681 | 8 | 122.0327 | 0.005848699 | Credible Set | purple |
| chr8:122035610 | 8 | 122.0356 | 0.005222523 | Credible Set | purple |
| chr8:122035650 | 8 | 122.0357 | 0.005191240 | Credible Set | purple |
| chr8:122035881 | 8 | 122.0359 | 0.005787015 | Credible Set | purple |
| chr8:122036007 | 8 | 122.0360 | 0.005672751 | Credible Set | purple |
| chr8:122036452 | 8 | 122.0365 | 0.006023736 | Credible Set | purple |
| chr8:122045025 | 8 | 122.0450 | 0.009645882 | Credible Set | purple |
| chr8:122045040 | 8 | 122.0450 | 0.006332939 | Credible Set | purple |
| chr8:122065233 | 8 | 122.0652 | 0.005731392 | Credible Set | purple |
| chr8:122065534 | 8 | 122.0655 | 0.005767254 | Credible Set | purple |
| chr8:122065719 | 8 | 122.0657 | 0.005978678 | Credible Set | purple |
| chr8:122074319 | 8 | 122.0743 | 0.005785234 | Credible Set | purple |
| chr8:122074380 | 8 | 122.0744 | 0.005804973 | Credible Set | purple |
| chr8:122074409 | 8 | 122.0744 | 0.005804973 | Credible Set | purple |
| chr8:122075721 | 8 | 122.0757 | 0.007299219 | Credible Set | purple |
| chr8:122077089 | 8 | 122.0771 | 0.005975461 | Credible Set | purple |

Fine−mapping Regions

| snp | chr | pos (Mb) | pp | group | color |
|---|---|---|---|---|---|
| chr8:122077746 | 8 | 122.0777 | 0.005863109 | Credible Set | purple |
| chr8:122078705 | 8 | 122.0787 | 0.007793853 | Credible Set | purple |
| chr8:122078711 | 8 | 122.0787 | 0.008626213 | Credible Set | purple |
| chr8:122087021 | 8 | 122.0870 | 0.006740003 | Credible Set | purple |
| chr8:122087024 | 8 | 122.0870 | 0.006992745 | Credible Set | purple |
| chr8:122087068 | 8 | 122.0871 | 0.023857465 | Credible Set | purple |
| chr8:122087488 | 8 | 122.0875 | 0.006142591 | Credible Set | purple |
| chr8:122097829 | 8 | 122.0978 | 0.004729330 | Credible Set | purple |
| chr8:122097864 | 8 | 122.0979 | 0.004718729 | Credible Set | purple |
| chr8:122097914 | 8 | 122.0979 | 0.004718729 | Credible Set | purple |
| chr8:122098026 | 8 | 122.0980 | 0.005968375 | Credible Set | purple |
| chr8:122098264 | 8 | 122.0983 | 0.006030626 | Credible Set | purple |
| chr8:122098555 | 8 | 122.0986 | 0.006134521 | Credible Set | purple |
| chr8:122099136 | 8 | 122.0991 | 0.005595918 | Credible Set | purple |
| chr8:122099269 | 8 | 122.0993 | 0.006116376 | Credible Set | purple |
| chr8:122099407 | 8 | 122.0994 | 0.005954153 | Credible Set | purple |
| chr8:122119850 | 8 | 122.1198 | 0.006174390 | Credible Set | purple |
| chr8:122126372 | 8 | 122.1264 | 0.005781633 | Credible Set | purple |
| chr8:122126643 | 8 | 122.1266 | 0.005675720 | Credible Set | purple |
| chr8:122126657 | 8 | 122.1267 | 0.005643272 | Credible Set | purple |
| chr8:122126675 | 8 | 122.1267 | 0.005655547 | Credible Set | purple |
| chr8:122129779 | 8 | 122.1298 | 0.014201591 | Credible Set | purple |
| chr8:122129781 | 8 | 122.1298 | 0.013935105 | Credible Set | purple |
| chr8:122132490 | 8 | 122.1325 | 0.007783198 | Credible Set | purple |
| chr8:122132563 | 8 | 122.1326 | 0.003411458 | Credible Set | purple |
| chr8:122132629 | 8 | 122.1326 | 0.003026721 | Credible Set | purple |
| chr8:122132657 | 8 | 122.1327 | 0.003039609 | Credible Set | purple |
| chr8:122132660 | 8 | 122.1327 | 0.002988825 | Credible Set | purple |
| chr8:122132700 | 8 | 122.1327 | 0.002998279 | Credible Set | purple |
| chr8:122140064 | 8 | 122.1401 | 0.002430579 | Credible Set | purple |
| chr8:122140617 | 8 | 122.1406 | 0.004967122 | Credible Set | purple |
| chr8:122140668 | 8 | 122.1407 | 0.003327326 | Credible Set | purple |
| chr8:122143194 | 8 | 122.1432 | 0.004096054 | Credible Set | purple |
| chr8:122143198 | 8 | 122.1432 | 0.003989126 | Credible Set | purple |
| chr8:122143647 | 8 | 122.1436 | 0.006227860 | Credible Set | purple |
| chr8:122161138 | 8 | 122.1611 | 0.004311153 | Credible Set | purple |

Fine−mapping Regions

| snp | chr | pos (Mb) | pp | group | color |
|---|---|---|---|---|---|
| chr8:122161157 | 8 | 122.1612 | 0.005598723 | Credible Set | purple |
| chr8:122161598 | 8 | 122.1616 | 0.002221463 | Credible Set | purple |
| chr8:122161622 | 8 | 122.1616 | 0.002230187 | Credible Set | purple |
| chr8:122161625 | 8 | 122.1616 | 0.002206512 | Credible Set | purple |
| chr8:122161665 | 8 | 122.1617 | 0.006208114 | Credible Set | purple |
| chr8:122161683 | 8 | 122.1617 | 0.006208114 | Credible Set | purple |
| chr8:122161737 | 8 | 122.1617 | 0.006208114 | Credible Set | purple |
| chr8:122163079 | 8 | 122.1631 | 0.003866526 | Credible Set | purple |
| chr8:122163130 | 8 | 122.1631 | 0.002933719 | Credible Set | purple |
| chr8:122178957 | 8 | 122.1790 | 0.004595634 | Credible Set | purple |
| chr8:122179254 | 8 | 122.1793 | 0.002057829 | Credible Set | purple |
| chr8:122179449 | 8 | 122.1794 | 0.005394876 | Credible Set | purple |
| chr8:122179473 | 8 | 122.1795 | 0.005412446 | Credible Set | purple |
| chr8:122185374 | 8 | 122.1854 | 0.002147277 | Credible Set | purple |
| chr8:122185458 | 8 | 122.1855 | 0.005866642 | Credible Set | purple |
| chr8:122186770 | 8 | 122.1868 | 0.005470198 | Credible Set | purple |
| chr8:122186780 | 8 | 122.1868 | 0.005649025 | Credible Set | purple |
| chr8:122192954 | 8 | 122.1930 | 0.005758913 | Credible Set | purple |
| chr8:122193526 | 8 | 122.1935 | 0.005802336 | Credible Set | purple |
| chr8:122193537 | 8 | 122.1935 | 0.005826553 | Credible Set | purple |
| chr8:122197166 | 8 | 122.1972 | 0.005812274 | Credible Set | purple |
| chr8:122197187 | 8 | 122.1972 | 0.006483091 | Credible Set | purple |
| chr8:122210438 | 8 | 122.2104 | 0.006972384 | Credible Set | purple |
| chr8:122211178 | 8 | 122.2112 | 0.006122677 | Credible Set | purple |
| chr8:122222775 | 8 | 122.2228 | 0.006030626 | Credible Set | purple |
| chr8:122223010 | 8 | 122.2230 | 0.003135265 | Credible Set | purple |
| chr8:122223079 | 8 | 122.2231 | 0.003072647 | Credible Set | purple |
| chr8:122224941 | 8 | 122.2249 | 0.006214862 | Credible Set | purple |
| chr8:122233190 | 8 | 122.2332 | 0.007433642 | Credible Set | purple |
| chr8:122235012 | 8 | 122.2350 | 0.006592882 | Credible Set | purple |
| chr8:122235350 | 8 | 122.2353 | 0.002366069 | Credible Set | purple |
| chr8:122235368 | 8 | 122.2354 | 0.002348658 | Credible Set | purple |
| chr8:122235394 | 8 | 122.2354 | 0.002400298 | Credible Set | purple |
| chr8:122236097 | 8 | 122.2361 | 0.004212245 | Credible Set | purple |
| chr8:122237812 | 8 | 122.2378 | 0.006386226 | Credible Set | purple |
| chr8:122237817 | 8 | 122.2378 | 0.006368699 | Credible Set | purple |

Fine−mapping Regions

| snp | chr | pos (Mb) | pp | group | color |
|---|---|---|---|---|---|
| chr8:122240202 | 8 | 122.2402 | 0.006331737 | Credible Set | purple |
| chr8:122240296 | 8 | 122.2403 | 0.005980477 | Credible Set | purple |
| chr8:122246861 | 8 | 122.2469 | 0.006659267 | Credible Set | purple |
| chr8:122259808 | 8 | 122.2598 | 0.007559009 | Credible Set | purple |
| chr8:122262163 | 8 | 122.2622 | 0.009314276 | Credible Set | purple |
| chr8:122262214 | 8 | 122.2622 | 0.007610542 | Credible Set | purple |
| chr8:122262948 | 8 | 122.2629 | 0.005477178 | Credible Set | purple |
| chr8:122265327 | 8 | 122.2653 | 0.007413144 | Credible Set | purple |
| chr8:122265814 | 8 | 122.2658 | 0.006834875 | Credible Set | purple |
| chr8:122265865 | 8 | 122.2659 | 0.006839467 | Credible Set | purple |
| chr8:122271333 | 8 | 122.2713 | 0.007855758 | Credible Set | purple |
| chr8:122273143 | 8 | 122.2731 | 0.007807822 | Credible Set | purple |
| chr8:122273765 | 8 | 122.2738 | 0.003506738 | Credible Set | purple |
| chr8:122274216 | 8 | 122.2742 | 0.003266308 | Credible Set | purple |
| chr8:122275678 | 8 | 122.2757 | 0.003175994 | Credible Set | purple |
| chr8:122275695 | 8 | 122.2757 | 0.003198128 | Credible Set | purple |

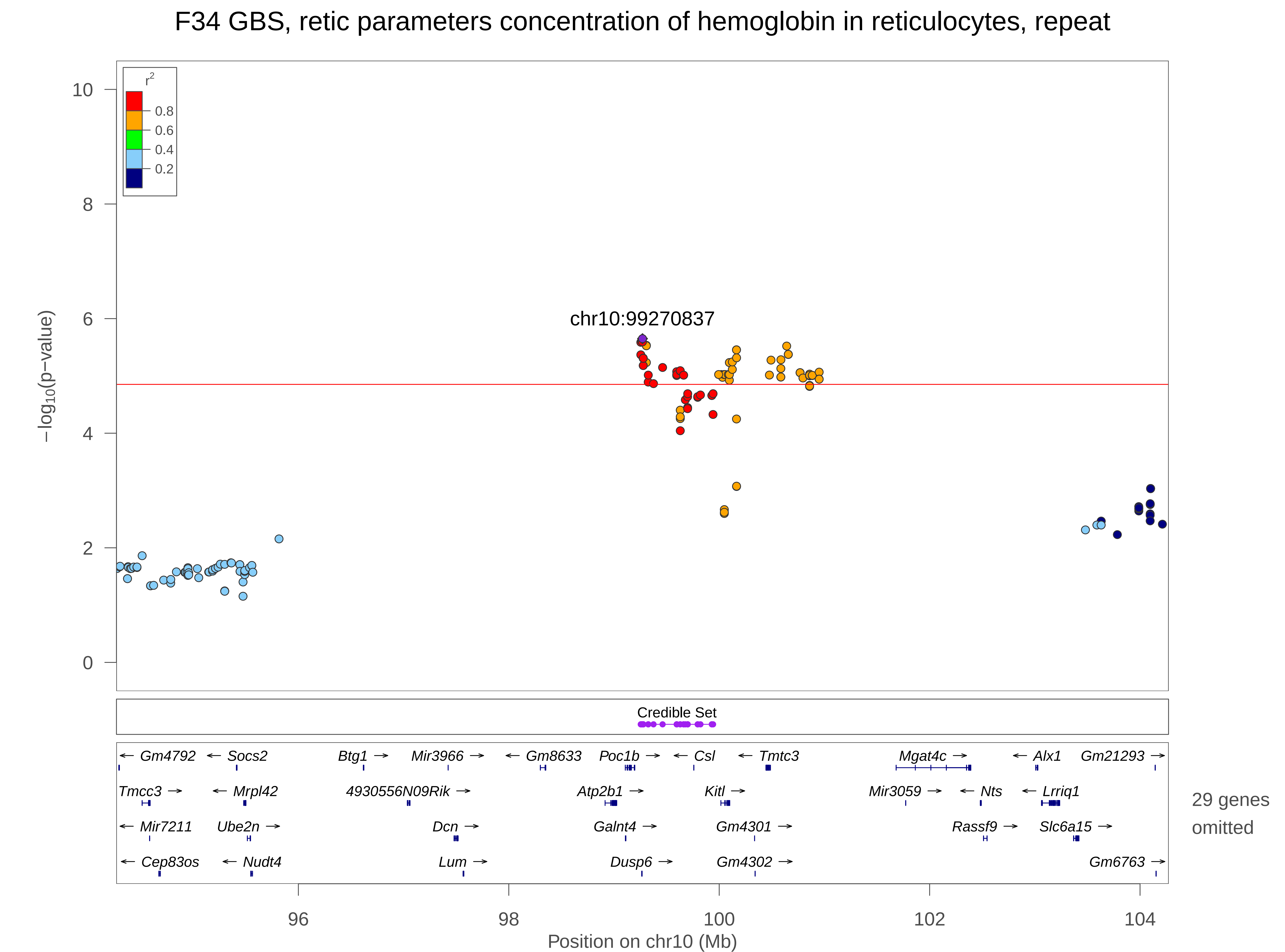

F34 GBS, retic parameters concentration of hemoglobin in reticulocytes, repeat
r2
0.8
0.6
0.4
0.2
10
8
6
4
2
0
−log10(p−value)
chr10:99270837
Credible Set
← Gm4792
← Socs2
Btg1 →
Mir3966 →
← Gm8633
Poc1b →
← Csl
← Tmtc3
Mgat4c →
← Alx1
Gm21293 →
Tmcc3 →
← Mrpl42
4930556N09Rik →
Atp2b1 →
Kitl →
Mir3059 →
← Nts
← Lrriq1
← Mir7211
Ube2n →
Dcn →
Galnt4 →
Gm4301 →
Rassf9 →
Slc6a15 →
← Cep83os
← Nudt4
Lum →
Dusp6 →
Gm4302 →
Gm6763 →
29 genes omitted
96
98
100
102
104
Position on chr10 (Mb)

| | |
|---|---|
| date: | Thu Oct 25 17:06:08 2018 |
| build: | F34GBS |
| display range: | chr10:94270837−104270837 [94270837−104270837] |
| hilite range: | 0 − 0 [ 0 − 0 ] |
| reference SNP: | chr10:99270837 |
| number of SNPs plotted: | 184 |
| min P−value: | 2.24E−6 [chr10:99270837] |
| max P−value: | 7E−2 [chr10:95474045] |
| omitted Genes: | Cep83, Plxnc1, Gm29684 |
| omitted Genes: | Cradd, 2310039L15Rik, 5730420D15Rik |
| omitted Genes: | Mir3058, Eea1, 4930459C07Rik |
| omitted Genes: | 4930473O22Rik, Kera, Epyc |
| omitted Genes: | Gm10754, Ccer1, 4930525C09Rik |
| omitted Genes: | Gm20110, Gm18409, Gm4303 |
| omitted Genes: | Gm4305, Gm4307, Gm4312 |
| omitted Genes: | Cep290, 4930430F08Rik, 1700017N19Rik |
| omitted Genes: | Gm8764, Gm21304, Gm21312 |
| omitted Genes: | Gm20765, Gm4340 |

Make more plots at http://csg.sph.umich.edu/locuszoom/

Fine−mapping Regions

| snp | chr | pos (Mb) | pp | group | color |
|---|---|---|---|---|---|
| chr10:99254791 | 10 | 99.25479 | 0.065606388 | Credible Set | purple |
| chr10:99254830 | 10 | 99.25483 | 0.049790182 | Credible Set | purple |
| chr10:99263835 | 10 | 99.26384 | 0.096825995 | Credible Set | purple |
| chr10:99263868 | 10 | 99.26387 | 0.096825995 | Credible Set | purple |
| chr10:99270417 | 10 | 99.27042 | 0.065606388 | Credible Set | purple |
| chr10:99270837 | 10 | 99.27084 | 0.096825995 | Credible Set | purple |
| chr10:99277813 | 10 | 99.27781 | 0.029137103 | Credible Set | purple |
| chr10:99277816 | 10 | 99.27782 | 0.029137103 | Credible Set | purple |
| chr10:99277972 | 10 | 99.27797 | 0.040206862 | Credible Set | purple |
| chr10:99277973 | 10 | 99.27797 | 0.040206862 | Credible Set | purple |
| chr10:99278026 | 10 | 99.27803 | 0.040206862 | Credible Set | purple |
| chr10:99325139 | 10 | 99.32514 | 0.016129540 | Credible Set | purple |
| chr10:99325722 | 10 | 99.32572 | 0.020720109 | Credible Set | purple |
| chr10:99375519 | 10 | 99.37552 | 0.015028662 | Credible Set | purple |
| chr10:99461777 | 10 | 99.46178 | 0.029137103 | Credible Set | purple |
| chr10:99596119 | 10 | 99.59612 | 0.020720109 | Credible Set | purple |
| chr10:99596142 | 10 | 99.59614 | 0.025644468 | Credible Set | purple |
| chr10:99596164 | 10 | 99.59616 | 0.022914216 | Credible Set | purple |
| chr10:99629121 | 10 | 99.62912 | 0.025644468 | Credible Set | purple |
| chr10:99629701 | 10 | 99.62970 | 0.025644468 | Credible Set | purple |
| chr10:99661770 | 10 | 99.66177 | 0.020720109 | Credible Set | purple |
| chr10:99661829 | 10 | 99.66183 | 0.020720109 | Credible Set | purple |
| chr10:99678204 | 10 | 99.67820 | 0.008334112 | Credible Set | purple |
| chr10:99699271 | 10 | 99.69927 | 0.009365149 | Credible Set | purple |
| chr10:99699649 | 10 | 99.69965 | 0.006117200 | Credible Set | purple |
| chr10:99699681 | 10 | 99.69968 | 0.005960117 | Credible Set | purple |
| chr10:99700646 | 10 | 99.70065 | 0.010697830 | Credible Set | purple |
| chr10:99795502 | 10 | 99.79550 | 0.008993465 | Credible Set | purple |
| chr10:99795653 | 10 | 99.79565 | 0.009365149 | Credible Set | purple |
| chr10:99821621 | 10 | 99.82162 | 0.010212225 | Credible Set | purple |
| chr10:99929872 | 10 | 99.92987 | 0.009769853 | Credible Set | purple |
| chr10:99930010 | 10 | 99.93001 | 0.009769853 | Credible Set | purple |
| chr10:99941475 | 10 | 99.94147 | 0.010697830 | Credible Set | purple |

F34 GBS, retic parameters concentration of hemoglobin in reticulocytes, repeat

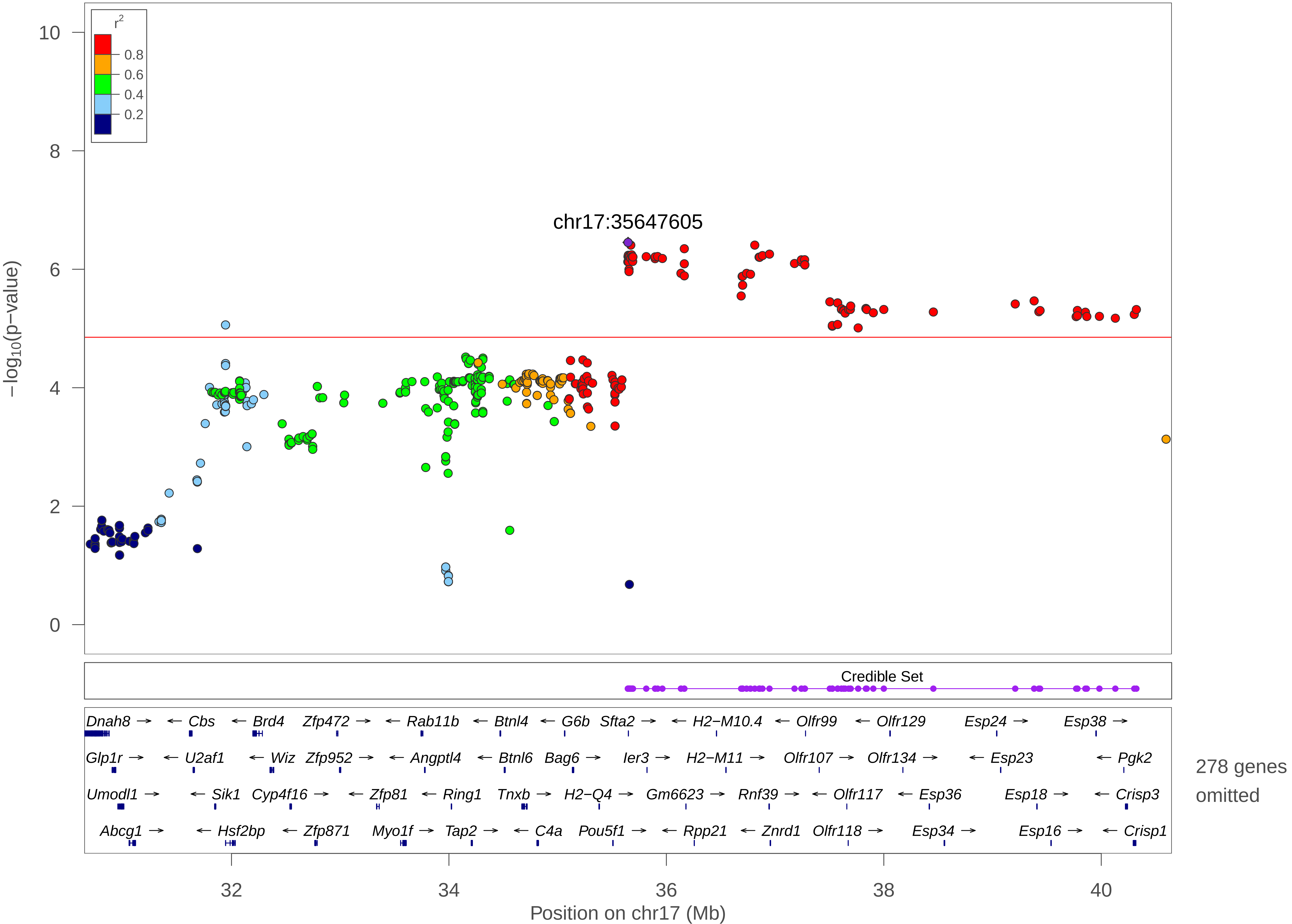

date: Thu Oct 25 17:06:23 2018
build: F34GBS
display range: chr17:30647605−40647605 [30647605−40647605]
hilite range: 0 − 0 [ 0 − 0 ]
reference SNP: chr17:35647605
number of SNPs plotted: 518
min P−value: 3.52E−7 [chr17:35647605]
max P−value: 2.09E−1 [chr17:35659711]
omitted Genes: Tff3, Tff2, LOC102631757
omitted Genes: Tff1, Tmprss3, Ubash3a
omitted Genes: Rsph1, Slc37a1, Pde9a
omitted Genes: Wdr4, Ndufv3, Pknox1
omitted Genes: Cryaa, Rrp1b, Pdxk−ps
omitted Genes: Notch3, Ephx3, Gm4432
omitted Genes: Akap8, Akap8l, Rasal3
omitted Genes: A530088E08Rik, Pglyrp2, Cyp4f39
omitted Genes: Cyp4f17, Cyp4f37, Cyp4f40
omitted Genes: Cyp4f15, Zfp811, Zfp799
omitted Genes: Zfp870, Cyp4f14, Cyp4f13
omitted Genes: Cyp4f41−ps, Zfp763, 4921501E09Rik
omitted Genes: Zfp563, Morc2b, Olfr55
omitted Genes: Olfr239, Gm4461, Zfp955a
omitted Genes: Olfr63, Zfp955b, Zfp101
omitted Genes: Actl9, Adamts10, Zfp414
omitted Genes: Pram1, Hnrnpm, March2
omitted Genes: Kank3, Rps28, Ndufa7
omitted Genes: Cd320, Kifc1, Gm27740
omitted Genes: Platr17, BC051226, Daxx
omitted Genes: Zbtb22, Tapbp, Rgl2
omitted Genes: Pfdn6, Wdr46, B3galt4
omitted Genes: Rps18, Vps52, H2−K2
omitted Genes: AA388235, H2−K1, Mir219a−1
omitted Genes: Mir219c, H2−Ke6, Slc39a7
omitted Genes: Rxrb, Col11a2, BC051537
omitted Genes: H2−Oa, Brd2, H2−DMa
omitted Genes: H2−DMb2, H2−DMb1, Psmb9
omitted Genes: Tap1, Psmb8, H2−Ob
omitted Genes: H2−Ab1, H2−Aa, H2−Eb1
omitted Genes: H2−Eb2, Btnl2, Btnl1
Make more plots at http://csg.sph.umich.edu/locuszoom/
omitted Genes: BC051142, Btnl5−ps, Notch4

Fine−mapping Regions

| snp | chr | pos (Mb) | pp | group | color |
|---|---|---|---|---|---|
| chr17:35645542 | 17 | 35.64554 | 0.015959227 | Credible Set | purple |
| chr17:35647446 | 17 | 35.64745 | 0.019403871 | Credible Set | purple |
| chr17:35647569 | 17 | 35.64757 | 0.019403871 | Credible Set | purple |
| chr17:35647605 | 17 | 35.64760 | 0.032938940 | Credible Set | purple |
| chr17:35647647 | 17 | 35.64765 | 0.032864243 | Credible Set | purple |
| chr17:35649675 | 17 | 35.64968 | 0.020347768 | Credible Set | purple |
| chr17:35653842 | 17 | 35.65384 | 0.015835848 | Credible Set | purple |
| chr17:35656227 | 17 | 35.65623 | 0.011953328 | Credible Set | purple |
| chr17:35656237 | 17 | 35.65624 | 0.011049252 | Credible Set | purple |
| chr17:35657043 | 17 | 35.65704 | 0.019403871 | Credible Set | purple |
| chr17:35659591 | 17 | 35.65959 | 0.019403871 | Credible Set | purple |
| chr17:35659790 | 17 | 35.65979 | 0.019285867 | Credible Set | purple |
| chr17:35661925 | 17 | 35.66192 | 0.020261720 | Credible Set | purple |
| chr17:35661953 | 17 | 35.66195 | 0.020235492 | Credible Set | purple |
| chr17:35662044 | 17 | 35.66204 | 0.019403871 | Credible Set | purple |
| chr17:35671886 | 17 | 35.67189 | 0.018244197 | Credible Set | purple |
| chr17:35672608 | 17 | 35.67261 | 0.029769228 | Credible Set | purple |
| chr17:35680171 | 17 | 35.68017 | 0.020699720 | Credible Set | purple |
| chr17:35685462 | 17 | 35.68546 | 0.019287142 | Credible Set | purple |
| chr17:35690117 | 17 | 35.69012 | 0.016166071 | Credible Set | purple |
| chr17:35692366 | 17 | 35.69237 | 0.019114064 | Credible Set | purple |
| chr17:35814797 | 17 | 35.81480 | 0.019338733 | Credible Set | purple |
| chr17:35896309 | 17 | 35.89631 | 0.017933541 | Credible Set | purple |
| chr17:35896336 | 17 | 35.89634 | 0.019157881 | Credible Set | purple |
| chr17:35919532 | 17 | 35.91953 | 0.019403871 | Credible Set | purple |
| chr17:35963419 | 17 | 35.96342 | 0.018046527 | Credible Set | purple |
| chr17:36134373 | 17 | 36.13437 | 0.010351821 | Credible Set | purple |
| chr17:36134388 | 17 | 36.13439 | 0.010351821 | Credible Set | purple |
| chr17:36164938 | 17 | 36.16494 | 0.026020321 | Credible Set | purple |
| chr17:36164968 | 17 | 36.16497 | 0.014785564 | Credible Set | purple |
| chr17:36165054 | 17 | 36.16505 | 0.009423360 | Credible Set | purple |
| chr17:36687888 | 17 | 36.68789 | 0.004445141 | Credible Set | purple |
| chr17:36696968 | 17 | 36.69697 | 0.009233952 | Credible Set | purple |
| chr17:36701296 | 17 | 36.70130 | 0.006640878 | Credible Set | purple |
| chr17:36701348 | 17 | 36.70135 | 0.006652416 | Credible Set | purple |
| chr17:36701690 | 17 | 36.70169 | 0.009295105 | Credible Set | purple |

Fine−mapping Regions

| snp | chr | pos (Mb) | pp | group | color |
|---|---|---|---|---|---|
| chr17:36737772 | 17 | 36.73777 | 0.010347145 | Credible Set | purple |
| chr17:36773709 | 17 | 36.77371 | 0.009993692 | Credible Set | purple |
| chr17:36813020 | 17 | 36.81302 | 0.029796440 | Credible Set | purple |
| chr17:36852594 | 17 | 36.85259 | 0.018974525 | Credible Set | purple |
| chr17:36857085 | 17 | 36.85708 | 0.018820602 | Credible Set | purple |
| chr17:36858903 | 17 | 36.85890 | 0.018974525 | Credible Set | purple |
| chr17:36858955 | 17 | 36.85896 | 0.018974525 | Credible Set | purple |
| chr17:36883876 | 17 | 36.88388 | 0.020112576 | Credible Set | purple |
| chr17:36948500 | 17 | 36.94850 | 0.021247355 | Credible Set | purple |
| chr17:37178457 | 17 | 37.17846 | 0.014950300 | Credible Set | purple |
| chr17:37241973 | 17 | 37.24197 | 0.015828467 | Credible Set | purple |
| chr17:37242059 | 17 | 37.24206 | 0.017167668 | Credible Set | purple |
| chr17:37271534 | 17 | 37.27153 | 0.017227009 | Credible Set | purple |
| chr17:37273627 | 17 | 37.27363 | 0.014371745 | Credible Set | purple |
| chr17:37273628 | 17 | 37.27363 | 0.014141616 | Credible Set | purple |
| chr17:37273636 | 17 | 37.27364 | 0.014230016 | Credible Set | purple |
| chr17:37503566 | 17 | 37.50357 | 0.003575854 | Credible Set | purple |
| chr17:37526332 | 17 | 37.52633 | 0.001443878 | Credible Set | purple |
| chr17:37526408 | 17 | 37.52641 | 0.001448747 | Credible Set | purple |
| chr17:37526413 | 17 | 37.52641 | 0.001480667 | Credible Set | purple |
| chr17:37574863 | 17 | 37.57486 | 0.001537708 | Credible Set | purple |
| chr17:37575322 | 17 | 37.57532 | 0.003429711 | Credible Set | purple |
| chr17:37575334 | 17 | 37.57533 | 0.003429711 | Credible Set | purple |
| chr17:37575341 | 17 | 37.57534 | 0.003429711 | Credible Set | purple |
| chr17:37575344 | 17 | 37.57534 | 0.003429711 | Credible Set | purple |
| chr17:37610173 | 17 | 37.61017 | 0.002685950 | Credible Set | purple |
| chr17:37610452 | 17 | 37.61045 | 0.002748952 | Credible Set | purple |
| chr17:37631225 | 17 | 37.63123 | 0.002599248 | Credible Set | purple |
| chr17:37631428 | 17 | 37.63143 | 0.002545401 | Credible Set | purple |
| chr17:37644696 | 17 | 37.64470 | 0.002348741 | Credible Set | purple |
| chr17:37676096 | 17 | 37.67610 | 0.002685890 | Credible Set | purple |
| chr17:37692132 | 17 | 37.69213 | 0.002685890 | Credible Set | purple |
| chr17:37695484 | 17 | 37.69548 | 0.003058205 | Credible Set | purple |
| chr17:37763996 | 17 | 37.76400 | 0.001353188 | Credible Set | purple |
| chr17:37835420 | 17 | 37.83542 | 0.002791536 | Credible Set | purple |
| chr17:37835736 | 17 | 37.83574 | 0.002756783 | Credible Set | purple |

Fine−mapping Regions

| snp | chr | pos (Mb) | pp | group | color |
|---|---|---|---|---|---|
| chr17:37842881 | 17 | 37.84288 | 0.002680426 | Credible Set | purple |
| chr17:37903355 | 17 | 37.90335 | 0.002376706 | Credible Set | purple |
| chr17:37999788 | 17 | 37.99979 | 0.002685890 | Credible Set | purple |
| chr17:38455586 | 17 | 38.45559 | 0.002444826 | Credible Set | purple |
| chr17:39208793 | 17 | 39.20879 | 0.003297060 | Credible Set | purple |
| chr17:39382632 | 17 | 39.38263 | 0.003698378 | Credible Set | purple |
| chr17:39426246 | 17 | 39.42625 | 0.002470239 | Credible Set | purple |
| chr17:39437308 | 17 | 39.43731 | 0.002579331 | Credible Set | purple |
| chr17:39769349 | 17 | 39.76935 | 0.002059838 | Credible Set | purple |
| chr17:39780605 | 17 | 39.78061 | 0.002591185 | Credible Set | purple |
| chr17:39780934 | 17 | 39.78093 | 0.002126763 | Credible Set | purple |
| chr17:39853992 | 17 | 39.85399 | 0.002427523 | Credible Set | purple |
| chr17:39867231 | 17 | 39.86723 | 0.002069375 | Credible Set | purple |
| chr17:39983068 | 17 | 39.98307 | 0.002078934 | Credible Set | purple |
| chr17:40129553 | 17 | 40.12955 | 0.001939989 | Credible Set | purple |
| chr17:40303781 | 17 | 40.30378 | 0.002233659 | Credible Set | purple |
| chr17:40322995 | 17 | 40.32299 | 0.002678436 | Credible Set | purple |

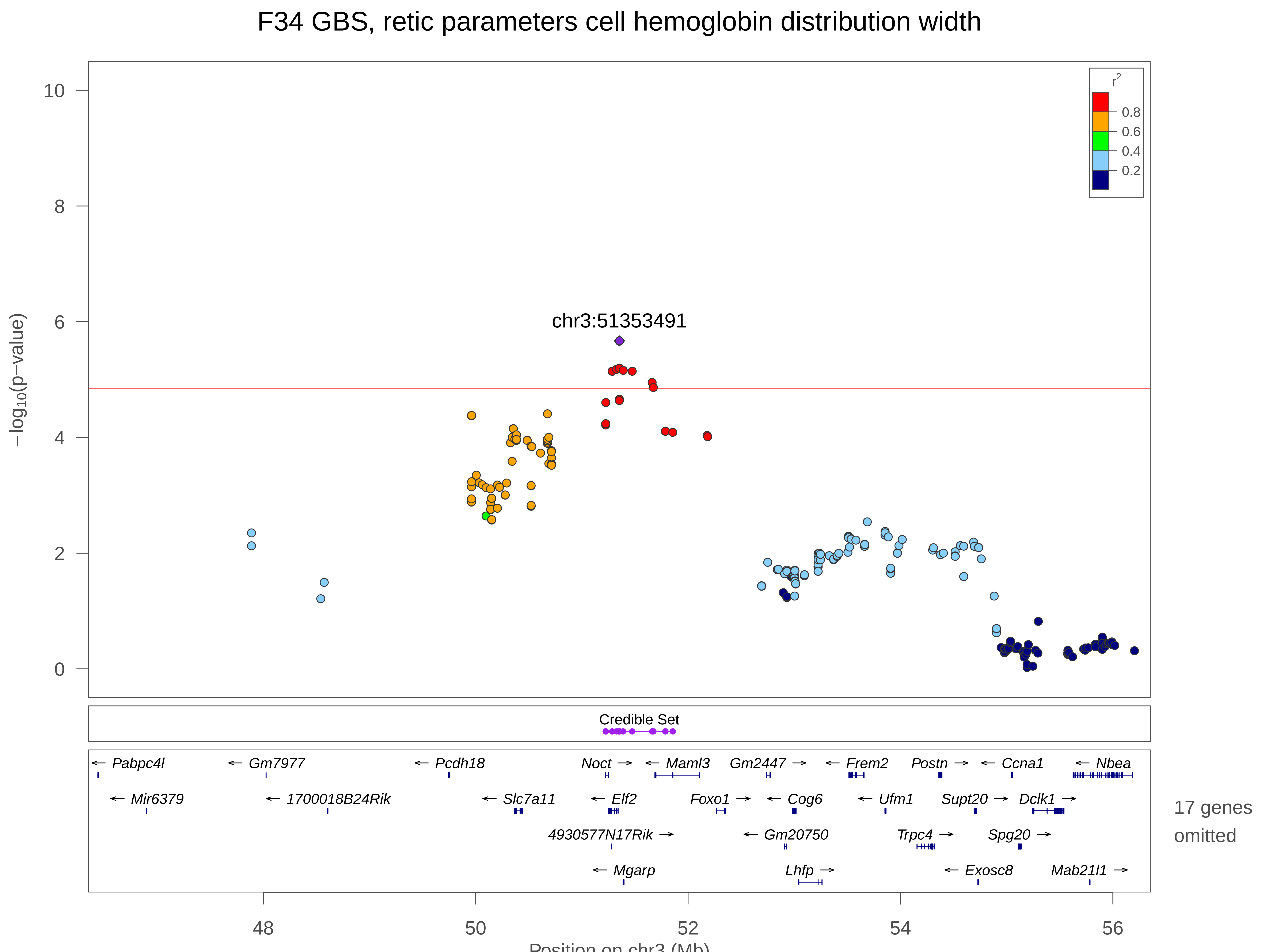
F34 GBS, retic parameters cell hemoglobin distribution width
chr3:51353491
r2
0.8
0.6
0.4
0.2
−log10(p−value)
10
8
6
4
2
0
Credible Set
← Pabpc4l
← Gm7977
← Pcdh18
Noct →
← Maml3
Gm2447 →
← Frem2
Postn →
← Ccna1
← Nbea
← Mir6379
← 1700018B24Rik
← Slc7a11
← Elf2
Foxo1 →
← Cog6
← Ufm1
Supt20 →
Dclk1 →
4930577N17Rik →
← Gm20750
Trpc4 →
Spg20 →
← Mgarp
Lhfp →
← Exosc8
Mab21l1 →
17 genes omitted
48
50
52
54
56
Position on chr3 (Mb)

| | |
|---|---|
| date: | Thu Oct 25 17:07:29 2018 |
| build: | F34GBS |
| display range: | chr3:46353491−56353491 [46353491−56353491] |
| hilite range: | 0 − 0 [ 0 − 0 ] |
| reference SNP: | chr3:51353491 |
| number of SNPs plotted: | 283 |
| min P−value: | 2.14E−6 [chr3:51353491] |
| max P−value: | 9.46E−1 [chr3:55192362] |
| omitted Genes: | Ndufc1, Naa15, Rab33b |
| omitted Genes: | Setd7, 5031434O11Rik, Mgst2 |
| omitted Genes: | Nhlrc3, Proser1, Stoml3 |
| omitted Genes: | Alg5, Smad9, Rfxap |
| omitted Genes: | Sertm1, 4931419H13Rik, Ccdc169 |
| omitted Genes: | Sohlh2, 4933417G07Rik |

Make more plots at http://csg.sph.umich.edu/locuszoom/

Fine−mapping Regions

| snp | chr | pos (Mb) | pp | group | color |
|---|---|---|---|---|---|
| chr3:51224536 | 3 | 51.22454 | 0.006549880 | Credible Set | purple |
| chr3:51224538 | 3 | 51.22454 | 0.006862850 | Credible Set | purple |
| chr3:51224542 | 3 | 51.22454 | 0.006880732 | Credible Set | purple |
| chr3:51224949 | 3 | 51.22495 | 0.015320677 | Credible Set | purple |
| chr3:51284978 | 3 | 51.28498 | 0.050073355 | Credible Set | purple |
| chr3:51285004 | 3 | 51.28500 | 0.050469758 | Credible Set | purple |
| chr3:51326496 | 3 | 51.32650 | 0.053967627 | Credible Set | purple |
| chr3:51353269 | 3 | 51.35327 | 0.056741272 | Credible Set | purple |
| chr3:51353361 | 3 | 51.35336 | 0.017369834 | Credible Set | purple |
| chr3:51353422 | 3 | 51.35342 | 0.016532821 | Credible Set | purple |
| chr3:51353461 | 3 | 51.35346 | 0.159007360 | Credible Set | purple |
| chr3:51353490 | 3 | 51.35349 | 0.158323691 | Credible Set | purple |
| chr3:51353491 | 3 | 51.35349 | 0.159783900 | Credible Set | purple |
| chr3:51353741 | 3 | 51.35374 | 0.056139059 | Credible Set | purple |
| chr3:51389257 | 3 | 51.38926 | 0.051905555 | Credible Set | purple |
| chr3:51473939 | 3 | 51.47394 | 0.050270008 | Credible Set | purple |
| chr3:51661337 | 3 | 51.66134 | 0.032781117 | Credible Set | purple |
| chr3:51673564 | 3 | 51.67356 | 0.027059421 | Credible Set | purple |
| chr3:51785724 | 3 | 51.78572 | 0.005167142 | Credible Set | purple |
| chr3:51786977 | 3 | 51.78698 | 0.005170855 | Credible Set | purple |
| chr3:51855671 | 3 | 51.85567 | 0.004975115 | Credible Set | purple |

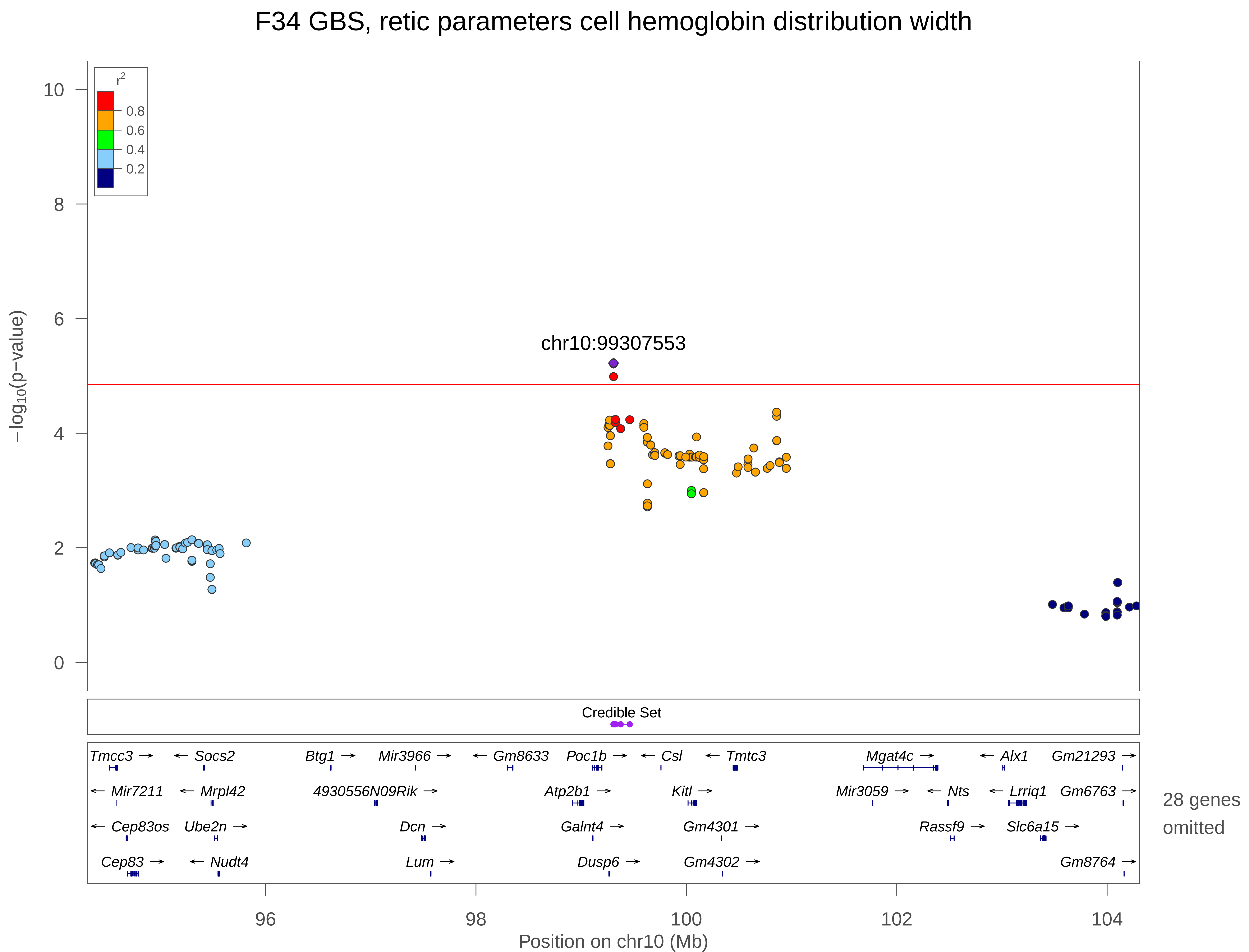

F34 GBS, retic parameters cell hemoglobin distribution width
r2
0.8
0.6
0.4
0.2
10
8
6
4
2
0
−log10(p−value)
chr10:99307553
Credible Set
Tmcc3 →
← Socs2
Btg1 →
Mir3966 →
← Gm8633
Poc1b →
← Csl
← Tmtc3
Mgat4c →
← Alx1
Gm21293 →
← Mir7211
← Mrpl42
4930556N09Rik →
Atp2b1 →
Kitl →
Mir3059 →
← Nts
← Lrriq1
Gm6763 →
← Cep83os
Ube2n →
Dcn →
Galnt4 →
Gm4301 →
Rassf9 →
Slc6a15 →
Cep83 →
← Nudt4
Lum →
Dusp6 →
Gm4302 →
Gm8764 →
28 genes omitted
96
98
100
102
104
Position on chr10 (Mb)

| | |
|---|---|
| date: | Thu Oct 25 17:07:06 2018 |
| build: | F34GBS |
| display range: | chr10:94307553−104307553 [94307553−104307553] |
| hilite range: | 0 − 0 [ 0 − 0 ] |
| reference SNP: | chr10:99307553 |
| number of SNPs plotted: | 180 |
| min P−value: | 5.98E−6 [chr10:99307553] |
| max P−value: | 1.57E−1 [chr10:103988206] |
| omitted Genes: | Plxnc1, Gm29684, Cradd |
| omitted Genes: | 2310039L15Rik, 5730420D15Rik, Mir3058 |
| omitted Genes: | Eea1, 4930459C07Rik, 4930473O22Rik |
| omitted Genes: | Kera, Epyc, Gm10754 |
| omitted Genes: | Ccer1, 4930525C09Rik, Gm20110 |
| omitted Genes: | Gm18409, Gm4303, Gm4305 |
| omitted Genes: | Gm4307, Gm4312, Cep290 |
| omitted Genes: | 4930430F08Rik, 1700017N19Rik, Gm21304 |
| omitted Genes: | Gm21312, Gm20765, Gm4340 |
| omitted Genes: | Mir6411 |

Make more plots at http://csg.sph.umich.edu/locuszoom/

Fine−mapping Regions

| snp | chr | pos (Mb) | pp | group | color |
|---|---|---|---|---|---|
| chr10:99307553 | 10 | 99.30755 | 0.33002136 | Credible Set | purple |
| chr10:99307573 | 10 | 99.30757 | 0.33002136 | Credible Set | purple |
| chr10:99307596 | 10 | 99.30760 | 0.20250564 | Credible Set | purple |
| chr10:99325139 | 10 | 99.32514 | 0.03365487 | Credible Set | purple |
| chr10:99325722 | 10 | 99.32572 | 0.03866595 | Credible Set | purple |
| chr10:99375519 | 10 | 99.37552 | 0.02709656 | Credible Set | purple |
| chr10:99461777 | 10 | 99.46178 | 0.03803425 | Credible Set | purple |

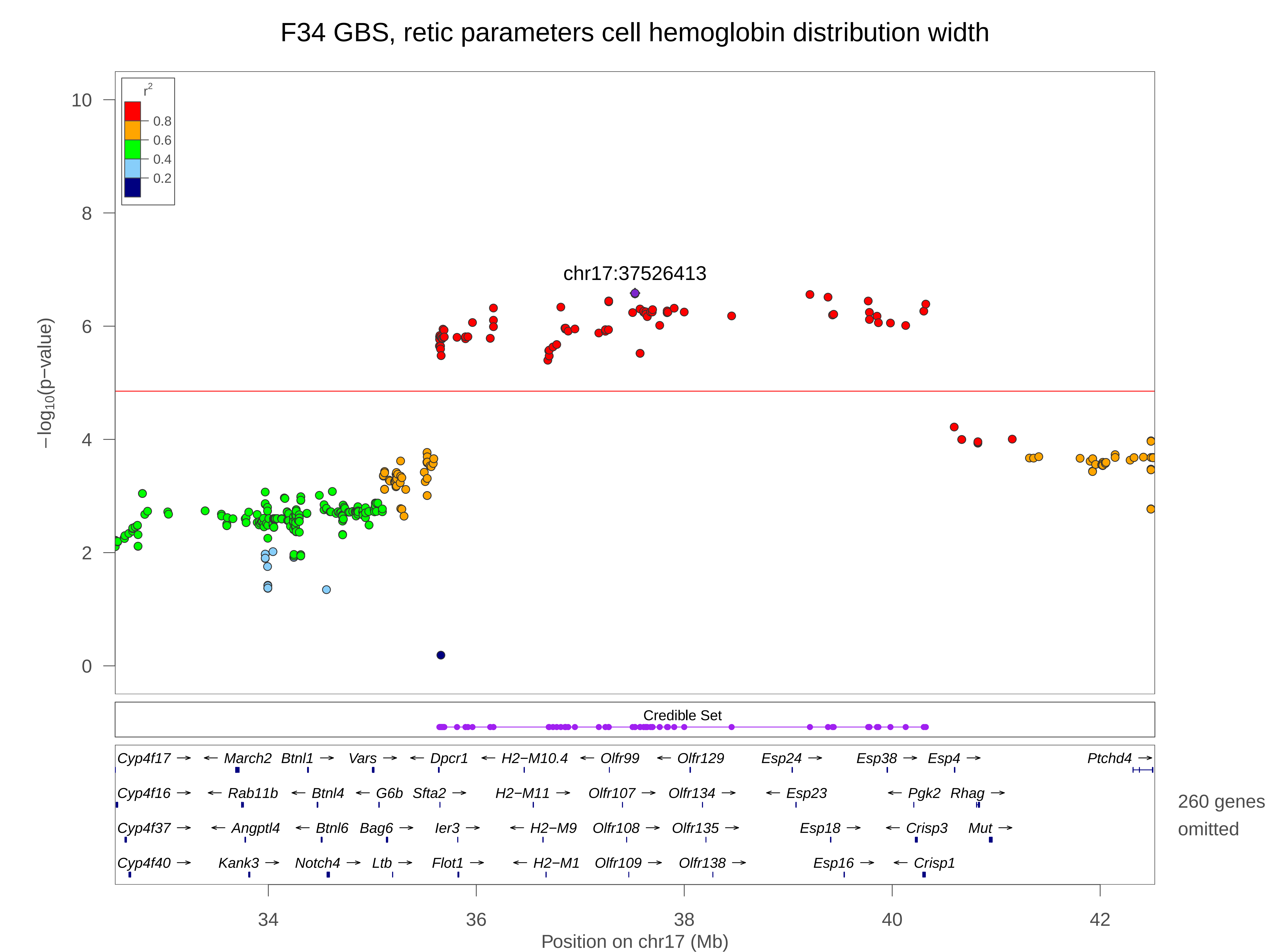
F34 GBS, retic parameters cell hemoglobin distribution width
r2
0.8
0.6
0.4
0.2
chr17:37526413
−log10(p−value)
0
2
4
6
8
10
Credible Set
Cyp4f17 →
← March2
Btnl1 →
Vars →
← Dpcr1
← H2−M10.4
← Olfr99
← Olfr129
Esp24 →
Esp38 →
Esp4 →
Ptchd4 →
Cyp4f16 →
← Rab11b
← Btnl4
← G6b
Sfta2 →
H2−M11 →
Olfr107 →
Olfr134 →
← Esp23
← Pgk2
Rhag →
Cyp4f37 →
← Angptl4
← Btnl6
Bag6 →
Ier3 →
← H2−M9
Olfr108 →
Olfr135 →
Esp18 →
← Crisp3
Mut →
Cyp4f40 →
Kank3 →
Notch4 →
Ltb →
Flot1 →
← H2−M1
Olfr109 →
Olfr138 →
Esp16 →
← Crisp1
260 genes omitted
34
36
38
40
42
Position on chr17 (Mb)

date: Thu Oct 25 17:07:20 2018
build: F34GBS
display range: chr17:32526413−42526413 [32526413−42526413]
hilite range: 0 − 0 [ 0 − 0 ]
reference SNP: chr17:37526413
number of SNPs plotted: 428
min P−value: 2.61E−7 [chr17:37526413]
max P−value: 6.46E−1 [chr17:35659711]
omitted Genes: Cyp4f15, Zfp871, Zfp811
omitted Genes: Zfp799, Zfp870, Cyp4f14
omitted Genes: Cyp4f13, Cyp4f41−ps, Zfp472
omitted Genes: Zfp952, Zfp763, 4921501E09Rik
omitted Genes: Zfp563, Morc2b, Olfr55
omitted Genes: Olfr239, Gm4461, Zfp955a
omitted Genes: Olfr63, Zfp955b, Zfp81
omitted Genes: Zfp101, Actl9, Adamts10
omitted Genes: Myo1f, Zfp414, Pram1
omitted Genes: Hnrnpm, Rps28, Ndufa7
omitted Genes: Cd320, Kifc1, Gm27740
omitted Genes: Platr17, BC051226, Daxx
omitted Genes: Zbtb22, Tapbp, Rgl2
omitted Genes: Pfdn6, Wdr46, B3galt4
omitted Genes: Rps18, Vps52, H2−K2
omitted Genes: AA388235, H2−K1, Ring1
omitted Genes: Mir219a−1, Mir219c, H2−Ke6
omitted Genes: Slc39a7, Rxrb, Col11a2
omitted Genes: BC051537, H2−Oa, Brd2
omitted Genes: H2−DMa, H2−DMb2, H2−DMb1
omitted Genes: Psmb9, Tap1, Psmb8
omitted Genes: Tap2, H2−Ob, H2−Ab1
omitted Genes: H2−Aa, H2−Eb1, H2−Eb2
omitted Genes: Btnl2, BC051142, Btnl5−ps
omitted Genes: Gpsm3, Pbx2, Ager
omitted Genes: Rnf5, Agpat1, Egfl8
omitted Genes: Ppt2, Prrt1, Fkbpl
omitted Genes: Atf6b, Tnxb, C4b
omitted Genes: Cyp21a1, C4a, Stk19
omitted Genes: Dxo, Skiv2l, Mir6971
omitted Genes: Mir6970, Nelfe, Cfb
Make more plots at http://csg.sph.umich.edu/locuszoom/
omitted Genes: Mir6972, C2, Zbtb12

Fine−mapping Regions

| snp | chr | pos (Mb) | pp | group | color |
|---|---|---|---|---|---|
| chr17:35645542 | 17 | 35.64554 | 0.004231237 | Credible Set | purple |
| chr17:35647446 | 17 | 35.64745 | 0.006073973 | Credible Set | purple |
| chr17:35647569 | 17 | 35.64757 | 0.006073973 | Credible Set | purple |
| chr17:35647605 | 17 | 35.64760 | 0.005547333 | Credible Set | purple |
| chr17:35647647 | 17 | 35.64765 | 0.005475530 | Credible Set | purple |
| chr17:35649675 | 17 | 35.64968 | 0.006384669 | Credible Set | purple |
| chr17:35653842 | 17 | 35.65384 | 0.003890164 | Credible Set | purple |
| chr17:35656227 | 17 | 35.65623 | 0.004226838 | Credible Set | purple |
| chr17:35656237 | 17 | 35.65624 | 0.003767926 | Credible Set | purple |
| chr17:35657043 | 17 | 35.65704 | 0.006073973 | Credible Set | purple |
| chr17:35659591 | 17 | 35.65959 | 0.006073973 | Credible Set | purple |
| chr17:35659790 | 17 | 35.65979 | 0.006053563 | Credible Set | purple |
| chr17:35661925 | 17 | 35.66192 | 0.002905546 | Credible Set | purple |
| chr17:35661953 | 17 | 35.66195 | 0.002908672 | Credible Set | purple |
| chr17:35662044 | 17 | 35.66204 | 0.006073973 | Credible Set | purple |
| chr17:35671886 | 17 | 35.67189 | 0.005621736 | Credible Set | purple |
| chr17:35672608 | 17 | 35.67261 | 0.005780892 | Credible Set | purple |
| chr17:35680171 | 17 | 35.68017 | 0.008183165 | Credible Set | purple |
| chr17:35685462 | 17 | 35.68546 | 0.006015166 | Credible Set | purple |
| chr17:35690117 | 17 | 35.69012 | 0.007886011 | Credible Set | purple |
| chr17:35692366 | 17 | 35.69237 | 0.006019554 | Credible Set | purple |
| chr17:35814797 | 17 | 35.81480 | 0.005943021 | Credible Set | purple |
| chr17:35896309 | 17 | 35.89631 | 0.005615817 | Credible Set | purple |
| chr17:35896336 | 17 | 35.89634 | 0.006072269 | Credible Set | purple |
| chr17:35919532 | 17 | 35.91953 | 0.006073973 | Credible Set | purple |
| chr17:35963419 | 17 | 35.96342 | 0.010599888 | Credible Set | purple |
| chr17:36134373 | 17 | 36.13437 | 0.005718160 | Credible Set | purple |
| chr17:36134388 | 17 | 36.13439 | 0.005718160 | Credible Set | purple |
| chr17:36164938 | 17 | 36.16494 | 0.018723323 | Credible Set | purple |
| chr17:36164968 | 17 | 36.16497 | 0.011608797 | Credible Set | purple |
| chr17:36165054 | 17 | 36.16505 | 0.008997770 | Credible Set | purple |
| chr17:36696968 | 17 | 36.69697 | 0.003519335 | Credible Set | purple |
| chr17:36701690 | 17 | 36.70169 | 0.003606588 | Credible Set | purple |
| chr17:36737772 | 17 | 36.73777 | 0.004082546 | Credible Set | purple |
| chr17:36773709 | 17 | 36.77371 | 0.004483641 | Credible Set | purple |
| chr17:36813020 | 17 | 36.81302 | 0.019401763 | Credible Set | purple |

Fine−mapping Regions

| snp | chr | pos (Mb) | pp | group | color |
|---|---|---|---|---|---|
| chr17:36852594 | 17 | 36.85259 | 0.008506659 | Credible Set | purple |
| chr17:36857085 | 17 | 36.85708 | 0.008186433 | Credible Set | purple |
| chr17:36858903 | 17 | 36.85890 | 0.008506659 | Credible Set | purple |
| chr17:36858955 | 17 | 36.85896 | 0.008506659 | Credible Set | purple |
| chr17:36883876 | 17 | 36.88388 | 0.007572203 | Credible Set | purple |
| chr17:36948500 | 17 | 36.94850 | 0.008241514 | Credible Set | purple |
| chr17:37178457 | 17 | 37.17846 | 0.007036269 | Credible Set | purple |
| chr17:37241973 | 17 | 37.24197 | 0.007529506 | Credible Set | purple |
| chr17:37242059 | 17 | 37.24206 | 0.008017601 | Credible Set | purple |
| chr17:37271534 | 17 | 37.27153 | 0.008004896 | Credible Set | purple |
| chr17:37273627 | 17 | 37.27363 | 0.024742759 | Credible Set | purple |
| chr17:37273628 | 17 | 37.27363 | 0.023882663 | Credible Set | purple |
| chr17:37273636 | 17 | 37.27364 | 0.024462362 | Credible Set | purple |
| chr17:37503566 | 17 | 37.50357 | 0.015651385 | Credible Set | purple |
| chr17:37526332 | 17 | 37.52633 | 0.032673092 | Credible Set | purple |
| chr17:37526408 | 17 | 37.52641 | 0.033201054 | Credible Set | purple |
| chr17:37526413 | 17 | 37.52641 | 0.033618526 | Credible Set | purple |
| chr17:37574863 | 17 | 37.57486 | 0.018025221 | Credible Set | purple |
| chr17:37575322 | 17 | 37.57532 | 0.003180852 | Credible Set | purple |
| chr17:37575334 | 17 | 37.57533 | 0.003180852 | Credible Set | purple |
| chr17:37575341 | 17 | 37.57534 | 0.003180852 | Credible Set | purple |
| chr17:37575344 | 17 | 37.57534 | 0.003180852 | Credible Set | purple |
| chr17:37610173 | 17 | 37.61017 | 0.015890002 | Credible Set | purple |
| chr17:37610452 | 17 | 37.61045 | 0.016510792 | Credible Set | purple |
| chr17:37631225 | 17 | 37.63123 | 0.016137959 | Credible Set | purple |
| chr17:37631428 | 17 | 37.63143 | 0.014707317 | Credible Set | purple |
| chr17:37644696 | 17 | 37.64470 | 0.013352743 | Credible Set | purple |
| chr17:37676096 | 17 | 37.67610 | 0.015988342 | Credible Set | purple |
| chr17:37692132 | 17 | 37.69213 | 0.015988342 | Credible Set | purple |
| chr17:37695484 | 17 | 37.69548 | 0.017572581 | Credible Set | purple |
| chr17:37763996 | 17 | 37.76400 | 0.009488744 | Credible Set | purple |
| chr17:37835420 | 17 | 37.83542 | 0.015582281 | Credible Set | purple |
| chr17:37835736 | 17 | 37.83574 | 0.016721185 | Credible Set | purple |
| chr17:37842881 | 17 | 37.84288 | 0.015929333 | Credible Set | purple |
| chr17:37903355 | 17 | 37.90335 | 0.018594999 | Credible Set | purple |
| chr17:37999788 | 17 | 37.99979 | 0.015988342 | Credible Set | purple |

Fine−mapping Regions

| snp | chr | pos (Mb) | pp | group | color |
|---|---|---|---|---|---|
| chr17:38455586 | 17 | 38.45559 | 0.013732756 | Credible Set | purple |
| chr17:39208793 | 17 | 39.20879 | 0.031852409 | Credible Set | purple |
| chr17:39382632 | 17 | 39.38263 | 0.028676486 | Credible Set | purple |
| chr17:39426246 | 17 | 39.42625 | 0.014247864 | Credible Set | purple |
| chr17:39437308 | 17 | 39.43731 | 0.014662066 | Credible Set | purple |
| chr17:39769349 | 17 | 39.76935 | 0.024653296 | Credible Set | purple |
| chr17:39780605 | 17 | 39.78061 | 0.015799966 | Credible Set | purple |
| chr17:39780934 | 17 | 39.78093 | 0.011937208 | Credible Set | purple |
| chr17:39853992 | 17 | 39.85399 | 0.013589919 | Credible Set | purple |
| chr17:39867231 | 17 | 39.86723 | 0.010518197 | Credible Set | purple |
| chr17:39983068 | 17 | 39.98307 | 0.010399961 | Credible Set | purple |
| chr17:40129553 | 17 | 40.12955 | 0.009439661 | Credible Set | purple |
| chr17:40303781 | 17 | 40.30378 | 0.016558534 | Credible Set | purple |
| chr17:40322995 | 17 | 40.32299 | 0.021851332 | Credible Set | purple |

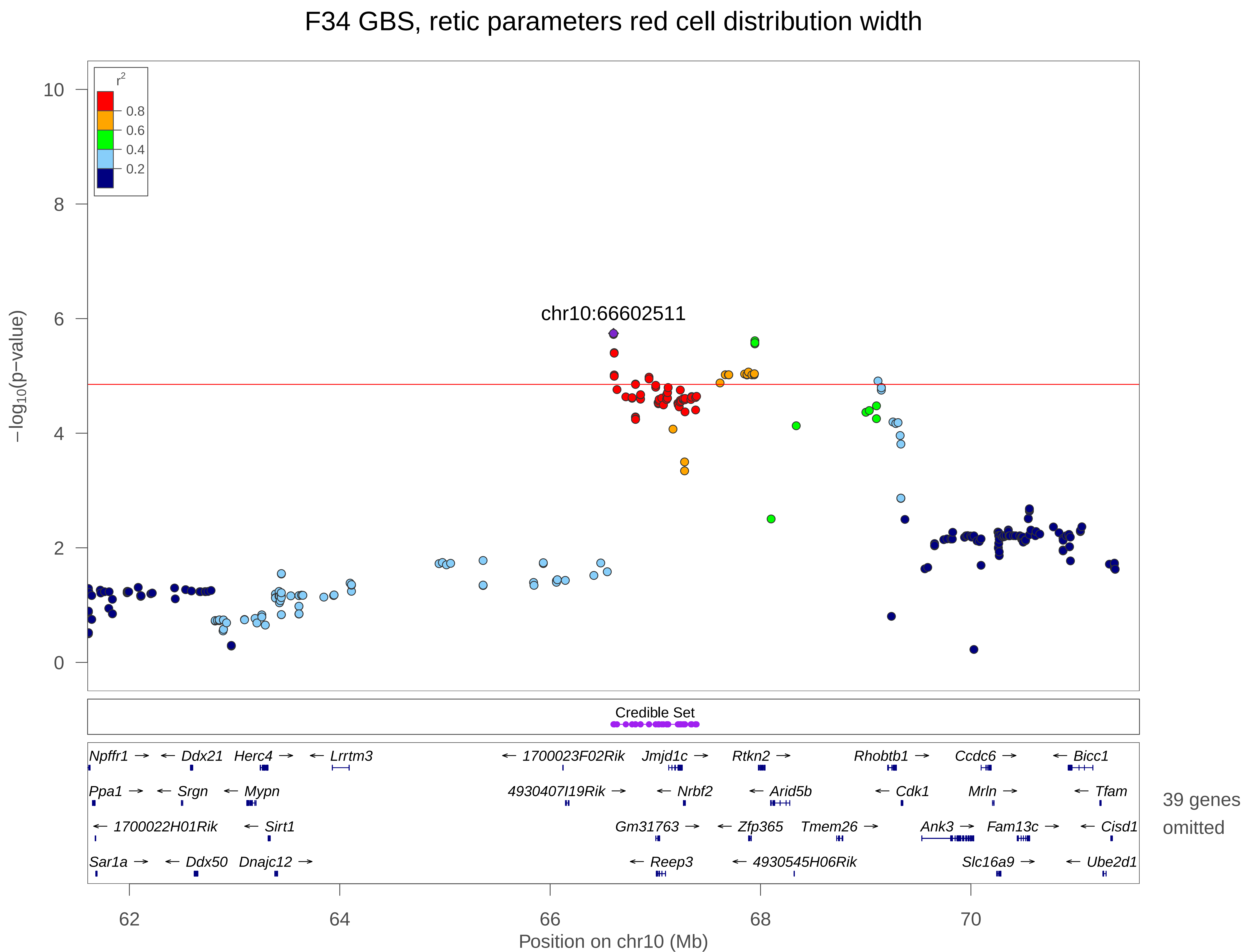

F34 GBS, retic parameters red cell distribution width
r2
0.8
0.6
0.4
0.2
10
8
6
4
2
0
−log10(p−value)
chr10:66602511
Credible Set
Npffr1 →
← Ddx21
Herc4 →
← Lrrtm3
← 1700023F02Rik
Jmjd1c →
Rtkn2 →
Rhobtb1 →
Ccdc6 →
← Bicc1
Ppa1 →
← Srgn
← Mypn
4930407I19Rik →
← Nrbf2
← Arid5b
← Cdk1
Mrln →
← Tfam
← 1700022H01Rik
← Sirt1
Gm31763 →
← Zfp365
Tmem26 →
Ank3 →
Fam13c →
← Cisd1
Sar1a →
← Ddx50
Dnajc12 →
← Reep3
← 4930545H06Rik
Slc16a9 →
← Ube2d1
39 genes omitted
62
64
66
68
70
Position on chr10 (Mb)

| | |
|---|---|
| date: | Thu Oct 25 17:07:38 2018 |
| build: | F34GBS |
| display range: | chr10:61602511−71602511 [61602511−71602511] |
| hilite range: | 0 − 0 [ 0 − 0 ] |
| reference SNP: | chr10:66602511 |
| number of SNPs plotted: | 346 |
| min P−value: | 1.81E−6 [chr10:66602511] |
| max P−value: | 5.94E−1 [chr10:70028591] |
| omitted Genes: | Tysnd1, Aifm2, H2afy2 |
| omitted Genes: | Mir5108, Col13a1, Gm5424 |
| omitted Genes: | 2010107G23Rik, Neurog3, Tspan15 |
| omitted Genes: | Mir7662, Tacr2, Hk1 |
| omitted Genes: | Hk1os, Hkdc1, Supv3l1 |
| omitted Genes: | 4930507D05Rik, Vps26a, Kif1bp |
| omitted Genes: | Stox1, Mir7215, Ccar1 |
| omitted Genes: | Snord98, Tet1, Slc25a16 |
| omitted Genes: | Dna2, Rufy2, Hnrnph3 |
| omitted Genes: | Pbld2, Pbld1, Atoh7 |
| omitted Genes: | Ctnna3, Egr2, Ado |
| omitted Genes: | 1700040L02Rik, Phyhipl, D630013N20Rik |
| omitted Genes: | C730027H18Rik, 1700113B09Rik, Ipmk |

Make more plots at http://csg.sph.umich.edu/locuszoom/

Fine−mapping Regions

| snp | chr | pos (Mb) | pp | group | color |
|---|---|---|---|---|---|
| chr10:66602509 | 10 | 66.60251 | 0.114957231 | Credible Set | purple |
| chr10:66602511 | 10 | 66.60251 | 0.114957231 | Credible Set | purple |
| chr10:66608856 | 10 | 66.60886 | 0.059113686 | Credible Set | purple |
| chr10:66608885 | 10 | 66.60889 | 0.059113686 | Credible Set | purple |
| chr10:66609052 | 10 | 66.60905 | 0.024600070 | Credible Set | purple |
| chr10:66609080 | 10 | 66.60908 | 0.024600070 | Credible Set | purple |
| chr10:66609124 | 10 | 66.60912 | 0.024600070 | Credible Set | purple |
| chr10:66635369 | 10 | 66.63537 | 0.014827777 | Credible Set | purple |
| chr10:66719524 | 10 | 66.71952 | 0.011118828 | Credible Set | purple |
| chr10:66778997 | 10 | 66.77900 | 0.010270821 | Credible Set | purple |
| chr10:66779040 | 10 | 66.77904 | 0.010677544 | Credible Set | purple |
| chr10:66811101 | 10 | 66.81110 | 0.005123930 | Credible Set | purple |
| chr10:66811116 | 10 | 66.81112 | 0.005032271 | Credible Set | purple |
| chr10:66811497 | 10 | 66.81150 | 0.017842867 | Credible Set | purple |
| chr10:66811529 | 10 | 66.81153 | 0.017842867 | Credible Set | purple |
| chr10:66811581 | 10 | 66.81158 | 0.004697080 | Credible Set | purple |
| chr10:66811611 | 10 | 66.81161 | 0.004697080 | Credible Set | purple |
| chr10:66811619 | 10 | 66.81162 | 0.004697080 | Credible Set | purple |
| chr10:66811684 | 10 | 66.81168 | 0.004697080 | Credible Set | purple |
| chr10:66859150 | 10 | 66.85915 | 0.010270821 | Credible Set | purple |
| chr10:66859163 | 10 | 66.85916 | 0.010270821 | Credible Set | purple |
| chr10:66859201 | 10 | 66.85920 | 0.012124524 | Credible Set | purple |
| chr10:66859671 | 10 | 66.85967 | 0.012124524 | Credible Set | purple |
| chr10:66940094 | 10 | 66.94009 | 0.024600070 | Credible Set | purple |
| chr10:66940121 | 10 | 66.94012 | 0.022459976 | Credible Set | purple |
| chr10:67003583 | 10 | 67.00358 | 0.015709767 | Credible Set | purple |
| chr10:67003588 | 10 | 67.00359 | 0.016706709 | Credible Set | purple |
| chr10:67026213 | 10 | 67.02621 | 0.008918899 | Credible Set | purple |
| chr10:67029219 | 10 | 67.02922 | 0.008371125 | Credible Set | purple |
| chr10:67029237 | 10 | 67.02924 | 0.008918899 | Credible Set | purple |
| chr10:67034406 | 10 | 67.03441 | 0.009545893 | Credible Set | purple |
| chr10:67036263 | 10 | 67.03626 | 0.009894723 | Credible Set | purple |
| chr10:67061736 | 10 | 67.06174 | 0.010677544 | Credible Set | purple |
| chr10:67077389 | 10 | 67.07739 | 0.008122366 | Credible Set | purple |
| chr10:67107858 | 10 | 67.10786 | 0.010270821 | Credible Set | purple |
| chr10:67107945 | 10 | 67.10795 | 0.009894723 | Credible Set | purple |

Fine−mapping Regions

| snp | chr | pos (Mb) | pp | group | color |
|---|---|---|---|---|---|
| chr10:67108562 | 10 | 67.10856 | 0.012124524 | Credible Set | purple |
| chr10:67115847 | 10 | 67.11585 | 0.010677544 | Credible Set | purple |
| chr10:67116164 | 10 | 67.11616 | 0.012701061 | Credible Set | purple |
| chr10:67121830 | 10 | 67.12183 | 0.015709767 | Credible Set | purple |
| chr10:67121898 | 10 | 67.12190 | 0.015709767 | Credible Set | purple |
| chr10:67214197 | 10 | 67.21420 | 0.008636086 | Credible Set | purple |
| chr10:67217456 | 10 | 67.21746 | 0.008122366 | Credible Set | purple |
| chr10:67217505 | 10 | 67.21751 | 0.008122366 | Credible Set | purple |
| chr10:67226747 | 10 | 67.22675 | 0.007459588 | Credible Set | purple |
| chr10:67237234 | 10 | 67.23723 | 0.009545893 | Credible Set | purple |
| chr10:67237236 | 10 | 67.23724 | 0.008918899 | Credible Set | purple |
| chr10:67237389 | 10 | 67.23739 | 0.014041816 | Credible Set | purple |
| chr10:67246731 | 10 | 67.24673 | 0.009221448 | Credible Set | purple |
| chr10:67264982 | 10 | 67.26498 | 0.009894723 | Credible Set | purple |
| chr10:67265380 | 10 | 67.26538 | 0.010270821 | Credible Set | purple |
| chr10:67281545 | 10 | 67.28154 | 0.009894723 | Credible Set | purple |
| chr10:67281940 | 10 | 67.28194 | 0.010270821 | Credible Set | purple |
| chr10:67282089 | 10 | 67.28209 | 0.006274143 | Credible Set | purple |
| chr10:67338317 | 10 | 67.33832 | 0.009894723 | Credible Set | purple |
| chr10:67345731 | 10 | 67.34573 | 0.011118828 | Credible Set | purple |
| chr10:67346184 | 10 | 67.34618 | 0.011118828 | Credible Set | purple |
| chr10:67382855 | 10 | 67.38286 | 0.006731257 | Credible Set | purple |
| chr10:67383022 | 10 | 67.38302 | 0.010677544 | Credible Set | purple |
| chr10:67391783 | 10 | 67.39178 | 0.011118828 | Credible Set | purple |

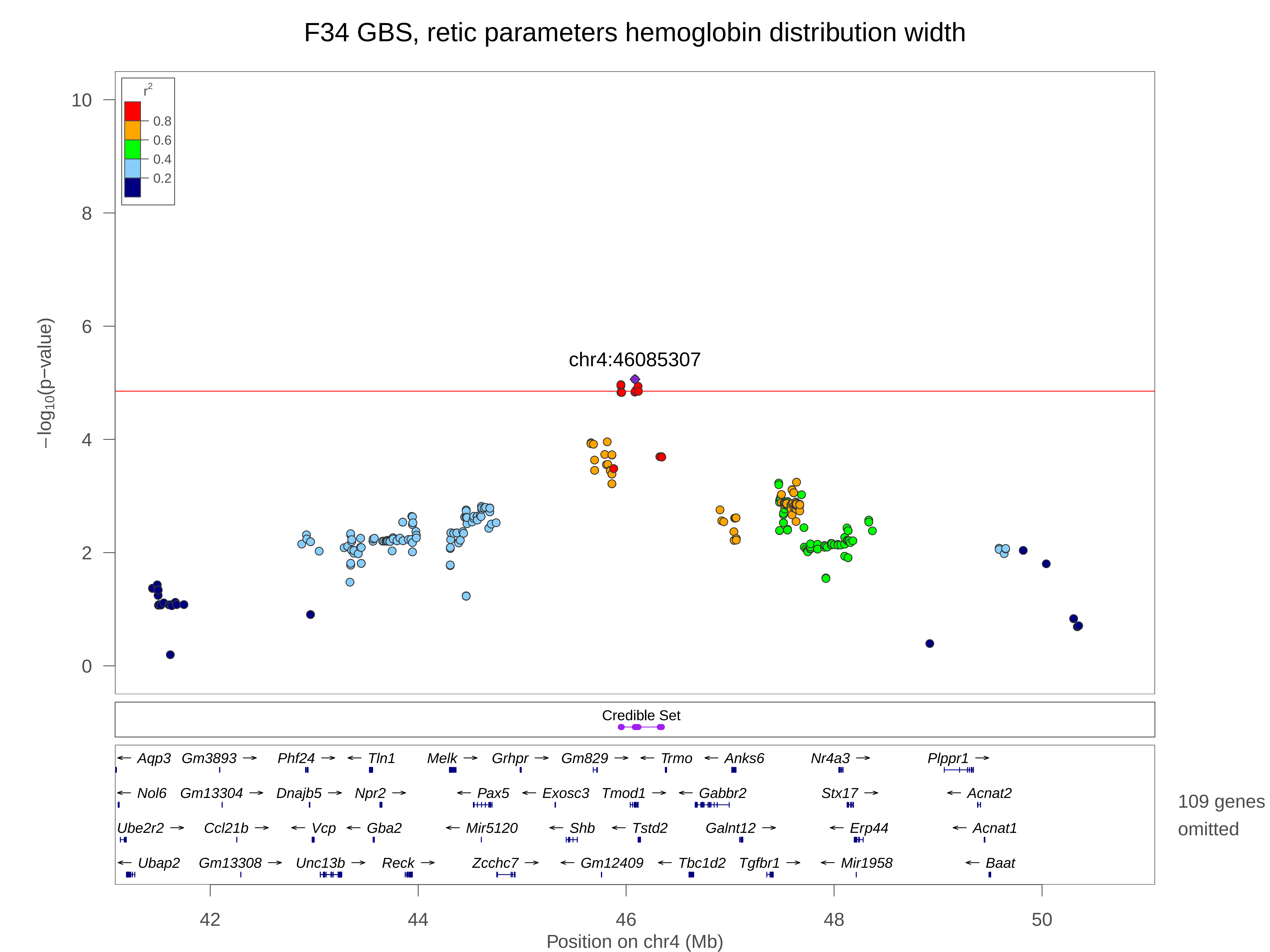
F34 GBS, retic parameters hemoglobin distribution width
r2
0.8
0.6
0.4
0.2
10
8
6
4
2
0
−log10(p−value)
chr4:46085307
Credible Set
← Aqp3 Gm3893 → Phf24 → ← Tln1 Melk → Grhpr → Gm829 → ← Trmo ← Anks6 Nr4a3 → Plppr1 →
← Nol6 Gm13304 → Dnajb5 → Npr2 → ← Pax5 ← Exosc3 Tmod1 → ← Gabbr2 Stx17 → ← Acnat2
Ube2r2 → Ccl21b → ← Vcp ← Gba2 ← Mir5120 ← Shb ← Tstd2 Galnt12 → ← Erp44 ← Acnat1
← Ubap2 Gm13308 → Unc13b → Reck → Zcchc7 → ← Gm12409 ← Tbc1d2 Tgfbr1 → ← Mir1958 ← Baat
109 genes omitted
42
44
46
48
50
Position on chr4 (Mb)

date: Thu Oct 25 17:07:57 2018
build: F34GBS
display range: chr4:41085307−51085307 [41085307−51085307]
hilite range: 0 − 0 [ 0 − 0 ]
reference SNP: chr4:46085307
number of SNPs plotted: 346
min P−value: 8.69E−6 [chr4:46085307]
max P−value: 6.35E−1 [chr4:41616530]
omitted Genes: Dcaf12, Ubap1, Kif24
omitted Genes: Nudt2, AI464131, 1110017D15Rik
omitted Genes: Fam219aos, Fam219a, Dnaic1
omitted Genes: Enho, Cntfr, Rpp25l
omitted Genes: Dctn3, Arid3c, Sigmar1
omitted Genes: Galt, Il11ra1, Ccl27a
omitted Genes: Gm21541, Gm20878, Gm13306
omitted Genes: Gm13305, Gm21586, Fam205a2
omitted Genes: Gm10591, Ccl27b, Il11ra2
omitted Genes: Ccl19, Ccl21a, Fam205a1
omitted Genes: Fam205c, 1700022I11Rik, Fancg
omitted Genes: Pigo, Stoml2, Fam214b
omitted Genes: Atp8b5, Rusc2, Fam166b
omitted Genes: Tesk1, Cd72, Sit1
omitted Genes: Rmrp, Ccdc107, Arhgef39
omitted Genes: Car9, Tpm2, Creb3
omitted Genes: Rgp1, Msmp, Spag8
omitted Genes: Hint2, Fam221b, Tmem8b
omitted Genes: Olfr70, Olfr71, Hrct1
omitted Genes: 5430416O09Rik, A630077J23Rik, Olfr159
omitted Genes: Olfr29−ps1, Olfr156, Olfr157
omitted Genes: Olfr155, Glipr2, Ccin
omitted Genes: Clta, Gne, Gm12504
omitted Genes: Rnf38, Mir5106, 5730488B01Rik
omitted Genes: Zbtb5, Polr1e, Fbxo10
omitted Genes: Tomm5, Frmpd1, Frmpd1os
omitted Genes: Trmt10b, Dcaf10, Slc25a51
omitted Genes: Aldh1b1, Igfbpl1, Stra6l
omitted Genes: Ccdc180, Tdrd7, Ncbp1
omitted Genes: Xpa, Foxe1, Hemgn
omitted Genes: Anp32b, Nans, Trim14

omitted Genes: Coro2a, Col15a1, Alg2

Fine−mapping Regions

| snp | chr | pos (Mb) | pp | group | color |
|---|---|---|---|---|---|
| chr4:45949623 | 4 | 45.94962 | 0.089762626 | Credible Set | purple |
| chr4:45949656 | 4 | 45.94966 | 0.093939720 | Credible Set | purple |
| chr4:45949665 | 4 | 45.94967 | 0.092953049 | Credible Set | purple |
| chr4:45950119 | 4 | 45.95012 | 0.069606476 | Credible Set | purple |
| chr4:45950209 | 4 | 45.95021 | 0.069509485 | Credible Set | purple |
| chr4:45951631 | 4 | 45.95163 | 0.069346826 | Credible Set | purple |
| chr4:45957405 | 4 | 45.95741 | 0.069129650 | Credible Set | purple |
| chr4:46084049 | 4 | 46.08405 | 0.070167667 | Credible Set | purple |
| chr4:46085307 | 4 | 46.08531 | 0.115350828 | Credible Set | purple |
| chr4:46091992 | 4 | 46.09199 | 0.074918273 | Credible Set | purple |
| chr4:46115063 | 4 | 46.11506 | 0.087995678 | Credible Set | purple |
| chr4:46118124 | 4 | 46.11812 | 0.072463060 | Credible Set | purple |
| chr4:46324082 | 4 | 46.32408 | 0.005837550 | Credible Set | purple |
| chr4:46340473 | 4 | 46.34047 | 0.005854421 | Credible Set | purple |
| chr4:46343171 | 4 | 46.34317 | 0.005746680 | Credible Set | purple |

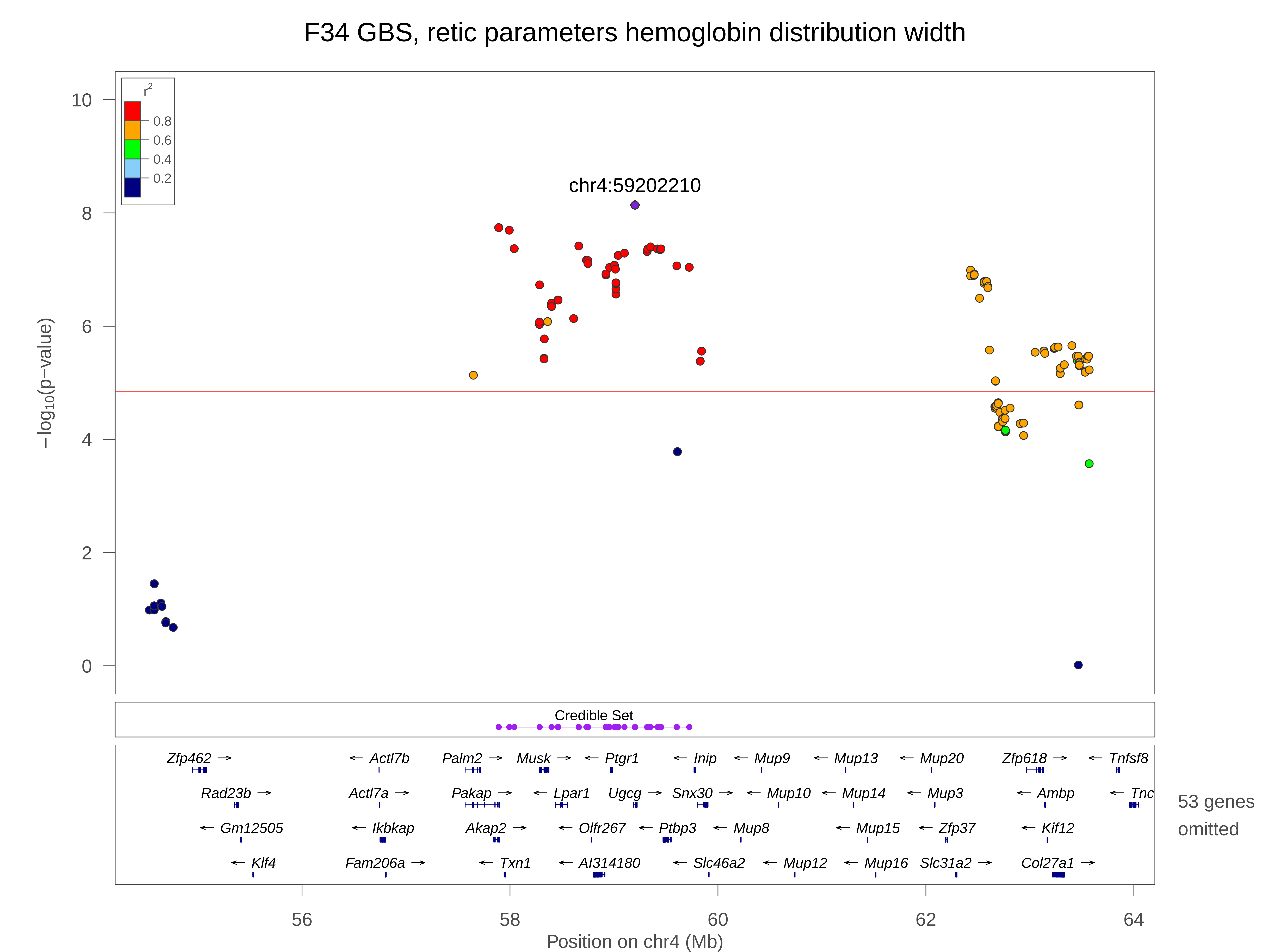
F34 GBS, retic parameters hemoglobin distribution width
r2
0.8
0.6
0.4
0.2
chr4:59202210
−log10(p−value)
10
8
6
4
2
0
Credible Set
Zfp462 →
← Actl7b
Palm2 →
Musk →
← Ptgr1
← Inip
← Mup9
← Mup13
← Mup20
Zfp618 →
← Tnfsf8
Rad23b →
Actl7a →
Pakap →
← Lpar1
Ugcg →
Snx30 →
← Mup10
← Mup14
← Mup3
← Ambp
← Tnc
← Gm12505
← Ikbkap
Akap2 →
← Olfr267
← Ptbp3
← Mup8
← Mup15
← Zfp37
← Kif12
← Klf4
Fam206a →
← Txn1
← AI314180
← Slc46a2
← Mup12
← Mup16
Slc31a2 →
Col27a1 →
53 genes omitted
56
58
60
62
64
Position on chr4 (Mb)

| | |
|---|---|
| date: | Thu Oct 25 17:08:05 2018 |
| build: | F34GBS |
| display range: | chr4:54202210−64202210 [54202210−64202210] |
| hilite range: | 0 − 0 [ 0 − 0 ] |
| reference SNP: | chr4:59202210 |
| number of SNPs plotted: | 157 |
| min P−value: | 7.28E−9 [chr4:59202210] |
| max P−value: | 9.66E−1 [chr4:63465549] |
| omitted Genes: | Ctnnal1, Tmem245, Mir32 |
| omitted Genes: | Frrs1l, Epb41l4b, Gm12530 |
| omitted Genes: | Ptpn3, 1700042G15Rik, D630039A03Rik |
| omitted Genes: | Txndc8, Svep1, Mir3095 |
| omitted Genes: | Zkscan16, Dnajc25, Gng10 |
| omitted Genes: | Gm12596, Susd1, Hsdl2 |
| omitted Genes: | E130308A19Rik, Mup4, Mup6 |
| omitted Genes: | Mup7, Mup2, Mup1 |
| omitted Genes: | Mup11, Mup17, Mup18 |
| omitted Genes: | Mup19, Mup5, Mup21 |
| omitted Genes: | Fkbp15, Slc31a1, Cdc26 |
| omitted Genes: | Prpf4, Rnf183, Wdr31 |
| omitted Genes: | Bspry, Hdhd3, Alad |
| omitted Genes: | Pole3, 4933430I17Rik, Rgs3 |
| omitted Genes: | Mir455, Orm1, Orm3 |
| omitted Genes: | Orm2, Akna, Whrn |
| omitted Genes: | Atp6v1g1, 6330416G13Rik, Gm11213 |
| omitted Genes: | 1700018C11Rik, Tnfsf15 |

Make more plots at http://csg.sph.umich.edu/locuszoom/

Fine−mapping Regions

| snp | chr | pos (Mb) | pp | group | color |
|---|---|---|---|---|---|
| chr4:57891427 | 4 | 57.89143 | 0.074789361 | Credible Set | purple |
| chr4:57891451 | 4 | 57.89145 | 0.074789361 | Credible Set | purple |
| chr4:57993173 | 4 | 57.99317 | 0.067416927 | Credible Set | purple |
| chr4:58041091 | 4 | 58.04109 | 0.032637673 | Credible Set | purple |
| chr4:58285477 | 4 | 58.28548 | 0.007854700 | Credible Set | purple |
| chr4:58399947 | 4 | 58.39995 | 0.003810993 | Credible Set | purple |
| chr4:58400011 | 4 | 58.40001 | 0.003350692 | Credible Set | purple |
| chr4:58461854 | 4 | 58.46185 | 0.004337250 | Credible Set | purple |
| chr4:58462401 | 4 | 58.46240 | 0.004311110 | Credible Set | purple |
| chr4:58662154 | 4 | 58.66215 | 0.036220962 | Credible Set | purple |
| chr4:58735042 | 4 | 58.73504 | 0.020669850 | Credible Set | purple |
| chr4:58748774 | 4 | 58.74877 | 0.020454574 | Credible Set | purple |
| chr4:58749245 | 4 | 58.74925 | 0.018065857 | Credible Set | purple |
| chr4:58923167 | 4 | 58.92317 | 0.011568694 | Credible Set | purple |
| chr4:58923190 | 4 | 58.92319 | 0.012046201 | Credible Set | purple |
| chr4:58958681 | 4 | 58.95868 | 0.015628801 | Credible Set | purple |
| chr4:59004314 | 4 | 59.00431 | 0.016952064 | Credible Set | purple |
| chr4:59013917 | 4 | 59.01392 | 0.014537933 | Credible Set | purple |
| chr4:59019003 | 4 | 59.01900 | 0.005457270 | Credible Set | purple |
| chr4:59019032 | 4 | 59.01903 | 0.006668589 | Credible Set | purple |
| chr4:59019072 | 4 | 59.01907 | 0.008243187 | Credible Set | purple |
| chr4:59019113 | 4 | 59.01911 | 0.008487489 | Credible Set | purple |
| chr4:59040591 | 4 | 59.04059 | 0.024982138 | Credible Set | purple |
| chr4:59100640 | 4 | 59.10064 | 0.027250456 | Credible Set | purple |
| chr4:59202210 | 4 | 59.20221 | 0.181971130 | Credible Set | purple |
| chr4:59319020 | 4 | 59.31902 | 0.029047920 | Credible Set | purple |
| chr4:59324851 | 4 | 59.32485 | 0.032108606 | Credible Set | purple |
| chr4:59352342 | 4 | 59.35234 | 0.035044679 | Credible Set | purple |
| chr4:59417447 | 4 | 59.41745 | 0.032336464 | Credible Set | purple |
| chr4:59417451 | 4 | 59.41745 | 0.032254525 | Credible Set | purple |
| chr4:59417505 | 4 | 59.41750 | 0.032172049 | Credible Set | purple |
| chr4:59444538 | 4 | 59.44454 | 0.031259187 | Credible Set | purple |
| chr4:59451945 | 4 | 59.45195 | 0.032339242 | Credible Set | purple |
| chr4:59604913 | 4 | 59.60491 | 0.016542001 | Credible Set | purple |
| chr4:59724135 | 4 | 59.72413 | 0.015635485 | Credible Set | purple |

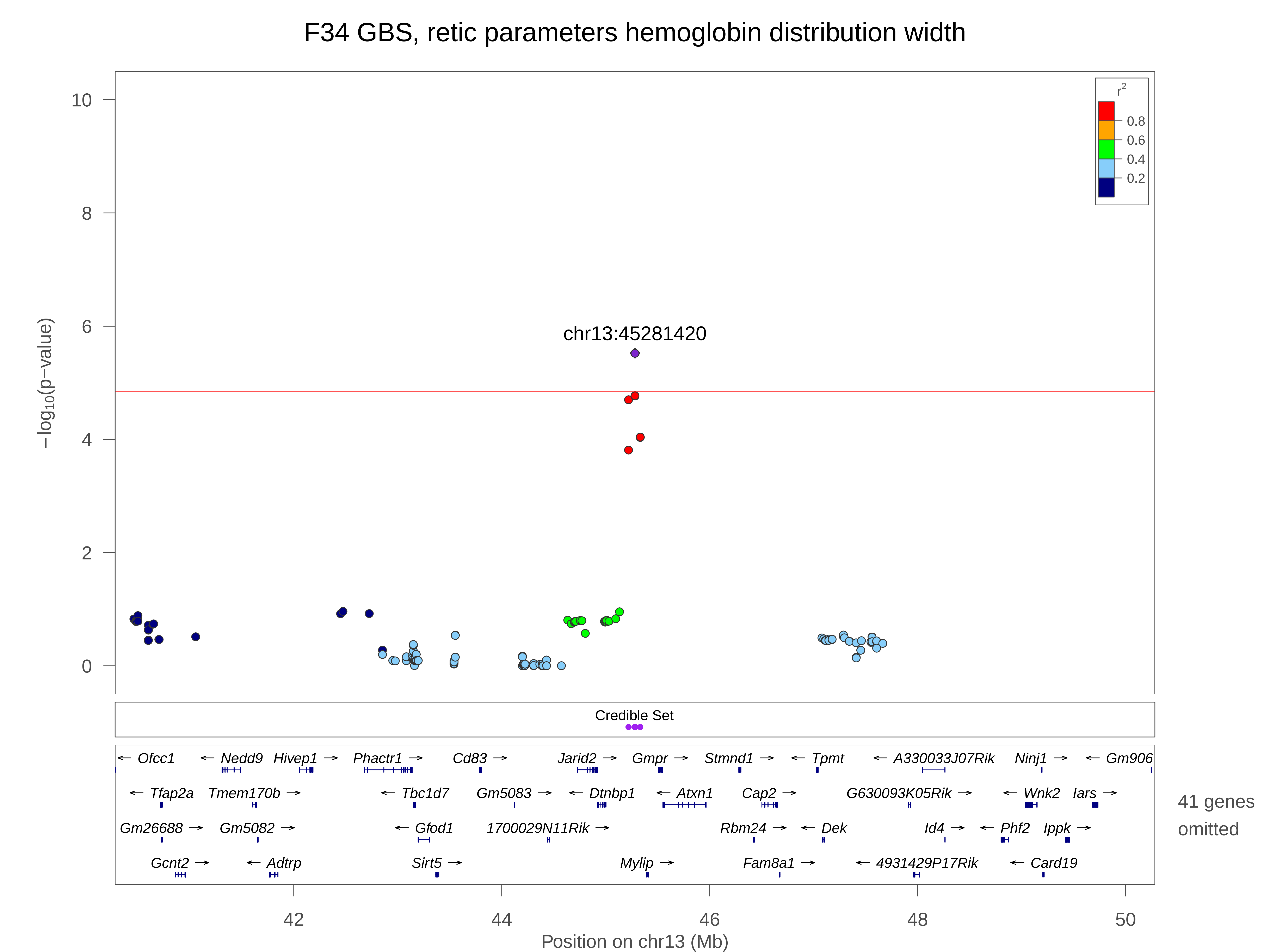
F34 GBS, retic parameters hemoglobin distribution width
chr13:45281420
$-\log_{10}$(p−value)
Position on chr13 (Mb)
r²
0.8
0.6
0.4
0.2
Credible Set
Ofcc1
Nedd9
Hivep1
Phactr1
Cd83
Jarid2
Gmpr
Stmnd1
Tpmt
A330033J07Rik
Ninj1
Gm906
Tfap2a
Tmem170b
Tbc1d7
Gm5083
Dtnbp1
Atxn1
Cap2
G630093K05Rik
Wnk2
Iars
Gm26688
Gm5082
Gfod1
1700029N11Rik
Rbm24
Dek
Id4
Phf2
Ippk
Gcnt2
Adtrp
Sirt5
Mylip
Fam8a1
4931429P17Rik
Card19
41 genes omitted

| | |
|---|---|
| date: | Thu Oct 25 17:07:47 2018 |
| build: | F34GBS |
| display range: | chr13:40281420−50281420 [40281420−50281420] |
| hilite range: | 0 − 0 [ 0 − 0 ] |
| reference SNP: | chr13:45281420 |
| number of SNPs plotted: | 173 |
| min P−value: | 3.01E−6 [chr13:45281420] |
| max P−value: | 10E−1 [chr13:44396308] |
| omitted Genes: | Mir5124a, Pak1ip1, Tmem14c |
| omitted Genes: | Mak, Gcm2, Sycp2l |
| omitted Genes: | Elovl2, Smim13, Gm10790 |
| omitted Genes: | Edn1, Nol7, Ranbp9 |
| omitted Genes: | Mcur1, Rnf182, Gm20751 |
| omitted Genes: | A330076C08Rik, Nup153, Kif13a |
| omitted Genes: | Nhlrc1, Kdm1b, Rnf144b |
| omitted Genes: | A930002C04Rik, A330048O09Rik, Zfp169 |
| omitted Genes: | Mirlet7d, Mirlet7f−1, Mirlet7a−1 |
| omitted Genes: | Ptpdc1, Barx1, Fam120a |
| omitted Genes: | Fam120aos, Susd3, Fgd3 |
| omitted Genes: | Bicd2, Cenpp, Ecm2 |
| omitted Genes: | Aspn, Omd, Ogn |
| omitted Genes: | Nol8, 2310081J21Rik |

Make more plots at http://csg.sph.umich.edu/locuszoom/

Fine−mapping Regions

| snp | chr | pos (Mb) | pp | group | color |
|---|---|---|---|---|---|
| chr13:45219307 | 13 | 45.21931 | 0.01259701 | Credible Set | purple |
| chr13:45219356 | 13 | 45.21936 | 0.08755971 | Credible Set | purple |
| chr13:45281289 | 13 | 45.28129 | 0.10158012 | Credible Set | purple |
| chr13:45281291 | 13 | 45.28129 | 0.10114708 | Credible Set | purple |
| chr13:45281309 | 13 | 45.28131 | 0.10177082 | Credible Set | purple |
| chr13:45281420 | 13 | 45.28142 | 0.53331775 | Credible Set | purple |
| chr13:45331781 | 13 | 45.33178 | 0.02081501 | Credible Set | purple |
| chr13:45331853 | 13 | 45.33185 | 0.02045599 | Credible Set | purple |
| chr13:45331868 | 13 | 45.33187 | 0.02075652 | Credible Set | purple |

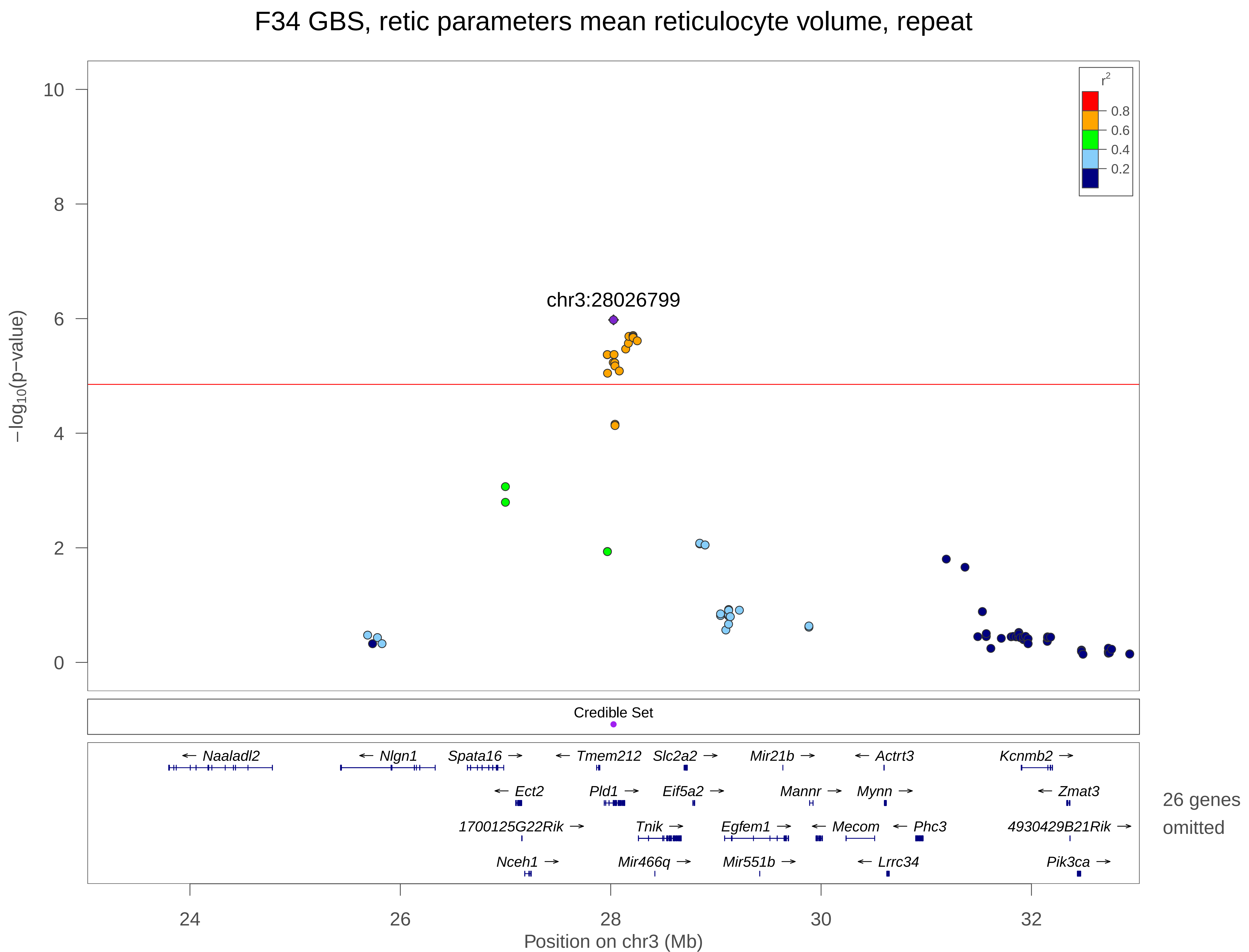
F34 GBS, retic parameters mean reticulocyte volume, repeat
r2
0.8
0.6
0.4
0.2
chr3:28026799
−log10(p−value)
10
8
6
4
2
0
Credible Set
← Naaladl2
← Nlgn1
Spata16 →
← Tmem212
Slc2a2 →
Mir21b →
← Actrt3
Kcnmb2 →
← Ect2
Pld1 →
Eif5a2 →
Mannr →
Mynn →
← Zmat3
1700125G22Rik →
Tnik →
Egfem1 →
← Mecom
← Phc3
4930429B21Rik →
Nceh1 →
Mir466q →
Mir551b →
← Lrrc34
Pik3ca →
26 genes omitted
24
26
28
30
32
Position on chr3 (Mb)

| | |
|---|---|
| date: | Thu Oct 25 17:08:37 2018 |
| build: | F34GBS |
| display range: | chr3:23026799−33026799 [23026799−33026799] |
| hilite range: | 0 − 0 [ 0 − 0 ] |
| reference SNP: | chr3:28026799 |
| number of SNPs plotted: | 116 |
| min P−value: | 1.05E−6 [chr3:28026799] |
| max P−value: | 7.21E−1 [chr3:32490030] |
| omitted Genes: | Tnfsf10, Ghsr, Fndc3b |
| omitted Genes: | Mir3092, Rpl22l1, Gm1527 |
| omitted Genes: | Lrriq4, Lrrc31, Samd7 |
| omitted Genes: | Sec62, Gpr160, Prkci |
| omitted Genes: | Mir7008, Skil, Cldn11 |
| omitted Genes: | Slc7a14, Kcnmb3, Zfp639 |
| omitted Genes: | Mfn1, Gnb4, Actl6a |
| omitted Genes: | Mrpl47, Ndufb5, Usp13 |
| omitted Genes: | Pex5l, 4930419G24Rik |

Make more plots at http://csg.sph.umich.edu/locuszoom/

Fine−mapping Regions

| snp | chr | pos (Mb) | pp | group | color |
| --- | --- | --- | --- | --- | --- |
| chr3:28026799 | 3 | 28.0268 | 1 | Credible Set | purple |

### F34 GBS, retic parameters mean reticulocyte volume, repeat

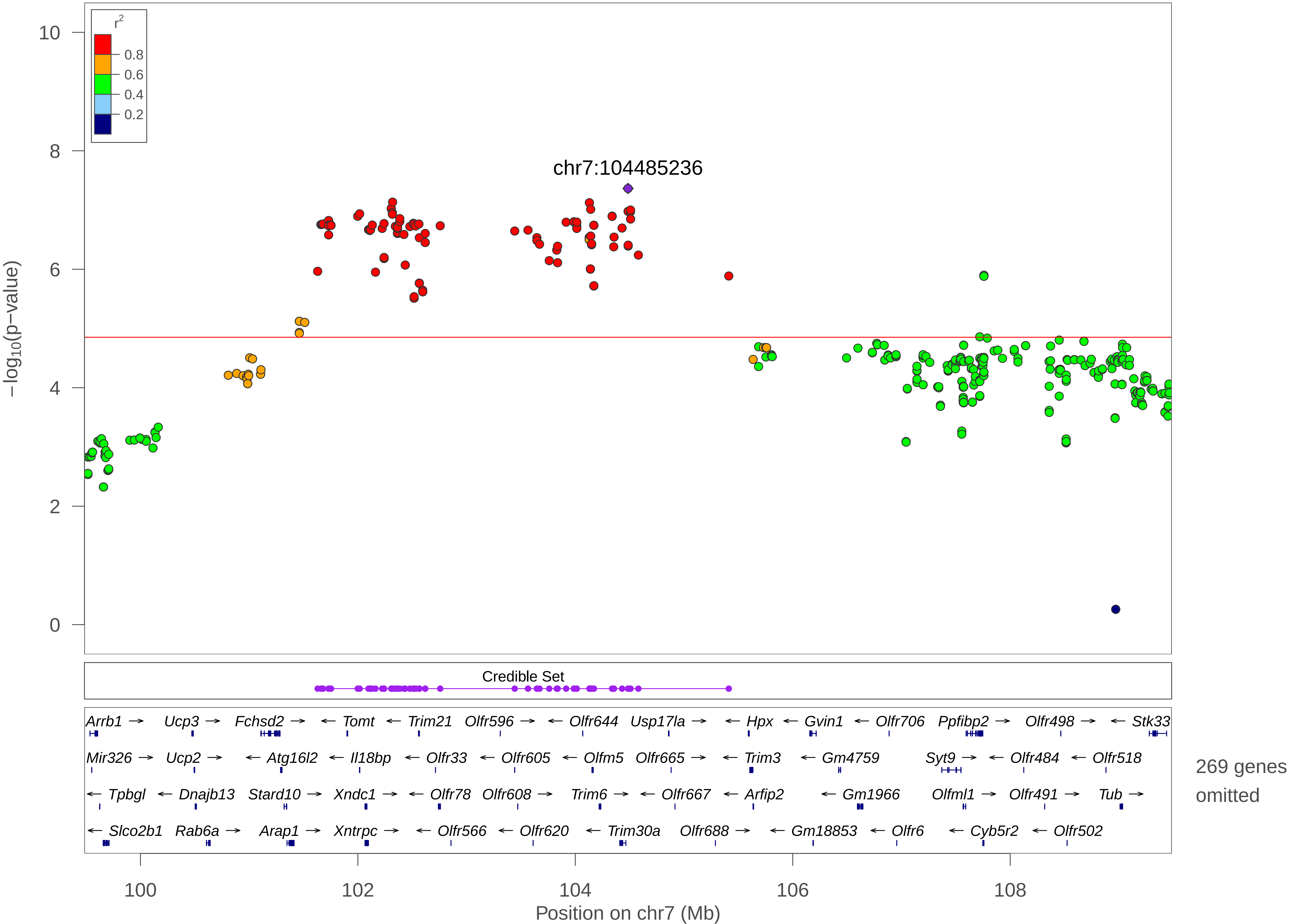

date: Thu Oct 25 17:08:51 2018
build: F34GBS
display range: chr7:99485236−109485236 [99485236−109485236]
hilite range: 0 − 0 [ 0 − 0 ]
reference SNP: chr7:104485236
number of SNPs plotted: 448
min P−value: 4.32E−8 [chr7:104485236]
max P−value: 5.53E−1 [chr7:108970722]
omitted Genes: Olfr520, Olfr521, Neu3
omitted Genes: Spcs2, Xrra1, Rnf169
omitted Genes: Chrdl2, Pold3, Lipt2
omitted Genes: Kcne3, Pgm2l1, Gpx2−ps1
omitted Genes: P4ha3, Ppme1, C2cd3
omitted Genes: Coa4, Mrpl48, Plekhb1
omitted Genes: Fam168a, Relt, Arhgef17
omitted Genes: Mir3102, P2ry6, P2ry2
omitted Genes: Pde2a, Mir139, Art2a−ps
omitted Genes: Art2b, Clpb, Phox2a
omitted Genes: Inppl1, Folr2, Folr1
omitted Genes: Anapc15, Lamtor1, Lrrc51
omitted Genes: Numa1, Rnf121, Trpc2
omitted Genes: Art5, Art1, Chrna10
omitted Genes: Nup98, Pgap2, Rhog
omitted Genes: Stim1, Rrm1, Olfr543
omitted Genes: Olfr544, Olfr545, Olfr547
omitted Genes: Olfr549, Olfr550, Olfr551
omitted Genes: Olfr552, Olfr553, Olfr554
omitted Genes: Olfr555, Olfr556, Trim68
omitted Genes: Olfr557, Olfr558, Olfr559
omitted Genes: Olfr560, Olfr561, Olfr564
omitted Genes: Olfr568, Olfr569, Olfr570
omitted Genes: Olfr571, Olfr572, Olfr574
omitted Genes: Olfr575, Olfr576, Olfr577
omitted Genes: Olfr578, Olfr582, Olfr583
omitted Genes: Olfr584, Olfr585, Olfr586
omitted Genes: Olfr589, Olfr591, Olfr592
omitted Genes: Olfr593, Olfr594, Usp17ld
omitted Genes: Olfr597, Olfr598, Olfr599
omitted Genes: Olfr600, Olfr601, Olfr603
Make more plots at http://csg.sph.umich.edu/locuszoom/
omitted Genes: Usp17le, Olfr606, Olfr609

Fine−mapping Regions

| snp | chr | pos (Mb) | pp | group | color |
|---|---|---|---|---|---|
| chr7:101630176 | 7 | 101.6302 | 0.001789679 | Credible Set | purple |
| chr7:101660383 | 7 | 101.6604 | 0.010341918 | Credible Set | purple |
| chr7:101669186 | 7 | 101.6692 | 0.010354086 | Credible Set | purple |
| chr7:101673483 | 7 | 101.6735 | 0.010512090 | Credible Set | purple |
| chr7:101680140 | 7 | 101.6801 | 0.010567581 | Credible Set | purple |
| chr7:101731061 | 7 | 101.7311 | 0.012046240 | Credible Set | purple |
| chr7:101731110 | 7 | 101.7311 | 0.010042784 | Credible Set | purple |
| chr7:101731130 | 7 | 101.7311 | 0.007006876 | Credible Set | purple |
| chr7:101731658 | 7 | 101.7317 | 0.009875469 | Credible Set | purple |
| chr7:101751790 | 7 | 101.7518 | 0.010045918 | Credible Set | purple |
| chr7:101752580 | 7 | 101.7526 | 0.010142363 | Credible Set | purple |
| chr7:101752971 | 7 | 101.7530 | 0.010001728 | Credible Set | purple |
| chr7:101998679 | 7 | 101.9987 | 0.014114434 | Credible Set | purple |
| chr7:102016307 | 7 | 102.0163 | 0.015472808 | Credible Set | purple |
| chr7:102098054 | 7 | 102.0981 | 0.008537204 | Credible Set | purple |
| chr7:102110109 | 7 | 102.1101 | 0.008275023 | Credible Set | purple |
| chr7:102110501 | 7 | 102.1105 | 0.008757857 | Credible Set | purple |
| chr7:102116318 | 7 | 102.1163 | 0.008549278 | Credible Set | purple |
| chr7:102116430 | 7 | 102.1164 | 0.008549278 | Credible Set | purple |
| chr7:102116478 | 7 | 102.1165 | 0.008485512 | Credible Set | purple |
| chr7:102132562 | 7 | 102.1326 | 0.010142489 | Credible Set | purple |
| chr7:102162058 | 7 | 102.1621 | 0.001736170 | Credible Set | purple |
| chr7:102223396 | 7 | 102.2234 | 0.008946636 | Credible Set | purple |
| chr7:102240695 | 7 | 102.2407 | 0.010738925 | Credible Set | purple |
| chr7:102241191 | 7 | 102.2412 | 0.002888969 | Credible Set | purple |
| chr7:102241211 | 7 | 102.2412 | 0.002928719 | Credible Set | purple |
| chr7:102241212 | 7 | 102.2412 | 0.003006739 | Credible Set | purple |
| chr7:102306968 | 7 | 102.3070 | 0.018668175 | Credible Set | purple |
| chr7:102306978 | 7 | 102.3070 | 0.019080554 | Credible Set | purple |
| chr7:102317213 | 7 | 102.3172 | 0.016102328 | Credible Set | purple |
| chr7:102317246 | 7 | 102.3172 | 0.015242451 | Credible Set | purple |
| chr7:102319586 | 7 | 102.3196 | 0.024041039 | Credible Set | purple |
| chr7:102344970 | 7 | 102.3450 | 0.009674340 | Credible Set | purple |
| chr7:102363897 | 7 | 102.3639 | 0.009626705 | Credible Set | purple |
| chr7:102364204 | 7 | 102.3642 | 0.007395796 | Credible Set | purple |
| chr7:102364223 | 7 | 102.3642 | 0.007656479 | Credible Set | purple |

Fine−mapping Regions

| snp | chr | pos (Mb) | pp | group | color |
|---|---|---|---|---|---|
| chr7:102364315 | 7 | 102.3643 | 0.008959549 | Credible Set | purple |
| chr7:102364326 | 7 | 102.3643 | 0.008959549 | Credible Set | purple |
| chr7:102364411 | 7 | 102.3644 | 0.009244659 | Credible Set | purple |
| chr7:102385520 | 7 | 102.3855 | 0.011613288 | Credible Set | purple |
| chr7:102385536 | 7 | 102.3855 | 0.012917652 | Credible Set | purple |
| chr7:102423165 | 7 | 102.4232 | 0.007131428 | Credible Set | purple |
| chr7:102435692 | 7 | 102.4357 | 0.002263594 | Credible Set | purple |
| chr7:102435720 | 7 | 102.4357 | 0.002263594 | Credible Set | purple |
| chr7:102478469 | 7 | 102.4785 | 0.009598491 | Credible Set | purple |
| chr7:102510475 | 7 | 102.5105 | 0.010682947 | Credible Set | purple |
| chr7:102510607 | 7 | 102.5106 | 0.010835669 | Credible Set | purple |
| chr7:102516878 | 7 | 102.5169 | 0.010503596 | Credible Set | purple |
| chr7:102531652 | 7 | 102.5317 | 0.009797808 | Credible Set | purple |
| chr7:102561708 | 7 | 102.5617 | 0.010550785 | Credible Set | purple |
| chr7:102565262 | 7 | 102.5653 | 0.001148009 | Credible Set | purple |
| chr7:102565292 | 7 | 102.5653 | 0.001154438 | Credible Set | purple |
| chr7:102565307 | 7 | 102.5653 | 0.001140220 | Credible Set | purple |
| chr7:102565323 | 7 | 102.5653 | 0.001146602 | Credible Set | purple |
| chr7:102565329 | 7 | 102.5653 | 0.001146602 | Credible Set | purple |
| chr7:102565410 | 7 | 102.5654 | 0.006305334 | Credible Set | purple |
| chr7:102619418 | 7 | 102.6194 | 0.005272459 | Credible Set | purple |
| chr7:102619431 | 7 | 102.6194 | 0.005270062 | Credible Set | purple |
| chr7:102619466 | 7 | 102.6195 | 0.007407272 | Credible Set | purple |
| chr7:102757499 | 7 | 102.7575 | 0.009879928 | Credible Set | purple |
| chr7:103442570 | 7 | 103.4426 | 0.008109336 | Credible Set | purple |
| chr7:103564836 | 7 | 103.5648 | 0.008394521 | Credible Set | purple |
| chr7:103646439 | 7 | 103.6464 | 0.006321513 | Credible Set | purple |
| chr7:103646709 | 7 | 103.6467 | 0.005661752 | Credible Set | purple |
| chr7:103671033 | 7 | 103.6710 | 0.004942555 | Credible Set | purple |
| chr7:103760301 | 7 | 103.7603 | 0.002669575 | Credible Set | purple |
| chr7:103829200 | 7 | 103.8292 | 0.003963801 | Credible Set | purple |
| chr7:103837099 | 7 | 103.8371 | 0.002471471 | Credible Set | purple |
| chr7:103837213 | 7 | 103.8372 | 0.004586112 | Credible Set | purple |
| chr7:103915503 | 7 | 103.9155 | 0.011309668 | Credible Set | purple |
| chr7:103986792 | 7 | 103.9868 | 0.011456842 | Credible Set | purple |
| chr7:103986795 | 7 | 103.9868 | 0.011456842 | Credible Set | purple |

Fine−mapping Regions

| snp | chr | pos (Mb) | pp | group | color |
|---|---|---|---|---|---|
| chr7:103986857 | 7 | 103.9869 | 0.011456499 | Credible Set | purple |
| chr7:103986860 | 7 | 103.9869 | 0.011456499 | Credible Set | purple |
| chr7:104012808 | 7 | 104.0128 | 0.010878150 | Credible Set | purple |
| chr7:104013370 | 7 | 104.0134 | 0.009686578 | Credible Set | purple |
| chr7:104013395 | 7 | 104.0134 | 0.008931380 | Credible Set | purple |
| chr7:104013467 | 7 | 104.0135 | 0.011329190 | Credible Set | purple |
| chr7:104129734 | 7 | 104.1297 | 0.023467857 | Credible Set | purple |
| chr7:104129736 | 7 | 104.1297 | 0.023467857 | Credible Set | purple |
| chr7:104138934 | 7 | 104.1389 | 0.001940021 | Credible Set | purple |
| chr7:104138936 | 7 | 104.1389 | 0.001961076 | Credible Set | purple |
| chr7:104138968 | 7 | 104.1390 | 0.001961076 | Credible Set | purple |
| chr7:104141343 | 7 | 104.1413 | 0.018365898 | Credible Set | purple |
| chr7:104141705 | 7 | 104.1417 | 0.006700518 | Credible Set | purple |
| chr7:104141768 | 7 | 104.1418 | 0.006704119 | Credible Set | purple |
| chr7:104149893 | 7 | 104.1499 | 0.004845480 | Credible Set | purple |
| chr7:104149911 | 7 | 104.1499 | 0.005032014 | Credible Set | purple |
| chr7:104170588 | 7 | 104.1706 | 0.010091899 | Credible Set | purple |
| chr7:104170594 | 7 | 104.1706 | 0.010042296 | Credible Set | purple |
| chr7:104170618 | 7 | 104.1706 | 0.010049479 | Credible Set | purple |
| chr7:104170662 | 7 | 104.1707 | 0.010049479 | Credible Set | purple |
| chr7:104338784 | 7 | 104.3388 | 0.014201135 | Credible Set | purple |
| chr7:104338810 | 7 | 104.3388 | 0.014048140 | Credible Set | purple |
| chr7:104353491 | 7 | 104.3535 | 0.004481911 | Credible Set | purple |
| chr7:104356230 | 7 | 104.3562 | 0.006449043 | Credible Set | purple |
| chr7:104430205 | 7 | 104.4302 | 0.009070232 | Credible Set | purple |
| chr7:104485236 | 7 | 104.4852 | 0.040217316 | Credible Set | purple |
| chr7:104485721 | 7 | 104.4857 | 0.016996983 | Credible Set | purple |
| chr7:104485727 | 7 | 104.4857 | 0.016932289 | Credible Set | purple |
| chr7:104486022 | 7 | 104.4860 | 0.004611366 | Credible Set | purple |
| chr7:104486041 | 7 | 104.4860 | 0.004771599 | Credible Set | purple |
| chr7:104507724 | 7 | 104.5077 | 0.016806775 | Credible Set | purple |
| chr7:104508127 | 7 | 104.5081 | 0.017848907 | Credible Set | purple |
| chr7:104508173 | 7 | 104.5082 | 0.012718918 | Credible Set | purple |
| chr7:104580091 | 7 | 104.5801 | 0.003289964 | Credible Set | purple |
| chr7:105411752 | 7 | 105.4118 | 0.001502488 | Credible Set | purple |

F34 GBS, retic parameters mean reticulocyte volume, repeat

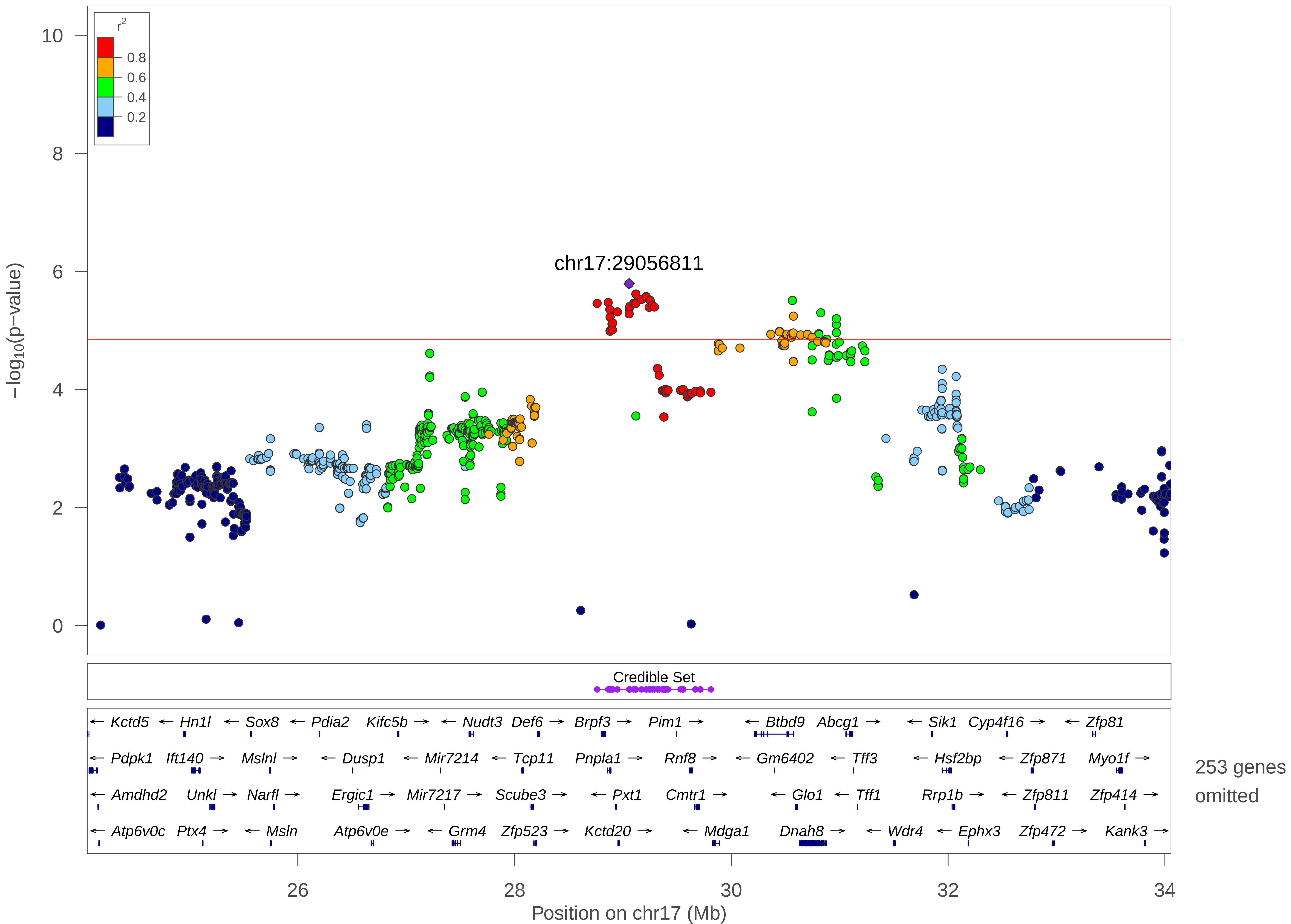

date: Thu Oct 25 17:08:18 2018
build: F34GBS
display range: chr17:24056811−34056811 [24056811−34056811]
hilite range: 0 − 0 [ 0 − 0 ]
reference SNP: chr17:29056811
number of SNPs plotted: 1034
min P−value: 1.62E−6 [chr17:29056811]
max P−value: 9.79E−1 [chr17:24180740]
omitted Genes: Tbc1d24, Ntn3, 1600002H07Rik
omitted Genes: Ccnf, Mir5134, Mir6965
omitted Genes: Abca17, Abca3, D330041H03Rik
omitted Genes: Rnps1, Eci1, Dnase1l2
omitted Genes: E4f1, Pgp, Mlst8
omitted Genes: Bricd5, Caskin1, Traf7
omitted Genes: Rab26os, Rab26, Pkd1
omitted Genes: Tsc2, Nthl1, Slc9a3r2
omitted Genes: Npw, Zfp598, Syngr3
omitted Genes: Gfer, Noxo1, Tbl3
omitted Genes: Rnf151, Snhg9, Snora78
omitted Genes: Rps2, Snora64, Ndufb10
omitted Genes: Rpl3l, Msrb1, Hs3st6
omitted Genes: Meiob, Fahd1, Hagh
omitted Genes: Igfals, Nubp2, Spsb3
omitted Genes: Eme2, Mrps34, Nme3
omitted Genes: Mapk8ip3, Cramp1l, Tmem204
omitted Genes: Telo2, Clcn7, Ccdc154
omitted Genes: BC003965, Gnptg, Tsr3
omitted Genes: Baiap3, Mir3547, Ube2i
omitted Genes: Gm17801, Prss34, Prss28
omitted Genes: Prss29, Tpsab1, Tpsb2
omitted Genes: Tpsg1, Cacna1h, Tekt4
omitted Genes: Sstr5, 2810468N07Rik, Lmf1
omitted Genes: Gng13, Chtf18, Rpusd1
omitted Genes: Haghl, Mir6966, Ccdc78
omitted Genes: Fam173a, Metrn, Fbxl16
omitted Genes: Wdr24, Jmjd8, Stub1
omitted Genes: Mir3082, Rhbdl1, Rhot2
omitted Genes: Wdr90, Fam195a, 0610011F06Rik
omitted Genes: Wfikkn1, Rab40c, Gm10012
Make more plots at http://csg.sph.umich.edu/locuszoom/
omitted Genes: Pigq, Nhlrc4, Capn15

Fine−mapping Regions

| snp | chr | pos (Mb) | pp | group | color |
|---|---|---|---|---|---|
| chr17:28761178 | 17 | 28.76118 | 0.039188938 | Credible Set | purple |
| chr17:28863817 | 17 | 28.86382 | 0.040517206 | Credible Set | purple |
| chr17:28878343 | 17 | 28.87834 | 0.031303527 | Credible Set | purple |
| chr17:28881088 | 17 | 28.88109 | 0.023421851 | Credible Set | purple |
| chr17:28881365 | 17 | 28.88136 | 0.014063971 | Credible Set | purple |
| chr17:28881407 | 17 | 28.88141 | 0.014064568 | Credible Set | purple |
| chr17:28881418 | 17 | 28.88142 | 0.013837047 | Credible Set | purple |
| chr17:28898789 | 17 | 28.89879 | 0.017992089 | Credible Set | purple |
| chr17:28902111 | 17 | 28.90211 | 0.014593218 | Credible Set | purple |
| chr17:28902126 | 17 | 28.90213 | 0.014663526 | Credible Set | purple |
| chr17:28905249 | 17 | 28.90525 | 0.018805268 | Credible Set | purple |
| chr17:28948304 | 17 | 28.94830 | 0.028528967 | Credible Set | purple |
| chr17:29056811 | 17 | 29.05681 | 0.081720674 | Credible Set | purple |
| chr17:29056940 | 17 | 29.05694 | 0.032169094 | Credible Set | purple |
| chr17:29057099 | 17 | 29.05710 | 0.026402913 | Credible Set | purple |
| chr17:29063670 | 17 | 29.06367 | 0.034972664 | Credible Set | purple |
| chr17:29098722 | 17 | 29.09872 | 0.039188938 | Credible Set | purple |
| chr17:29115818 | 17 | 29.11582 | 0.039188938 | Credible Set | purple |
| chr17:29118933 | 17 | 29.11893 | 0.055623342 | Credible Set | purple |
| chr17:29119041 | 17 | 29.11904 | 0.040175332 | Credible Set | purple |
| chr17:29119131 | 17 | 29.11913 | 0.039139174 | Credible Set | purple |
| chr17:29170505 | 17 | 29.17050 | 0.045422919 | Credible Set | purple |
| chr17:29213919 | 17 | 29.21392 | 0.050655738 | Credible Set | purple |
| chr17:29213920 | 17 | 29.21392 | 0.049427962 | Credible Set | purple |
| chr17:29241654 | 17 | 29.24165 | 0.033809733 | Credible Set | purple |
| chr17:29250546 | 17 | 29.25055 | 0.043869269 | Credible Set | purple |
| chr17:29271835 | 17 | 29.27183 | 0.036699318 | Credible Set | purple |
| chr17:29290714 | 17 | 29.29071 | 0.034151497 | Credible Set | purple |
| chr17:29319189 | 17 | 29.31919 | 0.003470127 | Credible Set | purple |
| chr17:29334046 | 17 | 29.33405 | 0.002703281 | Credible Set | purple |
| chr17:29362178 | 17 | 29.36218 | 0.001526830 | Credible Set | purple |
| chr17:29377491 | 17 | 29.37749 | 0.001550025 | Credible Set | purple |
| chr17:29378435 | 17 | 29.37843 | 0.001550025 | Credible Set | purple |
| chr17:29378474 | 17 | 29.37847 | 0.001557110 | Credible Set | purple |
| chr17:29378507 | 17 | 29.37851 | 0.001560914 | Credible Set | purple |
| chr17:29378548 | 17 | 29.37855 | 0.001563726 | Credible Set | purple |

Fine−mapping Regions

| snp | chr | pos (Mb) | pp | group | color |
|---|---|---|---|---|---|
| chr17:29393566 | 17 | 29.39357 | 0.001618165 | Credible Set | purple |
| chr17:29395215 | 17 | 29.39522 | 0.001405790 | Credible Set | purple |
| chr17:29395231 | 17 | 29.39523 | 0.001457827 | Credible Set | purple |
| chr17:29398643 | 17 | 29.39864 | 0.001529801 | Credible Set | purple |
| chr17:29415986 | 17 | 29.41599 | 0.001550025 | Credible Set | purple |
| chr17:29531394 | 17 | 29.53139 | 0.001550025 | Credible Set | purple |
| chr17:29553602 | 17 | 29.55360 | 0.001603443 | Credible Set | purple |
| chr17:29554963 | 17 | 29.55496 | 0.001515483 | Credible Set | purple |
| chr17:29555542 | 17 | 29.55554 | 0.001524576 | Credible Set | purple |
| chr17:29555787 | 17 | 29.55579 | 0.001593175 | Credible Set | purple |
| chr17:29667798 | 17 | 29.66780 | 0.001496360 | Credible Set | purple |
| chr17:29712556 | 17 | 29.71256 | 0.001518835 | Credible Set | purple |
| chr17:29714755 | 17 | 29.71476 | 0.001408293 | Credible Set | purple |
| chr17:29811310 | 17 | 29.81131 | 0.001443688 | Credible Set | purple |

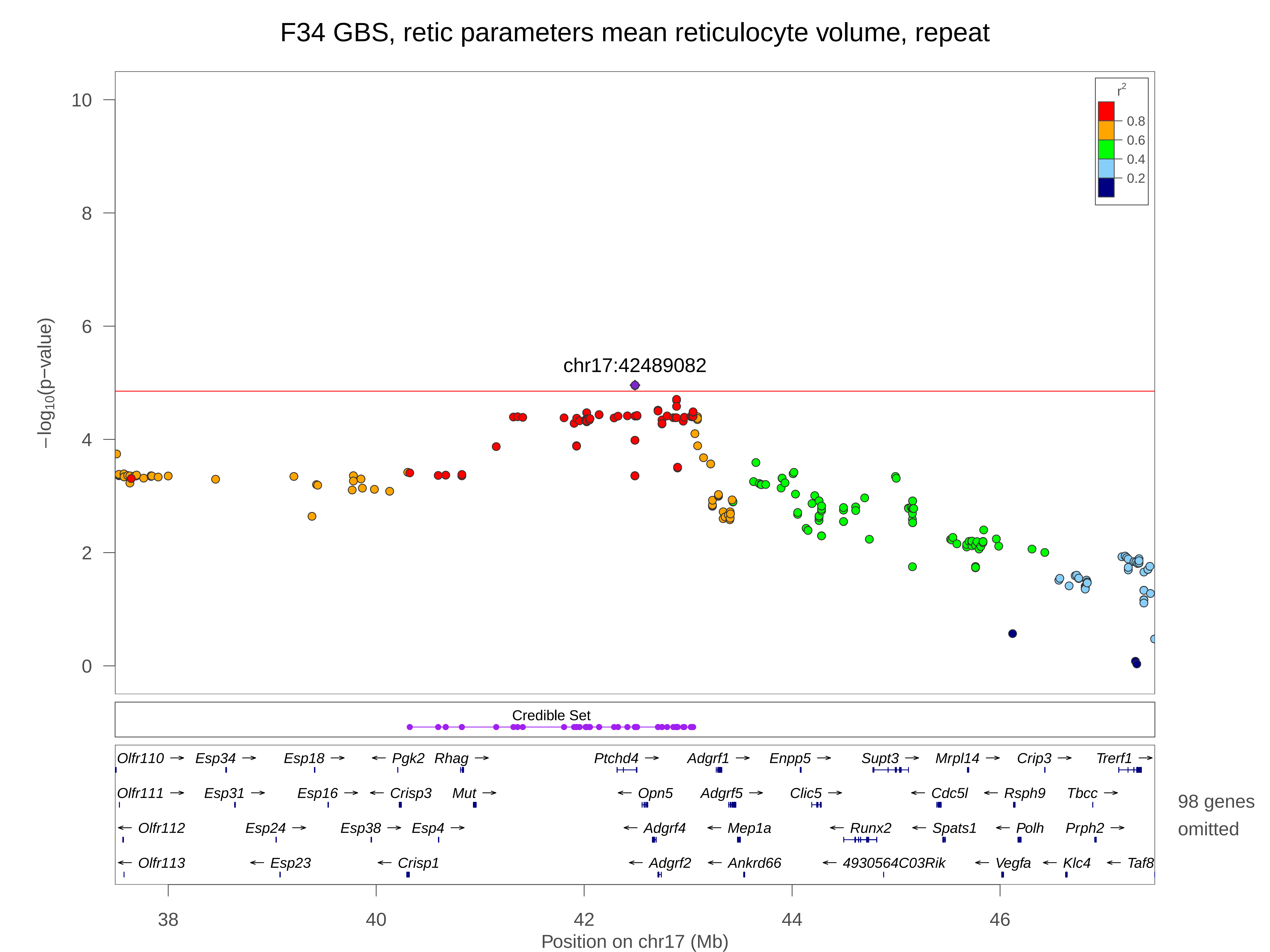
F34 GBS, retic parameters mean reticulocyte volume, repeat
−log10(p−value)
chr17:42489082
r2
0.8
0.6
0.4
0.2
Credible Set
Olfr110 → Esp34 → Esp18 → ← Pgk2 Rhag → Ptchd4 → Adgrf1 → Enpp5 → Supt3 → Mrpl14 → Crip3 → Trerf1 →
Olfr111 → Esp31 → Esp16 → ← Crisp3 Mut → ← Opn5 Adgrf5 → Clic5 → ← Cdc5l ← Rsph9 Tbcc →
← Olfr112 Esp24 → Esp38 → Esp4 → ← Adgrf4 ← Mep1a ← Runx2 ← Spats1 ← Polh Prph2 →
← Olfr113 ← Esp23 ← Crisp1 ← Adgrf2 ← Ankrd66 ← 4930564C03Rik ← Vegfa ← Klc4 ← Taf8
98 genes omitted
Position on chr17 (Mb)

date: Thu Oct 25 17:08:28 2018
build: F34GBS
display range: chr17:37489082−47489082 [37489082−47489082]
hilite range: 0 − 0 [ 0 − 0 ]
reference SNP: chr17:42489082
number of SNPs plotted: 292
min P−value: 1.1E−5 [chr17:42489082]
max P−value: 9.26E−1 [chr17:47315006]
omitted Genes: Olfr114, Olfr115, Olfr116
omitted Genes: Olfr117, Olfr118, Olfr119
omitted Genes: Olfr120, Olfr121, Olfr122
omitted Genes: Olfr123, Olfr124, Olfr125
omitted Genes: Olfr126, Olfr127, Olfr128
omitted Genes: Olfr761, Olfr129, Olfr130
omitted Genes: Olfr131, Olfr132, Olfr133
omitted Genes: Olfr134, Olfr135, Olfr138
omitted Genes: Olfr137, Olfr136, Esp36
omitted Genes: Esp15, Esp8, Esp6
omitted Genes: Gm44501, Esp5, Esp3
omitted Genes: Esp1, Crisp2, 9130008F23Rik
omitted Genes: Glyatl3, Cenpq, Cd2ap
omitted Genes: Tnfrsf21, E130008D07Rik, Pla2g7
omitted Genes: 1700071M16Rik, Tdrd6, Slc25a27
omitted Genes: Cyp39a1, Rcan2, Enpp4
omitted Genes: B230354K17Rik, Aars2, Tcte1
omitted Genes: Tmem151b, Nfkbie, Slc35b2
omitted Genes: Hsp90ab1, Slc29a1, Gm7325
omitted Genes: Capn11, Tmem63b, 1600014C23Rik
omitted Genes: Mrps18a, Mad2l1bp, Gtpbp2
omitted Genes: Xpo5, Mir693, Polr1c
omitted Genes: Yipf3, Lrrc73, Tjap1
omitted Genes: Dlk2, Abcc10, Zfp318
omitted Genes: Slc22a7, Ttbk1, Dnph1
omitted Genes: Cul9, Srf, Mir6976
omitted Genes: Ptk7, Mrpl2, Cul7
omitted Genes: Rrp36, Klhdc3, Mea1
omitted Genes: Ppp2r5d, Pex6, Gnmt
omitted Genes: Cnpy3, Ptcra, 2310039H08Rik
omitted Genes: Rpl7l1, Gltscr1l, Ubr2
Make more plots at http://csg.sph.umich.edu/locuszoom/
omitted Genes: Mrps10, Guca1b, Guca1a

Fine−mapping Regions

| snp | chr | pos (Mb) | pp | group | color |
|---|---|---|---|---|---|
| chr17:40322995 | 17 | 40.32299 | 0.001925994 | Credible Set | purple |
| chr17:40596316 | 17 | 40.59632 | 0.001747708 | Credible Set | purple |
| chr17:40668615 | 17 | 40.66862 | 0.001767481 | Credible Set | purple |
| chr17:40668642 | 17 | 40.66864 | 0.001767481 | Credible Set | purple |
| chr17:40668650 | 17 | 40.66865 | 0.001767481 | Credible Set | purple |
| chr17:40823871 | 17 | 40.82387 | 0.001728030 | Credible Set | purple |
| chr17:40824165 | 17 | 40.82417 | 0.001813987 | Credible Set | purple |
| chr17:41154429 | 17 | 41.15443 | 0.005256147 | Credible Set | purple |
| chr17:41320112 | 17 | 41.32011 | 0.016393635 | Credible Set | purple |
| chr17:41359808 | 17 | 41.35981 | 0.016568127 | Credible Set | purple |
| chr17:41409714 | 17 | 41.40971 | 0.016183259 | Credible Set | purple |
| chr17:41806772 | 17 | 41.80677 | 0.015882965 | Credible Set | purple |
| chr17:41903568 | 17 | 41.90357 | 0.012834321 | Credible Set | purple |
| chr17:41926884 | 17 | 41.92688 | 0.005444641 | Credible Set | purple |
| chr17:41926899 | 17 | 41.92690 | 0.005334929 | Credible Set | purple |
| chr17:41928375 | 17 | 41.92838 | 0.015676900 | Credible Set | purple |
| chr17:41956148 | 17 | 41.95615 | 0.014177500 | Credible Set | purple |
| chr17:42013920 | 17 | 42.01392 | 0.014708796 | Credible Set | purple |
| chr17:42023895 | 17 | 42.02390 | 0.013943435 | Credible Set | purple |
| chr17:42024466 | 17 | 42.02447 | 0.013623376 | Credible Set | purple |
| chr17:42024515 | 17 | 42.02452 | 0.019357133 | Credible Set | purple |
| chr17:42024583 | 17 | 42.02458 | 0.019312048 | Credible Set | purple |
| chr17:42024700 | 17 | 42.02470 | 0.015320345 | Credible Set | purple |
| chr17:42024755 | 17 | 42.02475 | 0.014674524 | Credible Set | purple |
| chr17:42024810 | 17 | 42.02481 | 0.014743765 | Credible Set | purple |
| chr17:42025311 | 17 | 42.02531 | 0.014105814 | Credible Set | purple |
| chr17:42050395 | 17 | 42.05040 | 0.014971741 | Credible Set | purple |
| chr17:42050673 | 17 | 42.05067 | 0.014596565 | Credible Set | purple |
| chr17:42056191 | 17 | 42.05619 | 0.015407444 | Credible Set | purple |
| chr17:42143893 | 17 | 42.14389 | 0.017848640 | Credible Set | purple |
| chr17:42143896 | 17 | 42.14390 | 0.018014846 | Credible Set | purple |
| chr17:42288066 | 17 | 42.28807 | 0.015812708 | Credible Set | purple |
| chr17:42325798 | 17 | 42.32580 | 0.016958613 | Credible Set | purple |
| chr17:42416069 | 17 | 42.41607 | 0.017164316 | Credible Set | purple |
| chr17:42488210 | 17 | 42.48821 | 0.001731401 | Credible Set | purple |
| chr17:42488216 | 17 | 42.48822 | 0.001728455 | Credible Set | purple |

Fine−mapping Regions

| snp | chr | pos (Mb) | pp | group | color |
|---|---|---|---|---|---|
| chr17:42488241 | 17 | 42.48824 | 0.001738098 | Credible Set | purple |
| chr17:42488701 | 17 | 42.48870 | 0.006698897 | Credible Set | purple |
| chr17:42488808 | 17 | 42.48881 | 0.006713565 | Credible Set | purple |
| chr17:42488822 | 17 | 42.48882 | 0.006699569 | Credible Set | purple |
| chr17:42489082 | 17 | 42.48908 | 0.056233892 | Credible Set | purple |
| chr17:42489130 | 17 | 42.48913 | 0.055350303 | Credible Set | purple |
| chr17:42489886 | 17 | 42.48989 | 0.016997208 | Credible Set | purple |
| chr17:42508272 | 17 | 42.50827 | 0.017063167 | Credible Set | purple |
| chr17:42508276 | 17 | 42.50828 | 0.017063167 | Credible Set | purple |
| chr17:42508349 | 17 | 42.50835 | 0.017412448 | Credible Set | purple |
| chr17:42710063 | 17 | 42.71006 | 0.021358708 | Credible Set | purple |
| chr17:42710132 | 17 | 42.71013 | 0.020640815 | Credible Set | purple |
| chr17:42748429 | 17 | 42.74843 | 0.014648457 | Credible Set | purple |
| chr17:42748810 | 17 | 42.74881 | 0.012528564 | Credible Set | purple |
| chr17:42748868 | 17 | 42.74887 | 0.012720279 | Credible Set | purple |
| chr17:42798191 | 17 | 42.79819 | 0.017098002 | Credible Set | purple |
| chr17:42857918 | 17 | 42.85792 | 0.015971822 | Credible Set | purple |
| chr17:42879412 | 17 | 42.87941 | 0.015951100 | Credible Set | purple |
| chr17:42888048 | 17 | 42.88805 | 0.015954438 | Credible Set | purple |
| chr17:42888694 | 17 | 42.88869 | 0.031411853 | Credible Set | purple |
| chr17:42888751 | 17 | 42.88875 | 0.032480984 | Credible Set | purple |
| chr17:42888758 | 17 | 42.88876 | 0.024729093 | Credible Set | purple |
| chr17:42898383 | 17 | 42.89838 | 0.002394195 | Credible Set | purple |
| chr17:42898392 | 17 | 42.89839 | 0.002318784 | Credible Set | purple |
| chr17:42898421 | 17 | 42.89842 | 0.002388373 | Credible Set | purple |
| chr17:42953111 | 17 | 42.95311 | 0.013939538 | Credible Set | purple |
| chr17:42963409 | 17 | 42.96341 | 0.016270304 | Credible Set | purple |
| chr17:43026300 | 17 | 43.02630 | 0.016705390 | Credible Set | purple |
| chr17:43026388 | 17 | 43.02639 | 0.016705390 | Credible Set | purple |
| chr17:43037723 | 17 | 43.03772 | 0.016470016 | Credible Set | purple |
| chr17:43047345 | 17 | 43.04734 | 0.016767498 | Credible Set | purple |
| chr17:43047599 | 17 | 43.04760 | 0.019782062 | Credible Set | purple |
| chr17:43047622 | 17 | 43.04762 | 0.020063233 | Credible Set | purple |

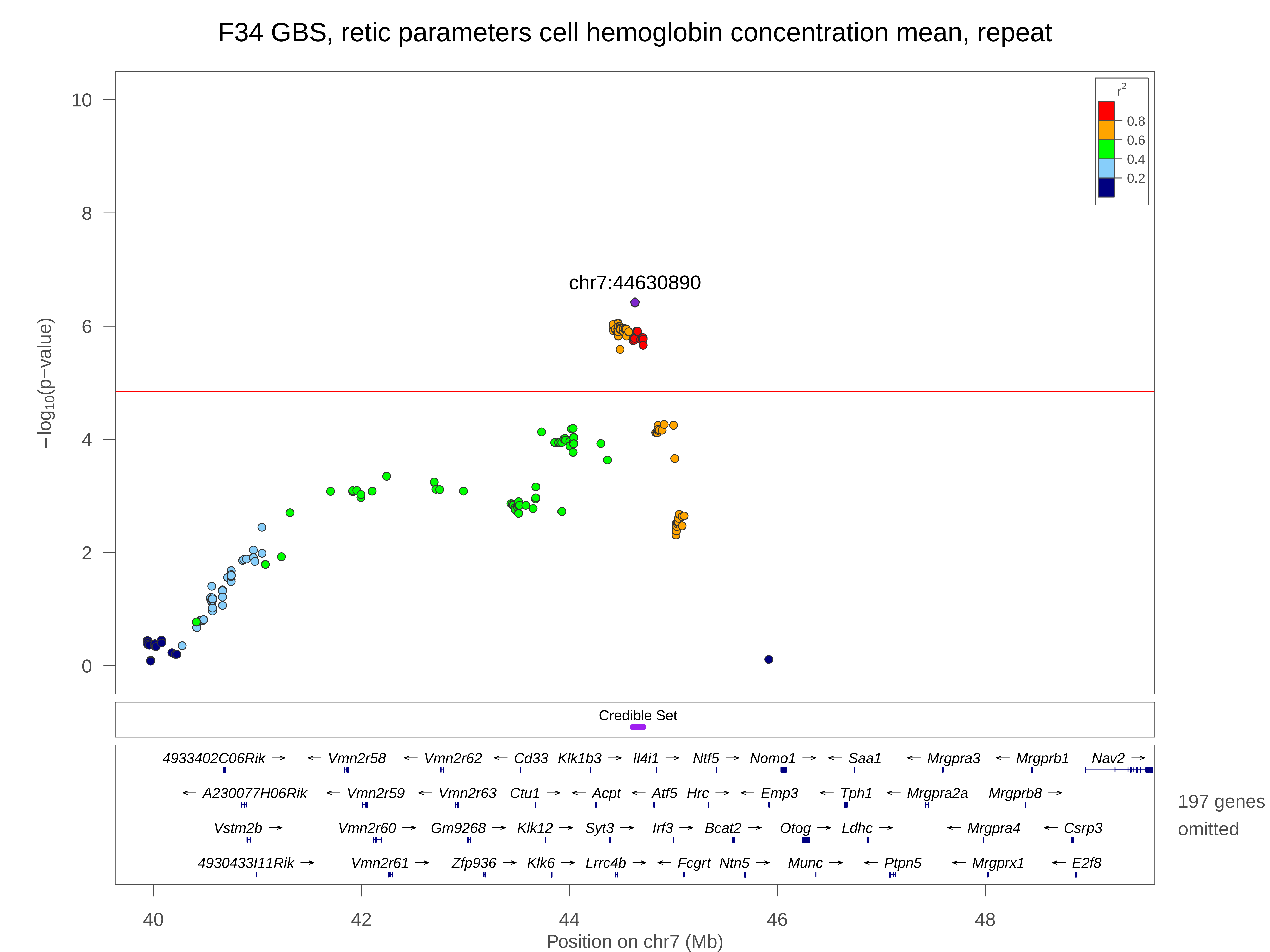

F34 GBS, retic parameters cell hemoglobin concentration mean, repeat
chr7:44630890
r2
0.8
0.6
0.4
0.2
−log10(p−value)
10
8
6
4
2
0
Credible Set
4933402C06Rik → ← Vmn2r58 ← Vmn2r62 ← Cd33 Klk1b3 → Il4i1 → Ntf5 → Nomo1 → ← Saa1 ← Mrgpra3 ← Mrgprb1 Nav2 →
← A230077H06Rik ← Vmn2r59 ← Vmn2r63 Ctu1 → ← Acpt ← Atf5 Hrc → ← Emp3 ← Tph1 ← Mrgpra2a Mrgprb8 →
Vstm2b → Vmn2r60 → Gm9268 → Klk12 → Syt3 → Irf3 → Bcat2 → Otog → Ldhc → ← Mrgpra4 ← Csrp3
4930433I11Rik → Vmn2r61 → Zfp936 → Klk6 → Lrrc4b → ← Fcgrt Ntn5 → Munc → ← Ptpn5 ← Mrgprx1 ← E2f8
197 genes omitted
40
42
44
46
48
Position on chr7 (Mb)

date: Thu Oct 25 17:09:29 2018
build: F34GBS
display range: chr7:39630890−49630890 [39630890−49630890]
hilite range: 0 − 0 [ 0 − 0 ]
reference SNP: chr7:44630890
number of SNPs plotted: 247
min P−value: 3.81E−7 [chr7:44630890]
max P−value: 8.26E−1 [chr7:39972874]
omitted Genes: Gm4884, 1700110I07Rik, Gm5592
omitted Genes: AI987944, Vmn2r57, AW146154
omitted Genes: Zfp788, Vmn2r−ps54, Vmn2r−ps60
omitted Genes: 4933421I07Rik, Zfp141, 9830147E19Rik
omitted Genes: Gm5595, Gm2381, EU599041
omitted Genes: Zfp715, Siglecf, Siglecg
omitted Genes: Lim2, Nkg7, Cldnd2
omitted Genes: Etfb, Mir7051, Vsig10l
omitted Genes: Iglon5, 4931406B18Rik, Zfp658
omitted Genes: Zfp719, Zfp819, Ceacam18
omitted Genes: Siglece, Klk14, Klk13
omitted Genes: 2310002F09Rik, Klk11, Klk10
omitted Genes: Klk9, Klk8, Klk7
omitted Genes: Klk5, Klk4, Klk15
omitted Genes: Klk1b7−ps, Klk1b8, Klk1b1
omitted Genes: Klk1b9, Klk1b11, Klk1b26
omitted Genes: Klk1b27, Klk1b21, Klk1b22
omitted Genes: Klk1b16, Klk1b24, Klk1b4
omitted Genes: Klk1b5, Klk1, 1700028J19Rik
omitted Genes: 2410002F23Rik, Snord88c, Snord88a
omitted Genes: Clec11a, Shank1, 1700008O03Rik
omitted Genes: Aspdh, Josd2, Mir7052
omitted Genes: 5430431A17Rik, Emc10, Fam71e1
omitted Genes: Mybpc2, Spib, Pold1
omitted Genes: Mir7053, Nr1h2, Napsa
omitted Genes: Kcnc3, Myh14, Izumo2
omitted Genes: Zfp473, Vrk3, Nup62
omitted Genes: Gm21948, Tbc1d17, Akt1s1
omitted Genes: Mir707, Pnkp, Ptov1
omitted Genes: Med25, Fuz, Ap2a1
omitted Genes: Tsks, Cpt1c, Prmt1
omitted Genes: Gm15545, Bcl2l12, Scaf1

Make more plots at http://csg.sph.umich.edu/locuszoom/

Fine−mapping Regions

| snp | chr | pos (Mb) | pp | group | color |
|---|---|---|---|---|---|
| chr7:44612740 | 7 | 44.61274 | 0.04036894 | Credible Set | purple |
| chr7:44613080 | 7 | 44.61308 | 0.03742242 | Credible Set | purple |
| chr7:44623189 | 7 | 44.62319 | 0.04048152 | Credible Set | purple |
| chr7:44625757 | 7 | 44.62576 | 0.03838420 | Credible Set | purple |
| chr7:44628444 | 7 | 44.62844 | 0.04138907 | Credible Set | purple |
| chr7:44630865 | 7 | 44.63087 | 0.16463846 | Credible Set | purple |
| chr7:44630890 | 7 | 44.63089 | 0.16806321 | Credible Set | purple |
| chr7:44647914 | 7 | 44.64791 | 0.05468613 | Credible Set | purple |
| chr7:44655619 | 7 | 44.65562 | 0.05331551 | Credible Set | purple |
| chr7:44655640 | 7 | 44.65564 | 0.05403463 | Credible Set | purple |
| chr7:44685749 | 7 | 44.68575 | 0.03949794 | Credible Set | purple |
| chr7:44700610 | 7 | 44.70061 | 0.04246376 | Credible Set | purple |
| chr7:44700692 | 7 | 44.70069 | 0.04002805 | Credible Set | purple |
| chr7:44708823 | 7 | 44.70882 | 0.04002805 | Credible Set | purple |
| chr7:44708981 | 7 | 44.70898 | 0.04215076 | Credible Set | purple |
| chr7:44708990 | 7 | 44.70899 | 0.04006775 | Credible Set | purple |
| chr7:44708997 | 7 | 44.70900 | 0.03148981 | Credible Set | purple |
| chr7:44709016 | 7 | 44.70902 | 0.03148981 | Credible Set | purple |

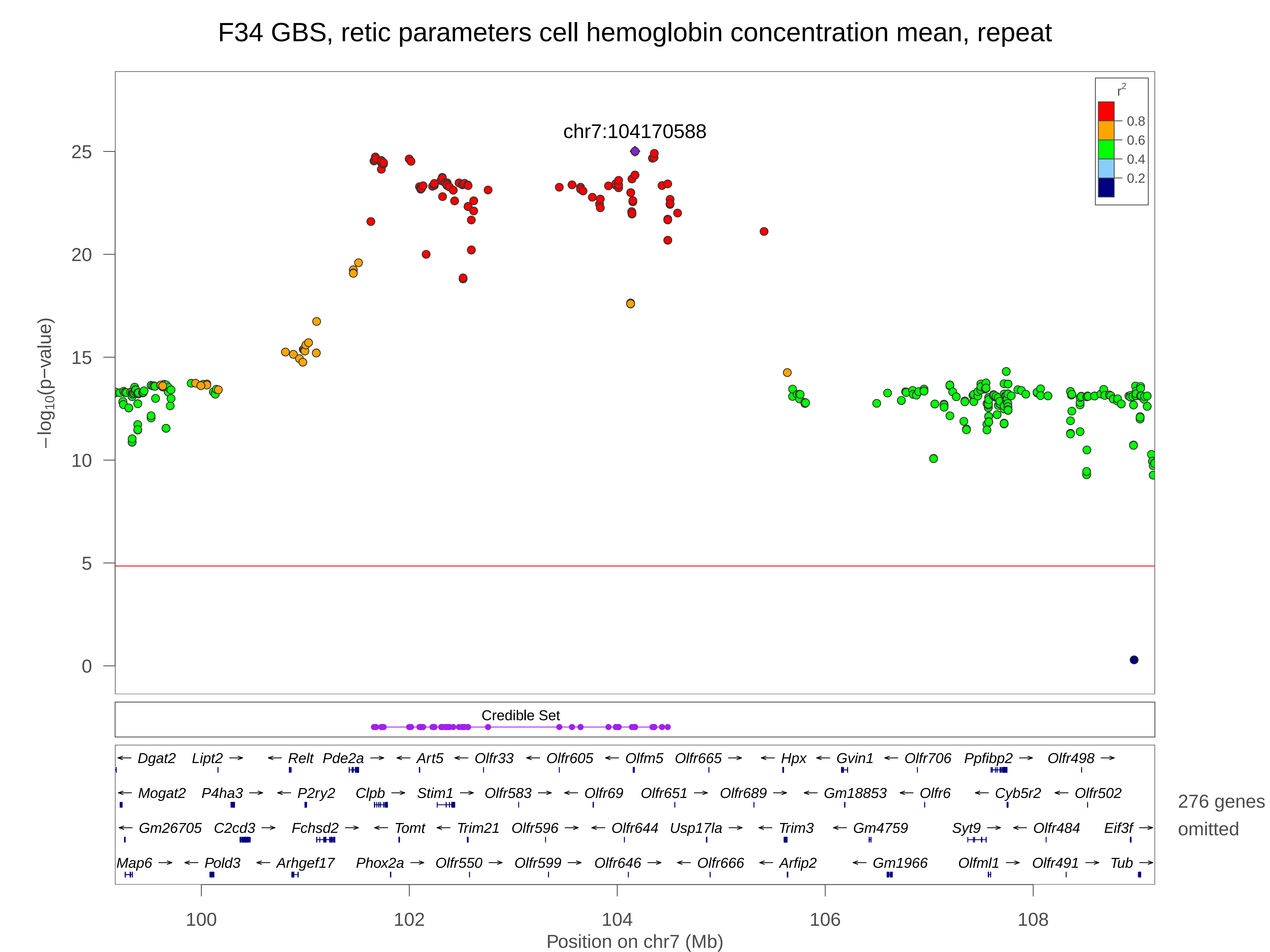
F34 GBS, retic parameters cell hemoglobin concentration mean, repeat
chr7:104170588
r2
0.8
0.6
0.4
0.2
−log10(p−value)
25
20
15
10
5
0
Credible Set
← Dgat2
Lipt2 →
← Relt
Pde2a →
← Art5
← Olfr33
← Olfr605
← Olfm5
Olfr665 →
← Hpx
← Gvin1
← Olfr706
Ppfibp2 →
Olfr498 →
← Mogat2
P4ha3 →
← P2ry2
Clpb →
Stim1 →
Olfr583 →
← Olfr69
Olfr651 →
Olfr689 →
← Gm18853
← Olfr6
← Cyb5r2
← Olfr502
← Gm26705
C2cd3 →
Fchsd2 →
← Tomt
← Trim21
Olfr596 →
← Olfr644
Usp17la →
← Trim3
← Gm4759
Syt9 →
← Olfr484
Eif3f →
Map6 →
← Pold3
← Arhgef17
Phox2a →
Olfr550 →
Olfr599 →
Olfr646 →
← Olfr666
← Arfip2
← Gm1966
Olfml1 →
Olfr491 →
Tub →
276 genes omitted
100
102
104
106
108
Position on chr7 (Mb)

date: Thu Oct 25 17:09:16 2018
build: F34GBS
display range: chr7:99170588−109170588 [99170588−109170588]
hilite range: 0 − 0 [ 0 − 0 ]
reference SNP: chr7:104170588
number of SNPs plotted: 464
min P−value: 9.7E−26 [chr7:104170588]
max P−value: 5.14E−1 [chr7:108970724]
omitted Genes: Serpinh1, Gdpd5, Klhl35
omitted Genes: Rps3, Snord15b, Snord15a
omitted Genes: Arrb1, Mir326, Tpbgl
omitted Genes: Slco2b1, Olfr520, Olfr521
omitted Genes: Neu3, Spcs2, Xrra1
omitted Genes: Rnf169, Chrdl2, Kcne3
omitted Genes: Pgm2l1, Gpx2−ps1, Ppme1
omitted Genes: Ucp3, Ucp2, Dnajb13
omitted Genes: Coa4, Mrpl48, Rab6a
omitted Genes: Plekhb1, Fam168a, Mir3102
omitted Genes: P2ry6, Atg16l2, Stard10
omitted Genes: Arap1, Mir139, Art2a−ps
omitted Genes: Art2b, Inppl1, Folr2
omitted Genes: Folr1, Anapc15, Lamtor1
omitted Genes: Lrrc51, Numa1, Il18bp
omitted Genes: Rnf121, Xndc1, Xntrpc
omitted Genes: Trpc2, Art1, Chrna10
omitted Genes: Nup98, Pgap2, Rhog
omitted Genes: Rrm1, Olfr543, Olfr544
omitted Genes: Olfr545, Olfr547, Olfr549
omitted Genes: Olfr551, Olfr552, Olfr553
omitted Genes: Olfr554, Olfr555, Olfr556
omitted Genes: Trim68, Olfr557, Olfr558
omitted Genes: Olfr559, Olfr78, Olfr560
omitted Genes: Olfr561, Olfr564, Olfr566
omitted Genes: Olfr568, Olfr569, Olfr570
omitted Genes: Olfr571, Olfr572, Olfr574
omitted Genes: Olfr575, Olfr576, Olfr577
omitted Genes: Olfr578, Olfr582, Olfr584
omitted Genes: Olfr585, Olfr586, Olfr589
omitted Genes: Olfr591, Olfr592, Olfr593
Make more plots at http://csg.sph.umich.edu/locuszoom/
omitted Genes: Olfr594, Usp17ld, Olfr597

Fine−mapping Regions

| snp | chr | pos (Mb) | pp | group | color |
|---|---|---|---|---|---|
| chr7:101660383 | 7 | 101.6604 | 0.027801217 | Credible Set | purple |
| chr7:101669186 | 7 | 101.6692 | 0.031049930 | Credible Set | purple |
| chr7:101673483 | 7 | 101.6735 | 0.043524010 | Credible Set | purple |
| chr7:101680140 | 7 | 101.6801 | 0.035559081 | Credible Set | purple |
| chr7:101731061 | 7 | 101.7311 | 0.029442197 | Credible Set | purple |
| chr7:101731110 | 7 | 101.7311 | 0.022726636 | Credible Set | purple |
| chr7:101731130 | 7 | 101.7311 | 0.011029991 | Credible Set | purple |
| chr7:101731658 | 7 | 101.7317 | 0.028189047 | Credible Set | purple |
| chr7:101751790 | 7 | 101.7518 | 0.019767205 | Credible Set | purple |
| chr7:101752580 | 7 | 101.7526 | 0.022135716 | Credible Set | purple |
| chr7:101752971 | 7 | 101.7530 | 0.022819663 | Credible Set | purple |
| chr7:101998679 | 7 | 101.9987 | 0.034974170 | Credible Set | purple |
| chr7:102016307 | 7 | 102.0163 | 0.026659422 | Credible Set | purple |
| chr7:102098054 | 7 | 102.0981 | 0.001606431 | Credible Set | purple |
| chr7:102110109 | 7 | 102.1101 | 0.001218167 | Credible Set | purple |
| chr7:102110501 | 7 | 102.1105 | 0.001359069 | Credible Set | purple |
| chr7:102116318 | 7 | 102.1163 | 0.001317094 | Credible Set | purple |
| chr7:102116430 | 7 | 102.1164 | 0.001317094 | Credible Set | purple |
| chr7:102116478 | 7 | 102.1165 | 0.001341070 | Credible Set | purple |
| chr7:102132562 | 7 | 102.1326 | 0.001789863 | Credible Set | purple |
| chr7:102223396 | 7 | 102.2234 | 0.001684376 | Credible Set | purple |
| chr7:102240695 | 7 | 102.2407 | 0.001845753 | Credible Set | purple |
| chr7:102241191 | 7 | 102.2412 | 0.002162235 | Credible Set | purple |
| chr7:102241211 | 7 | 102.2412 | 0.002304696 | Credible Set | purple |
| chr7:102241212 | 7 | 102.2412 | 0.002300427 | Credible Set | purple |
| chr7:102306968 | 7 | 102.3070 | 0.003118473 | Credible Set | purple |
| chr7:102306978 | 7 | 102.3070 | 0.003347757 | Credible Set | purple |
| chr7:102317213 | 7 | 102.3172 | 0.004528046 | Credible Set | purple |
| chr7:102317246 | 7 | 102.3172 | 0.004161109 | Credible Set | purple |
| chr7:102344970 | 7 | 102.3450 | 0.002447528 | Credible Set | purple |
| chr7:102363897 | 7 | 102.3639 | 0.001811027 | Credible Set | purple |
| chr7:102364204 | 7 | 102.3642 | 0.001991464 | Credible Set | purple |
| chr7:102364223 | 7 | 102.3642 | 0.002469608 | Credible Set | purple |
| chr7:102364315 | 7 | 102.3643 | 0.002114340 | Credible Set | purple |
| chr7:102364326 | 7 | 102.3643 | 0.002114340 | Credible Set | purple |
| chr7:102364411 | 7 | 102.3644 | 0.001962401 | Credible Set | purple |

Fine−mapping Regions

| snp | chr | pos (Mb) | pp | group | color |
|---|---|---|---|---|---|
| chr7:102385520 | 7 | 102.3855 | 0.001560069 | Credible Set | purple |
| chr7:102385536 | 7 | 102.3855 | 0.001573554 | Credible Set | purple |
| chr7:102423165 | 7 | 102.4232 | 0.001084442 | Credible Set | purple |
| chr7:102478469 | 7 | 102.4785 | 0.002439839 | Credible Set | purple |
| chr7:102510475 | 7 | 102.5105 | 0.002079327 | Credible Set | purple |
| chr7:102510607 | 7 | 102.5106 | 0.001969508 | Credible Set | purple |
| chr7:102516878 | 7 | 102.5169 | 0.002144741 | Credible Set | purple |
| chr7:102531652 | 7 | 102.5317 | 0.002332695 | Credible Set | purple |
| chr7:102561708 | 7 | 102.5617 | 0.001958134 | Credible Set | purple |
| chr7:102565410 | 7 | 102.5654 | 0.001808287 | Credible Set | purple |
| chr7:102757499 | 7 | 102.7575 | 0.001115733 | Credible Set | purple |
| chr7:103442570 | 7 | 103.4426 | 0.001511749 | Credible Set | purple |
| chr7:103564836 | 7 | 103.5648 | 0.001968672 | Credible Set | purple |
| chr7:103646439 | 7 | 103.6464 | 0.001519819 | Credible Set | purple |
| chr7:103646709 | 7 | 103.6467 | 0.001225704 | Credible Set | purple |
| chr7:103915503 | 7 | 103.9155 | 0.001757102 | Credible Set | purple |
| chr7:103986792 | 7 | 103.9868 | 0.001891418 | Credible Set | purple |
| chr7:103986795 | 7 | 103.9868 | 0.001891418 | Credible Set | purple |
| chr7:103986857 | 7 | 103.9869 | 0.002271127 | Credible Set | purple |
| chr7:103986860 | 7 | 103.9869 | 0.002271127 | Credible Set | purple |
| chr7:104012808 | 7 | 104.0128 | 0.001422940 | Credible Set | purple |
| chr7:104013370 | 7 | 104.0134 | 0.001937644 | Credible Set | purple |
| chr7:104013395 | 7 | 104.0134 | 0.001946171 | Credible Set | purple |
| chr7:104013467 | 7 | 104.0135 | 0.003250796 | Credible Set | purple |
| chr7:104141343 | 7 | 104.1413 | 0.003794442 | Credible Set | purple |
| chr7:104170479 | 7 | 104.1705 | 0.005874467 | Credible Set | purple |
| chr7:104170506 | 7 | 104.1705 | 0.005802691 | Credible Set | purple |
| chr7:104170518 | 7 | 104.1705 | 0.005834985 | Credible Set | purple |
| chr7:104170522 | 7 | 104.1705 | 0.005815978 | Credible Set | purple |
| chr7:104170543 | 7 | 104.1705 | 0.005836372 | Credible Set | purple |
| chr7:104170588 | 7 | 104.1706 | 0.082301601 | Credible Set | purple |
| chr7:104170594 | 7 | 104.1706 | 0.081799969 | Credible Set | purple |
| chr7:104170618 | 7 | 104.1706 | 0.079054313 | Credible Set | purple |
| chr7:104170662 | 7 | 104.1707 | 0.079054313 | Credible Set | purple |
| chr7:104338784 | 7 | 104.3388 | 0.037642915 | Credible Set | purple |
| chr7:104338810 | 7 | 104.3388 | 0.037589759 | Credible Set | purple |

Fine−mapping Regions

| snp | chr | pos (Mb) | pp | group | color |
|---|---|---|---|---|---|
| chr7:104353491 | 7 | 104.3535 | 0.040828316 | Credible Set | purple |
| chr7:104356230 | 7 | 104.3562 | 0.064752467 | Credible Set | purple |
| chr7:104430205 | 7 | 104.4302 | 0.001824304 | Credible Set | purple |
| chr7:104485236 | 7 | 104.4852 | 0.002191823 | Credible Set | purple |

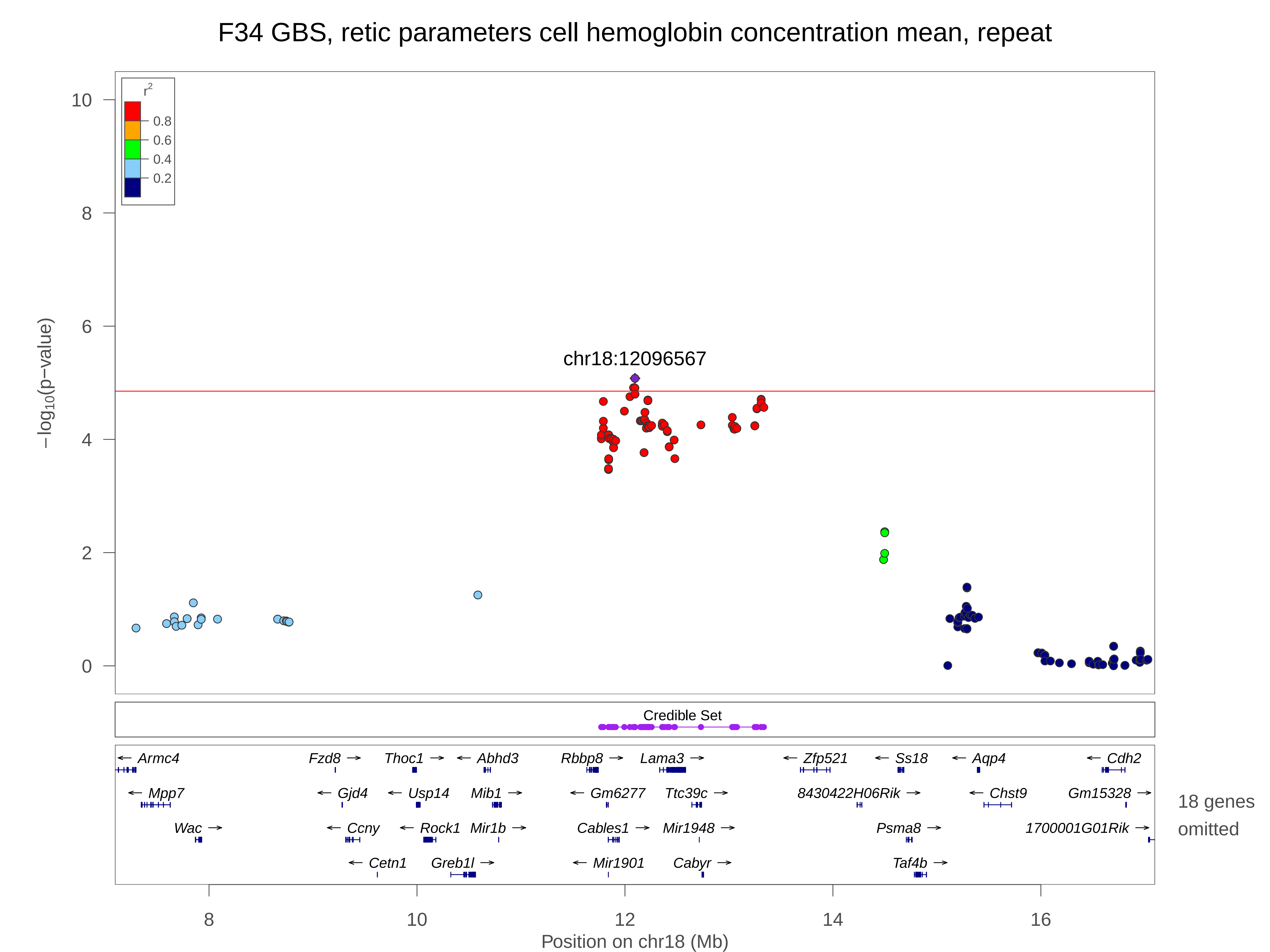

F34 GBS, retic parameters cell hemoglobin concentration mean, repeat
r2
0.8
0.6
0.4
0.2
10
8
6
4
2
0
−log10(p−value)
chr18:12096567
Credible Set
Armc4
Fzd8
Thoc1
Abhd3
Rbbp8
Lama3
Zfp521
Ss18
Aqp4
Cdh2
Mpp7
Gjd4
Usp14
Mib1
Gm6277
Ttc39c
8430422H06Rik
Chst9
Gm15328
Wac
Ccny
Rock1
Mir1b
Cables1
Mir1948
Psma8
1700001G01Rik
Cetn1
Greb1l
Mir1901
Cabyr
Taf4b
18 genes omitted
8
10
12
14
16
Position on chr18 (Mb)

| | |
|---|---|
| date: | Thu Oct 25 17:09:01 2018 |
| build: | F34GBS |
| display range: | chr18:7096567−17096567 [7096567−17096567] |
| hilite range: | 0 − 0 [ 0 − 0 ] |
| reference SNP: | chr18:12096567 |
| number of SNPs plotted: | 193 |
| min P−value: | 8.35E−6 [chr18:12096567] |
| max P−value: | 9.99E−1 [chr18:16699294] |
| omitted Genes: | Colec12, Esco1, Snrpd1 |
| omitted Genes: | 4930563E18Rik, Mir133a−1, Mir1a−2 |
| omitted Genes: | 1010001N08Rik, Gata6, Tmem241 |
| omitted Genes: | Riok3, 3110002H16Rik, Npc1 |
| omitted Genes: | Ankrd29, Osbpl1a, Impact |
| omitted Genes: | Hrh4, Kctd1, 4933424G05Rik |

Make more plots at http://csg.sph.umich.edu/locuszoom/

Fine−mapping Regions

| snp | chr | pos (Mb) | pp | group | color |
|---|---|---|---|---|---|
| chr18:11772906 | 18 | 11.77291 | 0.006021185 | Credible Set | purple |
| chr18:11772936 | 18 | 11.77294 | 0.006021185 | Credible Set | purple |
| chr18:11773024 | 18 | 11.77302 | 0.005824685 | Credible Set | purple |
| chr18:11773062 | 18 | 11.77306 | 0.006746175 | Credible Set | purple |
| chr18:11792180 | 18 | 11.79218 | 0.008775780 | Credible Set | purple |
| chr18:11792281 | 18 | 11.79228 | 0.024634261 | Credible Set | purple |
| chr18:11792295 | 18 | 11.79229 | 0.011480610 | Credible Set | purple |
| chr18:11841074 | 18 | 11.84107 | 0.006244823 | Credible Set | purple |
| chr18:11843471 | 18 | 11.84347 | 0.002693669 | Credible Set | purple |
| chr18:11843539 | 18 | 11.84354 | 0.002718494 | Credible Set | purple |
| chr18:11843691 | 18 | 11.84369 | 0.006799667 | Credible Set | purple |
| chr18:11849087 | 18 | 11.84909 | 0.005863858 | Credible Set | purple |
| chr18:11868302 | 18 | 11.86830 | 0.005950931 | Credible Set | purple |
| chr18:11868573 | 18 | 11.86857 | 0.005835239 | Credible Set | purple |
| chr18:11884565 | 18 | 11.88457 | 0.005253571 | Credible Set | purple |
| chr18:11888824 | 18 | 11.88882 | 0.005746699 | Credible Set | purple |
| chr18:11890738 | 18 | 11.89074 | 0.004179073 | Credible Set | purple |
| chr18:11890742 | 18 | 11.89074 | 0.004133836 | Credible Set | purple |
| chr18:11910882 | 18 | 11.91088 | 0.005415126 | Credible Set | purple |
| chr18:11994328 | 18 | 11.99433 | 0.016922237 | Credible Set | purple |
| chr18:12047734 | 18 | 12.04773 | 0.029654538 | Credible Set | purple |
| chr18:12082652 | 18 | 12.08265 | 0.041930306 | Credible Set | purple |
| chr18:12095502 | 18 | 12.09550 | 0.041391823 | Credible Set | purple |
| chr18:12095506 | 18 | 12.09551 | 0.041537419 | Credible Set | purple |
| chr18:12096567 | 18 | 12.09657 | 0.060323772 | Credible Set | purple |
| chr18:12096649 | 18 | 12.09665 | 0.032626185 | Credible Set | purple |
| chr18:12147962 | 18 | 12.14796 | 0.011623939 | Credible Set | purple |
| chr18:12159851 | 18 | 12.15985 | 0.011641308 | Credible Set | purple |
| chr18:12166070 | 18 | 12.16607 | 0.011794411 | Credible Set | purple |
| chr18:12183803 | 18 | 12.18380 | 0.003432600 | Credible Set | purple |
| chr18:12183818 | 18 | 12.18382 | 0.003432600 | Credible Set | purple |
| chr18:12183819 | 18 | 12.18382 | 0.003432600 | Credible Set | purple |
| chr18:12183957 | 18 | 12.18396 | 0.012010534 | Credible Set | purple |
| chr18:12188720 | 18 | 12.18872 | 0.012321028 | Credible Set | purple |
| chr18:12192191 | 18 | 12.19219 | 0.016185138 | Credible Set | purple |
| chr18:12205743 | 18 | 12.20574 | 0.008773730 | Credible Set | purple |

Fine−mapping Regions

| snp | chr | pos (Mb) | pp | group | color |
|---|---|---|---|---|---|
| chr18:12209149 | 18 | 12.20915 | 0.010651880 | Credible Set | purple |
| chr18:12209157 | 18 | 12.20916 | 0.010651880 | Credible Set | purple |
| chr18:12220479 | 18 | 12.22048 | 0.025270426 | Credible Set | purple |
| chr18:12220534 | 18 | 12.22053 | 0.026067156 | Credible Set | purple |
| chr18:12220537 | 18 | 12.22054 | 0.025472409 | Credible Set | purple |
| chr18:12220581 | 18 | 12.22058 | 0.009689983 | Credible Set | purple |
| chr18:12228637 | 18 | 12.22864 | 0.009765648 | Credible Set | purple |
| chr18:12232882 | 18 | 12.23288 | 0.008945730 | Credible Set | purple |
| chr18:12232913 | 18 | 12.23291 | 0.008943798 | Credible Set | purple |
| chr18:12232928 | 18 | 12.23293 | 0.008943798 | Credible Set | purple |
| chr18:12232935 | 18 | 12.23293 | 0.008943798 | Credible Set | purple |
| chr18:12233114 | 18 | 12.23311 | 0.009390026 | Credible Set | purple |
| chr18:12233710 | 18 | 12.23371 | 0.009688850 | Credible Set | purple |
| chr18:12233723 | 18 | 12.23372 | 0.009167282 | Credible Set | purple |
| chr18:12256132 | 18 | 12.25613 | 0.009693287 | Credible Set | purple |
| chr18:12358152 | 18 | 12.35815 | 0.009697208 | Credible Set | purple |
| chr18:12359533 | 18 | 12.35953 | 0.010684856 | Credible Set | purple |
| chr18:12359979 | 18 | 12.35998 | 0.009429262 | Credible Set | purple |
| chr18:12378538 | 18 | 12.37854 | 0.009933191 | Credible Set | purple |
| chr18:12406944 | 18 | 12.40694 | 0.007668154 | Credible Set | purple |
| chr18:12406948 | 18 | 12.40695 | 0.008005364 | Credible Set | purple |
| chr18:12425352 | 18 | 12.42535 | 0.004307831 | Credible Set | purple |
| chr18:12425400 | 18 | 12.42540 | 0.004253809 | Credible Set | purple |
| chr18:12473083 | 18 | 12.47308 | 0.005566343 | Credible Set | purple |
| chr18:12479736 | 18 | 12.47974 | 0.002725865 | Credible Set | purple |
| chr18:12730645 | 18 | 12.73065 | 0.009953065 | Credible Set | purple |
| chr18:13032150 | 18 | 13.03215 | 0.013298070 | Credible Set | purple |
| chr18:13032510 | 18 | 13.03251 | 0.009773893 | Credible Set | purple |
| chr18:13049016 | 18 | 13.04902 | 0.008479338 | Credible Set | purple |
| chr18:13049169 | 18 | 13.04917 | 0.008908733 | Credible Set | purple |
| chr18:13053875 | 18 | 13.05387 | 0.008425772 | Credible Set | purple |
| chr18:13058650 | 18 | 13.05865 | 0.009346800 | Credible Set | purple |
| chr18:13076442 | 18 | 13.07644 | 0.008689557 | Credible Set | purple |
| chr18:13248719 | 18 | 13.24872 | 0.009578204 | Credible Set | purple |
| chr18:13248733 | 18 | 13.24873 | 0.009684267 | Credible Set | purple |
| chr18:13248742 | 18 | 13.24874 | 0.009578879 | Credible Set | purple |

Fine−mapping Regions

| snp | chr | pos (Mb) | pp | group | color |
|---|---|---|---|---|---|
| chr18:13269113 | 18 | 13.26911 | 0.01849536 | Credible Set | purple |
| chr18:13269114 | 18 | 13.26911 | 0.01892845 | Credible Set | purple |
| chr18:13309638 | 18 | 13.30964 | 0.02681659 | Credible Set | purple |
| chr18:13310189 | 18 | 13.31019 | 0.02614172 | Credible Set | purple |
| chr18:13310299 | 18 | 13.31030 | 0.02332506 | Credible Set | purple |
| chr18:13335693 | 18 | 13.33569 | 0.01955042 | Credible Set | purple |

### F34 GBS, retic parameters concentration of hemoglobin in reticulocytes, repeat

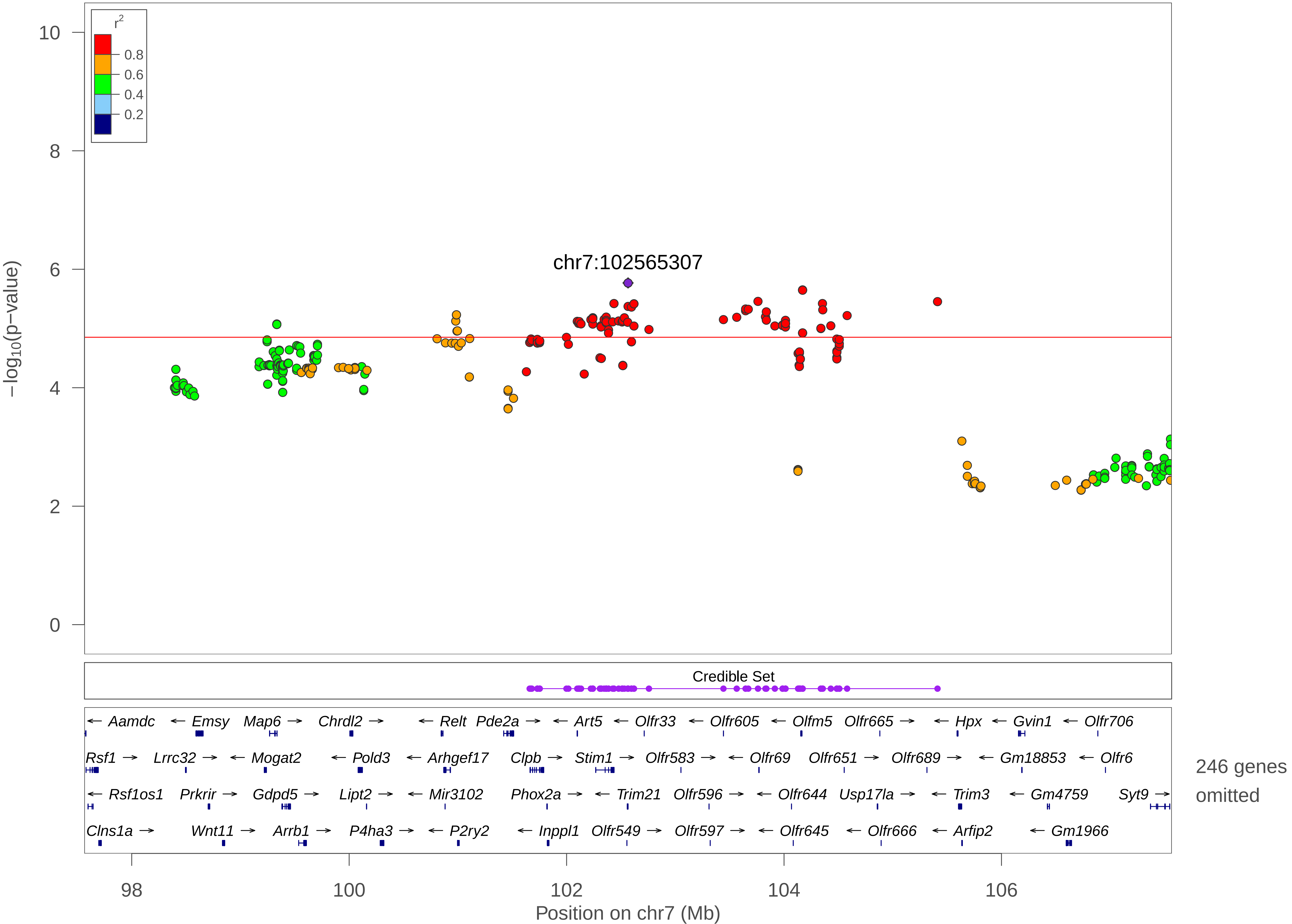

date: Thu Oct 25 17:09:52 2018

build: F34GBS

display range: chr7:97565307−107565307 [97565307−107565307]

hilite range: 0 − 0 [ 0 − 0 ]

reference SNP: chr7:102565307

number of SNPs plotted: 328

min P−value: 1.7E−6 [chr7:102565307]

max P−value: 5.37E−3 [chr7:106732278]

omitted Genes: Aqp11, Pak1, Gdpd4

omitted Genes: Myo7a, Capn5, Omp

omitted Genes: Gm16938, B3gnt6, Acer3

omitted Genes: Tsku, Gucy2d, Uvrag

omitted Genes: Dgat2, Gm26705, Serpinh1

omitted Genes: Klhl35, Rps3, Snord15b

omitted Genes: Snord15a, Mir326, Tpbgl

omitted Genes: Slco2b1, Olfr520, Olfr521

omitted Genes: Neu3, Spcs2, Xrra1

omitted Genes: Rnf169, Kcne3, Pgm2l1

omitted Genes: Gpx2−ps1, Ppme1, C2cd3

omitted Genes: Ucp3, Ucp2, Dnajb13

omitted Genes: Coa4, Mrpl48, Rab6a

omitted Genes: Plekhb1, Fam168a, P2ry6

omitted Genes: Fchsd2, Atg16l2, Stard10

omitted Genes: Arap1, Mir139, Art2a−ps

omitted Genes: Art2b, Folr2, Folr1

omitted Genes: Anapc15, Tomt, Lamtor1

omitted Genes: Lrrc51, Numa1, Il18bp

omitted Genes: Rnf121, Xndc1, Xntrpc

omitted Genes: Trpc2, Art1, Chrna10

omitted Genes: Nup98, Pgap2, Rhog

omitted Genes: Rrm1, Olfr543, Olfr544

omitted Genes: Olfr545, Olfr547, Olfr550

omitted Genes: Olfr551, Olfr552, Olfr553

omitted Genes: Olfr554, Olfr555, Olfr556

omitted Genes: Trim68, Olfr557, Olfr558

omitted Genes: Olfr559, Olfr78, Olfr560

omitted Genes: Olfr561, Olfr564, Olfr566

omitted Genes: Olfr568, Olfr569, Olfr570

omitted Genes: Olfr571, Olfr572, Olfr574

omitted Genes: Olfr575, Olfr576, Olfr577

Make more plots at http://csg.sph.umich.edu/locuszoom/

Fine−mapping Regions

| snp | chr | pos (Mb) | pp | group | color |
|---|---|---|---|---|---|
| chr7:101660383 | 7 | 101.6604 | 0.003627064 | Credible Set | purple |
| chr7:101669186 | 7 | 101.6692 | 0.003798908 | Credible Set | purple |
| chr7:101673483 | 7 | 101.6735 | 0.004132179 | Credible Set | purple |
| chr7:101680140 | 7 | 101.6801 | 0.003933425 | Credible Set | purple |
| chr7:101731061 | 7 | 101.7311 | 0.003544125 | Credible Set | purple |
| chr7:101731110 | 7 | 101.7311 | 0.003750966 | Credible Set | purple |
| chr7:101731130 | 7 | 101.7311 | 0.004069510 | Credible Set | purple |
| chr7:101731658 | 7 | 101.7317 | 0.003561151 | Credible Set | purple |
| chr7:101751790 | 7 | 101.7518 | 0.003597891 | Credible Set | purple |
| chr7:101752580 | 7 | 101.7526 | 0.003753693 | Credible Set | purple |
| chr7:101752971 | 7 | 101.7530 | 0.003846105 | Credible Set | purple |
| chr7:101998679 | 7 | 101.9987 | 0.004387980 | Credible Set | purple |
| chr7:102016307 | 7 | 102.0163 | 0.003382431 | Credible Set | purple |
| chr7:102098054 | 7 | 102.0981 | 0.007941011 | Credible Set | purple |
| chr7:102110109 | 7 | 102.1101 | 0.007486550 | Credible Set | purple |
| chr7:102110501 | 7 | 102.1105 | 0.007364089 | Credible Set | purple |
| chr7:102116318 | 7 | 102.1163 | 0.007691085 | Credible Set | purple |
| chr7:102116430 | 7 | 102.1164 | 0.007691085 | Credible Set | purple |
| chr7:102116478 | 7 | 102.1165 | 0.007875812 | Credible Set | purple |
| chr7:102132562 | 7 | 102.1326 | 0.007226850 | Credible Set | purple |
| chr7:102223396 | 7 | 102.2234 | 0.008459887 | Credible Set | purple |
| chr7:102240695 | 7 | 102.2407 | 0.007171970 | Credible Set | purple |
| chr7:102241191 | 7 | 102.2412 | 0.009094255 | Credible Set | purple |
| chr7:102241211 | 7 | 102.2412 | 0.008875089 | Credible Set | purple |
| chr7:102241212 | 7 | 102.2412 | 0.008744206 | Credible Set | purple |
| chr7:102306968 | 7 | 102.3070 | 0.002045277 | Credible Set | purple |
| chr7:102306978 | 7 | 102.3070 | 0.002058335 | Credible Set | purple |
| chr7:102317213 | 7 | 102.3172 | 0.006754915 | Credible Set | purple |
| chr7:102317246 | 7 | 102.3172 | 0.006440559 | Credible Set | purple |
| chr7:102319586 | 7 | 102.3196 | 0.002006137 | Credible Set | purple |
| chr7:102344970 | 7 | 102.3450 | 0.008527618 | Credible Set | purple |
| chr7:102363897 | 7 | 102.3639 | 0.007541180 | Credible Set | purple |
| chr7:102364204 | 7 | 102.3642 | 0.008696024 | Credible Set | purple |
| chr7:102364223 | 7 | 102.3642 | 0.009309070 | Credible Set | purple |
| chr7:102364315 | 7 | 102.3643 | 0.008205520 | Credible Set | purple |
| chr7:102364326 | 7 | 102.3643 | 0.008205520 | Credible Set | purple |

Fine−mapping Regions

| snp | chr | pos (Mb) | pp | group | color |
|---|---|---|---|---|---|
| chr7:102364411 | 7 | 102.3644 | 0.007791140 | Credible Set | purple |
| chr7:102385520 | 7 | 102.3855 | 0.005772189 | Credible Set | purple |
| chr7:102385536 | 7 | 102.3855 | 0.005140265 | Credible Set | purple |
| chr7:102423165 | 7 | 102.4232 | 0.007792384 | Credible Set | purple |
| chr7:102435692 | 7 | 102.4357 | 0.015407884 | Credible Set | purple |
| chr7:102435720 | 7 | 102.4357 | 0.015407884 | Credible Set | purple |
| chr7:102478469 | 7 | 102.4785 | 0.008058391 | Credible Set | purple |
| chr7:102510475 | 7 | 102.5105 | 0.007789352 | Credible Set | purple |
| chr7:102510607 | 7 | 102.5106 | 0.007777162 | Credible Set | purple |
| chr7:102516878 | 7 | 102.5169 | 0.008097979 | Credible Set | purple |
| chr7:102531652 | 7 | 102.5317 | 0.009075918 | Credible Set | purple |
| chr7:102561708 | 7 | 102.5617 | 0.007676879 | Credible Set | purple |
| chr7:102565262 | 7 | 102.5653 | 0.033181471 | Credible Set | purple |
| chr7:102565292 | 7 | 102.5653 | 0.033077596 | Credible Set | purple |
| chr7:102565307 | 7 | 102.5653 | 0.033239383 | Credible Set | purple |
| chr7:102565323 | 7 | 102.5653 | 0.033135343 | Credible Set | purple |
| chr7:102565329 | 7 | 102.5653 | 0.033135343 | Credible Set | purple |
| chr7:102565410 | 7 | 102.5654 | 0.013851517 | Credible Set | purple |
| chr7:102596442 | 7 | 102.5964 | 0.013508339 | Credible Set | purple |
| chr7:102596578 | 7 | 102.5966 | 0.003709545 | Credible Set | purple |
| chr7:102596636 | 7 | 102.5966 | 0.003731657 | Credible Set | purple |
| chr7:102619418 | 7 | 102.6194 | 0.015282756 | Credible Set | purple |
| chr7:102619431 | 7 | 102.6194 | 0.015245355 | Credible Set | purple |
| chr7:102619466 | 7 | 102.6195 | 0.006671488 | Credible Set | purple |
| chr7:102757499 | 7 | 102.7575 | 0.005869542 | Credible Set | purple |
| chr7:103442570 | 7 | 103.4426 | 0.008474361 | Credible Set | purple |
| chr7:103564836 | 7 | 103.5648 | 0.009243209 | Credible Set | purple |
| chr7:103646439 | 7 | 103.6464 | 0.011785582 | Credible Set | purple |
| chr7:103646709 | 7 | 103.6467 | 0.012579214 | Credible Set | purple |
| chr7:103671033 | 7 | 103.6710 | 0.012439283 | Credible Set | purple |
| chr7:103760301 | 7 | 103.7603 | 0.016704753 | Credible Set | purple |
| chr7:103829200 | 7 | 103.8292 | 0.009393752 | Credible Set | purple |
| chr7:103837099 | 7 | 103.8371 | 0.011314582 | Credible Set | purple |
| chr7:103837213 | 7 | 103.8372 | 0.008296113 | Credible Set | purple |
| chr7:103915503 | 7 | 103.9155 | 0.006691495 | Credible Set | purple |
| chr7:103986792 | 7 | 103.9868 | 0.006921892 | Credible Set | purple |

Fine−mapping Regions

| snp | chr | pos (Mb) | pp | group | color |
|---|---|---|---|---|---|
| chr7:103986795 | 7 | 103.9868 | 0.006921892 | Credible Set | purple |
| chr7:103986857 | 7 | 103.9869 | 0.006799078 | Credible Set | purple |
| chr7:103986860 | 7 | 103.9869 | 0.006799078 | Credible Set | purple |
| chr7:104012808 | 7 | 104.0128 | 0.006426842 | Credible Set | purple |
| chr7:104013370 | 7 | 104.0134 | 0.007306969 | Credible Set | purple |
| chr7:104013395 | 7 | 104.0134 | 0.008257676 | Credible Set | purple |
| chr7:104013467 | 7 | 104.0135 | 0.007341014 | Credible Set | purple |
| chr7:104129734 | 7 | 104.1297 | 0.002420744 | Credible Set | purple |
| chr7:104129736 | 7 | 104.1297 | 0.002420744 | Credible Set | purple |
| chr7:104138934 | 7 | 104.1389 | 0.001563108 | Credible Set | purple |
| chr7:104138936 | 7 | 104.1389 | 0.001569981 | Credible Set | purple |
| chr7:104138968 | 7 | 104.1390 | 0.001569981 | Credible Set | purple |
| chr7:104141705 | 7 | 104.1417 | 0.002456362 | Credible Set | purple |
| chr7:104141768 | 7 | 104.1418 | 0.002557425 | Credible Set | purple |
| chr7:104149893 | 7 | 104.1499 | 0.001981653 | Credible Set | purple |
| chr7:104149911 | 7 | 104.1499 | 0.001956598 | Credible Set | purple |
| chr7:104170479 | 7 | 104.1705 | 0.025445446 | Credible Set | purple |
| chr7:104170506 | 7 | 104.1705 | 0.025541514 | Credible Set | purple |
| chr7:104170518 | 7 | 104.1705 | 0.025630712 | Credible Set | purple |
| chr7:104170522 | 7 | 104.1705 | 0.025495253 | Credible Set | purple |
| chr7:104170543 | 7 | 104.1705 | 0.025338308 | Credible Set | purple |
| chr7:104170588 | 7 | 104.1706 | 0.005141288 | Credible Set | purple |
| chr7:104170594 | 7 | 104.1706 | 0.005149984 | Credible Set | purple |
| chr7:104170618 | 7 | 104.1706 | 0.005173026 | Credible Set | purple |
| chr7:104170662 | 7 | 104.1707 | 0.005173026 | Credible Set | purple |
| chr7:104338784 | 7 | 104.3388 | 0.006073698 | Credible Set | purple |
| chr7:104338810 | 7 | 104.3388 | 0.006144848 | Credible Set | purple |
| chr7:104353491 | 7 | 104.3535 | 0.015423013 | Credible Set | purple |
| chr7:104356230 | 7 | 104.3562 | 0.012184128 | Credible Set | purple |
| chr7:104430205 | 7 | 104.4302 | 0.006738007 | Credible Set | purple |
| chr7:104485236 | 7 | 104.4852 | 0.004117339 | Credible Set | purple |
| chr7:104485721 | 7 | 104.4857 | 0.002084849 | Credible Set | purple |
| chr7:104485727 | 7 | 104.4857 | 0.001972437 | Credible Set | purple |
| chr7:104486022 | 7 | 104.4860 | 0.002643835 | Credible Set | purple |
| chr7:104486041 | 7 | 104.4860 | 0.002527621 | Credible Set | purple |
| chr7:104507724 | 7 | 104.5077 | 0.003161793 | Credible Set | purple |

Fine−mapping Regions

| snp | chr | pos (Mb) | pp | group | color |
|---|---|---|---|---|---|
| chr7:104508127 | 7 | 104.5081 | 0.003467483 | Credible Set | purple |
| chr7:104508173 | 7 | 104.5082 | 0.004039979 | Credible Set | purple |
| chr7:104580091 | 7 | 104.5801 | 0.009856371 | Credible Set | purple |
| chr7:105411752 | 7 | 105.4118 | 0.016563255 | Credible Set | purple |

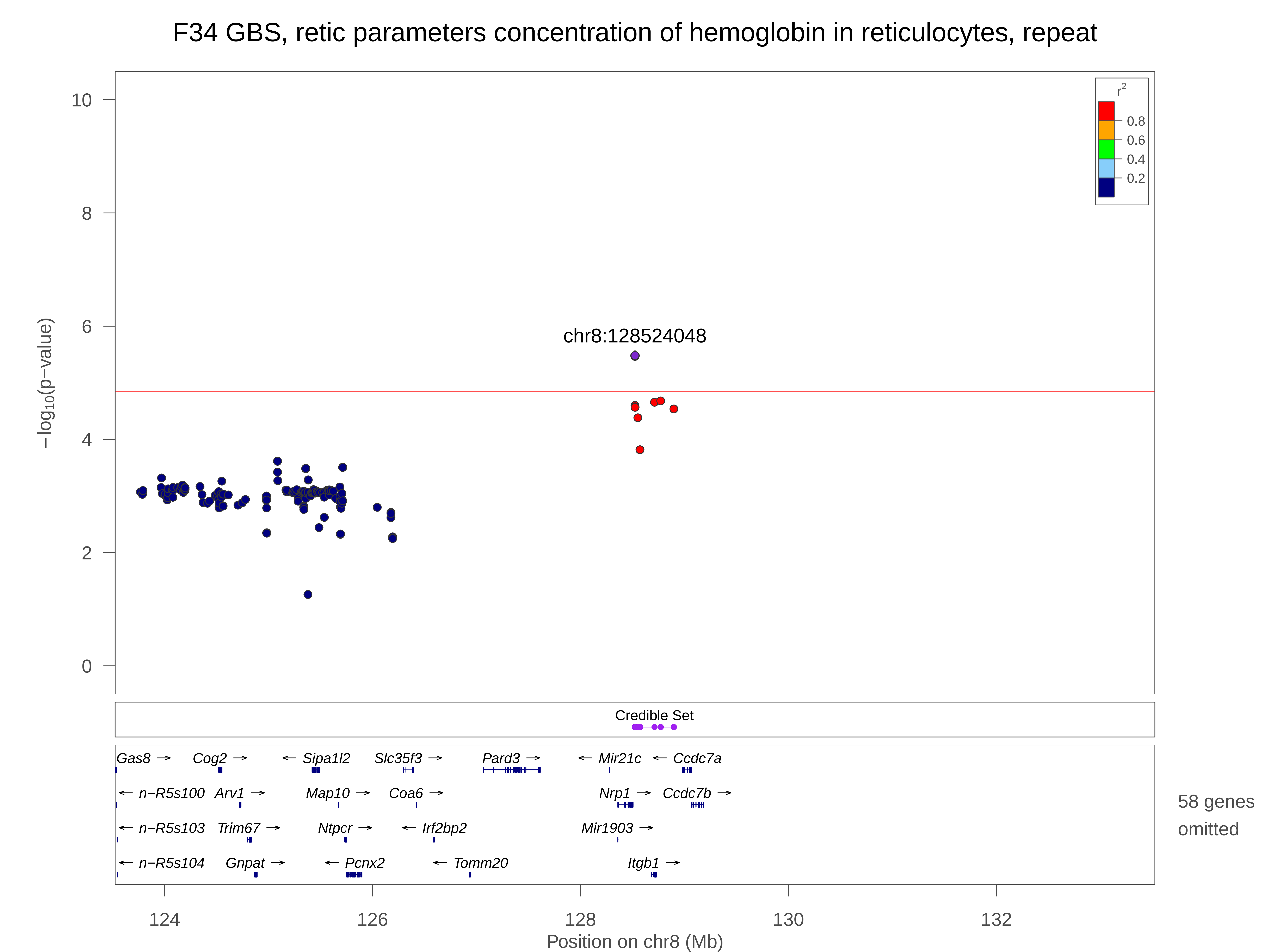

F34 GBS, retic parameters concentration of hemoglobin in reticulocytes, repeat
−log10(p−value)
chr8:128524048
r2
0.8
0.6
0.4
0.2
Credible Set
Gas8 →
Cog2 →
← Sipa1l2
Slc35f3 →
Pard3 →
← Mir21c
← Ccdc7a
← n−R5s100
Arv1 →
Map10 →
Coa6 →
Nrp1 →
Ccdc7b →
← n−R5s103
Trim67 →
Ntpcr →
← Irf2bp2
Mir1903 →
← n−R5s104
Gnpat →
← Pcnx2
← Tomm20
Itgb1 →
58 genes omitted
Position on chr8 (Mb)

| | |
|---|---|
| date: | Thu Oct 25 17:10:03 2018 |
| build: | F34GBS |
| display range: | chr8:123524048−133524048 [123524048−133524048] |
| hilite range: | 0 − 0 [ 0 − 0 ] |
| reference SNP: | chr8:128524048 |
| number of SNPs plotted: | 191 |
| min P−value: | 3.3E−6 [chr8:128524048] |
| max P−value: | 5.49E−2 [chr8:125378859] |
| omitted Genes: | n−R5s105, n−R5s108, n−R5s110 |
| omitted Genes: | n−R5s111, n−R5s113, Gm25018 |
| omitted Genes: | n−R5s117, n−R5s121, n−R5s122 |
| omitted Genes: | n−R5s123, n−R5s124, n−R5s128 |
| omitted Genes: | Gm26391, n−R5s133, n−R5s134 |
| omitted Genes: | Rn5s, n−R5s136, n−R5s138 |
| omitted Genes: | n−R5s139, Gm22109, n−R5s141 |
| omitted Genes: | n−R5s142, n−R5s143, n−R5s144 |
| omitted Genes: | Gm25212, n−R5s146, Gm22291 |
| omitted Genes: | Gm23284, n−R5s149, Rhou |
| omitted Genes: | 6030466F02Rik, Rab4a, Ccsap |
| omitted Genes: | Acta1, Nup133, Abcb10 |
| omitted Genes: | Taf5l, Mir1967, Urb2 |
| omitted Genes: | Galnt2, Pgbd5, Agt |
| omitted Genes: | Capn9, 2310022B05Rik, Ttc13 |
| omitted Genes: | Fam89a, 2810004N23Rik, Exoc8 |
| omitted Genes: | Sprtn, Egln1, Tsnax |
| omitted Genes: | Disc1, 4930567H12Rik, BC021891 |
| omitted Genes: | Kcnk1, Gm17296, Rbm34 |
| omitted Genes: | 2610044O15Rik8 |

Make more plots at http://csg.sph.umich.edu/locuszoom/

Fine−mapping Regions

| snp | chr | pos (Mb) | pp | group | color |
|---|---|---|---|---|---|
| chr8:128523972 | 8 | 128.5240 | 0.034880237 | Credible Set | purple |
| chr8:128524047 | 8 | 128.5240 | 0.239175550 | Credible Set | purple |
| chr8:128524048 | 8 | 128.5240 | 0.241911261 | Credible Set | purple |
| chr8:128524081 | 8 | 128.5241 | 0.234379335 | Credible Set | purple |
| chr8:128524424 | 8 | 128.5244 | 0.032999835 | Credible Set | purple |
| chr8:128524425 | 8 | 128.5244 | 0.032459243 | Credible Set | purple |
| chr8:128524441 | 8 | 128.5244 | 0.032395507 | Credible Set | purple |
| chr8:128551483 | 8 | 128.5515 | 0.021594105 | Credible Set | purple |
| chr8:128571869 | 8 | 128.5719 | 0.006315692 | Credible Set | purple |
| chr8:128571887 | 8 | 128.5719 | 0.006317955 | Credible Set | purple |
| chr8:128711717 | 8 | 128.7117 | 0.039387863 | Credible Set | purple |
| chr8:128771719 | 8 | 128.7717 | 0.041469323 | Credible Set | purple |
| chr8:128898075 | 8 | 128.8981 | 0.030398403 | Credible Set | purple |

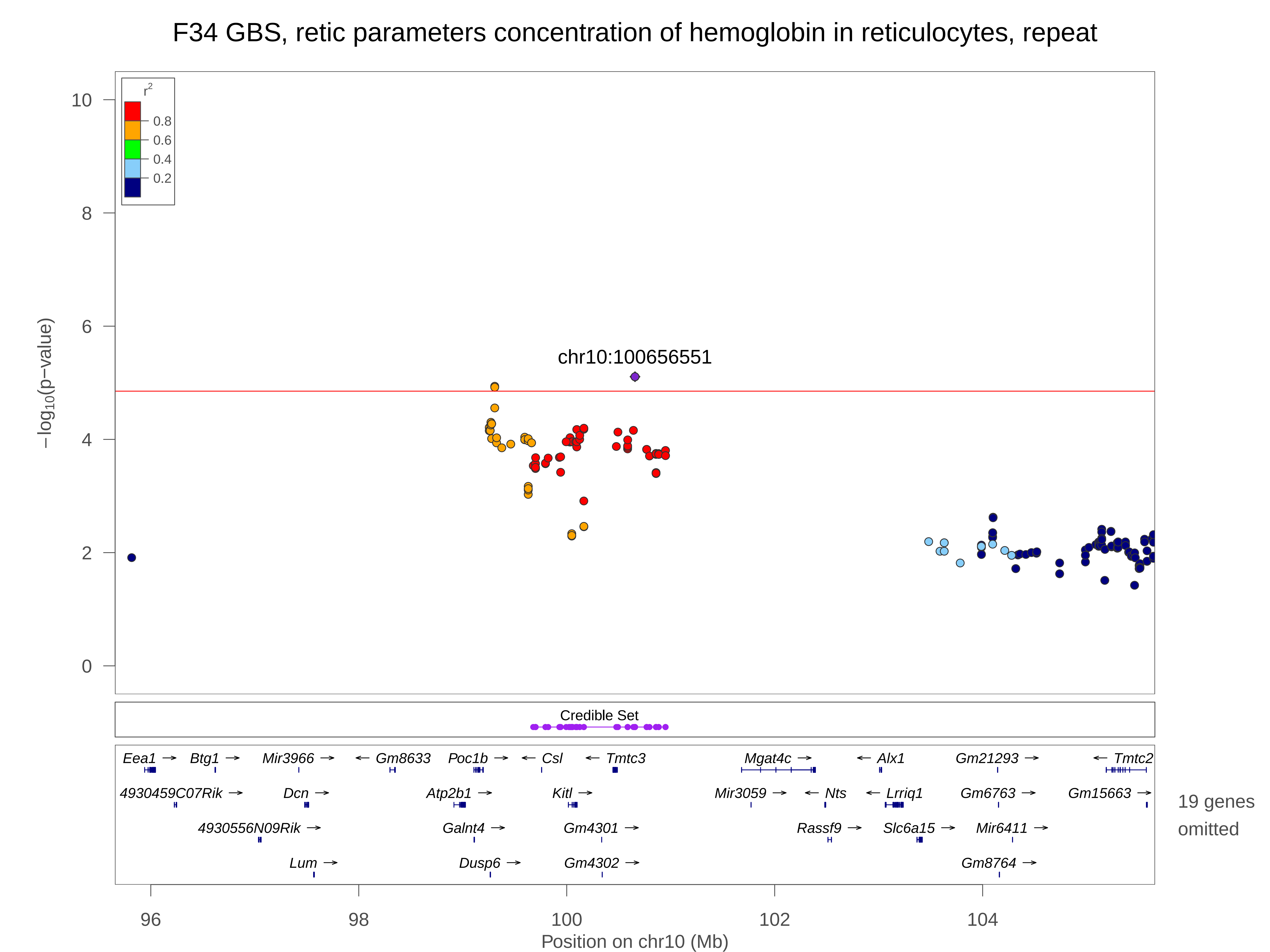

F34 GBS, retic parameters concentration of hemoglobin in reticulocytes, repeat
r2
0.8
0.6
0.4
0.2
10
8
6
4
2
0
−log10(p−value)
chr10:100656551
Credible Set
Eea1
Btg1
Mir3966
Gm8633
Poc1b
Csl
Tmtc3
Mgat4c
Alx1
Gm21293
Tmtc2
4930459C07Rik
Dcn
Atp2b1
Kitl
Mir3059
Nts
Lrriq1
Gm6763
Gm15663
4930556N09Rik
Galnt4
Gm4301
Rassf9
Slc6a15
Mir6411
Lum
Dusp6
Gm4302
Gm8764
19 genes omitted
96
98
100
102
104
Position on chr10 (Mb)

| | |
|---|---|
| date: | Thu Oct 25 17:09:37 2018 |
| build: | F34GBS |
| display range: | chr10:95656551−105656551 [95656551−105656551] |
| hilite range: | 0 − 0 [ 0 − 0 ] |
| reference SNP: | chr10:100656551 |
| number of SNPs plotted: | 225 |
| min P−value: | 7.81E−6 [chr10:100656551] |
| max P−value: | 3.77E−2 [chr10:105460999] |
| omitted Genes: | 4930473O22Rik, Kera, Epyc |
| omitted Genes: | Gm10754, Ccer1, 4930525C09Rik |
| omitted Genes: | Gm20110, Gm18409, Gm4303 |
| omitted Genes: | Gm4305, Gm4307, Gm4312 |
| omitted Genes: | Cep290, 4930430F08Rik, 1700017N19Rik |
| omitted Genes: | Gm21304, Gm21312, Gm20765 |
| omitted Genes: | Gm4340 |

Make more plots at http://csg.sph.umich.edu/locuszoom/

Fine−mapping Regions

| snp | chr | pos (Mb) | pp | group | color |
|---|---|---|---|---|---|
| chr10:99678204 | 10 | 99.67820 | 0.004298973 | Credible Set | purple |
| chr10:99699271 | 10 | 99.69927 | 0.004660097 | Credible Set | purple |
| chr10:99699649 | 10 | 99.69965 | 0.003854211 | Credible Set | purple |
| chr10:99699681 | 10 | 99.69968 | 0.004015350 | Credible Set | purple |
| chr10:99700646 | 10 | 99.70065 | 0.005837839 | Credible Set | purple |
| chr10:99795502 | 10 | 99.79550 | 0.004627611 | Credible Set | purple |
| chr10:99795653 | 10 | 99.79565 | 0.004743422 | Credible Set | purple |
| chr10:99821621 | 10 | 99.82162 | 0.005760589 | Credible Set | purple |
| chr10:99929872 | 10 | 99.92987 | 0.005890544 | Credible Set | purple |
| chr10:99930010 | 10 | 99.93001 | 0.005890544 | Credible Set | purple |
| chr10:99941189 | 10 | 99.94119 | 0.003340836 | Credible Set | purple |
| chr10:99941475 | 10 | 99.94147 | 0.006026726 | Credible Set | purple |
| chr10:99994442 | 10 | 99.99444 | 0.010730144 | Credible Set | purple |
| chr10:100020266 | 10 | 100.02027 | 0.010730144 | Credible Set | purple |
| chr10:100031418 | 10 | 100.03142 | 0.010638926 | Credible Set | purple |
| chr10:100031691 | 10 | 100.03169 | 0.012572651 | Credible Set | purple |
| chr10:100048262 | 10 | 100.04826 | 0.010730144 | Credible Set | purple |
| chr10:100048350 | 10 | 100.04835 | 0.010730144 | Credible Set | purple |
| chr10:100048362 | 10 | 100.04836 | 0.010730144 | Credible Set | purple |
| chr10:100055412 | 10 | 100.05541 | 0.010730144 | Credible Set | purple |
| chr10:100055418 | 10 | 100.05542 | 0.010730144 | Credible Set | purple |
| chr10:100086663 | 10 | 100.08666 | 0.010730144 | Credible Set | purple |
| chr10:100096296 | 10 | 100.09630 | 0.008784845 | Credible Set | purple |
| chr10:100096708 | 10 | 100.09671 | 0.010730144 | Credible Set | purple |
| chr10:100096785 | 10 | 100.09678 | 0.017141778 | Credible Set | purple |
| chr10:100096807 | 10 | 100.09681 | 0.017141778 | Credible Set | purple |
| chr10:100124479 | 10 | 100.12448 | 0.011740000 | Credible Set | purple |
| chr10:100125049 | 10 | 100.12505 | 0.013687694 | Credible Set | purple |
| chr10:100164149 | 10 | 100.16415 | 0.017387467 | Credible Set | purple |
| chr10:100164180 | 10 | 100.16418 | 0.017387467 | Credible Set | purple |
| chr10:100164914 | 10 | 100.16491 | 0.018170173 | Credible Set | purple |
| chr10:100477167 | 10 | 100.47717 | 0.008971417 | Credible Set | purple |
| chr10:100492319 | 10 | 100.49232 | 0.015603728 | Credible Set | purple |
| chr10:100585564 | 10 | 100.58556 | 0.008164326 | Credible Set | purple |
| chr10:100585784 | 10 | 100.58578 | 0.008664857 | Credible Set | purple |
| chr10:100585854 | 10 | 100.58585 | 0.009166375 | Credible Set | purple |

Fine−mapping Regions

| snp | chr | pos (Mb) | pp | group | color |
|---|---|---|---|---|---|
| chr10:100586507 | 10 | 100.5865 | 0.011522590 | Credible Set | purple |
| chr10:100641295 | 10 | 100.6413 | 0.016671266 | Credible Set | purple |
| chr10:100656540 | 10 | 100.6565 | 0.129487316 | Credible Set | purple |
| chr10:100656551 | 10 | 100.6566 | 0.129487316 | Credible Set | purple |
| chr10:100656588 | 10 | 100.6566 | 0.129487316 | Credible Set | purple |
| chr10:100656608 | 10 | 100.6566 | 0.129487316 | Credible Set | purple |
| chr10:100768688 | 10 | 100.7687 | 0.008010499 | Credible Set | purple |
| chr10:100768699 | 10 | 100.7687 | 0.008010499 | Credible Set | purple |
| chr10:100768710 | 10 | 100.7687 | 0.008010499 | Credible Set | purple |
| chr10:100768727 | 10 | 100.7687 | 0.008010499 | Credible Set | purple |
| chr10:100795827 | 10 | 100.7958 | 0.006228734 | Credible Set | purple |
| chr10:100858643 | 10 | 100.8586 | 0.006609795 | Credible Set | purple |
| chr10:100858698 | 10 | 100.8587 | 0.006855285 | Credible Set | purple |
| chr10:100858701 | 10 | 100.8587 | 0.006783238 | Credible Set | purple |
| chr10:100858992 | 10 | 100.8590 | 0.003308385 | Credible Set | purple |
| chr10:100884062 | 10 | 100.8841 | 0.006819065 | Credible Set | purple |
| chr10:100884636 | 10 | 100.8846 | 0.006609795 | Credible Set | purple |
| chr10:100949966 | 10 | 100.9500 | 0.007673860 | Credible Set | purple |
| chr10:100950077 | 10 | 100.9501 | 0.006319673 | Credible Set | purple |
| chr10:100950118 | 10 | 100.9501 | 0.006319673 | Credible Set | purple |

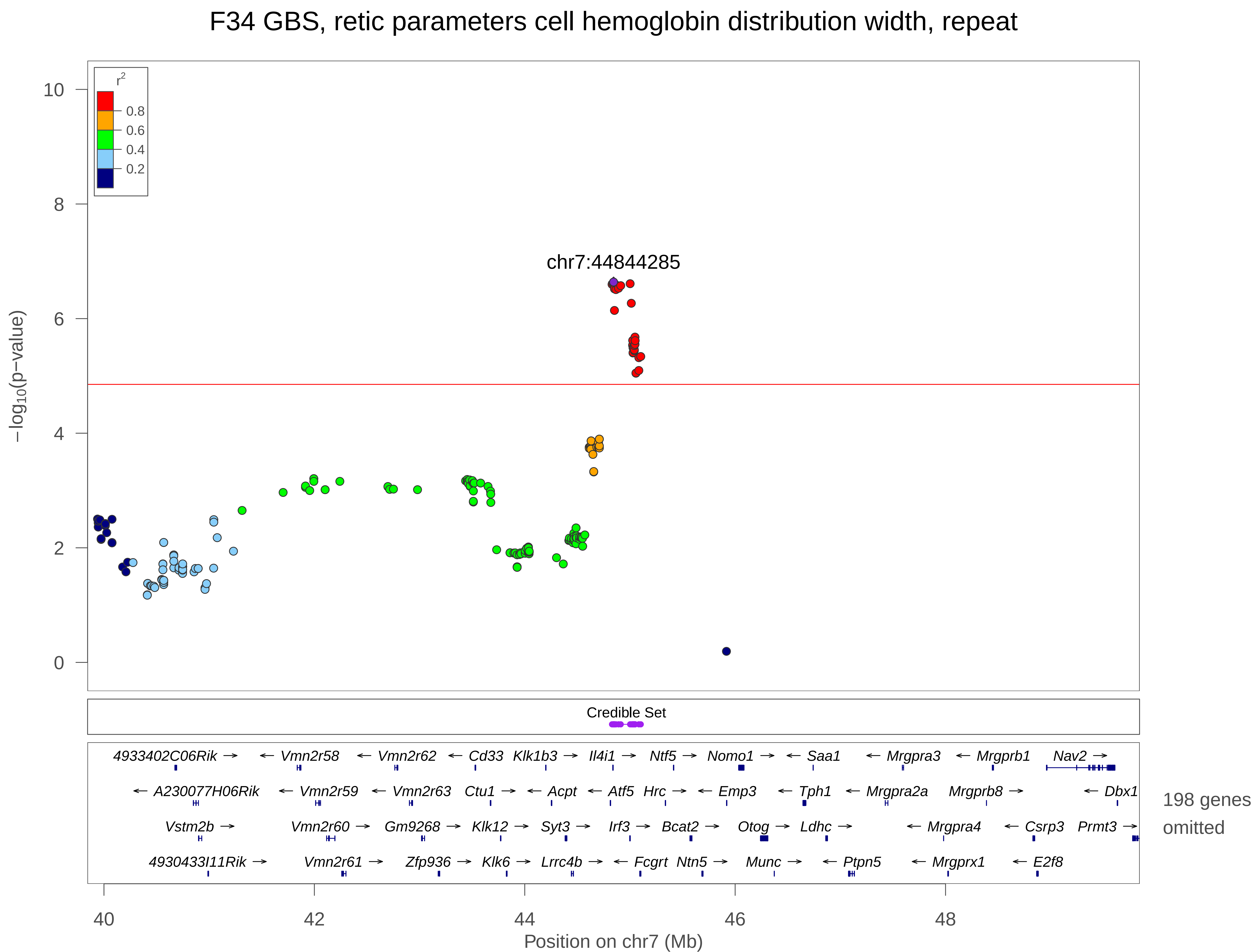
F34 GBS, retic parameters cell hemoglobin distribution width, repeat
r2
0.8
0.6
0.4
0.2
10
8
6
4
2
0
−log10(p−value)
chr7:44844285
Credible Set
4933402C06Rik →
← Vmn2r58
← Vmn2r62
← Cd33
Klk1b3 →
Il4i1 →
Ntf5 →
Nomo1 →
← Saa1
← Mrgpra3
← Mrgprb1
Nav2 →
← A230077H06Rik
← Vmn2r59
← Vmn2r63
Ctu1 →
← Acpt
← Atf5
Hrc →
← Emp3
← Tph1
← Mrgpra2a
Mrgprb8 →
← Dbx1
Vstm2b →
Vmn2r60 →
Gm9268 →
Klk12 →
Syt3 →
Irf3 →
Bcat2 →
Otog →
Ldhc →
← Mrgpra4
← Csrp3
Prmt3 →
4930433I11Rik →
Vmn2r61 →
Zfp936 →
Klk6 →
Lrrc4b →
← Fcgrt
Ntn5 →
Munc →
← Ptpn5
← Mrgprx1
← E2f8
198 genes omitted
40
42
44
46
48
Position on chr7 (Mb)

date: Thu Oct 25 17:10:30 2018
build: F34GBS
display range: chr7:39844285−49844285 [39844285−49844285]
hilite range: 0 − 0 [ 0 − 0 ]
reference SNP: chr7:44844285
number of SNPs plotted: 247
min P−value: 2.3E−7 [chr7:44844285]
max P−value: 6.41E−1 [chr7:45917340]
omitted Genes: Gm4884, 1700110I07Rik, Gm5592
omitted Genes: AI987944, Vmn2r57, AW146154
omitted Genes: Zfp788, Vmn2r−ps54, Vmn2r−ps60
omitted Genes: 4933421I07Rik, Zfp141, 9830147E19Rik
omitted Genes: Gm5595, Gm2381, EU599041
omitted Genes: Zfp715, Siglecf, Siglecg
omitted Genes: Lim2, Nkg7, Cldnd2
omitted Genes: Etfb, Mir7051, Vsig10l
omitted Genes: Iglon5, 4931406B18Rik, Zfp658
omitted Genes: Zfp719, Zfp819, Ceacam18
omitted Genes: Siglece, Klk14, Klk13
omitted Genes: 2310002F09Rik, Klk11, Klk10
omitted Genes: Klk9, Klk8, Klk7
omitted Genes: Klk5, Klk4, Klk15
omitted Genes: Klk1b7−ps, Klk1b8, Klk1b1
omitted Genes: Klk1b9, Klk1b11, Klk1b26
omitted Genes: Klk1b27, Klk1b21, Klk1b22
omitted Genes: Klk1b16, Klk1b24, Klk1b4
omitted Genes: Klk1b5, Klk1, 1700028J19Rik
omitted Genes: 2410002F23Rik, Snord88c, Snord88a
omitted Genes: Clec11a, Shank1, 1700008O03Rik
omitted Genes: Aspdh, Josd2, Mir7052
omitted Genes: 5430431A17Rik, Emc10, Fam71e1
omitted Genes: Mybpc2, Spib, Pold1
omitted Genes: Mir7053, Nr1h2, Napsa
omitted Genes: Kcnc3, Myh14, Izumo2
omitted Genes: Zfp473, Vrk3, Nup62
omitted Genes: Gm21948, Tbc1d17, Akt1s1
omitted Genes: Mir707, Pnkp, Ptov1
omitted Genes: Med25, Fuz, Ap2a1
omitted Genes: Tsks, Cpt1c, Prmt1
Make more plots at http://csg.sph.umich.edu/locuszoom/
omitted Genes: Gm15545, Bcl2l12, Scaf1

Fine−mapping Regions

| snp | chr | pos (Mb) | pp | group | color |
|---|---|---|---|---|---|
| chr7:44829933 | 7 | 44.82993 | 0.076255075 | Credible Set | purple |
| chr7:44839966 | 7 | 44.83997 | 0.077914501 | Credible Set | purple |
| chr7:44844016 | 7 | 44.84402 | 0.077689067 | Credible Set | purple |
| chr7:44844285 | 7 | 44.84428 | 0.083580561 | Credible Set | purple |
| chr7:44844322 | 7 | 44.84432 | 0.077689067 | Credible Set | purple |
| chr7:44851630 | 7 | 44.85163 | 0.070642061 | Credible Set | purple |
| chr7:44852639 | 7 | 44.85264 | 0.027791039 | Credible Set | purple |
| chr7:44852965 | 7 | 44.85296 | 0.064054774 | Credible Set | purple |
| chr7:44866590 | 7 | 44.86659 | 0.062306105 | Credible Set | purple |
| chr7:44891663 | 7 | 44.89166 | 0.065608028 | Credible Set | purple |
| chr7:44912209 | 7 | 44.91221 | 0.072611945 | Credible Set | purple |
| chr7:45001761 | 7 | 45.00176 | 0.078409708 | Credible Set | purple |
| chr7:45012901 | 7 | 45.01290 | 0.036685318 | Credible Set | purple |
| chr7:45025410 | 7 | 45.02541 | 0.007242901 | Credible Set | purple |
| chr7:45025443 | 7 | 45.02544 | 0.007244986 | Credible Set | purple |
| chr7:45026544 | 7 | 45.02654 | 0.008733339 | Credible Set | purple |
| chr7:45026548 | 7 | 45.02655 | 0.008700908 | Credible Set | purple |
| chr7:45029371 | 7 | 45.02937 | 0.006564970 | Credible Set | purple |
| chr7:45029402 | 7 | 45.02940 | 0.005385265 | Credible Set | purple |
| chr7:45029407 | 7 | 45.02941 | 0.005390352 | Credible Set | purple |
| chr7:45029413 | 7 | 45.02941 | 0.005391354 | Credible Set | purple |
| chr7:45029777 | 7 | 45.02978 | 0.005455005 | Credible Set | purple |
| chr7:45033529 | 7 | 45.03353 | 0.006564970 | Credible Set | purple |
| chr7:45041503 | 7 | 45.04150 | 0.005897525 | Credible Set | purple |
| chr7:45041522 | 7 | 45.04152 | 0.006080683 | Credible Set | purple |
| chr7:45041933 | 7 | 45.04193 | 0.007166194 | Credible Set | purple |
| chr7:45047834 | 7 | 45.04783 | 0.007495305 | Credible Set | purple |
| chr7:45047913 | 7 | 45.04791 | 0.009921599 | Credible Set | purple |
| chr7:45049496 | 7 | 45.04950 | 0.008655283 | Credible Set | purple |
| chr7:45084764 | 7 | 45.08476 | 0.004482703 | Credible Set | purple |
| chr7:45102598 | 7 | 45.10260 | 0.004690384 | Credible Set | purple |

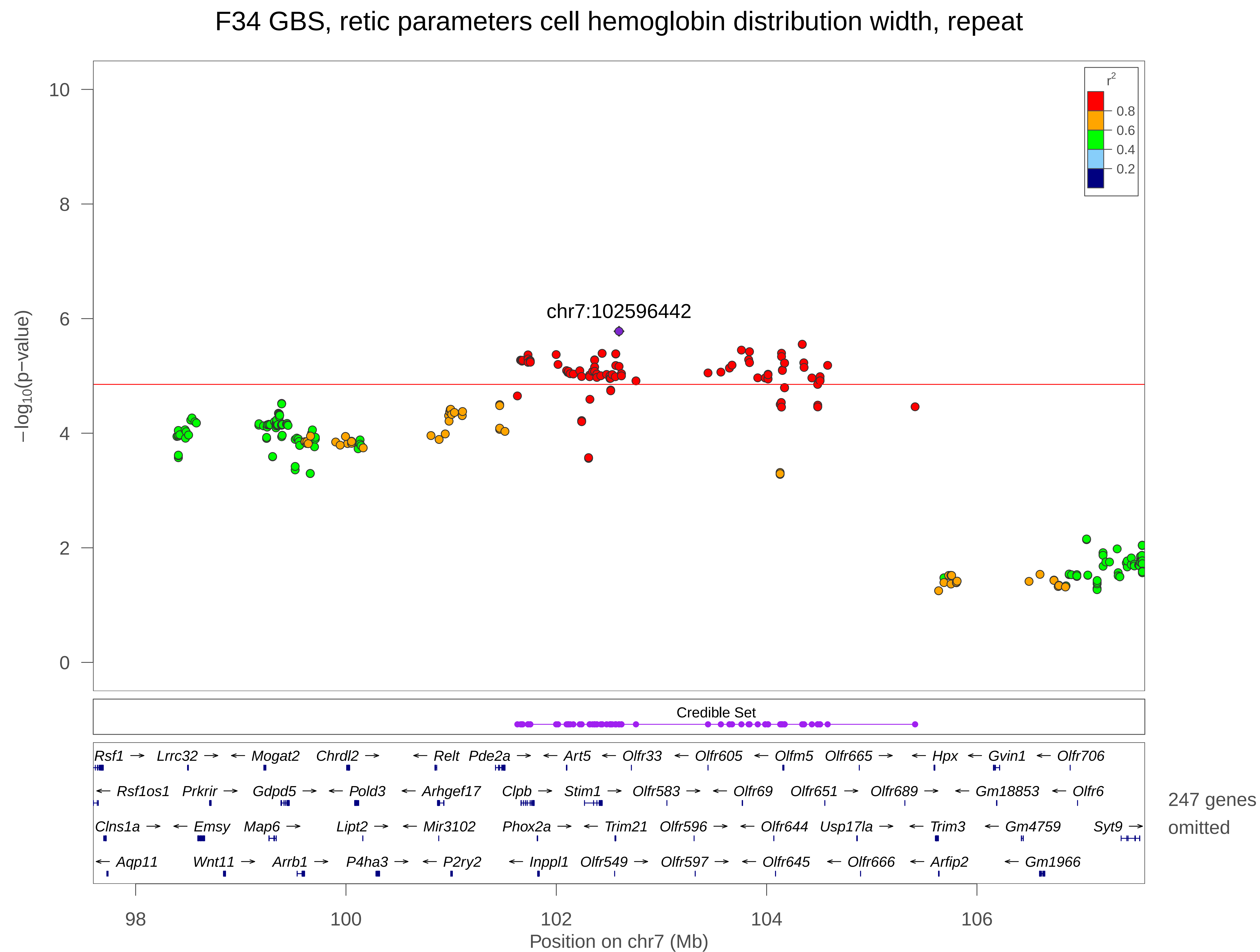
F34 GBS, retic parameters cell hemoglobin distribution width, repeat
−log10(p−value)
chr7:102596442
r2
0.8
0.6
0.4
0.2
Credible Set
Rsf1 → Lrrc32 → ← Mogat2 Chrdl2 → ← Relt Pde2a → ← Art5 ← Olfr33 ← Olfr605 ← Olfm5 Olfr665 → ← Hpx ← Gvin1 ← Olfr706
← Rsf1os1 Prkrir → Gdpd5 → ← Pold3 ← Arhgef17 Clpb → Stim1 → Olfr583 → ← Olfr69 Olfr651 → Olfr689 → ← Gm18853 ← Olfr6
Clns1a → ← Emsy Map6 → Lipt2 → ← Mir3102 Phox2a → ← Trim21 Olfr596 → ← Olfr644 Usp17la → ← Trim3 ← Gm4759 Syt9 →
← Aqp11 Wnt11 → Arrb1 → P4ha3 → ← P2ry2 ← Inppl1 Olfr549 → Olfr597 → ← Olfr645 ← Olfr666 ← Arfip2 ← Gm1966
247 genes omitted
Position on chr7 (Mb)

date: Thu Oct 25 17:10:16 2018
build: F34GBS
display range: chr7:97596442−107596442 [97596442−107596442]
hilite range: 0 − 0 [ 0 − 0 ]
reference SNP: chr7:102596442
number of SNPs plotted: 347
min P−value: 1.66E−6 [chr7:102596442]
max P−value: 5.62E−2 [chr7:105635493]
omitted Genes: Pak1, Gdpd4, Myo7a
omitted Genes: Capn5, Omp, Gm16938
omitted Genes: B3gnt6, Acer3, Tsku
omitted Genes: Gucy2d, Uvrag, Dgat2
omitted Genes: Gm26705, Serpinh1, Klhl35
omitted Genes: Rps3, Snord15b, Snord15a
omitted Genes: Mir326, Tpbgl, Slco2b1
omitted Genes: Olfr520, Olfr521, Neu3
omitted Genes: Spcs2, Xrra1, Rnf169
omitted Genes: Kcne3, Pgm2l1, Gpx2−ps1
omitted Genes: Ppme1, C2cd3, Ucp3
omitted Genes: Ucp2, Dnajb13, Coa4
omitted Genes: Mrpl48, Rab6a, Plekhb1
omitted Genes: Fam168a, P2ry6, Fchsd2
omitted Genes: Atg16l2, Stard10, Arap1
omitted Genes: Mir139, Art2a−ps, Art2b
omitted Genes: Folr2, Folr1, Anapc15
omitted Genes: Tomt, Lamtor1, Lrrc51
omitted Genes: Numa1, Il18bp, Rnf121
omitted Genes: Xndc1, Xntrpc, Trpc2
omitted Genes: Art1, Chrna10, Nup98
omitted Genes: Pgap2, Rhog, Rrm1
omitted Genes: Olfr543, Olfr544, Olfr545
omitted Genes: Olfr547, Olfr550, Olfr551
omitted Genes: Olfr552, Olfr553, Olfr554
omitted Genes: Olfr555, Olfr556, Trim68
omitted Genes: Olfr557, Olfr558, Olfr559
omitted Genes: Olfr78, Olfr560, Olfr561
omitted Genes: Olfr564, Olfr566, Olfr568
omitted Genes: Olfr569, Olfr570, Olfr571
omitted Genes: Olfr572, Olfr574, Olfr575
Make more plots at http://csg.sph.umich.edu/locuszoom/
omitted Genes: Olfr576, Olfr577, Olfr578

Fine−mapping Regions

| snp | chr | pos (Mb) | pp | group | color |
|---|---|---|---|---|---|
| chr7:101630176 | 7 | 101.6302 | 0.003096999 | Credible Set | purple |
| chr7:101660383 | 7 | 101.6604 | 0.012223216 | Credible Set | purple |
| chr7:101669186 | 7 | 101.6692 | 0.012120864 | Credible Set | purple |
| chr7:101673483 | 7 | 101.6735 | 0.011764260 | Credible Set | purple |
| chr7:101680140 | 7 | 101.6801 | 0.012162155 | Credible Set | purple |
| chr7:101731061 | 7 | 101.7311 | 0.013717504 | Credible Set | purple |
| chr7:101731110 | 7 | 101.7311 | 0.014983276 | Credible Set | purple |
| chr7:101731130 | 7 | 101.7311 | 0.012766398 | Credible Set | purple |
| chr7:101731658 | 7 | 101.7317 | 0.011235698 | Credible Set | purple |
| chr7:101751790 | 7 | 101.7518 | 0.012022773 | Credible Set | purple |
| chr7:101752580 | 7 | 101.7526 | 0.011764703 | Credible Set | purple |
| chr7:101752971 | 7 | 101.7530 | 0.011283295 | Credible Set | purple |
| chr7:101998679 | 7 | 101.9987 | 0.015148262 | Credible Set | purple |
| chr7:102016307 | 7 | 102.0163 | 0.010350693 | Credible Set | purple |
| chr7:102098054 | 7 | 102.0981 | 0.008168876 | Credible Set | purple |
| chr7:102110109 | 7 | 102.1101 | 0.007618489 | Credible Set | purple |
| chr7:102110501 | 7 | 102.1105 | 0.007761370 | Credible Set | purple |
| chr7:102116318 | 7 | 102.1163 | 0.007880212 | Credible Set | purple |
| chr7:102116430 | 7 | 102.1164 | 0.007880212 | Credible Set | purple |
| chr7:102116478 | 7 | 102.1165 | 0.007960892 | Credible Set | purple |
| chr7:102132562 | 7 | 102.1326 | 0.007255263 | Credible Set | purple |
| chr7:102162058 | 7 | 102.1621 | 0.007173808 | Credible Set | purple |
| chr7:102223396 | 7 | 102.2234 | 0.008138683 | Credible Set | purple |
| chr7:102240695 | 7 | 102.2407 | 0.006514444 | Credible Set | purple |
| chr7:102317213 | 7 | 102.3172 | 0.006918461 | Credible Set | purple |
| chr7:102317246 | 7 | 102.3172 | 0.006464993 | Credible Set | purple |
| chr7:102319586 | 7 | 102.3196 | 0.002721322 | Credible Set | purple |
| chr7:102344970 | 7 | 102.3450 | 0.007947491 | Credible Set | purple |
| chr7:102363897 | 7 | 102.3639 | 0.007849316 | Credible Set | purple |
| chr7:102364204 | 7 | 102.3642 | 0.012380383 | Credible Set | purple |
| chr7:102364223 | 7 | 102.3642 | 0.012234694 | Credible Set | purple |
| chr7:102364315 | 7 | 102.3643 | 0.009361449 | Credible Set | purple |
| chr7:102364326 | 7 | 102.3643 | 0.009361449 | Credible Set | purple |
| chr7:102364411 | 7 | 102.3644 | 0.007958604 | Credible Set | purple |
| chr7:102385520 | 7 | 102.3855 | 0.007151558 | Credible Set | purple |
| chr7:102385536 | 7 | 102.3855 | 0.006289323 | Credible Set | purple |

Fine−mapping Regions

| snp | chr | pos (Mb) | pp | group | color |
|---|---|---|---|---|---|
| chr7:102423165 | 7 | 102.4232 | 0.006711697 | Credible Set | purple |
| chr7:102435692 | 7 | 102.4357 | 0.015868556 | Credible Set | purple |
| chr7:102435720 | 7 | 102.4357 | 0.015868556 | Credible Set | purple |
| chr7:102478469 | 7 | 102.4785 | 0.007052608 | Credible Set | purple |
| chr7:102510475 | 7 | 102.5105 | 0.006648575 | Credible Set | purple |
| chr7:102510607 | 7 | 102.5106 | 0.006024644 | Credible Set | purple |
| chr7:102516878 | 7 | 102.5169 | 0.006090483 | Credible Set | purple |
| chr7:102517238 | 7 | 102.5172 | 0.003836833 | Credible Set | purple |
| chr7:102517241 | 7 | 102.5172 | 0.003878129 | Credible Set | purple |
| chr7:102517247 | 7 | 102.5172 | 0.003766622 | Credible Set | purple |
| chr7:102531652 | 7 | 102.5317 | 0.007025427 | Credible Set | purple |
| chr7:102561708 | 7 | 102.5617 | 0.006460720 | Credible Set | purple |
| chr7:102565262 | 7 | 102.5653 | 0.015471762 | Credible Set | purple |
| chr7:102565292 | 7 | 102.5653 | 0.015535507 | Credible Set | purple |
| chr7:102565307 | 7 | 102.5653 | 0.015520392 | Credible Set | purple |
| chr7:102565323 | 7 | 102.5653 | 0.015584350 | Credible Set | purple |
| chr7:102565329 | 7 | 102.5653 | 0.015584350 | Credible Set | purple |
| chr7:102565410 | 7 | 102.5654 | 0.009932812 | Credible Set | purple |
| chr7:102596442 | 7 | 102.5964 | 0.037183569 | Credible Set | purple |
| chr7:102596578 | 7 | 102.5966 | 0.009604558 | Credible Set | purple |
| chr7:102596636 | 7 | 102.5966 | 0.009599972 | Credible Set | purple |
| chr7:102619418 | 7 | 102.6194 | 0.007257376 | Credible Set | purple |
| chr7:102619431 | 7 | 102.6194 | 0.007256067 | Credible Set | purple |
| chr7:102619466 | 7 | 102.6195 | 0.006673185 | Credible Set | purple |
| chr7:102757499 | 7 | 102.7575 | 0.005518051 | Credible Set | purple |
| chr7:103442570 | 7 | 103.4426 | 0.007476220 | Credible Set | purple |
| chr7:103564836 | 7 | 103.5648 | 0.007716649 | Credible Set | purple |
| chr7:103646439 | 7 | 103.6464 | 0.009025333 | Credible Set | purple |
| chr7:103646709 | 7 | 103.6467 | 0.009004268 | Credible Set | purple |
| chr7:103671033 | 7 | 103.6710 | 0.010091982 | Credible Set | purple |
| chr7:103760301 | 7 | 103.7603 | 0.017991325 | Credible Set | purple |
| chr7:103829200 | 7 | 103.8292 | 0.012434995 | Credible Set | purple |
| chr7:103837099 | 7 | 103.8371 | 0.016944840 | Credible Set | purple |
| chr7:103837213 | 7 | 103.8372 | 0.011097493 | Credible Set | purple |
| chr7:103915503 | 7 | 103.9155 | 0.006196880 | Credible Set | purple |
| chr7:103986792 | 7 | 103.9868 | 0.006158547 | Credible Set | purple |

Fine−mapping Regions

| snp | chr | pos (Mb) | pp | group | color |
|---|---|---|---|---|---|
| chr7:103986795 | 7 | 103.9868 | 0.006158547 | Credible Set | purple |
| chr7:103986857 | 7 | 103.9869 | 0.006172090 | Credible Set | purple |
| chr7:103986860 | 7 | 103.9869 | 0.006172090 | Credible Set | purple |
| chr7:104012808 | 7 | 104.0128 | 0.006034451 | Credible Set | purple |
| chr7:104013370 | 7 | 104.0134 | 0.005918186 | Credible Set | purple |
| chr7:104013395 | 7 | 104.0134 | 0.007116710 | Credible Set | purple |
| chr7:104013467 | 7 | 104.0135 | 0.006974150 | Credible Set | purple |
| chr7:104129734 | 7 | 104.1297 | 0.002237892 | Credible Set | purple |
| chr7:104129736 | 7 | 104.1297 | 0.002237892 | Credible Set | purple |
| chr7:104138934 | 7 | 104.1389 | 0.002400300 | Credible Set | purple |
| chr7:104138936 | 7 | 104.1389 | 0.002408368 | Credible Set | purple |
| chr7:104138968 | 7 | 104.1390 | 0.002408368 | Credible Set | purple |
| chr7:104141705 | 7 | 104.1417 | 0.015962451 | Credible Set | purple |
| chr7:104141768 | 7 | 104.1418 | 0.014007024 | Credible Set | purple |
| chr7:104149893 | 7 | 104.1499 | 0.008394697 | Credible Set | purple |
| chr7:104149911 | 7 | 104.1499 | 0.008215429 | Credible Set | purple |
| chr7:104170479 | 7 | 104.1705 | 0.004239615 | Credible Set | purple |
| chr7:104170506 | 7 | 104.1705 | 0.004230559 | Credible Set | purple |
| chr7:104170518 | 7 | 104.1705 | 0.004268338 | Credible Set | purple |
| chr7:104170522 | 7 | 104.1705 | 0.004229257 | Credible Set | purple |
| chr7:104170543 | 7 | 104.1705 | 0.004227186 | Credible Set | purple |
| chr7:104170588 | 7 | 104.1706 | 0.010922302 | Credible Set | purple |
| chr7:104170594 | 7 | 104.1706 | 0.010925834 | Credible Set | purple |
| chr7:104170618 | 7 | 104.1706 | 0.010916495 | Credible Set | purple |
| chr7:104170662 | 7 | 104.1707 | 0.010916495 | Credible Set | purple |
| chr7:104338784 | 7 | 104.3388 | 0.022362521 | Credible Set | purple |
| chr7:104338810 | 7 | 104.3388 | 0.022579047 | Credible Set | purple |
| chr7:104353491 | 7 | 104.3535 | 0.011030919 | Credible Set | purple |
| chr7:104356230 | 7 | 104.3562 | 0.009230694 | Credible Set | purple |
| chr7:104430205 | 7 | 104.4302 | 0.006167265 | Credible Set | purple |
| chr7:104485236 | 7 | 104.4852 | 0.004809966 | Credible Set | purple |
| chr7:104485721 | 7 | 104.4857 | 0.002085735 | Credible Set | purple |
| chr7:104485727 | 7 | 104.4857 | 0.002038165 | Credible Set | purple |
| chr7:104486022 | 7 | 104.4860 | 0.002178204 | Credible Set | purple |
| chr7:104507724 | 7 | 104.5077 | 0.005871653 | Credible Set | purple |
| chr7:104508127 | 7 | 104.5081 | 0.006481453 | Credible Set | purple |

Fine−mapping Regions

| snp | chr | pos (Mb) | pp | group | color |
|---|---|---|---|---|---|
| chr7:104508173 | 7 | 104.5082 | 0.005525199 | Credible Set | purple |
| chr7:104580091 | 7 | 104.5801 | 0.010002260 | Credible Set | purple |
| chr7:105411752 | 7 | 105.4118 | 0.002046595 | Credible Set | purple |

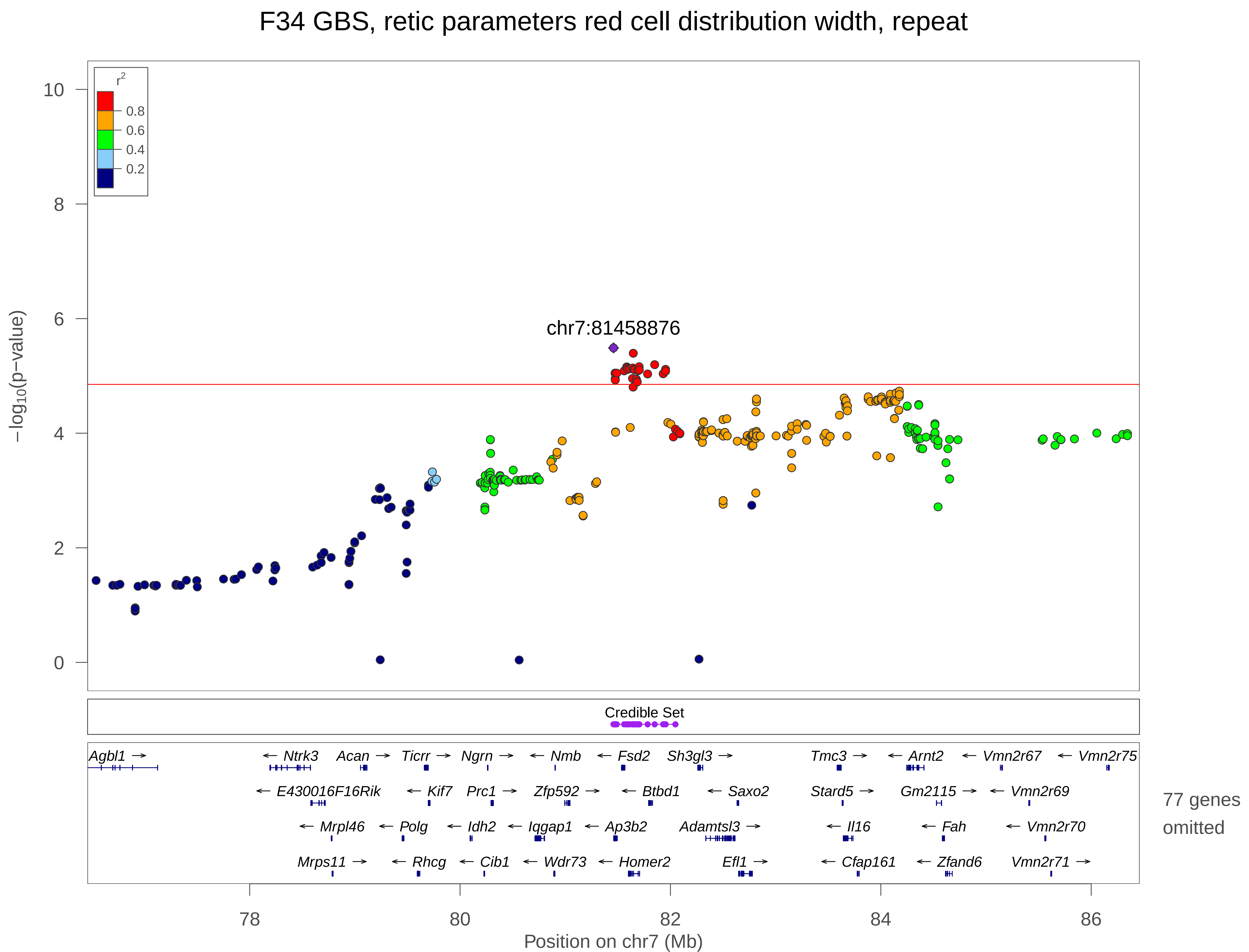

F34 GBS, retic parameters red cell distribution width, repeat
r2
0.8
0.6
0.4
0.2
10
8
6
4
2
0
−log10(p−value)
chr7:81458876
Credible Set
Agbl1 →
← Ntrk3
Acan →
Ticrr →
Ngrn →
← Nmb
← Fsd2
Sh3gl3 →
Tmc3 →
← Arnt2
← Vmn2r67
← Vmn2r75
← E430016F16Rik
← Kif7
Prc1 →
Zfp592 →
← Btbd1
← Saxo2
Stard5 →
Gm2115 →
← Vmn2r69
← Mrpl46
← Polg
← Idh2
← Iqgap1
← Ap3b2
Adamtsl3 →
← Il16
← Fah
← Vmn2r70
Mrps11 →
← Rhcg
← Cib1
← Wdr73
← Homer2
Efl1 →
← Cfap161
← Zfand6
Vmn2r71 →
77 genes omitted
78
80
82
84
86
Position on chr7 (Mb)

| | |
|---|---|
| date: | Thu Oct 25 17:10:40 2018 |
| build: | F34GBS |
| display range: | chr7:76458876−86458876 [76458876−86458876] |
| hilite range: | 0 − 0 [ 0 − 0 ] |
| reference SNP: | chr7:81458876 |
| number of SNPs plotted: | 417 |
| min P−value: | 3.25E−6 [chr7:81458876] |
| max P−value: | 9.08E−1 [chr7:80561830] |
| omitted Genes: | Det1, Mir7−2, Aen |
| omitted Genes: | Isg20, Gm26633, Hapln3 |
| omitted Genes: | Mfge8, Abhd2, Rlbp1 |
| omitted Genes: | Fanci, Mir9−3hg, Mir9−3 |
| omitted Genes: | Plin1, Pex11a, Wdr93 |
| omitted Genes: | Mesp1, Mesp2, Anpep |
| omitted Genes: | Ap3s2, Arpin, Zfp710 |
| omitted Genes: | Gm21057, Sema4b, Gdpgp1 |
| omitted Genes: | Ttll13, Vps33b, Rccd1 |
| omitted Genes: | Unc45a, Hddc3, Man2a2 |
| omitted Genes: | Fes, Furin, Blm |
| omitted Genes: | Crtc3, Gm15880, Zscan2 |
| omitted Genes: | Sec11a, Alpk3, Slc28a1 |
| omitted Genes: | Pde8a, Rps17, Cpeb1 |
| omitted Genes: | BC048679, 2900076A07Rik, Mir1839 |
| omitted Genes: | Whamm, Fam103a1, 3110040N11Rik |
| omitted Genes: | Tm6sf1, Hdgfrp3, Bnc1 |
| omitted Genes: | Gm20744, 4933406J10Rik, Mex3b |
| omitted Genes: | 4933430H16Rik, Mesdc1, Mesdc2 |
| omitted Genes: | Cemip, Abhd17c, 2610206C17Rik |
| omitted Genes: | Olfr291, Olfr290, Vmn2r65 |
| omitted Genes: | Vmn2r66, Vmn2r68, Vmn2r72 |
| omitted Genes: | Vmn2r73, Vmn2r74, Vmn2r76 |
| omitted Genes: | Olfr310, Olfr309, Olfr308 |
| omitted Genes: | Olfr307, Olfr305, Olfr304 |
| omitted Genes: | Olfr303, Olfr301 |

Make more plots at http://csg.sph.umich.edu/locuszoom/

Fine−mapping Regions

| snp | chr | pos (Mb) | pp | group | color |
|---|---|---|---|---|---|
| chr7:81458876 | 7 | 81.45888 | 0.06616771 | Credible Set | purple |
| chr7:81473821 | 7 | 81.47382 | 0.02512823 | Credible Set | purple |
| chr7:81473841 | 7 | 81.47384 | 0.02487803 | Credible Set | purple |
| chr7:81475123 | 7 | 81.47512 | 0.02016831 | Credible Set | purple |
| chr7:81475858 | 7 | 81.47586 | 0.01933686 | Credible Set | purple |
| chr7:81475869 | 7 | 81.47587 | 0.01932098 | Credible Set | purple |
| chr7:81489705 | 7 | 81.48971 | 0.02505795 | Credible Set | purple |
| chr7:81560306 | 7 | 81.56031 | 0.02734510 | Credible Set | purple |
| chr7:81584607 | 7 | 81.58461 | 0.03186561 | Credible Set | purple |
| chr7:81587028 | 7 | 81.58703 | 0.03051343 | Credible Set | purple |
| chr7:81587096 | 7 | 81.58710 | 0.03035124 | Credible Set | purple |
| chr7:81595380 | 7 | 81.59538 | 0.02904931 | Credible Set | purple |
| chr7:81614445 | 7 | 81.61445 | 0.02956262 | Credible Set | purple |
| chr7:81639723 | 7 | 81.63972 | 0.02040579 | Credible Set | purple |
| chr7:81641021 | 7 | 81.64102 | 0.03045086 | Credible Set | purple |
| chr7:81641166 | 7 | 81.64117 | 0.03051590 | Credible Set | purple |
| chr7:81645454 | 7 | 81.64545 | 0.01469745 | Credible Set | purple |
| chr7:81645464 | 7 | 81.64546 | 0.01470128 | Credible Set | purple |
| chr7:81646679 | 7 | 81.64668 | 0.05387744 | Credible Set | purple |
| chr7:81653902 | 7 | 81.65390 | 0.02886043 | Credible Set | purple |
| chr7:81660191 | 7 | 81.66019 | 0.02851127 | Credible Set | purple |
| chr7:81675727 | 7 | 81.67573 | 0.01983776 | Credible Set | purple |
| chr7:81675758 | 7 | 81.67576 | 0.01972593 | Credible Set | purple |
| chr7:81683805 | 7 | 81.68381 | 0.01807525 | Credible Set | purple |
| chr7:81683816 | 7 | 81.68382 | 0.01783724 | Credible Set | purple |
| chr7:81691936 | 7 | 81.69194 | 0.02749141 | Credible Set | purple |
| chr7:81691946 | 7 | 81.69195 | 0.02761668 | Credible Set | purple |
| chr7:81700477 | 7 | 81.70048 | 0.03110540 | Credible Set | purple |
| chr7:81702582 | 7 | 81.70258 | 0.03211692 | Credible Set | purple |
| chr7:81704380 | 7 | 81.70438 | 0.02835662 | Credible Set | purple |
| chr7:81783181 | 7 | 81.78318 | 0.02428726 | Credible Set | purple |
| chr7:81850247 | 7 | 81.85025 | 0.03473184 | Credible Set | purple |
| chr7:81931360 | 7 | 81.93136 | 0.02452354 | Credible Set | purple |
| chr7:81931538 | 7 | 81.93154 | 0.02456007 | Credible Set | purple |
| chr7:81953706 | 7 | 81.95371 | 0.02904836 | Credible Set | purple |
| chr7:81954104 | 7 | 81.95410 | 0.02705868 | Credible Set | purple |

Fine−mapping Regions

| snp | chr | pos (Mb) | pp | group | color |
|---|---|---|---|---|---|
| chr7:82047190 | 7 | 82.04719 | 0.002948056 | Credible Set | purple |

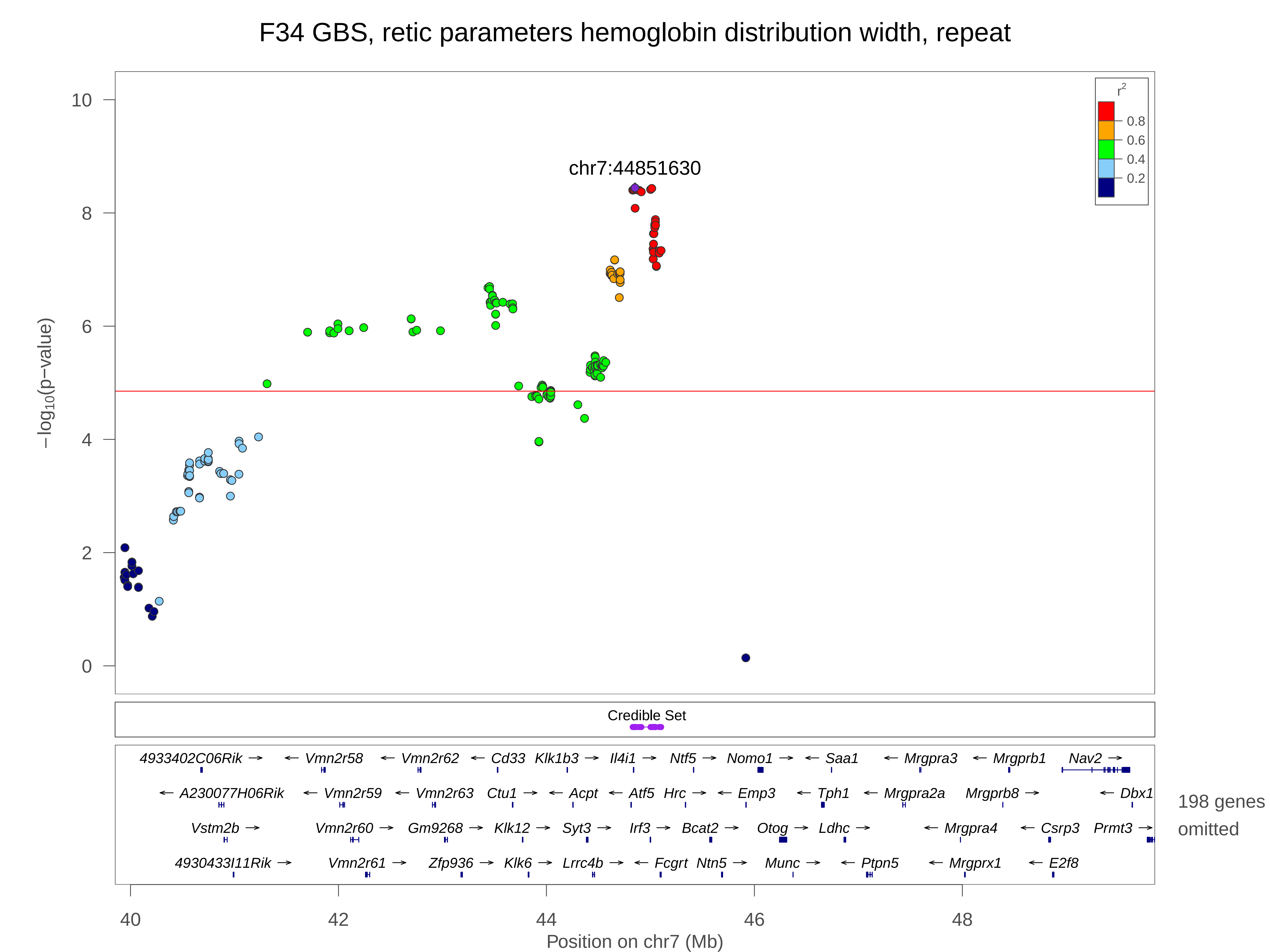

F34 GBS, retic parameters hemoglobin distribution width, repeat
chr7:44851630
r2
0.8
0.6
0.4
0.2
−log10(p−value)
10
8
6
4
2
0
Credible Set
4933402C06Rik → ← Vmn2r58 ← Vmn2r62 ← Cd33 Klk1b3 → Il4i1 → Ntf5 → Nomo1 → ← Saa1 ← Mrgpra3 ← Mrgprb1 Nav2 →
← A230077H06Rik ← Vmn2r59 ← Vmn2r63 Ctu1 → ← Acpt ← Atf5 Hrc → ← Emp3 ← Tph1 ← Mrgpra2a Mrgprb8 → ← Dbx1
Vstm2b → Vmn2r60 → Gm9268 → Klk12 → Syt3 → Irf3 → Bcat2 → Otog → Ldhc → ← Mrgpra4 ← Csrp3 Prmt3 →
4930433I11Rik → Vmn2r61 → Zfp936 → Klk6 → Lrrc4b → ← Fcgrt Ntn5 → Munc → ← Ptpn5 ← Mrgprx1 ← E2f8
198 genes omitted
40
42
44
46
48
Position on chr7 (Mb)

| | |
|---|---|
| date: | Thu Oct 25 17:11:16 2018 |
| build: | F34GBS |
| display range: | chr7:39851630−49851630 [39851630−49851630] |
| hilite range: | 0 − 0 [ 0 − 0 ] |
| reference SNP: | chr7:44851630 |
| number of SNPs plotted: | 247 |
| min P−value: | 3.6E−9 [chr7:44851630] |
| max P−value: | 7.22E−1 [chr7:45917340] |
| omitted Genes: | Gm4884, 1700110I07Rik, Gm5592 |
| omitted Genes: | AI987944, Vmn2r57, AW146154 |
| omitted Genes: | Zfp788, Vmn2r−ps54, Vmn2r−ps60 |
| omitted Genes: | 4933421I07Rik, Zfp141, 9830147E19Rik |
| omitted Genes: | Gm5595, Gm2381, EU599041 |
| omitted Genes: | Zfp715, Siglecf, Siglecg |
| omitted Genes: | Lim2, Nkg7, Cldnd2 |
| omitted Genes: | Etfb, Mir7051, Vsig10l |
| omitted Genes: | Iglon5, 4931406B18Rik, Zfp658 |
| omitted Genes: | Zfp719, Zfp819, Ceacam18 |
| omitted Genes: | Siglece, Klk14, Klk13 |
| omitted Genes: | 2310002F09Rik, Klk11, Klk10 |
| omitted Genes: | Klk9, Klk8, Klk7 |
| omitted Genes: | Klk5, Klk4, Klk15 |
| omitted Genes: | Klk1b7−ps, Klk1b8, Klk1b1 |
| omitted Genes: | Klk1b9, Klk1b11, Klk1b26 |
| omitted Genes: | Klk1b27, Klk1b21, Klk1b22 |
| omitted Genes: | Klk1b16, Klk1b24, Klk1b4 |
| omitted Genes: | Klk1b5, Klk1, 1700028J19Rik |
| omitted Genes: | 2410002F23Rik, Snord88c, Snord88a |
| omitted Genes: | Clec11a, Shank1, 1700008O03Rik |
| omitted Genes: | Aspdh, Josd2, Mir7052 |
| omitted Genes: | 5430431A17Rik, Emc10, Fam71e1 |
| omitted Genes: | Mybpc2, Spib, Pold1 |
| omitted Genes: | Mir7053, Nr1h2, Napsa |
| omitted Genes: | Kcnc3, Myh14, Izumo2 |
| omitted Genes: | Zfp473, Vrk3, Nup62 |
| omitted Genes: | Gm21948, Tbc1d17, Akt1s1 |
| omitted Genes: | Mir707, Pnkp, Ptov1 |
| omitted Genes: | Med25, Fuz, Ap2a1 |
| omitted Genes: | Tsks, Cpt1c, Prmt1 |
| omitted Genes: | Gm15545, Bcl2l12, Scaf1 |

Make more plots at http://csg.sph.umich.edu/locuszoom/

Fine−mapping Regions

| snp | chr | pos (Mb) | pp | group | color |
|---|---|---|---|---|---|
| chr7:44829933 | 7 | 44.82993 | 0.063242520 | Credible Set | purple |
| chr7:44839966 | 7 | 44.83997 | 0.064545439 | Credible Set | purple |
| chr7:44844016 | 7 | 44.84402 | 0.064505343 | Credible Set | purple |
| chr7:44844285 | 7 | 44.84428 | 0.065560645 | Credible Set | purple |
| chr7:44844322 | 7 | 44.84432 | 0.064505343 | Credible Set | purple |
| chr7:44851630 | 7 | 44.85163 | 0.069377128 | Credible Set | purple |
| chr7:44852639 | 7 | 44.85264 | 0.030886329 | Credible Set | purple |
| chr7:44852965 | 7 | 44.85296 | 0.067763560 | Credible Set | purple |
| chr7:44866590 | 7 | 44.86659 | 0.064281654 | Credible Set | purple |
| chr7:44891663 | 7 | 44.89166 | 0.062779023 | Credible Set | purple |
| chr7:44912209 | 7 | 44.91221 | 0.059203091 | Credible Set | purple |
| chr7:45001761 | 7 | 45.00176 | 0.065056667 | Credible Set | purple |
| chr7:45012901 | 7 | 45.01290 | 0.067751234 | Credible Set | purple |
| chr7:45025410 | 7 | 45.02541 | 0.006195526 | Credible Set | purple |
| chr7:45025443 | 7 | 45.02544 | 0.006214143 | Credible Set | purple |
| chr7:45026544 | 7 | 45.02654 | 0.004165466 | Credible Set | purple |
| chr7:45026548 | 7 | 45.02655 | 0.004157523 | Credible Set | purple |
| chr7:45029371 | 7 | 45.02937 | 0.011292601 | Credible Set | purple |
| chr7:45029402 | 7 | 45.02940 | 0.005576216 | Credible Set | purple |
| chr7:45029407 | 7 | 45.02941 | 0.005597764 | Credible Set | purple |
| chr7:45029413 | 7 | 45.02941 | 0.005455879 | Credible Set | purple |
| chr7:45029777 | 7 | 45.02978 | 0.007525295 | Credible Set | purple |
| chr7:45033529 | 7 | 45.03353 | 0.011292601 | Credible Set | purple |
| chr7:45041503 | 7 | 45.04150 | 0.016291766 | Credible Set | purple |
| chr7:45041522 | 7 | 45.04152 | 0.015598120 | Credible Set | purple |
| chr7:45041933 | 7 | 45.04193 | 0.014408712 | Credible Set | purple |
| chr7:45047834 | 7 | 45.04783 | 0.019762211 | Credible Set | purple |
| chr7:45047913 | 7 | 45.04791 | 0.018037235 | Credible Set | purple |
| chr7:45049496 | 7 | 45.04950 | 0.015855521 | Credible Set | purple |
| chr7:45084416 | 7 | 45.08442 | 0.005752337 | Credible Set | purple |
| chr7:45084764 | 7 | 45.08476 | 0.005257411 | Credible Set | purple |
| chr7:45102598 | 7 | 45.10260 | 0.005813696 | Credible Set | purple |

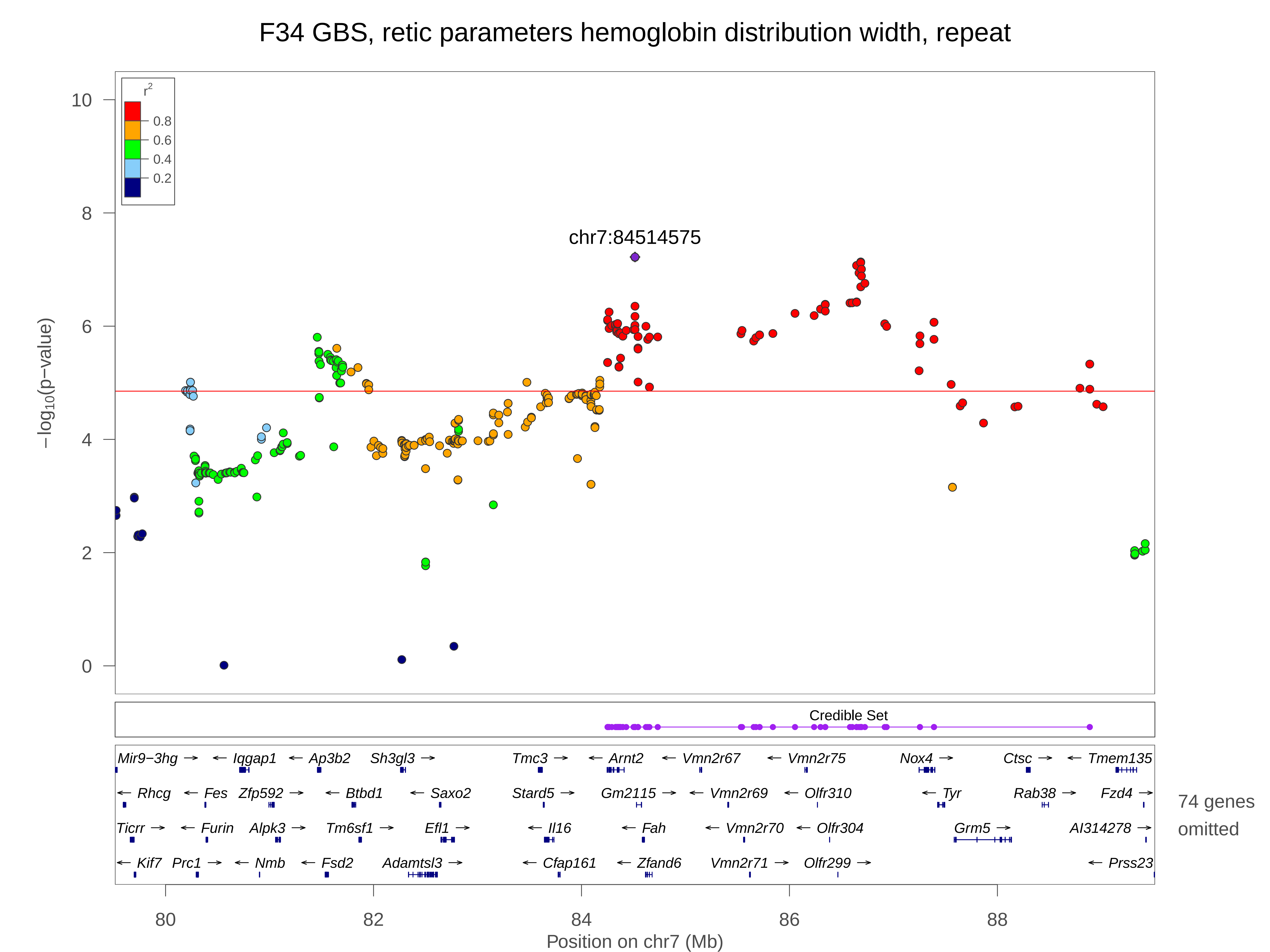
F34 GBS, retic parameters hemoglobin distribution width, repeat
r2
0.8
0.6
0.4
0.2
chr7:84514575
−log10(p−value)
10
8
6
4
2
0
Credible Set
Mir9−3hg → ← Iqgap1 ← Ap3b2 Sh3gl3 → Tmc3 → ← Arnt2 ← Vmn2r67 ← Vmn2r75 Nox4 → Ctsc → ← Tmem135
← Rhcg ← Fes Zfp592 → ← Btbd1 ← Saxo2 Stard5 → Gm2115 → ← Vmn2r69 ← Olfr310 ← Tyr Rab38 → Fzd4 →
Ticrr → ← Furin Alpk3 → Tm6sf1 → Efl1 → ← Il16 ← Fah ← Vmn2r70 ← Olfr304 Grm5 → AI314278 →
← Kif7 Prc1 → ← Nmb ← Fsd2 Adamtsl3 → ← Cfap161 ← Zfand6 Vmn2r71 → Olfr299 → ← Prss23
74 genes omitted
80
82
84
86
88
Position on chr7 (Mb)

| | |
|---|---|
| date: | Thu Oct 25 17:11:26 2018 |
| build: | F34GBS |
| display range: | chr7:79514575−89514575 [79514575−89514575] |
| hilite range: | 0 − 0 [ 0 − 0 ] |
| reference SNP: | chr7:84514575 |
| number of SNPs plotted: | 389 |
| min P−value: | 6E−8 [chr7:84514575] |
| max P−value: | 9.75E−1 [chr7:80561830] |
| omitted Genes: | Plin1, Pex11a, Wdr93 |
| omitted Genes: | Mesp1, Mesp2, Anpep |
| omitted Genes: | Ap3s2, Arpin, Zfp710 |
| omitted Genes: | Gm21057, Idh2, Sema4b |
| omitted Genes: | Cib1, Gdpgp1, Ttll13 |
| omitted Genes: | Ngrn, Vps33b, Rccd1 |
| omitted Genes: | Unc45a, Hddc3, Man2a2 |
| omitted Genes: | Blm, Crtc3, Gm15880 |
| omitted Genes: | Zscan2, Wdr73, Sec11a |
| omitted Genes: | Slc28a1, Pde8a, Rps17 |
| omitted Genes: | Cpeb1, BC048679, 2900076A07Rik |
| omitted Genes: | Mir1839, Whamm, Homer2 |
| omitted Genes: | Fam103a1, 3110040N11Rik, Hdgfrp3 |
| omitted Genes: | Bnc1, Gm20744, 4933406J10Rik |
| omitted Genes: | Mex3b, 4933430H16Rik, Mesdc1 |
| omitted Genes: | Mesdc2, Cemip, Abhd17c |
| omitted Genes: | 2610206C17Rik, Olfr291, Olfr290 |
| omitted Genes: | Vmn2r65, Vmn2r66, Vmn2r68 |
| omitted Genes: | Vmn2r72, Vmn2r73, Vmn2r74 |
| omitted Genes: | Vmn2r76, Olfr309, Olfr308 |
| omitted Genes: | Olfr307, Olfr305, Olfr303 |
| omitted Genes: | Olfr301, Olfr298, Olfr297 |
| omitted Genes: | Olfr295, Olfr294, Olfr293 |
| omitted Genes: | Olfr292, Folh1, Vmn2r77 |
| omitted Genes: | Vmn2r78, Vmn2r79 |

Make more plots at http://csg.sph.umich.edu/locuszoom/

Fine−mapping Regions

| snp | chr | pos (Mb) | pp | group | color |
|---|---|---|---|---|---|
| chr7:84250985 | 7 | 84.25099 | 0.001360020 | Credible Set | purple |
| chr7:84251011 | 7 | 84.25101 | 0.001346588 | Credible Set | purple |
| chr7:84251159 | 7 | 84.25116 | 0.006956519 | Credible Set | purple |
| chr7:84251209 | 7 | 84.25121 | 0.007221524 | Credible Set | purple |
| chr7:84265410 | 7 | 84.26541 | 0.009742708 | Credible Set | purple |
| chr7:84265685 | 7 | 84.26569 | 0.005104929 | Credible Set | purple |
| chr7:84291540 | 7 | 84.29154 | 0.005729120 | Credible Set | purple |
| chr7:84327288 | 7 | 84.32729 | 0.005562292 | Credible Set | purple |
| chr7:84327808 | 7 | 84.32781 | 0.005978483 | Credible Set | purple |
| chr7:84327850 | 7 | 84.32785 | 0.005967179 | Credible Set | purple |
| chr7:84338595 | 7 | 84.33859 | 0.004445736 | Credible Set | purple |
| chr7:84343118 | 7 | 84.34312 | 0.004730137 | Credible Set | purple |
| chr7:84344805 | 7 | 84.34480 | 0.006031629 | Credible Set | purple |
| chr7:84347423 | 7 | 84.34742 | 0.006204436 | Credible Set | purple |
| chr7:84360038 | 7 | 84.36004 | 0.004234925 | Credible Set | purple |
| chr7:84360224 | 7 | 84.36022 | 0.001160061 | Credible Set | purple |
| chr7:84360626 | 7 | 84.36063 | 0.004172698 | Credible Set | purple |
| chr7:84360642 | 7 | 84.36064 | 0.004172698 | Credible Set | purple |
| chr7:84360703 | 7 | 84.36070 | 0.004172698 | Credible Set | purple |
| chr7:84360714 | 7 | 84.36071 | 0.004172698 | Credible Set | purple |
| chr7:84360716 | 7 | 84.36072 | 0.004172698 | Credible Set | purple |
| chr7:84375526 | 7 | 84.37553 | 0.004291484 | Credible Set | purple |
| chr7:84375577 | 7 | 84.37558 | 0.001605662 | Credible Set | purple |
| chr7:84375642 | 7 | 84.37564 | 0.001605662 | Credible Set | purple |
| chr7:84397575 | 7 | 84.39758 | 0.003784376 | Credible Set | purple |
| chr7:84397597 | 7 | 84.39760 | 0.003784376 | Credible Set | purple |
| chr7:84430102 | 7 | 84.43010 | 0.004736942 | Credible Set | purple |
| chr7:84501082 | 7 | 84.50108 | 0.004881938 | Credible Set | purple |
| chr7:84514082 | 7 | 84.51408 | 0.005771754 | Credible Set | purple |
| chr7:84514444 | 7 | 84.51444 | 0.012233069 | Credible Set | purple |
| chr7:84514505 | 7 | 84.51450 | 0.008203041 | Credible Set | purple |
| chr7:84514564 | 7 | 84.51456 | 0.083415916 | Credible Set | purple |
| chr7:84514575 | 7 | 84.51457 | 0.084641734 | Credible Set | purple |
| chr7:84514734 | 7 | 84.51473 | 0.004890651 | Credible Set | purple |
| chr7:84543346 | 7 | 84.54335 | 0.002387437 | Credible Set | purple |
| chr7:84543355 | 7 | 84.54336 | 0.002286908 | Credible Set | purple |

Fine−mapping Regions

| snp | chr | pos (Mb) | pp | group | color |
|---|---|---|---|---|---|
| chr7:84543782 | 7 | 84.54378 | 0.003726403 | Credible Set | purple |
| chr7:84619632 | 7 | 84.61963 | 0.005569132 | Credible Set | purple |
| chr7:84636759 | 7 | 84.63676 | 0.003331992 | Credible Set | purple |
| chr7:84653095 | 7 | 84.65309 | 0.003658194 | Credible Set | purple |
| chr7:84733186 | 7 | 84.73319 | 0.003667514 | Credible Set | purple |
| chr7:85533207 | 7 | 85.53321 | 0.004150514 | Credible Set | purple |
| chr7:85543843 | 7 | 85.54384 | 0.004733558 | Credible Set | purple |
| chr7:85656243 | 7 | 85.65624 | 0.003123987 | Credible Set | purple |
| chr7:85677494 | 7 | 85.67749 | 0.003564857 | Credible Set | purple |
| chr7:85712663 | 7 | 85.71266 | 0.003985331 | Credible Set | purple |
| chr7:85712975 | 7 | 85.71298 | 0.003962875 | Credible Set | purple |
| chr7:85840665 | 7 | 85.84067 | 0.004199307 | Credible Set | purple |
| chr7:86053386 | 7 | 86.05339 | 0.009230214 | Credible Set | purple |
| chr7:86237374 | 7 | 86.23737 | 0.008469903 | Credible Set | purple |
| chr7:86298979 | 7 | 86.29898 | 0.010928221 | Credible Set | purple |
| chr7:86344420 | 7 | 86.34442 | 0.013185160 | Credible Set | purple |
| chr7:86344437 | 7 | 86.34444 | 0.012975268 | Credible Set | purple |
| chr7:86344488 | 7 | 86.34449 | 0.010088022 | Credible Set | purple |
| chr7:86582139 | 7 | 86.58214 | 0.013905831 | Credible Set | purple |
| chr7:86603770 | 7 | 86.60377 | 0.013969827 | Credible Set | purple |
| chr7:86645719 | 7 | 86.64572 | 0.014498747 | Credible Set | purple |
| chr7:86645792 | 7 | 86.64579 | 0.014256802 | Credible Set | purple |
| chr7:86645947 | 7 | 86.64595 | 0.060698996 | Credible Set | purple |
| chr7:86645990 | 7 | 86.64599 | 0.060628029 | Credible Set | purple |
| chr7:86668998 | 7 | 86.66900 | 0.045196830 | Credible Set | purple |
| chr7:86685426 | 7 | 86.68543 | 0.069735103 | Credible Set | purple |
| chr7:86685507 | 7 | 86.68551 | 0.067735636 | Credible Set | purple |
| chr7:86685648 | 7 | 86.68565 | 0.026167191 | Credible Set | purple |
| chr7:86692214 | 7 | 86.69221 | 0.052430898 | Credible Set | purple |
| chr7:86692491 | 7 | 86.69249 | 0.040021818 | Credible Set | purple |
| chr7:86725518 | 7 | 86.72552 | 0.030008906 | Credible Set | purple |
| chr7:86916161 | 7 | 86.91616 | 0.006155316 | Credible Set | purple |
| chr7:86934710 | 7 | 86.93471 | 0.005532621 | Credible Set | purple |
| chr7:87254690 | 7 | 87.25469 | 0.002814049 | Credible Set | purple |
| chr7:87255156 | 7 | 87.25516 | 0.003831057 | Credible Set | purple |
| chr7:87389952 | 7 | 87.38995 | 0.003341701 | Credible Set | purple |

Fine−mapping Regions

| snp | chr | pos (Mb) | pp | group | color |
|---|---|---|---|---|---|
| chr7:87389989 | 7 | 87.38999 | 0.006501179 | Credible Set | purple |
| chr7:87390014 | 7 | 87.39001 | 0.006521963 | Credible Set | purple |
| chr7:88888100 | 7 | 88.88810 | 0.001272919 | Credible Set | purple |
| chr7:88888124 | 7 | 88.88812 | 0.001272919 | Credible Set | purple |

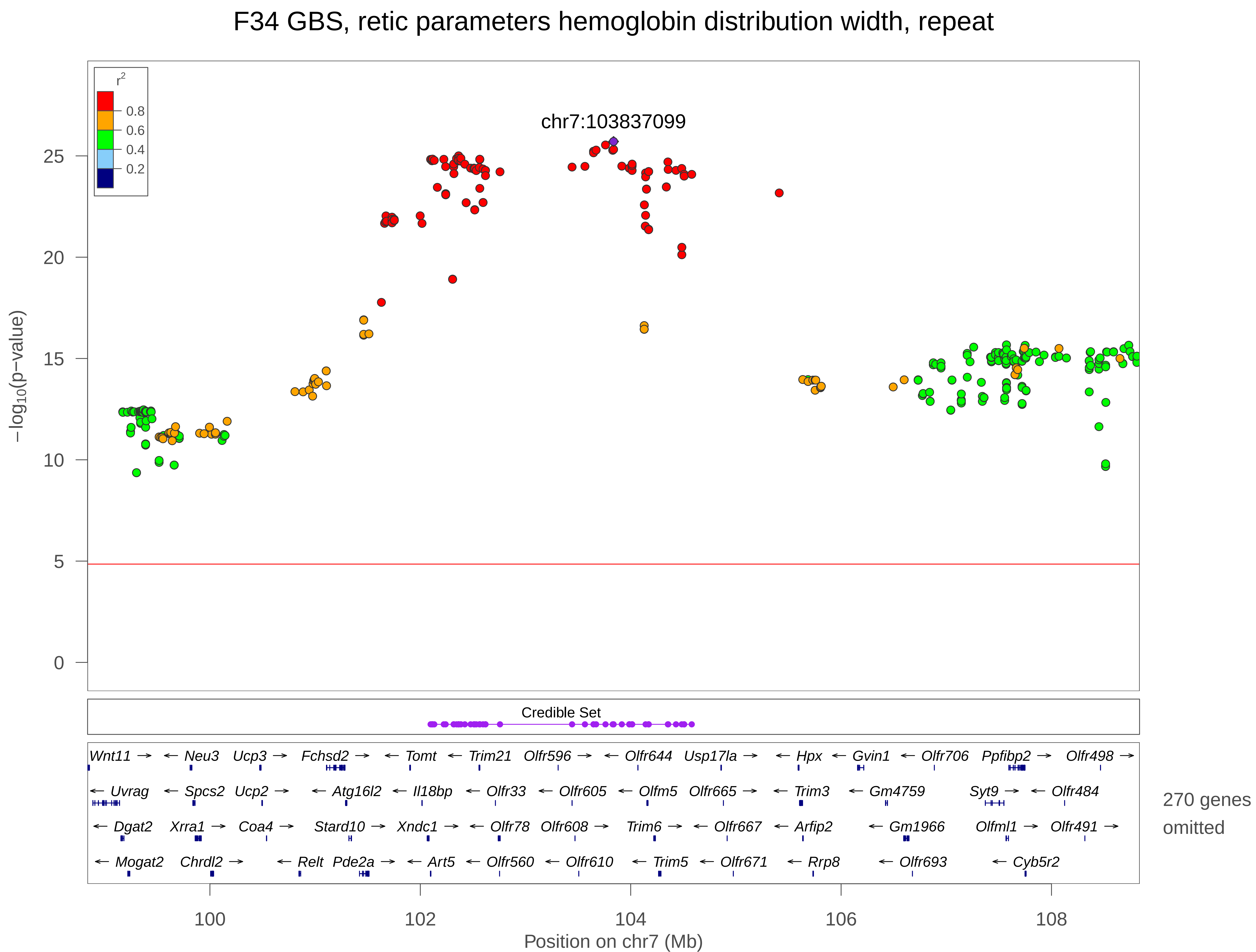

F34 GBS, retic parameters hemoglobin distribution width, repeat
r2
0.8
0.6
0.4
0.2
chr7:103837099
25
20
15
10
5
0
−log10(p−value)
Credible Set
Wnt11 → ← Neu3 Ucp3 → Fchsd2 → ← Tomt ← Trim21 Olfr596 → ← Olfr644 Usp17la → ← Hpx ← Gvin1 ← Olfr706 Ppfibp2 → Olfr498 →
← Uvrag ← Spcs2 Ucp2 → ← Atg16l2 ← Il18bp ← Olfr33 ← Olfr605 ← Olfm5 Olfr665 → ← Trim3 ← Gm4759 Syt9 → ← Olfr484
← Dgat2 Xrra1 → Coa4 → Stard10 → Xndc1 → ← Olfr78 Olfr608 → Trim6 → ← Olfr667 ← Arfip2 ← Gm1966 Olfml1 → Olfr491 →
← Mogat2 Chrdl2 → ← Relt Pde2a → ← Art5 ← Olfr560 ← Olfr610 ← Trim5 ← Olfr671 ← Rrp8 ← Olfr693 ← Cyb5r2
270 genes omitted
100
102
104
106
108
Position on chr7 (Mb)

date: Thu Oct 25 17:11:03 2018
build: F34GBS
display range: chr7:98837099−108837099 [98837099−108837099]
hilite range: 0 − 0 [ 0 − 0 ]
reference SNP: chr7:103837099
number of SNPs plotted: 422
min P−value: 1.95E−26 [chr7:103837099]
max P−value: 4.36E−10 [chr7:99302296]
omitted Genes: Gm26705, Map6, Serpinh1
omitted Genes: Gdpd5, Klhl35, Rps3
omitted Genes: Snord15b, Snord15a, Arrb1
omitted Genes: Mir326, Tpbgl, Slco2b1
omitted Genes: Olfr520, Olfr521, Rnf169
omitted Genes: Pold3, Lipt2, Kcne3
omitted Genes: Pgm2l1, Gpx2−ps1, P4ha3
omitted Genes: Ppme1, C2cd3, Dnajb13
omitted Genes: Mrpl48, Rab6a, Plekhb1
omitted Genes: Fam168a, Arhgef17, Mir3102
omitted Genes: P2ry6, P2ry2, Arap1
omitted Genes: Mir139, Art2a−ps, Art2b
omitted Genes: Clpb, Phox2a, Inppl1
omitted Genes: Folr2, Folr1, Anapc15
omitted Genes: Lamtor1, Lrrc51, Numa1
omitted Genes: Rnf121, Xntrpc, Trpc2
omitted Genes: Art1, Chrna10, Nup98
omitted Genes: Pgap2, Rhog, Stim1
omitted Genes: Rrm1, Olfr543, Olfr544
omitted Genes: Olfr545, Olfr547, Olfr549
omitted Genes: Olfr550, Olfr551, Olfr552
omitted Genes: Olfr553, Olfr554, Olfr555
omitted Genes: Olfr556, Trim68, Olfr557
omitted Genes: Olfr558, Olfr559, Olfr561
omitted Genes: Olfr564, Olfr566, Olfr568
omitted Genes: Olfr569, Olfr570, Olfr571
omitted Genes: Olfr572, Olfr574, Olfr575
omitted Genes: Olfr576, Olfr577, Olfr578
omitted Genes: Olfr582, Olfr583, Olfr584
omitted Genes: Olfr585, Olfr586, Olfr589
omitted Genes: Olfr591, Olfr592, Olfr593
omitted Genes: Olfr594, Usp17ld, Olfr597

Fine−mapping Regions

| snp | chr | pos (Mb) | pp | group | color |
|---|---|---|---|---|---|
| chr7:102098054 | 7 | 102.0981 | 0.015832258 | Credible Set | purple |
| chr7:102110109 | 7 | 102.1101 | 0.015679915 | Credible Set | purple |
| chr7:102110501 | 7 | 102.1105 | 0.013548429 | Credible Set | purple |
| chr7:102116318 | 7 | 102.1163 | 0.015078950 | Credible Set | purple |
| chr7:102116430 | 7 | 102.1164 | 0.015078950 | Credible Set | purple |
| chr7:102116478 | 7 | 102.1165 | 0.016308699 | Credible Set | purple |
| chr7:102132562 | 7 | 102.1326 | 0.014142236 | Credible Set | purple |
| chr7:102223396 | 7 | 102.2234 | 0.016016491 | Credible Set | purple |
| chr7:102240695 | 7 | 102.2407 | 0.007016629 | Credible Set | purple |
| chr7:102317213 | 7 | 102.3172 | 0.006918154 | Credible Set | purple |
| chr7:102317246 | 7 | 102.3172 | 0.009417055 | Credible Set | purple |
| chr7:102319586 | 7 | 102.3196 | 0.003176024 | Credible Set | purple |
| chr7:102344970 | 7 | 102.3450 | 0.017546832 | Credible Set | purple |
| chr7:102363897 | 7 | 102.3639 | 0.013224617 | Credible Set | purple |
| chr7:102364204 | 7 | 102.3642 | 0.023364889 | Credible Set | purple |
| chr7:102364223 | 7 | 102.3642 | 0.023594202 | Credible Set | purple |
| chr7:102364315 | 7 | 102.3643 | 0.017988068 | Credible Set | purple |
| chr7:102364326 | 7 | 102.3643 | 0.017988068 | Credible Set | purple |
| chr7:102364411 | 7 | 102.3644 | 0.014440388 | Credible Set | purple |
| chr7:102385520 | 7 | 102.3855 | 0.013034527 | Credible Set | purple |
| chr7:102385536 | 7 | 102.3855 | 0.017930915 | Credible Set | purple |
| chr7:102423165 | 7 | 102.4232 | 0.009215167 | Credible Set | purple |
| chr7:102478469 | 7 | 102.4785 | 0.005923028 | Credible Set | purple |
| chr7:102510475 | 7 | 102.5105 | 0.005932441 | Credible Set | purple |
| chr7:102510607 | 7 | 102.5106 | 0.005242781 | Credible Set | purple |
| chr7:102516878 | 7 | 102.5169 | 0.005652553 | Credible Set | purple |
| chr7:102531652 | 7 | 102.5317 | 0.004499354 | Credible Set | purple |
| chr7:102561708 | 7 | 102.5617 | 0.006361622 | Credible Set | purple |
| chr7:102565262 | 7 | 102.5653 | 0.015584216 | Credible Set | purple |
| chr7:102565292 | 7 | 102.5653 | 0.015993636 | Credible Set | purple |
| chr7:102565307 | 7 | 102.5653 | 0.015632984 | Credible Set | purple |
| chr7:102565323 | 7 | 102.5653 | 0.016043661 | Credible Set | purple |
| chr7:102565329 | 7 | 102.5653 | 0.016043661 | Credible Set | purple |
| chr7:102596442 | 7 | 102.5964 | 0.005353120 | Credible Set | purple |
| chr7:102619418 | 7 | 102.6194 | 0.004574513 | Credible Set | purple |
| chr7:102619431 | 7 | 102.6194 | 0.004584162 | Credible Set | purple |

Fine−mapping Regions

| snp | chr | pos (Mb) | pp | group | color |
|---|---|---|---|---|---|
| chr7:102619466 | 7 | 102.6195 | 0.002539171 | Credible Set | purple |
| chr7:102757499 | 7 | 102.7575 | 0.003908190 | Credible Set | purple |
| chr7:103442570 | 7 | 103.4426 | 0.006669963 | Credible Set | purple |
| chr7:103564836 | 7 | 103.5648 | 0.007212812 | Credible Set | purple |
| chr7:103646439 | 7 | 103.6464 | 0.039655634 | Credible Set | purple |
| chr7:103646709 | 7 | 103.6467 | 0.033269131 | Credible Set | purple |
| chr7:103671033 | 7 | 103.6710 | 0.045122825 | Credible Set | purple |
| chr7:103760301 | 7 | 103.7603 | 0.080303862 | Credible Set | purple |
| chr7:103829200 | 7 | 103.8292 | 0.044125024 | Credible Set | purple |
| chr7:103837099 | 7 | 103.8371 | 0.118562548 | Credible Set | purple |
| chr7:103837213 | 7 | 103.8372 | 0.048463508 | Credible Set | purple |
| chr7:103915503 | 7 | 103.9155 | 0.007395257 | Credible Set | purple |
| chr7:103986792 | 7 | 103.9868 | 0.005860667 | Credible Set | purple |
| chr7:103986795 | 7 | 103.9868 | 0.005860667 | Credible Set | purple |
| chr7:103986857 | 7 | 103.9869 | 0.005916115 | Credible Set | purple |
| chr7:103986860 | 7 | 103.9869 | 0.005916115 | Credible Set | purple |
| chr7:104012808 | 7 | 104.0128 | 0.007223316 | Credible Set | purple |
| chr7:104013370 | 7 | 104.0134 | 0.004539445 | Credible Set | purple |
| chr7:104013395 | 7 | 104.0134 | 0.007895879 | Credible Set | purple |
| chr7:104013467 | 7 | 104.0135 | 0.009328578 | Credible Set | purple |
| chr7:104141705 | 7 | 104.1417 | 0.003438599 | Credible Set | purple |
| chr7:104170588 | 7 | 104.1706 | 0.004024190 | Credible Set | purple |
| chr7:104170594 | 7 | 104.1706 | 0.003994168 | Credible Set | purple |
| chr7:104170618 | 7 | 104.1706 | 0.004003845 | Credible Set | purple |
| chr7:104170662 | 7 | 104.1707 | 0.004003845 | Credible Set | purple |
| chr7:104353491 | 7 | 104.3535 | 0.011885581 | Credible Set | purple |
| chr7:104356230 | 7 | 104.3562 | 0.005107791 | Credible Set | purple |
| chr7:104430205 | 7 | 104.4302 | 0.004595604 | Credible Set | purple |
| chr7:104485236 | 7 | 104.4852 | 0.005695110 | Credible Set | purple |
| chr7:104507724 | 7 | 104.5077 | 0.002769857 | Credible Set | purple |
| chr7:104508127 | 7 | 104.5081 | 0.002948166 | Credible Set | purple |
| chr7:104508173 | 7 | 104.5082 | 0.002399265 | Credible Set | purple |
| chr7:104580091 | 7 | 104.5801 | 0.002936084 | Credible Set | purple |

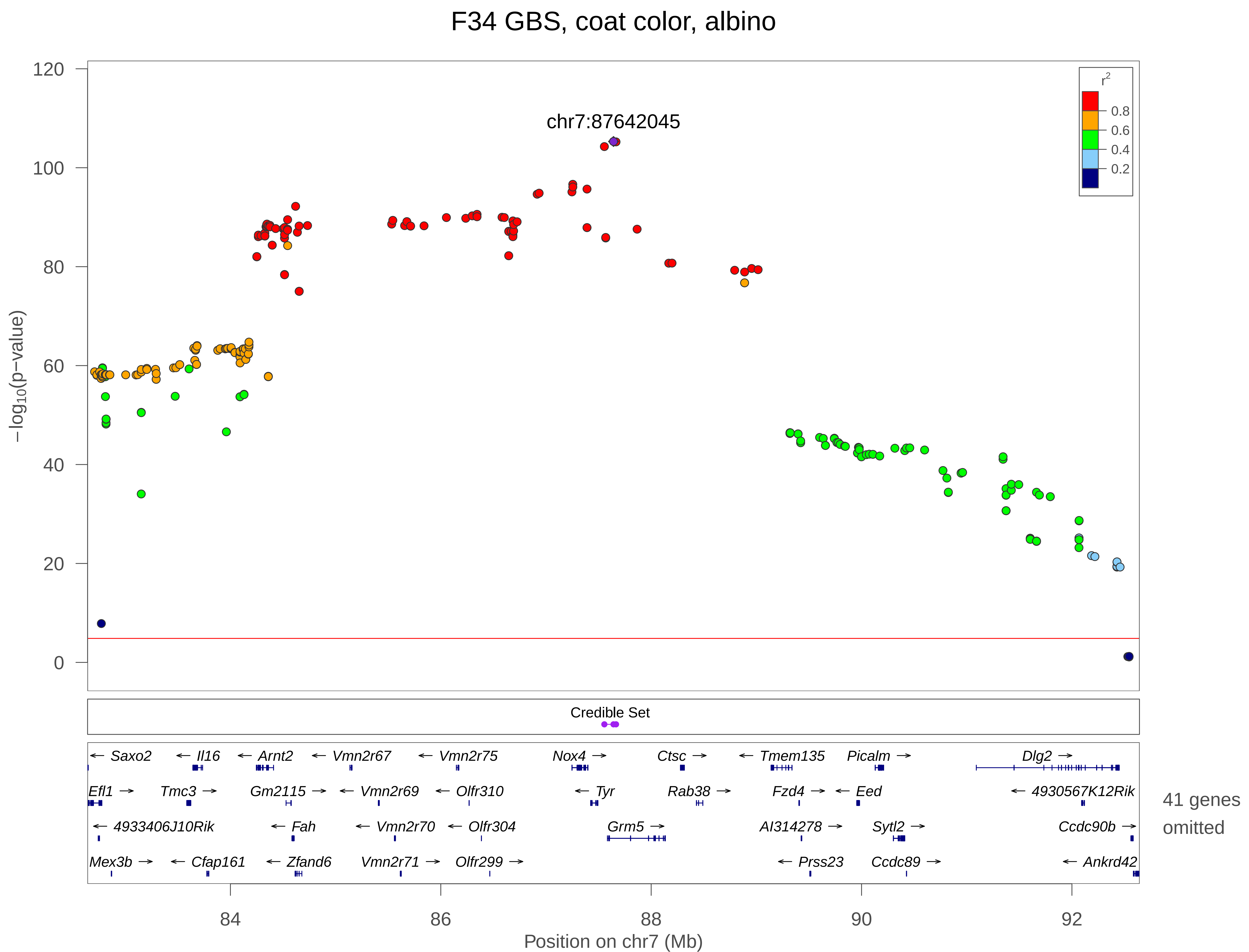

F34 GBS, coat color, albino
chr7:87642045
r2
0.8
0.6
0.4
0.2
120
100
80
60
40
20
0
−log10(p−value)
Credible Set
← Saxo2
← Il16
← Arnt2
← Vmn2r67
← Vmn2r75
Nox4 →
Ctsc →
← Tmem135
Picalm →
Dlg2 →
Efl1 →
Tmc3 →
Gm2115 →
← Vmn2r69
← Olfr310
← Tyr
Rab38 →
Fzd4 →
← Eed
← 4930567K12Rik
← 4933406J10Rik
← Fah
← Vmn2r70
← Olfr304
Grm5 →
AI314278 →
Sytl2 →
Ccdc90b →
Mex3b →
← Cfap161
← Zfand6
Vmn2r71 →
Olfr299 →
← Prss23
Ccdc89 →
← Ankrd42
41 genes omitted
84
86
88
90
92
Position on chr7 (Mb)

| | |
|---|---|
| date: | Thu Oct 25 17:13:07 2018 |
| build: | F34GBS |
| display range: | chr7:82642045−92642045 [82642045−92642045] |
| hilite range: | 0 − 0 [ 0 − 0 ] |
| reference SNP: | chr7:87642045 |
| number of SNPs plotted: | 282 |
| min P−value: | 5E−106 [chr7:87642045] |
| max P−value: | 7.84E−2 [chr7:92543164] |
| omitted Genes: | 4933430H16Rik, Stard5, Mesdc1 |
| omitted Genes: | Mesdc2, Cemip, Abhd17c |
| omitted Genes: | 2610206C17Rik, Olfr291, Olfr290 |
| omitted Genes: | Vmn2r65, Vmn2r66, Vmn2r68 |
| omitted Genes: | Vmn2r72, Vmn2r73, Vmn2r74 |
| omitted Genes: | Vmn2r76, Olfr309, Olfr308 |
| omitted Genes: | Olfr307, Olfr305, Olfr303 |
| omitted Genes: | Olfr301, Olfr298, Olfr297 |
| omitted Genes: | Olfr295, Olfr294, Olfr293 |
| omitted Genes: | Olfr292, Folh1, Vmn2r77 |
| omitted Genes: | Vmn2r78, Vmn2r79, Me3 |
| omitted Genes: | Ccdc81, I7Rn6, E230029C05Rik |
| omitted Genes: | 2310010J17Rik, Ccdc83, Crebzf |
| omitted Genes: | Tmem126a, Tmem126b |

Make more plots at http://csg.sph.umich.edu/locuszoom/

Fine−mapping Regions

| snp | chr | pos (Mb) | pp | group | color |
|---|---|---|---|---|---|
| chr7:87554961 | 7 | 87.55496 | 0.04890514 | Credible Set | purple |
| chr7:87642045 | 7 | 87.64204 | 0.51626891 | Credible Set | purple |
| chr7:87665727 | 7 | 87.66573 | 0.43482595 | Credible Set | purple |

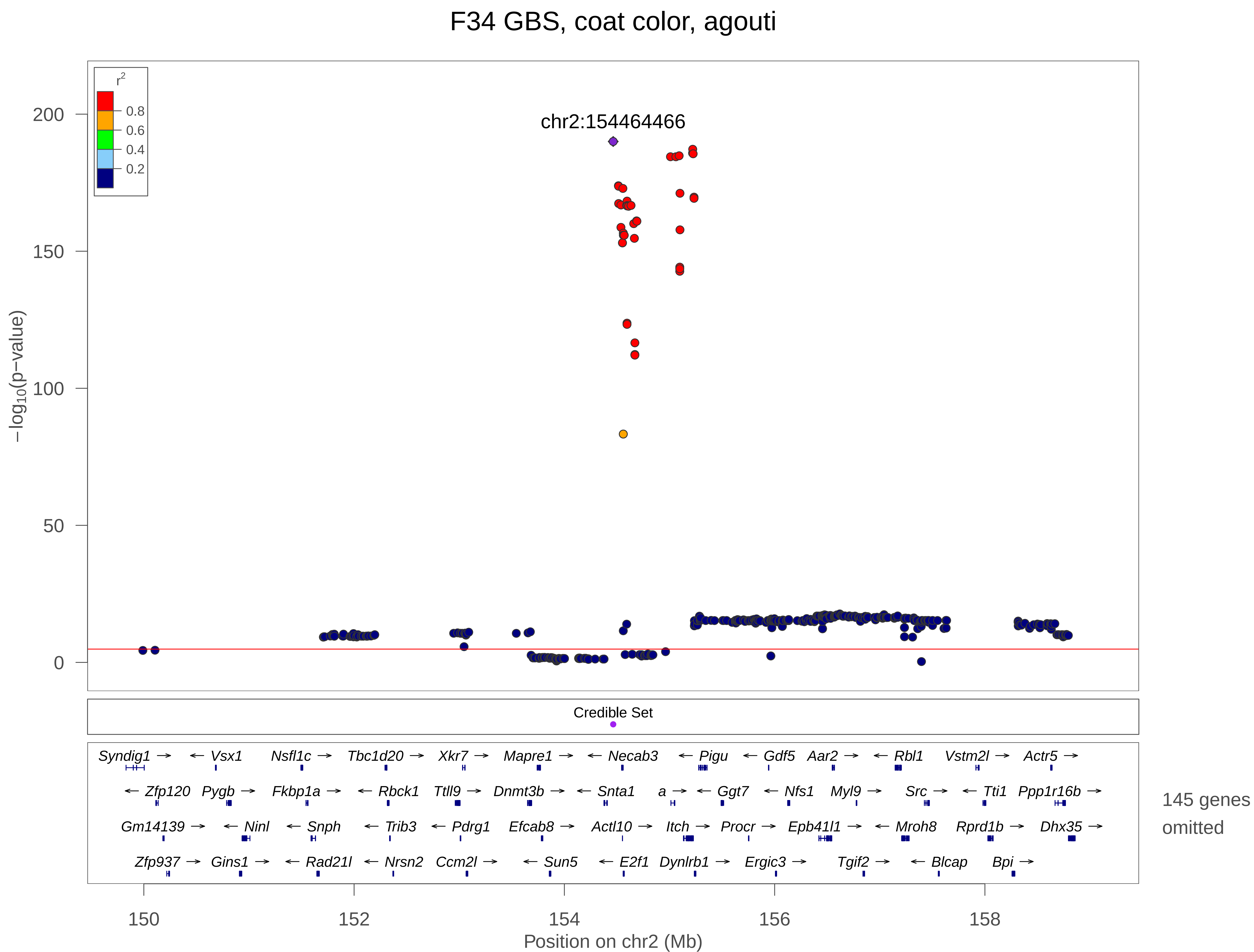

F34 GBS, coat color, agouti
r2
0.8
0.6
0.4
0.2
200
150
100
50
0
−log10(p−value)
chr2:154464466
Credible Set
Syndig1 →
← Vsx1
Nsfl1c →
Tbc1d20 →
Xkr7 →
Mapre1 →
← Necab3
← Pigu
← Gdf5
Aar2 →
← Rbl1
Vstm2l →
Actr5 →
← Zfp120
Pygb →
Fkbp1a →
← Rbck1
Ttll9 →
Dnmt3b →
← Snta1
a →
← Ggt7
← Nfs1
Myl9 →
Src →
← Tti1
Ppp1r16b →
Gm14139 →
← Ninl
← Snph
← Trib3
← Pdrg1
Efcab8 →
Actl10 →
Itch →
Procr →
Epb41l1 →
← Mroh8
Rprd1b →
Dhx35 →
Zfp937 →
Gins1 →
← Rad21l
← Nrsn2
Ccm2l →
← Sun5
← E2f1
Dynlrb1 →
Ergic3 →
Tgif2 →
← Blcap
Bpi →
145 genes omitted
150
152
154
156
158
Position on chr2 (Mb)

date: Thu Oct 25 17:13:18 2018
build: F34GBS
display range: chr2:149464466−159464466 [149464466−159464466]
hilite range: 0 − 0 [ 0 − 0 ]
reference SNP: chr2:154464466
number of SNPs plotted: 748
min P−value: 9.43E−191 [chr2:154464466]
max P−value: 4.9E−1 [chr2:157396497]
omitted Genes: Gm14124, 3300002I08Rik, Zfp442
omitted Genes: Zfp345, Cst7, Apmap
omitted Genes: Acss1, E130215H24Rik, Entpd6
omitted Genes: Abhd12, Nanp, Gm14151
omitted Genes: 4930442J19Rik, LOC100125594, 4921509C19Rik
omitted Genes: Sdcbp2, Tmem74b, Psmf1
omitted Genes: Rspo4, Angpt4, Mir1953
omitted Genes: Fam110a, Slc52a3, Scrt2
omitted Genes: Srxn1, Tcf15, Csnk2a1
omitted Genes: Gm14164, Sox12, Zcchc3
omitted Genes: 6820408C15Rik, Defb23, Defb20
omitted Genes: Defb22, Defb26, Defb28
omitted Genes: Defb29, Defb21, Defb19
omitted Genes: Defb45, Defb36, Defb25
omitted Genes: Rem1, H13, Mcts2
omitted Genes: Id1, Cox4i2, Bcl2l1
omitted Genes: Tpx2, Mir5622, Mylk2
omitted Genes: Foxs1, Dusp15, Hck
omitted Genes: Tm9sf4, Tspyl3, Plagl2
omitted Genes: Pofut1, Kif3b, 2500004C02Rik
omitted Genes: Asxl1, Nol4l, Commd7
omitted Genes: Bpifb2, Bpifb6, Bpifb3
omitted Genes: Bpifb4, Bpifa6, Bpifa2
omitted Genes: Bpifa3, Bpifa1, Bpifa5
omitted Genes: Bpifb1, Bpifb5, Bpifb9a
omitted Genes: Bpifb9b, Cdk5rap1, Cbfa2t2
omitted Genes: 1700003F12Rik, Pxmp4, Zfp341
omitted Genes: Chmp4b, Raly, Eif2s2
omitted Genes: Ahcy, 2310005A03Rik, Map1lc3a
omitted Genes: Mir695, Trp53inp2, Ncoa6
omitted Genes: Acss2, Acss2os, Gss

Make more plots at http://csg.sph.umich.edu/locuszoom/

omitted Genes: Myh7b, Mir499, Trpc4ap

Fine−mapping Regions

| snp | chr | pos (Mb) | pp | group | color |
|---|---|---|---|---|---|
| chr2:154464466 | 2 | 154.4645 | 0.9982817 | Credible Set | purple |

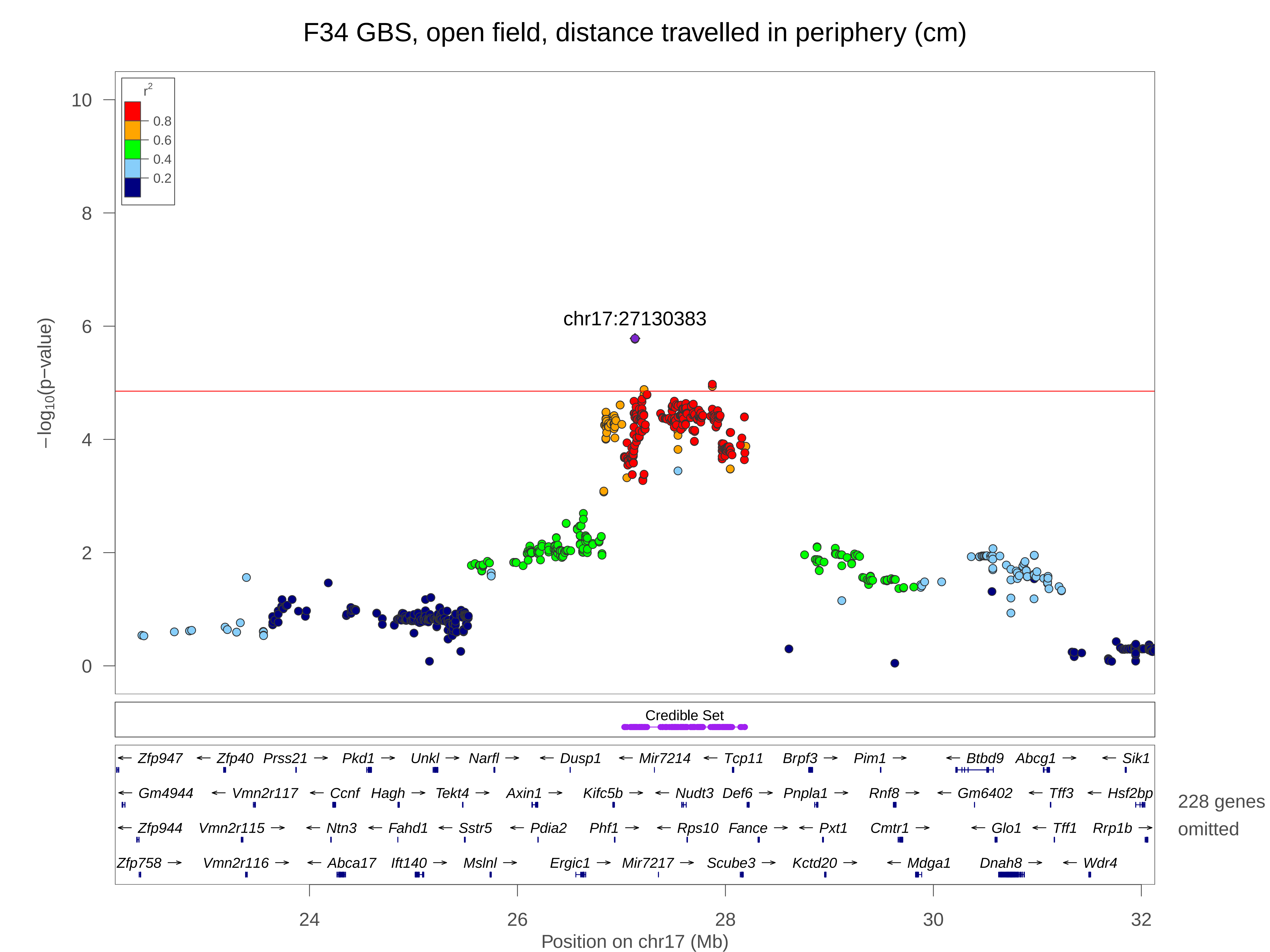

F34 GBS, open field, distance travelled in periphery (cm)
r2
0.8
0.6
0.4
0.2
10
8
6
4
2
0
−log10(p−value)
chr17:27130383
Credible Set
← Zfp947 ← Zfp40 Prss21 → Pkd1 → Unkl → Narfl → ← Dusp1 ← Mir7214 ← Tcp11 Brpf3 → Pim1 → ← Btbd9 Abcg1 → ← Sik1
← Gm4944 ← Vmn2r117 ← Ccnf Hagh → Tekt4 → Axin1 → Kifc5b → ← Nudt3 Def6 → Pnpla1 → Rnf8 → ← Gm6402 ← Tff3 ← Hsf2bp
← Zfp944 Vmn2r115 → ← Ntn3 ← Fahd1 ← Sstr5 ← Pdia2 Phf1 → ← Rps10 Fance → ← Pxt1 Cmtr1 → ← Glo1 ← Tff1 Rrp1b →
Zfp758 → Vmn2r116 → ← Abca17 Ift140 → Mslnl → Ergic1 → Mir7217 → Scube3 → Kctd20 → ← Mdga1 Dnah8 → ← Wdr4
228 genes omitted
24
26
28
30
32
Position on chr17 (Mb)

date: Thu Oct 25 17:13:54 2018
build: F34GBS
display range: chr17:22130383−32130383 [22130383−32130383]
hilite range: 0 − 0 [ 0 − 0 ]
reference SNP: chr17:27130383
number of SNPs plotted: 975
min P−value: 1.65E−6 [chr17:27130383]
max P−value: 8.98E−1 [chr17:29629020]
omitted Genes: Zfp946, Vmn2r111, Vmn2r112
omitted Genes: Gm16386, Zfp945, Vmn2r113
omitted Genes: Vmn2r−ps129, Vmn2r114, 6330415G19Rik
omitted Genes: Zfp213, Zfp13, Zscan10
omitted Genes: Mmp25, Ccdc64b, Thoc6
omitted Genes: Hcfc1r1, Tnfrsf12a, Cldn6
omitted Genes: Cldn9, Pkmyt1, Paqr4
omitted Genes: Kremen2, 9530082P21Rik, Flywch1
omitted Genes: Flywch2, Srrm2, Mir5125
omitted Genes: Tceb2, Prss33, Prss41
omitted Genes: Prss32, Dcpp1, Dcpp2
omitted Genes: Dcpp3, Sbp, Sbpl
omitted Genes: Prss30, Prss22, Prss27
omitted Genes: Kctd5, Pdpk1, Amdhd2
omitted Genes: Atp6v0c, Tbc1d24, 1600002H07Rik
omitted Genes: Mir5134, Mir6965, Abca3
omitted Genes: D330041H03Rik, Rnps1, Eci1
omitted Genes: Dnase1l2, E4f1, Pgp
omitted Genes: Mlst8, Bricd5, Caskin1
omitted Genes: Traf7, Rab26os, Rab26
omitted Genes: Tsc2, Nthl1, Slc9a3r2
omitted Genes: Npw, Zfp598, Syngr3
omitted Genes: Gfer, Noxo1, Tbl3
omitted Genes: Rnf151, Snhg9, Snora78
omitted Genes: Rps2, Snora64, Ndufb10
omitted Genes: Rpl3l, Msrb1, Hs3st6
omitted Genes: Meiob, Igfals, Nubp2
omitted Genes: Spsb3, Eme2, Mrps34
omitted Genes: Nme3, Mapk8ip3, Hn1l
omitted Genes: Cramp1l, Tmem204, Telo2
omitted Genes: Ptx4, Clcn7, Ccdc154
Make more plots at http://csg.sph.umich.edu/locuszoom/
omitted Genes: BC003965, Gnptg, Tsr3

Fine−mapping Regions

| snp | chr | pos (Mb) | pp | group | color |
|---|---|---|---|---|---|
| chr17:27028210 | 17 | 27.02821 | 0.0006905196 | Credible Set | purple |
| chr17:27028984 | 17 | 27.02898 | 0.0006622292 | Credible Set | purple |
| chr17:27035894 | 17 | 27.03589 | 0.0006679480 | Credible Set | purple |
| chr17:27038183 | 17 | 27.03818 | 0.0006708332 | Credible Set | purple |
| chr17:27053285 | 17 | 27.05328 | 0.0011704250 | Credible Set | purple |
| chr17:27083567 | 17 | 27.08357 | 0.0007396722 | Credible Set | purple |
| chr17:27090253 | 17 | 27.09025 | 0.0006679941 | Credible Set | purple |
| chr17:27090308 | 17 | 27.09031 | 0.0006625982 | Credible Set | purple |
| chr17:27092250 | 17 | 27.09225 | 0.0006679236 | Credible Set | purple |
| chr17:27099867 | 17 | 27.09987 | 0.0009607726 | Credible Set | purple |
| chr17:27100129 | 17 | 27.10013 | 0.0007180446 | Credible Set | purple |
| chr17:27104557 | 17 | 27.10456 | 0.0006643762 | Credible Set | purple |
| chr17:27114211 | 17 | 27.11421 | 0.0007016296 | Credible Set | purple |
| chr17:27114427 | 17 | 27.11443 | 0.0007430355 | Credible Set | purple |
| chr17:27119229 | 17 | 27.11923 | 0.0009431507 | Credible Set | purple |
| chr17:27119262 | 17 | 27.11926 | 0.0009350070 | Credible Set | purple |
| chr17:27119650 | 17 | 27.11965 | 0.0016022245 | Credible Set | purple |
| chr17:27119751 | 17 | 27.11975 | 0.0008576135 | Credible Set | purple |
| chr17:27119767 | 17 | 27.11977 | 0.0008532212 | Credible Set | purple |
| chr17:27119789 | 17 | 27.11979 | 0.0008576565 | Credible Set | purple |
| chr17:27120639 | 17 | 27.12064 | 0.0021446983 | Credible Set | purple |
| chr17:27120789 | 17 | 27.12079 | 0.0058109205 | Credible Set | purple |
| chr17:27120982 | 17 | 27.12098 | 0.0036397830 | Credible Set | purple |
| chr17:27124710 | 17 | 27.12471 | 0.0036067340 | Credible Set | purple |
| chr17:27127355 | 17 | 27.12736 | 0.0010606908 | Credible Set | purple |
| chr17:27129483 | 17 | 27.12948 | 0.0031674687 | Credible Set | purple |
| chr17:27129530 | 17 | 27.12953 | 0.0031098902 | Credible Set | purple |
| chr17:27130351 | 17 | 27.13035 | 0.0655034163 | Credible Set | purple |
| chr17:27130379 | 17 | 27.13038 | 0.0650354071 | Credible Set | purple |
| chr17:27130383 | 17 | 27.13038 | 0.0668860613 | Credible Set | purple |
| chr17:27130402 | 17 | 27.13040 | 0.0663849614 | Credible Set | purple |
| chr17:27139266 | 17 | 27.13927 | 0.0035516860 | Credible Set | purple |
| chr17:27140390 | 17 | 27.14039 | 0.0046340897 | Credible Set | purple |
| chr17:27143270 | 17 | 27.14327 | 0.0035863041 | Credible Set | purple |
| chr17:27143279 | 17 | 27.14328 | 0.0035118431 | Credible Set | purple |
| chr17:27143309 | 17 | 27.14331 | 0.0035118431 | Credible Set | purple |

Fine−mapping Regions

| snp | chr | pos (Mb) | pp | group | color |
|---|---|---|---|---|---|
| chr17:27143399 | 17 | 27.14340 | 0.003172901 | Credible Set | purple |
| chr17:27144798 | 17 | 27.14480 | 0.001214051 | Credible Set | purple |
| chr17:27146163 | 17 | 27.14616 | 0.002860352 | Credible Set | purple |
| chr17:27146360 | 17 | 27.14636 | 0.001412914 | Credible Set | purple |
| chr17:27146370 | 17 | 27.14637 | 0.001412914 | Credible Set | purple |
| chr17:27156143 | 17 | 27.15614 | 0.002889728 | Credible Set | purple |
| chr17:27171213 | 17 | 27.17121 | 0.004283235 | Credible Set | purple |
| chr17:27171685 | 17 | 27.17169 | 0.001593529 | Credible Set | purple |
| chr17:27171811 | 17 | 27.17181 | 0.001441306 | Credible Set | purple |
| chr17:27171813 | 17 | 27.17181 | 0.001480138 | Credible Set | purple |
| chr17:27172497 | 17 | 27.17250 | 0.001865232 | Credible Set | purple |
| chr17:27174542 | 17 | 27.17454 | 0.002877318 | Credible Set | purple |
| chr17:27184013 | 17 | 27.18401 | 0.003011507 | Credible Set | purple |
| chr17:27188020 | 17 | 27.18802 | 0.003018626 | Credible Set | purple |
| chr17:27188575 | 17 | 27.18858 | 0.002968736 | Credible Set | purple |
| chr17:27191517 | 17 | 27.19152 | 0.003559769 | Credible Set | purple |
| chr17:27191542 | 17 | 27.19154 | 0.003689981 | Credible Set | purple |
| chr17:27191945 | 17 | 27.19195 | 0.003307282 | Credible Set | purple |
| chr17:27193635 | 17 | 27.19364 | 0.003334340 | Credible Set | purple |
| chr17:27194000 | 17 | 27.19400 | 0.003052373 | Credible Set | purple |
| chr17:27195141 | 17 | 27.19514 | 0.003586952 | Credible Set | purple |
| chr17:27195293 | 17 | 27.19529 | 0.002622497 | Credible Set | purple |
| chr17:27195309 | 17 | 27.19531 | 0.002565096 | Credible Set | purple |
| chr17:27195731 | 17 | 27.19573 | 0.003445560 | Credible Set | purple |
| chr17:27195769 | 17 | 27.19577 | 0.003557021 | Credible Set | purple |
| chr17:27196042 | 17 | 27.19604 | 0.003417150 | Credible Set | purple |
| chr17:27197618 | 17 | 27.19762 | 0.003400012 | Credible Set | purple |
| chr17:27197752 | 17 | 27.19775 | 0.005670945 | Credible Set | purple |
| chr17:27197791 | 17 | 27.19779 | 0.006310012 | Credible Set | purple |
| chr17:27197888 | 17 | 27.19789 | 0.004344106 | Credible Set | purple |
| chr17:27200193 | 17 | 27.20019 | 0.001802654 | Credible Set | purple |
| chr17:27200651 | 17 | 27.20065 | 0.003594198 | Credible Set | purple |
| chr17:27215443 | 17 | 27.21544 | 0.003497046 | Credible Set | purple |
| chr17:27215484 | 17 | 27.21548 | 0.003359707 | Credible Set | purple |
| chr17:27223950 | 17 | 27.22395 | 0.001996354 | Credible Set | purple |
| chr17:27224285 | 17 | 27.22428 | 0.001976200 | Credible Set | purple |

Fine−mapping Regions

| snp | chr | pos (Mb) | pp | group | color |
|---|---|---|---|---|---|
| chr17:27227746 | 17 | 27.22775 | 0.001980280 | Credible Set | purple |
| chr17:27228977 | 17 | 27.22898 | 0.001973063 | Credible Set | purple |
| chr17:27230139 | 17 | 27.23014 | 0.002345358 | Credible Set | purple |
| chr17:27245666 | 17 | 27.24567 | 0.007424828 | Credible Set | purple |
| chr17:27245737 | 17 | 27.24574 | 0.007536836 | Credible Set | purple |
| chr17:27376234 | 17 | 27.37623 | 0.003618079 | Credible Set | purple |
| chr17:27393680 | 17 | 27.39368 | 0.003122487 | Credible Set | purple |
| chr17:27397461 | 17 | 27.39746 | 0.003069704 | Credible Set | purple |
| chr17:27397469 | 17 | 27.39747 | 0.003016427 | Credible Set | purple |
| chr17:27419536 | 17 | 27.41954 | 0.002970358 | Credible Set | purple |
| chr17:27428199 | 17 | 27.42820 | 0.002948887 | Credible Set | purple |
| chr17:27428246 | 17 | 27.42825 | 0.002947730 | Credible Set | purple |
| chr17:27437777 | 17 | 27.43778 | 0.002896468 | Credible Set | purple |
| chr17:27459951 | 17 | 27.45995 | 0.003030526 | Credible Set | purple |
| chr17:27475870 | 17 | 27.47587 | 0.002657809 | Credible Set | purple |
| chr17:27486422 | 17 | 27.48642 | 0.002808491 | Credible Set | purple |
| chr17:27486466 | 17 | 27.48647 | 0.002782808 | Credible Set | purple |
| chr17:27486535 | 17 | 27.48653 | 0.002937368 | Credible Set | purple |
| chr17:27486605 | 17 | 27.48661 | 0.002937368 | Credible Set | purple |
| chr17:27486642 | 17 | 27.48664 | 0.002937368 | Credible Set | purple |
| chr17:27486705 | 17 | 27.48671 | 0.002848904 | Credible Set | purple |
| chr17:27486720 | 17 | 27.48672 | 0.002939042 | Credible Set | purple |
| chr17:27486902 | 17 | 27.48690 | 0.003999480 | Credible Set | purple |
| chr17:27486957 | 17 | 27.48696 | 0.004749891 | Credible Set | purple |
| chr17:27492720 | 17 | 27.49272 | 0.004829780 | Credible Set | purple |
| chr17:27492775 | 17 | 27.49278 | 0.004951286 | Credible Set | purple |
| chr17:27507520 | 17 | 27.50752 | 0.002705067 | Credible Set | purple |
| chr17:27507982 | 17 | 27.50798 | 0.002902714 | Credible Set | purple |
| chr17:27508332 | 17 | 27.50833 | 0.002136315 | Credible Set | purple |
| chr17:27509007 | 17 | 27.50901 | 0.005797119 | Credible Set | purple |
| chr17:27509013 | 17 | 27.50901 | 0.005791401 | Credible Set | purple |
| chr17:27509015 | 17 | 27.50902 | 0.005835558 | Credible Set | purple |
| chr17:27509253 | 17 | 27.50925 | 0.003007173 | Credible Set | purple |
| chr17:27517180 | 17 | 27.51718 | 0.002751952 | Credible Set | purple |
| chr17:27517282 | 17 | 27.51728 | 0.004603356 | Credible Set | purple |
| chr17:27517332 | 17 | 27.51733 | 0.004587039 | Credible Set | purple |

Fine−mapping Regions

| snp | chr | pos (Mb) | pp | group | color |
|---|---|---|---|---|---|
| chr17:27520735 | 17 | 27.52073 | 0.002748236 | Credible Set | purple |
| chr17:27528952 | 17 | 27.52895 | 0.005080409 | Credible Set | purple |
| chr17:27529039 | 17 | 27.52904 | 0.002319879 | Credible Set | purple |
| chr17:27543951 | 17 | 27.54395 | 0.004917564 | Credible Set | purple |
| chr17:27543956 | 17 | 27.54396 | 0.004930713 | Credible Set | purple |
| chr17:27543999 | 17 | 27.54400 | 0.004880749 | Credible Set | purple |
| chr17:27544604 | 17 | 27.54460 | 0.003168622 | Credible Set | purple |
| chr17:27564093 | 17 | 27.56409 | 0.003352451 | Credible Set | purple |
| chr17:27564105 | 17 | 27.56411 | 0.003352451 | Credible Set | purple |
| chr17:27571562 | 17 | 27.57156 | 0.003344775 | Credible Set | purple |
| chr17:27571636 | 17 | 27.57164 | 0.001950998 | Credible Set | purple |
| chr17:27571643 | 17 | 27.57164 | 0.001950998 | Credible Set | purple |
| chr17:27571700 | 17 | 27.57170 | 0.005012142 | Credible Set | purple |
| chr17:27572004 | 17 | 27.57200 | 0.003371118 | Credible Set | purple |
| chr17:27580805 | 17 | 27.58081 | 0.003349890 | Credible Set | purple |
| chr17:27581132 | 17 | 27.58113 | 0.003368510 | Credible Set | purple |
| chr17:27587053 | 17 | 27.58705 | 0.004286323 | Credible Set | purple |
| chr17:27587087 | 17 | 27.58709 | 0.004261357 | Credible Set | purple |
| chr17:27587178 | 17 | 27.58718 | 0.003875413 | Credible Set | purple |
| chr17:27587179 | 17 | 27.58718 | 0.003875413 | Credible Set | purple |
| chr17:27587180 | 17 | 27.58718 | 0.003875413 | Credible Set | purple |
| chr17:27587187 | 17 | 27.58719 | 0.003875413 | Credible Set | purple |
| chr17:27587188 | 17 | 27.58719 | 0.003875413 | Credible Set | purple |
| chr17:27587192 | 17 | 27.58719 | 0.003875413 | Credible Set | purple |
| chr17:27587496 | 17 | 27.58750 | 0.004352865 | Credible Set | purple |
| chr17:27587576 | 17 | 27.58758 | 0.002329053 | Credible Set | purple |
| chr17:27590733 | 17 | 27.59073 | 0.003468632 | Credible Set | purple |
| chr17:27590765 | 17 | 27.59077 | 0.003241280 | Credible Set | purple |
| chr17:27590778 | 17 | 27.59078 | 0.003351669 | Credible Set | purple |
| chr17:27590796 | 17 | 27.59080 | 0.003354960 | Credible Set | purple |
| chr17:27596229 | 17 | 27.59623 | 0.002935597 | Credible Set | purple |
| chr17:27596270 | 17 | 27.59627 | 0.002935317 | Credible Set | purple |
| chr17:27596658 | 17 | 27.59666 | 0.004164313 | Credible Set | purple |
| chr17:27596666 | 17 | 27.59667 | 0.004279920 | Credible Set | purple |
| chr17:27596669 | 17 | 27.59667 | 0.004412048 | Credible Set | purple |
| chr17:27597174 | 17 | 27.59717 | 0.004340178 | Credible Set | purple |

Fine−mapping Regions

| snp | chr | pos (Mb) | pp | group | color |
|---|---|---|---|---|---|
| chr17:27612703 | 17 | 27.61270 | 0.004305540 | Credible Set | purple |
| chr17:27613227 | 17 | 27.61323 | 0.004387607 | Credible Set | purple |
| chr17:27613228 | 17 | 27.61323 | 0.004394670 | Credible Set | purple |
| chr17:27616766 | 17 | 27.61677 | 0.002437642 | Credible Set | purple |
| chr17:27616769 | 17 | 27.61677 | 0.002408529 | Credible Set | purple |
| chr17:27617069 | 17 | 27.61707 | 0.002351613 | Credible Set | purple |
| chr17:27619823 | 17 | 27.61982 | 0.005331907 | Credible Set | purple |
| chr17:27621865 | 17 | 27.62186 | 0.004478968 | Credible Set | purple |
| chr17:27630316 | 17 | 27.63032 | 0.003688785 | Credible Set | purple |
| chr17:27630364 | 17 | 27.63036 | 0.003610154 | Credible Set | purple |
| chr17:27630797 | 17 | 27.63080 | 0.003566265 | Credible Set | purple |
| chr17:27665592 | 17 | 27.66559 | 0.003077830 | Credible Set | purple |
| chr17:27665593 | 17 | 27.66559 | 0.003083298 | Credible Set | purple |
| chr17:27665645 | 17 | 27.66565 | 0.003083298 | Credible Set | purple |
| chr17:27671877 | 17 | 27.67188 | 0.004880048 | Credible Set | purple |
| chr17:27671883 | 17 | 27.67188 | 0.004759104 | Credible Set | purple |
| chr17:27686117 | 17 | 27.68612 | 0.003396868 | Credible Set | purple |
| chr17:27686262 | 17 | 27.68626 | 0.001893958 | Credible Set | purple |
| chr17:27686263 | 17 | 27.68626 | 0.001893958 | Credible Set | purple |
| chr17:27690758 | 17 | 27.69076 | 0.003696990 | Credible Set | purple |
| chr17:27690904 | 17 | 27.69090 | 0.005126885 | Credible Set | purple |
| chr17:27690906 | 17 | 27.69091 | 0.005200503 | Credible Set | purple |
| chr17:27701425 | 17 | 27.70143 | 0.001227921 | Credible Set | purple |
| chr17:27701456 | 17 | 27.70146 | 0.001243876 | Credible Set | purple |
| chr17:27701864 | 17 | 27.70186 | 0.003349890 | Credible Set | purple |
| chr17:27702067 | 17 | 27.70207 | 0.003336597 | Credible Set | purple |
| chr17:27704331 | 17 | 27.70433 | 0.003499133 | Credible Set | purple |
| chr17:27704567 | 17 | 27.70457 | 0.001803896 | Credible Set | purple |
| chr17:27704630 | 17 | 27.70463 | 0.001888245 | Credible Set | purple |
| chr17:27728842 | 17 | 27.72884 | 0.002851367 | Credible Set | purple |
| chr17:27728885 | 17 | 27.72888 | 0.002934453 | Credible Set | purple |
| chr17:27728889 | 17 | 27.72889 | 0.002952041 | Credible Set | purple |
| chr17:27728897 | 17 | 27.72890 | 0.002934453 | Credible Set | purple |
| chr17:27743592 | 17 | 27.74359 | 0.003192428 | Credible Set | purple |
| chr17:27743637 | 17 | 27.74364 | 0.004117684 | Credible Set | purple |
| chr17:27748732 | 17 | 27.74873 | 0.003225522 | Credible Set | purple |

Fine−mapping Regions

| snp | chr | pos (Mb) | pp | group | color |
|---|---|---|---|---|---|
| chr17:27748758 | 17 | 27.74876 | 0.003377620 | Credible Set | purple |
| chr17:27761092 | 17 | 27.76109 | 0.003389158 | Credible Set | purple |
| chr17:27761141 | 17 | 27.76114 | 0.003367948 | Credible Set | purple |
| chr17:27761632 | 17 | 27.76163 | 0.003350055 | Credible Set | purple |
| chr17:27762683 | 17 | 27.76268 | 0.003481436 | Credible Set | purple |
| chr17:27762853 | 17 | 27.76285 | 0.003122038 | Credible Set | purple |
| chr17:27763066 | 17 | 27.76307 | 0.003682703 | Credible Set | purple |
| chr17:27763276 | 17 | 27.76328 | 0.002590740 | Credible Set | purple |
| chr17:27771695 | 17 | 27.77170 | 0.003067641 | Credible Set | purple |
| chr17:27778534 | 17 | 27.77853 | 0.003265639 | Credible Set | purple |
| chr17:27782597 | 17 | 27.78260 | 0.003343349 | Credible Set | purple |
| chr17:27782890 | 17 | 27.78289 | 0.003344144 | Credible Set | purple |
| chr17:27782953 | 17 | 27.78295 | 0.003344144 | Credible Set | purple |
| chr17:27856821 | 17 | 27.85682 | 0.003255148 | Credible Set | purple |
| chr17:27874099 | 17 | 27.87410 | 0.004287132 | Credible Set | purple |
| chr17:27874161 | 17 | 27.87416 | 0.011245136 | Credible Set | purple |
| chr17:27878639 | 17 | 27.87864 | 0.003368408 | Credible Set | purple |
| chr17:27889423 | 17 | 27.88942 | 0.002781855 | Credible Set | purple |
| chr17:27889454 | 17 | 27.88945 | 0.002783606 | Credible Set | purple |
| chr17:27894408 | 17 | 27.89441 | 0.002759443 | Credible Set | purple |
| chr17:27894461 | 17 | 27.89446 | 0.002743438 | Credible Set | purple |
| chr17:27894647 | 17 | 27.89465 | 0.003489548 | Credible Set | purple |
| chr17:27894825 | 17 | 27.89483 | 0.003009436 | Credible Set | purple |
| chr17:27895648 | 17 | 27.89565 | 0.003226592 | Credible Set | purple |
| chr17:27908870 | 17 | 27.90887 | 0.003346538 | Credible Set | purple |
| chr17:27909014 | 17 | 27.90901 | 0.002139331 | Credible Set | purple |
| chr17:27909047 | 17 | 27.90905 | 0.002137497 | Credible Set | purple |
| chr17:27909083 | 17 | 27.90908 | 0.002154310 | Credible Set | purple |
| chr17:27924683 | 17 | 27.92468 | 0.002380880 | Credible Set | purple |
| chr17:27924726 | 17 | 27.92473 | 0.002423893 | Credible Set | purple |
| chr17:27925657 | 17 | 27.92566 | 0.004041239 | Credible Set | purple |
| chr17:27925757 | 17 | 27.92576 | 0.003349890 | Credible Set | purple |
| chr17:27925847 | 17 | 27.92585 | 0.003349890 | Credible Set | purple |
| chr17:27929518 | 17 | 27.92952 | 0.003212652 | Credible Set | purple |
| chr17:27938148 | 17 | 27.93815 | 0.003001811 | Credible Set | purple |
| chr17:27945251 | 17 | 27.94525 | 0.003188269 | Credible Set | purple |

Fine−mapping Regions

| snp | chr | pos (Mb) | pp | group | color |
|---|---|---|---|---|---|
| chr17:27945722 | 17 | 27.94572 | 0.0033441442 | Credible Set | purple |
| chr17:27945734 | 17 | 27.94573 | 0.0033441442 | Credible Set | purple |
| chr17:27945792 | 17 | 27.94579 | 0.0033498899 | Credible Set | purple |
| chr17:27949604 | 17 | 27.94960 | 0.0033498899 | Credible Set | purple |
| chr17:27967905 | 17 | 27.96790 | 0.0008648146 | Credible Set | purple |
| chr17:27968095 | 17 | 27.96810 | 0.0011365013 | Credible Set | purple |
| chr17:27968486 | 17 | 27.96849 | 0.0006912941 | Credible Set | purple |
| chr17:27969063 | 17 | 27.96906 | 0.0009828777 | Credible Set | purple |
| chr17:27978189 | 17 | 27.97819 | 0.0009987248 | Credible Set | purple |
| chr17:27978217 | 17 | 27.97822 | 0.0009987248 | Credible Set | purple |
| chr17:27978305 | 17 | 27.97830 | 0.0008644625 | Credible Set | purple |
| chr17:27982311 | 17 | 27.98231 | 0.0011086752 | Credible Set | purple |
| chr17:27982320 | 17 | 27.98232 | 0.0011252062 | Credible Set | purple |
| chr17:27986901 | 17 | 27.98690 | 0.0009202298 | Credible Set | purple |
| chr17:27996164 | 17 | 27.99616 | 0.0009316572 | Credible Set | purple |
| chr17:27996409 | 17 | 27.99641 | 0.0007122434 | Credible Set | purple |
| chr17:27996433 | 17 | 27.99643 | 0.0007061831 | Credible Set | purple |
| chr17:27998199 | 17 | 27.99820 | 0.0009372187 | Credible Set | purple |
| chr17:28005852 | 17 | 28.00585 | 0.0008597242 | Credible Set | purple |
| chr17:28006277 | 17 | 28.00628 | 0.0009220173 | Credible Set | purple |
| chr17:28009409 | 17 | 28.00941 | 0.0009203198 | Credible Set | purple |
| chr17:28013892 | 17 | 28.01389 | 0.0008496057 | Credible Set | purple |
| chr17:28016695 | 17 | 28.01669 | 0.0009987248 | Credible Set | purple |
| chr17:28016697 | 17 | 28.01670 | 0.0009987248 | Credible Set | purple |
| chr17:28016702 | 17 | 28.01670 | 0.0009987248 | Credible Set | purple |
| chr17:28016770 | 17 | 28.01677 | 0.0009653182 | Credible Set | purple |
| chr17:28023882 | 17 | 28.02388 | 0.0008816039 | Credible Set | purple |
| chr17:28024097 | 17 | 28.02410 | 0.0010087990 | Credible Set | purple |
| chr17:28026770 | 17 | 28.02677 | 0.0009805809 | Credible Set | purple |
| chr17:28036756 | 17 | 28.03676 | 0.0010088315 | Credible Set | purple |
| chr17:28037425 | 17 | 28.03742 | 0.0008502947 | Credible Set | purple |
| chr17:28046103 | 17 | 28.04610 | 0.0017489299 | Credible Set | purple |
| chr17:28046109 | 17 | 28.04611 | 0.0017300678 | Credible Set | purple |
| chr17:28046746 | 17 | 28.04675 | 0.0008583626 | Credible Set | purple |
| chr17:28046753 | 17 | 28.04675 | 0.0008987162 | Credible Set | purple |
| chr17:28050079 | 17 | 28.05008 | 0.0008890477 | Credible Set | purple |

Fine−mapping Regions

| snp | chr | pos (Mb) | pp | group | color |
|---|---|---|---|---|---|
| chr17:28050266 | 17 | 28.05027 | 0.0007842086 | Credible Set | purple |
| chr17:28050607 | 17 | 28.05061 | 0.0017435338 | Credible Set | purple |
| chr17:28063789 | 17 | 28.06379 | 0.0007356959 | Credible Set | purple |
| chr17:28144309 | 17 | 28.14431 | 0.0010739984 | Credible Set | purple |
| chr17:28157995 | 17 | 28.15799 | 0.0014091523 | Credible Set | purple |
| chr17:28182743 | 17 | 28.18274 | 0.0031813313 | Credible Set | purple |
| chr17:28182775 | 17 | 28.18277 | 0.0031455848 | Credible Set | purple |
| chr17:28186934 | 17 | 28.18693 | 0.0007944738 | Credible Set | purple |
| chr17:28186976 | 17 | 28.18698 | 0.0007971036 | Credible Set | purple |

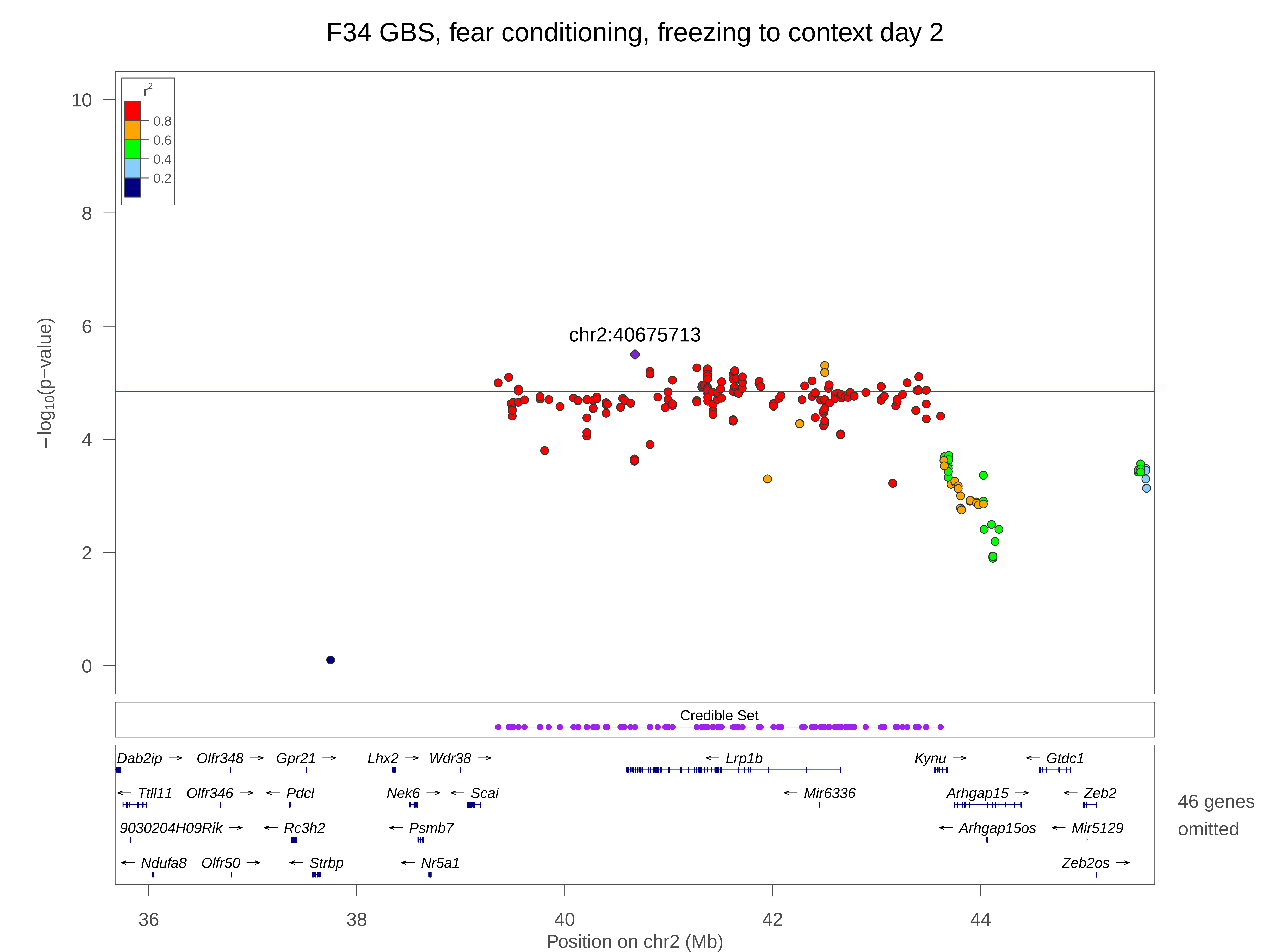
F34 GBS, fear conditioning, freezing to context day 2
r2
0.8
0.6
0.4
0.2
10
8
6
4
2
0
−log10(p−value)
chr2:40675713
Credible Set
Dab2ip
Olfr348
Gpr21
Lhx2
Wdr38
Lrp1b
Kynu
Gtdc1
Ttll1
Olfr346
Pdcl
Nek6
Scai
Mir6336
Arhgap15
Zeb2
9030204H09Rik
Rc3h2
Psmb7
Arhgap15os
Mir5129
Ndufa8
Olfr50
Strbp
Nr5a1
Zeb2os
46 genes omitted
36
38
40
42
44
Position on chr2 (Mb)

| | |
|---|---|
| date: | Thu Oct 25 17:14:07 2018 |
| build: | F34GBS |
| display range: | chr2:35675713−45675713 [35675713−45675713] |
| hilite range: | 0 − 0 [ 0 − 0 ] |
| reference SNP: | chr2:40675713 |
| number of SNPs plotted: | 274 |
| min P−value: | 3.16E−6 [chr2:40675713] |
| max P−value: | 7.84E−1 [chr2:37748995] |
| omitted Genes: | Morn5, Lhx6, Rbm18 |
| omitted Genes: | Mrrf, Ptgs1, Olfr338 |
| omitted Genes: | Olfr339, Olfr340, Olfr341 |
| omitted Genes: | Olfr342, Olfr344, Olfr345 |
| omitted Genes: | Olfr347, Olfr3, Olfr350 |
| omitted Genes: | Olfr351, Olfr352, Olfr353 |
| omitted Genes: | Olfr354, Olfr355, Olfr356 |
| omitted Genes: | Olfr357, Olfr358, Olfr360 |
| omitted Genes: | Olfr361, Olfr362, Olfr365 |
| omitted Genes: | Olfr366, Olfr367−ps, Olfr368 |
| omitted Genes: | Snord90, Zbtb6, Zbtb26 |
| omitted Genes: | Rabgap1, Mir5128, Crb2 |
| omitted Genes: | Dennd1a, Nr6a1, Mir181a−2 |
| omitted Genes: | Mir181b−2, Olfml2a, Rpl35 |
| omitted Genes: | Arpc5l, Golga1, Gm16523 |
| omitted Genes: | Ppp6c |

Make more plots at http://csg.sph.umich.edu/locuszoom/

Fine−mapping Regions

| snp | chr | pos (Mb) | pp | group | color |
|---|---|---|---|---|---|
| chr2:39359208 | 2 | 39.35921 | 0.007548244 | Credible Set | purple |
| chr2:39460505 | 2 | 39.46050 | 0.009372509 | Credible Set | purple |
| chr2:39460561 | 2 | 39.46056 | 0.009375488 | Credible Set | purple |
| chr2:39484371 | 2 | 39.48437 | 0.003359347 | Credible Set | purple |
| chr2:39493906 | 2 | 39.49391 | 0.002756126 | Credible Set | purple |
| chr2:39494797 | 2 | 39.49480 | 0.002087167 | Credible Set | purple |
| chr2:39494848 | 2 | 39.49485 | 0.002597478 | Credible Set | purple |
| chr2:39494860 | 2 | 39.49486 | 0.002591278 | Credible Set | purple |
| chr2:39494876 | 2 | 39.49488 | 0.002597478 | Credible Set | purple |
| chr2:39505733 | 2 | 39.50573 | 0.003555626 | Credible Set | purple |
| chr2:39505816 | 2 | 39.50582 | 0.003554107 | Credible Set | purple |
| chr2:39554087 | 2 | 39.55409 | 0.003557820 | Credible Set | purple |
| chr2:39554255 | 2 | 39.55425 | 0.005954087 | Credible Set | purple |
| chr2:39554302 | 2 | 39.55430 | 0.005469651 | Credible Set | purple |
| chr2:39554330 | 2 | 39.55433 | 0.005477828 | Credible Set | purple |
| chr2:39612884 | 2 | 39.61288 | 0.003911576 | Credible Set | purple |
| chr2:39762768 | 2 | 39.76277 | 0.004035344 | Credible Set | purple |
| chr2:39762904 | 2 | 39.76290 | 0.004457605 | Credible Set | purple |
| chr2:39847233 | 2 | 39.84723 | 0.003951008 | Credible Set | purple |
| chr2:39954028 | 2 | 39.95403 | 0.003014100 | Credible Set | purple |
| chr2:40082594 | 2 | 40.08259 | 0.004188936 | Credible Set | purple |
| chr2:40127679 | 2 | 40.12768 | 0.003760456 | Credible Set | purple |
| chr2:40127769 | 2 | 40.12777 | 0.003834860 | Credible Set | purple |
| chr2:40127812 | 2 | 40.12781 | 0.003834860 | Credible Set | purple |
| chr2:40212829 | 2 | 40.21283 | 0.001945604 | Credible Set | purple |
| chr2:40213202 | 2 | 40.21320 | 0.003902104 | Credible Set | purple |
| chr2:40213736 | 2 | 40.21374 | 0.003976946 | Credible Set | purple |
| chr2:40273583 | 2 | 40.27358 | 0.003834860 | Credible Set | purple |
| chr2:40273640 | 2 | 40.27364 | 0.003834860 | Credible Set | purple |
| chr2:40273652 | 2 | 40.27365 | 0.003834860 | Credible Set | purple |
| chr2:40273690 | 2 | 40.27369 | 0.002784294 | Credible Set | purple |
| chr2:40273730 | 2 | 40.27373 | 0.002846471 | Credible Set | purple |
| chr2:40309721 | 2 | 40.30972 | 0.004378585 | Credible Set | purple |
| chr2:40309754 | 2 | 40.30975 | 0.004040999 | Credible Set | purple |
| chr2:40397532 | 2 | 40.39753 | 0.002335952 | Credible Set | purple |
| chr2:40397564 | 2 | 40.39756 | 0.003250080 | Credible Set | purple |

Fine−mapping Regions

| snp | chr | pos (Mb) | pp | group | color |
|---|---|---|---|---|---|
| chr2:40397616 | 2 | 40.39762 | 0.003511113 | Credible Set | purple |
| chr2:40410289 | 2 | 40.41029 | 0.003298775 | Credible Set | purple |
| chr2:40536999 | 2 | 40.53700 | 0.002959671 | Credible Set | purple |
| chr2:40537001 | 2 | 40.53700 | 0.002927420 | Credible Set | purple |
| chr2:40558241 | 2 | 40.55824 | 0.004128112 | Credible Set | purple |
| chr2:40558249 | 2 | 40.55825 | 0.004128112 | Credible Set | purple |
| chr2:40558252 | 2 | 40.55825 | 0.004126635 | Credible Set | purple |
| chr2:40576355 | 2 | 40.57635 | 0.003828971 | Credible Set | purple |
| chr2:40633282 | 2 | 40.63328 | 0.003418566 | Credible Set | purple |
| chr2:40633327 | 2 | 40.63333 | 0.003427448 | Credible Set | purple |
| chr2:40675713 | 2 | 40.67571 | 0.022796704 | Credible Set | purple |
| chr2:40819654 | 2 | 40.81965 | 0.011948699 | Credible Set | purple |
| chr2:40819655 | 2 | 40.81965 | 0.010590288 | Credible Set | purple |
| chr2:40819681 | 2 | 40.81968 | 0.010670779 | Credible Set | purple |
| chr2:40895540 | 2 | 40.89554 | 0.004354632 | Credible Set | purple |
| chr2:40966785 | 2 | 40.96679 | 0.002890867 | Credible Set | purple |
| chr2:40993125 | 2 | 40.99312 | 0.003964297 | Credible Set | purple |
| chr2:40993261 | 2 | 40.99326 | 0.005339714 | Credible Set | purple |
| chr2:40993264 | 2 | 40.99326 | 0.005300705 | Credible Set | purple |
| chr2:41034946 | 2 | 41.03495 | 0.003133902 | Credible Set | purple |
| chr2:41034956 | 2 | 41.03496 | 0.003370645 | Credible Set | purple |
| chr2:41035611 | 2 | 41.03561 | 0.008381406 | Credible Set | purple |
| chr2:41035612 | 2 | 41.03561 | 0.008381406 | Credible Set | purple |
| chr2:41035617 | 2 | 41.03562 | 0.008381406 | Credible Set | purple |
| chr2:41270329 | 2 | 41.27033 | 0.003800026 | Credible Set | purple |
| chr2:41270331 | 2 | 41.27033 | 0.003799849 | Credible Set | purple |
| chr2:41270370 | 2 | 41.27037 | 0.003596583 | Credible Set | purple |
| chr2:41270733 | 2 | 41.27073 | 0.013531669 | Credible Set | purple |
| chr2:41318161 | 2 | 41.31816 | 0.006420104 | Credible Set | purple |
| chr2:41325733 | 2 | 41.32573 | 0.006979490 | Credible Set | purple |
| chr2:41344260 | 2 | 41.34426 | 0.006809245 | Credible Set | purple |
| chr2:41373379 | 2 | 41.37338 | 0.012061739 | Credible Set | purple |
| chr2:41373395 | 2 | 41.37340 | 0.013062810 | Credible Set | purple |
| chr2:41373397 | 2 | 41.37340 | 0.010593303 | Credible Set | purple |
| chr2:41373418 | 2 | 41.37342 | 0.009852093 | Credible Set | purple |
| chr2:41373858 | 2 | 41.37386 | 0.008776557 | Credible Set | purple |

Fine−mapping Regions

| snp | chr | pos (Mb) | pp | group | color |
|---|---|---|---|---|---|
| chr2:41374907 | 2 | 41.37491 | 0.006334603 | Credible Set | purple |
| chr2:41374921 | 2 | 41.37492 | 0.006011556 | Credible Set | purple |
| chr2:41375382 | 2 | 41.37538 | 0.005007241 | Credible Set | purple |
| chr2:41375442 | 2 | 41.37544 | 0.005575538 | Credible Set | purple |
| chr2:41375546 | 2 | 41.37555 | 0.003718006 | Credible Set | purple |
| chr2:41375551 | 2 | 41.37555 | 0.004310690 | Credible Set | purple |
| chr2:41420007 | 2 | 41.42001 | 0.005285896 | Credible Set | purple |
| chr2:41426404 | 2 | 41.42640 | 0.002615679 | Credible Set | purple |
| chr2:41426414 | 2 | 41.42641 | 0.002614234 | Credible Set | purple |
| chr2:41426429 | 2 | 41.42643 | 0.003302609 | Credible Set | purple |
| chr2:41426432 | 2 | 41.42643 | 0.002520385 | Credible Set | purple |
| chr2:41426436 | 2 | 41.42644 | 0.002221909 | Credible Set | purple |
| chr2:41467642 | 2 | 41.46764 | 0.003955334 | Credible Set | purple |
| chr2:41468064 | 2 | 41.46806 | 0.004911632 | Credible Set | purple |
| chr2:41471314 | 2 | 41.47131 | 0.005155447 | Credible Set | purple |
| chr2:41498784 | 2 | 41.49878 | 0.005987205 | Credible Set | purple |
| chr2:41507457 | 2 | 41.50746 | 0.004159746 | Credible Set | purple |
| chr2:41508402 | 2 | 41.50840 | 0.007893868 | Credible Set | purple |
| chr2:41619931 | 2 | 41.61993 | 0.001714713 | Credible Set | purple |
| chr2:41619935 | 2 | 41.61993 | 0.001814831 | Credible Set | purple |
| chr2:41620001 | 2 | 41.62000 | 0.001783550 | Credible Set | purple |
| chr2:41622012 | 2 | 41.62201 | 0.005378711 | Credible Set | purple |
| chr2:41622721 | 2 | 41.62272 | 0.008975935 | Credible Set | purple |
| chr2:41622751 | 2 | 41.62275 | 0.010758238 | Credible Set | purple |
| chr2:41622763 | 2 | 41.62276 | 0.008683312 | Credible Set | purple |
| chr2:41633469 | 2 | 41.63347 | 0.006472353 | Credible Set | purple |
| chr2:41633904 | 2 | 41.63390 | 0.012039397 | Credible Set | purple |
| chr2:41633967 | 2 | 41.63397 | 0.011723121 | Credible Set | purple |
| chr2:41633971 | 2 | 41.63397 | 0.012185559 | Credible Set | purple |
| chr2:41652829 | 2 | 41.65283 | 0.006009589 | Credible Set | purple |
| chr2:41653372 | 2 | 41.65337 | 0.008902364 | Credible Set | purple |
| chr2:41661623 | 2 | 41.66162 | 0.005124683 | Credible Set | purple |
| chr2:41666380 | 2 | 41.66638 | 0.005305685 | Credible Set | purple |
| chr2:41672293 | 2 | 41.67229 | 0.005022459 | Credible Set | purple |
| chr2:41672310 | 2 | 41.67231 | 0.005022459 | Credible Set | purple |
| chr2:41672317 | 2 | 41.67232 | 0.005022459 | Credible Set | purple |

Fine−mapping Regions

| snp | chr | pos (Mb) | pp | group | color |
|---|---|---|---|---|---|
| chr2:41706826 | 2 | 41.70683 | 0.006108973 | Credible Set | purple |
| chr2:41706866 | 2 | 41.70687 | 0.006094456 | Credible Set | purple |
| chr2:41707052 | 2 | 41.70705 | 0.007974998 | Credible Set | purple |
| chr2:41708898 | 2 | 41.70890 | 0.007697465 | Credible Set | purple |
| chr2:41708916 | 2 | 41.70892 | 0.007697465 | Credible Set | purple |
| chr2:41709000 | 2 | 41.70900 | 0.009530093 | Credible Set | purple |
| chr2:41868150 | 2 | 41.86815 | 0.007425278 | Credible Set | purple |
| chr2:41868765 | 2 | 41.86877 | 0.008051082 | Credible Set | purple |
| chr2:41884228 | 2 | 41.88423 | 0.006524157 | Credible Set | purple |
| chr2:41884645 | 2 | 41.88464 | 0.006507704 | Credible Set | purple |
| chr2:42008023 | 2 | 42.00802 | 0.003406870 | Credible Set | purple |
| chr2:42008027 | 2 | 42.00803 | 0.003288098 | Credible Set | purple |
| chr2:42008064 | 2 | 42.00806 | 0.003342753 | Credible Set | purple |
| chr2:42008225 | 2 | 42.00823 | 0.003054640 | Credible Set | purple |
| chr2:42008231 | 2 | 42.00823 | 0.003083756 | Credible Set | purple |
| chr2:42059626 | 2 | 42.05963 | 0.004146350 | Credible Set | purple |
| chr2:42080261 | 2 | 42.08026 | 0.004579822 | Credible Set | purple |
| chr2:42282211 | 2 | 42.28221 | 0.003923834 | Credible Set | purple |
| chr2:42307494 | 2 | 42.30749 | 0.006697496 | Credible Set | purple |
| chr2:42307501 | 2 | 42.30750 | 0.006733021 | Credible Set | purple |
| chr2:42379072 | 2 | 42.37907 | 0.008141966 | Credible Set | purple |
| chr2:42379092 | 2 | 42.37909 | 0.004466849 | Credible Set | purple |
| chr2:42409383 | 2 | 42.40938 | 0.001969462 | Credible Set | purple |
| chr2:42409544 | 2 | 42.40954 | 0.004994454 | Credible Set | purple |
| chr2:42409580 | 2 | 42.40958 | 0.005162732 | Credible Set | purple |
| chr2:42409609 | 2 | 42.40961 | 0.005165707 | Credible Set | purple |
| chr2:42461317 | 2 | 42.46132 | 0.003880943 | Credible Set | purple |
| chr2:42489186 | 2 | 42.48919 | 0.001444107 | Credible Set | purple |
| chr2:42489404 | 2 | 42.48940 | 0.002345778 | Credible Set | purple |
| chr2:42489412 | 2 | 42.48941 | 0.002554452 | Credible Set | purple |
| chr2:42500142 | 2 | 42.50014 | 0.003736525 | Credible Set | purple |
| chr2:42500166 | 2 | 42.50017 | 0.003950726 | Credible Set | purple |
| chr2:42500195 | 2 | 42.50019 | 0.003939746 | Credible Set | purple |
| chr2:42500470 | 2 | 42.50047 | 0.002793201 | Credible Set | purple |
| chr2:42500907 | 2 | 42.50091 | 0.001526252 | Credible Set | purple |
| chr2:42500955 | 2 | 42.50095 | 0.001733242 | Credible Set | purple |

Fine−mapping Regions

| snp | chr | pos (Mb) | pp | group | color |
|---|---|---|---|---|---|
| chr2:42535122 | 2 | 42.53512 | 0.006117644 | Credible Set | purple |
| chr2:42543271 | 2 | 42.54327 | 0.007025324 | Credible Set | purple |
| chr2:42547701 | 2 | 42.54770 | 0.003463079 | Credible Set | purple |
| chr2:42599483 | 2 | 42.59948 | 0.004837144 | Credible Set | purple |
| chr2:42599828 | 2 | 42.59983 | 0.004104847 | Credible Set | purple |
| chr2:42627072 | 2 | 42.62707 | 0.005084259 | Credible Set | purple |
| chr2:42652766 | 2 | 42.65277 | 0.004721332 | Credible Set | purple |
| chr2:42661197 | 2 | 42.66120 | 0.004217076 | Credible Set | purple |
| chr2:42661577 | 2 | 42.66158 | 0.004799634 | Credible Set | purple |
| chr2:42698140 | 2 | 42.69814 | 0.004414216 | Credible Set | purple |
| chr2:42724203 | 2 | 42.72420 | 0.004300084 | Credible Set | purple |
| chr2:42745376 | 2 | 42.74538 | 0.005243831 | Credible Set | purple |
| chr2:42781323 | 2 | 42.78132 | 0.004516269 | Credible Set | purple |
| chr2:42783202 | 2 | 42.78320 | 0.004580945 | Credible Set | purple |
| chr2:42783471 | 2 | 42.78347 | 0.004501612 | Credible Set | purple |
| chr2:42895360 | 2 | 42.89536 | 0.005193245 | Credible Set | purple |
| chr2:43043152 | 2 | 43.04315 | 0.003998648 | Credible Set | purple |
| chr2:43043214 | 2 | 43.04321 | 0.003852870 | Credible Set | purple |
| chr2:43043633 | 2 | 43.04363 | 0.006509506 | Credible Set | purple |
| chr2:43043675 | 2 | 43.04368 | 0.006607507 | Credible Set | purple |
| chr2:43043682 | 2 | 43.04368 | 0.006499695 | Credible Set | purple |
| chr2:43073408 | 2 | 43.07341 | 0.004494607 | Credible Set | purple |
| chr2:43183452 | 2 | 43.18345 | 0.003104119 | Credible Set | purple |
| chr2:43183517 | 2 | 43.18352 | 0.003102081 | Credible Set | purple |
| chr2:43196679 | 2 | 43.19668 | 0.003643281 | Credible Set | purple |
| chr2:43196709 | 2 | 43.19671 | 0.003624458 | Credible Set | purple |
| chr2:43196873 | 2 | 43.19687 | 0.003986405 | Credible Set | purple |
| chr2:43249424 | 2 | 43.24942 | 0.004822599 | Credible Set | purple |
| chr2:43292233 | 2 | 43.29223 | 0.007569541 | Credible Set | purple |
| chr2:43375733 | 2 | 43.37573 | 0.002581464 | Credible Set | purple |
| chr2:43375782 | 2 | 43.37578 | 0.002605897 | Credible Set | purple |
| chr2:43386699 | 2 | 43.38670 | 0.005667022 | Credible Set | purple |
| chr2:43399155 | 2 | 43.39916 | 0.005849502 | Credible Set | purple |
| chr2:43405381 | 2 | 43.40538 | 0.005670873 | Credible Set | purple |
| chr2:43405453 | 2 | 43.40545 | 0.009534120 | Credible Set | purple |
| chr2:43405489 | 2 | 43.40549 | 0.009630139 | Credible Set | purple |

Fine−mapping Regions

| snp | chr | pos (Mb) | pp | group | color |
|---|---|---|---|---|---|
| chr2:43476031 | 2 | 43.47603 | 0.003346023 | Credible Set | purple |
| chr2:43476032 | 2 | 43.47603 | 0.003329906 | Credible Set | purple |
| chr2:43476033 | 2 | 43.47603 | 0.003329906 | Credible Set | purple |
| chr2:43476096 | 2 | 43.47610 | 0.005667022 | Credible Set | purple |
| chr2:43476171 | 2 | 43.47617 | 0.001867513 | Credible Set | purple |
| chr2:43476172 | 2 | 43.47617 | 0.001867513 | Credible Set | purple |
| chr2:43476173 | 2 | 43.47617 | 0.001867513 | Credible Set | purple |
| chr2:43476189 | 2 | 43.47619 | 0.001867513 | Credible Set | purple |
| chr2:43476235 | 2 | 43.47624 | 0.005667022 | Credible Set | purple |
| chr2:43476254 | 2 | 43.47625 | 0.005667022 | Credible Set | purple |
| chr2:43476258 | 2 | 43.47626 | 0.005667022 | Credible Set | purple |
| chr2:43614807 | 2 | 43.61481 | 0.002081936 | Credible Set | purple |

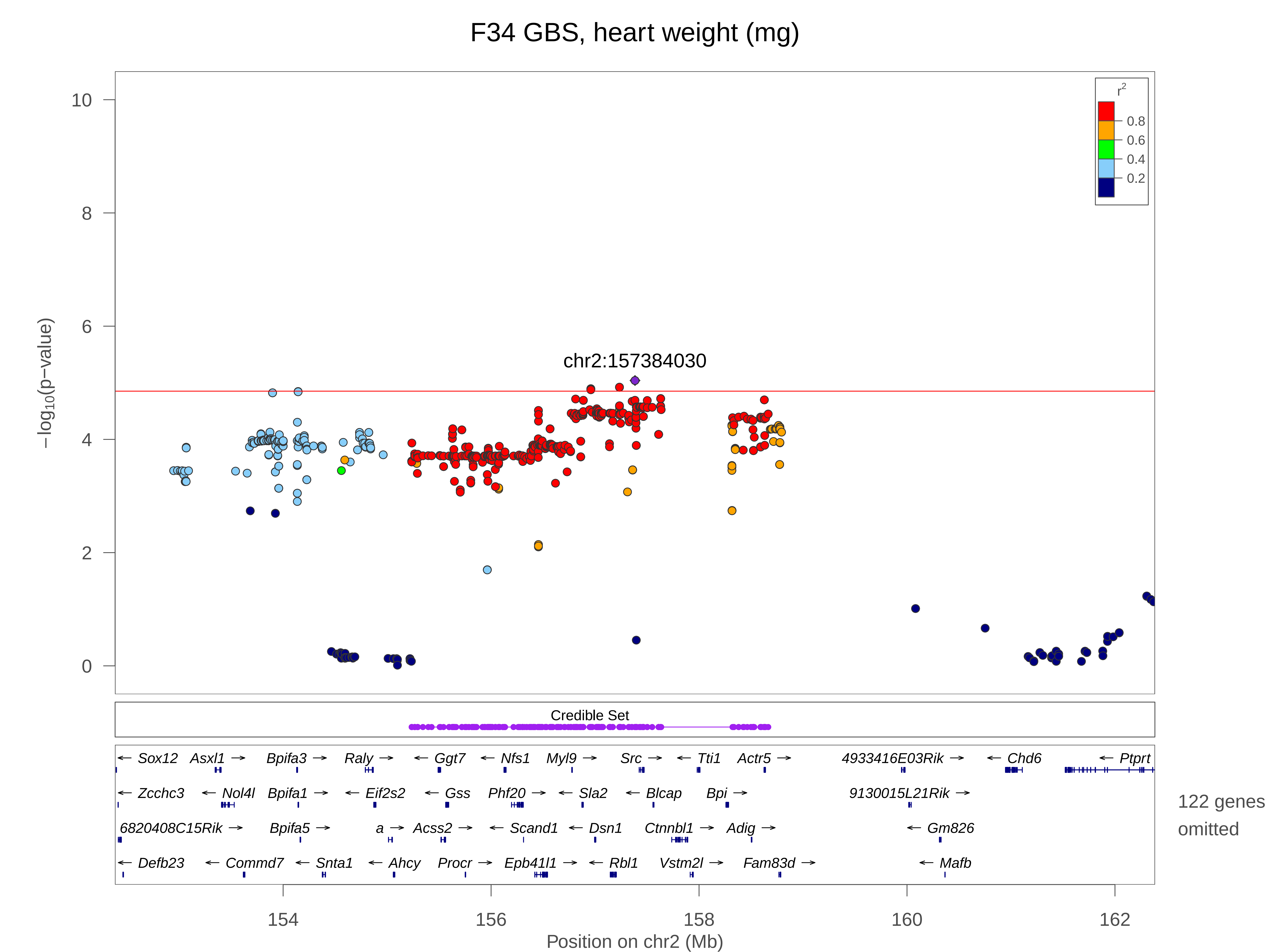

F34 GBS, heart weight (mg)
r2
0.8
0.6
0.4
0.2
chr2:157384030
−log10(p−value)
0
2
4
6
8
10
Credible Set
← Sox12
Asxl1 →
Bpifa3 →
Raly →
← Ggt7
← Nfs1
Myl9 →
Src →
← Tti1
Actr5 →
4933416E03Rik →
← Chd6
← Ptprt
← Zcchc3
← Nol4l
Bpifa1 →
← Eif2s2
← Gss
Phf20 →
← Sla2
← Blcap
Bpi →
9130015L21Rik →
6820408C15Rik →
Bpifa5 →
a →
Acss2 →
← Scand1
← Dsn1
Ctnnbl1 →
Adig →
← Gm826
← Defb23
← Commd7
← Snta1
← Ahcy
Procr →
Epb41l1 →
← Rbl1
Vstm2l →
Fam83d →
← Mafb
122 genes omitted
154
156
158
160
162
Position on chr2 (Mb)

date: Thu Oct 25 17:14:22 2018
build: F34GBS
display range: chr2:152384030−162384030 [152384030−162384030]
hilite range: 0 − 0 [ 0 − 0 ]
reference SNP: chr2:157384030
number of SNPs plotted: 729
min P−value: 9.12E−6 [chr2:157384030]
max P−value: 9.73E−1 [chr2:155099522]
omitted Genes: Defb20, Defb22, Defb26
omitted Genes: Defb28, Defb29, Defb21
omitted Genes: Defb19, Defb45, Defb36
omitted Genes: Defb25, Rem1, H13
omitted Genes: Mcts2, Id1, Cox4i2
omitted Genes: Bcl2l1, Tpx2, Mir5622
omitted Genes: Mylk2, Foxs1, Dusp15
omitted Genes: Ttll9, Pdrg1, Xkr7
omitted Genes: Ccm2l, Hck, Tm9sf4
omitted Genes: Tspyl3, Plagl2, Pofut1
omitted Genes: Kif3b, 2500004C02Rik, Dnmt3b
omitted Genes: Mapre1, Efcab8, Sun5
omitted Genes: Bpifb2, Bpifb6, Bpifb3
omitted Genes: Bpifb4, Bpifa6, Bpifa2
omitted Genes: Bpifb1, Bpifb5, Bpifb9a
omitted Genes: Bpifb9b, Cdk5rap1, Cbfa2t2
omitted Genes: Necab3, 1700003F12Rik, Actl10
omitted Genes: E2f1, Pxmp4, Zfp341
omitted Genes: Chmp4b, 2310005A03Rik, Itch
omitted Genes: Dynlrb1, Map1lc3a, Pigu
omitted Genes: Mir695, Trp53inp2, Ncoa6
omitted Genes: Acss2os, Myh7b, Mir499
omitted Genes: Trpc4ap, Edem2, Mmp24
omitted Genes: BC029722, Eif6, Fam83c
omitted Genes: Uqcc1, Gdf5, Cep250
omitted Genes: 6430550D23Rik, Ergic3, Fer1l4
omitted Genes: Spag4, Cpne1, Rbm12
omitted Genes: Romo1, Rbm39, Cnbd2
omitted Genes: 2900097C17Rik, Aar2, Gm14169
omitted Genes: Dlgap4, 4930405A21Rik, Tgif2
omitted Genes: 5430405H02Rik, 1110008F13Rik, Ndrg3
Make more plots at http://csg.sph.umich.edu/locuszoom/
omitted Genes: Soga1, Tldc2, Samhd1

Fine−mapping Regions

| snp | chr | pos (Mb) | pp | group | color |
|---|---|---|---|---|---|
| chr2:155236637 | 2 | 155.2366 | 0.0007199393 | Credible Set | purple |
| chr2:155236657 | 2 | 155.2367 | 0.0007136158 | Credible Set | purple |
| chr2:155236987 | 2 | 155.2370 | 0.0006939292 | Credible Set | purple |
| chr2:155237968 | 2 | 155.2380 | 0.0014328956 | Credible Set | purple |
| chr2:155264835 | 2 | 155.2648 | 0.0009474776 | Credible Set | purple |
| chr2:155265639 | 2 | 155.2656 | 0.0008729028 | Credible Set | purple |
| chr2:155266429 | 2 | 155.2664 | 0.0008427265 | Credible Set | purple |
| chr2:155266459 | 2 | 155.2665 | 0.0008720977 | Credible Set | purple |
| chr2:155291486 | 2 | 155.2915 | 0.0008908124 | Credible Set | purple |
| chr2:155294582 | 2 | 155.2946 | 0.0009336505 | Credible Set | purple |
| chr2:155294781 | 2 | 155.2948 | 0.0008242966 | Credible Set | purple |
| chr2:155294817 | 2 | 155.2948 | 0.0008071419 | Credible Set | purple |
| chr2:155344176 | 2 | 155.3442 | 0.0008784186 | Credible Set | purple |
| chr2:155396284 | 2 | 155.3963 | 0.0008830479 | Credible Set | purple |
| chr2:155427716 | 2 | 155.4277 | 0.0008786755 | Credible Set | purple |
| chr2:155506374 | 2 | 155.5064 | 0.0008902493 | Credible Set | purple |
| chr2:155514315 | 2 | 155.5143 | 0.0008729584 | Credible Set | purple |
| chr2:155542937 | 2 | 155.5429 | 0.0008682669 | Credible Set | purple |
| chr2:155596243 | 2 | 155.5962 | 0.0008729375 | Credible Set | purple |
| chr2:155596245 | 2 | 155.5962 | 0.0008729375 | Credible Set | purple |
| chr2:155596254 | 2 | 155.5963 | 0.0008729375 | Credible Set | purple |
| chr2:155626668 | 2 | 155.6267 | 0.0008729584 | Credible Set | purple |
| chr2:155626706 | 2 | 155.6267 | 0.0008729584 | Credible Set | purple |
| chr2:155626733 | 2 | 155.6267 | 0.0008284808 | Credible Set | purple |
| chr2:155626741 | 2 | 155.6267 | 0.0008240148 | Credible Set | purple |
| chr2:155627928 | 2 | 155.6279 | 0.0017096500 | Credible Set | purple |
| chr2:155629232 | 2 | 155.6292 | 0.0020257733 | Credible Set | purple |
| chr2:155629273 | 2 | 155.6293 | 0.0020068561 | Credible Set | purple |
| chr2:155629287 | 2 | 155.6293 | 0.0020151674 | Credible Set | purple |
| chr2:155629299 | 2 | 155.6293 | 0.0020162105 | Credible Set | purple |
| chr2:155630405 | 2 | 155.6304 | 0.0008729584 | Credible Set | purple |
| chr2:155630630 | 2 | 155.6306 | 0.0008729584 | Credible Set | purple |
| chr2:155630746 | 2 | 155.6307 | 0.0008729584 | Credible Set | purple |
| chr2:155631110 | 2 | 155.6311 | 0.0008612540 | Credible Set | purple |
| chr2:155631278 | 2 | 155.6313 | 0.0024809687 | Credible Set | purple |
| chr2:155631284 | 2 | 155.6313 | 0.0024809687 | Credible Set | purple |

Fine−mapping Regions

| snp | chr | pos (Mb) | pp | group | color |
|---|---|---|---|---|---|
| chr2:155631311 | 2 | 155.6313 | 0.0024809687 | Credible Set | purple |
| chr2:155632202 | 2 | 155.6322 | 0.0008729584 | Credible Set | purple |
| chr2:155632358 | 2 | 155.6324 | 0.0008969433 | Credible Set | purple |
| chr2:155632880 | 2 | 155.6329 | 0.0008530782 | Credible Set | purple |
| chr2:155633328 | 2 | 155.6333 | 0.0008750734 | Credible Set | purple |
| chr2:155640583 | 2 | 155.6406 | 0.0008721286 | Credible Set | purple |
| chr2:155640694 | 2 | 155.6407 | 0.0008825599 | Credible Set | purple |
| chr2:155640700 | 2 | 155.6407 | 0.0008825599 | Credible Set | purple |
| chr2:155641730 | 2 | 155.6417 | 0.0011207552 | Credible Set | purple |
| chr2:155647032 | 2 | 155.6470 | 0.0006982295 | Credible Set | purple |
| chr2:155647066 | 2 | 155.6471 | 0.0008546250 | Credible Set | purple |
| chr2:155659679 | 2 | 155.6597 | 0.0007542431 | Credible Set | purple |
| chr2:155664276 | 2 | 155.6643 | 0.0008553533 | Credible Set | purple |
| chr2:155718717 | 2 | 155.7187 | 0.0008729584 | Credible Set | purple |
| chr2:155718886 | 2 | 155.7189 | 0.0023705272 | Credible Set | purple |
| chr2:155718943 | 2 | 155.7189 | 0.0023801897 | Credible Set | purple |
| chr2:155748773 | 2 | 155.7488 | 0.0008729584 | Credible Set | purple |
| chr2:155754530 | 2 | 155.7545 | 0.0012327921 | Credible Set | purple |
| chr2:155754728 | 2 | 155.7547 | 0.0012082383 | Credible Set | purple |
| chr2:155765708 | 2 | 155.7657 | 0.0008729584 | Credible Set | purple |
| chr2:155787135 | 2 | 155.7871 | 0.0012369428 | Credible Set | purple |
| chr2:155787323 | 2 | 155.7873 | 0.0009387213 | Credible Set | purple |
| chr2:155813877 | 2 | 155.8139 | 0.0008614602 | Credible Set | purple |
| chr2:155816627 | 2 | 155.8166 | 0.0008729584 | Credible Set | purple |
| chr2:155818442 | 2 | 155.8184 | 0.0008729584 | Credible Set | purple |
| chr2:155819513 | 2 | 155.8195 | 0.0008729584 | Credible Set | purple |
| chr2:155819633 | 2 | 155.8196 | 0.0007644669 | Credible Set | purple |
| chr2:155819676 | 2 | 155.8197 | 0.0007721204 | Credible Set | purple |
| chr2:155828085 | 2 | 155.8281 | 0.0008668923 | Credible Set | purple |
| chr2:155828345 | 2 | 155.8283 | 0.0008751667 | Credible Set | purple |
| chr2:155828819 | 2 | 155.8288 | 0.0008233466 | Credible Set | purple |
| chr2:155830487 | 2 | 155.8305 | 0.0008729584 | Credible Set | purple |
| chr2:155830972 | 2 | 155.8310 | 0.0008404790 | Credible Set | purple |
| chr2:155843797 | 2 | 155.8438 | 0.0008729584 | Credible Set | purple |
| chr2:155843802 | 2 | 155.8438 | 0.0008729584 | Credible Set | purple |
| chr2:155845413 | 2 | 155.8454 | 0.0008658399 | Credible Set | purple |

Fine−mapping Regions

| snp | chr | pos (Mb) | pp | group | color |
|---|---|---|---|---|---|
| chr2:155859749 | 2 | 155.8597 | 0.0008729584 | Credible Set | purple |
| chr2:155860022 | 2 | 155.8600 | 0.0008248830 | Credible Set | purple |
| chr2:155917786 | 2 | 155.9178 | 0.0006886365 | Credible Set | purple |
| chr2:155917800 | 2 | 155.9178 | 0.0006855513 | Credible Set | purple |
| chr2:155934881 | 2 | 155.9349 | 0.0008749813 | Credible Set | purple |
| chr2:155935242 | 2 | 155.9352 | 0.0008471720 | Credible Set | purple |
| chr2:155941490 | 2 | 155.9415 | 0.0008729584 | Credible Set | purple |
| chr2:155941536 | 2 | 155.9415 | 0.0008729584 | Credible Set | purple |
| chr2:155962070 | 2 | 155.9621 | 0.0008729584 | Credible Set | purple |
| chr2:155968871 | 2 | 155.9689 | 0.0008455660 | Credible Set | purple |
| chr2:155968907 | 2 | 155.9689 | 0.0008455660 | Credible Set | purple |
| chr2:155969145 | 2 | 155.9691 | 0.0009201868 | Credible Set | purple |
| chr2:155969338 | 2 | 155.9693 | 0.0008729584 | Credible Set | purple |
| chr2:155973018 | 2 | 155.9730 | 0.0011764984 | Credible Set | purple |
| chr2:155973035 | 2 | 155.9730 | 0.0011544100 | Credible Set | purple |
| chr2:155973441 | 2 | 155.9734 | 0.0010746205 | Credible Set | purple |
| chr2:155973467 | 2 | 155.9735 | 0.0010746205 | Credible Set | purple |
| chr2:155973697 | 2 | 155.9737 | 0.0008706290 | Credible Set | purple |
| chr2:155974320 | 2 | 155.9743 | 0.0009072750 | Credible Set | purple |
| chr2:155974339 | 2 | 155.9743 | 0.0008180992 | Credible Set | purple |
| chr2:155974623 | 2 | 155.9746 | 0.0008729584 | Credible Set | purple |
| chr2:155985744 | 2 | 155.9857 | 0.0008525414 | Credible Set | purple |
| chr2:155986052 | 2 | 155.9861 | 0.0008729584 | Credible Set | purple |
| chr2:155988485 | 2 | 155.9885 | 0.0008757936 | Credible Set | purple |
| chr2:155989053 | 2 | 155.9891 | 0.0007539538 | Credible Set | purple |
| chr2:155991302 | 2 | 155.9913 | 0.0008729584 | Credible Set | purple |
| chr2:155991810 | 2 | 155.9918 | 0.0008729584 | Credible Set | purple |
| chr2:155992021 | 2 | 155.9920 | 0.0008729584 | Credible Set | purple |
| chr2:155997923 | 2 | 155.9979 | 0.0008554985 | Credible Set | purple |
| chr2:155998479 | 2 | 155.9985 | 0.0008859869 | Credible Set | purple |
| chr2:156011028 | 2 | 156.0110 | 0.0007276252 | Credible Set | purple |
| chr2:156011085 | 2 | 156.0111 | 0.0007311140 | Credible Set | purple |
| chr2:156011672 | 2 | 156.0117 | 0.0008479509 | Credible Set | purple |
| chr2:156039455 | 2 | 156.0395 | 0.0008729584 | Credible Set | purple |
| chr2:156041520 | 2 | 156.0415 | 0.0008752613 | Credible Set | purple |
| chr2:156043677 | 2 | 156.0437 | 0.0008729584 | Credible Set | purple |

Fine−mapping Regions

| snp | chr | pos (Mb) | pp | group | color |
|---|---|---|---|---|---|
| chr2:156071586 | 2 | 156.0716 | 0.0008715943 | Credible Set | purple |
| chr2:156071629 | 2 | 156.0716 | 0.0006392670 | Credible Set | purple |
| chr2:156071635 | 2 | 156.0716 | 0.0006391099 | Credible Set | purple |
| chr2:156071639 | 2 | 156.0716 | 0.0006392670 | Credible Set | purple |
| chr2:156071668 | 2 | 156.0717 | 0.0006361231 | Credible Set | purple |
| chr2:156073463 | 2 | 156.0735 | 0.0006715235 | Credible Set | purple |
| chr2:156077894 | 2 | 156.0779 | 0.0008686683 | Credible Set | purple |
| chr2:156078317 | 2 | 156.0783 | 0.0008841489 | Credible Set | purple |
| chr2:156078406 | 2 | 156.0784 | 0.0008729375 | Credible Set | purple |
| chr2:156078707 | 2 | 156.0787 | 0.0008729375 | Credible Set | purple |
| chr2:156078913 | 2 | 156.0789 | 0.0012663797 | Credible Set | purple |
| chr2:156082428 | 2 | 156.0824 | 0.0008729375 | Credible Set | purple |
| chr2:156082588 | 2 | 156.0826 | 0.0008729375 | Credible Set | purple |
| chr2:156082682 | 2 | 156.0827 | 0.0008729375 | Credible Set | purple |
| chr2:156111908 | 2 | 156.1119 | 0.0008606281 | Credible Set | purple |
| chr2:156111910 | 2 | 156.1119 | 0.0008547668 | Credible Set | purple |
| chr2:156124158 | 2 | 156.1242 | 0.0008773532 | Credible Set | purple |
| chr2:156124843 | 2 | 156.1248 | 0.0008729584 | Credible Set | purple |
| chr2:156133370 | 2 | 156.1334 | 0.0008986091 | Credible Set | purple |
| chr2:156133395 | 2 | 156.1334 | 0.0009067489 | Credible Set | purple |
| chr2:156133510 | 2 | 156.1335 | 0.0010201303 | Credible Set | purple |
| chr2:156215789 | 2 | 156.2158 | 0.0008729584 | Credible Set | purple |
| chr2:156259843 | 2 | 156.2598 | 0.0008729584 | Credible Set | purple |
| chr2:156267477 | 2 | 156.2675 | 0.0008613064 | Credible Set | purple |
| chr2:156269831 | 2 | 156.2698 | 0.0009066928 | Credible Set | purple |
| chr2:156282307 | 2 | 156.2823 | 0.0008729584 | Credible Set | purple |
| chr2:156302628 | 2 | 156.3026 | 0.0008729584 | Credible Set | purple |
| chr2:156302714 | 2 | 156.3027 | 0.0007702117 | Credible Set | purple |
| chr2:156302728 | 2 | 156.3027 | 0.0007702117 | Credible Set | purple |
| chr2:156302741 | 2 | 156.3027 | 0.0008729584 | Credible Set | purple |
| chr2:156303023 | 2 | 156.3030 | 0.0008794303 | Credible Set | purple |
| chr2:156305013 | 2 | 156.3050 | 0.0007059660 | Credible Set | purple |
| chr2:156323079 | 2 | 156.3231 | 0.0008523750 | Credible Set | purple |
| chr2:156343221 | 2 | 156.3432 | 0.0007941062 | Credible Set | purple |
| chr2:156343324 | 2 | 156.3433 | 0.0007956754 | Credible Set | purple |
| chr2:156368682 | 2 | 156.3687 | 0.0008417875 | Credible Set | purple |

Fine−mapping Regions

| snp | chr | pos (Mb) | pp | group | color |
|---|---|---|---|---|---|
| chr2:156368698 | 2 | 156.3687 | 0.0008552391 | Credible Set | purple |
| chr2:156378531 | 2 | 156.3785 | 0.0007349361 | Credible Set | purple |
| chr2:156383403 | 2 | 156.3834 | 0.0010412354 | Credible Set | purple |
| chr2:156383404 | 2 | 156.3834 | 0.0010412354 | Credible Set | purple |
| chr2:156383417 | 2 | 156.3834 | 0.0010412354 | Credible Set | purple |
| chr2:156387857 | 2 | 156.3879 | 0.0008729584 | Credible Set | purple |
| chr2:156401411 | 2 | 156.4014 | 0.0013159819 | Credible Set | purple |
| chr2:156407061 | 2 | 156.4071 | 0.0010562196 | Credible Set | purple |
| chr2:156407080 | 2 | 156.4071 | 0.0011150560 | Credible Set | purple |
| chr2:156407368 | 2 | 156.4074 | 0.0013323397 | Credible Set | purple |
| chr2:156417925 | 2 | 156.4179 | 0.0012908080 | Credible Set | purple |
| chr2:156419006 | 2 | 156.4190 | 0.0012792463 | Credible Set | purple |
| chr2:156423785 | 2 | 156.4238 | 0.0012331930 | Credible Set | purple |
| chr2:156449123 | 2 | 156.4491 | 0.0013182908 | Credible Set | purple |
| chr2:156449144 | 2 | 156.4491 | 0.0013182908 | Credible Set | purple |
| chr2:156453473 | 2 | 156.4535 | 0.0010382545 | Credible Set | purple |
| chr2:156453552 | 2 | 156.4536 | 0.0008230674 | Credible Set | purple |
| chr2:156453983 | 2 | 156.4540 | 0.0016877984 | Credible Set | purple |
| chr2:156455282 | 2 | 156.4553 | 0.0049765780 | Credible Set | purple |
| chr2:156455298 | 2 | 156.4553 | 0.0050296727 | Credible Set | purple |
| chr2:156455671 | 2 | 156.4557 | 0.0033185505 | Credible Set | purple |
| chr2:156455717 | 2 | 156.4557 | 0.0042939876 | Credible Set | purple |
| chr2:156465034 | 2 | 156.4650 | 0.0013182908 | Credible Set | purple |
| chr2:156465082 | 2 | 156.4651 | 0.0013182908 | Credible Set | purple |
| chr2:156465086 | 2 | 156.4651 | 0.0013182908 | Credible Set | purple |
| chr2:156465109 | 2 | 156.4651 | 0.0013182908 | Credible Set | purple |
| chr2:156465137 | 2 | 156.4651 | 0.0013182908 | Credible Set | purple |
| chr2:156465975 | 2 | 156.4660 | 0.0013182908 | Credible Set | purple |
| chr2:156468230 | 2 | 156.4682 | 0.0014013567 | Credible Set | purple |
| chr2:156474927 | 2 | 156.4749 | 0.0013183978 | Credible Set | purple |
| chr2:156483143 | 2 | 156.4831 | 0.0013082891 | Credible Set | purple |
| chr2:156488289 | 2 | 156.4883 | 0.0012884571 | Credible Set | purple |
| chr2:156491527 | 2 | 156.4915 | 0.0012903486 | Credible Set | purple |
| chr2:156491541 | 2 | 156.4915 | 0.0015609781 | Credible Set | purple |
| chr2:156491557 | 2 | 156.4916 | 0.0015453479 | Credible Set | purple |
| chr2:156521595 | 2 | 156.5216 | 0.0011542601 | Credible Set | purple |

Fine−mapping Regions

| snp | chr | pos (Mb) | pp | group | color |
|---|---|---|---|---|---|
| chr2:156521639 | 2 | 156.5216 | 0.0011669190 | Credible Set | purple |
| chr2:156530706 | 2 | 156.5307 | 0.0012401035 | Credible Set | purple |
| chr2:156532009 | 2 | 156.5320 | 0.0012860338 | Credible Set | purple |
| chr2:156532011 | 2 | 156.5320 | 0.0012772788 | Credible Set | purple |
| chr2:156532178 | 2 | 156.5322 | 0.0013163567 | Credible Set | purple |
| chr2:156567043 | 2 | 156.5670 | 0.0013236906 | Credible Set | purple |
| chr2:156567147 | 2 | 156.5671 | 0.0024815428 | Credible Set | purple |
| chr2:156567188 | 2 | 156.5672 | 0.0024638254 | Credible Set | purple |
| chr2:156567685 | 2 | 156.5677 | 0.0013753975 | Credible Set | purple |
| chr2:156576405 | 2 | 156.5764 | 0.0013780037 | Credible Set | purple |
| chr2:156576851 | 2 | 156.5769 | 0.0012061116 | Credible Set | purple |
| chr2:156577083 | 2 | 156.5771 | 0.0013186227 | Credible Set | purple |
| chr2:156585381 | 2 | 156.5854 | 0.0013195477 | Credible Set | purple |
| chr2:156587373 | 2 | 156.5874 | 0.0012676667 | Credible Set | purple |
| chr2:156587398 | 2 | 156.5874 | 0.0012214752 | Credible Set | purple |
| chr2:156592202 | 2 | 156.5922 | 0.0013162598 | Credible Set | purple |
| chr2:156595258 | 2 | 156.5953 | 0.0011612599 | Credible Set | purple |
| chr2:156632710 | 2 | 156.6327 | 0.0012542357 | Credible Set | purple |
| chr2:156632738 | 2 | 156.6327 | 0.0012339589 | Credible Set | purple |
| chr2:156632859 | 2 | 156.6329 | 0.0012190144 | Credible Set | purple |
| chr2:156641079 | 2 | 156.6411 | 0.0012057540 | Credible Set | purple |
| chr2:156641293 | 2 | 156.6413 | 0.0012257382 | Credible Set | purple |
| chr2:156649922 | 2 | 156.6499 | 0.0010137839 | Credible Set | purple |
| chr2:156649929 | 2 | 156.6499 | 0.0010224134 | Credible Set | purple |
| chr2:156650908 | 2 | 156.6509 | 0.0012558604 | Credible Set | purple |
| chr2:156668062 | 2 | 156.6681 | 0.0009524346 | Credible Set | purple |
| chr2:156668618 | 2 | 156.6686 | 0.0012820443 | Credible Set | purple |
| chr2:156704128 | 2 | 156.7041 | 0.0011191801 | Credible Set | purple |
| chr2:156704178 | 2 | 156.7042 | 0.0011197156 | Credible Set | purple |
| chr2:156711128 | 2 | 156.7111 | 0.0013186286 | Credible Set | purple |
| chr2:156743106 | 2 | 156.7431 | 0.0011996008 | Credible Set | purple |
| chr2:156743119 | 2 | 156.7431 | 0.0012283247 | Credible Set | purple |
| chr2:156764741 | 2 | 156.7647 | 0.0010280520 | Credible Set | purple |
| chr2:156764772 | 2 | 156.7648 | 0.0010593779 | Credible Set | purple |
| chr2:156770397 | 2 | 156.7704 | 0.0045005840 | Credible Set | purple |
| chr2:156795214 | 2 | 156.7952 | 0.0041958350 | Credible Set | purple |

Fine−mapping Regions

| snp | chr | pos (Mb) | pp | group | color |
|---|---|---|---|---|---|
| chr2:156795236 | 2 | 156.7952 | 0.0040398591 | Credible Set | purple |
| chr2:156803978 | 2 | 156.8040 | 0.0045005840 | Credible Set | purple |
| chr2:156812752 | 2 | 156.8128 | 0.0078204314 | Credible Set | purple |
| chr2:156815749 | 2 | 156.8157 | 0.0037639566 | Credible Set | purple |
| chr2:156815751 | 2 | 156.8158 | 0.0036573116 | Credible Set | purple |
| chr2:156815769 | 2 | 156.8158 | 0.0036428636 | Credible Set | purple |
| chr2:156825758 | 2 | 156.8258 | 0.0042301617 | Credible Set | purple |
| chr2:156825771 | 2 | 156.8258 | 0.0042233612 | Credible Set | purple |
| chr2:156825785 | 2 | 156.8258 | 0.0042233612 | Credible Set | purple |
| chr2:156825839 | 2 | 156.8258 | 0.0041361878 | Credible Set | purple |
| chr2:156825844 | 2 | 156.8258 | 0.0041361878 | Credible Set | purple |
| chr2:156825876 | 2 | 156.8259 | 0.0041361878 | Credible Set | purple |
| chr2:156845905 | 2 | 156.8459 | 0.0043854695 | Credible Set | purple |
| chr2:156861736 | 2 | 156.8617 | 0.0015350425 | Credible Set | purple |
| chr2:156861749 | 2 | 156.8617 | 0.0015350425 | Credible Set | purple |
| chr2:156861787 | 2 | 156.8618 | 0.0015350425 | Credible Set | purple |
| chr2:156861800 | 2 | 156.8618 | 0.0015350425 | Credible Set | purple |
| chr2:156861931 | 2 | 156.8619 | 0.0043039530 | Credible Set | purple |
| chr2:156862039 | 2 | 156.8620 | 0.0008416619 | Credible Set | purple |
| chr2:156862050 | 2 | 156.8621 | 0.0008416619 | Credible Set | purple |
| chr2:156862066 | 2 | 156.8621 | 0.0008477391 | Credible Set | purple |
| chr2:156862185 | 2 | 156.8622 | 0.0041150550 | Credible Set | purple |
| chr2:156862198 | 2 | 156.8622 | 0.0041150550 | Credible Set | purple |
| chr2:156882805 | 2 | 156.8828 | 0.0044848486 | Credible Set | purple |
| chr2:156882992 | 2 | 156.8830 | 0.0041668362 | Credible Set | purple |
| chr2:156883081 | 2 | 156.8831 | 0.0043785543 | Credible Set | purple |
| chr2:156883502 | 2 | 156.8835 | 0.0045005840 | Credible Set | purple |
| chr2:156886515 | 2 | 156.8865 | 0.0047444442 | Credible Set | purple |
| chr2:156886654 | 2 | 156.8867 | 0.0048164039 | Credible Set | purple |
| chr2:156887147 | 2 | 156.8871 | 0.0073998449 | Credible Set | purple |
| chr2:156887169 | 2 | 156.8872 | 0.0074553226 | Credible Set | purple |
| chr2:156948001 | 2 | 156.9480 | 0.0051733739 | Credible Set | purple |
| chr2:156959200 | 2 | 156.9592 | 0.0114860151 | Credible Set | purple |
| chr2:156959224 | 2 | 156.9592 | 0.0116728039 | Credible Set | purple |
| chr2:156959230 | 2 | 156.9592 | 0.0115102273 | Credible Set | purple |
| chr2:156959240 | 2 | 156.9592 | 0.0111861194 | Credible Set | purple |

Fine−mapping Regions

| snp | chr | pos (Mb) | pp | group | color |
|---|---|---|---|---|---|
| chr2:156973206 | 2 | 156.9732 | 0.004671987 | Credible Set | purple |
| chr2:156973473 | 2 | 156.9735 | 0.004701081 | Credible Set | purple |
| chr2:157010815 | 2 | 157.0108 | 0.004521964 | Credible Set | purple |
| chr2:157016739 | 2 | 157.0167 | 0.004048277 | Credible Set | purple |
| chr2:157017028 | 2 | 157.0170 | 0.004618923 | Credible Set | purple |
| chr2:157017101 | 2 | 157.0171 | 0.005369631 | Credible Set | purple |
| chr2:157017479 | 2 | 157.0175 | 0.005066103 | Credible Set | purple |
| chr2:157017498 | 2 | 157.0175 | 0.004966103 | Credible Set | purple |
| chr2:157027114 | 2 | 157.0271 | 0.004757518 | Credible Set | purple |
| chr2:157027115 | 2 | 157.0271 | 0.004757518 | Credible Set | purple |
| chr2:157027346 | 2 | 157.0273 | 0.004534067 | Credible Set | purple |
| chr2:157027552 | 2 | 157.0276 | 0.004500584 | Credible Set | purple |
| chr2:157027594 | 2 | 157.0276 | 0.004500584 | Credible Set | purple |
| chr2:157033406 | 2 | 157.0334 | 0.004500584 | Credible Set | purple |
| chr2:157033457 | 2 | 157.0335 | 0.004500584 | Credible Set | purple |
| chr2:157033569 | 2 | 157.0336 | 0.004500584 | Credible Set | purple |
| chr2:157033589 | 2 | 157.0336 | 0.004500584 | Credible Set | purple |
| chr2:157037970 | 2 | 157.0380 | 0.004941586 | Credible Set | purple |
| chr2:157040364 | 2 | 157.0404 | 0.003861107 | Credible Set | purple |
| chr2:157040379 | 2 | 157.0404 | 0.003985516 | Credible Set | purple |
| chr2:157040564 | 2 | 157.0406 | 0.004500584 | Credible Set | purple |
| chr2:157059250 | 2 | 157.0592 | 0.004500584 | Credible Set | purple |
| chr2:157059408 | 2 | 157.0594 | 0.004498129 | Credible Set | purple |
| chr2:157059810 | 2 | 157.0598 | 0.004499053 | Credible Set | purple |
| chr2:157059893 | 2 | 157.0599 | 0.004089039 | Credible Set | purple |
| chr2:157063511 | 2 | 157.0635 | 0.004500584 | Credible Set | purple |
| chr2:157065641 | 2 | 157.0656 | 0.004378791 | Credible Set | purple |
| chr2:157075626 | 2 | 157.0756 | 0.004500584 | Credible Set | purple |
| chr2:157075934 | 2 | 157.0759 | 0.004538454 | Credible Set | purple |
| chr2:157138758 | 2 | 157.1388 | 0.001397930 | Credible Set | purple |
| chr2:157138759 | 2 | 157.1388 | 0.001238887 | Credible Set | purple |
| chr2:157138913 | 2 | 157.1389 | 0.004518281 | Credible Set | purple |
| chr2:157144291 | 2 | 157.1443 | 0.004500584 | Credible Set | purple |
| chr2:157145837 | 2 | 157.1458 | 0.004500834 | Credible Set | purple |
| chr2:157149696 | 2 | 157.1497 | 0.004536776 | Credible Set | purple |
| chr2:157170168 | 2 | 157.1702 | 0.004499389 | Credible Set | purple |

Fine−mapping Regions

| snp | chr | pos (Mb) | pp | group | color |
|---|---|---|---|---|---|
| chr2:157170289 | 2 | 157.1703 | 0.003327649 | Credible Set | purple |
| chr2:157170311 | 2 | 157.1703 | 0.003320108 | Credible Set | purple |
| chr2:157234145 | 2 | 157.2341 | 0.004378489 | Credible Set | purple |
| chr2:157234194 | 2 | 157.2342 | 0.004244260 | Credible Set | purple |
| chr2:157234584 | 2 | 157.2346 | 0.005846621 | Credible Set | purple |
| chr2:157234605 | 2 | 157.2346 | 0.006023225 | Credible Set | purple |
| chr2:157234716 | 2 | 157.2347 | 0.012394370 | Credible Set | purple |
| chr2:157234718 | 2 | 157.2347 | 0.012313502 | Credible Set | purple |
| chr2:157234722 | 2 | 157.2347 | 0.012368836 | Credible Set | purple |
| chr2:157244801 | 2 | 157.2448 | 0.003053452 | Credible Set | purple |
| chr2:157245205 | 2 | 157.2452 | 0.004377639 | Credible Set | purple |
| chr2:157245270 | 2 | 157.2453 | 0.004323443 | Credible Set | purple |
| chr2:157271470 | 2 | 157.2715 | 0.004546292 | Credible Set | purple |
| chr2:157323455 | 2 | 157.3235 | 0.003731176 | Credible Set | purple |
| chr2:157323490 | 2 | 157.3235 | 0.004142781 | Credible Set | purple |
| chr2:157323542 | 2 | 157.3235 | 0.004164396 | Credible Set | purple |
| chr2:157328889 | 2 | 157.3289 | 0.003226906 | Credible Set | purple |
| chr2:157350851 | 2 | 157.3509 | 0.003485881 | Credible Set | purple |
| chr2:157356604 | 2 | 157.3566 | 0.007146490 | Credible Set | purple |
| chr2:157383750 | 2 | 157.3837 | 0.007447453 | Credible Set | purple |
| chr2:157384030 | 2 | 157.3840 | 0.016024599 | Credible Set | purple |
| chr2:157384046 | 2 | 157.3840 | 0.016005888 | Credible Set | purple |
| chr2:157393248 | 2 | 157.3932 | 0.002529089 | Credible Set | purple |
| chr2:157393326 | 2 | 157.3933 | 0.003629860 | Credible Set | purple |
| chr2:157393328 | 2 | 157.3933 | 0.003109633 | Credible Set | purple |
| chr2:157393885 | 2 | 157.3939 | 0.003844495 | Credible Set | purple |
| chr2:157393913 | 2 | 157.3939 | 0.003905813 | Credible Set | purple |
| chr2:157393914 | 2 | 157.3939 | 0.003855055 | Credible Set | purple |
| chr2:157393915 | 2 | 157.3939 | 0.003859032 | Credible Set | purple |
| chr2:157396118 | 2 | 157.3961 | 0.004921267 | Credible Set | purple |
| chr2:157396325 | 2 | 157.3963 | 0.001303472 | Credible Set | purple |
| chr2:157396340 | 2 | 157.3963 | 0.001310087 | Credible Set | purple |
| chr2:157396357 | 2 | 157.3964 | 0.001310087 | Credible Set | purple |
| chr2:157396513 | 2 | 157.3965 | 0.005767159 | Credible Set | purple |
| chr2:157401494 | 2 | 157.4015 | 0.005643284 | Credible Set | purple |
| chr2:157425583 | 2 | 157.4256 | 0.005767159 | Credible Set | purple |

Fine−mapping Regions

| snp | chr | pos (Mb) | pp | group | color |
|---|---|---|---|---|---|
| chr2:157434588 | 2 | 157.4346 | 0.005767159 | Credible Set | purple |
| chr2:157439371 | 2 | 157.4394 | 0.005767159 | Credible Set | purple |
| chr2:157455578 | 2 | 157.4556 | 0.005520168 | Credible Set | purple |
| chr2:157458440 | 2 | 157.4584 | 0.005767159 | Credible Set | purple |
| chr2:157464790 | 2 | 157.4648 | 0.003979447 | Credible Set | purple |
| chr2:157464811 | 2 | 157.4648 | 0.003979447 | Credible Set | purple |
| chr2:157464835 | 2 | 157.4648 | 0.003979447 | Credible Set | purple |
| chr2:157464946 | 2 | 157.4649 | 0.005768230 | Credible Set | purple |
| chr2:157465338 | 2 | 157.4653 | 0.005786666 | Credible Set | purple |
| chr2:157502041 | 2 | 157.5020 | 0.007384423 | Credible Set | purple |
| chr2:157502044 | 2 | 157.5020 | 0.005572784 | Credible Set | purple |
| chr2:157549374 | 2 | 157.5494 | 0.005672009 | Credible Set | purple |
| chr2:157611240 | 2 | 157.6112 | 0.001999621 | Credible Set | purple |
| chr2:157611267 | 2 | 157.6113 | 0.001994993 | Credible Set | purple |
| chr2:157611281 | 2 | 157.6113 | 0.001994993 | Credible Set | purple |
| chr2:157611292 | 2 | 157.6113 | 0.001994993 | Credible Set | purple |
| chr2:157611294 | 2 | 157.6113 | 0.001994993 | Credible Set | purple |
| chr2:157630394 | 2 | 157.6304 | 0.005983166 | Credible Set | purple |
| chr2:157630611 | 2 | 157.6306 | 0.005899822 | Credible Set | purple |
| chr2:157630938 | 2 | 157.6309 | 0.007912554 | Credible Set | purple |
| chr2:157630968 | 2 | 157.6310 | 0.007977824 | Credible Set | purple |
| chr2:157630998 | 2 | 157.6310 | 0.007909786 | Credible Set | purple |
| chr2:157635749 | 2 | 157.6357 | 0.005178926 | Credible Set | purple |
| chr2:158322238 | 2 | 158.3222 | 0.003802500 | Credible Set | purple |
| chr2:158334991 | 2 | 158.3350 | 0.003391315 | Credible Set | purple |
| chr2:158335036 | 2 | 158.3350 | 0.002878618 | Credible Set | purple |
| chr2:158380870 | 2 | 158.3809 | 0.003885663 | Credible Set | purple |
| chr2:158380953 | 2 | 158.3810 | 0.003892789 | Credible Set | purple |
| chr2:158424885 | 2 | 158.4249 | 0.001092579 | Credible Set | purple |
| chr2:158424892 | 2 | 158.4249 | 0.001092579 | Credible Set | purple |
| chr2:158432408 | 2 | 158.4324 | 0.004030962 | Credible Set | purple |
| chr2:158463993 | 2 | 158.4640 | 0.003605254 | Credible Set | purple |
| chr2:158499568 | 2 | 158.4996 | 0.003532919 | Credible Set | purple |
| chr2:158499611 | 2 | 158.4996 | 0.003603675 | Credible Set | purple |
| chr2:158518637 | 2 | 158.5186 | 0.002402589 | Credible Set | purple |
| chr2:158518964 | 2 | 158.5190 | 0.003409110 | Credible Set | purple |

Fine−mapping Regions

| snp | chr | pos (Mb) | pp | group | color |
|---|---|---|---|---|---|
| chr2:158519003 | 2 | 158.5190 | 0.003409110 | Credible Set | purple |
| chr2:158523085 | 2 | 158.5231 | 0.001070763 | Credible Set | purple |
| chr2:158523125 | 2 | 158.5231 | 0.001083386 | Credible Set | purple |
| chr2:158523133 | 2 | 158.5231 | 0.001075110 | Credible Set | purple |
| chr2:158523135 | 2 | 158.5231 | 0.001069148 | Credible Set | purple |
| chr2:158531021 | 2 | 158.5310 | 0.001788435 | Credible Set | purple |
| chr2:158591403 | 2 | 158.5914 | 0.003891379 | Credible Set | purple |
| chr2:158591550 | 2 | 158.5916 | 0.001241109 | Credible Set | purple |
| chr2:158591613 | 2 | 158.5916 | 0.001225800 | Credible Set | purple |
| chr2:158601272 | 2 | 158.6013 | 0.003826526 | Credible Set | purple |
| chr2:158625324 | 2 | 158.6253 | 0.003893113 | Credible Set | purple |
| chr2:158625366 | 2 | 158.6254 | 0.003893113 | Credible Set | purple |
| chr2:158627877 | 2 | 158.6279 | 0.007562548 | Credible Set | purple |
| chr2:158627920 | 2 | 158.6279 | 0.007562548 | Credible Set | purple |
| chr2:158628144 | 2 | 158.6281 | 0.003620250 | Credible Set | purple |
| chr2:158628165 | 2 | 158.6282 | 0.003632114 | Credible Set | purple |
| chr2:158631398 | 2 | 158.6314 | 0.001314150 | Credible Set | purple |
| chr2:158631624 | 2 | 158.6316 | 0.001912639 | Credible Set | purple |
| chr2:158640014 | 2 | 158.6400 | 0.003732603 | Credible Set | purple |
| chr2:158665126 | 2 | 158.6651 | 0.004378129 | Credible Set | purple |

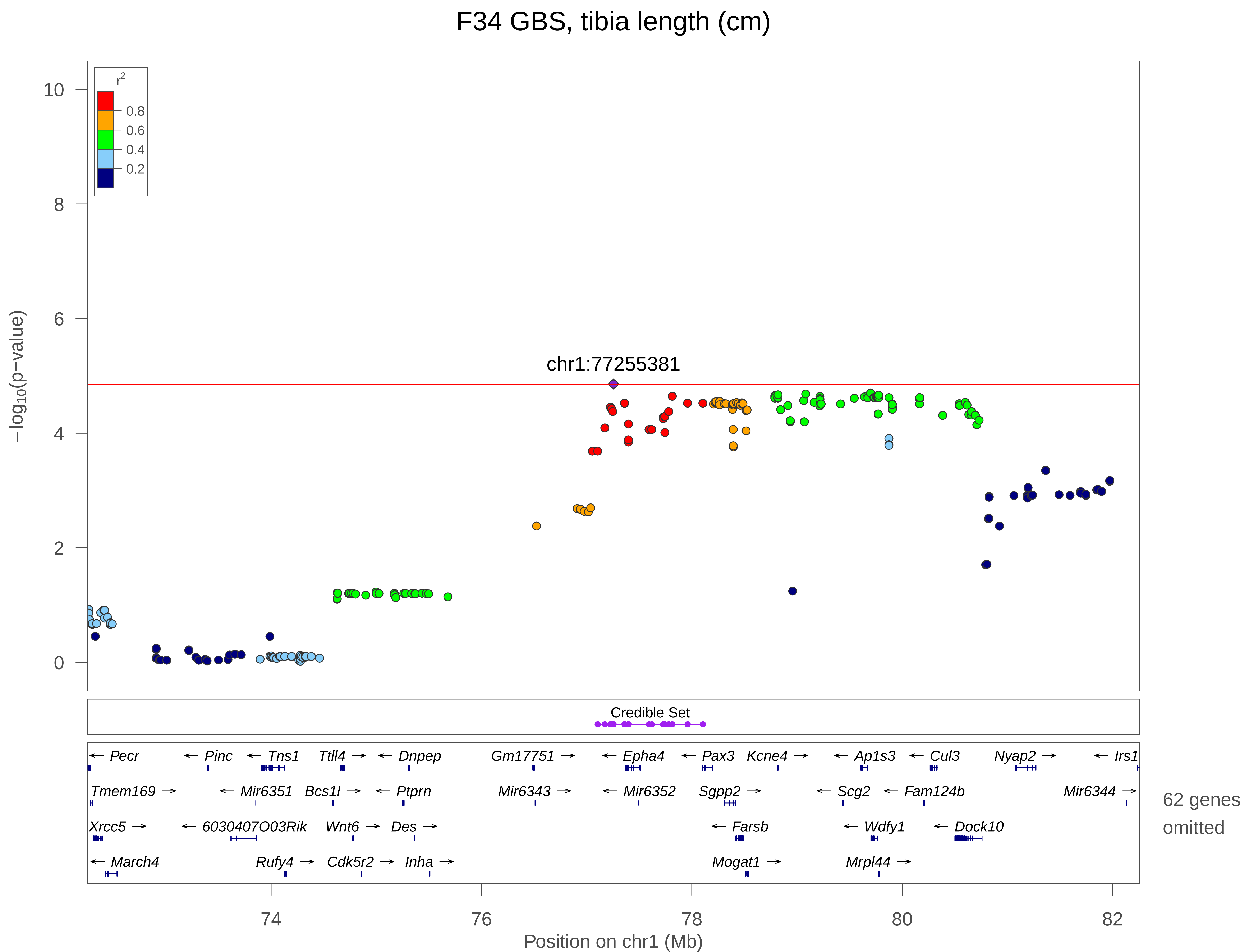

F34 GBS, tibia length (cm)
r2
0.8
0.6
0.4
0.2
10
8
6
4
2
0
−log10(p−value)
chr1:77255381
Credible Set
Pecr
Pinc
Tns1
Ttll4
Dnpep
Gm17751
Epha4
Pax3
Kcne4
Ap1s3
Cul3
Nyap2
Irs1
Tmem169
Mir6351
Bcs1l
Ptprn
Mir6343
Mir6352
Sgpp2
Scg2
Fam124b
Mir6344
Xrcc5
6030407O03Rik
Wnt6
Des
Farsb
Wdfy1
Dock10
March4
Rufy4
Cdk5r2
Inha
Mogat1
Mrpl44
62 genes omitted
74
76
78
80
82
Position on chr1 (Mb)

| | |
|---|---|
| date: | Thu Oct 25 17:14:39 2018 |
| build: | F34GBS |
| display range: | chr1:72255381−82255381 [72255381−82255381] |
| hilite range: | 0 − 0 [ 0 − 0 ] |
| reference SNP: | chr1:77255381 |
| number of SNPs plotted: | 308 |
| min P−value: | 1.39E−5 [chr1:77255381] |
| max P−value: | 9.52E−1 [chr1:74278512] |
| omitted Genes: | Smarcal1, Ankar, Rpl37a |
| omitted Genes: | Igfbp2, Igfbp5, Tnp1 |
| omitted Genes: | 1700027A15Rik, D530049I02Rik, Cxcr2 |
| omitted Genes: | Cxcr1, Mir1928, Arpc2 |
| omitted Genes: | Gpbar1, Aamp, Pnkd |
| omitted Genes: | Tmbim1, Catip, Slc11a1 |
| omitted Genes: | Ctdsp1, Mir26b, Vil1 |
| omitted Genes: | Usp37, Cnot9, Plcd4 |
| omitted Genes: | Zfp142, Rnf25, Stk36 |
| omitted Genes: | Cyp27a1, Prkag3, Wnt10a |
| omitted Genes: | Fev, Cryba2, Mir375 |
| omitted Genes: | Ccdc108, Ihh, Nhej1 |
| omitted Genes: | Slc23a3, Cnppd1, Fam134a |
| omitted Genes: | Zfand2b, Abcb6, Atg9a |
| omitted Genes: | Ankzf1, Glb1l, Stk16 |
| omitted Genes: | Tuba4a, A630095N17Rik, Dnajb2 |
| omitted Genes: | Resp18, Gm15179, Speg |
| omitted Genes: | Gmppa, Asic4, Chpf |
| omitted Genes: | Tmem198, Obsl1, Stk11ip |
| omitted Genes: | Slc4a3, Utp14b, Acsl3 |
| omitted Genes: | Serpine2, 1700016L21Rik |

Make more plots at http://csg.sph.umich.edu/locuszoom/

Fine−mapping Regions

| snp | chr | pos (Mb) | pp | group | color |
|---|---|---|---|---|---|
| chr1:77104885 | 1 | 77.10488 | 0.008126436 | Credible Set | purple |
| chr1:77172984 | 1 | 77.17298 | 0.019564894 | Credible Set | purple |
| chr1:77226748 | 1 | 77.22675 | 0.042605449 | Credible Set | purple |
| chr1:77226771 | 1 | 77.22677 | 0.043033112 | Credible Set | purple |
| chr1:77236123 | 1 | 77.23612 | 0.040789617 | Credible Set | purple |
| chr1:77247183 | 1 | 77.24718 | 0.036464543 | Credible Set | purple |
| chr1:77255381 | 1 | 77.25538 | 0.104374844 | Credible Set | purple |
| chr1:77360230 | 1 | 77.36023 | 0.049821912 | Credible Set | purple |
| chr1:77360231 | 1 | 77.36023 | 0.049911801 | Credible Set | purple |
| chr1:77360235 | 1 | 77.36024 | 0.050042131 | Credible Set | purple |
| chr1:77396673 | 1 | 77.39667 | 0.011417742 | Credible Set | purple |
| chr1:77396674 | 1 | 77.39667 | 0.012455288 | Credible Set | purple |
| chr1:77396959 | 1 | 77.39696 | 0.022673149 | Credible Set | purple |
| chr1:77592858 | 1 | 77.59286 | 0.018331029 | Credible Set | purple |
| chr1:77592859 | 1 | 77.59286 | 0.018331029 | Credible Set | purple |
| chr1:77617819 | 1 | 77.61782 | 0.018389979 | Credible Set | purple |
| chr1:77728800 | 1 | 77.72880 | 0.029422136 | Credible Set | purple |
| chr1:77728804 | 1 | 77.72880 | 0.029670651 | Credible Set | purple |
| chr1:77728811 | 1 | 77.72881 | 0.027913013 | Credible Set | purple |
| chr1:77743267 | 1 | 77.74327 | 0.029821353 | Credible Set | purple |
| chr1:77743271 | 1 | 77.74327 | 0.029699371 | Credible Set | purple |
| chr1:77743297 | 1 | 77.74330 | 0.030277323 | Credible Set | purple |
| chr1:77743443 | 1 | 77.74344 | 0.016399557 | Credible Set | purple |
| chr1:77779980 | 1 | 77.77998 | 0.036476533 | Credible Set | purple |
| chr1:77814150 | 1 | 77.81415 | 0.065375684 | Credible Set | purple |
| chr1:77959635 | 1 | 77.95964 | 0.050164984 | Credible Set | purple |
| chr1:78104946 | 1 | 78.10495 | 0.050164984 | Credible Set | purple |
| chr1:78105022 | 1 | 78.10502 | 0.050164984 | Credible Set | purple |

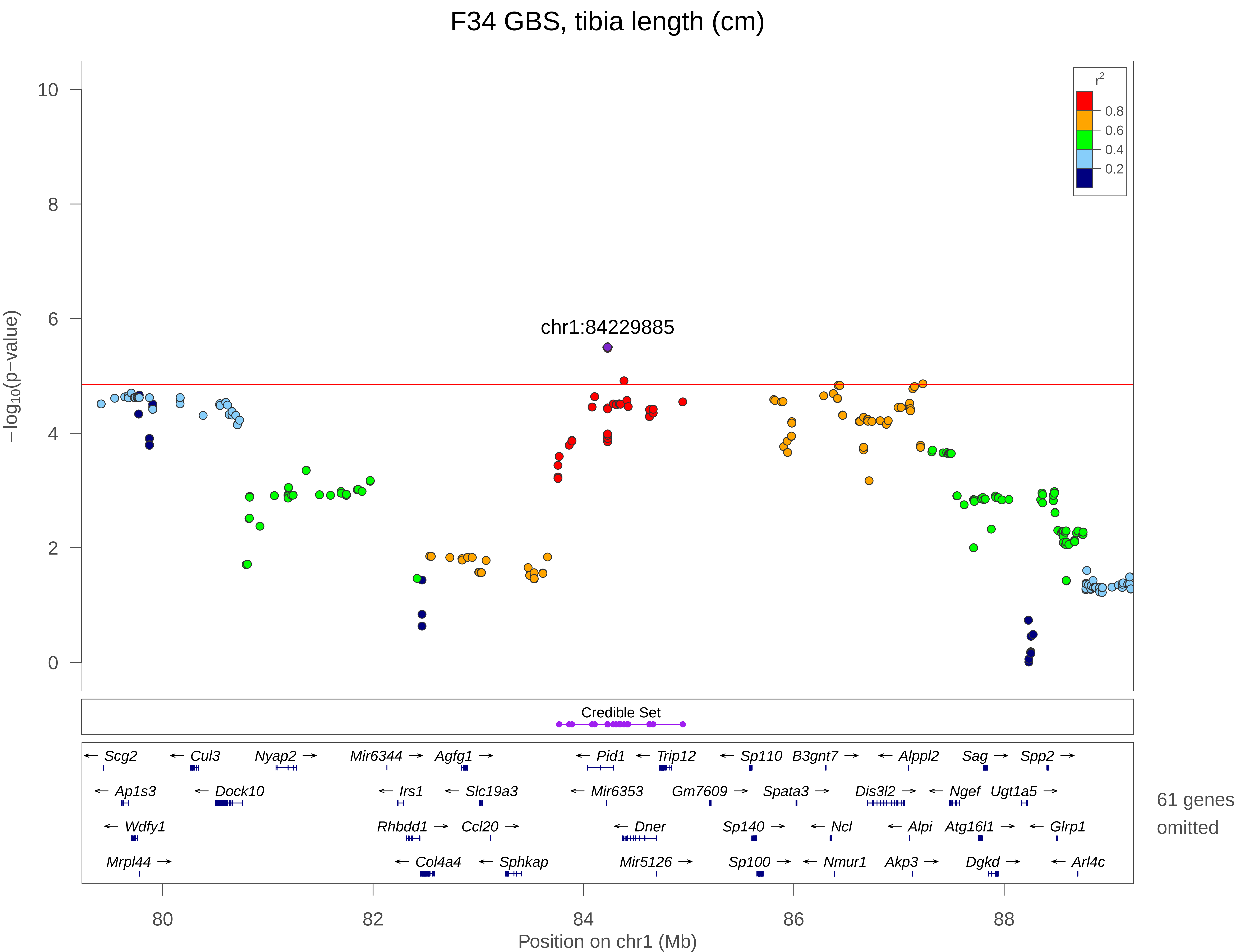
F34 GBS, tibia length (cm)
r2
0.8
0.6
0.4
0.2
10
8
6
4
2
0
−log10(p−value)
chr1:84229885
Credible Set
Scg2
Cul3
Nyap2
Mir6344
Agfg1
Pid1
Trip12
Sp110
B3gnt7
Alppl2
Sag
Spp2
Ap1s3
Dock10
Irs1
Slc19a3
Mir6353
Gm7609
Spata3
Dis3l2
Ngef
Ugt1a5
Wdfy1
Rhbdd1
Ccl20
Dner
Sp140
Ncl
Alpi
Atg16l1
Glrp1
Mrpl44
Col4a4
Sphkap
Mir5126
Sp100
Nmur1
Akp3
Dgkd
Arl4c
61 genes omitted
80
82
84
86
88
Position on chr1 (Mb)

| | |
|---|---|
| date: | Thu Oct 25 17:14:48 2018 |
| build: | F34GBS |
| display range: | chr1:79229885−89229885 [79229885−89229885] |
| hilite range: | 0 − 0 [ 0 − 0 ] |
| reference SNP: | chr1:84229885 |
| number of SNPs plotted: | 324 |
| min P−value: | 3.14E−6 [chr1:84229885] |
| max P−value: | 9.85E−1 [chr1:88235333] |
| omitted Genes: | Serpine2, Fam124b, 1700016L21Rik |
| omitted Genes: | Col4a3, Mff, Tm4sf20 |
| omitted Genes: | Krtap28−13, Daw1, Fbxo36 |
| omitted Genes: | Slc16a14, A530032D15Rik, C130026I21Rik |
| omitted Genes: | A630001G21Rik, Mir8096, 9930111H07Rik |
| omitted Genes: | Cab39, Itm2c, 4933407L21Rik |
| omitted Genes: | Gpr55, 2810459M11Rik, Psmd1 |
| omitted Genes: | Htr2b, Armc9, Snora75 |
| omitted Genes: | Gm24148, Snord82, C130036L24Rik |
| omitted Genes: | 1700019O17Rik, Ptma, Pde6d |
| omitted Genes: | Cops7b, Nppc, Ecel1 |
| omitted Genes: | Prss56, Chrnd, Chrng |
| omitted Genes: | Eif4e2, Efhd1, Gigyf2 |
| omitted Genes: | Kcnj13, 3110079O15Rik, Neu2 |
| omitted Genes: | Inpp5d, Scarna6, Gm19582 |
| omitted Genes: | Usp40, Ugt1a10, Ugt1a9 |
| omitted Genes: | Ugt1a7c, Ugt1a6b, Ugt1a6a |
| omitted Genes: | Ugt1a2, Dnajb3, Ugt1a1 |
| omitted Genes: | Mroh2a, Hjurp, Trpm8 |
| omitted Genes: | Gm19589, Platr5, 1700067G17Rik |
| omitted Genes: | Sh3bp4 |

Fine−mapping Regions

| snp | chr | pos (Mb) | pp | group | color |
|---|---|---|---|---|---|
| chr1:83769887 | 1 | 83.76989 | 0.003606986 | Credible Set | purple |
| chr1:83864780 | 1 | 83.86478 | 0.005513547 | Credible Set | purple |
| chr1:83891050 | 1 | 83.89105 | 0.006615451 | Credible Set | purple |
| chr1:83891408 | 1 | 83.89141 | 0.006462805 | Credible Set | purple |
| chr1:84082487 | 1 | 84.08249 | 0.023491739 | Credible Set | purple |
| chr1:84106829 | 1 | 84.10683 | 0.034932382 | Credible Set | purple |
| chr1:84229883 | 1 | 84.22988 | 0.223038567 | Credible Set | purple |
| chr1:84229885 | 1 | 84.22988 | 0.234056135 | Credible Set | purple |
| chr1:84230139 | 1 | 84.23014 | 0.022752797 | Credible Set | purple |
| chr1:84230140 | 1 | 84.23014 | 0.021803722 | Credible Set | purple |
| chr1:84230230 | 1 | 84.23023 | 0.006299284 | Credible Set | purple |
| chr1:84230232 | 1 | 84.23023 | 0.007095944 | Credible Set | purple |
| chr1:84230233 | 1 | 84.23023 | 0.008010994 | Credible Set | purple |
| chr1:84230245 | 1 | 84.23024 | 0.008408622 | Credible Set | purple |
| chr1:84230255 | 1 | 84.23025 | 0.008446651 | Credible Set | purple |
| chr1:84283668 | 1 | 84.28367 | 0.026427192 | Credible Set | purple |
| chr1:84311316 | 1 | 84.31132 | 0.026206138 | Credible Set | purple |
| chr1:84311384 | 1 | 84.31138 | 0.025543727 | Credible Set | purple |
| chr1:84340686 | 1 | 84.34069 | 0.026531472 | Credible Set | purple |
| chr1:84352584 | 1 | 84.35258 | 0.026185263 | Credible Set | purple |
| chr1:84385821 | 1 | 84.38582 | 0.064085697 | Credible Set | purple |
| chr1:84413844 | 1 | 84.41384 | 0.030348182 | Credible Set | purple |
| chr1:84425660 | 1 | 84.42566 | 0.023820802 | Credible Set | purple |
| chr1:84628469 | 1 | 84.62847 | 0.016422607 | Credible Set | purple |
| chr1:84628475 | 1 | 84.62847 | 0.016298809 | Credible Set | purple |
| chr1:84629053 | 1 | 84.62905 | 0.021283367 | Credible Set | purple |
| chr1:84663039 | 1 | 84.66304 | 0.018681984 | Credible Set | purple |
| chr1:84663099 | 1 | 84.66310 | 0.021628409 | Credible Set | purple |
| chr1:84944754 | 1 | 84.94475 | 0.028599845 | Credible Set | purple |

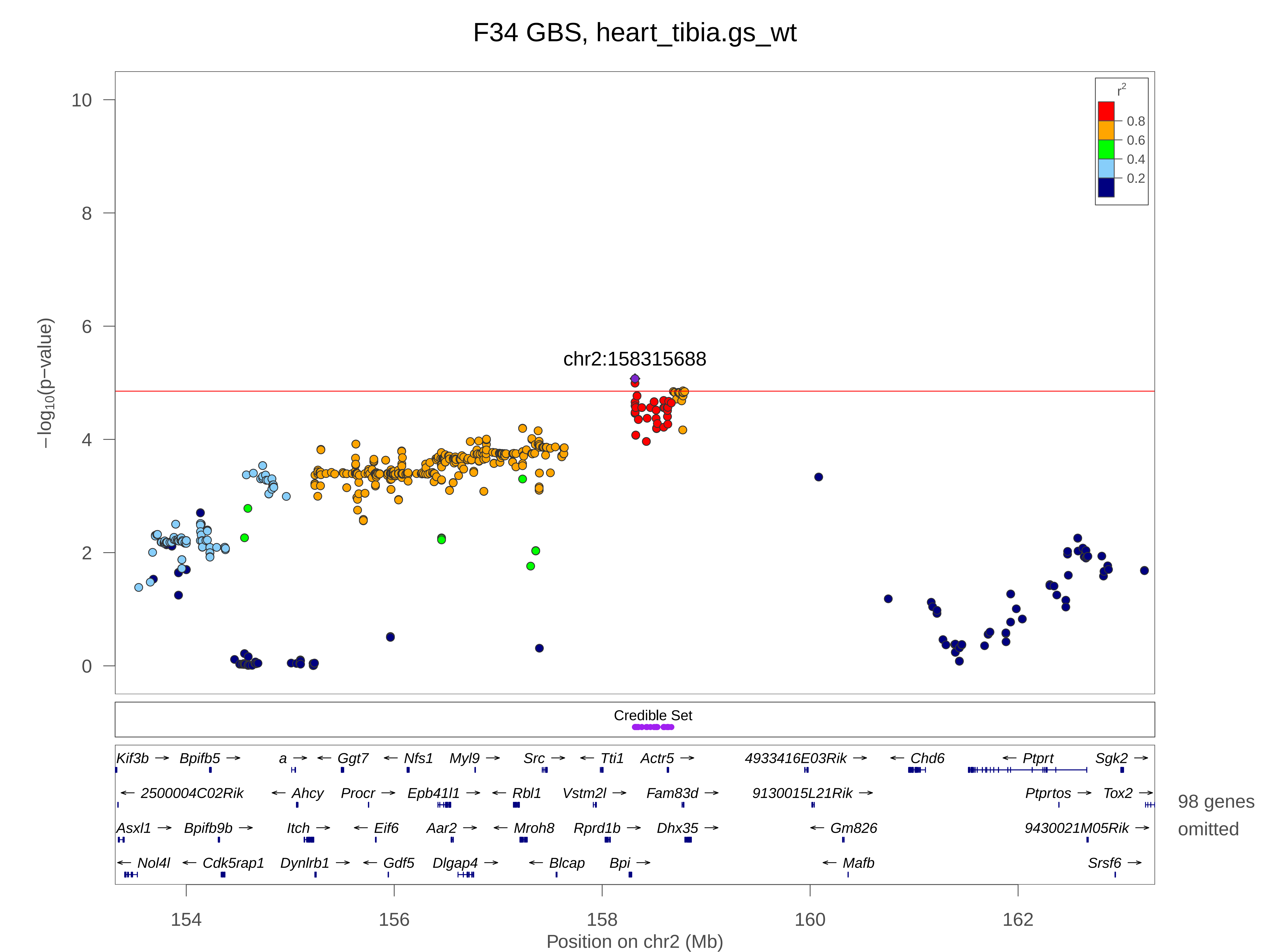

F34 GBS, heart_tibia.gs_wt
chr2:158315688
−log10(p−value)
10
8
6
4
2
0
r2
0.8
0.6
0.4
0.2
Credible Set
Kif3b →
Bpifb5 →
a →
← Ggt7
← Nfs1
Myl9 →
Src →
← Tti1
Actr5 →
4933416E03Rik →
← Chd6
← Ptprt
Sgk2 →
← 2500004C02Rik
← Ahcy
Procr →
Epb41l1 →
← Rbl1
Vstm2l →
Fam83d →
9130015L21Rik →
Ptprtos →
Tox2 →
Asxl1 →
Bpifb9b →
Itch →
← Eif6
Aar2 →
← Mroh8
Rprd1b →
Dhx35 →
← Gm826
9430021M05Rik →
← Nol4l
← Cdk5rap1
Dynlrb1 →
← Gdf5
Dlgap4 →
← Blcap
Bpi →
← Mafb
Srsf6 →
98 genes omitted
154
156
158
160
162
Position on chr2 (Mb)

date: Thu Oct 25 17:14:57 2018
build: F34GBS
display range: chr2:153315688−163315688 [153315688−163315688]
hilite range: 0 − 0 [ 0 − 0 ]
reference SNP: chr2:158315688
number of SNPs plotted: 737
min P−value: 8.45E−6 [chr2:158315688]
max P−value: 9.91E−1 [chr2:155220725]
omitted Genes: Commd7, Dnmt3b, Mapre1
omitted Genes: Efcab8, Sun5, Bpifb2
omitted Genes: Bpifb6, Bpifb3, Bpifb4
omitted Genes: Bpifa6, Bpifa2, Bpifa3
omitted Genes: Bpifa1, Bpifa5, Bpifb1
omitted Genes: Bpifb9a, Snta1, Cbfa2t2
omitted Genes: Necab3, 1700003F12Rik, Actl10
omitted Genes: E2f1, Pxmp4, Zfp341
omitted Genes: Chmp4b, Raly, Eif2s2
omitted Genes: 2310005A03Rik, Map1lc3a, Pigu
omitted Genes: Mir695, Trp53inp2, Ncoa6
omitted Genes: Acss2, Acss2os, Gss
omitted Genes: Myh7b, Mir499, Trpc4ap
omitted Genes: Edem2, Mmp24, BC029722
omitted Genes: Fam83c, Uqcc1, Cep250
omitted Genes: 6430550D23Rik, Ergic3, Fer1l4
omitted Genes: Spag4, Cpne1, Rbm12
omitted Genes: Romo1, Rbm39, Phf20
omitted Genes: Scand1, Cnbd2, 2900097C17Rik
omitted Genes: Gm14169, 4930405A21Rik, Tgif2
omitted Genes: 5430405H02Rik, 1110008F13Rik, Sla2
omitted Genes: Ndrg3, Dsn1, Soga1
omitted Genes: Tldc2, Samhd1, Rpn2
omitted Genes: Ghrh, Manbal, Nnat
omitted Genes: Ctnnbl1, 2010009K17Rik, Tgm2
omitted Genes: D630003M21Rik, 1700060C20Rik, Mir7685
omitted Genes: Lbp, Snhg17, Snhg11
omitted Genes: Ralgapb, Adig, Arhgap40
omitted Genes: Gm14204, Slc32a1, Mir3474
omitted Genes: Ppp1r16b, Top1, Plcg1
omitted Genes: Zhx3, Lpin3, Emilin3
Make more plots at http://csg.sph.umich.edu/locuszoom/
omitted Genes: L3mbtl1, Ift52, Mybl2

Fine−mapping Regions

| snp | chr | pos (Mb) | pp | group | color |
|---|---|---|---|---|---|
| chr2:158315688 | 2 | 158.3157 | 0.067553513 | Credible Set | purple |
| chr2:158315756 | 2 | 158.3158 | 0.056710404 | Credible Set | purple |
| chr2:158316223 | 2 | 158.3162 | 0.027192220 | Credible Set | purple |
| chr2:158316249 | 2 | 158.3162 | 0.023575944 | Credible Set | purple |
| chr2:158316290 | 2 | 158.3163 | 0.017644086 | Credible Set | purple |
| chr2:158316307 | 2 | 158.3163 | 0.017640453 | Credible Set | purple |
| chr2:158317181 | 2 | 158.3172 | 0.018433721 | Credible Set | purple |
| chr2:158317182 | 2 | 158.3172 | 0.018162421 | Credible Set | purple |
| chr2:158317215 | 2 | 158.3172 | 0.018356902 | Credible Set | purple |
| chr2:158322238 | 2 | 158.3222 | 0.022009191 | Credible Set | purple |
| chr2:158322761 | 2 | 158.3228 | 0.007553278 | Credible Set | purple |
| chr2:158322795 | 2 | 158.3228 | 0.007645301 | Credible Set | purple |
| chr2:158322809 | 2 | 158.3228 | 0.007625461 | Credible Set | purple |
| chr2:158334991 | 2 | 158.3350 | 0.034966947 | Credible Set | purple |
| chr2:158335036 | 2 | 158.3350 | 0.034419970 | Credible Set | purple |
| chr2:158348022 | 2 | 158.3480 | 0.014038046 | Credible Set | purple |
| chr2:158348028 | 2 | 158.3480 | 0.013985272 | Credible Set | purple |
| chr2:158348031 | 2 | 158.3480 | 0.013855400 | Credible Set | purple |
| chr2:158380870 | 2 | 158.3809 | 0.022022804 | Credible Set | purple |
| chr2:158380953 | 2 | 158.3810 | 0.021986754 | Credible Set | purple |
| chr2:158424885 | 2 | 158.4249 | 0.005965392 | Credible Set | purple |
| chr2:158432408 | 2 | 158.4324 | 0.014573331 | Credible Set | purple |
| chr2:158463993 | 2 | 158.4640 | 0.021983874 | Credible Set | purple |
| chr2:158499568 | 2 | 158.4996 | 0.027661448 | Credible Set | purple |
| chr2:158499611 | 2 | 158.4996 | 0.027510236 | Credible Set | purple |
| chr2:158518637 | 2 | 158.5186 | 0.014467513 | Credible Set | purple |
| chr2:158518964 | 2 | 158.5190 | 0.019963248 | Credible Set | purple |
| chr2:158519003 | 2 | 158.5190 | 0.019963248 | Credible Set | purple |
| chr2:158523085 | 2 | 158.5231 | 0.010001658 | Credible Set | purple |
| chr2:158523125 | 2 | 158.5231 | 0.009970331 | Credible Set | purple |
| chr2:158523133 | 2 | 158.5231 | 0.009764408 | Credible Set | purple |
| chr2:158523135 | 2 | 158.5231 | 0.009871917 | Credible Set | purple |
| chr2:158531021 | 2 | 158.5310 | 0.011888153 | Credible Set | purple |
| chr2:158591403 | 2 | 158.5914 | 0.021944319 | Credible Set | purple |
| chr2:158591550 | 2 | 158.5916 | 0.010313254 | Credible Set | purple |
| chr2:158591613 | 2 | 158.5916 | 0.010307309 | Credible Set | purple |

Fine−mapping Regions

| snp | chr | pos (Mb) | pp | group | color |
|---|---|---|---|---|---|
| chr2:158591734 | 2 | 158.5917 | 0.02899447 | Credible Set | purple |
| chr2:158591782 | 2 | 158.5918 | 0.02896804 | Credible Set | purple |
| chr2:158601272 | 2 | 158.6013 | 0.02186655 | Credible Set | purple |
| chr2:158625324 | 2 | 158.6253 | 0.02200009 | Credible Set | purple |
| chr2:158625366 | 2 | 158.6254 | 0.02200009 | Credible Set | purple |
| chr2:158627877 | 2 | 158.6279 | 0.01556789 | Credible Set | purple |
| chr2:158627920 | 2 | 158.6279 | 0.01556789 | Credible Set | purple |
| chr2:158628144 | 2 | 158.6281 | 0.01959211 | Credible Set | purple |
| chr2:158628165 | 2 | 158.6282 | 0.01967698 | Credible Set | purple |
| chr2:158631398 | 2 | 158.6314 | 0.01160882 | Credible Set | purple |
| chr2:158631624 | 2 | 158.6316 | 0.02223644 | Credible Set | purple |
| chr2:158640014 | 2 | 158.6400 | 0.02809391 | Credible Set | purple |
| chr2:158665126 | 2 | 158.6651 | 0.02633361 | Credible Set | purple |

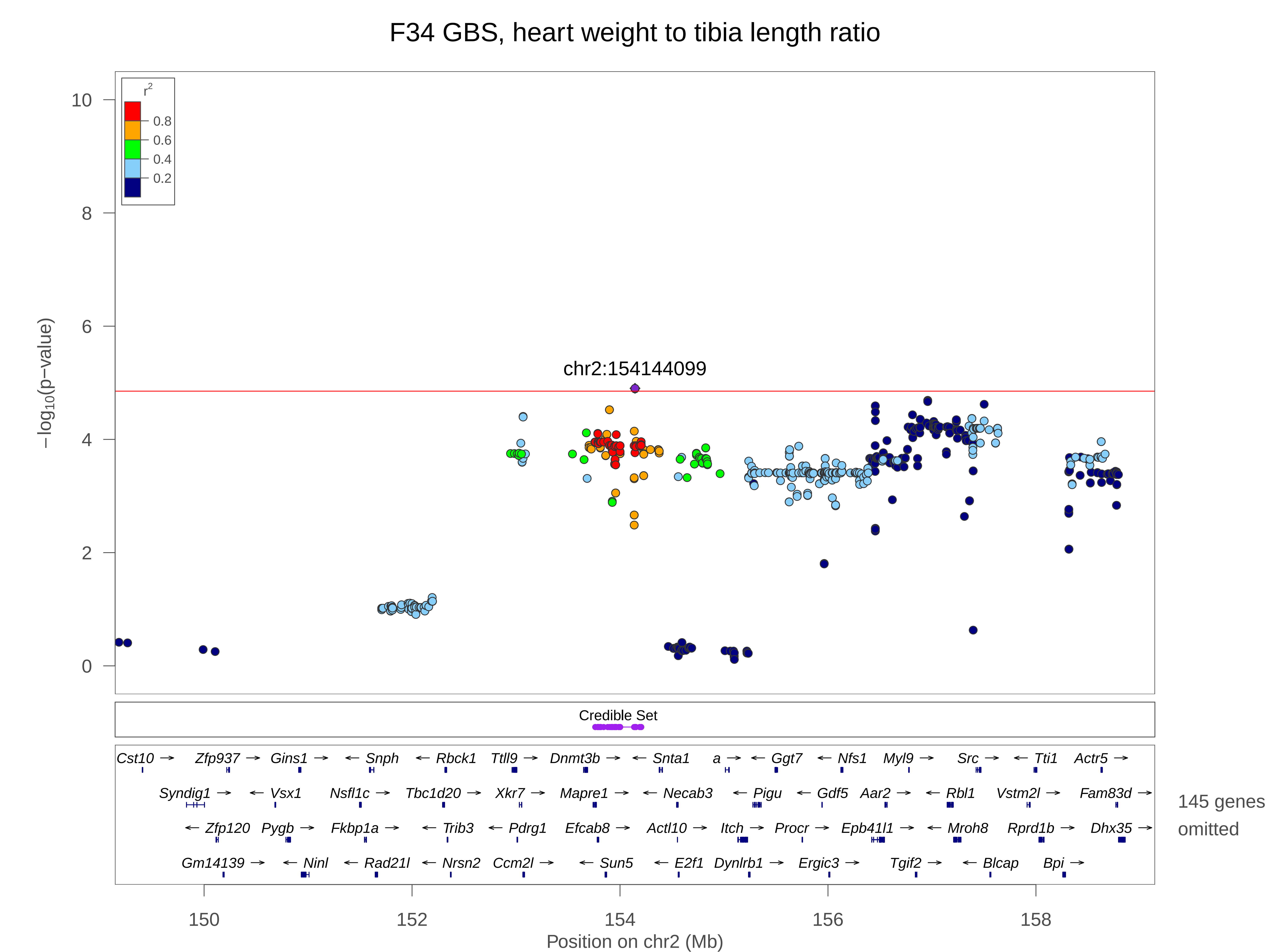
F34 GBS, heart weight to tibia length ratio
r2
0.8
0.6
0.4
0.2
10
8
6
4
2
0
−log10(p−value)
chr2:154144099
Credible Set
Cst10 →
Zfp937 →
Gins1 →
← Snph
← Rbck1
Ttll9 →
Dnmt3b →
← Snta1
a →
← Ggt7
← Nfs1
Myl9 →
Src →
← Tti1
Actr5 →
Syndig1 →
← Vsx1
Nsfl1c →
Tbc1d20 →
Xkr7 →
Mapre1 →
← Necab3
← Pigu
← Gdf5
Aar2 →
← Rbl1
Vstm2l →
Fam83d →
← Zfp120
Pygb →
Fkbp1a →
← Trib3
← Pdrg1
Efcab8 →
Actl10 →
Itch →
Procr →
Epb41l1 →
← Mroh8
Rprd1b →
Dhx35 →
Gm14139 →
← Ninl
← Rad21l
← Nrsn2
Ccm2l →
← Sun5
← E2f1
Dynlrb1 →
Ergic3 →
Tgif2 →
← Blcap
Bpi →
145 genes omitted
150
152
154
156
158
Position on chr2 (Mb)

date: Thu Oct 25 17:15:09 2018
build: F34GBS
display range: chr2:149144099−159144099 [149144099−159144099]
hilite range: 0 − 0 [ 0 − 0 ]
reference SNP: chr2:154144099
number of SNPs plotted: 751
min P−value: 1.25E−5 [chr2:154144099]
max P−value: 7.71E−1 [chr2:155099522]
omitted Genes: Gm14124, 3300002I08Rik, Zfp442
omitted Genes: Zfp345, Cst7, Apmap
omitted Genes: Acss1, E130215H24Rik, Entpd6
omitted Genes: Abhd12, Nanp, Gm14151
omitted Genes: 4930442J19Rik, LOC100125594, 4921509C19Rik
omitted Genes: Sdcbp2, Tmem74b, Psmf1
omitted Genes: Rspo4, Angpt4, Mir1953
omitted Genes: Fam110a, Slc52a3, Scrt2
omitted Genes: Srxn1, Tcf15, Csnk2a1
omitted Genes: Gm14164, Sox12, Zcchc3
omitted Genes: 6820408C15Rik, Defb23, Defb20
omitted Genes: Defb22, Defb26, Defb28
omitted Genes: Defb29, Defb21, Defb19
omitted Genes: Defb45, Defb36, Defb25
omitted Genes: Rem1, H13, Mcts2
omitted Genes: Id1, Cox4i2, Bcl2l1
omitted Genes: Tpx2, Mir5622, Mylk2
omitted Genes: Foxs1, Dusp15, Hck
omitted Genes: Tm9sf4, Tspyl3, Plagl2
omitted Genes: Pofut1, Kif3b, 2500004C02Rik
omitted Genes: Asxl1, Nol4l, Commd7
omitted Genes: Bpifb2, Bpifb6, Bpifb3
omitted Genes: Bpifb4, Bpifa6, Bpifa2
omitted Genes: Bpifa3, Bpifa1, Bpifa5
omitted Genes: Bpifb1, Bpifb5, Bpifb9a
omitted Genes: Bpifb9b, Cdk5rap1, Cbfa2t2
omitted Genes: 1700003F12Rik, Pxmp4, Zfp341
omitted Genes: Chmp4b, Raly, Eif2s2
omitted Genes: Ahcy, 2310005A03Rik, Map1lc3a
omitted Genes: Mir695, Trp53inp2, Ncoa6
omitted Genes: Acss2, Acss2os, Gss
Make more plots at http://csg.sph.umich.edu/locuszoom/
omitted Genes: Myh7b, Mir499, Trpc4ap

Fine−mapping Regions

| snp | chr | pos (Mb) | pp | group | color |
|---|---|---|---|---|---|
| chr2:153762531 | 2 | 153.7625 | 0.014340562 | Credible Set | purple |
| chr2:153762617 | 2 | 153.7626 | 0.014736630 | Credible Set | purple |
| chr2:153786486 | 2 | 153.7865 | 0.020275435 | Credible Set | purple |
| chr2:153786488 | 2 | 153.7865 | 0.020518988 | Credible Set | purple |
| chr2:153786535 | 2 | 153.7865 | 0.020202108 | Credible Set | purple |
| chr2:153787701 | 2 | 153.7877 | 0.014736630 | Credible Set | purple |
| chr2:153792798 | 2 | 153.7928 | 0.014736630 | Credible Set | purple |
| chr2:153792843 | 2 | 153.7928 | 0.013812327 | Credible Set | purple |
| chr2:153807997 | 2 | 153.8080 | 0.014736630 | Credible Set | purple |
| chr2:153808035 | 2 | 153.8080 | 0.014736630 | Credible Set | purple |
| chr2:153810771 | 2 | 153.8108 | 0.014719244 | Credible Set | purple |
| chr2:153816574 | 2 | 153.8166 | 0.014736630 | Credible Set | purple |
| chr2:153843032 | 2 | 153.8430 | 0.014736630 | Credible Set | purple |
| chr2:153881660 | 2 | 153.8817 | 0.015113659 | Credible Set | purple |
| chr2:153898207 | 2 | 153.8982 | 0.012900073 | Credible Set | purple |
| chr2:153913470 | 2 | 153.9135 | 0.012865507 | Credible Set | purple |
| chr2:153913717 | 2 | 153.9137 | 0.012852460 | Credible Set | purple |
| chr2:153921306 | 2 | 153.9213 | 0.013110608 | Credible Set | purple |
| chr2:153928744 | 2 | 153.9287 | 0.011050302 | Credible Set | purple |
| chr2:153929140 | 2 | 153.9291 | 0.010017034 | Credible Set | purple |
| chr2:153947803 | 2 | 153.9478 | 0.012681654 | Credible Set | purple |
| chr2:153947918 | 2 | 153.9479 | 0.006271755 | Credible Set | purple |
| chr2:153947973 | 2 | 153.9480 | 0.006400102 | Credible Set | purple |
| chr2:153950405 | 2 | 153.9504 | 0.012336003 | Credible Set | purple |
| chr2:153950802 | 2 | 153.9508 | 0.012424706 | Credible Set | purple |
| chr2:153952188 | 2 | 153.9522 | 0.007447154 | Credible Set | purple |
| chr2:153952255 | 2 | 153.9523 | 0.007561241 | Credible Set | purple |
| chr2:153957833 | 2 | 153.9578 | 0.006147223 | Credible Set | purple |
| chr2:153957885 | 2 | 153.9579 | 0.006147223 | Credible Set | purple |
| chr2:153957890 | 2 | 153.9579 | 0.006147223 | Credible Set | purple |
| chr2:153963486 | 2 | 153.9635 | 0.012786933 | Credible Set | purple |
| chr2:153963504 | 2 | 153.9635 | 0.012786933 | Credible Set | purple |
| chr2:153963562 | 2 | 153.9636 | 0.019575345 | Credible Set | purple |
| chr2:153987087 | 2 | 153.9871 | 0.011947686 | Credible Set | purple |
| chr2:153987540 | 2 | 153.9875 | 0.012207191 | Credible Set | purple |
| chr2:154000310 | 2 | 154.0003 | 0.010154622 | Credible Set | purple |

Fine−mapping Regions

| snp | chr | pos (Mb) | pp | group | color |
|---|---|---|---|---|---|
| chr2:154000346 | 2 | 154.0003 | 0.010182767 | Credible Set | purple |
| chr2:154000704 | 2 | 154.0007 | 0.012786933 | Credible Set | purple |
| chr2:154000755 | 2 | 154.0008 | 0.012786933 | Credible Set | purple |
| chr2:154000791 | 2 | 154.0008 | 0.012786933 | Credible Set | purple |
| chr2:154135816 | 2 | 154.1358 | 0.012786933 | Credible Set | purple |
| chr2:154143827 | 2 | 154.1438 | 0.009769867 | Credible Set | purple |
| chr2:154144099 | 2 | 154.1441 | 0.116950121 | Credible Set | purple |
| chr2:154144100 | 2 | 154.1441 | 0.115191612 | Credible Set | purple |
| chr2:154144142 | 2 | 154.1441 | 0.114572675 | Credible Set | purple |
| chr2:154152208 | 2 | 154.1522 | 0.012786933 | Credible Set | purple |
| chr2:154152238 | 2 | 154.1522 | 0.012786933 | Credible Set | purple |
| chr2:154152320 | 2 | 154.1523 | 0.012249480 | Credible Set | purple |
| chr2:154189385 | 2 | 154.1894 | 0.012628698 | Credible Set | purple |
| chr2:154189574 | 2 | 154.1896 | 0.012778016 | Credible Set | purple |
| chr2:154203292 | 2 | 154.2033 | 0.013706541 | Credible Set | purple |
| chr2:154203302 | 2 | 154.2033 | 0.014920169 | Credible Set | purple |
| chr2:154203329 | 2 | 154.2033 | 0.013262199 | Credible Set | purple |
| chr2:154203418 | 2 | 154.2034 | 0.012961320 | Credible Set | purple |

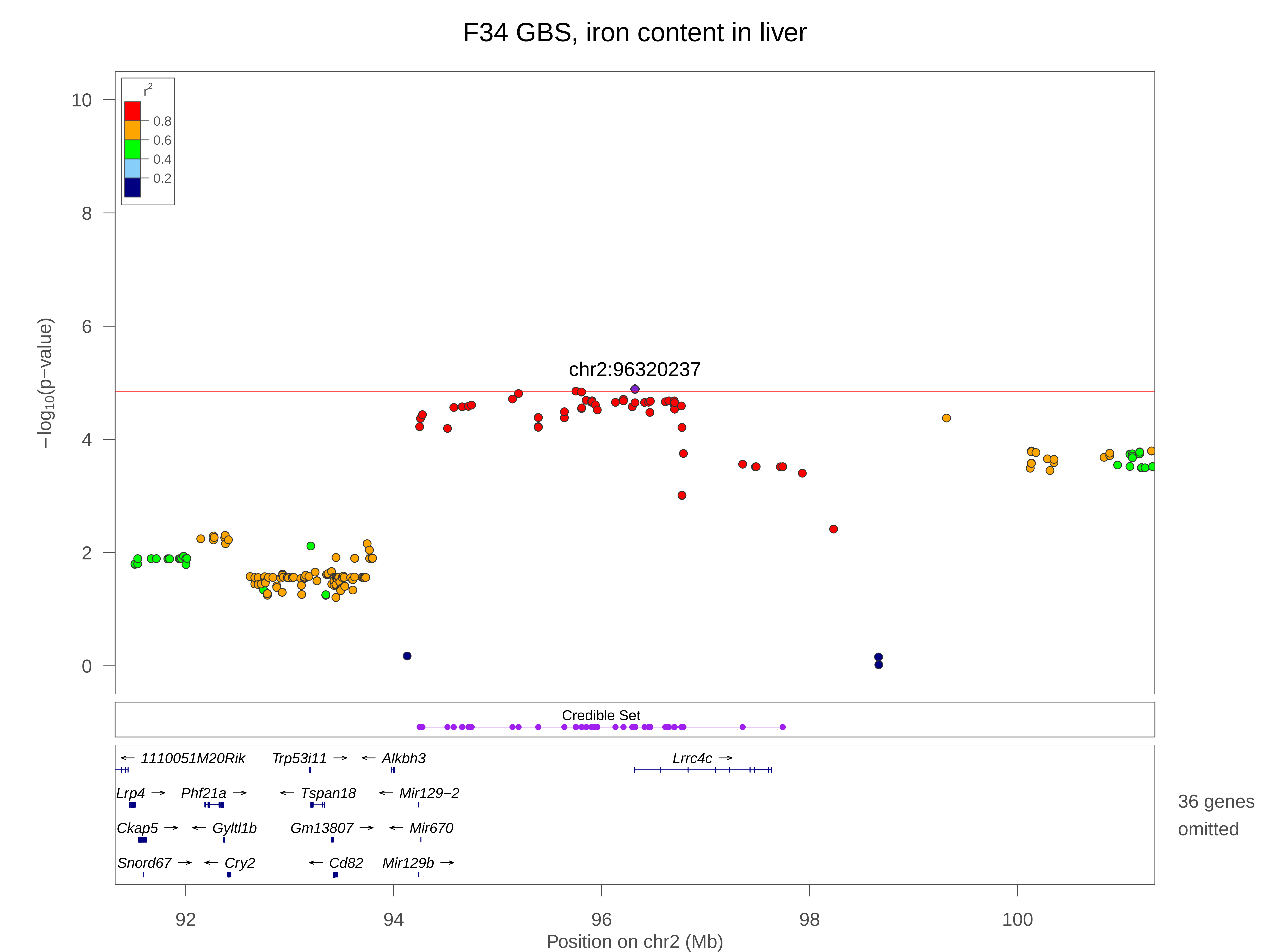

F34 GBS, iron content in liver
r2
0.8
0.6
0.4
0.2
10
8
6
4
2
0
−log10(p−value)
chr2:96320237
Credible Set
← 1110051M20Rik
Trp53i11 →
← Alkbh3
Lrrc4c →
Lrp4 →
Phf21a →
← Tspan18
← Mir129−2
Ckap5 →
← Gyltl1b
Gm13807 →
← Mir670
Snord67 →
← Cry2
← Cd82
Mir129b →
36 genes omitted
92
94
96
98
100
Position on chr2 (Mb)

| | |
|---|---|
| date: | Thu Oct 25 17:15:50 2018 |
| build: | F34GBS |
| display range: | chr2:91320237−101320237 [91320237−101320237] |
| hilite range: | 0 − 0 [ 0 − 0 ] |
| reference SNP: | chr2:96320237 |
| number of SNPs plotted: | 236 |
| min P−value: | 1.29E−5 [chr2:96320237] |
| max P−value: | 9.58E−1 [chr2:98665093] |
| omitted Genes: | F2, Zfp408, Arhgap1 |
| omitted Genes: | Atg13, Harbi1, Ambra1 |
| omitted Genes: | Chrm4, Mdk, Dgkz |
| omitted Genes: | Mir6999, Creb3l1, Mir1955 |
| omitted Genes: | Pex16, 1700029I15Rik, Mapk8ip1 |
| omitted Genes: | Mir7000, D930015M05Rik, Slc35c1 |
| omitted Genes: | Mir7221, Mir7222, Chst1 |
| omitted Genes: | Syt13, Prdm11, 4631405J19Rik |
| omitted Genes: | Mir7001, Gm10804, Alx4 |
| omitted Genes: | Ext2, Accs, Accsl |
| omitted Genes: | Gm13889, Alkbh3os1, Hsd17b12 |
| omitted Genes: | Ttc17, 2810002D19Rik, Api5 |

Make more plots at http://csg.sph.umich.edu/locuszoom/

Fine−mapping Regions

| snp | chr | pos (Mb) | pp | group | color |
|---|---|---|---|---|---|
| chr2:94249122 | 2 | 94.24912 | 0.008286008 | Credible Set | purple |
| chr2:94257379 | 2 | 94.25738 | 0.011351696 | Credible Set | purple |
| chr2:94276010 | 2 | 94.27601 | 0.013199881 | Credible Set | purple |
| chr2:94516977 | 2 | 94.51698 | 0.007744163 | Credible Set | purple |
| chr2:94577500 | 2 | 94.57750 | 0.017396457 | Credible Set | purple |
| chr2:94657941 | 2 | 94.65794 | 0.017833975 | Credible Set | purple |
| chr2:94718556 | 2 | 94.71856 | 0.018139036 | Credible Set | purple |
| chr2:94748712 | 2 | 94.74871 | 0.019053454 | Credible Set | purple |
| chr2:95141904 | 2 | 95.14190 | 0.024067265 | Credible Set | purple |
| chr2:95200398 | 2 | 95.20040 | 0.029835579 | Credible Set | purple |
| chr2:95390243 | 2 | 95.39024 | 0.008220943 | Credible Set | purple |
| chr2:95390247 | 2 | 95.39025 | 0.008090195 | Credible Set | purple |
| chr2:95390277 | 2 | 95.39028 | 0.008301131 | Credible Set | purple |
| chr2:95390284 | 2 | 95.39028 | 0.008117626 | Credible Set | purple |
| chr2:95390546 | 2 | 95.39055 | 0.011839594 | Credible Set | purple |
| chr2:95390592 | 2 | 95.39059 | 0.011751530 | Credible Set | purple |
| chr2:95641124 | 2 | 95.64112 | 0.011697477 | Credible Set | purple |
| chr2:95641477 | 2 | 95.64148 | 0.014740073 | Credible Set | purple |
| chr2:95752277 | 2 | 95.75228 | 0.032769533 | Credible Set | purple |
| chr2:95805867 | 2 | 95.80587 | 0.016687093 | Credible Set | purple |
| chr2:95806028 | 2 | 95.80603 | 0.031502905 | Credible Set | purple |
| chr2:95806038 | 2 | 95.80604 | 0.031743782 | Credible Set | purple |
| chr2:95808603 | 2 | 95.80860 | 0.017109349 | Credible Set | purple |
| chr2:95850674 | 2 | 95.85067 | 0.022997969 | Credible Set | purple |
| chr2:95900229 | 2 | 95.90023 | 0.021299943 | Credible Set | purple |
| chr2:95906738 | 2 | 95.90674 | 0.022491984 | Credible Set | purple |
| chr2:95906783 | 2 | 95.90678 | 0.021603954 | Credible Set | purple |
| chr2:95938782 | 2 | 95.93878 | 0.019239869 | Credible Set | purple |
| chr2:95957448 | 2 | 95.95745 | 0.015789929 | Credible Set | purple |
| chr2:96132587 | 2 | 96.13259 | 0.021193097 | Credible Set | purple |
| chr2:96209134 | 2 | 96.20913 | 0.023810661 | Credible Set | purple |
| chr2:96209157 | 2 | 96.20916 | 0.022461346 | Credible Set | purple |
| chr2:96293855 | 2 | 96.29385 | 0.017860711 | Credible Set | purple |
| chr2:96320222 | 2 | 96.32022 | 0.035016259 | Credible Set | purple |
| chr2:96320237 | 2 | 96.32024 | 0.035385938 | Credible Set | purple |
| chr2:96320641 | 2 | 96.32064 | 0.020864650 | Credible Set | purple |

Fine−mapping Regions

| snp | chr | pos (Mb) | pp | group | color |
|---|---|---|---|---|---|
| chr2:96412425 | 2 | 96.41242 | 0.021107363 | Credible Set | purple |
| chr2:96451438 | 2 | 96.45144 | 0.021299943 | Credible Set | purple |
| chr2:96462914 | 2 | 96.46291 | 0.014371330 | Credible Set | purple |
| chr2:96468064 | 2 | 96.46806 | 0.022194194 | Credible Set | purple |
| chr2:96611762 | 2 | 96.61176 | 0.021647155 | Credible Set | purple |
| chr2:96645727 | 2 | 96.64573 | 0.022445691 | Credible Set | purple |
| chr2:96697256 | 2 | 96.69726 | 0.022439873 | Credible Set | purple |
| chr2:96697871 | 2 | 96.69787 | 0.019417377 | Credible Set | purple |
| chr2:96697872 | 2 | 96.69787 | 0.019470185 | Credible Set | purple |
| chr2:96701045 | 2 | 96.70104 | 0.016285920 | Credible Set | purple |
| chr2:96701070 | 2 | 96.70107 | 0.016285920 | Credible Set | purple |
| chr2:96701077 | 2 | 96.70108 | 0.016285920 | Credible Set | purple |
| chr2:96701079 | 2 | 96.70108 | 0.016285920 | Credible Set | purple |
| chr2:96701166 | 2 | 96.70117 | 0.020975667 | Credible Set | purple |
| chr2:96766199 | 2 | 96.76620 | 0.018478019 | Credible Set | purple |
| chr2:96771761 | 2 | 96.77176 | 0.008045576 | Credible Set | purple |
| chr2:96771791 | 2 | 96.77179 | 0.008026753 | Credible Set | purple |
| chr2:96786046 | 2 | 96.78605 | 0.002958953 | Credible Set | purple |
| chr2:97355566 | 2 | 97.35557 | 0.001965050 | Credible Set | purple |
| chr2:97741014 | 2 | 97.74101 | 0.001788228 | Credible Set | purple |

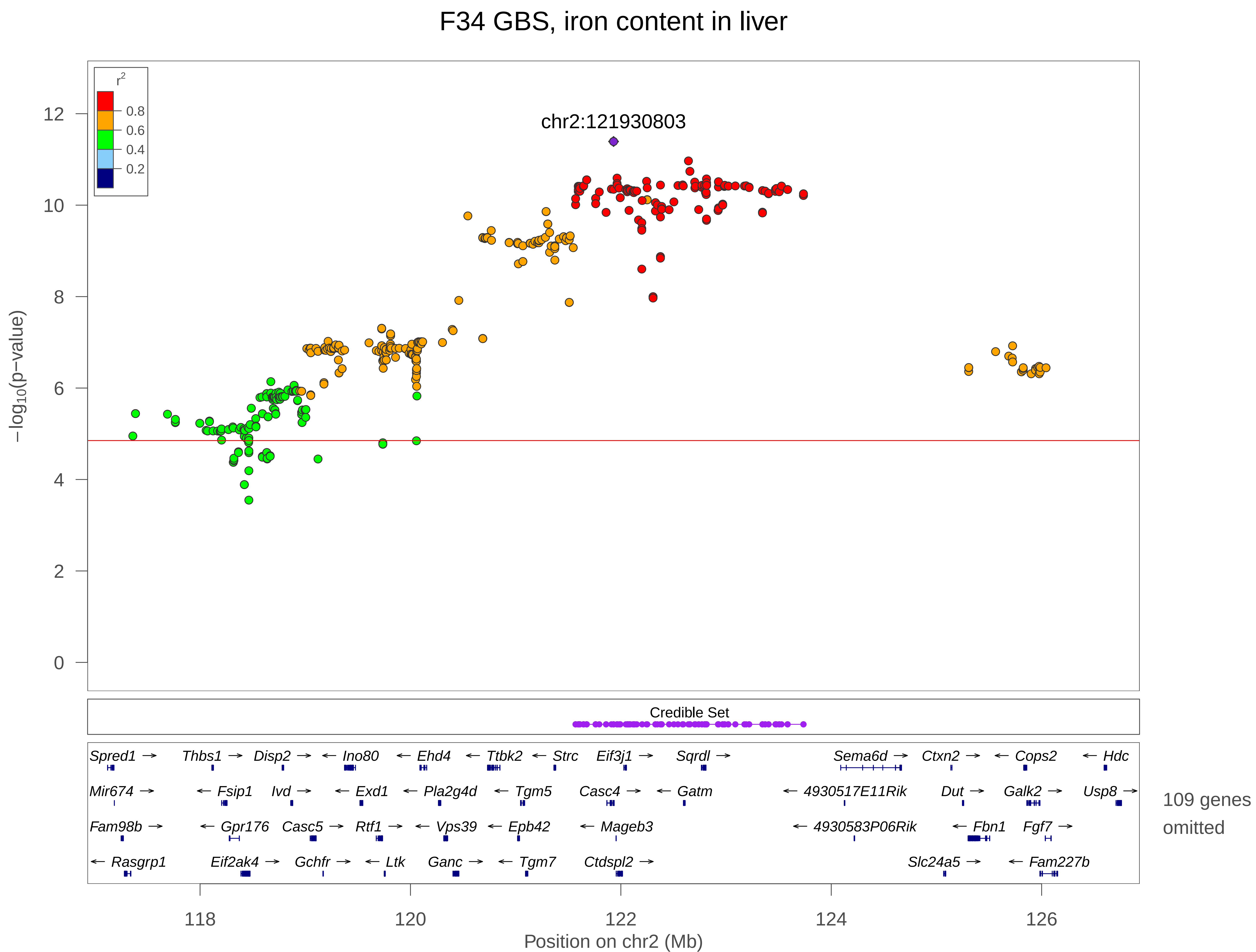
F34 GBS, iron content in liver
r2
0.8
0.6
0.4
0.2
chr2:121930803
−log10(p−value)
0
2
4
6
8
10
12
Credible Set
Spred1 →
Thbs1 →
Disp2 →
← Ino80
← Ehd4
← Ttbk2
← Strc
Eif3j1 →
Sqrdl →
Sema6d →
Ctxn2 →
← Cops2
← Hdc
Mir674 →
← Fsip1
Ivd →
← Exd1
← Pla2g4d
← Tgm5
Casc4 →
← Gatm
← 4930517E11Rik
Dut →
Galk2 →
Usp8 →
Fam98b →
← Gpr176
Casc5 →
Rtf1 →
← Vps39
← Epb42
← Mageb3
← 4930583P06Rik
← Fbn1
Fgf7 →
← Rasgrp1
Eif2ak4 →
Gchfr →
← Ltk
Ganc →
← Tgm7
Ctdspl2 →
Slc24a5 →
← Fam227b
109 genes omitted
118
120
122
124
126
Position on chr2 (Mb)

| | |
|---|---|
| date: | Thu Oct 25 17:15:40 2018 |
| build: | F34GBS |
| display range: | chr2:116930803−126930803 [116930803−126930803] |
| hilite range: | 0 − 0 [ 0 − 0 ] |
| reference SNP: | chr2:121930803 |
| number of SNPs plotted: | 530 |
| min P−value: | 4.06E−12 [chr2:121930803] |
| max P−value: | 2.83E−4 [chr2:118463923] |
| omitted Genes: | Gm13985, 4930412B13Rik, Gm13986 |
| omitted Genes: | 1700054M17Rik, Srp14, Bmf |
| omitted Genes: | Bub1b, Pak6, Ankrd63 |
| omitted Genes: | Plcb2, Inafm2, A430105I19Rik |
| omitted Genes: | Phgr1, Knstrn, Bahd1 |
| omitted Genes: | Chst14, Ccdc32, Rpusd2 |
| omitted Genes: | Rad51, Rmdn3, Dnajc17 |
| omitted Genes: | Gm14137, Zfyve19, Ppp1r14d |
| omitted Genes: | Spint1, Rhov, Vps18 |
| omitted Genes: | Gm14207, Dll4, Chac1 |
| omitted Genes: | Chp1, 1700020I14Rik, Oip5 |
| omitted Genes: | Nusap1, Ndufaf1, Itpka |
| omitted Genes: | Rpap1, Tyro3, Mga |
| omitted Genes: | Mapkbp1, Mir7665, Jmjd7 |
| omitted Genes: | Gm28042, Pla2g4b, Pla2g4e |
| omitted Genes: | Pla2g4f, Tmem87a, 4931402G19Rik |
| omitted Genes: | Capn3, Zfp106, Snap23 |
| omitted Genes: | Lrrc57, Haus2, Cdan1 |
| omitted Genes: | AV039307, Ubr1, Tmem62 |
| omitted Genes: | Ccndbp1, Lcmt2, Adal |
| omitted Genes: | Zscan29, Tubgcp4, Trp53bp1 |
| omitted Genes: | Map1a, Ppip5k1, Ckmt1 |
| omitted Genes: | Catsper2, Pdia3, Ell3 |
| omitted Genes: | Serf2, Serinc4, Hypk |
| omitted Genes: | Mfap1b, Mfap1a, Wdr76 |
| omitted Genes: | Frmd5, Spg11, Patl2 |
| omitted Genes: | B2m, Trim69, Terb2 |
| omitted Genes: | Sord, Duox2, Duoxa2 |
| omitted Genes: | Duoxa1, Duox1, Shf |
| omitted Genes: | Slc28a2, Bambi−ps1, Gm14085 |
| omitted Genes: | Spata5l1, AA467197, Mir147 |
| omitted Genes: | Slc30a4, Bloc1s6, Myef2 |

Make more plots at http://csg.sph.umich.edu/locuszoom/

Fine−mapping Regions

| snp | chr | pos (Mb) | pp | group | color |
|---|---|---|---|---|---|
| chr2:121569629 | 2 | 121.5696 | 0.003133483 | Credible Set | purple |
| chr2:121569951 | 2 | 121.5700 | 0.003145795 | Credible Set | purple |
| chr2:121570025 | 2 | 121.5700 | 0.004232924 | Credible Set | purple |
| chr2:121595218 | 2 | 121.5952 | 0.007755888 | Credible Set | purple |
| chr2:121595356 | 2 | 121.5954 | 0.007755888 | Credible Set | purple |
| chr2:121595424 | 2 | 121.5954 | 0.006025871 | Credible Set | purple |
| chr2:121595633 | 2 | 121.5956 | 0.006561241 | Credible Set | purple |
| chr2:121605090 | 2 | 121.6051 | 0.007755888 | Credible Set | purple |
| chr2:121605163 | 2 | 121.6052 | 0.007248706 | Credible Set | purple |
| chr2:121609496 | 2 | 121.6095 | 0.007629410 | Credible Set | purple |
| chr2:121609992 | 2 | 121.6100 | 0.006080155 | Credible Set | purple |
| chr2:121610054 | 2 | 121.6101 | 0.007039792 | Credible Set | purple |
| chr2:121644474 | 2 | 121.6445 | 0.007755888 | Credible Set | purple |
| chr2:121644502 | 2 | 121.6445 | 0.007755888 | Credible Set | purple |
| chr2:121647254 | 2 | 121.6473 | 0.007985426 | Credible Set | purple |
| chr2:121675210 | 2 | 121.6752 | 0.010662479 | Credible Set | purple |
| chr2:121675212 | 2 | 121.6752 | 0.010662479 | Credible Set | purple |
| chr2:121675249 | 2 | 121.6752 | 0.010662479 | Credible Set | purple |
| chr2:121675251 | 2 | 121.6753 | 0.010662479 | Credible Set | purple |
| chr2:121760589 | 2 | 121.7606 | 0.004296592 | Credible Set | purple |
| chr2:121760592 | 2 | 121.7606 | 0.004296592 | Credible Set | purple |
| chr2:121760608 | 2 | 121.7606 | 0.004296592 | Credible Set | purple |
| chr2:121760655 | 2 | 121.7607 | 0.003294660 | Credible Set | purple |
| chr2:121760675 | 2 | 121.7607 | 0.003294660 | Credible Set | purple |
| chr2:121795014 | 2 | 121.7950 | 0.005887328 | Credible Set | purple |
| chr2:121859536 | 2 | 121.8595 | 0.002151535 | Credible Set | purple |
| chr2:121859542 | 2 | 121.8595 | 0.002143376 | Credible Set | purple |
| chr2:121910457 | 2 | 121.9105 | 0.006793285 | Credible Set | purple |
| chr2:121930722 | 2 | 121.9307 | 0.006694281 | Credible Set | purple |
| chr2:121930803 | 2 | 121.9308 | 0.070714562 | Credible Set | purple |
| chr2:121930806 | 2 | 121.9308 | 0.070714562 | Credible Set | purple |
| chr2:121963958 | 2 | 121.9640 | 0.011635947 | Credible Set | purple |
| chr2:121964022 | 2 | 121.9640 | 0.008924639 | Credible Set | purple |
| chr2:121964577 | 2 | 121.9646 | 0.008234906 | Credible Set | purple |
| chr2:121982399 | 2 | 121.9824 | 0.007118213 | Credible Set | purple |
| chr2:121994315 | 2 | 121.9943 | 0.004434478 | Credible Set | purple |

Fine−mapping Regions

| snp | chr | pos (Mb) | pp | group | color |
|---|---|---|---|---|---|
| chr2:121994324 | 2 | 121.9943 | 0.004434478 | Credible Set | purple |
| chr2:122047663 | 2 | 122.0477 | 0.006586628 | Credible Set | purple |
| chr2:122060688 | 2 | 122.0607 | 0.006885602 | Credible Set | purple |
| chr2:122061099 | 2 | 122.0611 | 0.005927683 | Credible Set | purple |
| chr2:122065787 | 2 | 122.0658 | 0.006477903 | Credible Set | purple |
| chr2:122065870 | 2 | 122.0659 | 0.006301764 | Credible Set | purple |
| chr2:122077766 | 2 | 122.0778 | 0.002370989 | Credible Set | purple |
| chr2:122077798 | 2 | 122.0778 | 0.002370989 | Credible Set | purple |
| chr2:122083136 | 2 | 122.0831 | 0.006504421 | Credible Set | purple |
| chr2:122090229 | 2 | 122.0902 | 0.006422790 | Credible Set | purple |
| chr2:122115252 | 2 | 122.1153 | 0.005878040 | Credible Set | purple |
| chr2:122122941 | 2 | 122.1229 | 0.005690708 | Credible Set | purple |
| chr2:122123313 | 2 | 122.1233 | 0.006247165 | Credible Set | purple |
| chr2:122132763 | 2 | 122.1328 | 0.006087368 | Credible Set | purple |
| chr2:122151119 | 2 | 122.1511 | 0.006149327 | Credible Set | purple |
| chr2:122202473 | 2 | 122.2025 | 0.003852653 | Credible Set | purple |
| chr2:122244672 | 2 | 122.2447 | 0.009937406 | Credible Set | purple |
| chr2:122250299 | 2 | 122.2503 | 0.007191398 | Credible Set | purple |
| chr2:122328643 | 2 | 122.3286 | 0.002299839 | Credible Set | purple |
| chr2:122328650 | 2 | 122.3286 | 0.003475004 | Credible Set | purple |
| chr2:122346906 | 2 | 122.3469 | 0.003101151 | Credible Set | purple |
| chr2:122376352 | 2 | 122.3764 | 0.001713187 | Credible Set | purple |
| chr2:122376463 | 2 | 122.3765 | 0.008258864 | Credible Set | purple |
| chr2:122387979 | 2 | 122.3880 | 0.002883818 | Credible Set | purple |
| chr2:122388097 | 2 | 122.3881 | 0.002561703 | Credible Set | purple |
| chr2:122388208 | 2 | 122.3882 | 0.002514832 | Credible Set | purple |
| chr2:122458764 | 2 | 122.4588 | 0.002451155 | Credible Set | purple |
| chr2:122504055 | 2 | 122.5041 | 0.003608100 | Credible Set | purple |
| chr2:122542898 | 2 | 122.5429 | 0.008070659 | Credible Set | purple |
| chr2:122587284 | 2 | 122.5873 | 0.008272898 | Credible Set | purple |
| chr2:122587743 | 2 | 122.5877 | 0.008298741 | Credible Set | purple |
| chr2:122593432 | 2 | 122.5934 | 0.007882663 | Credible Set | purple |
| chr2:122642503 | 2 | 122.6425 | 0.027157517 | Credible Set | purple |
| chr2:122656435 | 2 | 122.6564 | 0.016242538 | Credible Set | purple |
| chr2:122656458 | 2 | 122.6565 | 0.016242538 | Credible Set | purple |
| chr2:122702118 | 2 | 122.7021 | 0.009577217 | Credible Set | purple |

Fine−mapping Regions

| snp | chr | pos (Mb) | pp | group | color |
|---|---|---|---|---|---|
| chr2:122704152 | 2 | 122.7042 | 0.007164023 | Credible Set | purple |
| chr2:122708239 | 2 | 122.7082 | 0.007755888 | Credible Set | purple |
| chr2:122739988 | 2 | 122.7400 | 0.002470998 | Credible Set | purple |
| chr2:122740035 | 2 | 122.7400 | 0.002469759 | Credible Set | purple |
| chr2:122771056 | 2 | 122.7711 | 0.007366102 | Credible Set | purple |
| chr2:122771465 | 2 | 122.7715 | 0.008149988 | Credible Set | purple |
| chr2:122797192 | 2 | 122.7972 | 0.007263408 | Credible Set | purple |
| chr2:122797216 | 2 | 122.7972 | 0.007219008 | Credible Set | purple |
| chr2:122797265 | 2 | 122.7973 | 0.008181448 | Credible Set | purple |
| chr2:122797266 | 2 | 122.7973 | 0.008197958 | Credible Set | purple |
| chr2:122804433 | 2 | 122.8044 | 0.005736508 | Credible Set | purple |
| chr2:122811212 | 2 | 122.8112 | 0.005168723 | Credible Set | purple |
| chr2:122811304 | 2 | 122.8113 | 0.005727219 | Credible Set | purple |
| chr2:122814314 | 2 | 122.8143 | 0.011175145 | Credible Set | purple |
| chr2:122814326 | 2 | 122.8143 | 0.008402303 | Credible Set | purple |
| chr2:122814381 | 2 | 122.8144 | 0.008531467 | Credible Set | purple |
| chr2:122814728 | 2 | 122.8147 | 0.009274010 | Credible Set | purple |
| chr2:122814902 | 2 | 122.8149 | 0.008132534 | Credible Set | purple |
| chr2:122814909 | 2 | 122.8149 | 0.008149358 | Credible Set | purple |
| chr2:122924669 | 2 | 122.9247 | 0.002361641 | Credible Set | purple |
| chr2:122924685 | 2 | 122.9247 | 0.002371527 | Credible Set | purple |
| chr2:122924695 | 2 | 122.9247 | 0.002361641 | Credible Set | purple |
| chr2:122924696 | 2 | 122.9247 | 0.002361641 | Credible Set | purple |
| chr2:122927536 | 2 | 122.9275 | 0.007485543 | Credible Set | purple |
| chr2:122927903 | 2 | 122.9279 | 0.009534543 | Credible Set | purple |
| chr2:122927980 | 2 | 122.9280 | 0.009752696 | Credible Set | purple |
| chr2:122927997 | 2 | 122.9280 | 0.002683199 | Credible Set | purple |
| chr2:122928025 | 2 | 122.9280 | 0.002539716 | Credible Set | purple |
| chr2:122968234 | 2 | 122.9682 | 0.003161310 | Credible Set | purple |
| chr2:122968260 | 2 | 122.9683 | 0.003230687 | Credible Set | purple |
| chr2:122968275 | 2 | 122.9683 | 0.003075386 | Credible Set | purple |
| chr2:122968296 | 2 | 122.9683 | 0.003088519 | Credible Set | purple |
| chr2:122983867 | 2 | 122.9839 | 0.007704921 | Credible Set | purple |
| chr2:122986477 | 2 | 122.9865 | 0.007941141 | Credible Set | purple |
| chr2:122986971 | 2 | 122.9870 | 0.008167152 | Credible Set | purple |
| chr2:123021071 | 2 | 123.0211 | 0.007846689 | Credible Set | purple |

Fine−mapping Regions

| snp | chr | pos (Mb) | pp | group | color |
|---|---|---|---|---|---|
| chr2:123021090 | 2 | 123.0211 | 0.007872734 | Credible Set | purple |
| chr2:123021532 | 2 | 123.0215 | 0.007823060 | Credible Set | purple |
| chr2:123088225 | 2 | 123.0882 | 0.007912931 | Credible Set | purple |
| chr2:123175096 | 2 | 123.1751 | 0.007879959 | Credible Set | purple |
| chr2:123190496 | 2 | 123.1905 | 0.007960532 | Credible Set | purple |
| chr2:123219436 | 2 | 123.2194 | 0.007469322 | Credible Set | purple |
| chr2:123219468 | 2 | 123.2195 | 0.007281566 | Credible Set | purple |
| chr2:123346239 | 2 | 123.3462 | 0.002171426 | Credible Set | purple |
| chr2:123346253 | 2 | 123.3463 | 0.002090464 | Credible Set | purple |
| chr2:123346704 | 2 | 123.3467 | 0.006300361 | Credible Set | purple |
| chr2:123371857 | 2 | 123.3719 | 0.006120310 | Credible Set | purple |
| chr2:123371991 | 2 | 123.3720 | 0.006120310 | Credible Set | purple |
| chr2:123404160 | 2 | 123.4042 | 0.005368147 | Credible Set | purple |
| chr2:123404632 | 2 | 123.4046 | 0.005499987 | Credible Set | purple |
| chr2:123470044 | 2 | 123.4700 | 0.006042828 | Credible Set | purple |
| chr2:123470368 | 2 | 123.4704 | 0.006689148 | Credible Set | purple |
| chr2:123470382 | 2 | 123.4704 | 0.006688751 | Credible Set | purple |
| chr2:123478720 | 2 | 123.4787 | 0.006981594 | Credible Set | purple |
| chr2:123505482 | 2 | 123.5055 | 0.005959121 | Credible Set | purple |
| chr2:123505557 | 2 | 123.5056 | 0.005930026 | Credible Set | purple |
| chr2:123527223 | 2 | 123.5272 | 0.007863122 | Credible Set | purple |
| chr2:123584403 | 2 | 123.5844 | 0.006645888 | Credible Set | purple |
| chr2:123585516 | 2 | 123.5855 | 0.006613768 | Credible Set | purple |
| chr2:123735848 | 2 | 123.7358 | 0.004937967 | Credible Set | purple |
| chr2:123735871 | 2 | 123.7359 | 0.004946098 | Credible Set | purple |
| chr2:123735958 | 2 | 123.7360 | 0.005374211 | Credible Set | purple |

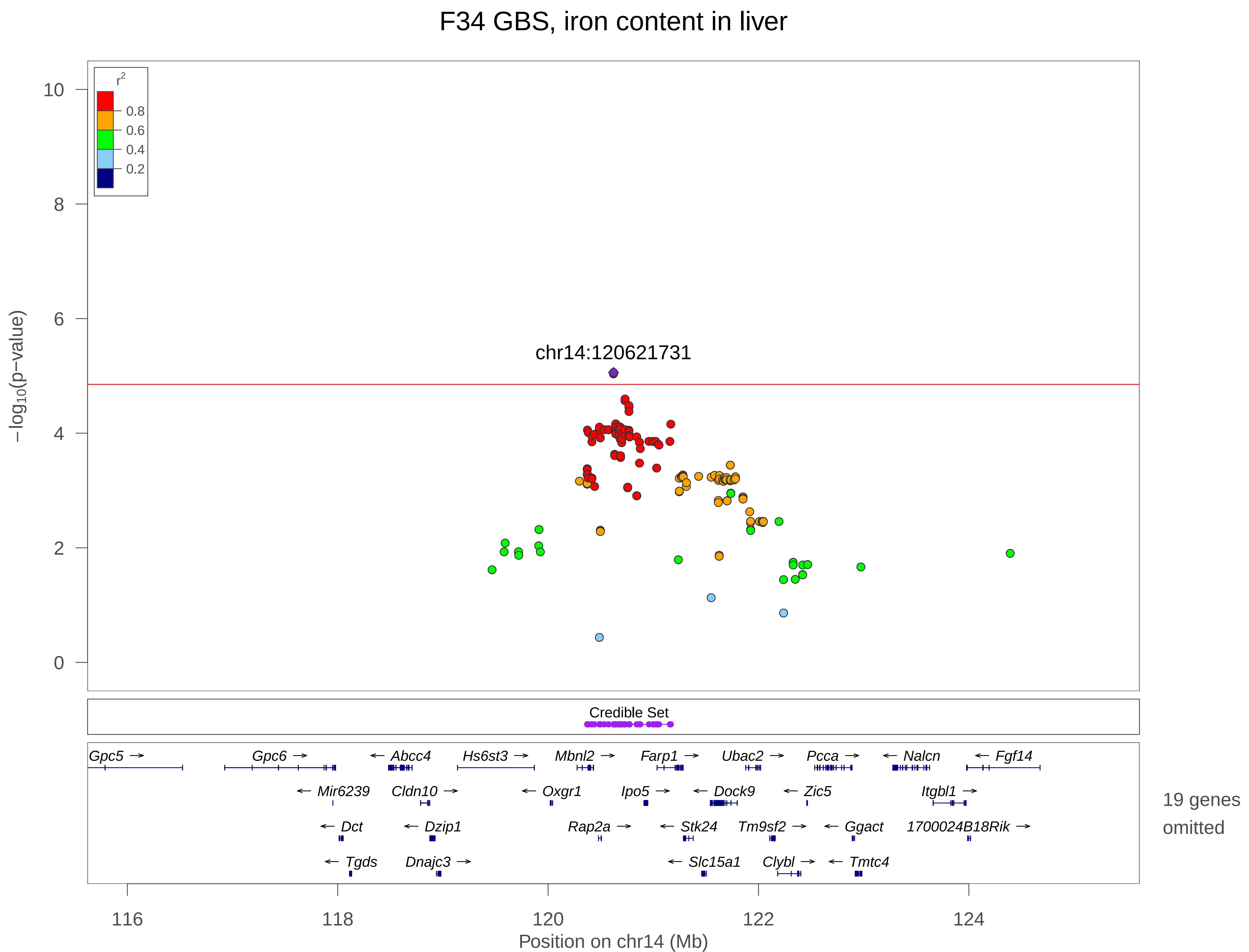
F34 GBS, iron content in liver
r2
0.8
0.6
0.4
0.2
10
8
6
4
2
0
−log10(p−value)
chr14:120621731
Credible Set
Gpc5 →
Gpc6 →
← Abcc4
Hs6st3 →
Mbnl2 →
Farp1 →
Ubac2 →
Pcca →
← Nalcn
← Fgf14
← Mir6239
Cldn10 →
← Oxgr1
Ipo5 →
← Dock9
← Zic5
Itgbl1 →
← Dct
← Dzip1
Rap2a →
← Stk24
Tm9sf2 →
← Ggact
1700024B18Rik →
← Tgds
Dnajc3 →
← Slc15a1
Clybl →
← Tmtc4
19 genes omitted
116
118
120
122
124
Position on chr14 (Mb)

| | |
|---|---|
| date: | Thu Oct 25 17:15:18 2018 |
| build: | F34GBS |
| display range: | chr14:115621731−125621731 [115621731−125621731] |
| hilite range: | 0 − 0 [ 0 − 0 ] |
| reference SNP: | chr14:120621731 |
| number of SNPs plotted: | 206 |
| min P−value: | 8.72E−6 [chr14:120621731] |
| max P−value: | 3.66E−1 [chr14:120487423] |
| omitted Genes: | Gpr180, Sox21, LOC105245869 |
| omitted Genes: | Mir6241, Gm9376, 1700044C05Rik |
| omitted Genes: | Mir6391, Uggt2, 1700006F04Rik |
| omitted Genes: | 1810041H14Rik, Gpr18, Gpr183 |
| omitted Genes: | Timm8a2, A330035P11Rik, 1700108J01Rik |
| omitted Genes: | Gm5089, 2610035F20Rik, Zic2 |
| omitted Genes: | AA536875 |

Make more plots at http://csg.sph.umich.edu/locuszoom/

Fine−mapping Regions

| snp | chr | pos (Mb) | pp | group | color |
|---|---|---|---|---|---|
| chr14:120371692 | 14 | 120.3717 | 0.002269808 | Credible Set | purple |
| chr14:120371706 | 14 | 120.3717 | 0.002191663 | Credible Set | purple |
| chr14:120371729 | 14 | 120.3717 | 0.001849816 | Credible Set | purple |
| chr14:120373900 | 14 | 120.3739 | 0.009793911 | Credible Set | purple |
| chr14:120374171 | 14 | 120.3742 | 0.001784498 | Credible Set | purple |
| chr14:120374181 | 14 | 120.3742 | 0.001837517 | Credible Set | purple |
| chr14:120379882 | 14 | 120.3799 | 0.001610230 | Credible Set | purple |
| chr14:120382338 | 14 | 120.3823 | 0.008887336 | Credible Set | purple |
| chr14:120416102 | 14 | 120.4161 | 0.001607522 | Credible Set | purple |
| chr14:120416136 | 14 | 120.4161 | 0.001558402 | Credible Set | purple |
| chr14:120416333 | 14 | 120.4163 | 0.006266201 | Credible Set | purple |
| chr14:120416347 | 14 | 120.4163 | 0.007748995 | Credible Set | purple |
| chr14:120439342 | 14 | 120.4393 | 0.008325976 | Credible Set | purple |
| chr14:120487392 | 14 | 120.4874 | 0.009982187 | Credible Set | purple |
| chr14:120487416 | 14 | 120.4874 | 0.009973058 | Credible Set | purple |
| chr14:120487566 | 14 | 120.4876 | 0.009973058 | Credible Set | purple |
| chr14:120487639 | 14 | 120.4876 | 0.010923959 | Credible Set | purple |
| chr14:120495956 | 14 | 120.4960 | 0.007265250 | Credible Set | purple |
| chr14:120495976 | 14 | 120.4960 | 0.007352010 | Credible Set | purple |
| chr14:120532718 | 14 | 120.5327 | 0.009935141 | Credible Set | purple |
| chr14:120532740 | 14 | 120.5327 | 0.009935141 | Credible Set | purple |
| chr14:120573821 | 14 | 120.5738 | 0.009873525 | Credible Set | purple |
| chr14:120573951 | 14 | 120.5740 | 0.009935141 | Credible Set | purple |
| chr14:120621718 | 14 | 120.6217 | 0.083468244 | Credible Set | purple |
| chr14:120621731 | 14 | 120.6217 | 0.088133759 | Credible Set | purple |
| chr14:120621749 | 14 | 120.6217 | 0.086407684 | Credible Set | purple |
| chr14:120632852 | 14 | 120.6329 | 0.003910726 | Credible Set | purple |
| chr14:120632882 | 14 | 120.6329 | 0.003722401 | Credible Set | purple |
| chr14:120633549 | 14 | 120.6335 | 0.009935141 | Credible Set | purple |
| chr14:120641466 | 14 | 120.6415 | 0.011887200 | Credible Set | purple |
| chr14:120641472 | 14 | 120.6415 | 0.011887200 | Credible Set | purple |
| chr14:120641546 | 14 | 120.6415 | 0.009935141 | Credible Set | purple |
| chr14:120642644 | 14 | 120.6426 | 0.012387622 | Credible Set | purple |
| chr14:120642688 | 14 | 120.6427 | 0.010961154 | Credible Set | purple |
| chr14:120642702 | 14 | 120.6427 | 0.008469508 | Credible Set | purple |
| chr14:120642723 | 14 | 120.6427 | 0.008469508 | Credible Set | purple |

Fine−mapping Regions

| snp | chr | pos (Mb) | pp | group | color |
|---|---|---|---|---|---|
| chr14:120665611 | 14 | 120.6656 | 0.009935141 | Credible Set | purple |
| chr14:120670727 | 14 | 120.6707 | 0.007891510 | Credible Set | purple |
| chr14:120671083 | 14 | 120.6711 | 0.009935141 | Credible Set | purple |
| chr14:120671092 | 14 | 120.6711 | 0.009935141 | Credible Set | purple |
| chr14:120671147 | 14 | 120.6711 | 0.009935141 | Credible Set | purple |
| chr14:120671180 | 14 | 120.6712 | 0.011159893 | Credible Set | purple |
| chr14:120671181 | 14 | 120.6712 | 0.011187145 | Credible Set | purple |
| chr14:120686008 | 14 | 120.6860 | 0.006905295 | Credible Set | purple |
| chr14:120686112 | 14 | 120.6861 | 0.010960143 | Credible Set | purple |
| chr14:120686362 | 14 | 120.6864 | 0.010143590 | Credible Set | purple |
| chr14:120686413 | 14 | 120.6864 | 0.010105737 | Credible Set | purple |
| chr14:120688462 | 14 | 120.6885 | 0.003459534 | Credible Set | purple |
| chr14:120688492 | 14 | 120.6885 | 0.003712945 | Credible Set | purple |
| chr14:120699639 | 14 | 120.6996 | 0.006046421 | Credible Set | purple |
| chr14:120699675 | 14 | 120.6997 | 0.008623701 | Credible Set | purple |
| chr14:120702426 | 14 | 120.7024 | 0.007042586 | Credible Set | purple |
| chr14:120728333 | 14 | 120.7283 | 0.009800183 | Credible Set | purple |
| chr14:120728361 | 14 | 120.7284 | 0.008240755 | Credible Set | purple |
| chr14:120730072 | 14 | 120.7301 | 0.009935141 | Credible Set | purple |
| chr14:120731135 | 14 | 120.7311 | 0.029942186 | Credible Set | purple |
| chr14:120731136 | 14 | 120.7311 | 0.031995432 | Credible Set | purple |
| chr14:120767991 | 14 | 120.7680 | 0.009722721 | Credible Set | purple |
| chr14:120768037 | 14 | 120.7680 | 0.009353242 | Credible Set | purple |
| chr14:120768301 | 14 | 120.7683 | 0.023343015 | Credible Set | purple |
| chr14:120768336 | 14 | 120.7683 | 0.024874429 | Credible Set | purple |
| chr14:120768349 | 14 | 120.7683 | 0.019802031 | Credible Set | purple |
| chr14:120768705 | 14 | 120.7687 | 0.009242886 | Credible Set | purple |
| chr14:120768743 | 14 | 120.7687 | 0.007934912 | Credible Set | purple |
| chr14:120768819 | 14 | 120.7688 | 0.007832997 | Credible Set | purple |
| chr14:120774259 | 14 | 120.7743 | 0.007683939 | Credible Set | purple |
| chr14:120774290 | 14 | 120.7743 | 0.007892917 | Credible Set | purple |
| chr14:120774318 | 14 | 120.7743 | 0.007530706 | Credible Set | purple |
| chr14:120842399 | 14 | 120.8424 | 0.007567607 | Credible Set | purple |
| chr14:120868390 | 14 | 120.8684 | 0.006115525 | Credible Set | purple |
| chr14:120868527 | 14 | 120.8685 | 0.006141997 | Credible Set | purple |
| chr14:120868538 | 14 | 120.8685 | 0.006141997 | Credible Set | purple |

Fine−mapping Regions

| snp | chr | pos (Mb) | pp | group | color |
|---|---|---|---|---|---|
| chr14:120868670 | 14 | 120.8687 | 0.002806309 | Credible Set | purple |
| chr14:120868693 | 14 | 120.8687 | 0.002806309 | Credible Set | purple |
| chr14:120875959 | 14 | 120.8760 | 0.004840687 | Credible Set | purple |
| chr14:120958410 | 14 | 120.9584 | 0.006378942 | Credible Set | purple |
| chr14:120996497 | 14 | 120.9965 | 0.006353278 | Credible Set | purple |
| chr14:120996551 | 14 | 120.9966 | 0.006353278 | Credible Set | purple |
| chr14:120996555 | 14 | 120.9966 | 0.006353278 | Credible Set | purple |
| chr14:120996573 | 14 | 120.9966 | 0.006353278 | Credible Set | purple |
| chr14:121021982 | 14 | 121.0220 | 0.006353278 | Credible Set | purple |
| chr14:121022009 | 14 | 121.0220 | 0.006353278 | Credible Set | purple |
| chr14:121022048 | 14 | 121.0220 | 0.006353278 | Credible Set | purple |
| chr14:121022050 | 14 | 121.0220 | 0.006353278 | Credible Set | purple |
| chr14:121031640 | 14 | 121.0316 | 0.002329522 | Credible Set | purple |
| chr14:121031672 | 14 | 121.0317 | 0.002329522 | Credible Set | purple |
| chr14:121047713 | 14 | 121.0477 | 0.005711483 | Credible Set | purple |
| chr14:121054065 | 14 | 121.0541 | 0.005540939 | Credible Set | purple |
| chr14:121157624 | 14 | 121.1576 | 0.006353278 | Credible Set | purple |
| chr14:121166355 | 14 | 121.1664 | 0.012231142 | Credible Set | purple |

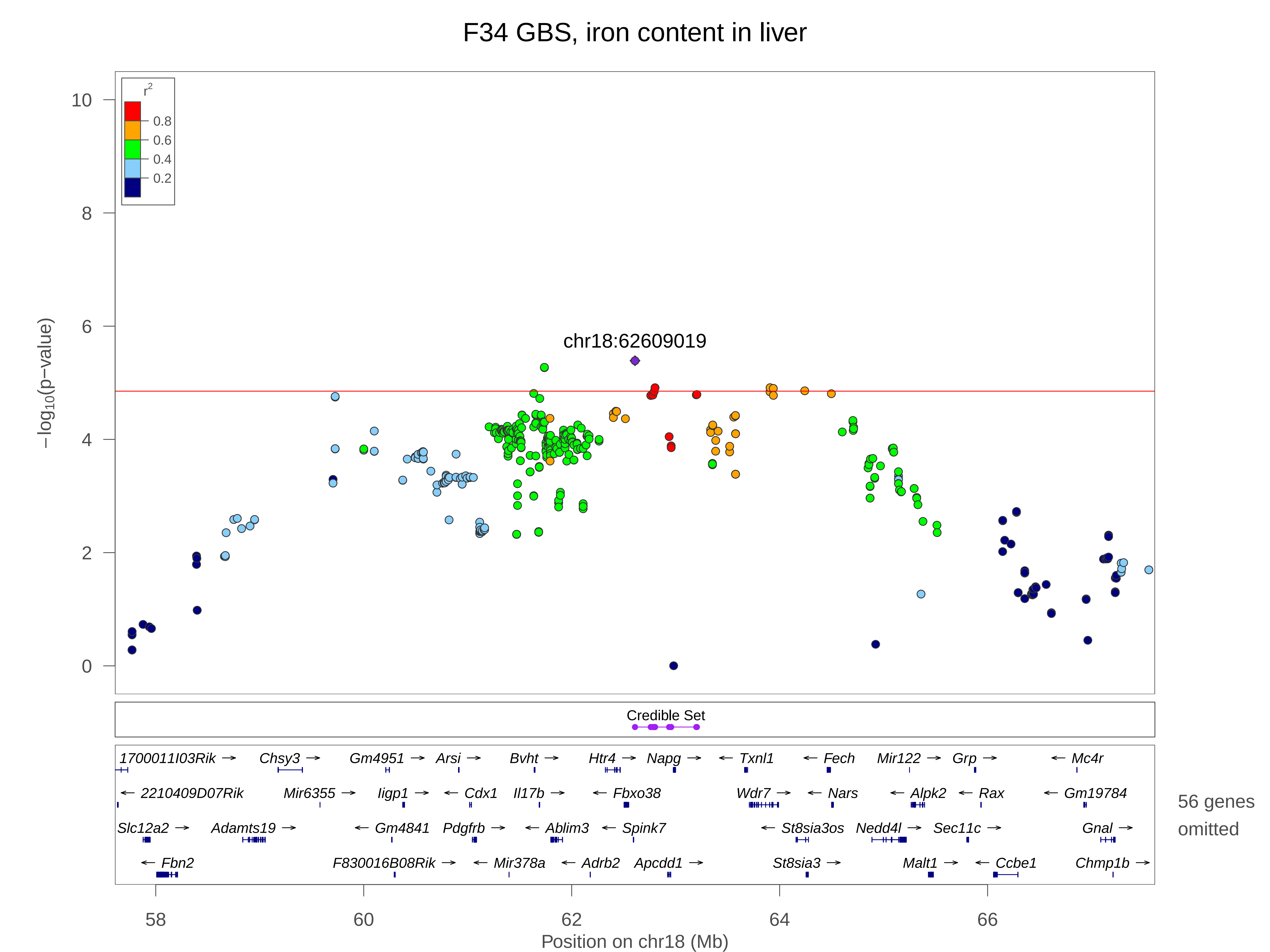
F34 GBS, iron content in liver
r2
0.8
0.6
0.4
0.2
10
8
6
4
2
0
−log10(p−value)
chr18:62609019
Credible Set
1700011I03Rik →
Chsy3 →
Gm4951 →
Arsi →
Bvht →
Htr4 →
Napg →
← Txnl1
← Fech
Mir122 →
Grp →
← Mc4r
← 2210409D07Rik
Mir6355 →
Iigp1 →
← Cdx1
Il17b →
← Fbxo38
Wdr7 →
← Nars
← Alpk2
← Rax
← Gm19784
Slc12a2 →
Adamts19 →
← Gm4841
Pdgfrb →
← Ablim3
← Spink7
← St8sia3os
Nedd4l →
Sec11c →
Gnal →
← Fbn2
F830016B08Rik →
← Mir378a
← Adrb2
Apcdd1 →
St8sia3 →
Malt1 →
← Ccbe1
Chmp1b →
56 genes
omitted
58
60
62
64
66
Position on chr18 (Mb)

| | |
|---|---|
| date: | Thu Oct 25 17:15:30 2018 |
| build: | F34GBS |
| display range: | chr18:57609019−67609019 [57609019−67609019] |
| hilite range: | 0 − 0 [ 0 − 0 ] |
| reference SNP: | chr18:62609019 |
| number of SNPs plotted: | 481 |
| min P−value: | 4.08E−6 [chr18:62609019] |
| max P−value: | 9.95E−1 [chr18:62980832] |
| omitted Genes: | Slc27a6, Isoc1, A730017C20Rik |
| omitted Genes: | Smim3, Dctn4, Rbm22 |
| omitted Genes: | Myoz3, Synpo, G630071F17Rik |
| omitted Genes: | Ndst1, Rps14, Cd74 |
| omitted Genes: | Mir5107, Tcof1, Camk2a |
| omitted Genes: | Mir7220, Mir6982, Slc6a7 |
| omitted Genes: | Csf1r, Mir6983, Hmgxb3 |
| omitted Genes: | Slc26a2, Pde6a, Ppargc1b |
| omitted Genes: | A530050N04Rik, Arhgef37, Csnk1a1 |
| omitted Genes: | Carmn, Mir145a, Mir143 |
| omitted Genes: | Pcyox1l, Grpel2, 1500015A07Rik |
| omitted Genes: | Afap1l1, Sh3tc2, Spink10 |
| omitted Genes: | Spink13, 2700046A07Rik, Piezo2 |
| omitted Genes: | Onecut2, Atp8b1, Zfp532 |
| omitted Genes: | Oacyl, Cplx4, Lman1 |
| omitted Genes: | Mir694, Pmaip1, Gm9926 |
| omitted Genes: | Mppe1, Impa2, B430212C06Rik |
| omitted Genes: | Cidea, Tubb6, Afg3l2 |
| omitted Genes: | Slmo1, Spire1 |

Make more plots at http://csg.sph.umich.edu/locuszoom/

Fine−mapping Regions

| snp | chr | pos (Mb) | pp | group | color |
|---|---|---|---|---|---|
| chr18:62609019 | 18 | 62.60902 | 0.31416088 | Credible Set | purple |
| chr18:62760743 | 18 | 62.76074 | 0.08133874 | Credible Set | purple |
| chr18:62780679 | 18 | 62.78068 | 0.08262774 | Credible Set | purple |
| chr18:62793979 | 18 | 62.79398 | 0.09517501 | Credible Set | purple |
| chr18:62801634 | 18 | 62.80163 | 0.10986009 | Credible Set | purple |
| chr18:62801661 | 18 | 62.80166 | 0.10986009 | Credible Set | purple |
| chr18:62937328 | 18 | 62.93733 | 0.01660355 | Credible Set | purple |
| chr18:62956832 | 18 | 62.95683 | 0.01167004 | Credible Set | purple |
| chr18:62956840 | 18 | 62.95684 | 0.01086881 | Credible Set | purple |
| chr18:63198665 | 18 | 63.19866 | 0.08343290 | Credible Set | purple |
| chr18:63204145 | 18 | 63.20414 | 0.08440215 | Credible Set | purple |

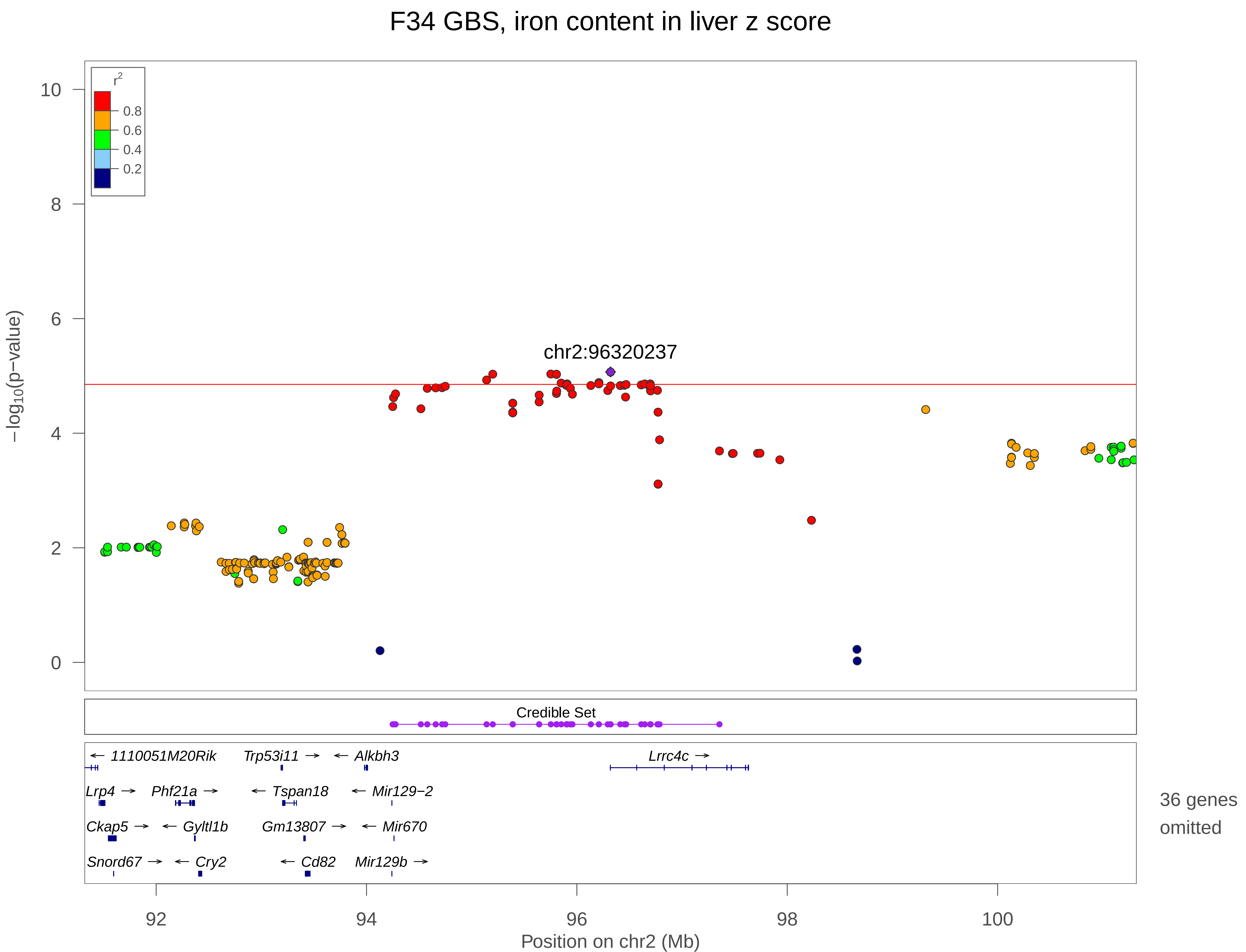
F34 GBS, iron content in liver z score
r2
0.8
0.6
0.4
0.2
10
8
6
4
2
0
−log10(p−value)
chr2:96320237
Credible Set
← 1110051M20Rik
Trp53i11 →
← Alkbh3
Lrrc4c →
Lrp4 →
Phf21a →
← Tspan18
← Mir129−2
Ckap5 →
← Gyltl1b
Gm13807 →
← Mir670
Snord67 →
← Cry2
← Cd82
Mir129b →
36 genes omitted
92
94
96
98
100
Position on chr2 (Mb)

| | |
|---|---|
| date: | Thu Oct 25 17:16:28 2018 |
| build: | F34GBS |
| display range: | chr2:91320237−101320237 [91320237−101320237] |
| hilite range: | 0 − 0 [ 0 − 0 ] |
| reference SNP: | chr2:96320237 |
| number of SNPs plotted: | 236 |
| min P−value: | 8.52E−6 [chr2:96320237] |
| max P−value: | 9.43E−1 [chr2:98665093] |
| omitted Genes: | F2, Zfp408, Arhgap1 |
| omitted Genes: | Atg13, Harbi1, Ambra1 |
| omitted Genes: | Chrm4, Mdk, Dgkz |
| omitted Genes: | Mir6999, Creb3l1, Mir1955 |
| omitted Genes: | Pex16, 1700029I15Rik, Mapk8ip1 |
| omitted Genes: | Mir7000, D930015M05Rik, Slc35c1 |
| omitted Genes: | Mir7221, Mir7222, Chst1 |
| omitted Genes: | Syt13, Prdm11, 4631405J19Rik |
| omitted Genes: | Mir7001, Gm10804, Alx4 |
| omitted Genes: | Ext2, Accs, Accsl |
| omitted Genes: | Gm13889, Alkbh3os1, Hsd17b12 |
| omitted Genes: | Ttc17, 2810002D19Rik, Api5 |

Make more plots at http://csg.sph.umich.edu/locuszoom/

Fine−mapping Regions

| snp | chr | pos (Mb) | pp | group | color |
|---|---|---|---|---|---|
| chr2:94249122 | 2 | 94.24912 | 0.009345889 | Credible Set | purple |
| chr2:94257379 | 2 | 94.25738 | 0.013149298 | Credible Set | purple |
| chr2:94276010 | 2 | 94.27601 | 0.015190090 | Credible Set | purple |
| chr2:94516977 | 2 | 94.51698 | 0.008596523 | Credible Set | purple |
| chr2:94577500 | 2 | 94.57750 | 0.018768120 | Credible Set | purple |
| chr2:94657941 | 2 | 94.65794 | 0.019106526 | Credible Set | purple |
| chr2:94718556 | 2 | 94.71856 | 0.019232070 | Credible Set | purple |
| chr2:94748712 | 2 | 94.74871 | 0.020318937 | Credible Set | purple |
| chr2:95141904 | 2 | 95.14190 | 0.025871490 | Credible Set | purple |
| chr2:95200398 | 2 | 95.20040 | 0.032322327 | Credible Set | purple |
| chr2:95390243 | 2 | 95.39024 | 0.007488085 | Credible Set | purple |
| chr2:95390247 | 2 | 95.39025 | 0.007341922 | Credible Set | purple |
| chr2:95390277 | 2 | 95.39028 | 0.007556063 | Credible Set | purple |
| chr2:95390284 | 2 | 95.39028 | 0.007394122 | Credible Set | purple |
| chr2:95390546 | 2 | 95.39055 | 0.010663314 | Credible Set | purple |
| chr2:95390592 | 2 | 95.39059 | 0.010574691 | Credible Set | purple |
| chr2:95641124 | 2 | 95.64112 | 0.011159002 | Credible Set | purple |
| chr2:95641477 | 2 | 95.64148 | 0.014491530 | Credible Set | purple |
| chr2:95752277 | 2 | 95.75228 | 0.032479006 | Credible Set | purple |
| chr2:95805867 | 2 | 95.80587 | 0.015405704 | Credible Set | purple |
| chr2:95806028 | 2 | 95.80603 | 0.031965628 | Credible Set | purple |
| chr2:95806038 | 2 | 95.80604 | 0.032299024 | Credible Set | purple |
| chr2:95808603 | 2 | 95.80860 | 0.016903695 | Credible Set | purple |
| chr2:95850674 | 2 | 95.85067 | 0.022992151 | Credible Set | purple |
| chr2:95900229 | 2 | 95.90023 | 0.021113571 | Credible Set | purple |
| chr2:95906738 | 2 | 95.90674 | 0.022263063 | Credible Set | purple |
| chr2:95906783 | 2 | 95.90678 | 0.021372052 | Credible Set | purple |
| chr2:95938782 | 2 | 95.93878 | 0.018709679 | Credible Set | purple |
| chr2:95957448 | 2 | 95.95745 | 0.014994804 | Credible Set | purple |
| chr2:96132587 | 2 | 96.13259 | 0.020911104 | Credible Set | purple |
| chr2:96209134 | 2 | 96.20913 | 0.023419246 | Credible Set | purple |
| chr2:96209157 | 2 | 96.20916 | 0.022268999 | Credible Set | purple |
| chr2:96293855 | 2 | 96.29385 | 0.017241174 | Credible Set | purple |
| chr2:96320222 | 2 | 96.32022 | 0.034860342 | Credible Set | purple |
| chr2:96320237 | 2 | 96.32024 | 0.035235075 | Credible Set | purple |
| chr2:96320641 | 2 | 96.32064 | 0.020592921 | Credible Set | purple |

Fine−mapping Regions

| snp | chr | pos (Mb) | pp | group | color |
|---|---|---|---|---|---|
| chr2:96412425 | 2 | 96.41242 | 0.020875290 | Credible Set | purple |
| chr2:96451438 | 2 | 96.45144 | 0.021113571 | Credible Set | purple |
| chr2:96462914 | 2 | 96.46291 | 0.013447390 | Credible Set | purple |
| chr2:96468064 | 2 | 96.46806 | 0.021712016 | Credible Set | purple |
| chr2:96611762 | 2 | 96.61176 | 0.021390367 | Credible Set | purple |
| chr2:96645727 | 2 | 96.64573 | 0.022222538 | Credible Set | purple |
| chr2:96697256 | 2 | 96.69726 | 0.022262601 | Credible Set | purple |
| chr2:96697871 | 2 | 96.69787 | 0.018980753 | Credible Set | purple |
| chr2:96697872 | 2 | 96.69787 | 0.019258598 | Credible Set | purple |
| chr2:96701045 | 2 | 96.70104 | 0.017070663 | Credible Set | purple |
| chr2:96701070 | 2 | 96.70107 | 0.017070663 | Credible Set | purple |
| chr2:96701077 | 2 | 96.70108 | 0.017070663 | Credible Set | purple |
| chr2:96701079 | 2 | 96.70108 | 0.017070663 | Credible Set | purple |
| chr2:96701166 | 2 | 96.70117 | 0.020818440 | Credible Set | purple |
| chr2:96766199 | 2 | 96.76620 | 0.017362363 | Credible Set | purple |
| chr2:96771761 | 2 | 96.77176 | 0.007564679 | Credible Set | purple |
| chr2:96771791 | 2 | 96.77179 | 0.007543133 | Credible Set | purple |
| chr2:96786046 | 2 | 96.78605 | 0.002639576 | Credible Set | purple |
| chr2:97355566 | 2 | 97.35557 | 0.001733072 | Credible Set | purple |

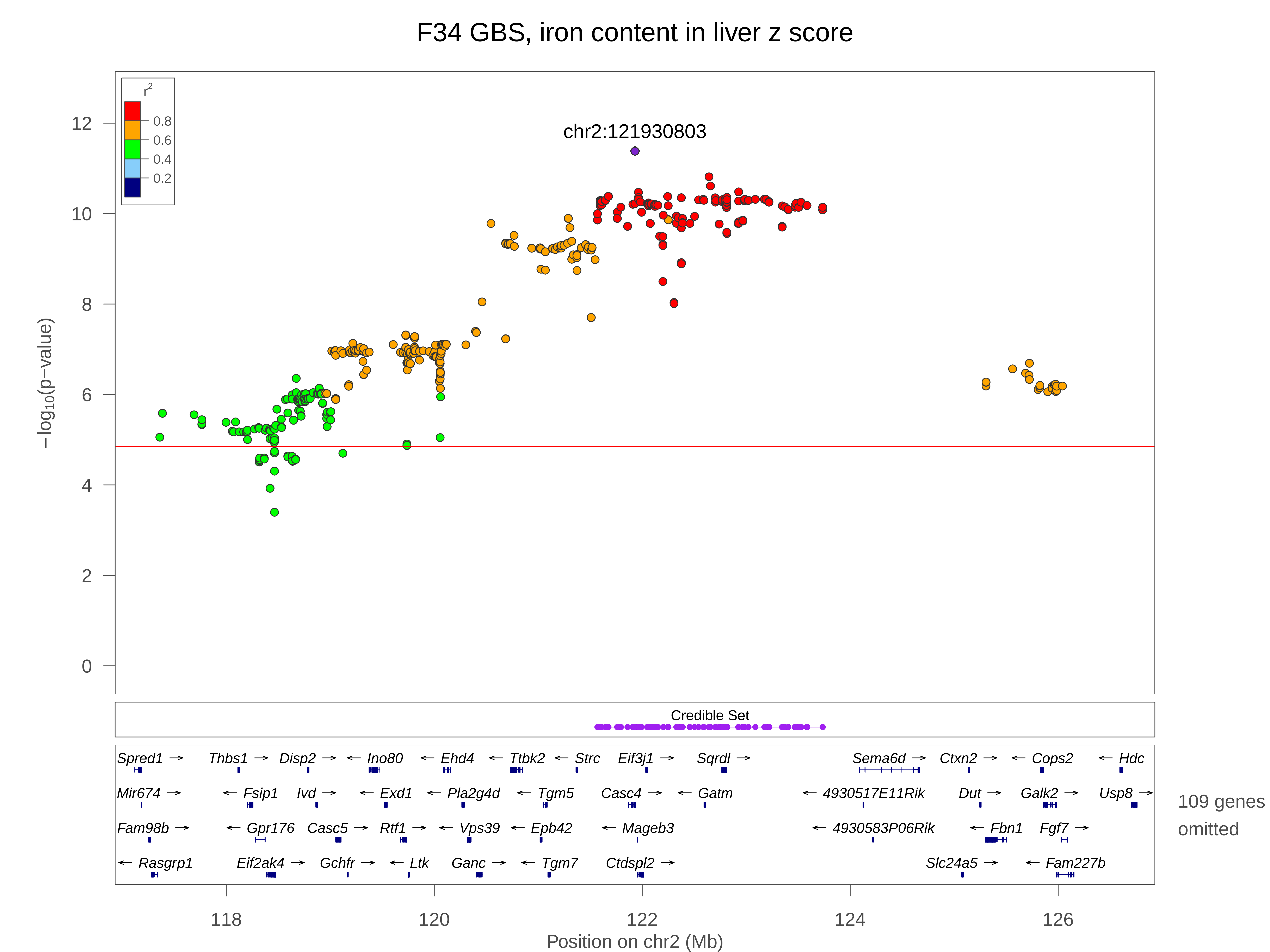
F34 GBS, iron content in liver z score
chr2:121930803
r2
0.8
0.6
0.4
0.2
−log10(p−value)
0
2
4
6
8
10
12
Credible Set
Spred1 →
Thbs1 →
Disp2 →
← Ino80
← Ehd4
← Ttbk2
← Strc
Eif3j1 →
Sqrdl →
Sema6d →
Ctxn2 →
← Cops2
← Hdc
Mir674 →
← Fsip1
Ivd →
← Exd1
← Pla2g4d
← Tgm5
Casc4 →
← Gatm
← 4930517E11Rik
Dut →
Galk2 →
Usp8 →
Fam98b →
← Gpr176
Casc5 →
Rtf1 →
← Vps39
← Epb42
← Mageb3
← 4930583P06Rik
← Fbn1
Fgf7 →
← Rasgrp1
Eif2ak4 →
Gchfr →
← Ltk
Ganc →
← Tgm7
Ctdspl2 →
Slc24a5 →
← Fam227b
109 genes omitted
118
120
122
124
126
Position on chr2 (Mb)

date: Thu Oct 25 17:16:19 2018
build: F34GBS
display range: chr2:116930803−126930803 [116930803−126930803]
hilite range: 0 − 0 [ 0 − 0 ]
reference SNP: chr2:121930803
number of SNPs plotted: 530
min P−value: 4.17E−12 [chr2:121930803]
max P−value: 4.02E−4 [chr2:118463923]
omitted Genes: Gm13985, 4930412B13Rik, Gm13986
omitted Genes: 1700054M17Rik, Srp14, Bmf
omitted Genes: Bub1b, Pak6, Ankrd63
omitted Genes: Plcb2, Inafm2, A430105I19Rik
omitted Genes: Phgr1, Knstrn, Bahd1
omitted Genes: Chst14, Ccdc32, Rpusd2
omitted Genes: Rad51, Rmdn3, Dnajc17
omitted Genes: Gm14137, Zfyve19, Ppp1r14d
omitted Genes: Spint1, Rhov, Vps18
omitted Genes: Gm14207, Dll4, Chac1
omitted Genes: Chp1, 1700020I14Rik, Oip5
omitted Genes: Nusap1, Ndufaf1, Itpka
omitted Genes: Rpap1, Tyro3, Mga
omitted Genes: Mapkbp1, Mir7665, Jmjd7
omitted Genes: Gm28042, Pla2g4b, Pla2g4e
omitted Genes: Pla2g4f, Tmem87a, 4931402G19Rik
omitted Genes: Capn3, Zfp106, Snap23
omitted Genes: Lrrc57, Haus2, Cdan1
omitted Genes: AV039307, Ubr1, Tmem62
omitted Genes: Ccndbp1, Lcmt2, Adal
omitted Genes: Zscan29, Tubgcp4, Trp53bp1
omitted Genes: Map1a, Ppip5k1, Ckmt1
omitted Genes: Catsper2, Pdia3, Ell3
omitted Genes: Serf2, Serinc4, Hypk
omitted Genes: Mfap1b, Mfap1a, Wdr76
omitted Genes: Frmd5, Spg11, Patl2
omitted Genes: B2m, Trim69, Terb2
omitted Genes: Sord, Duox2, Duoxa2
omitted Genes: Duoxa1, Duox1, Shf
omitted Genes: Slc28a2, Bambi−ps1, Gm14085
omitted Genes: Spata5l1, AA467197, Mir147
Make more plots at http://csg.sph.umich.edu/locuszoom/
omitted Genes: Slc30a4, Bloc1s6, Myef2

Fine−mapping Regions

| snp | chr | pos (Mb) | pp | group | color |
|---|---|---|---|---|---|
| chr2:121569629 | 2 | 121.5696 | 0.002857489 | Credible Set | purple |
| chr2:121569951 | 2 | 121.5700 | 0.002888488 | Credible Set | purple |
| chr2:121570025 | 2 | 121.5700 | 0.003934945 | Credible Set | purple |
| chr2:121595218 | 2 | 121.5952 | 0.007532776 | Credible Set | purple |
| chr2:121595356 | 2 | 121.5954 | 0.007532776 | Credible Set | purple |
| chr2:121595424 | 2 | 121.5954 | 0.005848599 | Credible Set | purple |
| chr2:121595633 | 2 | 121.5956 | 0.006371258 | Credible Set | purple |
| chr2:121605090 | 2 | 121.6051 | 0.007532776 | Credible Set | purple |
| chr2:121605163 | 2 | 121.6052 | 0.007356642 | Credible Set | purple |
| chr2:121609496 | 2 | 121.6095 | 0.007385239 | Credible Set | purple |
| chr2:121609992 | 2 | 121.6100 | 0.006101736 | Credible Set | purple |
| chr2:121610054 | 2 | 121.6101 | 0.007184117 | Credible Set | purple |
| chr2:121644474 | 2 | 121.6445 | 0.007532776 | Credible Set | purple |
| chr2:121644502 | 2 | 121.6445 | 0.007532776 | Credible Set | purple |
| chr2:121647254 | 2 | 121.6473 | 0.007687139 | Credible Set | purple |
| chr2:121675210 | 2 | 121.6752 | 0.009304945 | Credible Set | purple |
| chr2:121675212 | 2 | 121.6752 | 0.009304945 | Credible Set | purple |
| chr2:121675249 | 2 | 121.6752 | 0.009304945 | Credible Set | purple |
| chr2:121675251 | 2 | 121.6753 | 0.009304945 | Credible Set | purple |
| chr2:121760589 | 2 | 121.7606 | 0.004252916 | Credible Set | purple |
| chr2:121760592 | 2 | 121.7606 | 0.004252916 | Credible Set | purple |
| chr2:121760608 | 2 | 121.7606 | 0.004252916 | Credible Set | purple |
| chr2:121760655 | 2 | 121.7607 | 0.003110863 | Credible Set | purple |
| chr2:121760675 | 2 | 121.7607 | 0.003110863 | Credible Set | purple |
| chr2:121795014 | 2 | 121.7950 | 0.005449237 | Credible Set | purple |
| chr2:121859536 | 2 | 121.8595 | 0.002099820 | Credible Set | purple |
| chr2:121859542 | 2 | 121.8595 | 0.002090902 | Credible Set | purple |
| chr2:121910457 | 2 | 121.9105 | 0.006283734 | Credible Set | purple |
| chr2:121930722 | 2 | 121.9307 | 0.006480551 | Credible Set | purple |
| chr2:121930803 | 2 | 121.9308 | 0.088633203 | Credible Set | purple |
| chr2:121930806 | 2 | 121.9308 | 0.088633203 | Credible Set | purple |
| chr2:121963958 | 2 | 121.9640 | 0.011392752 | Credible Set | purple |
| chr2:121964022 | 2 | 121.9640 | 0.008828939 | Credible Set | purple |
| chr2:121964577 | 2 | 121.9646 | 0.008056022 | Credible Set | purple |
| chr2:121982399 | 2 | 121.9824 | 0.007056880 | Credible Set | purple |
| chr2:121994315 | 2 | 121.9943 | 0.004227053 | Credible Set | purple |

Fine−mapping Regions

| snp | chr | pos (Mb) | pp | group | color |
|---|---|---|---|---|---|
| chr2:121994324 | 2 | 121.9943 | 0.004227053 | Credible Set | purple |
| chr2:122047663 | 2 | 122.0477 | 0.006414192 | Credible Set | purple |
| chr2:122060688 | 2 | 122.0607 | 0.006732340 | Credible Set | purple |
| chr2:122061099 | 2 | 122.0611 | 0.005826115 | Credible Set | purple |
| chr2:122065787 | 2 | 122.0658 | 0.006442917 | Credible Set | purple |
| chr2:122065870 | 2 | 122.0659 | 0.006208577 | Credible Set | purple |
| chr2:122077766 | 2 | 122.0778 | 0.002419555 | Credible Set | purple |
| chr2:122077798 | 2 | 122.0778 | 0.002419555 | Credible Set | purple |
| chr2:122083136 | 2 | 122.0831 | 0.006330985 | Credible Set | purple |
| chr2:122090229 | 2 | 122.0902 | 0.006403112 | Credible Set | purple |
| chr2:122115252 | 2 | 122.1153 | 0.005833033 | Credible Set | purple |
| chr2:122122941 | 2 | 122.1229 | 0.005667775 | Credible Set | purple |
| chr2:122123313 | 2 | 122.1233 | 0.006215054 | Credible Set | purple |
| chr2:122132763 | 2 | 122.1328 | 0.006096369 | Credible Set | purple |
| chr2:122151119 | 2 | 122.1511 | 0.006015879 | Credible Set | purple |
| chr2:122202473 | 2 | 122.2025 | 0.003651499 | Credible Set | purple |
| chr2:122244672 | 2 | 122.2447 | 0.009251758 | Credible Set | purple |
| chr2:122250299 | 2 | 122.2503 | 0.005837905 | Credible Set | purple |
| chr2:122328643 | 2 | 122.3286 | 0.002437696 | Credible Set | purple |
| chr2:122328650 | 2 | 122.3286 | 0.003494080 | Credible Set | purple |
| chr2:122346906 | 2 | 122.3469 | 0.003192585 | Credible Set | purple |
| chr2:122376352 | 2 | 122.3764 | 0.001934156 | Credible Set | purple |
| chr2:122376463 | 2 | 122.3765 | 0.008709426 | Credible Set | purple |
| chr2:122387979 | 2 | 122.3880 | 0.003101788 | Credible Set | purple |
| chr2:122388097 | 2 | 122.3881 | 0.002523569 | Credible Set | purple |
| chr2:122388208 | 2 | 122.3882 | 0.002469119 | Credible Set | purple |
| chr2:122458764 | 2 | 122.4588 | 0.002420198 | Credible Set | purple |
| chr2:122504055 | 2 | 122.5041 | 0.003442057 | Credible Set | purple |
| chr2:122542898 | 2 | 122.5429 | 0.007834508 | Credible Set | purple |
| chr2:122587284 | 2 | 122.5873 | 0.008038345 | Credible Set | purple |
| chr2:122587743 | 2 | 122.5877 | 0.008006825 | Credible Set | purple |
| chr2:122593432 | 2 | 122.5934 | 0.007658566 | Credible Set | purple |
| chr2:122642503 | 2 | 122.6425 | 0.024688407 | Credible Set | purple |
| chr2:122656435 | 2 | 122.6564 | 0.015653215 | Credible Set | purple |
| chr2:122656458 | 2 | 122.6565 | 0.015653215 | Credible Set | purple |
| chr2:122702118 | 2 | 122.7021 | 0.008676101 | Credible Set | purple |

Fine−mapping Regions

| snp | chr | pos (Mb) | pp | group | color |
|---|---|---|---|---|---|
| chr2:122704152 | 2 | 122.7042 | 0.006983921 | Credible Set | purple |
| chr2:122708239 | 2 | 122.7082 | 0.007532776 | Credible Set | purple |
| chr2:122739988 | 2 | 122.7400 | 0.002346524 | Credible Set | purple |
| chr2:122740035 | 2 | 122.7400 | 0.002349438 | Credible Set | purple |
| chr2:122771056 | 2 | 122.7711 | 0.007171837 | Credible Set | purple |
| chr2:122771465 | 2 | 122.7715 | 0.007902549 | Credible Set | purple |
| chr2:122797192 | 2 | 122.7972 | 0.006787411 | Credible Set | purple |
| chr2:122797216 | 2 | 122.7972 | 0.006731594 | Credible Set | purple |
| chr2:122797265 | 2 | 122.7973 | 0.007873788 | Credible Set | purple |
| chr2:122797266 | 2 | 122.7973 | 0.007886566 | Credible Set | purple |
| chr2:122804433 | 2 | 122.8044 | 0.005962276 | Credible Set | purple |
| chr2:122811212 | 2 | 122.8112 | 0.005339439 | Credible Set | purple |
| chr2:122811304 | 2 | 122.8113 | 0.005928817 | Credible Set | purple |
| chr2:122814314 | 2 | 122.8143 | 0.008469307 | Credible Set | purple |
| chr2:122814326 | 2 | 122.8143 | 0.006895917 | Credible Set | purple |
| chr2:122814381 | 2 | 122.8144 | 0.007027315 | Credible Set | purple |
| chr2:122814728 | 2 | 122.8147 | 0.008880280 | Credible Set | purple |
| chr2:122814902 | 2 | 122.8149 | 0.007838651 | Credible Set | purple |
| chr2:122814909 | 2 | 122.8149 | 0.007851767 | Credible Set | purple |
| chr2:122924669 | 2 | 122.9247 | 0.002412749 | Credible Set | purple |
| chr2:122924685 | 2 | 122.9247 | 0.002421077 | Credible Set | purple |
| chr2:122924695 | 2 | 122.9247 | 0.002412749 | Credible Set | purple |
| chr2:122924696 | 2 | 122.9247 | 0.002412749 | Credible Set | purple |
| chr2:122927536 | 2 | 122.9275 | 0.007386738 | Credible Set | purple |
| chr2:122927903 | 2 | 122.9279 | 0.011619776 | Credible Set | purple |
| chr2:122927980 | 2 | 122.9280 | 0.011677971 | Credible Set | purple |
| chr2:122927997 | 2 | 122.9280 | 0.002624947 | Credible Set | purple |
| chr2:122928025 | 2 | 122.9280 | 0.002461347 | Credible Set | purple |
| chr2:122968234 | 2 | 122.9682 | 0.002820595 | Credible Set | purple |
| chr2:122968260 | 2 | 122.9683 | 0.002855143 | Credible Set | purple |
| chr2:122968275 | 2 | 122.9683 | 0.002718756 | Credible Set | purple |
| chr2:122968296 | 2 | 122.9683 | 0.002763713 | Credible Set | purple |
| chr2:122983867 | 2 | 122.9839 | 0.007483440 | Credible Set | purple |
| chr2:122986477 | 2 | 122.9865 | 0.007648956 | Credible Set | purple |
| chr2:122986971 | 2 | 122.9870 | 0.008076340 | Credible Set | purple |
| chr2:123021071 | 2 | 123.0211 | 0.007626697 | Credible Set | purple |

Fine−mapping Regions

| snp | chr | pos (Mb) | pp | group | color |
|---|---|---|---|---|---|
| chr2:123021090 | 2 | 123.0211 | 0.007647626 | Credible Set | purple |
| chr2:123021532 | 2 | 123.0215 | 0.007610671 | Credible Set | purple |
| chr2:123088225 | 2 | 123.0882 | 0.008023564 | Credible Set | purple |
| chr2:123175096 | 2 | 123.1751 | 0.008038758 | Credible Set | purple |
| chr2:123190496 | 2 | 123.1905 | 0.008073418 | Credible Set | purple |
| chr2:123219436 | 2 | 123.2194 | 0.007202137 | Credible Set | purple |
| chr2:123219468 | 2 | 123.2195 | 0.006968850 | Credible Set | purple |
| chr2:123346239 | 2 | 123.3462 | 0.002094342 | Credible Set | purple |
| chr2:123346253 | 2 | 123.3463 | 0.002015733 | Credible Set | purple |
| chr2:123346704 | 2 | 123.3467 | 0.005800515 | Credible Set | purple |
| chr2:123371857 | 2 | 123.3719 | 0.005505525 | Credible Set | purple |
| chr2:123371991 | 2 | 123.3720 | 0.005505525 | Credible Set | purple |
| chr2:123404160 | 2 | 123.4042 | 0.004761916 | Credible Set | purple |
| chr2:123404632 | 2 | 123.4046 | 0.004880475 | Credible Set | purple |
| chr2:123470044 | 2 | 123.4700 | 0.005432790 | Credible Set | purple |
| chr2:123470368 | 2 | 123.4704 | 0.005852956 | Credible Set | purple |
| chr2:123470382 | 2 | 123.4704 | 0.005867645 | Credible Set | purple |
| chr2:123478720 | 2 | 123.4787 | 0.006578271 | Credible Set | purple |
| chr2:123505482 | 2 | 123.5055 | 0.005411790 | Credible Set | purple |
| chr2:123505557 | 2 | 123.5056 | 0.005391510 | Credible Set | purple |
| chr2:123527223 | 2 | 123.5272 | 0.006990985 | Credible Set | purple |
| chr2:123584403 | 2 | 123.5844 | 0.005907269 | Credible Set | purple |
| chr2:123585516 | 2 | 123.5855 | 0.005911792 | Credible Set | purple |
| chr2:123735848 | 2 | 123.7358 | 0.004751842 | Credible Set | purple |
| chr2:123735871 | 2 | 123.7359 | 0.004766643 | Credible Set | purple |
| chr2:123735958 | 2 | 123.7360 | 0.005416055 | Credible Set | purple |

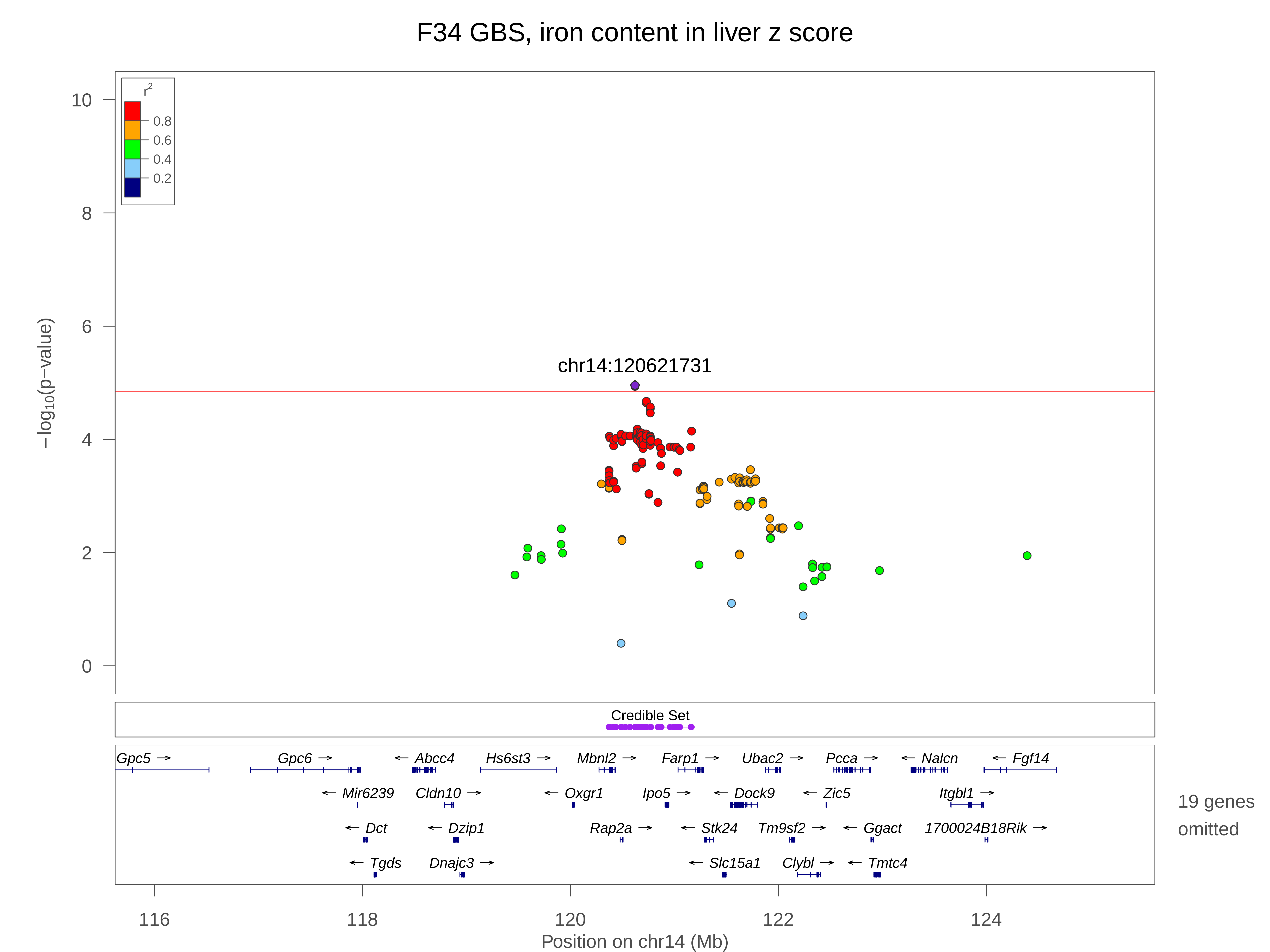
F34 GBS, iron content in liver z score
r2
0.8
0.6
0.4
0.2
chr14:120621731
−log10(p−value)
Credible Set
Gpc5
Gpc6
Abcc4
Hs6st3
Mbnl2
Farp1
Ubac2
Pcca
Nalcn
Fgf14
Mir6239
Cldn10
Oxgr1
Ipo5
Dock9
Zic5
Itgbl1
Dct
Dzip1
Rap2a
Stk24
Tm9sf2
Ggact
1700024B18Rik
Tgds
Dnajc3
Slc15a1
Clybl
Tmtc4
19 genes omitted
Position on chr14 (Mb)

| | |
|---|---|
| date: | Thu Oct 25 17:15:59 2018 |
| build: | F34GBS |
| display range: | chr14:115621731−125621731 [115621731−125621731] |
| hilite range: | 0 − 0 [ 0 − 0 ] |
| reference SNP: | chr14:120621731 |
| number of SNPs plotted: | 206 |
| min P−value: | 1.11E−5 [chr14:120621731] |
| max P−value: | 3.98E−1 [chr14:120487423] |
| omitted Genes: | Gpr180, Sox21, LOC105245869 |
| omitted Genes: | Mir6241, Gm9376, 1700044C05Rik |
| omitted Genes: | Mir6391, Uggt2, 1700006F04Rik |
| omitted Genes: | 1810041H14Rik, Gpr18, Gpr183 |
| omitted Genes: | Timm8a2, A330035P11Rik, 1700108J01Rik |
| omitted Genes: | Gm5089, 2610035F20Rik, Zic2 |
| omitted Genes: | AA536875 |

Fine−mapping Regions

| snp | chr | pos (Mb) | pp | group | color |
|---|---|---|---|---|---|
| chr14:120371692 | 14 | 120.3717 | 0.002717539 | Credible Set | purple |
| chr14:120371706 | 14 | 120.3717 | 0.002619874 | Credible Set | purple |
| chr14:120371729 | 14 | 120.3717 | 0.002193366 | Credible Set | purple |
| chr14:120373900 | 14 | 120.3739 | 0.009981640 | Credible Set | purple |
| chr14:120374171 | 14 | 120.3742 | 0.001788578 | Credible Set | purple |
| chr14:120374181 | 14 | 120.3742 | 0.001833904 | Credible Set | purple |
| chr14:120379882 | 14 | 120.3799 | 0.001675364 | Credible Set | purple |
| chr14:120382338 | 14 | 120.3823 | 0.009276992 | Credible Set | purple |
| chr14:120416102 | 14 | 120.4161 | 0.001784302 | Credible Set | purple |
| chr14:120416136 | 14 | 120.4161 | 0.001732416 | Credible Set | purple |
| chr14:120416333 | 14 | 120.4163 | 0.006921214 | Credible Set | purple |
| chr14:120416347 | 14 | 120.4163 | 0.008575546 | Credible Set | purple |
| chr14:120439342 | 14 | 120.4393 | 0.009203875 | Credible Set | purple |
| chr14:120487392 | 14 | 120.4874 | 0.010131800 | Credible Set | purple |
| chr14:120487416 | 14 | 120.4874 | 0.010123622 | Credible Set | purple |
| chr14:120487566 | 14 | 120.4876 | 0.010123622 | Credible Set | purple |
| chr14:120487639 | 14 | 120.4876 | 0.010750502 | Credible Set | purple |
| chr14:120495956 | 14 | 120.4960 | 0.008102774 | Credible Set | purple |
| chr14:120495976 | 14 | 120.4960 | 0.008247406 | Credible Set | purple |
| chr14:120532718 | 14 | 120.5327 | 0.010089466 | Credible Set | purple |
| chr14:120532740 | 14 | 120.5327 | 0.010089466 | Credible Set | purple |
| chr14:120573821 | 14 | 120.5738 | 0.010060872 | Credible Set | purple |
| chr14:120573951 | 14 | 120.5740 | 0.010089466 | Credible Set | purple |
| chr14:120621718 | 14 | 120.6217 | 0.068360725 | Credible Set | purple |
| chr14:120621731 | 14 | 120.6217 | 0.071436539 | Credible Set | purple |
| chr14:120621749 | 14 | 120.6217 | 0.070173723 | Credible Set | purple |
| chr14:120632852 | 14 | 120.6329 | 0.003183660 | Credible Set | purple |
| chr14:120632882 | 14 | 120.6329 | 0.002939877 | Credible Set | purple |
| chr14:120633549 | 14 | 120.6335 | 0.010089466 | Credible Set | purple |
| chr14:120641466 | 14 | 120.6415 | 0.012419721 | Credible Set | purple |
| chr14:120641472 | 14 | 120.6415 | 0.012419721 | Credible Set | purple |
| chr14:120641546 | 14 | 120.6415 | 0.010089466 | Credible Set | purple |
| chr14:120642644 | 14 | 120.6426 | 0.013126447 | Credible Set | purple |
| chr14:120642688 | 14 | 120.6427 | 0.011413870 | Credible Set | purple |
| chr14:120642702 | 14 | 120.6427 | 0.008687413 | Credible Set | purple |
| chr14:120642723 | 14 | 120.6427 | 0.008687413 | Credible Set | purple |

Fine−mapping Regions

| snp | chr | pos (Mb) | pp | group | color |
|---|---|---|---|---|---|
| chr14:120665611 | 14 | 120.6656 | 0.010089466 | Credible Set | purple |
| chr14:120670727 | 14 | 120.6707 | 0.007788504 | Credible Set | purple |
| chr14:120671083 | 14 | 120.6711 | 0.010089466 | Credible Set | purple |
| chr14:120671092 | 14 | 120.6711 | 0.010089466 | Credible Set | purple |
| chr14:120671147 | 14 | 120.6711 | 0.010089466 | Credible Set | purple |
| chr14:120671180 | 14 | 120.6712 | 0.011455704 | Credible Set | purple |
| chr14:120671181 | 14 | 120.6712 | 0.011486631 | Credible Set | purple |
| chr14:120686008 | 14 | 120.6860 | 0.007088228 | Credible Set | purple |
| chr14:120686112 | 14 | 120.6861 | 0.011316934 | Credible Set | purple |
| chr14:120686362 | 14 | 120.6864 | 0.010234701 | Credible Set | purple |
| chr14:120686413 | 14 | 120.6864 | 0.010201053 | Credible Set | purple |
| chr14:120688462 | 14 | 120.6885 | 0.003473863 | Credible Set | purple |
| chr14:120688492 | 14 | 120.6885 | 0.003703304 | Credible Set | purple |
| chr14:120699639 | 14 | 120.6996 | 0.006226111 | Credible Set | purple |
| chr14:120699675 | 14 | 120.6997 | 0.008755304 | Credible Set | purple |
| chr14:120702426 | 14 | 120.7024 | 0.007134272 | Credible Set | purple |
| chr14:120728333 | 14 | 120.7283 | 0.010881628 | Credible Set | purple |
| chr14:120728361 | 14 | 120.7284 | 0.009139025 | Credible Set | purple |
| chr14:120730072 | 14 | 120.7301 | 0.010089466 | Credible Set | purple |
| chr14:120731135 | 14 | 120.7311 | 0.035957368 | Credible Set | purple |
| chr14:120731136 | 14 | 120.7311 | 0.038301247 | Credible Set | purple |
| chr14:120767991 | 14 | 120.7680 | 0.010028215 | Credible Set | purple |
| chr14:120768037 | 14 | 120.7680 | 0.007032670 | Credible Set | purple |
| chr14:120768301 | 14 | 120.7683 | 0.028774894 | Credible Set | purple |
| chr14:120768336 | 14 | 120.7683 | 0.030890411 | Credible Set | purple |
| chr14:120768349 | 14 | 120.7683 | 0.024382310 | Credible Set | purple |
| chr14:120768705 | 14 | 120.7687 | 0.009643288 | Credible Set | purple |
| chr14:120768743 | 14 | 120.7687 | 0.008240744 | Credible Set | purple |
| chr14:120768819 | 14 | 120.7688 | 0.008134422 | Credible Set | purple |
| chr14:120774259 | 14 | 120.7743 | 0.008671610 | Credible Set | purple |
| chr14:120774290 | 14 | 120.7743 | 0.008888689 | Credible Set | purple |
| chr14:120774318 | 14 | 120.7743 | 0.008465055 | Credible Set | purple |
| chr14:120842399 | 14 | 120.8424 | 0.007803704 | Credible Set | purple |
| chr14:120868390 | 14 | 120.8684 | 0.006284490 | Credible Set | purple |
| chr14:120868527 | 14 | 120.8685 | 0.006321294 | Credible Set | purple |
| chr14:120868538 | 14 | 120.8685 | 0.006321294 | Credible Set | purple |

Fine−mapping Regions

| snp | chr | pos (Mb) | pp | group | color |
|---|---|---|---|---|---|
| chr14:120868670 | 14 | 120.8687 | 0.003218845 | Credible Set | purple |
| chr14:120868693 | 14 | 120.8687 | 0.003218845 | Credible Set | purple |
| chr14:120875959 | 14 | 120.8760 | 0.005142616 | Credible Set | purple |
| chr14:120958410 | 14 | 120.9584 | 0.006586249 | Credible Set | purple |
| chr14:120996497 | 14 | 120.9965 | 0.006559802 | Credible Set | purple |
| chr14:120996551 | 14 | 120.9966 | 0.006559802 | Credible Set | purple |
| chr14:120996555 | 14 | 120.9966 | 0.006559802 | Credible Set | purple |
| chr14:120996573 | 14 | 120.9966 | 0.006559802 | Credible Set | purple |
| chr14:121021982 | 14 | 121.0220 | 0.006559802 | Credible Set | purple |
| chr14:121022009 | 14 | 121.0220 | 0.006559802 | Credible Set | purple |
| chr14:121022048 | 14 | 121.0220 | 0.006559802 | Credible Set | purple |
| chr14:121022050 | 14 | 121.0220 | 0.006559802 | Credible Set | purple |
| chr14:121031640 | 14 | 121.0316 | 0.002521834 | Credible Set | purple |
| chr14:121031672 | 14 | 121.0317 | 0.002521834 | Credible Set | purple |
| chr14:121047713 | 14 | 121.0477 | 0.005965178 | Credible Set | purple |
| chr14:121054065 | 14 | 121.0541 | 0.005742071 | Credible Set | purple |
| chr14:121157624 | 14 | 121.1576 | 0.006559802 | Credible Set | purple |
| chr14:121166355 | 14 | 121.1664 | 0.012117052 | Credible Set | purple |

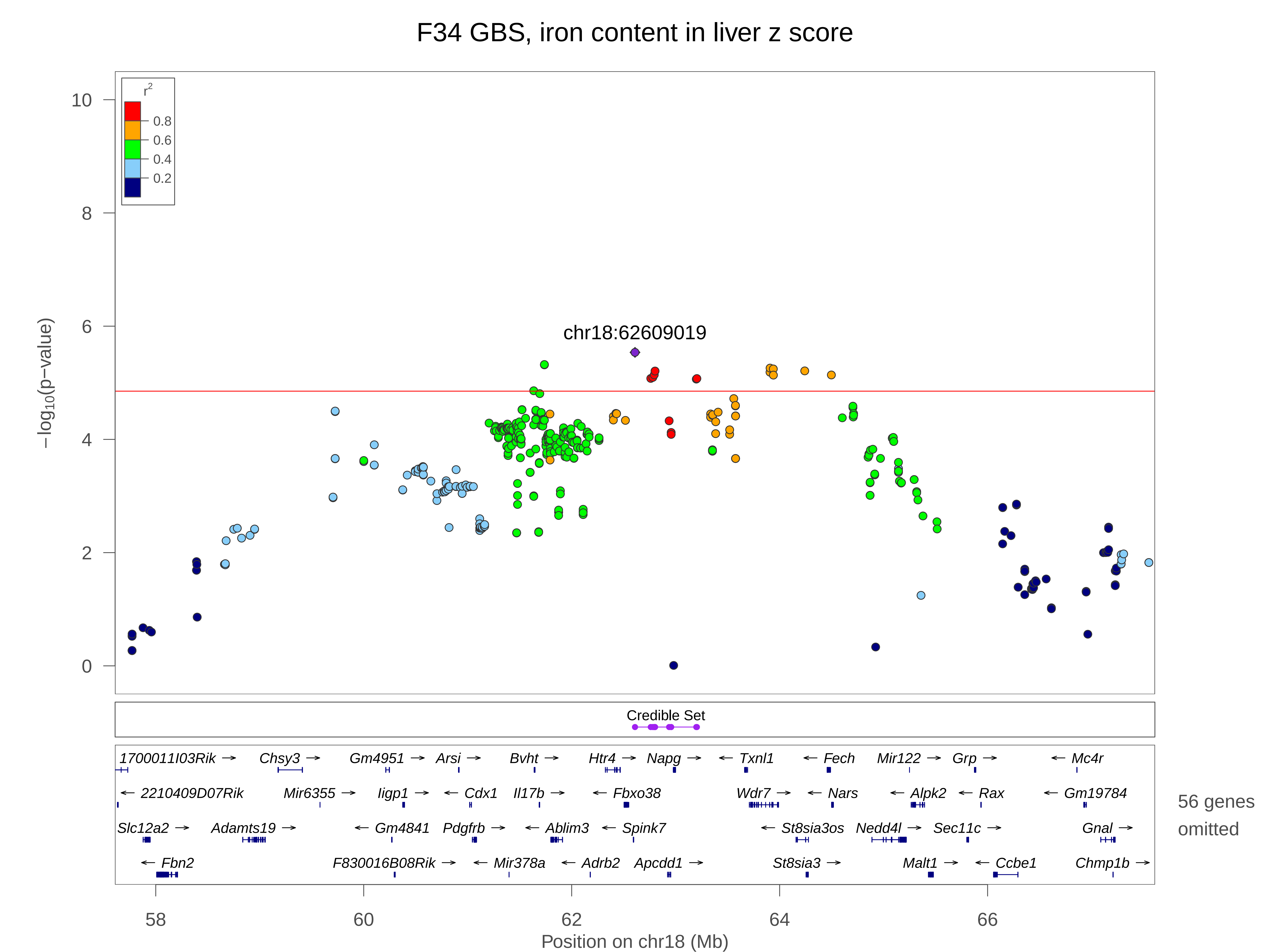

F34 GBS, iron content in liver z score
r2
0.8
0.6
0.4
0.2
10
8
6
4
2
0
−log10(p−value)
chr18:62609019
Credible Set
1700011I03Rik →
Chsy3 →
Gm4951 →
Arsi →
Bvht →
Htr4 →
Napg →
← Txnl1
← Fech
Mir122 →
Grp →
← Mc4r
← 2210409D07Rik
Mir6355 →
Iigp1 →
← Cdx1
Il17b →
← Fbxo38
Wdr7 →
← Nars
← Alpk2
← Rax
← Gm19784
Slc12a2 →
Adamts19 →
← Gm4841
Pdgfrb →
← Ablim3
← Spink7
← St8sia3os
Nedd4l →
Sec11c →
Gnal →
← Fbn2
F830016B08Rik →
← Mir378a
← Adrb2
Apcdd1 →
St8sia3 →
Malt1 →
← Ccbe1
Chmp1b →
56 genes omitted
58
60
62
64
66
Position on chr18 (Mb)

| | |
|---|---|
| date: | Thu Oct 25 17:16:09 2018 |
| build: | F34GBS |
| display range: | chr18:57609019−67609019 [57609019−67609019] |
| hilite range: | 0 − 0 [ 0 − 0 ] |
| reference SNP: | chr18:62609019 |
| number of SNPs plotted: | 481 |
| min P−value: | 2.9E−6 [chr18:62609019] |
| max P−value: | 9.81E−1 [chr18:62980832] |
| omitted Genes: | Slc27a6, Isoc1, A730017C20Rik |
| omitted Genes: | Smim3, Dctn4, Rbm22 |
| omitted Genes: | Myoz3, Synpo, G630071F17Rik |
| omitted Genes: | Ndst1, Rps14, Cd74 |
| omitted Genes: | Mir5107, Tcof1, Camk2a |
| omitted Genes: | Mir7220, Mir6982, Slc6a7 |
| omitted Genes: | Csf1r, Mir6983, Hmgxb3 |
| omitted Genes: | Slc26a2, Pde6a, Ppargc1b |
| omitted Genes: | A530050N04Rik, Arhgef37, Csnk1a1 |
| omitted Genes: | Carmn, Mir145a, Mir143 |
| omitted Genes: | Pcyox1l, Grpel2, 1500015A07Rik |
| omitted Genes: | Afap1l1, Sh3tc2, Spink10 |
| omitted Genes: | Spink13, 2700046A07Rik, Piezo2 |
| omitted Genes: | Onecut2, Atp8b1, Zfp532 |
| omitted Genes: | Oacyl, Cplx4, Lman1 |
| omitted Genes: | Mir694, Pmaip1, Gm9926 |
| omitted Genes: | Mppe1, Impa2, B430212C06Rik |
| omitted Genes: | Cidea, Tubb6, Afg3l2 |
| omitted Genes: | Slmo1, Spire1 |

Make more plots at http://csg.sph.umich.edu/locuszoom/

Fine−mapping Regions

| snp | chr | pos (Mb) | pp | group | color |
|---|---|---|---|---|---|
| chr18:62609019 | 18 | 62.60902 | 0.25031263 | Credible Set | purple |
| chr18:62760743 | 18 | 62.76074 | 0.09083502 | Credible Set | purple |
| chr18:62780679 | 18 | 62.78068 | 0.09443162 | Credible Set | purple |
| chr18:62793979 | 18 | 62.79398 | 0.10439724 | Credible Set | purple |
| chr18:62801634 | 18 | 62.80163 | 0.12100760 | Credible Set | purple |
| chr18:62801661 | 18 | 62.80166 | 0.12100760 | Credible Set | purple |
| chr18:62937328 | 18 | 62.93733 | 0.01757442 | Credible Set | purple |
| chr18:62956832 | 18 | 62.95683 | 0.01123247 | Credible Set | purple |
| chr18:62956840 | 18 | 62.95684 | 0.01041890 | Credible Set | purple |
| chr18:63198665 | 18 | 63.19866 | 0.08841425 | Credible Set | purple |
| chr18:63204145 | 18 | 63.20414 | 0.09036824 | Credible Set | purple |

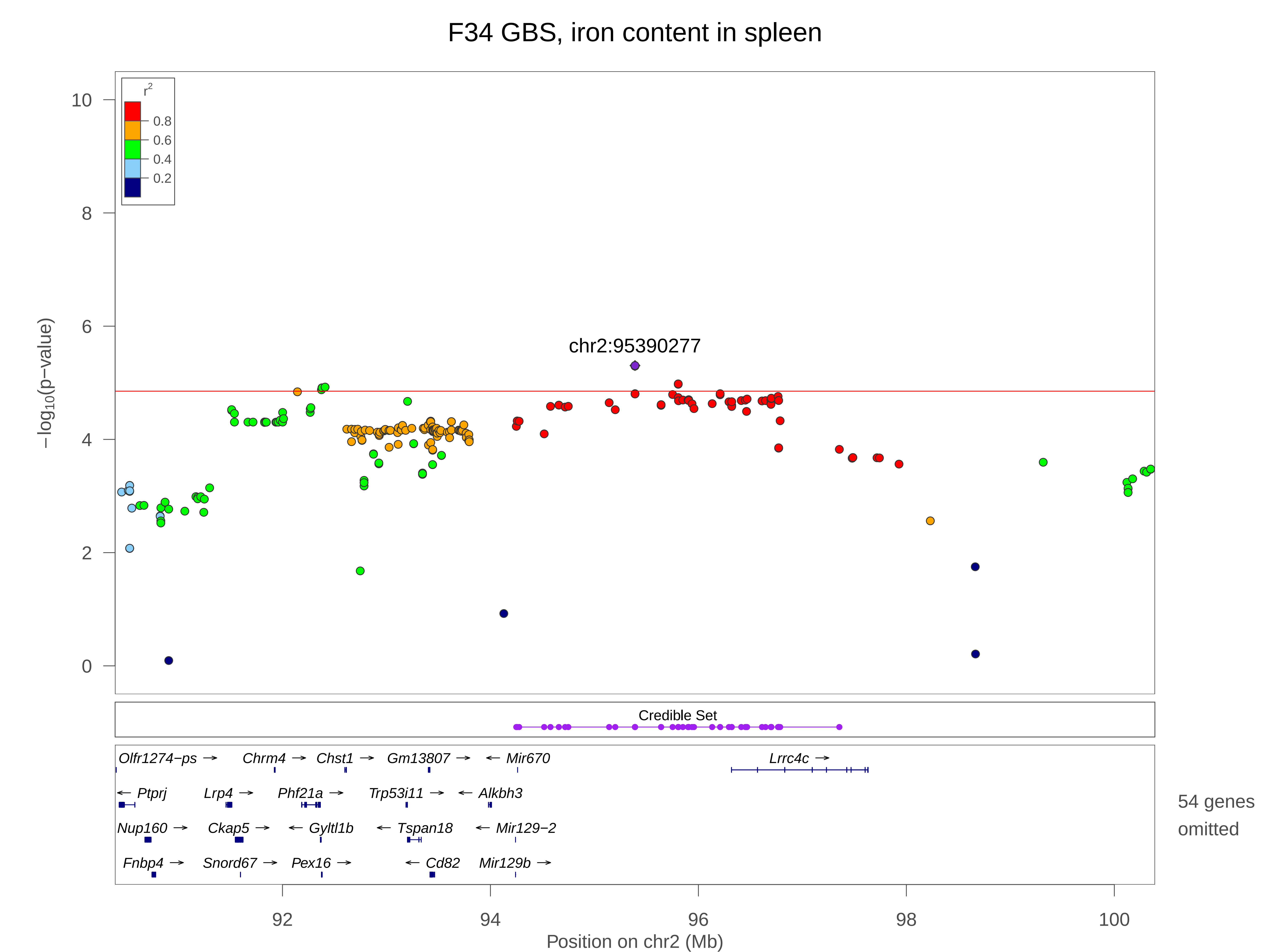

F34 GBS, iron content in spleen
r2
0.8
0.6
0.4
0.2
−log10(p−value)
chr2:95390277
Credible Set
Olfr1274−ps →
Chrm4 →
Chst1 →
Gm13807 →
← Mir670
Lrrc4c →
← Ptprj
Lrp4 →
Phf21a →
Trp53i11 →
← Alkbh3
Nup160 →
Ckap5 →
← Gyltl1b
← Tspan18
← Mir129−2
Fnbp4 →
Snord67 →
Pex16 →
← Cd82
Mir129b →
54 genes omitted
Position on chr2 (Mb)

| | |
|---|---|
| date: | Thu Oct 25 17:16:48 2018 |
| build: | F34GBS |
| display range: | chr2:90390277−100390277 [90390277−100390277] |
| hilite range: | 0 − 0 [ 0 − 0 ] |
| reference SNP: | chr2:95390277 |
| number of SNPs plotted: | 248 |
| min P−value: | 4.96E−6 [chr2:95390277] |
| max P−value: | 8.05E−1 [chr2:90906135] |
| omitted Genes: | Agbl2, Mtch2, C1qtnf4 |
| omitted Genes: | Ndufs3, Kbtbd4, Ptpmt1 |
| omitted Genes: | Celf1, Rapsn, Psmc3 |
| omitted Genes: | Slc39a13, Spi1, Mybpc3 |
| omitted Genes: | Madd, Nr1h3, Acp2 |
| omitted Genes: | Ddb2, A330069E16Rik, Pacsin3 |
| omitted Genes: | Arfgap2, 1110051M20Rik, F2 |
| omitted Genes: | Zfp408, Arhgap1, Atg13 |
| omitted Genes: | Harbi1, Ambra1, Mdk |
| omitted Genes: | Dgkz, Mir6999, Creb3l1 |
| omitted Genes: | Mir1955, 1700029I15Rik, Mapk8ip1 |
| omitted Genes: | Mir7000, Cry2, D930015M05Rik |
| omitted Genes: | Slc35c1, Mir7221, Mir7222 |
| omitted Genes: | Syt13, Prdm11, 4631405J19Rik |
| omitted Genes: | Mir7001, Gm10804, Alx4 |
| omitted Genes: | Ext2, Accs, Accsl |
| omitted Genes: | Gm13889, Alkbh3os1, Hsd17b12 |
| omitted Genes: | Ttc17, 2810002D19Rik, Api5 |

Make more plots at http://csg.sph.umich.edu/locuszoom/

Fine−mapping Regions

| snp | chr | pos (Mb) | pp | group | color |
|---|---|---|---|---|---|
| chr2:94249122 | 2 | 94.24912 | 0.005706935 | Credible Set | purple |
| chr2:94257379 | 2 | 94.25738 | 0.007037982 | Credible Set | purple |
| chr2:94276010 | 2 | 94.27601 | 0.006959189 | Credible Set | purple |
| chr2:94516977 | 2 | 94.51698 | 0.004273585 | Credible Set | purple |
| chr2:94577500 | 2 | 94.57750 | 0.012341045 | Credible Set | purple |
| chr2:94657941 | 2 | 94.65794 | 0.012961878 | Credible Set | purple |
| chr2:94718556 | 2 | 94.71856 | 0.012015751 | Credible Set | purple |
| chr2:94748712 | 2 | 94.74871 | 0.012380196 | Credible Set | purple |
| chr2:95141904 | 2 | 95.14190 | 0.014227263 | Credible Set | purple |
| chr2:95200398 | 2 | 95.20040 | 0.010844635 | Credible Set | purple |
| chr2:95390243 | 2 | 95.39024 | 0.059728141 | Credible Set | purple |
| chr2:95390247 | 2 | 95.39025 | 0.059770866 | Credible Set | purple |
| chr2:95390277 | 2 | 95.39028 | 0.060230096 | Credible Set | purple |
| chr2:95390284 | 2 | 95.39028 | 0.058906829 | Credible Set | purple |
| chr2:95390546 | 2 | 95.39055 | 0.020184125 | Credible Set | purple |
| chr2:95390592 | 2 | 95.39059 | 0.019936029 | Credible Set | purple |
| chr2:95641124 | 2 | 95.64112 | 0.012846967 | Credible Set | purple |
| chr2:95641477 | 2 | 95.64148 | 0.013193742 | Credible Set | purple |
| chr2:95752277 | 2 | 95.75228 | 0.019523766 | Credible Set | purple |
| chr2:95805867 | 2 | 95.80587 | 0.017373588 | Credible Set | purple |
| chr2:95806028 | 2 | 95.80603 | 0.029073764 | Credible Set | purple |
| chr2:95806038 | 2 | 95.80604 | 0.029529119 | Credible Set | purple |
| chr2:95808603 | 2 | 95.80860 | 0.015342283 | Credible Set | purple |
| chr2:95850674 | 2 | 95.85067 | 0.015788549 | Credible Set | purple |
| chr2:95900229 | 2 | 95.90023 | 0.015717564 | Credible Set | purple |
| chr2:95906738 | 2 | 95.90674 | 0.015978506 | Credible Set | purple |
| chr2:95906783 | 2 | 95.90678 | 0.015482285 | Credible Set | purple |
| chr2:95938782 | 2 | 95.93878 | 0.013640948 | Credible Set | purple |
| chr2:95957448 | 2 | 95.95745 | 0.011315624 | Credible Set | purple |
| chr2:96132587 | 2 | 96.13259 | 0.013714611 | Credible Set | purple |
| chr2:96209134 | 2 | 96.20913 | 0.019358695 | Credible Set | purple |
| chr2:96209157 | 2 | 96.20916 | 0.020182630 | Credible Set | purple |
| chr2:96293855 | 2 | 96.29385 | 0.014727956 | Credible Set | purple |
| chr2:96320222 | 2 | 96.32022 | 0.012895956 | Credible Set | purple |
| chr2:96320237 | 2 | 96.32024 | 0.012286318 | Credible Set | purple |
| chr2:96320641 | 2 | 96.32064 | 0.014788430 | Credible Set | purple |

Fine−mapping Regions

| snp | chr | pos (Mb) | pp | group | color |
|---|---|---|---|---|---|
| chr2:96412425 | 2 | 96.41242 | 0.015456362 | Credible Set | purple |
| chr2:96451438 | 2 | 96.45144 | 0.015717564 | Credible Set | purple |
| chr2:96462914 | 2 | 96.46291 | 0.010161779 | Credible Set | purple |
| chr2:96468064 | 2 | 96.46806 | 0.016410181 | Credible Set | purple |
| chr2:96611762 | 2 | 96.61176 | 0.015225620 | Credible Set | purple |
| chr2:96645727 | 2 | 96.64573 | 0.015411943 | Credible Set | purple |
| chr2:96697256 | 2 | 96.69726 | 0.015923691 | Credible Set | purple |
| chr2:96697871 | 2 | 96.69787 | 0.013982276 | Credible Set | purple |
| chr2:96697872 | 2 | 96.69787 | 0.013340510 | Credible Set | purple |
| chr2:96701045 | 2 | 96.70104 | 0.016735851 | Credible Set | purple |
| chr2:96701070 | 2 | 96.70107 | 0.016735851 | Credible Set | purple |
| chr2:96701077 | 2 | 96.70108 | 0.016735851 | Credible Set | purple |
| chr2:96701079 | 2 | 96.70108 | 0.016735851 | Credible Set | purple |
| chr2:96701166 | 2 | 96.70117 | 0.016956979 | Credible Set | purple |
| chr2:96766199 | 2 | 96.76620 | 0.018132457 | Credible Set | purple |
| chr2:96771761 | 2 | 96.77176 | 0.015495858 | Credible Set | purple |
| chr2:96771791 | 2 | 96.77179 | 0.015502934 | Credible Set | purple |
| chr2:96771900 | 2 | 96.77190 | 0.002487796 | Credible Set | purple |
| chr2:96771931 | 2 | 96.77193 | 0.002470562 | Credible Set | purple |
| chr2:96771968 | 2 | 96.77197 | 0.002505934 | Credible Set | purple |
| chr2:96786046 | 2 | 96.78605 | 0.007082194 | Credible Set | purple |
| chr2:97355566 | 2 | 97.35557 | 0.002361193 | Credible Set | purple |

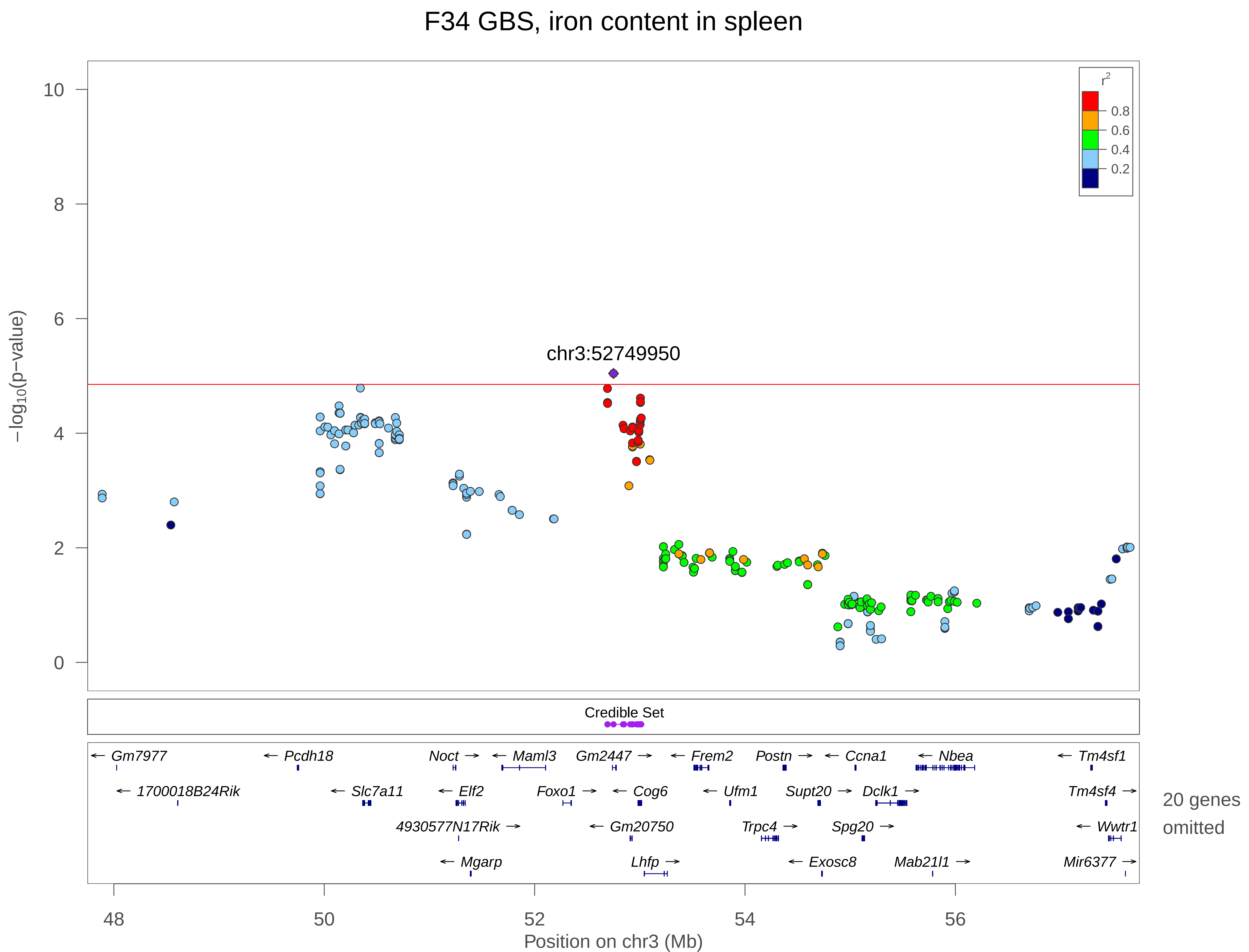

F34 GBS, iron content in spleen
r2
0.8
0.6
0.4
0.2
chr3:52749950
−log10(p−value)
10
8
6
4
2
0
Credible Set
Gm7977
1700018B24Rik
Pcdh18
Slc7a11
Noct
4930577N17Rik
Elf2
Mgarp
Maml3
Foxo1
Gm2447
Gm20750
Cog6
Lhfp
Frem2
Ufm1
Trpc4
Postn
Supt20
Exosc8
Ccna1
Spg20
Dclk1
Nbea
Mab21l1
Tm4sf1
Tm4sf4
Wwtr1
Mir6377
20 genes omitted
48
50
52
54
56
Position on chr3 (Mb)

| | |
|---|---|
| date: | Thu Oct 25 17:16:57 2018 |
| build: | F34GBS |
| display range: | chr3:47749950−57749950 [47749950−57749950] |
| hilite range: | 0 − 0 [ 0 − 0 ] |
| reference SNP: | chr3:52749950 |
| number of SNPs plotted: | 317 |
| min P−value: | 9.06E−6 [chr3:52749950] |
| max P−value: | 5.15E−1 [chr3:54904358] |
| omitted Genes: | Ndufc1, Naa15, Rab33b |
| omitted Genes: | Setd7, 5031434O11Rik, Mgst2 |
| omitted Genes: | Nhlrc3, Proser1, Stoml3 |
| omitted Genes: | Alg5, Smad9, Rfxap |
| omitted Genes: | Sertm1, 4931419H13Rik, Ccdc169 |
| omitted Genes: | Sohlh2, 4933417G07Rik, Commd2 |
| omitted Genes: | Ankub1, Rnf13 |

Make more plots at http://csg.sph.umich.edu/locuszoom/

Fine−mapping Regions

| snp | chr | pos (Mb) | pp | group | color |
|---|---|---|---|---|---|
| chr3:52692409 | 3 | 52.69241 | 0.091070052 | Credible Set | purple |
| chr3:52693102 | 3 | 52.69310 | 0.053261125 | Credible Set | purple |
| chr3:52693118 | 3 | 52.69312 | 0.051424485 | Credible Set | purple |
| chr3:52749950 | 3 | 52.74995 | 0.162176484 | Credible Set | purple |
| chr3:52840387 | 3 | 52.84039 | 0.022454062 | Credible Set | purple |
| chr3:52850138 | 3 | 52.85014 | 0.019591585 | Credible Set | purple |
| chr3:52909024 | 3 | 52.90902 | 0.018062705 | Credible Set | purple |
| chr3:52930454 | 3 | 52.93045 | 0.011449708 | Credible Set | purple |
| chr3:52930496 | 3 | 52.93050 | 0.021017474 | Credible Set | purple |
| chr3:52930984 | 3 | 52.93098 | 0.020415498 | Credible Set | purple |
| chr3:52968213 | 3 | 52.96821 | 0.005732275 | Credible Set | purple |
| chr3:52984307 | 3 | 52.98431 | 0.012393474 | Credible Set | purple |
| chr3:52984901 | 3 | 52.98490 | 0.011875211 | Credible Set | purple |
| chr3:52984912 | 3 | 52.98491 | 0.012712593 | Credible Set | purple |
| chr3:52990872 | 3 | 52.99087 | 0.017261375 | Credible Set | purple |
| chr3:52991306 | 3 | 52.99131 | 0.018288001 | Credible Set | purple |
| chr3:53003556 | 3 | 53.00356 | 0.022880283 | Credible Set | purple |
| chr3:53003614 | 3 | 53.00361 | 0.025714821 | Credible Set | purple |
| chr3:53005849 | 3 | 53.00585 | 0.053327565 | Credible Set | purple |
| chr3:53005874 | 3 | 53.00587 | 0.053759283 | Credible Set | purple |
| chr3:53005885 | 3 | 53.00588 | 0.063184534 | Credible Set | purple |
| chr3:53005909 | 3 | 53.00591 | 0.054663344 | Credible Set | purple |
| chr3:53007126 | 3 | 53.00713 | 0.028248747 | Credible Set | purple |
| chr3:53007129 | 3 | 53.00713 | 0.028100592 | Credible Set | purple |
| chr3:53007177 | 3 | 53.00718 | 0.028100592 | Credible Set | purple |
| chr3:53007214 | 3 | 53.00721 | 0.028152758 | Credible Set | purple |
| chr3:53013134 | 3 | 53.01313 | 0.029551286 | Credible Set | purple |
| chr3:53013153 | 3 | 53.01315 | 0.029486145 | Credible Set | purple |

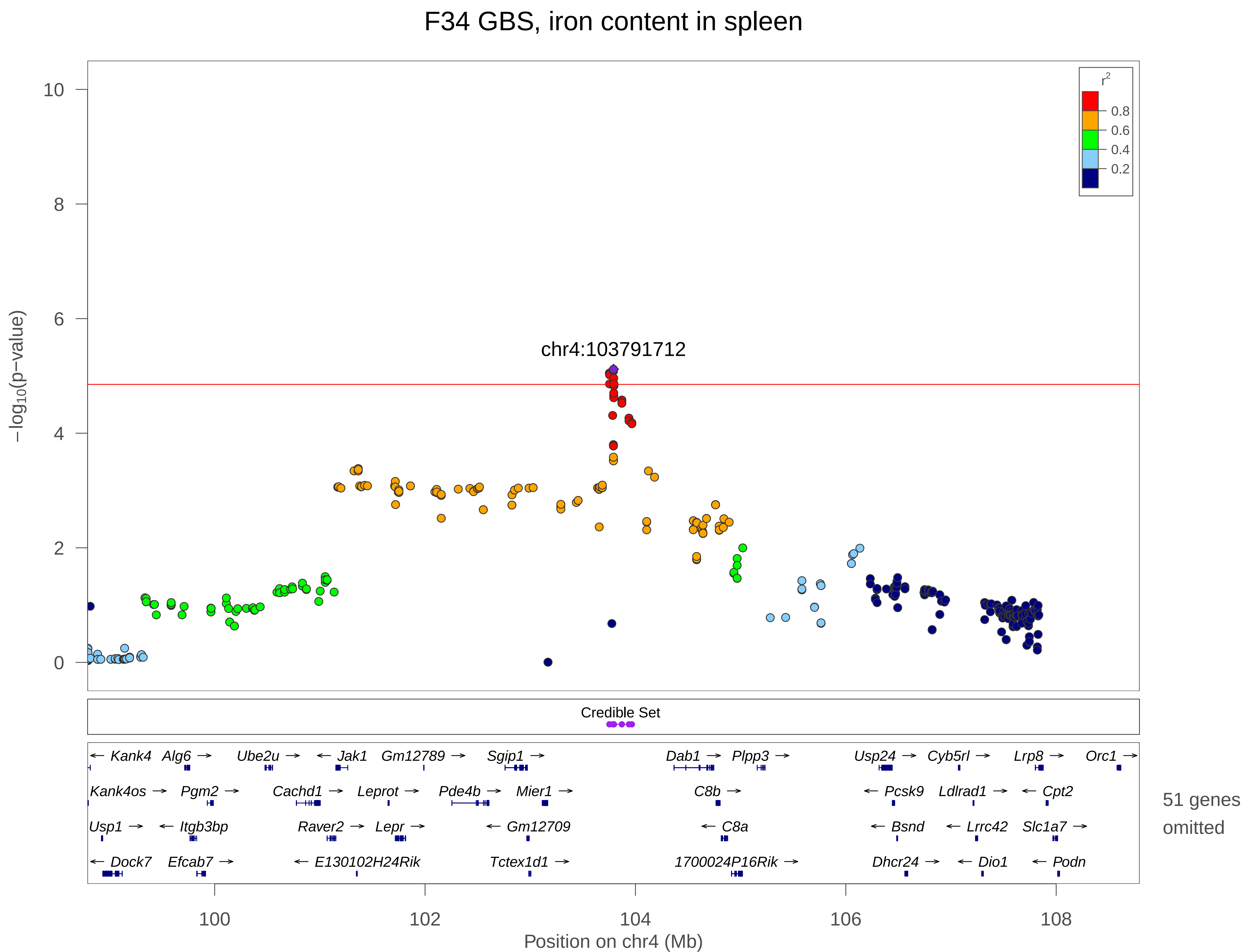

F34 GBS, iron content in spleen
r2
0.8
0.6
0.4
0.2
10
8
6
4
2
0
−log10(p−value)
chr4:103791712
Credible Set
← Kank4 Alg6 → Ube2u → ← Jak1 Gm12789 → Sgip1 → Dab1 → Plpp3 → Usp24 → Cyb5rl → Lrp8 → Orc1 →
Kank4os → Pgm2 → Cachd1 → Leprot → Pde4b → Mier1 → C8b → ← Pcsk9 Ldlrad1 → ← Cpt2
Usp1 → ← Itgb3bp Raver2 → Lepr → ← Gm12709 ← C8a ← Bsnd ← Lrrc42 Slc1a7 →
← Dock7 Efcab7 → ← E130102H24Rik Tctex1d1 → 1700024P16Rik → Dhcr24 → ← Dio1 ← Podn
51 genes omitted
100
102
104
106
108
Position on chr4 (Mb)

| | |
|---|---|
| date: | Thu Oct 25 17:17:06 2018 |
| build: | F34GBS |
| display range: | chr4:98791712−108791712 [98791712−108791712] |
| hilite range: | 0 − 0 [ 0 − 0 ] |
| reference SNP: | chr4:103791712 |
| number of SNPs plotted: | 490 |
| min P−value: | 7.66E−6 [chr4:103791712] |
| max P−value: | 9.9E−1 [chr4:103168898] |
| omitted Genes: | Angptl3, Atg4c, Foxd3 |
| omitted Genes: | Ror1, Mir101a, 0610043K17Rik |
| omitted Genes: | Ak4, Dnajc6, B020004J07Rik |
| omitted Genes: | Gm12794, BB031773, Insl5 |
| omitted Genes: | Wdr78, Slc35d1, 4921539E11Rik |
| omitted Genes: | 4930456L15Rik, Oma1, Gm12718 |
| omitted Genes: | Prkaa2, Lexm, Ttc22 |
| omitted Genes: | Pars2, Ttc4, Mroh7 |
| omitted Genes: | Fam151a, Acot11, Ssbp3 |
| omitted Genes: | Mrpl37, Cdcp2, Tceanc2 |
| omitted Genes: | Tmem59, Hspb11, Yipf1 |
| omitted Genes: | Ndc1, Glis1, Dmrtb1 |
| omitted Genes: | Magoh, 0610037L13Rik, 4930407G08Rik |
| omitted Genes: | Scp2, Mir6397, Echdc2 |
| omitted Genes: | Zyg11a, Zyg11b, Coa7 |
| omitted Genes: | Fam159a, Gpx7, Zcchc11 |
| omitted Genes: | Prpf38a, Cc2d1b, Zfyve9 |

Make more plots at http://csg.sph.umich.edu/locuszoom/

Fine−mapping Regions

| snp | chr | pos (Mb) | pp | group | color |
|---|---|---|---|---|---|
| chr4:103751840 | 4 | 103.7518 | 0.060248995 | Credible Set | purple |
| chr4:103753885 | 4 | 103.7539 | 0.059873440 | Credible Set | purple |
| chr4:103754141 | 4 | 103.7541 | 0.058482338 | Credible Set | purple |
| chr4:103754184 | 4 | 103.7542 | 0.056405548 | Credible Set | purple |
| chr4:103754202 | 4 | 103.7542 | 0.039877523 | Credible Set | purple |
| chr4:103783541 | 4 | 103.7835 | 0.011960620 | Credible Set | purple |
| chr4:103790658 | 4 | 103.7907 | 0.003950926 | Credible Set | purple |
| chr4:103791594 | 4 | 103.7916 | 0.039534427 | Credible Set | purple |
| chr4:103791682 | 4 | 103.7917 | 0.065301368 | Credible Set | purple |
| chr4:103791711 | 4 | 103.7917 | 0.067833988 | Credible Set | purple |
| chr4:103791712 | 4 | 103.7917 | 0.069999100 | Credible Set | purple |
| chr4:103791722 | 4 | 103.7917 | 0.068503483 | Credible Set | purple |
| chr4:103793565 | 4 | 103.7936 | 0.025606589 | Credible Set | purple |
| chr4:103793571 | 4 | 103.7936 | 0.025606589 | Credible Set | purple |
| chr4:103793605 | 4 | 103.7936 | 0.024249422 | Credible Set | purple |
| chr4:103793645 | 4 | 103.7936 | 0.023516073 | Credible Set | purple |
| chr4:103793861 | 4 | 103.7939 | 0.028138877 | Credible Set | purple |
| chr4:103793969 | 4 | 103.7940 | 0.049739800 | Credible Set | purple |
| chr4:103795720 | 4 | 103.7957 | 0.037046784 | Credible Set | purple |
| chr4:103795724 | 4 | 103.7957 | 0.037566074 | Credible Set | purple |
| chr4:103795744 | 4 | 103.7957 | 0.039998714 | Credible Set | purple |
| chr4:103871016 | 4 | 103.8710 | 0.021578348 | Credible Set | purple |
| chr4:103871045 | 4 | 103.8710 | 0.020066791 | Credible Set | purple |
| chr4:103871082 | 4 | 103.8711 | 0.019014715 | Credible Set | purple |
| chr4:103938428 | 4 | 103.9384 | 0.010822195 | Credible Set | purple |
| chr4:103938894 | 4 | 103.9389 | 0.009735167 | Credible Set | purple |
| chr4:103966010 | 4 | 103.9660 | 0.009009925 | Credible Set | purple |
| chr4:103966020 | 4 | 103.9660 | 0.008705388 | Credible Set | purple |

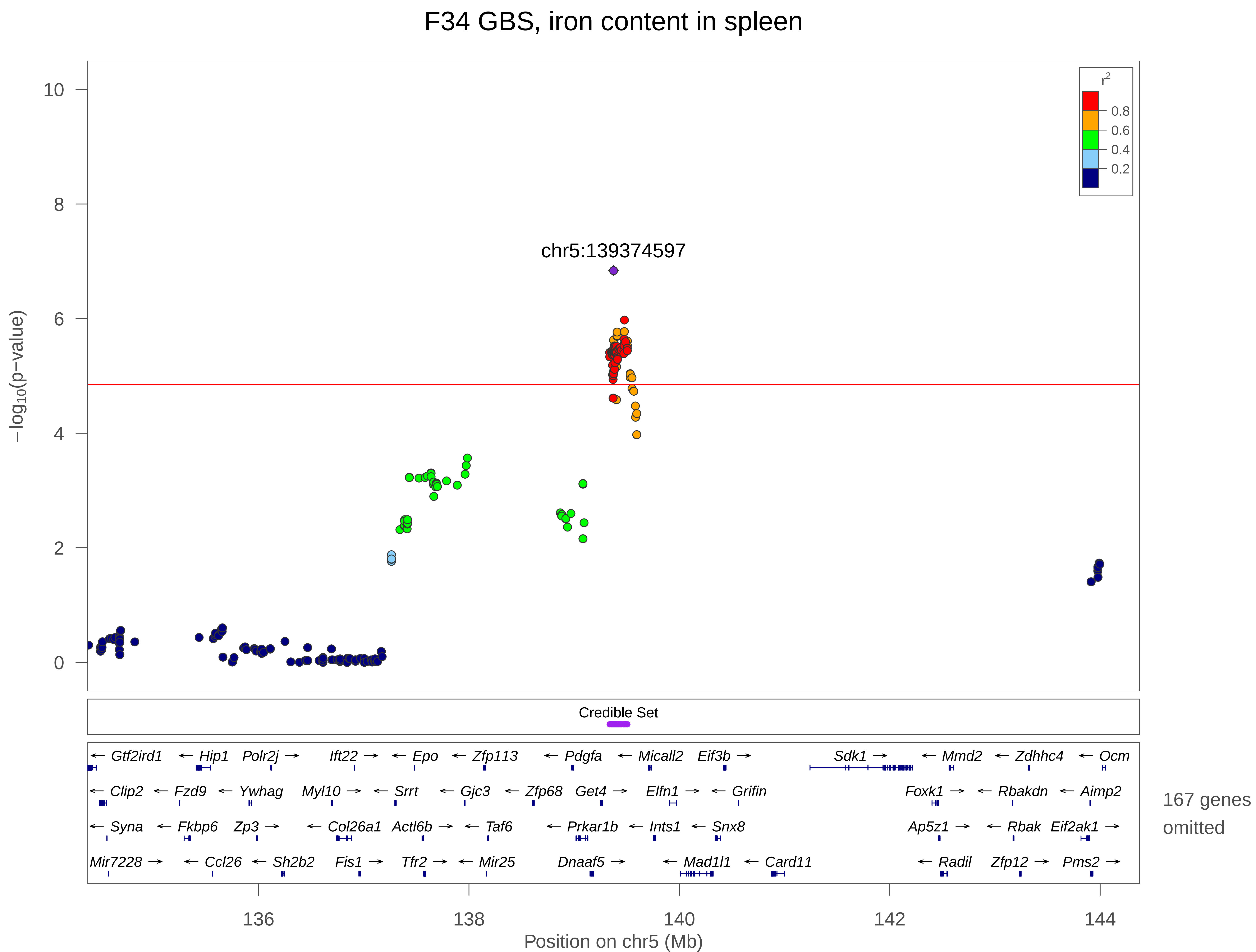

F34 GBS, iron content in spleen
chr5:139374597
−log10(p−value)
r2
0.8
0.6
0.4
0.2
10
8
6
4
2
0
Credible Set
Gtf2ird1
Hip1
Polr2j
Ift22
Epo
Zfp113
Pdgfa
Micall2
Eif3b
Sdk1
Mmd2
Zdhhc4
Ocm
Clip2
Fzd9
Ywhag
Myl10
Srrt
Gjc3
Zfp68
Get4
Elfn1
Grifin
Foxk1
Rbakdn
Aimp2
Syna
Fkbp6
Zp3
Col26a1
Actl6b
Taf6
Prkar1b
Ints1
Snx8
Ap5z1
Rbak
Eif2ak1
Mir7228
Ccl26
Sh2b2
Fis1
Tfr2
Mir25
Dnaaf5
Mad1l1
Card11
Radil
Zfp12
Pms2
167 genes omitted
136
138
140
142
144
Position on chr5 (Mb)

date: Thu Oct 25 17:17:18 2018
build: F34GBS
display range: chr5:134374597−144374597 [134374597−144374597]
hilite range: 0 − 0 [ 0 − 0 ]
reference SNP: chr5:139374597
number of SNPs plotted: 310
min P−value: 1.45E−7 [chr5:139374597]
max P−value: 10E−1 [chr5:136841971]
omitted Genes: Rfc2, Lat2, Eif4h
omitted Genes: Limk1, Eln, Wbscr28
omitted Genes: Cldn13, Wbscr27, Cldn4
omitted Genes: Cldn3, Wbscr25, Abhd11
omitted Genes: Abhd11os, Stx1a, Wbscr22
omitted Genes: Dnajc30, Vps37d, Mlxipl
omitted Genes: Tbl2, Bcl7b, Baz1b
omitted Genes: Trim50, Nsun5, Pom121
omitted Genes: Mir7033, Ccl24, Rhbdd2
omitted Genes: Por, Tmem120a, Mir7034
omitted Genes: Styxl1, Mdh2, Srrm3
omitted Genes: Hspb1, Ssc4d, Dtx2
omitted Genes: Upk3b, Upk3bl, Rasa4
omitted Genes: Mir7035, Lrwd1, Alkbh4
omitted Genes: Orai2, Prkrip1, Cux1
omitted Genes: Mir721, 4933404O12Rik, Cldn15
omitted Genes: Znhit1, Plod3, Mir702
omitted Genes: Vgf, Ap1s1, Serpine1
omitted Genes: Trim56, Ache, Mir8116
omitted Genes: Ufsp1, Mir7036, Trip6
omitted Genes: Slc12a9, Ephb4, Zan
omitted Genes: Pop7, Gigyf1, Gnb2
omitted Genes: Mir6418, Mospd3, Pcolce
omitted Genes: Fbxo24, Lrch4, Gm20605
omitted Genes: Sap25, Irs3, Agfg2
omitted Genes: Nyap1, Tsc22d4, Ppp1r35
omitted Genes: Mepce, Zcwpw1, Pilra
omitted Genes: Pilrb1, Pilrb2, Cyp3a13
omitted Genes: Azgp1, Smok3a, Smok3b
omitted Genes: Smok3c, Zkscan1, Zscan21
omitted Genes: Cops6, Mcm7, Mir93
omitted Genes: Mir106b, Ap4m1, [illegible]

Fine−mapping Regions

| snp | chr | pos (Mb) | pp | group | color |
|---|---|---|---|---|---|
| chr5:139337717 | 5 | 139.3377 | 0.010560657 | Credible Set | purple |
| chr5:139338124 | 5 | 139.3381 | 0.008919521 | Credible Set | purple |
| chr5:139356059 | 5 | 139.3561 | 0.010727733 | Credible Set | purple |
| chr5:139357023 | 5 | 139.3570 | 0.010724215 | Credible Set | purple |
| chr5:139357028 | 5 | 139.3570 | 0.010724215 | Credible Set | purple |
| chr5:139359983 | 5 | 139.3600 | 0.010197570 | Credible Set | purple |
| chr5:139360537 | 5 | 139.3605 | 0.010396517 | Credible Set | purple |
| chr5:139365262 | 5 | 139.3653 | 0.009568568 | Credible Set | purple |
| chr5:139365292 | 5 | 139.3653 | 0.011368721 | Credible Set | purple |
| chr5:139365343 | 5 | 139.3653 | 0.009718053 | Credible Set | purple |
| chr5:139365434 | 5 | 139.3654 | 0.006460345 | Credible Set | purple |
| chr5:139365448 | 5 | 139.3654 | 0.006468110 | Credible Set | purple |
| chr5:139365601 | 5 | 139.3656 | 0.009732867 | Credible Set | purple |
| chr5:139365659 | 5 | 139.3657 | 0.010562277 | Credible Set | purple |
| chr5:139365700 | 5 | 139.3657 | 0.004471959 | Credible Set | purple |
| chr5:139370400 | 5 | 139.3704 | 0.004984709 | Credible Set | purple |
| chr5:139370538 | 5 | 139.3705 | 0.004315281 | Credible Set | purple |
| chr5:139370586 | 5 | 139.3706 | 0.004772895 | Credible Set | purple |
| chr5:139370648 | 5 | 139.3706 | 0.004692687 | Credible Set | purple |
| chr5:139374544 | 5 | 139.3745 | 0.004711422 | Credible Set | purple |
| chr5:139374549 | 5 | 139.3745 | 0.004679596 | Credible Set | purple |
| chr5:139374597 | 5 | 139.3746 | 0.250932238 | Credible Set | purple |
| chr5:139374918 | 5 | 139.3749 | 0.010693262 | Credible Set | purple |
| chr5:139382014 | 5 | 139.3820 | 0.009692099 | Credible Set | purple |
| chr5:139382197 | 5 | 139.3822 | 0.013432980 | Credible Set | purple |
| chr5:139382215 | 5 | 139.3822 | 0.013432980 | Credible Set | purple |
| chr5:139382343 | 5 | 139.3823 | 0.013583832 | Credible Set | purple |
| chr5:139382347 | 5 | 139.3823 | 0.013583832 | Credible Set | purple |
| chr5:139382775 | 5 | 139.3828 | 0.010735423 | Credible Set | purple |
| chr5:139382789 | 5 | 139.3828 | 0.010735423 | Credible Set | purple |
| chr5:139384477 | 5 | 139.3845 | 0.005512212 | Credible Set | purple |
| chr5:139387948 | 5 | 139.3879 | 0.007107830 | Credible Set | purple |
| chr5:139388094 | 5 | 139.3881 | 0.011374631 | Credible Set | purple |
| chr5:139388781 | 5 | 139.3888 | 0.010788820 | Credible Set | purple |
| chr5:139389536 | 5 | 139.3895 | 0.013290725 | Credible Set | purple |
| chr5:139389540 | 5 | 139.3895 | 0.013290725 | Credible Set | purple |

Fine−mapping Regions

| snp | chr | pos (Mb) | pp | group | color |
|---|---|---|---|---|---|
| chr5:139389572 | 5 | 139.3896 | 0.013290725 | Credible Set | purple |
| chr5:139389578 | 5 | 139.3896 | 0.013290725 | Credible Set | purple |
| chr5:139391234 | 5 | 139.3912 | 0.010693262 | Credible Set | purple |
| chr5:139391358 | 5 | 139.3914 | 0.010693262 | Credible Set | purple |
| chr5:139395030 | 5 | 139.3950 | 0.010717465 | Credible Set | purple |
| chr5:139402752 | 5 | 139.4028 | 0.013387738 | Credible Set | purple |
| chr5:139403170 | 5 | 139.4032 | 0.010694234 | Credible Set | purple |
| chr5:139404712 | 5 | 139.4047 | 0.010693262 | Credible Set | purple |
| chr5:139404757 | 5 | 139.4048 | 0.010693262 | Credible Set | purple |
| chr5:139404767 | 5 | 139.4048 | 0.010693262 | Credible Set | purple |
| chr5:139411750 | 5 | 139.4118 | 0.009044341 | Credible Set | purple |
| chr5:139411803 | 5 | 139.4118 | 0.008416576 | Credible Set | purple |
| chr5:139411845 | 5 | 139.4118 | 0.008416576 | Credible Set | purple |
| chr5:139411920 | 5 | 139.4119 | 0.008458430 | Credible Set | purple |
| chr5:139412108 | 5 | 139.4121 | 0.008087735 | Credible Set | purple |
| chr5:139412109 | 5 | 139.4121 | 0.008087735 | Credible Set | purple |
| chr5:139422570 | 5 | 139.4226 | 0.012139100 | Credible Set | purple |
| chr5:139426676 | 5 | 139.4267 | 0.011789144 | Credible Set | purple |
| chr5:139437340 | 5 | 139.4373 | 0.012597787 | Credible Set | purple |
| chr5:139445073 | 5 | 139.4451 | 0.011395209 | Credible Set | purple |
| chr5:139445415 | 5 | 139.4454 | 0.010683502 | Credible Set | purple |
| chr5:139449959 | 5 | 139.4500 | 0.011440936 | Credible Set | purple |
| chr5:139472776 | 5 | 139.4728 | 0.011360873 | Credible Set | purple |
| chr5:139472992 | 5 | 139.4730 | 0.010093789 | Credible Set | purple |
| chr5:139472993 | 5 | 139.4730 | 0.010117467 | Credible Set | purple |
| chr5:139474788 | 5 | 139.4748 | 0.014378625 | Credible Set | purple |
| chr5:139474790 | 5 | 139.4748 | 0.013770989 | Credible Set | purple |
| chr5:139474824 | 5 | 139.4748 | 0.013724204 | Credible Set | purple |
| chr5:139474889 | 5 | 139.4749 | 0.017667987 | Credible Set | purple |
| chr5:139477643 | 5 | 139.4776 | 0.036929212 | Credible Set | purple |
| chr5:139488214 | 5 | 139.4882 | 0.015959674 | Credible Set | purple |
| chr5:139504674 | 5 | 139.5047 | 0.012514153 | Credible Set | purple |
| chr5:139505164 | 5 | 139.5052 | 0.012395512 | Credible Set | purple |
| chr5:139505384 | 5 | 139.5054 | 0.011326291 | Credible Set | purple |

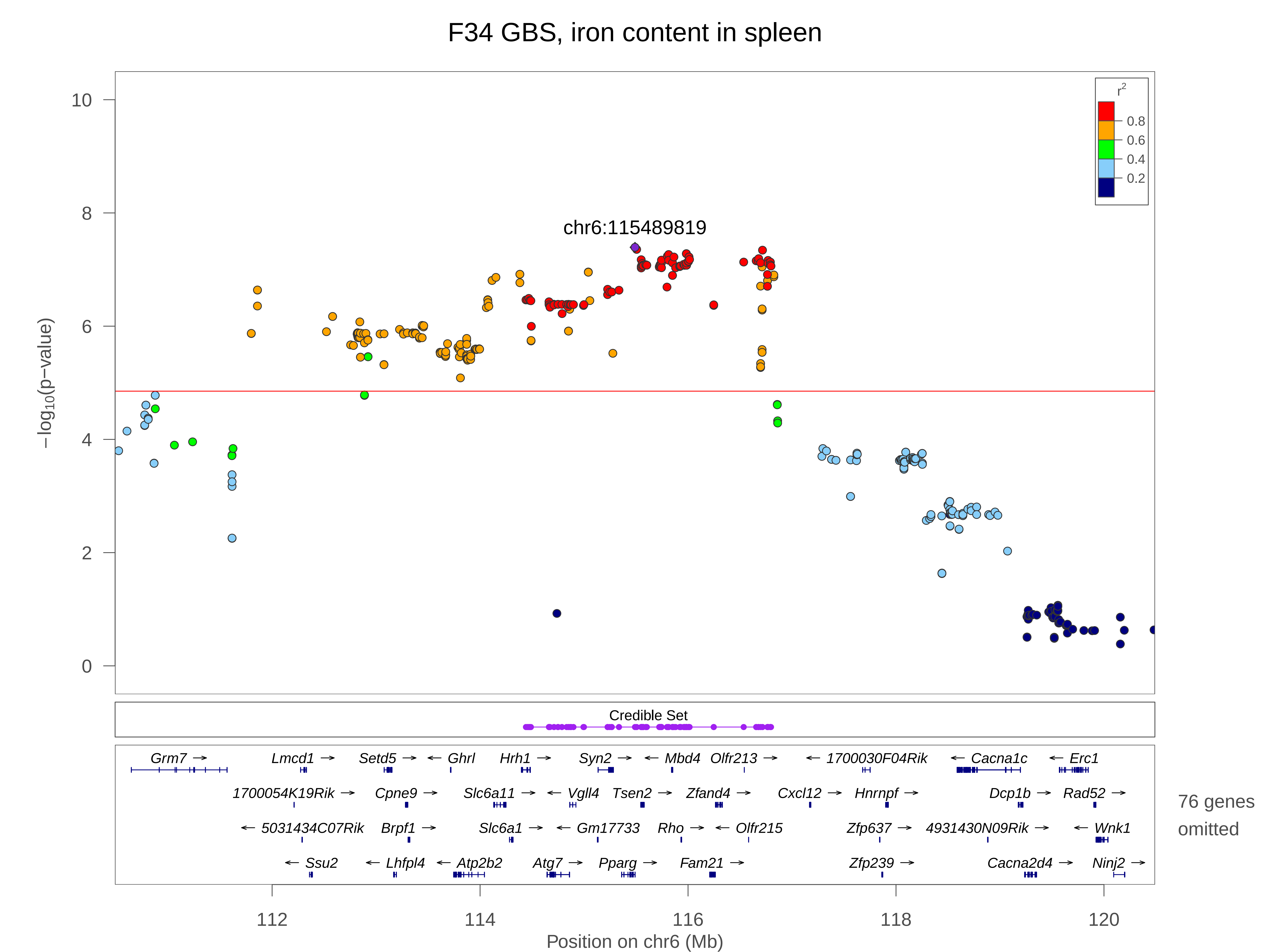

F34 GBS, iron content in spleen
−log10(p−value)
10
8
6
4
2
0
r2
0.8
0.6
0.4
0.2
chr6:115489819
Credible Set
Grm7 →
Lmcd1 →
Setd5 →
← Ghrl
Hrh1 →
Syn2 →
← Mbd4
Olfr213 →
← 1700030F04Rik
← Cacna1c
← Erc1
1700054K19Rik →
Cpne9 →
Slc6a11 →
← Vgll4
Tsen2 →
Zfand4 →
Cxcl12 →
Hnrnpf →
Dcp1b →
Rad52 →
← 5031434C07Rik
Brpf1 →
Slc6a1 →
← Gm17733
Rho →
← Olfr215
Zfp637 →
4931430N09Rik →
← Wnk1
← Ssu2
← Lhfpl4
← Atp2b2
Atg7 →
Pparg →
Fam21 →
Zfp239 →
Cacna2d4 →
Ninj2 →
76 genes omitted
112
114
116
118
120
Position on chr6 (Mb)

| | |
|---|---|
| date: | Thu Oct 25 17:17:29 2018 |
| build: | F34GBS |
| display range: | chr6:110489819−120489819 [110489819−120489819] |
| hilite range: | 0 − 0 [ 0 − 0 ] |
| reference SNP: | chr6:115489819 |
| number of SNPs plotted: | 413 |
| min P−value: | 4.05E−8 [chr6:115489819] |
| max P−value: | 4.11E−1 [chr6:120157431] |
| omitted Genes: | Cav3, Oxtr, Rad18 |
| omitted Genes: | Srgap3, Thumpd3, Gt(ROSA)26Sor |
| omitted Genes: | Mtmr14, Ogg1, Camk1 |
| omitted Genes: | Tada3, Arpc4, Ttll3 |
| omitted Genes: | Rpusd3, Cidec, Jagn1 |
| omitted Genes: | Il17re, Il17rc, Creld1 |
| omitted Genes: | Prrt3, Emc3, Fancd2 |
| omitted Genes: | Fancd2os, Brk1, Vhl |
| omitted Genes: | Irak2, Tatdn2, Mir7042 |
| omitted Genes: | Sec13, Tamm41, Timp4 |
| omitted Genes: | Mkrn2os, Mkrn2, Raf1 |
| omitted Genes: | Tmem40, Cand2, Mir7660 |
| omitted Genes: | Rpl32, Snora7a, Efcab12 |
| omitted Genes: | Ift122, H1foo, Plxnd1 |
| omitted Genes: | Tmcc1, 9530062K07Rik, March8 |
| omitted Genes: | Alox5, Olfr211, Olfr212 |
| omitted Genes: | Olfr214, Zfp422, Rassf4 |
| omitted Genes: | Mir7043, 8430408G22Rik, Tmem72 |
| omitted Genes: | Fxyd4, Rasgef1a, Mir7044 |
| omitted Genes: | Csgalnact2, Ret, 1700069P05Rik |
| omitted Genes: | Gm30498, Bms1, Zfp248 |
| omitted Genes: | Zfp9, Ankrd26, Lrtm2 |
| omitted Genes: | Adipor2, Wnt5b, Fbxl14 |
| omitted Genes: | 3110021A11Rik, Mir706, B4galnt3 |
| omitted Genes: | Ccdc77, Kdm5a, Il17ra |
| omitted Genes: | Cecr6 |

Make more plots at http://csg.sph.umich.edu/locuszoom/

Fine−mapping Regions

| snp | chr | pos (Mb) | pp | group | color |
|---|---|---|---|---|---|
| chr6:114441337 | 6 | 114.4413 | 0.003131998 | Credible Set | purple |
| chr6:114455822 | 6 | 114.4558 | 0.003142268 | Credible Set | purple |
| chr6:114468631 | 6 | 114.4686 | 0.003325717 | Credible Set | purple |
| chr6:114478350 | 6 | 114.4784 | 0.003062969 | Credible Set | purple |
| chr6:114488365 | 6 | 114.4884 | 0.003031376 | Credible Set | purple |
| chr6:114662685 | 6 | 114.6627 | 0.002909267 | Credible Set | purple |
| chr6:114662687 | 6 | 114.6627 | 0.002678002 | Credible Set | purple |
| chr6:114662831 | 6 | 114.6628 | 0.002911317 | Credible Set | purple |
| chr6:114663572 | 6 | 114.6636 | 0.002590470 | Credible Set | purple |
| chr6:114672224 | 6 | 114.6722 | 0.002466579 | Credible Set | purple |
| chr6:114710879 | 6 | 114.7109 | 0.002571449 | Credible Set | purple |
| chr6:114748520 | 6 | 114.7485 | 0.002634972 | Credible Set | purple |
| chr6:114750218 | 6 | 114.7502 | 0.002634972 | Credible Set | purple |
| chr6:114786887 | 6 | 114.7869 | 0.002628276 | Credible Set | purple |
| chr6:114832604 | 6 | 114.8326 | 0.002620164 | Credible Set | purple |
| chr6:114850118 | 6 | 114.8501 | 0.002634972 | Credible Set | purple |
| chr6:114865580 | 6 | 114.8656 | 0.002607757 | Credible Set | purple |
| chr6:114875411 | 6 | 114.8754 | 0.002607757 | Credible Set | purple |
| chr6:114897837 | 6 | 114.8978 | 0.002607757 | Credible Set | purple |
| chr6:114994269 | 6 | 114.9943 | 0.002517055 | Credible Set | purple |
| chr6:114998565 | 6 | 114.9986 | 0.002604341 | Credible Set | purple |
| chr6:115226366 | 6 | 115.2264 | 0.003853860 | Credible Set | purple |
| chr6:115226669 | 6 | 115.2267 | 0.004757316 | Credible Set | purple |
| chr6:115247801 | 6 | 115.2478 | 0.004285612 | Credible Set | purple |
| chr6:115266235 | 6 | 115.2662 | 0.004280605 | Credible Set | purple |
| chr6:115266236 | 6 | 115.2662 | 0.004280605 | Credible Set | purple |
| chr6:115335743 | 6 | 115.3357 | 0.004584876 | Credible Set | purple |
| chr6:115489819 | 6 | 115.4898 | 0.024827641 | Credible Set | purple |
| chr6:115505788 | 6 | 115.5058 | 0.022858162 | Credible Set | purple |
| chr6:115549490 | 6 | 115.5495 | 0.015233174 | Credible Set | purple |
| chr6:115549730 | 6 | 115.5497 | 0.010993232 | Credible Set | purple |
| chr6:115549829 | 6 | 115.5498 | 0.011657837 | Credible Set | purple |
| chr6:115555634 | 6 | 115.5556 | 0.012298781 | Credible Set | purple |
| chr6:115559836 | 6 | 115.5598 | 0.013108472 | Credible Set | purple |
| chr6:115567453 | 6 | 115.5675 | 0.012346112 | Credible Set | purple |
| chr6:115569261 | 6 | 115.5693 | 0.011983096 | Credible Set | purple |

Fine−mapping Regions

| snp | chr | pos (Mb) | pp | group | color |
|---|---|---|---|---|---|
| chr6:115593481 | 6 | 115.5935 | 0.012497097 | Credible Set | purple |
| chr6:115603445 | 6 | 115.6034 | 0.012270603 | Credible Set | purple |
| chr6:115723557 | 6 | 115.7236 | 0.011626473 | Credible Set | purple |
| chr6:115723602 | 6 | 115.7236 | 0.011370495 | Credible Set | purple |
| chr6:115729470 | 6 | 115.7295 | 0.011895550 | Credible Set | purple |
| chr6:115729519 | 6 | 115.7295 | 0.012059737 | Credible Set | purple |
| chr6:115730587 | 6 | 115.7306 | 0.012469945 | Credible Set | purple |
| chr6:115736586 | 6 | 115.7366 | 0.012746512 | Credible Set | purple |
| chr6:115743744 | 6 | 115.7437 | 0.013425921 | Credible Set | purple |
| chr6:115744308 | 6 | 115.7443 | 0.011092578 | Credible Set | purple |
| chr6:115744336 | 6 | 115.7443 | 0.011125369 | Credible Set | purple |
| chr6:115744487 | 6 | 115.7445 | 0.014961390 | Credible Set | purple |
| chr6:115744493 | 6 | 115.7445 | 0.014961390 | Credible Set | purple |
| chr6:115797352 | 6 | 115.7974 | 0.005191679 | Credible Set | purple |
| chr6:115801564 | 6 | 115.8016 | 0.017575283 | Credible Set | purple |
| chr6:115813387 | 6 | 115.8134 | 0.018687205 | Credible Set | purple |
| chr6:115813904 | 6 | 115.8139 | 0.014919512 | Credible Set | purple |
| chr6:115850524 | 6 | 115.8505 | 0.008200616 | Credible Set | purple |
| chr6:115850541 | 6 | 115.8505 | 0.008200616 | Credible Set | purple |
| chr6:115850586 | 6 | 115.8506 | 0.013474106 | Credible Set | purple |
| chr6:115864297 | 6 | 115.8643 | 0.016939179 | Credible Set | purple |
| chr6:115880648 | 6 | 115.8806 | 0.011075584 | Credible Set | purple |
| chr6:115919848 | 6 | 115.9198 | 0.011599903 | Credible Set | purple |
| chr6:115920287 | 6 | 115.9203 | 0.011878528 | Credible Set | purple |
| chr6:115928586 | 6 | 115.9286 | 0.011849687 | Credible Set | purple |
| chr6:115929540 | 6 | 115.9295 | 0.011907240 | Credible Set | purple |
| chr6:115957904 | 6 | 115.9579 | 0.012676989 | Credible Set | purple |
| chr6:115967362 | 6 | 115.9674 | 0.012196402 | Credible Set | purple |
| chr6:115967880 | 6 | 115.9679 | 0.012247326 | Credible Set | purple |
| chr6:115983898 | 6 | 115.9839 | 0.019331309 | Credible Set | purple |
| chr6:115983910 | 6 | 115.9839 | 0.019279810 | Credible Set | purple |
| chr6:115983923 | 6 | 115.9839 | 0.019326769 | Credible Set | purple |
| chr6:115983929 | 6 | 115.9839 | 0.019268930 | Credible Set | purple |
| chr6:115985523 | 6 | 115.9855 | 0.012243389 | Credible Set | purple |
| chr6:115985527 | 6 | 115.9855 | 0.012243389 | Credible Set | purple |
| chr6:115985543 | 6 | 115.9855 | 0.012243389 | Credible Set | purple |

Fine−mapping Regions

| snp | chr | pos (Mb) | pp | group | color |
|---|---|---|---|---|---|
| chr6:115985717 | 6 | 115.9857 | 0.013400852 | Credible Set | purple |
| chr6:116003809 | 6 | 116.0038 | 0.014130965 | Credible Set | purple |
| chr6:116004432 | 6 | 116.0044 | 0.014339466 | Credible Set | purple |
| chr6:116009041 | 6 | 116.0090 | 0.017016046 | Credible Set | purple |
| chr6:116009081 | 6 | 116.0091 | 0.016958752 | Credible Set | purple |
| chr6:116014484 | 6 | 116.0145 | 0.015354426 | Credible Set | purple |
| chr6:116246960 | 6 | 116.2470 | 0.002524237 | Credible Set | purple |
| chr6:116246991 | 6 | 116.2470 | 0.002584386 | Credible Set | purple |
| chr6:116535049 | 6 | 116.5350 | 0.013881652 | Credible Set | purple |
| chr6:116655672 | 6 | 116.6557 | 0.014593999 | Credible Set | purple |
| chr6:116655715 | 6 | 116.6557 | 0.014593999 | Credible Set | purple |
| chr6:116675286 | 6 | 116.6753 | 0.014579158 | Credible Set | purple |
| chr6:116679618 | 6 | 116.6796 | 0.015884492 | Credible Set | purple |
| chr6:116698225 | 6 | 116.6982 | 0.013505759 | Credible Set | purple |
| chr6:116716073 | 6 | 116.7161 | 0.022190232 | Credible Set | purple |
| chr6:116763618 | 6 | 116.7636 | 0.008509094 | Credible Set | purple |
| chr6:116763629 | 6 | 116.7636 | 0.005349393 | Credible Set | purple |
| chr6:116769198 | 6 | 116.7692 | 0.014822402 | Credible Set | purple |
| chr6:116769221 | 6 | 116.7692 | 0.013191775 | Credible Set | purple |
| chr6:116791717 | 6 | 116.7917 | 0.013875766 | Credible Set | purple |
| chr6:116791844 | 6 | 116.7918 | 0.012414677 | Credible Set | purple |
| chr6:116791847 | 6 | 116.7918 | 0.012414677 | Credible Set | purple |
| chr6:116797564 | 6 | 116.7976 | 0.011861166 | Credible Set | purple |

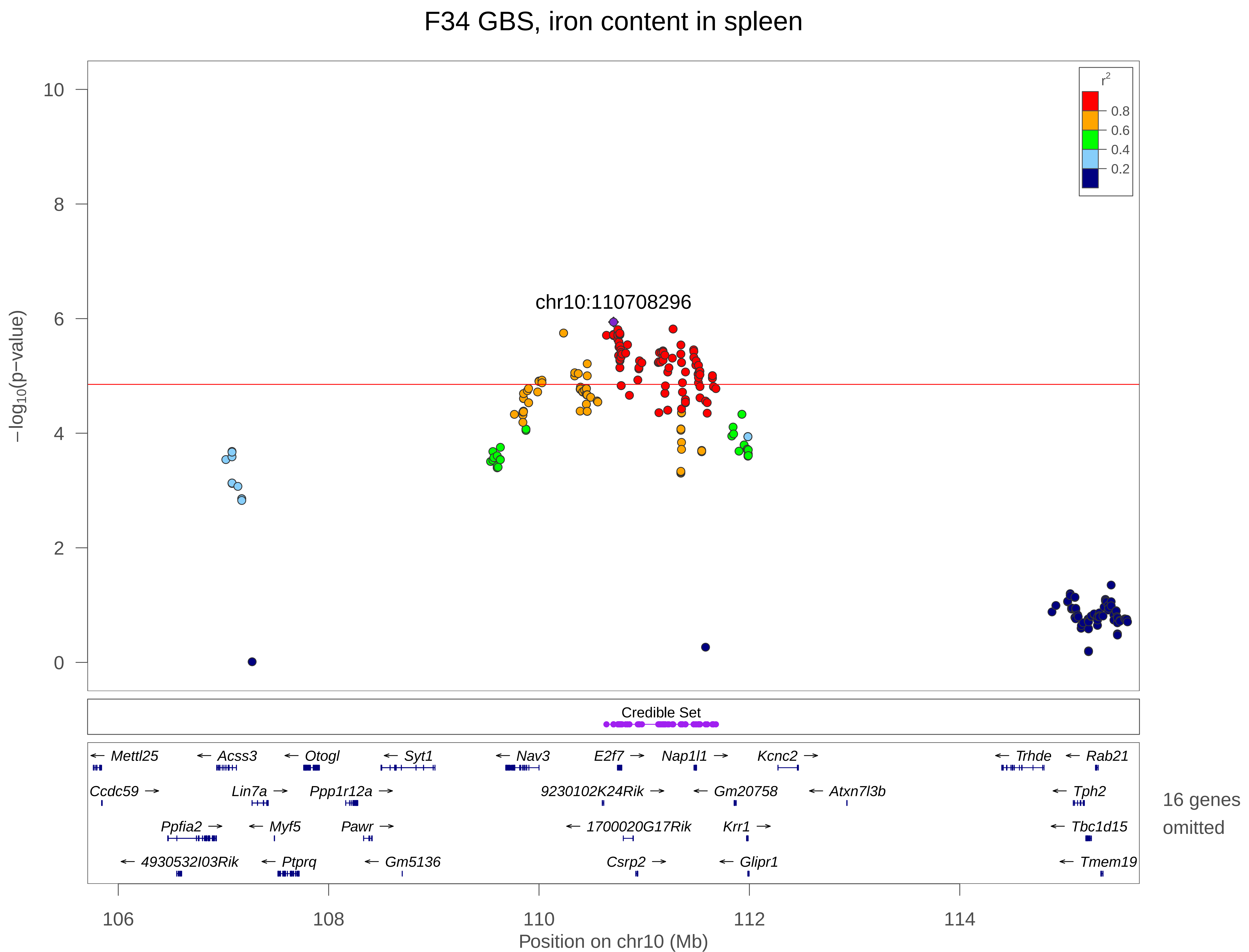

F34 GBS, iron content in spleen
−log10(p−value)
10
8
6
4
2
0
chr10:110708296
r2
0.8
0.6
0.4
0.2
Credible Set
← Mettl25
← Acss3
← Otogl
← Syt1
← Nav3
E2f7 →
Nap1l1 →
Kcnc2 →
← Trhde
← Rab21
Ccdc59 →
Lin7a →
Ppp1r12a →
9230102K24Rik →
← Gm20758
← Atxn7l3b
← Tph2
Ppfia2 →
← Myf5
Pawr →
← 1700020G17Rik
Krr1 →
← Tbc1d15
← 4930532I03Rik
← Ptprq
← Gm5136
Csrp2 →
← Glipr1
← Tmem19
16 genes omitted
106
108
110
112
114
Position on chr10 (Mb)

| | |
|---|---|
| date: | Thu Oct 25 17:16:38 2018 |
| build: | F34GBS |
| display range: | chr10:105708296−115708296 [105708296−115708296] |
| hilite range: | 0 − 0 [ 0 − 0 ] |
| reference SNP: | chr10:110708296 |
| number of SNPs plotted: | 326 |
| min P−value: | 1.15E−6 [chr10:110708296] |
| max P−value: | 9.74E−1 [chr10:107272725] |
| omitted Genes: | Myf6, Gm29685, Zdhhc17 |
| omitted Genes: | Osbpl8, Bbs10, Gm5176 |
| omitted Genes: | Phlda1, 4933440J02Rik, Glipr1l1 |
| omitted Genes: | Glipr1l2, Caps2, 1700010J16Rik |
| omitted Genes: | Thap2, Zfc3h1, Lgr5 |
| omitted Genes: | A930009A15Rik |

Make more plots at http://csg.sph.umich.edu/locuszoom/

Fine−mapping Regions

| snp | chr | pos (Mb) | pp | group | color |
|---|---|---|---|---|---|
| chr10:110641142 | 10 | 110.6411 | 0.021106003 | Credible Set | purple |
| chr10:110708014 | 10 | 110.7080 | 0.021106003 | Credible Set | purple |
| chr10:110708268 | 10 | 110.7083 | 0.021106003 | Credible Set | purple |
| chr10:110708269 | 10 | 110.7083 | 0.021106003 | Credible Set | purple |
| chr10:110708296 | 10 | 110.7083 | 0.041103762 | Credible Set | purple |
| chr10:110708628 | 10 | 110.7086 | 0.021106003 | Credible Set | purple |
| chr10:110749349 | 10 | 110.7493 | 0.021106003 | Credible Set | purple |
| chr10:110749883 | 10 | 110.7499 | 0.021106003 | Credible Set | purple |
| chr10:110749995 | 10 | 110.7500 | 0.021106003 | Credible Set | purple |
| chr10:110750029 | 10 | 110.7500 | 0.021106003 | Credible Set | purple |
| chr10:110750810 | 10 | 110.7508 | 0.021106003 | Credible Set | purple |
| chr10:110756900 | 10 | 110.7569 | 0.010853198 | Credible Set | purple |
| chr10:110757011 | 10 | 110.7570 | 0.010853198 | Credible Set | purple |
| chr10:110759826 | 10 | 110.7598 | 0.014300794 | Credible Set | purple |
| chr10:110760381 | 10 | 110.7604 | 0.014300794 | Credible Set | purple |
| chr10:110768547 | 10 | 110.7685 | 0.014300794 | Credible Set | purple |
| chr10:110768747 | 10 | 110.7687 | 0.006351267 | Credible Set | purple |
| chr10:110768762 | 10 | 110.7688 | 0.006351267 | Credible Set | purple |
| chr10:110768830 | 10 | 110.7688 | 0.021106003 | Credible Set | purple |
| chr10:110768858 | 10 | 110.7689 | 0.021106003 | Credible Set | purple |
| chr10:110768862 | 10 | 110.7689 | 0.021106003 | Credible Set | purple |
| chr10:110768863 | 10 | 110.7689 | 0.021106003 | Credible Set | purple |
| chr10:110768865 | 10 | 110.7689 | 0.021106003 | Credible Set | purple |
| chr10:110769342 | 10 | 110.7693 | 0.008764239 | Credible Set | purple |
| chr10:110769400 | 10 | 110.7694 | 0.008764239 | Credible Set | purple |
| chr10:110769437 | 10 | 110.7694 | 0.008764239 | Credible Set | purple |
| chr10:110769467 | 10 | 110.7695 | 0.008764239 | Credible Set | purple |
| chr10:110769523 | 10 | 110.7695 | 0.008764239 | Credible Set | purple |
| chr10:110769585 | 10 | 110.7696 | 0.008764239 | Credible Set | purple |
| chr10:110778078 | 10 | 110.7781 | 0.010853198 | Credible Set | purple |
| chr10:110778106 | 10 | 110.7781 | 0.010853198 | Credible Set | purple |
| chr10:110778366 | 10 | 110.7784 | 0.008764239 | Credible Set | purple |
| chr10:110780233 | 10 | 110.7802 | 0.008764239 | Credible Set | purple |
| chr10:110781079 | 10 | 110.7811 | 0.003067331 | Credible Set | purple |
| chr10:110781085 | 10 | 110.7811 | 0.003067331 | Credible Set | purple |
| chr10:110781088 | 10 | 110.7811 | 0.003067331 | Credible Set | purple |

Fine−mapping Regions

| snp | chr | pos (Mb) | pp | group | color |
|---|---|---|---|---|---|
| chr10:110782063 | 10 | 110.7821 | 0.010853198 | Credible Set | purple |
| chr10:110790601 | 10 | 110.7906 | 0.010853198 | Credible Set | purple |
| chr10:110824052 | 10 | 110.8241 | 0.010853198 | Credible Set | purple |
| chr10:110841372 | 10 | 110.8414 | 0.014300794 | Credible Set | purple |
| chr10:110859622 | 10 | 110.8596 | 0.002129620 | Credible Set | purple |
| chr10:110939251 | 10 | 110.9393 | 0.003794960 | Credible Set | purple |
| chr10:110949219 | 10 | 110.9492 | 0.005589947 | Credible Set | purple |
| chr10:110949220 | 10 | 110.9492 | 0.005589947 | Credible Set | purple |
| chr10:110949581 | 10 | 110.9496 | 0.006351267 | Credible Set | purple |
| chr10:110955372 | 10 | 110.9554 | 0.008764239 | Credible Set | purple |
| chr10:110977680 | 10 | 110.9777 | 0.007360562 | Credible Set | purple |
| chr10:111134114 | 10 | 111.1341 | 0.007360562 | Credible Set | purple |
| chr10:111139942 | 10 | 111.1399 | 0.007360562 | Credible Set | purple |
| chr10:111139969 | 10 | 111.1400 | 0.007360562 | Credible Set | purple |
| chr10:111145119 | 10 | 111.1451 | 0.010853198 | Credible Set | purple |
| chr10:111156216 | 10 | 111.1562 | 0.007360562 | Credible Set | purple |
| chr10:111162654 | 10 | 111.1627 | 0.010853198 | Credible Set | purple |
| chr10:111178371 | 10 | 111.1784 | 0.010853198 | Credible Set | purple |
| chr10:111178404 | 10 | 111.1784 | 0.010853198 | Credible Set | purple |
| chr10:111179231 | 10 | 111.1792 | 0.010853198 | Credible Set | purple |
| chr10:111179272 | 10 | 111.1793 | 0.008764239 | Credible Set | purple |
| chr10:111180485 | 10 | 111.1805 | 0.010853198 | Credible Set | purple |
| chr10:111180489 | 10 | 111.1805 | 0.010853198 | Credible Set | purple |
| chr10:111196665 | 10 | 111.1967 | 0.010853198 | Credible Set | purple |
| chr10:111196850 | 10 | 111.1968 | 0.002331899 | Credible Set | purple |
| chr10:111196872 | 10 | 111.1969 | 0.002331899 | Credible Set | purple |
| chr10:111201273 | 10 | 111.2013 | 0.003067331 | Credible Set | purple |
| chr10:111223748 | 10 | 111.2237 | 0.004994810 | Credible Set | purple |
| chr10:111234804 | 10 | 111.2348 | 0.006351267 | Credible Set | purple |
| chr10:111267516 | 10 | 111.2675 | 0.008764239 | Credible Set | purple |
| chr10:111274460 | 10 | 111.2745 | 0.021106003 | Credible Set | purple |
| chr10:111274480 | 10 | 111.2745 | 0.021106003 | Credible Set | purple |
| chr10:111347458 | 10 | 111.3475 | 0.010853198 | Credible Set | purple |
| chr10:111348661 | 10 | 111.3487 | 0.010853198 | Credible Set | purple |
| chr10:111348802 | 10 | 111.3488 | 0.014300794 | Credible Set | purple |
| chr10:111354708 | 10 | 111.3547 | 0.007360562 | Credible Set | purple |

Fine−mapping Regions

| snp | chr | pos (Mb) | pp | group | color |
|---|---|---|---|---|---|
| chr10:111363662 | 10 | 111.3637 | 0.003515896 | Credible Set | purple |
| chr10:111363665 | 10 | 111.3637 | 0.002448640 | Credible Set | purple |
| chr10:111391890 | 10 | 111.3919 | 0.001752614 | Credible Set | purple |
| chr10:111391901 | 10 | 111.3919 | 0.001637499 | Credible Set | purple |
| chr10:111391954 | 10 | 111.3920 | 0.001816659 | Credible Set | purple |
| chr10:111391955 | 10 | 111.3920 | 0.001637499 | Credible Set | purple |
| chr10:111392557 | 10 | 111.3926 | 0.004994810 | Credible Set | purple |
| chr10:111470144 | 10 | 111.4701 | 0.010853198 | Credible Set | purple |
| chr10:111471816 | 10 | 111.4718 | 0.010853198 | Credible Set | purple |
| chr10:111471933 | 10 | 111.4719 | 0.008764239 | Credible Set | purple |
| chr10:111490761 | 10 | 111.4908 | 0.007360562 | Credible Set | purple |
| chr10:111494112 | 10 | 111.4941 | 0.008764239 | Credible Set | purple |
| chr10:111512151 | 10 | 111.5122 | 0.004994810 | Credible Set | purple |
| chr10:111515685 | 10 | 111.5157 | 0.003515896 | Credible Set | purple |
| chr10:111516056 | 10 | 111.5161 | 0.004123623 | Credible Set | purple |
| chr10:111516061 | 10 | 111.5161 | 0.006351267 | Credible Set | purple |
| chr10:111522846 | 10 | 111.5228 | 0.005589947 | Credible Set | purple |
| chr10:111529740 | 10 | 111.5297 | 0.005589947 | Credible Set | purple |
| chr10:111529759 | 10 | 111.5298 | 0.004994810 | Credible Set | purple |
| chr10:111530393 | 10 | 111.5304 | 0.003067331 | Credible Set | purple |
| chr10:111530397 | 10 | 111.5304 | 0.001960384 | Credible Set | purple |
| chr10:111583278 | 10 | 111.5833 | 0.001752614 | Credible Set | purple |
| chr10:111598785 | 10 | 111.5988 | 0.001637499 | Credible Set | purple |
| chr10:111648540 | 10 | 111.6485 | 0.004123623 | Credible Set | purple |
| chr10:111648550 | 10 | 111.6486 | 0.004123623 | Credible Set | purple |
| chr10:111648631 | 10 | 111.6486 | 0.004516542 | Credible Set | purple |
| chr10:111658438 | 10 | 111.6584 | 0.003067331 | Credible Set | purple |
| chr10:111658509 | 10 | 111.6585 | 0.003067331 | Credible Set | purple |
| chr10:111681517 | 10 | 111.6815 | 0.002722361 | Credible Set | purple |

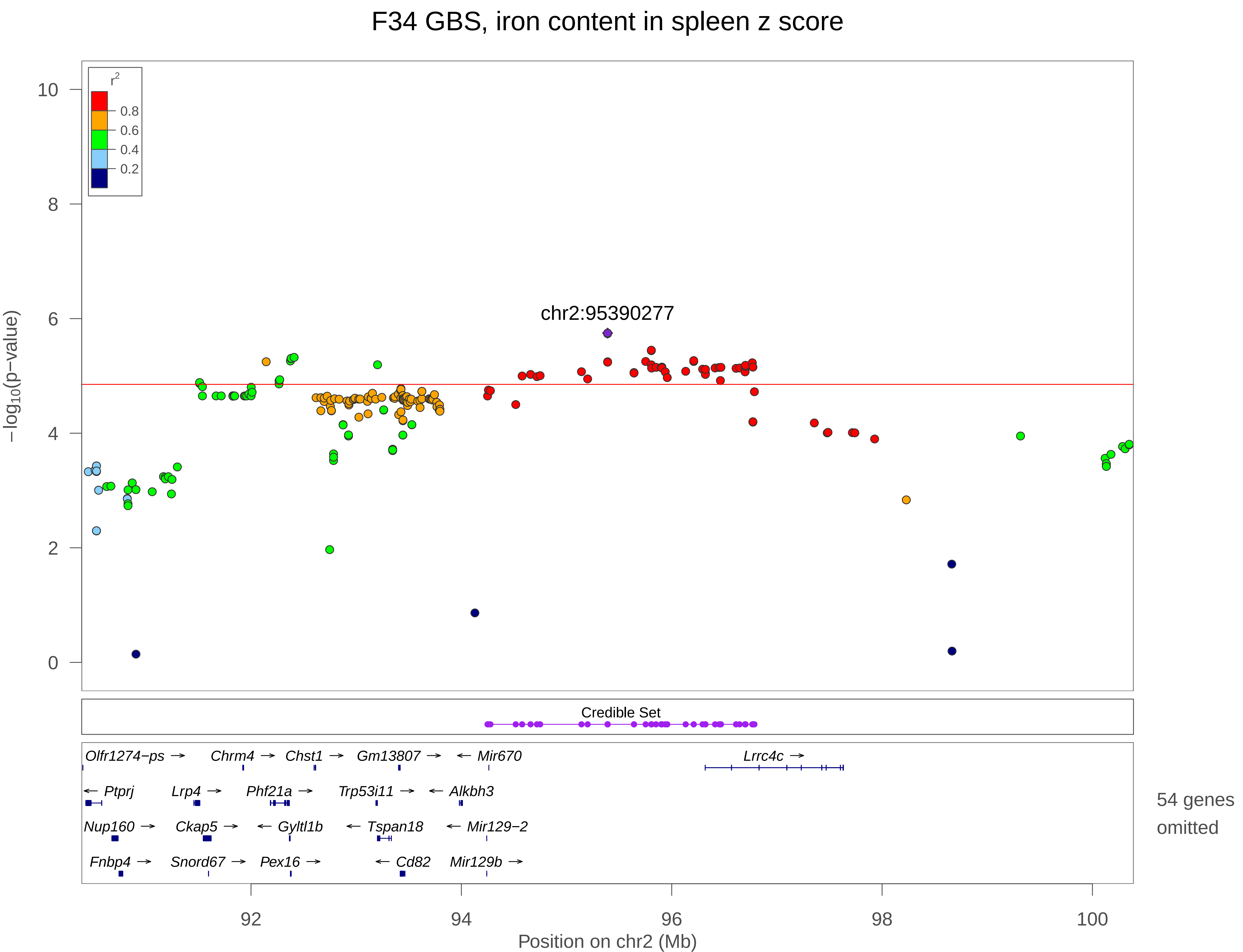

F34 GBS, iron content in spleen z score
r2
0.8
0.6
0.4
0.2
chr2:95390277
−log10(p−value)
Credible Set
Olfr1274−ps →
Chrm4 →
Chst1 →
Gm13807 →
← Mir670
Lrrc4c →
← Ptprj
Lrp4 →
Phf21a →
Trp53i11 →
← Alkbh3
Nup160 →
Ckap5 →
← Gyltl1b
← Tspan18
← Mir129−2
Fnbp4 →
Snord67 →
Pex16 →
← Cd82
Mir129b →
54 genes omitted
Position on chr2 (Mb)

| | |
|---|---|
| date: | Thu Oct 25 17:17:49 2018 |
| build: | F34GBS |
| display range: | chr2:90390277−100390277 [90390277−100390277] |
| hilite range: | 0 − 0 [ 0 − 0 ] |
| reference SNP: | chr2:95390277 |
| number of SNPs plotted: | 248 |
| min P−value: | 1.79E−6 [chr2:95390277] |
| max P−value: | 7.17E−1 [chr2:90906135] |
| omitted Genes: | Agbl2, Mtch2, C1qtnf4 |
| omitted Genes: | Ndufs3, Kbtbd4, Ptpmt1 |
| omitted Genes: | Celf1, Rapsn, Psmc3 |
| omitted Genes: | Slc39a13, Spi1, Mybpc3 |
| omitted Genes: | Madd, Nr1h3, Acp2 |
| omitted Genes: | Ddb2, A330069E16Rik, Pacsin3 |
| omitted Genes: | Arfgap2, 1110051M20Rik, F2 |
| omitted Genes: | Zfp408, Arhgap1, Atg13 |
| omitted Genes: | Harbi1, Ambra1, Mdk |
| omitted Genes: | Dgkz, Mir6999, Creb3l1 |
| omitted Genes: | Mir1955, 1700029I15Rik, Mapk8ip1 |
| omitted Genes: | Mir7000, Cry2, D930015M05Rik |
| omitted Genes: | Slc35c1, Mir7221, Mir7222 |
| omitted Genes: | Syt13, Prdm11, 4631405J19Rik |
| omitted Genes: | Mir7001, Gm10804, Alx4 |
| omitted Genes: | Ext2, Accs, Accsl |
| omitted Genes: | Gm13889, Alkbh3os1, Hsd17b12 |
| omitted Genes: | Ttc17, 2810002D19Rik, Api5 |

Make more plots at http://csg.sph.umich.edu/locuszoom/

Fine−mapping Regions

| snp | chr | pos (Mb) | pp | group | color |
|---|---|---|---|---|---|
| chr2:94249122 | 2 | 94.24912 | 0.005363639 | Credible Set | purple |
| chr2:94257379 | 2 | 94.25738 | 0.006731559 | Credible Set | purple |
| chr2:94276010 | 2 | 94.27601 | 0.006572847 | Credible Set | purple |
| chr2:94516977 | 2 | 94.51698 | 0.003868297 | Credible Set | purple |
| chr2:94577500 | 2 | 94.57750 | 0.011545684 | Credible Set | purple |
| chr2:94657941 | 2 | 94.65794 | 0.012263420 | Credible Set | purple |
| chr2:94718556 | 2 | 94.71856 | 0.011267592 | Credible Set | purple |
| chr2:94748712 | 2 | 94.74871 | 0.011701742 | Credible Set | purple |
| chr2:95141904 | 2 | 95.14190 | 0.013610171 | Credible Set | purple |
| chr2:95200398 | 2 | 95.20040 | 0.010328782 | Credible Set | purple |
| chr2:95390243 | 2 | 95.39024 | 0.059766693 | Credible Set | purple |
| chr2:95390247 | 2 | 95.39025 | 0.059840800 | Credible Set | purple |
| chr2:95390277 | 2 | 95.39028 | 0.060192977 | Credible Set | purple |
| chr2:95390284 | 2 | 95.39028 | 0.058968519 | Credible Set | purple |
| chr2:95390546 | 2 | 95.39055 | 0.019940281 | Credible Set | purple |
| chr2:95390592 | 2 | 95.39059 | 0.019636641 | Credible Set | purple |
| chr2:95641124 | 2 | 95.64112 | 0.013175078 | Credible Set | purple |
| chr2:95641477 | 2 | 95.64148 | 0.012917785 | Credible Set | purple |
| chr2:95752277 | 2 | 95.75228 | 0.020177296 | Credible Set | purple |
| chr2:95805867 | 2 | 95.80587 | 0.017731199 | Credible Set | purple |
| chr2:95806028 | 2 | 95.80603 | 0.030582477 | Credible Set | purple |
| chr2:95806038 | 2 | 95.80604 | 0.031166796 | Credible Set | purple |
| chr2:95808603 | 2 | 95.80860 | 0.015643848 | Credible Set | purple |
| chr2:95850674 | 2 | 95.85067 | 0.016109648 | Credible Set | purple |
| chr2:95900229 | 2 | 95.90023 | 0.016002886 | Credible Set | purple |
| chr2:95906738 | 2 | 95.90674 | 0.016267898 | Credible Set | purple |
| chr2:95906783 | 2 | 95.90678 | 0.015718414 | Credible Set | purple |
| chr2:95938782 | 2 | 95.93878 | 0.013502048 | Credible Set | purple |
| chr2:95957448 | 2 | 95.95745 | 0.010847978 | Credible Set | purple |
| chr2:96132587 | 2 | 96.13259 | 0.013828976 | Credible Set | purple |
| chr2:96209134 | 2 | 96.20913 | 0.020164104 | Credible Set | purple |
| chr2:96209157 | 2 | 96.20916 | 0.020860168 | Credible Set | purple |
| chr2:96293855 | 2 | 96.29385 | 0.014998462 | Credible Set | purple |
| chr2:96320222 | 2 | 96.32022 | 0.012844909 | Credible Set | purple |
| chr2:96320237 | 2 | 96.32024 | 0.012264114 | Credible Set | purple |
| chr2:96320641 | 2 | 96.32064 | 0.014907873 | Credible Set | purple |

Fine−mapping Regions

| snp | chr | pos (Mb) | pp | group | color |
|---|---|---|---|---|---|
| chr2:96412425 | 2 | 96.41242 | 0.015702216 | Credible Set | purple |
| chr2:96451438 | 2 | 96.45144 | 0.016002886 | Credible Set | purple |
| chr2:96462914 | 2 | 96.46291 | 0.009700138 | Credible Set | purple |
| chr2:96468064 | 2 | 96.46806 | 0.016103254 | Credible Set | purple |
| chr2:96611762 | 2 | 96.61176 | 0.015506139 | Credible Set | purple |
| chr2:96645727 | 2 | 96.64573 | 0.015691358 | Credible Set | purple |
| chr2:96697256 | 2 | 96.69726 | 0.016234165 | Credible Set | purple |
| chr2:96697871 | 2 | 96.69787 | 0.013936852 | Credible Set | purple |
| chr2:96697872 | 2 | 96.69787 | 0.013527542 | Credible Set | purple |
| chr2:96701045 | 2 | 96.70104 | 0.016647624 | Credible Set | purple |
| chr2:96701070 | 2 | 96.70107 | 0.016647624 | Credible Set | purple |
| chr2:96701077 | 2 | 96.70108 | 0.016647624 | Credible Set | purple |
| chr2:96701079 | 2 | 96.70108 | 0.016647624 | Credible Set | purple |
| chr2:96701166 | 2 | 96.70117 | 0.017405925 | Credible Set | purple |
| chr2:96766199 | 2 | 96.76620 | 0.019191454 | Credible Set | purple |
| chr2:96771761 | 2 | 96.77176 | 0.016261856 | Credible Set | purple |
| chr2:96771791 | 2 | 96.77179 | 0.016262277 | Credible Set | purple |
| chr2:96771900 | 2 | 96.77190 | 0.001994975 | Credible Set | purple |
| chr2:96771931 | 2 | 96.77193 | 0.001982158 | Credible Set | purple |
| chr2:96771968 | 2 | 96.77197 | 0.002010158 | Credible Set | purple |
| chr2:96786046 | 2 | 96.78605 | 0.006313408 | Credible Set | purple |

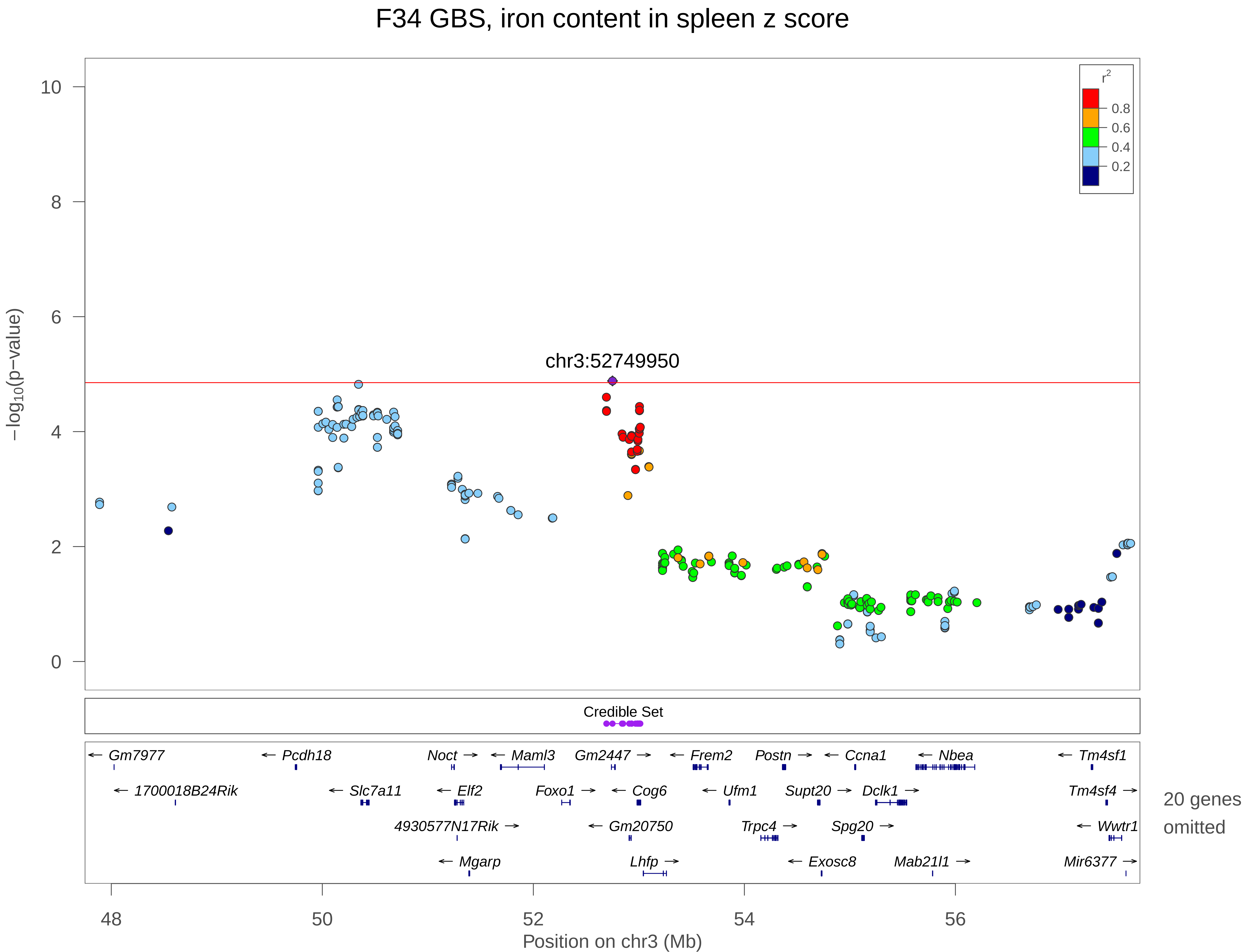
F34 GBS, iron content in spleen z score
chr3:52749950
−log10(p−value)
r2
0.8
0.6
0.4
0.2
Credible Set
Gm7977
Pcdh18
Noct
Maml3
Gm2447
Frem2
Postn
Ccna1
Nbea
Tm4sf1
1700018B24Rik
Slc7a11
Elf2
Foxo1
Cog6
Ufm1
Supt20
Dclk1
Tm4sf4
4930577N17Rik
Gm20750
Trpc4
Spg20
Wwtr1
Mgarp
Lhfp
Exosc8
Mab21l1
Mir6377
20 genes omitted
48
50
52
54
56
Position on chr3 (Mb)

| | |
|---|---|
| date: | Thu Oct 25 17:17:58 2018 |
| build: | F34GBS |
| display range: | chr3:47749950−57749950 [47749950−57749950] |
| hilite range: | 0 − 0 [ 0 − 0 ] |
| reference SNP: | chr3:52749950 |
| number of SNPs plotted: | 317 |
| min P−value: | 1.31E−5 [chr3:52749950] |
| max P−value: | 4.95E−1 [chr3:54904358] |
| omitted Genes: | Ndufc1, Naa15, Rab33b |
| omitted Genes: | Setd7, 5031434O11Rik, Mgst2 |
| omitted Genes: | Nhlrc3, Proser1, Stoml3 |
| omitted Genes: | Alg5, Smad9, Rfxap |
| omitted Genes: | Sertm1, 4931419H13Rik, Ccdc169 |
| omitted Genes: | Sohlh2, 4933417G07Rik, Commd2 |
| omitted Genes: | Ankub1, Rnf13 |

Make more plots at http://csg.sph.umich.edu/locuszoom/

Fine−mapping Regions

| snp | chr | pos (Mb) | pp | group | color |
|---|---|---|---|---|---|
| chr3:52692409 | 3 | 52.69241 | 0.089838719 | Credible Set | purple |
| chr3:52693102 | 3 | 52.69310 | 0.054132032 | Credible Set | purple |
| chr3:52693118 | 3 | 52.69312 | 0.052301075 | Credible Set | purple |
| chr3:52749950 | 3 | 52.74995 | 0.167096637 | Credible Set | purple |
| chr3:52840387 | 3 | 52.84039 | 0.022309578 | Credible Set | purple |
| chr3:52850138 | 3 | 52.85014 | 0.019643067 | Credible Set | purple |
| chr3:52909024 | 3 | 52.90902 | 0.018037015 | Credible Set | purple |
| chr3:52930454 | 3 | 52.93045 | 0.011306871 | Credible Set | purple |
| chr3:52930496 | 3 | 52.93050 | 0.021032229 | Credible Set | purple |
| chr3:52930984 | 3 | 52.93098 | 0.020376718 | Credible Set | purple |
| chr3:52968213 | 3 | 52.96821 | 0.005893668 | Credible Set | purple |
| chr3:52984307 | 3 | 52.98431 | 0.011980952 | Credible Set | purple |
| chr3:52984901 | 3 | 52.98490 | 0.011558553 | Credible Set | purple |
| chr3:52984912 | 3 | 52.98491 | 0.012424554 | Credible Set | purple |
| chr3:52990872 | 3 | 52.99087 | 0.017071957 | Credible Set | purple |
| chr3:52991306 | 3 | 52.99131 | 0.018133903 | Credible Set | purple |
| chr3:53003556 | 3 | 53.00356 | 0.022927130 | Credible Set | purple |
| chr3:53003614 | 3 | 53.00361 | 0.026843355 | Credible Set | purple |
| chr3:53005849 | 3 | 53.00585 | 0.053732447 | Credible Set | purple |
| chr3:53005874 | 3 | 53.00587 | 0.054050290 | Credible Set | purple |
| chr3:53005885 | 3 | 53.00588 | 0.063157304 | Credible Set | purple |
| chr3:53005909 | 3 | 53.00591 | 0.054810868 | Credible Set | purple |
| chr3:53007126 | 3 | 53.00713 | 0.027090896 | Credible Set | purple |
| chr3:53007129 | 3 | 53.00713 | 0.026948953 | Credible Set | purple |
| chr3:53007177 | 3 | 53.00718 | 0.026948953 | Credible Set | purple |
| chr3:53007214 | 3 | 53.00721 | 0.026987237 | Credible Set | purple |
| chr3:53013134 | 3 | 53.01313 | 0.028823993 | Credible Set | purple |
| chr3:53013153 | 3 | 53.01315 | 0.028734960 | Credible Set | purple |

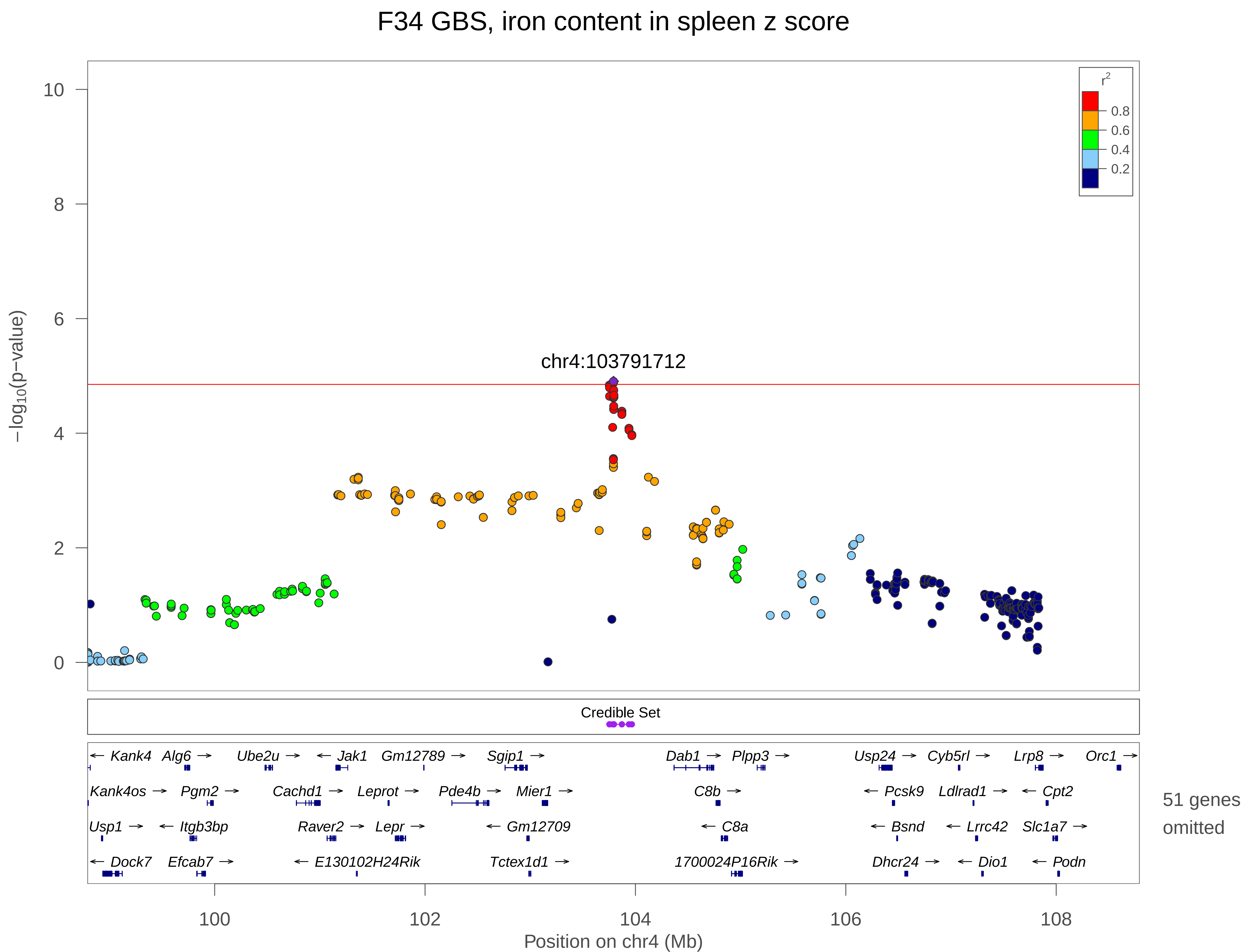

F34 GBS, iron content in spleen z score
r2
0.8
0.6
0.4
0.2
chr4:103791712
−log10(p−value)
0
2
4
6
8
10
Credible Set
← Kank4 Alg6 → Ube2u → ← Jak1 Gm12789 → Sgip1 → Dab1 → Plpp3 → Usp24 → Cyb5rl → Lrp8 → Orc1 →
Kank4os → Pgm2 → Cachd1 → Leprot → Pde4b → Mier1 → C8b → ← Pcsk9 Ldlrad1 → ← Cpt2
Usp1 → ← Itgb3bp Raver2 → Lepr → ← Gm12709 ← C8a ← Bsnd ← Lrrc42 Slc1a7 →
← Dock7 Efcab7 → ← E130102H24Rik Tctex1d1 → 1700024P16Rik → Dhcr24 → ← Dio1 ← Podn
51 genes omitted
100
102
104
106
108
Position on chr4 (Mb)

| | |
|---|---|
| date: | Thu Oct 25 17:18:07 2018 |
| build: | F34GBS |
| display range: | chr4:98791712−108791712 [98791712−108791712] |
| hilite range: | 0 − 0 [ 0 − 0 ] |
| reference SNP: | chr4:103791712 |
| number of SNPs plotted: | 490 |
| min P−value: | 1.24E−5 [chr4:103791712] |
| max P−value: | 9.9E−1 [chr4:98796618] |
| omitted Genes: | Angptl3, Atg4c, Foxd3 |
| omitted Genes: | Ror1, Mir101a, 0610043K17Rik |
| omitted Genes: | Ak4, Dnajc6, B020004J07Rik |
| omitted Genes: | Gm12794, BB031773, Insl5 |
| omitted Genes: | Wdr78, Slc35d1, 4921539E11Rik |
| omitted Genes: | 4930456L15Rik, Oma1, Gm12718 |
| omitted Genes: | Prkaa2, Lexm, Ttc22 |
| omitted Genes: | Pars2, Ttc4, Mroh7 |
| omitted Genes: | Fam151a, Acot11, Ssbp3 |
| omitted Genes: | Mrpl37, Cdcp2, Tceanc2 |
| omitted Genes: | Tmem59, Hspb11, Yipf1 |
| omitted Genes: | Ndc1, Glis1, Dmrtb1 |
| omitted Genes: | Magoh, 0610037L13Rik, 4930407G08Rik |
| omitted Genes: | Scp2, Mir6397, Echdc2 |
| omitted Genes: | Zyg11a, Zyg11b, Coa7 |
| omitted Genes: | Fam159a, Gpx7, Zcchc11 |
| omitted Genes: | Prpf38a, Cc2d1b, Zfyve9 |

Make more plots at http://csg.sph.umich.edu/locuszoom/

Fine−mapping Regions

| snp | chr | pos (Mb) | pp | group | color |
|---|---|---|---|---|---|
| chr4:103751840 | 4 | 103.7518 | 0.060477508 | Credible Set | purple |
| chr4:103753885 | 4 | 103.7539 | 0.060155313 | Credible Set | purple |
| chr4:103754141 | 4 | 103.7541 | 0.056628681 | Credible Set | purple |
| chr4:103754184 | 4 | 103.7542 | 0.054737834 | Credible Set | purple |
| chr4:103754202 | 4 | 103.7542 | 0.039516345 | Credible Set | purple |
| chr4:103783541 | 4 | 103.7835 | 0.012088639 | Credible Set | purple |
| chr4:103790658 | 4 | 103.7907 | 0.003708715 | Credible Set | purple |
| chr4:103791594 | 4 | 103.7916 | 0.037183308 | Credible Set | purple |
| chr4:103791682 | 4 | 103.7917 | 0.065847398 | Credible Set | purple |
| chr4:103791711 | 4 | 103.7917 | 0.068015582 | Credible Set | purple |
| chr4:103791712 | 4 | 103.7917 | 0.070110846 | Credible Set | purple |
| chr4:103791722 | 4 | 103.7917 | 0.068550001 | Credible Set | purple |
| chr4:103793565 | 4 | 103.7936 | 0.024540983 | Credible Set | purple |
| chr4:103793571 | 4 | 103.7936 | 0.024540983 | Credible Set | purple |
| chr4:103793605 | 4 | 103.7936 | 0.024616382 | Credible Set | purple |
| chr4:103793645 | 4 | 103.7936 | 0.023839493 | Credible Set | purple |
| chr4:103793861 | 4 | 103.7939 | 0.027104748 | Credible Set | purple |
| chr4:103793969 | 4 | 103.7940 | 0.049967021 | Credible Set | purple |
| chr4:103795720 | 4 | 103.7957 | 0.038070727 | Credible Set | purple |
| chr4:103795724 | 4 | 103.7957 | 0.038649946 | Credible Set | purple |
| chr4:103795744 | 4 | 103.7957 | 0.041067485 | Credible Set | purple |
| chr4:103871016 | 4 | 103.8710 | 0.022372760 | Credible Set | purple |
| chr4:103871045 | 4 | 103.8710 | 0.020746898 | Credible Set | purple |
| chr4:103871082 | 4 | 103.8711 | 0.019764792 | Credible Set | purple |
| chr4:103938428 | 4 | 103.9384 | 0.011676744 | Credible Set | purple |
| chr4:103938894 | 4 | 103.9389 | 0.010866450 | Credible Set | purple |
| chr4:103966010 | 4 | 103.9660 | 0.009113775 | Credible Set | purple |
| chr4:103966020 | 4 | 103.9660 | 0.008833631 | Credible Set | purple |

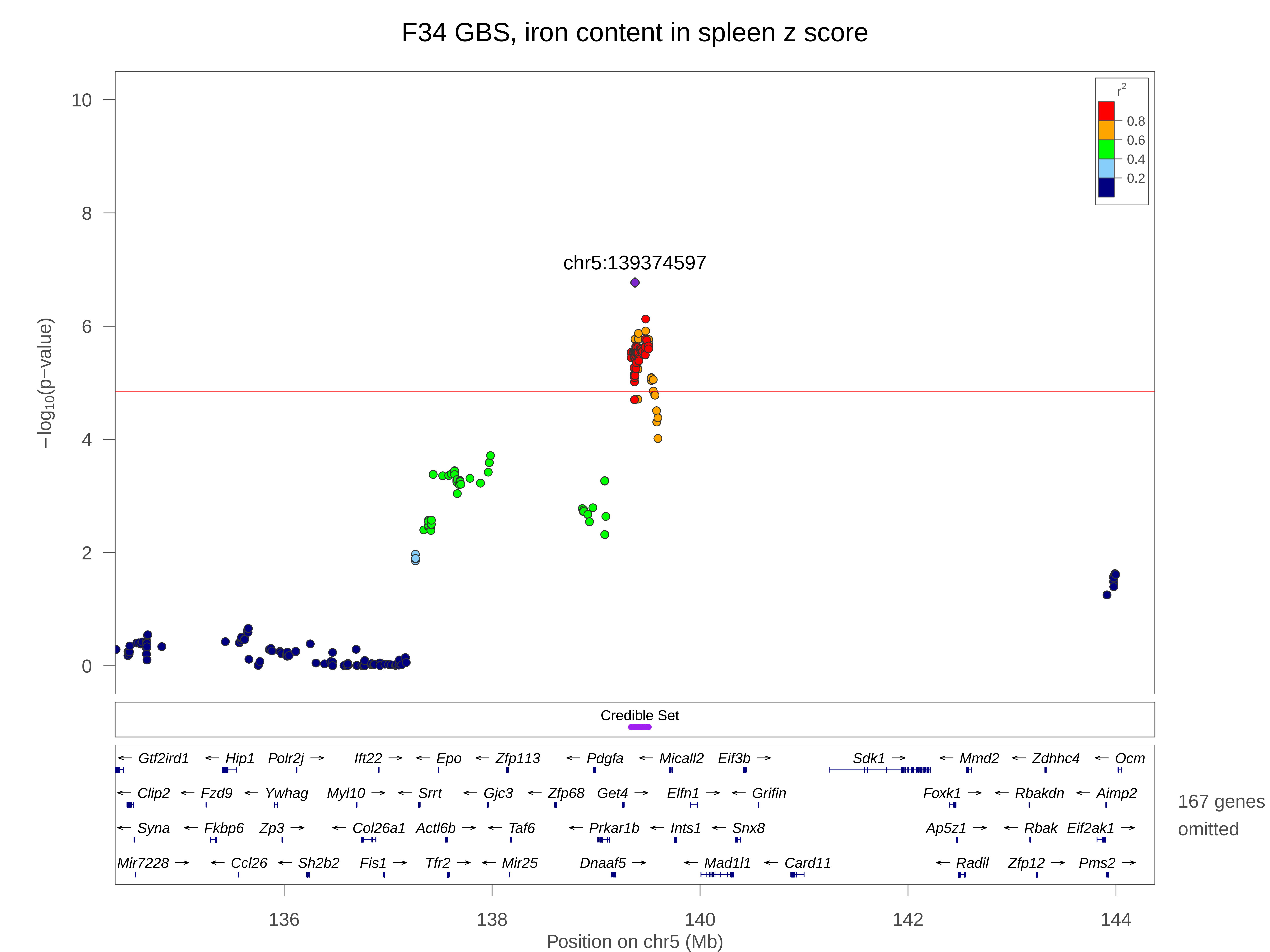

F34 GBS, iron content in spleen z score
chr5:139374597
−log10(p−value)
r2
0.8
0.6
0.4
0.2
Credible Set
← Gtf2ird1 ← Hip1 Polr2j → Ift22 → ← Epo ← Zfp113 ← Pdgfa ← Micall2 Eif3b → Sdk1 → ← Mmd2 ← Zdhhc4 ← Ocm
← Clip2 ← Fzd9 ← Ywhag Myl10 → ← Srrt ← Gjc3 ← Zfp68 Get4 → Elfn1 → ← Grifin Foxk1 → ← Rbakdn ← Aimp2
← Syna ← Fkbp6 Zp3 → ← Col26a1 Actl6b → ← Taf6 ← Prkar1b ← Ints1 ← Snx8 Ap5z1 → ← Rbak Eif2ak1 →
Mir7228 → ← Ccl26 ← Sh2b2 Fis1 → Tfr2 → ← Mir25 Dnaaf5 → ← Mad1l1 ← Card11 ← Radil Zfp12 → Pms2 →
167 genes omitted
Position on chr5 (Mb)

date: Thu Oct 25 17:18:18 2018
build: F34GBS
display range: chr5:134374597−144374597 [134374597−144374597]
hilite range: 0 − 0 [ 0 − 0 ]
reference SNP: chr5:139374597
number of SNPs plotted: 310
min P−value: 1.7E−7 [chr5:139374597]
max P−value: 10E−1 [chr5:136774656]
omitted Genes: Rfc2, Lat2, Eif4h
omitted Genes: Limk1, Eln, Wbscr28
omitted Genes: Cldn13, Wbscr27, Cldn4
omitted Genes: Cldn3, Wbscr25, Abhd11
omitted Genes: Abhd11os, Stx1a, Wbscr22
omitted Genes: Dnajc30, Vps37d, Mlxipl
omitted Genes: Tbl2, Bcl7b, Baz1b
omitted Genes: Trim50, Nsun5, Pom121
omitted Genes: Mir7033, Ccl24, Rhbdd2
omitted Genes: Por, Tmem120a, Mir7034
omitted Genes: Styxl1, Mdh2, Srrm3
omitted Genes: Hspb1, Ssc4d, Dtx2
omitted Genes: Upk3b, Upk3bl, Rasa4
omitted Genes: Mir7035, Lrwd1, Alkbh4
omitted Genes: Orai2, Prkrip1, Cux1
omitted Genes: Mir721, 4933404O12Rik, Cldn15
omitted Genes: Znhit1, Plod3, Mir702
omitted Genes: Vgf, Ap1s1, Serpine1
omitted Genes: Trim56, Ache, Mir8116
omitted Genes: Ufsp1, Mir7036, Trip6
omitted Genes: Slc12a9, Ephb4, Zan
omitted Genes: Pop7, Gigyf1, Gnb2
omitted Genes: Mir6418, Mospd3, Pcolce
omitted Genes: Fbxo24, Lrch4, Gm20605
omitted Genes: Sap25, Irs3, Agfg2
omitted Genes: Nyap1, Tsc22d4, Ppp1r35
omitted Genes: Mepce, Zcwpw1, Pilra
omitted Genes: Pilrb1, Pilrb2, Cyp3a13
omitted Genes: Azgp1, Smok3a, Smok3b
omitted Genes: Smok3c, Zkscan1, Zscan21
omitted Genes: Cops6, Mcm7, Mir93
Make more plots at http://csg.sph.umich.edu/locuszoom/
omitted Genes: Mir106b, Ap4m1, Cnpy4

Fine−mapping Regions

| snp | chr | pos (Mb) | pp | group | color |
|---|---|---|---|---|---|
| chr5:139337717 | 5 | 139.3377 | 0.011843849 | Credible Set | purple |
| chr5:139338124 | 5 | 139.3381 | 0.009604424 | Credible Set | purple |
| chr5:139356059 | 5 | 139.3561 | 0.011678110 | Credible Set | purple |
| chr5:139357023 | 5 | 139.3570 | 0.011676185 | Credible Set | purple |
| chr5:139357028 | 5 | 139.3570 | 0.011676185 | Credible Set | purple |
| chr5:139359983 | 5 | 139.3600 | 0.011213880 | Credible Set | purple |
| chr5:139360537 | 5 | 139.3605 | 0.011337767 | Credible Set | purple |
| chr5:139365262 | 5 | 139.3653 | 0.009836049 | Credible Set | purple |
| chr5:139365292 | 5 | 139.3653 | 0.011818085 | Credible Set | purple |
| chr5:139365343 | 5 | 139.3653 | 0.010006669 | Credible Set | purple |
| chr5:139365434 | 5 | 139.3654 | 0.006449839 | Credible Set | purple |
| chr5:139365448 | 5 | 139.3654 | 0.006455095 | Credible Set | purple |
| chr5:139365601 | 5 | 139.3656 | 0.010551312 | Credible Set | purple |
| chr5:139365659 | 5 | 139.3657 | 0.011511756 | Credible Set | purple |
| chr5:139365700 | 5 | 139.3657 | 0.004588735 | Credible Set | purple |
| chr5:139370400 | 5 | 139.3704 | 0.005162146 | Credible Set | purple |
| chr5:139370538 | 5 | 139.3705 | 0.004410022 | Credible Set | purple |
| chr5:139370586 | 5 | 139.3706 | 0.004888566 | Credible Set | purple |
| chr5:139370648 | 5 | 139.3706 | 0.004803120 | Credible Set | purple |
| chr5:139374544 | 5 | 139.3745 | 0.004822292 | Credible Set | purple |
| chr5:139374549 | 5 | 139.3745 | 0.004789127 | Credible Set | purple |
| chr5:139374597 | 5 | 139.3746 | 0.182402395 | Credible Set | purple |
| chr5:139374918 | 5 | 139.3749 | 0.011643184 | Credible Set | purple |
| chr5:139382014 | 5 | 139.3820 | 0.010570673 | Credible Set | purple |
| chr5:139382197 | 5 | 139.3822 | 0.014689264 | Credible Set | purple |
| chr5:139382215 | 5 | 139.3822 | 0.014689264 | Credible Set | purple |
| chr5:139382343 | 5 | 139.3823 | 0.014915282 | Credible Set | purple |
| chr5:139382347 | 5 | 139.3823 | 0.014915282 | Credible Set | purple |
| chr5:139382775 | 5 | 139.3828 | 0.011690070 | Credible Set | purple |
| chr5:139382789 | 5 | 139.3828 | 0.011690070 | Credible Set | purple |
| chr5:139384477 | 5 | 139.3845 | 0.006220628 | Credible Set | purple |
| chr5:139387948 | 5 | 139.3879 | 0.007921164 | Credible Set | purple |
| chr5:139388094 | 5 | 139.3881 | 0.012455759 | Credible Set | purple |
| chr5:139388781 | 5 | 139.3888 | 0.011743918 | Credible Set | purple |
| chr5:139389536 | 5 | 139.3895 | 0.014359155 | Credible Set | purple |
| chr5:139389540 | 5 | 139.3895 | 0.014359155 | Credible Set | purple |

Fine−mapping Regions

| snp | chr | pos (Mb) | pp | group | color |
|---|---|---|---|---|---|
| chr5:139389572 | 5 | 139.3896 | 0.014359155 | Credible Set | purple |
| chr5:139389578 | 5 | 139.3896 | 0.014359155 | Credible Set | purple |
| chr5:139391234 | 5 | 139.3912 | 0.011643184 | Credible Set | purple |
| chr5:139391358 | 5 | 139.3914 | 0.011643184 | Credible Set | purple |
| chr5:139395030 | 5 | 139.3950 | 0.011664814 | Credible Set | purple |
| chr5:139402752 | 5 | 139.4028 | 0.014563760 | Credible Set | purple |
| chr5:139403170 | 5 | 139.4032 | 0.011682145 | Credible Set | purple |
| chr5:139404712 | 5 | 139.4047 | 0.011643184 | Credible Set | purple |
| chr5:139404757 | 5 | 139.4048 | 0.011643184 | Credible Set | purple |
| chr5:139404767 | 5 | 139.4048 | 0.011643184 | Credible Set | purple |
| chr5:139411750 | 5 | 139.4118 | 0.009698102 | Credible Set | purple |
| chr5:139411803 | 5 | 139.4118 | 0.008992399 | Credible Set | purple |
| chr5:139411845 | 5 | 139.4118 | 0.008992399 | Credible Set | purple |
| chr5:139411920 | 5 | 139.4119 | 0.009042781 | Credible Set | purple |
| chr5:139412108 | 5 | 139.4121 | 0.008464481 | Credible Set | purple |
| chr5:139412109 | 5 | 139.4121 | 0.008464481 | Credible Set | purple |
| chr5:139422570 | 5 | 139.4226 | 0.013812241 | Credible Set | purple |
| chr5:139426676 | 5 | 139.4267 | 0.013119564 | Credible Set | purple |
| chr5:139437340 | 5 | 139.4373 | 0.013299241 | Credible Set | purple |
| chr5:139445073 | 5 | 139.4451 | 0.012367835 | Credible Set | purple |
| chr5:139445415 | 5 | 139.4454 | 0.011544452 | Credible Set | purple |
| chr5:139449959 | 5 | 139.4500 | 0.012377106 | Credible Set | purple |
| chr5:139472776 | 5 | 139.4728 | 0.012039033 | Credible Set | purple |
| chr5:139472992 | 5 | 139.4730 | 0.010674549 | Credible Set | purple |
| chr5:139472993 | 5 | 139.4730 | 0.010696332 | Credible Set | purple |
| chr5:139474788 | 5 | 139.4748 | 0.015301490 | Credible Set | purple |
| chr5:139474790 | 5 | 139.4748 | 0.014598204 | Credible Set | purple |
| chr5:139474824 | 5 | 139.4748 | 0.014545417 | Credible Set | purple |
| chr5:139474889 | 5 | 139.4749 | 0.019783865 | Credible Set | purple |
| chr5:139477643 | 5 | 139.4776 | 0.043516286 | Credible Set | purple |
| chr5:139488214 | 5 | 139.4882 | 0.019088989 | Credible Set | purple |
| chr5:139504674 | 5 | 139.5047 | 0.015149475 | Credible Set | purple |
| chr5:139505164 | 5 | 139.5052 | 0.015175055 | Credible Set | purple |
| chr5:139505384 | 5 | 139.5054 | 0.013522652 | Credible Set | purple |

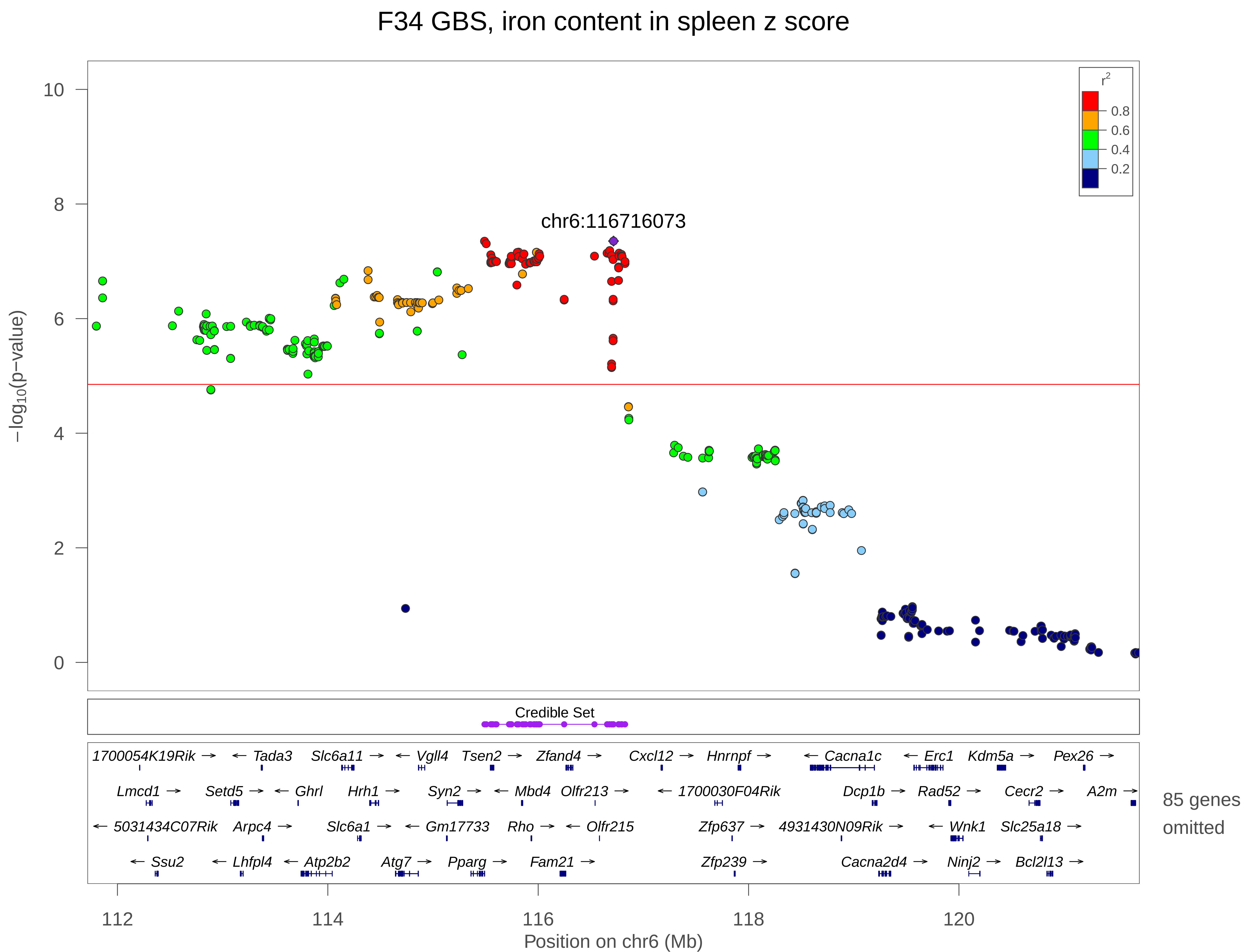

F34 GBS, iron content in spleen z score
chr6:116716073
−log10(p−value)
r2
0.8
0.6
0.4
0.2
Credible Set
1700054K19Rik → ← Tada3 Slc6a11 → ← Vgll4 Tsen2 → Zfand4 → Cxcl12 → Hnrnpf → ← Cacna1c ← Erc1 Kdm5a → Pex26 →
Lmcd1 → Setd5 → ← Ghrl Hrh1 → Syn2 → ← Mbd4 Olfr213 → ← 1700030F04Rik Dcp1b → Rad52 → Cecr2 → A2m →
← 5031434C07Rik Arpc4 → Slc6a1 → ← Gm17733 Rho → ← Olfr215 Zfp637 → 4931430N09Rik → ← Wnk1 Slc25a18 →
← Ssu2 ← Lhfpl4 ← Atp2b2 Atg7 → Pparg → Fam21 → Zfp239 → Cacna2d4 → Ninj2 → Bcl2l13 →
85 genes omitted
Position on chr6 (Mb)

| | |
|---|---|
| date: | Thu Oct 25 17:18:29 2018 |
| build: | F34GBS |
| display range: | chr6:111716073−121716073 [111716073−121716073] |
| hilite range: | 0 − 0 [ 0 − 0 ] |
| reference SNP: | chr6:116716073 |
| number of SNPs plotted: | 445 |
| min P−value: | 4.43E−8 [chr6:116716073] |
| max P−value: | 7.14E−1 [chr6:121678889] |
| omitted Genes: | Cav3, Oxtr, Rad18 |
| omitted Genes: | Srgap3, Thumpd3, Gt(ROSA)26Sor |
| omitted Genes: | Mtmr14, Cpne9, Brpf1 |
| omitted Genes: | Ogg1, Camk1, Ttll3 |
| omitted Genes: | Rpusd3, Cidec, Jagn1 |
| omitted Genes: | Il17re, Il17rc, Creld1 |
| omitted Genes: | Prrt3, Emc3, Fancd2 |
| omitted Genes: | Fancd2os, Brk1, Vhl |
| omitted Genes: | Irak2, Tatdn2, Mir7042 |
| omitted Genes: | Sec13, Tamm41, Timp4 |
| omitted Genes: | Mkrn2os, Mkrn2, Raf1 |
| omitted Genes: | Tmem40, Cand2, Mir7660 |
| omitted Genes: | Rpl32, Snora7a, Efcab12 |
| omitted Genes: | Ift122, H1foo, Plxnd1 |
| omitted Genes: | Tmcc1, 9530062K07Rik, March8 |
| omitted Genes: | Alox5, Olfr211, Olfr212 |
| omitted Genes: | Olfr214, Zfp422, Rassf4 |
| omitted Genes: | Mir7043, 8430408G22Rik, Tmem72 |
| omitted Genes: | Fxyd4, Rasgef1a, Mir7044 |
| omitted Genes: | Csgalnact2, Ret, 1700069P05Rik |
| omitted Genes: | Gm30498, Bms1, Zfp248 |
| omitted Genes: | Zfp9, Ankrd26, Lrtm2 |
| omitted Genes: | Adipor2, Wnt5b, Fbxl14 |
| omitted Genes: | 3110021A11Rik, Mir706, B4galnt3 |
| omitted Genes: | Ccdc77, Il17ra, Cecr6 |
| omitted Genes: | Cecr5, 1700072O05Rik, Atp6v1e1 |
| omitted Genes: | Bid, Mical3, Tuba8 |
| omitted Genes: | Usp18, Slc6a13, Slc6a12 |
| omitted Genes: | Iqsec3 |

Make more plots at http://csg.sph.umich.edu/locuszoom/

Fine−mapping Regions

| snp | chr | pos (Mb) | pp | group | color |
|---|---|---|---|---|---|
| chr6:115489819 | 6 | 115.4898 | 0.029911876 | Credible Set | purple |
| chr6:115505788 | 6 | 115.5058 | 0.027122258 | Credible Set | purple |
| chr6:115549490 | 6 | 115.5495 | 0.017619729 | Credible Set | purple |
| chr6:115549730 | 6 | 115.5497 | 0.012877132 | Credible Set | purple |
| chr6:115549829 | 6 | 115.5498 | 0.013403440 | Credible Set | purple |
| chr6:115555634 | 6 | 115.5556 | 0.013556316 | Credible Set | purple |
| chr6:115559836 | 6 | 115.5598 | 0.015448422 | Credible Set | purple |
| chr6:115567453 | 6 | 115.5675 | 0.013576450 | Credible Set | purple |
| chr6:115569261 | 6 | 115.5693 | 0.013183497 | Credible Set | purple |
| chr6:115593481 | 6 | 115.5935 | 0.013760669 | Credible Set | purple |
| chr6:115603445 | 6 | 115.6034 | 0.013530179 | Credible Set | purple |
| chr6:115723557 | 6 | 115.7236 | 0.012711679 | Credible Set | purple |
| chr6:115723602 | 6 | 115.7236 | 0.012434817 | Credible Set | purple |
| chr6:115729470 | 6 | 115.7295 | 0.013075719 | Credible Set | purple |
| chr6:115729519 | 6 | 115.7295 | 0.013184296 | Credible Set | purple |
| chr6:115730587 | 6 | 115.7306 | 0.013741989 | Credible Set | purple |
| chr6:115736586 | 6 | 115.7366 | 0.014052210 | Credible Set | purple |
| chr6:115743744 | 6 | 115.7437 | 0.014894984 | Credible Set | purple |
| chr6:115744308 | 6 | 115.7443 | 0.012471193 | Credible Set | purple |
| chr6:115744336 | 6 | 115.7443 | 0.012511485 | Credible Set | purple |
| chr6:115744487 | 6 | 115.7445 | 0.016616915 | Credible Set | purple |
| chr6:115744493 | 6 | 115.7445 | 0.016616915 | Credible Set | purple |
| chr6:115797352 | 6 | 115.7974 | 0.005438448 | Credible Set | purple |
| chr6:115801564 | 6 | 115.8016 | 0.019346817 | Credible Set | purple |
| chr6:115813904 | 6 | 115.8139 | 0.016444940 | Credible Set | purple |
| chr6:115850586 | 6 | 115.8506 | 0.015064290 | Credible Set | purple |
| chr6:115864297 | 6 | 115.8643 | 0.018189298 | Credible Set | purple |
| chr6:115880648 | 6 | 115.8806 | 0.012217102 | Credible Set | purple |
| chr6:115919848 | 6 | 115.9198 | 0.012772538 | Credible Set | purple |
| chr6:115920287 | 6 | 115.9203 | 0.013075212 | Credible Set | purple |
| chr6:115928586 | 6 | 115.9286 | 0.013054236 | Credible Set | purple |
| chr6:115929540 | 6 | 115.9295 | 0.012981674 | Credible Set | purple |
| chr6:115957904 | 6 | 115.9579 | 0.013880932 | Credible Set | purple |
| chr6:115967362 | 6 | 115.9674 | 0.013395694 | Credible Set | purple |
| chr6:115967880 | 6 | 115.9679 | 0.013446363 | Credible Set | purple |
| chr6:115985523 | 6 | 115.9855 | 0.013433663 | Credible Set | purple |

Fine−mapping Regions

| snp | chr | pos (Mb) | pp | group | color |
|---|---|---|---|---|---|
| chr6:115985527 | 6 | 115.9855 | 0.013433663 | Credible Set | purple |
| chr6:115985543 | 6 | 115.9855 | 0.013433663 | Credible Set | purple |
| chr6:115985717 | 6 | 115.9857 | 0.014755059 | Credible Set | purple |
| chr6:116003809 | 6 | 116.0038 | 0.015313769 | Credible Set | purple |
| chr6:116004432 | 6 | 116.0044 | 0.015529522 | Credible Set | purple |
| chr6:116009041 | 6 | 116.0090 | 0.018413782 | Credible Set | purple |
| chr6:116009081 | 6 | 116.0091 | 0.018293016 | Credible Set | purple |
| chr6:116014484 | 6 | 116.0145 | 0.016682609 | Credible Set | purple |
| chr6:116246991 | 6 | 116.2470 | 0.003127944 | Credible Set | purple |
| chr6:116535049 | 6 | 116.5350 | 0.016733290 | Credible Set | purple |
| chr6:116655672 | 6 | 116.6557 | 0.018804842 | Credible Set | purple |
| chr6:116655715 | 6 | 116.6557 | 0.018804842 | Credible Set | purple |
| chr6:116675286 | 6 | 116.6753 | 0.018754154 | Credible Set | purple |
| chr6:116679618 | 6 | 116.6796 | 0.020765511 | Credible Set | purple |
| chr6:116697627 | 6 | 116.6976 | 0.006264746 | Credible Set | purple |
| chr6:116698225 | 6 | 116.6982 | 0.017080051 | Credible Set | purple |
| chr6:116712020 | 6 | 116.7120 | 0.014661507 | Credible Set | purple |
| chr6:116712593 | 6 | 116.7126 | 0.003130955 | Credible Set | purple |
| chr6:116716073 | 6 | 116.7161 | 0.030067886 | Credible Set | purple |
| chr6:116763618 | 6 | 116.7636 | 0.010980677 | Credible Set | purple |
| chr6:116763629 | 6 | 116.7636 | 0.006501999 | Credible Set | purple |
| chr6:116764660 | 6 | 116.7647 | 0.010655808 | Credible Set | purple |
| chr6:116764667 | 6 | 116.7647 | 0.010565951 | Credible Set | purple |
| chr6:116769198 | 6 | 116.7692 | 0.018775783 | Credible Set | purple |
| chr6:116769221 | 6 | 116.7692 | 0.016754096 | Credible Set | purple |
| chr6:116791717 | 6 | 116.7917 | 0.017909708 | Credible Set | purple |
| chr6:116791844 | 6 | 116.7918 | 0.016713787 | Credible Set | purple |
| chr6:116791847 | 6 | 116.7918 | 0.016713787 | Credible Set | purple |
| chr6:116797564 | 6 | 116.7976 | 0.016373768 | Credible Set | purple |
| chr6:116825061 | 6 | 116.8251 | 0.012552009 | Credible Set | purple |
| chr6:116825105 | 6 | 116.8251 | 0.013529920 | Credible Set | purple |

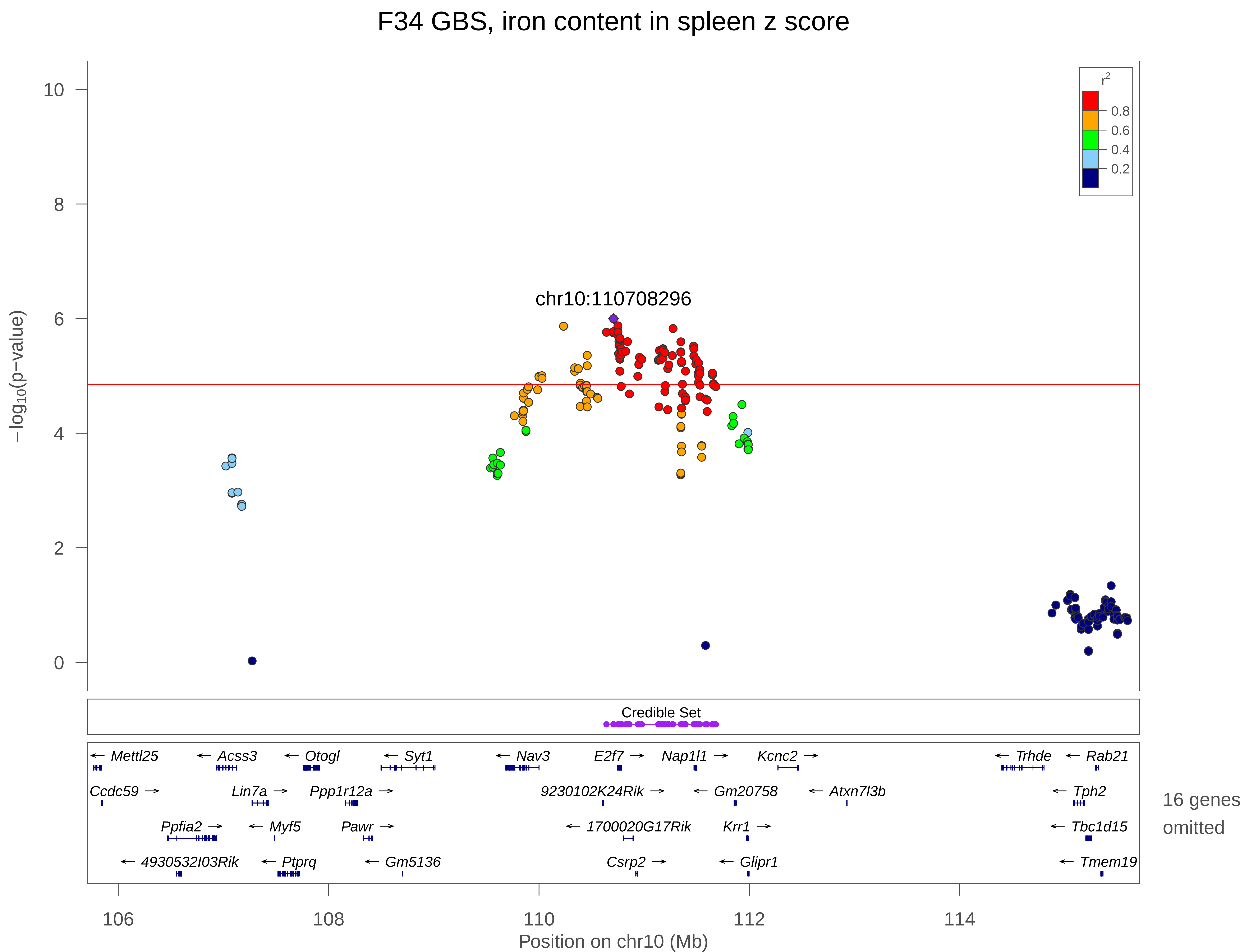

F34 GBS, iron content in spleen z score
chr10:110708296
−log10(p−value)
r2
0.8
0.6
0.4
0.2
Credible Set
Mettl25
Acss3
Otogl
Syt1
Nav3
E2f7
Nap1l1
Kcnc2
Trhde
Rab21
Ccdc59
Lin7a
Ppp1r12a
9230102K24Rik
Gm20758
Atxn7l3b
Tph2
Ppfia2
Myf5
Pawr
1700020G17Rik
Krr1
Tbc1d15
4930532I03Rik
Ptprq
Gm5136
Csrp2
Glipr1
Tmem19
16 genes omitted
Position on chr10 (Mb)

| | |
|---|---|
| date: | Thu Oct 25 17:17:39 2018 |
| build: | F34GBS |
| display range: | chr10:105708296−115708296 [105708296−115708296] |
| hilite range: | 0 − 0 [ 0 − 0 ] |
| reference SNP: | chr10:110708296 |
| number of SNPs plotted: | 326 |
| min P−value: | 1E−6 [chr10:110708296] |
| max P−value: | 9.41E−1 [chr10:107272725] |
| omitted Genes: | Myf6, Gm29685, Zdhhc17 |
| omitted Genes: | Osbpl8, Bbs10, Gm5176 |
| omitted Genes: | Phlda1, 4933440J02Rik, Glipr1l1 |
| omitted Genes: | Glipr1l2, Caps2, 1700010J16Rik |
| omitted Genes: | Thap2, Zfc3h1, Lgr5 |
| omitted Genes: | A930009A15Rik |

Make more plots at http://csg.sph.umich.edu/locuszoom/

Fine−mapping Regions

| snp | chr | pos (Mb) | pp | group | color |
|---|---|---|---|---|---|
| chr10:110641142 | 10 | 110.6411 | 0.018986892 | Credible Set | purple |
| chr10:110708014 | 10 | 110.7080 | 0.018986892 | Credible Set | purple |
| chr10:110708268 | 10 | 110.7083 | 0.018986892 | Credible Set | purple |
| chr10:110708269 | 10 | 110.7083 | 0.018986892 | Credible Set | purple |
| chr10:110708296 | 10 | 110.7083 | 0.036976812 | Credible Set | purple |
| chr10:110708628 | 10 | 110.7086 | 0.018986892 | Credible Set | purple |
| chr10:110749349 | 10 | 110.7493 | 0.018986892 | Credible Set | purple |
| chr10:110749883 | 10 | 110.7499 | 0.018986892 | Credible Set | purple |
| chr10:110749995 | 10 | 110.7500 | 0.018986892 | Credible Set | purple |
| chr10:110750029 | 10 | 110.7500 | 0.036976812 | Credible Set | purple |
| chr10:110750810 | 10 | 110.7508 | 0.018986892 | Credible Set | purple |
| chr10:110756900 | 10 | 110.7569 | 0.009763502 | Credible Set | purple |
| chr10:110757011 | 10 | 110.7570 | 0.009763502 | Credible Set | purple |
| chr10:110759826 | 10 | 110.7598 | 0.012864948 | Credible Set | purple |
| chr10:110760381 | 10 | 110.7604 | 0.012864948 | Credible Set | purple |
| chr10:110768547 | 10 | 110.7685 | 0.012864948 | Credible Set | purple |
| chr10:110768747 | 10 | 110.7687 | 0.005028699 | Credible Set | purple |
| chr10:110768762 | 10 | 110.7688 | 0.005028699 | Credible Set | purple |
| chr10:110768830 | 10 | 110.7688 | 0.018986892 | Credible Set | purple |
| chr10:110768858 | 10 | 110.7689 | 0.018986892 | Credible Set | purple |
| chr10:110768862 | 10 | 110.7689 | 0.018986892 | Credible Set | purple |
| chr10:110768863 | 10 | 110.7689 | 0.018986892 | Credible Set | purple |
| chr10:110768865 | 10 | 110.7689 | 0.018986892 | Credible Set | purple |
| chr10:110769342 | 10 | 110.7693 | 0.007884281 | Credible Set | purple |
| chr10:110769400 | 10 | 110.7694 | 0.007884281 | Credible Set | purple |
| chr10:110769437 | 10 | 110.7694 | 0.007884281 | Credible Set | purple |
| chr10:110769467 | 10 | 110.7695 | 0.007884281 | Credible Set | purple |
| chr10:110769523 | 10 | 110.7695 | 0.007884281 | Credible Set | purple |
| chr10:110769585 | 10 | 110.7696 | 0.007884281 | Credible Set | purple |
| chr10:110778078 | 10 | 110.7781 | 0.012864948 | Credible Set | purple |
| chr10:110778106 | 10 | 110.7781 | 0.012864948 | Credible Set | purple |
| chr10:110778366 | 10 | 110.7784 | 0.009763502 | Credible Set | purple |
| chr10:110780233 | 10 | 110.7802 | 0.009763502 | Credible Set | purple |
| chr10:110781079 | 10 | 110.7811 | 0.002759361 | Credible Set | purple |
| chr10:110781085 | 10 | 110.7811 | 0.002759361 | Credible Set | purple |
| chr10:110781088 | 10 | 110.7811 | 0.002759361 | Credible Set | purple |

Fine−mapping Regions

| snp | chr | pos (Mb) | pp | group | color |
|---|---|---|---|---|---|
| chr10:110782063 | 10 | 110.7821 | 0.012864948 | Credible Set | purple |
| chr10:110790601 | 10 | 110.7906 | 0.009763502 | Credible Set | purple |
| chr10:110824052 | 10 | 110.8241 | 0.009763502 | Credible Set | purple |
| chr10:110841372 | 10 | 110.8414 | 0.012864948 | Credible Set | purple |
| chr10:110859622 | 10 | 110.8596 | 0.002002545 | Credible Set | purple |
| chr10:110939251 | 10 | 110.9393 | 0.004063067 | Credible Set | purple |
| chr10:110949219 | 10 | 110.9492 | 0.006621538 | Credible Set | purple |
| chr10:110949220 | 10 | 110.9492 | 0.006621538 | Credible Set | purple |
| chr10:110949581 | 10 | 110.9496 | 0.006621538 | Credible Set | purple |
| chr10:110955372 | 10 | 110.9554 | 0.007884281 | Credible Set | purple |
| chr10:110977680 | 10 | 110.9777 | 0.007884281 | Credible Set | purple |
| chr10:111134114 | 10 | 111.1341 | 0.007884281 | Credible Set | purple |
| chr10:111139942 | 10 | 111.1399 | 0.007884281 | Credible Set | purple |
| chr10:111139969 | 10 | 111.1400 | 0.007884281 | Credible Set | purple |
| chr10:111145119 | 10 | 111.1451 | 0.009763502 | Credible Set | purple |
| chr10:111156216 | 10 | 111.1562 | 0.007884281 | Credible Set | purple |
| chr10:111162654 | 10 | 111.1627 | 0.009763502 | Credible Set | purple |
| chr10:111178371 | 10 | 111.1784 | 0.012864948 | Credible Set | purple |
| chr10:111178404 | 10 | 111.1784 | 0.012864948 | Credible Set | purple |
| chr10:111179231 | 10 | 111.1792 | 0.009763502 | Credible Set | purple |
| chr10:111179272 | 10 | 111.1793 | 0.007884281 | Credible Set | purple |
| chr10:111180485 | 10 | 111.1805 | 0.009763502 | Credible Set | purple |
| chr10:111180489 | 10 | 111.1805 | 0.009763502 | Credible Set | purple |
| chr10:111196665 | 10 | 111.1967 | 0.009763502 | Credible Set | purple |
| chr10:111196850 | 10 | 111.1968 | 0.002202789 | Credible Set | purple |
| chr10:111196872 | 10 | 111.1969 | 0.002202789 | Credible Set | purple |
| chr10:111201273 | 10 | 111.2013 | 0.002759361 | Credible Set | purple |
| chr10:111223748 | 10 | 111.2237 | 0.005713580 | Credible Set | purple |
| chr10:111234804 | 10 | 111.2348 | 0.006621538 | Credible Set | purple |
| chr10:111267516 | 10 | 111.2675 | 0.009763502 | Credible Set | purple |
| chr10:111274460 | 10 | 111.2745 | 0.036976812 | Credible Set | purple |
| chr10:111274480 | 10 | 111.2745 | 0.036976812 | Credible Set | purple |
| chr10:111347458 | 10 | 111.3475 | 0.009763502 | Credible Set | purple |
| chr10:111348661 | 10 | 111.3487 | 0.009763502 | Credible Set | purple |
| chr10:111348802 | 10 | 111.3488 | 0.012864948 | Credible Set | purple |
| chr10:111354708 | 10 | 111.3547 | 0.006621538 | Credible Set | purple |

Fine−mapping Regions

| snp | chr | pos (Mb) | pp | group | color |
|---|---|---|---|---|---|
| chr10:111363662 | 10 | 111.3637 | 0.002947014 | Credible Set | purple |
| chr10:111363665 | 10 | 111.3637 | 0.002097769 | Credible Set | purple |
| chr10:111391890 | 10 | 111.3919 | 0.001696378 | Credible Set | purple |
| chr10:111391901 | 10 | 111.3919 | 0.001576646 | Credible Set | purple |
| chr10:111391954 | 10 | 111.3920 | 0.001763555 | Credible Set | purple |
| chr10:111391955 | 10 | 111.3920 | 0.001634260 | Credible Set | purple |
| chr10:111392557 | 10 | 111.3926 | 0.005028699 | Credible Set | purple |
| chr10:111470144 | 10 | 111.4701 | 0.012864948 | Credible Set | purple |
| chr10:111471816 | 10 | 111.4718 | 0.012864948 | Credible Set | purple |
| chr10:111471933 | 10 | 111.4719 | 0.009763502 | Credible Set | purple |
| chr10:111490761 | 10 | 111.4908 | 0.006621538 | Credible Set | purple |
| chr10:111494112 | 10 | 111.4941 | 0.007884281 | Credible Set | purple |
| chr10:111512151 | 10 | 111.5122 | 0.004493315 | Credible Set | purple |
| chr10:111515685 | 10 | 111.5157 | 0.003162889 | Credible Set | purple |
| chr10:111516056 | 10 | 111.5161 | 0.004063067 | Credible Set | purple |
| chr10:111516061 | 10 | 111.5161 | 0.006621538 | Credible Set | purple |
| chr10:111522846 | 10 | 111.5228 | 0.005028699 | Credible Set | purple |
| chr10:111529740 | 10 | 111.5297 | 0.005028699 | Credible Set | purple |
| chr10:111529759 | 10 | 111.5298 | 0.004493315 | Credible Set | purple |
| chr10:111530393 | 10 | 111.5304 | 0.002759361 | Credible Set | purple |
| chr10:111530397 | 10 | 111.5304 | 0.001836439 | Credible Set | purple |
| chr10:111583278 | 10 | 111.5833 | 0.001696378 | Credible Set | purple |
| chr10:111598785 | 10 | 111.5988 | 0.001634260 | Credible Set | purple |
| chr10:111648540 | 10 | 111.6485 | 0.004493315 | Credible Set | purple |
| chr10:111648550 | 10 | 111.6486 | 0.004063067 | Credible Set | purple |
| chr10:111648631 | 10 | 111.6486 | 0.004493315 | Credible Set | purple |
| chr10:111658438 | 10 | 111.6584 | 0.002947014 | Credible Set | purple |
| chr10:111658509 | 10 | 111.6585 | 0.002947014 | Credible Set | purple |
| chr10:111681517 | 10 | 111.6815 | 0.002759361 | Credible Set | purple |

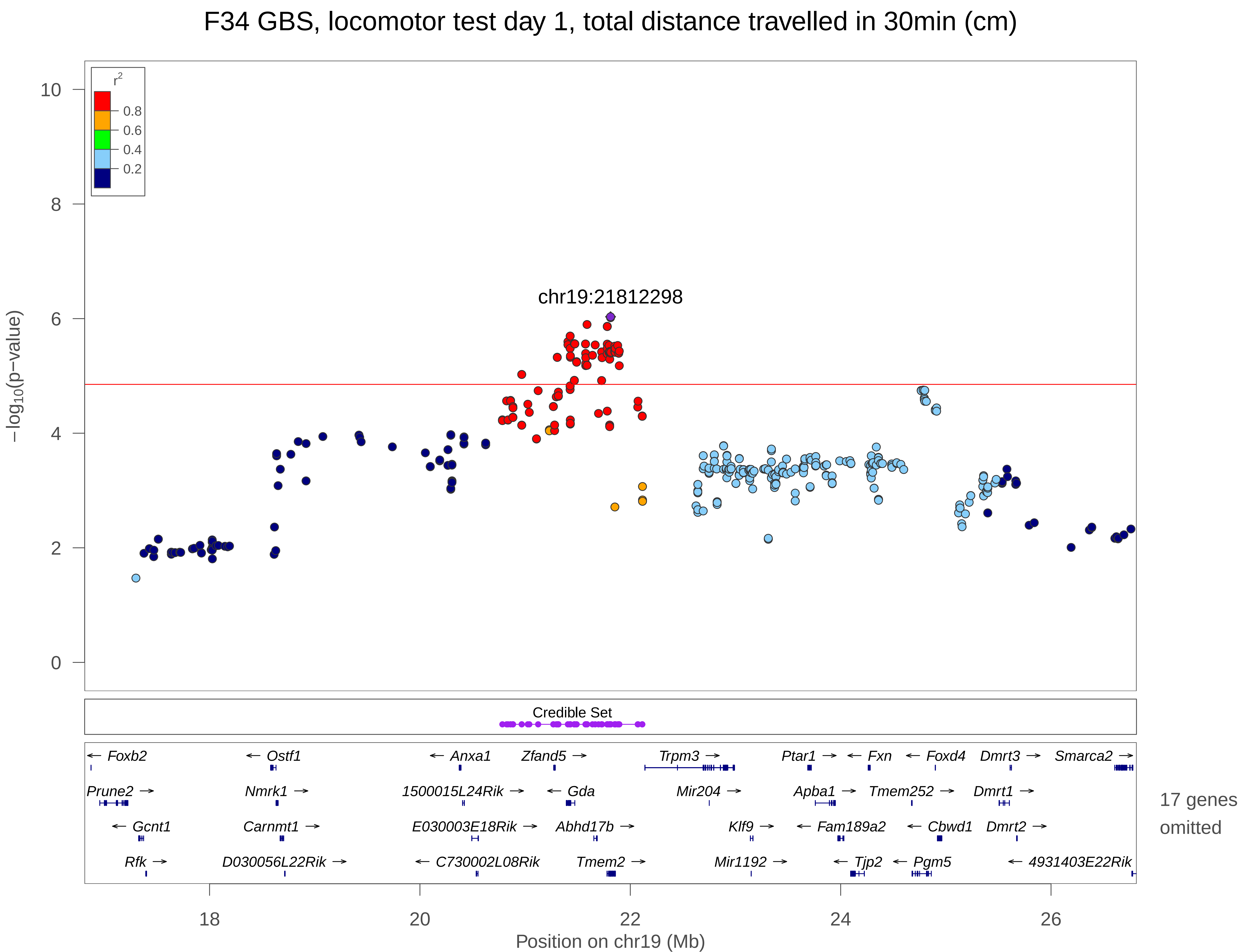

F34 GBS, locomotor test day 1, total distance travelled in 30min (cm)
r2
0.8
0.6
0.4
0.2
chr19:21812298
10
8
6
4
2
0
−log10(p−value)
Credible Set
← Foxb2
← Ostf1
← Anxa1
Zfand5 →
Trpm3 →
Ptar1 →
← Fxn
← Foxd4
Dmrt3 →
Smarca2 →
Prune2 →
Nmrk1 →
1500015L24Rik →
← Gda
Mir204 →
Apba1 →
Tmem252 →
Dmrt1 →
← Gcnt1
Carnmt1 →
E030003E18Rik →
Abhd17b →
Klf9 →
← Fam189a2
← Cbwd1
Dmrt2 →
Rfk →
D030056L22Rik →
← C730002L08Rik
Tmem2 →
Mir1192 →
← Tjp2
← Pgm5
← 4931403E22Rik
17 genes omitted
18
20
22
24
26
Position on chr19 (Mb)

| | |
|---|---|
| date: | Thu Oct 25 17:18:38 2018 |
| build: | F34GBS |
| display range: | chr19:16812298−26812298 [16812298−26812298] |
| hilite range: | 0 − 0 [ 0 − 0 ] |
| reference SNP: | chr19:21812298 |
| number of SNPs plotted: | 470 |
| min P−value: | 9.28E−7 [chr19:21812298] |
| max P−value: | 3.38E−2 [chr19:17299794] |
| omitted Genes: | Pcsk5, Trpm6, Rorb |
| omitted Genes: | Aldh1a1, Aldh1a7, Tmc1 |
| omitted Genes: | 4930554I06Rik, 1110059E24Rik, Smc5 |
| omitted Genes: | Mamdc2, 1700028P14Rik, Pip5k1b |
| omitted Genes: | Fam122a, Mir3084−1, Dock8 |
| omitted Genes: | Kank1, 2610016A17Rik |

Make more plots at http://csg.sph.umich.edu/locuszoom/

Fine−mapping Regions

| snp | chr | pos (Mb) | pp | group | color |
|---|---|---|---|---|---|
| chr19:20784050 | 19 | 20.78405 | 0.0008587211 | Credible Set | purple |
| chr19:20824062 | 19 | 20.82406 | 0.0017655107 | Credible Set | purple |
| chr19:20837342 | 19 | 20.83734 | 0.0008561549 | Credible Set | purple |
| chr19:20861811 | 19 | 20.86181 | 0.0018014019 | Credible Set | purple |
| chr19:20861821 | 19 | 20.86182 | 0.0017659141 | Credible Set | purple |
| chr19:20883962 | 19 | 20.88396 | 0.0014211894 | Credible Set | purple |
| chr19:20883975 | 19 | 20.88397 | 0.0009496508 | Credible Set | purple |
| chr19:20883979 | 19 | 20.88398 | 0.0009455779 | Credible Set | purple |
| chr19:20883980 | 19 | 20.88398 | 0.0009370136 | Credible Set | purple |
| chr19:20884530 | 19 | 20.88453 | 0.0013534603 | Credible Set | purple |
| chr19:20967742 | 19 | 20.96774 | 0.0048736412 | Credible Set | purple |
| chr19:20967772 | 19 | 20.96777 | 0.0048533035 | Credible Set | purple |
| chr19:21025668 | 19 | 21.02567 | 0.0015582333 | Credible Set | purple |
| chr19:21039744 | 19 | 21.03974 | 0.0011401914 | Credible Set | purple |
| chr19:21123769 | 19 | 21.12377 | 0.0026108852 | Credible Set | purple |
| chr19:21268525 | 19 | 21.26853 | 0.0014308116 | Credible Set | purple |
| chr19:21268527 | 19 | 21.26853 | 0.0014252921 | Credible Set | purple |
| chr19:21268550 | 19 | 21.26855 | 0.0014190158 | Credible Set | purple |
| chr19:21296098 | 19 | 21.29610 | 0.0020623460 | Credible Set | purple |
| chr19:21305518 | 19 | 21.30552 | 0.0094436402 | Credible Set | purple |
| chr19:21305568 | 19 | 21.30557 | 0.0093562265 | Credible Set | purple |
| chr19:21316234 | 19 | 21.31623 | 0.0024741063 | Credible Set | purple |
| chr19:21316268 | 19 | 21.31627 | 0.0021044591 | Credible Set | purple |
| chr19:21316305 | 19 | 21.31630 | 0.0021457242 | Credible Set | purple |
| chr19:21407108 | 19 | 21.40711 | 0.0171734453 | Credible Set | purple |
| chr19:21407182 | 19 | 21.40718 | 0.0152564050 | Credible Set | purple |
| chr19:21427800 | 19 | 21.42780 | 0.0030720723 | Credible Set | purple |
| chr19:21427805 | 19 | 21.42780 | 0.0027222168 | Credible Set | purple |
| chr19:21427851 | 19 | 21.42785 | 0.0031403069 | Credible Set | purple |
| chr19:21428151 | 19 | 21.42815 | 0.0132728482 | Credible Set | purple |
| chr19:21428338 | 19 | 21.42834 | 0.0213259592 | Credible Set | purple |
| chr19:21428343 | 19 | 21.42834 | 0.0213259592 | Credible Set | purple |
| chr19:21429788 | 19 | 21.42979 | 0.0008550118 | Credible Set | purple |
| chr19:21430361 | 19 | 21.43036 | 0.0094097845 | Credible Set | purple |
| chr19:21430412 | 19 | 21.43041 | 0.0099109711 | Credible Set | purple |
| chr19:21467882 | 19 | 21.46788 | 0.0159382825 | Credible Set | purple |

Fine−mapping Regions

| snp | chr | pos (Mb) | pp | group | color |
|---|---|---|---|---|---|
| chr19:21467967 | 19 | 21.46797 | 0.003887071 | Credible Set | purple |
| chr19:21467982 | 19 | 21.46798 | 0.003853277 | Credible Set | purple |
| chr19:21467985 | 19 | 21.46798 | 0.003853277 | Credible Set | purple |
| chr19:21467987 | 19 | 21.46799 | 0.003870160 | Credible Set | purple |
| chr19:21472509 | 19 | 21.47251 | 0.015775053 | Credible Set | purple |
| chr19:21472526 | 19 | 21.47253 | 0.015757697 | Credible Set | purple |
| chr19:21489550 | 19 | 21.48955 | 0.007955430 | Credible Set | purple |
| chr19:21489579 | 19 | 21.48958 | 0.007779354 | Credible Set | purple |
| chr19:21574198 | 19 | 21.57420 | 0.015685100 | Credible Set | purple |
| chr19:21574202 | 19 | 21.57420 | 0.015732871 | Credible Set | purple |
| chr19:21575782 | 19 | 21.57578 | 0.010901563 | Credible Set | purple |
| chr19:21576364 | 19 | 21.57636 | 0.006832973 | Credible Set | purple |
| chr19:21576365 | 19 | 21.57636 | 0.007386746 | Credible Set | purple |
| chr19:21576374 | 19 | 21.57637 | 0.009238239 | Credible Set | purple |
| chr19:21588462 | 19 | 21.58846 | 0.006909232 | Credible Set | purple |
| chr19:21588470 | 19 | 21.58847 | 0.006924589 | Credible Set | purple |
| chr19:21588527 | 19 | 21.58853 | 0.006942255 | Credible Set | purple |
| chr19:21588528 | 19 | 21.58853 | 0.006918987 | Credible Set | purple |
| chr19:21588608 | 19 | 21.58861 | 0.033350689 | Credible Set | purple |
| chr19:21640403 | 19 | 21.64040 | 0.010165871 | Credible Set | purple |
| chr19:21666239 | 19 | 21.66624 | 0.015151776 | Credible Set | purple |
| chr19:21697884 | 19 | 21.69788 | 0.001094068 | Credible Set | purple |
| chr19:21727318 | 19 | 21.72732 | 0.003848370 | Credible Set | purple |
| chr19:21727391 | 19 | 21.72739 | 0.003854631 | Credible Set | purple |
| chr19:21727561 | 19 | 21.72756 | 0.011584296 | Credible Set | purple |
| chr19:21731633 | 19 | 21.73163 | 0.009265556 | Credible Set | purple |
| chr19:21780788 | 19 | 21.78079 | 0.010764350 | Credible Set | purple |
| chr19:21780850 | 19 | 21.78085 | 0.001196689 | Credible Set | purple |
| chr19:21780922 | 19 | 21.78092 | 0.030986314 | Credible Set | purple |
| chr19:21780928 | 19 | 21.78093 | 0.031077481 | Credible Set | purple |
| chr19:21780934 | 19 | 21.78093 | 0.030806278 | Credible Set | purple |
| chr19:21780935 | 19 | 21.78093 | 0.030806278 | Credible Set | purple |
| chr19:21781005 | 19 | 21.78101 | 0.015642938 | Credible Set | purple |
| chr19:21781019 | 19 | 21.78102 | 0.013330150 | Credible Set | purple |
| chr19:21798450 | 19 | 21.79845 | 0.011441822 | Credible Set | purple |
| chr19:21798541 | 19 | 21.79854 | 0.014854567 | Credible Set | purple |

Fine−mapping Regions

| snp | chr | pos (Mb) | pp | group | color |
|---|---|---|---|---|---|
| chr19:21804356 | 19 | 21.80436 | 0.0087403512 | Credible Set | purple |
| chr19:21804420 | 19 | 21.80442 | 0.0117742911 | Credible Set | purple |
| chr19:21804582 | 19 | 21.80458 | 0.0117204809 | Credible Set | purple |
| chr19:21812258 | 19 | 21.81226 | 0.0434339587 | Credible Set | purple |
| chr19:21812298 | 19 | 21.81230 | 0.0449686000 | Credible Set | purple |
| chr19:21812594 | 19 | 21.81259 | 0.0110246233 | Credible Set | purple |
| chr19:21814790 | 19 | 21.81479 | 0.0115665461 | Credible Set | purple |
| chr19:21814806 | 19 | 21.81481 | 0.0114621002 | Credible Set | purple |
| chr19:21851795 | 19 | 21.85179 | 0.0135232085 | Credible Set | purple |
| chr19:21852236 | 19 | 21.85224 | 0.0115893310 | Credible Set | purple |
| chr19:21852273 | 19 | 21.85227 | 0.0120023002 | Credible Set | purple |
| chr19:21852296 | 19 | 21.85230 | 0.0144371770 | Credible Set | purple |
| chr19:21857059 | 19 | 21.85706 | 0.0121634027 | Credible Set | purple |
| chr19:21857128 | 19 | 21.85713 | 0.0113823313 | Credible Set | purple |
| chr19:21857704 | 19 | 21.85770 | 0.0133088652 | Credible Set | purple |
| chr19:21857707 | 19 | 21.85771 | 0.0132335871 | Credible Set | purple |
| chr19:21880097 | 19 | 21.88010 | 0.0148329875 | Credible Set | purple |
| chr19:21889764 | 19 | 21.88976 | 0.0110209332 | Credible Set | purple |
| chr19:21889768 | 19 | 21.88977 | 0.0109843231 | Credible Set | purple |
| chr19:21889777 | 19 | 21.88978 | 0.0110209332 | Credible Set | purple |
| chr19:21889831 | 19 | 21.88983 | 0.0110592602 | Credible Set | purple |
| chr19:21889832 | 19 | 21.88983 | 0.0111010108 | Credible Set | purple |
| chr19:21889838 | 19 | 21.88984 | 0.0110536282 | Credible Set | purple |
| chr19:21895163 | 19 | 21.89516 | 0.0068073283 | Credible Set | purple |
| chr19:21895174 | 19 | 21.89517 | 0.0119227998 | Credible Set | purple |
| chr19:22070939 | 19 | 22.07094 | 0.0013942169 | Credible Set | purple |
| chr19:22073618 | 19 | 22.07362 | 0.0017512896 | Credible Set | purple |
| chr19:22113626 | 19 | 22.11363 | 0.0009951584 | Credible Set | purple |
| chr19:22113628 | 19 | 22.11363 | 0.0009812304 | Credible Set | purple |
| chr19:22113687 | 19 | 22.11369 | 0.0009817933 | Credible Set | purple |

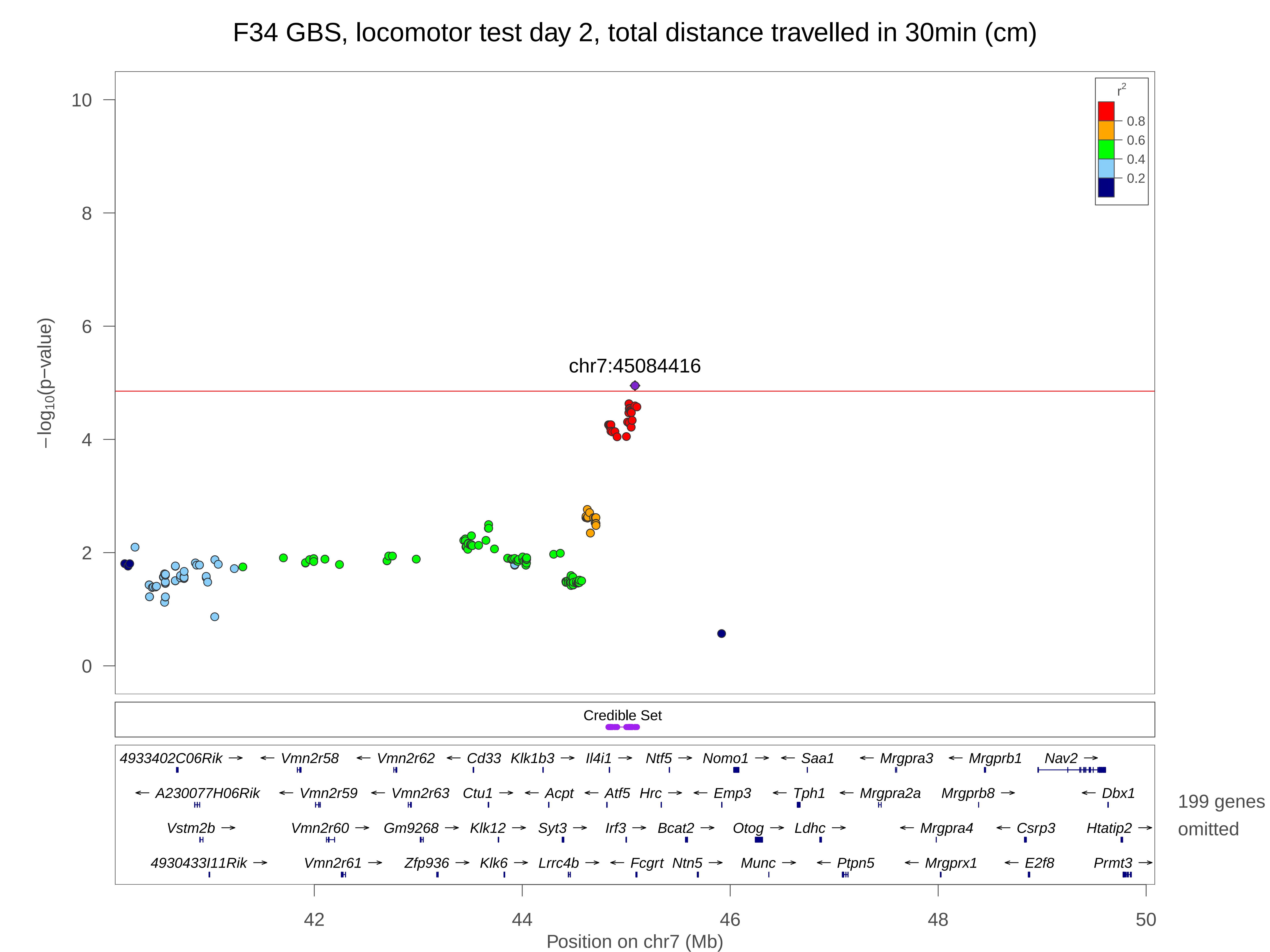
F34 GBS, locomotor test day 2, total distance travelled in 30min (cm)
chr7:45084416
−log10(p−value)
r2
0.8
0.6
0.4
0.2
Credible Set
4933402C06Rik → ← Vmn2r58 ← Vmn2r62 ← Cd33 Klk1b3 → Il4i1 → Ntf5 → Nomo1 → ← Saa1 ← Mrgpra3 ← Mrgprb1 Nav2 →
← A230077H06Rik ← Vmn2r59 ← Vmn2r63 Ctu1 → ← Acpt ← Atf5 Hrc → ← Emp3 ← Tph1 ← Mrgpra2a Mrgprb8 → ← Dbx1
Vstm2b → Vmn2r60 → Gm9268 → Klk12 → Syt3 → Irf3 → Bcat2 → Otog → Ldhc → ← Mrgpra4 ← Csrp3 Htatip2 →
4930433I11Rik → Vmn2r61 → Zfp936 → Klk6 → Lrrc4b → ← Fcgrt Ntn5 → Munc → ← Ptpn5 ← Mrgprx1 ← E2f8 Prmt3 →
199 genes omitted
Position on chr7 (Mb)

date: Thu Oct 25 17:18:53 2018
build: F34GBS
display range: chr7:40084416−50084416 [40084416−50084416]
hilite range: 0 − 0 [ 0 − 0 ]
reference SNP: chr7:45084416
number of SNPs plotted: 225
min P−value: 1.12E−5 [chr7:45084416]
max P−value: 2.69E−1 [chr7:45917340]
omitted Genes: Gm4884, 1700110I07Rik, Gm5592
omitted Genes: AI987944, Vmn2r57, AW146154
omitted Genes: Zfp788, Vmn2r−ps54, Vmn2r−ps60
omitted Genes: 4933421I07Rik, Zfp141, 9830147E19Rik
omitted Genes: Gm5595, Gm2381, EU599041
omitted Genes: Zfp715, Siglecf, Siglecg
omitted Genes: Lim2, Nkg7, Cldnd2
omitted Genes: Etfb, Mir7051, Vsig10l
omitted Genes: Iglon5, 4931406B18Rik, Zfp658
omitted Genes: Zfp719, Zfp819, Ceacam18
omitted Genes: Siglece, Klk14, Klk13
omitted Genes: 2310002F09Rik, Klk11, Klk10
omitted Genes: Klk9, Klk8, Klk7
omitted Genes: Klk5, Klk4, Klk15
omitted Genes: Klk1b7−ps, Klk1b8, Klk1b1
omitted Genes: Klk1b9, Klk1b11, Klk1b26
omitted Genes: Klk1b27, Klk1b21, Klk1b22
omitted Genes: Klk1b16, Klk1b24, Klk1b4
omitted Genes: Klk1b5, Klk1, 1700028J19Rik
omitted Genes: 2410002F23Rik, Snord88c, Snord88a
omitted Genes: Clec11a, Shank1, 1700008O03Rik
omitted Genes: Aspdh, Josd2, Mir7052
omitted Genes: 5430431A17Rik, Emc10, Fam71e1
omitted Genes: Mybpc2, Spib, Pold1
omitted Genes: Mir7053, Nr1h2, Napsa
omitted Genes: Kcnc3, Myh14, Izumo2
omitted Genes: Zfp473, Vrk3, Nup62
omitted Genes: Gm21948, Tbc1d17, Akt1s1
omitted Genes: Mir707, Pnkp, Ptov1
omitted Genes: Med25, Fuz, Ap2a1
omitted Genes: Tsks, Cpt1c, Prmt1
Make more plots at http://csg.sph.umich.edu/locuszoom/
omitted Genes: Gm15545, Bcl2l12, Scaf1

Fine−mapping Regions

| snp | chr | pos (Mb) | pp | group | color |
|---|---|---|---|---|---|
| chr7:44829933 | 7 | 44.82993 | 0.02073332 | Credible Set | purple |
| chr7:44839966 | 7 | 44.83997 | 0.02069526 | Credible Set | purple |
| chr7:44844016 | 7 | 44.84402 | 0.02073662 | Credible Set | purple |
| chr7:44844285 | 7 | 44.84428 | 0.01890285 | Credible Set | purple |
| chr7:44844322 | 7 | 44.84432 | 0.02073662 | Credible Set | purple |
| chr7:44851630 | 7 | 44.85163 | 0.01976668 | Credible Set | purple |
| chr7:44852639 | 7 | 44.85264 | 0.02076204 | Credible Set | purple |
| chr7:44852965 | 7 | 44.85296 | 0.01619501 | Credible Set | purple |
| chr7:44866590 | 7 | 44.86659 | 0.01570153 | Credible Set | purple |
| chr7:44891663 | 7 | 44.89166 | 0.01589941 | Credible Set | purple |
| chr7:44912209 | 7 | 44.91221 | 0.01302882 | Credible Set | purple |
| chr7:45001761 | 7 | 45.00176 | 0.01316522 | Credible Set | purple |
| chr7:45012901 | 7 | 45.01290 | 0.02304829 | Credible Set | purple |
| chr7:45025410 | 7 | 45.02541 | 0.03286698 | Credible Set | purple |
| chr7:45025443 | 7 | 45.02544 | 0.03289392 | Credible Set | purple |
| chr7:45026544 | 7 | 45.02654 | 0.04672684 | Credible Set | purple |
| chr7:45026548 | 7 | 45.02655 | 0.04655413 | Credible Set | purple |
| chr7:45029371 | 7 | 45.02937 | 0.03831053 | Credible Set | purple |
| chr7:45029402 | 7 | 45.02940 | 0.02185794 | Credible Set | purple |
| chr7:45029407 | 7 | 45.02941 | 0.02234440 | Credible Set | purple |
| chr7:45029413 | 7 | 45.02941 | 0.02251314 | Credible Set | purple |
| chr7:45029777 | 7 | 45.02978 | 0.03912933 | Credible Set | purple |
| chr7:45033529 | 7 | 45.03353 | 0.03831053 | Credible Set | purple |
| chr7:45041503 | 7 | 45.04150 | 0.03493138 | Credible Set | purple |
| chr7:45041522 | 7 | 45.04152 | 0.03623031 | Credible Set | purple |
| chr7:45041933 | 7 | 45.04193 | 0.03478022 | Credible Set | purple |
| chr7:45047834 | 7 | 45.04783 | 0.03364742 | Credible Set | purple |
| chr7:45047913 | 7 | 45.04791 | 0.01884979 | Credible Set | purple |
| chr7:45049496 | 7 | 45.04950 | 0.03310522 | Credible Set | purple |
| chr7:45057062 | 7 | 45.05706 | 0.02453473 | Credible Set | purple |
| chr7:45057083 | 7 | 45.05708 | 0.02459962 | Credible Set | purple |
| chr7:45084416 | 7 | 45.08442 | 0.09421839 | Credible Set | purple |
| chr7:45084764 | 7 | 45.08476 | 0.04295456 | Credible Set | purple |
| chr7:45102598 | 7 | 45.10260 | 0.04126896 | Credible Set | purple |

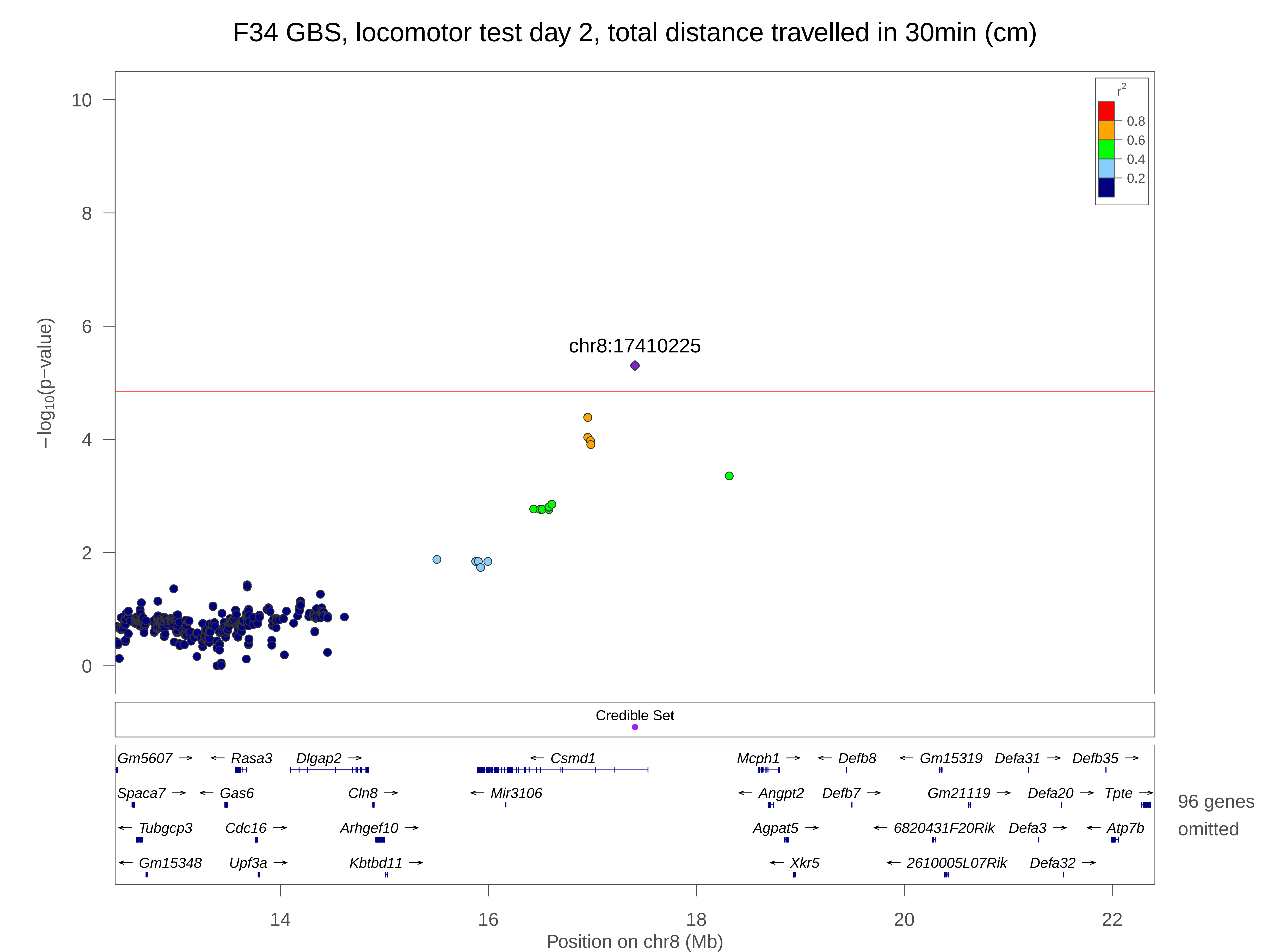

F34 GBS, locomotor test day 2, total distance travelled in 30min (cm)
10
8
6
4
2
0
−log10(p−value)
chr8:17410225
r2
0.8
0.6
0.4
0.2
Credible Set
Gm5607 →
← Rasa3
Dlgap2 →
← Csmd1
Mcph1 →
← Defb8
← Gm15319
Defa31 →
Defb35 →
Spaca7 →
← Gas6
Cln8 →
← Mir3106
← Angpt2
Defb7 →
Gm21119 →
Defa20 →
Tpte →
← Tubgcp3
Cdc16 →
Arhgef10 →
Agpat5 →
← 6820431F20Rik
Defa3 →
← Atp7b
← Gm15348
Upf3a →
Kbtbd11 →
← Xkr5
← 2610005L07Rik
Defa32 →
96 genes omitted
14
16
18
20
22
Position on chr8 (Mb)

| | |
|---|---|
| date: | Thu Oct 25 17:19:03 2018 |
| build: | F34GBS |
| display range: | chr8:12410225−22410225 [12410225−22410225] |
| hilite range: | 0 − 0 [ 0 − 0 ] |
| reference SNP: | chr8:17410225 |
| number of SNPs plotted: | 596 |
| min P−value: | 4.96E−6 [chr8:17410225] |
| max P−value: | 9.99E−1 [chr8:13390417] |
| omitted Genes: | Atp11a, Gm15350, Mcf2l |
| omitted Genes: | Gm38671, F7, F10 |
| omitted Genes: | Proz, Pcid2, Cul4a |
| omitted Genes: | Mir7065, Lamp1, Grtp1 |
| omitted Genes: | Mir1968, Adprhl1, Dcun1d2 |
| omitted Genes: | Tmco3, Tfdp1, Atp4b |
| omitted Genes: | Grk1, Tmem255b, 1700029H14Rik |
| omitted Genes: | 4932443I19Rik, AF366264, Champ1 |
| omitted Genes: | Coprs, Fbxo25, Tdrp |
| omitted Genes: | Gm5907, Erich1, BB014433 |
| omitted Genes: | Myom2, Defb40, Defb37 |
| omitted Genes: | Defb38, Defb39, Defb12 |
| omitted Genes: | Defb34, Spag11b, Spag11a |
| omitted Genes: | Defb14, Defb4, Defa−ps12 |
| omitted Genes: | Defb6, Defb46, Defb5 |
| omitted Genes: | Defb3, Defa−ps13, 4930467E23Rik |
| omitted Genes: | Gm21944, Defb33, Gm15056 |
| omitted Genes: | Gm6040, AY761185, Defa21 |
| omitted Genes: | Defa23, Defa35, Defa25 |
| omitted Genes: | Gm14851, Defa30, Defa22 |
| omitted Genes: | Gm15293, Gm15292, Defa5 |
| omitted Genes: | Defa27, Defa29, Defa2 |
| omitted Genes: | Defa33, Defa36, Gm7849 |
| omitted Genes: | Defa37, Defa28, Gm7861 |
| omitted Genes: | Defa26, Defa17, Defa34 |
| omitted Genes: | Defa−ps1, AY761184, Defa24 |
| omitted Genes: | Defb1, Defb50, Defb2 |
| omitted Genes: | Defb10, Defb9, Defb11 |
| omitted Genes: | Defb15, Defb13, Fam90a1a |
| omitted Genes: | Ccdc70, Alg11, Nek5 |
| omitted Genes: | Nek3, Ckap2, Vps36 |
| omitted Genes: | Thsd1, Slc25a15, 1810012K16Rik |

Make more plots at http://csg.sph.umich.edu/locuszoom/

Fine−mapping Regions

| snp | chr | pos (Mb) | pp | group | color |
|---|---|---|---|---|---|
| chr8:17410225 | 8 | 17.41023 | 1 | Credible Set | purple |

### F34 GBS, quadriceps femoris muscle weight (mg)

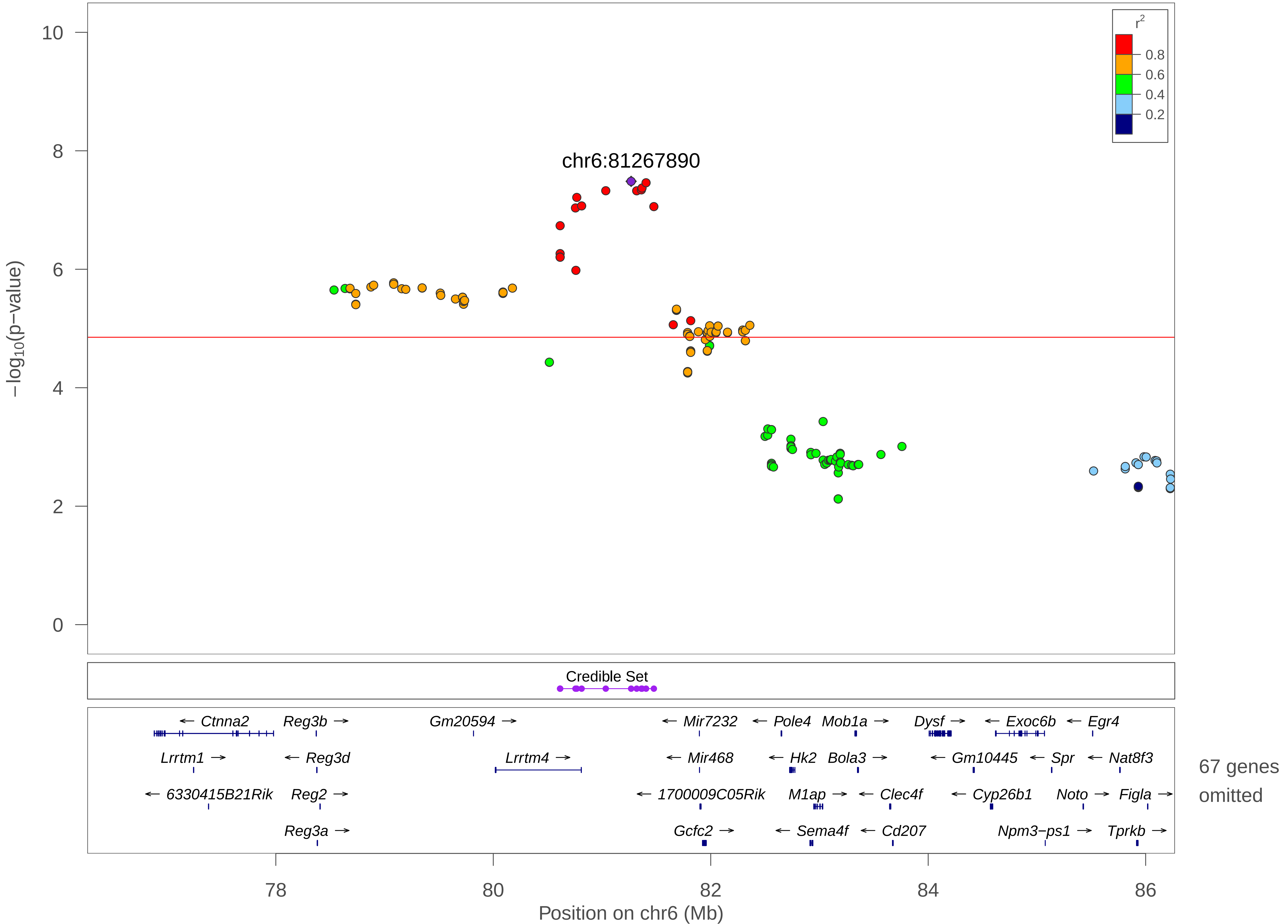

| | |
|---|---|
| date: | Thu Oct 25 17:19:28 2018 |
| build: | F34GBS |
| display range: | chr6:76267890−86267890 [76267890−86267890] |
| hilite range: | 0 − 0 [ 0 − 0 ] |
| reference SNP: | chr6:81267890 |
| number of SNPs plotted: | 166 |
| min P−value: | 3.28E−8 [chr6:81267890] |
| max P−value: | 7.57E−3 [chr6:83172373] |
| omitted Genes: | Reg1, Reg3g, Gm20362 |
| omitted Genes: | Mrpl19, Eva1a, Tacr1 |
| omitted Genes: | 2310069B03Rik, Dok1, Loxl3 |
| omitted Genes: | Mir7040, Htra2, Aup1 |
| omitted Genes: | Dqx1, Tlx2, Pcgf1 |
| omitted Genes: | Lbx2, Mir3470a, Ccdc142 |
| omitted Genes: | Mrpl53, Mogs, Wbp1 |
| omitted Genes: | Ino80b, Rtkn, Wdr54 |
| omitted Genes: | 1700003E16Rik, Dctn1, Slc4a5 |
| omitted Genes: | Mthfd2, Tet3, Mir6374 |
| omitted Genes: | B230319C09Rik, Dguok, Actg2 |
| omitted Genes: | Stambp, Gm21284, Vax2os |
| omitted Genes: | Vax2, Atp6v1b1, 1700124L16Rik |
| omitted Genes: | Ankrd53, Tex261, Gm7443 |
| omitted Genes: | Nagk, Paip2b, Zfp638 |
| omitted Genes: | Gm5878, Emx1, Sfxn5 |
| omitted Genes: | Rab11fip5, Mir705, Smyd5 |
| omitted Genes: | Pradc1, Cct7, Fbxo41 |
| omitted Genes: | Alms1, Nat8f7, Mir6375 |
| omitted Genes: | Alms1−ps2, Nat8f6, Nat8f5 |
| omitted Genes: | Nat8, Nat8f2, Nat8f4 |
| omitted Genes: | Nat8f1, Dusp11, Add2 |
| omitted Genes: | Tgfa |

Make more plots at http://csg.sph.umich.edu/locuszoom/

Fine−mapping Regions

| snp | chr | pos (Mb) | pp | group | color |
|---|---|---|---|---|---|
| chr6:80614710 | 6 | 80.61471 | 0.009415797 | Credible Set | purple |
| chr6:80614725 | 6 | 80.61473 | 0.008218335 | Credible Set | purple |
| chr6:80614740 | 6 | 80.61474 | 0.008298367 | Credible Set | purple |
| chr6:80614750 | 6 | 80.61475 | 0.008218335 | Credible Set | purple |
| chr6:80614907 | 6 | 80.61491 | 0.026822237 | Credible Set | purple |
| chr6:80755939 | 6 | 80.75594 | 0.052295576 | Credible Set | purple |
| chr6:80767933 | 6 | 80.76793 | 0.077712822 | Credible Set | purple |
| chr6:80812582 | 6 | 80.81258 | 0.056469459 | Credible Set | purple |
| chr6:81034488 | 6 | 81.03449 | 0.099977371 | Credible Set | purple |
| chr6:81267890 | 6 | 81.26789 | 0.142429680 | Credible Set | purple |
| chr6:81318989 | 6 | 81.31899 | 0.099719868 | Credible Set | purple |
| chr6:81362217 | 6 | 81.36222 | 0.103292735 | Credible Set | purple |
| chr6:81368112 | 6 | 81.36811 | 0.110689845 | Credible Set | purple |
| chr6:81405109 | 6 | 81.40511 | 0.134991257 | Credible Set | purple |
| chr6:81476902 | 6 | 81.47690 | 0.054991179 | Credible Set | purple |

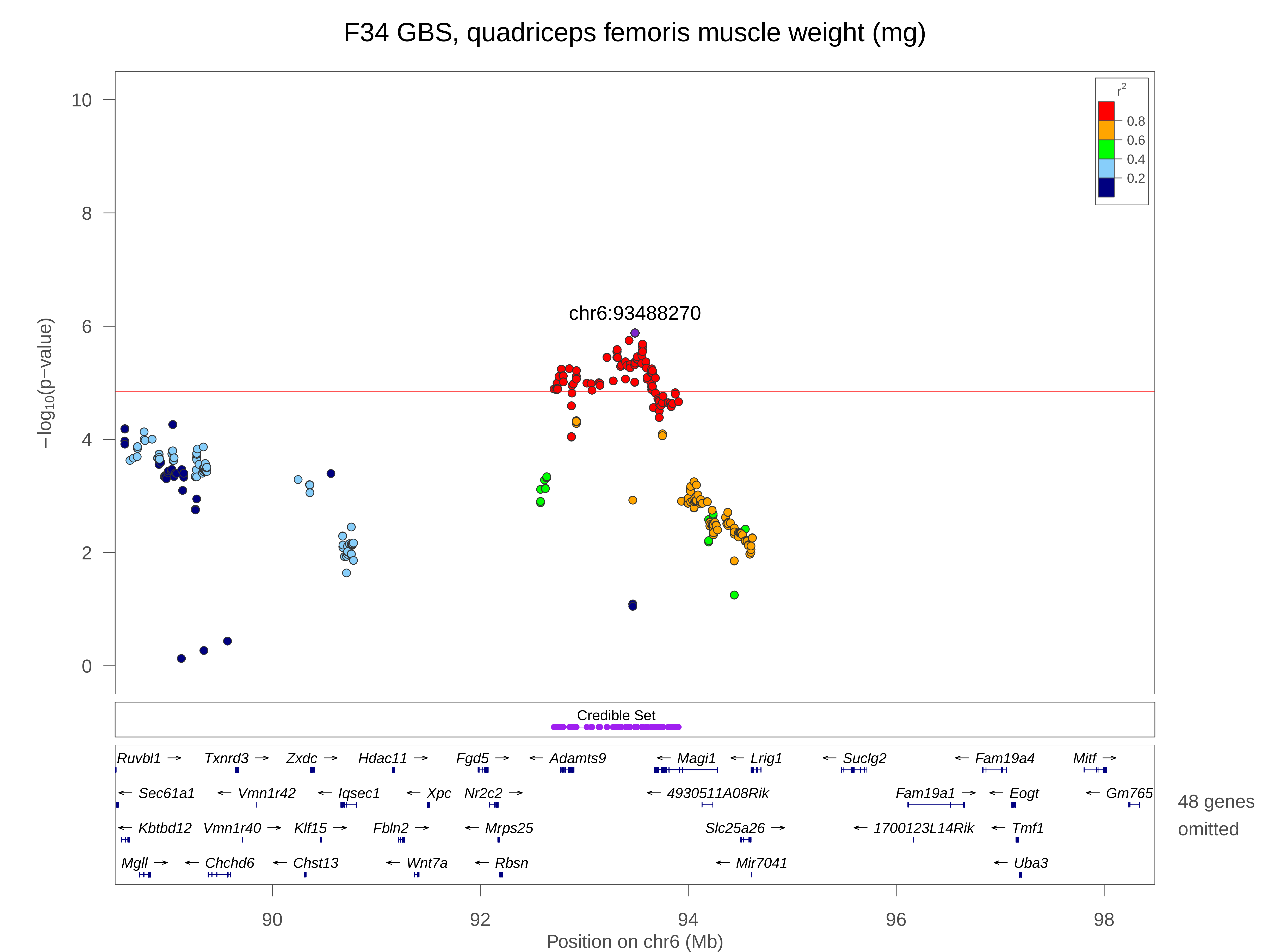

F34 GBS, quadriceps femoris muscle weight (mg)
r2
0.8
0.6
0.4
0.2
chr6:93488270
−log10(p−value)
10
8
6
4
2
0
Credible Set
Ruvbl1 →
Txnrd3 →
Zxdc →
Hdac11 →
Fgd5 →
← Adamts9
← Magi1
← Lrig1
← Suclg2
← Fam19a4
Mitf →
← Sec61a1
← Vmn1r42
← Iqsec1
← Xpc
Nr2c2 →
← 4930511A08Rik
Fam19a1 →
← Eogt
← Gm765
← Kbtbd12
Vmn1r40 →
Klf15 →
Fbln2 →
← Mrps25
Slc25a26 →
← 1700123L14Rik
← Tmf1
Mgll →
← Chchd6
← Chst13
← Wnt7a
← Rbsn
← Mir7041
← Uba3
48 genes omitted
90
92
94
96
98
Position on chr6 (Mb)

| | |
|---|---|
| date: | Thu Oct 25 17:19:37 2018 |
| build: | F34GBS |
| display range: | chr6:88488270−98488270 [88488270−98488270] |
| hilite range: | 0 − 0 [ 0 − 0 ] |
| reference SNP: | chr6:93488270 |
| number of SNPs plotted: | 407 |
| min P−value: | 1.32E−6 [chr6:93488270] |
| max P−value: | 7.41E−1 [chr6:89125612] |
| omitted Genes: | Abtb1, Podxl2, Gm15612 |
| omitted Genes: | Mcm2, Tpra1, 4933427D06Rik |
| omitted Genes: | Gm1965, Gm839, 4930512J16Rik |
| omitted Genes: | Plxna1, Vmn1r41, Vmn1r43 |
| omitted Genes: | Vmn1r44, Vmn1r45, Vmn1r46 |
| omitted Genes: | Vmn1r47, Vmn1r48, Vmn1r49 |
| omitted Genes: | Vmn1r50, Vmn1r51, Vmn1r52 |
| omitted Genes: | V1ra8, Vmn1r53, Vmn1r54 |
| omitted Genes: | BC048671, Uroc1, Cfap100 |
| omitted Genes: | Aldh1l1, Slc41a3, Nup210 |
| omitted Genes: | 4930471M09Rik, 1810044D09Rik, Chchd4 |
| omitted Genes: | Tmem43, Lsm3, Slc6a6 |
| omitted Genes: | Grip2, Ccdc174, 4930590J08Rik |
| omitted Genes: | Trh, Prickle2, 9530026P05Rik |
| omitted Genes: | 4930511E03Rik, Kbtbd8, AY512915 |
| omitted Genes: | Arl6ip5, Lmod3, Frmd4b |

Make more plots at http://csg.sph.umich.edu/locuszoom/

Fine−mapping Regions

| snp | chr | pos (Mb) | pp | group | color |
|---|---|---|---|---|---|
| chr6:92709583 | 6 | 92.70958 | 0.003651370 | Credible Set | purple |
| chr6:92727807 | 6 | 92.72781 | 0.003592184 | Credible Set | purple |
| chr6:92738290 | 6 | 92.73829 | 0.004579701 | Credible Set | purple |
| chr6:92738574 | 6 | 92.73857 | 0.003748828 | Credible Set | purple |
| chr6:92738612 | 6 | 92.73861 | 0.003572039 | Credible Set | purple |
| chr6:92744636 | 6 | 92.74464 | 0.003644646 | Credible Set | purple |
| chr6:92757091 | 6 | 92.75709 | 0.005968885 | Credible Set | purple |
| chr6:92779233 | 6 | 92.77923 | 0.007931589 | Credible Set | purple |
| chr6:92794752 | 6 | 92.79475 | 0.006126065 | Credible Set | purple |
| chr6:92798221 | 6 | 92.79822 | 0.006060561 | Credible Set | purple |
| chr6:92798236 | 6 | 92.79824 | 0.006060561 | Credible Set | purple |
| chr6:92798407 | 6 | 92.79841 | 0.004834126 | Credible Set | purple |
| chr6:92798441 | 6 | 92.79844 | 0.004803612 | Credible Set | purple |
| chr6:92857145 | 6 | 92.85715 | 0.008072631 | Credible Set | purple |
| chr6:92877050 | 6 | 92.87705 | 0.001897090 | Credible Set | purple |
| chr6:92877107 | 6 | 92.87711 | 0.001912643 | Credible Set | purple |
| chr6:92882118 | 6 | 92.88212 | 0.003122060 | Credible Set | purple |
| chr6:92882395 | 6 | 92.88240 | 0.004168587 | Credible Set | purple |
| chr6:92894714 | 6 | 92.89471 | 0.004447249 | Credible Set | purple |
| chr6:92924648 | 6 | 92.92465 | 0.006008710 | Credible Set | purple |
| chr6:92924684 | 6 | 92.92468 | 0.005369544 | Credible Set | purple |
| chr6:92925230 | 6 | 92.92523 | 0.007483354 | Credible Set | purple |
| chr6:93025394 | 6 | 93.02539 | 0.004592582 | Credible Set | purple |
| chr6:93065645 | 6 | 93.06565 | 0.004487367 | Credible Set | purple |
| chr6:93074839 | 6 | 93.07484 | 0.003487255 | Credible Set | purple |
| chr6:93141109 | 6 | 93.14111 | 0.004673172 | Credible Set | purple |
| chr6:93150410 | 6 | 93.15041 | 0.004505086 | Credible Set | purple |
| chr6:93150661 | 6 | 93.15066 | 0.004538087 | Credible Set | purple |
| chr6:93150825 | 6 | 93.15082 | 0.004210537 | Credible Set | purple |
| chr6:93150870 | 6 | 93.15087 | 0.004219324 | Credible Set | purple |
| chr6:93218675 | 6 | 93.21868 | 0.012614793 | Credible Set | purple |
| chr6:93218685 | 6 | 93.21868 | 0.012615602 | Credible Set | purple |
| chr6:93218689 | 6 | 93.21869 | 0.012408312 | Credible Set | purple |
| chr6:93278268 | 6 | 93.27827 | 0.005004540 | Credible Set | purple |
| chr6:93278280 | 6 | 93.27828 | 0.005004540 | Credible Set | purple |
| chr6:93278290 | 6 | 93.27829 | 0.005004540 | Credible Set | purple |

Fine−mapping Regions

| snp | chr | pos (Mb) | pp | group | color |
|---|---|---|---|---|---|
| chr6:93313781 | 6 | 93.31378 | 0.012508707 | Credible Set | purple |
| chr6:93313998 | 6 | 93.31400 | 0.015701627 | Credible Set | purple |
| chr6:93314001 | 6 | 93.31400 | 0.015749337 | Credible Set | purple |
| chr6:93316695 | 6 | 93.31669 | 0.016994413 | Credible Set | purple |
| chr6:93316707 | 6 | 93.31671 | 0.015605240 | Credible Set | purple |
| chr6:93316769 | 6 | 93.31677 | 0.016935319 | Credible Set | purple |
| chr6:93321239 | 6 | 93.32124 | 0.012427973 | Credible Set | purple |
| chr6:93349184 | 6 | 93.34918 | 0.008792167 | Credible Set | purple |
| chr6:93358542 | 6 | 93.35854 | 0.009266523 | Credible Set | purple |
| chr6:93396517 | 6 | 93.39652 | 0.005376651 | Credible Set | purple |
| chr6:93396531 | 6 | 93.39653 | 0.005376651 | Credible Set | purple |
| chr6:93396614 | 6 | 93.39661 | 0.010572450 | Credible Set | purple |
| chr6:93410238 | 6 | 93.41024 | 0.009119871 | Credible Set | purple |
| chr6:93431052 | 6 | 93.43105 | 0.024395752 | Credible Set | purple |
| chr6:93431078 | 6 | 93.43108 | 0.024135995 | Credible Set | purple |
| chr6:93431080 | 6 | 93.43108 | 0.024135995 | Credible Set | purple |
| chr6:93431081 | 6 | 93.43108 | 0.024135995 | Credible Set | purple |
| chr6:93431095 | 6 | 93.43109 | 0.024135995 | Credible Set | purple |
| chr6:93440971 | 6 | 93.44097 | 0.009437307 | Credible Set | purple |
| chr6:93440986 | 6 | 93.44099 | 0.008295711 | Credible Set | purple |
| chr6:93485590 | 6 | 93.48559 | 0.009333079 | Credible Set | purple |
| chr6:93485603 | 6 | 93.48560 | 0.009311901 | Credible Set | purple |
| chr6:93485684 | 6 | 93.48568 | 0.004734015 | Credible Set | purple |
| chr6:93485761 | 6 | 93.48576 | 0.004782209 | Credible Set | purple |
| chr6:93487880 | 6 | 93.48788 | 0.010479311 | Credible Set | purple |
| chr6:93488270 | 6 | 93.48827 | 0.032496395 | Credible Set | purple |
| chr6:93488305 | 6 | 93.48830 | 0.032485504 | Credible Set | purple |
| chr6:93488315 | 6 | 93.48832 | 0.032113205 | Credible Set | purple |
| chr6:93488485 | 6 | 93.48848 | 0.010037475 | Credible Set | purple |
| chr6:93510428 | 6 | 93.51043 | 0.011547964 | Credible Set | purple |
| chr6:93510476 | 6 | 93.51048 | 0.011652741 | Credible Set | purple |
| chr6:93510483 | 6 | 93.51048 | 0.011705898 | Credible Set | purple |
| chr6:93510591 | 6 | 93.51059 | 0.012851228 | Credible Set | purple |
| chr6:93552797 | 6 | 93.55280 | 0.009915162 | Credible Set | purple |
| chr6:93552882 | 6 | 93.55288 | 0.009929088 | Credible Set | purple |
| chr6:93552911 | 6 | 93.55291 | 0.014300970 | Credible Set | purple |

Fine−mapping Regions

| snp | chr | pos (Mb) | pp | group | color |
|---|---|---|---|---|---|
| chr6:93555049 | 6 | 93.55505 | 0.013290510 | Credible Set | purple |
| chr6:93560069 | 6 | 93.56007 | 0.016617122 | Credible Set | purple |
| chr6:93560082 | 6 | 93.56008 | 0.019143733 | Credible Set | purple |
| chr6:93560724 | 6 | 93.56072 | 0.018899195 | Credible Set | purple |
| chr6:93560726 | 6 | 93.56073 | 0.015685733 | Credible Set | purple |
| chr6:93560747 | 6 | 93.56075 | 0.021029916 | Credible Set | purple |
| chr6:93593460 | 6 | 93.59346 | 0.010368451 | Credible Set | purple |
| chr6:93593464 | 6 | 93.59346 | 0.010523745 | Credible Set | purple |
| chr6:93598219 | 6 | 93.59822 | 0.008243905 | Credible Set | purple |
| chr6:93604482 | 6 | 93.60448 | 0.005360362 | Credible Set | purple |
| chr6:93604527 | 6 | 93.60453 | 0.005721503 | Credible Set | purple |
| chr6:93641872 | 6 | 93.64187 | 0.006908577 | Credible Set | purple |
| chr6:93648881 | 6 | 93.64888 | 0.004038566 | Credible Set | purple |
| chr6:93648927 | 6 | 93.64893 | 0.003935323 | Credible Set | purple |
| chr6:93648944 | 6 | 93.64894 | 0.006768429 | Credible Set | purple |
| chr6:93649344 | 6 | 93.64934 | 0.004467091 | Credible Set | purple |
| chr6:93649429 | 6 | 93.64943 | 0.008010358 | Credible Set | purple |
| chr6:93649450 | 6 | 93.64945 | 0.008010358 | Credible Set | purple |
| chr6:93649512 | 6 | 93.64951 | 0.003559046 | Credible Set | purple |
| chr6:93655527 | 6 | 93.65553 | 0.003559189 | Credible Set | purple |
| chr6:93655546 | 6 | 93.65555 | 0.004045917 | Credible Set | purple |
| chr6:93656134 | 6 | 93.65613 | 0.006752322 | Credible Set | purple |
| chr6:93656164 | 6 | 93.65616 | 0.007472931 | Credible Set | purple |
| chr6:93684217 | 6 | 93.68422 | 0.005596328 | Credible Set | purple |
| chr6:93684273 | 6 | 93.68427 | 0.005601768 | Credible Set | purple |
| chr6:93684716 | 6 | 93.68472 | 0.003108959 | Credible Set | purple |
| chr6:93708258 | 6 | 93.70826 | 0.002505461 | Credible Set | purple |
| chr6:93716266 | 6 | 93.71627 | 0.002378404 | Credible Set | purple |
| chr6:93718483 | 6 | 93.71848 | 0.002087457 | Credible Set | purple |
| chr6:93721362 | 6 | 93.72136 | 0.002257798 | Credible Set | purple |
| chr6:93737282 | 6 | 93.73728 | 0.001918222 | Credible Set | purple |
| chr6:93752696 | 6 | 93.75270 | 0.002155863 | Credible Set | purple |
| chr6:93757437 | 6 | 93.75744 | 0.002767976 | Credible Set | purple |
| chr6:93757440 | 6 | 93.75744 | 0.002773412 | Credible Set | purple |
| chr6:93757443 | 6 | 93.75744 | 0.002771953 | Credible Set | purple |
| chr6:93806708 | 6 | 93.80671 | 0.002114076 | Credible Set | purple |

Fine−mapping Regions

| snp | chr | pos (Mb) | pp | group | color |
|---|---|---|---|---|---|
| chr6:93806963 | 6 | 93.80696 | 0.002148832 | Credible Set | purple |
| chr6:93806994 | 6 | 93.80699 | 0.002143401 | Credible Set | purple |
| chr6:93828162 | 6 | 93.82816 | 0.002106132 | Credible Set | purple |
| chr6:93835869 | 6 | 93.83587 | 0.001893876 | Credible Set | purple |
| chr6:93835874 | 6 | 93.83587 | 0.001878328 | Credible Set | purple |
| chr6:93836339 | 6 | 93.83634 | 0.001924572 | Credible Set | purple |
| chr6:93836347 | 6 | 93.83635 | 0.001867839 | Credible Set | purple |
| chr6:93850047 | 6 | 93.85005 | 0.002077286 | Credible Set | purple |
| chr6:93875308 | 6 | 93.87531 | 0.003169621 | Credible Set | purple |
| chr6:93875350 | 6 | 93.87535 | 0.002993688 | Credible Set | purple |
| chr6:93907106 | 6 | 93.90711 | 0.002233728 | Credible Set | purple |

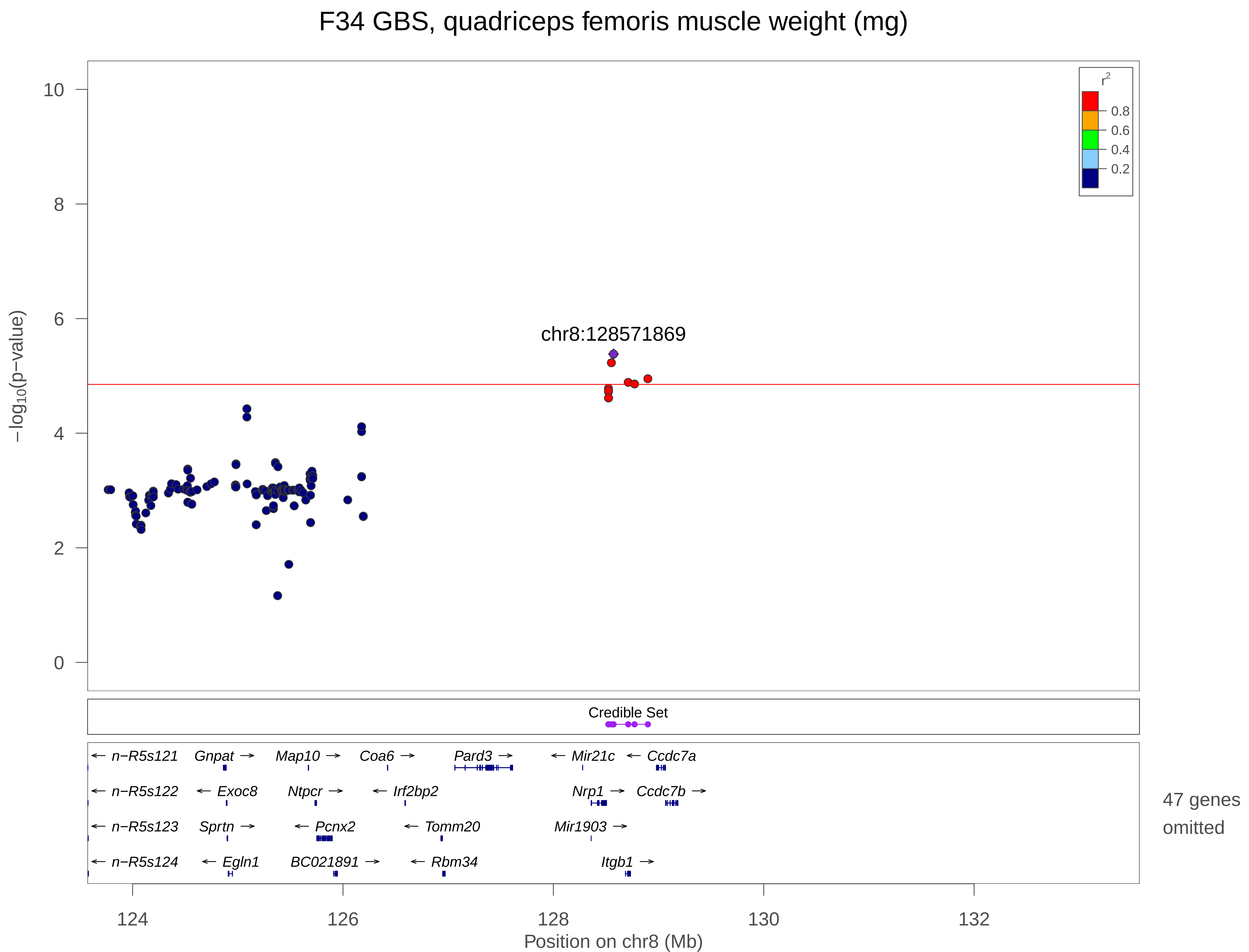
F34 GBS, quadriceps femoris muscle weight (mg)
chr8:128571869
r2
0.8
0.6
0.4
0.2
−log10(p−value)
0
2
4
6
8
10
Credible Set
← n−R5s121
Gnpat →
Map10 →
Coa6 →
Pard3 →
← Mir21c
← Ccdc7a
← n−R5s122
← Exoc8
Ntpcr →
← Irf2bp2
Nrp1 →
Ccdc7b →
← n−R5s123
Sprtn →
← Pcnx2
← Tomm20
Mir1903 →
← n−R5s124
← Egln1
BC021891 →
← Rbm34
Itgb1 →
47 genes omitted
124
126
128
130
132
Position on chr8 (Mb)

| | |
|---|---|
| date: | Thu Oct 25 17:19:48 2018 |
| build: | F34GBS |
| display range: | chr8:123571869−133571869 [123571869−133571869] |
| hilite range: | 0 − 0 [ 0 − 0 ] |
| reference SNP: | chr8:128571869 |
| number of SNPs plotted: | 191 |
| min P−value: | 4.15E−6 [chr8:128571869] |
| max P−value: | 6.86E−2 [chr8:125378859] |
| omitted Genes: | n−R5s128, Gm26391, n−R5s133 |
| omitted Genes: | n−R5s134, Rn5s, n−R5s136 |
| omitted Genes: | n−R5s138, n−R5s139, Gm22109 |
| omitted Genes: | n−R5s141, n−R5s142, n−R5s143 |
| omitted Genes: | n−R5s144, Gm25212, n−R5s146 |
| omitted Genes: | Gm22291, Gm23284, n−R5s149 |
| omitted Genes: | Rhou, 6030466F02Rik, Rab4a |
| omitted Genes: | Ccsap, Acta1, Nup133 |
| omitted Genes: | Abcb10, Taf5l, Mir1967 |
| omitted Genes: | Urb2, Galnt2, Pgbd5 |
| omitted Genes: | Cog2, Agt, Capn9 |
| omitted Genes: | 2310022B05Rik, Ttc13, Arv1 |
| omitted Genes: | Fam89a, Trim67, 2810004N23Rik |
| omitted Genes: | Tsnax, Disc1, Sipa1l2 |
| omitted Genes: | 4930567H12Rik, Kcnk1, Slc35f3 |
| omitted Genes: | Gm17296, 2610044O15Rik8 |

Make more plots at http://csg.sph.umich.edu/locuszoom/

Fine−mapping Regions

| snp | chr | pos (Mb) | pp | group | color |
|---|---|---|---|---|---|
| chr8:128523972 | 8 | 128.5240 | 0.03751086 | Credible Set | purple |
| chr8:128524047 | 8 | 128.5240 | 0.03967617 | Credible Set | purple |
| chr8:128524048 | 8 | 128.5240 | 0.04232385 | Credible Set | purple |
| chr8:128524081 | 8 | 128.5241 | 0.03856074 | Credible Set | purple |
| chr8:128524424 | 8 | 128.5244 | 0.02941370 | Credible Set | purple |
| chr8:128524425 | 8 | 128.5244 | 0.02936529 | Credible Set | purple |
| chr8:128524441 | 8 | 128.5244 | 0.02942450 | Credible Set | purple |
| chr8:128551483 | 8 | 128.5515 | 0.11342310 | Credible Set | purple |
| chr8:128571869 | 8 | 128.5719 | 0.15873044 | Credible Set | purple |
| chr8:128571875 | 8 | 128.5719 | 0.15873044 | Credible Set | purple |
| chr8:128571887 | 8 | 128.5719 | 0.15788269 | Credible Set | purple |
| chr8:128711717 | 8 | 128.7117 | 0.05340323 | Credible Set | purple |
| chr8:128771719 | 8 | 128.7717 | 0.05017147 | Credible Set | purple |
| chr8:128898075 | 8 | 128.8981 | 0.06138352 | Credible Set | purple |

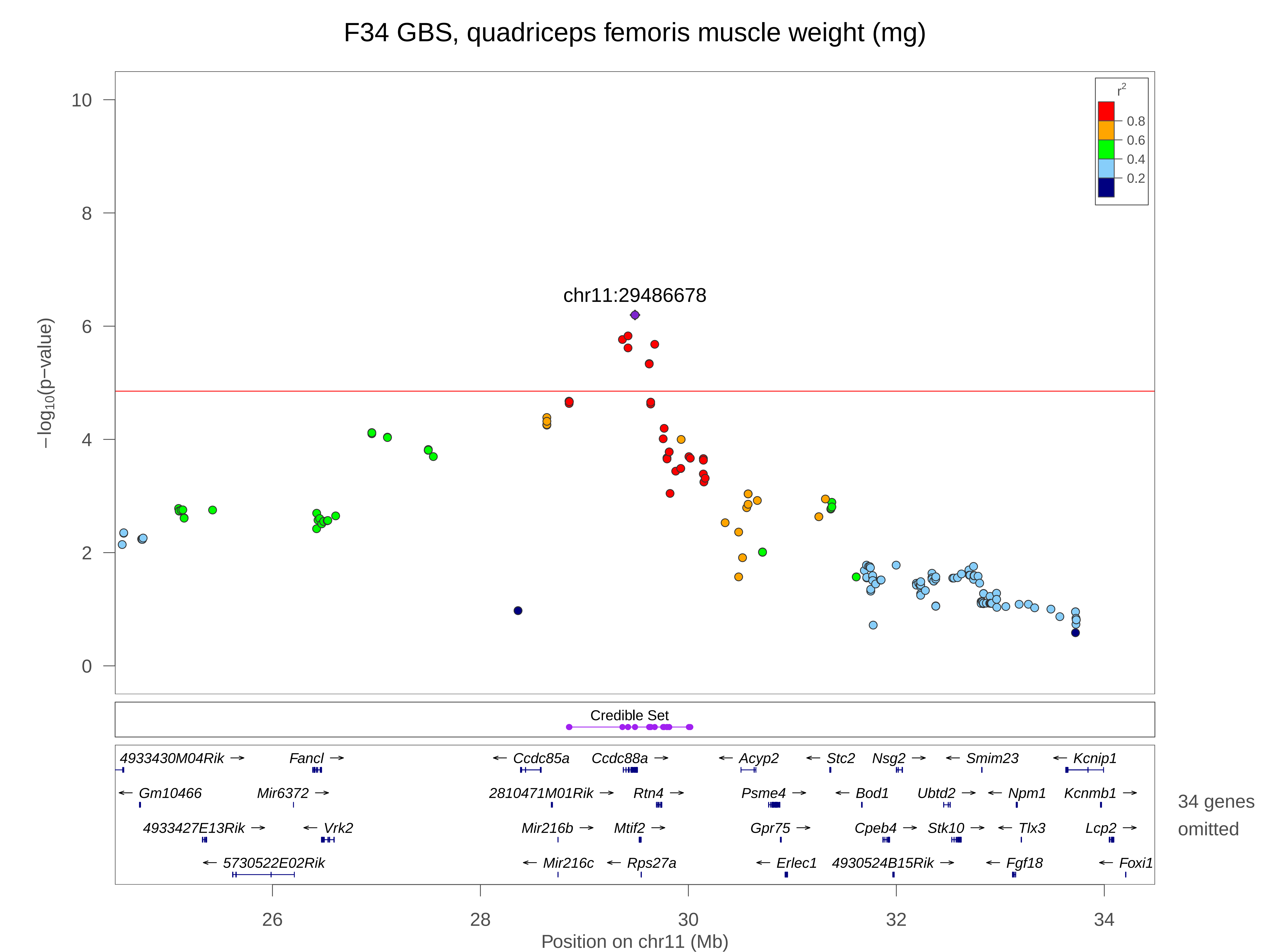

F34 GBS, quadriceps femoris muscle weight (mg)
−log10(p−value)
chr11:29486678
r2
0.8
0.6
0.4
0.2
Credible Set
4933430M04Rik
Fancl
Ccdc85a
Ccdc88a
Acyp2
Stc2
Nsg2
Smim23
Kcnip1
Gm10466
Mir6372
2810471M01Rik
Rtn4
Psme4
Bod1
Ubtd2
Npm1
Kcnmb1
4933427E13Rik
Vrk2
Mir216b
Mtif2
Gpr75
Cpeb4
Stk10
Tlx3
Lcp2
5730522E02Rik
Mir216c
Rps27a
Erlec1
4930524B15Rik
Fgf18
Foxi1
34 genes omitted
Position on chr11 (Mb)

| | |
|---|---|
| date: | Thu Oct 25 17:19:11 2018 |
| build: | F34GBS |
| display range: | chr11:24486678−34486678 [24486678−34486678] |
| hilite range: | 0 − 0 [ 0 − 0 ] |
| reference SNP: | chr11:29486678 |
| number of SNPs plotted: | 208 |
| min P−value: | 6.35E−7 [chr11:29486678] |
| max P−value: | 2.6E−1 [chr11:33723026] |
| omitted Genes: | Gm12070, Mir216a, Mir217 |
| omitted Genes: | Efemp1, Pnpt1, Smek2 |
| omitted Genes: | Cfap36, Prorsd1, Clhc1 |
| omitted Genes: | Eml6, 4931440F15Rik, Sptbn1 |
| omitted Genes: | 4930505A04Rik, Asb3, Chac2 |
| omitted Genes: | D630024D03Rik, Gm12108, Il9r |
| omitted Genes: | Snrnp25, Rhbdf1, Mpg |
| omitted Genes: | Nprl3, Hba−x, Hba−a1 |
| omitted Genes: | Hbq1b, Hba−a2, Hbq1a |
| omitted Genes: | Sh3pxd2b, Efcab9, Fbxw11 |
| omitted Genes: | Ranbp17, Gabrp, Dock2 |
| omitted Genes: | Fam196b |

Fine−mapping Regions

| snp | chr | pos (Mb) | pp | group | color |
|---|---|---|---|---|---|
| chr11:28851860 | 11 | 28.85186 | 0.011196072 | Credible Set | purple |
| chr11:28853308 | 11 | 28.85331 | 0.010233745 | Credible Set | purple |
| chr11:28855496 | 11 | 28.85550 | 0.010963145 | Credible Set | purple |
| chr11:29365866 | 11 | 29.36587 | 0.123345163 | Credible Set | purple |
| chr11:29419484 | 11 | 29.41948 | 0.088878362 | Credible Set | purple |
| chr11:29419643 | 11 | 29.41964 | 0.142119135 | Credible Set | purple |
| chr11:29486678 | 11 | 29.48668 | 0.322110318 | Credible Set | purple |
| chr11:29623528 | 11 | 29.62353 | 0.048248804 | Credible Set | purple |
| chr11:29623531 | 11 | 29.62353 | 0.048135344 | Credible Set | purple |
| chr11:29623542 | 11 | 29.62354 | 0.047430361 | Credible Set | purple |
| chr11:29637584 | 11 | 29.63758 | 0.010014850 | Credible Set | purple |
| chr11:29637586 | 11 | 29.63759 | 0.010803516 | Credible Set | purple |
| chr11:29676398 | 11 | 29.67640 | 0.102343124 | Credible Set | purple |
| chr11:29757518 | 11 | 29.75752 | 0.002624170 | Credible Set | purple |
| chr11:29767130 | 11 | 29.76713 | 0.003921785 | Credible Set | purple |
| chr11:29794101 | 11 | 29.79410 | 0.001266856 | Credible Set | purple |
| chr11:29794488 | 11 | 29.79449 | 0.001248625 | Credible Set | purple |
| chr11:29815423 | 11 | 29.81542 | 0.001587720 | Credible Set | purple |
| chr11:29815516 | 11 | 29.81552 | 0.001587720 | Credible Set | purple |
| chr11:30005177 | 11 | 30.00518 | 0.001324759 | Credible Set | purple |
| chr11:30018452 | 11 | 30.01845 | 0.001247503 | Credible Set | purple |

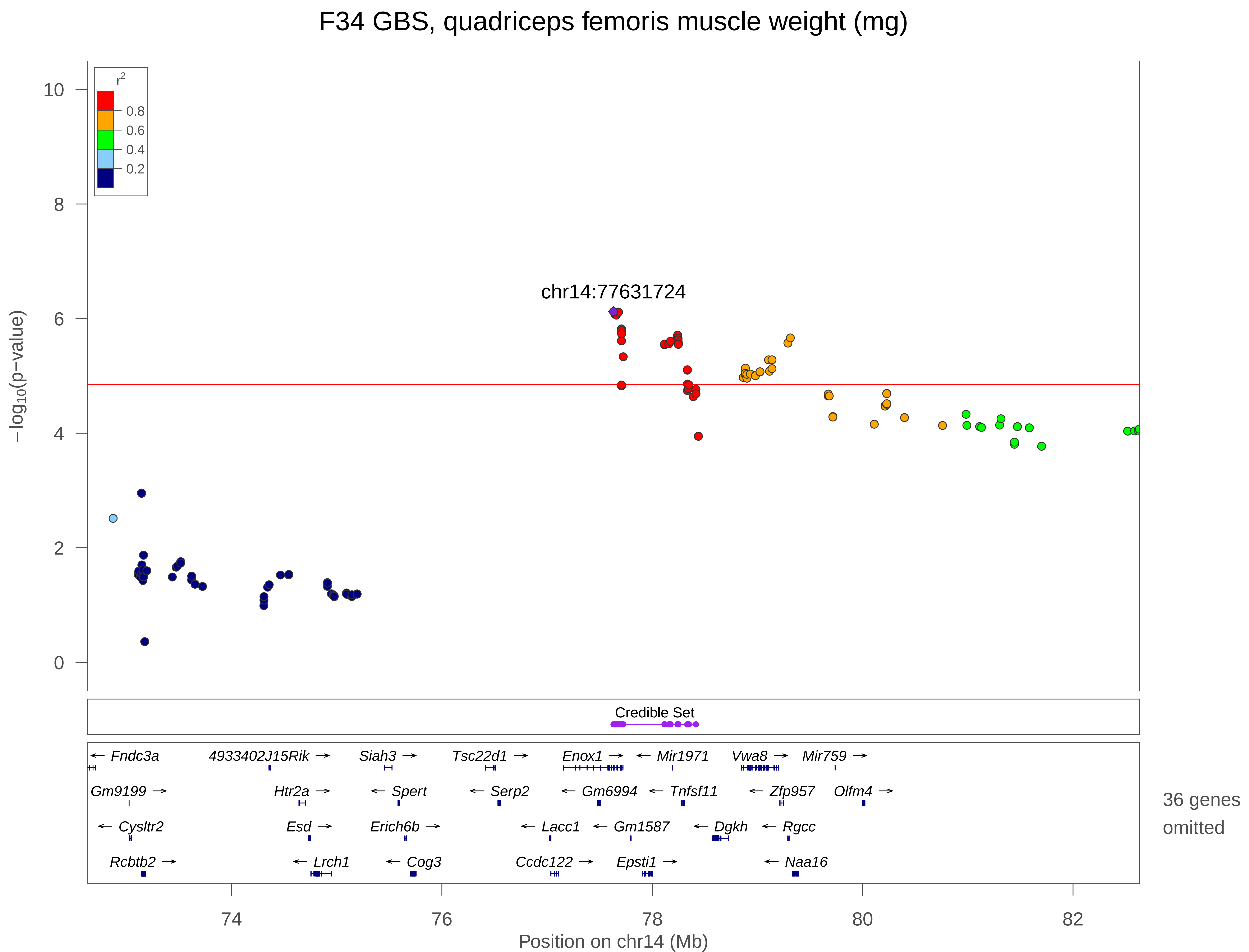

F34 GBS, quadriceps femoris muscle weight (mg)
r2
0.8
0.6
0.4
0.2
chr14:77631724
−log10(p−value)
Credible Set
Fndc3a
4933402J15Rik
Siah3
Tsc22d1
Enox1
Mir1971
Vwa8
Mir759
Gm9199
Htr2a
Spert
Serp2
Gm6994
Tnfsf11
Zfp957
Olfm4
Cysltr2
Esd
Erich6b
Lacc1
Gm1587
Dgkh
Rgcc
Rcbtb2
Lrch1
Cog3
Ccdc122
Epsti1
Naa16
36 genes omitted
Position on chr14 (Mb)

| | |
|---|---|
| date: | Thu Oct 25 17:19:19 2018 |
| build: | F34GBS |
| display range: | chr14:72631724−82631724 [72631724−82631724] |
| hilite range: | 0 − 0 [ 0 − 0 ] |
| reference SNP: | chr14:77631724 |
| number of SNPs plotted: | 152 |
| min P−value: | 7.58E−7 [chr14:77631724] |
| max P−value: | 4.34E−1 [chr14:73175203] |
| omitted Genes: | Rb1, Mir687, Lpar6 |
| omitted Genes: | Itm2b, Med4, Nudt15 |
| omitted Genes: | Sucla2, Gm4278, 5031414D18Rik |
| omitted Genes: | Lrrc63, Lcp1, Cpb2 |
| omitted Genes: | Zc3h13, Slc25a30, Gm4285 |
| omitted Genes: | Tpt1, Snora31, Gtf2f2 |
| omitted Genes: | Kctd4, Gpalpp1, Nufip1 |
| omitted Genes: | 4930444M15Rik, 1700108F19Rik, Dnajc15 |
| omitted Genes: | Fam216b, Akap11, Mtrf1 |
| omitted Genes: | Kbtbd7, Kbtbd6, Wbp4 |
| omitted Genes: | Elf1, Sugt1, Lect1 |
| omitted Genes: | 4930452G13Rik, Pcdh8, Gm10845 |

Make more plots at http://csg.sph.umich.edu/locuszoom/

Fine−mapping Regions

| snp | chr | pos (Mb) | pp | group | color |
|---|---|---|---|---|---|
| chr14:77631724 | 14 | 77.63172 | 0.065951423 | Credible Set | purple |
| chr14:77642597 | 14 | 77.64260 | 0.065442836 | Credible Set | purple |
| chr14:77642802 | 14 | 77.64280 | 0.065442836 | Credible Set | purple |
| chr14:77647283 | 14 | 77.64728 | 0.059571500 | Credible Set | purple |
| chr14:77647687 | 14 | 77.64769 | 0.061712381 | Credible Set | purple |
| chr14:77658356 | 14 | 77.65836 | 0.057734634 | Credible Set | purple |
| chr14:77678847 | 14 | 77.67885 | 0.065206095 | Credible Set | purple |
| chr14:77678929 | 14 | 77.67893 | 0.064608702 | Credible Set | purple |
| chr14:77707033 | 14 | 77.70703 | 0.033959058 | Credible Set | purple |
| chr14:77707351 | 14 | 77.70735 | 0.021432720 | Credible Set | purple |
| chr14:77707392 | 14 | 77.70739 | 0.021526237 | Credible Set | purple |
| chr14:77707431 | 14 | 77.70743 | 0.003874422 | Credible Set | purple |
| chr14:77707491 | 14 | 77.70749 | 0.003776912 | Credible Set | purple |
| chr14:77707493 | 14 | 77.70749 | 0.003911421 | Credible Set | purple |
| chr14:77707780 | 14 | 77.70778 | 0.030662519 | Credible Set | purple |
| chr14:77707832 | 14 | 77.70783 | 0.031648653 | Credible Set | purple |
| chr14:77708407 | 14 | 77.70841 | 0.028103002 | Credible Set | purple |
| chr14:77724180 | 14 | 77.72418 | 0.011581226 | Credible Set | purple |
| chr14:78116598 | 14 | 78.11660 | 0.018378782 | Credible Set | purple |
| chr14:78116913 | 14 | 78.11691 | 0.018911253 | Credible Set | purple |
| chr14:78157408 | 14 | 78.15741 | 0.019006437 | Credible Set | purple |
| chr14:78173886 | 14 | 78.17389 | 0.021007088 | Credible Set | purple |
| chr14:78240912 | 14 | 78.24091 | 0.024711263 | Credible Set | purple |
| chr14:78240962 | 14 | 78.24096 | 0.025038119 | Credible Set | purple |
| chr14:78241501 | 14 | 78.24150 | 0.026369333 | Credible Set | purple |
| chr14:78241810 | 14 | 78.24181 | 0.026750039 | Credible Set | purple |
| chr14:78245017 | 14 | 78.24502 | 0.023761908 | Credible Set | purple |
| chr14:78245066 | 14 | 78.24507 | 0.022104762 | Credible Set | purple |
| chr14:78247459 | 14 | 78.24746 | 0.019855882 | Credible Set | purple |
| chr14:78247805 | 14 | 78.24780 | 0.018647975 | Credible Set | purple |
| chr14:78333693 | 14 | 78.33369 | 0.006912693 | Credible Set | purple |
| chr14:78333727 | 14 | 78.33373 | 0.007022265 | Credible Set | purple |
| chr14:78334235 | 14 | 78.33424 | 0.003168384 | Credible Set | purple |
| chr14:78334352 | 14 | 78.33435 | 0.004061125 | Credible Set | purple |
| chr14:78342935 | 14 | 78.34293 | 0.003405232 | Credible Set | purple |
| chr14:78349294 | 14 | 78.34929 | 0.003907212 | Credible Set | purple |

Fine−mapping Regions

| snp | chr | pos (Mb) | pp | group | color |
|---|---|---|---|---|---|
| chr14:78415035 | 14 | 78.41504 | 0.003307998 | Credible Set | purple |

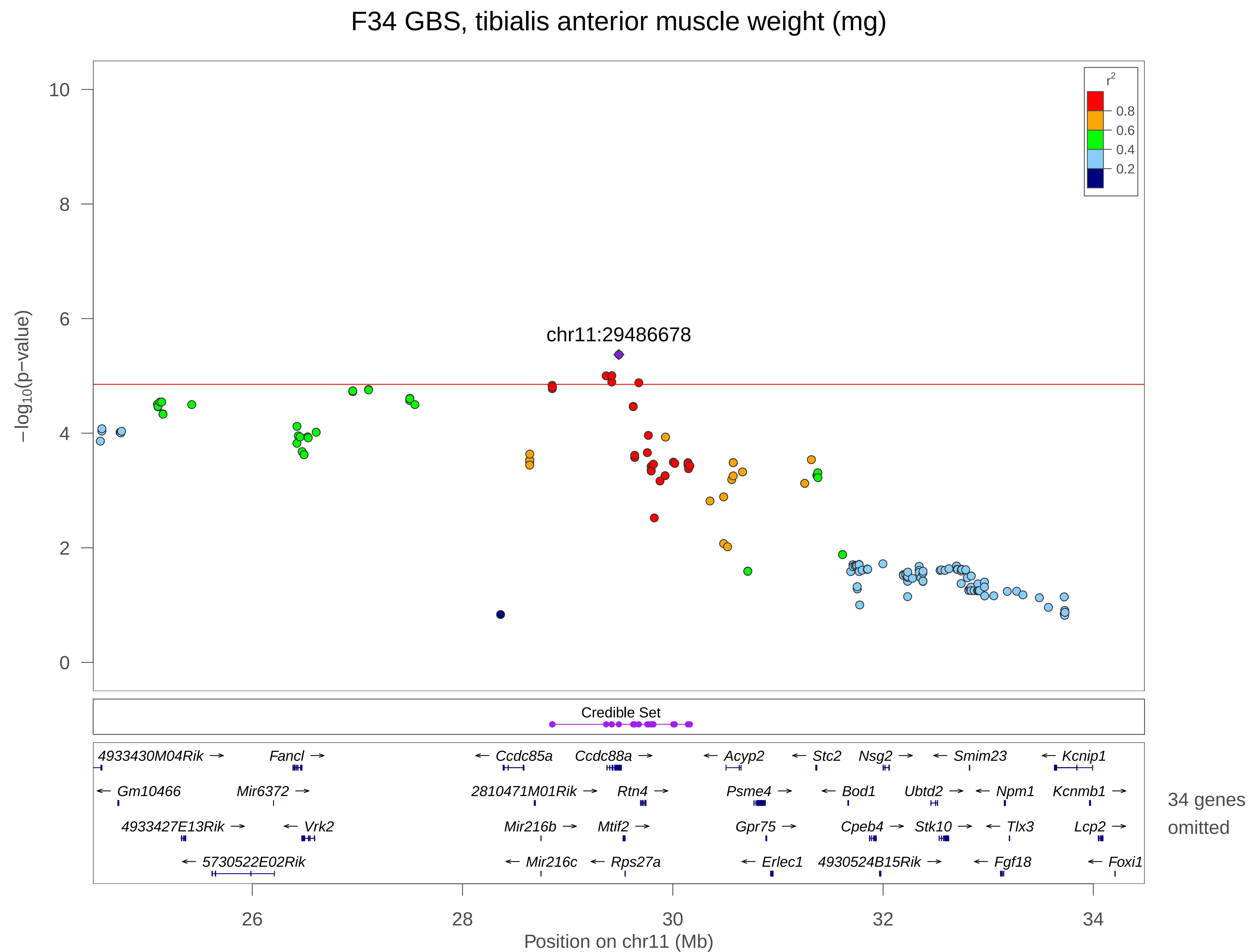

F34 GBS, tibialis anterior muscle weight (mg)
−log10(p−value)
10
8
6
4
2
0
r2
0.8
0.6
0.4
0.2
chr11:29486678
Credible Set
4933430M04Rik →
Fancl →
← Ccdc85a
Ccdc88a →
← Acyp2
← Stc2
Nsg2 →
← Smim23
← Kcnip1
← Gm10466
Mir6372 →
2810471M01Rik →
Rtn4 →
Psme4 →
← Bod1
Ubtd2 →
← Npm1
Kcnmb1 →
4933427E13Rik →
← Vrk2
Mir216b →
Mtif2 →
Gpr75 →
Cpeb4 →
Stk10 →
← Tlx3
Lcp2 →
← 5730522E02Rik
← Mir216c
← Rps27a
← Erlec1
4930524B15Rik →
← Fgf18
← Foxi1
34 genes omitted
26
28
30
32
34
Position on chr11 (Mb)

| | |
|---|---|
| date: | Thu Oct 25 17:19:56 2018 |
| build: | F34GBS |
| display range: | chr11:24486678−34486678 [24486678−34486678] |
| hilite range: | 0 − 0 [ 0 − 0 ] |
| reference SNP: | chr11:29486678 |
| number of SNPs plotted: | 208 |
| min P−value: | 4.24E−6 [chr11:29486678] |
| max P−value: | 1.51E−1 [chr11:33727201] |
| omitted Genes: | Gm12070, Mir216a, Mir217 |
| omitted Genes: | Efemp1, Pnpt1, Smek2 |
| omitted Genes: | Cfap36, Prorsd1, Clhc1 |
| omitted Genes: | Eml6, 4931440F15Rik, Sptbn1 |
| omitted Genes: | 4930505A04Rik, Asb3, Chac2 |
| omitted Genes: | D630024D03Rik, Gm12108, Il9r |
| omitted Genes: | Snrnp25, Rhbdf1, Mpg |
| omitted Genes: | Nprl3, Hba−x, Hba−a1 |
| omitted Genes: | Hbq1b, Hba−a2, Hbq1a |
| omitted Genes: | Sh3pxd2b, Efcab9, Fbxw11 |
| omitted Genes: | Ranbp17, Gabrp, Dock2 |
| omitted Genes: | Fam196b |

Make more plots at http://csg.sph.umich.edu/locuszoom/

Fine−mapping Regions

| snp | chr | pos (Mb) | pp | group | color |
|---|---|---|---|---|---|
| chr11:28851860 | 11 | 28.85186 | 0.073389453 | Credible Set | purple |
| chr11:28853308 | 11 | 28.85331 | 0.064643672 | Credible Set | purple |
| chr11:28855496 | 11 | 28.85550 | 0.068857619 | Credible Set | purple |
| chr11:29365866 | 11 | 29.36587 | 0.105679620 | Credible Set | purple |
| chr11:29419484 | 11 | 29.41948 | 0.083318030 | Credible Set | purple |
| chr11:29419643 | 11 | 29.41964 | 0.106499999 | Credible Set | purple |
| chr11:29486678 | 11 | 29.48668 | 0.239200283 | Credible Set | purple |
| chr11:29623528 | 11 | 29.62353 | 0.032853206 | Credible Set | purple |
| chr11:29623531 | 11 | 29.62353 | 0.032494083 | Credible Set | purple |
| chr11:29623542 | 11 | 29.62354 | 0.032408279 | Credible Set | purple |
| chr11:29637584 | 11 | 29.63758 | 0.004718717 | Credible Set | purple |
| chr11:29637586 | 11 | 29.63759 | 0.005140145 | Credible Set | purple |
| chr11:29676398 | 11 | 29.67640 | 0.081041441 | Credible Set | purple |
| chr11:29757518 | 11 | 29.75752 | 0.005669447 | Credible Set | purple |
| chr11:29767130 | 11 | 29.76713 | 0.010892164 | Credible Set | purple |
| chr11:29794101 | 11 | 29.79410 | 0.003385191 | Credible Set | purple |
| chr11:29794488 | 11 | 29.79449 | 0.002970763 | Credible Set | purple |
| chr11:29794516 | 11 | 29.79452 | 0.002838729 | Credible Set | purple |
| chr11:29815423 | 11 | 29.81542 | 0.003663428 | Credible Set | purple |
| chr11:29815516 | 11 | 29.81552 | 0.003663428 | Credible Set | purple |
| chr11:30005177 | 11 | 30.00518 | 0.003967363 | Credible Set | purple |
| chr11:30018452 | 11 | 30.01845 | 0.003773369 | Credible Set | purple |
| chr11:30143941 | 11 | 30.14394 | 0.003605437 | Credible Set | purple |
| chr11:30144319 | 11 | 30.14432 | 0.003893775 | Credible Set | purple |
| chr11:30144348 | 11 | 30.14435 | 0.003827763 | Credible Set | purple |
| chr11:30144383 | 11 | 30.14438 | 0.003831083 | Credible Set | purple |
| chr11:30149181 | 11 | 30.14918 | 0.003098802 | Credible Set | purple |
| chr11:30161875 | 11 | 30.16187 | 0.003455055 | Credible Set | purple |

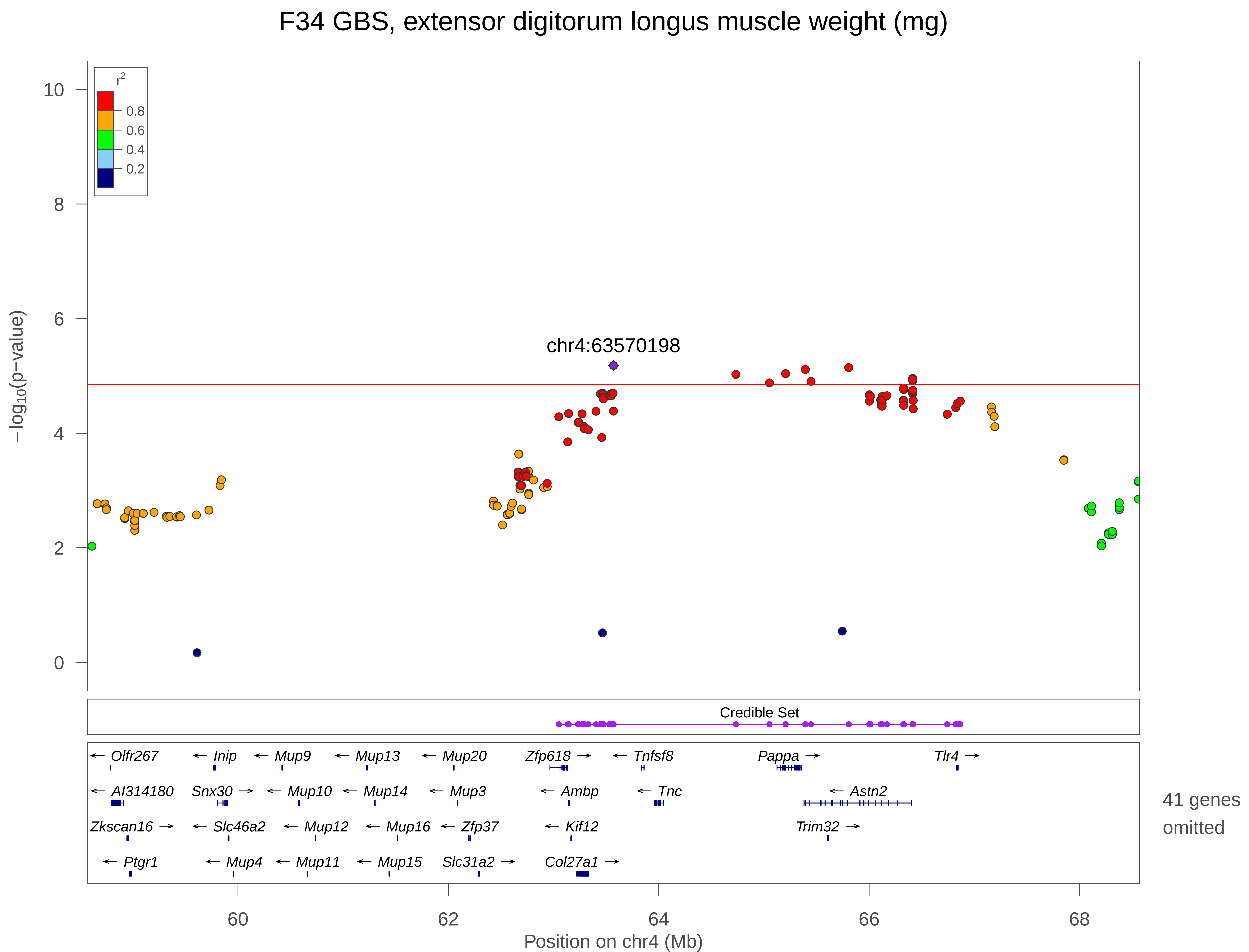
F34 GBS, extensor digitorum longus muscle weight (mg)
r²
0.8
0.6
0.4
0.2
10
8
6
4
2
0
−log10(p−value)
chr4:63570198
Credible Set
← Olfr267
← Inip
← Mup9
← Mup13
← Mup20
Zfp618 →
← Tnfsf8
Pappa →
Tlr4 →
← AI314180
Snx30 →
← Mup10
← Mup14
← Mup3
← Ambp
← Tnc
← Astn2
Zkscan16 →
← Slc46a2
← Mup12
← Mup16
← Zfp37
← Kif12
Trim32 →
← Ptgr1
← Mup4
← Mup11
← Mup15
Slc31a2 →
Col27a1 →
41 genes omitted
60
62
64
66
68
Position on chr4 (Mb)

| | |
|---|---|
| date: | Thu Oct 25 17:20:05 2018 |
| build: | F34GBS |
| display range: | chr4:58570198−68570198 [58570198−68570198] |
| hilite range: | 0 − 0 [ 0 − 0 ] |
| reference SNP: | chr4:63570198 |
| number of SNPs plotted: | 192 |
| min P−value: | 6.58E−6 [chr4:63570198] |
| max P−value: | 6.78E−1 [chr4:59610662] |
| omitted Genes: | Dnajc25, Gng10, Ugcg |
| omitted Genes: | Gm12596, Susd1, Ptbp3 |
| omitted Genes: | Hsdl2, E130308A19Rik, Mup6 |
| omitted Genes: | Mup7, Mup2, Mup8 |
| omitted Genes: | Mup1, Mup17, Mup18 |
| omitted Genes: | Mup19, Mup5, Mup21 |
| omitted Genes: | Fkbp15, Slc31a1, Cdc26 |
| omitted Genes: | Prpf4, Rnf183, Wdr31 |
| omitted Genes: | Bspry, Hdhd3, Alad |
| omitted Genes: | Pole3, 4933430I17Rik, Rgs3 |
| omitted Genes: | Mir455, Orm1, Orm3 |
| omitted Genes: | Orm2, Akna, Whrn |
| omitted Genes: | Atp6v1g1, 6330416G13Rik, Gm11213 |
| omitted Genes: | 1700018C11Rik, Tnfsf15 |

Make more plots at http://csg.sph.umich.edu/locuszoom/

Fine−mapping Regions

| snp | chr | pos (Mb) | pp | group | color |
|---|---|---|---|---|---|
| chr4:63050153 | 4 | 63.05015 | 0.005586469 | Credible Set | purple |
| chr4:63135285 | 4 | 63.13529 | 0.002155219 | Credible Set | purple |
| chr4:63143387 | 4 | 63.14339 | 0.006319390 | Credible Set | purple |
| chr4:63232655 | 4 | 63.23266 | 0.004506028 | Credible Set | purple |
| chr4:63239449 | 4 | 63.23945 | 0.004569768 | Credible Set | purple |
| chr4:63239821 | 4 | 63.23982 | 0.004525341 | Credible Set | purple |
| chr4:63239854 | 4 | 63.23985 | 0.004516917 | Credible Set | purple |
| chr4:63270812 | 4 | 63.27081 | 0.006242967 | Credible Set | purple |
| chr4:63291699 | 4 | 63.29170 | 0.003837657 | Credible Set | purple |
| chr4:63291735 | 4 | 63.29174 | 0.003837138 | Credible Set | purple |
| chr4:63292002 | 4 | 63.29200 | 0.003565385 | Credible Set | purple |
| chr4:63292021 | 4 | 63.29202 | 0.003582722 | Credible Set | purple |
| chr4:63331035 | 4 | 63.33103 | 0.003408827 | Credible Set | purple |
| chr4:63403594 | 4 | 63.40359 | 0.006898749 | Credible Set | purple |
| chr4:63445899 | 4 | 63.44590 | 0.013422221 | Credible Set | purple |
| chr4:63458203 | 4 | 63.45820 | 0.002543254 | Credible Set | purple |
| chr4:63465528 | 4 | 63.46553 | 0.013717511 | Credible Set | purple |
| chr4:63471028 | 4 | 63.47103 | 0.012115126 | Credible Set | purple |
| chr4:63471086 | 4 | 63.47109 | 0.011300514 | Credible Set | purple |
| chr4:63473150 | 4 | 63.47315 | 0.011770552 | Credible Set | purple |
| chr4:63473181 | 4 | 63.47318 | 0.011667117 | Credible Set | purple |
| chr4:63473231 | 4 | 63.47323 | 0.013077005 | Credible Set | purple |
| chr4:63473937 | 4 | 63.47394 | 0.011805191 | Credible Set | purple |
| chr4:63474009 | 4 | 63.47401 | 0.011026218 | Credible Set | purple |
| chr4:63531405 | 4 | 63.53140 | 0.012964723 | Credible Set | purple |
| chr4:63531436 | 4 | 63.53144 | 0.013007659 | Credible Set | purple |
| chr4:63535096 | 4 | 63.53510 | 0.012463435 | Credible Set | purple |
| chr4:63548238 | 4 | 63.54824 | 0.013125578 | Credible Set | purple |
| chr4:63548572 | 4 | 63.54857 | 0.012457840 | Credible Set | purple |
| chr4:63558825 | 4 | 63.55882 | 0.013715376 | Credible Set | purple |
| chr4:63565618 | 4 | 63.56562 | 0.013717511 | Credible Set | purple |
| chr4:63565712 | 4 | 63.56571 | 0.013717511 | Credible Set | purple |
| chr4:63570198 | 4 | 63.57020 | 0.039756613 | Credible Set | purple |
| chr4:63570284 | 4 | 63.57028 | 0.006920540 | Credible Set | purple |
| chr4:64733521 | 4 | 64.73352 | 0.028193873 | Credible Set | purple |
| chr4:65052703 | 4 | 65.05270 | 0.020407037 | Credible Set | purple |

Fine−mapping Regions

| snp | chr | pos (Mb) | pp | group | color |
|---|---|---|---|---|---|
| chr4:65205920 | 4 | 65.20592 | 0.029105020 | Credible Set | purple |
| chr4:65393970 | 4 | 65.39397 | 0.034087638 | Credible Set | purple |
| chr4:65447841 | 4 | 65.44784 | 0.021665351 | Credible Set | purple |
| chr4:65806420 | 4 | 65.80642 | 0.036665107 | Credible Set | purple |
| chr4:66002530 | 4 | 66.00253 | 0.012677452 | Credible Set | purple |
| chr4:66003029 | 4 | 66.00303 | 0.010115198 | Credible Set | purple |
| chr4:66003164 | 4 | 66.00316 | 0.012951976 | Credible Set | purple |
| chr4:66013512 | 4 | 66.01351 | 0.012199736 | Credible Set | purple |
| chr4:66110072 | 4 | 66.11007 | 0.010583302 | Credible Set | purple |
| chr4:66114163 | 4 | 66.11416 | 0.009706855 | Credible Set | purple |
| chr4:66114164 | 4 | 66.11416 | 0.008517105 | Credible Set | purple |
| chr4:66114176 | 4 | 66.11418 | 0.008502830 | Credible Set | purple |
| chr4:66123765 | 4 | 66.12377 | 0.009323297 | Credible Set | purple |
| chr4:66123768 | 4 | 66.12377 | 0.008359871 | Credible Set | purple |
| chr4:66123832 | 4 | 66.12383 | 0.012075522 | Credible Set | purple |
| chr4:66124242 | 4 | 66.12424 | 0.010537474 | Credible Set | purple |
| chr4:66169378 | 4 | 66.16938 | 0.012473269 | Credible Set | purple |
| chr4:66325857 | 4 | 66.32586 | 0.010470415 | Credible Set | purple |
| chr4:66325987 | 4 | 66.32599 | 0.010234159 | Credible Set | purple |
| chr4:66328189 | 4 | 66.32819 | 0.010171039 | Credible Set | purple |
| chr4:66328195 | 4 | 66.32819 | 0.010349660 | Credible Set | purple |
| chr4:66328436 | 4 | 66.32844 | 0.008640016 | Credible Set | purple |
| chr4:66328528 | 4 | 66.32853 | 0.016711908 | Credible Set | purple |
| chr4:66328560 | 4 | 66.32856 | 0.016651954 | Credible Set | purple |
| chr4:66329044 | 4 | 66.32904 | 0.015940153 | Credible Set | purple |
| chr4:66329065 | 4 | 66.32906 | 0.016703717 | Credible Set | purple |
| chr4:66414332 | 4 | 66.41433 | 0.014848635 | Credible Set | purple |
| chr4:66414345 | 4 | 66.41434 | 0.014937879 | Credible Set | purple |
| chr4:66414373 | 4 | 66.41437 | 0.014389080 | Credible Set | purple |
| chr4:66414374 | 4 | 66.41437 | 0.014462506 | Credible Set | purple |
| chr4:66414384 | 4 | 66.41438 | 0.013469720 | Credible Set | purple |
| chr4:66414387 | 4 | 66.41439 | 0.014718109 | Credible Set | purple |
| chr4:66414451 | 4 | 66.41445 | 0.015118442 | Credible Set | purple |
| chr4:66414476 | 4 | 66.41448 | 0.022146038 | Credible Set | purple |
| chr4:66414508 | 4 | 66.41451 | 0.024217351 | Credible Set | purple |
| chr4:66414513 | 4 | 66.41451 | 0.023099784 | Credible Set | purple |

Fine−mapping Regions

| snp | chr | pos (Mb) | pp | group | color |
|---|---|---|---|---|---|
| chr4:66414963 | 4 | 66.41496 | 0.015322361 | Credible Set | purple |
| chr4:66419064 | 4 | 66.41906 | 0.007564370 | Credible Set | purple |
| chr4:66419126 | 4 | 66.41913 | 0.010420861 | Credible Set | purple |
| chr4:66743103 | 4 | 66.74310 | 0.006150139 | Credible Set | purple |
| chr4:66823255 | 4 | 66.82326 | 0.007915557 | Credible Set | purple |
| chr4:66840696 | 4 | 66.84070 | 0.009284521 | Credible Set | purple |
| chr4:66866758 | 4 | 66.86676 | 0.010175646 | Credible Set | purple |

F34 GBS, gastrocnemius muscle weight (mg)

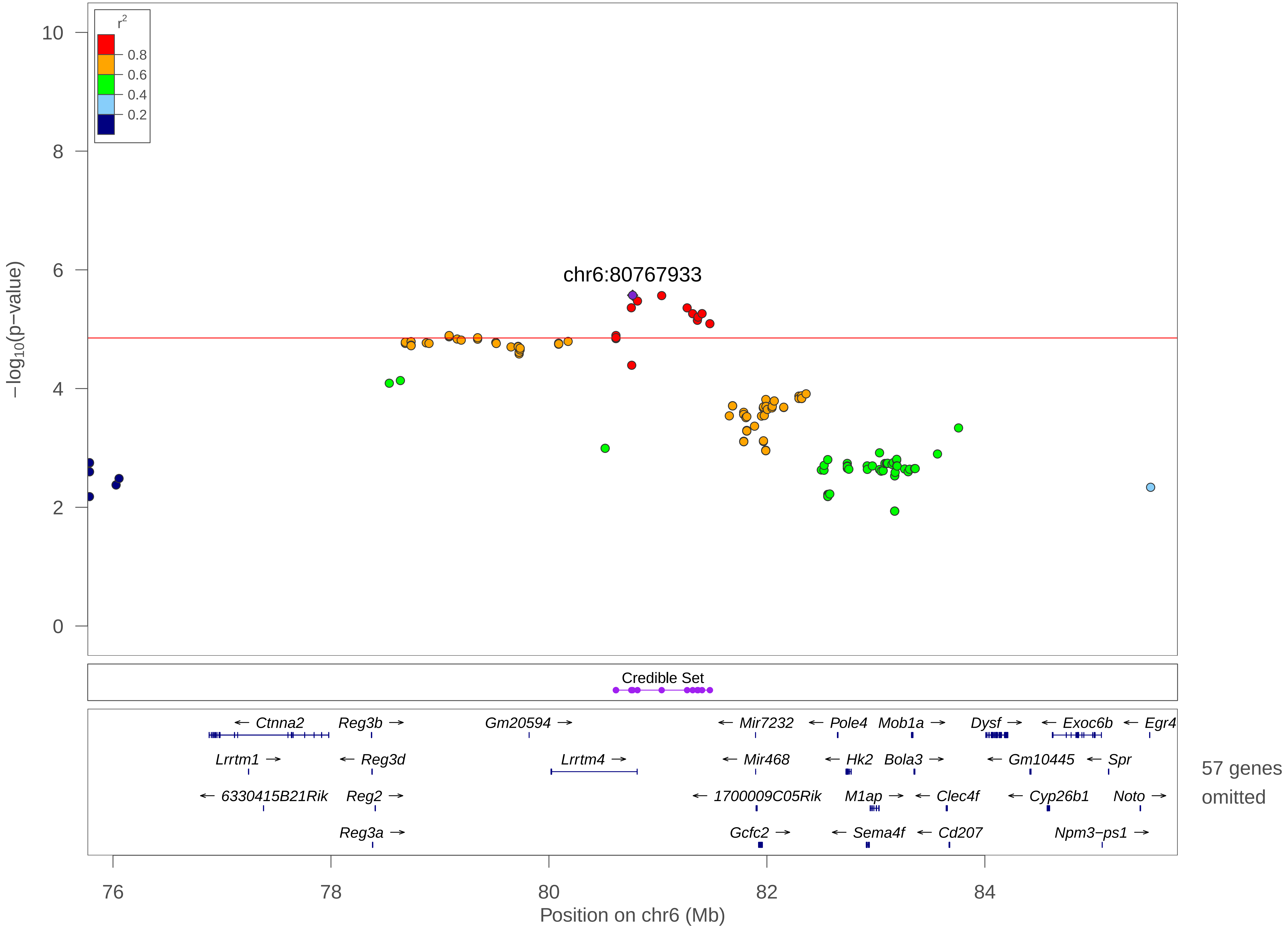

| | |
|---|---|
| date: | Thu Oct 25 17:20:16 2018 |
| build: | F34GBS |
| display range: | chr6:75767933−85767933 [75767933−85767933] |
| hilite range: | 0 − 0 [ 0 − 0 ] |
| reference SNP: | chr6:80767933 |
| number of SNPs plotted: | 156 |
| min P−value: | 2.68E−6 [chr6:80767933] |
| max P−value: | 1.16E−2 [chr6:83172373] |
| omitted Genes: | Reg1, Reg3g, Gm20362 |
| omitted Genes: | Mrpl19, Eva1a, Tacr1 |
| omitted Genes: | 2310069B03Rik, Dok1, Loxl3 |
| omitted Genes: | Mir7040, Htra2, Aup1 |
| omitted Genes: | Dqx1, Tlx2, Pcgf1 |
| omitted Genes: | Lbx2, Mir3470a, Ccdc142 |
| omitted Genes: | Mrpl53, Mogs, Wbp1 |
| omitted Genes: | Ino80b, Rtkn, Wdr54 |
| omitted Genes: | 1700003E16Rik, Dctn1, Slc4a5 |
| omitted Genes: | Mthfd2, Tet3, Mir6374 |
| omitted Genes: | B230319C09Rik, Dguok, Actg2 |
| omitted Genes: | Stambp, Gm21284, Vax2os |
| omitted Genes: | Vax2, Atp6v1b1, 1700124L16Rik |
| omitted Genes: | Ankrd53, Tex261, Gm7443 |
| omitted Genes: | Nagk, Paip2b, Zfp638 |
| omitted Genes: | Gm5878, Emx1, Sfxn5 |
| omitted Genes: | Rab11fip5, Mir705, Smyd5 |
| omitted Genes: | Pradc1, Cct7, Fbxo41 |
| omitted Genes: | Alms1, Nat8f7, Nat8f3 |

Make more plots at http://csg.sph.umich.edu/locuszoom/

Fine−mapping Regions

| snp | chr | pos (Mb) | pp | group | color |
|---|---|---|---|---|---|
| chr6:80614710 | 6 | 80.61471 | 0.03055088 | Credible Set | purple |
| chr6:80614725 | 6 | 80.61473 | 0.02725666 | Credible Set | purple |
| chr6:80614740 | 6 | 80.61474 | 0.02732058 | Credible Set | purple |
| chr6:80614750 | 6 | 80.61475 | 0.02725666 | Credible Set | purple |
| chr6:80614907 | 6 | 80.61491 | 0.02820462 | Credible Set | purple |
| chr6:80755939 | 6 | 80.75594 | 0.08540020 | Credible Set | purple |
| chr6:80759589 | 6 | 80.75959 | 0.01017886 | Credible Set | purple |
| chr6:80767933 | 6 | 80.76793 | 0.13610015 | Credible Set | purple |
| chr6:80812582 | 6 | 80.81258 | 0.10974954 | Credible Set | purple |
| chr6:81034488 | 6 | 81.03449 | 0.13422460 | Credible Set | purple |
| chr6:81267890 | 6 | 81.26789 | 0.08512251 | Credible Set | purple |
| chr6:81318989 | 6 | 81.31899 | 0.06852202 | Credible Set | purple |
| chr6:81362217 | 6 | 81.36222 | 0.05361753 | Credible Set | purple |
| chr6:81368112 | 6 | 81.36811 | 0.06044366 | Credible Set | purple |
| chr6:81405109 | 6 | 81.40511 | 0.06876887 | Credible Set | purple |
| chr6:81476902 | 6 | 81.47690 | 0.04728265 | Credible Set | purple |

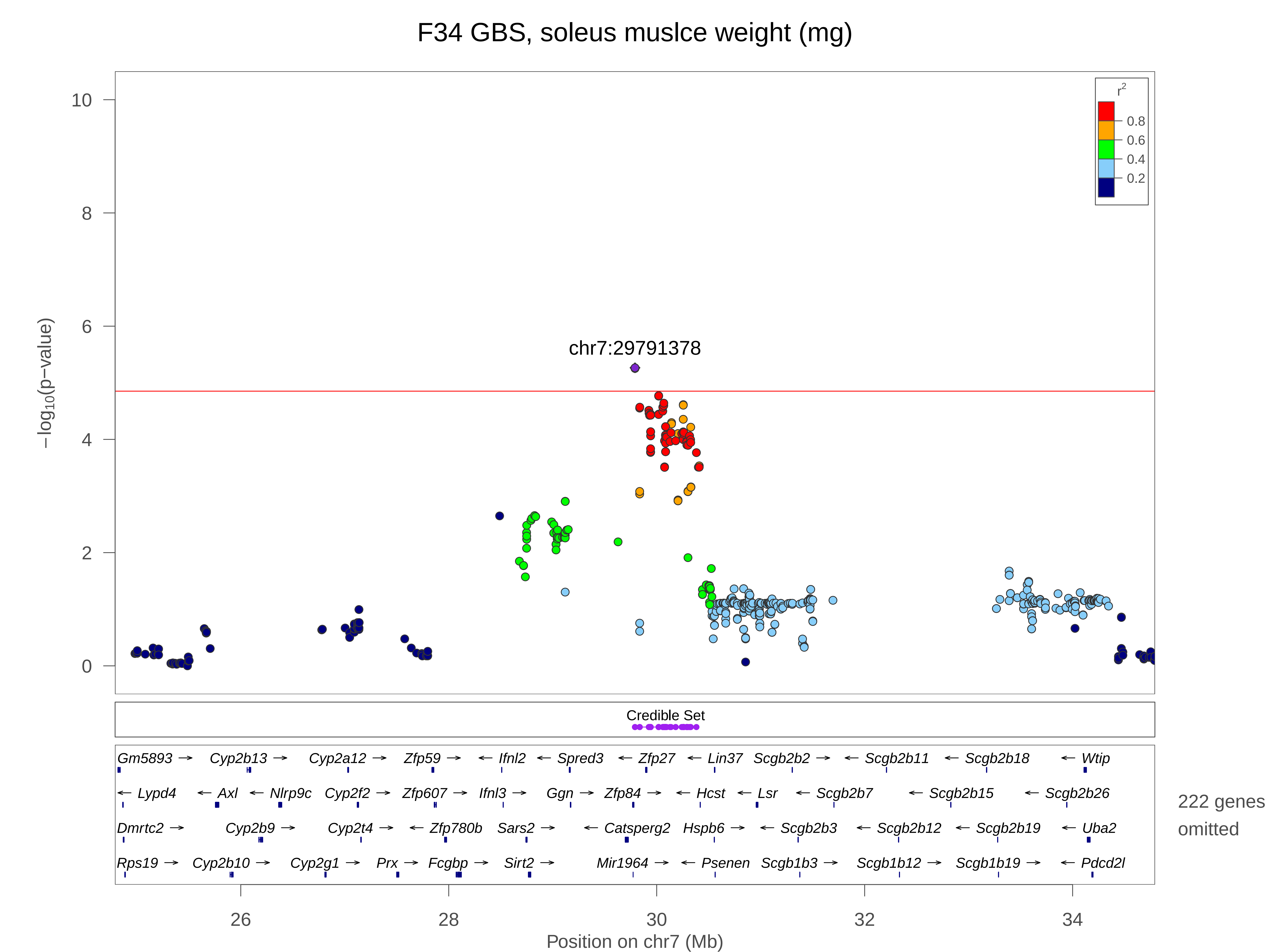

F34 GBS, soleus muslce weight (mg)
r2
0.8
0.6
0.4
0.2
chr7:29791378
−log10(p−value)
10
8
6
4
2
0
Credible Set
Gm5893 →
Cyp2b13 →
Cyp2a12 →
Zfp59 →
← Ifnl2
← Spred3
← Zfp27
← Lin37
Scgb2b2 →
← Scgb2b11
← Scgb2b18
← Wtip
← Lypd4
← Axl
← Nlrp9c
Cyp2f2 →
Zfp607 →
Ifnl3 →
Ggn →
Zfp84 →
← Hcst
← Lsr
← Scgb2b7
← Scgb2b15
← Scgb2b26
Dmrtc2 →
Cyp2b9 →
Cyp2t4 →
← Zfp780b
Sars2 →
← Catsperg2
Hspb6 →
← Scgb2b3
← Scgb2b12
← Scgb2b19
← Uba2
Rps19 →
Cyp2b10 →
Cyp2g1 →
Prx →
Fcgbp →
Sirt2 →
Mir1964 →
← Psenen
Scgb1b3 →
Scgb1b12 →
Scgb1b19 →
← Pdcd2l
222 genes omitted
26
28
30
32
34
Position on chr7 (Mb)

date: Thu Oct 25 17:20:27 2018
build: F34GBS
display range: chr7:24791378−34791378 [24791378−34791378]
hilite range: 0 − 0 [ 0 − 0 ]
reference SNP: chr7:29791378
number of SNPs plotted: 603
min P−value: 5.43E−6 [chr7:29791378]
max P−value: 9.98E−1 [chr7:25487376]
omitted Genes: Cd79a, Arhgef1, Rabac1
omitted Genes: Mir7046, Atp1a3, Mir7047
omitted Genes: Grik5, Zfp574, Pou2f2
omitted Genes: Mir6537, D930028M14Rik, Dedd2
omitted Genes: Mir7048, Zfp526, Gsk3a
omitted Genes: 9130221H12Rik, Erf, Cic
omitted Genes: Pafah1b3, Prr19, Tmem145
omitted Genes: Megf8, Cnfn, 4732471J01Rik
omitted Genes: Lipe, Cxcl17, Ceacam1
omitted Genes: Ceacam2, Erich4, Atp5sl
omitted Genes: B3gnt8, Bckdha, Exosc5
omitted Genes: Tmem91, B9d2, Tgfb1
omitted Genes: Ccdc97, Hnrnpul1, Cyp2s1
omitted Genes: Vmn1r184, Cyp2a4, Nlrp4a
omitted Genes: Nlrp9a, Vmn1r185, Cyp2b23
omitted Genes: Cyp2b19, Cyp2a5, Cyp2a22
omitted Genes: Egln2, Rab4b, Mir3101
omitted Genes: Mia, Snrpa, BC024978
omitted Genes: Mir1191, Itpkc, Adck4
omitted Genes: Numbl, Ltbp4, Shkbp1
omitted Genes: Sptbn4, Blvrb, Sertad3
omitted Genes: Sertad1, Hipk4, Pld3
omitted Genes: 2310022A10Rik, Akt2, Ttc9b
omitted Genes: Map3k10, C030039L03Rik, Zfp60
omitted Genes: Gm10046, Zfp626, 1700049G17Rik
omitted Genes: Zfp850, Psmc4, 9530053A07Rik
omitted Genes: Fbl, Dyrk1b, Eid2
omitted Genes: Eid2b, BC089491, Dll3
omitted Genes: Timm50, Supt5, Rps16
omitted Genes: AF357399, Plekhg2, Zfp36
omitted Genes: Med29, Paf1, Samd4b
Make more plots at http://csg.sph.umich.edu/locuszoom/
omitted Genes: Gmfg, Lrfn1, 1700028B04Rik

Fine−mapping Regions

| snp | chr | pos (Mb) | pp | group | color |
|---|---|---|---|---|---|
| chr7:29791322 | 7 | 29.79132 | 0.098275664 | Credible Set | purple |
| chr7:29791378 | 7 | 29.79138 | 0.100896825 | Credible Set | purple |
| chr7:29835954 | 7 | 29.83595 | 0.021094227 | Credible Set | purple |
| chr7:29837224 | 7 | 29.83722 | 0.021897496 | Credible Set | purple |
| chr7:29924583 | 7 | 29.92458 | 0.019392260 | Credible Set | purple |
| chr7:29924600 | 7 | 29.92460 | 0.019302108 | Credible Set | purple |
| chr7:29926689 | 7 | 29.92669 | 0.017338965 | Credible Set | purple |
| chr7:29931632 | 7 | 29.93163 | 0.015945916 | Credible Set | purple |
| chr7:29941228 | 7 | 29.94123 | 0.003838486 | Credible Set | purple |
| chr7:29941252 | 7 | 29.94125 | 0.003855683 | Credible Set | purple |
| chr7:29941259 | 7 | 29.94126 | 0.003905284 | Credible Set | purple |
| chr7:29941272 | 7 | 29.94127 | 0.003932245 | Credible Set | purple |
| chr7:29941287 | 7 | 29.94129 | 0.004422145 | Credible Set | purple |
| chr7:29941313 | 7 | 29.94131 | 0.016699891 | Credible Set | purple |
| chr7:29941361 | 7 | 29.94136 | 0.016476051 | Credible Set | purple |
| chr7:29941644 | 7 | 29.94164 | 0.016701635 | Credible Set | purple |
| chr7:29941657 | 7 | 29.94166 | 0.016024127 | Credible Set | purple |
| chr7:29941708 | 7 | 29.94171 | 0.008449947 | Credible Set | purple |
| chr7:29941716 | 7 | 29.94172 | 0.007239586 | Credible Set | purple |
| chr7:29941717 | 7 | 29.94172 | 0.008441506 | Credible Set | purple |
| chr7:29941765 | 7 | 29.94177 | 0.008502193 | Credible Set | purple |
| chr7:30018478 | 7 | 30.01848 | 0.016709972 | Credible Set | purple |
| chr7:30018532 | 7 | 30.01853 | 0.016455486 | Credible Set | purple |
| chr7:30018938 | 7 | 30.01894 | 0.034193123 | Credible Set | purple |
| chr7:30018947 | 7 | 30.01895 | 0.033532453 | Credible Set | purple |
| chr7:30018955 | 7 | 30.01895 | 0.033964231 | Credible Set | purple |
| chr7:30058592 | 7 | 30.05859 | 0.018773627 | Credible Set | purple |
| chr7:30058599 | 7 | 30.05860 | 0.021892512 | Credible Set | purple |
| chr7:30067674 | 7 | 30.06767 | 0.023566323 | Credible Set | purple |
| chr7:30067701 | 7 | 30.06770 | 0.023247881 | Credible Set | purple |
| chr7:30068385 | 7 | 30.06838 | 0.025573745 | Credible Set | purple |
| chr7:30075552 | 7 | 30.07555 | 0.005983281 | Credible Set | purple |
| chr7:30075596 | 7 | 30.07560 | 0.005980927 | Credible Set | purple |
| chr7:30084581 | 7 | 30.08458 | 0.007492523 | Credible Set | purple |
| chr7:30084617 | 7 | 30.08462 | 0.007377294 | Credible Set | purple |
| chr7:30085368 | 7 | 30.08537 | 0.010325017 | Credible Set | purple |

Fine−mapping Regions

| snp | chr | pos (Mb) | pp | group | color |
|---|---|---|---|---|---|
| chr7:30085376 | 7 | 30.08538 | 0.010325017 | Credible Set | purple |
| chr7:30085378 | 7 | 30.08538 | 0.010325017 | Credible Set | purple |
| chr7:30085382 | 7 | 30.08538 | 0.010325017 | Credible Set | purple |
| chr7:30085386 | 7 | 30.08539 | 0.010325017 | Credible Set | purple |
| chr7:30085398 | 7 | 30.08540 | 0.010325017 | Credible Set | purple |
| chr7:30085403 | 7 | 30.08540 | 0.010325017 | Credible Set | purple |
| chr7:30085427 | 7 | 30.08543 | 0.010325017 | Credible Set | purple |
| chr7:30085431 | 7 | 30.08543 | 0.010325017 | Credible Set | purple |
| chr7:30085517 | 7 | 30.08552 | 0.003945093 | Credible Set | purple |
| chr7:30085526 | 7 | 30.08553 | 0.003945093 | Credible Set | purple |
| chr7:30085548 | 7 | 30.08555 | 0.003945093 | Credible Set | purple |
| chr7:30085558 | 7 | 30.08556 | 0.003945093 | Credible Set | purple |
| chr7:30085567 | 7 | 30.08557 | 0.003935726 | Credible Set | purple |
| chr7:30085636 | 7 | 30.08564 | 0.006910810 | Credible Set | purple |
| chr7:30086094 | 7 | 30.08609 | 0.005463718 | Credible Set | purple |
| chr7:30095828 | 7 | 30.09583 | 0.006909874 | Credible Set | purple |
| chr7:30127954 | 7 | 30.12795 | 0.005821576 | Credible Set | purple |
| chr7:30137791 | 7 | 30.13779 | 0.008168606 | Credible Set | purple |
| chr7:30182363 | 7 | 30.18236 | 0.005985035 | Credible Set | purple |
| chr7:30182366 | 7 | 30.18237 | 0.006001099 | Credible Set | purple |
| chr7:30240571 | 7 | 30.24057 | 0.007864599 | Credible Set | purple |
| chr7:30255201 | 7 | 30.25520 | 0.008194478 | Credible Set | purple |
| chr7:30255202 | 7 | 30.25520 | 0.008162407 | Credible Set | purple |
| chr7:30255242 | 7 | 30.25524 | 0.008443790 | Credible Set | purple |
| chr7:30256110 | 7 | 30.25611 | 0.006266192 | Credible Set | purple |
| chr7:30262031 | 7 | 30.26203 | 0.007729693 | Credible Set | purple |
| chr7:30262034 | 7 | 30.26203 | 0.008253511 | Credible Set | purple |
| chr7:30289075 | 7 | 30.28908 | 0.005825148 | Credible Set | purple |
| chr7:30289086 | 7 | 30.28909 | 0.005893662 | Credible Set | purple |
| chr7:30289112 | 7 | 30.28911 | 0.005605923 | Credible Set | purple |
| chr7:30289270 | 7 | 30.28927 | 0.005751710 | Credible Set | purple |
| chr7:30289273 | 7 | 30.28927 | 0.005736705 | Credible Set | purple |
| chr7:30289902 | 7 | 30.28990 | 0.005662450 | Credible Set | purple |
| chr7:30289956 | 7 | 30.28996 | 0.005716106 | Credible Set | purple |
| chr7:30290355 | 7 | 30.29036 | 0.005092684 | Credible Set | purple |
| chr7:30301382 | 7 | 30.30138 | 0.005027048 | Credible Set | purple |

Fine−mapping Regions

| snp | chr | pos (Mb) | pp | group | color |
|---|---|---|---|---|---|
| chr7:30316542 | 7 | 30.31654 | 0.007256893 | Credible Set | purple |
| chr7:30326035 | 7 | 30.32604 | 0.006264187 | Credible Set | purple |
| chr7:30327435 | 7 | 30.32744 | 0.005610562 | Credible Set | purple |
| chr7:30381850 | 7 | 30.38185 | 0.003805361 | Credible Set | purple |

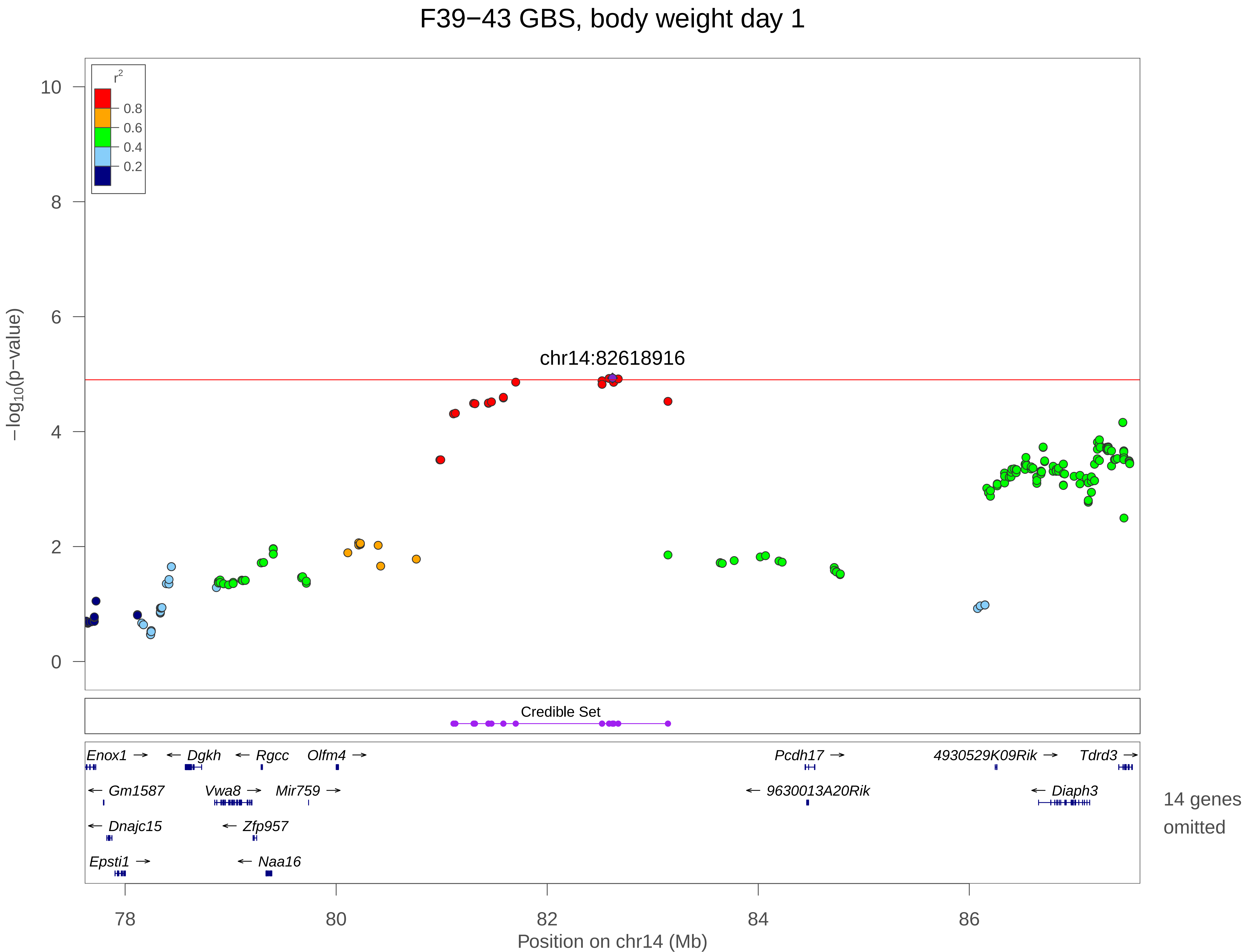
F39−43 GBS, body weight day 1
r2
0.8
0.6
0.4
0.2
10
8
6
4
2
0
−log10(p−value)
chr14:82618916
Credible Set
Enox1 →
← Dgkh
← Rgcc
Olfm4 →
Pcdh17 →
4930529K09Rik →
Tdrd3 →
← Gm1587
Vwa8 →
Mir759 →
← 9630013A20Rik
← Diaph3
← Dnajc15
← Zfp957
Epsti1 →
← Naa16
14 genes
omitted
78
80
82
84
86
Position on chr14 (Mb)

| | |
|---|---|
| date: | Thu Oct 25 17:20:37 2018 |
| build: | F3943GBS |
| display range: | chr14:77618916−87618916 [77618916−87618916] |
| hilite range: | 0 − 0 [ 0 − 0 ] |
| reference SNP: | chr14:82618916 |
| number of SNPs plotted: | 260 |
| min P−value: | 1.17E−5 [chr14:82618916] |
| max P−value: | 3.42E−1 [chr14:78241501] |
| omitted Genes: | Fam216b, Mir1971, Tnfsf11 |
| omitted Genes: | Akap11, Mtrf1, Kbtbd7 |
| omitted Genes: | Kbtbd6, Wbp4, Elf1 |
| omitted Genes: | Sugt1, Lect1, 4930452G13Rik |
| omitted Genes: | Pcdh8, Gm10845 |

Make more plots at http://csg.sph.umich.edu/locuszoom/

Fine−mapping Regions

| snp | chr | pos (Mb) | pp | group | color |
|---|---|---|---|---|---|
| chr14:81112053 | 14 | 81.11205 | 0.02402823 | Credible Set | purple |
| chr14:81129931 | 14 | 81.12993 | 0.02462348 | Credible Set | purple |
| chr14:81302522 | 14 | 81.30252 | 0.03584985 | Credible Set | purple |
| chr14:81315250 | 14 | 81.31525 | 0.03533937 | Credible Set | purple |
| chr14:81443053 | 14 | 81.44305 | 0.03593258 | Credible Set | purple |
| chr14:81443088 | 14 | 81.44309 | 0.03604954 | Credible Set | purple |
| chr14:81443103 | 14 | 81.44310 | 0.03636872 | Credible Set | purple |
| chr14:81471656 | 14 | 81.47166 | 0.03796048 | Credible Set | purple |
| chr14:81584557 | 14 | 81.58456 | 0.04401372 | Credible Set | purple |
| chr14:81584584 | 14 | 81.58458 | 0.04518519 | Credible Set | purple |
| chr14:81701500 | 14 | 81.70150 | 0.08054753 | Credible Set | purple |
| chr14:82519671 | 14 | 82.51967 | 0.08492962 | Credible Set | purple |
| chr14:82519834 | 14 | 82.51983 | 0.07406665 | Credible Set | purple |
| chr14:82586326 | 14 | 82.58633 | 0.09274759 | Credible Set | purple |
| chr14:82618916 | 14 | 82.61892 | 0.09409563 | Credible Set | purple |
| chr14:82629334 | 14 | 82.62933 | 0.08000369 | Credible Set | purple |
| chr14:82672838 | 14 | 82.67284 | 0.09111274 | Credible Set | purple |
| chr14:83144282 | 14 | 83.14428 | 0.03866860 | Credible Set | purple |

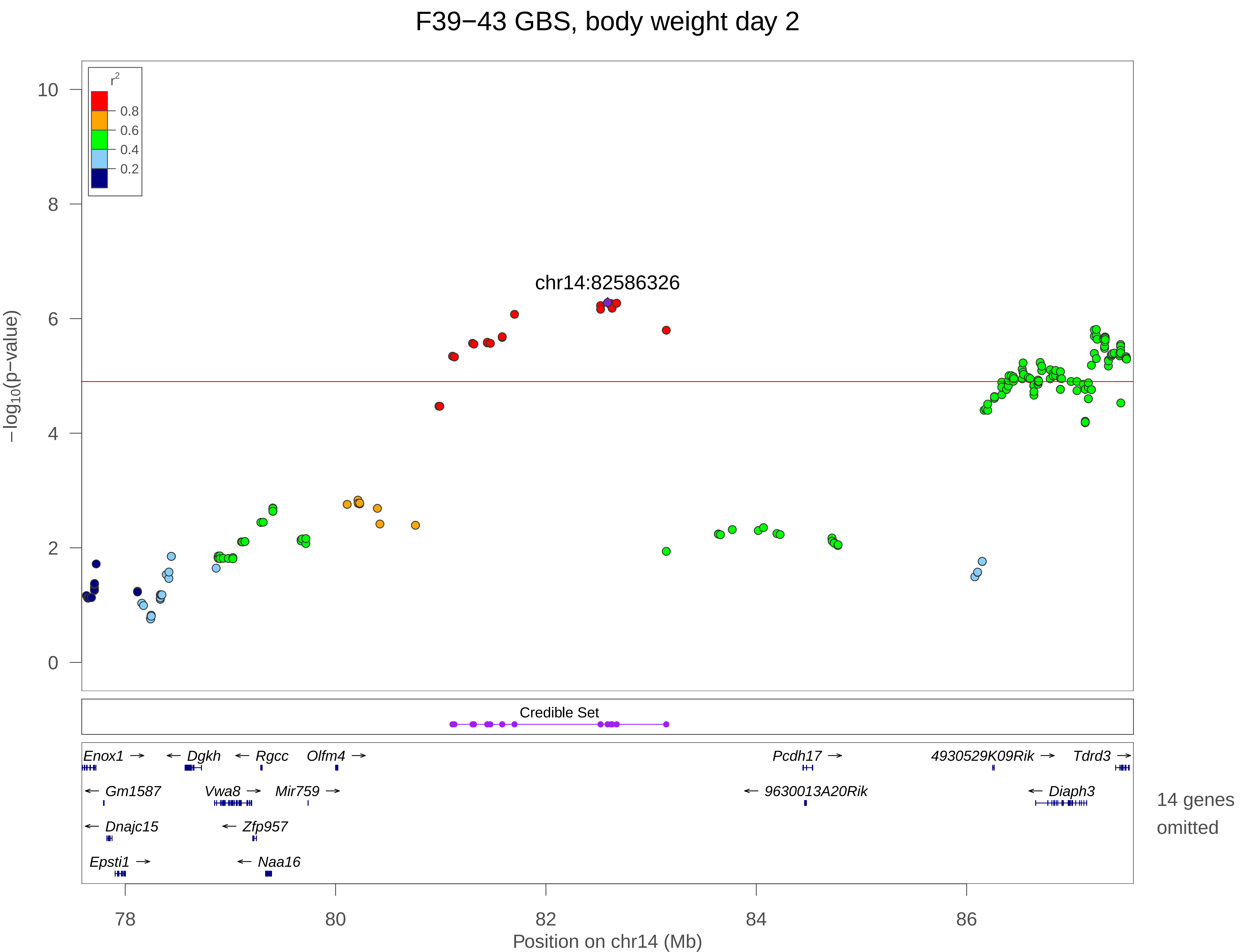
F39−43 GBS, body weight day 2
r2
0.8
0.6
0.4
0.2
10
8
6
4
2
0
−log10(p−value)
chr14:82586326
Credible Set
Enox1 →
← Dgkh
← Rgcc
Olfm4 →
Pcdh17 →
4930529K09Rik →
Tdrd3 →
← Gm1587
Vwa8 →
Mir759 →
← 9630013A20Rik
← Diaph3
← Dnajc15
← Zfp957
Epsti1 →
← Naa16
14 genes omitted
78
80
82
84
86
Position on chr14 (Mb)

| | |
|---|---|
| date: | Thu Oct 25 17:20:44 2018 |
| build: | F3943GBS |
| display range: | chr14:77586326−87586326 [77586326−87586326] |
| hilite range: | 0 − 0 [ 0 − 0 ] |
| reference SNP: | chr14:82586326 |
| number of SNPs plotted: | 260 |
| min P−value: | 5.25E−7 [chr14:82586326] |
| max P−value: | 1.74E−1 [chr14:78241501] |
| omitted Genes: | Fam216b, Mir1971, Tnfsf11 |
| omitted Genes: | Akap11, Mtrf1, Kbtbd7 |
| omitted Genes: | Kbtbd6, Wbp4, Elf1 |
| omitted Genes: | Sugt1, Lect1, 4930452G13Rik |
| omitted Genes: | Pcdh8, Gm10845 |

Make more plots at http://csg.sph.umich.edu/locuszoom/

Fine−mapping Regions

| snp | chr | pos (Mb) | pp | group | color |
|---|---|---|---|---|---|
| chr14:81112053 | 14 | 81.11205 | 0.01490137 | Credible Set | purple |
| chr14:81129931 | 14 | 81.12993 | 0.01447357 | Credible Set | purple |
| chr14:81302522 | 14 | 81.30252 | 0.02452853 | Credible Set | purple |
| chr14:81315250 | 14 | 81.31525 | 0.02373874 | Credible Set | purple |
| chr14:81443053 | 14 | 81.44305 | 0.02493733 | Credible Set | purple |
| chr14:81443088 | 14 | 81.44309 | 0.02516714 | Credible Set | purple |
| chr14:81443103 | 14 | 81.44310 | 0.02551083 | Credible Set | purple |
| chr14:81471656 | 14 | 81.47166 | 0.02445477 | Credible Set | purple |
| chr14:81584557 | 14 | 81.58456 | 0.03075131 | Credible Set | purple |
| chr14:81584584 | 14 | 81.58458 | 0.03163767 | Credible Set | purple |
| chr14:81701500 | 14 | 81.70150 | 0.07512201 | Credible Set | purple |
| chr14:82519671 | 14 | 82.51967 | 0.10542503 | Credible Set | purple |
| chr14:82519834 | 14 | 82.51983 | 0.09131098 | Credible Set | purple |
| chr14:82586326 | 14 | 82.58633 | 0.11823370 | Credible Set | purple |
| chr14:82618916 | 14 | 82.61892 | 0.11422741 | Credible Set | purple |
| chr14:82629334 | 14 | 82.62933 | 0.09506362 | Credible Set | purple |
| chr14:82672838 | 14 | 82.67284 | 0.11550022 | Credible Set | purple |
| chr14:83144282 | 14 | 83.14428 | 0.04063550 | Credible Set | purple |

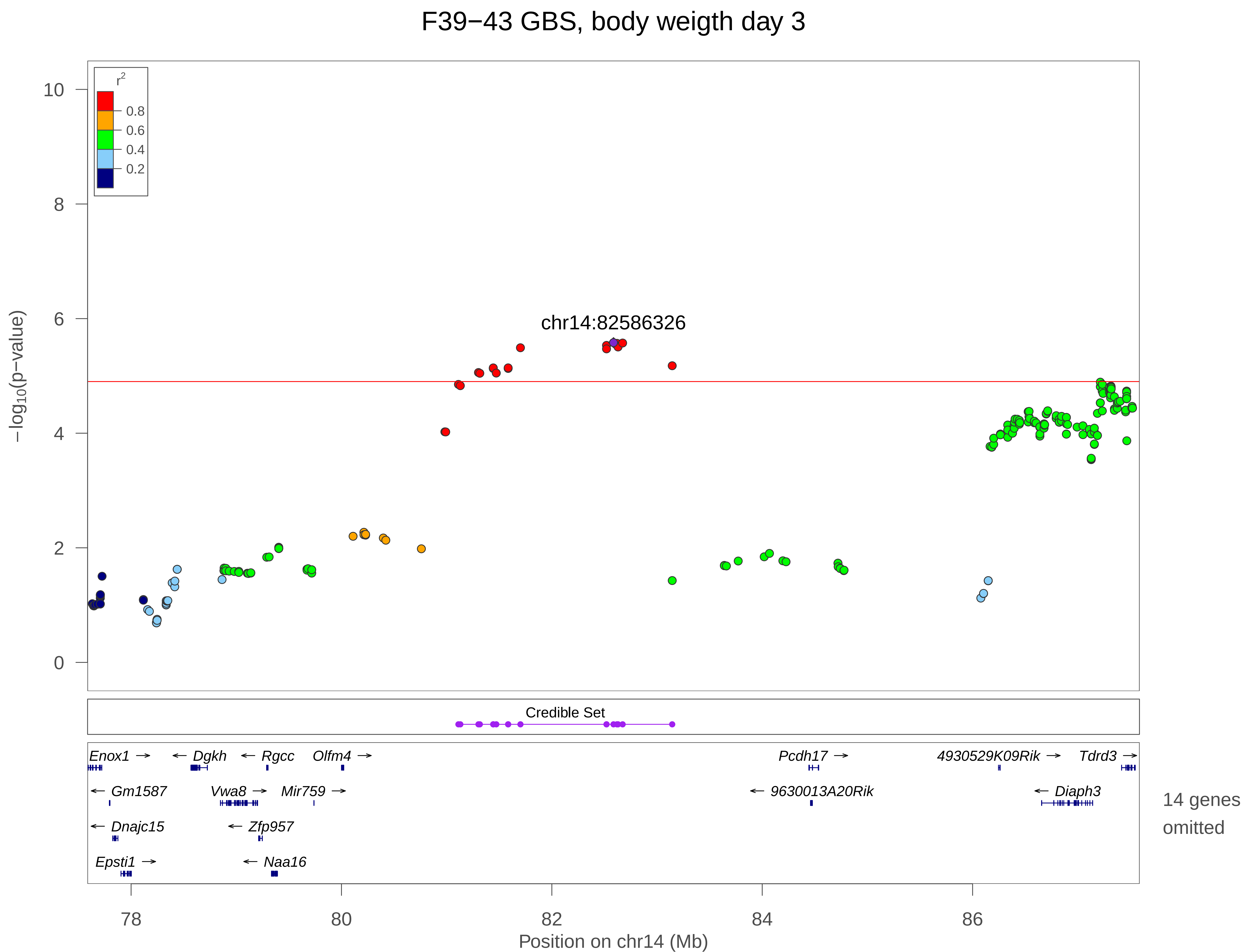

F39−43 GBS, body weigth day 3
r2
0.8
0.6
0.4
0.2
10
8
6
4
2
0
−log10(p−value)
chr14:82586326
Credible Set
Enox1 →
← Dgkh
← Rgcc
Olfm4 →
Pcdh17 →
4930529K09Rik →
Tdrd3 →
← Gm1587
Vwa8 →
Mir759 →
← 9630013A20Rik
← Diaph3
← Dnajc15
← Zfp957
Epsti1 →
← Naa16
14 genes omitted
78
80
82
84
86
Position on chr14 (Mb)

| | |
|---|---|
| date: | Thu Oct 25 17:20:51 2018 |
| build: | F3943GBS |
| display range: | chr14:77586326−87586326 [77586326−87586326] |
| hilite range: | 0 − 0 [ 0 − 0 ] |
| reference SNP: | chr14:82586326 |
| number of SNPs plotted: | 260 |
| min P−value: | 2.63E−6 [chr14:82586326] |
| max P−value: | 2.04E−1 [chr14:78241501] |
| omitted Genes: | Fam216b, Mir1971, Tnfsf11 |
| omitted Genes: | Akap11, Mtrf1, Kbtbd7 |
| omitted Genes: | Kbtbd6, Wbp4, Elf1 |
| omitted Genes: | Sugt1, Lect1, 4930452G13Rik |
| omitted Genes: | Pcdh8, Gm10845 |

Make more plots at http://csg.sph.umich.edu/locuszoom/

Fine−mapping Regions

| snp | chr | pos (Mb) | pp | group | color |
|---|---|---|---|---|---|
| chr14:81112053 | 14 | 81.11205 | 0.02010840 | Credible Set | purple |
| chr14:81129931 | 14 | 81.12993 | 0.01914174 | Credible Set | purple |
| chr14:81302522 | 14 | 81.30252 | 0.03170730 | Credible Set | purple |
| chr14:81315250 | 14 | 81.31525 | 0.03079619 | Credible Set | purple |
| chr14:81443053 | 14 | 81.44305 | 0.03699838 | Credible Set | purple |
| chr14:81443088 | 14 | 81.44309 | 0.03754149 | Credible Set | purple |
| chr14:81443103 | 14 | 81.44310 | 0.03788507 | Credible Set | purple |
| chr14:81471656 | 14 | 81.47166 | 0.03103994 | Credible Set | purple |
| chr14:81584557 | 14 | 81.58456 | 0.03707361 | Credible Set | purple |
| chr14:81584584 | 14 | 81.58458 | 0.03792188 | Credible Set | purple |
| chr14:81701500 | 14 | 81.70150 | 0.08223439 | Credible Set | purple |
| chr14:82519671 | 14 | 82.51967 | 0.09010403 | Credible Set | purple |
| chr14:82519834 | 14 | 82.51983 | 0.07874841 | Credible Set | purple |
| chr14:82586326 | 14 | 82.58633 | 0.10014396 | Credible Set | purple |
| chr14:82618916 | 14 | 82.61892 | 0.09747427 | Credible Set | purple |
| chr14:82629334 | 14 | 82.62933 | 0.08469180 | Credible Set | purple |
| chr14:82672838 | 14 | 82.67284 | 0.09867105 | Credible Set | purple |
| chr14:83144282 | 14 | 83.14428 | 0.04114562 | Credible Set | purple |

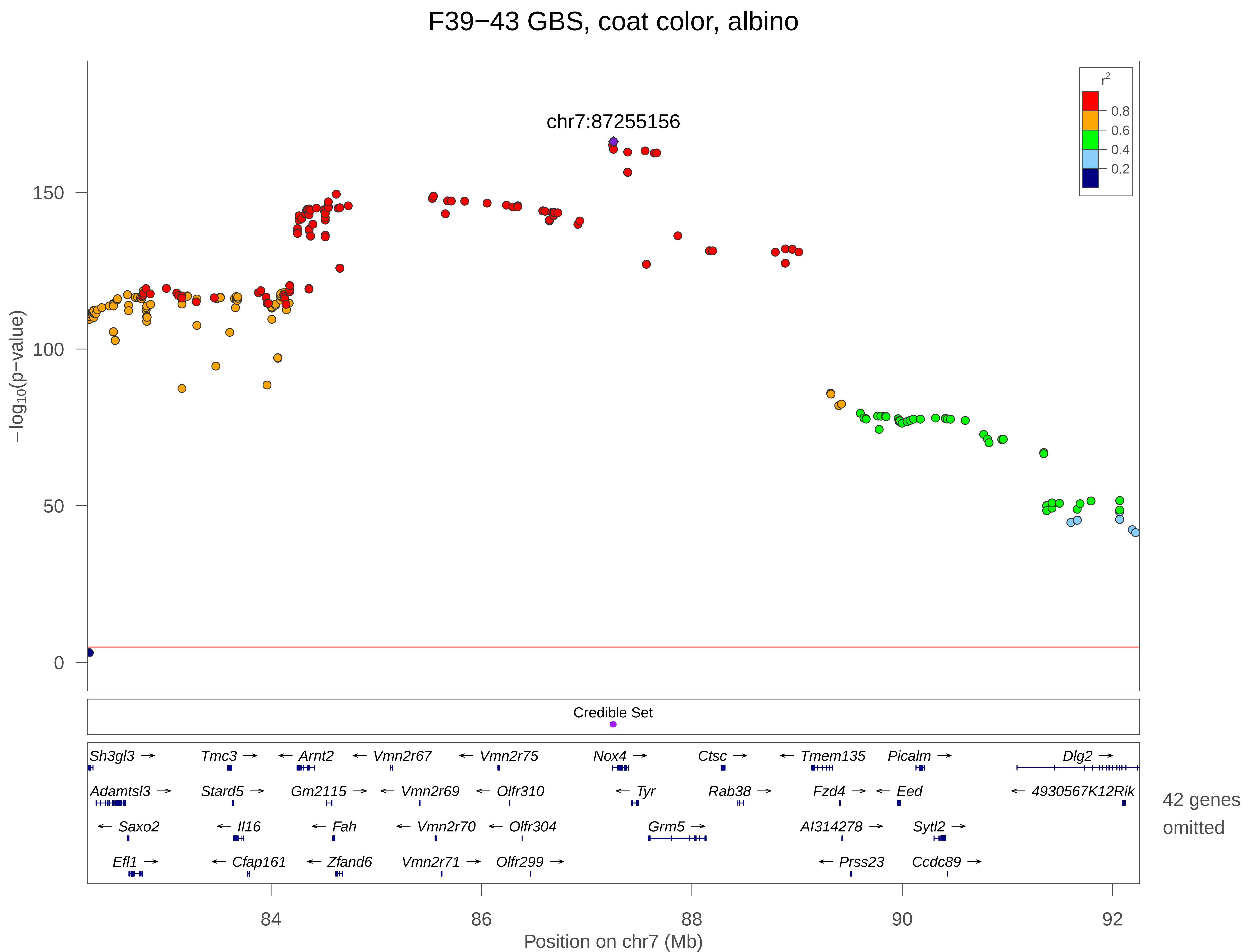

F39−43 GBS, coat color, albino
chr7:87255156
r²
0.8
0.6
0.4
0.2
150
100
50
0
−log₁₀(p−value)
Credible Set
Sh3gl3 →
Tmc3 →
← Arnt2
← Vmn2r67
← Vmn2r75
Nox4 →
Ctsc →
← Tmem135
Picalm →
Dlg2 →
Adamtsl3 →
Stard5 →
Gm2115 →
← Vmn2r69
← Olfr310
← Tyr
Rab38 →
Fzd4 →
← Eed
← 4930567K12Rik
← Saxo2
← Il16
← Fah
← Vmn2r70
← Olfr304
Grm5 →
AI314278 →
Sytl2 →
Efl1 →
← Cfap161
← Zfand6
Vmn2r71 →
Olfr299 →
← Prss23
Ccdc89 →
42 genes omitted
84
86
88
90
92
Position on chr7 (Mb)

| | |
|---|---|
| date: | Thu Oct 25 17:21:12 2018 |
| build: | F3943GBS |
| display range: | chr7:82255156−92255156 [82255156−92255156] |
| hilite range: | 0 − 0 [ 0 − 0 ] |
| reference SNP: | chr7:87255156 |
| number of SNPs plotted: | 313 |
| min P−value: | 5.91E−167 [chr7:87255156] |
| max P−value: | 7.79E−4 [chr7:82271732] |
| omitted Genes: | 4933406J10Rik, Mex3b, 4933430H16Rik |
| omitted Genes: | Mesdc1, Mesdc2, Cemip |
| omitted Genes: | Abhd17c, 2610206C17Rik, Olfr291 |
| omitted Genes: | Olfr290, Vmn2r65, Vmn2r66 |
| omitted Genes: | Vmn2r68, Vmn2r72, Vmn2r73 |
| omitted Genes: | Vmn2r74, Vmn2r76, Olfr309 |
| omitted Genes: | Olfr308, Olfr307, Olfr305 |
| omitted Genes: | Olfr303, Olfr301, Olfr298 |
| omitted Genes: | Olfr297, Olfr295, Olfr294 |
| omitted Genes: | Olfr293, Olfr292, Folh1 |
| omitted Genes: | Vmn2r77, Vmn2r78, Vmn2r79 |
| omitted Genes: | Me3, Ccdc81, l7Rn6 |
| omitted Genes: | E230029C05Rik, 2310010J17Rik, Ccdc83 |
| omitted Genes: | Crebzf, Tmem126a, Tmem126b |

Make more plots at http://csg.sph.umich.edu/locuszoom/

Fine−mapping Regions

| snp | chr | pos (Mb) | pp | group | color |
|---|---|---|---|---|---|
| chr7:87246786 | 7 | 87.24679 | 0.07055338 | Credible Set | purple |
| chr7:87255156 | 7 | 87.25516 | 0.92467500 | Credible Set | purple |

F39−43 GBS, coat color, agouti

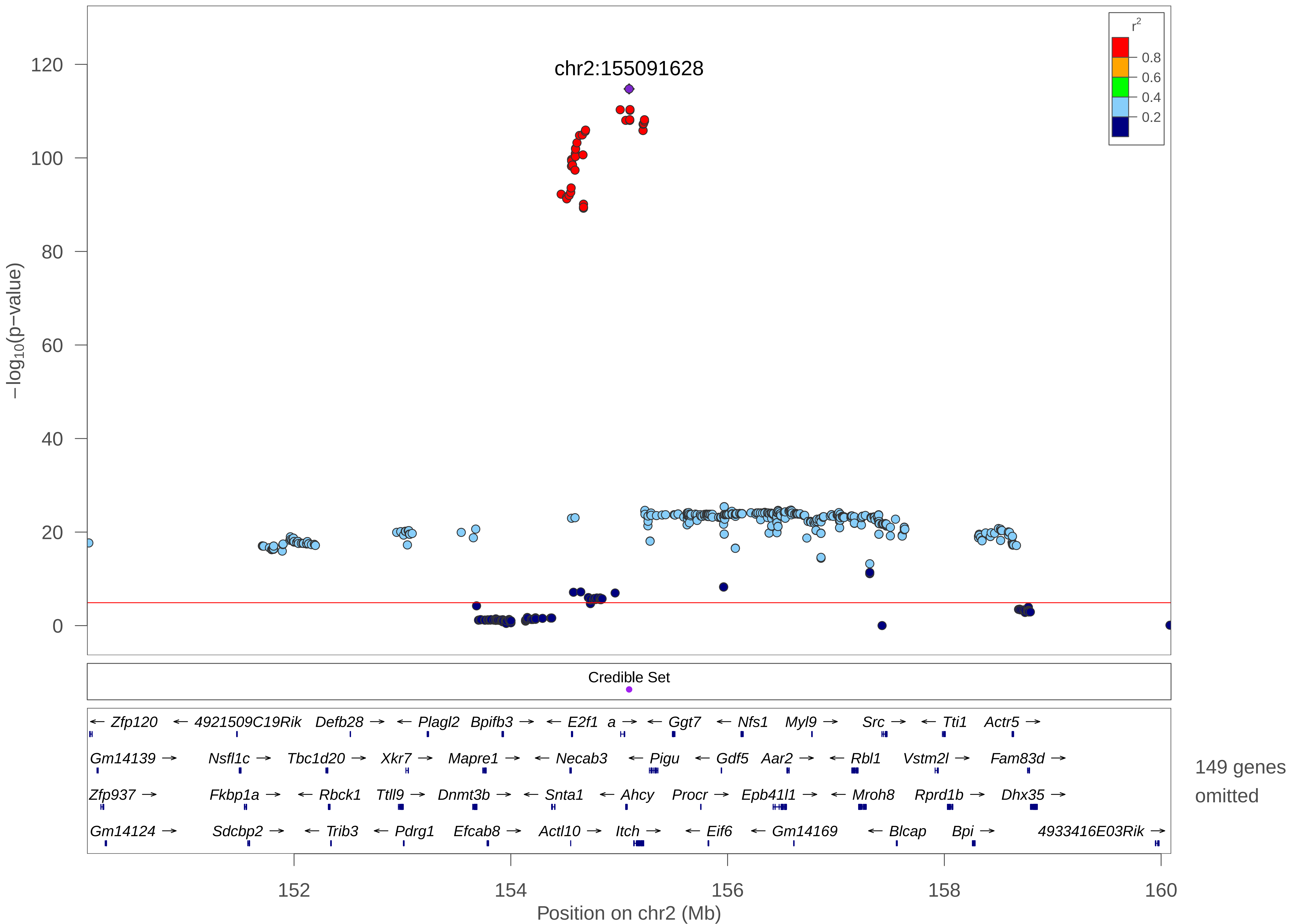

date: Thu Oct 25 17:21:58 2018

build: F3943GBS

display range: chr2:150091628−160091628 [150091628−160091628]

hilite range: 0 − 0 [ 0 − 0 ]

reference SNP: chr2:155091628

number of SNPs plotted: 758

min P−value: 1.78E−115 [chr2:155091628]

max P−value: 9.64E−1 [chr2:157425605]

omitted Genes: 3300002I08Rik, Zfp442, Zfp345

omitted Genes: Cst7, Apmap, Acss1

omitted Genes: E130215H24Rik, Vsx1, Entpd6

omitted Genes: Pygb, Abhd12, Gins1

omitted Genes: Ninl, Nanp, Gm14151

omitted Genes: 4930442J19Rik, LOC100125594, Snph

omitted Genes: Rad21l, Tmem74b, Psmf1

omitted Genes: Rspo4, Angpt4, Mir1953

omitted Genes: Fam110a, Slc52a3, Scrt2

omitted Genes: Srxn1, Tcf15, Csnk2a1

omitted Genes: Gm14164, Nrsn2, Sox12

omitted Genes: Zcchc3, 6820408C15Rik, Defb23

omitted Genes: Defb20, Defb22, Defb26

omitted Genes: Defb29, Defb21, Defb19

omitted Genes: Defb45, Defb36, Defb25

omitted Genes: Rem1, H13, Mcts2

omitted Genes: Id1, Cox4i2, Bcl2l1

omitted Genes: Tpx2, Mir5622, Mylk2

omitted Genes: Foxs1, Dusp15, Ccm2l

omitted Genes: Hck, Tm9sf4, Tspyl3

omitted Genes: Pofut1, Kif3b, 2500004C02Rik

omitted Genes: Asxl1, Nol4l, Commd7

omitted Genes: Sun5, Bpifb2, Bpifb6

omitted Genes: Bpifb4, Bpifa6, Bpifa2

omitted Genes: Bpifa3, Bpifa1, Bpifa5

omitted Genes: Bpifb1, Bpifb5, Bpifb9a

omitted Genes: Bpifb9b, Cdk5rap1, Cbfa2t2

omitted Genes: 1700003F12Rik, Pxmp4, Zfp341

omitted Genes: Chmp4b, Raly, Eif2s2

omitted Genes: 2310005A03Rik, Dynlrb1, Map1lc3a

omitted Genes: Mir695, Trp53inp2, Ncoa6

Make more plots at http://csg.sph.umich.edu/locuszoom/

omitted Genes: Acss2, Acss2os, Gss

Fine−mapping Regions

| snp | chr | pos (Mb) | pp | group | color |
|---|---|---|---|---|---|
| chr2:155091628 | 2 | 155.0916 | 0.9998944 | Credible Set | purple |

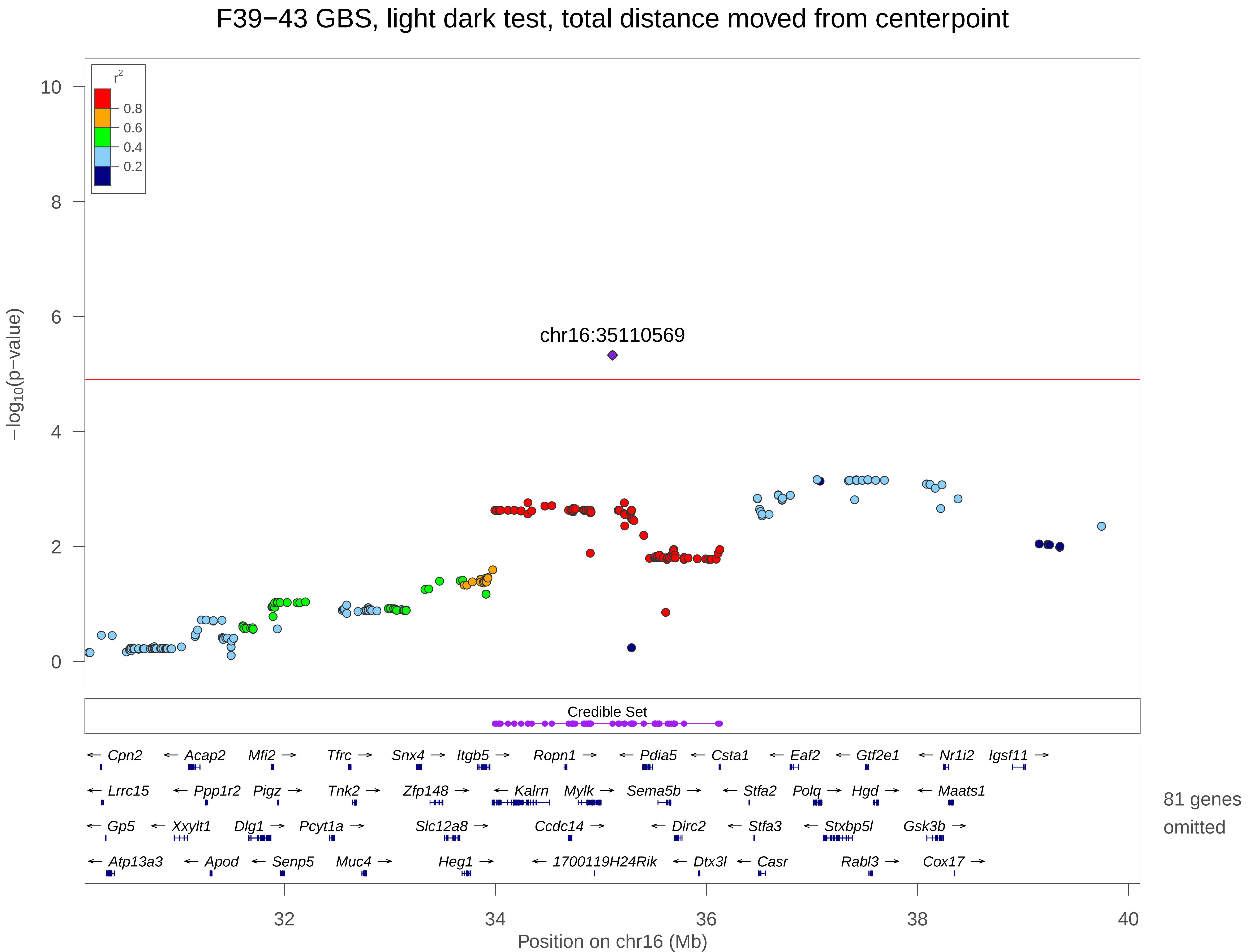

F39−43 GBS, light dark test, total distance moved from centerpoint
r2
0.8
0.6
0.4
0.2
10
8
6
4
2
0
−log10(p−value)
chr16:35110569
Credible Set
← Cpn2
← Acap2
Mfi2 →
Tfrc →
Snx4 →
Itgb5 →
Ropn1 →
← Pdia5
← Csta1
← Eaf2
← Gtf2e1
← Nr1i2
Igsf11 →
← Lrrc15
← Ppp1r2
Pigz →
Tnk2 →
Zfp148 →
← Kalrn
Mylk →
Sema5b →
← Stfa2
Polq →
Hgd →
← Maats1
← Gp5
← Xxylt1
Dlg1 →
Pcyt1a →
Slc12a8 →
Ccdc14 →
← Dirc2
← Stfa3
← Stxbp5l
Gsk3b →
← Atp13a3
← Apod
← Senp5
Muc4 →
Heg1 →
← 1700119H24Rik
← Dtx3l
← Casr
Rabl3 →
Cox17 →
81 genes omitted
32
34
36
38
40
Position on chr16 (Mb)

date: Thu Oct 25 17:22:08 2018
build: F3943GBS
display range: chr16:30110569−40110569 [30110569−40110569]
hilite range: 0 − 0 [ 0 − 0 ]
reference SNP: chr16:35110569
number of SNPs plotted: 353
min P−value: 4.67E−6 [chr16:35110569]
max P−value: 7.87E−1 [chr16:31495890]
omitted Genes: Tmem44, Lsg1, Fam43a
omitted Genes: Bdh1, 0610012G03Rik, Ncbp2
omitted Genes: Gm15694, Pak2, Pigx
omitted Genes: Cep19, Nrros, Bex6
omitted Genes: Fbxo45, Wdr53, Mir1946a
omitted Genes: Smco1, Rnf168, Ubxn7
omitted Genes: Tm4sf19, Tctex1d2, Slc51a
omitted Genes: Zdhhc19, Tnk2os, Muc20
omitted Genes: Gm933, Rubcn, Fyttd1
omitted Genes: Lrch3, Iqcg, Rpl35a
omitted Genes: Lmln, Mir1947, Osbpl11
omitted Genes: 1700007L15Rik, Muc13, Umps
omitted Genes: E130310I04Rik, Hacd2, Adcy5
omitted Genes: Sec22a, 1600019K03Rik, Hspbap1
omitted Genes: Parp14, Parp9, Kpna1
omitted Genes: Wdr5b, Fam162a, Ccdc58
omitted Genes: Stfa2l1, Gm5483, Gm6815
omitted Genes: Gm5416, 2010005H15Rik, Stfa1
omitted Genes: BC117090, BC100530, Cd86
omitted Genes: Ildr1, Slc15a2, Iqcb1
omitted Genes: 4930565N06Rik, Golgb1, Hcls1
omitted Genes: Fbxo40, Ndufb4, Fstl1
omitted Genes: Lrrc58, Gpr156, 4930542C21Rik
omitted Genes: BC031361, Popdc2, Pla1a
omitted Genes: Adprh, Cd80, Timmdc1
omitted Genes: Poglut1, Tmem39a, Arhgap31
omitted Genes: B4galt4, Upk1b, 4930435E12Rik

Make more plots at http://csg.sph.umich.edu/locuszoom/

Fine−mapping Regions

| snp | chr | pos (Mb) | pp | group | color |
|---|---|---|---|---|---|
| chr16:33995903 | 16 | 33.99590 | 0.0024368050 | Credible Set | purple |
| chr16:34010204 | 16 | 34.01020 | 0.0023962077 | Credible Set | purple |
| chr16:34035325 | 16 | 34.03533 | 0.0023962077 | Credible Set | purple |
| chr16:34036026 | 16 | 34.03603 | 0.0024368050 | Credible Set | purple |
| chr16:34050045 | 16 | 34.05004 | 0.0024368050 | Credible Set | purple |
| chr16:34120293 | 16 | 34.12029 | 0.0024368050 | Credible Set | purple |
| chr16:34179072 | 16 | 34.17907 | 0.0024368050 | Credible Set | purple |
| chr16:34179375 | 16 | 34.17938 | 0.0024368050 | Credible Set | purple |
| chr16:34243886 | 16 | 34.24389 | 0.0023809430 | Credible Set | purple |
| chr16:34307872 | 16 | 34.30787 | 0.0021342532 | Credible Set | purple |
| chr16:34308832 | 16 | 34.30883 | 0.0032088489 | Credible Set | purple |
| chr16:34345248 | 16 | 34.34525 | 0.0023771611 | Credible Set | purple |
| chr16:34469856 | 16 | 34.46986 | 0.0028430192 | Credible Set | purple |
| chr16:34535437 | 16 | 34.53544 | 0.0028795529 | Credible Set | purple |
| chr16:34694709 | 16 | 34.69471 | 0.0024368050 | Credible Set | purple |
| chr16:34723432 | 16 | 34.72343 | 0.0024368050 | Credible Set | purple |
| chr16:34736017 | 16 | 34.73602 | 0.0022995103 | Credible Set | purple |
| chr16:34736067 | 16 | 34.73607 | 0.0025934673 | Credible Set | purple |
| chr16:34736543 | 16 | 34.73654 | 0.0024084851 | Credible Set | purple |
| chr16:34736546 | 16 | 34.73655 | 0.0025316505 | Credible Set | purple |
| chr16:34758455 | 16 | 34.75845 | 0.0025754154 | Credible Set | purple |
| chr16:34837044 | 16 | 34.83704 | 0.0024368050 | Credible Set | purple |
| chr16:34845368 | 16 | 34.84537 | 0.0024368050 | Credible Set | purple |
| chr16:34848045 | 16 | 34.84804 | 0.0024368050 | Credible Set | purple |
| chr16:34862374 | 16 | 34.86237 | 0.0024368050 | Credible Set | purple |
| chr16:34878627 | 16 | 34.87863 | 0.0024368050 | Credible Set | purple |
| chr16:34879746 | 16 | 34.87975 | 0.0023766208 | Credible Set | purple |
| chr16:34880408 | 16 | 34.88041 | 0.0024368050 | Credible Set | purple |
| chr16:34880442 | 16 | 34.88044 | 0.0024368050 | Credible Set | purple |
| chr16:34899407 | 16 | 34.89941 | 0.0024368050 | Credible Set | purple |
| chr16:34899479 | 16 | 34.89948 | 0.0024368050 | Credible Set | purple |
| chr16:34899567 | 16 | 34.89957 | 0.0024368050 | Credible Set | purple |
| chr16:34899593 | 16 | 34.89959 | 0.0024319097 | Credible Set | purple |
| chr16:34899645 | 16 | 34.89964 | 0.0024319097 | Credible Set | purple |
| chr16:34899669 | 16 | 34.89967 | 0.0005164153 | Credible Set | purple |
| chr16:34899682 | 16 | 34.89968 | 0.0022494902 | Credible Set | purple |

Fine−mapping Regions

| snp | chr | pos (Mb) | pp | group | color |
|---|---|---|---|---|---|
| chr16:34899715 | 16 | 34.89972 | 0.0022174086 | Credible Set | purple |
| chr16:34907286 | 16 | 34.90729 | 0.0022782650 | Credible Set | purple |
| chr16:35110569 | 16 | 35.11057 | 0.8469245785 | Credible Set | purple |
| chr16:35165377 | 16 | 35.16538 | 0.0024337472 | Credible Set | purple |
| chr16:35165381 | 16 | 35.16538 | 0.0024337472 | Credible Set | purple |
| chr16:35176099 | 16 | 35.17610 | 0.0024368050 | Credible Set | purple |
| chr16:35222444 | 16 | 35.22244 | 0.0021450546 | Credible Set | purple |
| chr16:35222475 | 16 | 35.22248 | 0.0021450546 | Credible Set | purple |
| chr16:35222947 | 16 | 35.22295 | 0.0021160438 | Credible Set | purple |
| chr16:35223024 | 16 | 35.22302 | 0.0032081124 | Credible Set | purple |
| chr16:35226617 | 16 | 35.22662 | 0.0013740275 | Credible Set | purple |
| chr16:35226697 | 16 | 35.22670 | 0.0020796366 | Credible Set | purple |
| chr16:35282970 | 16 | 35.28297 | 0.0022315710 | Credible Set | purple |
| chr16:35286816 | 16 | 35.28682 | 0.0023115616 | Credible Set | purple |
| chr16:35290682 | 16 | 35.29068 | 0.0024184026 | Credible Set | purple |
| chr16:35290929 | 16 | 35.29093 | 0.0024368050 | Credible Set | purple |
| chr16:35291606 | 16 | 35.29161 | 0.0017806385 | Credible Set | purple |
| chr16:35301999 | 16 | 35.30200 | 0.0017074395 | Credible Set | purple |
| chr16:35303186 | 16 | 35.30319 | 0.0017229572 | Credible Set | purple |
| chr16:35313747 | 16 | 35.31375 | 0.0016643743 | Credible Set | purple |
| chr16:35406924 | 16 | 35.40692 | 0.0009783719 | Credible Set | purple |
| chr16:35407400 | 16 | 35.40740 | 0.0009728870 | Credible Set | purple |
| chr16:35509375 | 16 | 35.50937 | 0.0004403027 | Credible Set | purple |
| chr16:35512153 | 16 | 35.51215 | 0.0004403027 | Credible Set | purple |
| chr16:35514329 | 16 | 35.51433 | 0.0004403027 | Credible Set | purple |
| chr16:35514333 | 16 | 35.51433 | 0.0004403027 | Credible Set | purple |
| chr16:35514335 | 16 | 35.51434 | 0.0004403027 | Credible Set | purple |
| chr16:35514337 | 16 | 35.51434 | 0.0004403027 | Credible Set | purple |
| chr16:35519320 | 16 | 35.51932 | 0.0004440990 | Credible Set | purple |
| chr16:35520175 | 16 | 35.52018 | 0.0004616747 | Credible Set | purple |
| chr16:35535811 | 16 | 35.53581 | 0.0004527721 | Credible Set | purple |
| chr16:35552316 | 16 | 35.55232 | 0.0004454180 | Credible Set | purple |
| chr16:35552393 | 16 | 35.55239 | 0.0004475908 | Credible Set | purple |
| chr16:35552802 | 16 | 35.55280 | 0.0004429417 | Credible Set | purple |
| chr16:35554979 | 16 | 35.55498 | 0.0004531593 | Credible Set | purple |
| chr16:35555052 | 16 | 35.55505 | 0.0004513451 | Credible Set | purple |

Fine−mapping Regions

| snp | chr | pos (Mb) | pp | group | color |
|---|---|---|---|---|---|
| chr16:35555962 | 16 | 35.55596 | 0.0004550084 | Credible Set | purple |
| chr16:35555964 | 16 | 35.55596 | 0.0004815372 | Credible Set | purple |
| chr16:35632372 | 16 | 35.63237 | 0.0004447240 | Credible Set | purple |
| chr16:35648933 | 16 | 35.64893 | 0.0004452547 | Credible Set | purple |
| chr16:35668489 | 16 | 35.66849 | 0.0004763965 | Credible Set | purple |
| chr16:35688514 | 16 | 35.68851 | 0.0004971362 | Credible Set | purple |
| chr16:35690798 | 16 | 35.69080 | 0.0005898687 | Credible Set | purple |
| chr16:35691243 | 16 | 35.69124 | 0.0004954077 | Credible Set | purple |
| chr16:35691297 | 16 | 35.69130 | 0.0005585339 | Credible Set | purple |
| chr16:35704080 | 16 | 35.70408 | 0.0004846041 | Credible Set | purple |
| chr16:35788676 | 16 | 35.78868 | 0.0004442823 | Credible Set | purple |
| chr16:35789020 | 16 | 35.78902 | 0.0004427413 | Credible Set | purple |
| chr16:36111216 | 16 | 36.11122 | 0.0005108505 | Credible Set | purple |
| chr16:36127401 | 16 | 36.12740 | 0.0005881521 | Credible Set | purple |

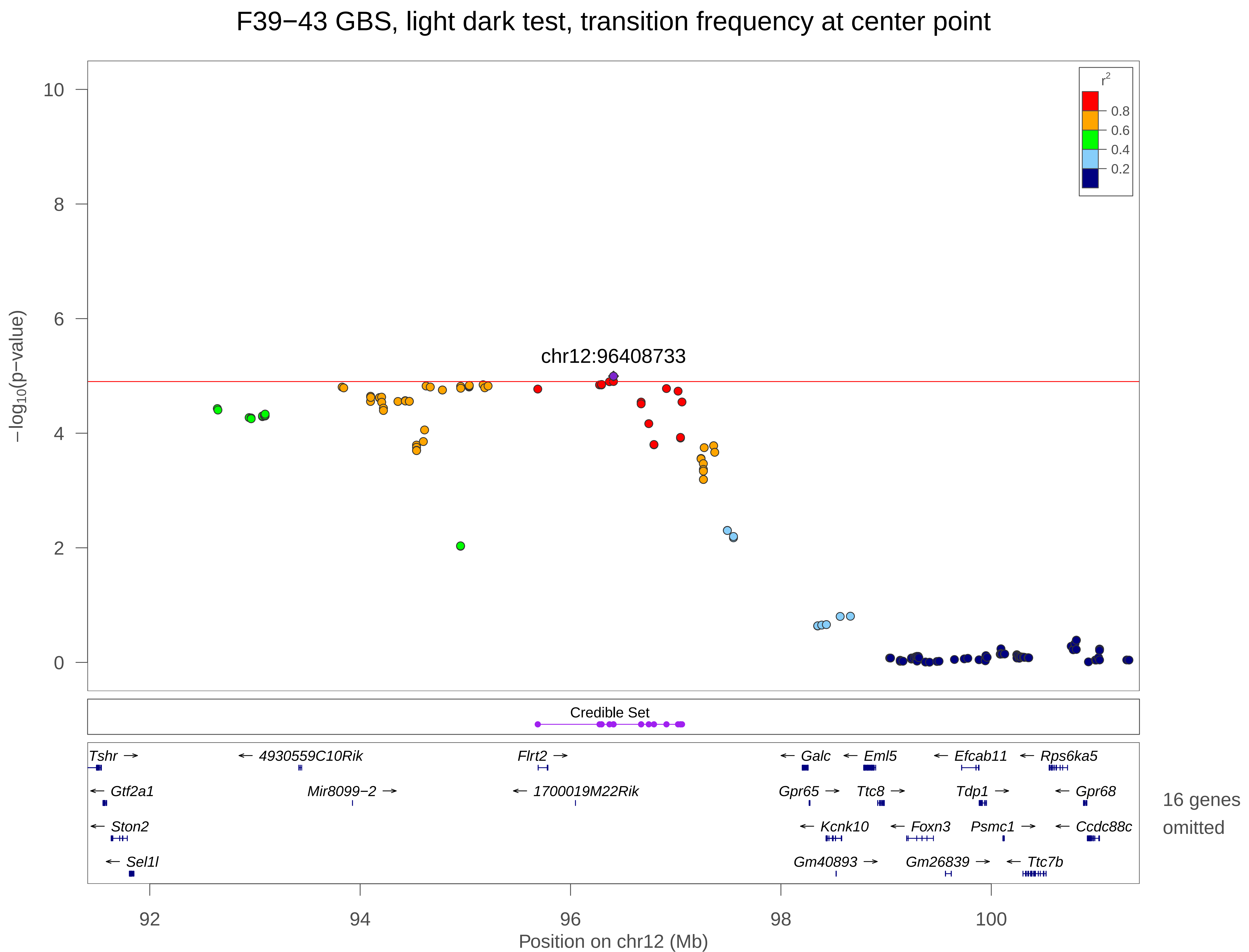
F39−43 GBS, light dark test, transition frequency at center point
−log10(p−value)
10
8
6
4
2
0
chr12:96408733
r2
0.8
0.6
0.4
0.2
Credible Set
Tshr
4930559C10Rik
Flrt2
Galc
Eml5
Efcab11
Rps6ka5
Gtf2a1
Mir8099−2
1700019M22Rik
Gpr65
Ttc8
Tdp1
Gpr68
Ston2
Kcnk10
Foxn3
Psmc1
Ccdc88c
Sel1l
Gm40893
Gm26839
Ttc7b
16 genes omitted
92
94
96
98
100
Position on chr12 (Mb)

| | |
|---|---|
| date: | Thu Oct 25 17:22:18 2018 |
| build: | F3943GBS |
| display range: | chr12:91408733−101408733 [91408733−101408733] |
| hilite range: | 0 − 0 [ 0 − 0 ] |
| reference SNP: | chr12:96408733 |
| number of SNPs plotted: | 187 |
| min P−value: | 1.01E−5 [chr12:96408733] |
| max P−value: | 9.91E−1 [chr12:99412684] |
| omitted Genes: | Spata7, Ptpn21, Zc3h14 |
| omitted Genes: | 4930474N09Rik, 1700064M15Rik, Kcnk13 |
| omitted Genes: | Nrde2, Gm10433, Calm1 |
| omitted Genes: | Gm10432, 9030617O03Rik, Mir1190 |
| omitted Genes: | Smek1, D130020L05Rik, Kif4−ps |
| omitted Genes: | Catsperb |

Make more plots at http://csg.sph.umich.edu/locuszoom/

Fine−mapping Regions

| snp | chr | pos (Mb) | pp | group | color |
|---|---|---|---|---|---|
| chr12:95688559 | 12 | 95.68856 | 0.066583928 | Credible Set | purple |
| chr12:96276115 | 12 | 96.27612 | 0.078063741 | Credible Set | purple |
| chr12:96293767 | 12 | 96.29377 | 0.077868747 | Credible Set | purple |
| chr12:96295488 | 12 | 96.29549 | 0.079269278 | Credible Set | purple |
| chr12:96370579 | 12 | 96.37058 | 0.088151618 | Credible Set | purple |
| chr12:96408510 | 12 | 96.40851 | 0.088922195 | Credible Set | purple |
| chr12:96408667 | 12 | 96.40867 | 0.109049512 | Credible Set | purple |
| chr12:96408733 | 12 | 96.40873 | 0.109283651 | Credible Set | purple |
| chr12:96671267 | 12 | 96.67127 | 0.040480551 | Credible Set | purple |
| chr12:96671328 | 12 | 96.67133 | 0.037775520 | Credible Set | purple |
| chr12:96743964 | 12 | 96.74396 | 0.017774380 | Credible Set | purple |
| chr12:96793184 | 12 | 96.79318 | 0.008065971 | Credible Set | purple |
| chr12:96912045 | 12 | 96.91205 | 0.067868114 | Credible Set | purple |
| chr12:97022371 | 12 | 97.02237 | 0.061429753 | Credible Set | purple |
| chr12:97045366 | 12 | 97.04537 | 0.010299811 | Credible Set | purple |
| chr12:97045367 | 12 | 97.04537 | 0.010556824 | Credible Set | purple |
| chr12:97059616 | 12 | 97.05962 | 0.040591987 | Credible Set | purple |

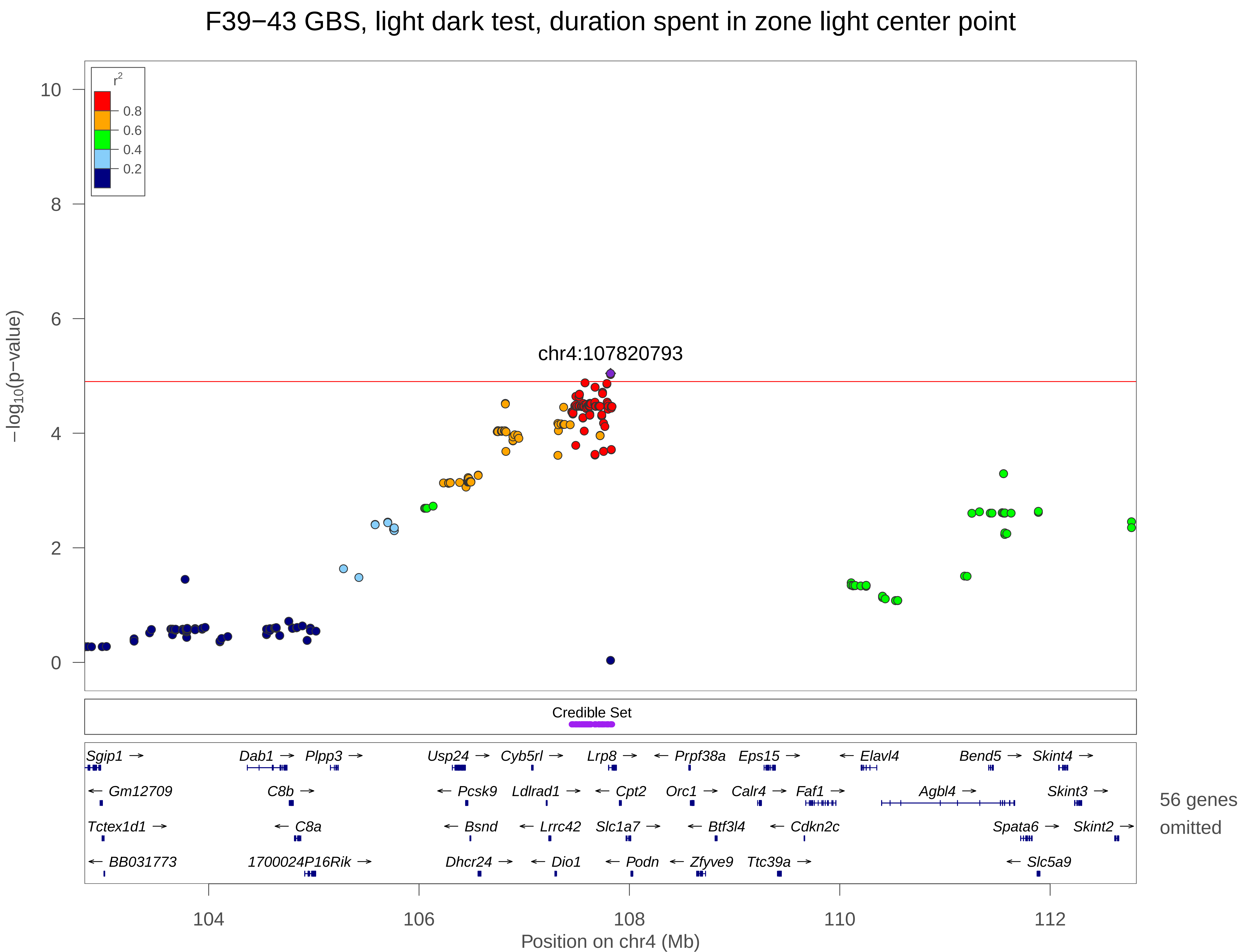
F39−43 GBS, light dark test, duration spent in zone light center point
r2
0.8
0.6
0.4
0.2
10
8
6
4
2
0
−log10(p−value)
chr4:107820793
Credible Set
Sgip1 →
Dab1 →
Plpp3 →
Usp24 →
Cyb5rl →
Lrp8 →
← Prpf38a
Eps15 →
← Elavl4
Bend5 →
Skint4 →
← Gm12709
C8b →
← Pcsk9
Ldlrad1 →
← Cpt2
Orc1 →
Calr4 →
Faf1 →
Agbl4 →
Skint3 →
Tctex1d1 →
← C8a
← Bsnd
← Lrrc42
Slc1a7 →
← Btf3l4
← Cdkn2c
Spata6 →
Skint2 →
← BB031773
1700024P16Rik →
Dhcr24 →
← Dio1
← Podn
← Zfyve9
Ttc39a →
← Slc5a9
56 genes omitted
104
106
108
110
112
Position on chr4 (Mb)

| | |
|---|---|
| date: | Thu Oct 25 17:22:26 2018 |
| build: | F3943GBS |
| display range: | chr4:102820793−112820793 [102820793−112820793] |
| hilite range: | 0 − 0 [ 0 − 0 ] |
| reference SNP: | chr4:107820793 |
| number of SNPs plotted: | 382 |
| min P−value: | 9.04E−6 [chr4:107820793] |
| max P−value: | 9.21E−1 [chr4:107820753] |
| omitted Genes: | Insl5, Wdr78, Mier1 |
| omitted Genes: | Slc35d1, 4921539E11Rik, 4930456L15Rik |
| omitted Genes: | Oma1, Gm12718, Prkaa2 |
| omitted Genes: | Lexm, Ttc22, Pars2 |
| omitted Genes: | Ttc4, Mroh7, Fam151a |
| omitted Genes: | Acot11, Ssbp3, Mrpl37 |
| omitted Genes: | Cdcp2, Tceanc2, Tmem59 |
| omitted Genes: | Hspb11, Yipf1, Ndc1 |
| omitted Genes: | Glis1, Dmrtb1, Magoh |
| omitted Genes: | 0610037L13Rik, 4930407G08Rik, Scp2 |
| omitted Genes: | Mir6397, Echdc2, Zyg11a |
| omitted Genes: | Zyg11b, Coa7, Fam159a |
| omitted Genes: | Gpx7, Zcchc11, Cc2d1b |
| omitted Genes: | Txndc12, Kti12, Rab3b |
| omitted Genes: | 8030443G20Rik, Nrd1, Mir761 |
| omitted Genes: | Osbpl9, Ttc39aos1, Rnf11 |
| omitted Genes: | 4930522H14Rik, Dmrta2, Skint8 |
| omitted Genes: | Skint7, Skint1, Skint9 |
| omitted Genes: | Skint10, Skint6 |

Make more plots at http://csg.sph.umich.edu/locuszoom/

Fine−mapping Regions

| snp | chr | pos (Mb) | pp | group | color |
|---|---|---|---|---|---|
| chr4:107452487 | 4 | 107.4525 | 0.006139849 | Credible Set | purple |
| chr4:107452522 | 4 | 107.4525 | 0.006210446 | Credible Set | purple |
| chr4:107460844 | 4 | 107.4608 | 0.005884823 | Credible Set | purple |
| chr4:107462435 | 4 | 107.4624 | 0.005929123 | Credible Set | purple |
| chr4:107462443 | 4 | 107.4624 | 0.005685471 | Credible Set | purple |
| chr4:107462966 | 4 | 107.4630 | 0.005927923 | Credible Set | purple |
| chr4:107463057 | 4 | 107.4631 | 0.006242026 | Credible Set | purple |
| chr4:107463550 | 4 | 107.4635 | 0.005804363 | Credible Set | purple |
| chr4:107463697 | 4 | 107.4637 | 0.005890754 | Credible Set | purple |
| chr4:107480817 | 4 | 107.4808 | 0.007801357 | Credible Set | purple |
| chr4:107489799 | 4 | 107.4898 | 0.001733295 | Credible Set | purple |
| chr4:107489889 | 4 | 107.4899 | 0.007645256 | Credible Set | purple |
| chr4:107491410 | 4 | 107.4914 | 0.008082270 | Credible Set | purple |
| chr4:107491522 | 4 | 107.4915 | 0.011163287 | Credible Set | purple |
| chr4:107491535 | 4 | 107.4915 | 0.011163287 | Credible Set | purple |
| chr4:107491555 | 4 | 107.4916 | 0.011163287 | Credible Set | purple |
| chr4:107502298 | 4 | 107.5023 | 0.007645256 | Credible Set | purple |
| chr4:107502806 | 4 | 107.5028 | 0.007645256 | Credible Set | purple |
| chr4:107502862 | 4 | 107.5029 | 0.007645256 | Credible Set | purple |
| chr4:107519045 | 4 | 107.5190 | 0.007645256 | Credible Set | purple |
| chr4:107520793 | 4 | 107.5208 | 0.010952576 | Credible Set | purple |
| chr4:107524390 | 4 | 107.5244 | 0.007655839 | Credible Set | purple |
| chr4:107524399 | 4 | 107.5244 | 0.007748733 | Credible Set | purple |
| chr4:107524747 | 4 | 107.5247 | 0.011850794 | Credible Set | purple |
| chr4:107524800 | 4 | 107.5248 | 0.012169599 | Credible Set | purple |
| chr4:107524808 | 4 | 107.5248 | 0.011839126 | Credible Set | purple |
| chr4:107524812 | 4 | 107.5248 | 0.012062379 | Credible Set | purple |
| chr4:107542864 | 4 | 107.5429 | 0.007645256 | Credible Set | purple |
| chr4:107543088 | 4 | 107.5431 | 0.007645256 | Credible Set | purple |
| chr4:107543347 | 4 | 107.5433 | 0.007621806 | Credible Set | purple |
| chr4:107548442 | 4 | 107.5484 | 0.007645256 | Credible Set | purple |
| chr4:107550049 | 4 | 107.5500 | 0.007539375 | Credible Set | purple |
| chr4:107550052 | 4 | 107.5501 | 0.007542804 | Credible Set | purple |
| chr4:107550153 | 4 | 107.5502 | 0.007645256 | Credible Set | purple |
| chr4:107558047 | 4 | 107.5580 | 0.004953255 | Credible Set | purple |
| chr4:107558061 | 4 | 107.5581 | 0.004872070 | Credible Set | purple |

Fine−mapping Regions

| snp | chr | pos (Mb) | pp | group | color |
|---|---|---|---|---|---|
| chr4:107558064 | 4 | 107.5581 | 0.004970102 | Credible Set | purple |
| chr4:107561625 | 4 | 107.5616 | 0.008519314 | Credible Set | purple |
| chr4:107563786 | 4 | 107.5638 | 0.007555548 | Credible Set | purple |
| chr4:107570421 | 4 | 107.5704 | 0.002980557 | Credible Set | purple |
| chr4:107570446 | 4 | 107.5704 | 0.008029709 | Credible Set | purple |
| chr4:107570783 | 4 | 107.5708 | 0.007555548 | Credible Set | purple |
| chr4:107578001 | 4 | 107.5780 | 0.018747346 | Credible Set | purple |
| chr4:107578013 | 4 | 107.5780 | 0.018747346 | Credible Set | purple |
| chr4:107589309 | 4 | 107.5893 | 0.007588680 | Credible Set | purple |
| chr4:107589365 | 4 | 107.5894 | 0.007564067 | Credible Set | purple |
| chr4:107589847 | 4 | 107.5898 | 0.007645256 | Credible Set | purple |
| chr4:107589899 | 4 | 107.5899 | 0.007645256 | Credible Set | purple |
| chr4:107590217 | 4 | 107.5902 | 0.007645256 | Credible Set | purple |
| chr4:107590304 | 4 | 107.5903 | 0.008225228 | Credible Set | purple |
| chr4:107590311 | 4 | 107.5903 | 0.008113060 | Credible Set | purple |
| chr4:107590318 | 4 | 107.5903 | 0.008204375 | Credible Set | purple |
| chr4:107590777 | 4 | 107.5908 | 0.007038290 | Credible Set | purple |
| chr4:107590782 | 4 | 107.5908 | 0.006994323 | Credible Set | purple |
| chr4:107590825 | 4 | 107.5908 | 0.007016611 | Credible Set | purple |
| chr4:107601123 | 4 | 107.6011 | 0.007645256 | Credible Set | purple |
| chr4:107601170 | 4 | 107.6012 | 0.007645256 | Credible Set | purple |
| chr4:107602251 | 4 | 107.6023 | 0.007541042 | Credible Set | purple |
| chr4:107603006 | 4 | 107.6030 | 0.007645256 | Credible Set | purple |
| chr4:107603359 | 4 | 107.6034 | 0.007217989 | Credible Set | purple |
| chr4:107603395 | 4 | 107.6034 | 0.007240987 | Credible Set | purple |
| chr4:107614922 | 4 | 107.6149 | 0.007645256 | Credible Set | purple |
| chr4:107619675 | 4 | 107.6197 | 0.007645256 | Credible Set | purple |
| chr4:107619763 | 4 | 107.6198 | 0.007756452 | Credible Set | purple |
| chr4:107623179 | 4 | 107.6232 | 0.007761225 | Credible Set | purple |
| chr4:107623470 | 4 | 107.6235 | 0.005729825 | Credible Set | purple |
| chr4:107623488 | 4 | 107.6235 | 0.005554516 | Credible Set | purple |
| chr4:107623496 | 4 | 107.6235 | 0.005426062 | Credible Set | purple |
| chr4:107623506 | 4 | 107.6235 | 0.005517344 | Credible Set | purple |
| chr4:107623515 | 4 | 107.6235 | 0.005488297 | Credible Set | purple |
| chr4:107623565 | 4 | 107.6236 | 0.007779348 | Credible Set | purple |
| chr4:107623613 | 4 | 107.6236 | 0.008569345 | Credible Set | purple |

Fine−mapping Regions

| snp | chr | pos (Mb) | pp | group | color |
|---|---|---|---|---|---|
| chr4:107624024 | 4 | 107.6240 | 0.005470502 | Credible Set | purple |
| chr4:107630955 | 4 | 107.6310 | 0.008527009 | Credible Set | purple |
| chr4:107636029 | 4 | 107.6360 | 0.008427555 | Credible Set | purple |
| chr4:107671009 | 4 | 107.6710 | 0.008181240 | Credible Set | purple |
| chr4:107671349 | 4 | 107.6713 | 0.008547164 | Credible Set | purple |
| chr4:107672292 | 4 | 107.6723 | 0.008584331 | Credible Set | purple |
| chr4:107672310 | 4 | 107.6723 | 0.008491017 | Credible Set | purple |
| chr4:107672333 | 4 | 107.6723 | 0.008952812 | Credible Set | purple |
| chr4:107673647 | 4 | 107.6736 | 0.015849244 | Credible Set | purple |
| chr4:107673650 | 4 | 107.6736 | 0.015849244 | Credible Set | purple |
| chr4:107673669 | 4 | 107.6737 | 0.015849244 | Credible Set | purple |
| chr4:107676434 | 4 | 107.6764 | 0.007645256 | Credible Set | purple |
| chr4:107677045 | 4 | 107.6770 | 0.007645256 | Credible Set | purple |
| chr4:107677381 | 4 | 107.6774 | 0.007645256 | Credible Set | purple |
| chr4:107680225 | 4 | 107.6802 | 0.007645256 | Credible Set | purple |
| chr4:107681615 | 4 | 107.6816 | 0.007645256 | Credible Set | purple |
| chr4:107707410 | 4 | 107.7074 | 0.007735884 | Credible Set | purple |
| chr4:107707549 | 4 | 107.7075 | 0.007645256 | Credible Set | purple |
| chr4:107707574 | 4 | 107.7076 | 0.007645256 | Credible Set | purple |
| chr4:107710876 | 4 | 107.7109 | 0.007645256 | Credible Set | purple |
| chr4:107719853 | 4 | 107.7199 | 0.007645256 | Credible Set | purple |
| chr4:107719899 | 4 | 107.7199 | 0.007645256 | Credible Set | purple |
| chr4:107736672 | 4 | 107.7367 | 0.005306097 | Credible Set | purple |
| chr4:107737167 | 4 | 107.7372 | 0.005521127 | Credible Set | purple |
| chr4:107744658 | 4 | 107.7447 | 0.013103814 | Credible Set | purple |
| chr4:107744663 | 4 | 107.7447 | 0.012946291 | Credible Set | purple |
| chr4:107744682 | 4 | 107.7447 | 0.012392523 | Credible Set | purple |
| chr4:107754250 | 4 | 107.7542 | 0.004010015 | Credible Set | purple |
| chr4:107766842 | 4 | 107.7668 | 0.003532624 | Credible Set | purple |
| chr4:107785888 | 4 | 107.7859 | 0.017900074 | Credible Set | purple |
| chr4:107785926 | 4 | 107.7859 | 0.018294844 | Credible Set | purple |
| chr4:107789013 | 4 | 107.7890 | 0.008962432 | Credible Set | purple |
| chr4:107797328 | 4 | 107.7973 | 0.006837035 | Credible Set | purple |
| chr4:107797813 | 4 | 107.7978 | 0.008409549 | Credible Set | purple |
| chr4:107800894 | 4 | 107.8009 | 0.007680796 | Credible Set | purple |
| chr4:107820793 | 4 | 107.8208 | 0.026928760 | Credible Set | purple |

Fine−mapping Regions

| snp | chr | pos (Mb) | pp | group | color |
|---|---|---|---|---|---|
| chr4:107820800 | 4 | 107.8208 | 0.025762219 | Credible Set | purple |
| chr4:107827933 | 4 | 107.8279 | 0.001462287 | Credible Set | purple |
| chr4:107827937 | 4 | 107.8279 | 0.001481504 | Credible Set | purple |
| chr4:107827946 | 4 | 107.8279 | 0.007385856 | Credible Set | purple |
| chr4:107828049 | 4 | 107.8280 | 0.007124388 | Credible Set | purple |
| chr4:107828736 | 4 | 107.8287 | 0.007645256 | Credible Set | purple |
| chr4:107828810 | 4 | 107.8288 | 0.007645256 | Credible Set | purple |
| chr4:107828843 | 4 | 107.8288 | 0.007645256 | Credible Set | purple |
| chr4:107828858 | 4 | 107.8289 | 0.007645256 | Credible Set | purple |
| chr4:107834883 | 4 | 107.8349 | 0.007575529 | Credible Set | purple |

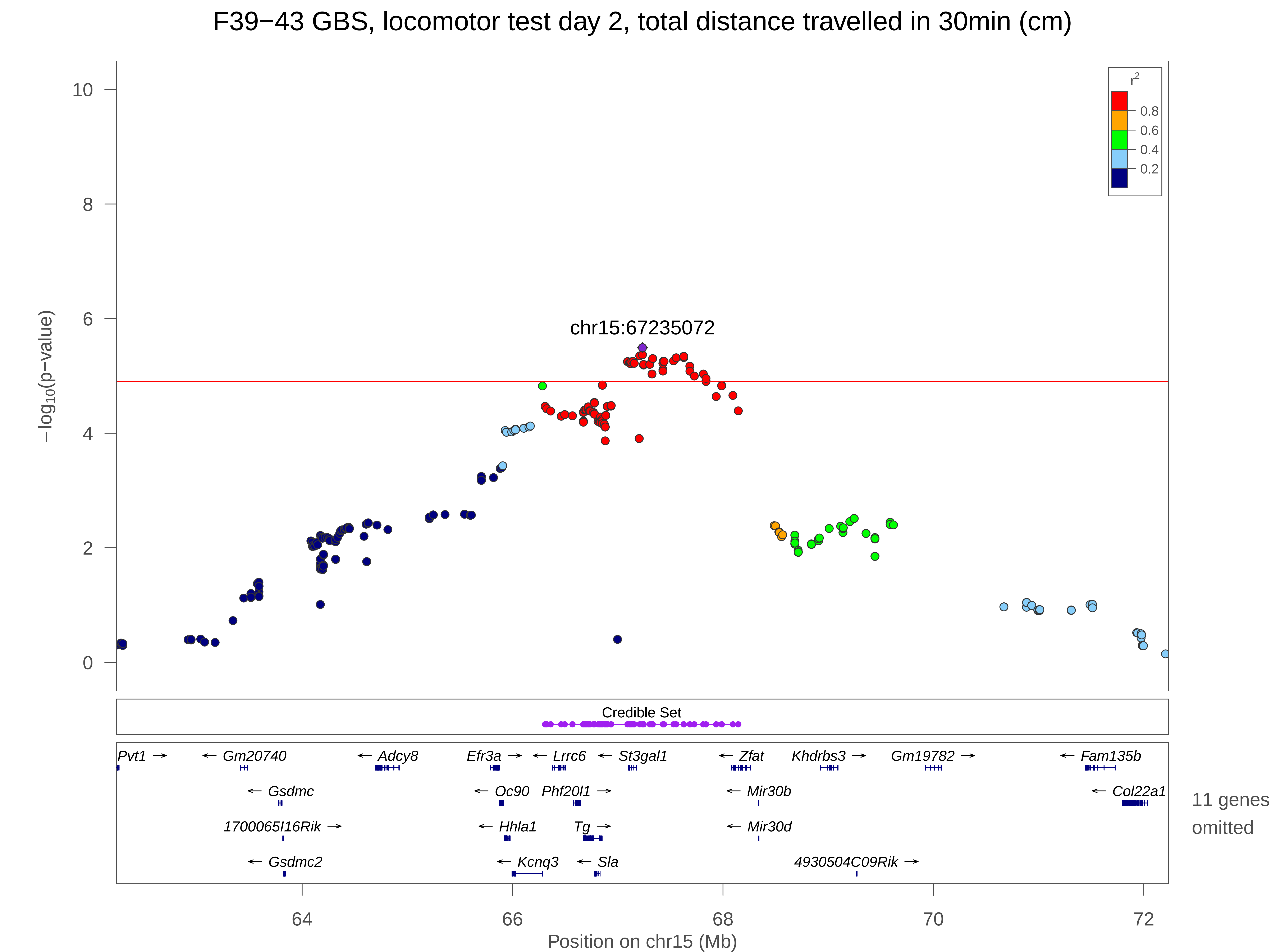

F39−43 GBS, locomotor test day 2, total distance travelled in 30min (cm)
−log10(p−value)
10
8
6
4
2
0
r2
0.8
0.6
0.4
0.2
chr15:67235072
Credible Set
Pvt1 →
← Gm20740
← Adcy8
Efr3a →
← Lrrc6
← St3gal1
← Zfat
Khdrbs3 →
Gm19782 →
← Fam135b
← Gsdmc
← Oc90
Phf20l1 →
← Mir30b
← Col22a1
1700065I16Rik →
← Hhla1
Tg →
← Mir30d
← Gsdmc2
← Kcnq3
← Sla
4930504C09Rik →
11 genes omitted
64
66
68
70
72
Position on chr15 (Mb)

| | |
|---|---|
| date: | Thu Oct 25 17:22:36 2018 |
| build: | F3943GBS |
| display range: | chr15:62235072−72235072 [62235072−72235072] |
| hilite range: | 0 − 0 [ 0 − 0 ] |
| reference SNP: | chr15:67235072 |
| number of SNPs plotted: | 310 |
| min P−value: | 3.21E−6 [chr15:67235072] |
| max P−value: | 7.1E−1 [chr15:72206582] |
| omitted Genes: | Gsdmcl1, Gsdmc3, Gsdmcl2 |
| omitted Genes: | Gsdmc4, Gsdmcl−ps, Fam49b |
| omitted Genes: | Asap1, Tmem71, Wisp1 |
| omitted Genes: | Ndrg1, 1700012I11Rik |

Make more plots at http://csg.sph.umich.edu/locuszoom/

Fine−mapping Regions

| snp | chr | pos (Mb) | pp | group | color |
|---|---|---|---|---|---|
| chr15:66309070 | 15 | 66.30907 | 0.004218721 | Credible Set | purple |
| chr15:66324506 | 15 | 66.32451 | 0.003847034 | Credible Set | purple |
| chr15:66361813 | 15 | 66.36181 | 0.003523650 | Credible Set | purple |
| chr15:66462727 | 15 | 66.46273 | 0.002893196 | Credible Set | purple |
| chr15:66495529 | 15 | 66.49553 | 0.003080992 | Credible Set | purple |
| chr15:66569471 | 15 | 66.56947 | 0.002943707 | Credible Set | purple |
| chr15:66672984 | 15 | 66.67298 | 0.002364374 | Credible Set | purple |
| chr15:66672997 | 15 | 66.67300 | 0.002396174 | Credible Set | purple |
| chr15:66673010 | 15 | 66.67301 | 0.002381686 | Credible Set | purple |
| chr15:66673015 | 15 | 66.67302 | 0.002304852 | Credible Set | purple |
| chr15:66673436 | 15 | 66.67344 | 0.003341858 | Credible Set | purple |
| chr15:66673541 | 15 | 66.67354 | 0.003372912 | Credible Set | purple |
| chr15:66673940 | 15 | 66.67394 | 0.003385618 | Credible Set | purple |
| chr15:66685097 | 15 | 66.68510 | 0.003667083 | Credible Set | purple |
| chr15:66696518 | 15 | 66.69652 | 0.003683460 | Credible Set | purple |
| chr15:66699723 | 15 | 66.69972 | 0.003729646 | Credible Set | purple |
| chr15:66719126 | 15 | 66.71913 | 0.003898221 | Credible Set | purple |
| chr15:66719241 | 15 | 66.71924 | 0.004140018 | Credible Set | purple |
| chr15:66720772 | 15 | 66.72077 | 0.004121078 | Credible Set | purple |
| chr15:66730424 | 15 | 66.73042 | 0.003560449 | Credible Set | purple |
| chr15:66737376 | 15 | 66.73738 | 0.003519368 | Credible Set | purple |
| chr15:66768701 | 15 | 66.76870 | 0.003341876 | Credible Set | purple |
| chr15:66777400 | 15 | 66.77740 | 0.003179822 | Credible Set | purple |
| chr15:66777425 | 15 | 66.77742 | 0.003127549 | Credible Set | purple |
| chr15:66777461 | 15 | 66.77746 | 0.003138400 | Credible Set | purple |
| chr15:66777706 | 15 | 66.77771 | 0.004896398 | Credible Set | purple |
| chr15:66777728 | 15 | 66.77773 | 0.004816430 | Credible Set | purple |
| chr15:66777729 | 15 | 66.77773 | 0.004843788 | Credible Set | purple |
| chr15:66777730 | 15 | 66.77773 | 0.004809500 | Credible Set | purple |
| chr15:66777731 | 15 | 66.77773 | 0.004763740 | Credible Set | purple |
| chr15:66777743 | 15 | 66.77774 | 0.004802494 | Credible Set | purple |
| chr15:66812941 | 15 | 66.81294 | 0.002370029 | Credible Set | purple |
| chr15:66826934 | 15 | 66.82693 | 0.002388166 | Credible Set | purple |
| chr15:66834506 | 15 | 66.83451 | 0.002747356 | Credible Set | purple |
| chr15:66834532 | 15 | 66.83453 | 0.002755779 | Credible Set | purple |
| chr15:66834574 | 15 | 66.83457 | 0.002766909 | Credible Set | purple |

Fine−mapping Regions

| snp | chr | pos (Mb) | pp | group | color |
|---|---|---|---|---|---|
| chr15:66834575 | 15 | 66.83458 | 0.002806045 | Credible Set | purple |
| chr15:66834600 | 15 | 66.83460 | 0.002326459 | Credible Set | purple |
| chr15:66834621 | 15 | 66.83462 | 0.002390184 | Credible Set | purple |
| chr15:66834624 | 15 | 66.83462 | 0.002336239 | Credible Set | purple |
| chr15:66834637 | 15 | 66.83464 | 0.002272719 | Credible Set | purple |
| chr15:66853166 | 15 | 66.85317 | 0.002560343 | Credible Set | purple |
| chr15:66853439 | 15 | 66.85344 | 0.002425835 | Credible Set | purple |
| chr15:66853460 | 15 | 66.85346 | 0.002412435 | Credible Set | purple |
| chr15:66853489 | 15 | 66.85349 | 0.002427979 | Credible Set | purple |
| chr15:66853520 | 15 | 66.85352 | 0.002184658 | Credible Set | purple |
| chr15:66853893 | 15 | 66.85389 | 0.009535500 | Credible Set | purple |
| chr15:66853894 | 15 | 66.85389 | 0.009432374 | Credible Set | purple |
| chr15:66853923 | 15 | 66.85392 | 0.009487307 | Credible Set | purple |
| chr15:66853954 | 15 | 66.85395 | 0.009436047 | Credible Set | purple |
| chr15:66872519 | 15 | 66.87252 | 0.002135683 | Credible Set | purple |
| chr15:66880635 | 15 | 66.88063 | 0.001921461 | Credible Set | purple |
| chr15:66886219 | 15 | 66.88622 | 0.002992645 | Credible Set | purple |
| chr15:66886266 | 15 | 66.88627 | 0.002980984 | Credible Set | purple |
| chr15:66901528 | 15 | 66.90153 | 0.004201679 | Credible Set | purple |
| chr15:66901548 | 15 | 66.90155 | 0.004201679 | Credible Set | purple |
| chr15:66901593 | 15 | 66.90159 | 0.004201679 | Credible Set | purple |
| chr15:66935376 | 15 | 66.93538 | 0.004259205 | Credible Set | purple |
| chr15:66935390 | 15 | 66.93539 | 0.004259205 | Credible Set | purple |
| chr15:66935391 | 15 | 66.93539 | 0.004259205 | Credible Set | purple |
| chr15:66935405 | 15 | 66.93541 | 0.004259205 | Credible Set | purple |
| chr15:66937231 | 15 | 66.93723 | 0.004348588 | Credible Set | purple |
| chr15:66937423 | 15 | 66.93742 | 0.004326168 | Credible Set | purple |
| chr15:67092666 | 15 | 67.09267 | 0.023397132 | Credible Set | purple |
| chr15:67111279 | 15 | 67.11128 | 0.022180952 | Credible Set | purple |
| chr15:67121570 | 15 | 67.12157 | 0.021663589 | Credible Set | purple |
| chr15:67121571 | 15 | 67.12157 | 0.021703182 | Credible Set | purple |
| chr15:67121756 | 15 | 67.12176 | 0.023177643 | Credible Set | purple |
| chr15:67126159 | 15 | 67.12616 | 0.023149580 | Credible Set | purple |
| chr15:67144054 | 15 | 67.14405 | 0.023635943 | Credible Set | purple |
| chr15:67156480 | 15 | 67.15648 | 0.021964536 | Credible Set | purple |
| chr15:67208761 | 15 | 67.20876 | 0.029211974 | Credible Set | purple |

Fine−mapping Regions

| snp | chr | pos (Mb) | pp | group | color |
|---|---|---|---|---|---|
| chr15:67234910 | 15 | 67.23491 | 0.030492843 | Credible Set | purple |
| chr15:67235072 | 15 | 67.23507 | 0.040065247 | Credible Set | purple |
| chr15:67245093 | 15 | 67.24509 | 0.020492326 | Credible Set | purple |
| chr15:67245123 | 15 | 67.24512 | 0.020986614 | Credible Set | purple |
| chr15:67304194 | 15 | 67.30419 | 0.021077940 | Credible Set | purple |
| chr15:67325463 | 15 | 67.32546 | 0.014546746 | Credible Set | purple |
| chr15:67331738 | 15 | 67.33174 | 0.026328169 | Credible Set | purple |
| chr15:67428857 | 15 | 67.42886 | 0.021852100 | Credible Set | purple |
| chr15:67429407 | 15 | 67.42941 | 0.017335645 | Credible Set | purple |
| chr15:67429420 | 15 | 67.42942 | 0.016288454 | Credible Set | purple |
| chr15:67433395 | 15 | 67.43340 | 0.023764279 | Credible Set | purple |
| chr15:67439974 | 15 | 67.43997 | 0.023625179 | Credible Set | purple |
| chr15:67530570 | 15 | 67.53057 | 0.024089841 | Credible Set | purple |
| chr15:67555683 | 15 | 67.55568 | 0.027120550 | Credible Set | purple |
| chr15:67627161 | 15 | 67.62716 | 0.027397710 | Credible Set | purple |
| chr15:67627183 | 15 | 67.62718 | 0.028716840 | Credible Set | purple |
| chr15:67627194 | 15 | 67.62719 | 0.028529663 | Credible Set | purple |
| chr15:67685224 | 15 | 67.68522 | 0.019665874 | Credible Set | purple |
| chr15:67685781 | 15 | 67.68578 | 0.016203904 | Credible Set | purple |
| chr15:67727448 | 15 | 67.72745 | 0.013448954 | Credible Set | purple |
| chr15:67812971 | 15 | 67.81297 | 0.014546955 | Credible Set | purple |
| chr15:67838708 | 15 | 67.83871 | 0.011277705 | Credible Set | purple |
| chr15:67838720 | 15 | 67.83872 | 0.010906976 | Credible Set | purple |
| chr15:67838721 | 15 | 67.83872 | 0.012305026 | Credible Set | purple |
| chr15:67935812 | 15 | 67.93581 | 0.006137792 | Credible Set | purple |
| chr15:67987913 | 15 | 67.98791 | 0.009345022 | Credible Set | purple |
| chr15:67987932 | 15 | 67.98793 | 0.009214896 | Credible Set | purple |
| chr15:68094377 | 15 | 68.09438 | 0.006410000 | Credible Set | purple |
| chr15:68145325 | 15 | 68.14532 | 0.003551565 | Credible Set | purple |

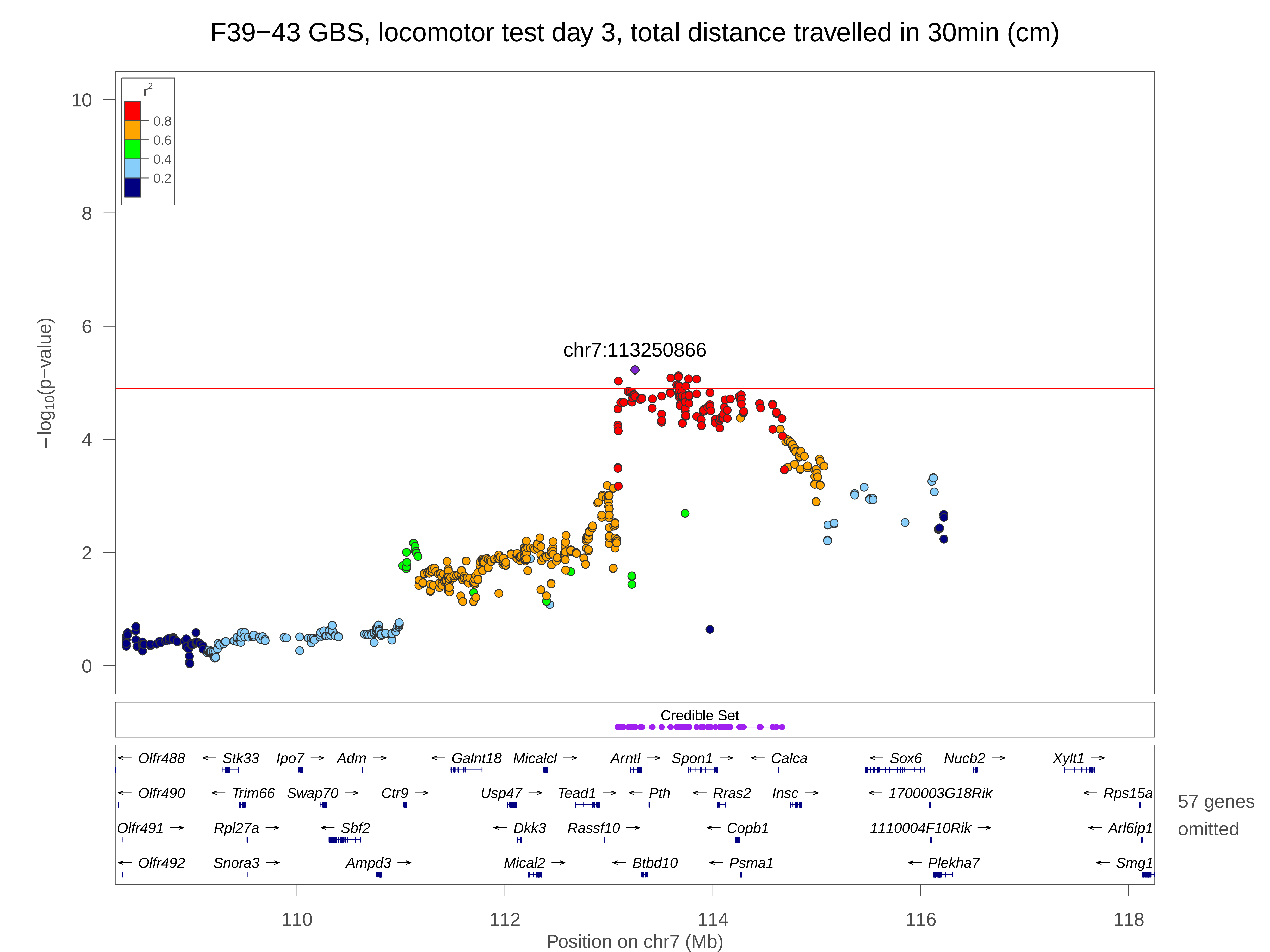

F39−43 GBS, locomotor test day 3, total distance travelled in 30min (cm)
r2
0.8
0.6
0.4
0.2
10
8
6
4
2
0
−log10(p−value)
chr7:113250866
Credible Set
← Olfr488
← Stk33
Ipo7 →
Adm →
← Galnt18
Micalcl →
Arntl →
Spon1 →
← Calca
← Sox6
Nucb2 →
Xylt1 →
← Olfr490
← Trim66
Swap70 →
Ctr9 →
Usp47 →
Tead1 →
← Pth
← Rras2
Insc →
← 1700003G18Rik
← Rps15a
Olfr491 →
Rpl27a →
← Sbf2
← Dkk3
Rassf10 →
← Copb1
1110004F10Rik →
← Arl6ip1
← Olfr492
Snora3 →
Ampd3 →
Mical2 →
← Btbd10
← Psma1
← Plekha7
← Smg1
57 genes omitted
110
112
114
116
118
Position on chr7 (Mb)

| | |
|---|---|
| date: | Thu Oct 25 17:22:48 2018 |
| build: | F3943GBS |
| display range: | chr7:108250866−118250866 [108250866−118250866] |
| hilite range: | 0 − 0 [ 0 − 0 ] |
| reference SNP: | chr7:113250866 |
| number of SNPs plotted: | 880 |
| min P−value: | 5.88E−6 [chr7:113250866] |
| max P−value: | 9.18E−1 [chr7:108970686] |
| omitted Genes: | Olfr493, Olfr494, Olfr495 |
| omitted Genes: | Olfr497, Olfr498, Olfr502 |
| omitted Genes: | Olfr503, Olfr504, Olfr506 |
| omitted Genes: | Olfr507, Olfr508, Olfr509 |
| omitted Genes: | Olfr510, Olfr512, Olfr513 |
| omitted Genes: | Olfr514, Olfr516, Olfr517 |
| omitted Genes: | Olfr518, Olfr519, Nlrp10 |
| omitted Genes: | Eif3f, Tub, Ric3 |
| omitted Genes: | Lmo1, St5, Akip1 |
| omitted Genes: | BC051019, Ascl3, Tmem9b |
| omitted Genes: | Nrip3, Scube2, Dennd5a |
| omitted Genes: | Tmem41b, 1600010M07Rik, Snora23 |
| omitted Genes: | Zfp143, Wee1, Rnf141 |
| omitted Genes: | Lyve1, Mrvi1, Gm16336 |
| omitted Genes: | Eif4g2, 1700012D14Rik, Parva |
| omitted Genes: | 4930543E12Rik, Far1, Far1os |
| omitted Genes: | 4933406I18Rik, Pde3b, Cyp2r1 |
| omitted Genes: | Calcb, A730082K24Rik, Rps13 |
| omitted Genes: | Snord14a, Pik3c2a, 4930583K01Rik |

Make more plots at http://csg.sph.umich.edu/locuszoom/

Fine−mapping Regions

| snp | chr | pos (Mb) | pp | group | color |
|---|---|---|---|---|---|
| chr7:113085544 | 7 | 113.0855 | 0.002796477 | Credible Set | purple |
| chr7:113085717 | 7 | 113.0857 | 0.002819713 | Credible Set | purple |
| chr7:113085735 | 7 | 113.0857 | 0.002568453 | Credible Set | purple |
| chr7:113086048 | 7 | 113.0860 | 0.005247494 | Credible Set | purple |
| chr7:113090456 | 7 | 113.0905 | 0.015481312 | Credible Set | purple |
| chr7:113114328 | 7 | 113.1143 | 0.006730116 | Credible Set | purple |
| chr7:113141980 | 7 | 113.1420 | 0.006781083 | Credible Set | purple |
| chr7:113184658 | 7 | 113.1847 | 0.010244835 | Credible Set | purple |
| chr7:113185148 | 7 | 113.1851 | 0.010362436 | Credible Set | purple |
| chr7:113203215 | 7 | 113.2032 | 0.010206688 | Credible Set | purple |
| chr7:113203253 | 7 | 113.2033 | 0.010182231 | Credible Set | purple |
| chr7:113220615 | 7 | 113.2206 | 0.010150578 | Credible Set | purple |
| chr7:113221228 | 7 | 113.2212 | 0.006839210 | Credible Set | purple |
| chr7:113229151 | 7 | 113.2292 | 0.008137461 | Credible Set | purple |
| chr7:113229207 | 7 | 113.2292 | 0.008079794 | Credible Set | purple |
| chr7:113235202 | 7 | 113.2352 | 0.009099652 | Credible Set | purple |
| chr7:113241920 | 7 | 113.2419 | 0.008951104 | Credible Set | purple |
| chr7:113242507 | 7 | 113.2425 | 0.008951104 | Credible Set | purple |
| chr7:113242523 | 7 | 113.2425 | 0.008951104 | Credible Set | purple |
| chr7:113245448 | 7 | 113.2454 | 0.009046167 | Credible Set | purple |
| chr7:113250866 | 7 | 113.2509 | 0.024015711 | Credible Set | purple |
| chr7:113251619 | 7 | 113.2516 | 0.008432309 | Credible Set | purple |
| chr7:113251923 | 7 | 113.2519 | 0.008431163 | Credible Set | purple |
| chr7:113300135 | 7 | 113.3001 | 0.007541746 | Credible Set | purple |
| chr7:113316522 | 7 | 113.3165 | 0.008003237 | Credible Set | purple |
| chr7:113316880 | 7 | 113.3169 | 0.007972719 | Credible Set | purple |
| chr7:113317049 | 7 | 113.3170 | 0.007820710 | Credible Set | purple |
| chr7:113416115 | 7 | 113.4161 | 0.005391595 | Credible Set | purple |
| chr7:113416126 | 7 | 113.4161 | 0.005415708 | Credible Set | purple |
| chr7:113419158 | 7 | 113.4192 | 0.007710598 | Credible Set | purple |
| chr7:113419175 | 7 | 113.4192 | 0.007776287 | Credible Set | purple |
| chr7:113506581 | 7 | 113.5066 | 0.004301741 | Credible Set | purple |
| chr7:113506703 | 7 | 113.5067 | 0.003137199 | Credible Set | purple |
| chr7:113506713 | 7 | 113.5067 | 0.003153893 | Credible Set | purple |
| chr7:113506834 | 7 | 113.5068 | 0.003366335 | Credible Set | purple |
| chr7:113507166 | 7 | 113.5072 | 0.008622940 | Credible Set | purple |

Fine−mapping Regions

| snp | chr | pos (Mb) | pp | group | color |
|---|---|---|---|---|---|
| chr7:113507171 | 7 | 113.5072 | 0.008695265 | Credible Set | purple |
| chr7:113507899 | 7 | 113.5079 | 0.008654981 | Credible Set | purple |
| chr7:113507930 | 7 | 113.5079 | 0.008654981 | Credible Set | purple |
| chr7:113591658 | 7 | 113.5917 | 0.009966100 | Credible Set | purple |
| chr7:113591661 | 7 | 113.5917 | 0.009599952 | Credible Set | purple |
| chr7:113595728 | 7 | 113.5957 | 0.017412622 | Credible Set | purple |
| chr7:113652590 | 7 | 113.6526 | 0.013349584 | Credible Set | purple |
| chr7:113668251 | 7 | 113.6683 | 0.013081478 | Credible Set | purple |
| chr7:113668346 | 7 | 113.6683 | 0.018986237 | Credible Set | purple |
| chr7:113668406 | 7 | 113.6684 | 0.018178367 | Credible Set | purple |
| chr7:113674001 | 7 | 113.6740 | 0.011109042 | Credible Set | purple |
| chr7:113674010 | 7 | 113.6740 | 0.012488081 | Credible Set | purple |
| chr7:113674041 | 7 | 113.6740 | 0.010135776 | Credible Set | purple |
| chr7:113674594 | 7 | 113.6746 | 0.008623906 | Credible Set | purple |
| chr7:113674603 | 7 | 113.6746 | 0.008342368 | Credible Set | purple |
| chr7:113685578 | 7 | 113.6856 | 0.008692391 | Credible Set | purple |
| chr7:113685734 | 7 | 113.6857 | 0.006284474 | Credible Set | purple |
| chr7:113685763 | 7 | 113.6858 | 0.005909508 | Credible Set | purple |
| chr7:113701178 | 7 | 113.7012 | 0.010191691 | Credible Set | purple |
| chr7:113701223 | 7 | 113.7012 | 0.010199906 | Credible Set | purple |
| chr7:113701622 | 7 | 113.7016 | 0.008505141 | Credible Set | purple |
| chr7:113701636 | 7 | 113.7016 | 0.008543398 | Credible Set | purple |
| chr7:113707091 | 7 | 113.7071 | 0.008729783 | Credible Set | purple |
| chr7:113707401 | 7 | 113.7074 | 0.008643357 | Credible Set | purple |
| chr7:113707540 | 7 | 113.7075 | 0.008720290 | Credible Set | purple |
| chr7:113707570 | 7 | 113.7076 | 0.003035583 | Credible Set | purple |
| chr7:113707711 | 7 | 113.7077 | 0.002992048 | Credible Set | purple |
| chr7:113732491 | 7 | 113.7325 | 0.008064347 | Credible Set | purple |
| chr7:113732536 | 7 | 113.7325 | 0.004901827 | Credible Set | purple |
| chr7:113732545 | 7 | 113.7325 | 0.004945548 | Credible Set | purple |
| chr7:113732598 | 7 | 113.7326 | 0.004834842 | Credible Set | purple |
| chr7:113732713 | 7 | 113.7327 | 0.008322569 | Credible Set | purple |
| chr7:113732753 | 7 | 113.7328 | 0.008443791 | Credible Set | purple |
| chr7:113732795 | 7 | 113.7328 | 0.005314738 | Credible Set | purple |
| chr7:113732809 | 7 | 113.7328 | 0.006881168 | Credible Set | purple |
| chr7:113733239 | 7 | 113.7332 | 0.003977621 | Credible Set | purple |

Fine−mapping Regions

| snp | chr | pos (Mb) | pp | group | color |
|---|---|---|---|---|---|
| chr7:113739150 | 7 | 113.7391 | 0.003957119 | Credible Set | purple |
| chr7:113739171 | 7 | 113.7392 | 0.004027427 | Credible Set | purple |
| chr7:113739179 | 7 | 113.7392 | 0.004065202 | Credible Set | purple |
| chr7:113739386 | 7 | 113.7394 | 0.012654550 | Credible Set | purple |
| chr7:113766388 | 7 | 113.7664 | 0.016892162 | Credible Set | purple |
| chr7:113766440 | 7 | 113.7664 | 0.016892162 | Credible Set | purple |
| chr7:113769934 | 7 | 113.7699 | 0.006550702 | Credible Set | purple |
| chr7:113770008 | 7 | 113.7700 | 0.009061291 | Credible Set | purple |
| chr7:113770031 | 7 | 113.7700 | 0.008562666 | Credible Set | purple |
| chr7:113770154 | 7 | 113.7702 | 0.008827398 | Credible Set | purple |
| chr7:113844950 | 7 | 113.8449 | 0.003918973 | Credible Set | purple |
| chr7:113844980 | 7 | 113.8450 | 0.009386433 | Credible Set | purple |
| chr7:113844999 | 7 | 113.8450 | 0.016683601 | Credible Set | purple |
| chr7:113887720 | 7 | 113.8877 | 0.003448300 | Credible Set | purple |
| chr7:113887792 | 7 | 113.8878 | 0.003495612 | Credible Set | purple |
| chr7:113890924 | 7 | 113.8909 | 0.002754082 | Credible Set | purple |
| chr7:113911095 | 7 | 113.9111 | 0.005141458 | Credible Set | purple |
| chr7:113911106 | 7 | 113.9111 | 0.004786978 | Credible Set | purple |
| chr7:113911113 | 7 | 113.9111 | 0.004989588 | Credible Set | purple |
| chr7:113950754 | 7 | 113.9508 | 0.005593657 | Credible Set | purple |
| chr7:113971635 | 7 | 113.9716 | 0.006208169 | Credible Set | purple |
| chr7:113971799 | 7 | 113.9718 | 0.009760761 | Credible Set | purple |
| chr7:113971835 | 7 | 113.9718 | 0.005758970 | Credible Set | purple |
| chr7:113971836 | 7 | 113.9718 | 0.005758970 | Credible Set | purple |
| chr7:113971886 | 7 | 113.9719 | 0.005758970 | Credible Set | purple |
| chr7:113980077 | 7 | 113.9801 | 0.004867257 | Credible Set | purple |
| chr7:114025271 | 7 | 114.0253 | 0.003474267 | Credible Set | purple |
| chr7:114025295 | 7 | 114.0253 | 0.003528738 | Credible Set | purple |
| chr7:114025329 | 7 | 114.0253 | 0.003488529 | Credible Set | purple |
| chr7:114025421 | 7 | 114.0254 | 0.003533123 | Credible Set | purple |
| chr7:114025449 | 7 | 114.0254 | 0.003031656 | Credible Set | purple |
| chr7:114064763 | 7 | 114.0648 | 0.003381599 | Credible Set | purple |
| chr7:114064790 | 7 | 114.0648 | 0.003381599 | Credible Set | purple |
| chr7:114065169 | 7 | 114.0652 | 0.003395443 | Credible Set | purple |
| chr7:114065223 | 7 | 114.0652 | 0.003379083 | Credible Set | purple |
| chr7:114065340 | 7 | 114.0653 | 0.003400781 | Credible Set | purple |

Fine−mapping Regions

| snp | chr | pos (Mb) | pp | group | color |
|---|---|---|---|---|---|
| chr7:114067633 | 7 | 114.0676 | 0.002516105 | Credible Set | purple |
| chr7:114072319 | 7 | 114.0723 | 0.003650765 | Credible Set | purple |
| chr7:114072371 | 7 | 114.0724 | 0.003649414 | Credible Set | purple |
| chr7:114087599 | 7 | 114.0876 | 0.003680860 | Credible Set | purple |
| chr7:114097503 | 7 | 114.0975 | 0.003679244 | Credible Set | purple |
| chr7:114103184 | 7 | 114.1032 | 0.004298548 | Credible Set | purple |
| chr7:114110302 | 7 | 114.1103 | 0.005565783 | Credible Set | purple |
| chr7:114117795 | 7 | 114.1178 | 0.007454142 | Credible Set | purple |
| chr7:114137930 | 7 | 114.1379 | 0.005002165 | Credible Set | purple |
| chr7:114138279 | 7 | 114.1383 | 0.003640964 | Credible Set | purple |
| chr7:114138316 | 7 | 114.1383 | 0.003649746 | Credible Set | purple |
| chr7:114167242 | 7 | 114.1672 | 0.007733764 | Credible Set | purple |
| chr7:114254780 | 7 | 114.2548 | 0.008477343 | Credible Set | purple |
| chr7:114254849 | 7 | 114.2548 | 0.008566051 | Credible Set | purple |
| chr7:114267072 | 7 | 114.2671 | 0.007429313 | Credible Set | purple |
| chr7:114272429 | 7 | 114.2724 | 0.009074710 | Credible Set | purple |
| chr7:114272945 | 7 | 114.2729 | 0.007657361 | Credible Set | purple |
| chr7:114272958 | 7 | 114.2730 | 0.006341523 | Credible Set | purple |
| chr7:114294236 | 7 | 114.2942 | 0.004606543 | Credible Set | purple |
| chr7:114294864 | 7 | 114.2949 | 0.004544244 | Credible Set | purple |
| chr7:114294958 | 7 | 114.2950 | 0.004716236 | Credible Set | purple |
| chr7:114295531 | 7 | 114.2955 | 0.004761816 | Credible Set | purple |
| chr7:114448685 | 7 | 114.4487 | 0.006479947 | Credible Set | purple |
| chr7:114459930 | 7 | 114.4599 | 0.005434514 | Credible Set | purple |
| chr7:114574313 | 7 | 114.5743 | 0.006357905 | Credible Set | purple |
| chr7:114574808 | 7 | 114.5748 | 0.006119974 | Credible Set | purple |
| chr7:114576812 | 7 | 114.5768 | 0.002409922 | Credible Set | purple |
| chr7:114611182 | 7 | 114.6112 | 0.004416014 | Credible Set | purple |
| chr7:114611200 | 7 | 114.6112 | 0.004596392 | Credible Set | purple |
| chr7:114664623 | 7 | 114.6646 | 0.003599431 | Credible Set | purple |

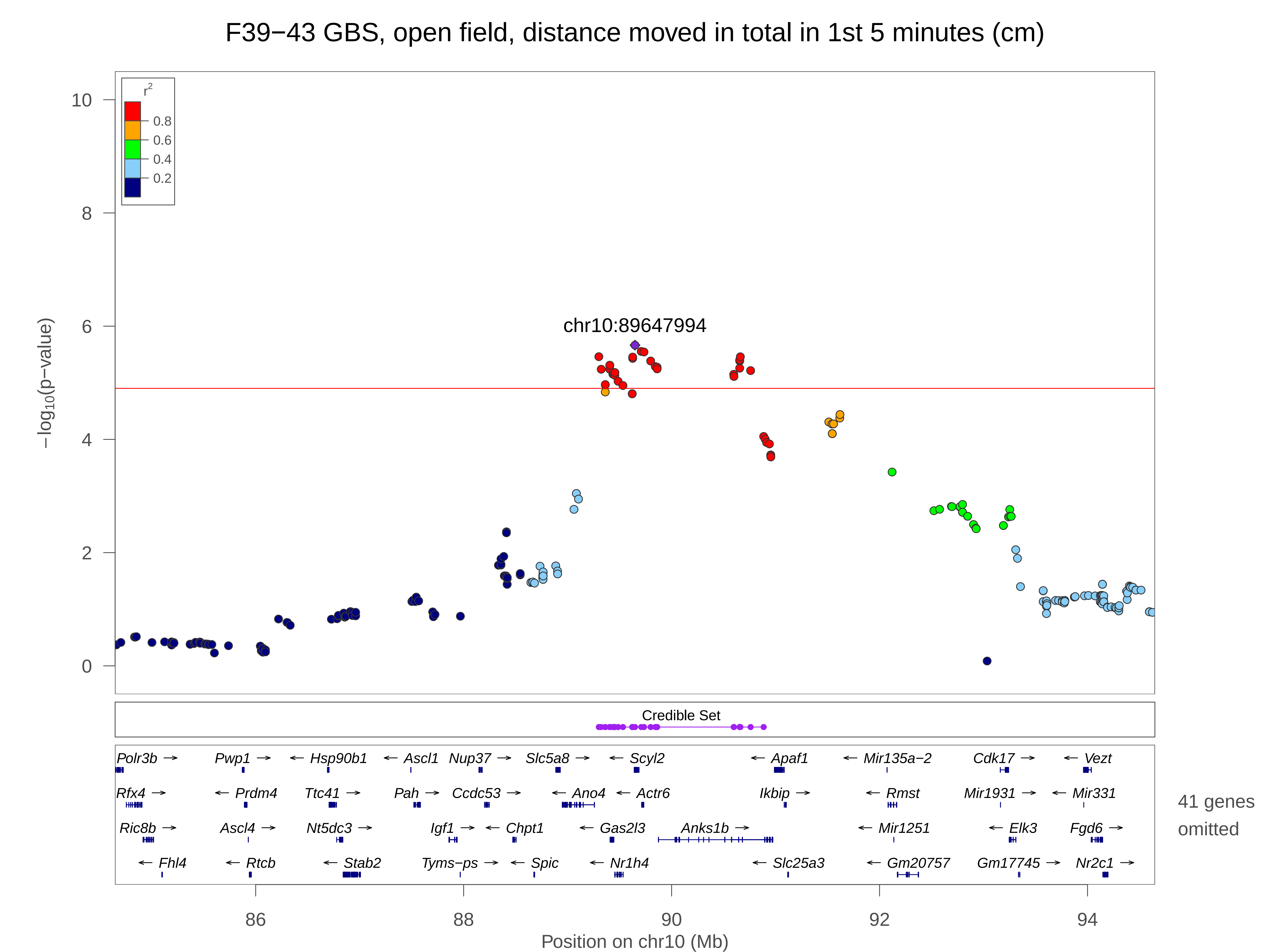
F39−43 GBS, open field, distance moved in total in 1st 5 minutes (cm)
r2
0.8
0.6
0.4
0.2
10
8
6
4
2
0
−log10(p−value)
chr10:89647994
Credible Set
Polr3b →
Pwp1 →
← Hsp90b1
← Ascl1
Nup37 →
Slc5a8 →
← Scyl2
← Apaf1
← Mir135a−2
Cdk17 →
← Vezt
Rfx4 →
← Prdm4
Ttc41 →
Pah →
Ccdc53 →
← Ano4
← Actr6
Ikbip →
← Rmst
Mir1931 →
← Mir331
Ric8b →
Ascl4 →
Nt5dc3 →
Igf1 →
← Chpt1
← Gas2l3
Anks1b →
← Mir1251
← Elk3
Fgd6 →
← Fhl4
← Rtcb
← Stab2
Tyms−ps →
← Spic
← Nr1h4
← Slc25a3
← Gm20757
Gm17745 →
Nr2c1 →
41 genes omitted
86
88
90
92
94
Position on chr10 (Mb)

| | |
|---|---|
| date: | Thu Oct 25 17:24:27 2018 |
| build: | F3943GBS |
| display range: | chr10:84647994−94647994 [84647994−94647994] |
| hilite range: | 0 − 0 [ 0 − 0 ] |
| reference SNP: | chr10:89647994 |
| number of SNPs plotted: | 282 |
| min P−value: | 2.17E−6 [chr10:89647994] |
| max P−value: | 8.21E−1 [chr10:93034582] |
| omitted Genes: | Tmem263, Mterf2, Cry1 |
| omitted Genes: | Btbd11, Bpifc, Fbxo7 |
| omitted Genes: | Syn3, 4930486F22Rik, Timp3 |
| omitted Genes: | 1810014B01Rik, 1700113H08Rik, Pmch |
| omitted Genes: | Parpbp, Dram1, Gnptab |
| omitted Genes: | Sycp3, Mybpc1, Gm4925 |
| omitted Genes: | Arl1, Utp20, Slc17a8 |
| omitted Genes: | 1500026H17Rik, Uhrf1bp1l, Tmpo |
| omitted Genes: | Nedd1, Mir7688, Lta4h |
| omitted Genes: | Hal, Amdhd1, Ccdc38 |
| omitted Genes: | Snrpf, 4933408J17Rik, Ntn4 |
| omitted Genes: | Gm8580, Gm15915, Usp44 |
| omitted Genes: | Metap2, Ndufa12, Gm4792 |
| omitted Genes: | Tmcc3, Mir7211 |

Make more plots at http://csg.sph.umich.edu/locuszoom/

Fine−mapping Regions

| snp | chr | pos (Mb) | pp | group | color |
|---|---|---|---|---|---|
| chr10:89300286 | 10 | 89.30029 | 0.039658176 | Credible Set | purple |
| chr10:89323084 | 10 | 89.32308 | 0.024301506 | Credible Set | purple |
| chr10:89362102 | 10 | 89.36210 | 0.013262560 | Credible Set | purple |
| chr10:89362162 | 10 | 89.36216 | 0.013402175 | Credible Set | purple |
| chr10:89405522 | 10 | 89.40552 | 0.028064254 | Credible Set | purple |
| chr10:89405533 | 10 | 89.40553 | 0.028049881 | Credible Set | purple |
| chr10:89405556 | 10 | 89.40556 | 0.024383361 | Credible Set | purple |
| chr10:89405564 | 10 | 89.40556 | 0.024383361 | Credible Set | purple |
| chr10:89405589 | 10 | 89.40559 | 0.028403912 | Credible Set | purple |
| chr10:89405627 | 10 | 89.40563 | 0.028468271 | Credible Set | purple |
| chr10:89435280 | 10 | 89.43528 | 0.020095981 | Credible Set | purple |
| chr10:89443390 | 10 | 89.44339 | 0.020508863 | Credible Set | purple |
| chr10:89454154 | 10 | 89.45415 | 0.019212661 | Credible Set | purple |
| chr10:89454848 | 10 | 89.45485 | 0.021379181 | Credible Set | purple |
| chr10:89485534 | 10 | 89.48553 | 0.015219465 | Credible Set | purple |
| chr10:89532131 | 10 | 89.53213 | 0.012905509 | Credible Set | purple |
| chr10:89621284 | 10 | 89.62128 | 0.009400969 | Credible Set | purple |
| chr10:89621319 | 10 | 89.62132 | 0.009334225 | Credible Set | purple |
| chr10:89625488 | 10 | 89.62549 | 0.037120864 | Credible Set | purple |
| chr10:89625559 | 10 | 89.62556 | 0.039059226 | Credible Set | purple |
| chr10:89647994 | 10 | 89.64799 | 0.062146943 | Credible Set | purple |
| chr10:89708192 | 10 | 89.70819 | 0.048669204 | Credible Set | purple |
| chr10:89734590 | 10 | 89.73459 | 0.047603407 | Credible Set | purple |
| chr10:89798643 | 10 | 89.79864 | 0.033517475 | Credible Set | purple |
| chr10:89843657 | 10 | 89.84366 | 0.026960457 | Credible Set | purple |
| chr10:89860498 | 10 | 89.86050 | 0.026344017 | Credible Set | purple |
| chr10:89860846 | 10 | 89.86085 | 0.024632428 | Credible Set | purple |
| chr10:90597510 | 10 | 90.59751 | 0.019974958 | Credible Set | purple |
| chr10:90599690 | 10 | 90.59969 | 0.018344301 | Credible Set | purple |
| chr10:90654718 | 10 | 90.65472 | 0.025324184 | Credible Set | purple |
| chr10:90654994 | 10 | 90.65499 | 0.033637282 | Credible Set | purple |
| chr10:90654998 | 10 | 90.65500 | 0.034502714 | Credible Set | purple |
| chr10:90655002 | 10 | 90.65500 | 0.033705251 | Credible Set | purple |
| chr10:90655039 | 10 | 90.65504 | 0.034927973 | Credible Set | purple |
| chr10:90661222 | 10 | 90.66122 | 0.039527761 | Credible Set | purple |
| chr10:90759813 | 10 | 90.75981 | 0.023054811 | Credible Set | purple |

Fine−mapping Regions

| snp | chr | pos (Mb) | pp | group | color |
|---|---|---|---|---|---|
| chr10:90885383 | 10 | 90.88538 | 0.001806502 | Credible Set | purple |

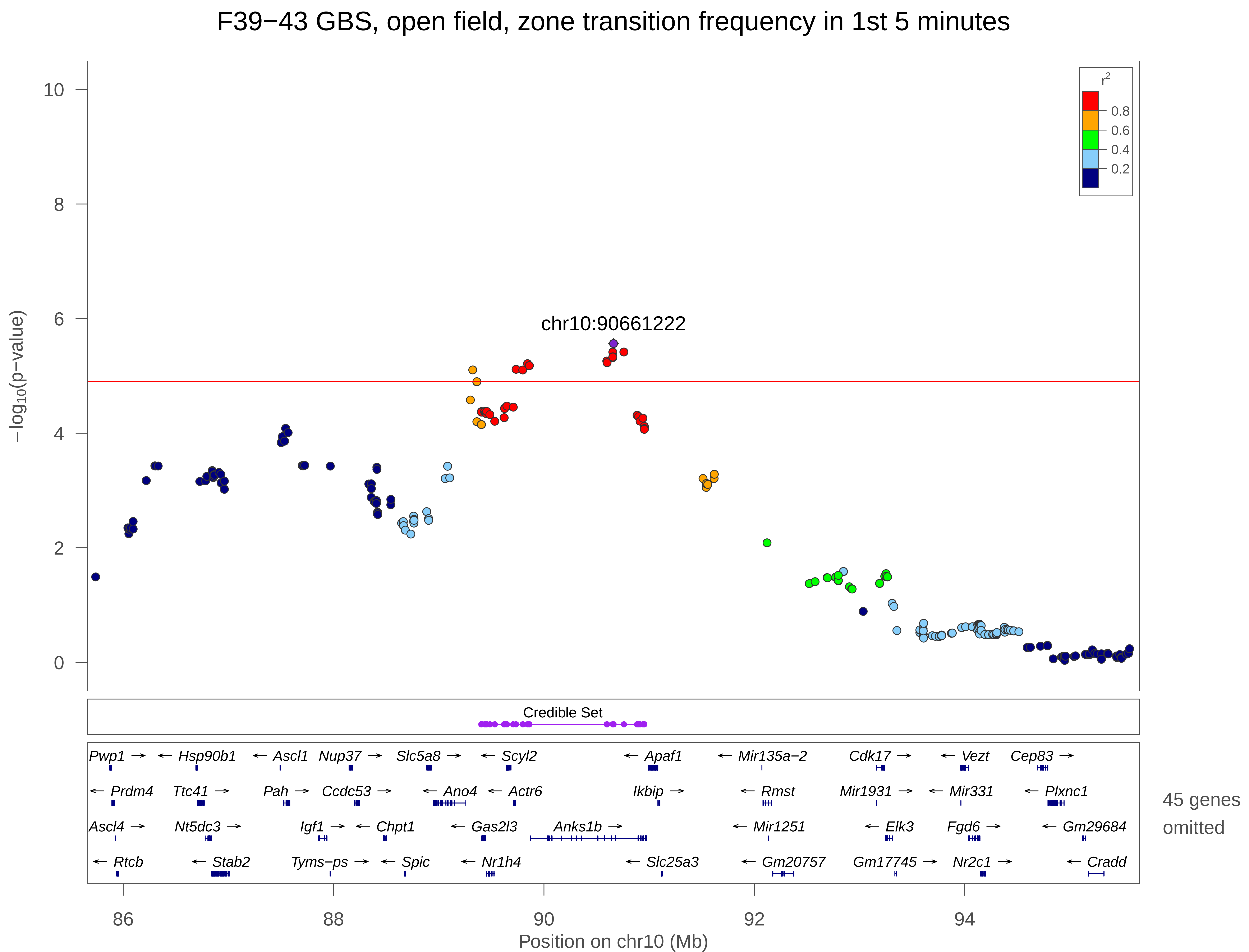
F39−43 GBS, open field, zone transition frequency in 1st 5 minutes
chr10:90661222
−log10(p−value)
r2
0.8
0.6
0.4
0.2
Credible Set
Pwp1 → ← Hsp90b1 ← Ascl1 Nup37 → Slc5a8 → ← Scyl2 ← Apaf1 ← Mir135a−2 Cdk17 → ← Vezt Cep83 →
← Prdm4 Ttc41 → Pah → Ccdc53 → ← Ano4 ← Actr6 Ikbip → ← Rmst Mir1931 → ← Mir331 ← Plxnc1
Ascl4 → Nt5dc3 → Igf1 → ← Chpt1 ← Gas2l3 Anks1b → ← Mir1251 ← Elk3 Fgd6 → ← Gm29684
← Rtcb ← Stab2 Tyms−ps → ← Spic ← Nr1h4 ← Slc25a3 ← Gm20757 Gm17745 → Nr2c1 → ← Cradd
45 genes omitted
Position on chr10 (Mb)

| | |
|---|---|
| date: | Thu Oct 25 17:24:36 2018 |
| build: | F3943GBS |
| display range: | chr10:85661222−95661222 [85661222−95661222] |
| hilite range: | 0 − 0 [ 0 − 0 ] |
| reference SNP: | chr10:90661222 |
| number of SNPs plotted: | 288 |
| min P−value: | 2.72E−6 [chr10:90661222] |
| max P−value: | 9.2E−1 [chr10:94950093] |
| omitted Genes: | Bpifc, Fbxo7, Syn3 |
| omitted Genes: | 4930486F22Rik, Timp3, 1810014B01Rik |
| omitted Genes: | 1700113H08Rik, Pmch, Parpbp |
| omitted Genes: | Dram1, Gnptab, Sycp3 |
| omitted Genes: | Mybpc1, Gm4925, Arl1 |
| omitted Genes: | Utp20, Slc17a8, 1500026H17Rik |
| omitted Genes: | Uhrf1bp1l, Tmpo, Nedd1 |
| omitted Genes: | Mir7688, Lta4h, Hal |
| omitted Genes: | Amdhd1, Ccdc38, Snrpf |
| omitted Genes: | 4933408J17Rik, Ntn4, Gm8580 |
| omitted Genes: | Gm15915, Usp44, Metap2 |
| omitted Genes: | Ndufa12, Gm4792, Tmcc3 |
| omitted Genes: | Mir7211, Cep83os, 2310039L15Rik |
| omitted Genes: | Socs2, 5730420D15Rik, Mrpl42 |
| omitted Genes: | Ube2n, Nudt4, Mir3058 |

Make more plots at http://csg.sph.umich.edu/locuszoom/

Fine−mapping Regions

| snp | chr | pos (Mb) | pp | group | color |
|---|---|---|---|---|---|
| chr10:89405556 | 10 | 89.40556 | 0.007778265 | Credible Set | purple |
| chr10:89405564 | 10 | 89.40556 | 0.007778265 | Credible Set | purple |
| chr10:89405589 | 10 | 89.40559 | 0.007649517 | Credible Set | purple |
| chr10:89405627 | 10 | 89.40563 | 0.007652497 | Credible Set | purple |
| chr10:89435280 | 10 | 89.43528 | 0.007793765 | Credible Set | purple |
| chr10:89443390 | 10 | 89.44339 | 0.007345316 | Credible Set | purple |
| chr10:89454154 | 10 | 89.45415 | 0.007184423 | Credible Set | purple |
| chr10:89454848 | 10 | 89.45485 | 0.007849913 | Credible Set | purple |
| chr10:89485534 | 10 | 89.48553 | 0.006939689 | Credible Set | purple |
| chr10:89532131 | 10 | 89.53213 | 0.005397014 | Credible Set | purple |
| chr10:89621284 | 10 | 89.62128 | 0.006175437 | Credible Set | purple |
| chr10:89621319 | 10 | 89.62132 | 0.006176909 | Credible Set | purple |
| chr10:89625488 | 10 | 89.62549 | 0.008845867 | Credible Set | purple |
| chr10:89625559 | 10 | 89.62556 | 0.008757609 | Credible Set | purple |
| chr10:89647994 | 10 | 89.64799 | 0.009630289 | Credible Set | purple |
| chr10:89708192 | 10 | 89.70819 | 0.009238231 | Credible Set | purple |
| chr10:89734590 | 10 | 89.73459 | 0.039442741 | Credible Set | purple |
| chr10:89798643 | 10 | 89.79864 | 0.038333385 | Credible Set | purple |
| chr10:89843657 | 10 | 89.84366 | 0.048954795 | Credible Set | purple |
| chr10:89860498 | 10 | 89.86050 | 0.045276879 | Credible Set | purple |
| chr10:89860846 | 10 | 89.86085 | 0.045378919 | Credible Set | purple |
| chr10:90597510 | 10 | 90.59751 | 0.054112353 | Credible Set | purple |
| chr10:90599690 | 10 | 90.59969 | 0.050624147 | Credible Set | purple |
| chr10:90654718 | 10 | 90.65472 | 0.076450300 | Credible Set | purple |
| chr10:90654994 | 10 | 90.65499 | 0.061975512 | Credible Set | purple |
| chr10:90654998 | 10 | 90.65500 | 0.063650490 | Credible Set | purple |
| chr10:90655002 | 10 | 90.65500 | 0.061862432 | Credible Set | purple |
| chr10:90655039 | 10 | 90.65504 | 0.063120913 | Credible Set | purple |
| chr10:90661222 | 10 | 90.66122 | 0.106070538 | Credible Set | purple |
| chr10:90759813 | 10 | 90.75981 | 0.076575883 | Credible Set | purple |
| chr10:90885383 | 10 | 90.88538 | 0.006802878 | Credible Set | purple |
| chr10:90899946 | 10 | 90.89995 | 0.006402570 | Credible Set | purple |
| chr10:90913583 | 10 | 90.91358 | 0.005417445 | Credible Set | purple |
| chr10:90940720 | 10 | 90.94072 | 0.006087540 | Credible Set | purple |
| chr10:90953385 | 10 | 90.95338 | 0.004347358 | Credible Set | purple |
| chr10:90953402 | 10 | 90.95340 | 0.004401991 | Credible Set | purple |

Fine−mapping Regions

| snp | chr | pos (Mb) | pp | group | color |
| --- | --- | --- | --- | --- | --- |
| chr10:90953425 | 10 | 90.95342 | 0.004457171 | Credible Set | purple |

F39−43 GBS, open field, immobile duration in 1st 5 minutes (s)

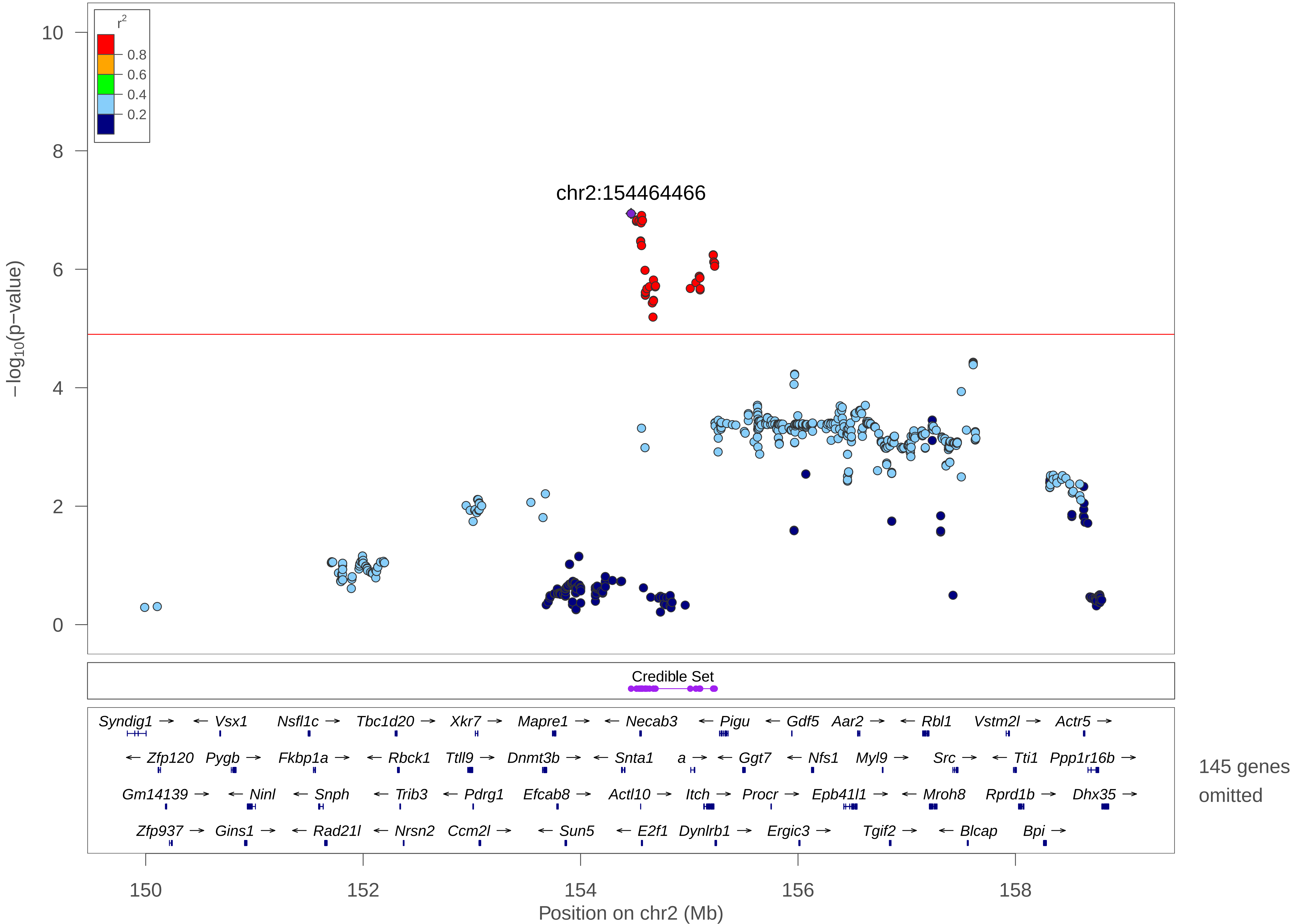

date: Thu Oct 25 17:24:08 2018
build: F3943GBS
display range: chr2:149464466−159464466 [149464466−159464466]
hilite range: 0 − 0 [ 0 − 0 ]
reference SNP: chr2:154464466
number of SNPs plotted: 757
min P−value: 1.15E−7 [chr2:154464466]
max P−value: 6.13E−1 [chr2:154734707]
omitted Genes: Gm14124, 3300002I08Rik, Zfp442
omitted Genes: Zfp345, Cst7, Apmap
omitted Genes: Acss1, E130215H24Rik, Entpd6
omitted Genes: Abhd12, Nanp, Gm14151
omitted Genes: 4930442J19Rik, LOC100125594, 4921509C19Rik
omitted Genes: Sdcbp2, Tmem74b, Psmf1
omitted Genes: Rspo4, Angpt4, Mir1953
omitted Genes: Fam110a, Slc52a3, Scrt2
omitted Genes: Srxn1, Tcf15, Csnk2a1
omitted Genes: Gm14164, Sox12, Zcchc3
omitted Genes: 6820408C15Rik, Defb23, Defb20
omitted Genes: Defb22, Defb26, Defb28
omitted Genes: Defb29, Defb21, Defb19
omitted Genes: Defb45, Defb36, Defb25
omitted Genes: Rem1, H13, Mcts2
omitted Genes: Id1, Cox4i2, Bcl2l1
omitted Genes: Tpx2, Mir5622, Mylk2
omitted Genes: Foxs1, Dusp15, Hck
omitted Genes: Tm9sf4, Tspyl3, Plagl2
omitted Genes: Pofut1, Kif3b, 2500004C02Rik
omitted Genes: Asxl1, Nol4l, Commd7
omitted Genes: Bpifb2, Bpifb6, Bpifb3
omitted Genes: Bpifb4, Bpifa6, Bpifa2
omitted Genes: Bpifa3, Bpifa1, Bpifa5
omitted Genes: Bpifb1, Bpifb5, Bpifb9a
omitted Genes: Bpifb9b, Cdk5rap1, Cbfa2t2
omitted Genes: 1700003F12Rik, Pxmp4, Zfp341
omitted Genes: Chmp4b, Raly, Eif2s2
omitted Genes: Ahcy, 2310005A03Rik, Map1lc3a
omitted Genes: Mir695, Trp53inp2, Ncoa6
omitted Genes: Acss2, Acss2os, Gss
Make more plots at http://csg.sph.umich.edu/locuszoom/
omitted Genes: Myh7b, Mir499, Trpc4ap

Fine−mapping Regions

| snp | chr | pos (Mb) | pp | group | color |
|---|---|---|---|---|---|
| chr2:154464466 | 2 | 154.4645 | 0.077380251 | Credible Set | purple |
| chr2:154513996 | 2 | 154.5140 | 0.057628986 | Credible Set | purple |
| chr2:154515752 | 2 | 154.5158 | 0.060925709 | Credible Set | purple |
| chr2:154516249 | 2 | 154.5162 | 0.059631478 | Credible Set | purple |
| chr2:154537995 | 2 | 154.5380 | 0.058400539 | Credible Set | purple |
| chr2:154538009 | 2 | 154.5380 | 0.058348012 | Credible Set | purple |
| chr2:154552607 | 2 | 154.5526 | 0.027660183 | Credible Set | purple |
| chr2:154552612 | 2 | 154.5526 | 0.027063118 | Credible Set | purple |
| chr2:154556468 | 2 | 154.5565 | 0.054353119 | Credible Set | purple |
| chr2:154560354 | 2 | 154.5604 | 0.023289374 | Credible Set | purple |
| chr2:154560360 | 2 | 154.5604 | 0.023197502 | Credible Set | purple |
| chr2:154560680 | 2 | 154.5607 | 0.064741829 | Credible Set | purple |
| chr2:154561127 | 2 | 154.5611 | 0.071957422 | Credible Set | purple |
| chr2:154569018 | 2 | 154.5690 | 0.059729276 | Credible Set | purple |
| chr2:154569036 | 2 | 154.5690 | 0.059430630 | Credible Set | purple |
| chr2:154592528 | 2 | 154.5925 | 0.009192320 | Credible Set | purple |
| chr2:154595741 | 2 | 154.5957 | 0.004013589 | Credible Set | purple |
| chr2:154595774 | 2 | 154.5958 | 0.003612895 | Credible Set | purple |
| chr2:154596294 | 2 | 154.5963 | 0.003638474 | Credible Set | purple |
| chr2:154597124 | 2 | 154.5971 | 0.003979535 | Credible Set | purple |
| chr2:154597483 | 2 | 154.5975 | 0.004092658 | Credible Set | purple |
| chr2:154610318 | 2 | 154.6103 | 0.004630510 | Credible Set | purple |
| chr2:154633945 | 2 | 154.6339 | 0.004990385 | Credible Set | purple |
| chr2:154670525 | 2 | 154.6705 | 0.002992188 | Credible Set | purple |
| chr2:154670528 | 2 | 154.6705 | 0.002995686 | Credible Set | purple |
| chr2:154670571 | 2 | 154.6706 | 0.006387038 | Credible Set | purple |
| chr2:154686815 | 2 | 154.6868 | 0.004948734 | Credible Set | purple |
| chr2:154689824 | 2 | 154.6898 | 0.005134110 | Credible Set | purple |
| chr2:155009299 | 2 | 155.0093 | 0.004663011 | Credible Set | purple |
| chr2:155060609 | 2 | 155.0606 | 0.005778488 | Credible Set | purple |
| chr2:155091628 | 2 | 155.0916 | 0.007372176 | Credible Set | purple |
| chr2:155097606 | 2 | 155.0976 | 0.007059231 | Credible Set | purple |
| chr2:155097628 | 2 | 155.0976 | 0.006897689 | Credible Set | purple |
| chr2:155097650 | 2 | 155.0976 | 0.006904636 | Credible Set | purple |
| chr2:155099477 | 2 | 155.0995 | 0.004414824 | Credible Set | purple |
| chr2:155099522 | 2 | 155.0995 | 0.004621923 | Credible Set | purple |

Fine−mapping Regions

| snp | chr | pos (Mb) | pp | group | color |
|---|---|---|---|---|---|
| chr2:155220007 | 2 | 155.2200 | 0.01639115 | Credible Set | purple |
| chr2:155220040 | 2 | 155.2200 | 0.01639115 | Credible Set | purple |
| chr2:155220051 | 2 | 155.2201 | 0.01639115 | Credible Set | purple |
| chr2:155220725 | 2 | 155.2207 | 0.01621069 | Credible Set | purple |
| chr2:155224800 | 2 | 155.2248 | 0.01256823 | Credible Set | purple |
| chr2:155233215 | 2 | 155.2332 | 0.01202755 | Credible Set | purple |
| chr2:155233265 | 2 | 155.2333 | 0.01070872 | Credible Set | purple |

F39−43 GBS, open field, immobile duration in 1st 5 minutes (s)

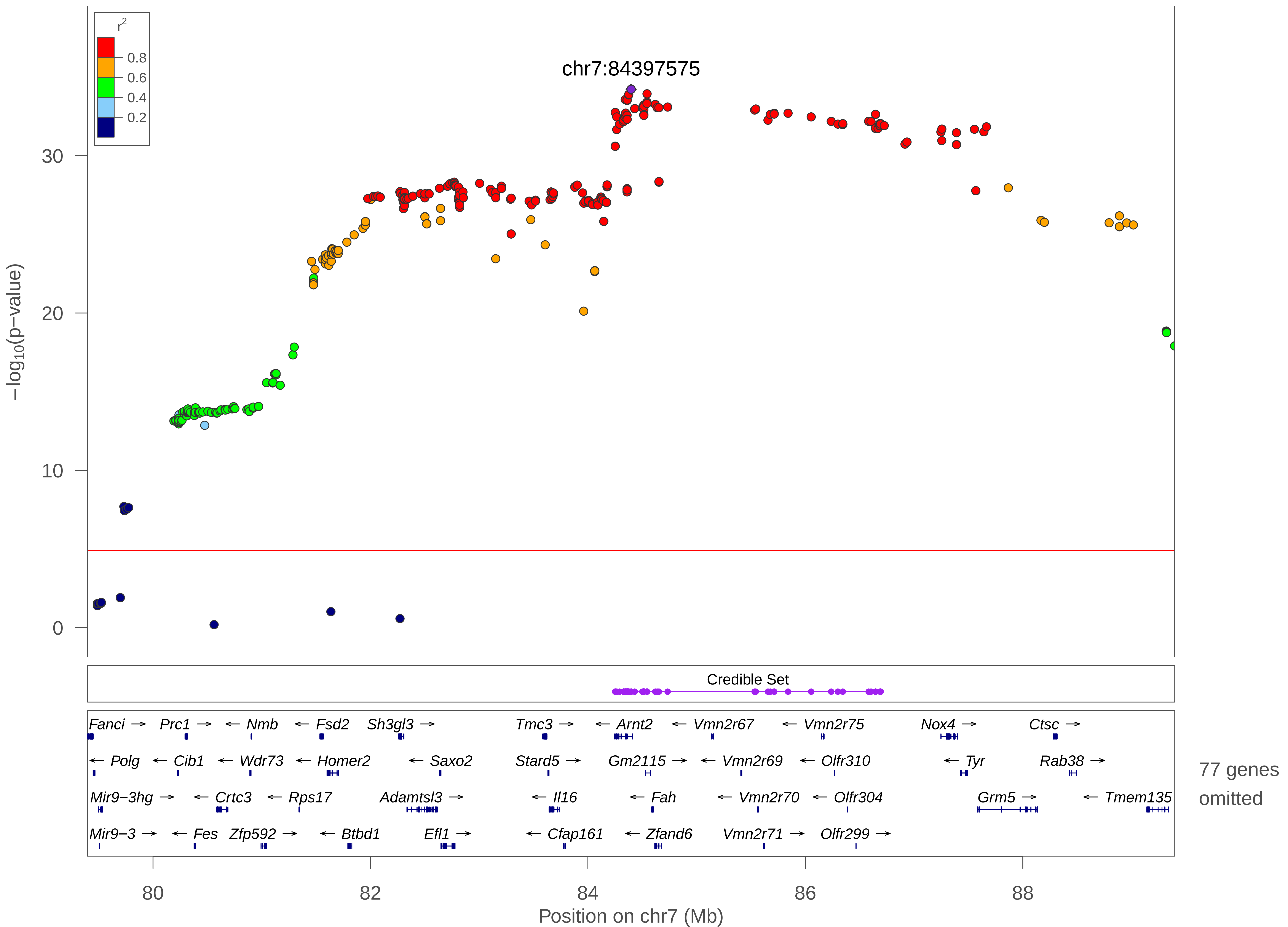

| | |
|---|---|
| date: | Thu Oct 25 17:24:18 2018 |
| build: | F3943GBS |
| display range: | chr7:79397575−89397575 [79397575−89397575] |
| hilite range: | 0 − 0 [ 0 − 0 ] |
| reference SNP: | chr7:84397575 |
| number of SNPs plotted: | 396 |
| min P−value: | 5.92E−35 [chr7:84397575] |
| max P−value: | 6.44E−1 [chr7:80561830] |
| omitted Genes: | Rhcg, Ticrr, Kif7 |
| omitted Genes: | Plin1, Pex11a, Wdr93 |
| omitted Genes: | Mesp1, Mesp2, Anpep |
| omitted Genes: | Ap3s2, Arpin, Zfp710 |
| omitted Genes: | Gm21057, Idh2, Sema4b |
| omitted Genes: | Gdpgp1, Ttll13, Ngrn |
| omitted Genes: | Vps33b, Rccd1, Unc45a |
| omitted Genes: | Hddc3, Man2a2, Furin |
| omitted Genes: | Blm, Gm15880, Iqgap1 |
| omitted Genes: | Zscan2, Sec11a, Alpk3 |
| omitted Genes: | Slc28a1, Pde8a, Cpeb1 |
| omitted Genes: | Ap3b2, BC048679, 2900076A07Rik |
| omitted Genes: | Mir1839, Whamm, Fam103a1 |
| omitted Genes: | 3110040N11Rik, Tm6sf1, Hdgfrp3 |
| omitted Genes: | Bnc1, Gm20744, 4933406J10Rik |
| omitted Genes: | Mex3b, 4933430H16Rik, Mesdc1 |
| omitted Genes: | Mesdc2, Cemip, Abhd17c |
| omitted Genes: | 2610206C17Rik, Olfr291, Olfr290 |
| omitted Genes: | Vmn2r65, Vmn2r66, Vmn2r68 |
| omitted Genes: | Vmn2r72, Vmn2r73, Vmn2r74 |
| omitted Genes: | Vmn2r76, Olfr309, Olfr308 |
| omitted Genes: | Olfr307, Olfr305, Olfr303 |
| omitted Genes: | Olfr301, Olfr298, Olfr297 |
| omitted Genes: | Olfr295, Olfr294, Olfr293 |
| omitted Genes: | Olfr292, Folh1, Vmn2r77 |
| omitted Genes: | Vmn2r78, Vmn2r79 |

Make more plots at http://csg.sph.umich.edu/locuszoom/

Fine−mapping Regions

| snp | chr | pos (Mb) | pp | group | color |
|---|---|---|---|---|---|
| chr7:84250985 | 7 | 84.25099 | 0.0052463340 | Credible Set | purple |
| chr7:84251011 | 7 | 84.25101 | 0.0053700774 | Credible Set | purple |
| chr7:84265685 | 7 | 84.26569 | 0.0027673352 | Credible Set | purple |
| chr7:84291540 | 7 | 84.29154 | 0.0009713545 | Credible Set | purple |
| chr7:84327288 | 7 | 84.32729 | 0.0014225509 | Credible Set | purple |
| chr7:84327808 | 7 | 84.32781 | 0.0020139860 | Credible Set | purple |
| chr7:84327850 | 7 | 84.32785 | 0.0017832356 | Credible Set | purple |
| chr7:84338595 | 7 | 84.33859 | 0.0027256758 | Credible Set | purple |
| chr7:84343118 | 7 | 84.34312 | 0.0345440729 | Credible Set | purple |
| chr7:84344805 | 7 | 84.34480 | 0.0036057811 | Credible Set | purple |
| chr7:84347423 | 7 | 84.34742 | 0.0048578487 | Credible Set | purple |
| chr7:84360038 | 7 | 84.36004 | 0.0035380346 | Credible Set | purple |
| chr7:84360626 | 7 | 84.36063 | 0.0019959784 | Credible Set | purple |
| chr7:84360642 | 7 | 84.36064 | 0.0019619519 | Credible Set | purple |
| chr7:84360703 | 7 | 84.36070 | 0.0302445200 | Credible Set | purple |
| chr7:84360714 | 7 | 84.36071 | 0.0327604981 | Credible Set | purple |
| chr7:84360716 | 7 | 84.36072 | 0.0327604981 | Credible Set | purple |
| chr7:84375526 | 7 | 84.37553 | 0.0716098398 | Credible Set | purple |
| chr7:84375577 | 7 | 84.37558 | 0.0682758895 | Credible Set | purple |
| chr7:84375642 | 7 | 84.37564 | 0.0728765690 | Credible Set | purple |
| chr7:84397575 | 7 | 84.39758 | 0.1562491568 | Credible Set | purple |
| chr7:84397597 | 7 | 84.39760 | 0.1519139229 | Credible Set | purple |
| chr7:84430102 | 7 | 84.43010 | 0.0093372654 | Credible Set | purple |
| chr7:84501082 | 7 | 84.50108 | 0.0104880653 | Credible Set | purple |
| chr7:84514082 | 7 | 84.51408 | 0.0157884220 | Credible Set | purple |
| chr7:84514444 | 7 | 84.51444 | 0.0081297329 | Credible Set | purple |
| chr7:84514505 | 7 | 84.51450 | 0.0081344474 | Credible Set | purple |
| chr7:84514564 | 7 | 84.51456 | 0.0039338436 | Credible Set | purple |
| chr7:84514575 | 7 | 84.51457 | 0.0034559980 | Credible Set | purple |
| chr7:84514734 | 7 | 84.51473 | 0.0129532981 | Credible Set | purple |
| chr7:84543346 | 7 | 84.54335 | 0.0235612271 | Credible Set | purple |
| chr7:84543355 | 7 | 84.54336 | 0.0195719118 | Credible Set | purple |
| chr7:84543782 | 7 | 84.54378 | 0.0800154583 | Credible Set | purple |
| chr7:84619632 | 7 | 84.61963 | 0.0169651351 | Credible Set | purple |
| chr7:84636759 | 7 | 84.63676 | 0.0106453849 | Credible Set | purple |
| chr7:84653095 | 7 | 84.65309 | 0.0105528889 | Credible Set | purple |

Fine−mapping Regions

| snp | chr | pos (Mb) | pp | group | color |
|---|---|---|---|---|---|
| chr7:84733186 | 7 | 84.73319 | 0.0117010324 | Credible Set | purple |
| chr7:85533207 | 7 | 85.53321 | 0.0075770159 | Credible Set | purple |
| chr7:85543843 | 7 | 85.54384 | 0.0088065401 | Credible Set | purple |
| chr7:85656243 | 7 | 85.65624 | 0.0017286559 | Credible Set | purple |
| chr7:85677494 | 7 | 85.67749 | 0.0039911693 | Credible Set | purple |
| chr7:85712663 | 7 | 85.71266 | 0.0047372285 | Credible Set | purple |
| chr7:85712975 | 7 | 85.71298 | 0.0042050829 | Credible Set | purple |
| chr7:85840665 | 7 | 85.84067 | 0.0047691707 | Credible Set | purple |
| chr7:86053386 | 7 | 86.05339 | 0.0027882394 | Credible Set | purple |
| chr7:86237374 | 7 | 86.23737 | 0.0014626507 | Credible Set | purple |
| chr7:86298979 | 7 | 86.29898 | 0.0009803139 | Credible Set | purple |
| chr7:86344488 | 7 | 86.34449 | 0.0010427187 | Credible Set | purple |
| chr7:86582139 | 7 | 86.58214 | 0.0014691844 | Credible Set | purple |
| chr7:86603770 | 7 | 86.60377 | 0.0014033521 | Credible Set | purple |
| chr7:86645719 | 7 | 86.64572 | 0.0040095552 | Credible Set | purple |
| chr7:86645792 | 7 | 86.64579 | 0.0040968628 | Credible Set | purple |
| chr7:86685648 | 7 | 86.68565 | 0.0010582767 | Credible Set | purple |
| chr7:86692214 | 7 | 86.69221 | 0.0010525011 | Credible Set | purple |
| chr7:86692491 | 7 | 86.69249 | 0.0009860399 | Credible Set | purple |

F39−43 GBS, open field, distance moved in total in 10 minutes (cm)

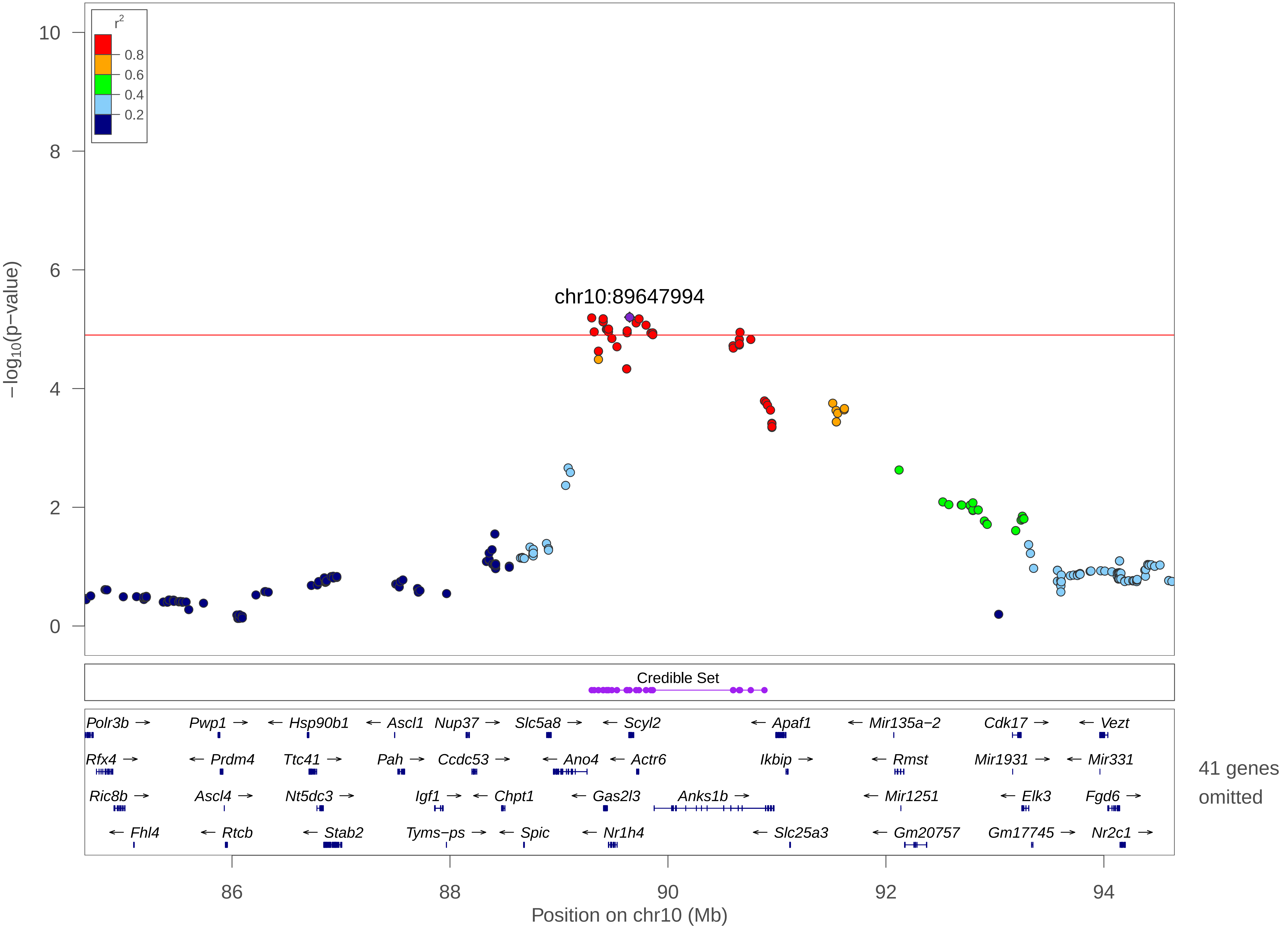

| | |
|---|---|
| date: | Thu Oct 25 17:23:20 2018 |
| build: | F3943GBS |
| display range: | chr10:84647994−94647994 [84647994−94647994] |
| hilite range: | 0 − 0 [ 0 − 0 ] |
| reference SNP: | chr10:89647994 |
| number of SNPs plotted: | 282 |
| min P−value: | 6.28E−6 [chr10:89647994] |
| max P−value: | 7.46E−1 [chr10:86053984] |
| omitted Genes: | Tmem263, Mterf2, Cry1 |
| omitted Genes: | Btbd11, Bpifc, Fbxo7 |
| omitted Genes: | Syn3, 4930486F22Rik, Timp3 |
| omitted Genes: | 1810014B01Rik, 1700113H08Rik, Pmch |
| omitted Genes: | Parpbp, Dram1, Gnptab |
| omitted Genes: | Sycp3, Mybpc1, Gm4925 |
| omitted Genes: | Arl1, Utp20, Slc17a8 |
| omitted Genes: | 1500026H17Rik, Uhrf1bp1l, Tmpo |
| omitted Genes: | Nedd1, Mir7688, Lta4h |
| omitted Genes: | Hal, Amdhd1, Ccdc38 |
| omitted Genes: | Snrpf, 4933408J17Rik, Ntn4 |
| omitted Genes: | Gm8580, Gm15915, Usp44 |
| omitted Genes: | Metap2, Ndufa12, Gm4792 |
| omitted Genes: | Tmcc3, Mir7211 |

Make more plots at http://csg.sph.umich.edu/locuszoom/

Fine−mapping Regions

| snp | chr | pos (Mb) | pp | group | color |
|---|---|---|---|---|---|
| chr10:89300286 | 10 | 89.30029 | 0.046195711 | Credible Set | purple |
| chr10:89323084 | 10 | 89.32308 | 0.027506654 | Credible Set | purple |
| chr10:89362102 | 10 | 89.36210 | 0.013374071 | Credible Set | purple |
| chr10:89362162 | 10 | 89.36216 | 0.013495842 | Credible Set | purple |
| chr10:89405522 | 10 | 89.40552 | 0.040687353 | Credible Set | purple |
| chr10:89405533 | 10 | 89.40553 | 0.040386541 | Credible Set | purple |
| chr10:89405556 | 10 | 89.40556 | 0.039848670 | Credible Set | purple |
| chr10:89405564 | 10 | 89.40556 | 0.039848670 | Credible Set | purple |
| chr10:89405589 | 10 | 89.40559 | 0.044574853 | Credible Set | purple |
| chr10:89405627 | 10 | 89.40563 | 0.044658488 | Credible Set | purple |
| chr10:89435280 | 10 | 89.43528 | 0.030235851 | Credible Set | purple |
| chr10:89443390 | 10 | 89.44339 | 0.029877098 | Credible Set | purple |
| chr10:89454154 | 10 | 89.45415 | 0.027594433 | Credible Set | purple |
| chr10:89454848 | 10 | 89.45485 | 0.030576768 | Credible Set | purple |
| chr10:89485534 | 10 | 89.48553 | 0.021521170 | Credible Set | purple |
| chr10:89532131 | 10 | 89.53213 | 0.015870127 | Credible Set | purple |
| chr10:89621284 | 10 | 89.62128 | 0.007052658 | Credible Set | purple |
| chr10:89621319 | 10 | 89.62132 | 0.007003513 | Credible Set | purple |
| chr10:89625488 | 10 | 89.62549 | 0.026605413 | Credible Set | purple |
| chr10:89625559 | 10 | 89.62556 | 0.028651320 | Credible Set | purple |
| chr10:89647994 | 10 | 89.64799 | 0.047396078 | Credible Set | purple |
| chr10:89708192 | 10 | 89.70819 | 0.038350982 | Credible Set | purple |
| chr10:89734590 | 10 | 89.73459 | 0.044560120 | Credible Set | purple |
| chr10:89798643 | 10 | 89.79864 | 0.035307665 | Credible Set | purple |
| chr10:89843657 | 10 | 89.84366 | 0.026328612 | Credible Set | purple |
| chr10:89860498 | 10 | 89.86050 | 0.026503471 | Credible Set | purple |
| chr10:89860846 | 10 | 89.86085 | 0.024741475 | Credible Set | purple |
| chr10:90597510 | 10 | 90.59751 | 0.016448636 | Credible Set | purple |
| chr10:90599690 | 10 | 90.59969 | 0.015101222 | Credible Set | purple |
| chr10:90654718 | 10 | 90.65472 | 0.020831486 | Credible Set | purple |
| chr10:90654994 | 10 | 90.65499 | 0.016918577 | Credible Set | purple |
| chr10:90654998 | 10 | 90.65500 | 0.017341253 | Credible Set | purple |
| chr10:90655002 | 10 | 90.65500 | 0.016952658 | Credible Set | purple |
| chr10:90655039 | 10 | 90.65504 | 0.017621113 | Credible Set | purple |
| chr10:90661222 | 10 | 90.66122 | 0.027101088 | Credible Set | purple |
| chr10:90759813 | 10 | 90.75981 | 0.020780488 | Credible Set | purple |

Fine−mapping Regions

| snp | chr | pos (Mb) | pp | group | color |
|---|---|---|---|---|---|
| chr10:90885383 | 10 | 90.88538 | 0.002169931 | Credible Set | purple |

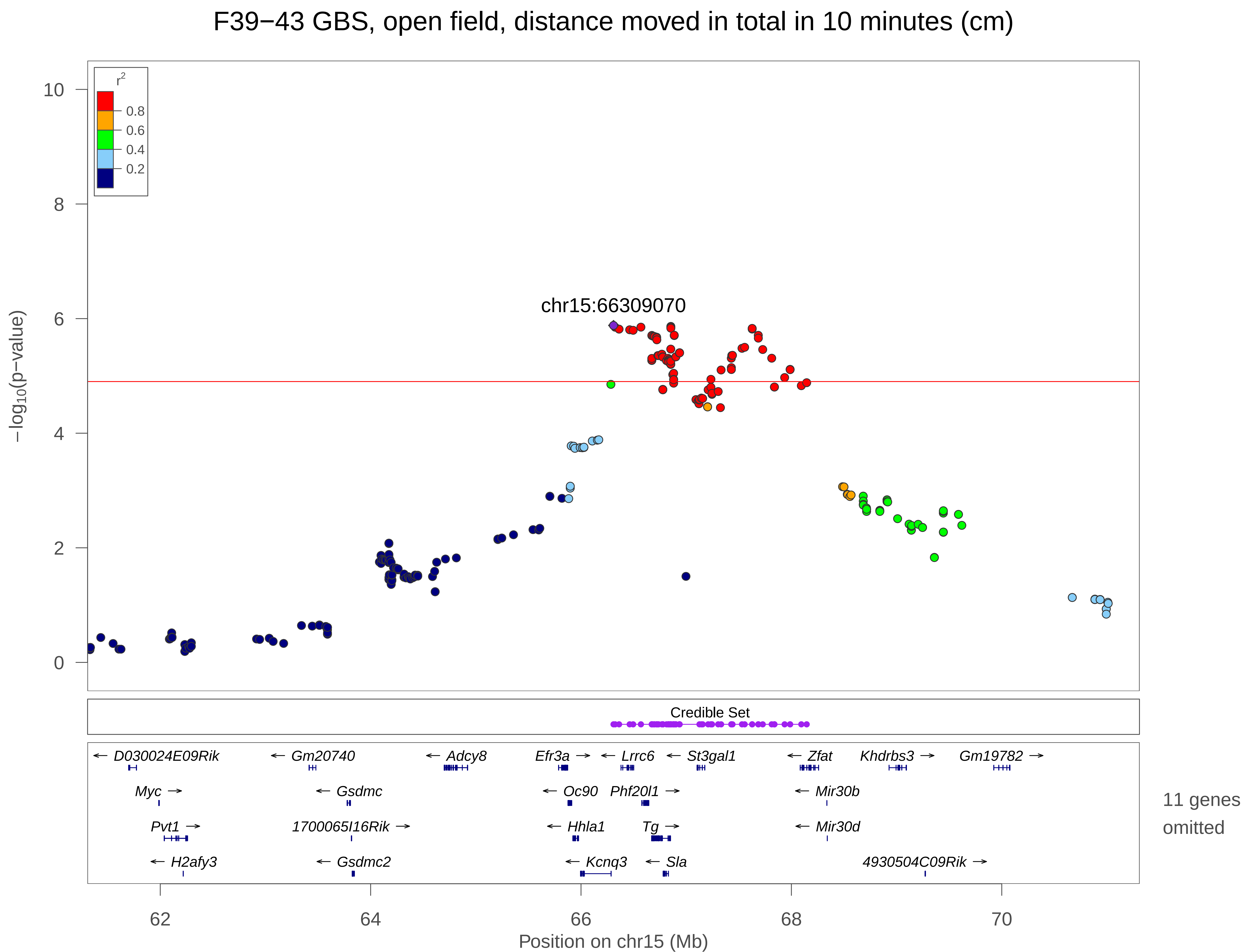

F39−43 GBS, open field, distance moved in total in 10 minutes (cm)
r2
0.8
0.6
0.4
0.2
10
8
6
4
2
0
−log10(p−value)
chr15:66309070
Credible Set
← D030024E09Rik
← Gm20740
← Adcy8
Efr3a →
← Lrrc6
← St3gal1
← Zfat
Khdrbs3 →
Gm19782 →
Myc →
← Gsdmc
← Oc90
Phf20l1 →
← Mir30b
Pvt1 →
1700065I16Rik →
← Hhla1
Tg →
← Mir30d
← H2afy3
← Gsdmc2
← Kcnq3
← Sla
4930504C09Rik →
11 genes omitted
62
64
66
68
70
Position on chr15 (Mb)

| | |
|---|---|
| date: | Thu Oct 25 17:23:28 2018 |
| build: | F3943GBS |
| display range: | chr15:61309070−71309070 [61309070−71309070] |
| hilite range: | 0 − 0 [ 0 − 0 ] |
| reference SNP: | chr15:66309070 |
| number of SNPs plotted: | 317 |
| min P−value: | 1.31E−6 [chr15:66309070] |
| max P−value: | 6.44E−1 [chr15:62233468] |
| omitted Genes: | Gsdmcl1, Gsdmc3, Gsdmcl2 |
| omitted Genes: | Gsdmc4, Gsdmcl−ps, Fam49b |
| omitted Genes: | Asap1, Tmem71, Wisp1 |
| omitted Genes: | Ndrg1, 1700012I11Rik |

Make more plots at http://csg.sph.umich.edu/locuszoom/

Fine−mapping Regions

| snp | chr | pos (Mb) | pp | group | color |
|---|---|---|---|---|---|
| chr15:66309070 | 15 | 66.30907 | 0.027602429 | Credible Set | purple |
| chr15:66324506 | 15 | 66.32451 | 0.025784362 | Credible Set | purple |
| chr15:66361813 | 15 | 66.36181 | 0.023925074 | Credible Set | purple |
| chr15:66462727 | 15 | 66.46273 | 0.023337518 | Credible Set | purple |
| chr15:66495529 | 15 | 66.49553 | 0.022875143 | Credible Set | purple |
| chr15:66569471 | 15 | 66.56947 | 0.025827651 | Credible Set | purple |
| chr15:66672984 | 15 | 66.67298 | 0.007175589 | Credible Set | purple |
| chr15:66672997 | 15 | 66.67300 | 0.007144867 | Credible Set | purple |
| chr15:66673010 | 15 | 66.67301 | 0.007238337 | Credible Set | purple |
| chr15:66673015 | 15 | 66.67302 | 0.007736400 | Credible Set | purple |
| chr15:66673436 | 15 | 66.67344 | 0.018768276 | Credible Set | purple |
| chr15:66673541 | 15 | 66.67354 | 0.018660582 | Credible Set | purple |
| chr15:66673940 | 15 | 66.67394 | 0.018616154 | Credible Set | purple |
| chr15:66685097 | 15 | 66.68510 | 0.018201349 | Credible Set | purple |
| chr15:66696518 | 15 | 66.69652 | 0.018000747 | Credible Set | purple |
| chr15:66699723 | 15 | 66.69972 | 0.017931702 | Credible Set | purple |
| chr15:66719126 | 15 | 66.71913 | 0.017530102 | Credible Set | purple |
| chr15:66719241 | 15 | 66.71924 | 0.017048919 | Credible Set | purple |
| chr15:66720772 | 15 | 66.72077 | 0.015837501 | Credible Set | purple |
| chr15:66730424 | 15 | 66.73042 | 0.008618150 | Credible Set | purple |
| chr15:66737376 | 15 | 66.73738 | 0.008641068 | Credible Set | purple |
| chr15:66768701 | 15 | 66.76870 | 0.009102203 | Credible Set | purple |
| chr15:66777400 | 15 | 66.77740 | 0.008198637 | Credible Set | purple |
| chr15:66777425 | 15 | 66.77742 | 0.008199840 | Credible Set | purple |
| chr15:66777461 | 15 | 66.77746 | 0.008138798 | Credible Set | purple |
| chr15:66777706 | 15 | 66.77771 | 0.002350850 | Credible Set | purple |
| chr15:66777728 | 15 | 66.77773 | 0.002315625 | Credible Set | purple |
| chr15:66777729 | 15 | 66.77773 | 0.002344350 | Credible Set | purple |
| chr15:66777730 | 15 | 66.77773 | 0.002356767 | Credible Set | purple |
| chr15:66777731 | 15 | 66.77773 | 0.002334958 | Credible Set | purple |
| chr15:66777743 | 15 | 66.77774 | 0.002333687 | Credible Set | purple |
| chr15:66812941 | 15 | 66.81294 | 0.007096851 | Credible Set | purple |
| chr15:66826934 | 15 | 66.82693 | 0.007768279 | Credible Set | purple |
| chr15:66834506 | 15 | 66.83451 | 0.007041661 | Credible Set | purple |
| chr15:66834532 | 15 | 66.83453 | 0.007221292 | Credible Set | purple |
| chr15:66834574 | 15 | 66.83457 | 0.007203856 | Credible Set | purple |

Fine−mapping Regions

| snp | chr | pos (Mb) | pp | group | color |
|---|---|---|---|---|---|
| chr15:66834575 | 15 | 66.83458 | 0.007380243 | Credible Set | purple |
| chr15:66834600 | 15 | 66.83460 | 0.006909807 | Credible Set | purple |
| chr15:66834621 | 15 | 66.83462 | 0.007256518 | Credible Set | purple |
| chr15:66834624 | 15 | 66.83462 | 0.006944240 | Credible Set | purple |
| chr15:66834637 | 15 | 66.83464 | 0.006724454 | Credible Set | purple |
| chr15:66853166 | 15 | 66.85317 | 0.011076730 | Credible Set | purple |
| chr15:66853439 | 15 | 66.85344 | 0.006141571 | Credible Set | purple |
| chr15:66853460 | 15 | 66.85346 | 0.006206811 | Credible Set | purple |
| chr15:66853489 | 15 | 66.85349 | 0.006216656 | Credible Set | purple |
| chr15:66853520 | 15 | 66.85352 | 0.006954034 | Credible Set | purple |
| chr15:66853893 | 15 | 66.85389 | 0.026569445 | Credible Set | purple |
| chr15:66853894 | 15 | 66.85389 | 0.025751834 | Credible Set | purple |
| chr15:66853923 | 15 | 66.85392 | 0.025749399 | Credible Set | purple |
| chr15:66853954 | 15 | 66.85395 | 0.024823326 | Credible Set | purple |
| chr15:66872519 | 15 | 66.87252 | 0.004148601 | Credible Set | purple |
| chr15:66880575 | 15 | 66.88057 | 0.002987713 | Credible Set | purple |
| chr15:66880583 | 15 | 66.88058 | 0.004361522 | Credible Set | purple |
| chr15:66880635 | 15 | 66.88063 | 0.003460220 | Credible Set | purple |
| chr15:66880638 | 15 | 66.88064 | 0.003412050 | Credible Set | purple |
| chr15:66886219 | 15 | 66.88622 | 0.018666227 | Credible Set | purple |
| chr15:66886266 | 15 | 66.88627 | 0.018934431 | Credible Set | purple |
| chr15:66901528 | 15 | 66.90153 | 0.008224937 | Credible Set | purple |
| chr15:66901548 | 15 | 66.90155 | 0.008224937 | Credible Set | purple |
| chr15:66901593 | 15 | 66.90159 | 0.008224937 | Credible Set | purple |
| chr15:66935376 | 15 | 66.93538 | 0.009558409 | Credible Set | purple |
| chr15:66935390 | 15 | 66.93539 | 0.009558409 | Credible Set | purple |
| chr15:66935391 | 15 | 66.93539 | 0.009558409 | Credible Set | purple |
| chr15:66935405 | 15 | 66.93541 | 0.009558409 | Credible Set | purple |
| chr15:66937231 | 15 | 66.93723 | 0.009632455 | Credible Set | purple |
| chr15:66937423 | 15 | 66.93742 | 0.009580685 | Credible Set | purple |
| chr15:67126159 | 15 | 67.12616 | 0.001608394 | Credible Set | purple |
| chr15:67144054 | 15 | 67.14405 | 0.001689761 | Credible Set | purple |
| chr15:67156480 | 15 | 67.15648 | 0.001663602 | Credible Set | purple |
| chr15:67208761 | 15 | 67.20876 | 0.002315708 | Credible Set | purple |
| chr15:67234910 | 15 | 67.23491 | 0.002553108 | Credible Set | purple |
| chr15:67235072 | 15 | 67.23507 | 0.003465870 | Credible Set | purple |

Fine−mapping Regions

| snp | chr | pos (Mb) | pp | group | color |
|---|---|---|---|---|---|
| chr15:67245093 | 15 | 67.24509 | 0.001948659 | Credible Set | purple |
| chr15:67245123 | 15 | 67.24512 | 0.002023520 | Credible Set | purple |
| chr15:67304194 | 15 | 67.30419 | 0.002169408 | Credible Set | purple |
| chr15:67331738 | 15 | 67.33174 | 0.004954677 | Credible Set | purple |
| chr15:67428857 | 15 | 67.42886 | 0.007784087 | Credible Set | purple |
| chr15:67429407 | 15 | 67.42941 | 0.005426222 | Credible Set | purple |
| chr15:67429420 | 15 | 67.42942 | 0.005044111 | Credible Set | purple |
| chr15:67433395 | 15 | 67.43340 | 0.008657028 | Credible Set | purple |
| chr15:67439974 | 15 | 67.43997 | 0.008716280 | Credible Set | purple |
| chr15:67530570 | 15 | 67.53057 | 0.011413170 | Credible Set | purple |
| chr15:67555683 | 15 | 67.55568 | 0.011864812 | Credible Set | purple |
| chr15:67627161 | 15 | 67.62716 | 0.023978263 | Credible Set | purple |
| chr15:67627183 | 15 | 67.62718 | 0.024568459 | Credible Set | purple |
| chr15:67627194 | 15 | 67.62719 | 0.024589519 | Credible Set | purple |
| chr15:67685224 | 15 | 67.68522 | 0.018843202 | Credible Set | purple |
| chr15:67685781 | 15 | 67.68578 | 0.016888870 | Credible Set | purple |
| chr15:67727448 | 15 | 67.72745 | 0.010895379 | Credible Set | purple |
| chr15:67812971 | 15 | 67.81297 | 0.007799587 | Credible Set | purple |
| chr15:67838708 | 15 | 67.83871 | 0.002574356 | Credible Set | purple |
| chr15:67838720 | 15 | 67.83872 | 0.002572597 | Credible Set | purple |
| chr15:67838721 | 15 | 67.83872 | 0.002581671 | Credible Set | purple |
| chr15:67935812 | 15 | 67.93581 | 0.003710205 | Credible Set | purple |
| chr15:67987913 | 15 | 67.98791 | 0.005080623 | Credible Set | purple |
| chr15:67987932 | 15 | 67.98793 | 0.005020532 | Credible Set | purple |
| chr15:68094377 | 15 | 68.09438 | 0.002712696 | Credible Set | purple |
| chr15:68145325 | 15 | 68.14532 | 0.003041452 | Credible Set | purple |

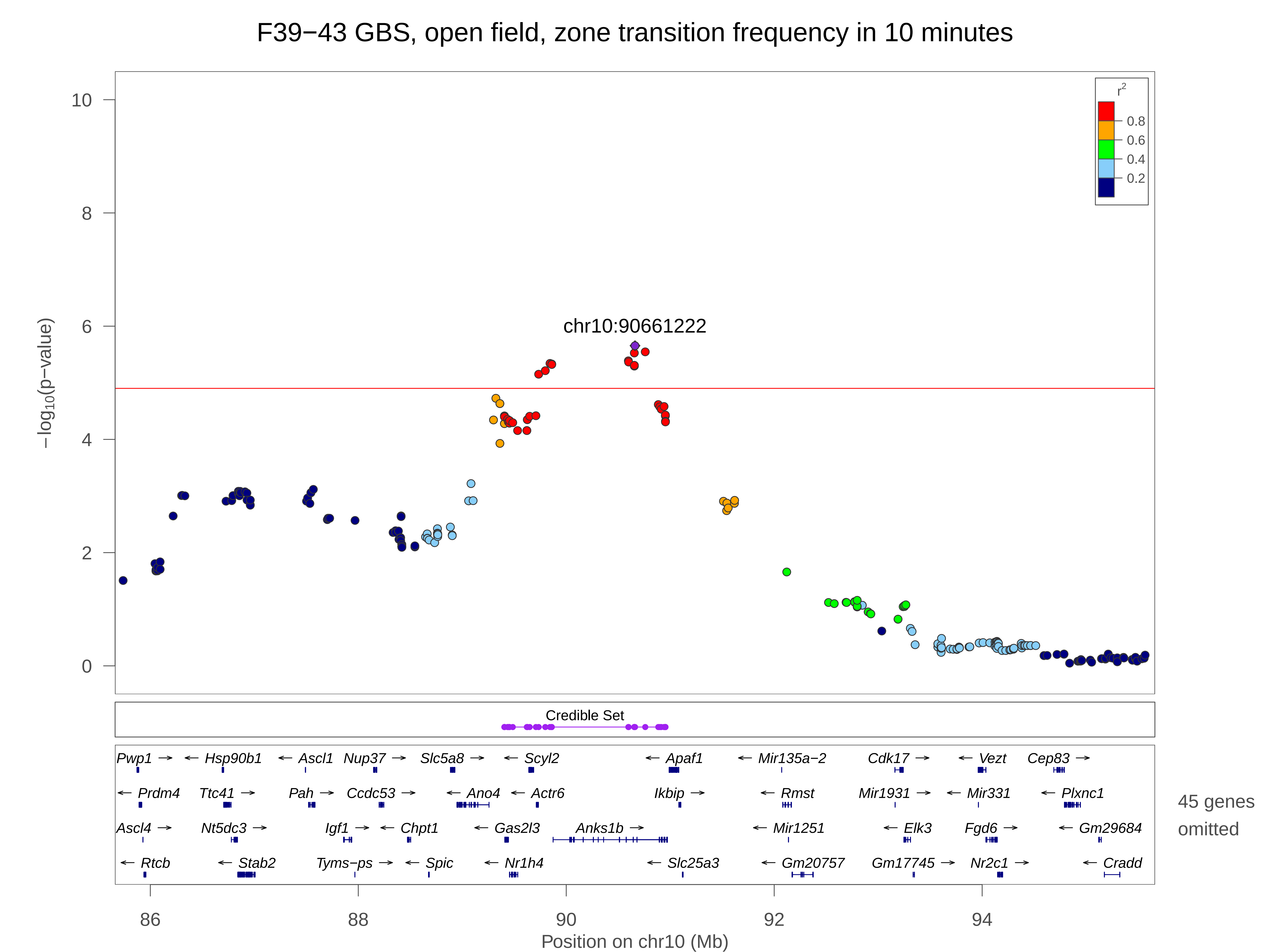
F39−43 GBS, open field, zone transition frequency in 10 minutes
chr10:90661222
r2
0.8
0.6
0.4
0.2
−log10(p−value)
Credible Set
Pwp1
Hsp90b1
Ascl1
Nup37
Slc5a8
Scyl2
Apaf1
Mir135a−2
Cdk17
Vezt
Cep83
Prdm4
Ttc41
Pah
Ccdc53
Ano4
Actr6
Ikbip
Rmst
Mir1931
Mir331
Plxnc1
Ascl4
Nt5dc3
Igf1
Chpt1
Gas2l3
Anks1b
Mir1251
Elk3
Fgd6
Gm29684
Rtcb
Stab2
Tyms−ps
Spic
Nr1h4
Slc25a3
Gm20757
Gm17745
Nr2c1
Cradd
45 genes omitted
Position on chr10 (Mb)

| | |
|---|---|
| date: | Thu Oct 25 17:23:37 2018 |
| build: | F3943GBS |
| display range: | chr10:85661222−95661222 [85661222−95661222] |
| hilite range: | 0 − 0 [ 0 − 0 ] |
| reference SNP: | chr10:90661222 |
| number of SNPs plotted: | 288 |
| min P−value: | 2.21E−6 [chr10:90661222] |
| max P−value: | 8.96E−1 [chr10:94840769] |
| omitted Genes: | Bpifc, Fbxo7, Syn3 |
| omitted Genes: | 4930486F22Rik, Timp3, 1810014B01Rik |
| omitted Genes: | 1700113H08Rik, Pmch, Parpbp |
| omitted Genes: | Dram1, Gnptab, Sycp3 |
| omitted Genes: | Mybpc1, Gm4925, Arl1 |
| omitted Genes: | Utp20, Slc17a8, 1500026H17Rik |
| omitted Genes: | Uhrf1bp1l, Tmpo, Nedd1 |
| omitted Genes: | Mir7688, Lta4h, Hal |
| omitted Genes: | Amdhd1, Ccdc38, Snrpf |
| omitted Genes: | 4933408J17Rik, Ntn4, Gm8580 |
| omitted Genes: | Gm15915, Usp44, Metap2 |
| omitted Genes: | Ndufa12, Gm4792, Tmcc3 |
| omitted Genes: | Mir7211, Cep83os, 2310039L15Rik |
| omitted Genes: | Socs2, 5730420D15Rik, Mrpl42 |
| omitted Genes: | Ube2n, Nudt4, Mir3058 |

Make more plots at http://csg.sph.umich.edu/locuszoom/

Fine−mapping Regions

| snp | chr | pos (Mb) | pp | group | color |
|---|---|---|---|---|---|
| chr10:89405556 | 10 | 89.40556 | 0.007110253 | Credible Set | purple |
| chr10:89405564 | 10 | 89.40556 | 0.007110253 | Credible Set | purple |
| chr10:89405589 | 10 | 89.40559 | 0.006873806 | Credible Set | purple |
| chr10:89405627 | 10 | 89.40563 | 0.006872709 | Credible Set | purple |
| chr10:89435280 | 10 | 89.43528 | 0.006237086 | Credible Set | purple |
| chr10:89443390 | 10 | 89.44339 | 0.005643714 | Credible Set | purple |
| chr10:89454154 | 10 | 89.45415 | 0.005374599 | Credible Set | purple |
| chr10:89454848 | 10 | 89.45485 | 0.005948185 | Credible Set | purple |
| chr10:89485534 | 10 | 89.48553 | 0.005483304 | Credible Set | purple |
| chr10:89621284 | 10 | 89.62128 | 0.004055150 | Credible Set | purple |
| chr10:89625488 | 10 | 89.62549 | 0.006149478 | Credible Set | purple |
| chr10:89625559 | 10 | 89.62556 | 0.006163075 | Credible Set | purple |
| chr10:89647994 | 10 | 89.64799 | 0.007025377 | Credible Set | purple |
| chr10:89708192 | 10 | 89.70819 | 0.007167567 | Credible Set | purple |
| chr10:89734590 | 10 | 89.73459 | 0.035816412 | Credible Set | purple |
| chr10:89798643 | 10 | 89.79864 | 0.041113810 | Credible Set | purple |
| chr10:89843657 | 10 | 89.84366 | 0.054575227 | Credible Set | purple |
| chr10:89860498 | 10 | 89.86050 | 0.053350253 | Credible Set | purple |
| chr10:89860846 | 10 | 89.86085 | 0.052304898 | Credible Set | purple |
| chr10:90597510 | 10 | 90.59751 | 0.060343620 | Credible Set | purple |
| chr10:90599690 | 10 | 90.59969 | 0.057550014 | Credible Set | purple |
| chr10:90654718 | 10 | 90.65472 | 0.081877350 | Credible Set | purple |
| chr10:90654994 | 10 | 90.65499 | 0.049181449 | Credible Set | purple |
| chr10:90654998 | 10 | 90.65500 | 0.050621256 | Credible Set | purple |
| chr10:90655002 | 10 | 90.65500 | 0.049224914 | Credible Set | purple |
| chr10:90655039 | 10 | 90.65504 | 0.050537434 | Credible Set | purple |
| chr10:90661222 | 10 | 90.66122 | 0.108813289 | Credible Set | purple |
| chr10:90759813 | 10 | 90.75981 | 0.085624559 | Credible Set | purple |
| chr10:90885383 | 10 | 90.88538 | 0.011035572 | Credible Set | purple |
| chr10:90899946 | 10 | 90.89995 | 0.010107830 | Credible Set | purple |
| chr10:90913583 | 10 | 90.91358 | 0.009231826 | Credible Set | purple |
| chr10:90940720 | 10 | 90.94072 | 0.010245749 | Credible Set | purple |
| chr10:90953385 | 10 | 90.95338 | 0.007127699 | Credible Set | purple |
| chr10:90953402 | 10 | 90.95340 | 0.007216611 | Credible Set | purple |
| chr10:90953425 | 10 | 90.95342 | 0.007360237 | Credible Set | purple |
| chr10:90954053 | 10 | 90.95405 | 0.005791753 | Credible Set | purple |

Fine−mapping Regions

| snp | chr | pos (Mb) | pp | group | color |
|---|---|---|---|---|---|
| chr10:90954116 | 10 | 90.95412 | 0.005647662 | Credible Set | purple |

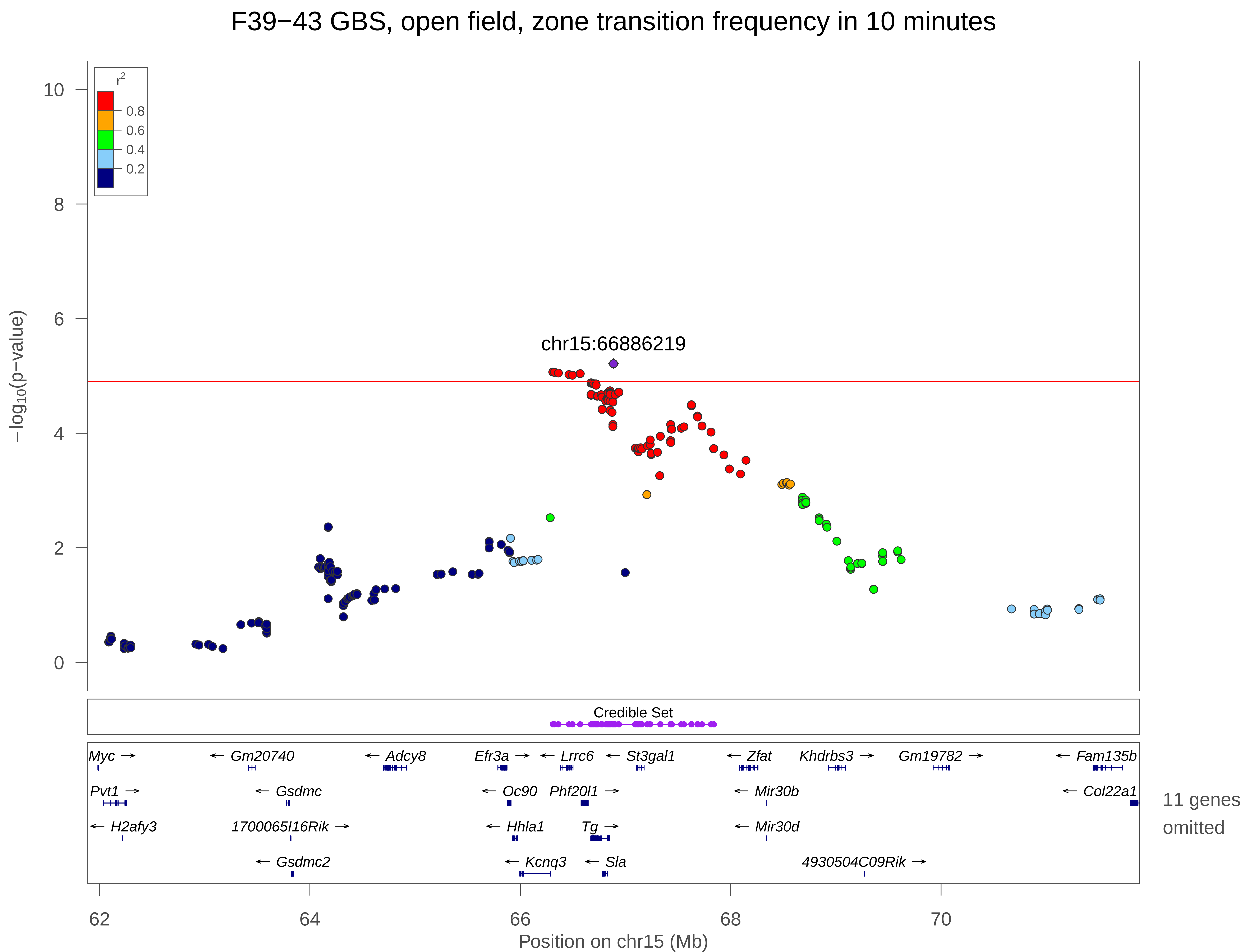
F39−43 GBS, open field, zone transition frequency in 10 minutes
r2
0.8
0.6
0.4
0.2
10
8
6
4
2
0
−log10(p−value)
chr15:66886219
Credible Set
Myc →
← Gm20740
← Adcy8
Efr3a →
← Lrrc6
← St3gal1
← Zfat
Khdrbs3 →
Gm19782 →
← Fam135b
Pvt1 →
← Gsdmc
← Oc90
Phf20l1 →
← Mir30b
← Col22a1
← H2afy3
1700065I16Rik →
← Hhla1
Tg →
← Mir30d
← Gsdmc2
← Kcnq3
← Sla
4930504C09Rik →
11 genes omitted
62
64
66
68
70
Position on chr15 (Mb)

| | |
|---|---|
| date: | Thu Oct 25 17:23:45 2018 |
| build: | F3943GBS |
| display range: | chr15:61886219−71886219 [61886219−71886219] |
| hilite range: | 0 − 0 [ 0 − 0 ] |
| reference SNP: | chr15:66886219 |
| number of SNPs plotted: | 313 |
| min P−value: | 6.11E−6 [chr15:66886219] |
| max P−value: | 5.75E−1 [chr15:63173066] |
| omitted Genes: | Gsdmcl1, Gsdmc3, Gsdmcl2 |
| omitted Genes: | Gsdmc4, Gsdmcl−ps, Fam49b |
| omitted Genes: | Asap1, Tmem71, Wisp1 |
| omitted Genes: | Ndrg1, 1700012I11Rik |

Make more plots at http://csg.sph.umich.edu/locuszoom/

Fine−mapping Regions

| snp | chr | pos (Mb) | pp | group | color |
|---|---|---|---|---|---|
| chr15:66309070 | 15 | 66.30907 | 0.028511659 | Credible Set | purple |
| chr15:66324506 | 15 | 66.32451 | 0.028181511 | Credible Set | purple |
| chr15:66361813 | 15 | 66.36181 | 0.027429878 | Credible Set | purple |
| chr15:66462727 | 15 | 66.46273 | 0.025826000 | Credible Set | purple |
| chr15:66495529 | 15 | 66.49553 | 0.025206244 | Credible Set | purple |
| chr15:66569471 | 15 | 66.56947 | 0.026786024 | Credible Set | purple |
| chr15:66672984 | 15 | 66.67298 | 0.012177363 | Credible Set | purple |
| chr15:66672997 | 15 | 66.67300 | 0.012049709 | Credible Set | purple |
| chr15:66673010 | 15 | 66.67301 | 0.011713852 | Credible Set | purple |
| chr15:66673015 | 15 | 66.67302 | 0.011966795 | Credible Set | purple |
| chr15:66673436 | 15 | 66.67344 | 0.018585760 | Credible Set | purple |
| chr15:66673541 | 15 | 66.67354 | 0.018767046 | Credible Set | purple |
| chr15:66673940 | 15 | 66.67394 | 0.018823625 | Credible Set | purple |
| chr15:66685097 | 15 | 66.68510 | 0.018420545 | Credible Set | purple |
| chr15:66696518 | 15 | 66.69652 | 0.018046317 | Credible Set | purple |
| chr15:66699723 | 15 | 66.69972 | 0.018160473 | Credible Set | purple |
| chr15:66719126 | 15 | 66.71913 | 0.018119210 | Credible Set | purple |
| chr15:66719241 | 15 | 66.71924 | 0.018033946 | Credible Set | purple |
| chr15:66720772 | 15 | 66.72077 | 0.017196761 | Credible Set | purple |
| chr15:66730424 | 15 | 66.73042 | 0.011300042 | Credible Set | purple |
| chr15:66737376 | 15 | 66.73738 | 0.011358314 | Credible Set | purple |
| chr15:66768701 | 15 | 66.76870 | 0.011949547 | Credible Set | purple |
| chr15:66777400 | 15 | 66.77740 | 0.010904753 | Credible Set | purple |
| chr15:66777425 | 15 | 66.77742 | 0.010900928 | Credible Set | purple |
| chr15:66777461 | 15 | 66.77746 | 0.010891382 | Credible Set | purple |
| chr15:66777706 | 15 | 66.77771 | 0.006879374 | Credible Set | purple |
| chr15:66777728 | 15 | 66.77773 | 0.006789628 | Credible Set | purple |
| chr15:66777729 | 15 | 66.77773 | 0.006877831 | Credible Set | purple |
| chr15:66777730 | 15 | 66.77773 | 0.006834784 | Credible Set | purple |
| chr15:66777731 | 15 | 66.77773 | 0.006834310 | Credible Set | purple |
| chr15:66777743 | 15 | 66.77774 | 0.006808240 | Credible Set | purple |
| chr15:66812941 | 15 | 66.81294 | 0.009413262 | Credible Set | purple |
| chr15:66826934 | 15 | 66.82693 | 0.010303910 | Credible Set | purple |
| chr15:66834506 | 15 | 66.83451 | 0.012647346 | Credible Set | purple |
| chr15:66834532 | 15 | 66.83453 | 0.012876972 | Credible Set | purple |
| chr15:66834574 | 15 | 66.83457 | 0.012504120 | Credible Set | purple |

Fine−mapping Regions

| snp | chr | pos (Mb) | pp | group | color |
|---|---|---|---|---|---|
| chr15:66834575 | 15 | 66.83458 | 0.012686151 | Credible Set | purple |
| chr15:66834600 | 15 | 66.83460 | 0.009679353 | Credible Set | purple |
| chr15:66834621 | 15 | 66.83462 | 0.010209681 | Credible Set | purple |
| chr15:66834624 | 15 | 66.83462 | 0.009817655 | Credible Set | purple |
| chr15:66834637 | 15 | 66.83464 | 0.009537756 | Credible Set | purple |
| chr15:66853166 | 15 | 66.85317 | 0.013866897 | Credible Set | purple |
| chr15:66853439 | 15 | 66.85344 | 0.006552096 | Credible Set | purple |
| chr15:66853460 | 15 | 66.85346 | 0.006624216 | Credible Set | purple |
| chr15:66853489 | 15 | 66.85349 | 0.006548992 | Credible Set | purple |
| chr15:66853520 | 15 | 66.85352 | 0.009333603 | Credible Set | purple |
| chr15:66853893 | 15 | 66.85389 | 0.012991772 | Credible Set | purple |
| chr15:66853894 | 15 | 66.85389 | 0.012766362 | Credible Set | purple |
| chr15:66853923 | 15 | 66.85392 | 0.012583236 | Credible Set | purple |
| chr15:66853954 | 15 | 66.85395 | 0.012142949 | Credible Set | purple |
| chr15:66872519 | 15 | 66.87252 | 0.006095154 | Credible Set | purple |
| chr15:66880575 | 15 | 66.88057 | 0.003844528 | Credible Set | purple |
| chr15:66880583 | 15 | 66.88058 | 0.009036405 | Credible Set | purple |
| chr15:66880635 | 15 | 66.88063 | 0.003543907 | Credible Set | purple |
| chr15:66880638 | 15 | 66.88064 | 0.003523287 | Credible Set | purple |
| chr15:66886219 | 15 | 66.88622 | 0.039337654 | Credible Set | purple |
| chr15:66886266 | 15 | 66.88627 | 0.038961971 | Credible Set | purple |
| chr15:66901528 | 15 | 66.90153 | 0.011970614 | Credible Set | purple |
| chr15:66901548 | 15 | 66.90155 | 0.011970614 | Credible Set | purple |
| chr15:66901593 | 15 | 66.90159 | 0.011970614 | Credible Set | purple |
| chr15:66935376 | 15 | 66.93538 | 0.013148876 | Credible Set | purple |
| chr15:66935390 | 15 | 66.93539 | 0.013148876 | Credible Set | purple |
| chr15:66935391 | 15 | 66.93539 | 0.013148876 | Credible Set | purple |
| chr15:66935405 | 15 | 66.93541 | 0.013148876 | Credible Set | purple |
| chr15:66937231 | 15 | 66.93723 | 0.013166389 | Credible Set | purple |
| chr15:66937423 | 15 | 66.93742 | 0.013168301 | Credible Set | purple |
| chr15:67092666 | 15 | 67.09267 | 0.001569620 | Credible Set | purple |
| chr15:67111279 | 15 | 67.11128 | 0.001467006 | Credible Set | purple |
| chr15:67121570 | 15 | 67.12157 | 0.001373203 | Credible Set | purple |
| chr15:67121756 | 15 | 67.12176 | 0.001554881 | Credible Set | purple |
| chr15:67126159 | 15 | 67.12616 | 0.001570026 | Credible Set | purple |
| chr15:67144054 | 15 | 67.14405 | 0.001585262 | Credible Set | purple |

Fine−mapping Regions

| snp | chr | pos (Mb) | pp | group | color |
|---|---|---|---|---|---|
| chr15:67156480 | 15 | 67.15648 | 0.001518148 | Credible Set | purple |
| chr15:67208761 | 15 | 67.20876 | 0.001703156 | Credible Set | purple |
| chr15:67234910 | 15 | 67.23491 | 0.001802725 | Credible Set | purple |
| chr15:67235072 | 15 | 67.23507 | 0.002136883 | Credible Set | purple |
| chr15:67331738 | 15 | 67.33174 | 0.002448117 | Credible Set | purple |
| chr15:67428857 | 15 | 67.42886 | 0.003831127 | Credible Set | purple |
| chr15:67429407 | 15 | 67.42941 | 0.002081463 | Credible Set | purple |
| chr15:67429420 | 15 | 67.42942 | 0.001935854 | Credible Set | purple |
| chr15:67433395 | 15 | 67.43340 | 0.003215775 | Credible Set | purple |
| chr15:67439974 | 15 | 67.43997 | 0.003212451 | Credible Set | purple |
| chr15:67530570 | 15 | 67.53057 | 0.003328201 | Credible Set | purple |
| chr15:67555683 | 15 | 67.55568 | 0.003506209 | Credible Set | purple |
| chr15:67627161 | 15 | 67.62716 | 0.007848421 | Credible Set | purple |
| chr15:67627183 | 15 | 67.62718 | 0.008139256 | Credible Set | purple |
| chr15:67627194 | 15 | 67.62719 | 0.008099851 | Credible Set | purple |
| chr15:67685224 | 15 | 67.68522 | 0.005299297 | Credible Set | purple |
| chr15:67685781 | 15 | 67.68578 | 0.005091978 | Credible Set | purple |
| chr15:67727448 | 15 | 67.72745 | 0.003622422 | Credible Set | purple |
| chr15:67812971 | 15 | 67.81297 | 0.002877637 | Credible Set | purple |
| chr15:67838708 | 15 | 67.83871 | 0.001529699 | Credible Set | purple |
| chr15:67838720 | 15 | 67.83872 | 0.001527159 | Credible Set | purple |
| chr15:67838721 | 15 | 67.83872 | 0.001538616 | Credible Set | purple |

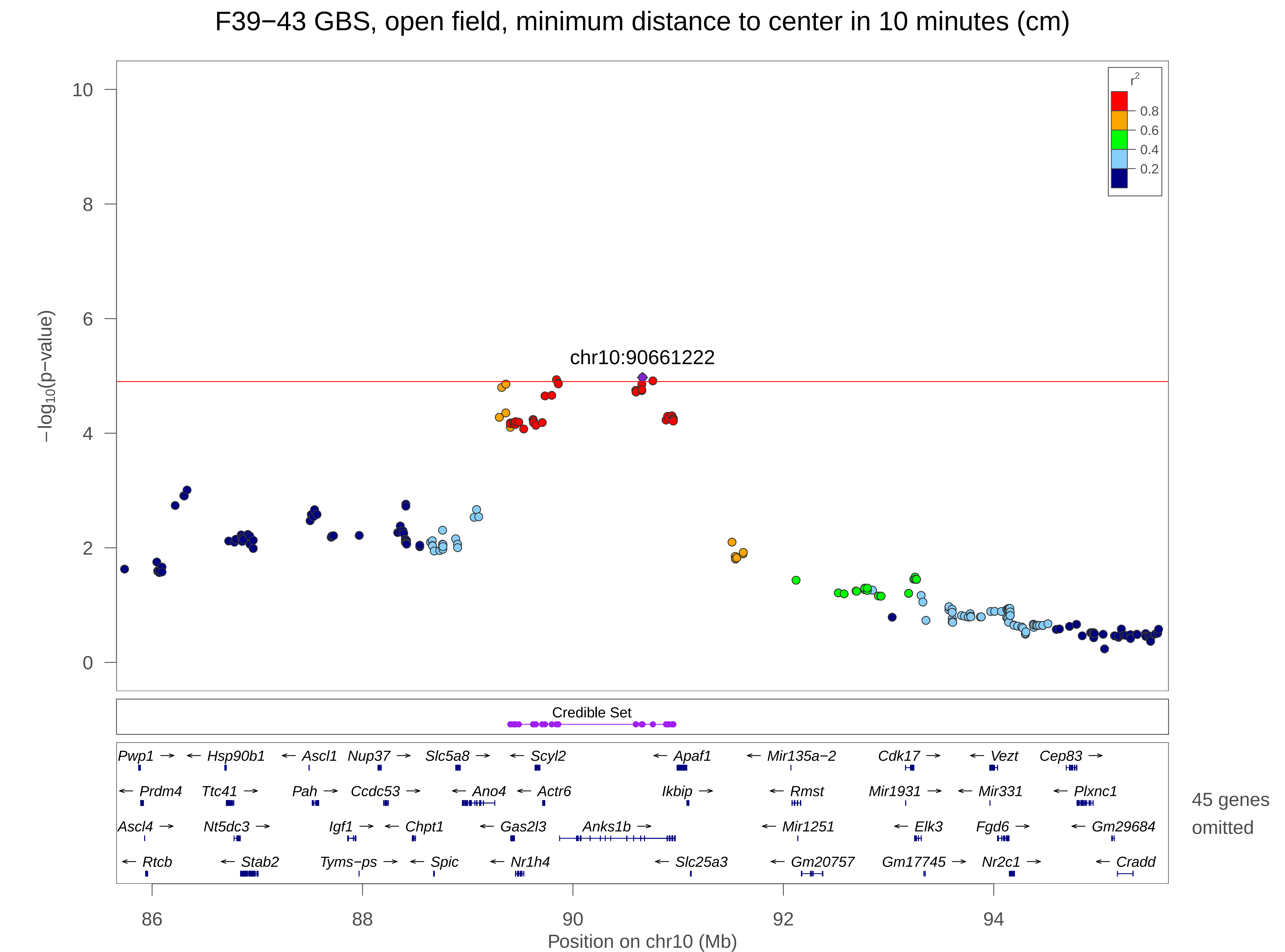

F39−43 GBS, open field, minimum distance to center in 10 minutes (cm)
−log10(p−value)
10
8
6
4
2
0
r2
0.8
0.6
0.4
0.2
chr10:90661222
Credible Set
Pwp1 →
← Hsp90b1
← Ascl1
Nup37 →
Slc5a8 →
← Scyl2
← Apaf1
← Mir135a−2
Cdk17 →
← Vezt
Cep83 →
← Prdm4
Ttc41 →
Pah →
Ccdc53 →
← Ano4
← Actr6
Ikbip →
← Rmst
Mir1931 →
← Mir331
← Plxnc1
Ascl4 →
Nt5dc3 →
Igf1 →
← Chpt1
← Gas2l3
Anks1b →
← Mir1251
← Elk3
Fgd6 →
← Gm29684
← Rtcb
← Stab2
Tyms−ps →
← Spic
← Nr1h4
← Slc25a3
← Gm20757
Gm17745 →
Nr2c1 →
← Cradd
45 genes
omitted
86
88
90
92
94
Position on chr10 (Mb)

| | |
|---|---|
| date: | Thu Oct 25 17:23:56 2018 |
| build: | F3943GBS |
| display range: | chr10:85661222−95661222 [85661222−95661222] |
| hilite range: | 0 − 0 [ 0 − 0 ] |
| reference SNP: | chr10:90661222 |
| number of SNPs plotted: | 288 |
| min P−value: | 1.06E−5 [chr10:90661222] |
| max P−value: | 5.81E−1 [chr10:95052984] |
| omitted Genes: | Bpifc, Fbxo7, Syn3 |
| omitted Genes: | 4930486F22Rik, Timp3, 1810014B01Rik |
| omitted Genes: | 1700113H08Rik, Pmch, Parpbp |
| omitted Genes: | Dram1, Gnptab, Sycp3 |
| omitted Genes: | Mybpc1, Gm4925, Arl1 |
| omitted Genes: | Utp20, Slc17a8, 1500026H17Rik |
| omitted Genes: | Uhrf1bp1l, Tmpo, Nedd1 |
| omitted Genes: | Mir7688, Lta4h, Hal |
| omitted Genes: | Amdhd1, Ccdc38, Snrpf |
| omitted Genes: | 4933408J17Rik, Ntn4, Gm8580 |
| omitted Genes: | Gm15915, Usp44, Metap2 |
| omitted Genes: | Ndufa12, Gm4792, Tmcc3 |
| omitted Genes: | Mir7211, Cep83os, 2310039L15Rik |
| omitted Genes: | Socs2, 5730420D15Rik, Mrpl42 |
| omitted Genes: | Ube2n, Nudt4, Mir3058 |

Make more plots at http://csg.sph.umich.edu/locuszoom/

Fine−mapping Regions

| snp | chr | pos (Mb) | pp | group | color |
|---|---|---|---|---|---|
| chr10:89405556 | 10 | 89.40556 | 0.01211623 | Credible Set | purple |
| chr10:89405564 | 10 | 89.40556 | 0.01211623 | Credible Set | purple |
| chr10:89405589 | 10 | 89.40559 | 0.01182010 | Credible Set | purple |
| chr10:89405627 | 10 | 89.40563 | 0.01180355 | Credible Set | purple |
| chr10:89435280 | 10 | 89.43528 | 0.01232855 | Credible Set | purple |
| chr10:89443390 | 10 | 89.44339 | 0.01156987 | Credible Set | purple |
| chr10:89454154 | 10 | 89.45415 | 0.01146621 | Credible Set | purple |
| chr10:89454848 | 10 | 89.45485 | 0.01277391 | Credible Set | purple |
| chr10:89485534 | 10 | 89.48553 | 0.01250510 | Credible Set | purple |
| chr10:89621284 | 10 | 89.62128 | 0.01382924 | Credible Set | purple |
| chr10:89621319 | 10 | 89.62132 | 0.01381266 | Credible Set | purple |
| chr10:89625488 | 10 | 89.62549 | 0.01323555 | Credible Set | purple |
| chr10:89625559 | 10 | 89.62556 | 0.01228437 | Credible Set | purple |
| chr10:89647994 | 10 | 89.64799 | 0.01109332 | Credible Set | purple |
| chr10:89708192 | 10 | 89.70819 | 0.01231649 | Credible Set | purple |
| chr10:89734590 | 10 | 89.73459 | 0.03405194 | Credible Set | purple |
| chr10:89798643 | 10 | 89.79864 | 0.03481205 | Credible Set | purple |
| chr10:89843657 | 10 | 89.84366 | 0.06339964 | Credible Set | purple |
| chr10:89860498 | 10 | 89.86050 | 0.05525050 | Credible Set | purple |
| chr10:89860846 | 10 | 89.86085 | 0.05396630 | Credible Set | purple |
| chr10:90597510 | 10 | 90.59751 | 0.04215795 | Credible Set | purple |
| chr10:90599690 | 10 | 90.59969 | 0.03941872 | Credible Set | purple |
| chr10:90654718 | 10 | 90.65472 | 0.05308422 | Credible Set | purple |
| chr10:90654994 | 10 | 90.65499 | 0.04179892 | Credible Set | purple |
| chr10:90654998 | 10 | 90.65500 | 0.04282740 | Credible Set | purple |
| chr10:90655002 | 10 | 90.65500 | 0.04181644 | Credible Set | purple |
| chr10:90655039 | 10 | 90.65504 | 0.04316454 | Credible Set | purple |
| chr10:90661222 | 10 | 90.66122 | 0.06913804 | Credible Set | purple |
| chr10:90759813 | 10 | 90.75981 | 0.06057082 | Credible Set | purple |
| chr10:90885383 | 10 | 90.88538 | 0.01353082 | Credible Set | purple |
| chr10:90899946 | 10 | 90.89995 | 0.01566368 | Credible Set | purple |
| chr10:90913583 | 10 | 90.91358 | 0.01451603 | Credible Set | purple |
| chr10:90940720 | 10 | 90.94072 | 0.01591151 | Credible Set | purple |
| chr10:90953385 | 10 | 90.95338 | 0.01444023 | Credible Set | purple |
| chr10:90953402 | 10 | 90.95340 | 0.01448270 | Credible Set | purple |
| chr10:90953425 | 10 | 90.95342 | 0.01446890 | Credible Set | purple |

Fine−mapping Regions

| snp | chr | pos (Mb) | pp | group | color |
|---|---|---|---|---|---|
| chr10:90954053 | 10 | 90.95405 | 0.01376738 | Credible Set | purple |
| chr10:90954116 | 10 | 90.95412 | 0.01304015 | Credible Set | purple |

F39−43 GBS, open field, immobile duration in 10 minutes (s)

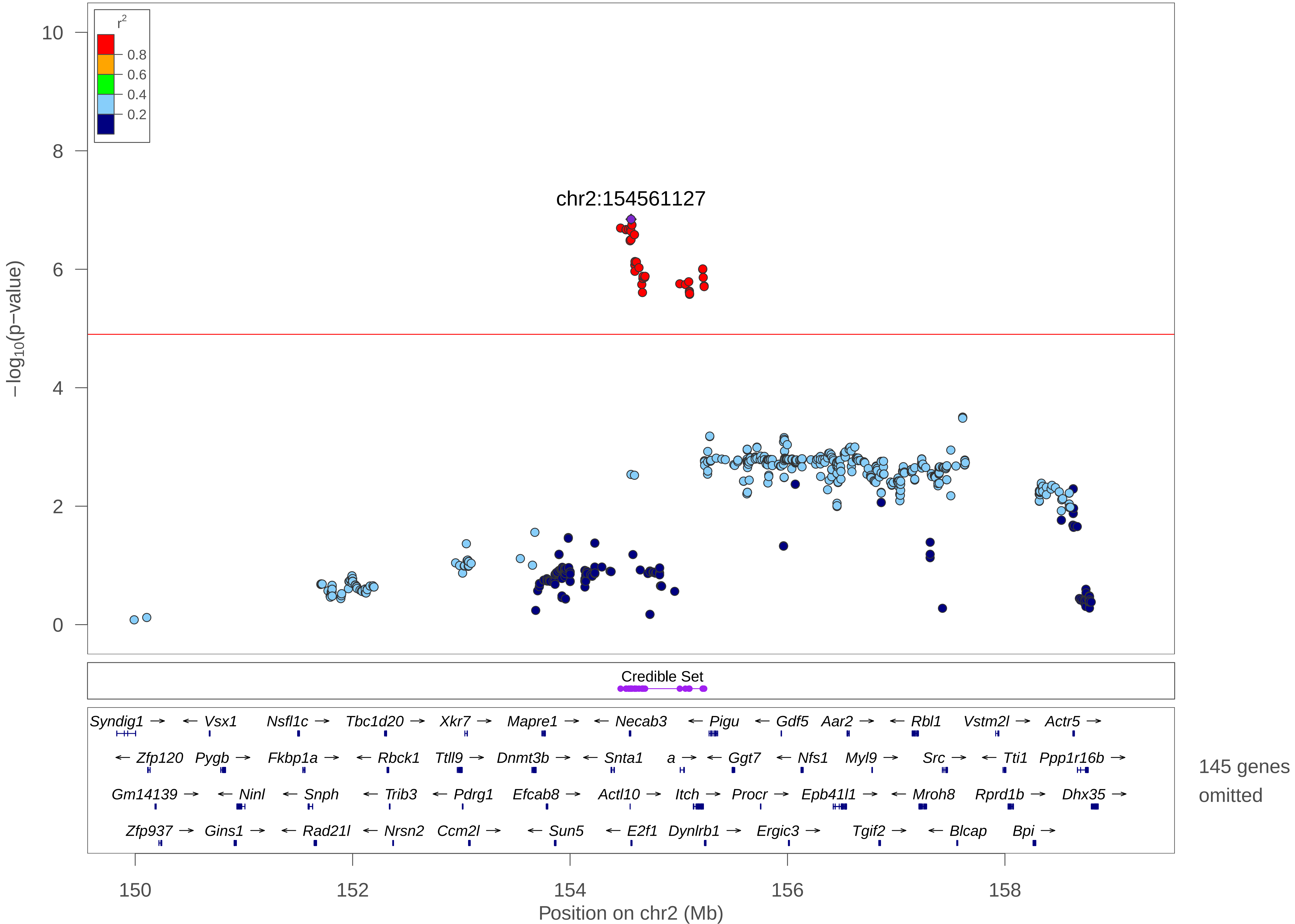

date: Thu Oct 25 17:23:01 2018
build: F3943GBS
display range: chr2:149561127−159561127 [149561127−159561127]
hilite range: 0 − 0 [ 0 − 0 ]
reference SNP: chr2:154561127
number of SNPs plotted: 757
min P−value: 1.43E−7 [chr2:154561127]
max P−value: 8.28E−1 [chr2:149990644]
omitted Genes: Gm14124, 3300002I08Rik, Zfp442
omitted Genes: Zfp345, Cst7, Apmap
omitted Genes: Acss1, E130215H24Rik, Entpd6
omitted Genes: Abhd12, Nanp, Gm14151
omitted Genes: 4930442J19Rik, LOC100125594, 4921509C19Rik
omitted Genes: Sdcbp2, Tmem74b, Psmf1
omitted Genes: Rspo4, Angpt4, Mir1953
omitted Genes: Fam110a, Slc52a3, Scrt2
omitted Genes: Srxn1, Tcf15, Csnk2a1
omitted Genes: Gm14164, Sox12, Zcchc3
omitted Genes: 6820408C15Rik, Defb23, Defb20
omitted Genes: Defb22, Defb26, Defb28
omitted Genes: Defb29, Defb21, Defb19
omitted Genes: Defb45, Defb36, Defb25
omitted Genes: Rem1, H13, Mcts2
omitted Genes: Id1, Cox4i2, Bcl2l1
omitted Genes: Tpx2, Mir5622, Mylk2
omitted Genes: Foxs1, Dusp15, Hck
omitted Genes: Tm9sf4, Tspyl3, Plagl2
omitted Genes: Pofut1, Kif3b, 2500004C02Rik
omitted Genes: Asxl1, Nol4l, Commd7
omitted Genes: Bpifb2, Bpifb6, Bpifb3
omitted Genes: Bpifb4, Bpifa6, Bpifa2
omitted Genes: Bpifa3, Bpifa1, Bpifa5
omitted Genes: Bpifb1, Bpifb5, Bpifb9a
omitted Genes: Bpifb9b, Cdk5rap1, Cbfa2t2
omitted Genes: 1700003F12Rik, Pxmp4, Zfp341
omitted Genes: Chmp4b, Raly, Eif2s2
omitted Genes: Ahcy, 2310005A03Rik, Map1lc3a
omitted Genes: Mir695, Trp53inp2, Ncoa6
omitted Genes: Acss2, Acss2os, Gss
Make more plots at http://csg.sph.umich.edu/locuszoom/
omitted Genes: Myh7b, Mir499, Trpc4ap

Fine−mapping Regions

| snp | chr | pos (Mb) | pp | group | color |
|---|---|---|---|---|---|
| chr2:154464466 | 2 | 154.4645 | 0.051164763 | Credible Set | purple |
| chr2:154513996 | 2 | 154.5140 | 0.048023099 | Credible Set | purple |
| chr2:154515752 | 2 | 154.5158 | 0.048448123 | Credible Set | purple |
| chr2:154516249 | 2 | 154.5162 | 0.048263220 | Credible Set | purple |
| chr2:154537995 | 2 | 154.5380 | 0.048967248 | Credible Set | purple |
| chr2:154538009 | 2 | 154.5380 | 0.048896921 | Credible Set | purple |
| chr2:154552607 | 2 | 154.5526 | 0.032573216 | Credible Set | purple |
| chr2:154552612 | 2 | 154.5526 | 0.031728222 | Credible Set | purple |
| chr2:154556468 | 2 | 154.5565 | 0.049374395 | Credible Set | purple |
| chr2:154560354 | 2 | 154.5604 | 0.033265595 | Credible Set | purple |
| chr2:154560360 | 2 | 154.5604 | 0.032895776 | Credible Set | purple |
| chr2:154560680 | 2 | 154.5607 | 0.045519233 | Credible Set | purple |
| chr2:154561127 | 2 | 154.5611 | 0.071188223 | Credible Set | purple |
| chr2:154569018 | 2 | 154.5690 | 0.057540698 | Credible Set | purple |
| chr2:154569036 | 2 | 154.5690 | 0.057432877 | Credible Set | purple |
| chr2:154592528 | 2 | 154.5925 | 0.039935441 | Credible Set | purple |
| chr2:154595741 | 2 | 154.5957 | 0.012837799 | Credible Set | purple |
| chr2:154595774 | 2 | 154.5958 | 0.012735284 | Credible Set | purple |
| chr2:154596294 | 2 | 154.5963 | 0.010112418 | Credible Set | purple |
| chr2:154597124 | 2 | 154.5971 | 0.014584592 | Credible Set | purple |
| chr2:154597483 | 2 | 154.5975 | 0.014398781 | Credible Set | purple |
| chr2:154610318 | 2 | 154.6103 | 0.014319167 | Credible Set | purple |
| chr2:154633945 | 2 | 154.6339 | 0.011567385 | Credible Set | purple |
| chr2:154659631 | 2 | 154.6596 | 0.006176648 | Credible Set | purple |
| chr2:154665835 | 2 | 154.6658 | 0.004574325 | Credible Set | purple |
| chr2:154670525 | 2 | 154.6705 | 0.007725612 | Credible Set | purple |
| chr2:154670528 | 2 | 154.6705 | 0.007729226 | Credible Set | purple |
| chr2:154670562 | 2 | 154.6706 | 0.007791349 | Credible Set | purple |
| chr2:154670571 | 2 | 154.6706 | 0.008385732 | Credible Set | purple |
| chr2:154686815 | 2 | 154.6868 | 0.008042347 | Credible Set | purple |
| chr2:154689824 | 2 | 154.6898 | 0.008385389 | Credible Set | purple |
| chr2:155009299 | 2 | 155.0093 | 0.006336677 | Credible Set | purple |
| chr2:155060609 | 2 | 155.0606 | 0.006174878 | Credible Set | purple |
| chr2:155091628 | 2 | 155.0916 | 0.006841165 | Credible Set | purple |
| chr2:155097606 | 2 | 155.0976 | 0.004820513 | Credible Set | purple |
| chr2:155097628 | 2 | 155.0976 | 0.004623856 | Credible Set | purple |

Fine−mapping Regions

| snp | chr | pos (Mb) | pp | group | color |
|---|---|---|---|---|---|
| chr2:155097650 | 2 | 155.0976 | 0.004634884 | Credible Set | purple |
| chr2:155220007 | 2 | 155.2200 | 0.010901354 | Credible Set | purple |
| chr2:155220040 | 2 | 155.2200 | 0.010901354 | Credible Set | purple |
| chr2:155220051 | 2 | 155.2201 | 0.010901354 | Credible Set | purple |
| chr2:155220725 | 2 | 155.2207 | 0.011057406 | Credible Set | purple |
| chr2:155224800 | 2 | 155.2248 | 0.008009014 | Credible Set | purple |
| chr2:155233215 | 2 | 155.2332 | 0.005844886 | Credible Set | purple |
| chr2:155233265 | 2 | 155.2333 | 0.005699991 | Credible Set | purple |

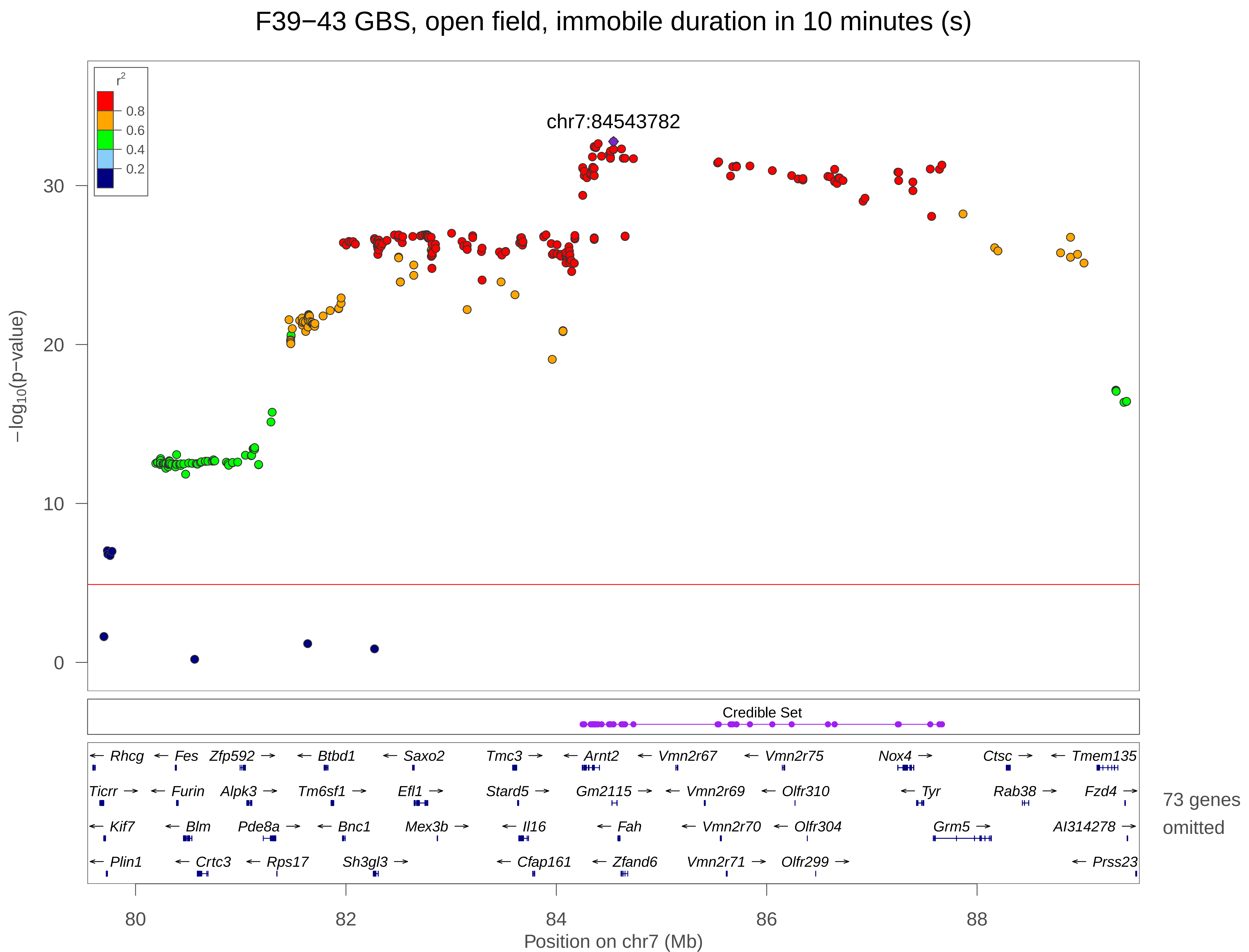
F39−43 GBS, open field, immobile duration in 10 minutes (s)
r2
0.8
0.6
0.4
0.2
chr7:84543782
−log10(p−value)
30
20
10
0
Credible Set
← Rhcg
← Fes
Zfp592 →
← Btbd1
← Saxo2
Tmc3 →
← Arnt2
← Vmn2r67
← Vmn2r75
Nox4 →
Ctsc →
← Tmem135
Ticrr →
← Furin
Alpk3 →
Tm6sf1 →
Efl1 →
Stard5 →
Gm2115 →
← Vmn2r69
← Olfr310
← Tyr
Rab38 →
Fzd4 →
← Kif7
← Blm
Pde8a →
← Bnc1
Mex3b →
← Il16
← Fah
← Vmn2r70
← Olfr304
Grm5 →
AI314278 →
← Plin1
← Crtc3
← Rps17
Sh3gl3 →
← Cfap161
← Zfand6
Vmn2r71 →
Olfr299 →
← Prss23
73 genes omitted
80
82
84
86
88
Position on chr7 (Mb)

| | |
|---|---|
| date: | Thu Oct 25 17:23:11 2018 |
| build: | F3943GBS |
| display range: | chr7:79543782−89543782 [79543782−89543782] |
| hilite range: | 0 − 0 [ 0 − 0 ] |
| reference SNP: | chr7:84543782 |
| number of SNPs plotted: | 390 |
| min P−value: | 1.69E−33 [chr7:84543782] |
| max P−value: | 6.35E−1 [chr7:80561830] |
| omitted Genes: | Pex11a, Wdr93, Mesp1 |
| omitted Genes: | Mesp2, Anpep, Ap3s2 |
| omitted Genes: | Arpin, Zfp710, Gm21057 |
| omitted Genes: | Idh2, Sema4b, Cib1 |
| omitted Genes: | Gdpgp1, Ttll13, Ngrn |
| omitted Genes: | Vps33b, Prc1, Rccd1 |
| omitted Genes: | Unc45a, Hddc3, Man2a2 |
| omitted Genes: | Gm15880, Iqgap1, Zscan2 |
| omitted Genes: | Wdr73, Nmb, Sec11a |
| omitted Genes: | Slc28a1, Cpeb1, Ap3b2 |
| omitted Genes: | BC048679, 2900076A07Rik, Mir1839 |
| omitted Genes: | Fsd2, Whamm, Homer2 |
| omitted Genes: | Fam103a1, 3110040N11Rik, Hdgfrp3 |
| omitted Genes: | Gm20744, Adamtsl3, 4933406J10Rik |
| omitted Genes: | 4933430H16Rik, Mesdc1, Mesdc2 |
| omitted Genes: | Cemip, Abhd17c, 2610206C17Rik |
| omitted Genes: | Olfr291, Olfr290, Vmn2r65 |
| omitted Genes: | Vmn2r66, Vmn2r68, Vmn2r72 |
| omitted Genes: | Vmn2r73, Vmn2r74, Vmn2r76 |
| omitted Genes: | Olfr309, Olfr308, Olfr307 |
| omitted Genes: | Olfr305, Olfr303, Olfr301 |
| omitted Genes: | Olfr298, Olfr297, Olfr295 |
| omitted Genes: | Olfr294, Olfr293, Olfr292 |
| omitted Genes: | Folh1, Vmn2r77, Vmn2r78 |
| omitted Genes: | Vmn2r79 |

Make more plots at http://csg.sph.umich.edu/locuszoom/

Fine−mapping Regions

| snp | chr | pos (Mb) | pp | group | color |
|---|---|---|---|---|---|
| chr7:84250985 | 7 | 84.25099 | 0.0025162275 | Credible Set | purple |
| chr7:84251011 | 7 | 84.25101 | 0.0028294597 | Credible Set | purple |
| chr7:84265410 | 7 | 84.26541 | 0.0008798686 | Credible Set | purple |
| chr7:84265685 | 7 | 84.26569 | 0.0016549308 | Credible Set | purple |
| chr7:84327288 | 7 | 84.32729 | 0.0010533763 | Credible Set | purple |
| chr7:84327808 | 7 | 84.32781 | 0.0012331951 | Credible Set | purple |
| chr7:84327850 | 7 | 84.32785 | 0.0012347214 | Credible Set | purple |
| chr7:84338595 | 7 | 84.33859 | 0.0019602362 | Credible Set | purple |
| chr7:84343118 | 7 | 84.34312 | 0.0131853721 | Credible Set | purple |
| chr7:84344805 | 7 | 84.34480 | 0.0025040726 | Credible Set | purple |
| chr7:84347423 | 7 | 84.34742 | 0.0029631230 | Credible Set | purple |
| chr7:84360038 | 7 | 84.36004 | 0.0025151078 | Credible Set | purple |
| chr7:84360626 | 7 | 84.36063 | 0.0008943498 | Credible Set | purple |
| chr7:84360642 | 7 | 84.36064 | 0.0008787910 | Credible Set | purple |
| chr7:84360703 | 7 | 84.36070 | 0.0541024657 | Credible Set | purple |
| chr7:84360714 | 7 | 84.36071 | 0.0582167848 | Credible Set | purple |
| chr7:84360716 | 7 | 84.36072 | 0.0582167848 | Credible Set | purple |
| chr7:84375526 | 7 | 84.37553 | 0.0491642099 | Credible Set | purple |
| chr7:84375577 | 7 | 84.37558 | 0.0522902104 | Credible Set | purple |
| chr7:84375642 | 7 | 84.37564 | 0.0543287044 | Credible Set | purple |
| chr7:84397575 | 7 | 84.39758 | 0.0877330639 | Credible Set | purple |
| chr7:84397597 | 7 | 84.39760 | 0.0880304180 | Credible Set | purple |
| chr7:84430102 | 7 | 84.43010 | 0.0145224597 | Credible Set | purple |
| chr7:84501082 | 7 | 84.50108 | 0.0180773249 | Credible Set | purple |
| chr7:84514082 | 7 | 84.51408 | 0.0280583866 | Credible Set | purple |
| chr7:84514444 | 7 | 84.51444 | 0.0112579762 | Credible Set | purple |
| chr7:84514505 | 7 | 84.51450 | 0.0106321952 | Credible Set | purple |
| chr7:84514564 | 7 | 84.51456 | 0.0130934303 | Credible Set | purple |
| chr7:84514575 | 7 | 84.51457 | 0.0110187035 | Credible Set | purple |
| chr7:84514734 | 7 | 84.51473 | 0.0302152868 | Credible Set | purple |
| chr7:84543346 | 7 | 84.54335 | 0.0373559777 | Credible Set | purple |
| chr7:84543355 | 7 | 84.54336 | 0.0389205248 | Credible Set | purple |
| chr7:84543782 | 7 | 84.54378 | 0.1203381884 | Credible Set | purple |
| chr7:84619632 | 7 | 84.61963 | 0.0413522640 | Credible Set | purple |
| chr7:84636759 | 7 | 84.63676 | 0.0107822552 | Credible Set | purple |
| chr7:84653095 | 7 | 84.65309 | 0.0107428651 | Credible Set | purple |

Fine−mapping Regions

| snp | chr | pos (Mb) | pp | group | color |
|---|---|---|---|---|---|
| chr7:84733186 | 7 | 84.73319 | 0.0101730164 | Credible Set | purple |
| chr7:85533207 | 7 | 85.53321 | 0.0055404757 | Credible Set | purple |
| chr7:85543843 | 7 | 85.54384 | 0.0064939262 | Credible Set | purple |
| chr7:85656243 | 7 | 85.65624 | 0.0008365762 | Credible Set | purple |
| chr7:85677494 | 7 | 85.67749 | 0.0032155380 | Credible Set | purple |
| chr7:85712663 | 7 | 85.71266 | 0.0035325306 | Credible Set | purple |
| chr7:85712975 | 7 | 85.71298 | 0.0031447501 | Credible Set | purple |
| chr7:85840665 | 7 | 85.84067 | 0.0035804567 | Credible Set | purple |
| chr7:86053386 | 7 | 86.05339 | 0.0018307574 | Credible Set | purple |
| chr7:86237374 | 7 | 86.23737 | 0.0008958302 | Credible Set | purple |
| chr7:86582139 | 7 | 86.58214 | 0.0007973163 | Credible Set | purple |
| chr7:86645719 | 7 | 86.64572 | 0.0021693658 | Credible Set | purple |
| chr7:86645792 | 7 | 86.64579 | 0.0022281328 | Credible Set | purple |
| chr7:87246786 | 7 | 87.24679 | 0.0014708473 | Credible Set | purple |
| chr7:87255156 | 7 | 87.25516 | 0.0014480939 | Credible Set | purple |
| chr7:87554961 | 7 | 87.55496 | 0.0022912610 | Credible Set | purple |
| chr7:87642045 | 7 | 87.64204 | 0.0022093291 | Credible Set | purple |
| chr7:87665727 | 7 | 87.66573 | 0.0040732812 | Credible Set | purple |

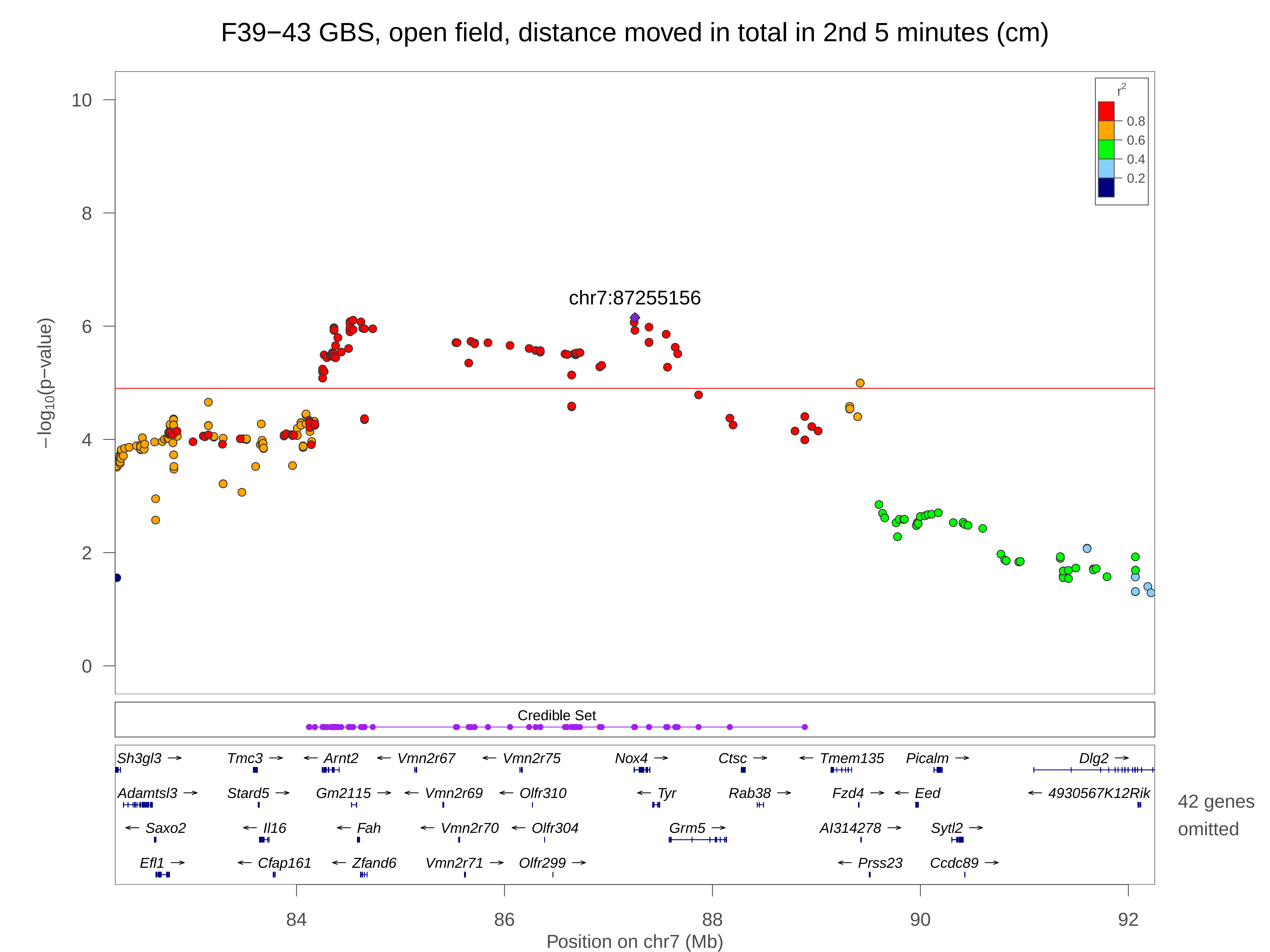
F39−43 GBS, open field, distance moved in total in 2nd 5 minutes (cm)
chr7:87255156
r2
0.8
0.6
0.4
0.2
−log10(p−value)
10
8
6
4
2
0
Credible Set
Sh3gl3 →
Tmc3 →
← Arnt2
← Vmn2r67
← Vmn2r75
Nox4 →
Ctsc →
← Tmem135
Picalm →
Dlg2 →
Adamtsl3 →
Stard5 →
Gm2115 →
← Vmn2r69
← Olfr310
← Tyr
Rab38 →
Fzd4 →
← Eed
← 4930567K12Rik
← Saxo2
← Il16
← Fah
← Vmn2r70
← Olfr304
Grm5 →
AI314278 →
Sytl2 →
Efl1 →
← Cfap161
← Zfand6
Vmn2r71 →
Olfr299 →
← Prss23
Ccdc89 →
42 genes omitted
84
86
88
90
92
Position on chr7 (Mb)

| | |
|---|---|
| date: | Thu Oct 25 17:25:16 2018 |
| build: | F3943GBS |
| display range: | chr7:82255156−92255156 [82255156−92255156] |
| hilite range: | 0 − 0 [ 0 − 0 ] |
| reference SNP: | chr7:87255156 |
| number of SNPs plotted: | 313 |
| min P−value: | 7.04E−7 [chr7:87255156] |
| max P−value: | 5.12E−2 [chr7:92218225] |
| omitted Genes: | 4933406J10Rik, Mex3b, 4933430H16Rik |
| omitted Genes: | Mesdc1, Mesdc2, Cemip |
| omitted Genes: | Abhd17c, 2610206C17Rik, Olfr291 |
| omitted Genes: | Olfr290, Vmn2r65, Vmn2r66 |
| omitted Genes: | Vmn2r68, Vmn2r72, Vmn2r73 |
| omitted Genes: | Vmn2r74, Vmn2r76, Olfr309 |
| omitted Genes: | Olfr308, Olfr307, Olfr305 |
| omitted Genes: | Olfr303, Olfr301, Olfr298 |
| omitted Genes: | Olfr297, Olfr295, Olfr294 |
| omitted Genes: | Olfr293, Olfr292, Folh1 |
| omitted Genes: | Vmn2r77, Vmn2r78, Vmn2r79 |
| omitted Genes: | Me3, Ccdc81, l7Rn6 |
| omitted Genes: | E230029C05Rik, 2310010J17Rik, Ccdc83 |
| omitted Genes: | Crebzf, Tmem126a, Tmem126b |

Fine−mapping Regions

| snp | chr | pos (Mb) | pp | group | color |
|---|---|---|---|---|---|
| chr7:84119855 | 7 | 84.11986 | 0.0005778993 | Credible Set | purple |
| chr7:84124190 | 7 | 84.12419 | 0.0005357889 | Credible Set | purple |
| chr7:84124515 | 7 | 84.12452 | 0.0005171650 | Credible Set | purple |
| chr7:84175856 | 7 | 84.17586 | 0.0004909846 | Credible Set | purple |
| chr7:84176482 | 7 | 84.17648 | 0.0005027755 | Credible Set | purple |
| chr7:84250985 | 7 | 84.25099 | 0.0037721754 | Credible Set | purple |
| chr7:84251011 | 7 | 84.25101 | 0.0042096789 | Credible Set | purple |
| chr7:84251159 | 7 | 84.25116 | 0.0029634419 | Credible Set | purple |
| chr7:84251209 | 7 | 84.25121 | 0.0029575140 | Credible Set | purple |
| chr7:84265410 | 7 | 84.26541 | 0.0038360642 | Credible Set | purple |
| chr7:84265685 | 7 | 84.26569 | 0.0073011626 | Credible Set | purple |
| chr7:84291540 | 7 | 84.29154 | 0.0065812100 | Credible Set | purple |
| chr7:84327288 | 7 | 84.32729 | 0.0070070876 | Credible Set | purple |
| chr7:84327808 | 7 | 84.32781 | 0.0070022368 | Credible Set | purple |
| chr7:84327850 | 7 | 84.32785 | 0.0071765186 | Credible Set | purple |
| chr7:84338595 | 7 | 84.33859 | 0.0074734726 | Credible Set | purple |
| chr7:84343118 | 7 | 84.34312 | 0.0072591545 | Credible Set | purple |
| chr7:84344805 | 7 | 84.34480 | 0.0079722874 | Credible Set | purple |
| chr7:84347423 | 7 | 84.34742 | 0.0077553426 | Credible Set | purple |
| chr7:84360038 | 7 | 84.36004 | 0.0075790774 | Credible Set | purple |
| chr7:84360224 | 7 | 84.36022 | 0.0209963132 | Credible Set | purple |
| chr7:84360247 | 7 | 84.36025 | 0.0206930607 | Credible Set | purple |
| chr7:84360258 | 7 | 84.36026 | 0.0211543126 | Credible Set | purple |
| chr7:84360279 | 7 | 84.36028 | 0.0205208054 | Credible Set | purple |
| chr7:84360626 | 7 | 84.36063 | 0.0067942350 | Credible Set | purple |
| chr7:84360642 | 7 | 84.36064 | 0.0067436055 | Credible Set | purple |
| chr7:84360703 | 7 | 84.36070 | 0.0190356718 | Credible Set | purple |
| chr7:84360714 | 7 | 84.36071 | 0.0194835692 | Credible Set | purple |
| chr7:84360716 | 7 | 84.36072 | 0.0194835692 | Credible Set | purple |
| chr7:84375526 | 7 | 84.37553 | 0.0105168391 | Credible Set | purple |
| chr7:84375577 | 7 | 84.37558 | 0.0065432131 | Credible Set | purple |
| chr7:84375642 | 7 | 84.37564 | 0.0065252941 | Credible Set | purple |
| chr7:84397575 | 7 | 84.39758 | 0.0143639053 | Credible Set | purple |
| chr7:84397597 | 7 | 84.39760 | 0.0143942156 | Credible Set | purple |
| chr7:84430102 | 7 | 84.43010 | 0.0081738215 | Credible Set | purple |
| chr7:84501082 | 7 | 84.50108 | 0.0093843197 | Credible Set | purple |

Fine−mapping Regions

| snp | chr | pos (Mb) | pp | group | color |
|---|---|---|---|---|---|
| chr7:84514082 | 7 | 84.51408 | 0.0201037678 | Credible Set | purple |
| chr7:84514444 | 7 | 84.51444 | 0.0188191681 | Credible Set | purple |
| chr7:84514505 | 7 | 84.51450 | 0.0180192757 | Credible Set | purple |
| chr7:84514564 | 7 | 84.51456 | 0.0268849195 | Credible Set | purple |
| chr7:84514575 | 7 | 84.51457 | 0.0249583567 | Credible Set | purple |
| chr7:84514734 | 7 | 84.51473 | 0.0211992357 | Credible Set | purple |
| chr7:84543346 | 7 | 84.54335 | 0.0196454568 | Credible Set | purple |
| chr7:84543355 | 7 | 84.54336 | 0.0195611893 | Credible Set | purple |
| chr7:84543782 | 7 | 84.54378 | 0.0284006070 | Credible Set | purple |
| chr7:84619632 | 7 | 84.61963 | 0.0265908744 | Credible Set | purple |
| chr7:84636759 | 7 | 84.63676 | 0.0207510453 | Credible Set | purple |
| chr7:84653095 | 7 | 84.65309 | 0.0203711319 | Credible Set | purple |
| chr7:84653660 | 7 | 84.65366 | 0.0006114912 | Credible Set | purple |
| chr7:84653702 | 7 | 84.65370 | 0.0005916057 | Credible Set | purple |
| chr7:84653707 | 7 | 84.65371 | 0.0006163577 | Credible Set | purple |
| chr7:84733186 | 7 | 84.73319 | 0.0202879621 | Credible Set | purple |
| chr7:85533207 | 7 | 85.53321 | 0.0118245423 | Credible Set | purple |
| chr7:85543843 | 7 | 85.54384 | 0.0117809342 | Credible Set | purple |
| chr7:85656243 | 7 | 85.65624 | 0.0053322504 | Credible Set | purple |
| chr7:85677494 | 7 | 85.67749 | 0.0124047503 | Credible Set | purple |
| chr7:85712663 | 7 | 85.71266 | 0.0115982331 | Credible Set | purple |
| chr7:85712975 | 7 | 85.71298 | 0.0112675652 | Credible Set | purple |
| chr7:85840665 | 7 | 85.84067 | 0.0117591812 | Credible Set | purple |
| chr7:86053386 | 7 | 86.05339 | 0.0105606946 | Credible Set | purple |
| chr7:86237374 | 7 | 86.23737 | 0.0093901368 | Credible Set | purple |
| chr7:86298979 | 7 | 86.29898 | 0.0086891344 | Credible Set | purple |
| chr7:86344420 | 7 | 86.34442 | 0.0082223415 | Credible Set | purple |
| chr7:86344437 | 7 | 86.34444 | 0.0081311776 | Credible Set | purple |
| chr7:86344488 | 7 | 86.34449 | 0.0086472449 | Credible Set | purple |
| chr7:86582139 | 7 | 86.58214 | 0.0075978445 | Credible Set | purple |
| chr7:86603770 | 7 | 86.60377 | 0.0074308701 | Credible Set | purple |
| chr7:86645719 | 7 | 86.64572 | 0.0033481407 | Credible Set | purple |
| chr7:86645792 | 7 | 86.64579 | 0.0033366141 | Credible Set | purple |
| chr7:86645947 | 7 | 86.64595 | 0.0009731313 | Credible Set | purple |
| chr7:86645990 | 7 | 86.64599 | 0.0010036619 | Credible Set | purple |
| chr7:86668998 | 7 | 86.66900 | 0.0076263605 | Credible Set | purple |

Fine−mapping Regions

| snp | chr | pos (Mb) | pp | group | color |
|---|---|---|---|---|---|
| chr7:86685426 | 7 | 86.68543 | 0.0073526957 | Credible Set | purple |
| chr7:86685507 | 7 | 86.68551 | 0.0073576016 | Credible Set | purple |
| chr7:86685648 | 7 | 86.68565 | 0.0076124420 | Credible Set | purple |
| chr7:86692214 | 7 | 86.69221 | 0.0077154897 | Credible Set | purple |
| chr7:86692491 | 7 | 86.69249 | 0.0078837695 | Credible Set | purple |
| chr7:86725518 | 7 | 86.72552 | 0.0079804769 | Credible Set | purple |
| chr7:86916161 | 7 | 86.91616 | 0.0045834015 | Credible Set | purple |
| chr7:86934710 | 7 | 86.93471 | 0.0048625197 | Credible Set | purple |
| chr7:87246786 | 7 | 87.24679 | 0.0261587611 | Credible Set | purple |
| chr7:87254690 | 7 | 87.25469 | 0.0191106937 | Credible Set | purple |
| chr7:87255156 | 7 | 87.25516 | 0.0315032437 | Credible Set | purple |
| chr7:87389952 | 7 | 87.38995 | 0.0216541466 | Credible Set | purple |
| chr7:87389989 | 7 | 87.38999 | 0.0118605169 | Credible Set | purple |
| chr7:87390014 | 7 | 87.39001 | 0.0120263294 | Credible Set | purple |
| chr7:87554961 | 7 | 87.55496 | 0.0163924310 | Credible Set | purple |
| chr7:87567685 | 7 | 87.56768 | 0.0045458375 | Credible Set | purple |
| chr7:87567686 | 7 | 87.56769 | 0.0045284478 | Credible Set | purple |
| chr7:87642045 | 7 | 87.64204 | 0.0098629187 | Credible Set | purple |
| chr7:87665727 | 7 | 87.66573 | 0.0076458768 | Credible Set | purple |
| chr7:87866323 | 7 | 87.86632 | 0.0015433559 | Credible Set | purple |
| chr7:88167216 | 7 | 88.16722 | 0.0006286716 | Credible Set | purple |
| chr7:88888200 | 7 | 88.88820 | 0.0006690654 | Credible Set | purple |

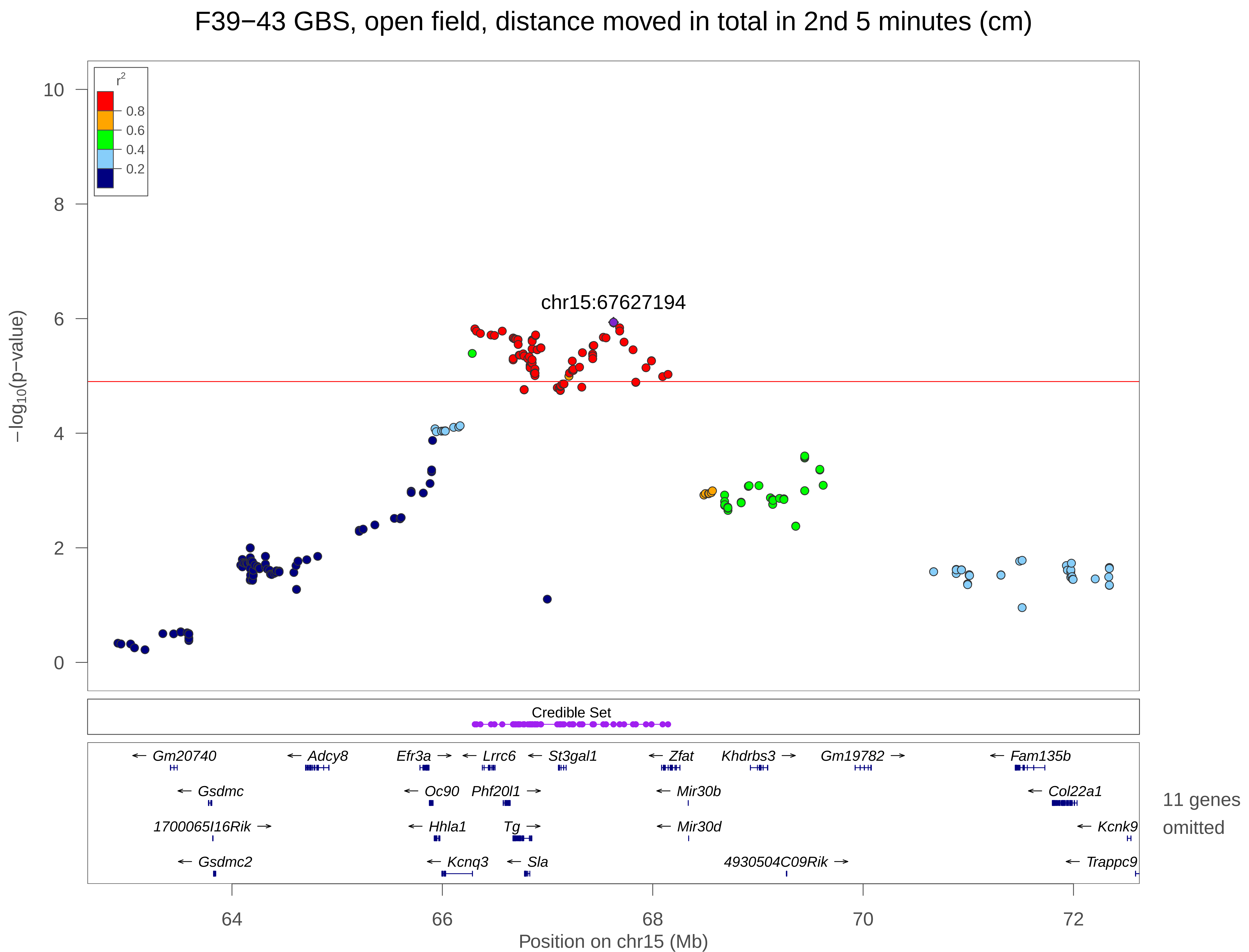
F39−43 GBS, open field, distance moved in total in 2nd 5 minutes (cm)
r2
0.8
0.6
0.4
0.2
chr15:67627194
−log10(p−value)
Credible Set
Gm20740
Adcy8
Efr3a
Lrrc6
St3gal1
Zfat
Khdrbs3
Gm19782
Fam135b
Gsdmc
Oc90
Phf20l1
Mir30b
Col22a1
1700065I16Rik
Hhla1
Tg
Mir30d
Kcnk9
Gsdmc2
Kcnq3
Sla
4930504C09Rik
Trappc9
11 genes omitted
Position on chr15 (Mb)

| | |
|---|---|
| date: | Thu Oct 25 17:25:06 2018 |
| build: | F3943GBS |
| display range: | chr15:62627194−72627194 [62627194−72627194] |
| hilite range: | 0 − 0 [ 0 − 0 ] |
| reference SNP: | chr15:67627194 |
| number of SNPs plotted: | 313 |
| min P−value: | 1.16E−6 [chr15:67627194] |
| max P−value: | 6.01E−1 [chr15:63173066] |
| omitted Genes: | Gsdmcl1, Gsdmc3, Gsdmcl2 |
| omitted Genes: | Gsdmc4, Gsdmcl−ps, Fam49b |
| omitted Genes: | Asap1, Tmem71, Wisp1 |
| omitted Genes: | Ndrg1, 1700012I11Rik |

Make more plots at http://csg.sph.umich.edu/locuszoom/

Fine−mapping Regions

| snp | chr | pos (Mb) | pp | group | color |
|---|---|---|---|---|---|
| chr15:66309070 | 15 | 66.30907 | 0.023207097 | Credible Set | purple |
| chr15:66324506 | 15 | 66.32451 | 0.021174756 | Credible Set | purple |
| chr15:66361813 | 15 | 66.36181 | 0.019390850 | Credible Set | purple |
| chr15:66462727 | 15 | 66.46273 | 0.018298992 | Credible Set | purple |
| chr15:66495529 | 15 | 66.49553 | 0.017979664 | Credible Set | purple |
| chr15:66569471 | 15 | 66.56947 | 0.021277214 | Credible Set | purple |
| chr15:66672984 | 15 | 66.67298 | 0.007050229 | Credible Set | purple |
| chr15:66672997 | 15 | 66.67300 | 0.007027169 | Credible Set | purple |
| chr15:66673010 | 15 | 66.67301 | 0.007009014 | Credible Set | purple |
| chr15:66673015 | 15 | 66.67302 | 0.007370279 | Credible Set | purple |
| chr15:66673436 | 15 | 66.67344 | 0.016272011 | Credible Set | purple |
| chr15:66673541 | 15 | 66.67354 | 0.016297356 | Credible Set | purple |
| chr15:66673940 | 15 | 66.67394 | 0.016276737 | Credible Set | purple |
| chr15:66685097 | 15 | 66.68510 | 0.015778110 | Credible Set | purple |
| chr15:66696518 | 15 | 66.69652 | 0.015588218 | Credible Set | purple |
| chr15:66699723 | 15 | 66.69972 | 0.015597573 | Credible Set | purple |
| chr15:66719126 | 15 | 66.71913 | 0.015330466 | Credible Set | purple |
| chr15:66719241 | 15 | 66.71924 | 0.015171761 | Credible Set | purple |
| chr15:66720772 | 15 | 66.72077 | 0.012702722 | Credible Set | purple |
| chr15:66730424 | 15 | 66.73042 | 0.008425202 | Credible Set | purple |
| chr15:66737376 | 15 | 66.73738 | 0.008431567 | Credible Set | purple |
| chr15:66768701 | 15 | 66.76870 | 0.008860071 | Credible Set | purple |
| chr15:66777400 | 15 | 66.77740 | 0.008145108 | Credible Set | purple |
| chr15:66777425 | 15 | 66.77742 | 0.008157646 | Credible Set | purple |
| chr15:66777461 | 15 | 66.77746 | 0.008131774 | Credible Set | purple |
| chr15:66777729 | 15 | 66.77773 | 0.002249629 | Credible Set | purple |
| chr15:66777730 | 15 | 66.77773 | 0.002246403 | Credible Set | purple |
| chr15:66777731 | 15 | 66.77773 | 0.002240508 | Credible Set | purple |
| chr15:66777743 | 15 | 66.77774 | 0.002235736 | Credible Set | purple |
| chr15:66812941 | 15 | 66.81294 | 0.007358563 | Credible Set | purple |
| chr15:66826934 | 15 | 66.82693 | 0.007915947 | Credible Set | purple |
| chr15:66834506 | 15 | 66.83451 | 0.005517590 | Credible Set | purple |
| chr15:66834532 | 15 | 66.83453 | 0.005630602 | Credible Set | purple |
| chr15:66834574 | 15 | 66.83457 | 0.005616528 | Credible Set | purple |
| chr15:66834575 | 15 | 66.83458 | 0.005757617 | Credible Set | purple |
| chr15:66834600 | 15 | 66.83460 | 0.005204227 | Credible Set | purple |

Fine−mapping Regions

| snp | chr | pos (Mb) | pp | group | color |
|---|---|---|---|---|---|
| chr15:66834621 | 15 | 66.83462 | 0.005392772 | Credible Set | purple |
| chr15:66834624 | 15 | 66.83462 | 0.005160620 | Credible Set | purple |
| chr15:66834637 | 15 | 66.83464 | 0.005175929 | Credible Set | purple |
| chr15:66853166 | 15 | 66.85317 | 0.010694344 | Credible Set | purple |
| chr15:66853439 | 15 | 66.85344 | 0.006159641 | Credible Set | purple |
| chr15:66853460 | 15 | 66.85346 | 0.006206085 | Credible Set | purple |
| chr15:66853489 | 15 | 66.85349 | 0.006225632 | Credible Set | purple |
| chr15:66853520 | 15 | 66.85352 | 0.007075779 | Credible Set | purple |
| chr15:66853893 | 15 | 66.85389 | 0.015165548 | Credible Set | purple |
| chr15:66853894 | 15 | 66.85389 | 0.014682533 | Credible Set | purple |
| chr15:66853923 | 15 | 66.85392 | 0.014734403 | Credible Set | purple |
| chr15:66853954 | 15 | 66.85395 | 0.014358853 | Credible Set | purple |
| chr15:66872519 | 15 | 66.87252 | 0.004252856 | Credible Set | purple |
| chr15:66880575 | 15 | 66.88057 | 0.003831402 | Credible Set | purple |
| chr15:66880583 | 15 | 66.88058 | 0.004922123 | Credible Set | purple |
| chr15:66880635 | 15 | 66.88063 | 0.004245451 | Credible Set | purple |
| chr15:66880638 | 15 | 66.88064 | 0.004176892 | Credible Set | purple |
| chr15:66886219 | 15 | 66.88622 | 0.017924386 | Credible Set | purple |
| chr15:66886266 | 15 | 66.88627 | 0.018358923 | Credible Set | purple |
| chr15:66901528 | 15 | 66.90153 | 0.010354701 | Credible Set | purple |
| chr15:66901548 | 15 | 66.90155 | 0.010354701 | Credible Set | purple |
| chr15:66901593 | 15 | 66.90159 | 0.010354701 | Credible Set | purple |
| chr15:66935376 | 15 | 66.93538 | 0.011140589 | Credible Set | purple |
| chr15:66935390 | 15 | 66.93539 | 0.011140589 | Credible Set | purple |
| chr15:66935391 | 15 | 66.93539 | 0.011140589 | Credible Set | purple |
| chr15:66935405 | 15 | 66.93541 | 0.011140589 | Credible Set | purple |
| chr15:66937231 | 15 | 66.93723 | 0.011182417 | Credible Set | purple |
| chr15:66937423 | 15 | 66.93742 | 0.011091471 | Credible Set | purple |
| chr15:67092666 | 15 | 67.09267 | 0.002405816 | Credible Set | purple |
| chr15:67111279 | 15 | 67.11128 | 0.002343101 | Credible Set | purple |
| chr15:67121756 | 15 | 67.12176 | 0.002523667 | Credible Set | purple |
| chr15:67126159 | 15 | 67.12616 | 0.002601266 | Credible Set | purple |
| chr15:67144054 | 15 | 67.14405 | 0.002779978 | Credible Set | purple |
| chr15:67156480 | 15 | 67.15648 | 0.002781250 | Credible Set | purple |
| chr15:67208761 | 15 | 67.20876 | 0.004234644 | Credible Set | purple |
| chr15:67234910 | 15 | 67.23491 | 0.004728429 | Credible Set | purple |

Fine−mapping Regions

| snp | chr | pos (Mb) | pp | group | color |
|---|---|---|---|---|---|
| chr15:67235072 | 15 | 67.23507 | 0.006718163 | Credible Set | purple |
| chr15:67245093 | 15 | 67.24509 | 0.004663734 | Credible Set | purple |
| chr15:67245123 | 15 | 67.24512 | 0.004892108 | Credible Set | purple |
| chr15:67304194 | 15 | 67.30419 | 0.005299837 | Credible Set | purple |
| chr15:67325463 | 15 | 67.32546 | 0.002461294 | Credible Set | purple |
| chr15:67331738 | 15 | 67.33174 | 0.009255784 | Credible Set | purple |
| chr15:67428857 | 15 | 67.42886 | 0.008783811 | Credible Set | purple |
| chr15:67429407 | 15 | 67.42941 | 0.008171220 | Credible Set | purple |
| chr15:67429420 | 15 | 67.42942 | 0.007334340 | Credible Set | purple |
| chr15:67433395 | 15 | 67.43340 | 0.012121725 | Credible Set | purple |
| chr15:67439974 | 15 | 67.43997 | 0.012223057 | Credible Set | purple |
| chr15:67530570 | 15 | 67.53057 | 0.016779644 | Credible Set | purple |
| chr15:67555683 | 15 | 67.55568 | 0.016388221 | Credible Set | purple |
| chr15:67627161 | 15 | 67.62716 | 0.029220941 | Credible Set | purple |
| chr15:67627183 | 15 | 67.62718 | 0.029871455 | Credible Set | purple |
| chr15:67627194 | 15 | 67.62719 | 0.029916701 | Credible Set | purple |
| chr15:67685224 | 15 | 67.68522 | 0.024198847 | Credible Set | purple |
| chr15:67685781 | 15 | 67.68578 | 0.021301057 | Credible Set | purple |
| chr15:67727448 | 15 | 67.72745 | 0.013939461 | Credible Set | purple |
| chr15:67812971 | 15 | 67.81297 | 0.010354803 | Credible Set | purple |
| chr15:67838708 | 15 | 67.83871 | 0.002956534 | Credible Set | purple |
| chr15:67838720 | 15 | 67.83872 | 0.002959211 | Credible Set | purple |
| chr15:67838721 | 15 | 67.83872 | 0.002986615 | Credible Set | purple |
| chr15:67935812 | 15 | 67.93581 | 0.005186890 | Credible Set | purple |
| chr15:67987913 | 15 | 67.98791 | 0.006800399 | Credible Set | purple |
| chr15:67987932 | 15 | 67.98793 | 0.006735037 | Credible Set | purple |
| chr15:68094377 | 15 | 68.09438 | 0.003697219 | Credible Set | purple |
| chr15:68145325 | 15 | 68.14532 | 0.004017346 | Credible Set | purple |

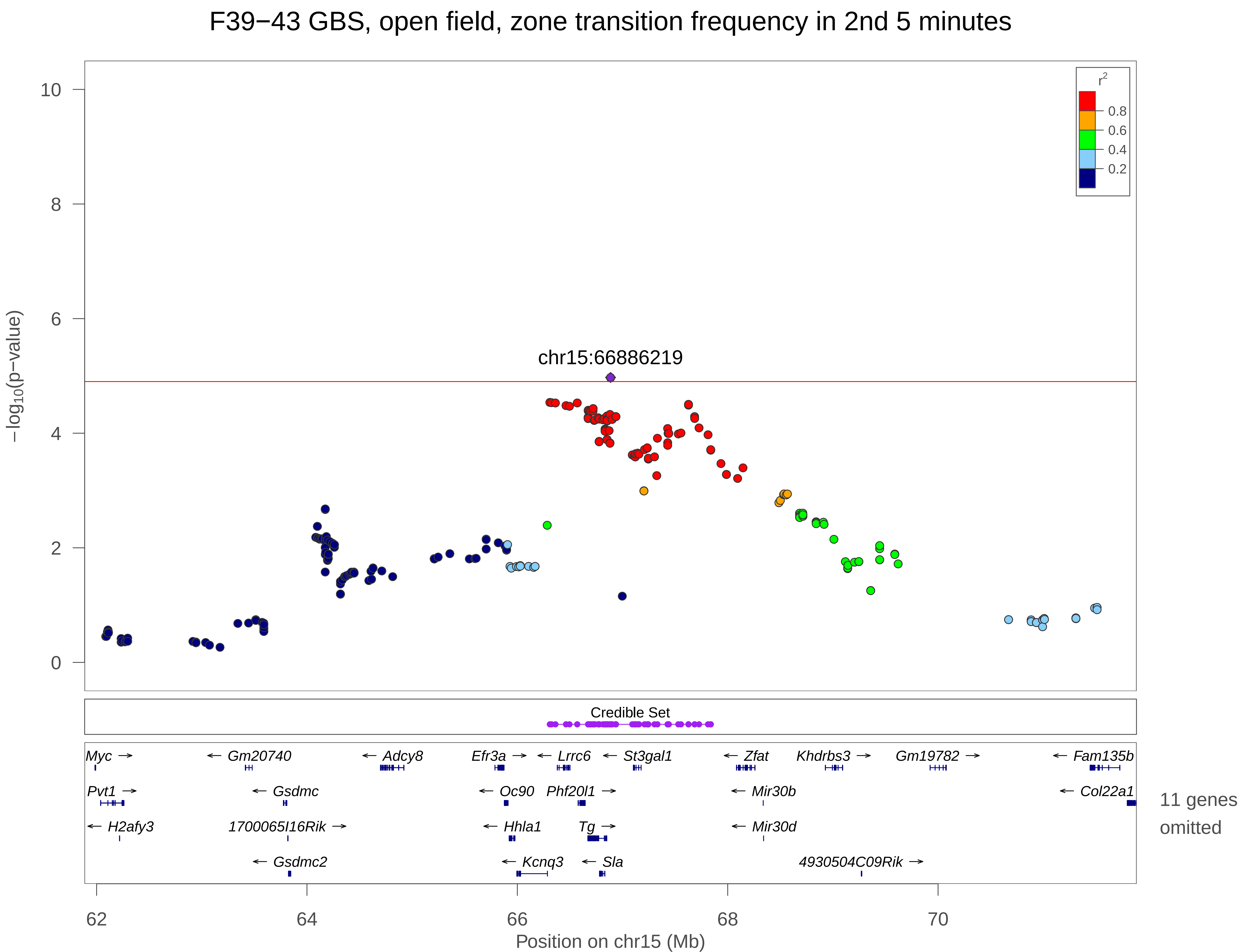
F39−43 GBS, open field, zone transition frequency in 2nd 5 minutes
−log10(p−value)
10
8
6
4
2
0
r2
0.8
0.6
0.4
0.2
chr15:66886219
Credible Set
Myc →
← Gm20740
← Adcy8
Efr3a →
← Lrrc6
← St3gal1
← Zfat
Khdrbs3 →
Gm19782 →
← Fam135b
Pvt1 →
← Gsdmc
← Oc90
Phf20l1 →
← Mir30b
← Col22a1
← H2afy3
1700065I16Rik →
← Hhla1
Tg →
← Mir30d
← Gsdmc2
← Kcnq3
← Sla
4930504C09Rik →
11 genes omitted
62
64
66
68
70
Position on chr15 (Mb)

| | |
|---|---|
| date: | Thu Oct 25 17:25:25 2018 |
| build: | F3943GBS |
| display range: | chr15:61886219−71886219 [61886219−71886219] |
| hilite range: | 0 − 0 [ 0 − 0 ] |
| reference SNP: | chr15:66886219 |
| number of SNPs plotted: | 313 |
| min P−value: | 1.07E−5 [chr15:66886219] |
| max P−value: | 5.45E−1 [chr15:63173066] |
| omitted Genes: | Gsdmcl1, Gsdmc3, Gsdmcl2 |
| omitted Genes: | Gsdmc4, Gsdmcl−ps, Fam49b |
| omitted Genes: | Asap1, Tmem71, Wisp1 |
| omitted Genes: | Ndrg1, 1700012I11Rik |

Make more plots at http://csg.sph.umich.edu/locuszoom/

Fine−mapping Regions

| snp | chr | pos (Mb) | pp | group | color |
|---|---|---|---|---|---|
| chr15:66309070 | 15 | 66.30907 | 0.020698958 | Credible Set | purple |
| chr15:66324506 | 15 | 66.32451 | 0.020509886 | Credible Set | purple |
| chr15:66361813 | 15 | 66.36181 | 0.020262884 | Credible Set | purple |
| chr15:66462727 | 15 | 66.46273 | 0.018331792 | Credible Set | purple |
| chr15:66495529 | 15 | 66.49553 | 0.017879978 | Credible Set | purple |
| chr15:66569471 | 15 | 66.56947 | 0.020294586 | Credible Set | purple |
| chr15:66672984 | 15 | 66.67298 | 0.011738641 | Credible Set | purple |
| chr15:66672997 | 15 | 66.67300 | 0.011658753 | Credible Set | purple |
| chr15:66673010 | 15 | 66.67301 | 0.011164878 | Credible Set | purple |
| chr15:66673015 | 15 | 66.67302 | 0.011209706 | Credible Set | purple |
| chr15:66673436 | 15 | 66.67344 | 0.015080409 | Credible Set | purple |
| chr15:66673541 | 15 | 66.67354 | 0.015274113 | Credible Set | purple |
| chr15:66673940 | 15 | 66.67394 | 0.015339929 | Credible Set | purple |
| chr15:66685097 | 15 | 66.68510 | 0.014987047 | Credible Set | purple |
| chr15:66696518 | 15 | 66.69652 | 0.014666886 | Credible Set | purple |
| chr15:66699723 | 15 | 66.69972 | 0.014803203 | Credible Set | purple |
| chr15:66719126 | 15 | 66.71913 | 0.014855476 | Credible Set | purple |
| chr15:66719241 | 15 | 66.71924 | 0.014872116 | Credible Set | purple |
| chr15:66720772 | 15 | 66.72077 | 0.016394163 | Credible Set | purple |
| chr15:66730424 | 15 | 66.73042 | 0.010415374 | Credible Set | purple |
| chr15:66737376 | 15 | 66.73738 | 0.010561024 | Credible Set | purple |
| chr15:66768701 | 15 | 66.76870 | 0.011528918 | Credible Set | purple |
| chr15:66777400 | 15 | 66.77740 | 0.010959676 | Credible Set | purple |
| chr15:66777425 | 15 | 66.77742 | 0.010990736 | Credible Set | purple |
| chr15:66777461 | 15 | 66.77746 | 0.011029969 | Credible Set | purple |
| chr15:66777706 | 15 | 66.77771 | 0.004662397 | Credible Set | purple |
| chr15:66777728 | 15 | 66.77773 | 0.004625887 | Credible Set | purple |
| chr15:66777729 | 15 | 66.77773 | 0.004709199 | Credible Set | purple |
| chr15:66777730 | 15 | 66.77773 | 0.004664560 | Credible Set | purple |
| chr15:66777731 | 15 | 66.77773 | 0.004679039 | Credible Set | purple |
| chr15:66777743 | 15 | 66.77774 | 0.004653623 | Credible Set | purple |
| chr15:66812941 | 15 | 66.81294 | 0.010659136 | Credible Set | purple |
| chr15:66826934 | 15 | 66.82693 | 0.011255160 | Credible Set | purple |
| chr15:66834506 | 15 | 66.83451 | 0.007469981 | Credible Set | purple |
| chr15:66834532 | 15 | 66.83453 | 0.007511882 | Credible Set | purple |
| chr15:66834574 | 15 | 66.83457 | 0.007328150 | Credible Set | purple |

Fine−mapping Regions

| snp | chr | pos (Mb) | pp | group | color |
|---|---|---|---|---|---|
| chr15:66834575 | 15 | 66.83458 | 0.007408892 | Credible Set | purple |
| chr15:66834600 | 15 | 66.83460 | 0.006935110 | Credible Set | purple |
| chr15:66834621 | 15 | 66.83462 | 0.007236619 | Credible Set | purple |
| chr15:66834624 | 15 | 66.83462 | 0.006981609 | Credible Set | purple |
| chr15:66834637 | 15 | 66.83464 | 0.006883963 | Credible Set | purple |
| chr15:66853166 | 15 | 66.85317 | 0.012349756 | Credible Set | purple |
| chr15:66853439 | 15 | 66.85344 | 0.004997227 | Credible Set | purple |
| chr15:66853460 | 15 | 66.85346 | 0.005061523 | Credible Set | purple |
| chr15:66853489 | 15 | 66.85349 | 0.005000614 | Credible Set | purple |
| chr15:66853520 | 15 | 66.85352 | 0.010203847 | Credible Set | purple |
| chr15:66853893 | 15 | 66.85389 | 0.011028860 | Credible Set | purple |
| chr15:66853894 | 15 | 66.85389 | 0.010888260 | Credible Set | purple |
| chr15:66853923 | 15 | 66.85392 | 0.010743687 | Credible Set | purple |
| chr15:66853954 | 15 | 66.85395 | 0.010387656 | Credible Set | purple |
| chr15:66872519 | 15 | 66.87252 | 0.007063003 | Credible Set | purple |
| chr15:66880575 | 15 | 66.88057 | 0.004445166 | Credible Set | purple |
| chr15:66880583 | 15 | 66.88058 | 0.013105398 | Credible Set | purple |
| chr15:66880635 | 15 | 66.88063 | 0.004416464 | Credible Set | purple |
| chr15:66880638 | 15 | 66.88064 | 0.004371283 | Credible Set | purple |
| chr15:66886219 | 15 | 66.88622 | 0.053782866 | Credible Set | purple |
| chr15:66886266 | 15 | 66.88627 | 0.052999560 | Credible Set | purple |
| chr15:66901528 | 15 | 66.90153 | 0.010841851 | Credible Set | purple |
| chr15:66901548 | 15 | 66.90155 | 0.010841851 | Credible Set | purple |
| chr15:66901593 | 15 | 66.90159 | 0.010841851 | Credible Set | purple |
| chr15:66935376 | 15 | 66.93538 | 0.012062147 | Credible Set | purple |
| chr15:66935390 | 15 | 66.93539 | 0.012062147 | Credible Set | purple |
| chr15:66935391 | 15 | 66.93539 | 0.012062147 | Credible Set | purple |
| chr15:66935405 | 15 | 66.93541 | 0.012062147 | Credible Set | purple |
| chr15:66937231 | 15 | 66.93723 | 0.012036848 | Credible Set | purple |
| chr15:66937423 | 15 | 66.93742 | 0.012043692 | Credible Set | purple |
| chr15:67092666 | 15 | 67.09267 | 0.002820092 | Credible Set | purple |
| chr15:67111279 | 15 | 67.11128 | 0.002696999 | Credible Set | purple |
| chr15:67121570 | 15 | 67.12157 | 0.002621619 | Credible Set | purple |
| chr15:67121571 | 15 | 67.12157 | 0.002611509 | Credible Set | purple |
| chr15:67121756 | 15 | 67.12176 | 0.002930503 | Credible Set | purple |
| chr15:67126159 | 15 | 67.12616 | 0.002958378 | Credible Set | purple |

Fine−mapping Regions

| snp | chr | pos (Mb) | pp | group | color |
|---|---|---|---|---|---|
| chr15:67144054 | 15 | 67.14405 | 0.003016865 | Credible Set | purple |
| chr15:67156480 | 15 | 67.15648 | 0.002902829 | Credible Set | purple |
| chr15:67208761 | 15 | 67.20876 | 0.003452345 | Credible Set | purple |
| chr15:67234910 | 15 | 67.23491 | 0.003626787 | Credible Set | purple |
| chr15:67235072 | 15 | 67.23507 | 0.003693117 | Credible Set | purple |
| chr15:67245093 | 15 | 67.24509 | 0.002409990 | Credible Set | purple |
| chr15:67245123 | 15 | 67.24512 | 0.002495326 | Credible Set | purple |
| chr15:67304194 | 15 | 67.30419 | 0.002621891 | Credible Set | purple |
| chr15:67331738 | 15 | 67.33174 | 0.005297626 | Credible Set | purple |
| chr15:67428857 | 15 | 67.42886 | 0.007662751 | Credible Set | purple |
| chr15:67429407 | 15 | 67.42941 | 0.004473028 | Credible Set | purple |
| chr15:67429420 | 15 | 67.42942 | 0.004049878 | Credible Set | purple |
| chr15:67433395 | 15 | 67.43340 | 0.006358279 | Credible Set | purple |
| chr15:67439974 | 15 | 67.43997 | 0.006350720 | Credible Set | purple |
| chr15:67530570 | 15 | 67.53057 | 0.006236362 | Credible Set | purple |
| chr15:67555683 | 15 | 67.55568 | 0.006458702 | Credible Set | purple |
| chr15:67627161 | 15 | 67.62716 | 0.018660436 | Credible Set | purple |
| chr15:67627183 | 15 | 67.62718 | 0.019240017 | Credible Set | purple |
| chr15:67627194 | 15 | 67.62719 | 0.019169201 | Credible Set | purple |
| chr15:67685224 | 15 | 67.68522 | 0.012050550 | Credible Set | purple |
| chr15:67685781 | 15 | 67.68578 | 0.011217577 | Credible Set | purple |
| chr15:67727448 | 15 | 67.72745 | 0.007843457 | Credible Set | purple |
| chr15:67812971 | 15 | 67.81297 | 0.006043895 | Credible Set | purple |
| chr15:67838708 | 15 | 67.83871 | 0.003403577 | Credible Set | purple |
| chr15:67838720 | 15 | 67.83872 | 0.003375711 | Credible Set | purple |
| chr15:67838721 | 15 | 67.83872 | 0.003438125 | Credible Set | purple |

F39−43 GBS, open field, minimum distance to center in 2nd 5 minutes (cm)

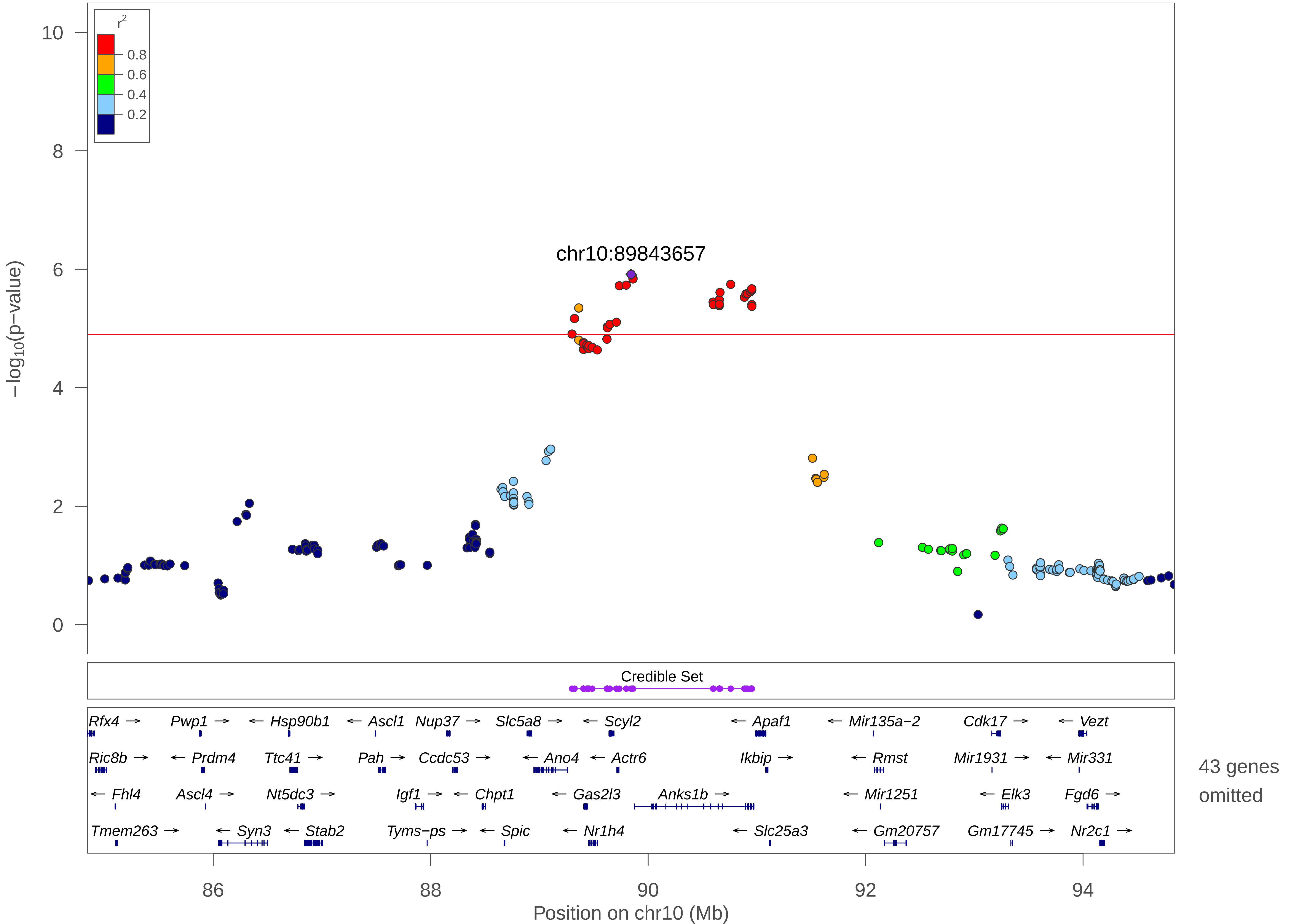

| | |
|---|---|
| date: | Thu Oct 25 17:25:35 2018 |
| build: | F3943GBS |
| display range: | chr10:84843657−94843657 [84843657−94843657] |
| hilite range: | 0 − 0 [ 0 − 0 ] |
| reference SNP: | chr10:89843657 |
| number of SNPs plotted: | 279 |
| min P−value: | 1.22E−6 [chr10:89843657] |
| max P−value: | 6.76E−1 [chr10:93034582] |
| omitted Genes: | Mterf2, Cry1, Btbd11 |
| omitted Genes: | Rtcb, Bpifc, Fbxo7 |
| omitted Genes: | 4930486F22Rik, Timp3, 1810014B01Rik |
| omitted Genes: | 1700113H08Rik, Pmch, Parpbp |
| omitted Genes: | Dram1, Gnptab, Sycp3 |
| omitted Genes: | Mybpc1, Gm4925, Arl1 |
| omitted Genes: | Utp20, Slc17a8, 1500026H17Rik |
| omitted Genes: | Uhrf1bp1l, Tmpo, Nedd1 |
| omitted Genes: | Mir7688, Lta4h, Hal |
| omitted Genes: | Amdhd1, Ccdc38, Snrpf |
| omitted Genes: | 4933408J17Rik, Ntn4, Gm8580 |
| omitted Genes: | Gm15915, Usp44, Metap2 |
| omitted Genes: | Ndufa12, Gm4792, Tmcc3 |
| omitted Genes: | Mir7211, Cep83os, Cep83 |
| omitted Genes: | Plxnc1 |

Make more plots at http://csg.sph.umich.edu/locuszoom/

Fine−mapping Regions

| snp | chr | pos (Mb) | pp | group | color |
|---|---|---|---|---|---|
| chr10:89300286 | 10 | 89.30029 | 0.007881034 | Credible Set | purple |
| chr10:89323084 | 10 | 89.32308 | 0.014030792 | Credible Set | purple |
| chr10:89405522 | 10 | 89.40552 | 0.004490975 | Credible Set | purple |
| chr10:89405556 | 10 | 89.40556 | 0.005729703 | Credible Set | purple |
| chr10:89405564 | 10 | 89.40556 | 0.005729703 | Credible Set | purple |
| chr10:89405589 | 10 | 89.40559 | 0.005479471 | Credible Set | purple |
| chr10:89405627 | 10 | 89.40563 | 0.005476568 | Credible Set | purple |
| chr10:89435280 | 10 | 89.43528 | 0.005212697 | Credible Set | purple |
| chr10:89443390 | 10 | 89.44339 | 0.004782635 | Credible Set | purple |
| chr10:89454154 | 10 | 89.45415 | 0.004580889 | Credible Set | purple |
| chr10:89454848 | 10 | 89.45485 | 0.005135398 | Credible Set | purple |
| chr10:89485534 | 10 | 89.48553 | 0.004825721 | Credible Set | purple |
| chr10:89621284 | 10 | 89.62128 | 0.006538935 | Credible Set | purple |
| chr10:89621319 | 10 | 89.62132 | 0.006507063 | Credible Set | purple |
| chr10:89625488 | 10 | 89.62549 | 0.010285437 | Credible Set | purple |
| chr10:89625559 | 10 | 89.62556 | 0.009912130 | Credible Set | purple |
| chr10:89647994 | 10 | 89.64799 | 0.011330864 | Credible Set | purple |
| chr10:89708192 | 10 | 89.70819 | 0.012229407 | Credible Set | purple |
| chr10:89734590 | 10 | 89.73459 | 0.047512984 | Credible Set | purple |
| chr10:89798643 | 10 | 89.79864 | 0.048713569 | Credible Set | purple |
| chr10:89843657 | 10 | 89.84366 | 0.072985672 | Credible Set | purple |
| chr10:89860498 | 10 | 89.86050 | 0.064329409 | Credible Set | purple |
| chr10:89860846 | 10 | 89.86085 | 0.061311481 | Credible Set | purple |
| chr10:90597510 | 10 | 90.59751 | 0.025879311 | Credible Set | purple |
| chr10:90599690 | 10 | 90.59969 | 0.023518083 | Credible Set | purple |
| chr10:90654718 | 10 | 90.65472 | 0.028128409 | Credible Set | purple |
| chr10:90654994 | 10 | 90.65499 | 0.022586308 | Credible Set | purple |
| chr10:90654998 | 10 | 90.65500 | 0.023434447 | Credible Set | purple |
| chr10:90655002 | 10 | 90.65500 | 0.022854374 | Credible Set | purple |
| chr10:90655039 | 10 | 90.65504 | 0.023867980 | Credible Set | purple |
| chr10:90661222 | 10 | 90.66122 | 0.037154221 | Credible Set | purple |
| chr10:90759813 | 10 | 90.75981 | 0.050097333 | Credible Set | purple |
| chr10:90885383 | 10 | 90.88538 | 0.030949421 | Credible Set | purple |
| chr10:90899946 | 10 | 90.89995 | 0.035095899 | Credible Set | purple |
| chr10:90913583 | 10 | 90.91358 | 0.035510922 | Credible Set | purple |
| chr10:90940720 | 10 | 90.94072 | 0.038119699 | Credible Set | purple |

Fine−mapping Regions

| snp | chr | pos (Mb) | pp | group | color |
|---|---|---|---|---|---|
| chr10:90953385 | 10 | 90.95338 | 0.04030947 | Credible Set | purple |
| chr10:90953402 | 10 | 90.95340 | 0.04063134 | Credible Set | purple |
| chr10:90953425 | 10 | 90.95342 | 0.04243464 | Credible Set | purple |
| chr10:90954053 | 10 | 90.95405 | 0.02357875 | Credible Set | purple |
| chr10:90954116 | 10 | 90.95412 | 0.02200545 | Credible Set | purple |

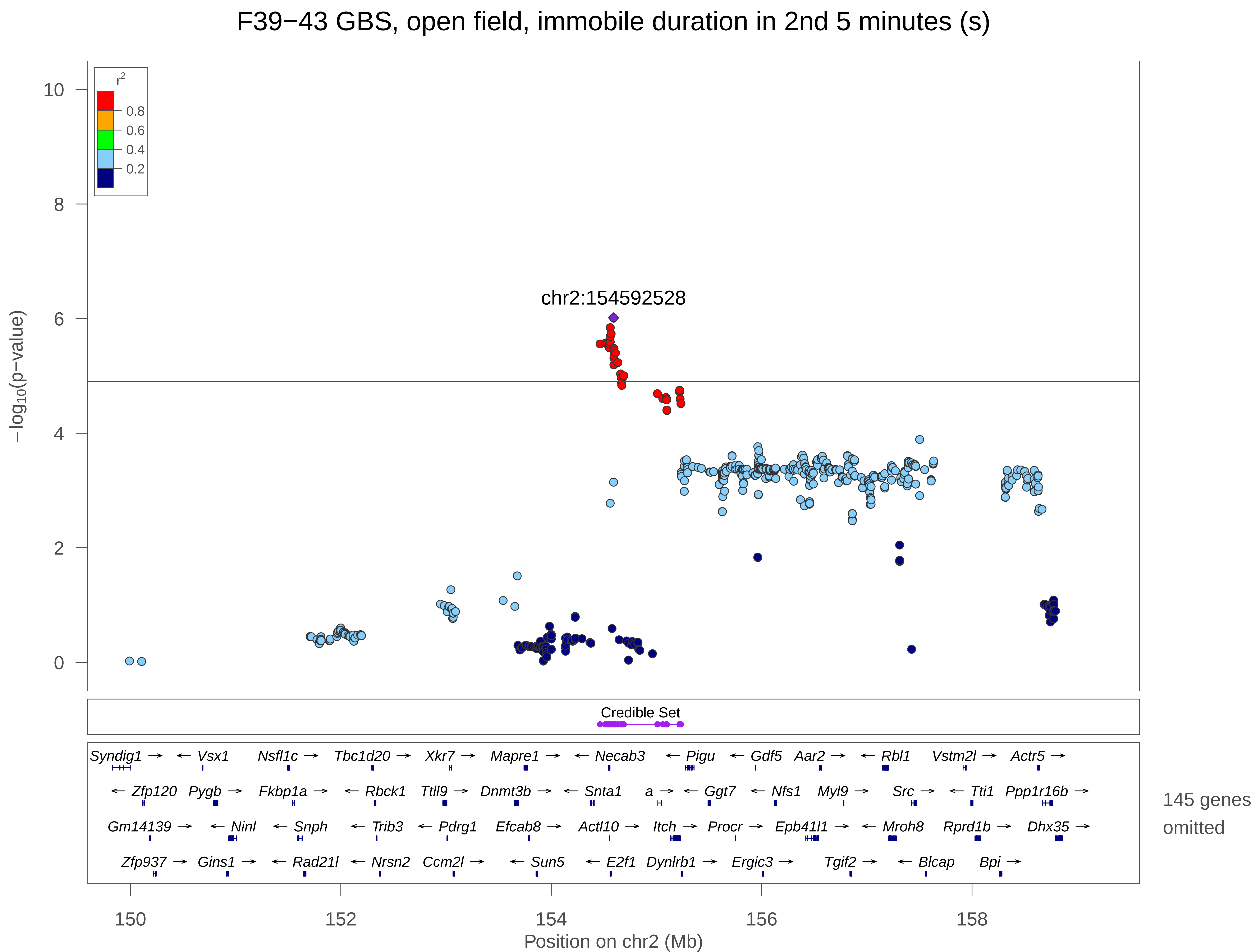

F39−43 GBS, open field, immobile duration in 2nd 5 minutes (s)
r2
0.8
0.6
0.4
0.2
chr2:154592528
−log10(p−value)
10
8
6
4
2
0
Credible Set
Syndig1 →
← Vsx1
Nsfl1c →
Tbc1d20 →
Xkr7 →
Mapre1 →
← Necab3
← Pigu
← Gdf5
Aar2 →
← Rbl1
Vstm2l →
Actr5 →
← Zfp120
Pygb →
Fkbp1a →
← Rbck1
Ttll9 →
Dnmt3b →
← Snta1
a →
← Ggt7
← Nfs1
Myl9 →
Src →
← Tti1
Ppp1r16b →
Gm14139 →
← Ninl
← Snph
← Trib3
← Pdrg1
Efcab8 →
Actl10 →
Itch →
Procr →
Epb41l1 →
← Mroh8
Rprd1b →
Dhx35 →
Zfp937 →
Gins1 →
← Rad21l
← Nrsn2
Ccm2l →
← Sun5
← E2f1
Dynlrb1 →
Ergic3 →
Tgif2 →
← Blcap
Bpi →
145 genes omitted
150
152
154
156
158
Position on chr2 (Mb)

date: Thu Oct 25 17:24:48 2018
build: F3943GBS
display range: chr2:149592528−159592528 [149592528−159592528]
hilite range: 0 − 0 [ 0 − 0 ]
reference SNP: chr2:154592528
number of SNPs plotted: 757
min P−value: 9.7E−7 [chr2:154592528]
max P−value: 9.63E−1 [chr2:150106309]
omitted Genes: Gm14124, 3300002I08Rik, Zfp442
omitted Genes: Zfp345, Cst7, Apmap
omitted Genes: Acss1, E130215H24Rik, Entpd6
omitted Genes: Abhd12, Nanp, Gm14151
omitted Genes: 4930442J19Rik, LOC100125594, 4921509C19Rik
omitted Genes: Sdcbp2, Tmem74b, Psmf1
omitted Genes: Rspo4, Angpt4, Mir1953
omitted Genes: Fam110a, Slc52a3, Scrt2
omitted Genes: Srxn1, Tcf15, Csnk2a1
omitted Genes: Gm14164, Sox12, Zcchc3
omitted Genes: 6820408C15Rik, Defb23, Defb20
omitted Genes: Defb22, Defb26, Defb28
omitted Genes: Defb29, Defb21, Defb19
omitted Genes: Defb45, Defb36, Defb25
omitted Genes: Rem1, H13, Mcts2
omitted Genes: Id1, Cox4i2, Bcl2l1
omitted Genes: Tpx2, Mir5622, Mylk2
omitted Genes: Foxs1, Dusp15, Hck
omitted Genes: Tm9sf4, Tspyl3, Plagl2
omitted Genes: Pofut1, Kif3b, 2500004C02Rik
omitted Genes: Asxl1, Nol4l, Commd7
omitted Genes: Bpifb2, Bpifb6, Bpifb3
omitted Genes: Bpifb4, Bpifa6, Bpifa2
omitted Genes: Bpifa3, Bpifa1, Bpifa5
omitted Genes: Bpifb1, Bpifb5, Bpifb9a
omitted Genes: Bpifb9b, Cdk5rap1, Cbfa2t2
omitted Genes: 1700003F12Rik, Pxmp4, Zfp341
omitted Genes: Chmp4b, Raly, Eif2s2
omitted Genes: Ahcy, 2310005A03Rik, Map1lc3a
omitted Genes: Mir695, Trp53inp2, Ncoa6
omitted Genes: Acss2, Acss2os, Gss
omitted Genes: Myh7b, Mir499, Trpc4ap

Fine−mapping Regions

| snp | chr | pos (Mb) | pp | group | color |
|---|---|---|---|---|---|
| chr2:154464466 | 2 | 154.4645 | 0.035286527 | Credible Set | purple |
| chr2:154513996 | 2 | 154.5140 | 0.036271254 | Credible Set | purple |
| chr2:154515752 | 2 | 154.5158 | 0.036417311 | Credible Set | purple |
| chr2:154516249 | 2 | 154.5162 | 0.036630633 | Credible Set | purple |
| chr2:154537995 | 2 | 154.5380 | 0.036352452 | Credible Set | purple |
| chr2:154538009 | 2 | 154.5380 | 0.036261717 | Credible Set | purple |
| chr2:154552607 | 2 | 154.5526 | 0.030876085 | Credible Set | purple |
| chr2:154552612 | 2 | 154.5526 | 0.030404513 | Credible Set | purple |
| chr2:154556468 | 2 | 154.5565 | 0.034983777 | Credible Set | purple |
| chr2:154560354 | 2 | 154.5604 | 0.038234556 | Credible Set | purple |
| chr2:154560360 | 2 | 154.5604 | 0.038191772 | Credible Set | purple |
| chr2:154560680 | 2 | 154.5607 | 0.047096613 | Credible Set | purple |
| chr2:154561127 | 2 | 154.5611 | 0.066223474 | Credible Set | purple |
| chr2:154569018 | 2 | 154.5690 | 0.052319012 | Credible Set | purple |
| chr2:154569036 | 2 | 154.5690 | 0.052334620 | Credible Set | purple |
| chr2:154592528 | 2 | 154.5925 | 0.096694605 | Credible Set | purple |
| chr2:154595741 | 2 | 154.5957 | 0.015770754 | Credible Set | purple |
| chr2:154595774 | 2 | 154.5958 | 0.020210672 | Credible Set | purple |
| chr2:154596294 | 2 | 154.5963 | 0.022143255 | Credible Set | purple |
| chr2:154597124 | 2 | 154.5971 | 0.029691904 | Credible Set | purple |
| chr2:154597483 | 2 | 154.5975 | 0.027252388 | Credible Set | purple |
| chr2:154610318 | 2 | 154.6103 | 0.024933804 | Credible Set | purple |
| chr2:154633945 | 2 | 154.6339 | 0.017170610 | Credible Set | purple |
| chr2:154659631 | 2 | 154.6596 | 0.011141211 | Credible Set | purple |
| chr2:154665835 | 2 | 154.6658 | 0.009651117 | Credible Set | purple |
| chr2:154670525 | 2 | 154.6705 | 0.007735798 | Credible Set | purple |
| chr2:154670528 | 2 | 154.6705 | 0.007793134 | Credible Set | purple |
| chr2:154670562 | 2 | 154.6706 | 0.008035652 | Credible Set | purple |
| chr2:154670571 | 2 | 154.6706 | 0.007175127 | Credible Set | purple |
| chr2:154686815 | 2 | 154.6868 | 0.010112313 | Credible Set | purple |
| chr2:154689824 | 2 | 154.6898 | 0.010371693 | Credible Set | purple |
| chr2:155009299 | 2 | 155.0093 | 0.005223306 | Credible Set | purple |
| chr2:155060609 | 2 | 155.0606 | 0.004315967 | Credible Set | purple |
| chr2:155091628 | 2 | 155.0916 | 0.004519797 | Credible Set | purple |
| chr2:155097606 | 2 | 155.0976 | 0.004250527 | Credible Set | purple |
| chr2:155097628 | 2 | 155.0976 | 0.004125484 | Credible Set | purple |

Fine−mapping Regions

| snp | chr | pos (Mb) | pp | group | color |
|---|---|---|---|---|---|
| chr2:155097650 | 2 | 155.0976 | 0.004125190 | Credible Set | purple |
| chr2:155220007 | 2 | 155.2200 | 0.005591515 | Credible Set | purple |
| chr2:155220040 | 2 | 155.2200 | 0.005591515 | Credible Set | purple |
| chr2:155220051 | 2 | 155.2201 | 0.005591515 | Credible Set | purple |
| chr2:155220725 | 2 | 155.2207 | 0.005946901 | Credible Set | purple |
| chr2:155224800 | 2 | 155.2248 | 0.004255826 | Credible Set | purple |
| chr2:155233265 | 2 | 155.2333 | 0.003591229 | Credible Set | purple |

F39−43 GBS, open field, immobile duration in 2nd 5 minutes (s)

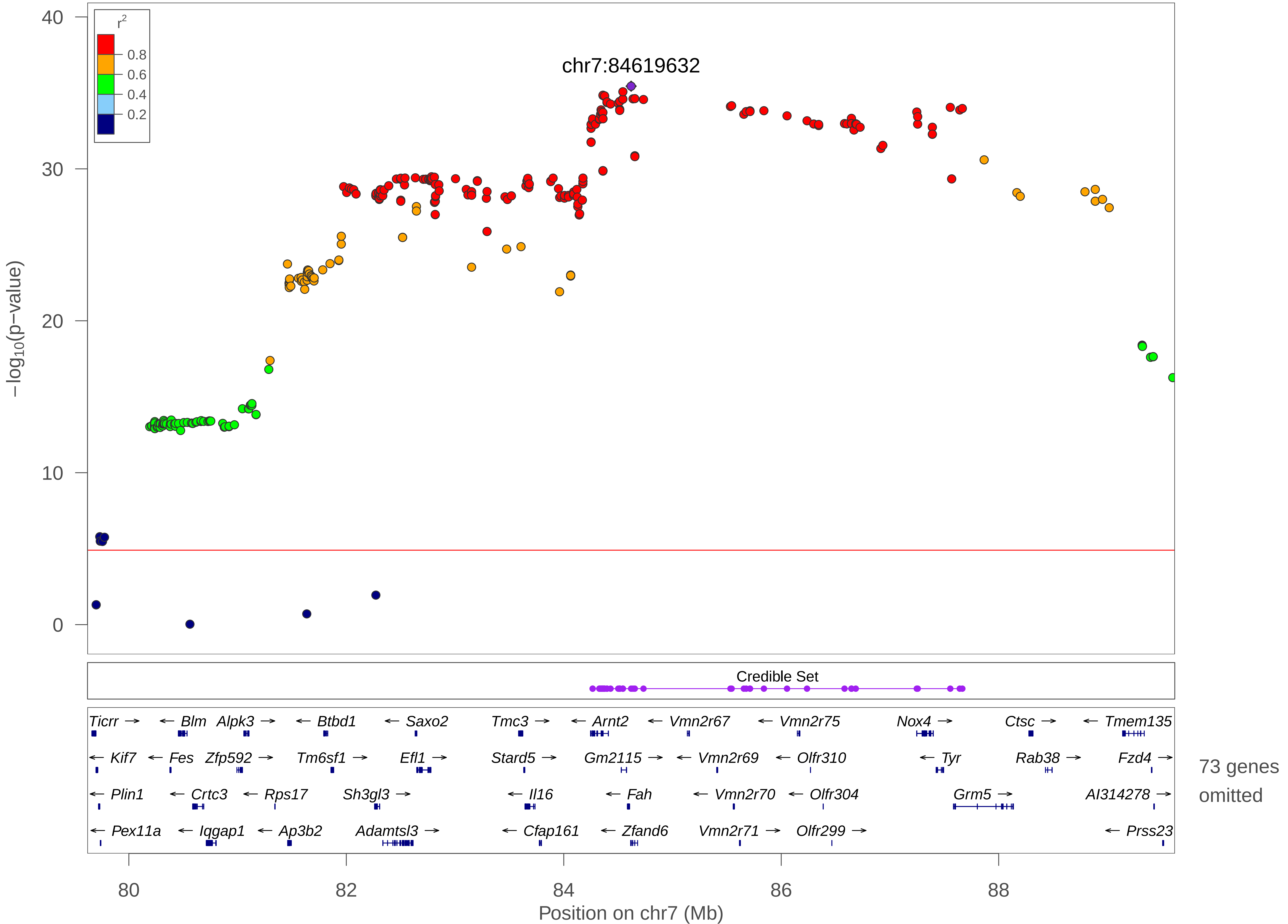

| | |
|---|---|
| date: | Thu Oct 25 17:24:57 2018 |
| build: | F3943GBS |
| display range: | chr7:79619632−89619632 [79619632−89619632] |
| hilite range: | 0 − 0 [ 0 − 0 ] |
| reference SNP: | chr7:84619632 |
| number of SNPs plotted: | 391 |
| min P−value: | 3.62E−36 [chr7:84619632] |
| max P−value: | 9.31E−1 [chr7:80561830] |
| omitted Genes: | Wdr93, Mesp1, Mesp2 |
| omitted Genes: | Anpep, Ap3s2, Arpin |
| omitted Genes: | Zfp710, Gm21057, Idh2 |
| omitted Genes: | Sema4b, Cib1, Gdpgp1 |
| omitted Genes: | Ttll13, Ngrn, Vps33b |
| omitted Genes: | Prc1, Rccd1, Unc45a |
| omitted Genes: | Hddc3, Man2a2, Furin |
| omitted Genes: | Gm15880, Zscan2, Wdr73 |
| omitted Genes: | Nmb, Sec11a, Slc28a1 |
| omitted Genes: | Pde8a, Cpeb1, BC048679 |
| omitted Genes: | 2900076A07Rik, Mir1839, Fsd2 |
| omitted Genes: | Whamm, Homer2, Fam103a1 |
| omitted Genes: | 3110040N11Rik, Hdgfrp3, Bnc1 |
| omitted Genes: | Gm20744, 4933406J10Rik, Mex3b |
| omitted Genes: | 4933430H16Rik, Mesdc1, Mesdc2 |
| omitted Genes: | Cemip, Abhd17c, 2610206C17Rik |
| omitted Genes: | Olfr291, Olfr290, Vmn2r65 |
| omitted Genes: | Vmn2r66, Vmn2r68, Vmn2r72 |
| omitted Genes: | Vmn2r73, Vmn2r74, Vmn2r76 |
| omitted Genes: | Olfr309, Olfr308, Olfr307 |
| omitted Genes: | Olfr305, Olfr303, Olfr301 |
| omitted Genes: | Olfr298, Olfr297, Olfr295 |
| omitted Genes: | Olfr294, Olfr293, Olfr292 |
| omitted Genes: | Folh1, Vmn2r77, Vmn2r78 |
| omitted Genes: | Vmn2r79 |

Make more plots at http://csg.sph.umich.edu/locuszoom/

Fine−mapping Regions

| snp | chr | pos (Mb) | pp | group | color |
|---|---|---|---|---|---|
| chr7:84265410 | 7 | 84.26541 | 0.001183419 | Credible Set | purple |
| chr7:84265685 | 7 | 84.26569 | 0.001509007 | Credible Set | purple |
| chr7:84327288 | 7 | 84.32729 | 0.001401618 | Credible Set | purple |
| chr7:84327808 | 7 | 84.32781 | 0.001552073 | Credible Set | purple |
| chr7:84327850 | 7 | 84.32785 | 0.001545965 | Credible Set | purple |
| chr7:84338595 | 7 | 84.33859 | 0.002998816 | Credible Set | purple |
| chr7:84343118 | 7 | 84.34312 | 0.005858123 | Credible Set | purple |
| chr7:84344805 | 7 | 84.34480 | 0.004029253 | Credible Set | purple |
| chr7:84347423 | 7 | 84.34742 | 0.004526111 | Credible Set | purple |
| chr7:84360038 | 7 | 84.36004 | 0.003963513 | Credible Set | purple |
| chr7:84360626 | 7 | 84.36063 | 0.001553613 | Credible Set | purple |
| chr7:84360642 | 7 | 84.36064 | 0.001525472 | Credible Set | purple |
| chr7:84360703 | 7 | 84.36070 | 0.050889386 | Credible Set | purple |
| chr7:84360714 | 7 | 84.36071 | 0.053201226 | Credible Set | purple |
| chr7:84360716 | 7 | 84.36072 | 0.053201226 | Credible Set | purple |
| chr7:84375526 | 7 | 84.37553 | 0.044215699 | Credible Set | purple |
| chr7:84375577 | 7 | 84.37558 | 0.047425076 | Credible Set | purple |
| chr7:84375642 | 7 | 84.37564 | 0.048536816 | Credible Set | purple |
| chr7:84397575 | 7 | 84.39758 | 0.017691865 | Credible Set | purple |
| chr7:84397597 | 7 | 84.39760 | 0.019455578 | Credible Set | purple |
| chr7:84430102 | 7 | 84.43010 | 0.014131985 | Credible Set | purple |
| chr7:84501082 | 7 | 84.50108 | 0.015769948 | Credible Set | purple |
| chr7:84514082 | 7 | 84.51408 | 0.020790519 | Credible Set | purple |
| chr7:84514444 | 7 | 84.51444 | 0.006400261 | Credible Set | purple |
| chr7:84514505 | 7 | 84.51450 | 0.006094613 | Credible Set | purple |
| chr7:84514564 | 7 | 84.51456 | 0.006123336 | Credible Set | purple |
| chr7:84514575 | 7 | 84.51457 | 0.005391904 | Credible Set | purple |
| chr7:84514734 | 7 | 84.51473 | 0.020708261 | Credible Set | purple |
| chr7:84543346 | 7 | 84.54335 | 0.030263351 | Credible Set | purple |
| chr7:84543355 | 7 | 84.54336 | 0.029230233 | Credible Set | purple |
| chr7:84543782 | 7 | 84.54378 | 0.089896894 | Credible Set | purple |
| chr7:84619632 | 7 | 84.61963 | 0.207457240 | Credible Set | purple |
| chr7:84636759 | 7 | 84.63676 | 0.030910316 | Credible Set | purple |
| chr7:84653095 | 7 | 84.65309 | 0.031909170 | Credible Set | purple |
| chr7:84733186 | 7 | 84.73319 | 0.027921200 | Credible Set | purple |
| chr7:85533207 | 7 | 85.53321 | 0.009769245 | Credible Set | purple |

Fine−mapping Regions

| snp | chr | pos (Mb) | pp | group | color |
|---|---|---|---|---|---|
| chr7:85543843 | 7 | 85.54384 | 0.0110083725 | Credible Set | purple |
| chr7:85656243 | 7 | 85.65624 | 0.0030240920 | Credible Set | purple |
| chr7:85677494 | 7 | 85.67749 | 0.0045119519 | Credible Set | purple |
| chr7:85712663 | 7 | 85.71266 | 0.0050950184 | Credible Set | purple |
| chr7:85712975 | 7 | 85.71298 | 0.0045901339 | Credible Set | purple |
| chr7:85840665 | 7 | 85.84067 | 0.0051785105 | Credible Set | purple |
| chr7:86053386 | 7 | 86.05339 | 0.0023920905 | Credible Set | purple |
| chr7:86237374 | 7 | 86.23737 | 0.0011361793 | Credible Set | purple |
| chr7:86582139 | 7 | 86.58214 | 0.0007483074 | Credible Set | purple |
| chr7:86645719 | 7 | 86.64572 | 0.0015866035 | Credible Set | purple |
| chr7:86645792 | 7 | 86.64579 | 0.0016436241 | Credible Set | purple |
| chr7:86645947 | 7 | 86.64595 | 0.0007312383 | Credible Set | purple |
| chr7:86645990 | 7 | 86.64599 | 0.0007466666 | Credible Set | purple |
| chr7:86685507 | 7 | 86.68551 | 0.0007236343 | Credible Set | purple |
| chr7:87246786 | 7 | 87.24679 | 0.0042594462 | Credible Set | purple |
| chr7:87255156 | 7 | 87.25516 | 0.0021102153 | Credible Set | purple |
| chr7:87554961 | 7 | 87.55496 | 0.0085326206 | Credible Set | purple |
| chr7:87642045 | 7 | 87.64204 | 0.0057800749 | Credible Set | purple |
| chr7:87665727 | 7 | 87.66573 | 0.0072430847 | Credible Set | purple |
