## Supplementary material for "Genome-wide association study in two cohorts from a multi-generational mouse advanced intercross line highlights the difficulty of replication": S8 Fig

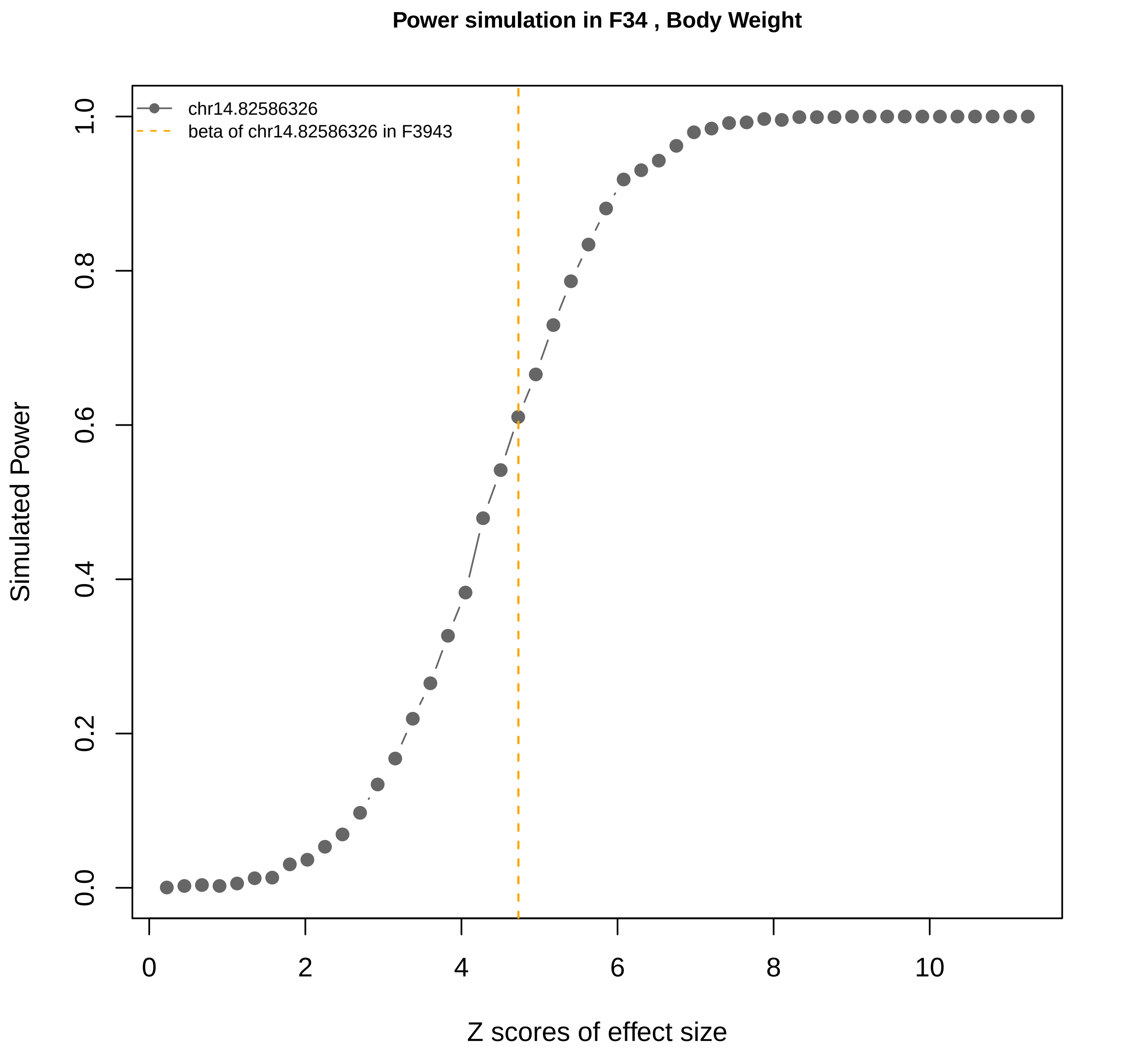

Power simulation in F34 , Body Weight
chr14.82586326
beta of chr14.82586326 in F3943
Simulated Power
Z scores of effect size
0.0
0.2
0.4
0.6
0.8
1.0
0
2
4
6
8
10

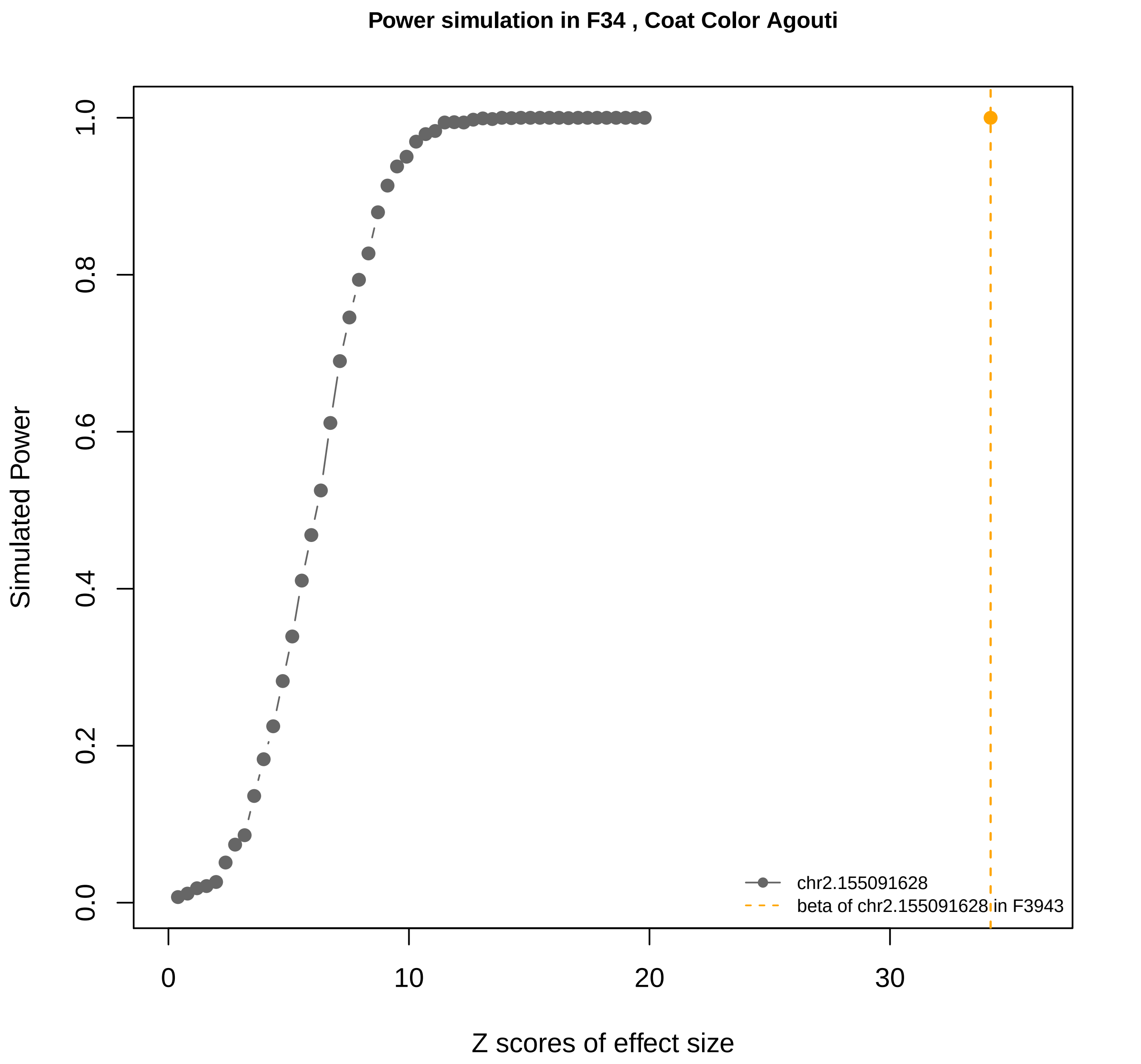

Power simulation in F34 , Coat Color Agouti
Simulated Power
0.0
0.2
0.4
0.6
0.8
1.0
0
10
20
30
Z scores of effect size
chr2.155091628
beta of chr2.155091628 in F3943

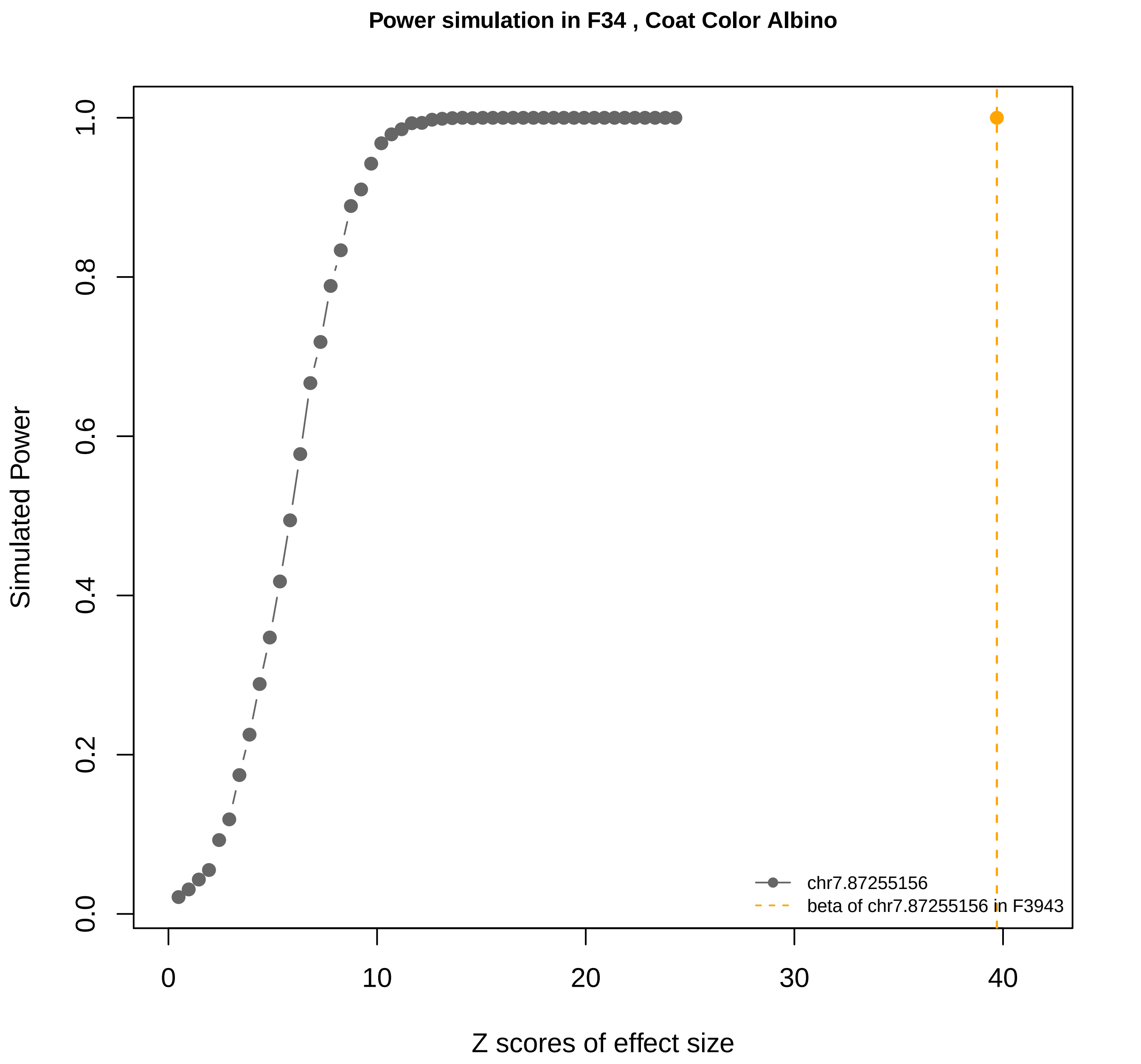
Power simulation in F34 , Coat Color Albino
Simulated Power
0.0
0.2
0.4
0.6
0.8
1.0
0
10
20
30
40
Z scores of effect size
chr7.87255156
beta of chr7.87255156 in F3943

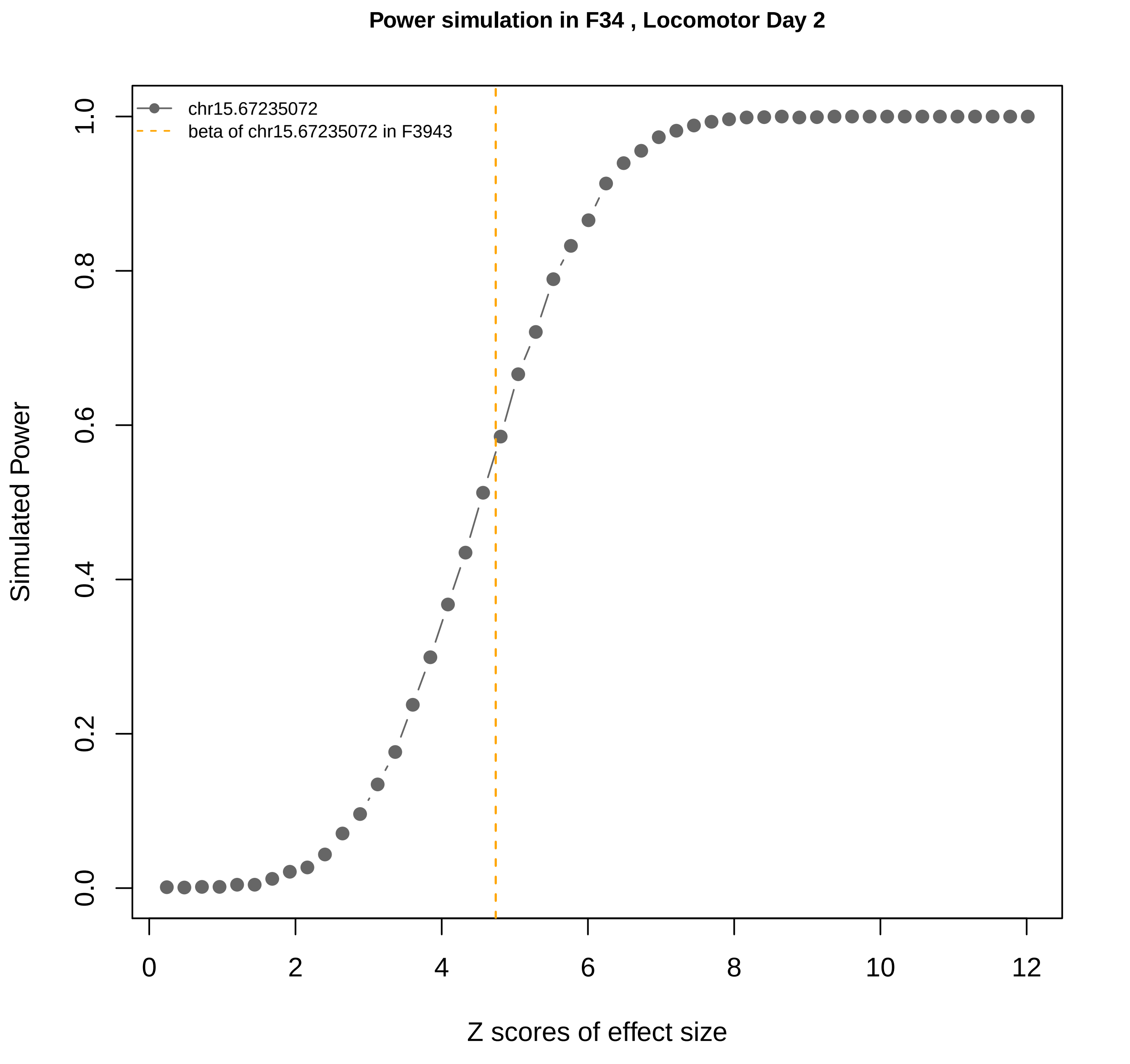

Power simulation in F34 , Locomotor Day 2
chr15.67235072
beta of chr15.67235072 in F3943
Simulated Power
Z scores of effect size
0.0
0.2
0.4
0.6
0.8
1.0
0
2
4
6
8
10
12

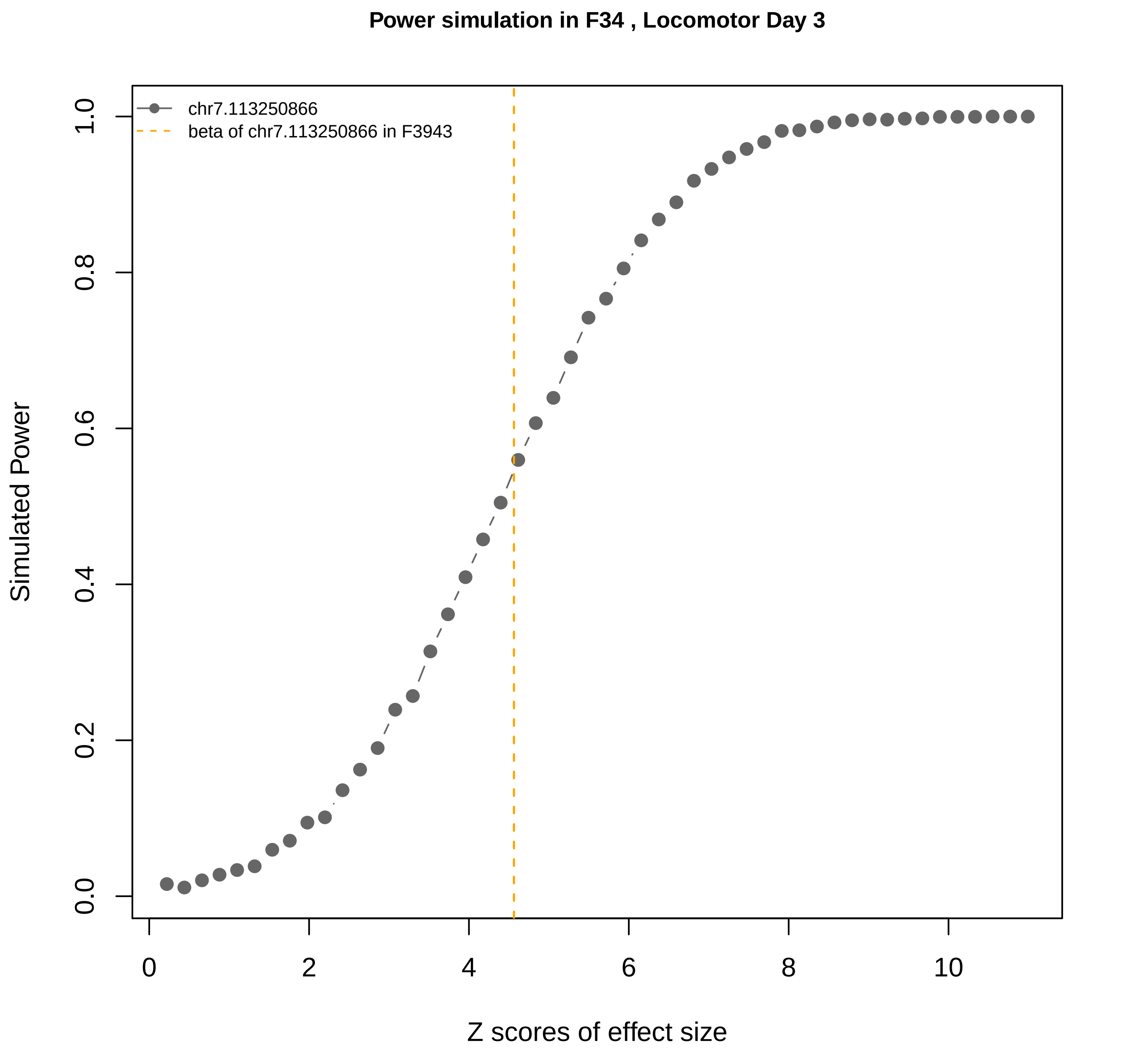

Power simulation in F34 , Locomotor Day 3
chr7.113250866
beta of chr7.113250866 in F3943
Simulated Power
0.0
0.2
0.4
0.6
0.8
1.0
0
2
4
6
8
10
Z scores of effect size

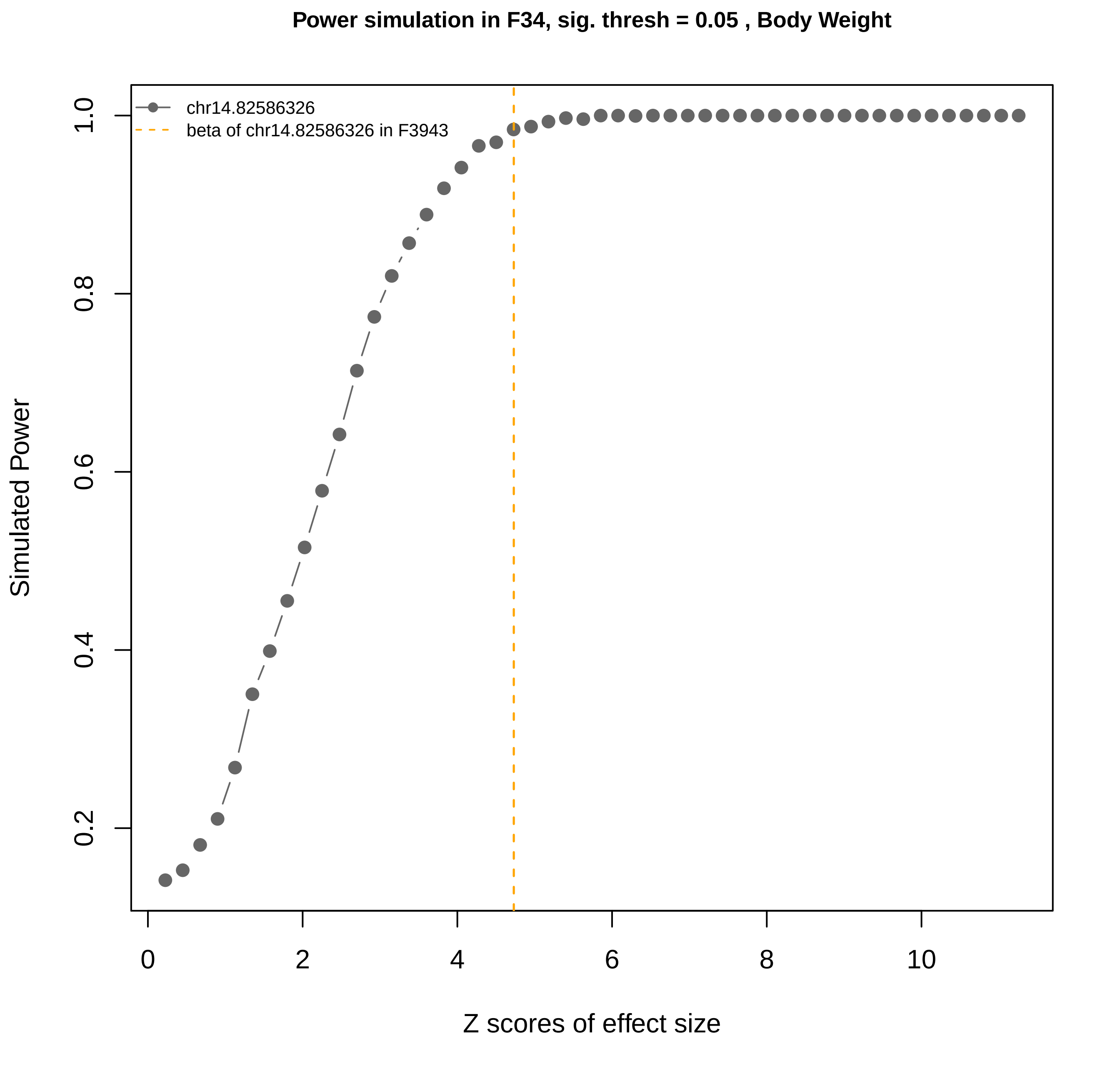

Power simulation in F34, sig. thresh = 0.05 , Body Weight
chr14.82586326
beta of chr14.82586326 in F3943
Simulated Power
0.2
0.4
0.6
0.8
1.0
0
2
4
6
8
10
Z scores of effect size

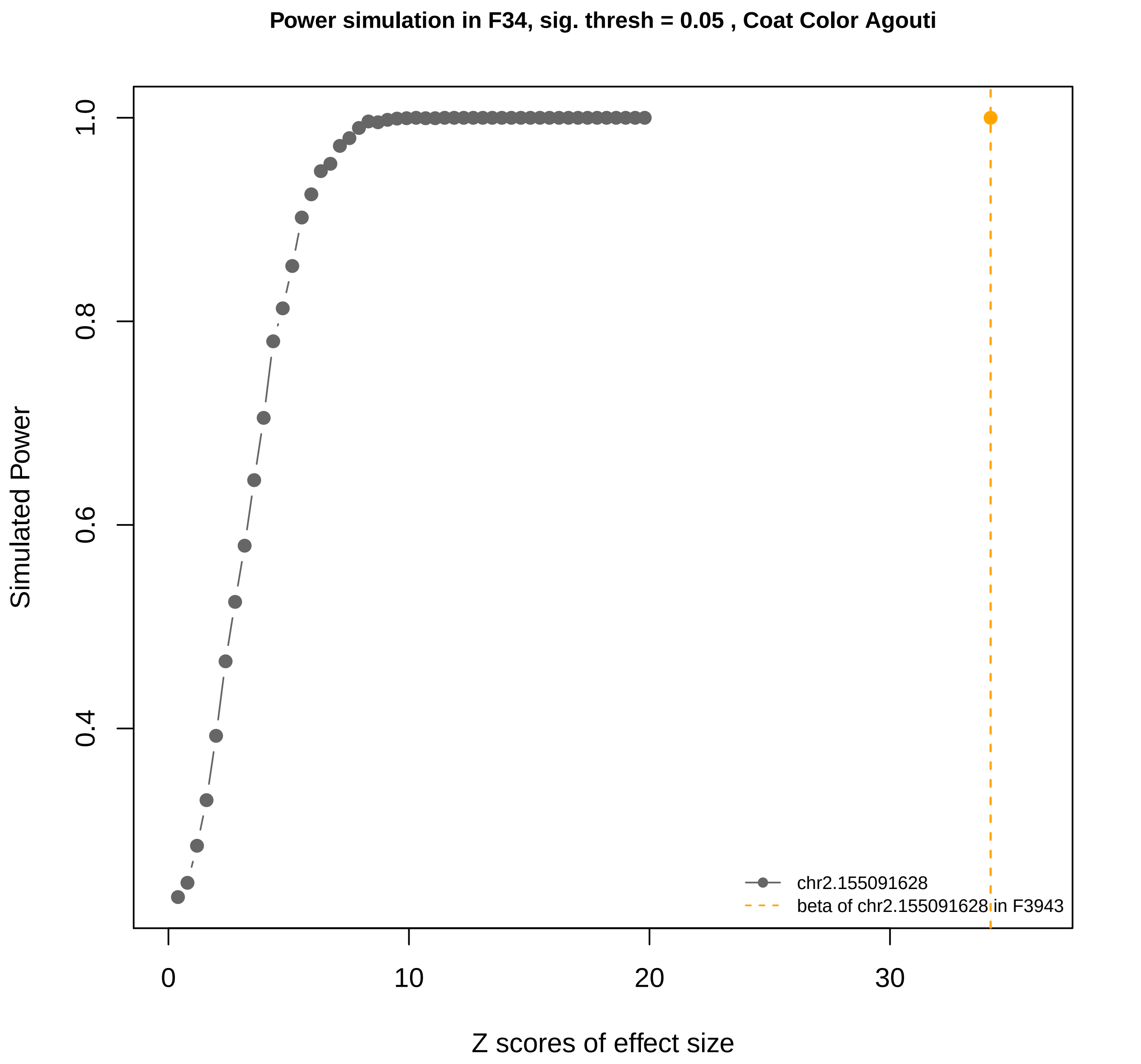
Power simulation in F34, sig. thresh = 0.05 , Coat Color Agouti
Simulated Power
Z scores of effect size
0.4
0.6
0.8
1.0
0
10
20
30
chr2.155091628
beta of chr2.155091628 in F3943

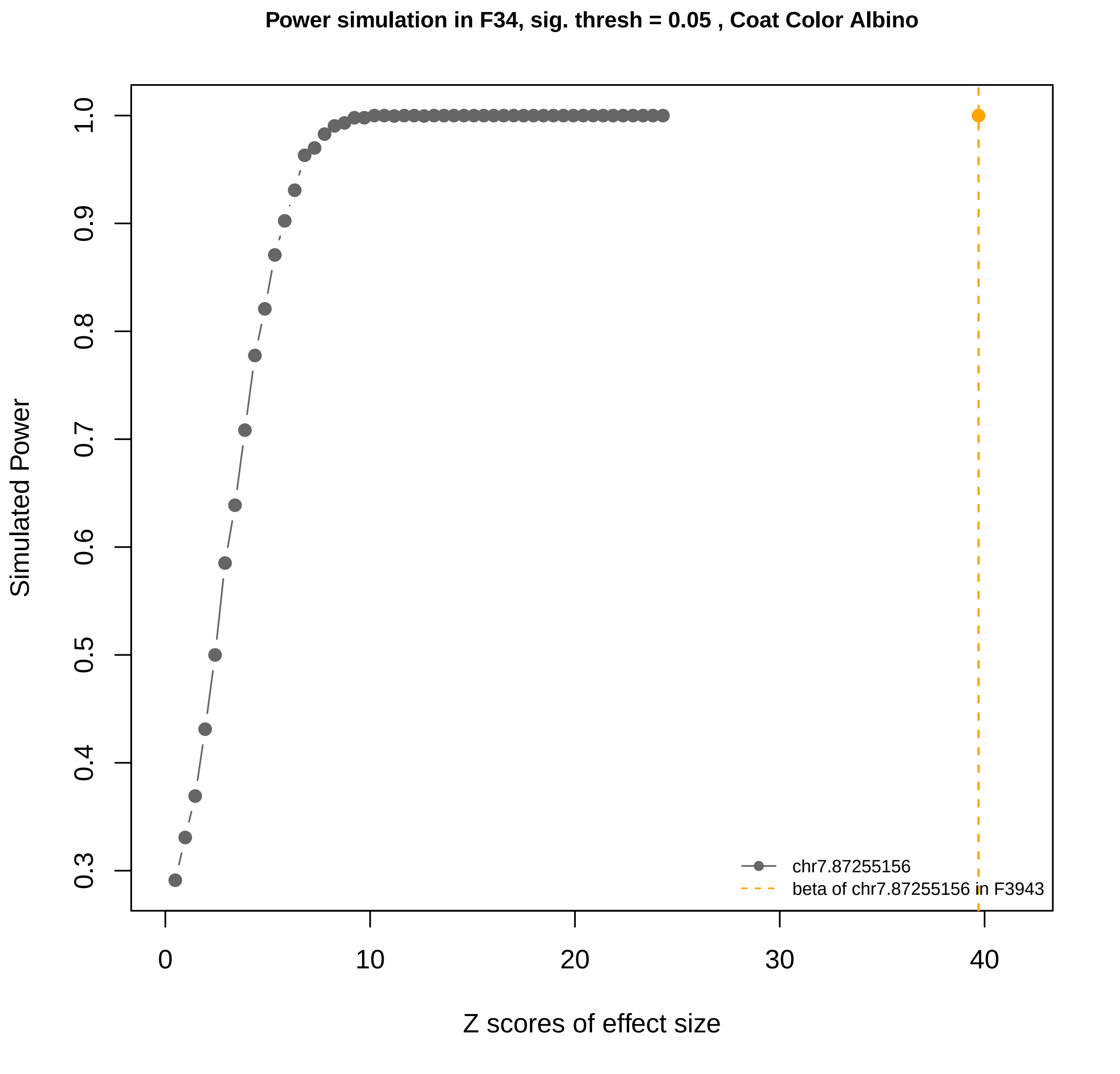

Power simulation in F34, sig. thresh = 0.05 , Coat Color Albino
Simulated Power
0.3
0.4
0.5
0.6
0.7
0.8
0.9
1.0
0
10
20
30
40
Z scores of effect size
chr7.87255156
beta of chr7.87255156 in F3943

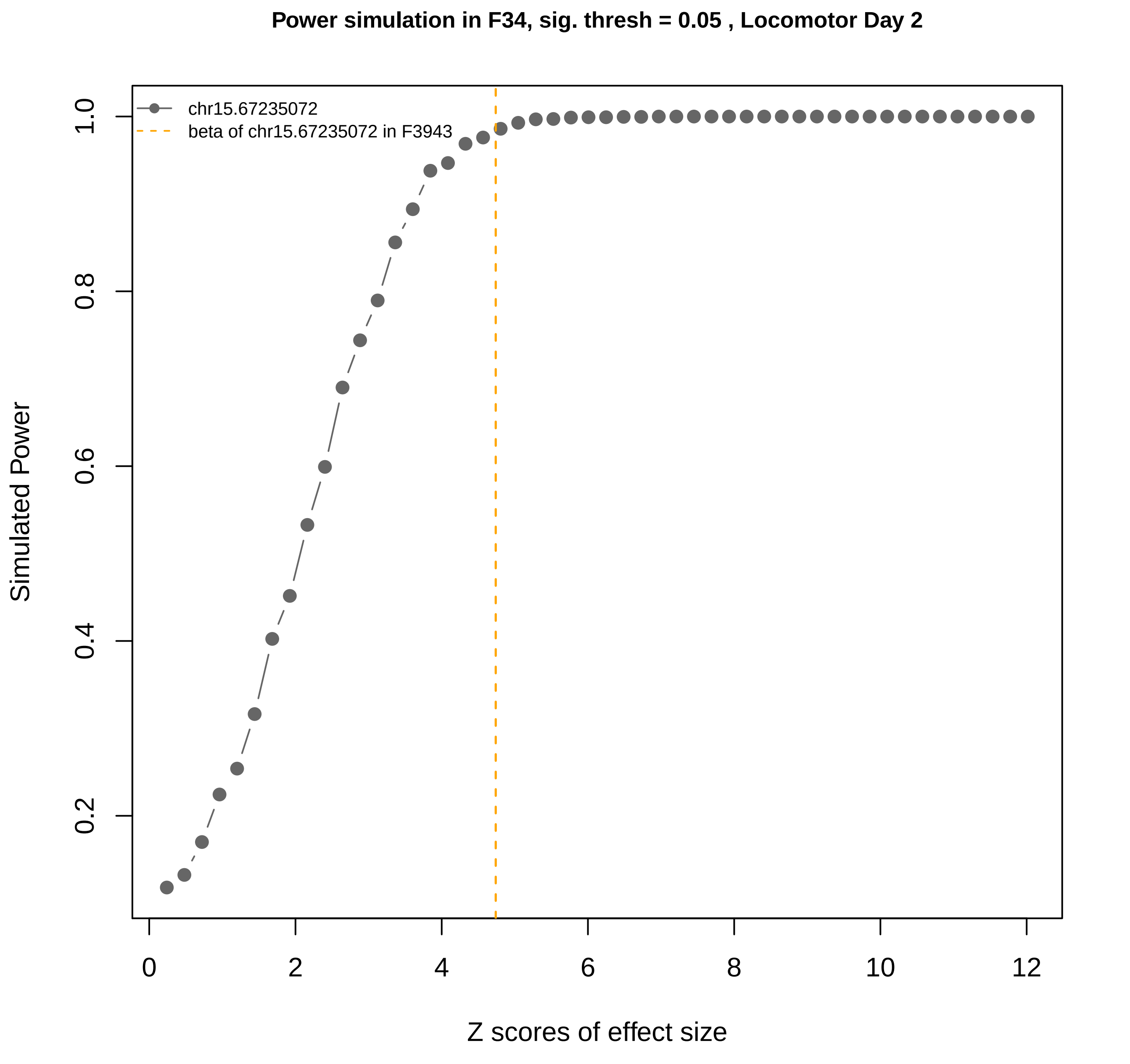

Power simulation in F34, sig. thresh = 0.05 , Locomotor Day 2
chr15.67235072
beta of chr15.67235072 in F3943
Simulated Power
Z scores of effect size
0.2
0.4
0.6
0.8
1.0
0
2
4
6
8
10
12

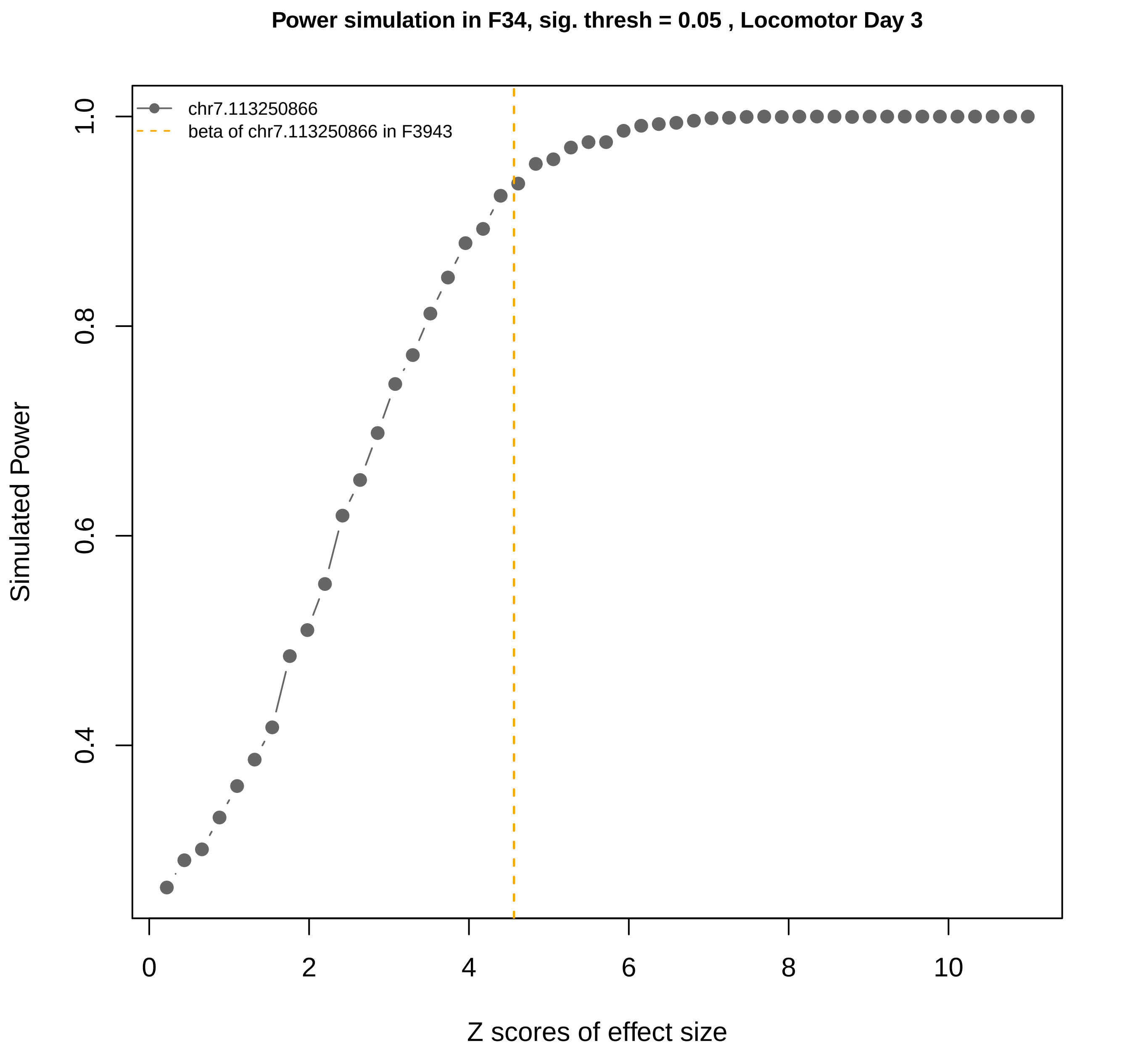

Power simulation in F34, sig. thresh = 0.05 , Locomotor Day 3
chr7.113250866
beta of chr7.113250866 in F3943
Simulated Power
1.0
0.8
0.6
0.4
0
2
4
6
8
10
Z scores of effect size

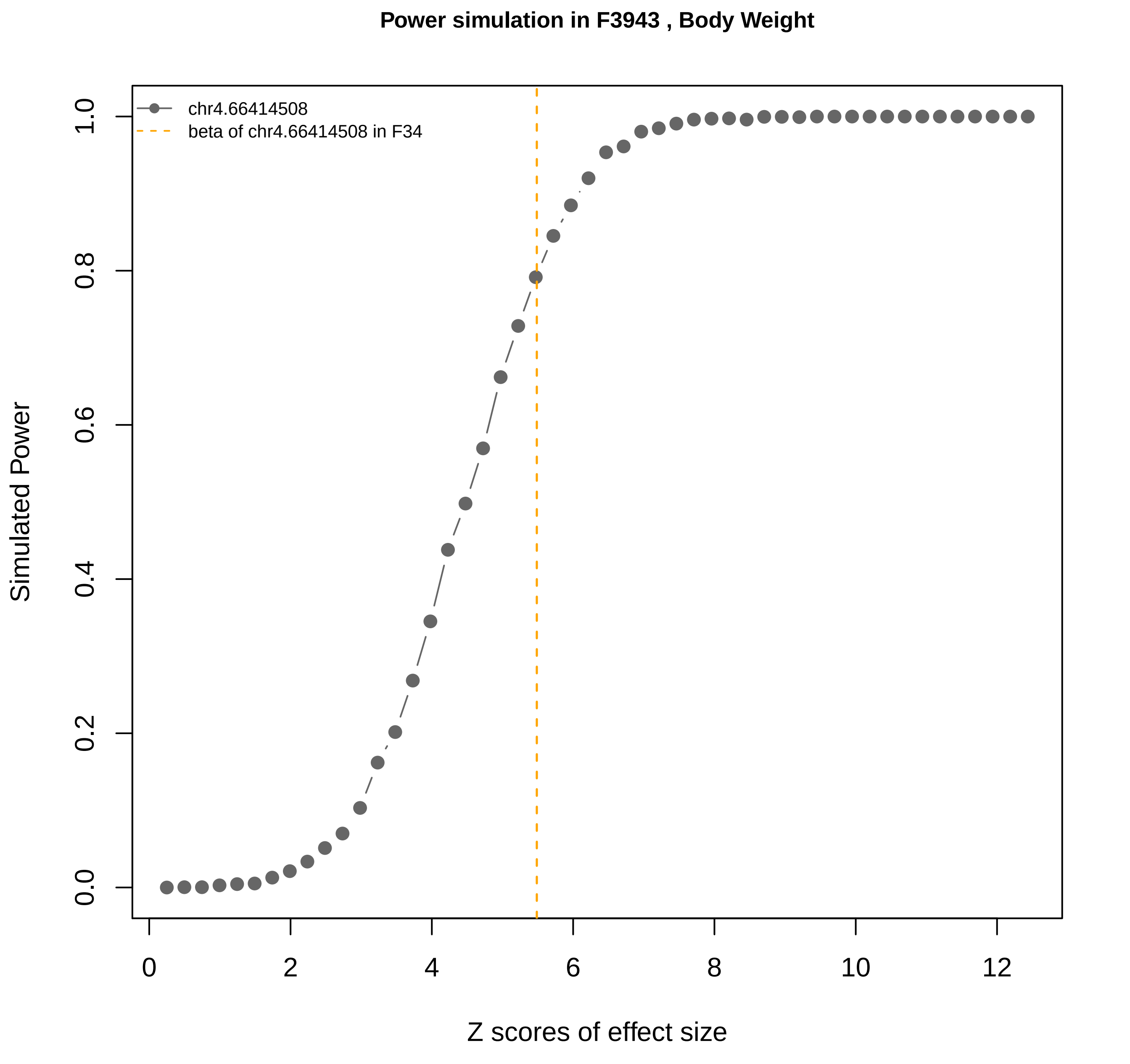
Power simulation in F3943 , Body Weight
chr4.66414508
beta of chr4.66414508 in F34
Simulated Power
Z scores of effect size
0.0
0.2
0.4
0.6
0.8
1.0
0
2
4
6
8
10
12

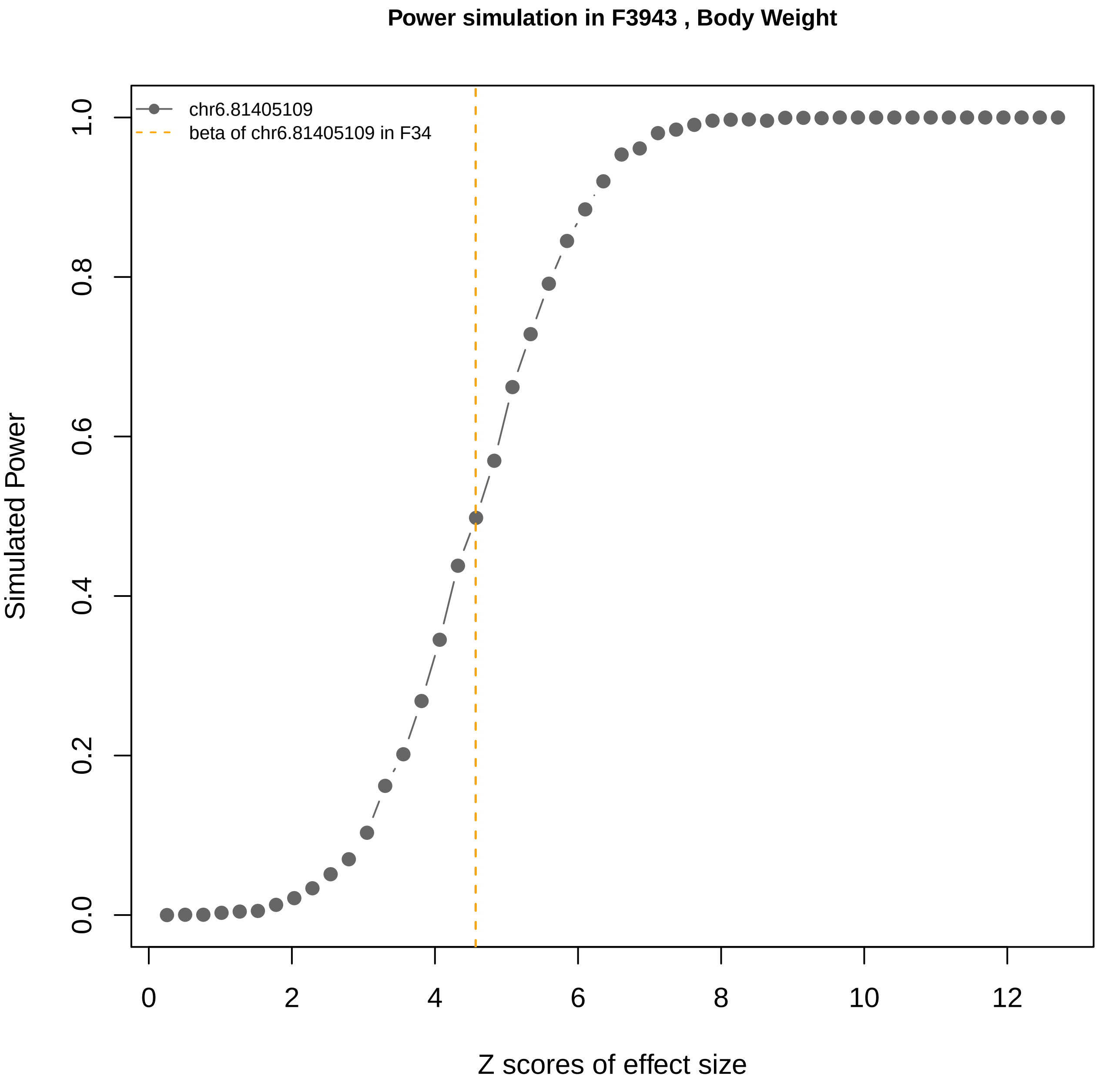
Power simulation in F3943 , Body Weight
chr6.81405109
beta of chr6.81405109 in F34
Simulated Power
Z scores of effect size
0.0
0.2
0.4
0.6
0.8
1.0
0
2
4
6
8
10
12

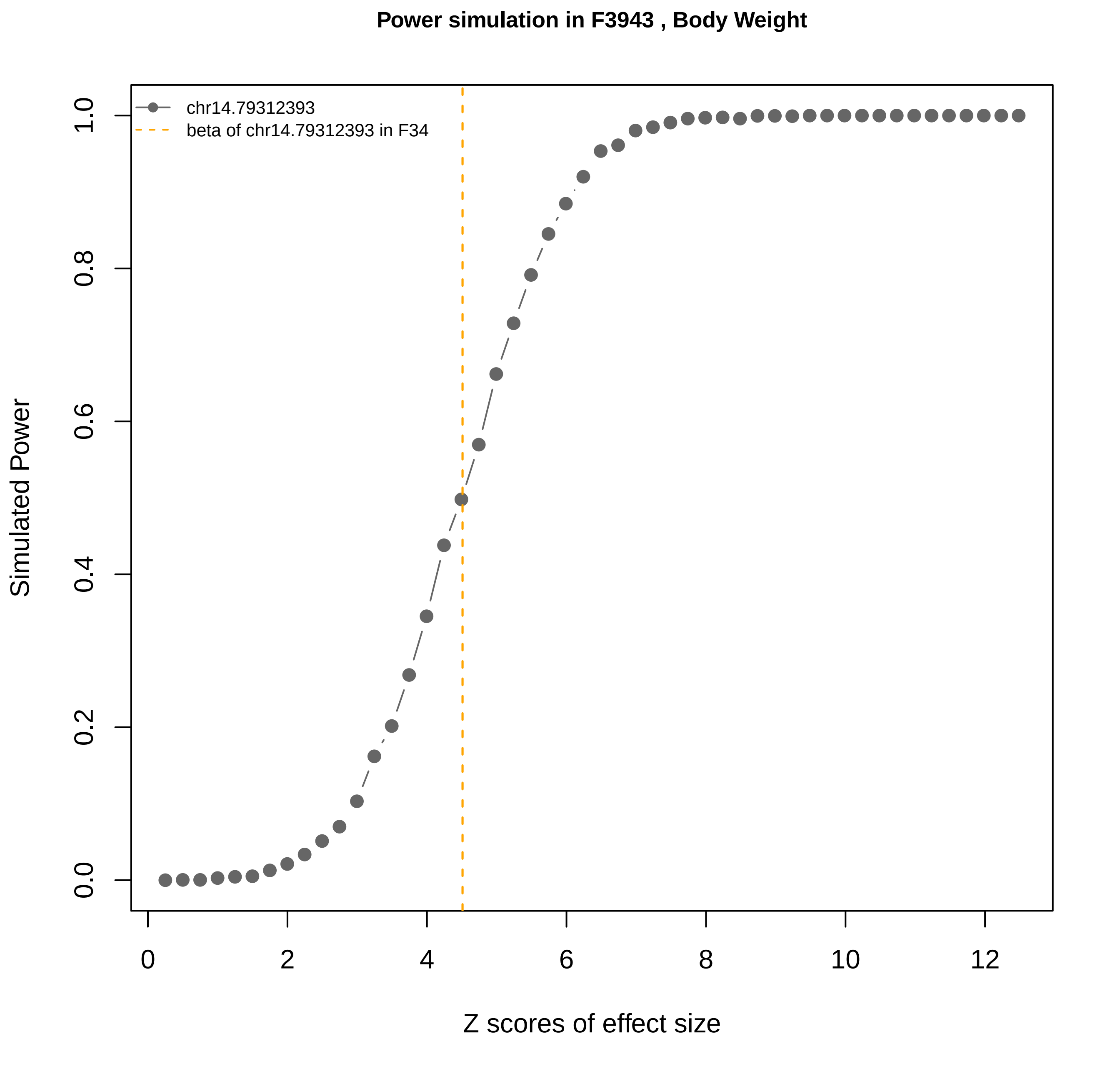
Power simulation in F3943 , Body Weight
chr14.79312393
beta of chr14.79312393 in F34
Simulated Power
Z scores of effect size
0.0
0.2
0.4
0.6
0.8
1.0
0
2
4
6
8
10
12

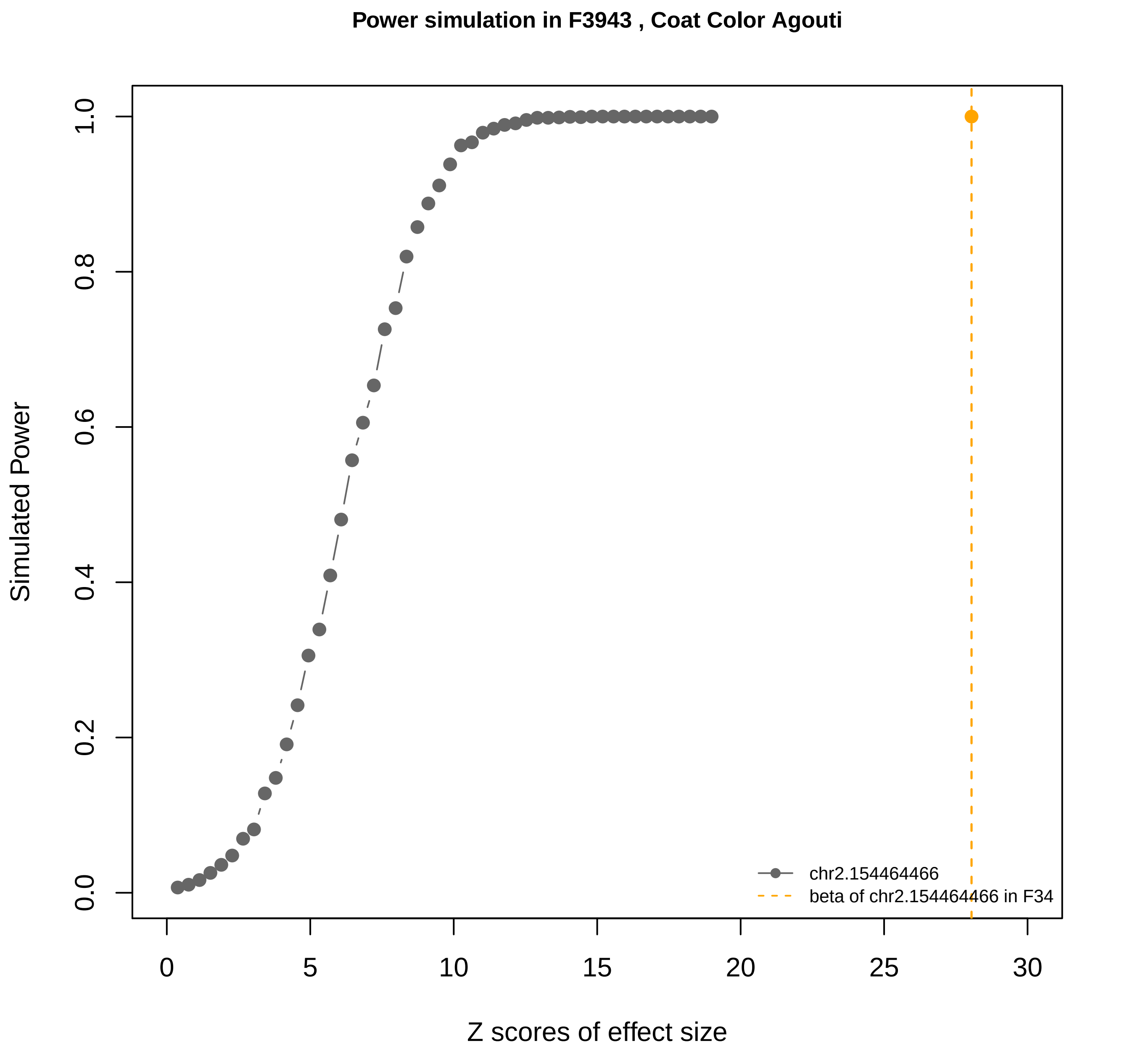
Power simulation in F3943 , Coat Color Agouti
Simulated Power
0.0
0.2
0.4
0.6
0.8
1.0
Z scores of effect size
0
5
10
15
20
25
30
chr2.154464466
beta of chr2.154464466 in F34

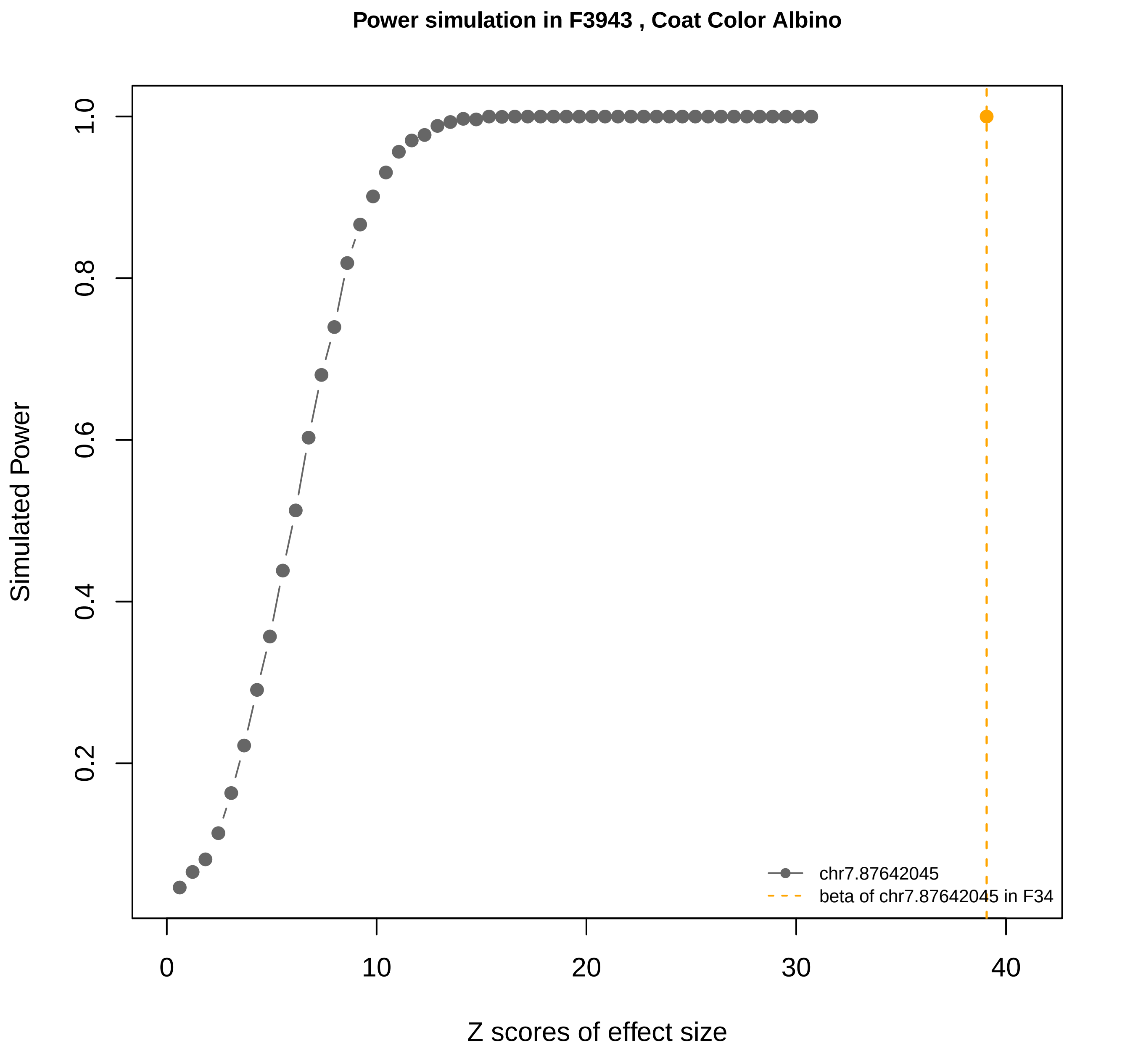

Power simulation in F3943 , Coat Color Albino
Simulated Power
0.2
0.4
0.6
0.8
1.0
0
10
20
30
40
Z scores of effect size
chr7.87642045
beta of chr7.87642045 in F34

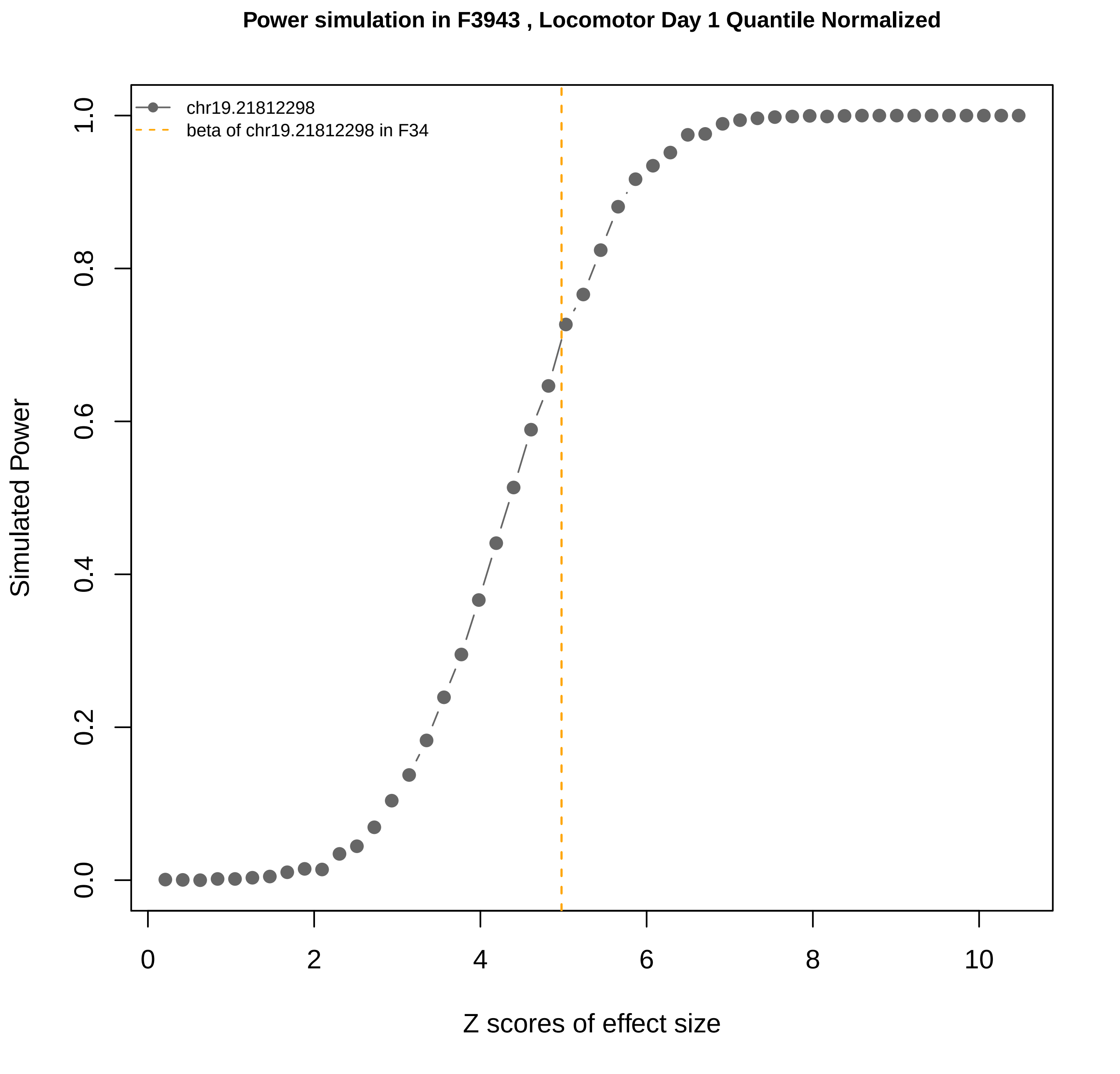

Power simulation in F3943 , Locomotor Day 1 Quantile Normalized
chr19.21812298
beta of chr19.21812298 in F34
Simulated Power
Z scores of effect size
0.0
0.2
0.4
0.6
0.8
1.0
0
2
4
6
8
10

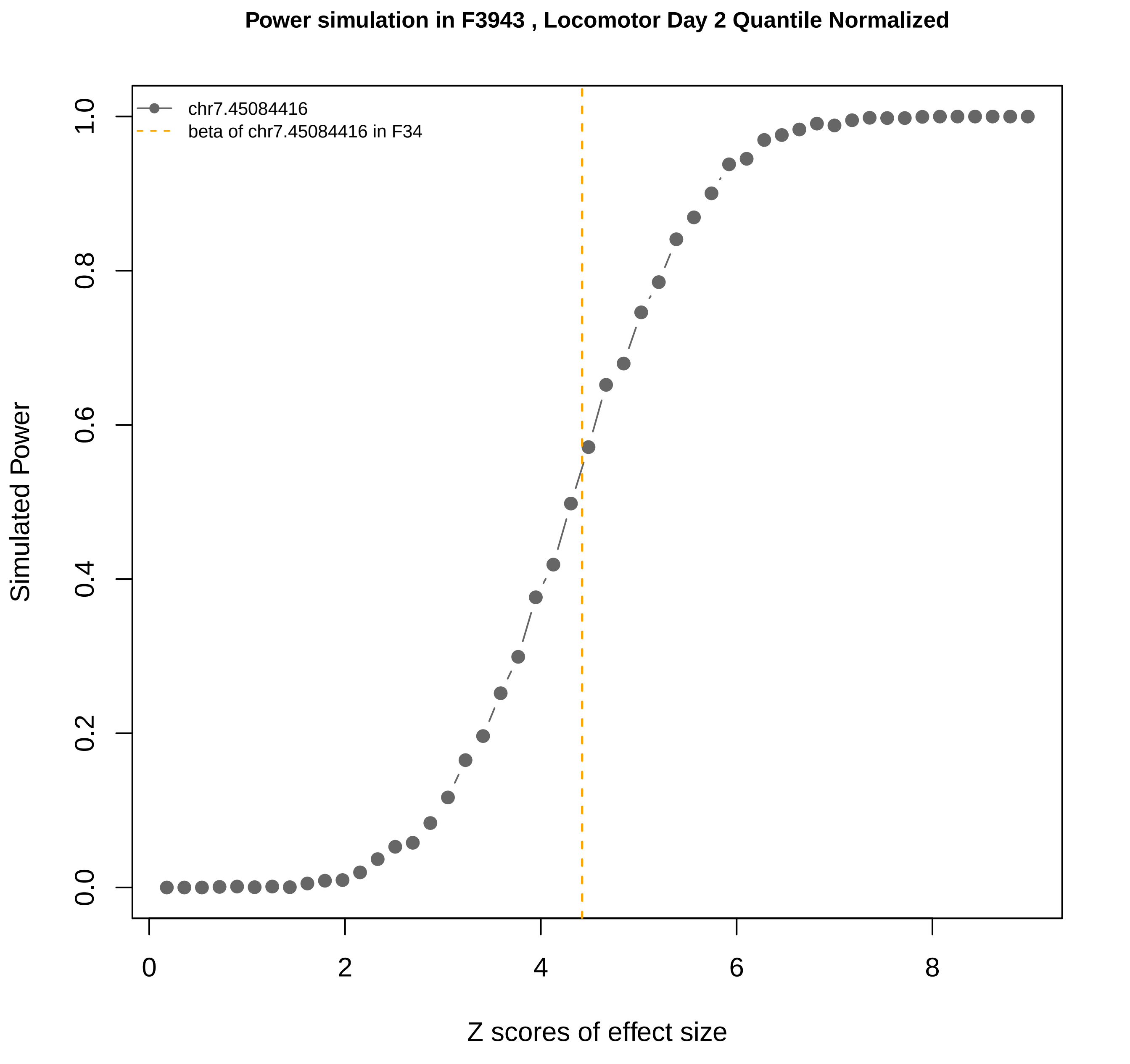

Power simulation in F3943 , Locomotor Day 2 Quantile Normalized
chr7.45084416
beta of chr7.45084416 in F34
Simulated Power
Z scores of effect size
0.0
0.2
0.4
0.6
0.8
1.0
0
2
4
6
8

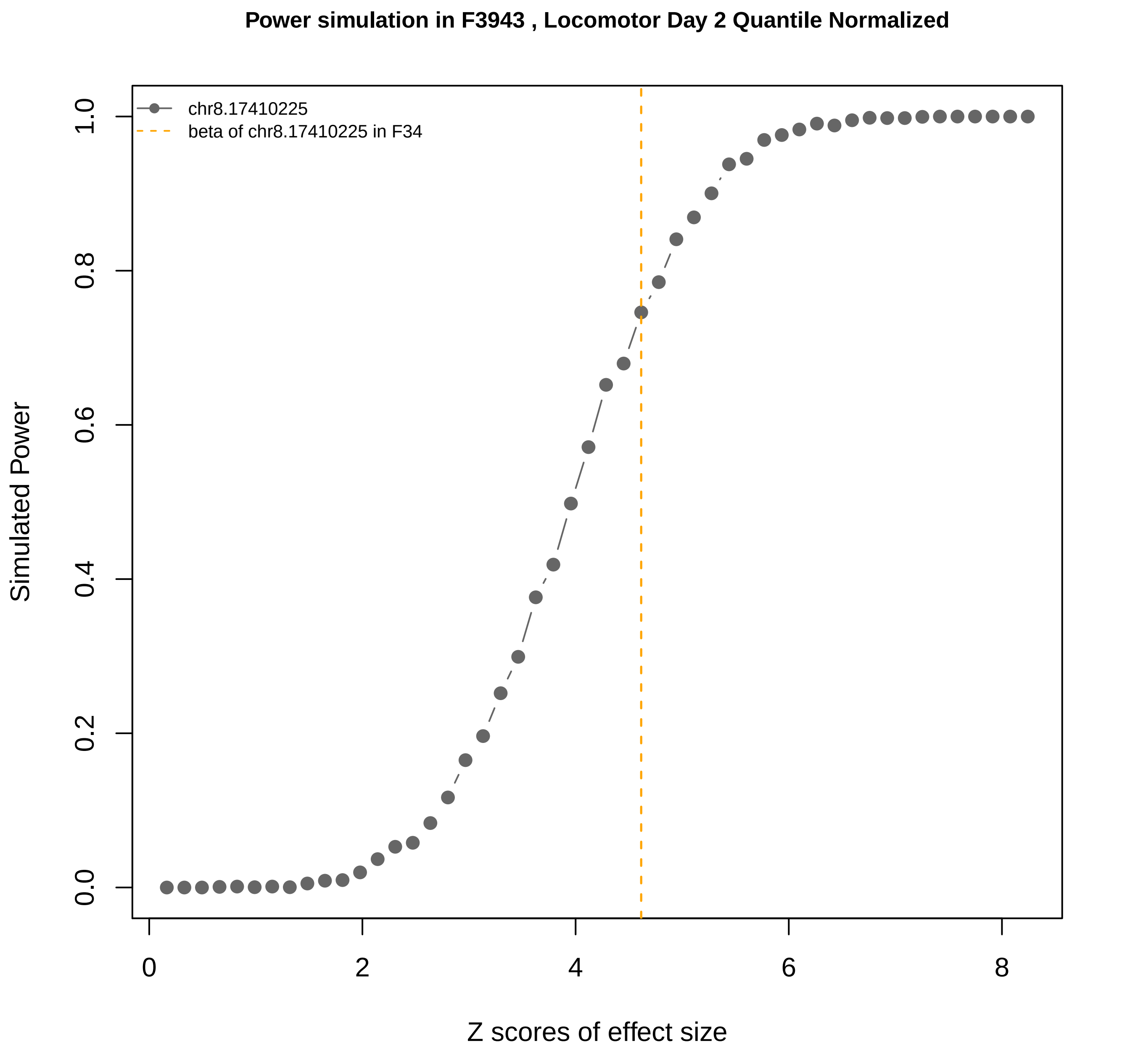

Power simulation in F3943 , Locomotor Day 2 Quantile Normalized
chr8.17410225
beta of chr8.17410225 in F34
Simulated Power
0.0
0.2
0.4
0.6
0.8
1.0
0
2
4
6
8
Z scores of effect size

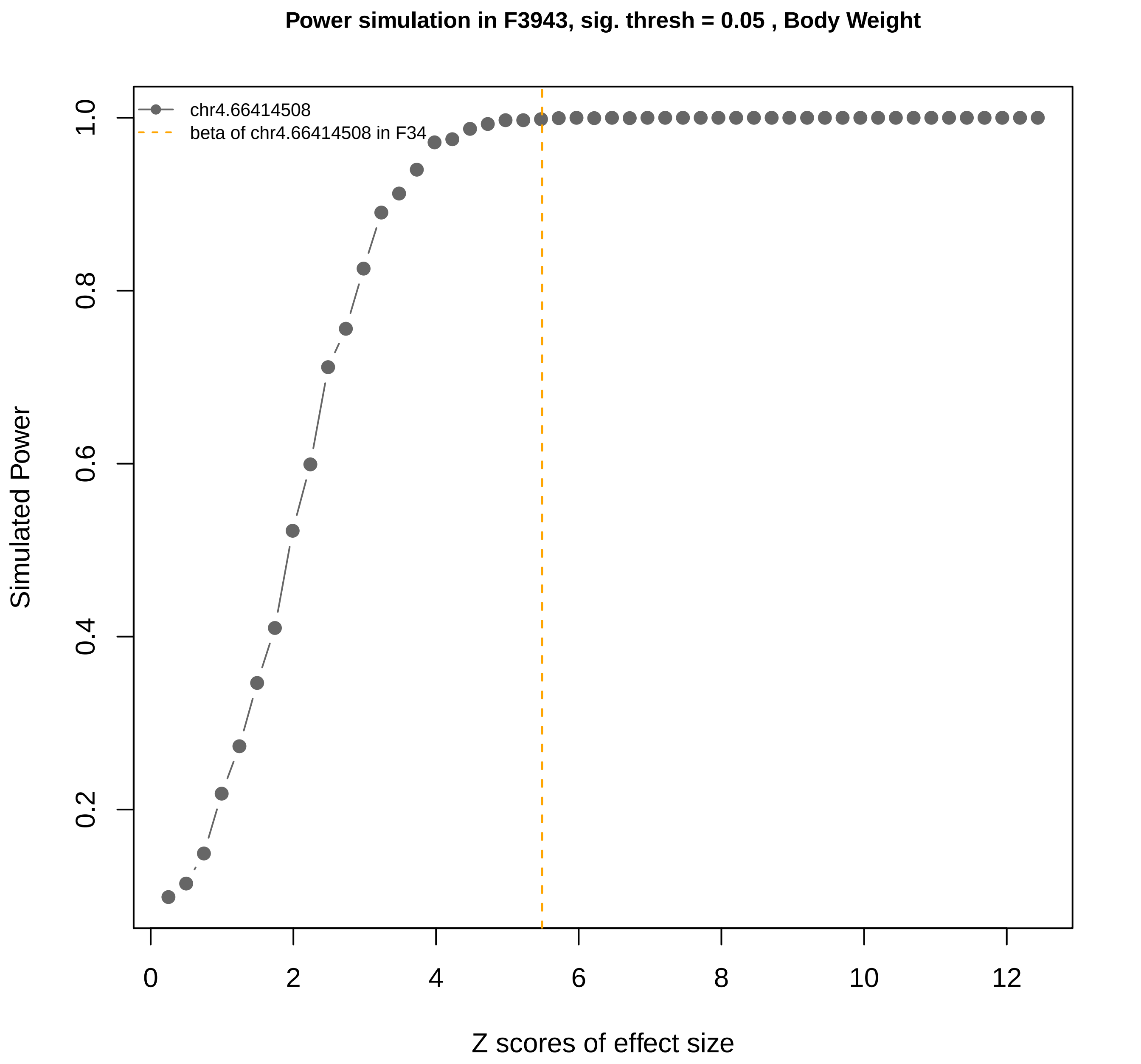
Power simulation in F3943, sig. thresh = 0.05 , Body Weight
chr4.66414508
beta of chr4.66414508 in F34
Simulated Power
1.0
0.8
0.6
0.4
0.2
0
2
4
6
8
10
12
Z scores of effect size

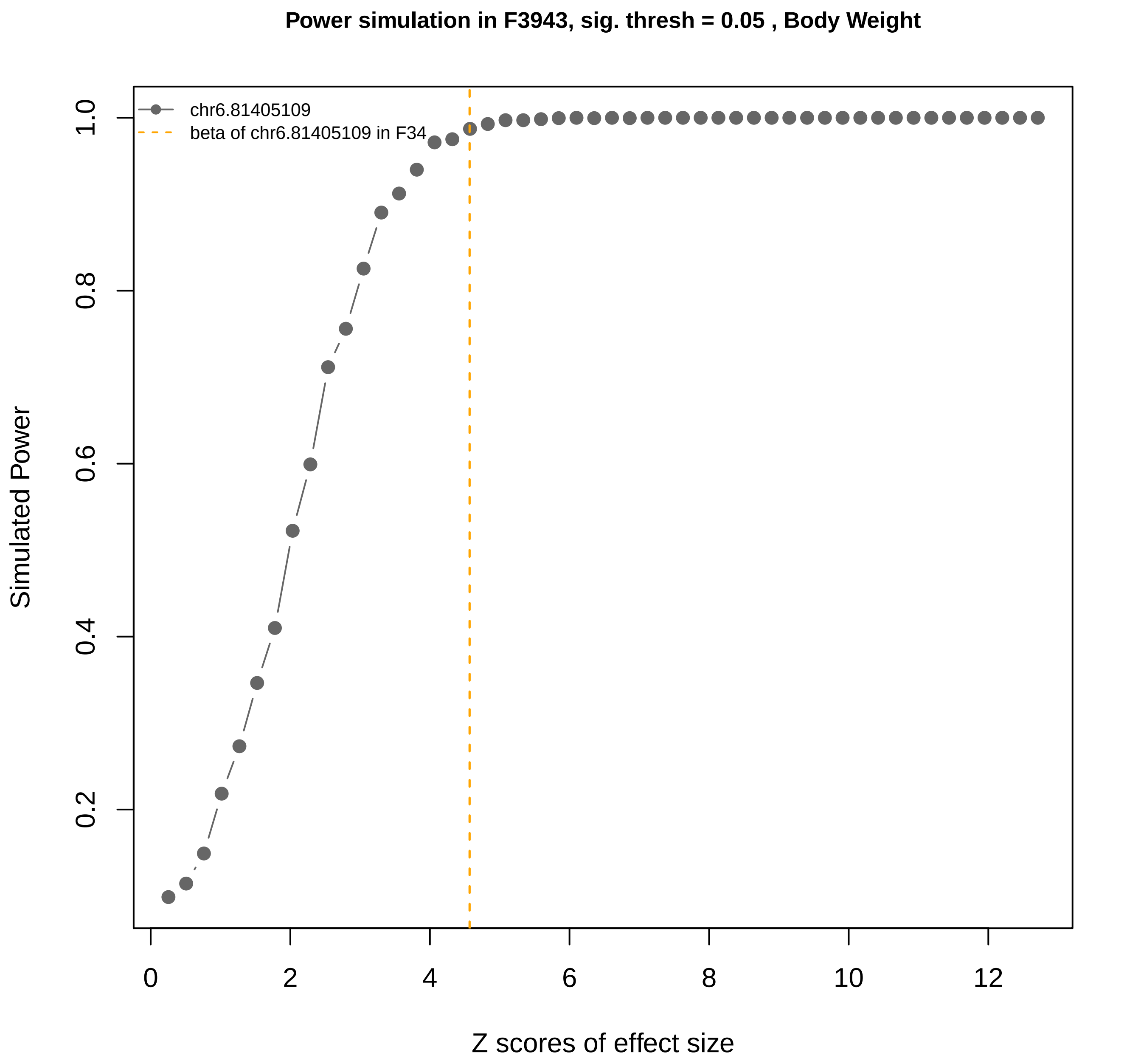
Power simulation in F3943, sig. thresh = 0.05 , Body Weight
chr6.81405109
beta of chr6.81405109 in F34
Simulated Power
Z scores of effect size
0.2
0.4
0.6
0.8
1.0
0
2
4
6
8
10
12

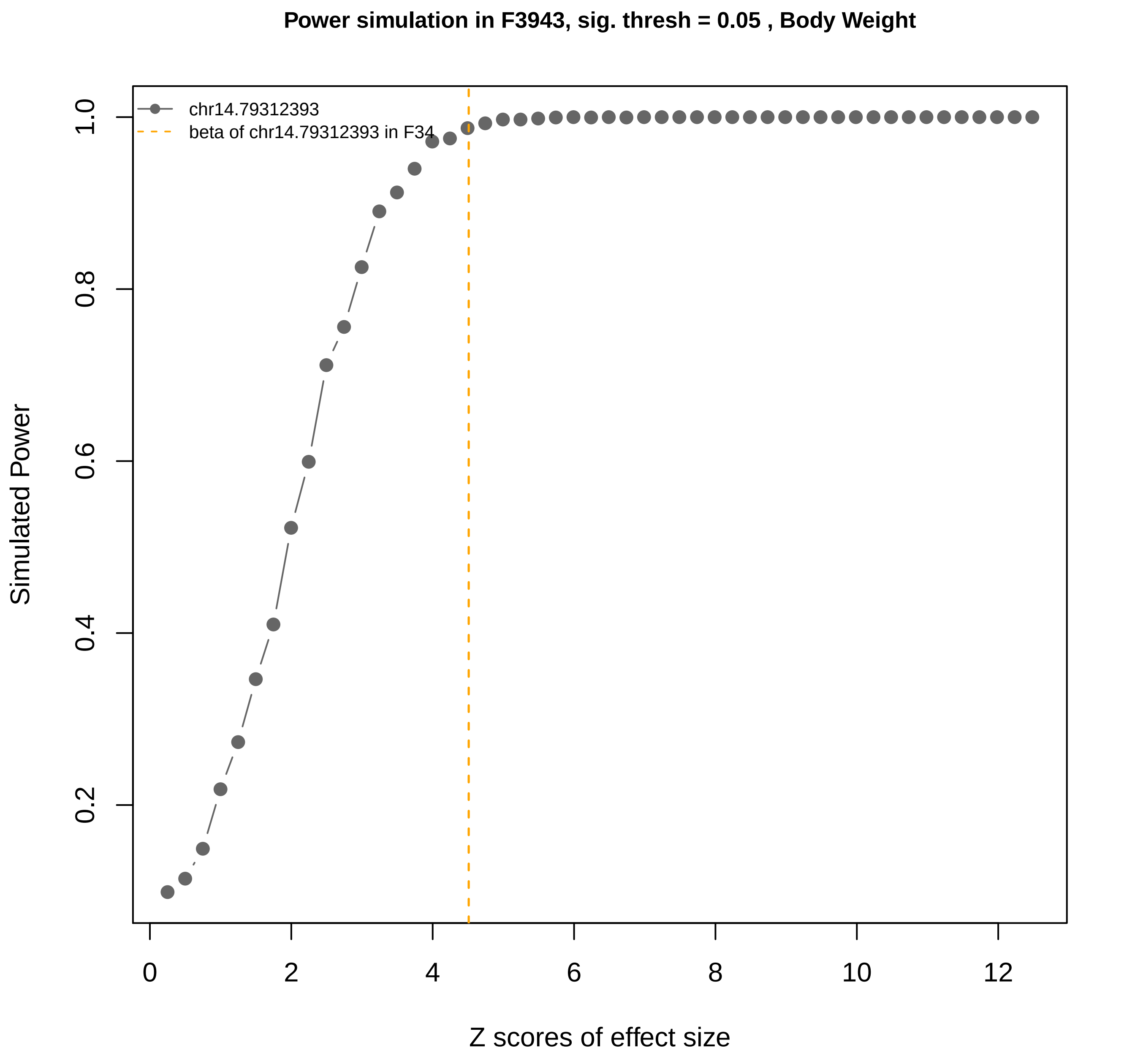

Power simulation in F3943, sig. thresh = 0.05 , Body Weight
chr14.79312393
beta of chr14.79312393 in F34
Simulated Power
0.2
0.4
0.6
0.8
1.0
0
2
4
6
8
10
12
Z scores of effect size

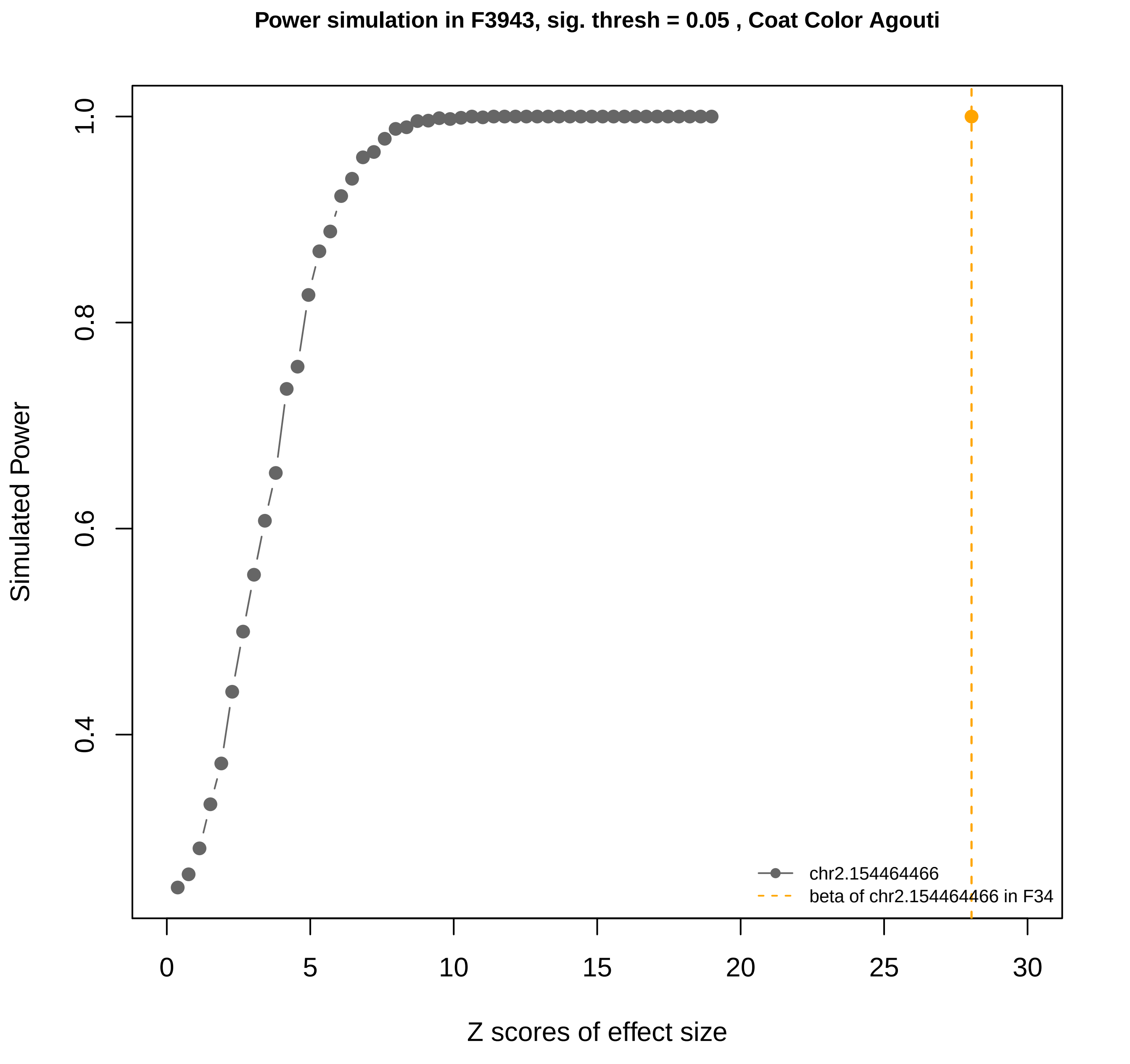

Power simulation in F3943, sig. thresh = 0.05 , Coat Color Agouti
Simulated Power
0.4
0.6
0.8
1.0
0
5
10
15
20
25
30
Z scores of effect size
chr2.154464466
beta of chr2.154464466 in F34

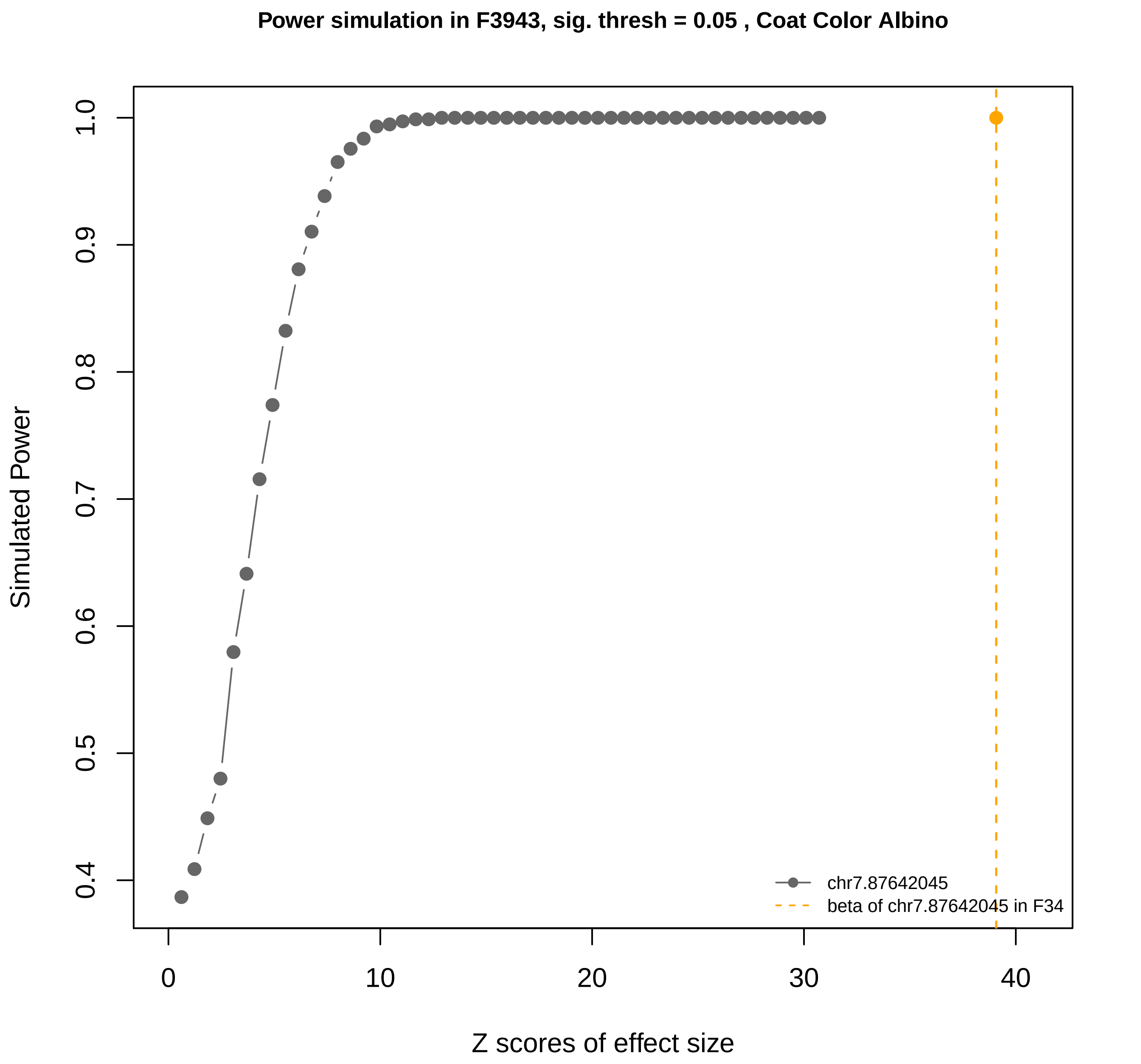
Power simulation in F3943, sig. thresh = 0.05 , Coat Color Albino
Simulated Power
Z scores of effect size
0.4
0.5
0.6
0.7
0.8
0.9
1.0
0
10
20
30
40
chr7.87642045
beta of chr7.87642045 in F34

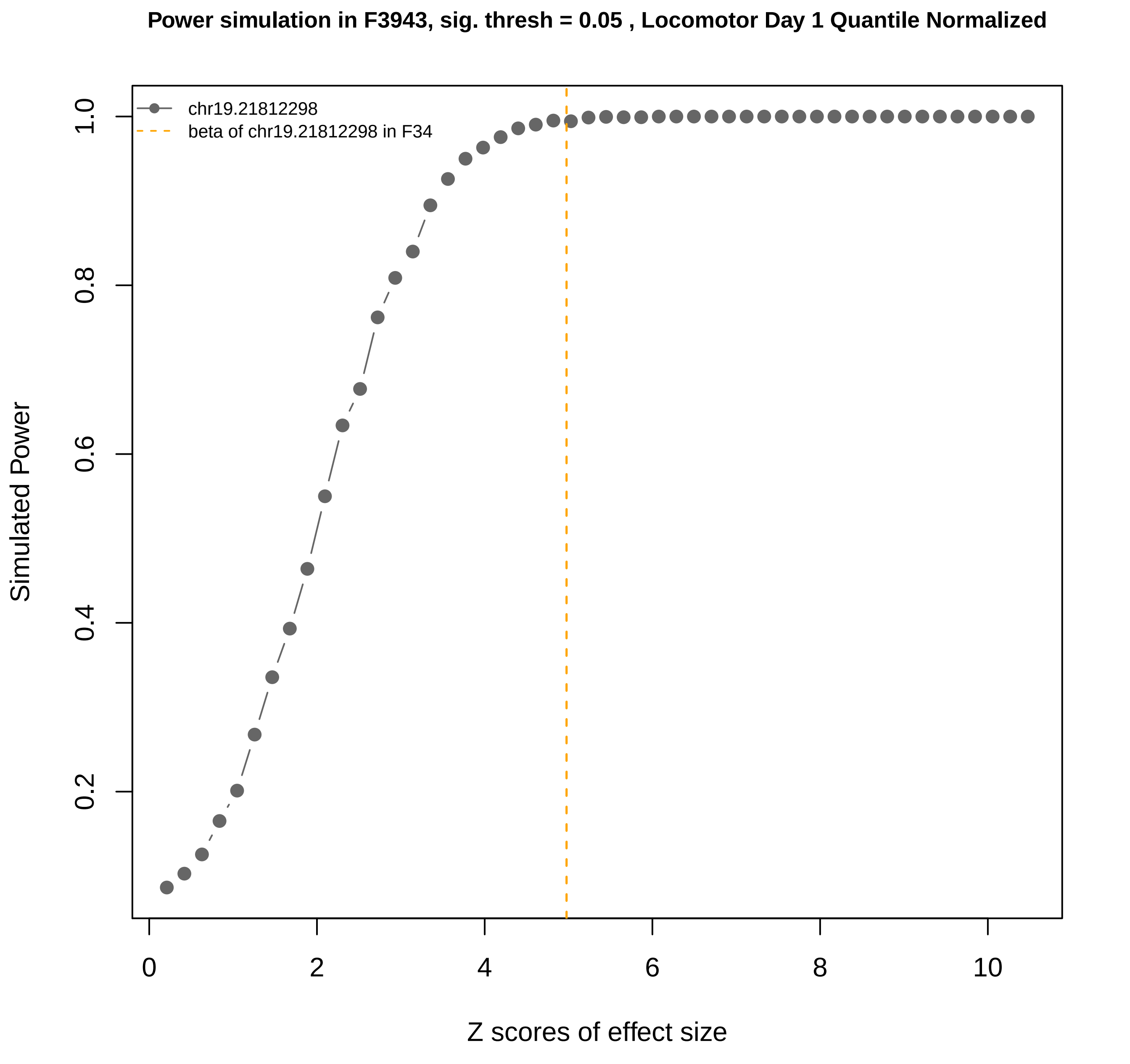

Power simulation in F3943, sig. thresh = 0.05 , Locomotor Day 1 Quantile Normalized
chr19.21812298
beta of chr19.21812298 in F34
Simulated Power
0.2
0.4
0.6
0.8
1.0
Z scores of effect size
0
2
4
6
8
10

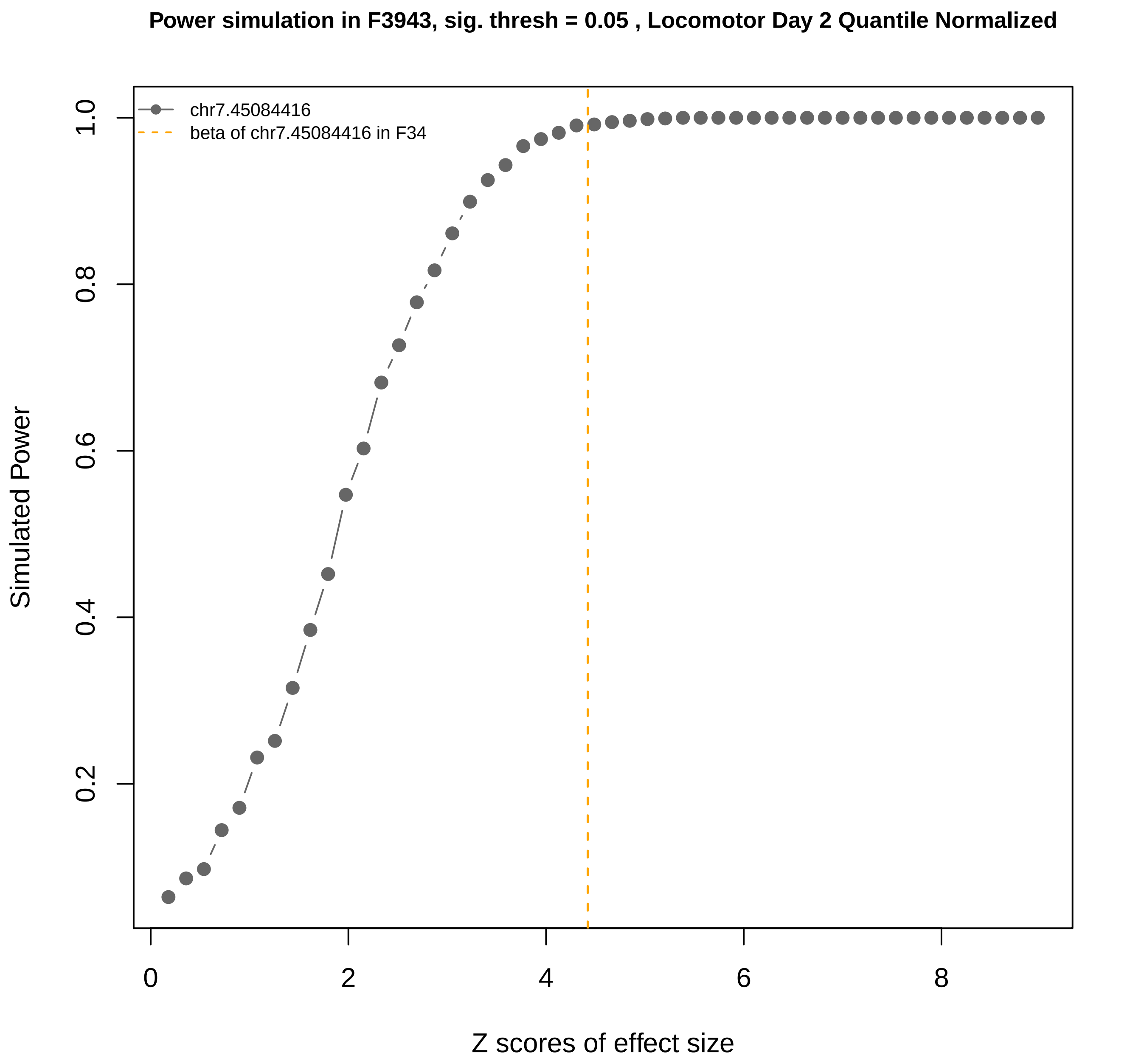

Power simulation in F3943, sig. thresh = 0.05 , Locomotor Day 2 Quantile Normalized
chr7.45084416
beta of chr7.45084416 in F34
Simulated Power
0.2
0.4
0.6
0.8
1.0
0
2
4
6
8
Z scores of effect size

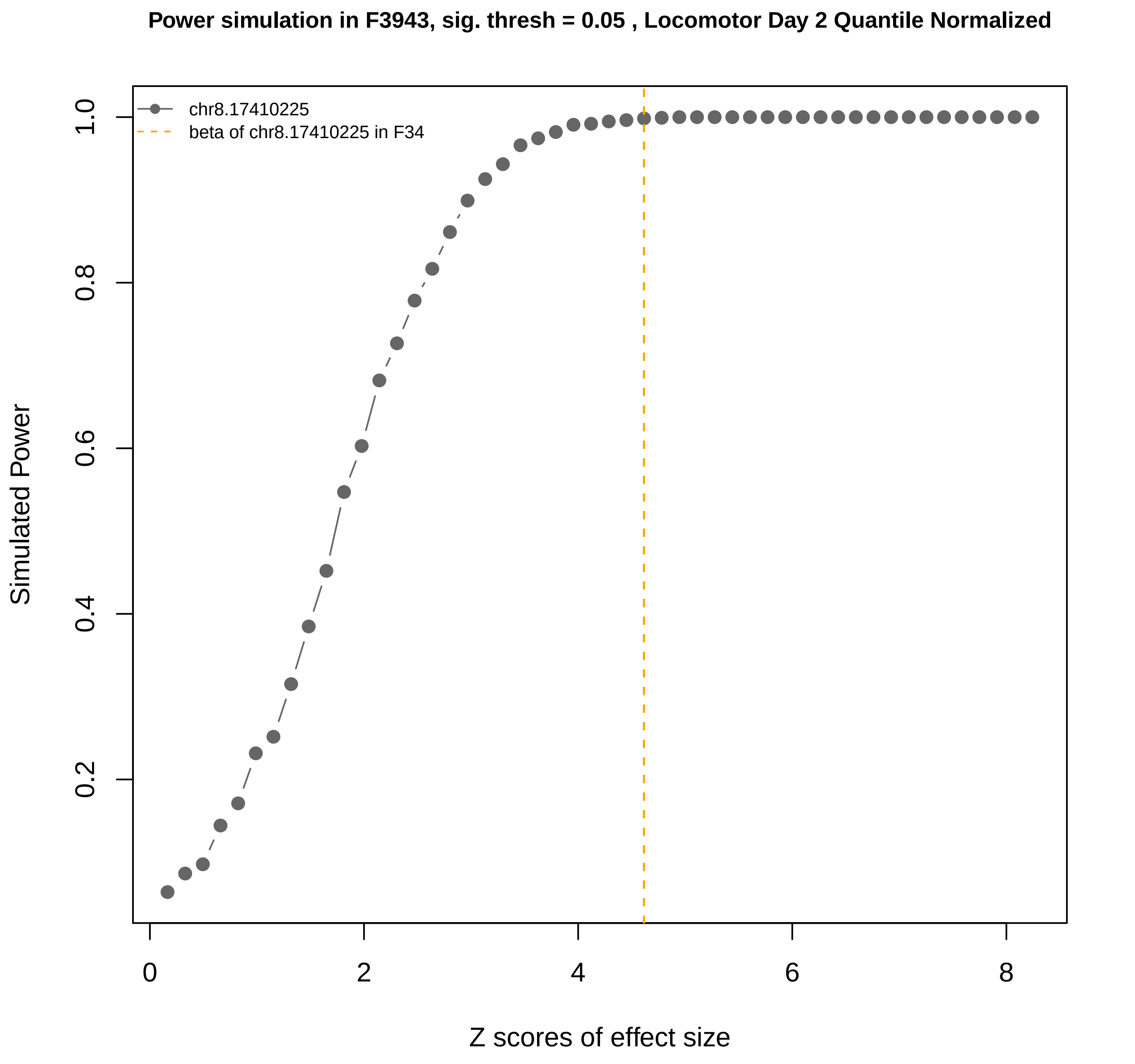

Power simulation in F3943, sig. thresh = 0.05 , Locomotor Day 2 Quantile Normalized
chr8.17410225
beta of chr8.17410225 in F34
Simulated Power
0.2
0.4
0.6
0.8
1.0
0
2
4
6
8
Z scores of effect size
